# statSuma: automated selection and performance of statistical comparisons for microbiome studies

**DOI:** 10.1101/2021.06.15.448299

**Authors:** R.J. Leigh, R.A. Murphy, F. Walsh

## Abstract

There is a reproducibility crisis in scientific studies. Some of these crises arise from incorrect application of statistical tests to data that follow inappropriate distributions, have inconsistent equivariance, or have very small sample sizes. As determining which test is most appropriate for all data in a multicategorical study (such as comparing taxa between sites in microbiome studies), we present statsSuma, an interactive Python notebook (which can be run from any desktop computer using the Google Colaboratory web service) and does not require a user to have any programming experience. This software assesses underlying data structures in a given dataset to advise what pairwise or listwise statistical procedure would be best suited for all data. As some users may be interested in further mining specific trends, statSuma performs 5 different two-tailed pairwise tests (Student’s *t*-test, Welch’s *t*-test, Mann-Whitney *U*-test, Brunner-Munzel test, and a pairwise Kruskal-Wallis *H*-test) and advises the best test for each comparison. This software also advises whether ANOVA or a multicategorical Kruskal-Wallis *H-*test is most appropriate for a given dataset and performs both procedures. A data distribution-*vs*-Gaussian distribution plot is produced for each taxon at each site and a variance plot between all combinations of 2 taxa at each site are produced so Gaussian tests and variance tests can be visually confirmed alongside associated statistical determinants.

## Introduction

The advent of readily available, low-cost high throughput DNA sequencing has promoted the rampant growth of microbiome and metagenome studies. These studies, while informative and insightful, are often fraught by well-meaning but ultimately incorrect data assumptions and statistical test applications (Martin, 2019; Free, 2020). The selection of statistical tests requires knowledge of underlying data characteristics such as sample distribution, sample size and equivariance (Figure 1), which can be easily misinterpreted especially for complex studies with a lot of groups and relatively small sample sizes per group (Makin and De Xivry, 2019; Konietschke, Schwab and Pauly, 2021). Statistical analyses are of considerable importance in microbiology and microbiome research to ensure accurate result reporting and optimal experimental design (Adams-Huet and Ahn, 2009). Statistical analysis for non-statisticians is often cumbersome, requiring a researcher to purchase paid statistical software or to use a programming language (*eg*. Python or R) for analyses not available in such packages.

**Figure 1:**
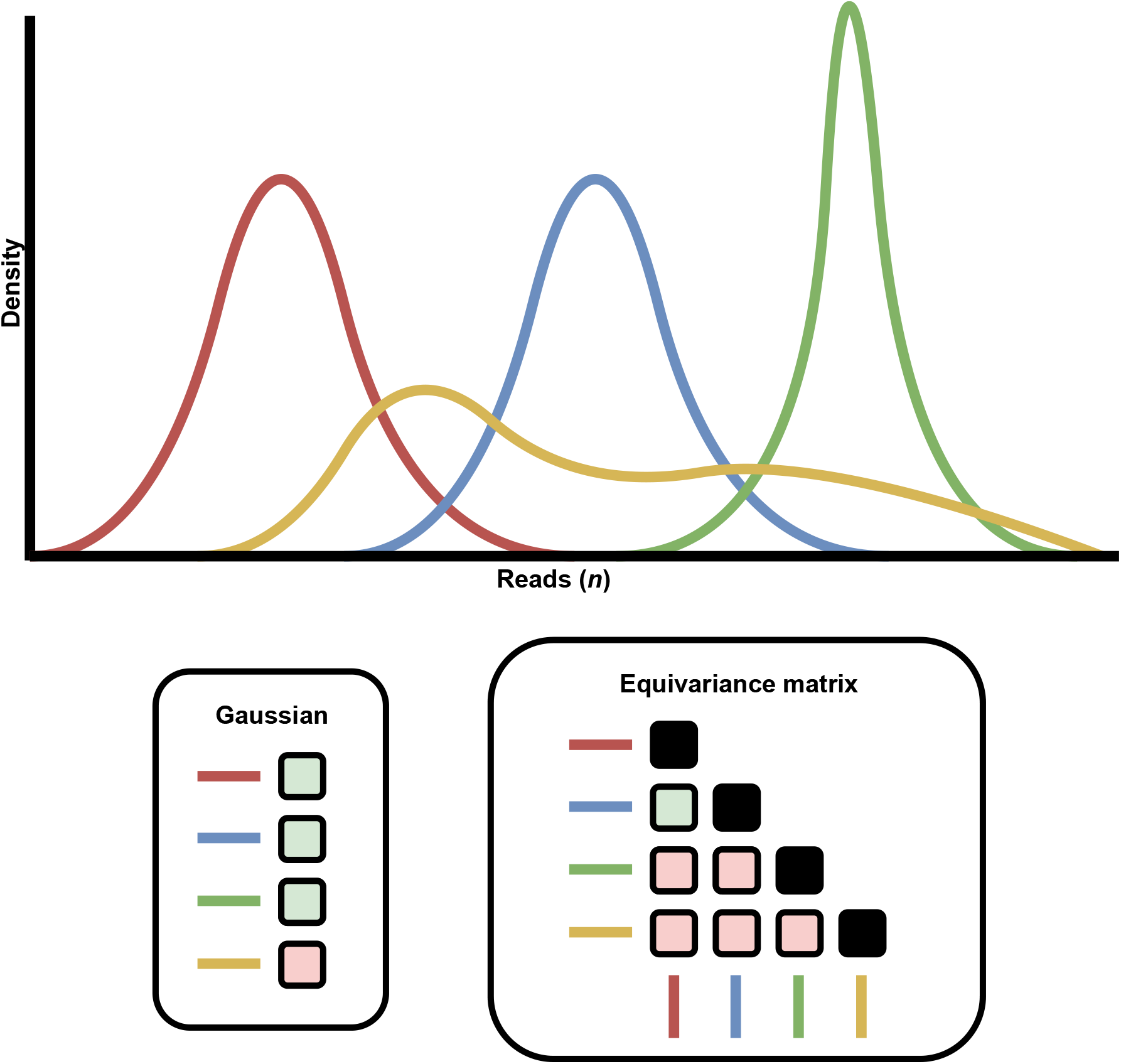
Example of distributions and equivariance. In this example, 4 distributions are presented, 3 of which are Gaussian (as determined by the green box in the Gaussian grid) and one is non-Gaussian. There is one instance of equivariance in this example (between the red and blue distributions; indicated by the green box in the equivariance matrix) and the rest are non-equivariant (denoted as red boxes in the equivariance matrix). Black boxes in the equivariance matrix indicate a non-comparator.

Here, we present statSuma, an interactive Python notebook (which can be run from any desktop computer using the Google Colaboratory web service) for the automatic selection (and reasoning behind the selection) of statistical comparisons and performance of said comparisons for a user. The software is most concerned with the some of the most popular comparison procedures: Student’s (Gosset’s) *t*-test (Student, 1908), Welch’s *t*-test (Welch, 1947), Mann-Whitney *U*-test (Mann and Whitney, 1947), Brunner-Munzel test (Brunner and Munzel, 2000), analysis-of-variance (ANOVA) (Fisher, 1921), and the Kruskal-Wallis H-test (Kruskal and Wallis, 1952). Due to the excessive variance associated with microbiome studies (Falony *et al*., 2016; Leigh, Murphy and Walsh, 2021), it is anticipated that comparisons utilising equivariance will be rare.

## Methods

### Test selection

The software focusses on the most used comparisons in microbiome studies where all comparisons discussed will focus on two-tailed analysis of independent data. For the purposes of these explanations, it is assumed that all groups used for a given comparison have more than one sample > 0 and have a standard deviation > 0. Selection of a particular test is predicated on 6 main criteria:

(*a*.): The number of groups (*n*_groups_) to be analysed (*n*_groups_ = 2; *n*_groups_ ≥ 2)

(*b*.): The minimum number of samples (*n*_samples_) within given groups

(*c*.): Whether groups have equal sample sizes

(*d*.): Whether samples are equivariant

(*e*.): The underlying distribution of the data (Gaussian or non-Gaussian)

(*f*.): Underlying hypothesis to be tested (discussed below)

Once all these criteria are satisfactorily addressed, the most appropriate statistical comparison is determined. The user selects whether a test is to be pairwise (*n*_groups_ = 2) or listwise (*n*_groups_ > 2) and statSuma determines the minimum *n*_samples_ for each comparison and whether sample sizes are equal between all groups, thus satisfying these requirements.

A Levene’s test (Levene, 1960) is employed to assess equivariance (H_0_:σ^2^_(*a*)_ =σ^2^_(*b*)_ ;H_A_:σ^2^_(*a*)_ ≠σ^2^_(*b*)_), where a *P* ≥ α is used to determine equivariance. Traditionally, α = 0.05 so a *P* ≥ 0.05 determines equivariance (a 5% likelihood of equivariance is determined to be equivariant).

A Shapiro-Wilk test (Shapiro and Wilk, 1965) is employed to determine whether both datasets follow a Gaussian distribution (H_0_:*X*∼*N*(μ,σ^2^);H_A_:*X*≁*N*(μ,σ^2^)). Again, a *P* ≥ α (α = 0.05) is used to determine whether a Gaussian distribution is observed. As with the Levene’s test, statSuma sets α to 0.5 to determine if a distribution is Gaussian by default (which can be changed by a user).

For pairwise tests, if both distributions are determined to be equivariant and Gaussian, a Student’s *t*-test is most appropriate (Student, 1908). In this scenario, the means of two groups are compared (H_0_:μ_(*a*)_=μ_(*b*)_;H_A_:μ_(*a*)_≠μ_(*b*)_). If both distributions are assumed to be Gaussian but not equivariant the Welch’s *t*-test is most appropriate (Welch, 1947), again this is a comparison of means (H_0_:μ_(*a*)_=μ_(*b*)_;H_A_:μ_(*a*)_≠μ_(*b*)_). Both *t*-tests can technically be performed with a minimum sample size of 2, however larger sample sizes are strongly recommended to ensure statistical power (Rusticus and Lovato, 2014). If either distribution is non-Gaussian but are equivariant, a Mann-Whitney *U*-test is most appropriate. Generally speaking, the Mann-Whitney *U*-test is a comparison of medians which is also sensitive differences in data distributions (H_0_:η_(*a*)_=η_(*b*)_;H_A_:η_(*a*)_≠η_(*b*)_). A Mann-Whitney U-test requires at least 8 samples per group to correctly function (Mann and Whitney, 1947; Cheung and Klotz, 1997). While the Student’s *t*-test, Welch’s *t*-test, and Mann-Whitney *U*-test do not require equal sample sizes, losses in statistical power are proportionate to differences in sample sizes between groups (Rusticus and Lovato, 2014).

If either distribution is non-Gaussian and equivariance is not observed, a Brunner-Munzel test is most appropriate if a minimum *n*_sample_ ≥ 10 is observed. The Brunner-Munzel test assesses stochastic equality (H_0_:*B*=0.5;H_A_:*B*≠0.5), which determines whether samples in one group are larger than samples in another (Brunner and Munzel, 2000). Finally, if either distribution is non-Gaussian and non-equivariance is observed, a Kruskal-Wallis H test is performed if the minimum *n*_sample_ ≥ 5. Generally speaking, like the Mann-Whitney U-test, the Kruskal-Wallis *H*-test is a comparison of medians (H_0_:η_(*a*)_=η_(*b*)_;H_A_:η_(*a*)_≠η_(*b*)_).

Once all pairwise data examinations have been completed, one test is recommended for all comparisons to ensure an appropriate standard approach to all comparisons to a given study. As comparisons that do not require underlying Gaussian distributions provide accurate results for non-Gaussian data and as comparisons that do not require underlying equivariance provide accurate results for equivariant data (Nahm, 2016; Delacre, Lakens and Leys, 2017), these methods are ranked as follows:

(1.): Kruskal-Wallis *H*-test

(2.): Brunner-Munzel test

(3.): Welch’s *t*-test

(4.): Student’s *t*-test

(5.): Mann-Whitney U-test

These rankings are considered by statSuma to be absolute, and the highest ranked recommendation for any test is recommended for all tests. For example, if the Kruskal-Wallis *H*-test (ranked highest) is recommended for any test, it is recommended for all tests.

Conversely, all recommendations must be Mann-Whitney U-test for the Mann-Whitney U-test to be the overall recommendation. The low ranking of the Mann-Whitney U-test is commiserate with its multifactorial requirements. For transparency, statSuma computes all comparisons provides descriptive statistics (mean, standard deviation, variance, median, maximum, and minimum) using all 5 tests. This is so individual comparisons of interests may be examined (for example, in instances where the minimum *n*_sample_ = 2 in some comparisons; referred to as “Size Student’s *t*-tests” and “Size Welch’s *t*-tests” in statSuma’s outputs). For completeness, *P*-values derived from pairwise tests are presented as naïve *P*-values, Bonferroni-Dunn corrected *P*-values (*P*_BD_) to control for false discovery rates (FDR).

### Listwise tests

For listwise tests, if all distributions are equivariant, have equal sample sizes, and each follow a Gaussian distribution, the most appropriate comparison is an ANOVA. An ANOVA is a comparison of means across 3 or more samples (H_0_:μ_(*a*)_=μ_(*b*)_=…=μ_(*x*)_;H_A_:μ_(*a*)_≠μ_(*b*)_≠…≠μ_(*x*)_). Comparatively, if a ubiquitous Gaussian distribution or non-equivariance are observed, the Kruskal-Wallis H-test may be used to compare the medians (H_0_:η_(*a*)_=η_(*b*)_=…=η_(*x*)_;H_A_:η_(*a*)_≠η_(*b*)_≠…≠η_(*x*)_). The Kruskal-Wallis *H*-test is ranked higher than ANOVA for recommendations. Again, *P*-values derived from listwise tests are presented as naïve *P*-values, Bonferroni-Dunn corrected *P*-values (*P*_BD_) to control for false discovery rates (FDR).

### Plots

For visual analysis, statSuma offers an optional plot for the standardised distribution of each taxon with a generated (standardised) Gaussian distribution for comparison. For these. The *y*-axis is given as probability as the cumulative scores of each bin for each taxon (and associated generated Gaussian curve) each equated to 1 (*eg*. Figure 2). For completeness, QQ-plots are also produced for each taxon (*eg*. Figure 3). Comparatively, optional plots are offered to visually assess equivariance between two taxon distributions. Taxon distributions are each cumulatively standardised so samples with unequal sample sizes can be visualised together (*eg*. Figure 4). Again, *y*-axis is given as probability as the cumulative of each taxon equated to 1.

**Figure 2:**
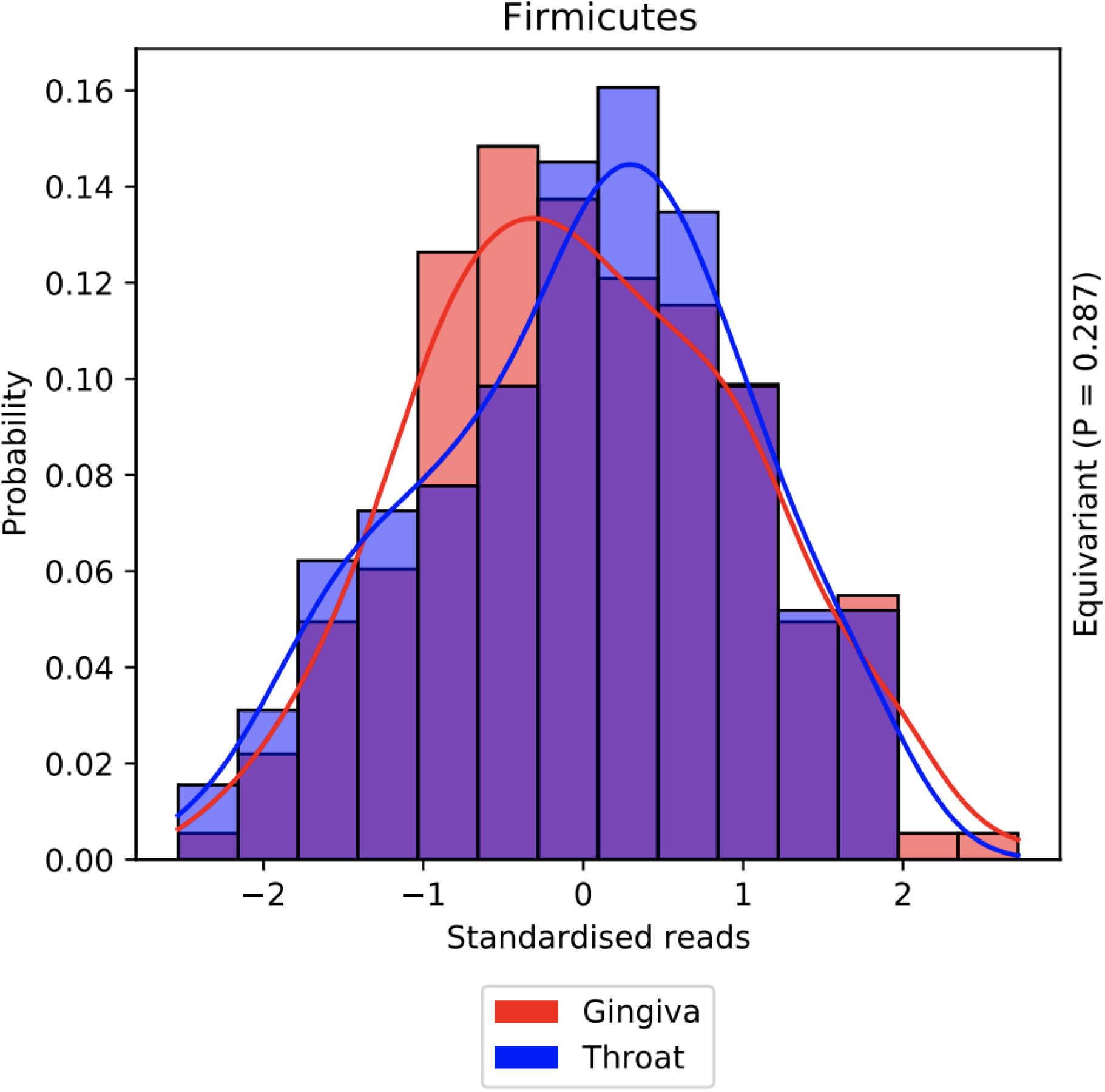
Comparison of two Firmicute distributions from two anatomical sites.

**Figure 3:**
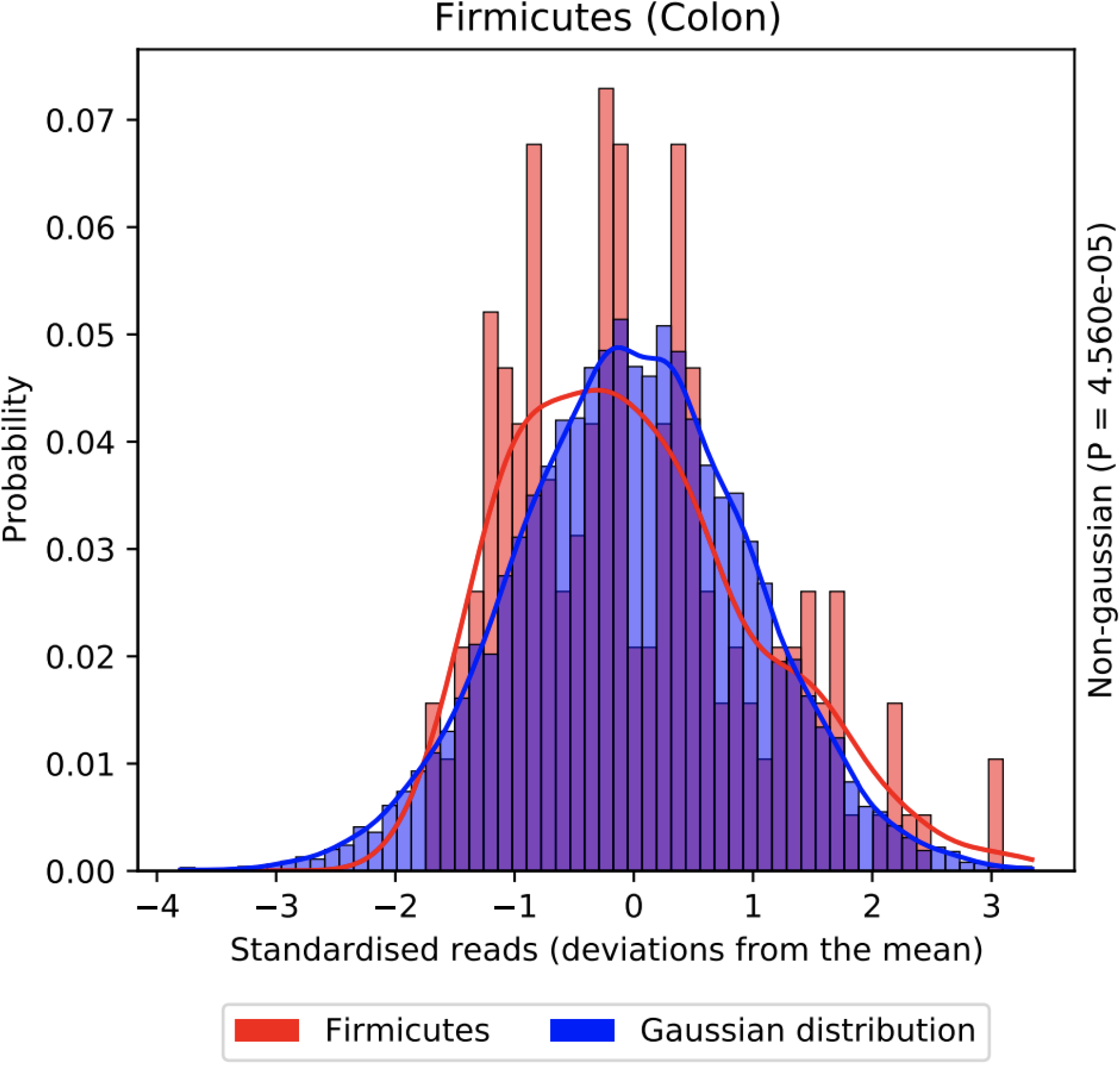
Comparison of colon isolated Firmicute distribution to a Gaussian distribution.

**Figure 4:**
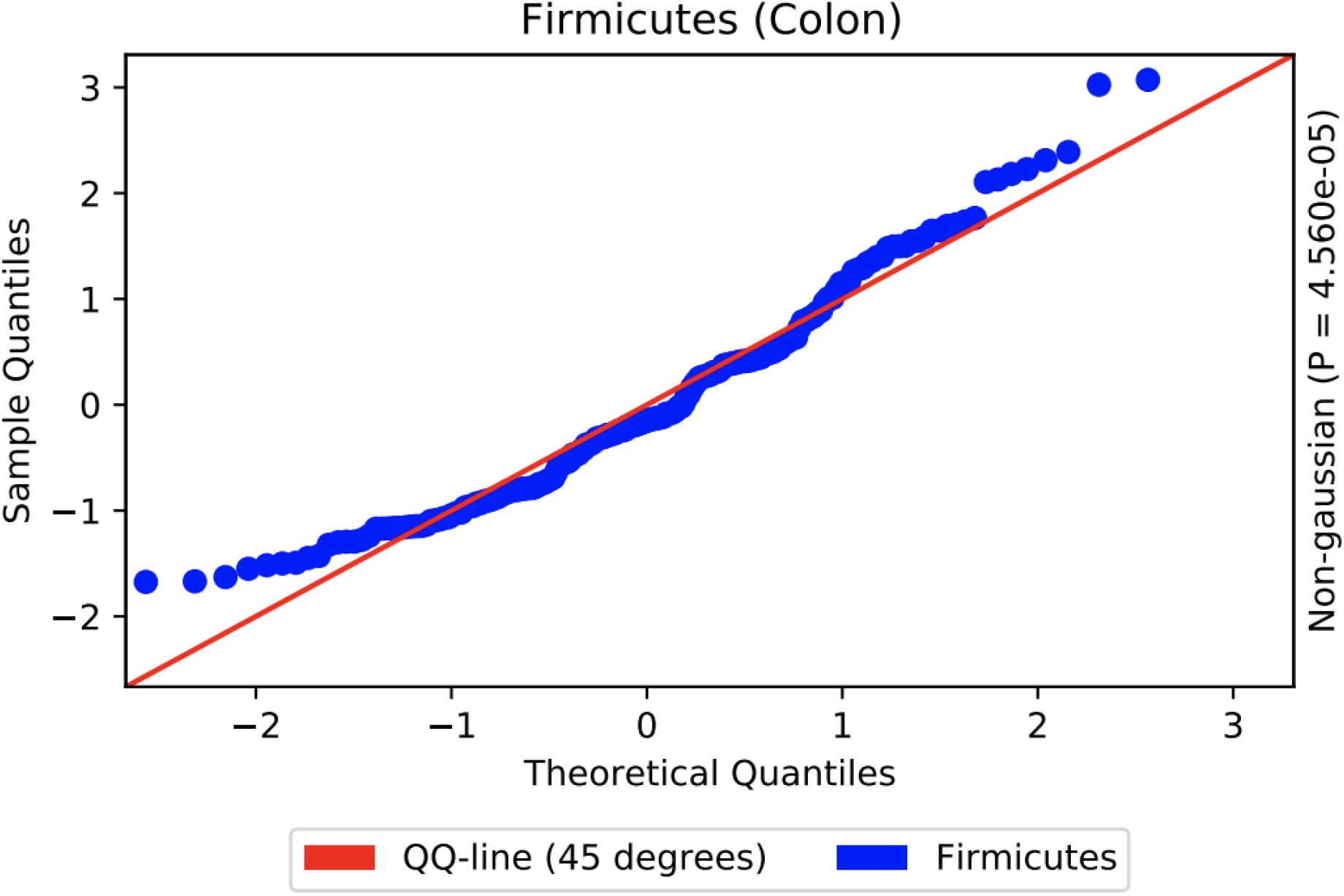
Comparison of colon isolated Firmicute distribution to a Gaussian distribution (QQ-plot)

### Application to real-world data

The 16S V3-V5 region dataset of Healthy Human Microbiome project (Huttenhower *et al*., 2012) was downloaded from MicrobiomeDB (Oliveira *et al*., 2018) and was processed through statSuma using default settings (SI Tables 1-4; SI Figures 1-3). A result was considered significant if *P*_BD_ ≤ 0.005 (SI Tables 3-4).

## Results

### Performance

Using Google Chrome *v*. 90.0.4430.212 (Official Build) (x86_64), statSuma was able to process the Healthy Human Microbiome dataset in 11 minutes and 15 seconds where pairwise analyses were completed in 6 seconds, listwise in 1 second, optional variance plots in 8 minutes and 45 seconds, optional Gaussian plots in 2 minutes and 0 seconds, and optional QQ-plots in 0 minutes and 23 seconds. If plots are not required, this process takes 7 seconds to complete.

### Recommended results

Of the 1,266 pairwise tests performed, there were no situations where a Gaussian distribution was observed in both datasets, and equivariance was observed in 118 cases. As such, Brunner-Munzel tests were advised in 873 cases (68.957%), Mann-Whitney U-tests were advised in 376 cases (30.49%) of cases, Welch’s *t*-test in 5 cases (0.395%), and Student’s *t*-test in 2 cases (0.158%). Using the test ranking criteria specified above, the recommended test for all comparisons is the Brunner-Munzel test (SI Table 1).

Of the 12 listwise tests performed, while all were observed to be equivariant (*P* ≥ 0.05) when observing a single taxon across all anatomical sites none were observed to contain ubiquitous Gaussian distributions, therefore the Kruskal-Wallis H-test was advised in all cases (SI Table 2).

## Discussion

This software is designed to guide researchers in choosing the most appropriate statistical comparators in microbiome studies and to provide access to some statistical tests that are not yet commonly available on professional statistics analysis package (such as the Brunner-Munzel test). Due to the variability and cyclical shifts associated with microbiomes (Falony *et al*., 2016; Leigh, Murphy and Walsh, 2021), many pairwise tests requiring equivariance (Student’s *t*-test and Mann-Whitney *U*-test) or pairwise tests requiring ubiquitous Gaussian distributions (Student’s *t*-test and Welch’s *t*-test) were not expected. In line with these expectations, we observed 376 of 1,266 (30.49%) of tests to be equivariant and none to be ubiquitously Gaussian. Strikingly, *t*-tests were only deemed appropriate for 7 comparisons in the entire dataset using our criteria. For listwise comparisons, ubiquitous Gaussian distributions and equivariance were also expected to be rare. In practice, equivariance and ubiquitous Gaussian distribution were not observed in any instance.

We advise against statSuma to be used blindly by researchers without ensuring that the provided tests are appropriate for the hypotheses they intend to explore. It is provided as a helpful guide in choosing and performing statistical analyses with the aim of reducing time spent on these tasks for researchers.

## Supporting information

SI Figure 1 (Variance comparisons)

SI Figure 2 (Gaussian comparisons)

SI Figure 3 (QQ plots)

## Code availability

The software and dataset used for this publication are fully available at https://github.com/RobLeighBioinformatics/statSuma

## Competing interests

This project was funded by Alltech. RL received funding in the form of a postdoctoral research fellowship from Alltech and RM received a salary from Alltech during this project.

## Notes

https://github.com/RobLeighBioinformatics/statSuma/

