## Supplementary material for "statSuma: automated selection and performance of statistical comparisons for microbiome studies": SI Figure 1 (Variance comparisons)

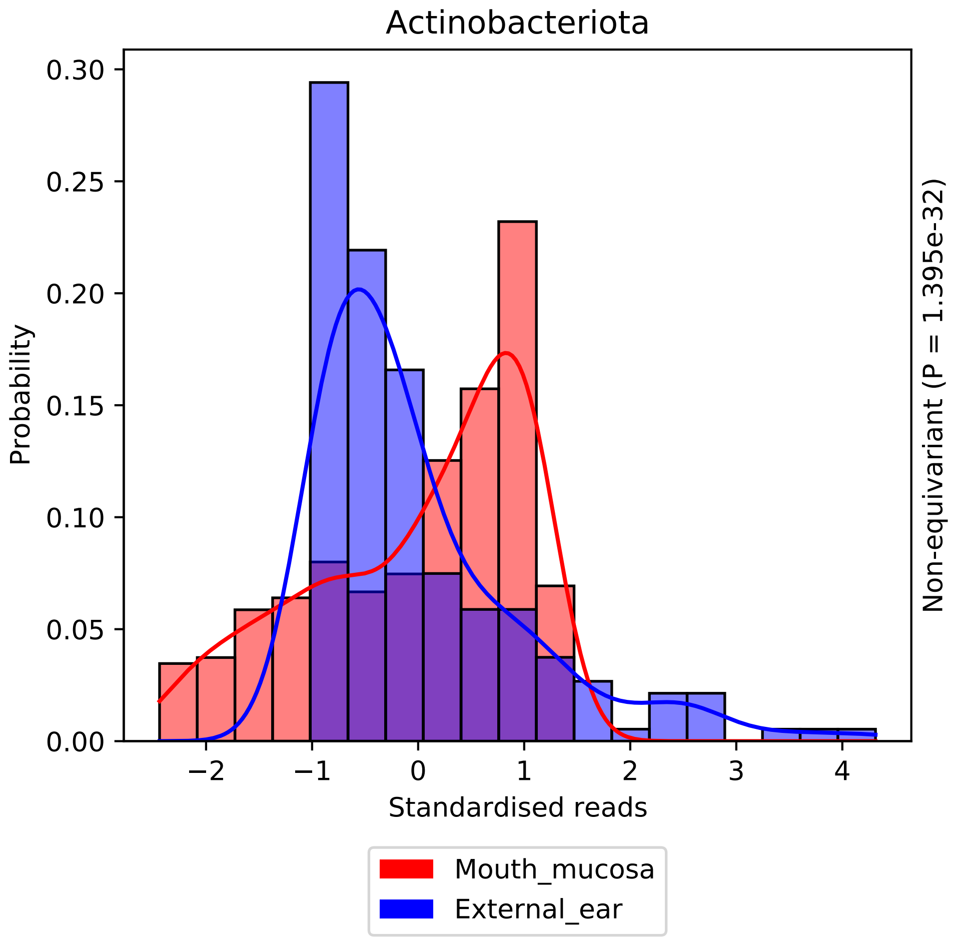
Actinobacteriota
Probability
0.30
0.25
0.20
0.15
0.10
0.05
0.00
−2
−1
0
1
2
3
4
Standardised reads
Non-equivariant (P = 1.395e-32)
Mouth_mucosa
External_ear

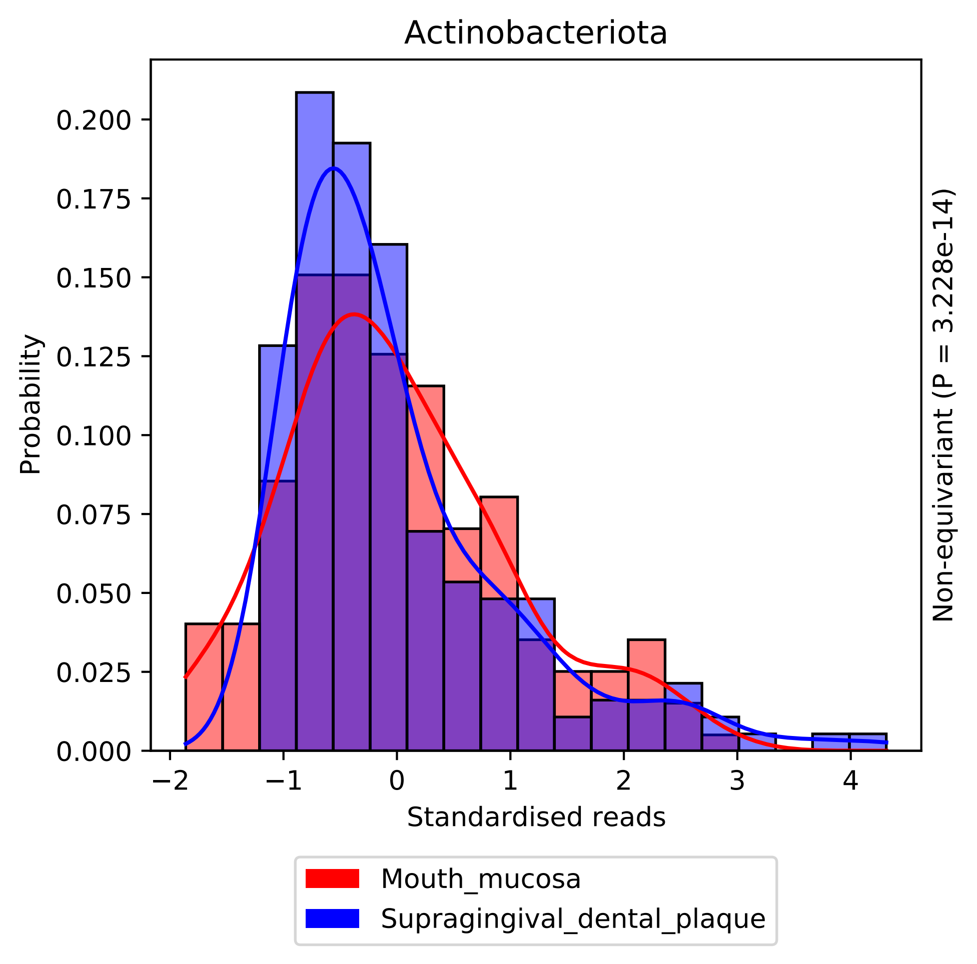
Actinobacteriota
Probability
0.000
0.025
0.050
0.075
0.100
0.125
0.150
0.175
0.200
−2
−1
0
1
2
3
4
Standardised reads
Non-equivariant (P = 3.228e-14)
Mouth_mucosa
Supragingival_dental_plaque

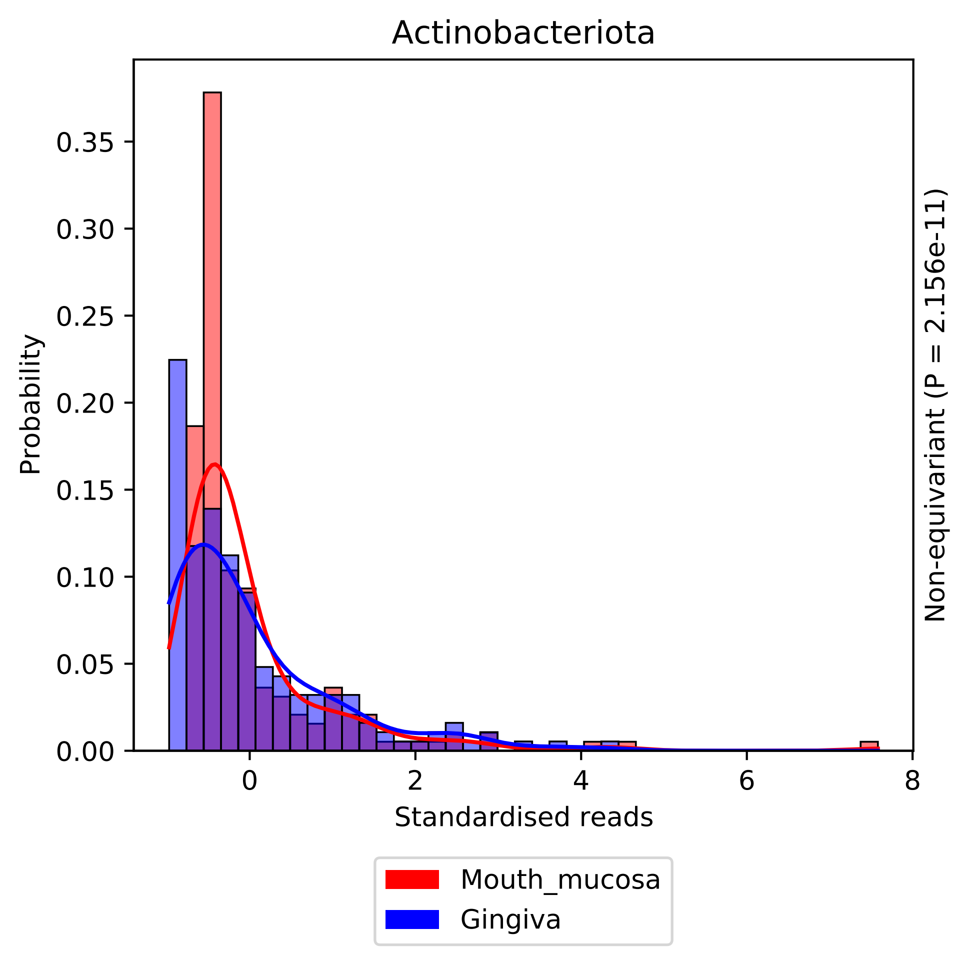

Actinobacteriota
Probability
0.00
0.05
0.10
0.15
0.20
0.25
0.30
0.35
0
2
4
6
8
Standardised reads
Non-equivariant (P = 2.156e-11)
Mouth_mucosa
Gingiva

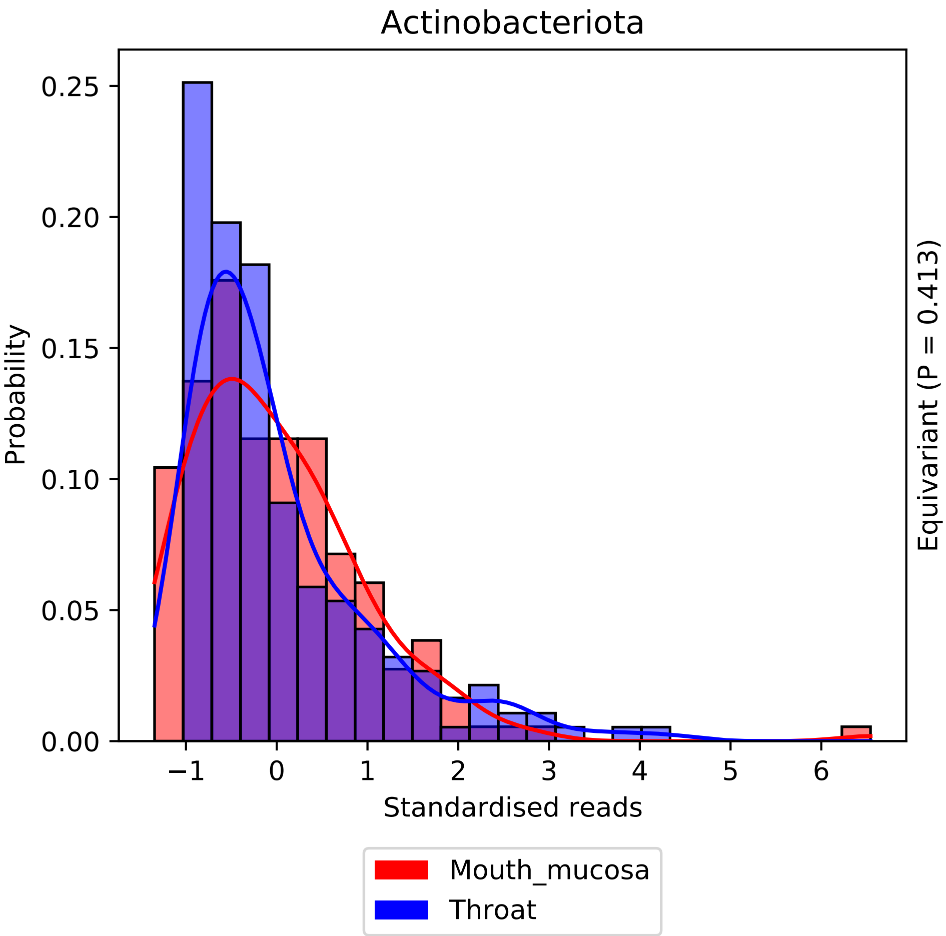

Actinobacteriota
Probability
Standardised reads
Equivariant (P = 0.413)
0.00
0.05
0.10
0.15
0.20
0.25
−1
0
1
2
3
4
5
6
Mouth_mucosa
Throat

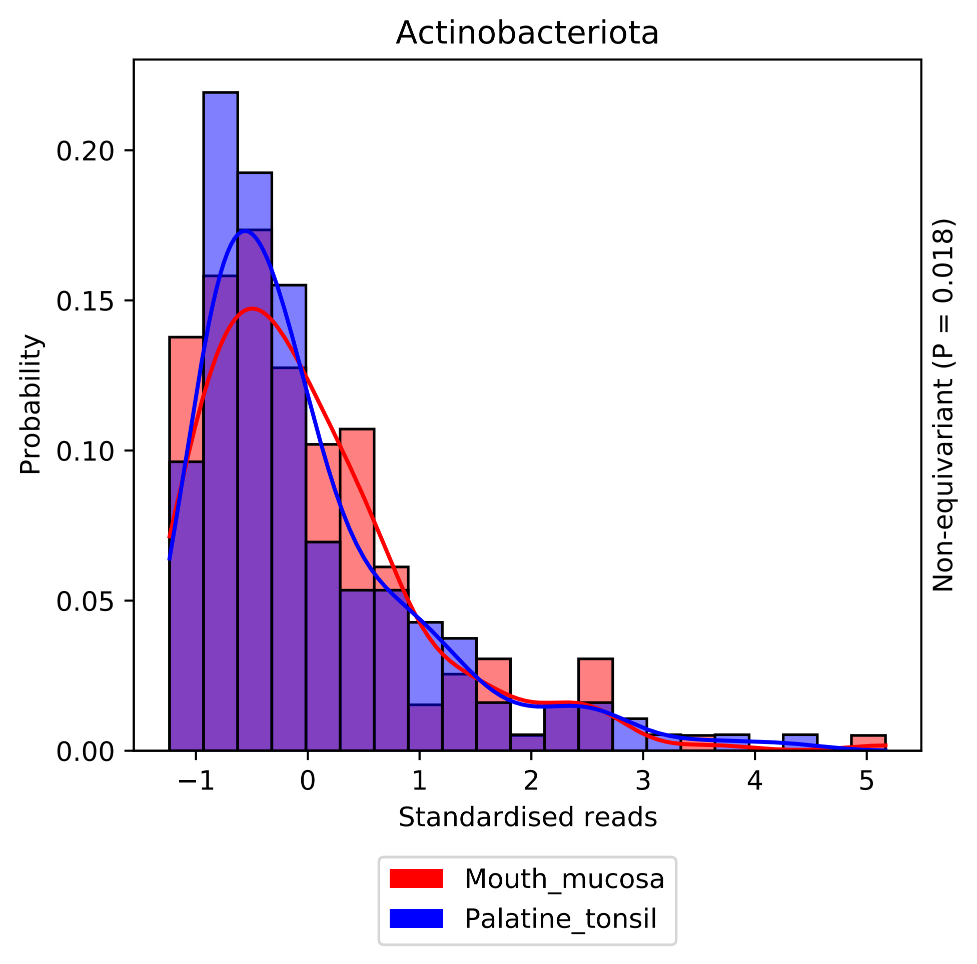

Actinobacteriota
Probability
0.00
0.05
0.10
0.15
0.20
−1
0
1
2
3
4
5
Standardised reads
Non-equivariant (P = 0.018)
Mouth_mucosa
Palatine_tonsil

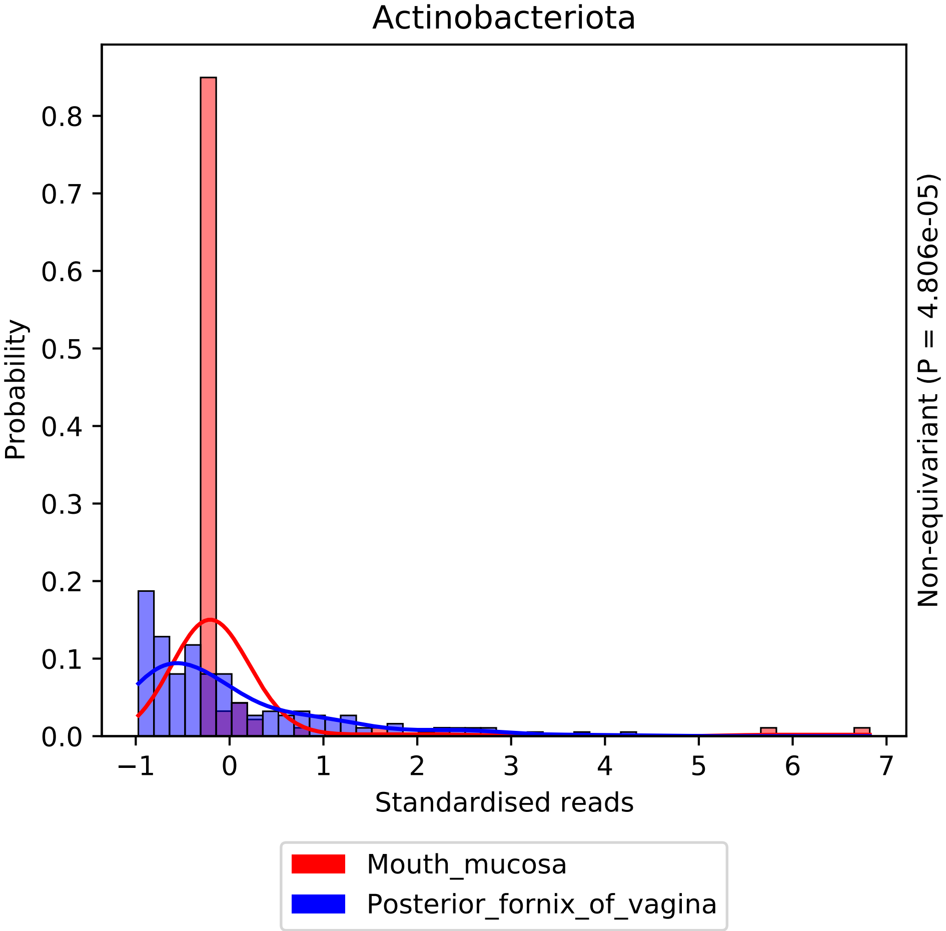

Actinobacteriota
Probability
0.0
0.1
0.2
0.3
0.4
0.5
0.6
0.7
0.8
−1
0
1
2
3
4
5
6
7
Standardised reads
Non-equivariant (P = 4.806e-05)
Mouth_mucosa
Posterior_fornix_of_vagina

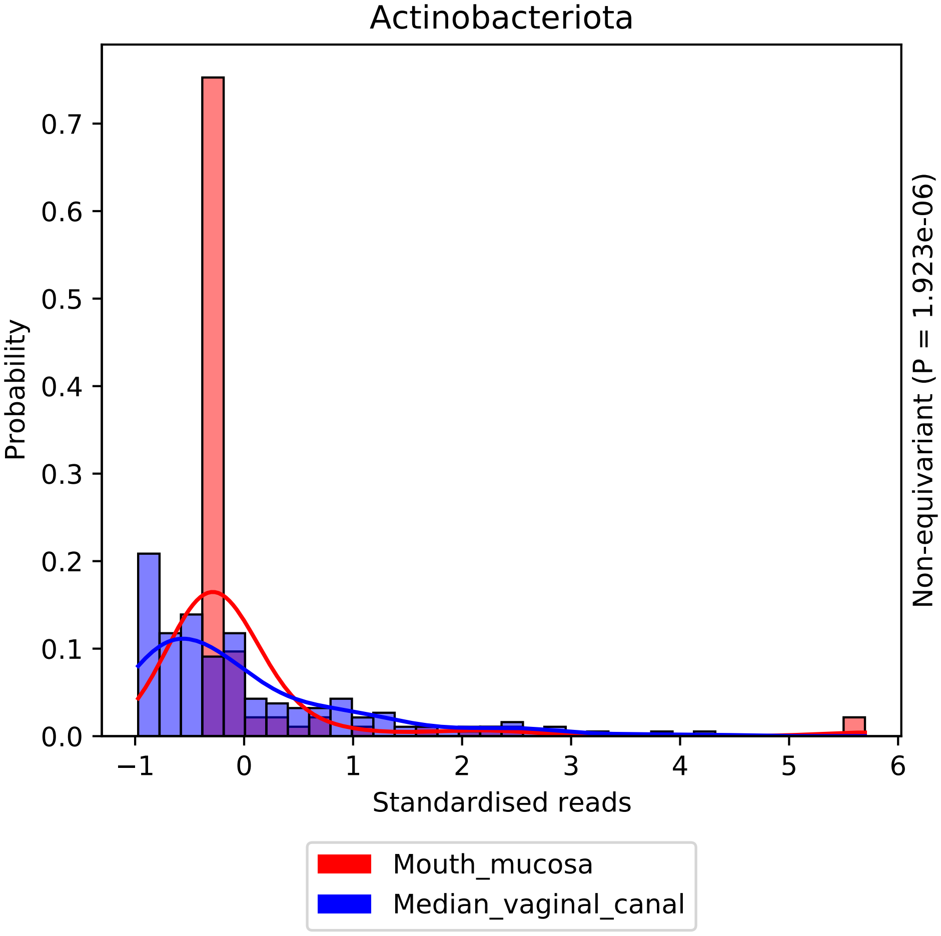

Actinobacteriota
Probability
Standardised reads
Non-equivariant (P = 1.923e-06)
0.0
0.1
0.2
0.3
0.4
0.5
0.6
0.7
−1
0
1
2
3
4
5
6
Mouth_mucosa
Median_vaginal_canal

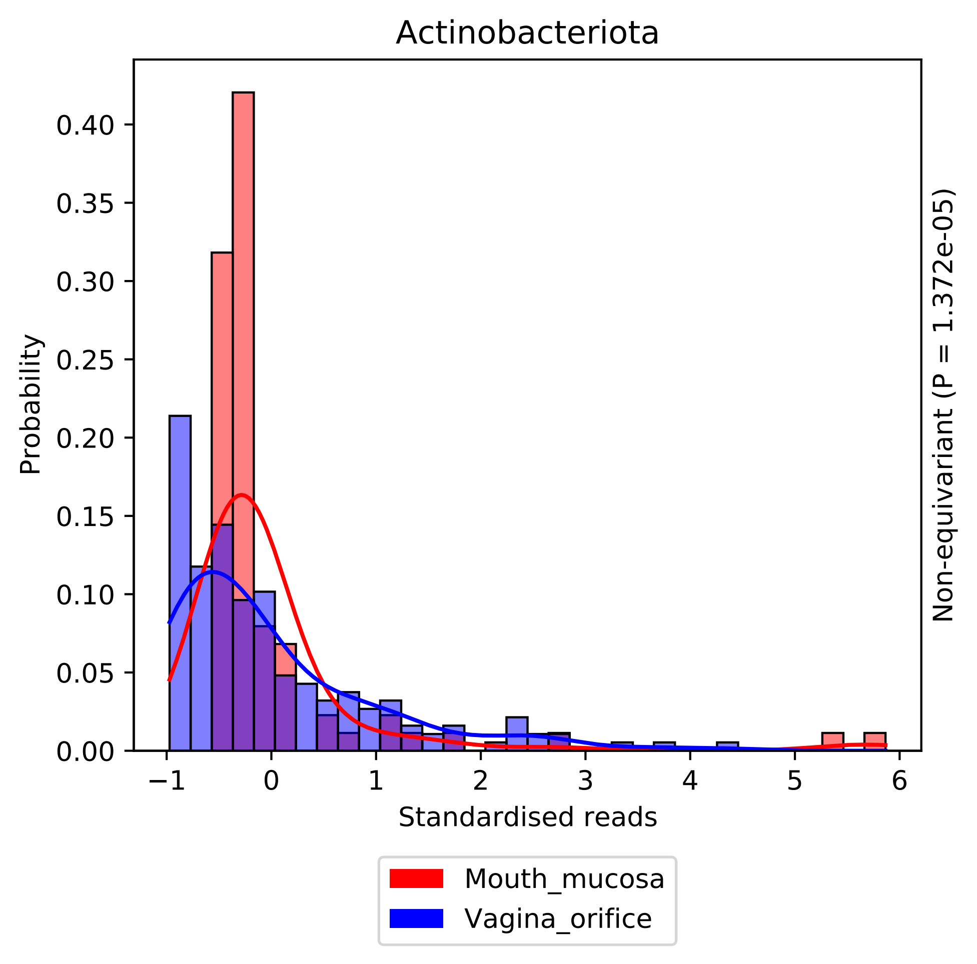

Actinobacteriota
Probability
0.00
0.05
0.10
0.15
0.20
0.25
0.30
0.35
0.40
−1
0
1
2
3
4
5
6
Standardised reads
Non-equivariant (P = 1.372e-05)
Mouth_mucosa
Vagina_orifice

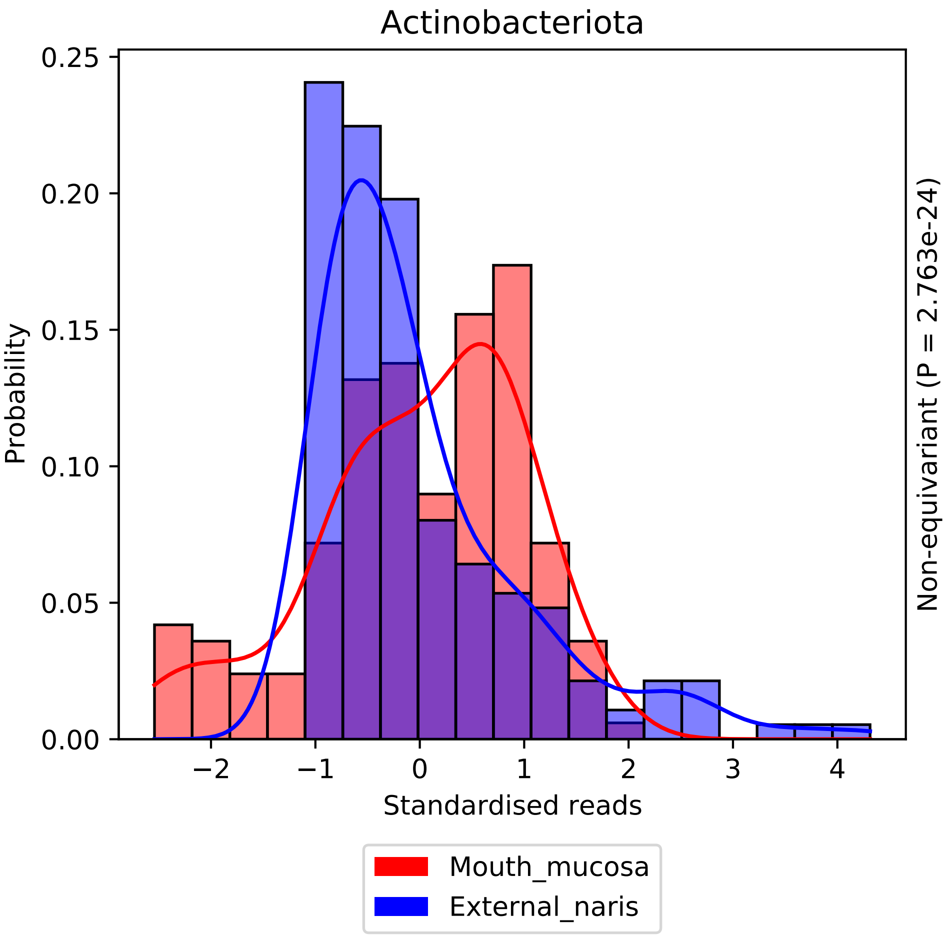
Actinobacteriota
Probability
0.25
0.20
0.15
0.10
0.05
0.00
−2
−1
0
1
2
3
4
Standardised reads
Non-equivariant (P = 2.763e-24)
Mouth_mucosa
External_naris

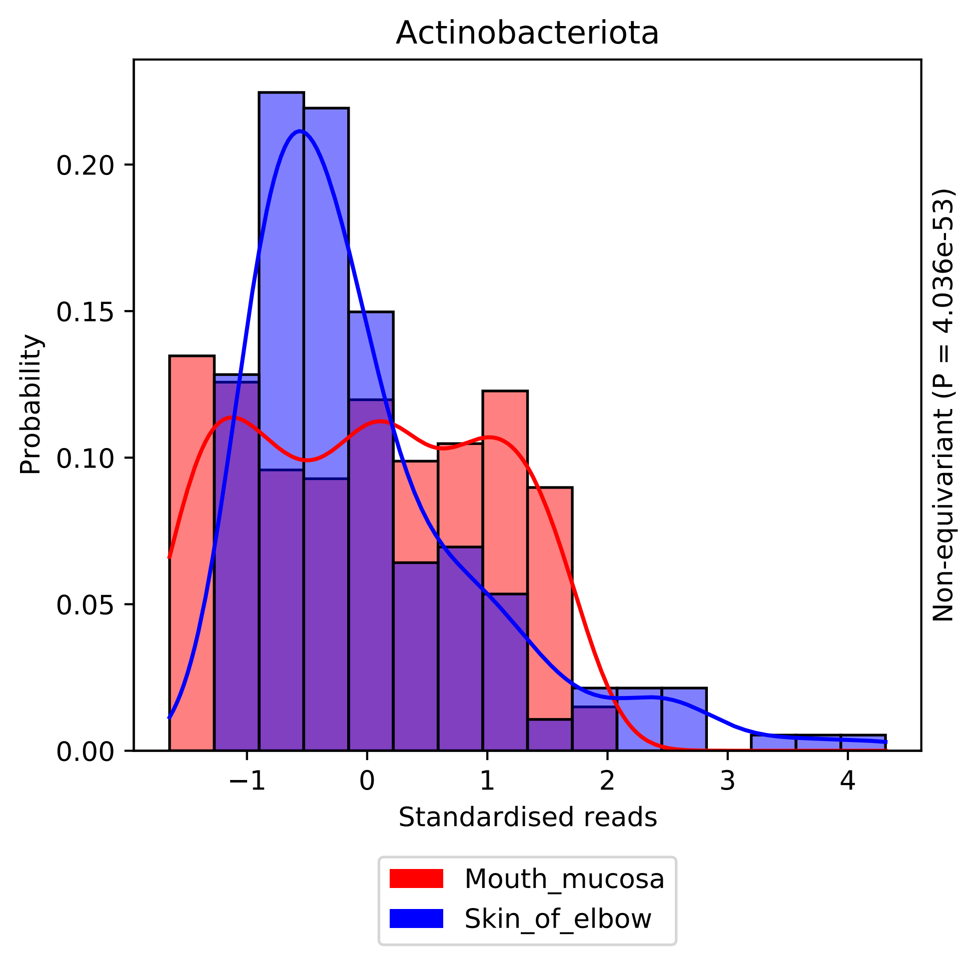
Actinobacteriota
Probability
0.00
0.05
0.10
0.15
0.20
−1
0
1
2
3
4
Standardised reads
Non-equivariant (P = 4.036e-53)
Mouth_mucosa
Skin_of_elbow

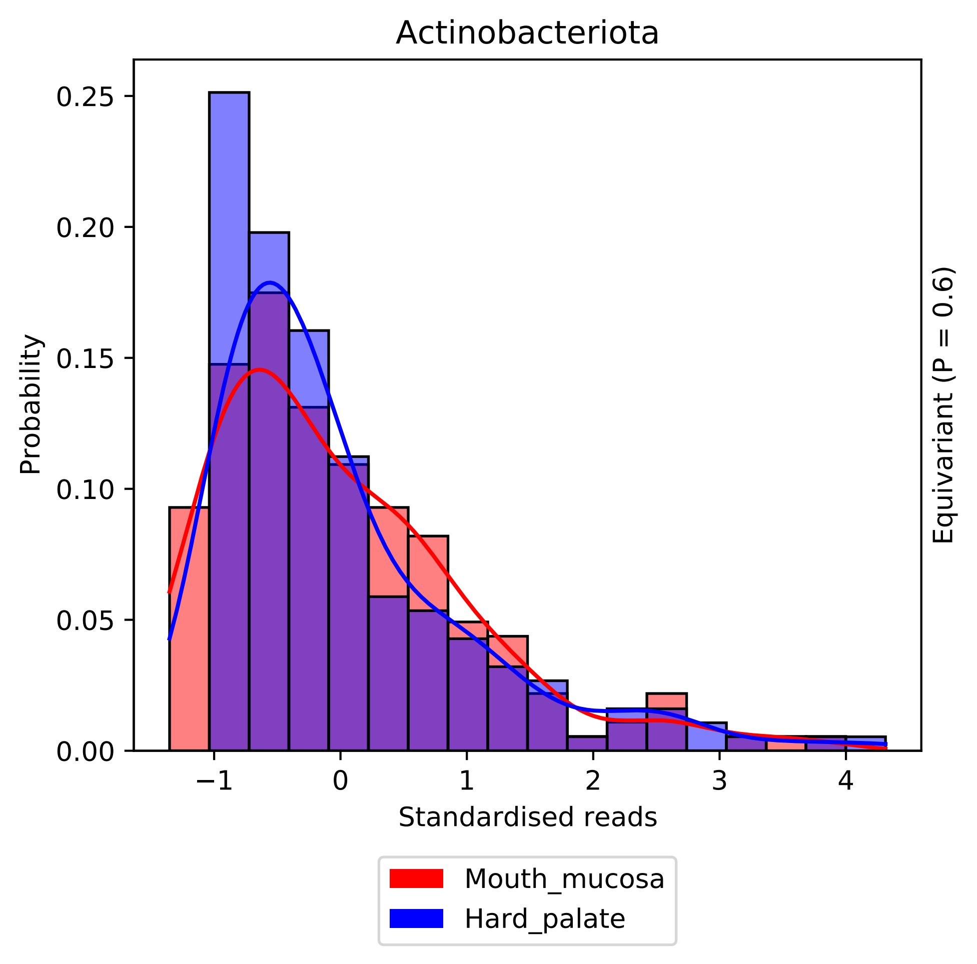

Actinobacteriota
Probability
0.25
0.20
0.15
0.10
0.05
0.00
−1
0
1
2
3
4
Standardised reads
Equivariant (P = 0.6)
Mouth_mucosa
Hard_palate

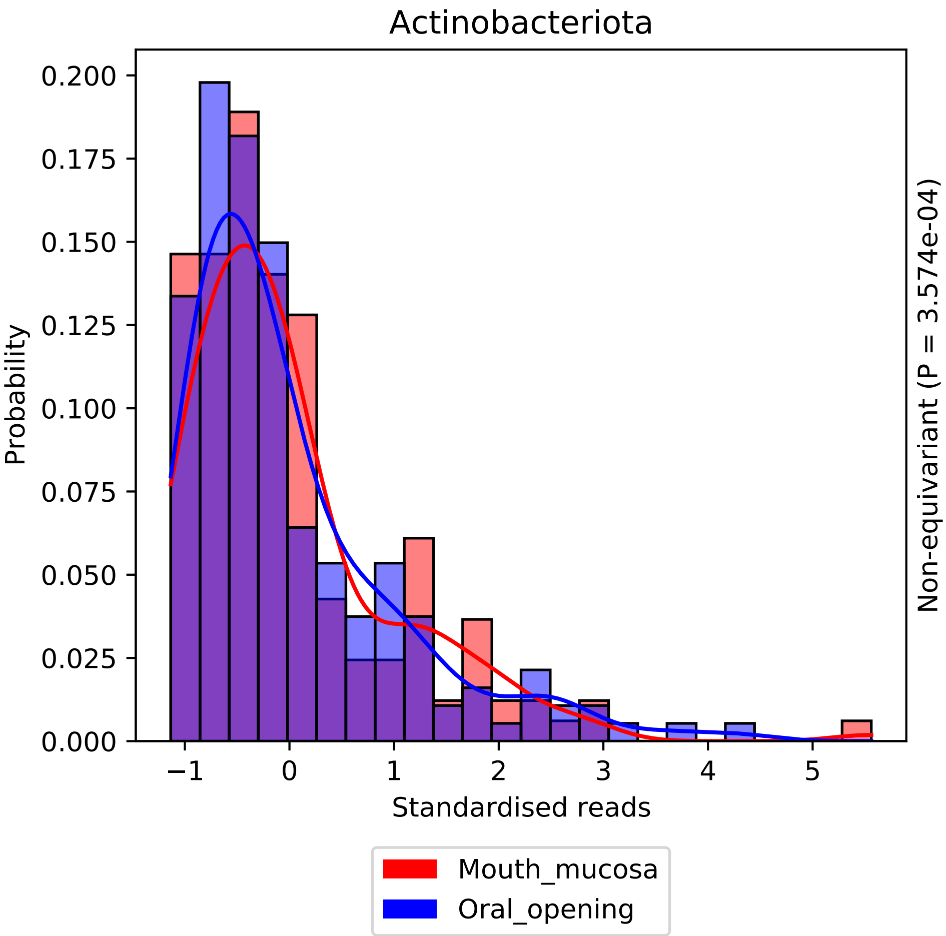

Actinobacteriota
Probability
0.000
0.025
0.050
0.075
0.100
0.125
0.150
0.175
0.200
−1
0
1
2
3
4
5
Standardised reads
Non-equivariant (P = 3.574e-04)
Mouth_mucosa
Oral_opening

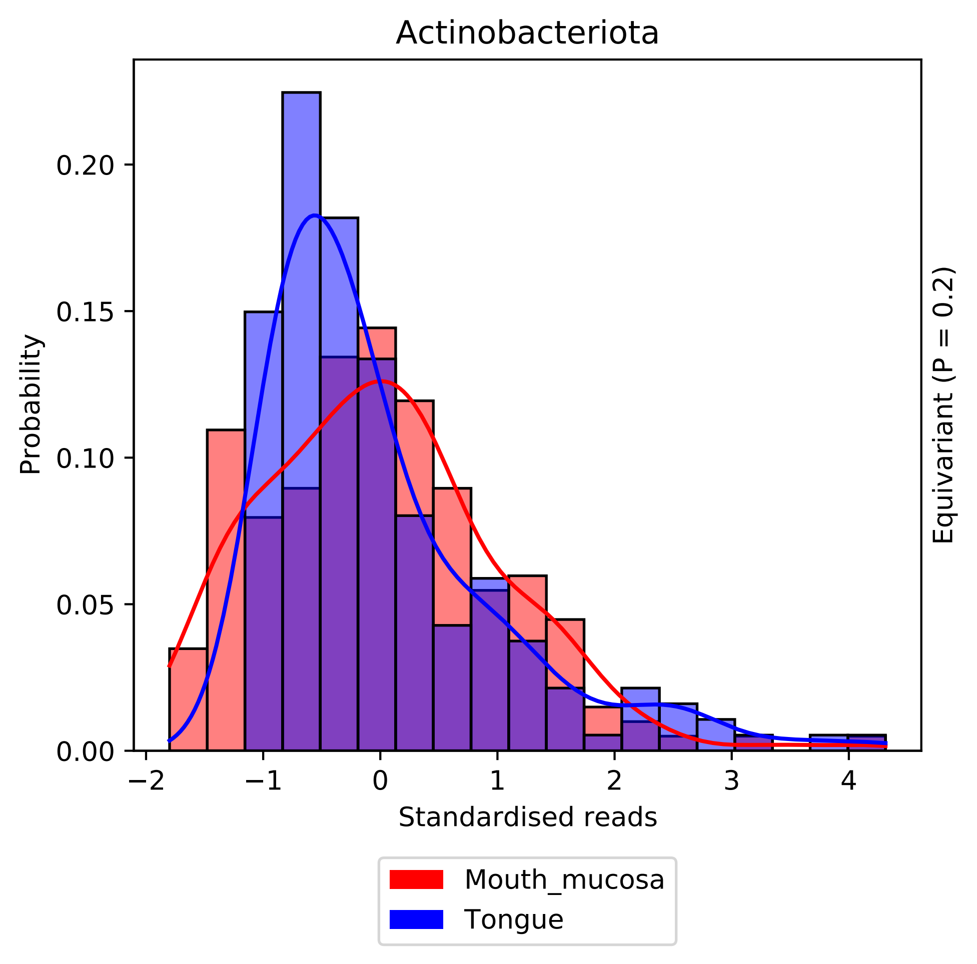

Actinobacteriota
Probability
0.00
0.05
0.10
0.15
0.20
−2
−1
0
1
2
3
4
Standardised reads
Equivariant (P = 0.2)
Mouth_mucosa
Tongue

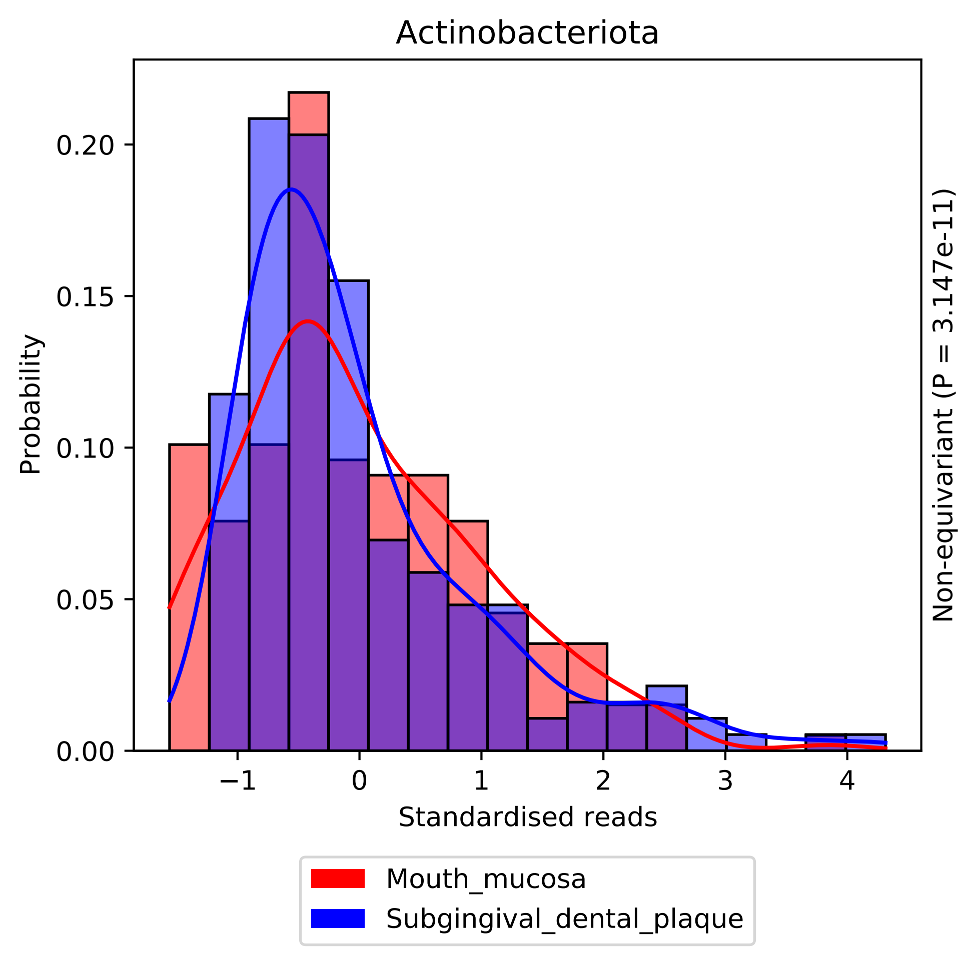

Actinobacteriota
Probability
Standardised reads
Non-equivariant (P = 3.147e-11)
0.00
0.05
0.10
0.15
0.20
−1
0
1
2
3
4
Mouth_mucosa
Subgingival_dental_plaque

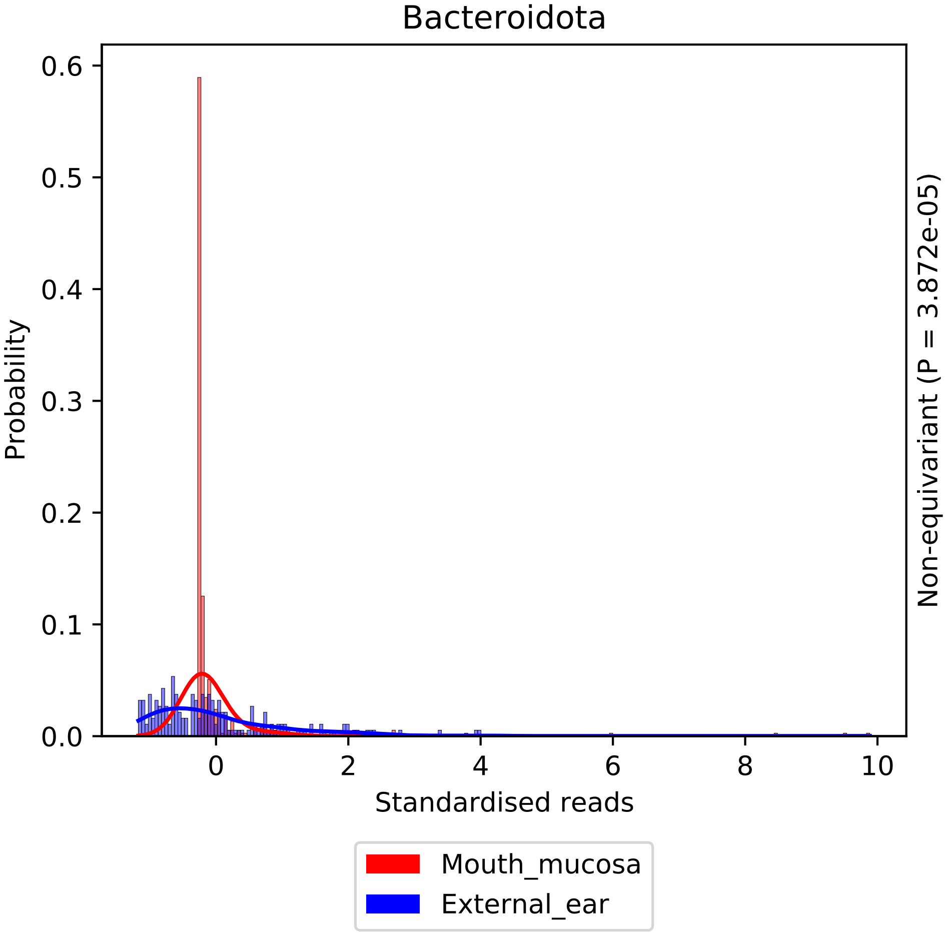

Bacteroidota
Probability
0.0
0.1
0.2
0.3
0.4
0.5
0.6
0
2
4
6
8
10
Standardised reads
Non-equivariant (P = 3.872e-05)
Mouth_mucosa
External_ear

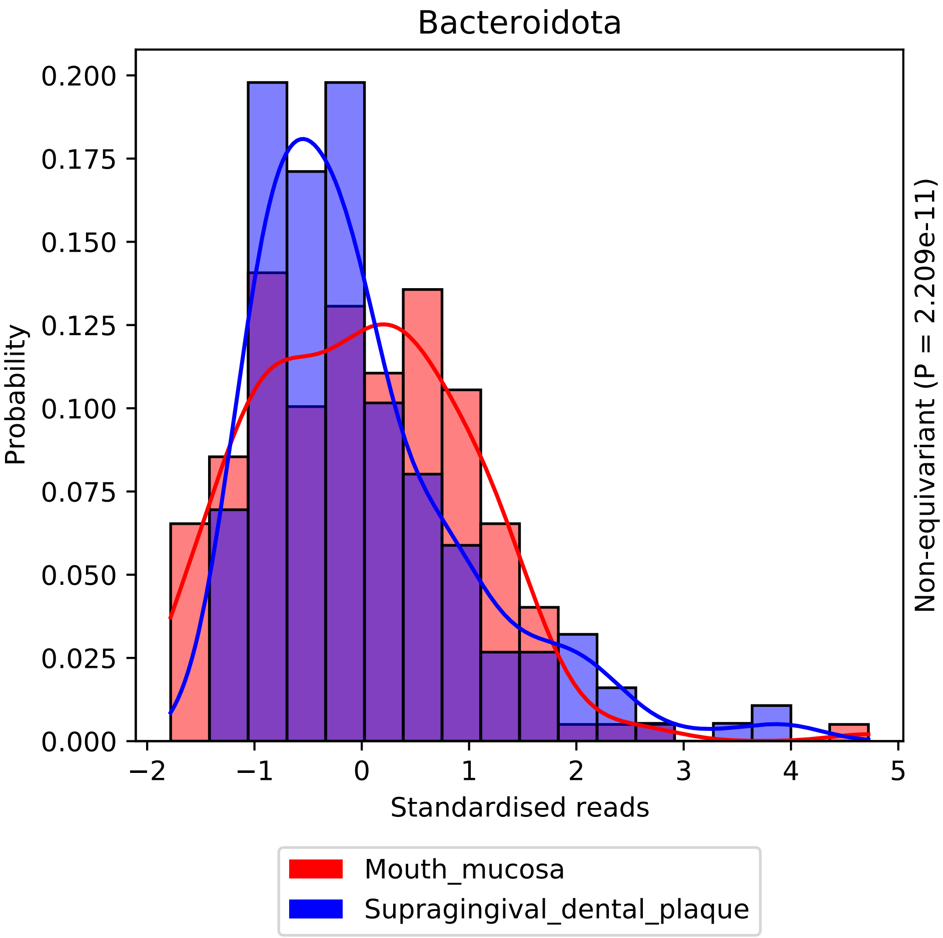
Bacteroidota
Probability
0.200
0.175
0.150
0.125
0.100
0.075
0.050
0.025
0.000
−2
−1
0
1
2
3
4
5
Standardised reads
Non-equivariant (P = 2.209e-11)
Mouth_mucosa
Supragingival_dental_plaque

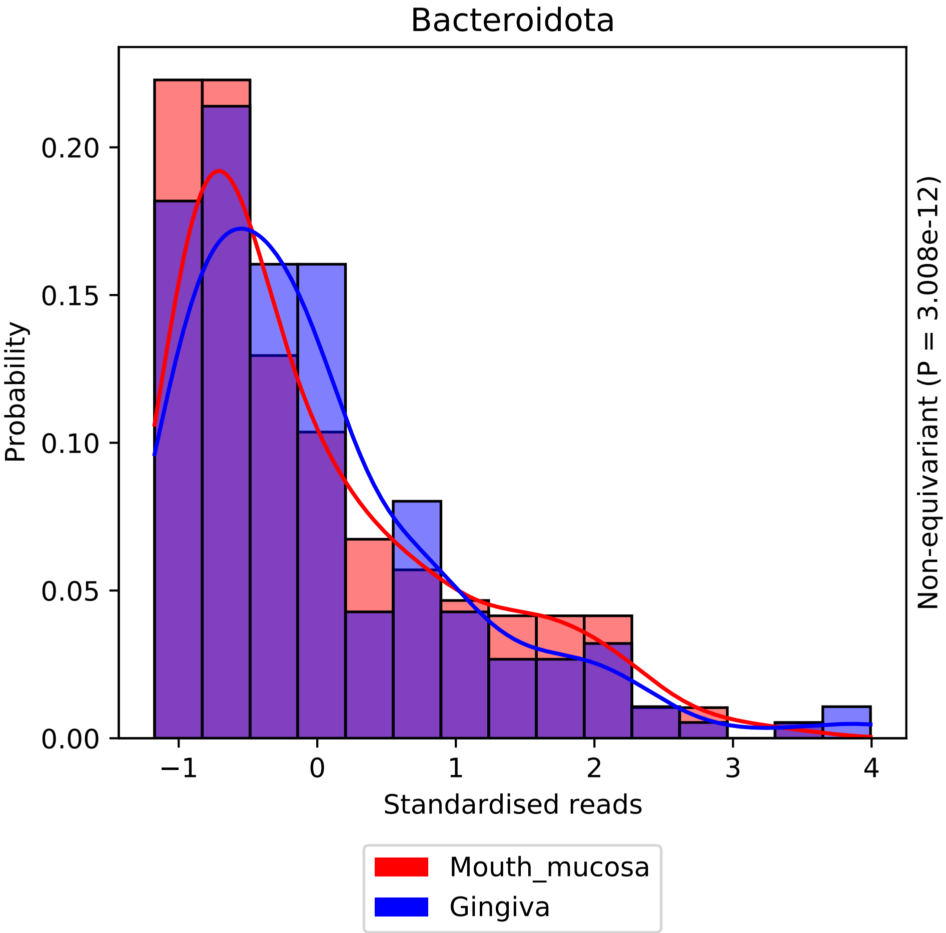

Bacteroidota
Probability
0.00
0.05
0.10
0.15
0.20
−1
0
1
2
3
4
Standardised reads
Non-equivariant (P = 3.008e-12)
Mouth_mucosa
Gingiva

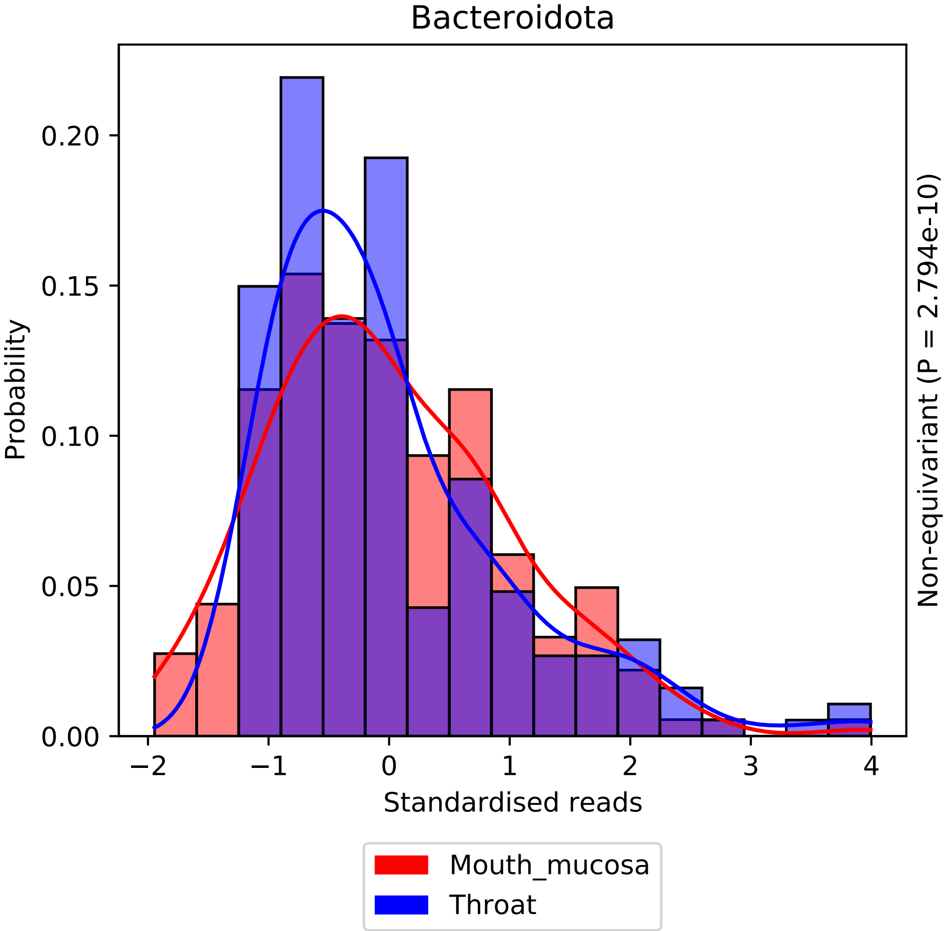

Bacteroidota
Probability
0.00
0.05
0.10
0.15
0.20
−2
−1
0
1
2
3
4
Standardised reads
Non-equivariant (P = 2.794e-10)
Mouth_mucosa
Throat

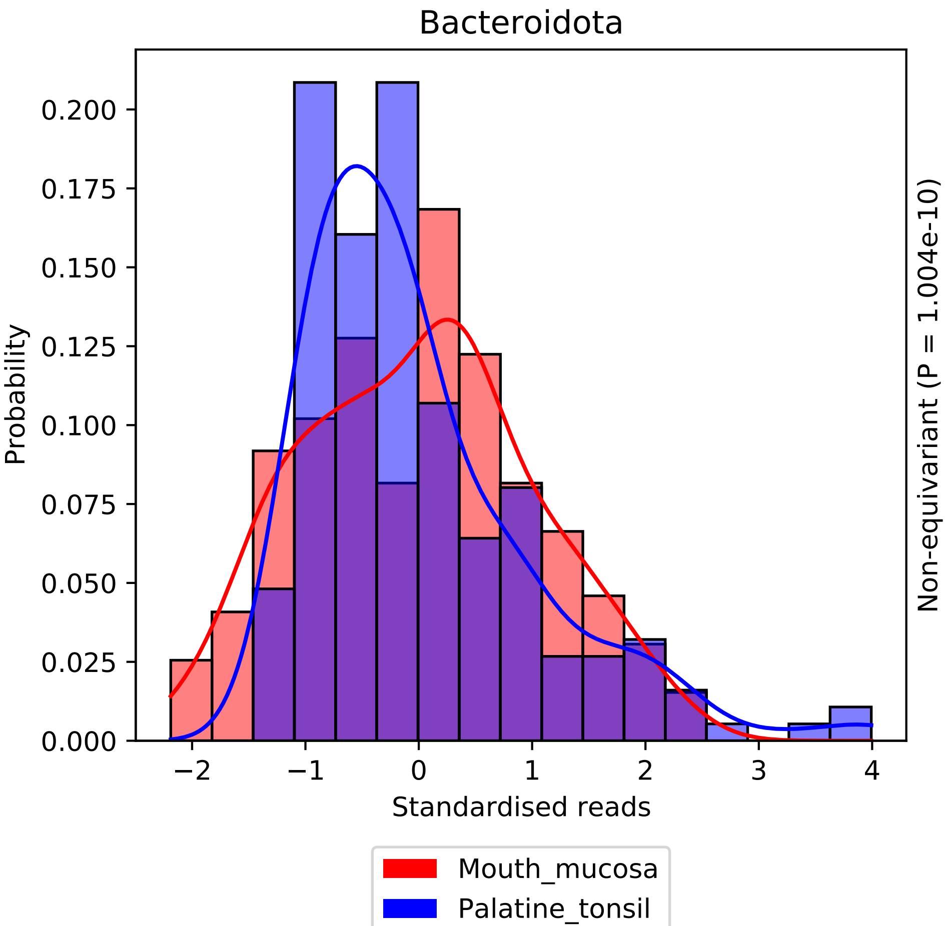
Bacteroidota
Probability
0.200
0.175
0.150
0.125
0.100
0.075
0.050
0.025
0.000
−2
−1
0
1
2
3
4
Standardised reads
Non-equivariant (P = 1.004e-10)
Mouth_mucosa
Palatine_tonsil

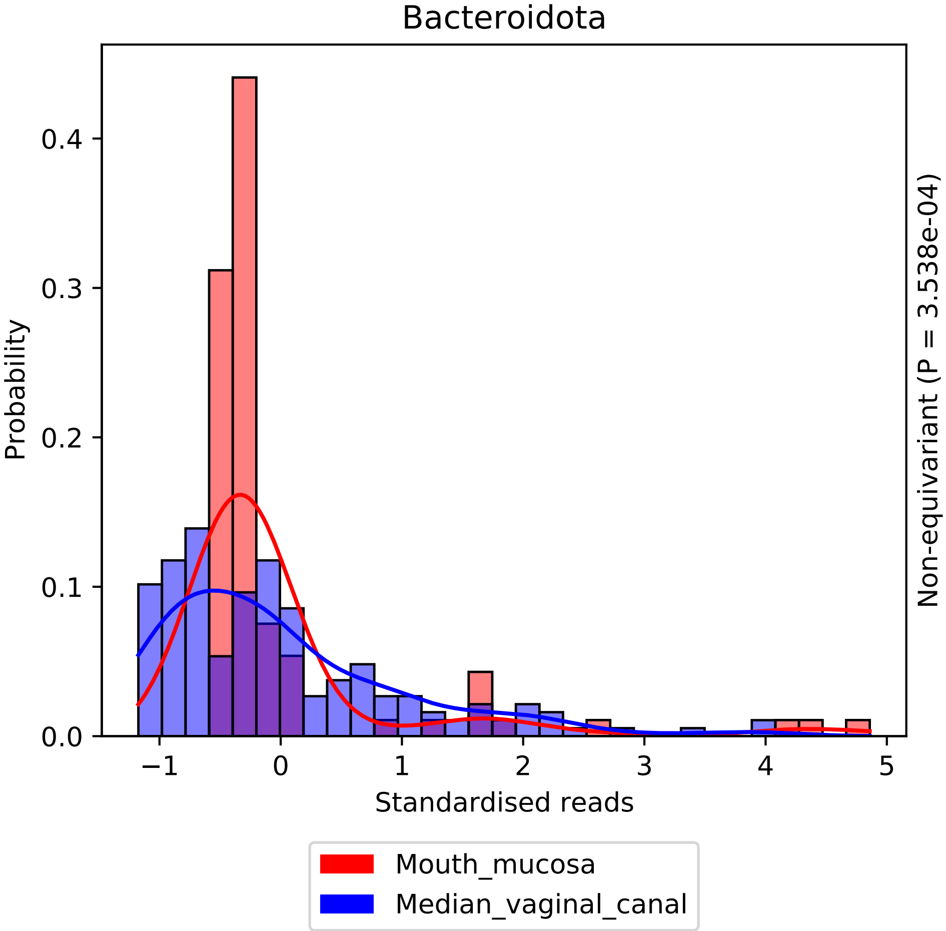
Bacteroidota
Probability
0.0
0.1
0.2
0.3
0.4
−1
0
1
2
3
4
5
Standardised reads
Non-equivariant (P = 3.538e-04)
Mouth_mucosa
Median_vaginal_canal

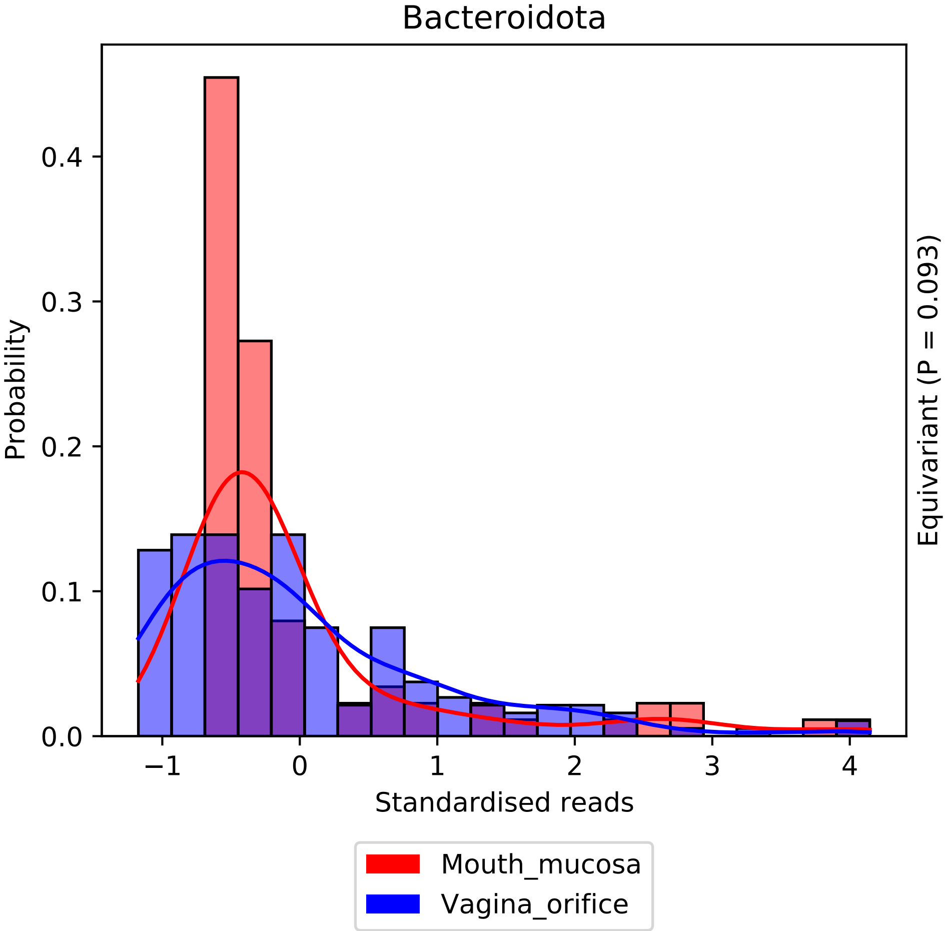

Bacteroidota
Probability
0.4
0.3
0.2
0.1
0.0
−1
0
1
2
3
4
Standardised reads
Equivariant (P = 0.093)
Mouth_mucosa
Vagina_orifice

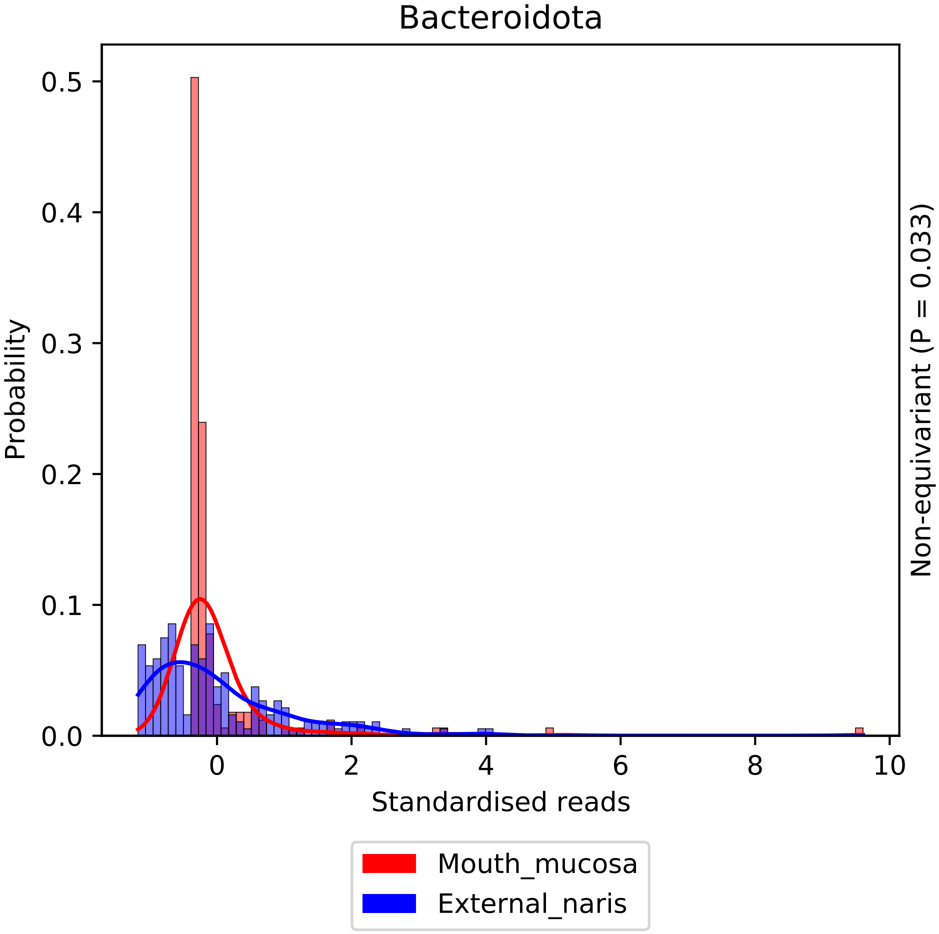
Bacteroidota
Probability
0.0
0.1
0.2
0.3
0.4
0.5
0
2
4
6
8
10
Standardised reads
Non-equivariant (P = 0.033)
Mouth_mucosa
External_naris

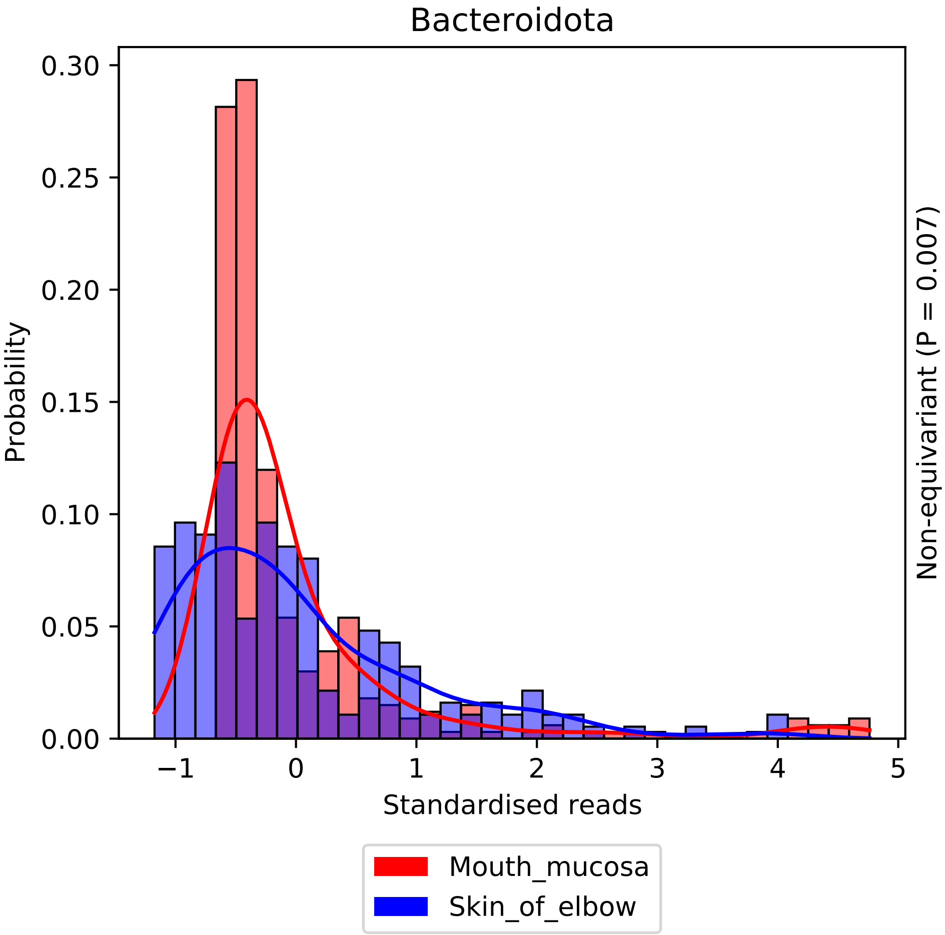
Bacteroidota
Probability
0.00
0.05
0.10
0.15
0.20
0.25
0.30
−1
0
1
2
3
4
5
Standardised reads
Non-equivariant (P = 0.007)
Mouth_mucosa
Skin_of_elbow

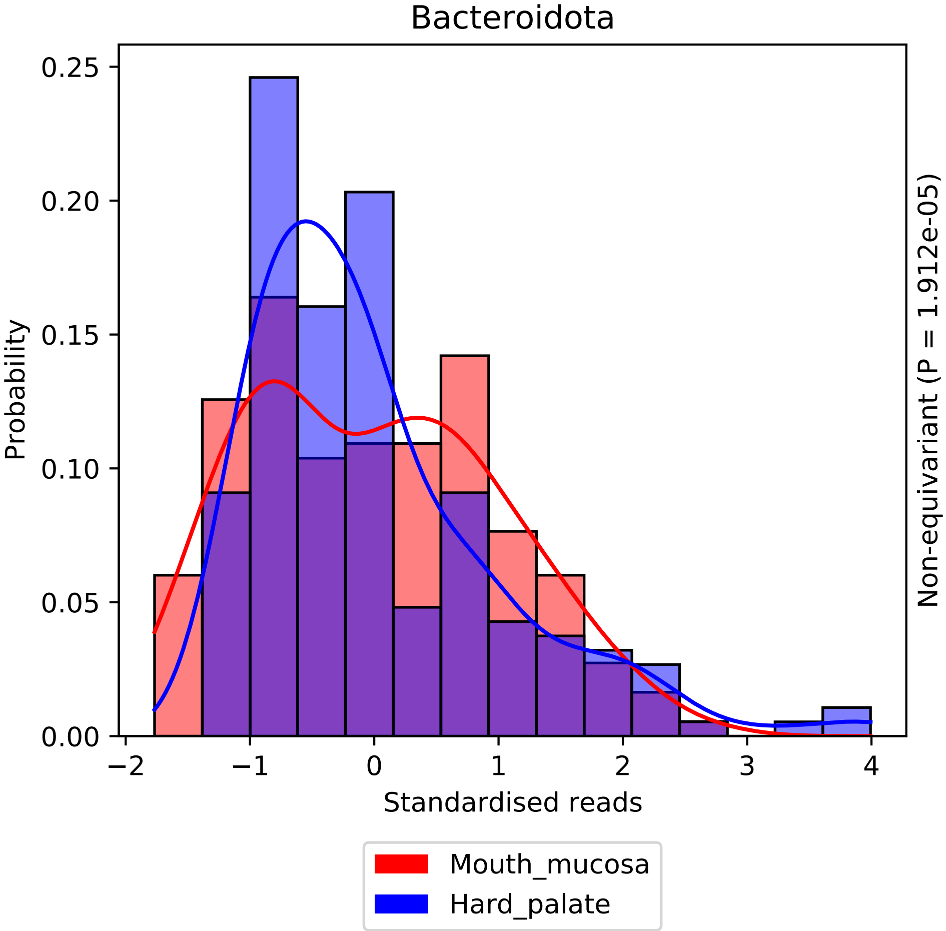

Bacteroidota
Probability
0.25
0.20
0.15
0.10
0.05
0.00
−2
−1
0
1
2
3
4
Standardised reads
Non-equivariant (P = 1.912e-05)
Mouth_mucosa
Hard_palate

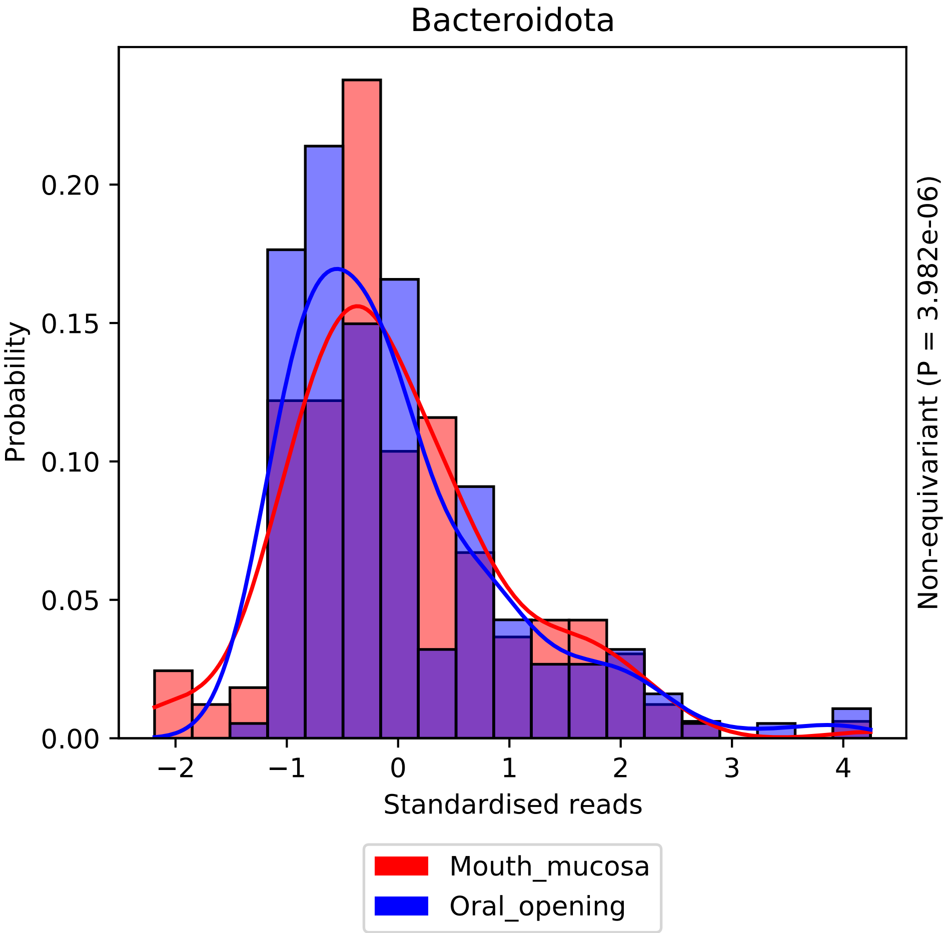

Bacteroidota
Probability
0.00
0.05
0.10
0.15
0.20
−2
−1
0
1
2
3
4
Standardised reads
Non-equivariant (P = 3.982e-06)
Mouth_mucosa
Oral_opening

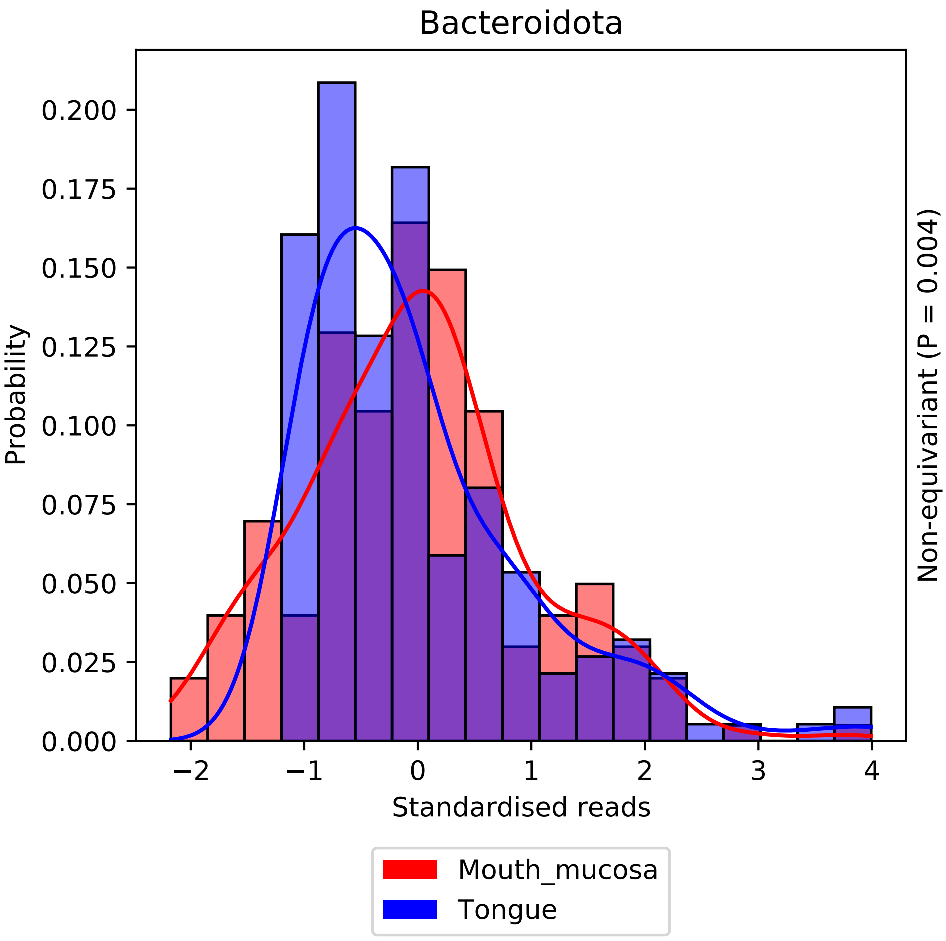

Bacteroidota
Probability
0.200
0.175
0.150
0.125
0.100
0.075
0.050
0.025
0.000
−2
−1
0
1
2
3
4
Standardised reads
Non-equivariant (P = 0.004)
Mouth_mucosa
Tongue

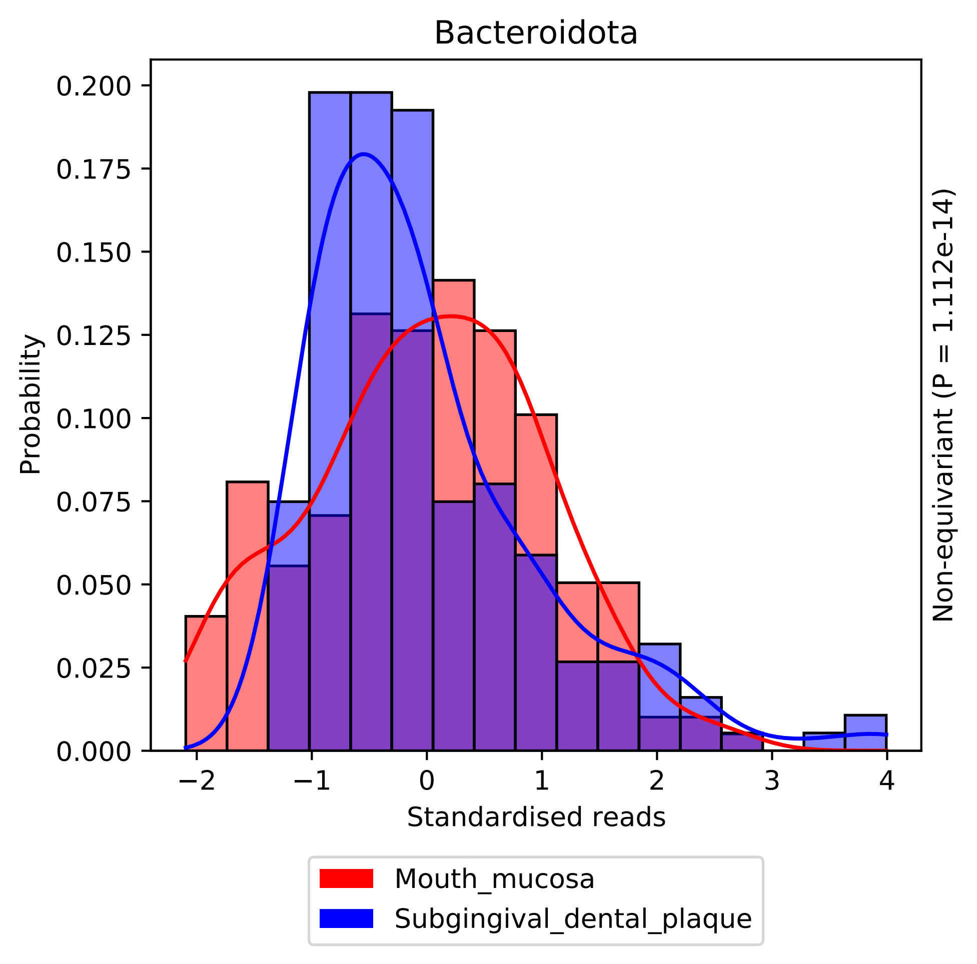

Bacteroidota
Probability
0.200
0.175
0.150
0.125
0.100
0.075
0.050
0.025
0.000
−2
−1
0
1
2
3
4
Standardised reads
Non-equivariant (P = 1.112e-14)
Mouth_mucosa
Subgingival_dental_plaque

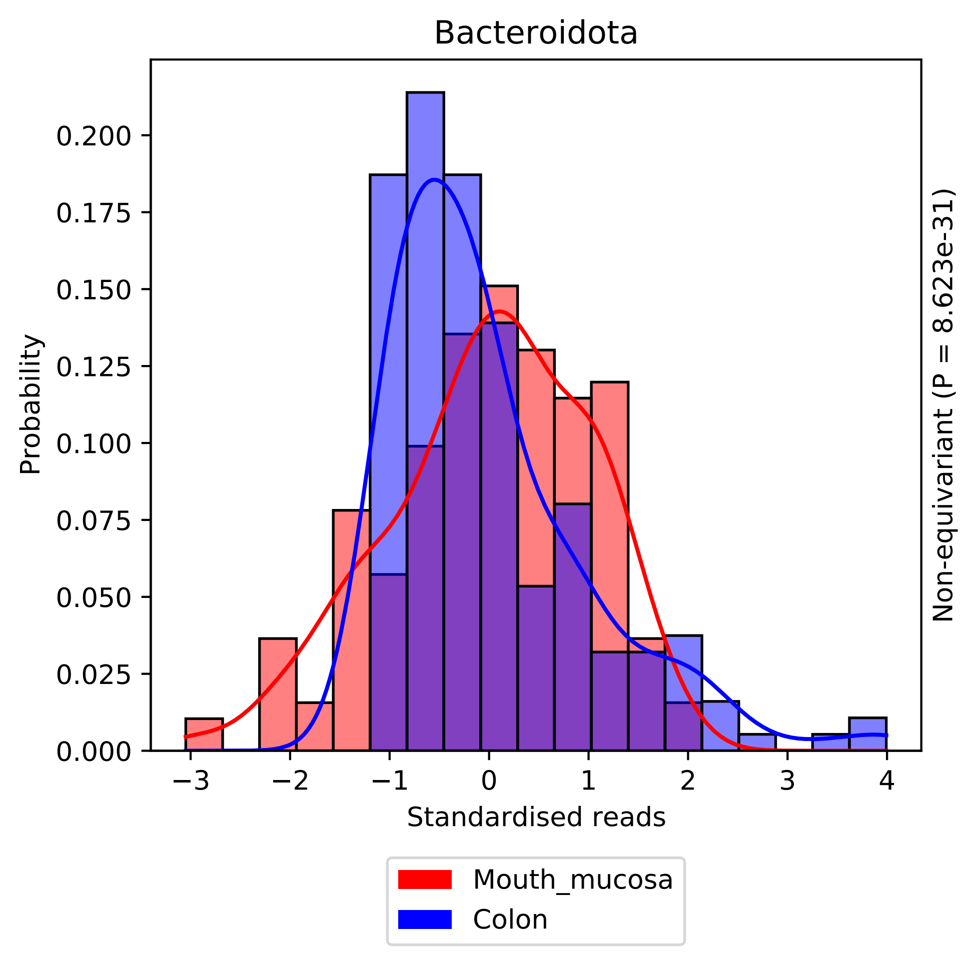
Bacteroidota
Probability
0.000
0.025
0.050
0.075
0.100
0.125
0.150
0.175
0.200
−3
−2
−1
0
1
2
3
4
Standardised reads
Non-equivariant (P = 8.623e-31)
Mouth_mucosa
Colon

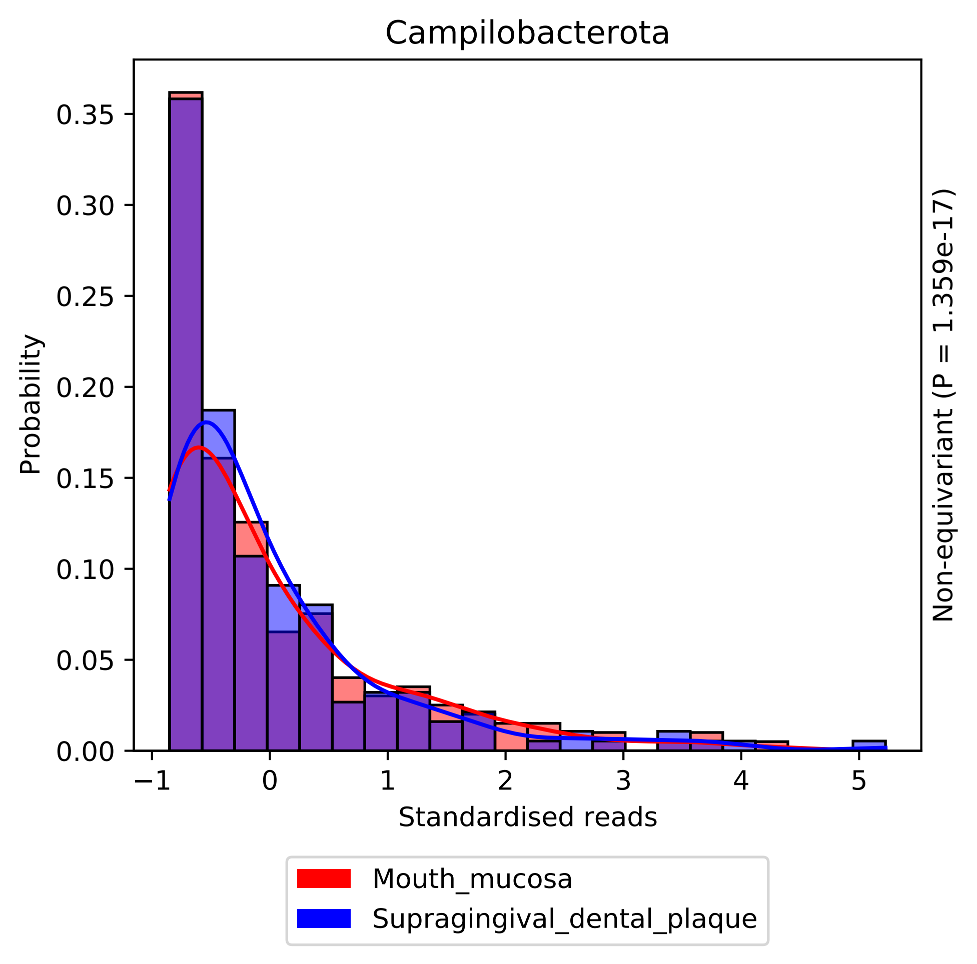
Campilobacterota
Probability
Standardised reads
Non-equivariant (P = 1.359e-17)
Mouth_mucosa
Supragingival_dental_plaque
0.00
0.05
0.10
0.15
0.20
0.25
0.30
0.35
−1
0
1
2
3
4
5

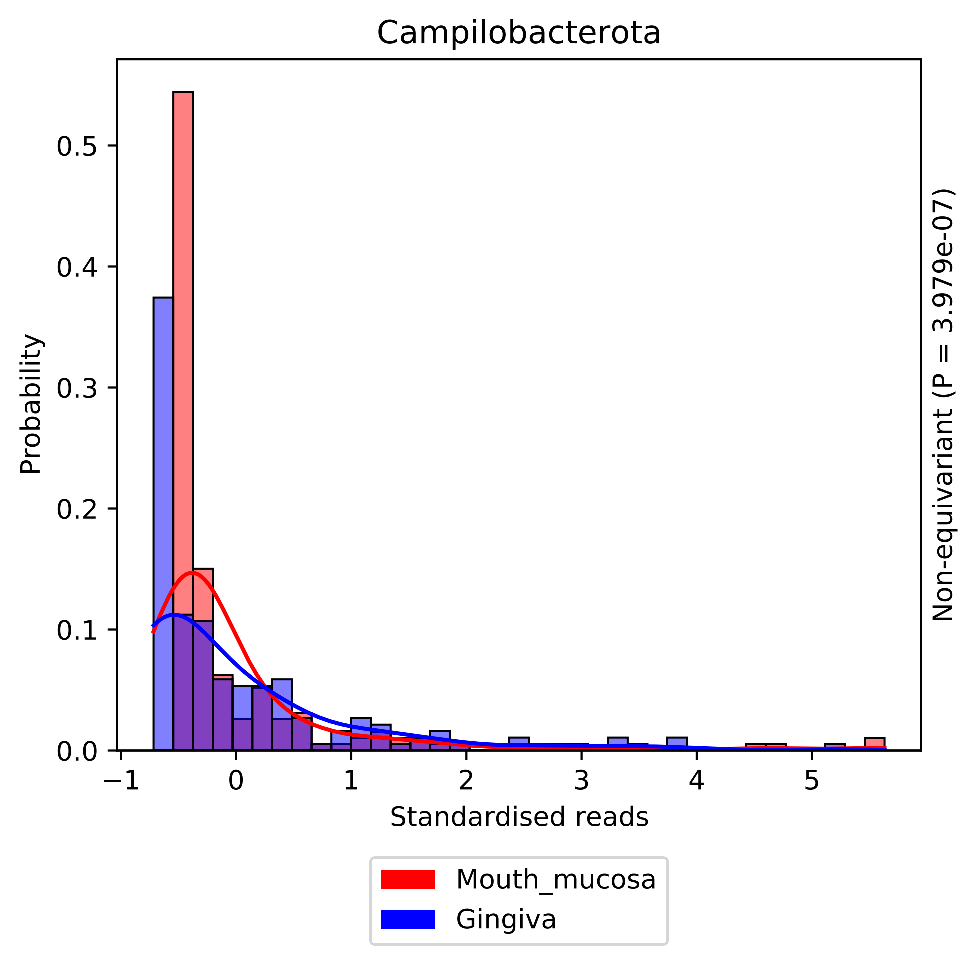
Campilobacterota
Probability
Standardised reads
Non-equivariant (P = 3.979e-07)
Mouth_mucosa
Gingiva
0.0
0.1
0.2
0.3
0.4
0.5
−1
0
1
2
3
4
5

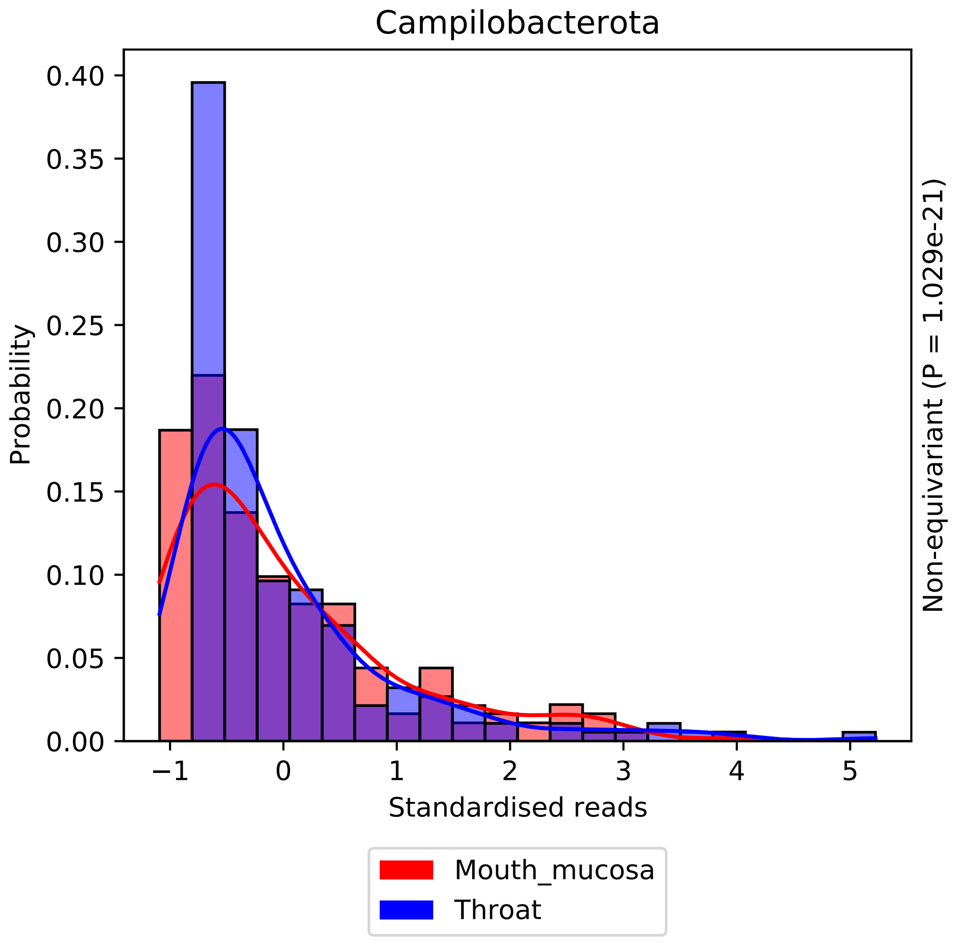
Campilobacterota
Probability
Standardised reads
Non-equivariant (P = 1.029e-21)
Mouth_mucosa
Throat
0.00
0.05
0.10
0.15
0.20
0.25
0.30
0.35
0.40
−1
0
1
2
3
4
5

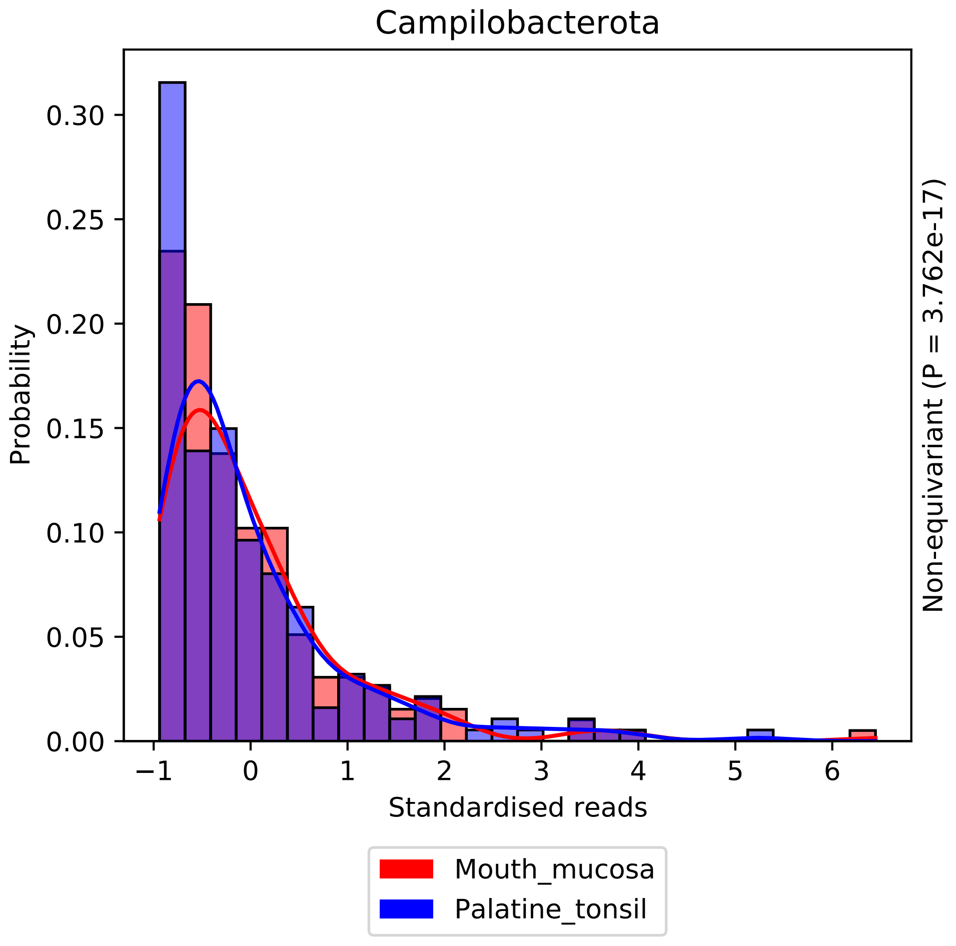

Campilobacterota
Probability
0.00
0.05
0.10
0.15
0.20
0.25
0.30
−1
0
1
2
3
4
5
6
Standardised reads
Non-equivariant (P = 3.762e-17)
Mouth_mucosa
Palatine_tonsil

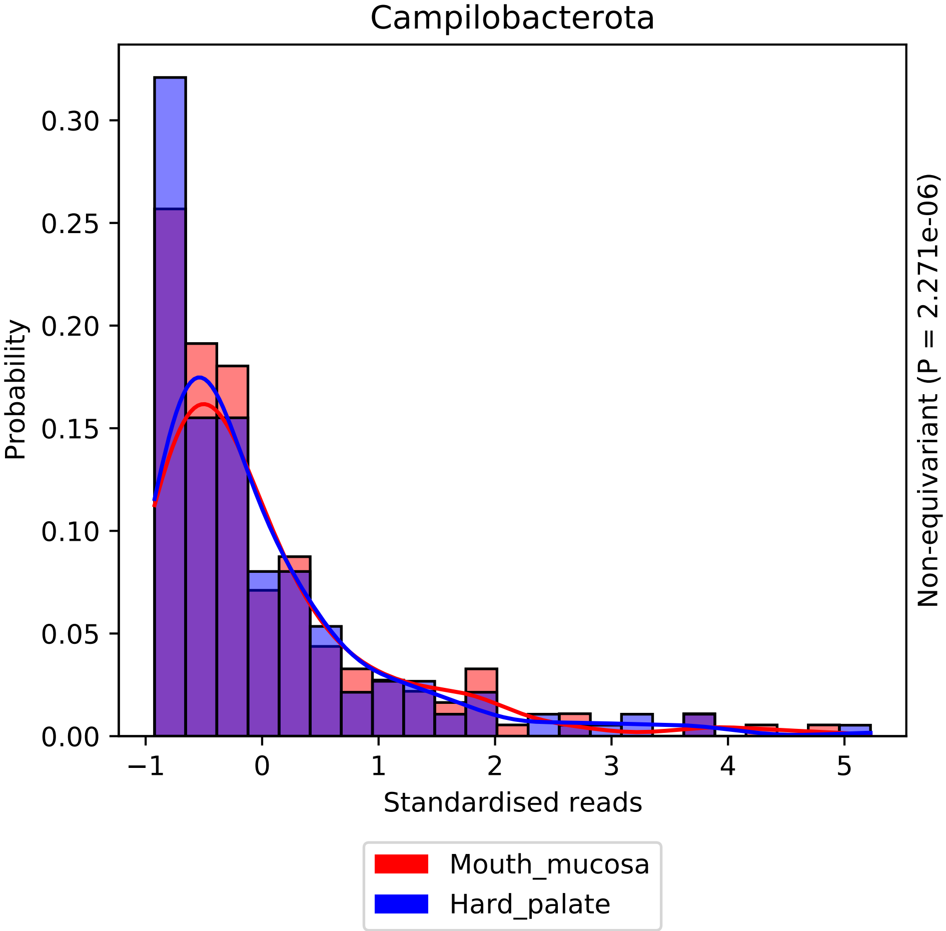

Campilobacterota
Probability
0.00
0.05
0.10
0.15
0.20
0.25
0.30
−1
0
1
2
3
4
5
Standardised reads
Non-equivariant (P = 2.271e-06)
Mouth_mucosa
Hard_palate

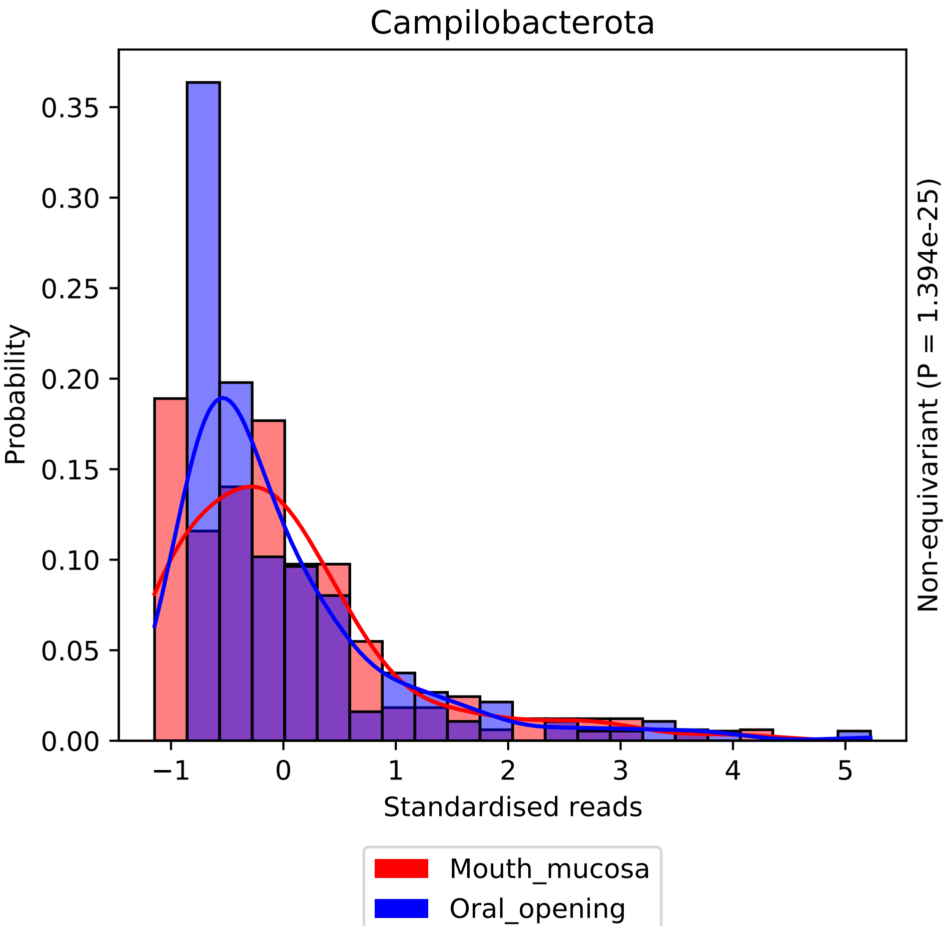

Campilobacterota
Probability
0.00
0.05
0.10
0.15
0.20
0.25
0.30
0.35
−1
0
1
2
3
4
5
Standardised reads
Non-equivariant (P = 1.394e-25)
Mouth_mucosa
Oral_opening

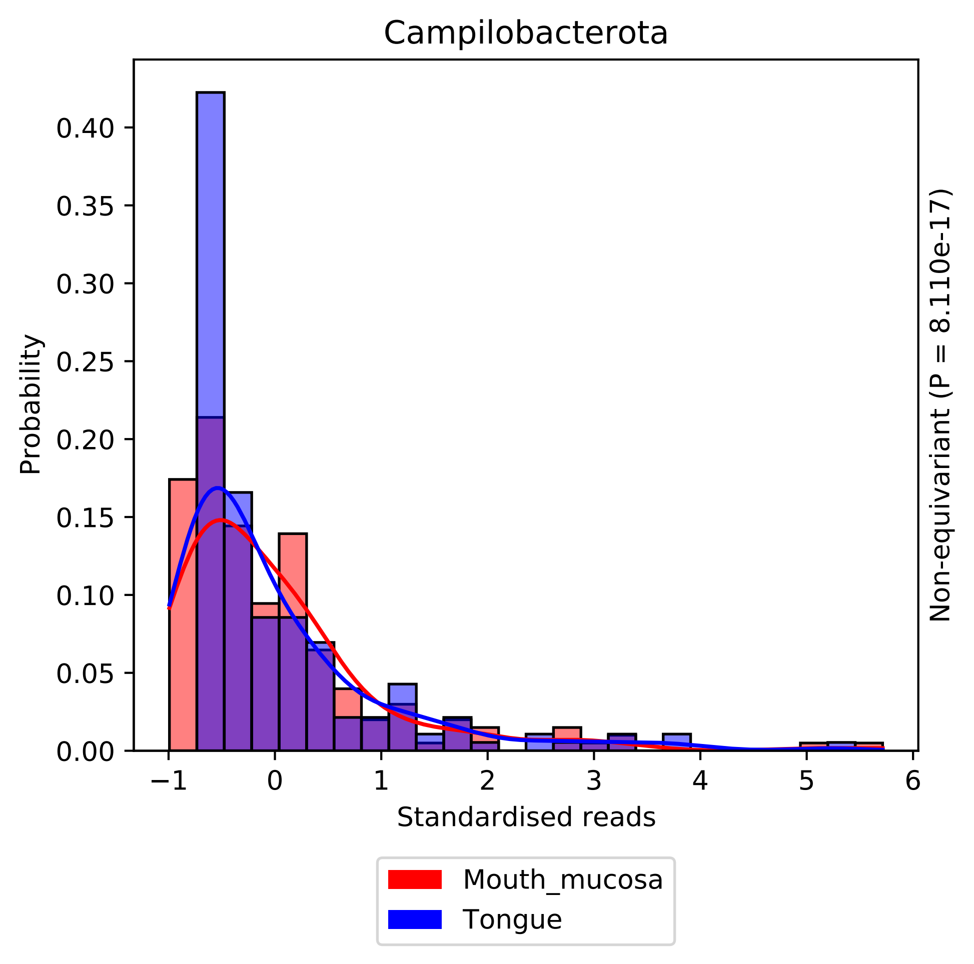
Campilobacterota
Probability
0.00
0.05
0.10
0.15
0.20
0.25
0.30
0.35
0.40
−1
0
1
2
3
4
5
6
Standardised reads
Non-equivariant (P = 8.110e-17)
Mouth_mucosa
Tongue

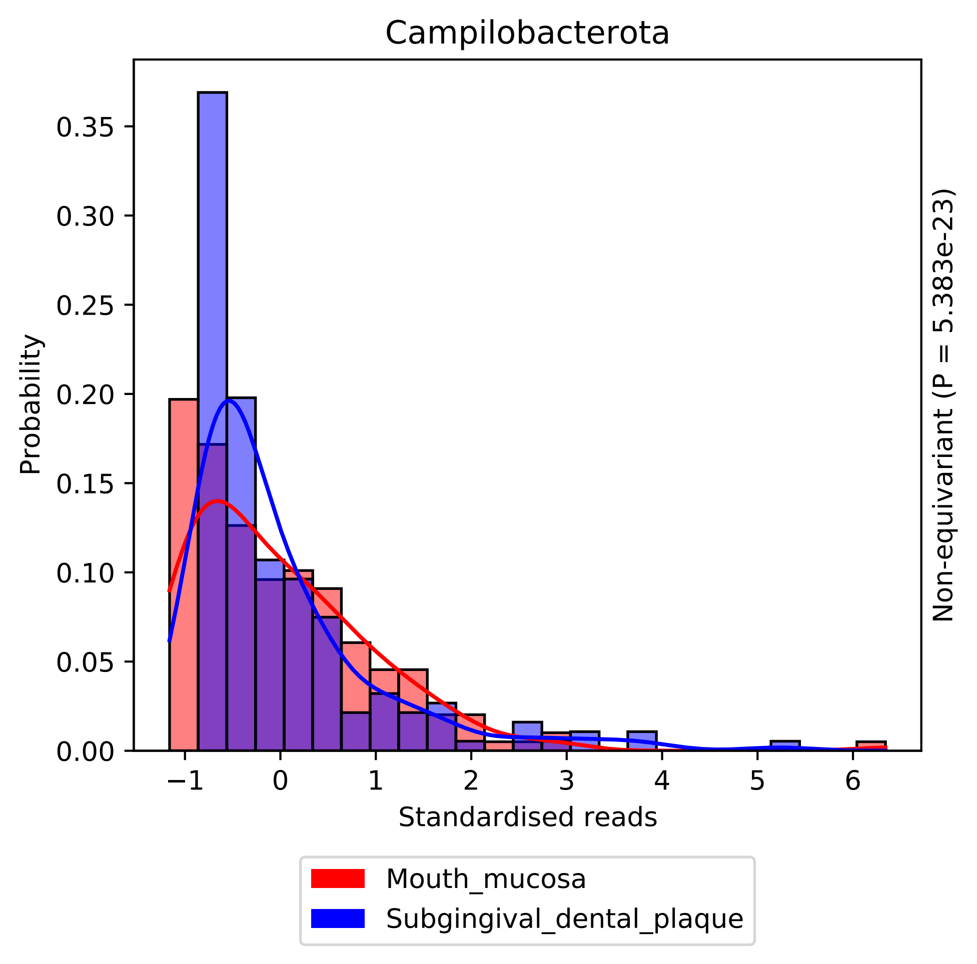

Campilobacterota
Probability
0.00
0.05
0.10
0.15
0.20
0.25
0.30
0.35
−1
0
1
2
3
4
5
6
Standardised reads
Non-equivariant (P = 5.383e-23)
Mouth_mucosa
Subgingival_dental_plaque

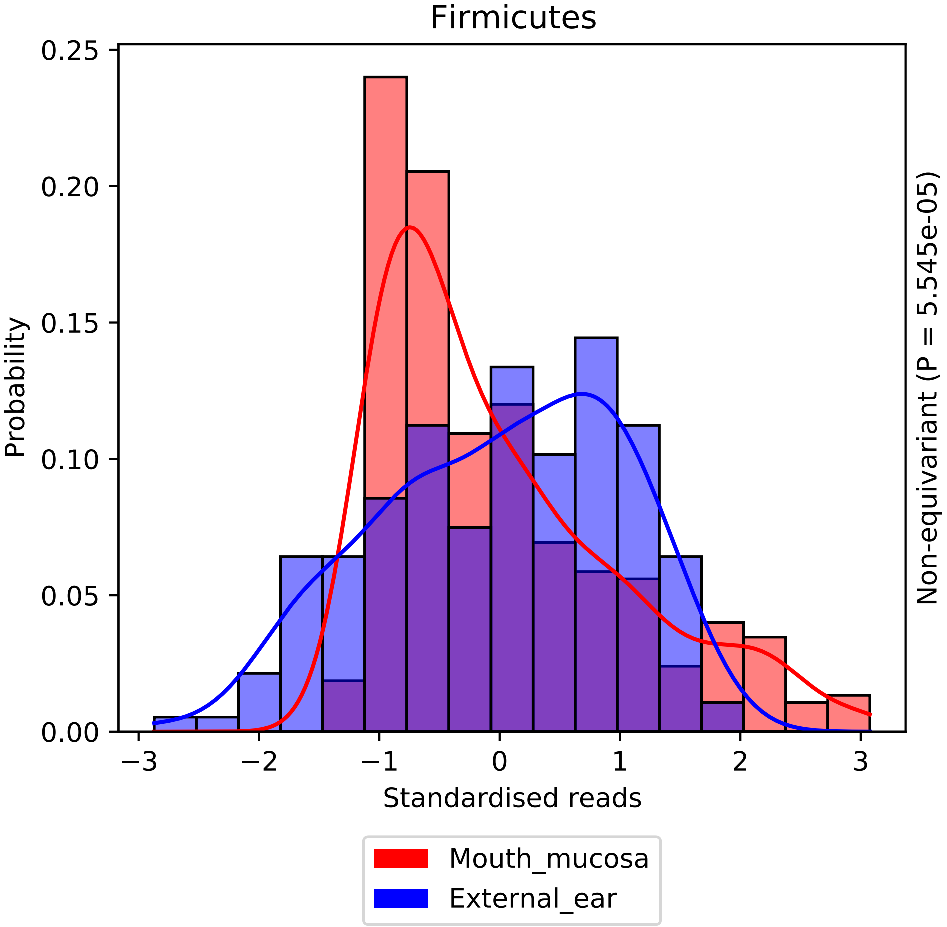

Firmicutes
0.25
0.20
0.15
0.10
0.05
0.00
Probability
−3
−2
−1
0
1
2
3
Standardised reads
Non-equivariant (P = 5.545e-05)
Mouth_mucosa
External_ear

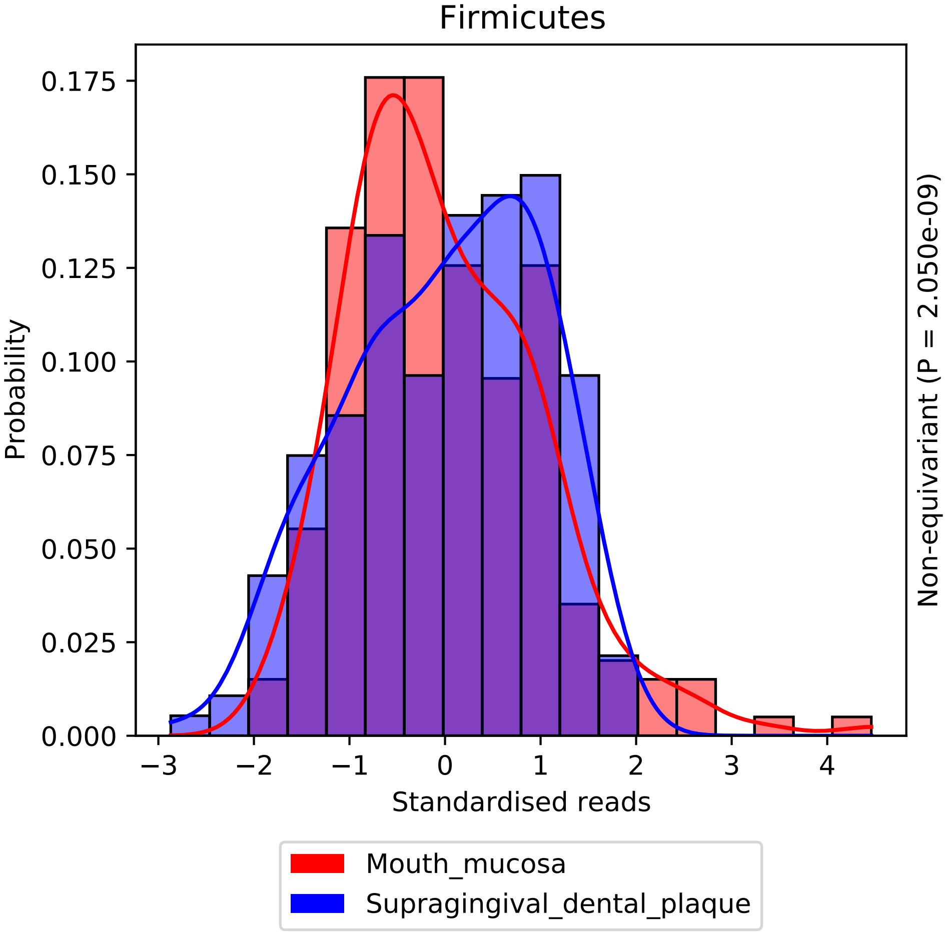

Firmicutes
Probability
0.000
0.025
0.050
0.075
0.100
0.125
0.150
0.175
−3
−2
−1
0
1
2
3
4
Standardised reads
Non-equivariant (P = 2.050e-09)
Mouth_mucosa
Supragingival_dental_plaque

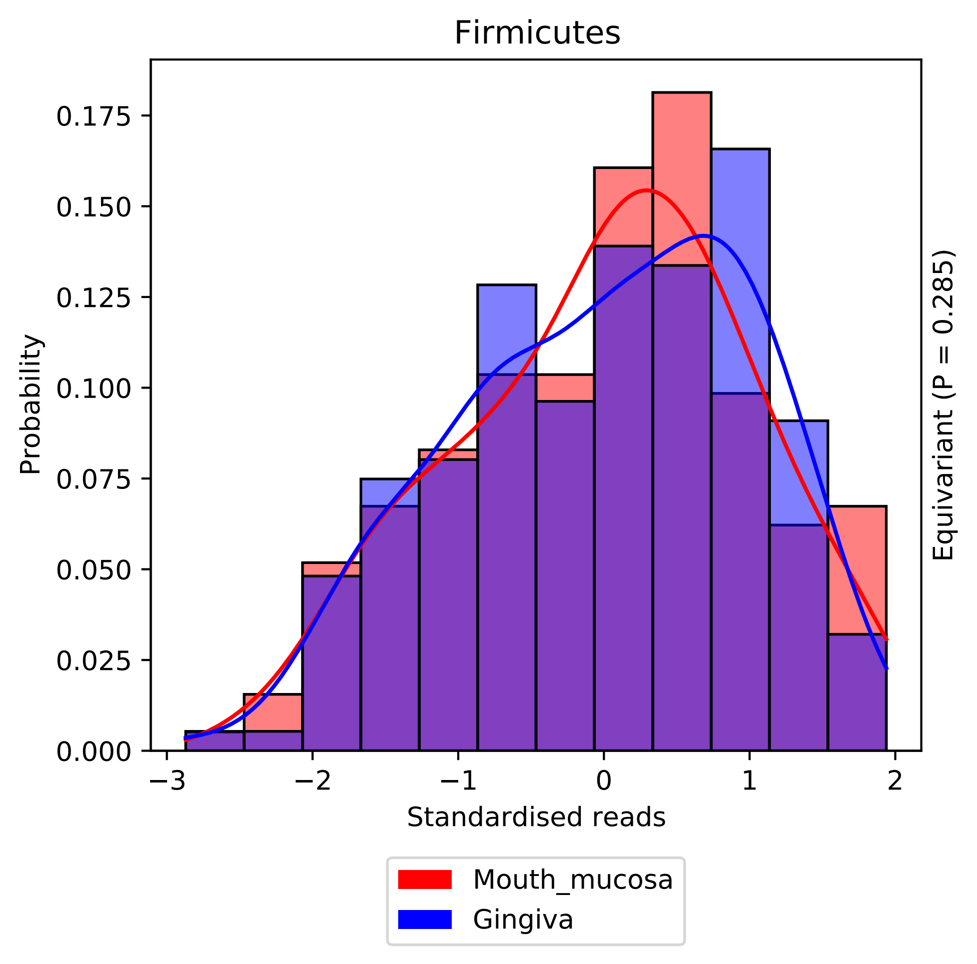

Firmicutes
Probability
0.000
0.025
0.050
0.075
0.100
0.125
0.150
0.175
−3
−2
−1
0
1
2
Standardised reads
Equivariant (P = 0.285)
Mouth_mucosa
Gingiva

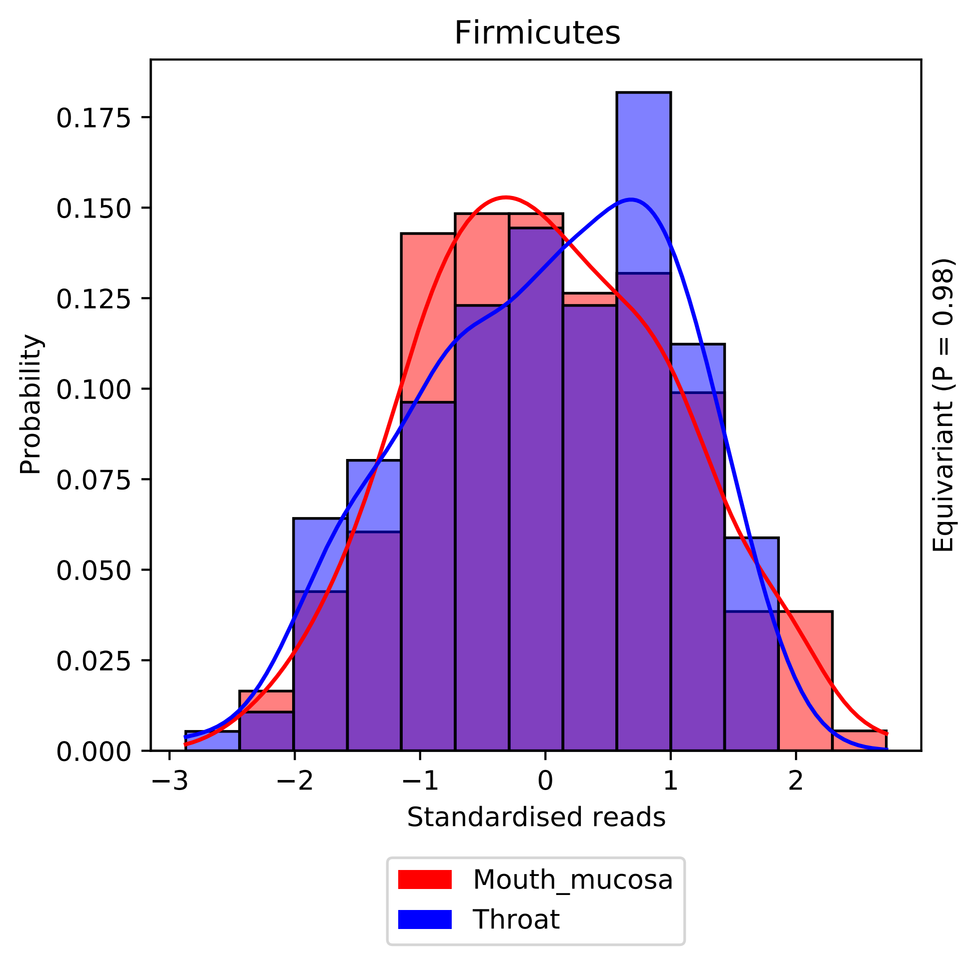
Firmicutes
0.175
0.150
0.125
0.100
0.075
0.050
0.025
0.000
Probability
−3
−2
−1
0
1
2
Standardised reads
Equivariant (P = 0.98)
Mouth_mucosa
Throat

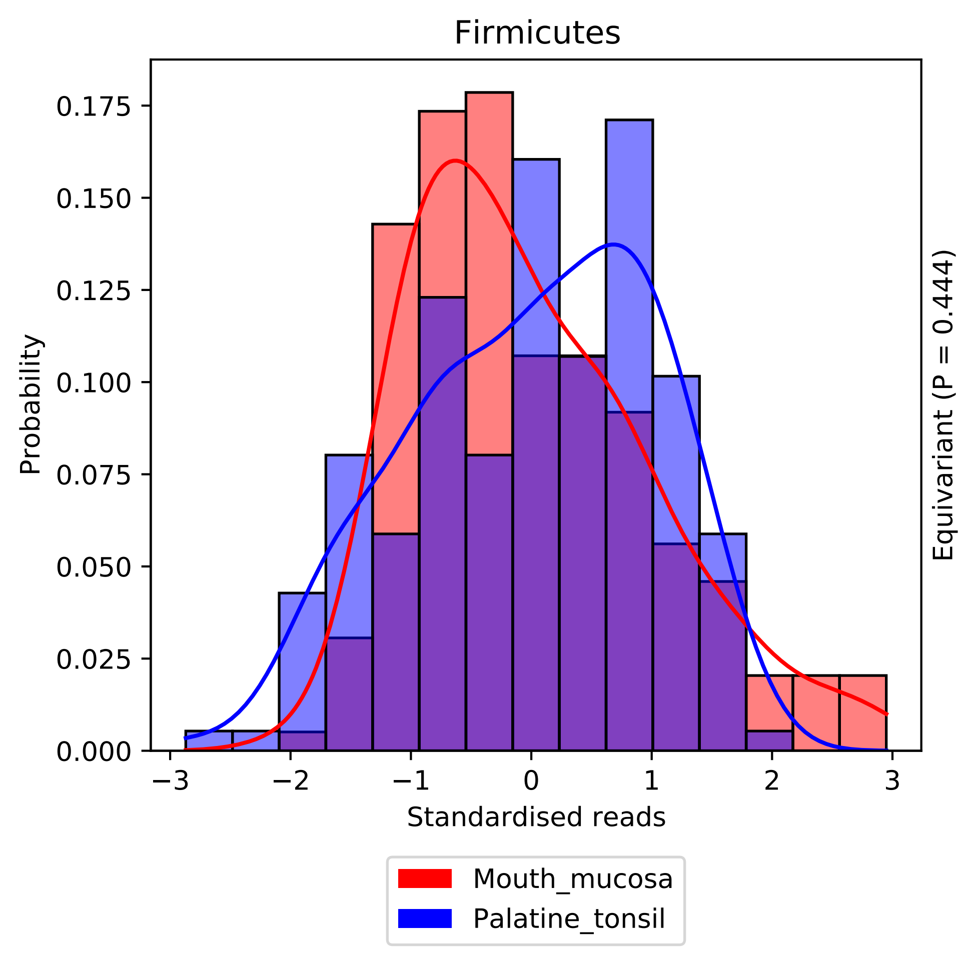

Firmicutes
Probability
0.000
0.025
0.050
0.075
0.100
0.125
0.150
0.175
−3
−2
−1
0
1
2
3
Standardised reads
Equivariant (P = 0.444)
Mouth_mucosa
Palatine_tonsil

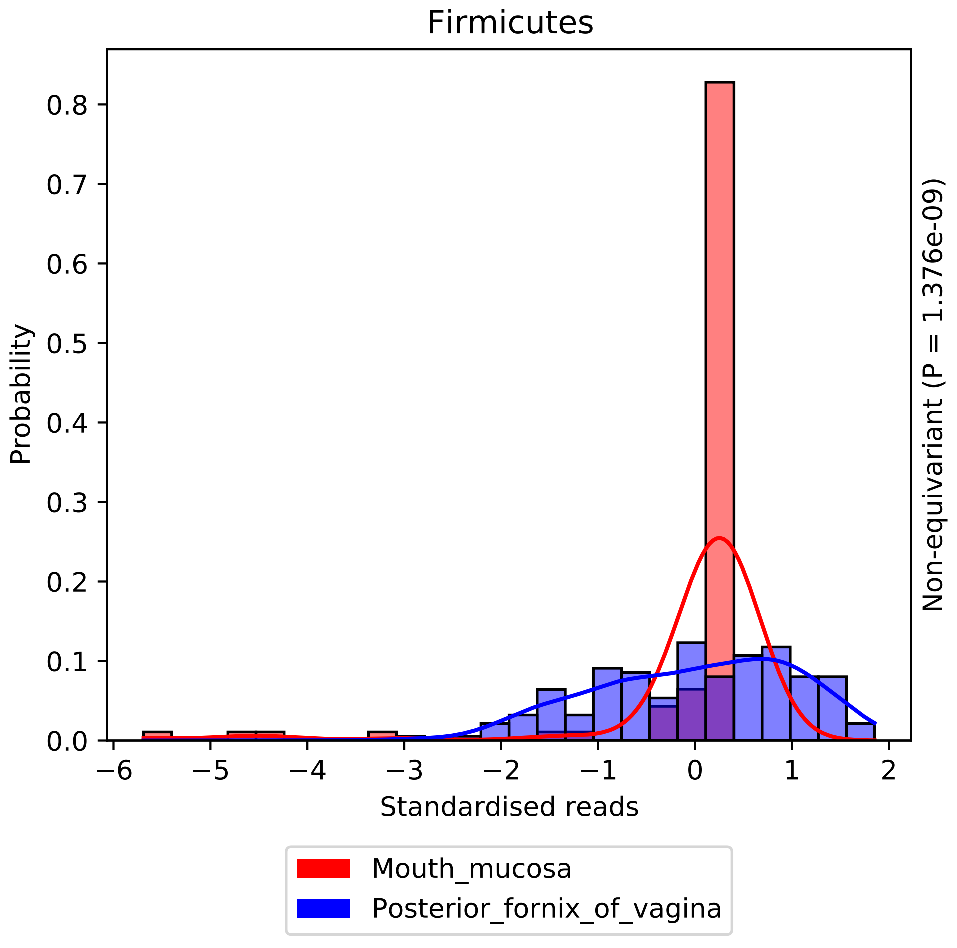
Firmicutes
Probability
0.0
0.1
0.2
0.3
0.4
0.5
0.6
0.7
0.8
−6
−5
−4
−3
−2
−1
0
1
2
Standardised reads
Non-equivariant (P = 1.376e-09)
Mouth_mucosa
Posterior_fornix_of_vagina

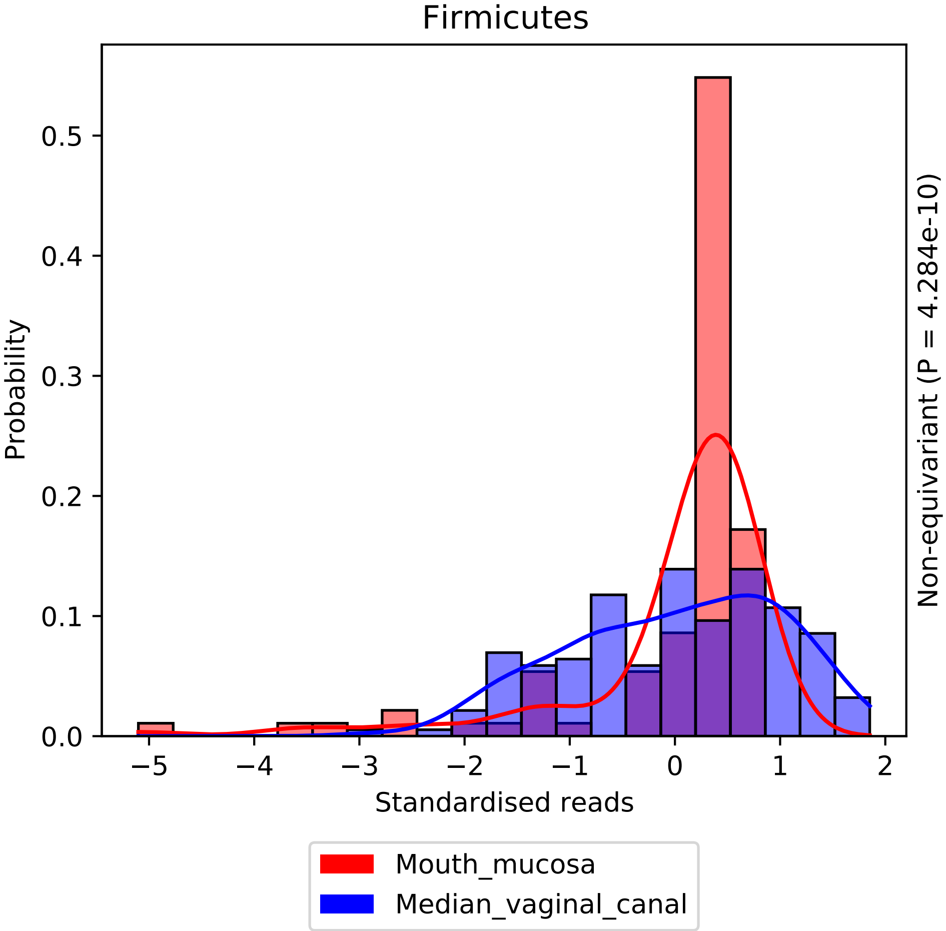

Firmicutes
Probability
0.0
0.1
0.2
0.3
0.4
0.5
−5
−4
−3
−2
−1
0
1
2
Standardised reads
Non-equivariant (P = 4.284e-10)
Mouth_mucosa
Median_vaginal_canal

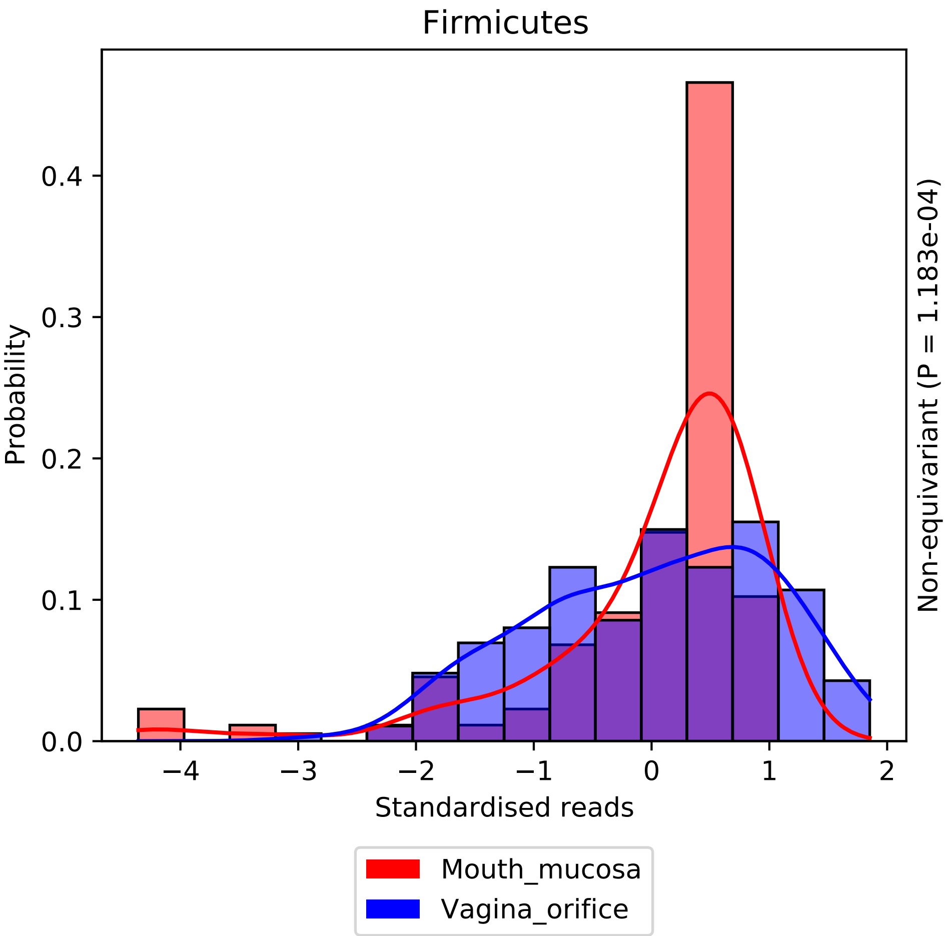

Firmicutes
Probability
0.0
0.1
0.2
0.3
0.4
−4
−3
−2
−1
0
1
2
Standardised reads
Non-equivariant (P = 1.183e-04)
Mouth_mucosa
Vagina_orifice

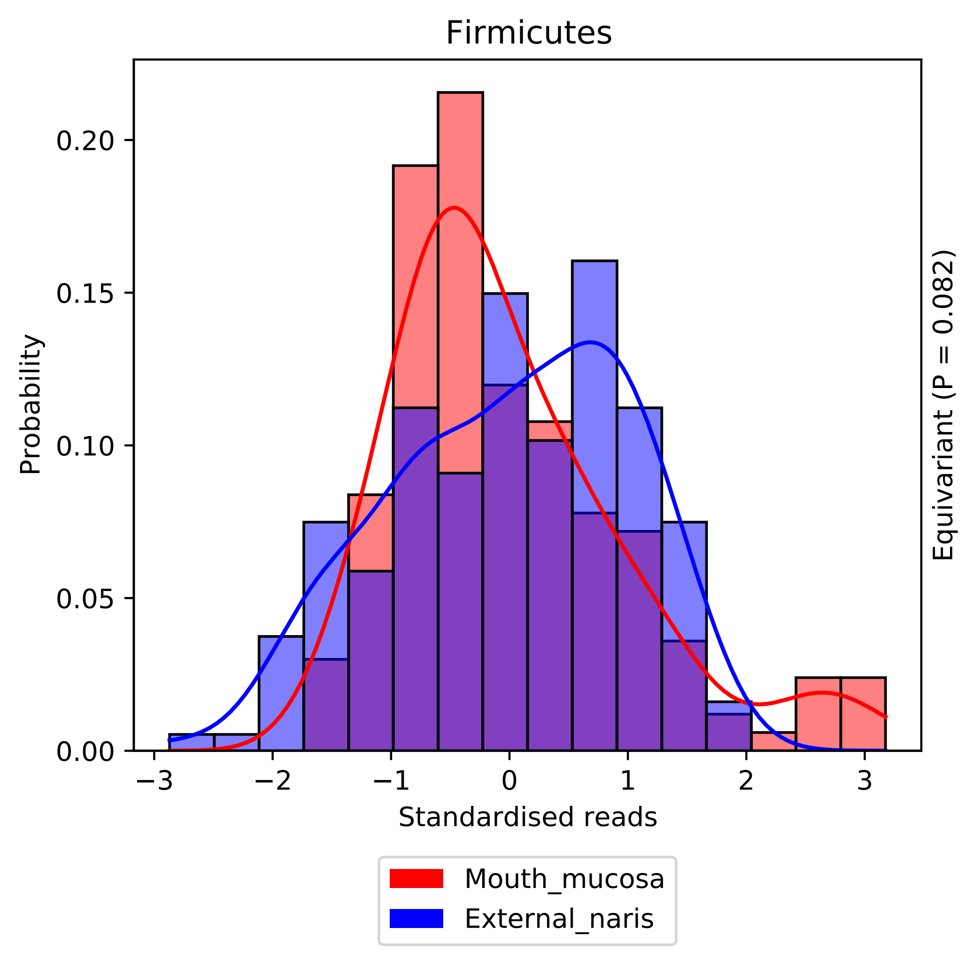

Firmicutes
Probability
0.00
0.05
0.10
0.15
0.20
−3
−2
−1
0
1
2
3
Standardised reads
Equivariant (P = 0.082)
Mouth_mucosa
External_naris

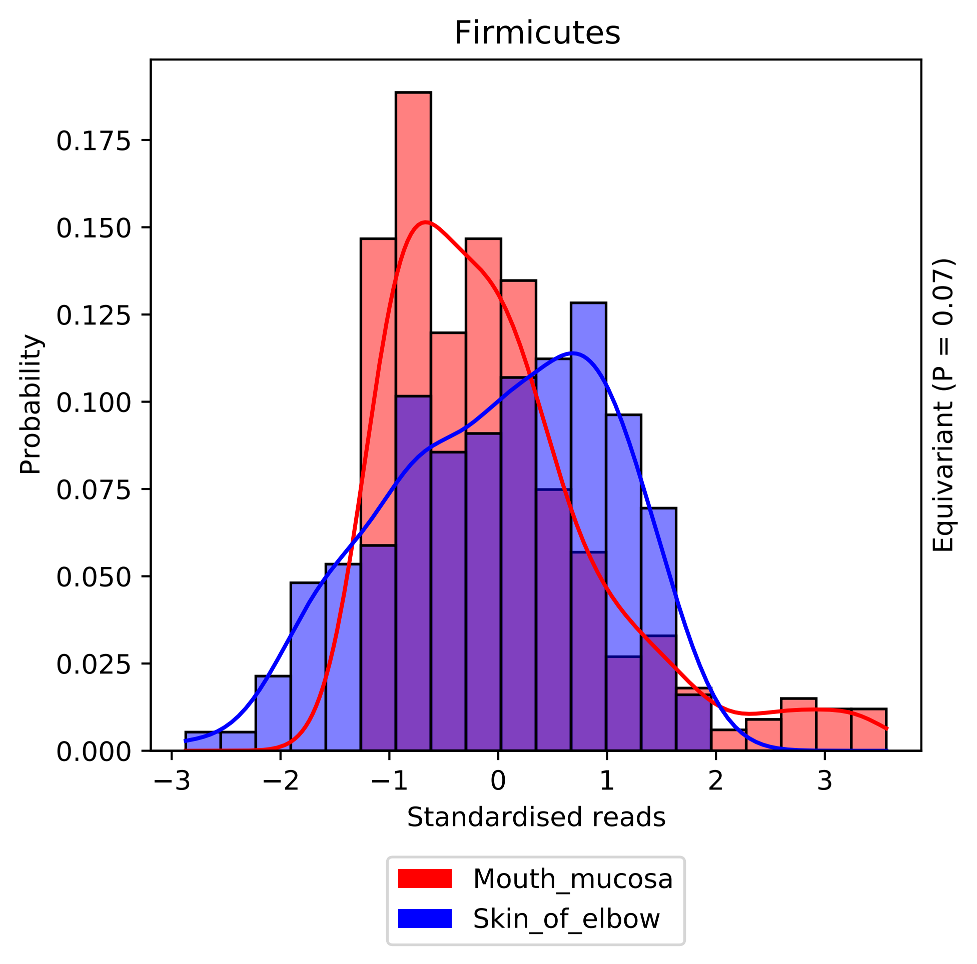

Firmicutes
0.000
0.025
0.050
0.075
0.100
0.125
0.150
0.175
Probability
−3
−2
−1
0
1
2
3
Standardised reads
Equivariant (P = 0.07)
Mouth_mucosa
Skin_of_elbow

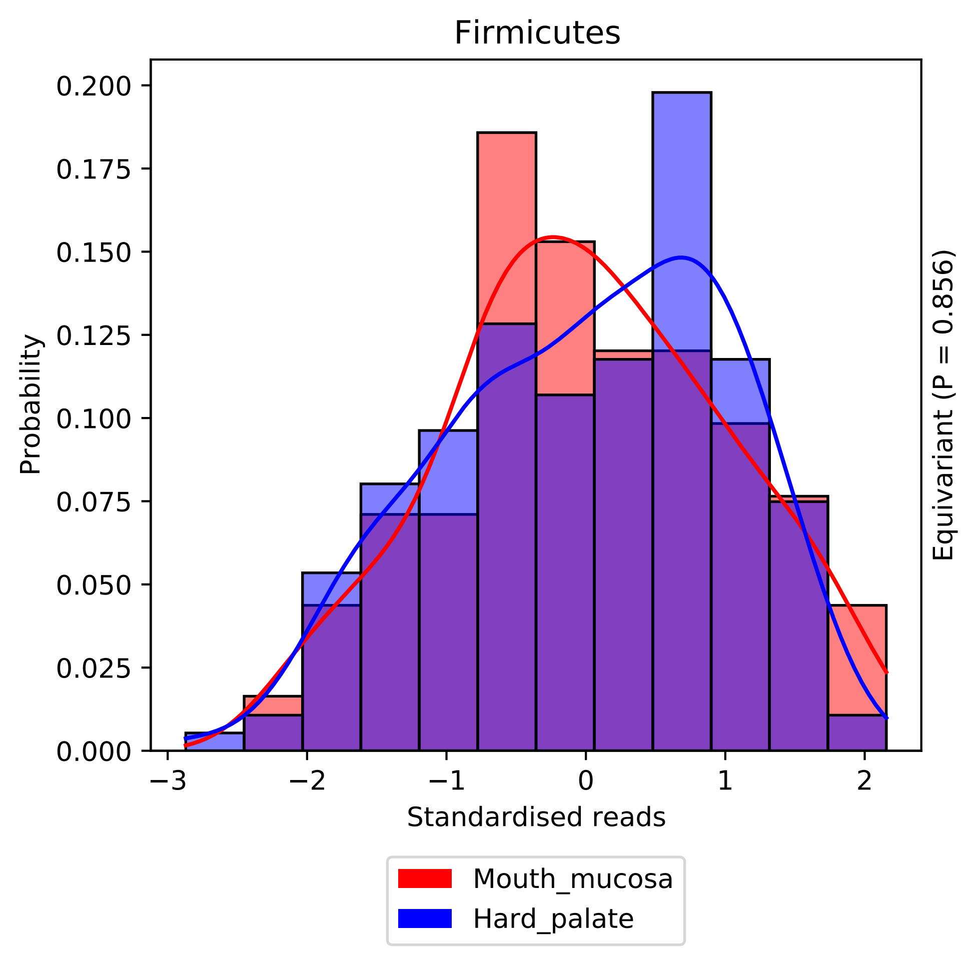

Firmicutes
Probability
0.200
0.175
0.150
0.125
0.100
0.075
0.050
0.025
0.000
−3
−2
−1
0
1
2
Standardised reads
Equivariant (P = 0.856)
Mouth_mucosa
Hard_palate

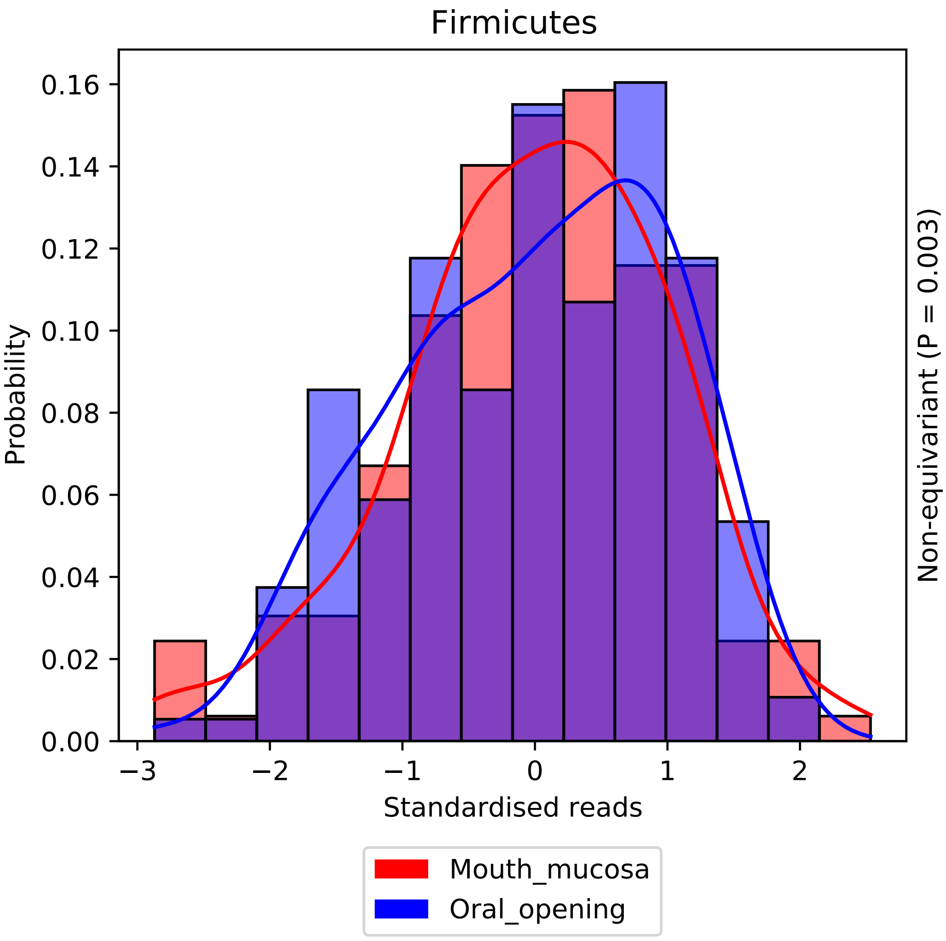
Firmicutes
Probability
0.16
0.14
0.12
0.10
0.08
0.06
0.04
0.02
0.00
−3
−2
−1
0
1
2
Standardised reads
Non-equivariant (P = 0.003)
Mouth_mucosa
Oral_opening

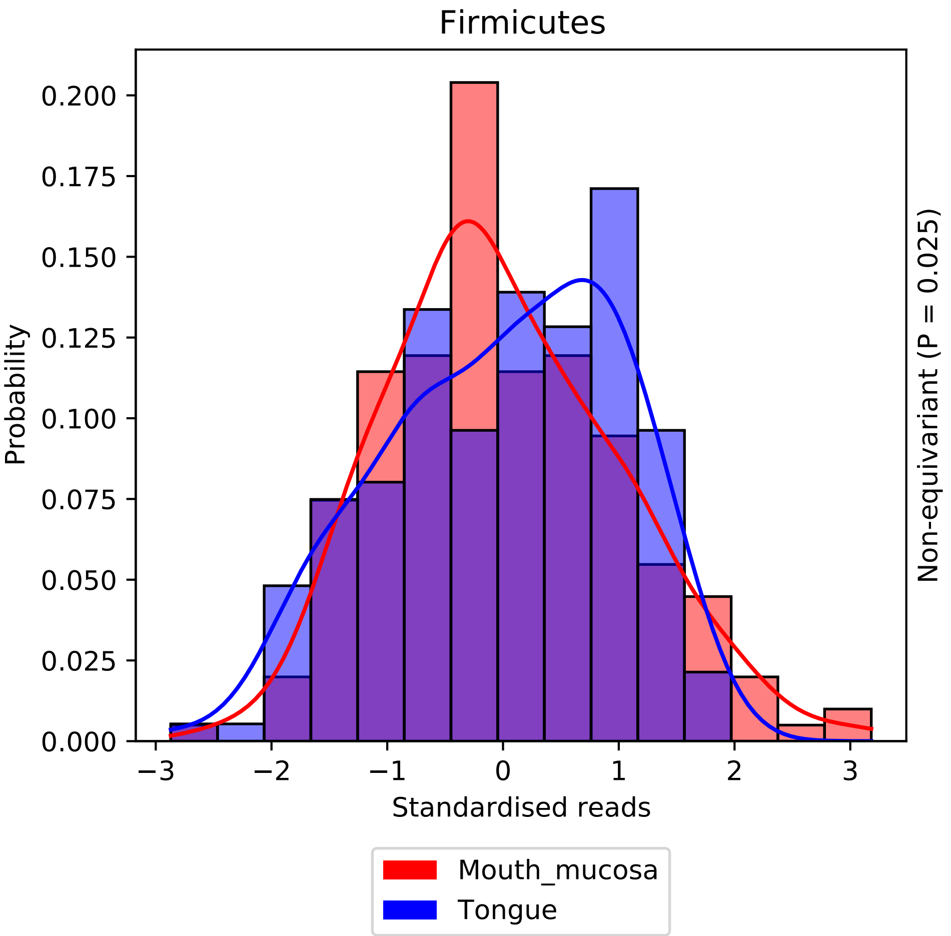
Firmicutes
Probability
0.200
0.175
0.150
0.125
0.100
0.075
0.050
0.025
0.000
−3
−2
−1
0
1
2
3
Standardised reads
Non-equivariant (P = 0.025)
Mouth_mucosa
Tongue

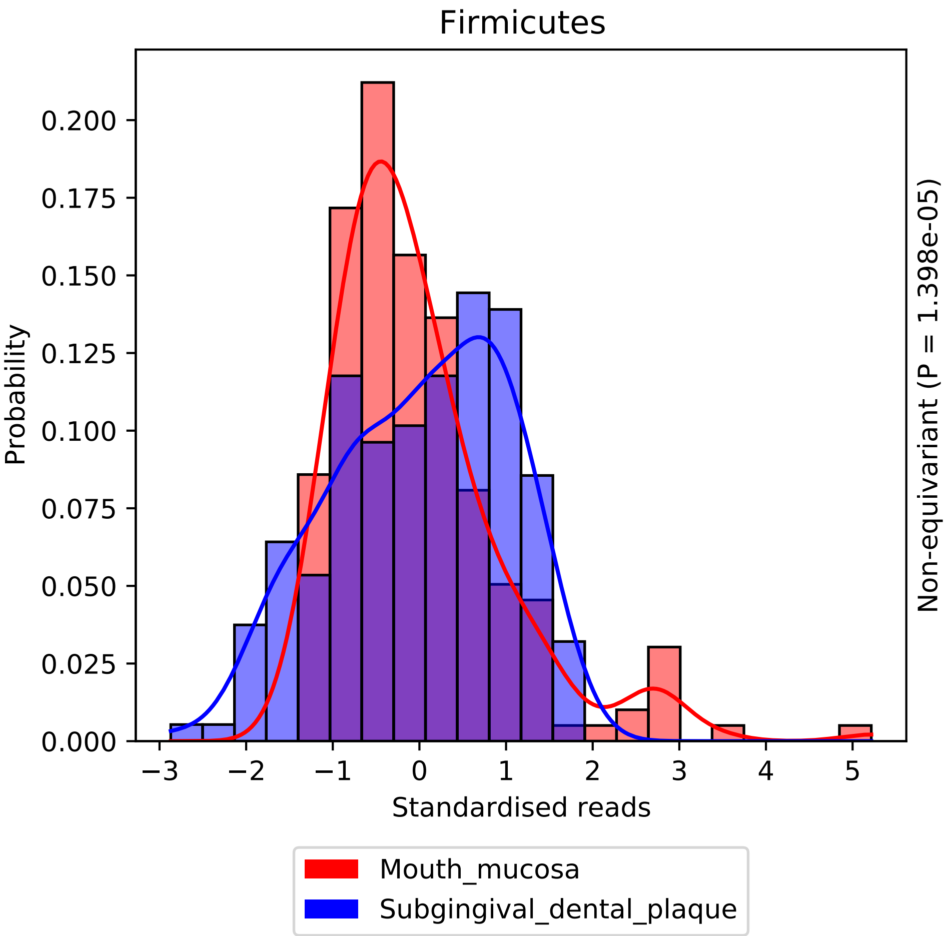
Firmicutes
Probability
Standardised reads
Non-equivariant (P = 1.398e-05)
0.000
0.025
0.050
0.075
0.100
0.125
0.150
0.175
0.200
−3
−2
−1
0
1
2
3
4
5
Mouth_mucosa
Subgingival_dental_plaque

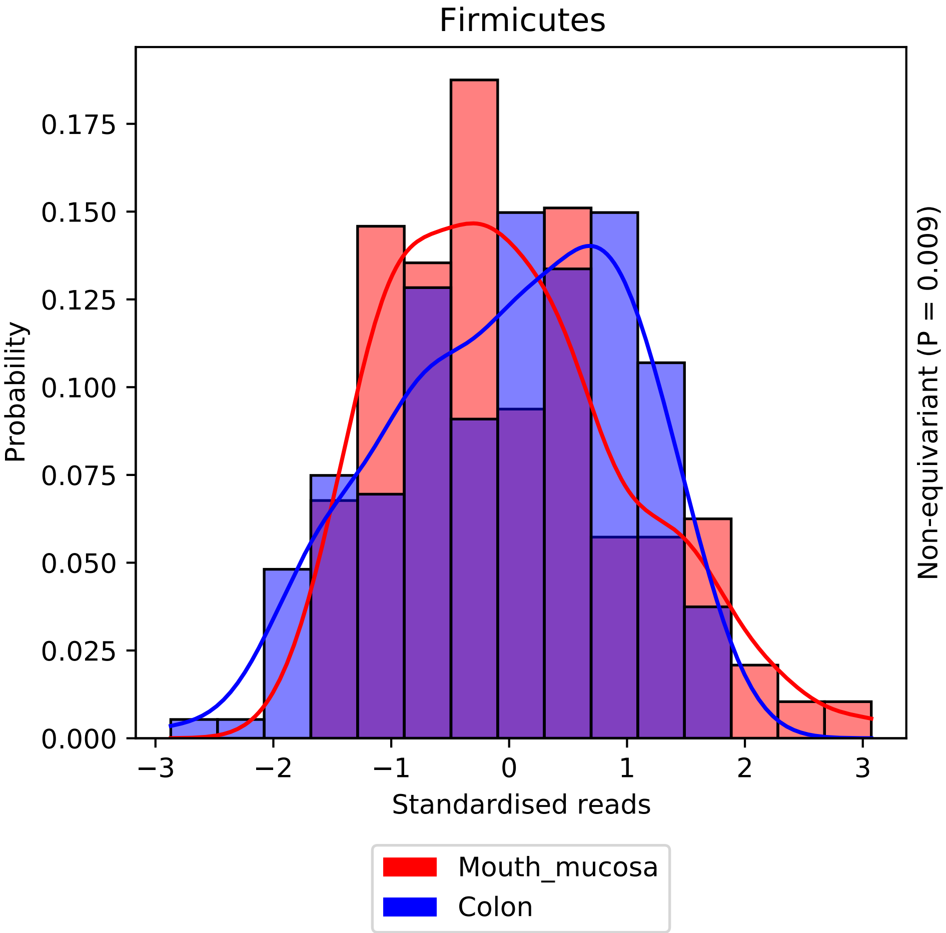
Firmicutes
0.000
0.025
0.050
0.075
0.100
0.125
0.150
0.175
Probability
−3
−2
−1
0
1
2
3
Standardised reads
Non-equivariant (P = 0.009)
Mouth_mucosa
Colon

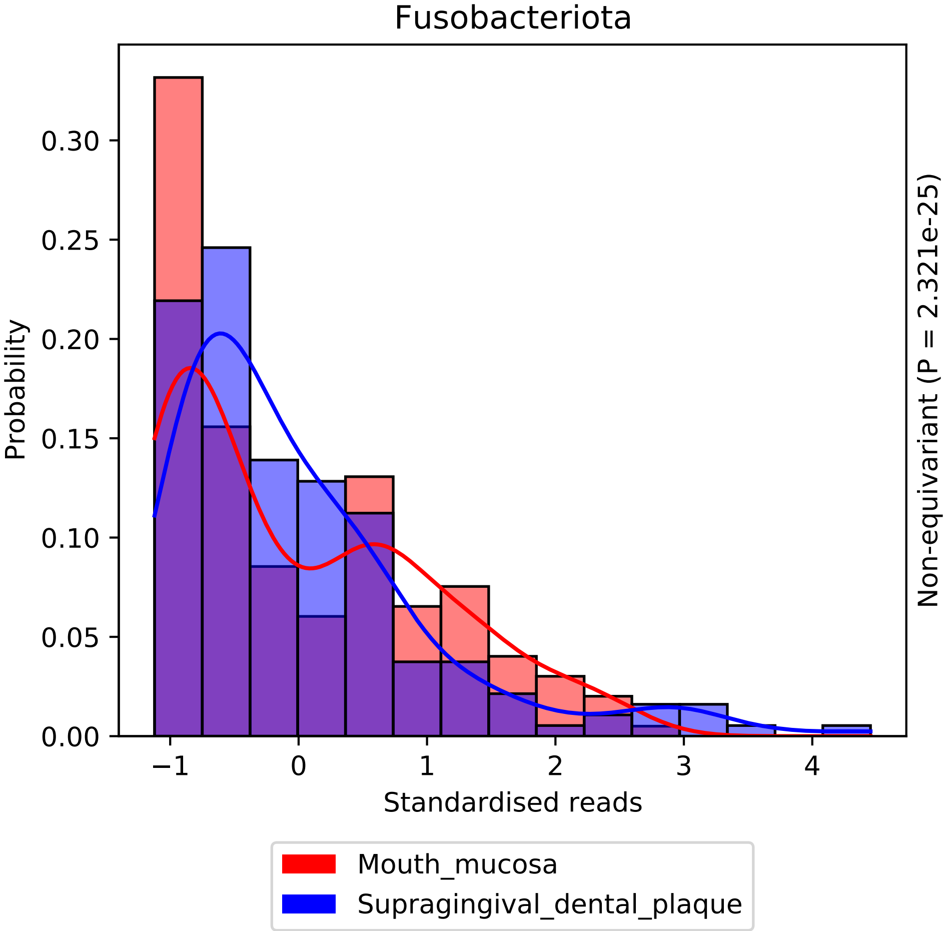
Fusobacteriota
Probability
Standardised reads
Non-equivariant (P = 2.321e-25)
0.00
0.05
0.10
0.15
0.20
0.25
0.30
−1
0
1
2
3
4
Mouth_mucosa
Supragingival_dental_plaque

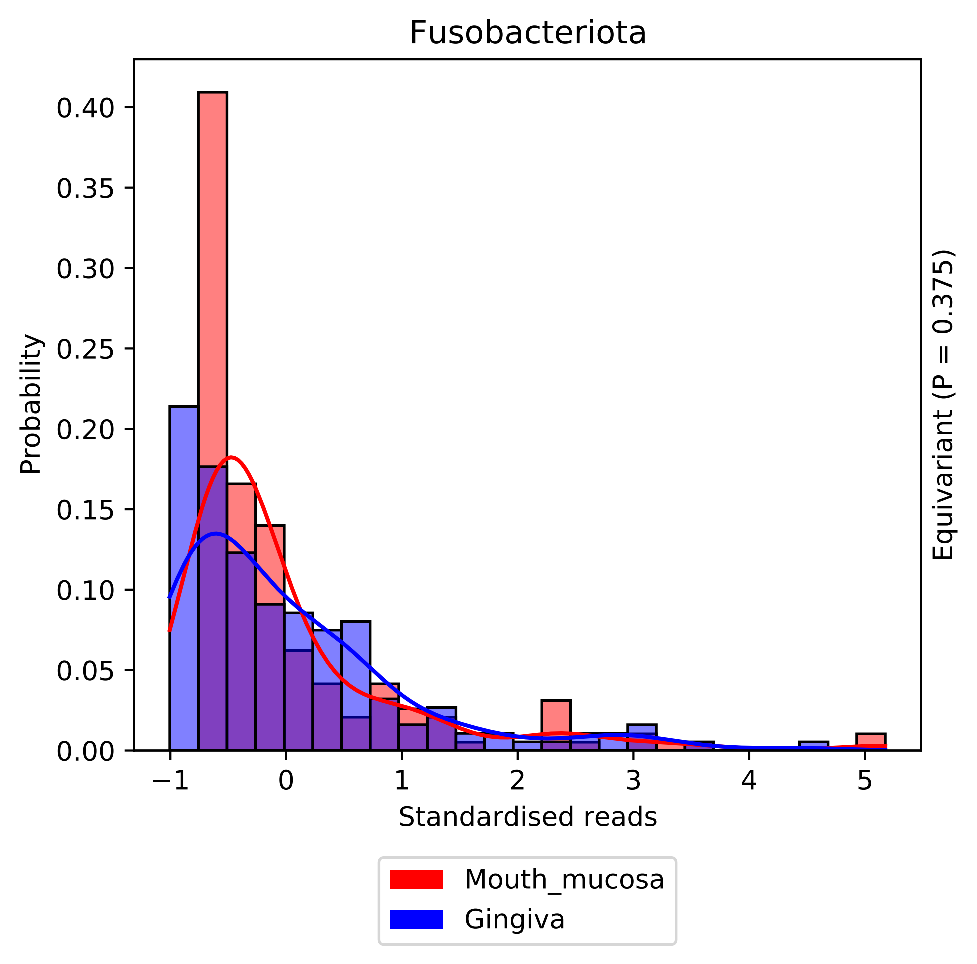
Fusobacteriota
Probability
Standardised reads
Equivariant (P = 0.375)
Mouth_mucosa
Gingiva
0.00
0.05
0.10
0.15
0.20
0.25
0.30
0.35
0.40
−1
0
1
2
3
4
5

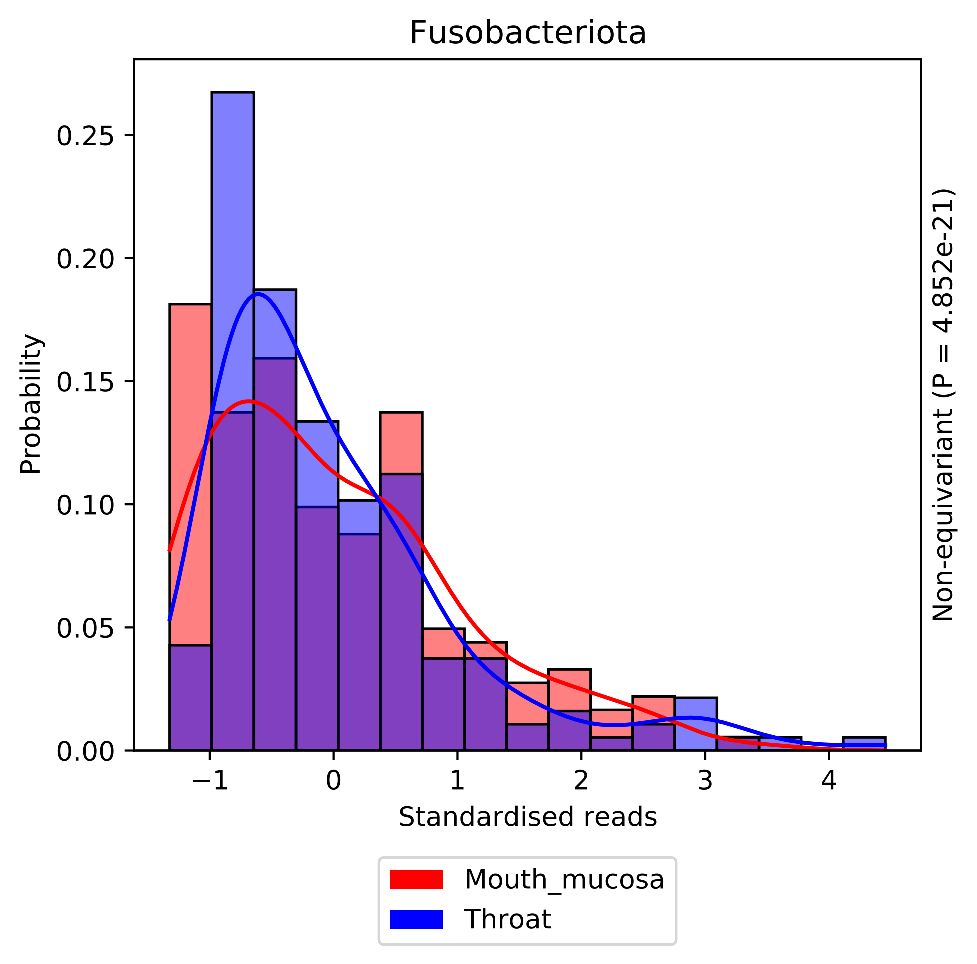

Fusobacteriota
Probability
0.00
0.05
0.10
0.15
0.20
0.25
−1
0
1
2
3
4
Standardised reads
Non-equivariant (P = 4.852e-21)
Mouth_mucosa
Throat

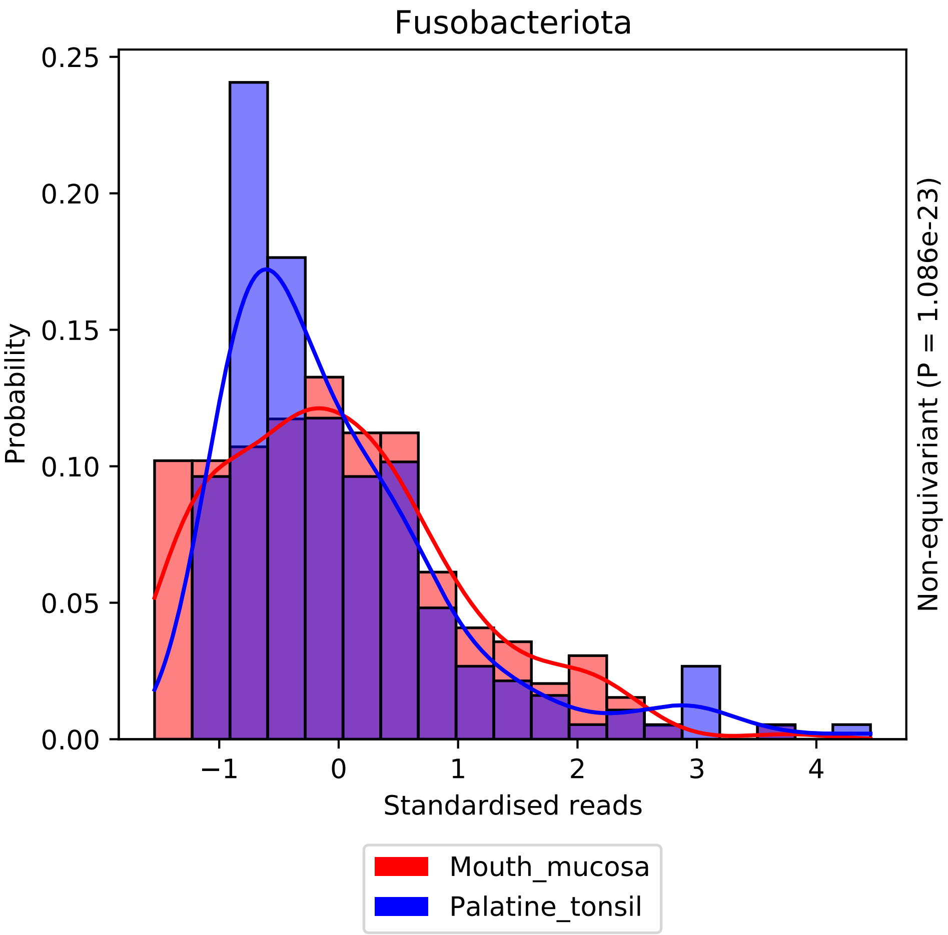
Fusobacteriota
Probability
Standardised reads
Non-equivariant (P = 1.086e-23)
0.25
0.20
0.15
0.10
0.05
0.00
−1
0
1
2
3
4
Mouth_mucosa
Palatine_tonsil

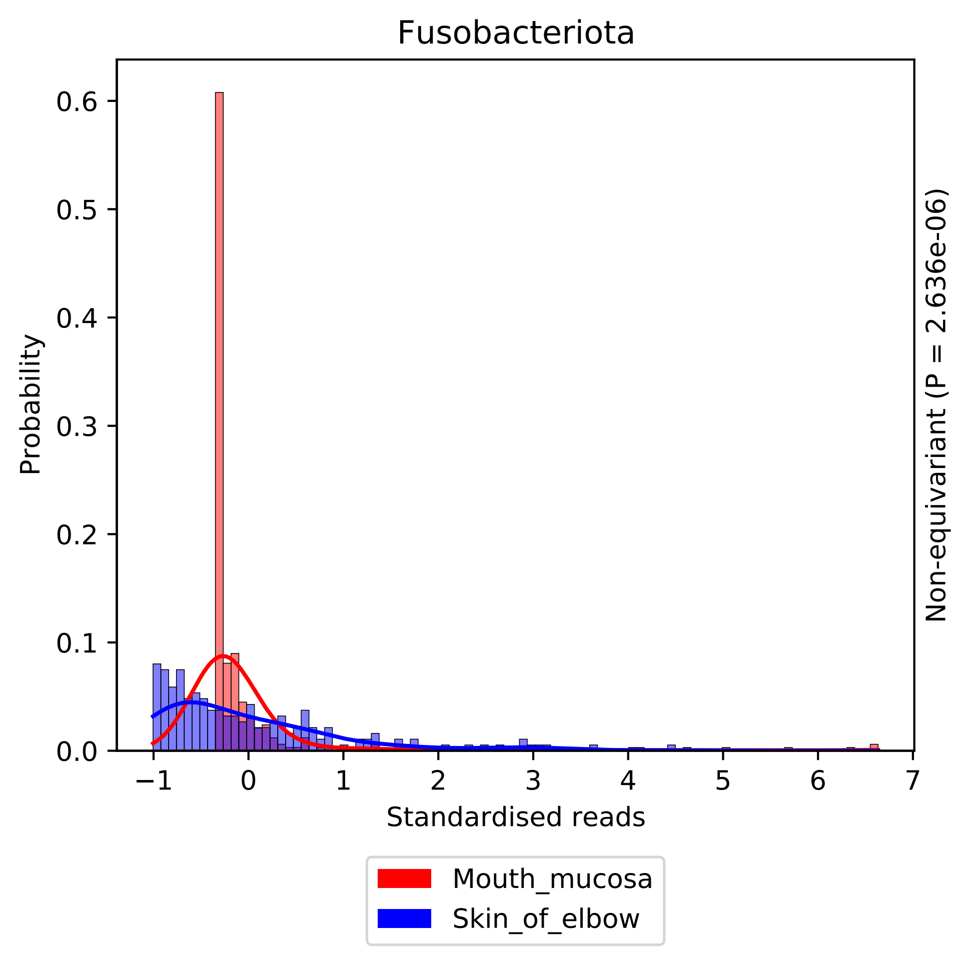
Fusobacteriota
Probability
0.0
0.1
0.2
0.3
0.4
0.5
0.6
−1
0
1
2
3
4
5
6
7
Standardised reads
Non-equivariant (P = 2.636e-06)
Mouth_mucosa
Skin_of_elbow

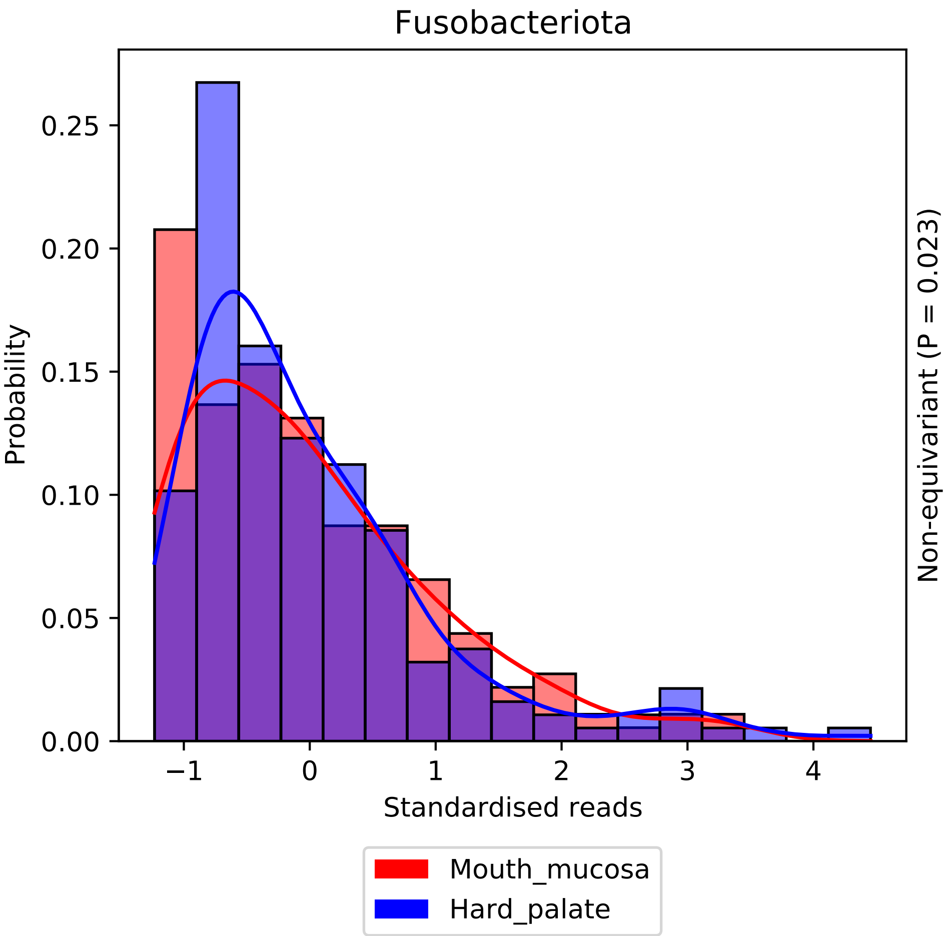

Fusobacteriota
Probability
0.00
0.05
0.10
0.15
0.20
0.25
−1
0
1
2
3
4
Standardised reads
Non-equivariant (P = 0.023)
Mouth_mucosa
Hard_palate

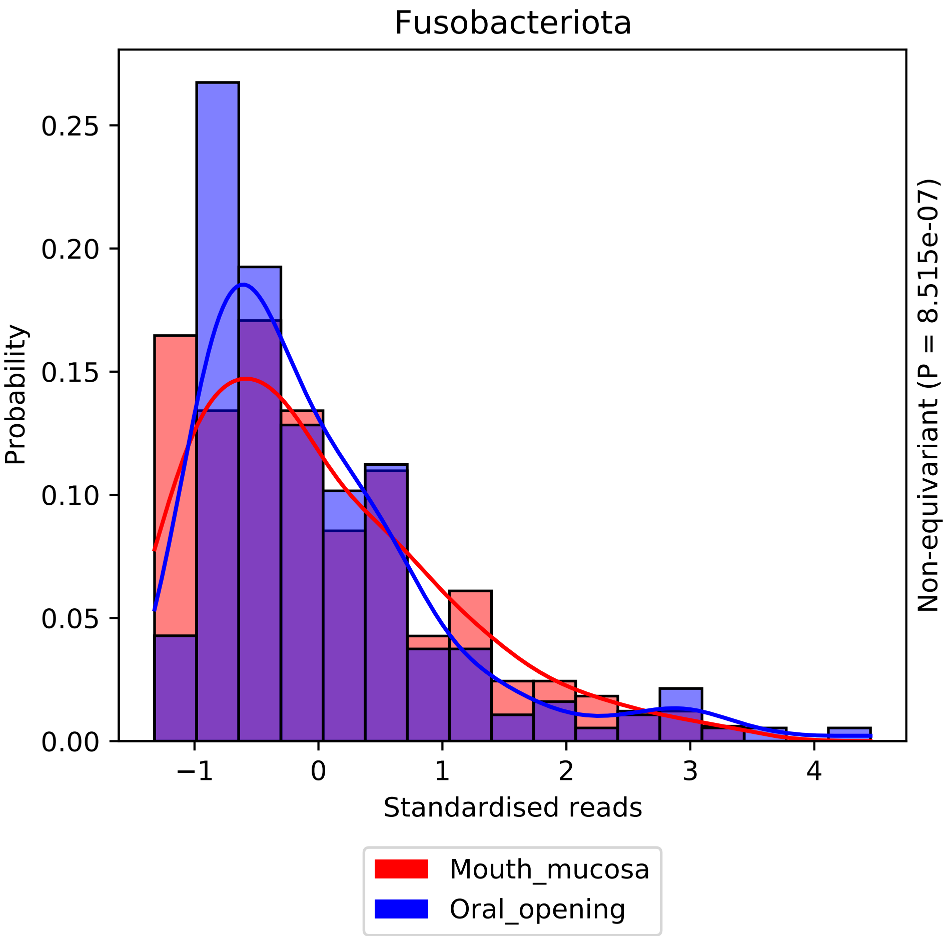

Fusobacteriota
Probability
0.00
0.05
0.10
0.15
0.20
0.25
−1
0
1
2
3
4
Standardised reads
Non-equivariant (P = 8.515e-07)
Mouth_mucosa
Oral_opening

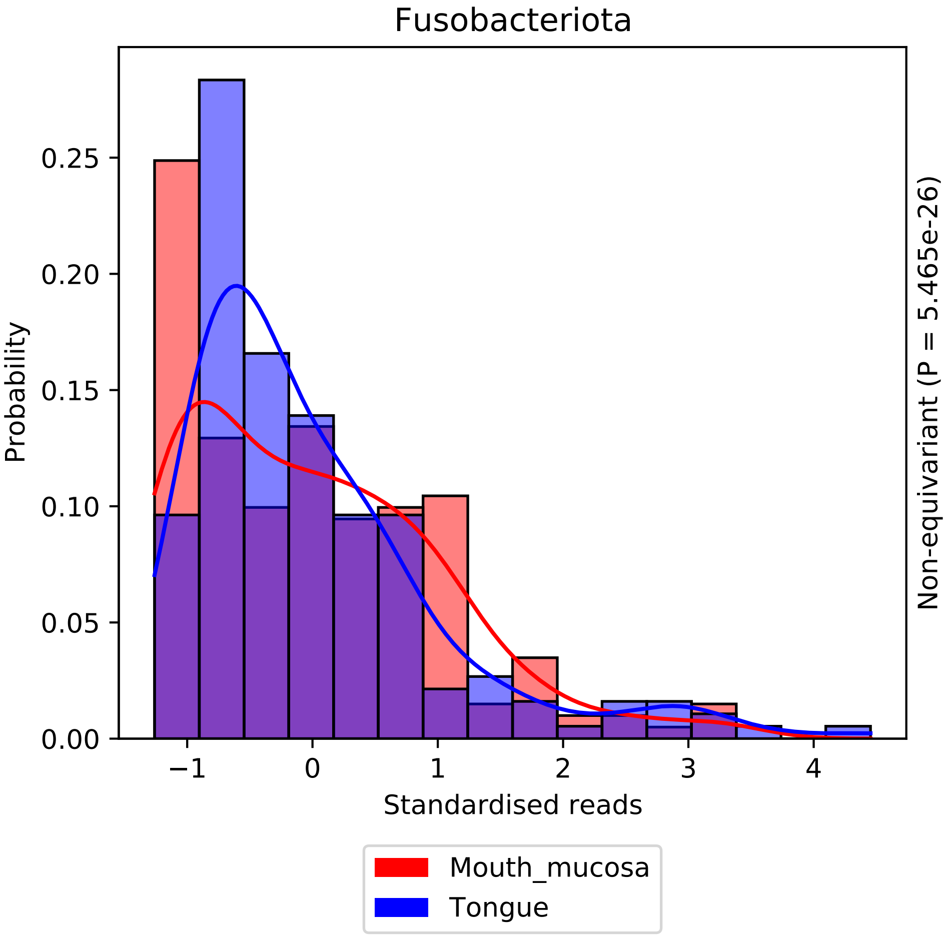
Fusobacteriota
Probability
0.25
0.20
0.15
0.10
0.05
0.00
−1
0
1
2
3
4
Standardised reads
Non-equivariant (P = 5.465e-26)
Mouth_mucosa
Tongue

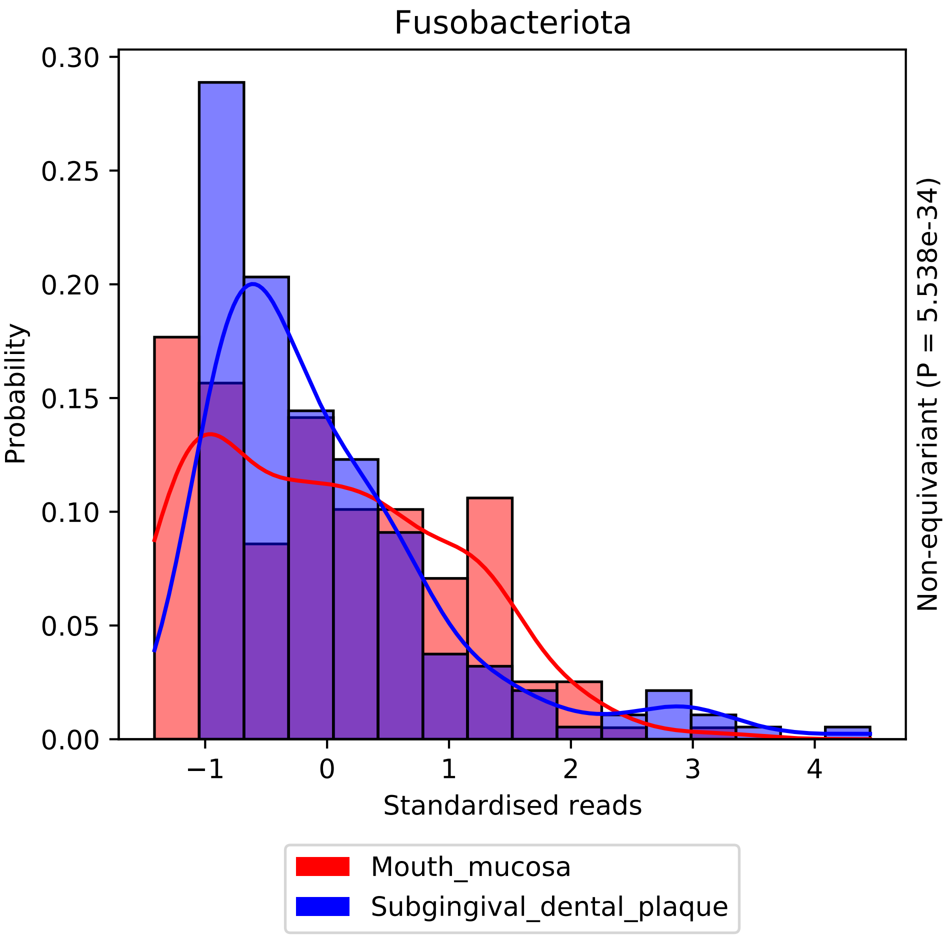
Fusobacteriota
Probability
0.00
0.05
0.10
0.15
0.20
0.25
0.30
−1
0
1
2
3
4
Standardised reads
Non-equivariant (P = 5.538e-34)
Mouth_mucosa
Subgingival_dental_plaque

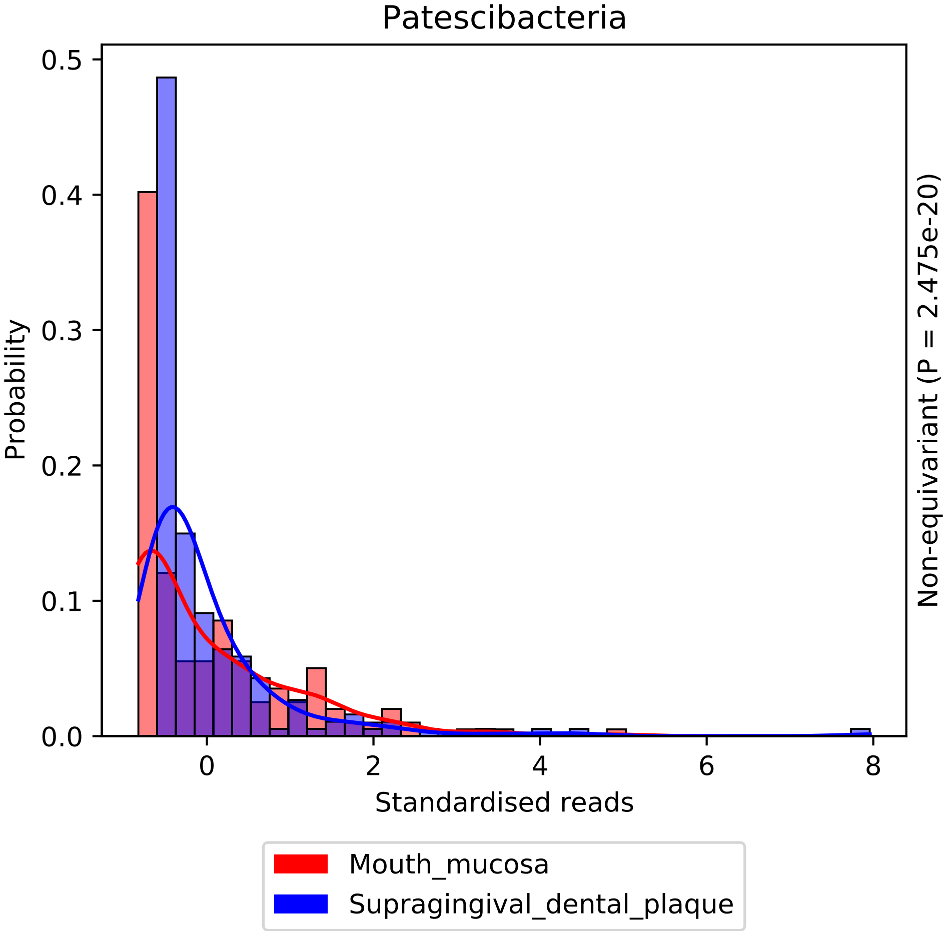
Patescibacteria
Probability
0.0
0.1
0.2
0.3
0.4
0.5
0
2
4
6
8
Standardised reads
Non-equivariant (P = 2.475e-20)
Mouth_mucosa
Supragingival_dental_plaque

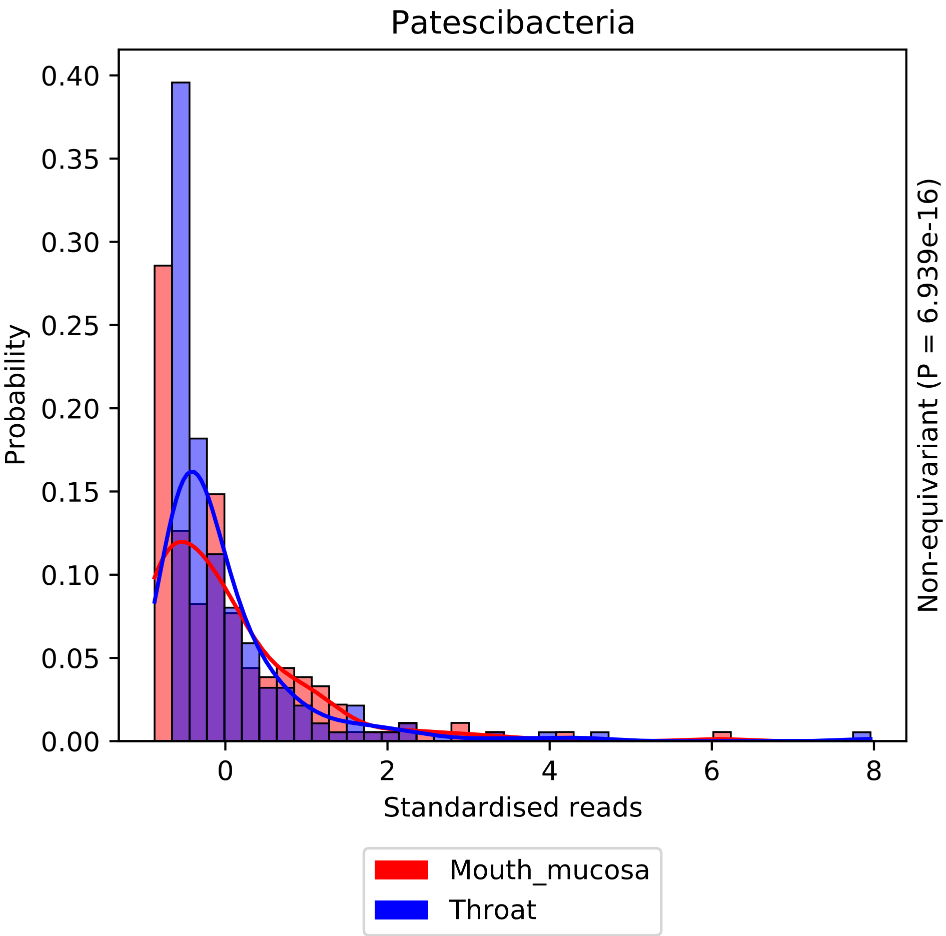

Patescibacteria
Probability
0.00
0.05
0.10
0.15
0.20
0.25
0.30
0.35
0.40
0
2
4
6
8
Standardised reads
Non-equivariant (P = 6.939e-16)
Mouth_mucosa
Throat

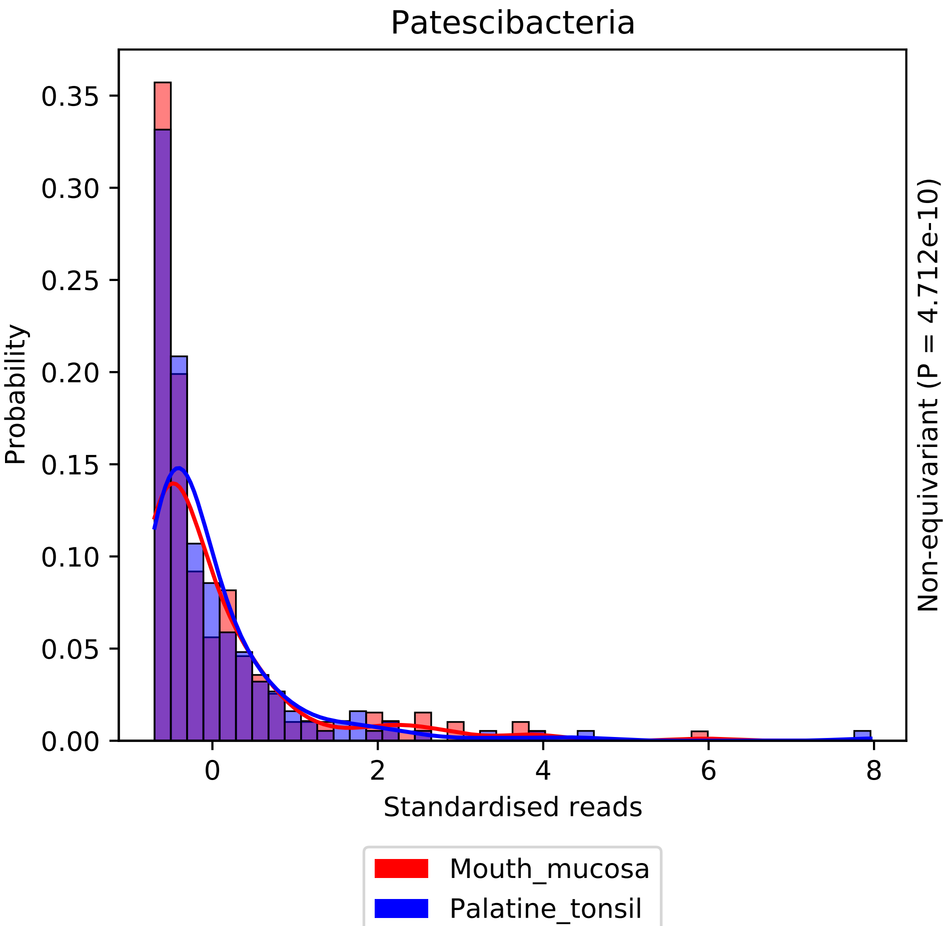
Patescibacteria
Probability
0.00
0.05
0.10
0.15
0.20
0.25
0.30
0.35
0
2
4
6
8
Standardised reads
Non-equivariant (P = 4.712e-10)
Mouth_mucosa
Palatine_tonsil

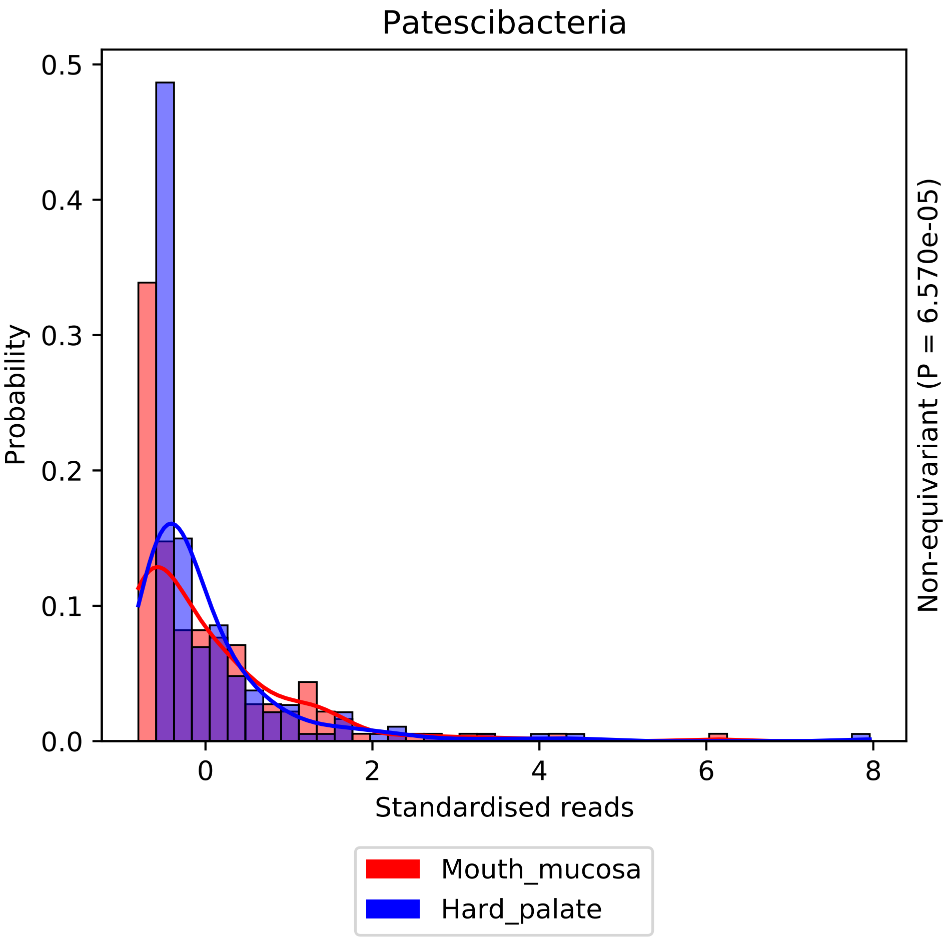

Patescibacteria
Probability
0.5
0.4
0.3
0.2
0.1
0.0
0
2
4
6
8
Standardised reads
Non-equivariant (P = 6.570e-05)
Mouth_mucosa
Hard_palate

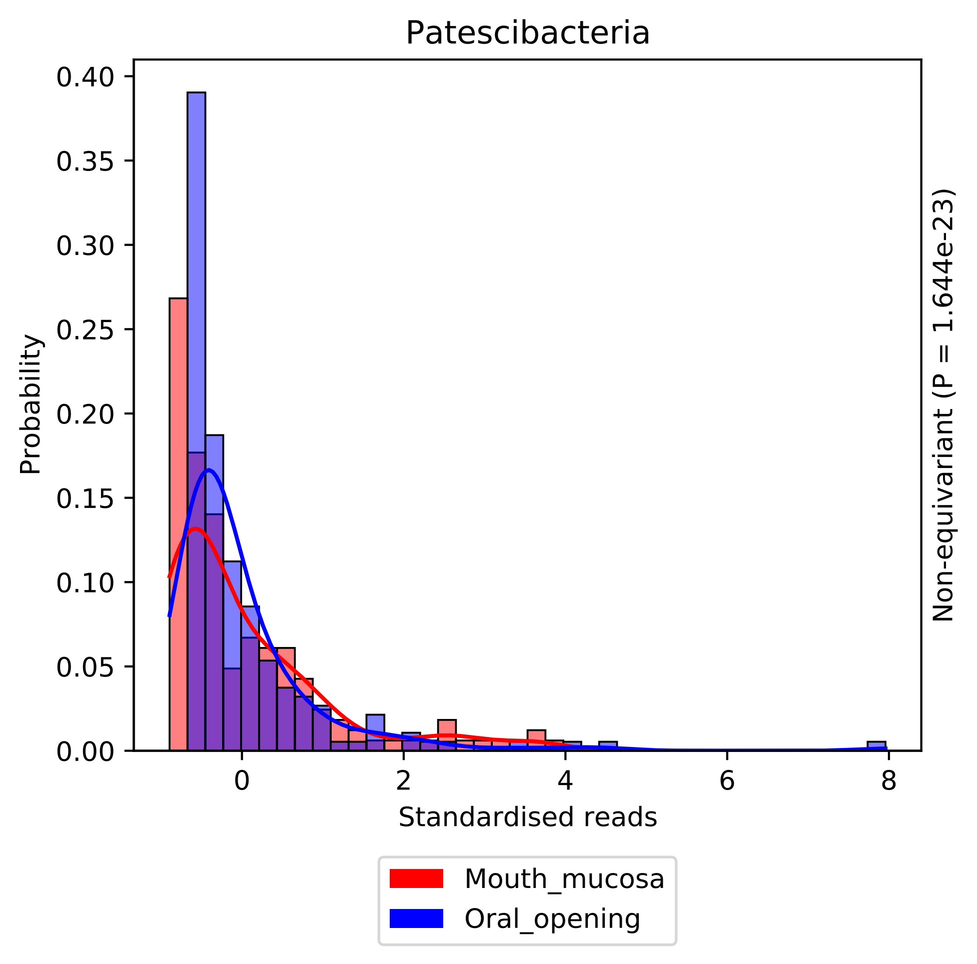

Patescibacteria
Probability
0.40
0.35
0.30
0.25
0.20
0.15
0.10
0.05
0.00
0
2
4
6
8
Standardised reads
Non-equivariant (P = 1.644e-23)
Mouth_mucosa
Oral_opening

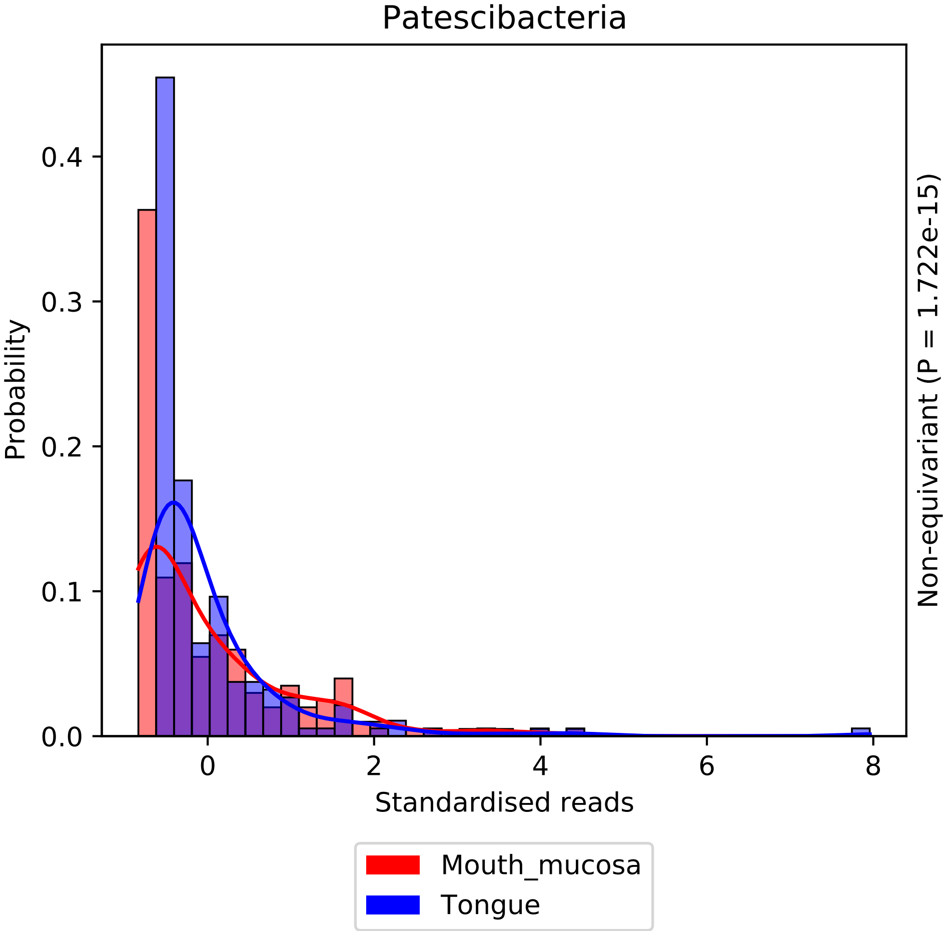

Patescibacteria
Probability
0.0
0.1
0.2
0.3
0.4
0
2
4
6
8
Standardised reads
Non-equivariant (P = 1.722e-15)
Mouth_mucosa
Tongue

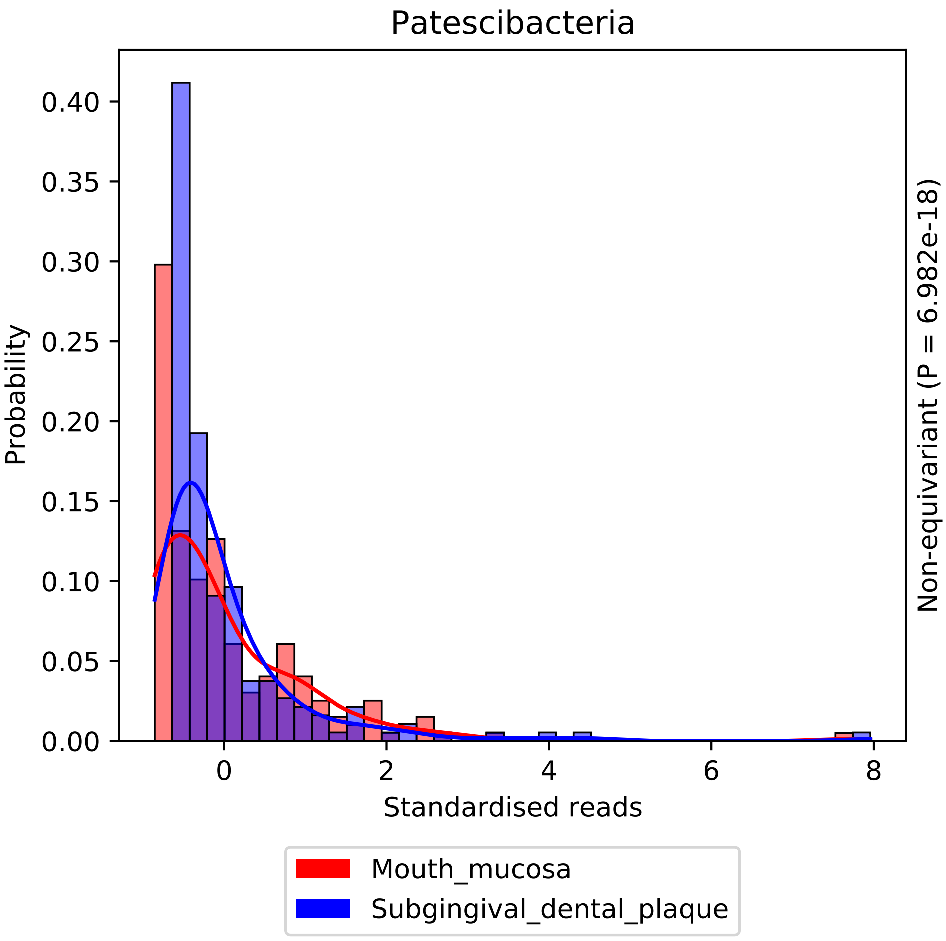

Patescibacteria
Probability
0.00
0.05
0.10
0.15
0.20
0.25
0.30
0.35
0.40
0
2
4
6
8
Standardised reads
Non-equivariant (P = 6.982e-18)
Mouth_mucosa
Subgingival_dental_plaque

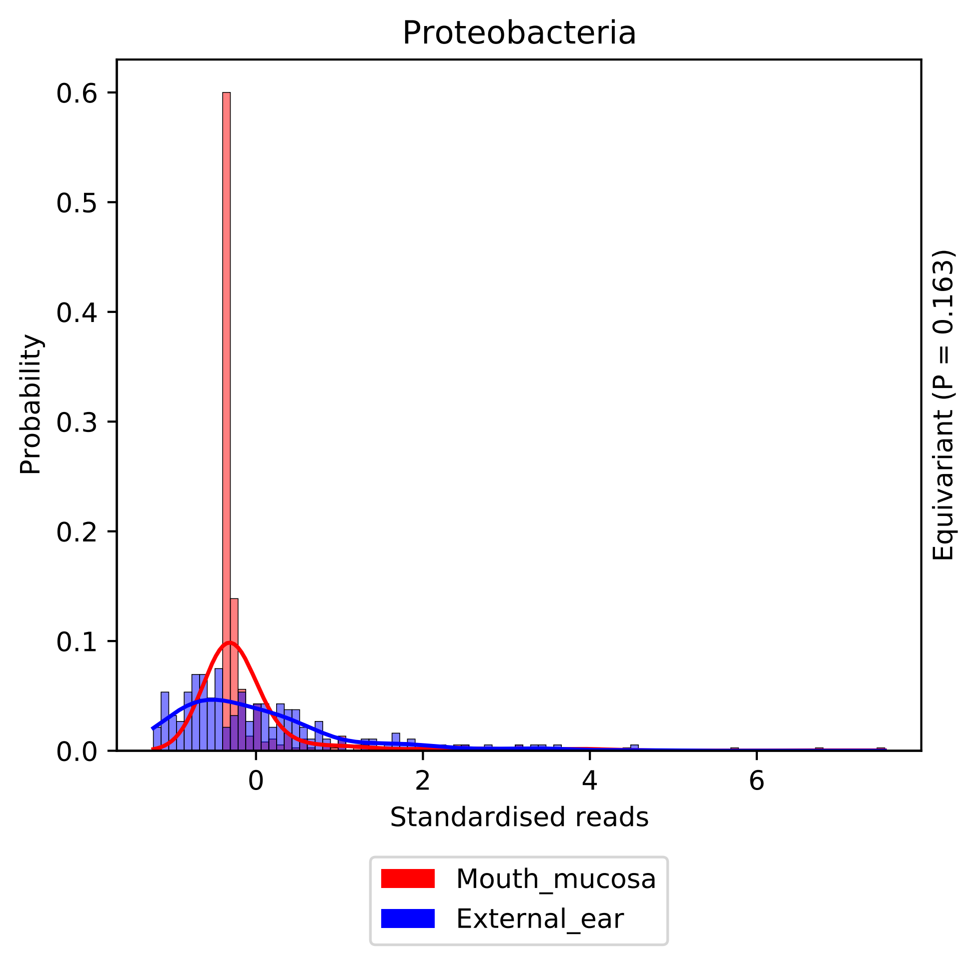
Proteobacteria
Probability
0.0
0.1
0.2
0.3
0.4
0.5
0.6
0
2
4
6
Standardised reads
Equivariant (P = 0.163)
Mouth_mucosa
External_ear

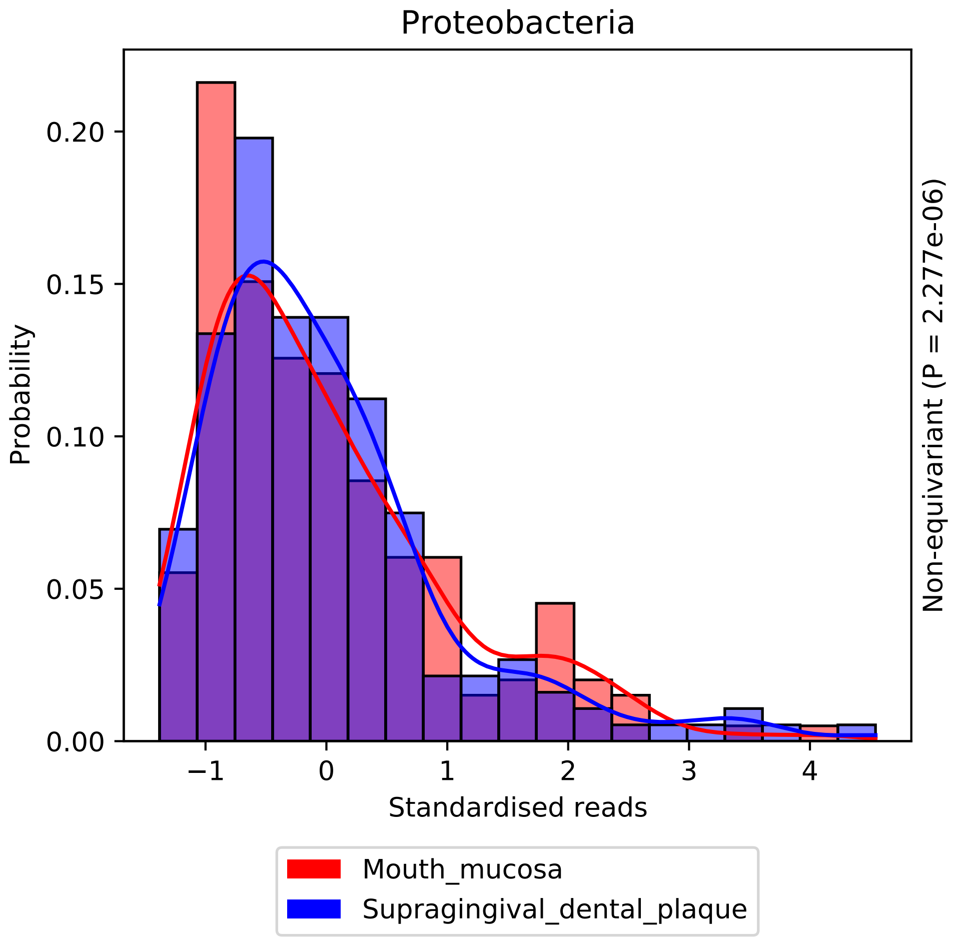
Proteobacteria
Probability
0.00
0.05
0.10
0.15
0.20
−1
0
1
2
3
4
Standardised reads
Non-equivariant (P = 2.277e-06)
Mouth_mucosa
Supragingival_dental_plaque

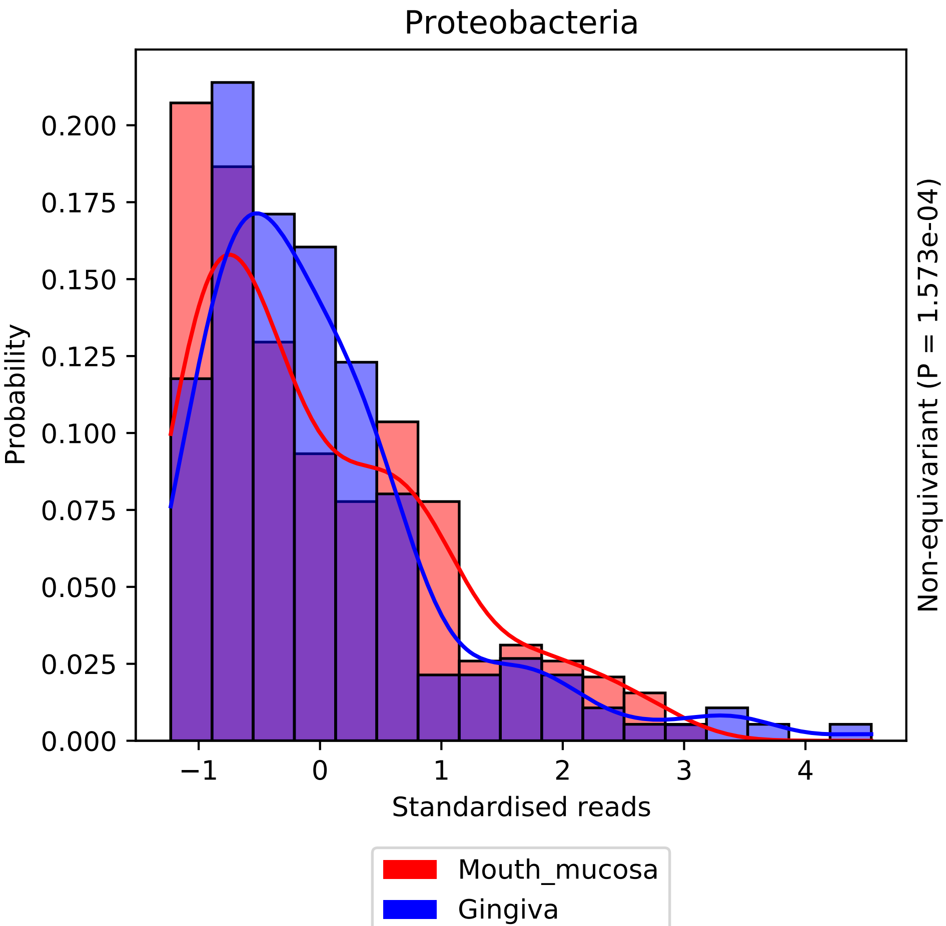

Proteobacteria
Probability
0.200
0.175
0.150
0.125
0.100
0.075
0.050
0.025
0.000
−1
0
1
2
3
4
Standardised reads
Non-equivariant (P = 1.573e-04)
Mouth_mucosa
Gingiva

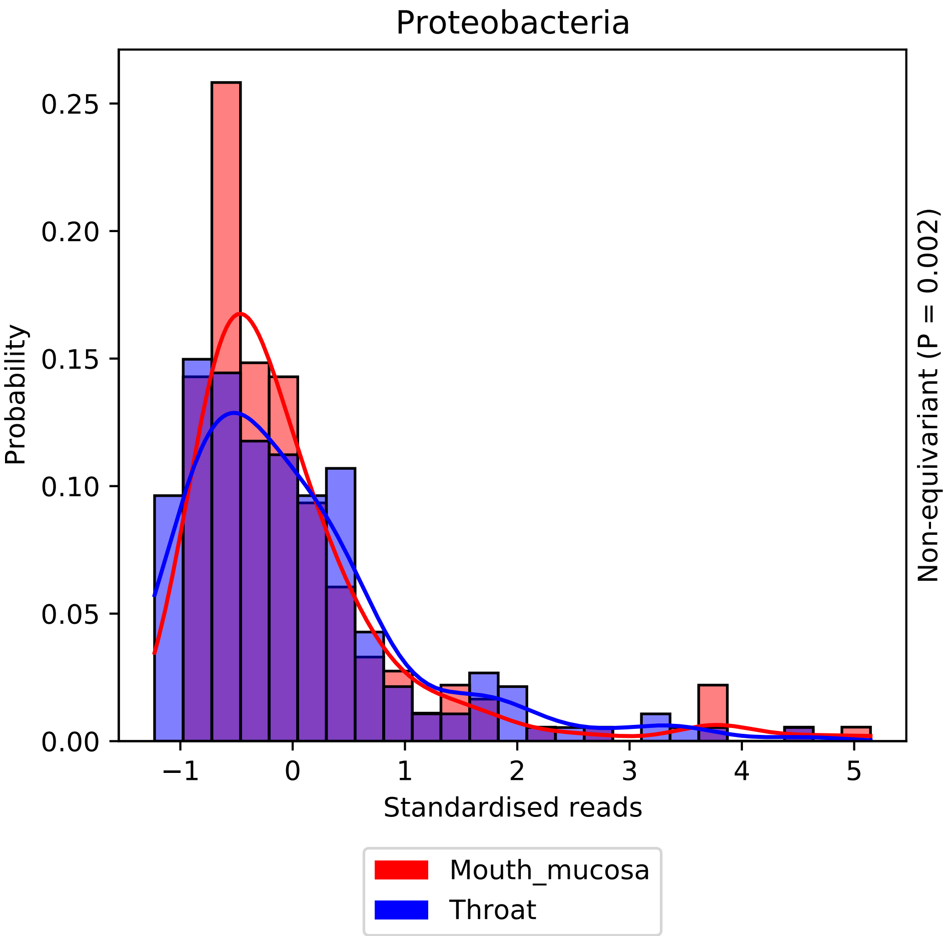

Proteobacteria
Probability
0.25
0.20
0.15
0.10
0.05
0.00
−1
0
1
2
3
4
5
Standardised reads
Non-equivariant (P = 0.002)
Mouth_mucosa
Throat

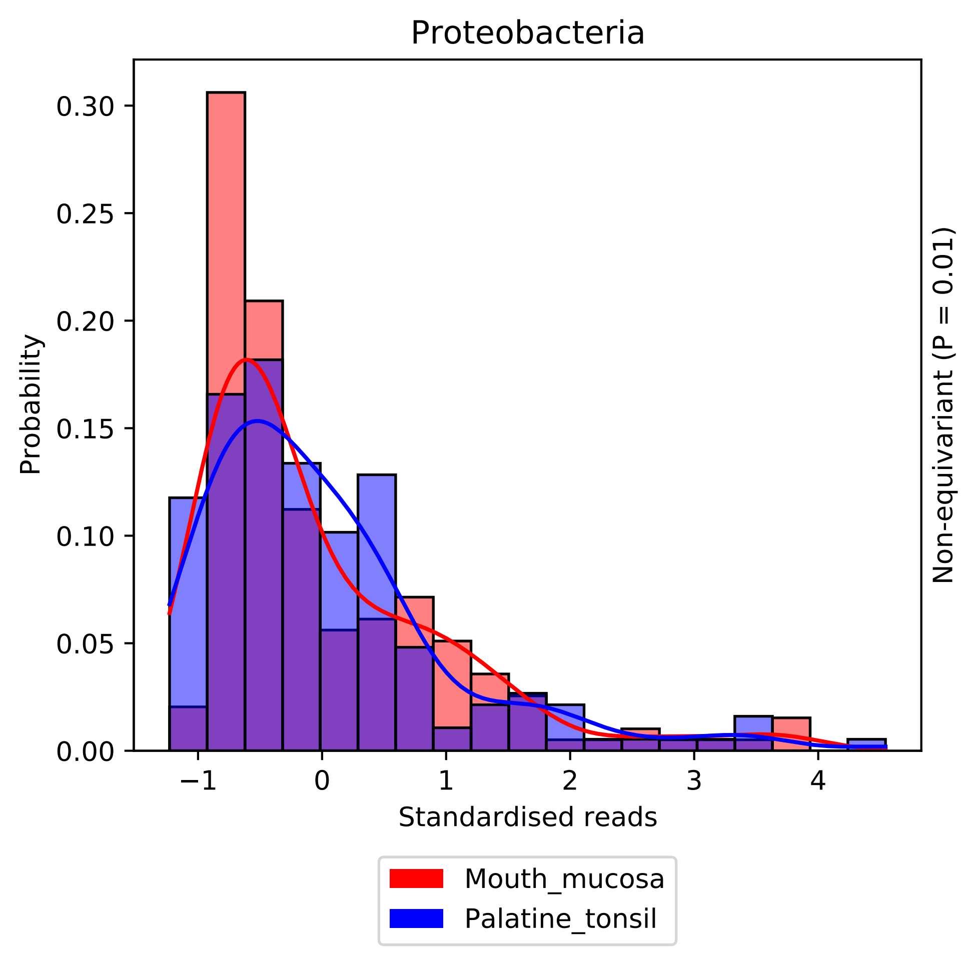
Proteobacteria
Probability
0.30
0.25
0.20
0.15
0.10
0.05
0.00
−1
0
1
2
3
4
Standardised reads
Non-equivariant (P = 0.01)
Mouth_mucosa
Palatine_tonsil

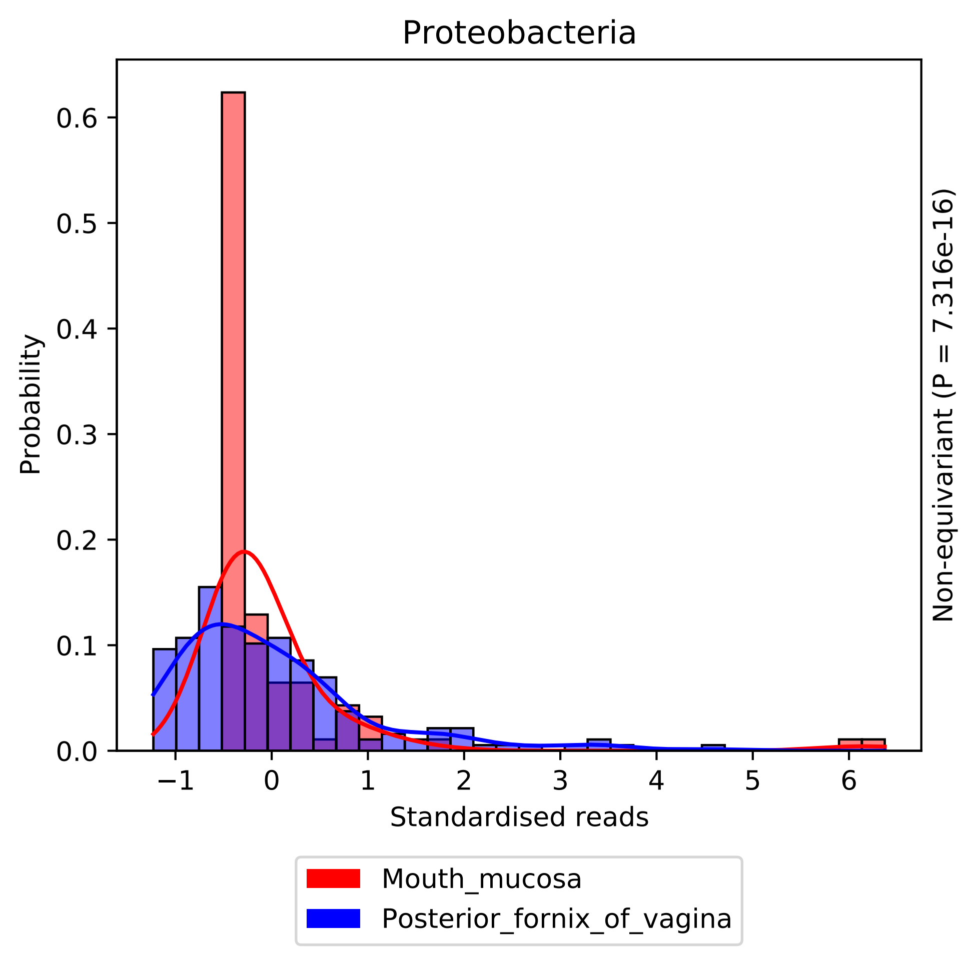

Proteobacteria
Probability
0.0
0.1
0.2
0.3
0.4
0.5
0.6
−1
0
1
2
3
4
5
6
Standardised reads
Non-equivariant (P = 7.316e-16)
Mouth_mucosa
Posterior_fornix_of_vagina

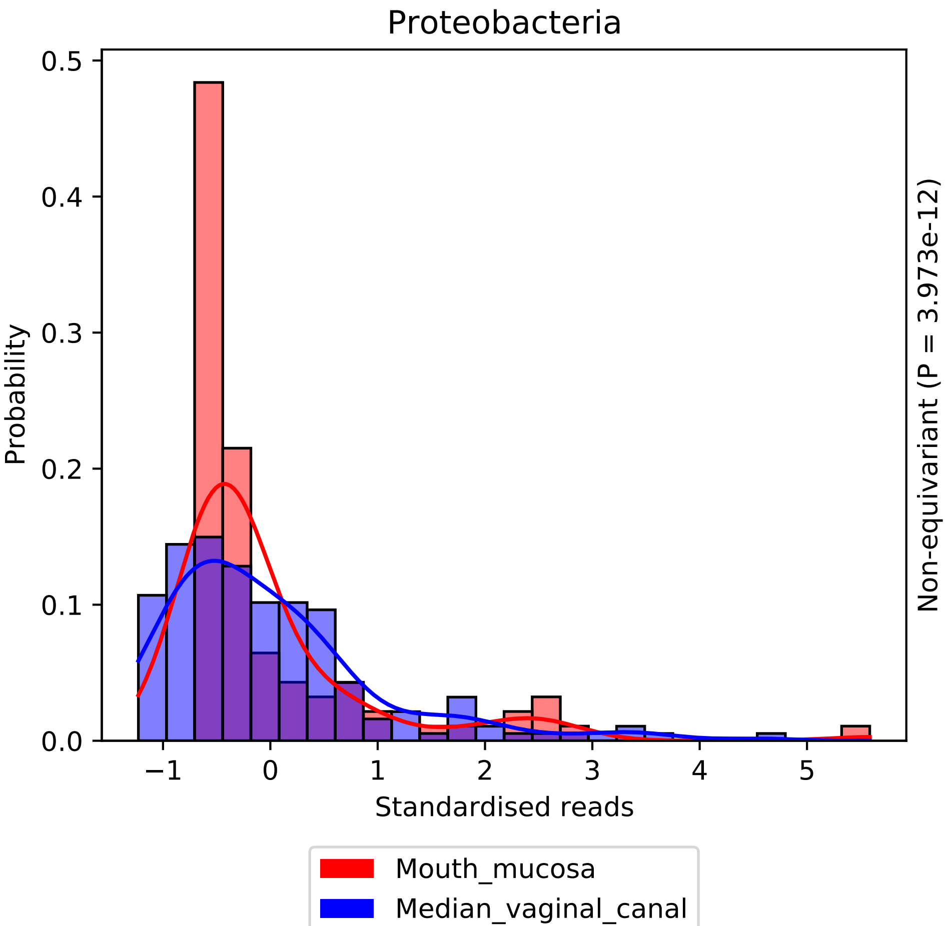

Proteobacteria
Probability
0.0
0.1
0.2
0.3
0.4
0.5
−1
0
1
2
3
4
5
Standardised reads
Non-equivariant (P = 3.973e-12)
Mouth_mucosa
Median_vaginal_canal

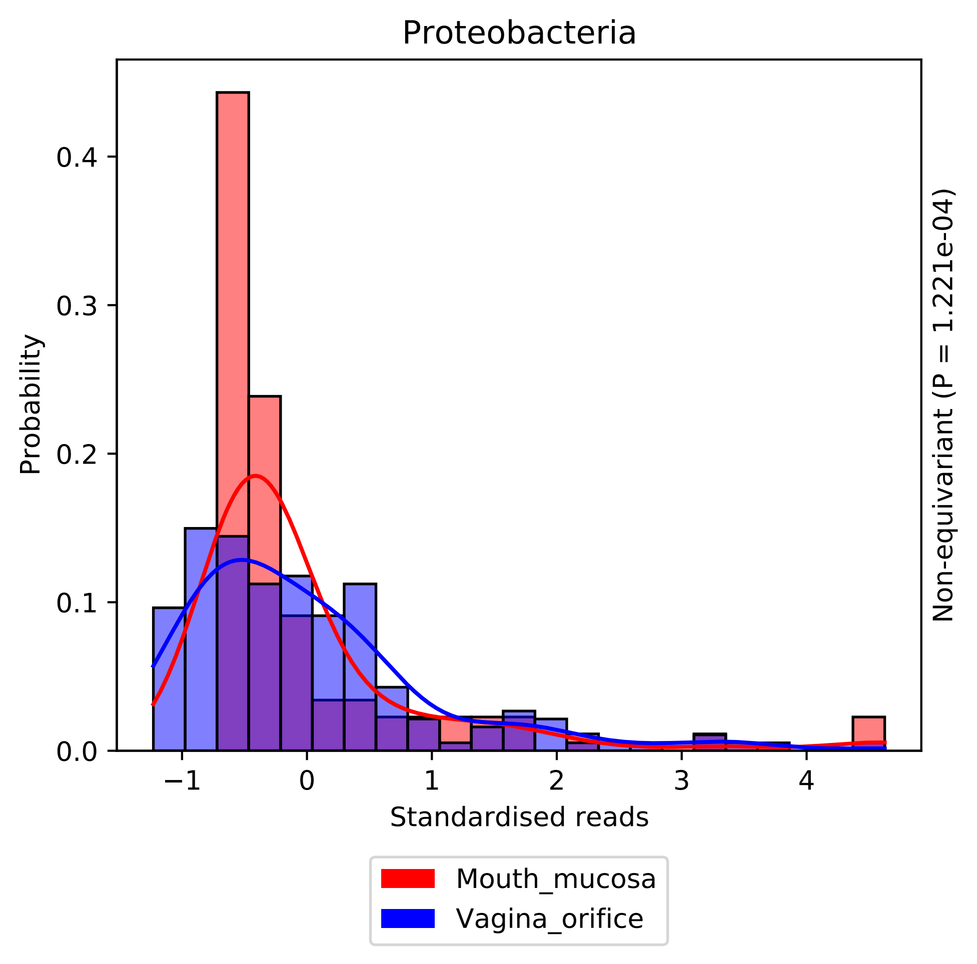

Proteobacteria
Probability
0.0
0.1
0.2
0.3
0.4
−1
0
1
2
3
4
Standardised reads
Non-equivariant (P = 1.221e-04)
Mouth_mucosa
Vagina_orifice

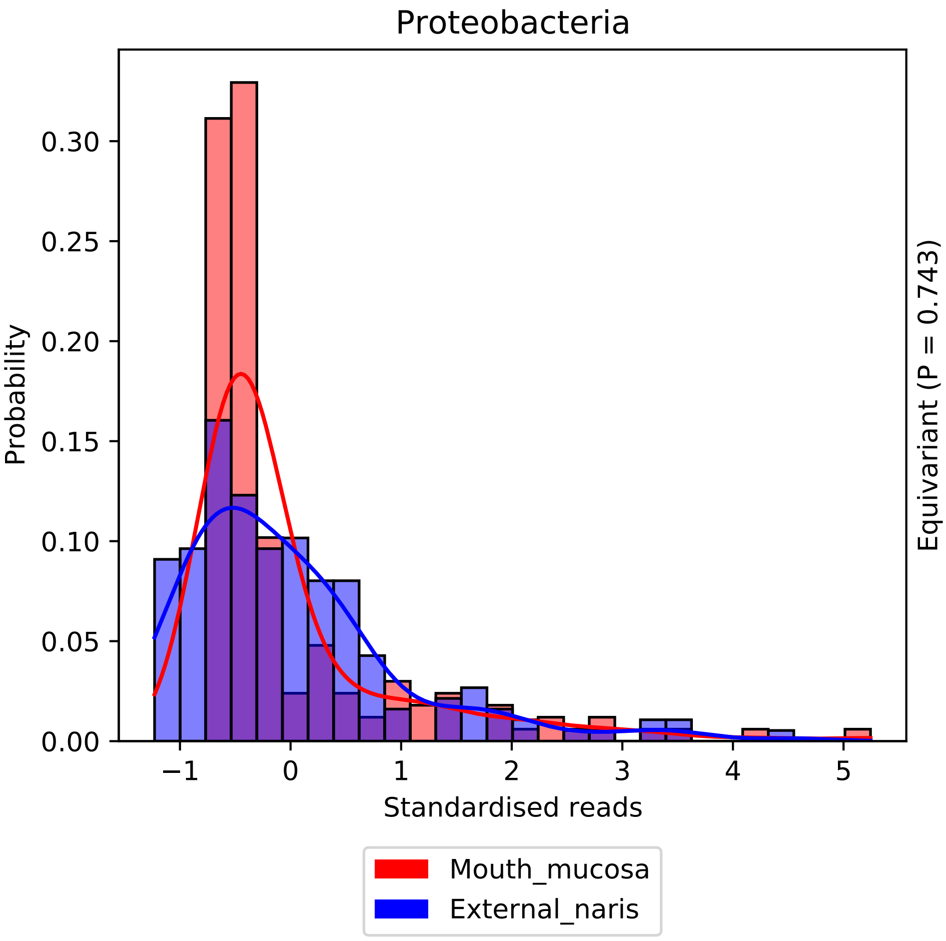
Proteobacteria
0.30
0.25
0.20
0.15
0.10
0.05
0.00
Probability
−1
0
1
2
3
4
5
Standardised reads
Equivariant (P = 0.743)
Mouth_mucosa
External_naris

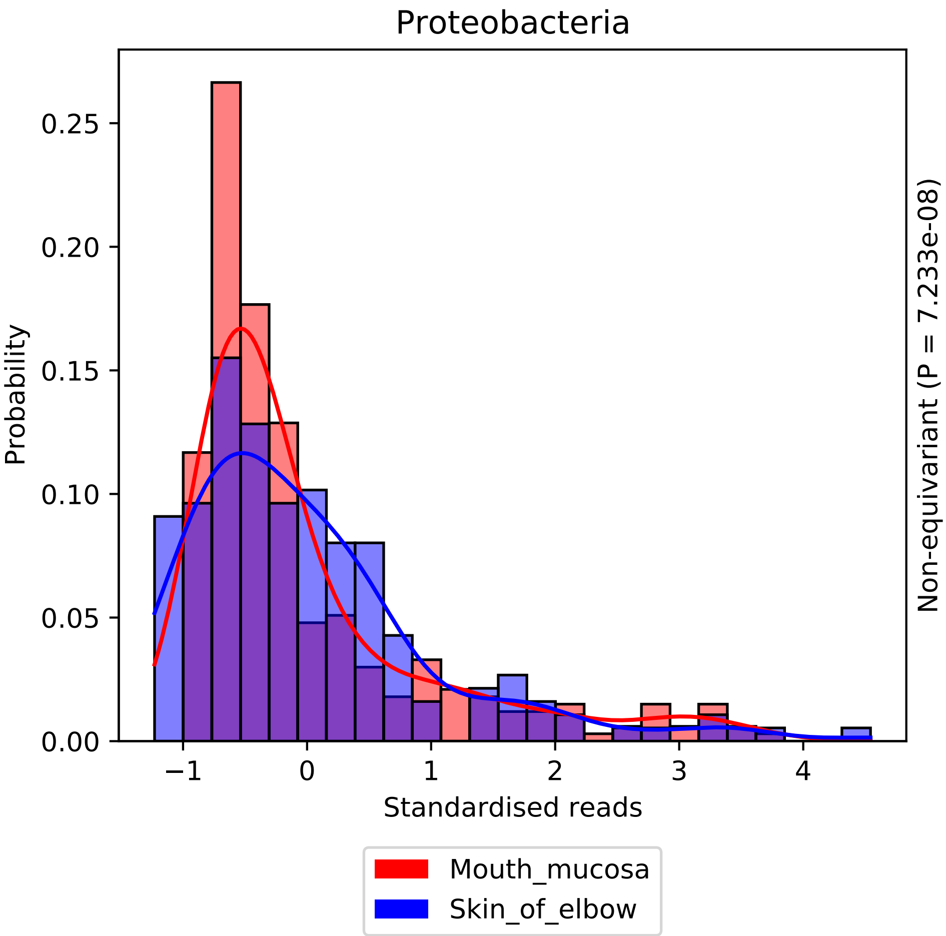
Proteobacteria
Probability
0.00
0.05
0.10
0.15
0.20
0.25
−1
0
1
2
3
4
Standardised reads
Non-equivariant (P = 7.233e-08)
Mouth_mucosa
Skin_of_elbow

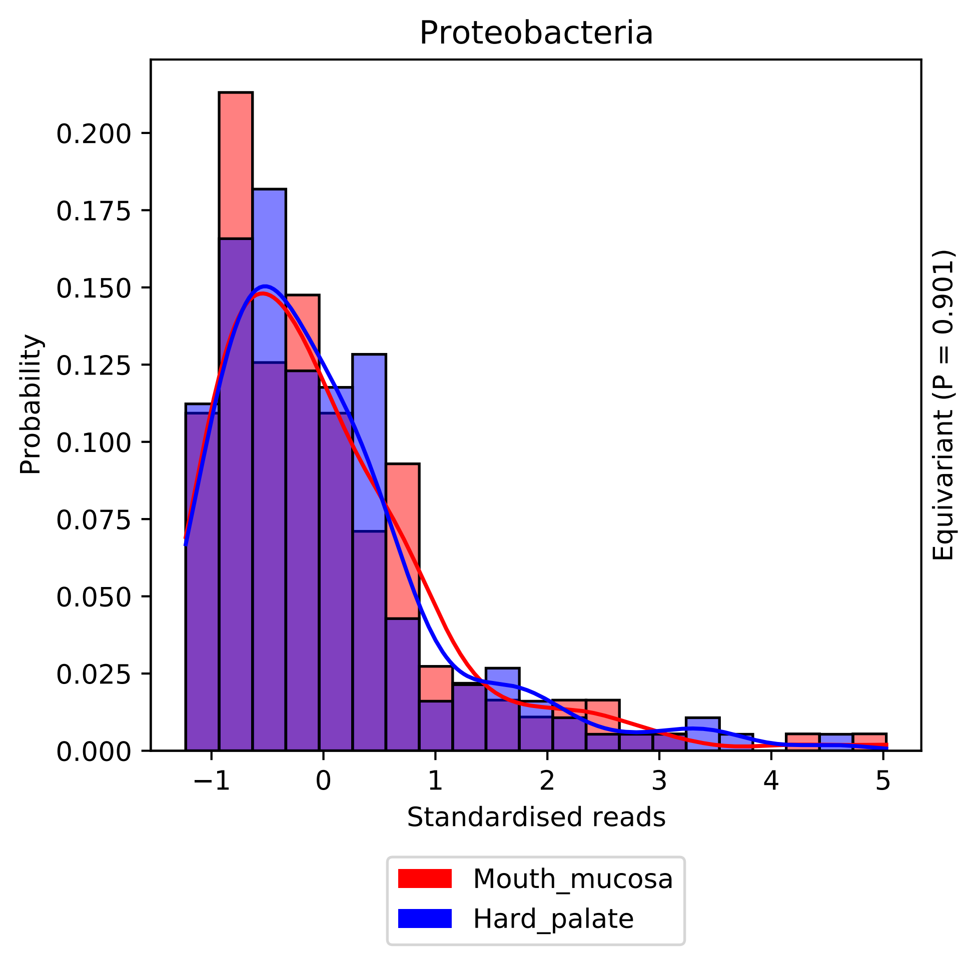
Proteobacteria
Probability
0.000
0.025
0.050
0.075
0.100
0.125
0.150
0.175
0.200
−1
0
1
2
3
4
5
Standardised reads
Equivariant (P = 0.901)
Mouth_mucosa
Hard_palate

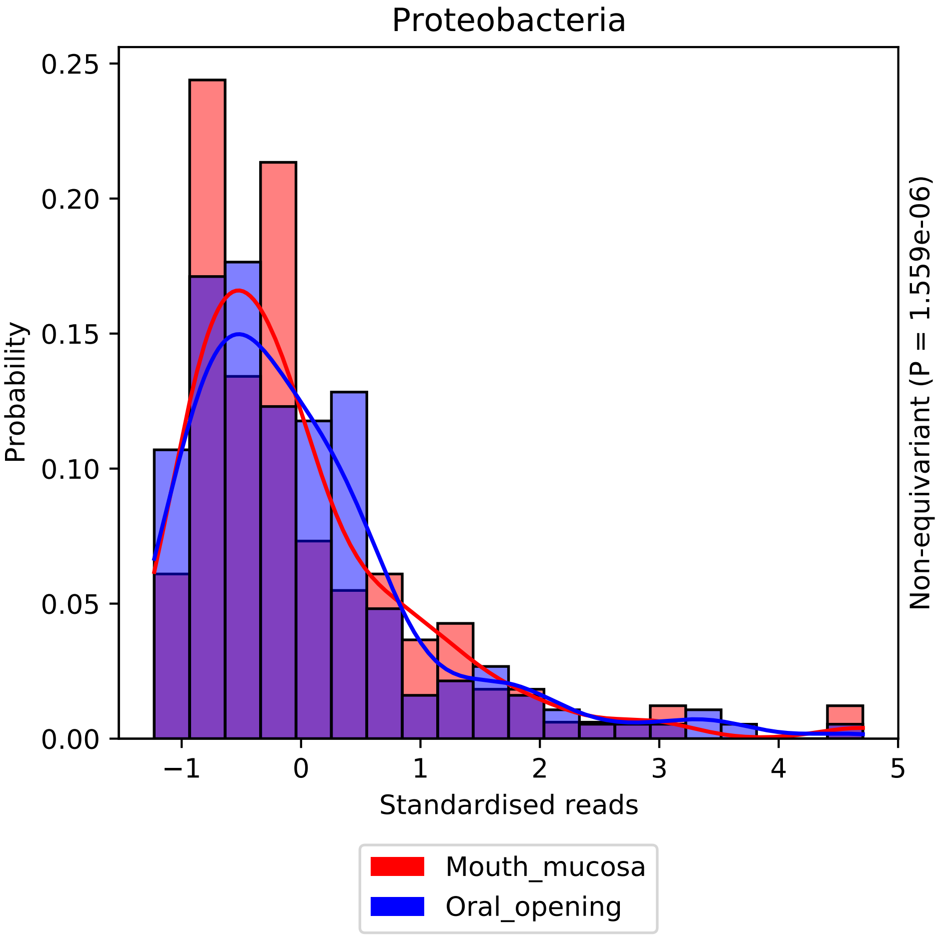

Proteobacteria
Probability
0.25
0.20
0.15
0.10
0.05
0.00
−1
0
1
2
3
4
5
Standardised reads
Non-equivariant (P = 1.559e-06)
Mouth_mucosa
Oral_opening

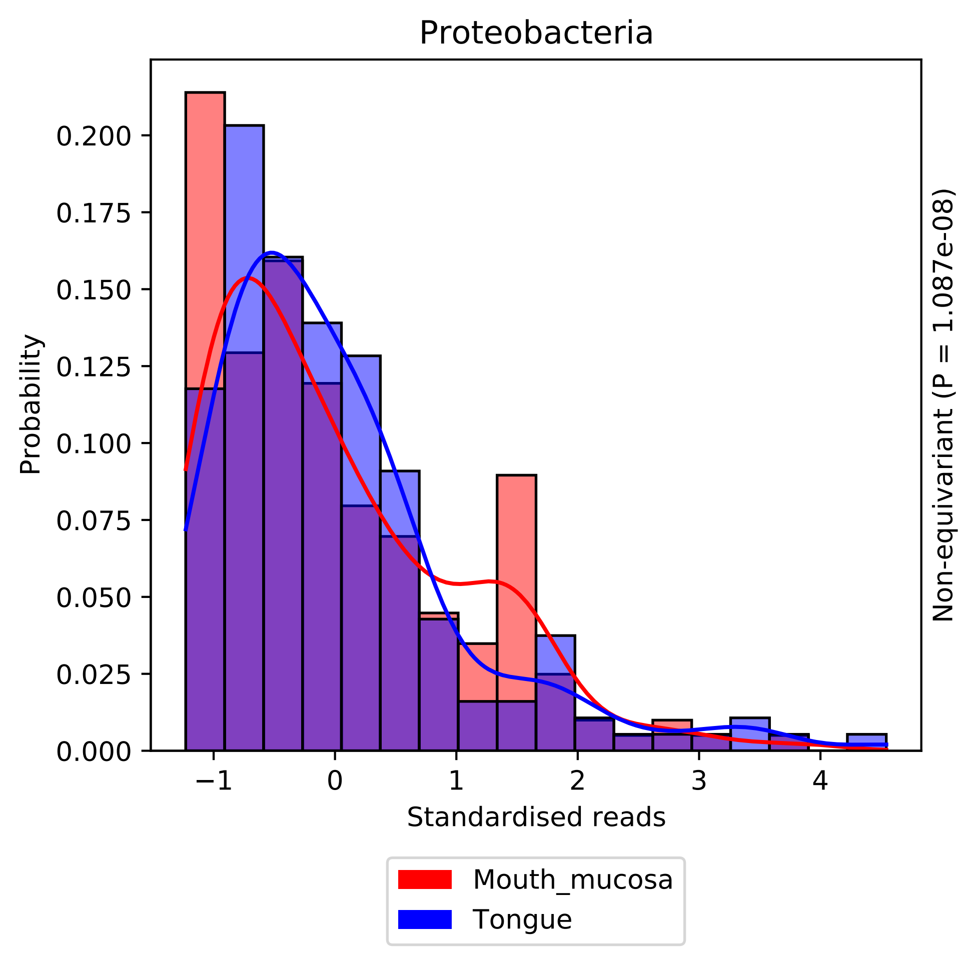

Proteobacteria
Probability
Standardised reads
Non-equivariant (P = 1.087e-08)
Mouth_mucosa
Tongue
0.000
0.025
0.050
0.075
0.100
0.125
0.150
0.175
0.200
−1
0
1
2
3
4

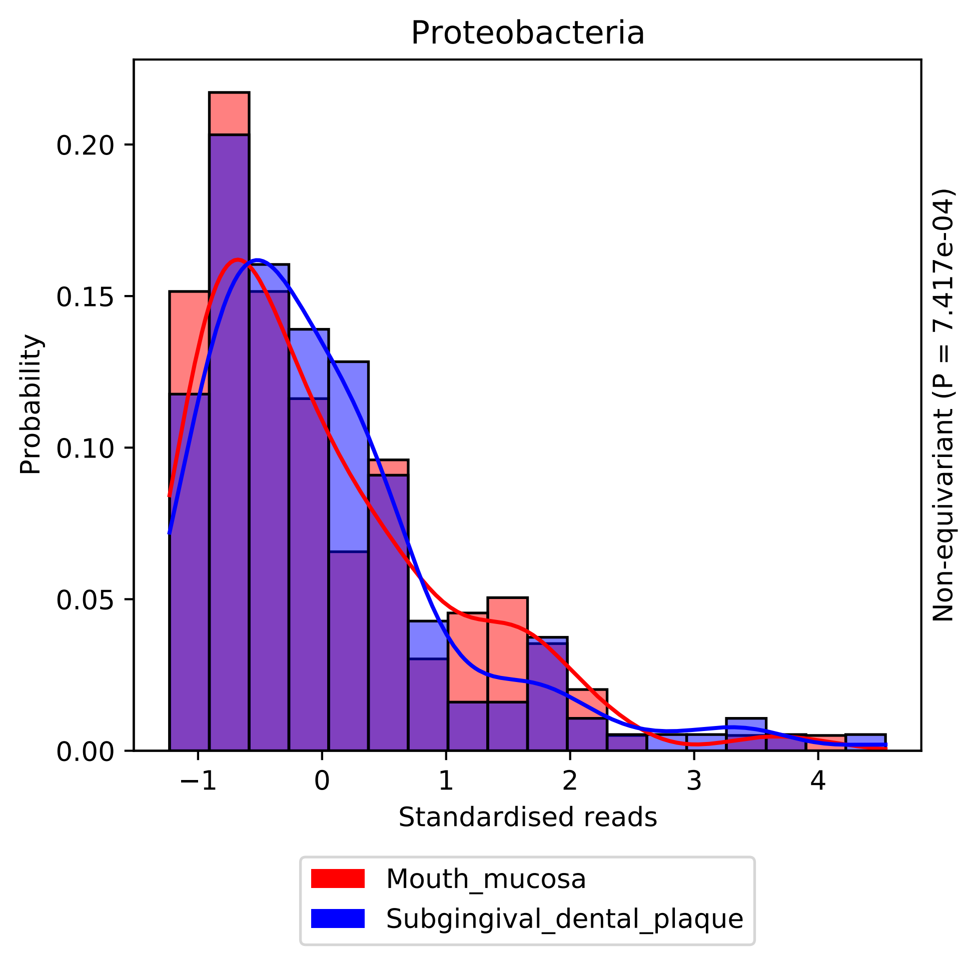

Proteobacteria
Probability
0.00
0.05
0.10
0.15
0.20
−1
0
1
2
3
4
Standardised reads
Non-equivariant (P = 7.417e-04)
Mouth_mucosa
Subgingival_dental_plaque

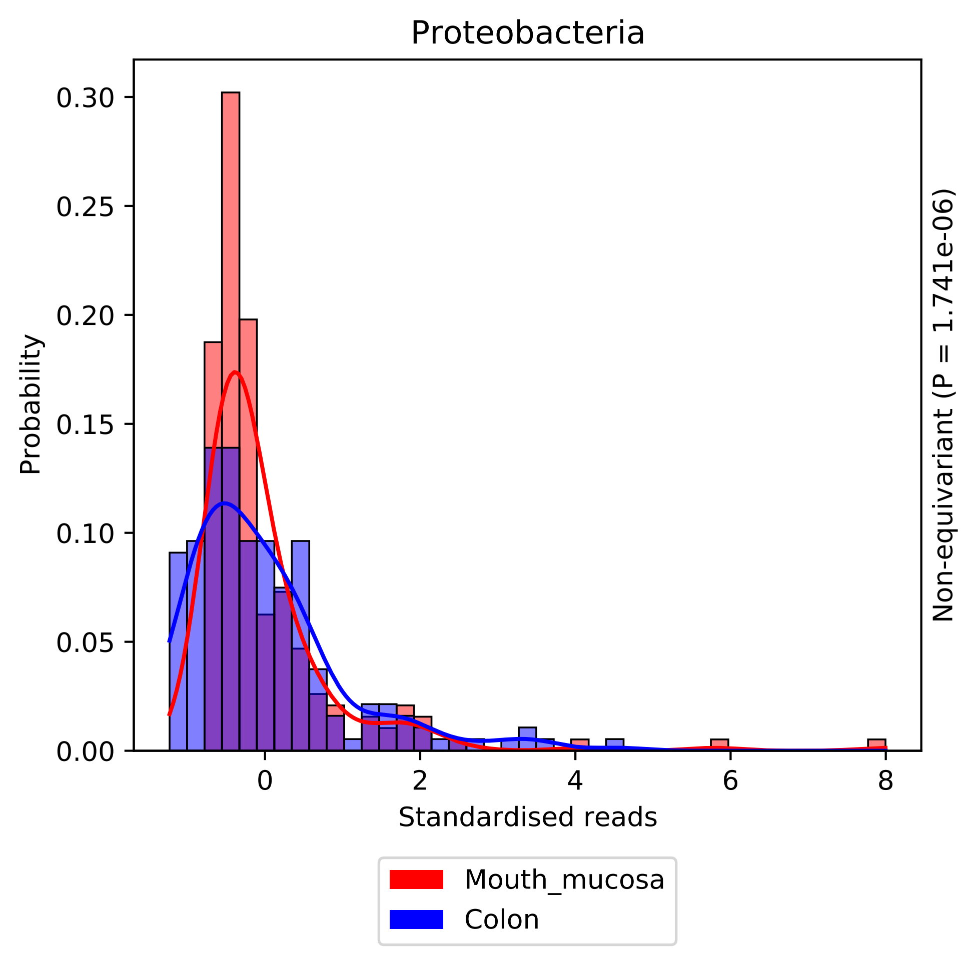

Proteobacteria
0.30
0.25
0.20
0.15
0.10
0.05
0.00
Probability
0
2
4
6
8
Standardised reads
Non-equivariant (P = 1.741e-06)
Mouth_mucosa
Colon

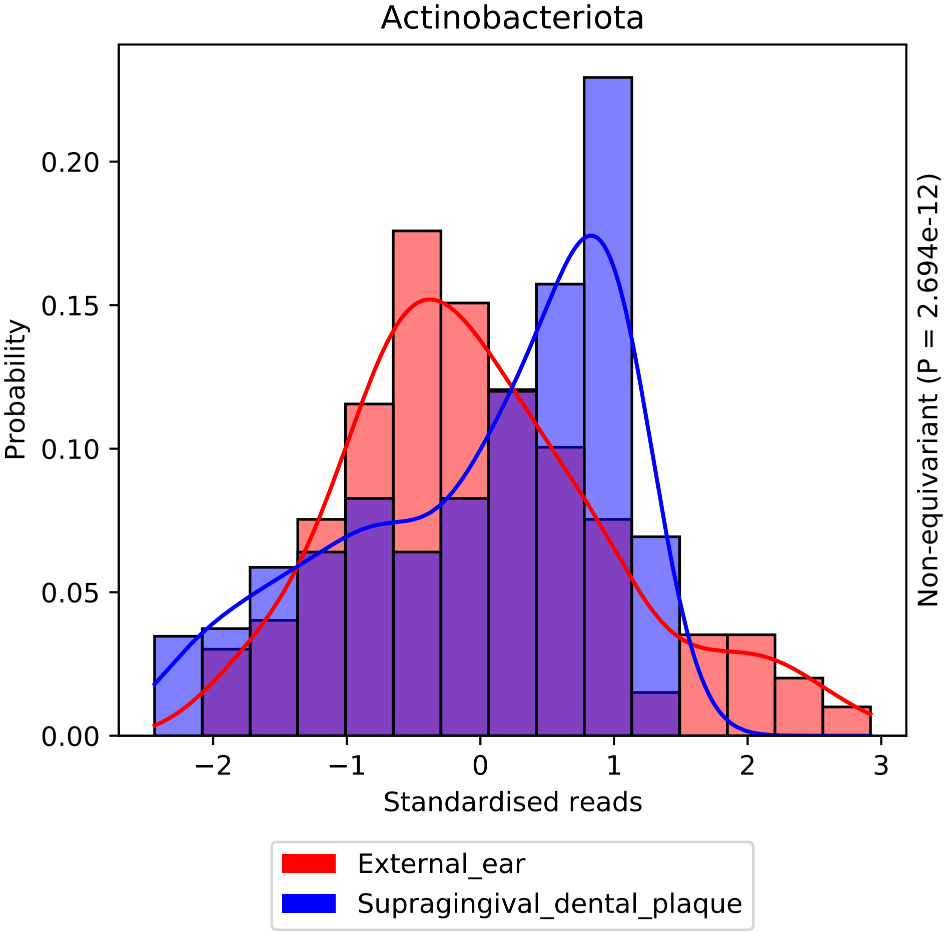

Actinobacteriota
Probability
0.00
0.05
0.10
0.15
0.20
−2
−1
0
1
2
3
Standardised reads
Non-equivariant (P = 2.694e-12)
External_ear
Supragingival_dental_plaque

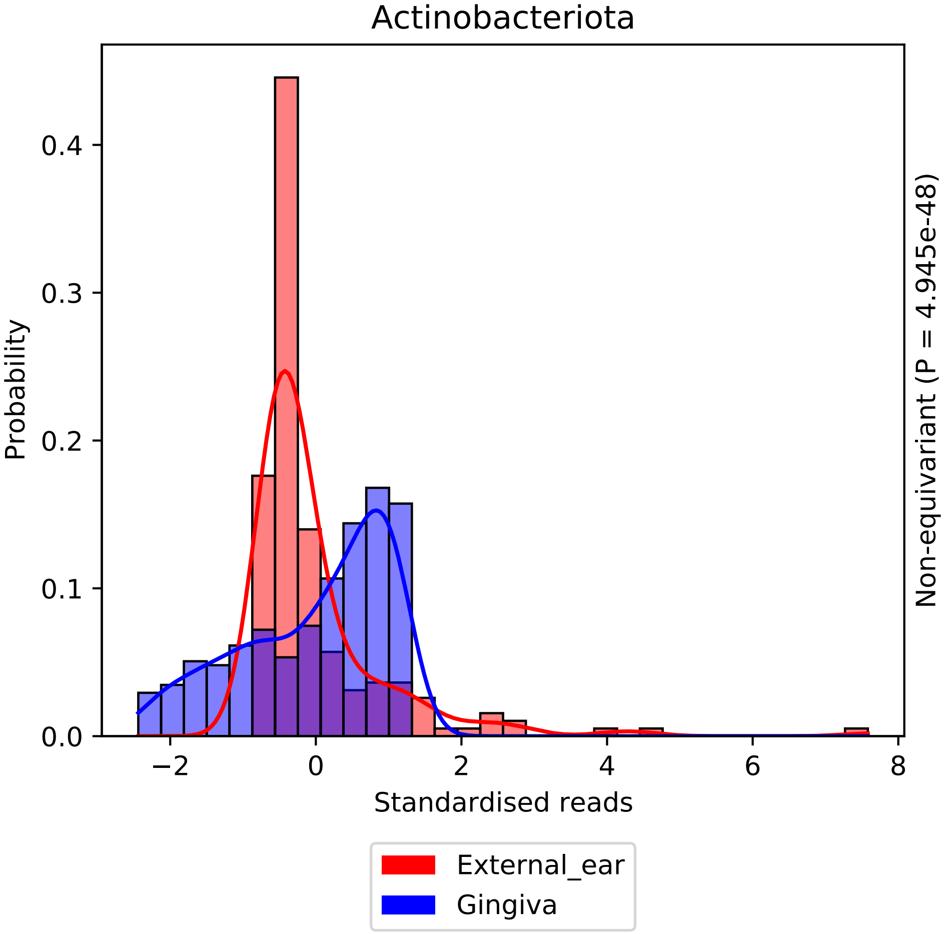

Actinobacteriota
Probability
0.0
0.1
0.2
0.3
0.4
−2
0
2
4
6
8
Standardised reads
Non-equivariant (P = 4.945e-48)
External_ear
Gingiva

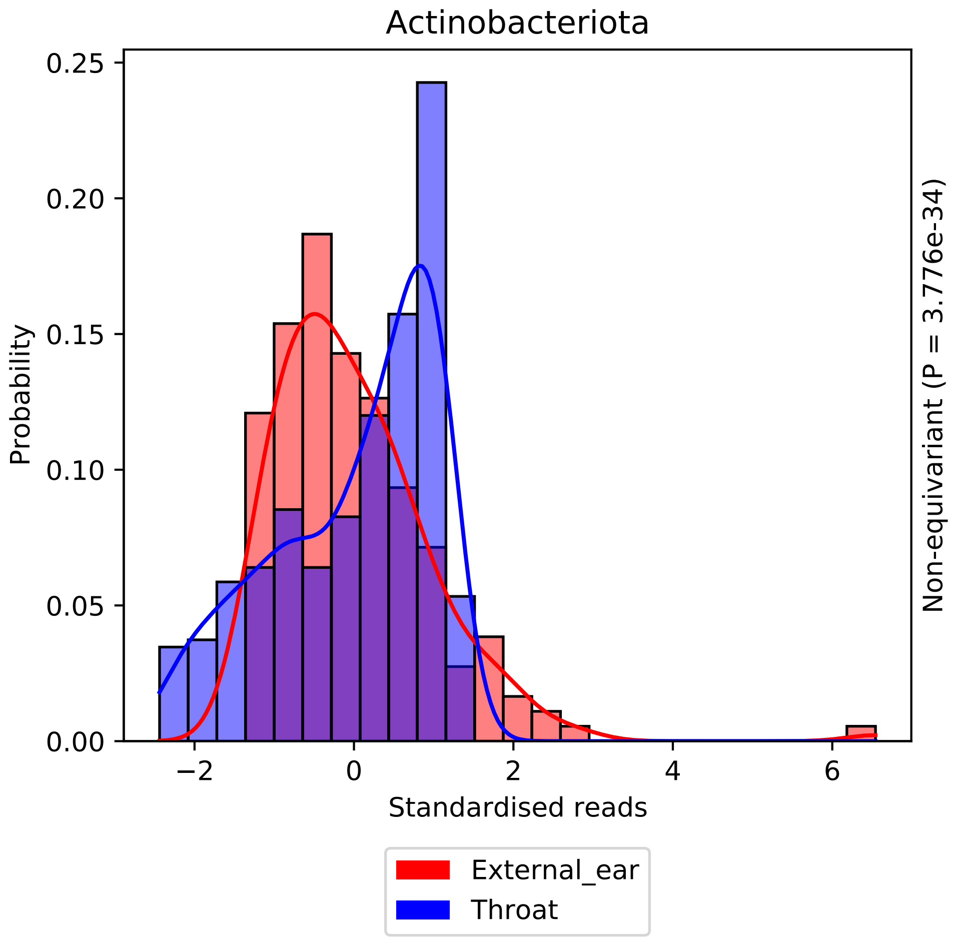
Actinobacteriota
Probability
0.25
0.20
0.15
0.10
0.05
0.00
−2
0
2
4
6
Standardised reads
Non-equivariant (P = 3.776e-34)
External_ear
Throat

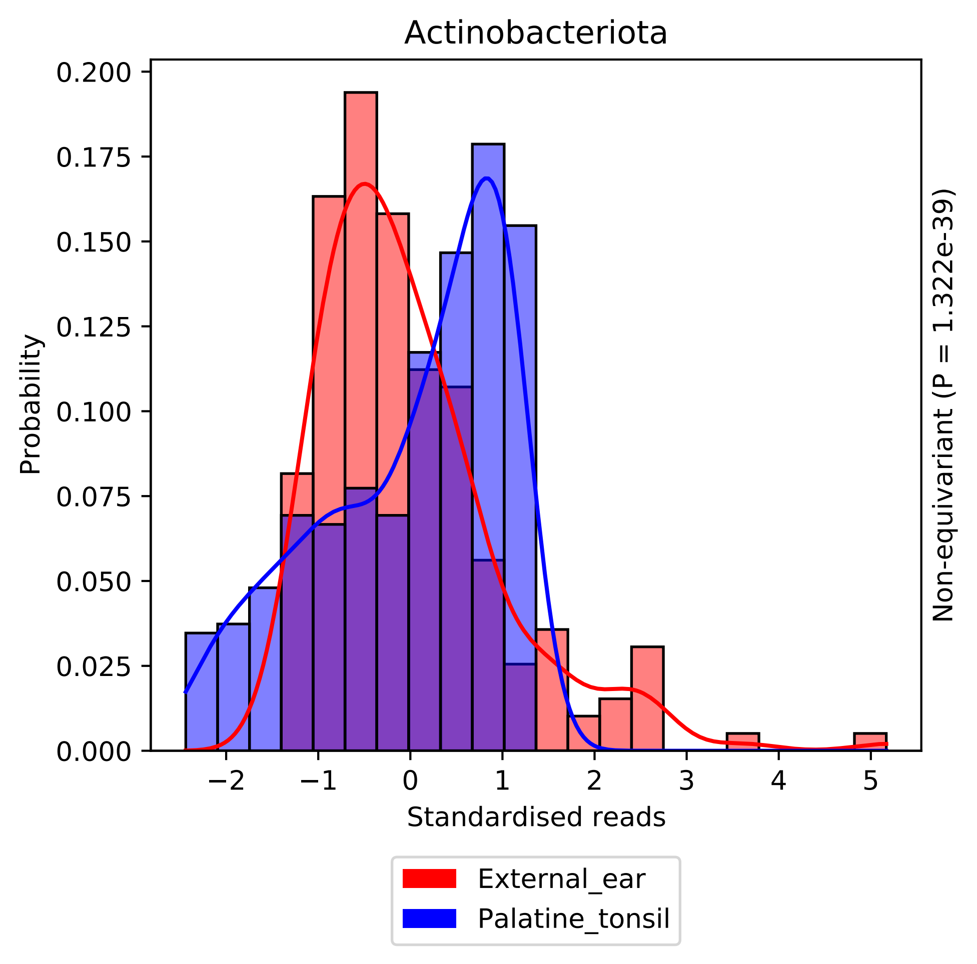

Actinobacteriota
Probability
0.200
0.175
0.150
0.125
0.100
0.075
0.050
0.025
0.000
−2
−1
0
1
2
3
4
5
Standardised reads
Non-equivariant (P = 1.322e-39)
External_ear
Palatine_tonsil

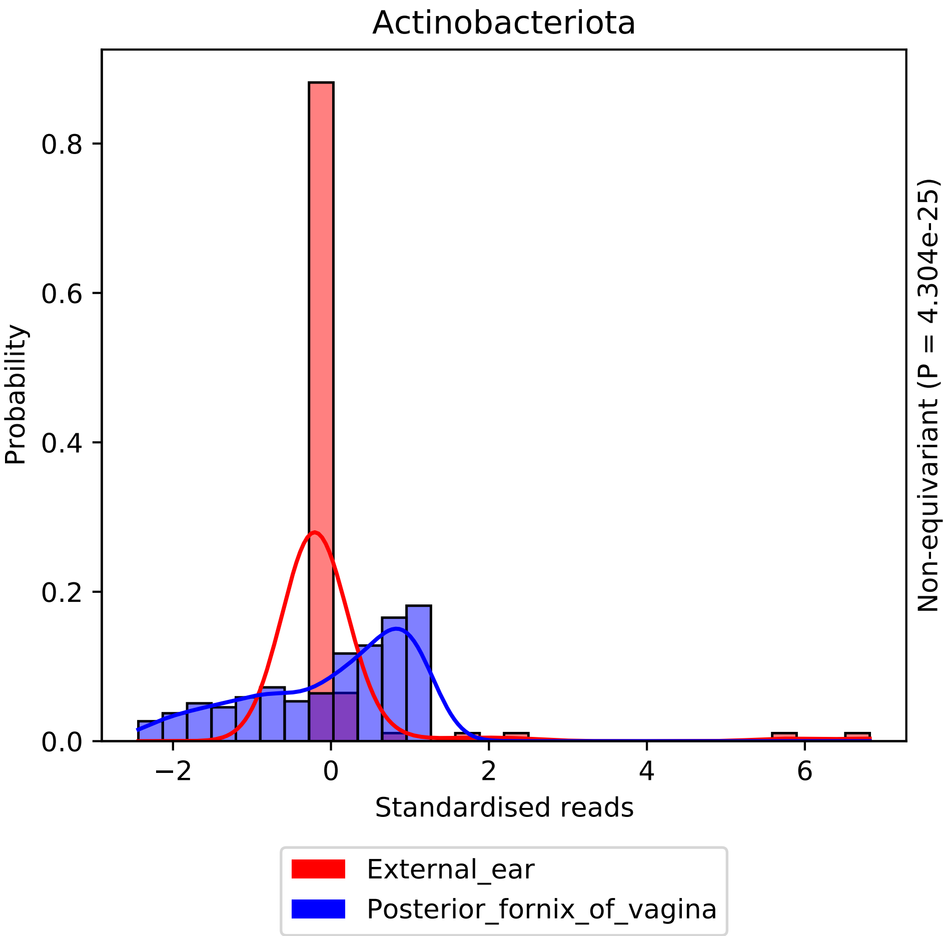

Actinobacteriota
Probability
0.0
0.2
0.4
0.6
0.8
−2
0
2
4
6
Standardised reads
Non-equivariant (P = 4.304e-25)
External_ear
Posterior_fornix_of_vagina

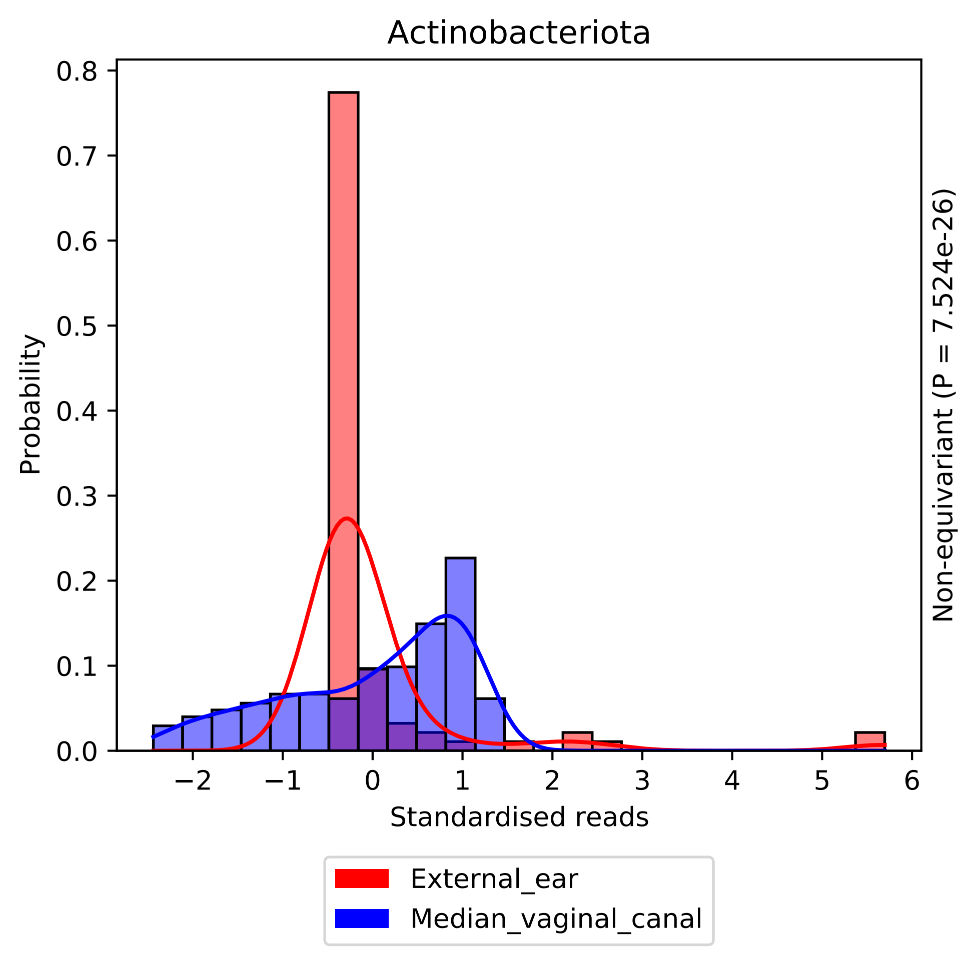

Actinobacteriota
Probability
Standardised reads
Non-equivariant (P = 7.524e-26)
0.0
0.1
0.2
0.3
0.4
0.5
0.6
0.7
0.8
−2
−1
0
1
2
3
4
5
6
External_ear
Median_vaginal_canal

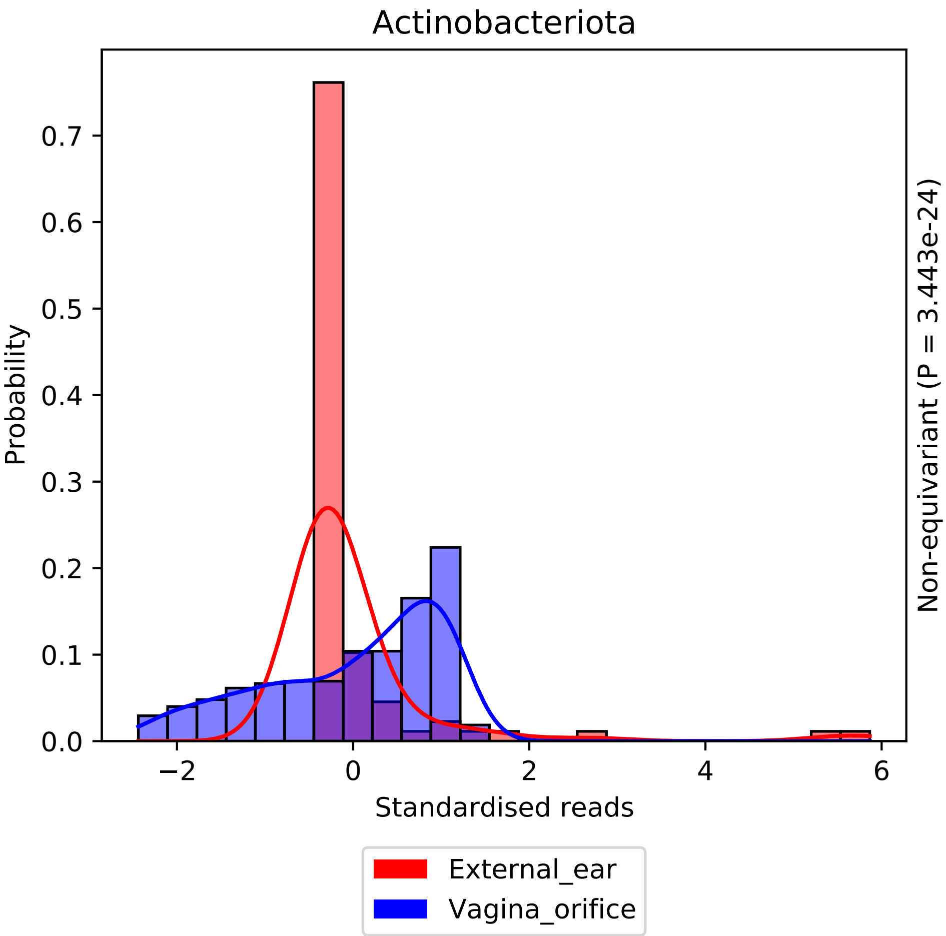

Actinobacteriota
Probability
0.0
0.1
0.2
0.3
0.4
0.5
0.6
0.7
−2
0
2
4
6
Standardised reads
Non-equivariant (P = 3.443e-24)
External_ear
Vagina_orifice

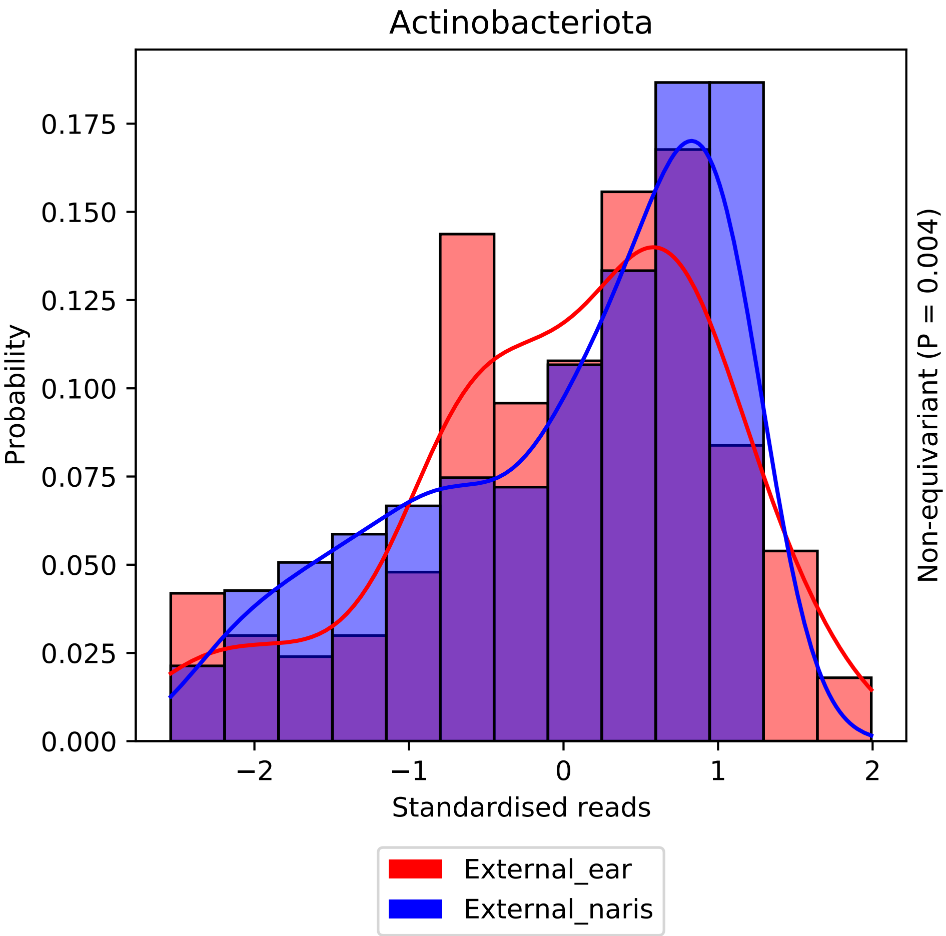

Actinobacteriota
Probability
0.000
0.025
0.050
0.075
0.100
0.125
0.150
0.175
−2
−1
0
1
2
Standardised reads
Non-equivariant (P = 0.004)
External_ear
External_naris

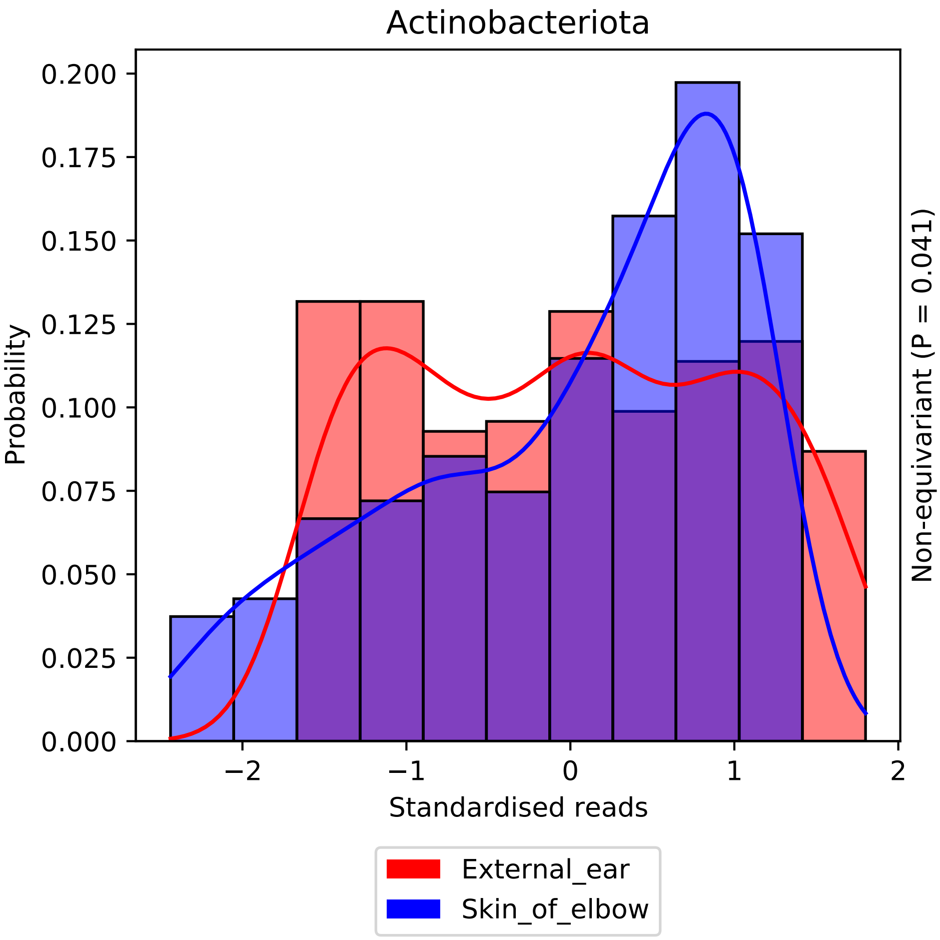
Actinobacteriota
Probability
0.200
0.175
0.150
0.125
0.100
0.075
0.050
0.025
0.000
−2
−1
0
1
2
Standardised reads
Non-equivariant (P = 0.041)
External_ear
Skin_of_elbow

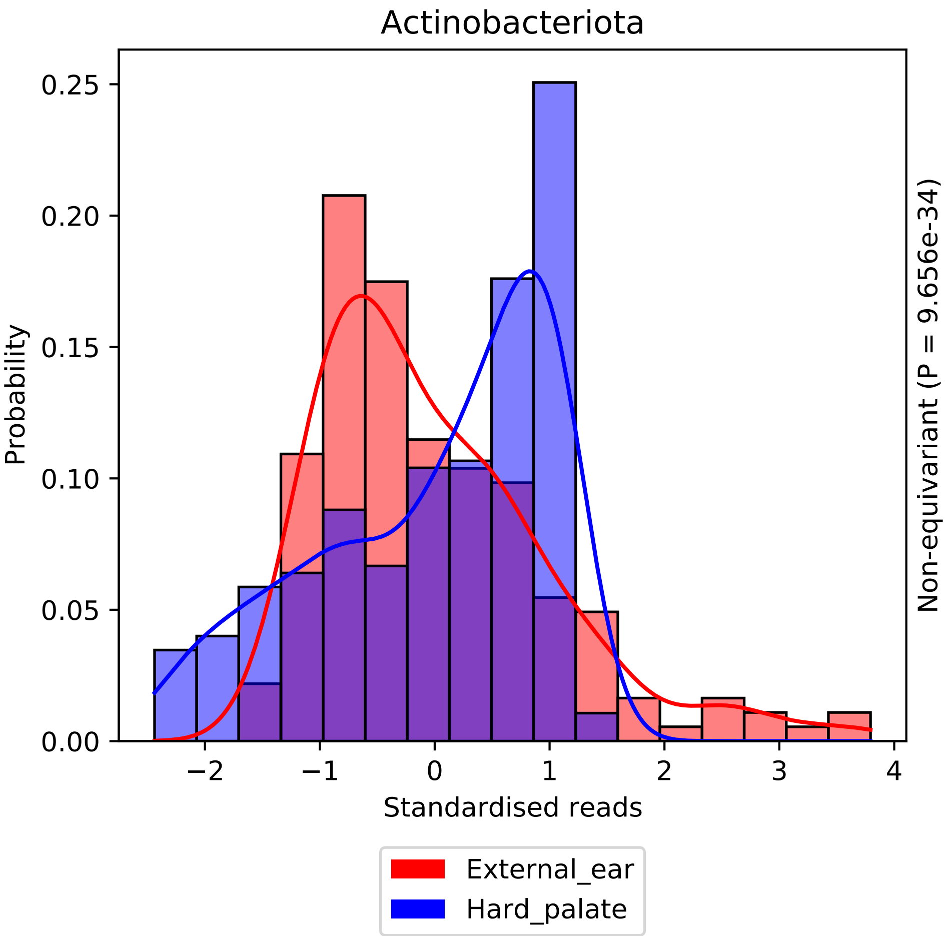
Actinobacteriota
Probability
0.25
0.20
0.15
0.10
0.05
0.00
−2
−1
0
1
2
3
4
Standardised reads
Non-equivariant (P = 9.656e-34)
External_ear
Hard_palate

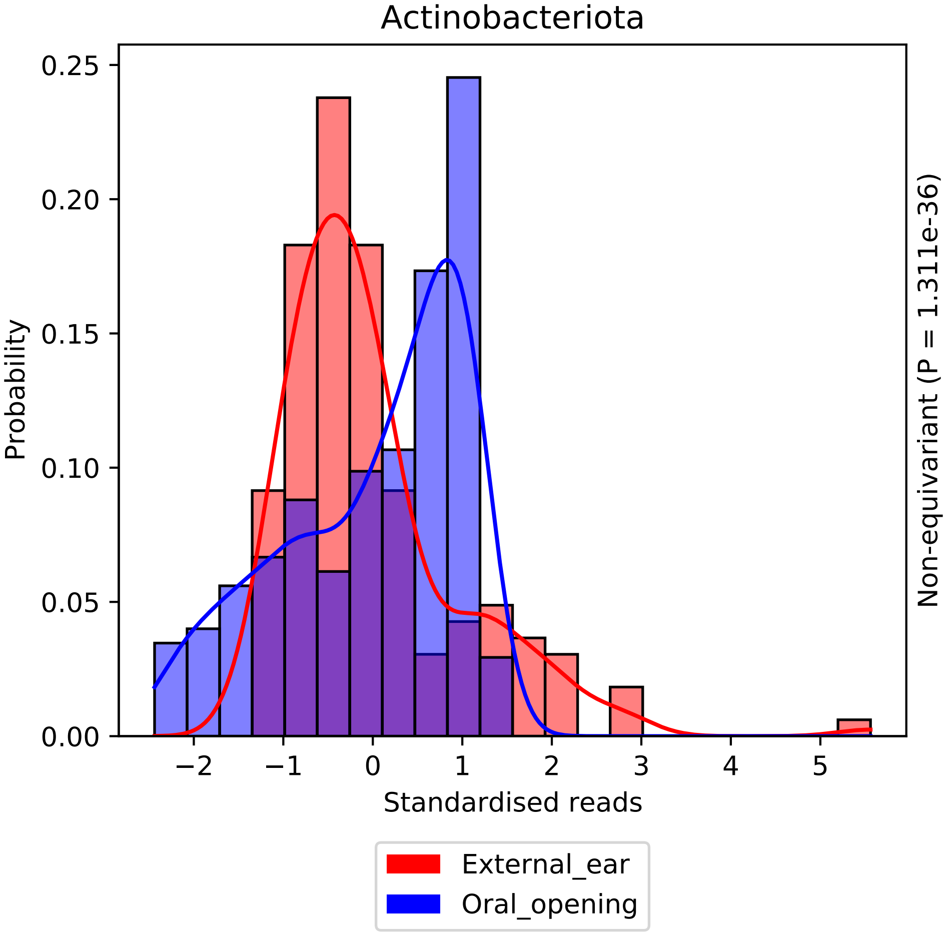
Actinobacteriota
Probability
0.25
0.20
0.15
0.10
0.05
0.00
−2
−1
0
1
2
3
4
5
Standardised reads
Non-equivariant (P = 1.311e-36)
External_ear
Oral_opening

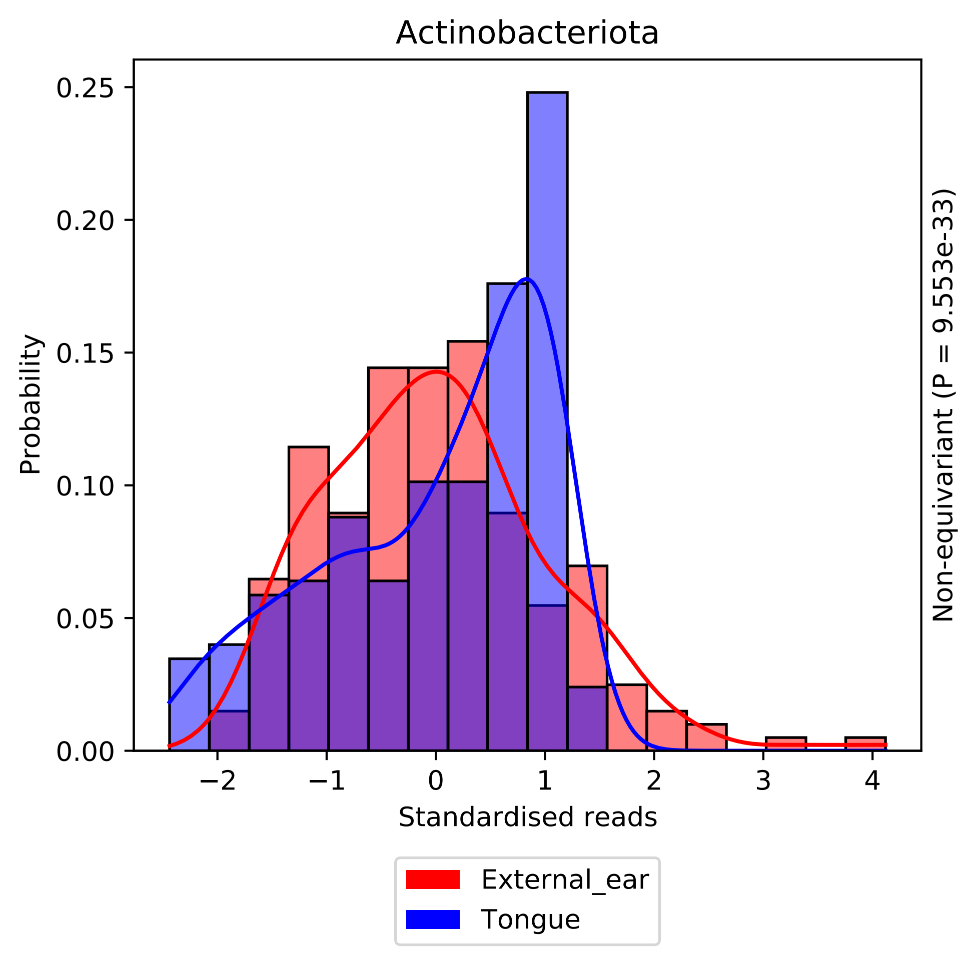
Actinobacteriota
Probability
0.25
0.20
0.15
0.10
0.05
0.00
−2
−1
0
1
2
3
4
Standardised reads
Non-equivariant (P = 9.553e-33)
External_ear
Tongue

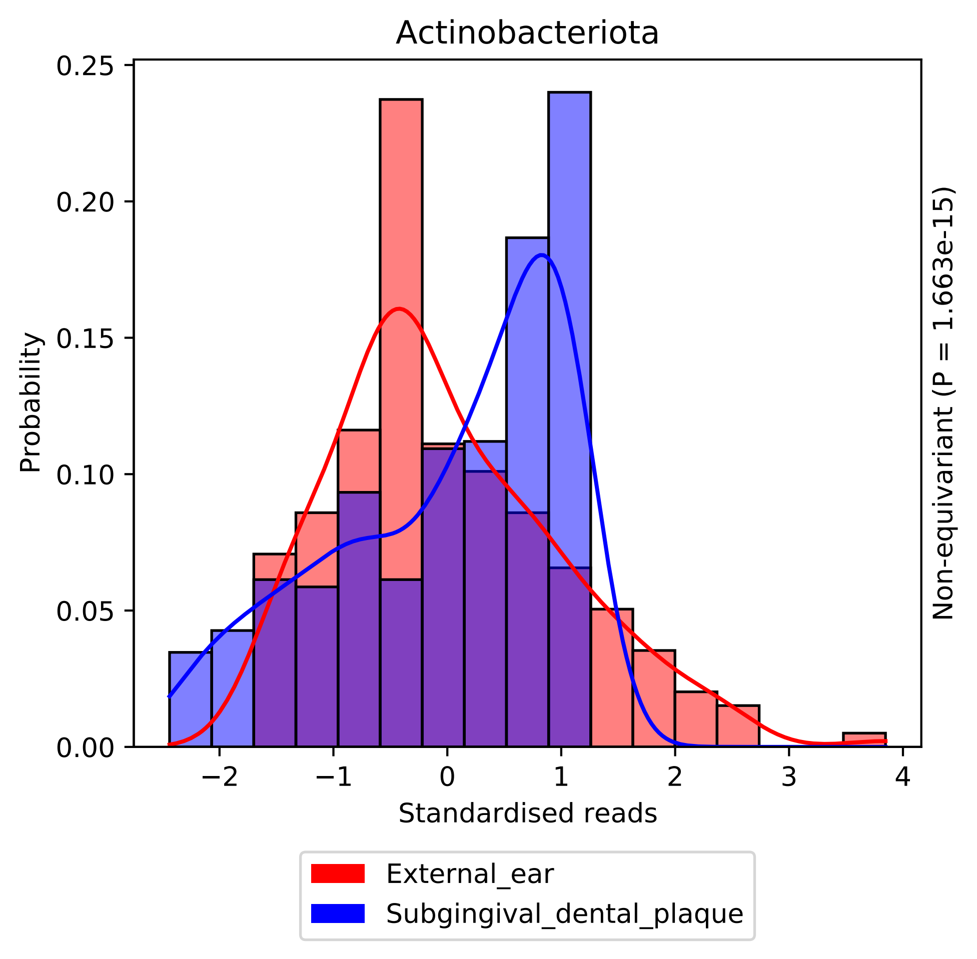
Actinobacteriota
0.25
0.20
0.15
0.10
0.05
0.00
Probability
−2
−1
0
1
2
3
4
Standardised reads
Non-equivariant (P = 1.663e-15)
External_ear
Subgingival_dental_plaque

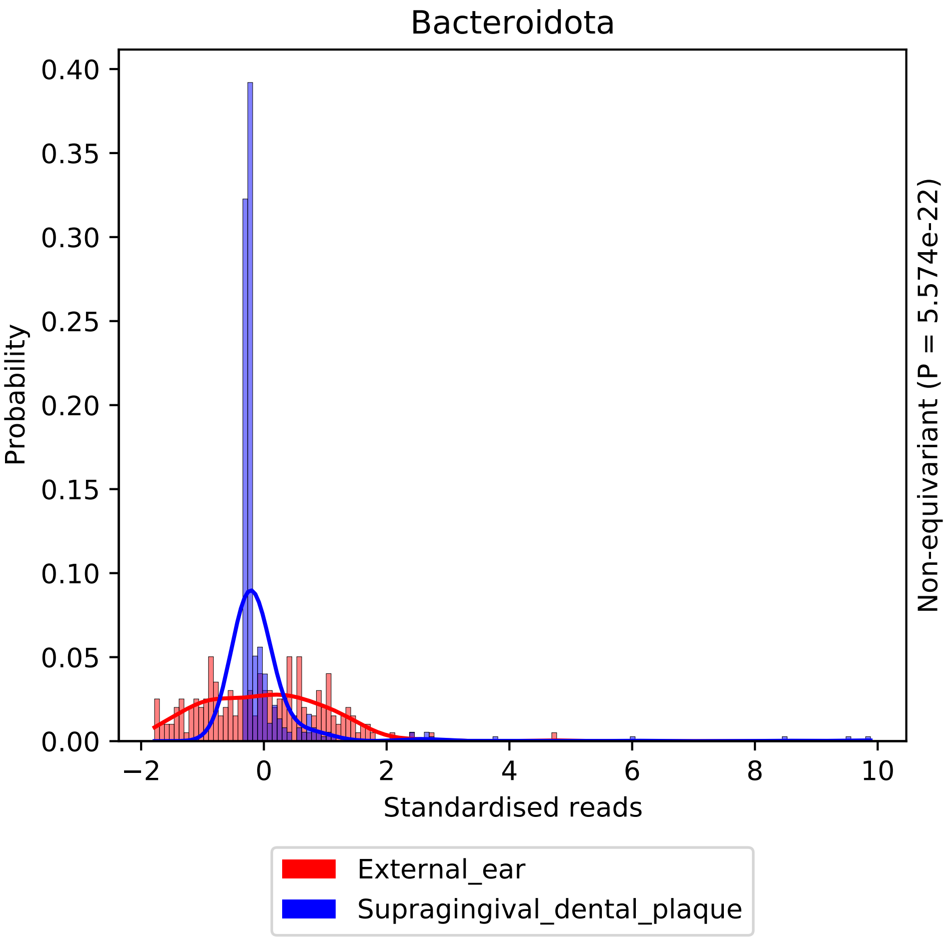
Bacteroidota
Probability
Standardised reads
Non-equivariant (P = 5.574e-22)
0.00
0.05
0.10
0.15
0.20
0.25
0.30
0.35
0.40
−2
0
2
4
6
8
10
External_ear
Supragingival_dental_plaque

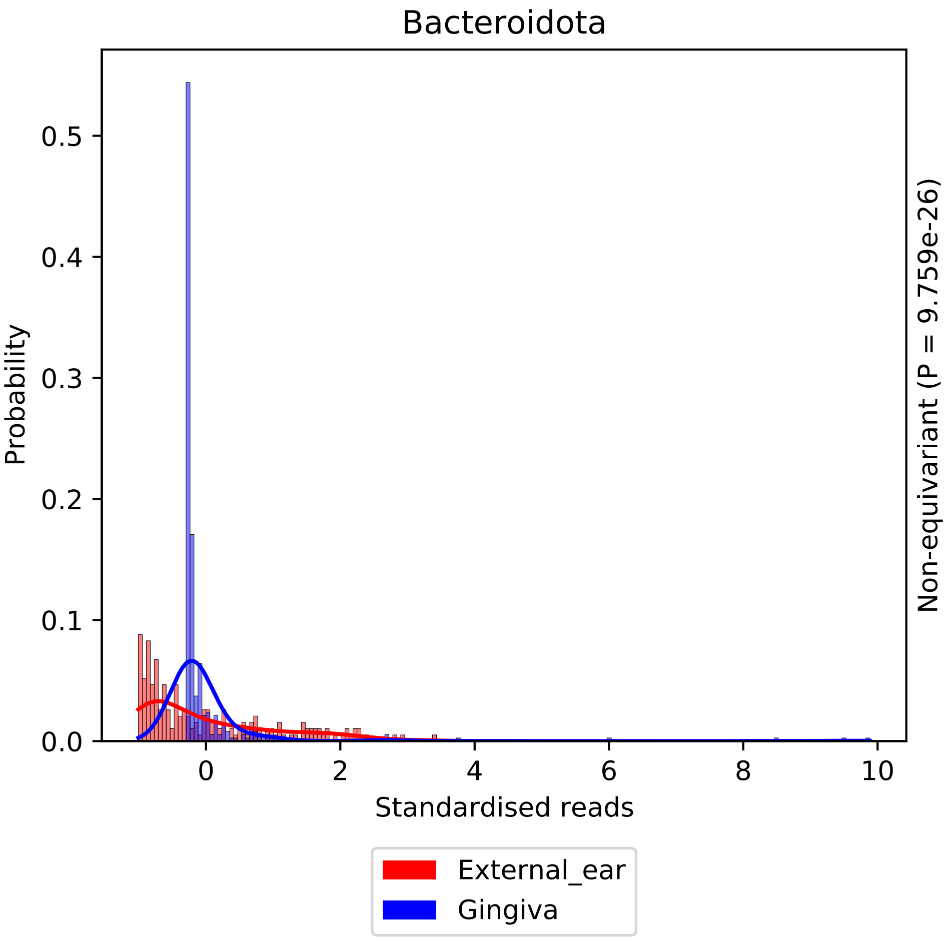

Bacteroidota
Probability
Standardised reads
0.0
0.1
0.2
0.3
0.4
0.5
0
2
4
6
8
10
Non-equivariant (P = 9.759e-26)
External_ear
Gingiva

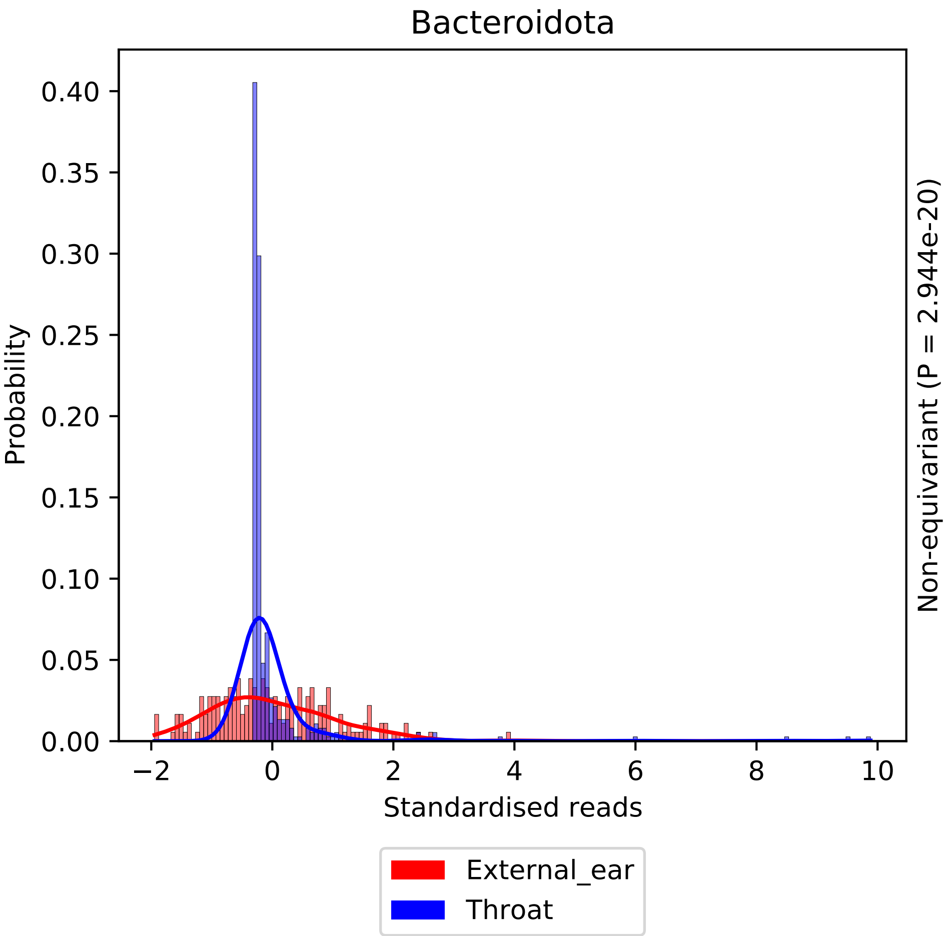

Bacteroidota
Probability
0.40
0.35
0.30
0.25
0.20
0.15
0.10
0.05
0.00
−2
0
2
4
6
8
10
Standardised reads
Non-equivariant (P = 2.944e-20)
External_ear
Throat

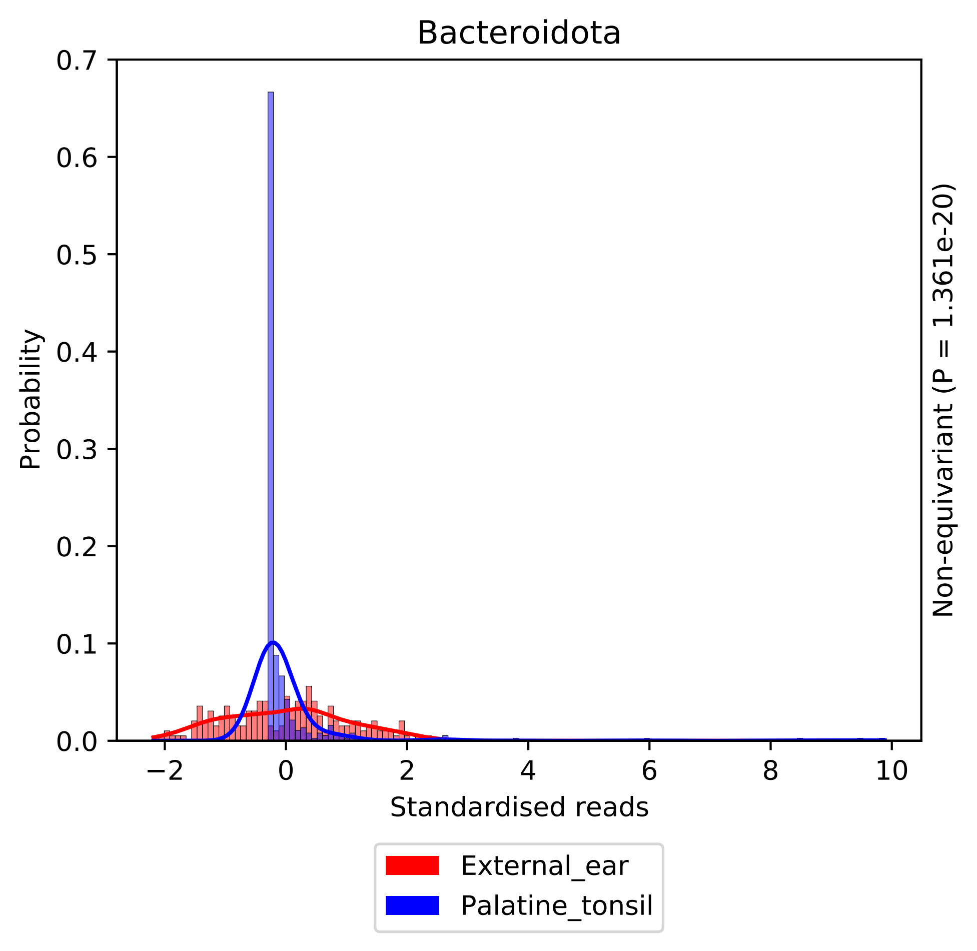
Bacteroidota
Probability
0.7
0.6
0.5
0.4
0.3
0.2
0.1
0.0
−2
0
2
4
6
8
10
Standardised reads
Non-equivariant (P = 1.361e-20)
External_ear
Palatine_tonsil

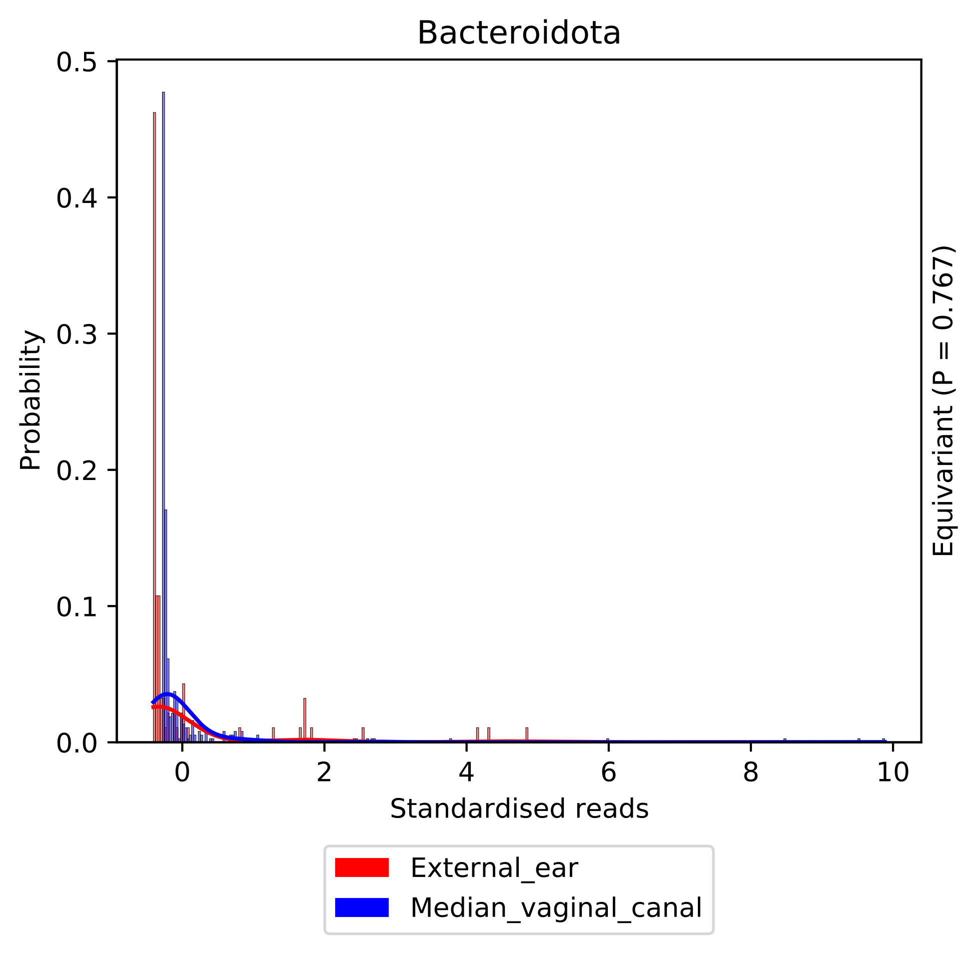
Bacteroidota
Probability
0.5
0.4
0.3
0.2
0.1
0.0
0
2
4
6
8
10
Standardised reads
Equivariant (P = 0.767)
External_ear
Median_vaginal_canal

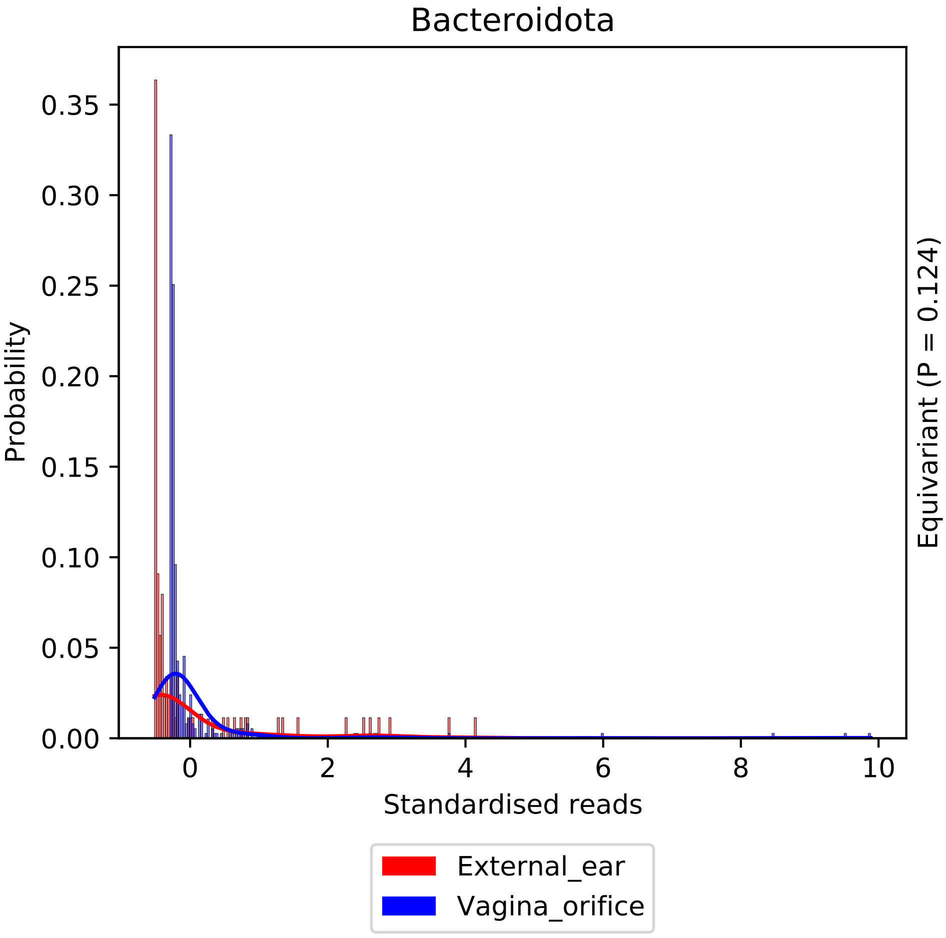

Bacteroidota
0.35
0.30
0.25
0.20
0.15
0.10
0.05
0.00
Probability
0
2
4
6
8
10
Standardised reads
Equivariant (P = 0.124)
External_ear
Vagina_orifice

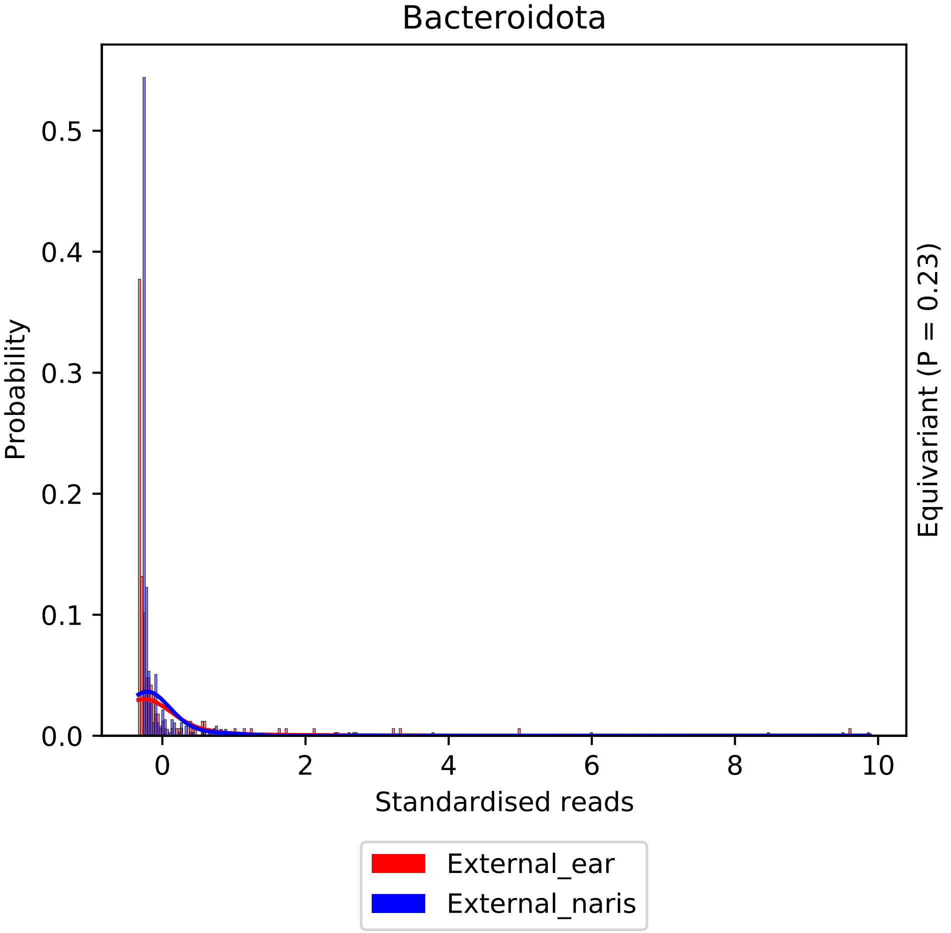

Bacteroidota
Probability
0.0
0.1
0.2
0.3
0.4
0.5
0
2
4
6
8
10
Standardised reads
Equivariant (P = 0.23)
External_ear
External_naris

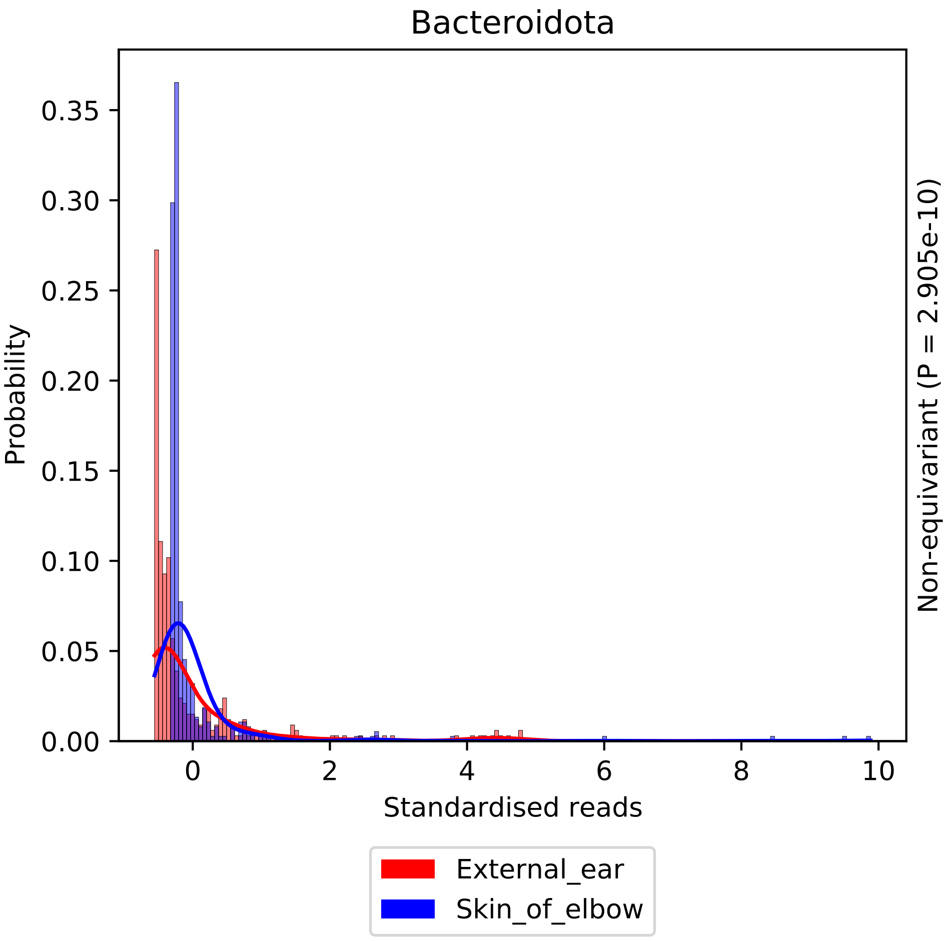

Bacteroidota
Probability
0.00
0.05
0.10
0.15
0.20
0.25
0.30
0.35
0
2
4
6
8
10
Standardised reads
Non-equivariant (P = 2.905e-10)
External_ear
Skin_of_elbow

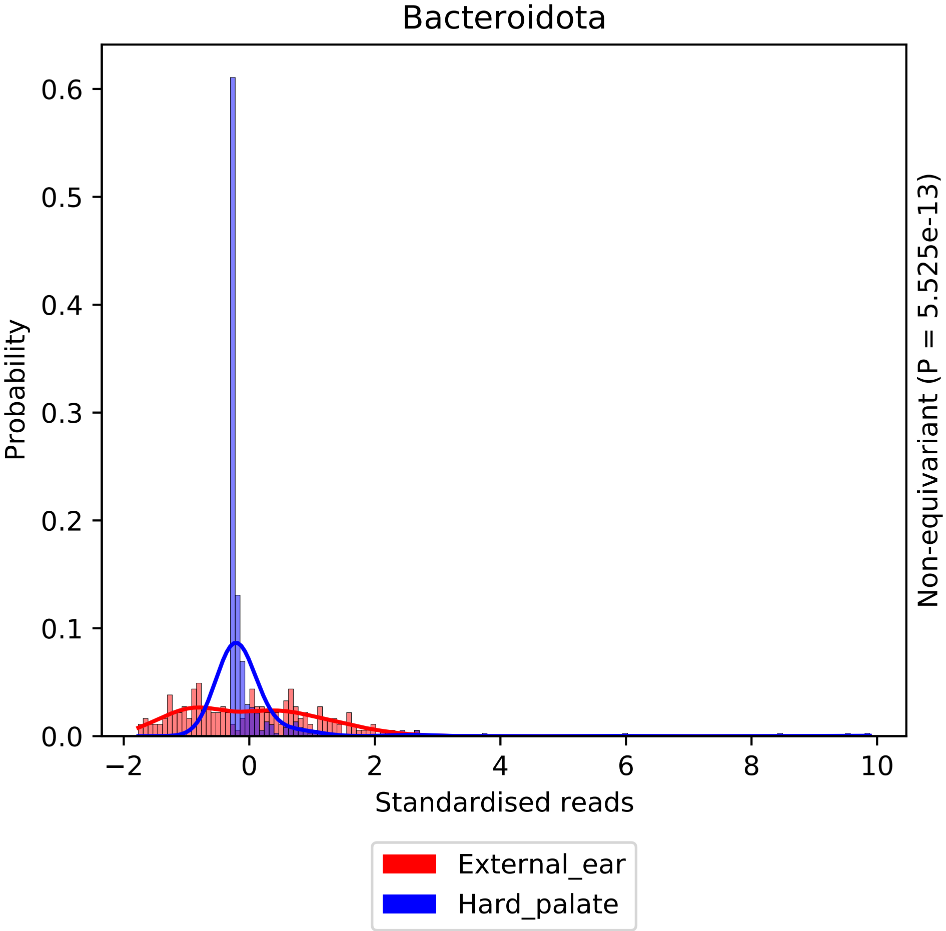

Bacteroidota
Probability
0.0
0.1
0.2
0.3
0.4
0.5
0.6
−2
0
2
4
6
8
10
Standardised reads
Non-equivariant (P = 5.525e-13)
External_ear
Hard_palate

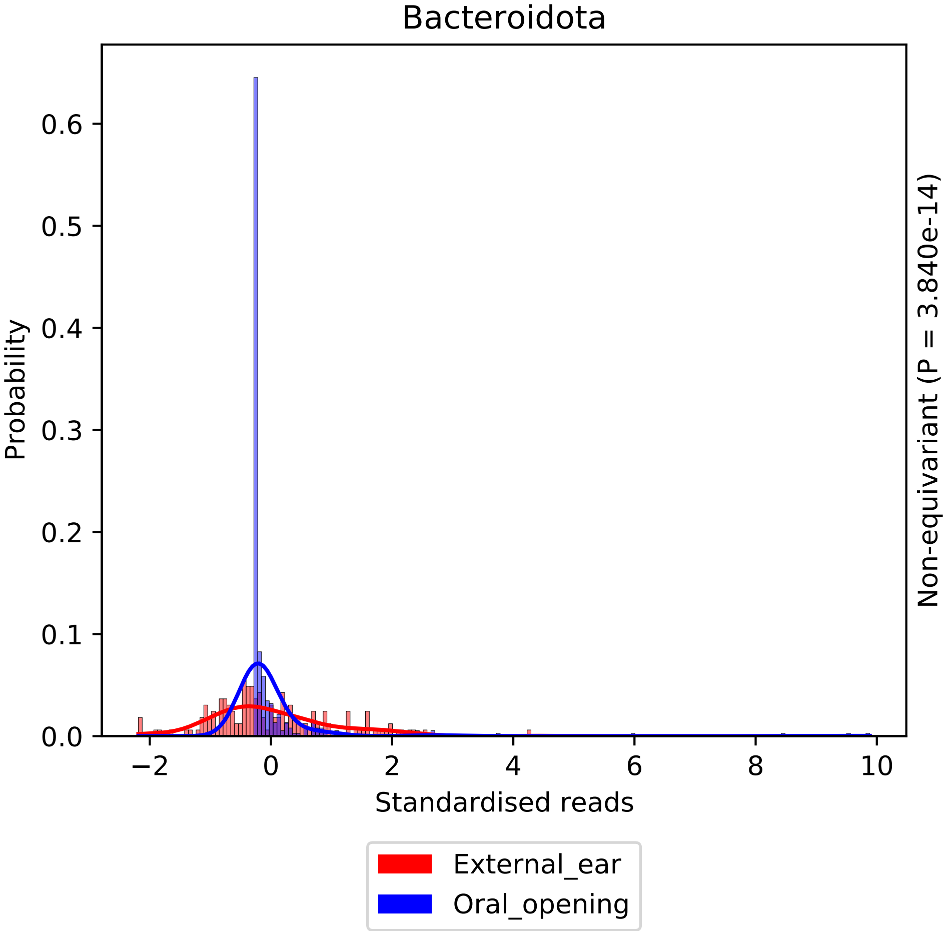

Bacteroidota
Probability
Standardised reads
Non-equivariant (P = 3.840e-14)
0.0
0.1
0.2
0.3
0.4
0.5
0.6
−2
0
2
4
6
8
10
External_ear
Oral_opening

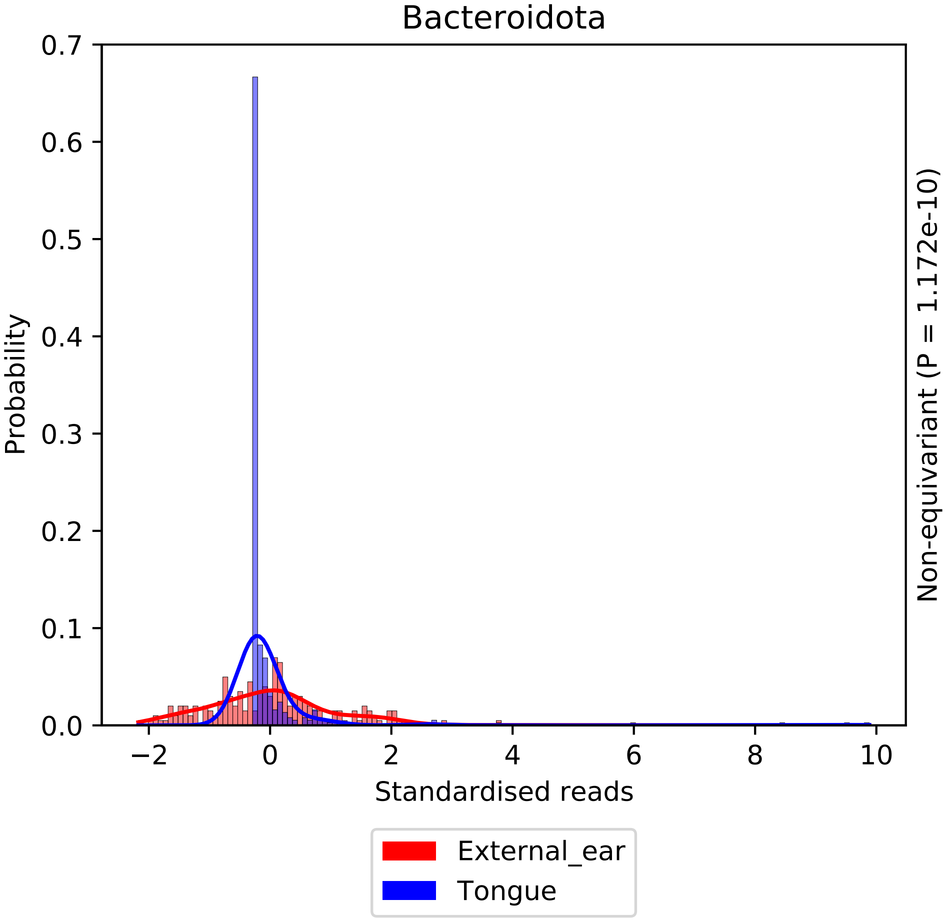
Bacteroidota
0.7
0.6
0.5
0.4
0.3
0.2
0.1
0.0
Probability
−2
0
2
4
6
8
10
Standardised reads
Non-equivariant (P = 1.172e-10)
External_ear
Tongue

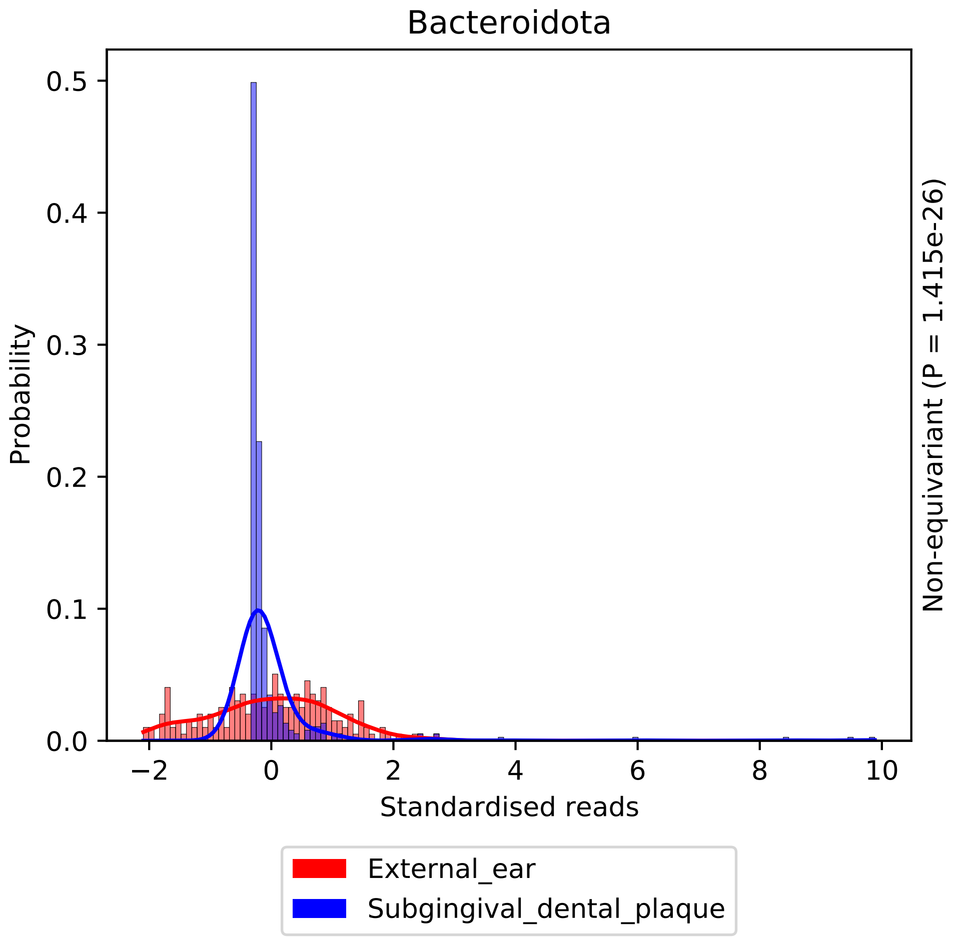

Bacteroidota
Probability
0.5
0.4
0.3
0.2
0.1
0.0
−2
0
2
4
6
8
10
Standardised reads
Non-equivariant (P = 1.415e-26)
External_ear
Subgingival_dental_plaque

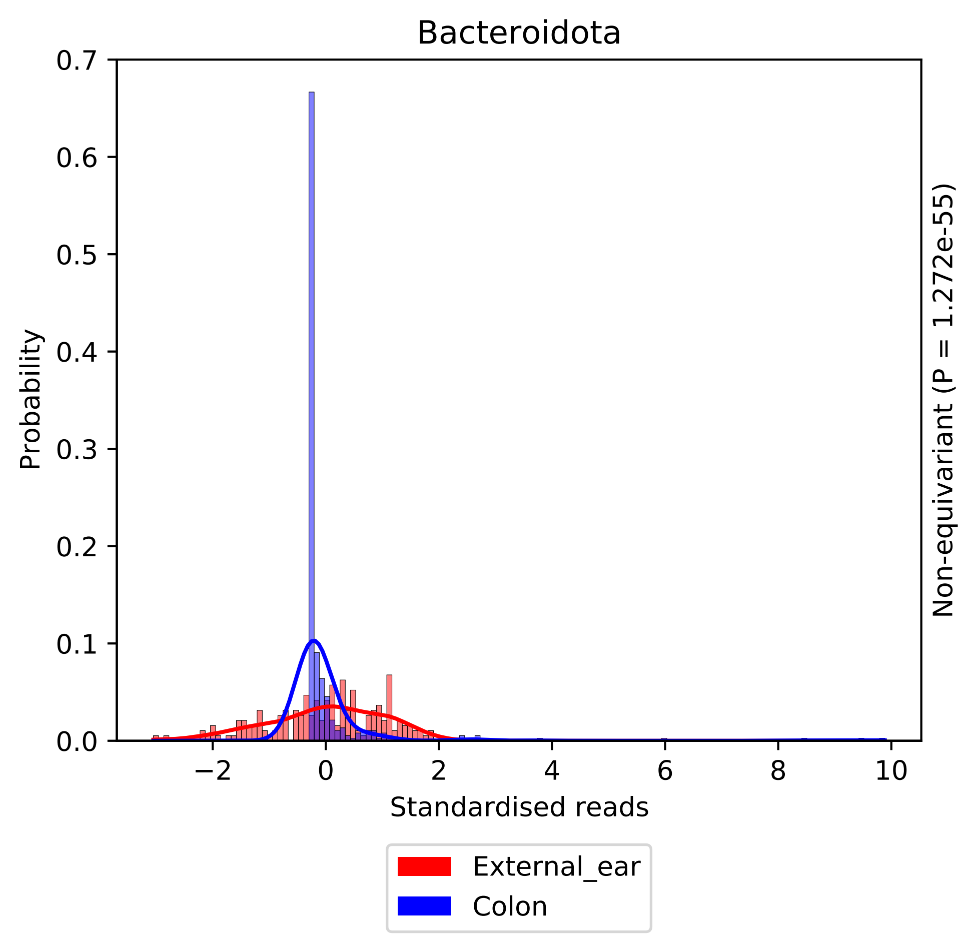

Bacteroidota
Probability
0.0
0.1
0.2
0.3
0.4
0.5
0.6
0.7
−2
0
2
4
6
8
10
Standardised reads
Non-equivariant (P = 1.272e-55)
External_ear
Colon

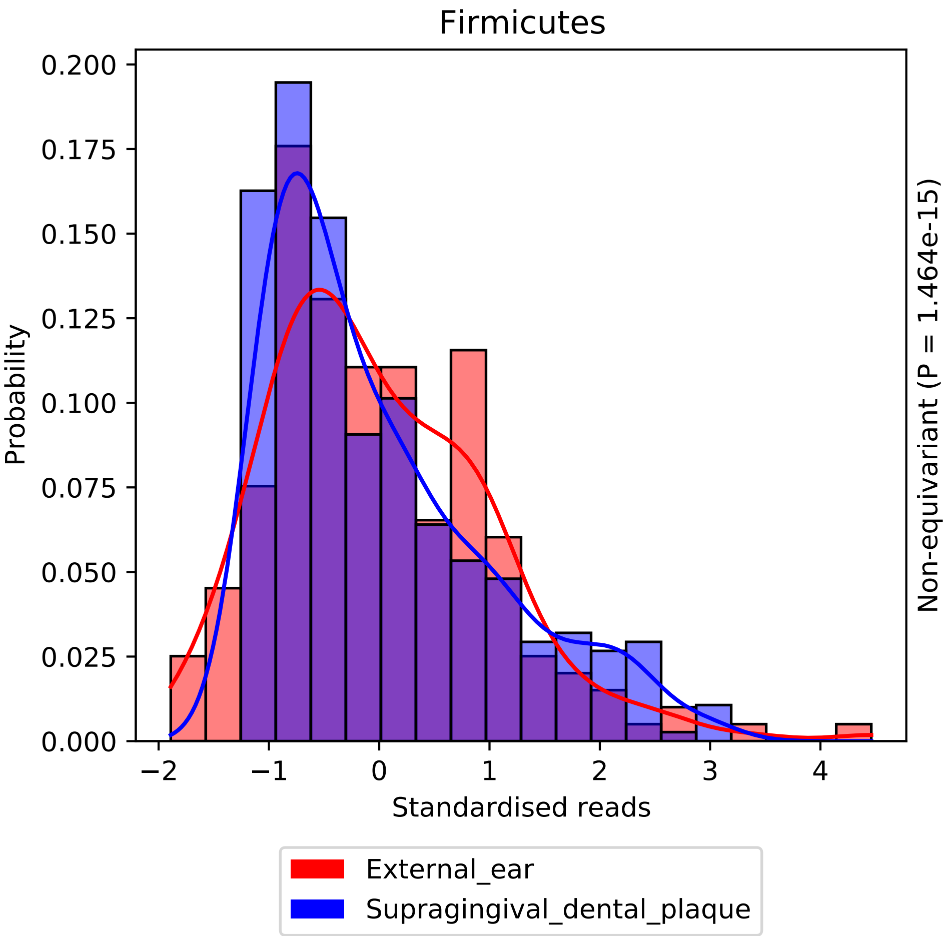

Firmicutes
Probability
0.000
0.025
0.050
0.075
0.100
0.125
0.150
0.175
0.200
−2
−1
0
1
2
3
4
Standardised reads
Non-equivariant (P = 1.464e-15)
External_ear
Supragingival_dental_plaque

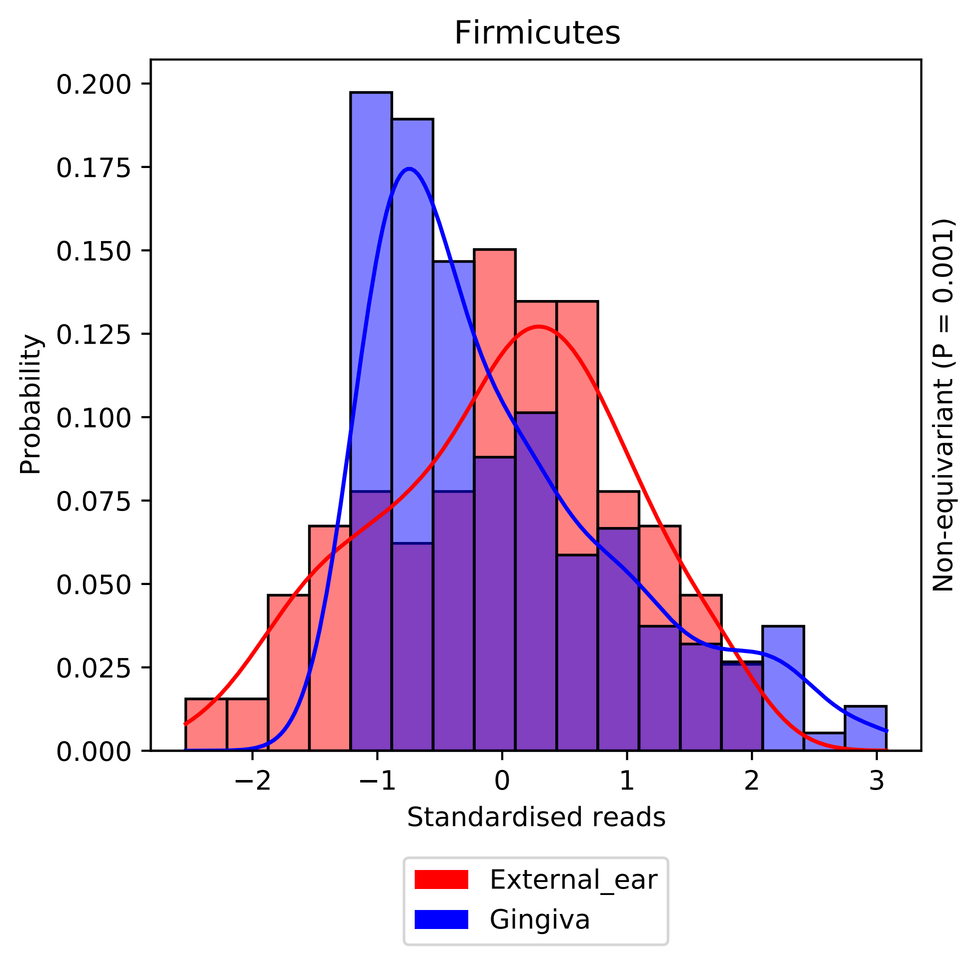

Firmicutes
Probability
0.000
0.025
0.050
0.075
0.100
0.125
0.150
0.175
0.200
−2
−1
0
1
2
3
Standardised reads
Non-equivariant (P = 0.001)
External_ear
Gingiva

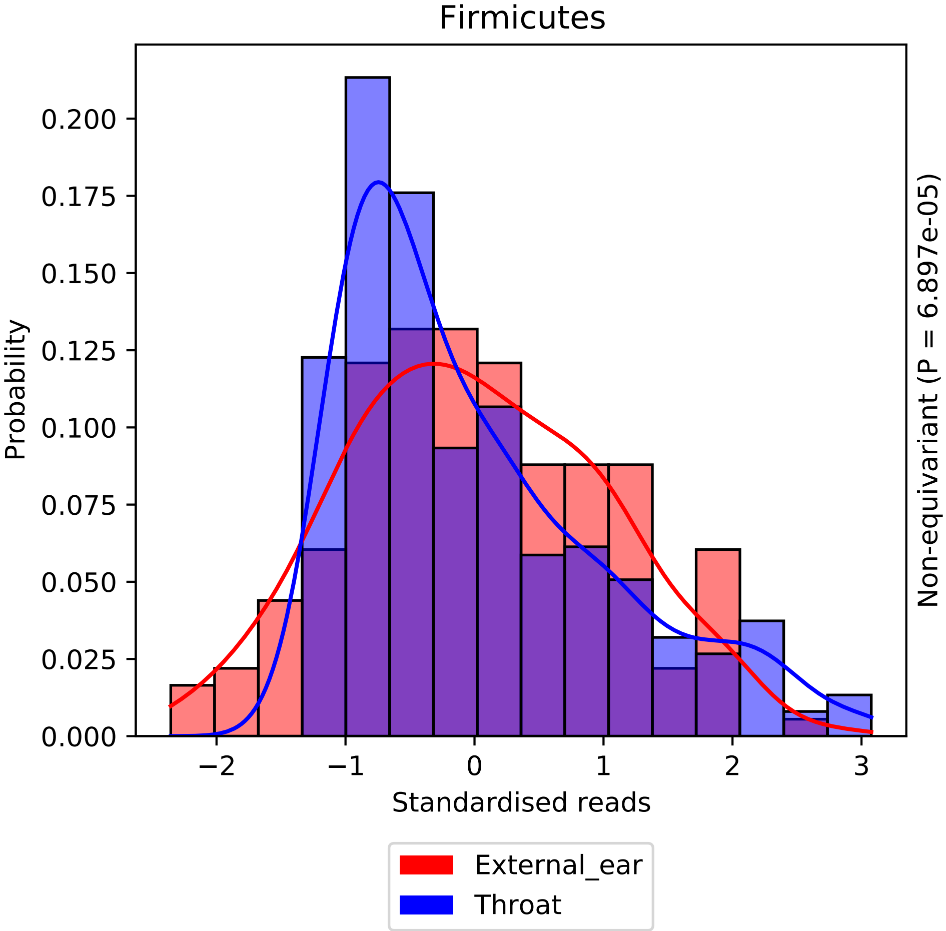

Firmicutes
Probability
0.000
0.025
0.050
0.075
0.100
0.125
0.150
0.175
0.200
−2
−1
0
1
2
3
Standardised reads
Non-equivariant (P = 6.897e-05)
External_ear
Throat

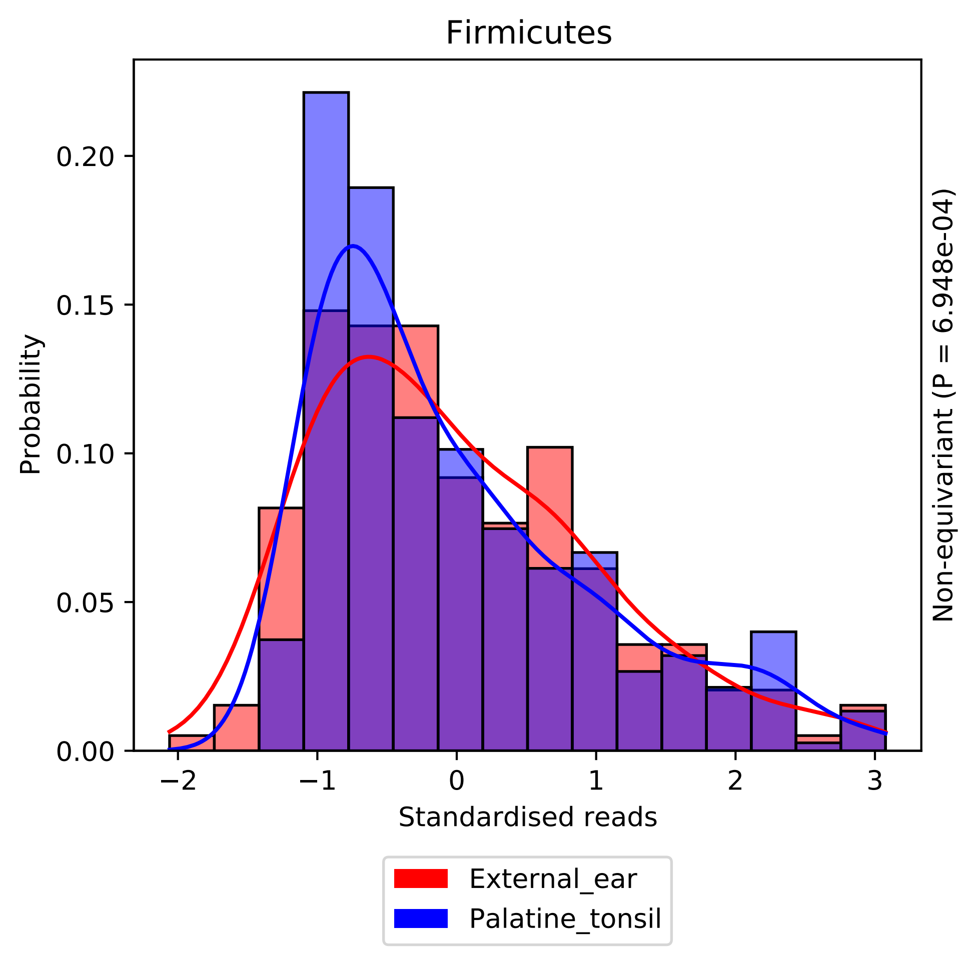

Firmicutes
Probability
0.00
0.05
0.10
0.15
0.20
−2
−1
0
1
2
3
Standardised reads
Non-equivariant (P = 6.948e-04)
External_ear
Palatine_tonsil

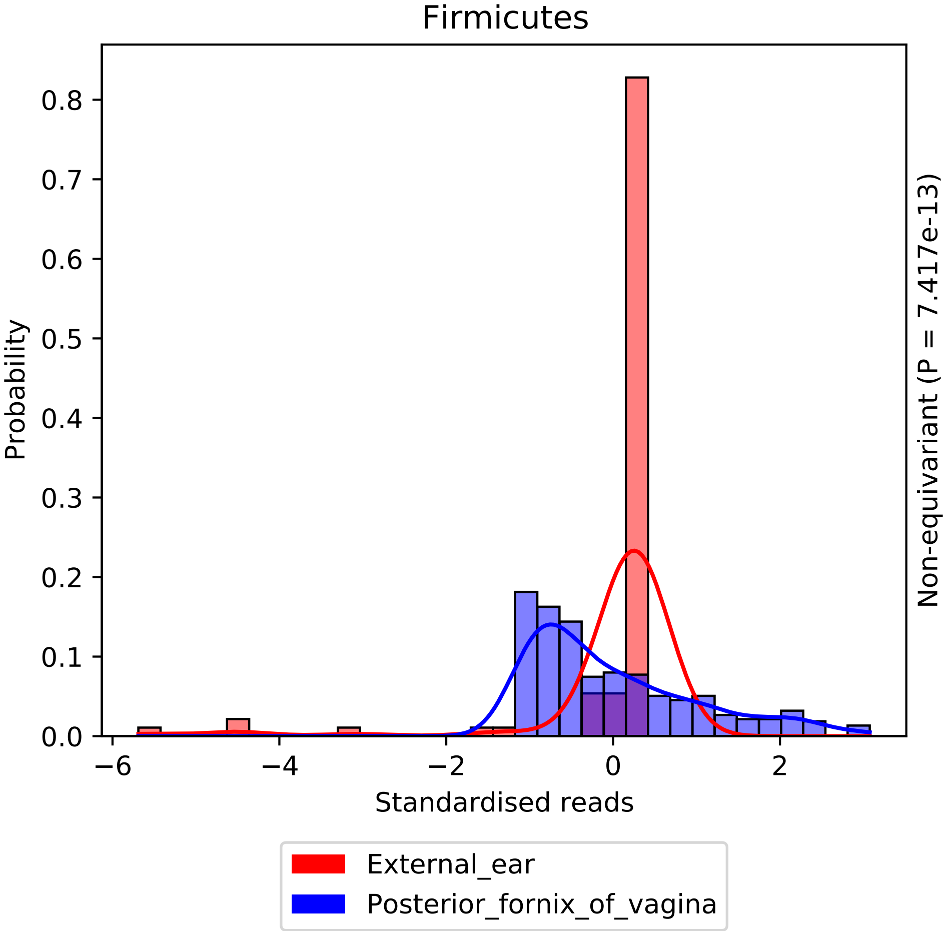
Firmicutes
Probability
Standardised reads
Non-equivariant (P = 7.417e-13)
0.0
0.1
0.2
0.3
0.4
0.5
0.6
0.7
0.8
−6
−4
−2
0
2
External_ear
Posterior_fornix_of_vagina

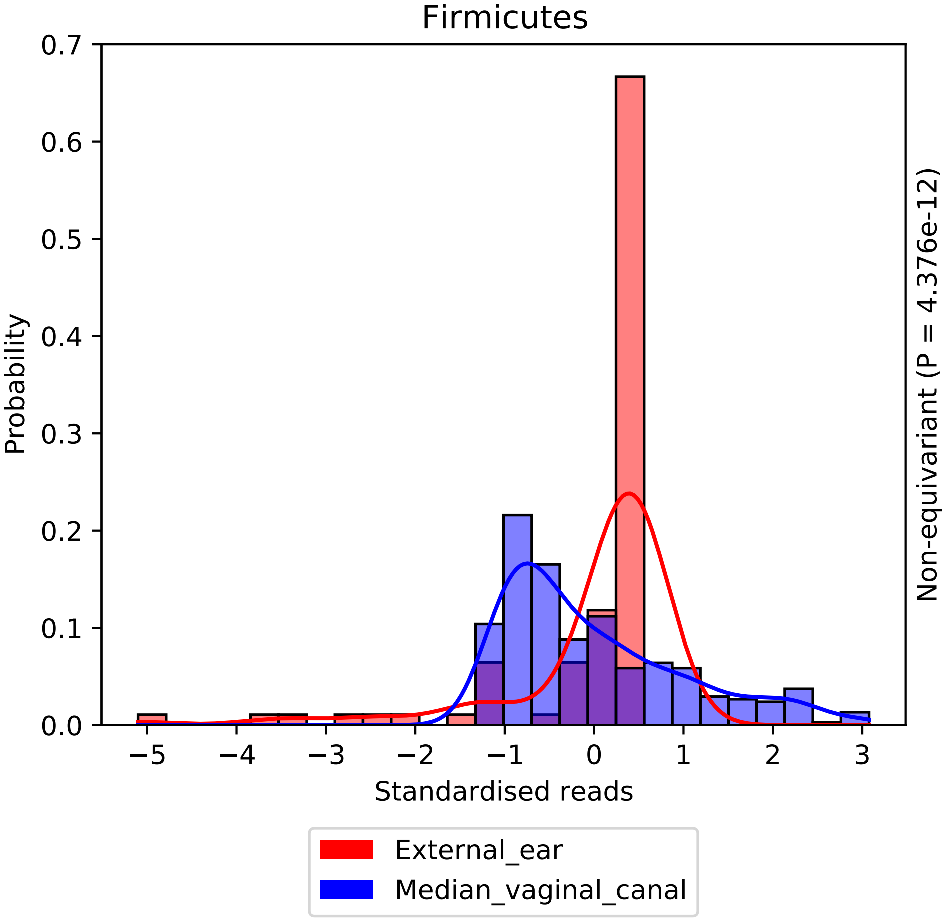

Firmicutes
0.7
0.6
0.5
0.4
0.3
0.2
0.1
0.0
Probability
−5
−4
−3
−2
−1
0
1
2
3
Standardised reads
Non-equivariant (P = 4.376e-12)
External_ear
Median_vaginal_canal

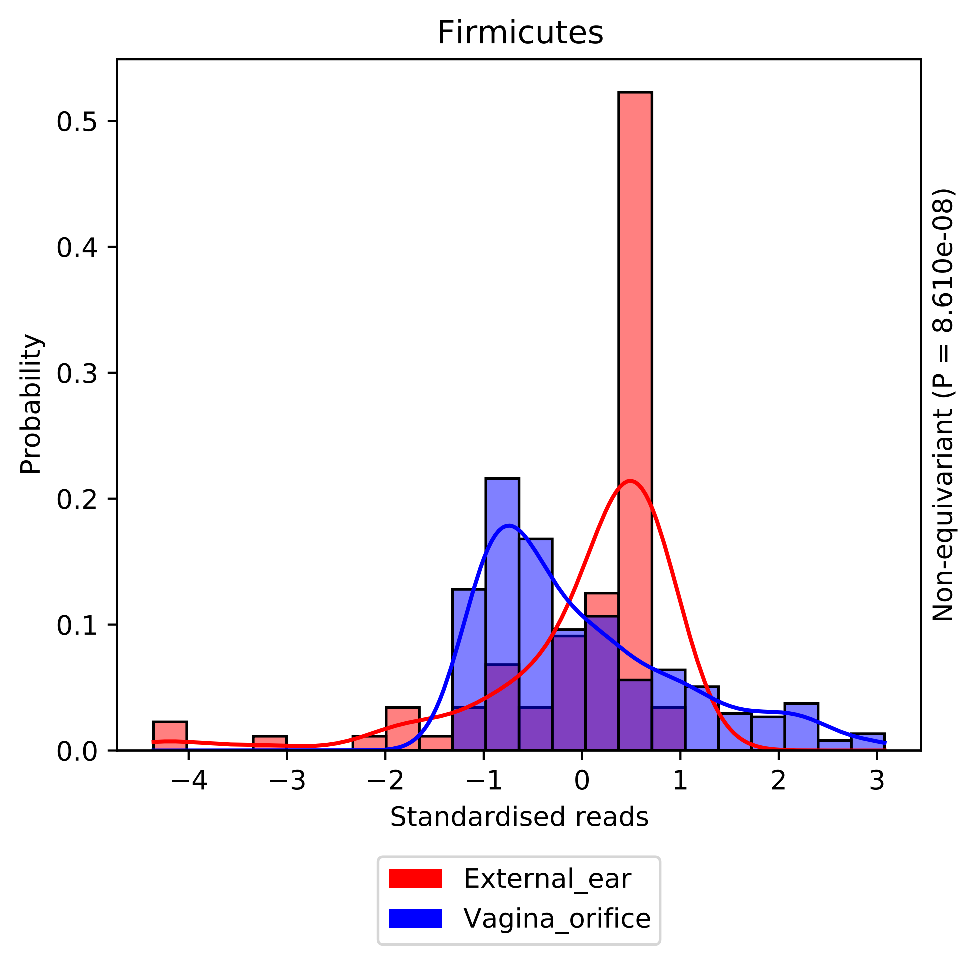

Firmicutes
Probability
0.0
0.1
0.2
0.3
0.4
0.5
−4
−3
−2
−1
0
1
2
3
Standardised reads
Non-equivariant (P = 8.610e-08)
External_ear
Vagina_orifice

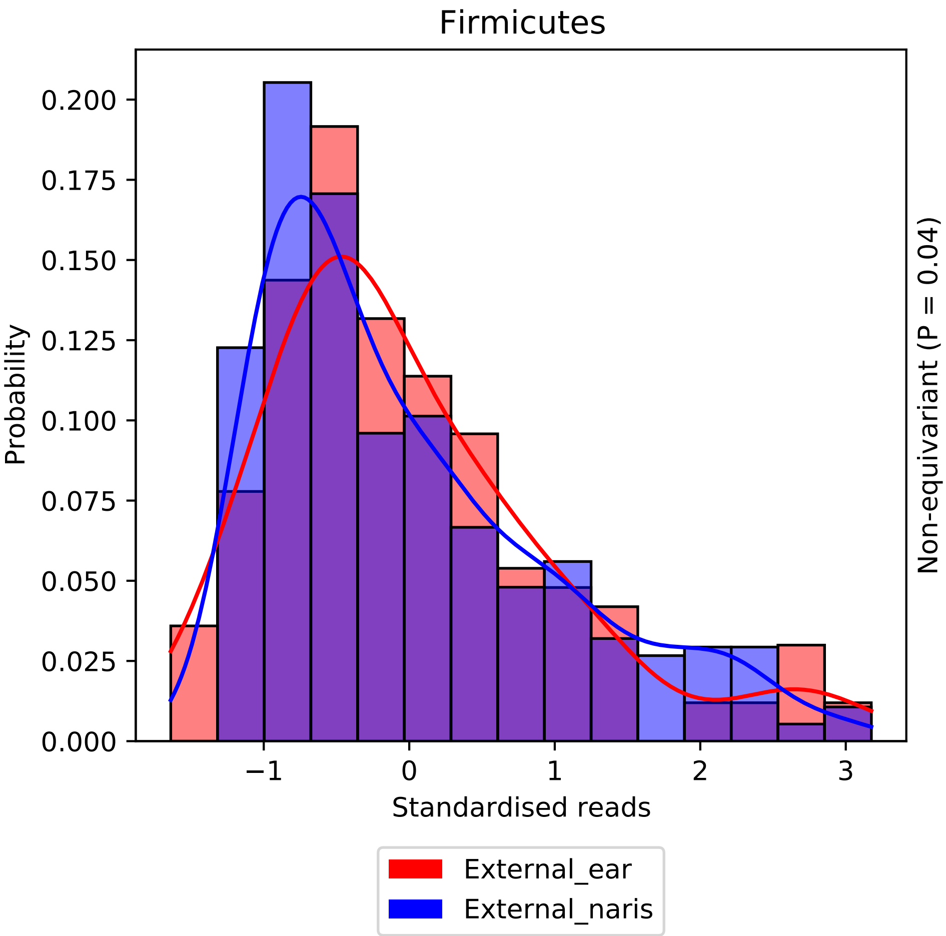

Firmicutes
Probability
0.000
0.025
0.050
0.075
0.100
0.125
0.150
0.175
0.200
−1
0
1
2
3
Standardised reads
Non-equivariant (P = 0.04)
External_ear
External_naris

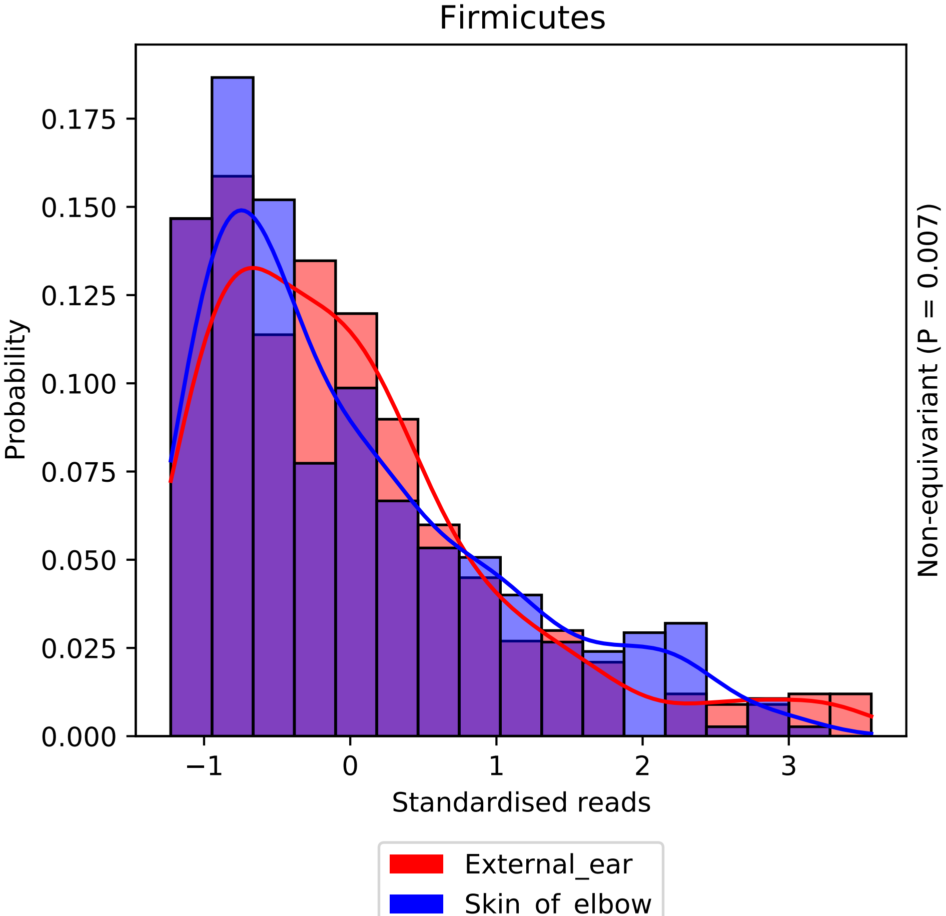
Firmicutes
Probability
0.000
0.025
0.050
0.075
0.100
0.125
0.150
0.175
−1
0
1
2
3
Standardised reads
Non-equivariant (P = 0.007)
External_ear
Skin_of_elbow

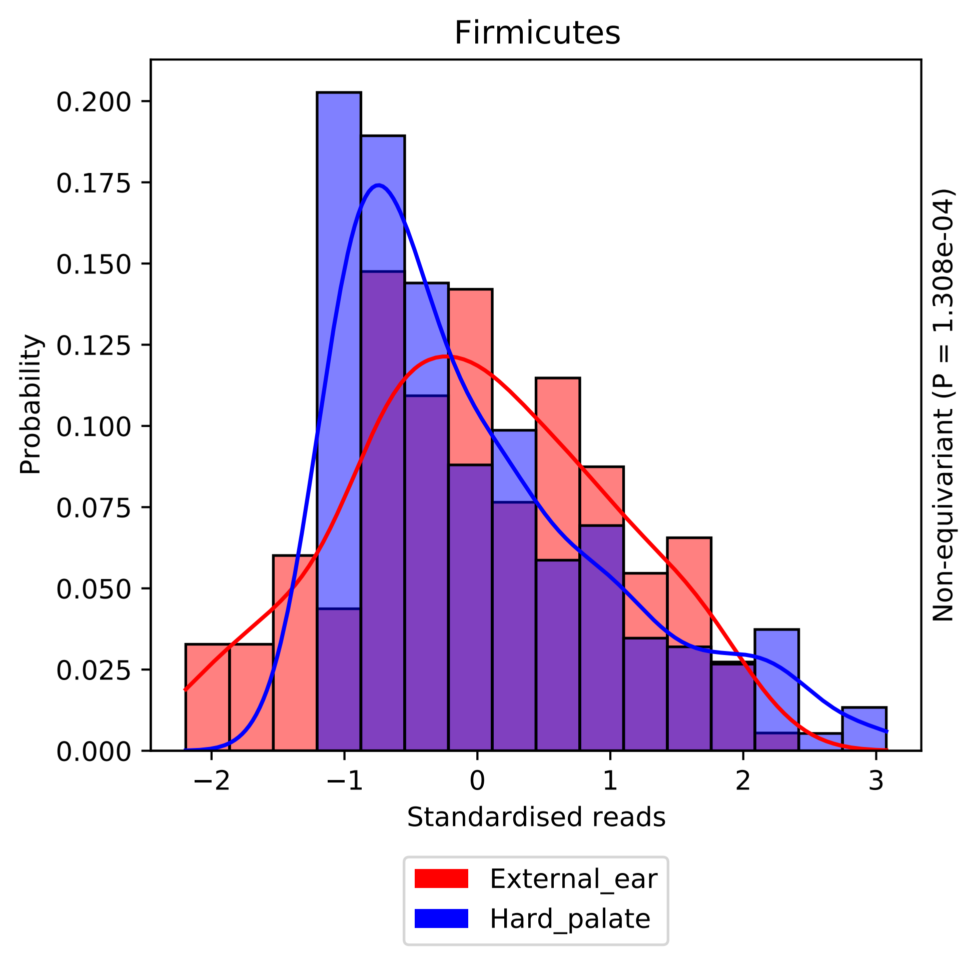

Firmicutes
Probability
0.000
0.025
0.050
0.075
0.100
0.125
0.150
0.175
0.200
−2
−1
0
1
2
3
Standardised reads
Non-equivariant (P = 1.308e-04)
External_ear
Hard_palate

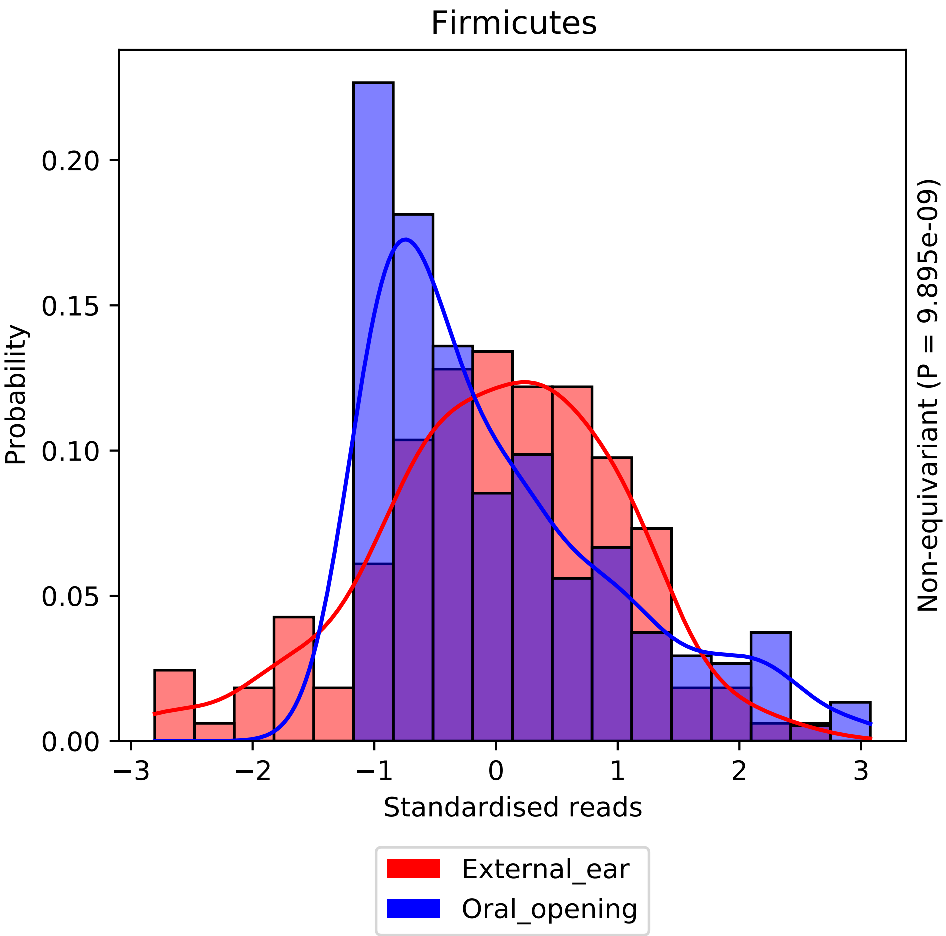

Firmicutes
Probability
0.00
0.05
0.10
0.15
0.20
−3
−2
−1
0
1
2
3
Standardised reads
Non-equivariant (P = 9.895e-09)
External_ear
Oral_opening

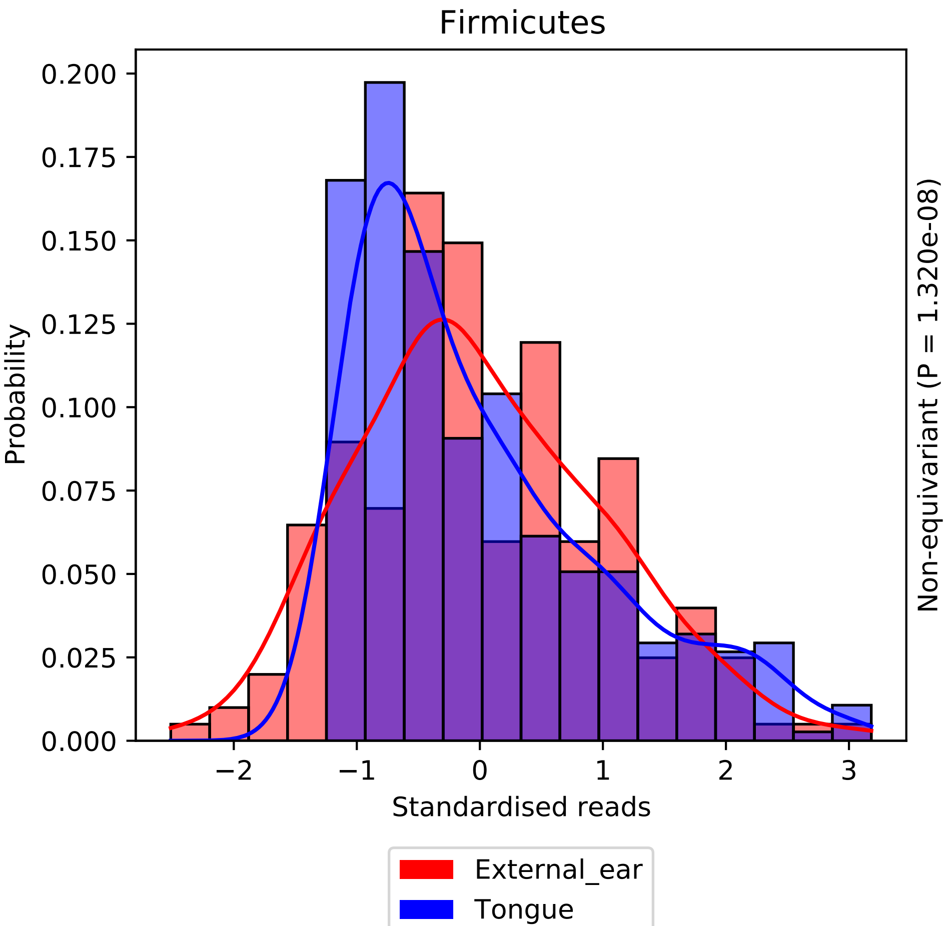

Firmicutes
Probability
0.200
0.175
0.150
0.125
0.100
0.075
0.050
0.025
0.000
−2
−1
0
1
2
3
Standardised reads
Non-equivariant (P = 1.320e-08)
External_ear
Tongue

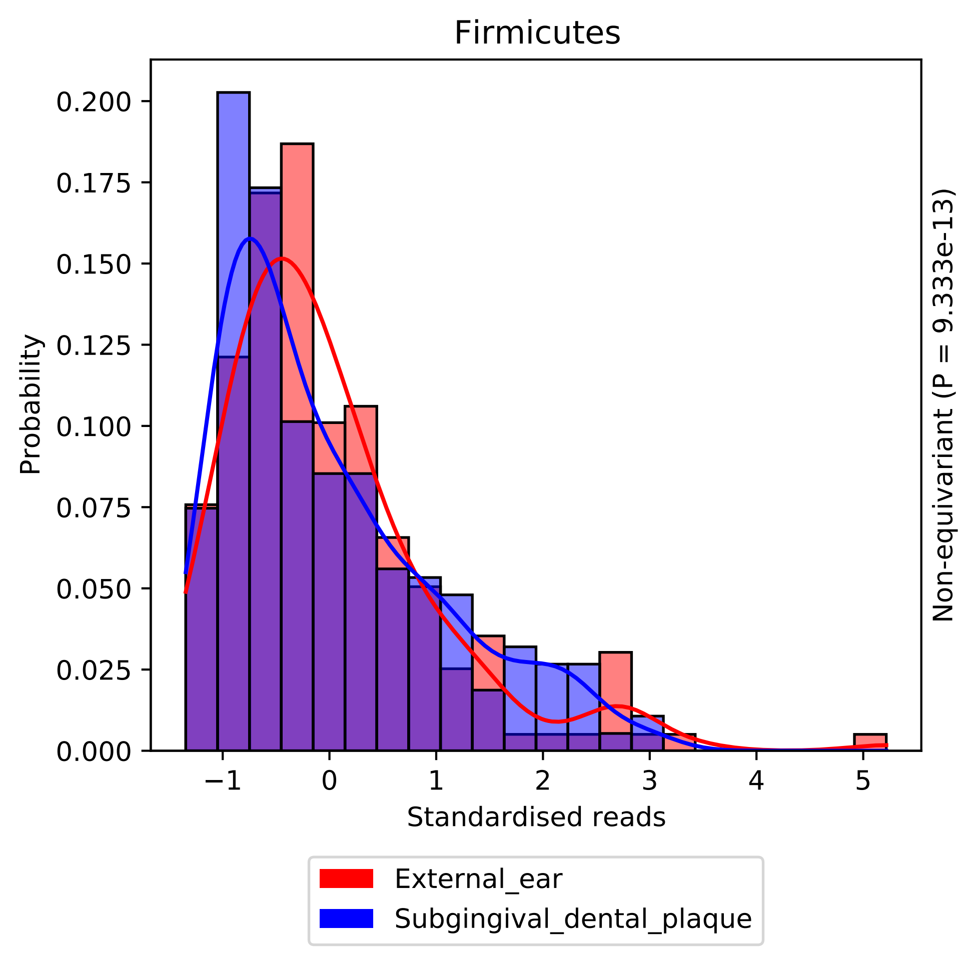

Firmicutes
0.200
0.175
0.150
0.125
0.100
0.075
0.050
0.025
0.000
Probability
−1
0
1
2
3
4
5
Standardised reads
Non-equivariant (P = 9.333e-13)
External_ear
Subgingival_dental_plaque

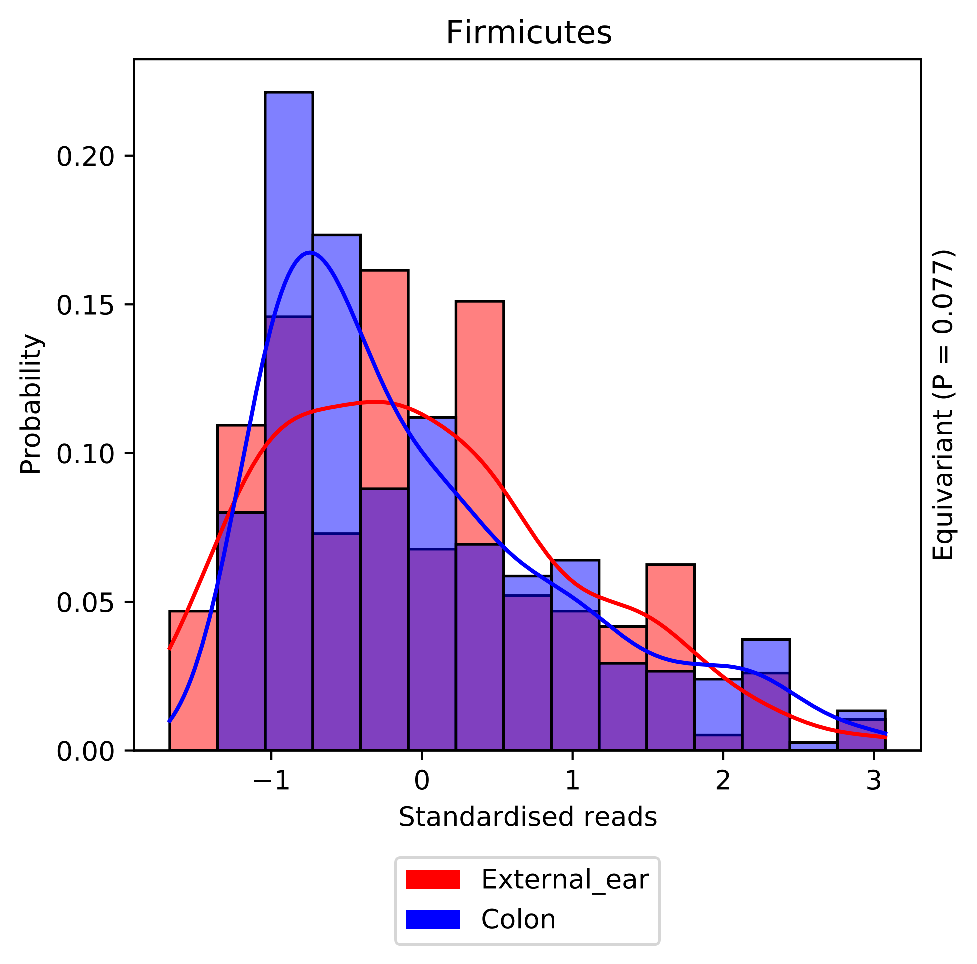

Firmicutes
Probability
0.00
0.05
0.10
0.15
0.20
−1
0
1
2
3
Standardised reads
Equivariant (P = 0.077)
External_ear
Colon

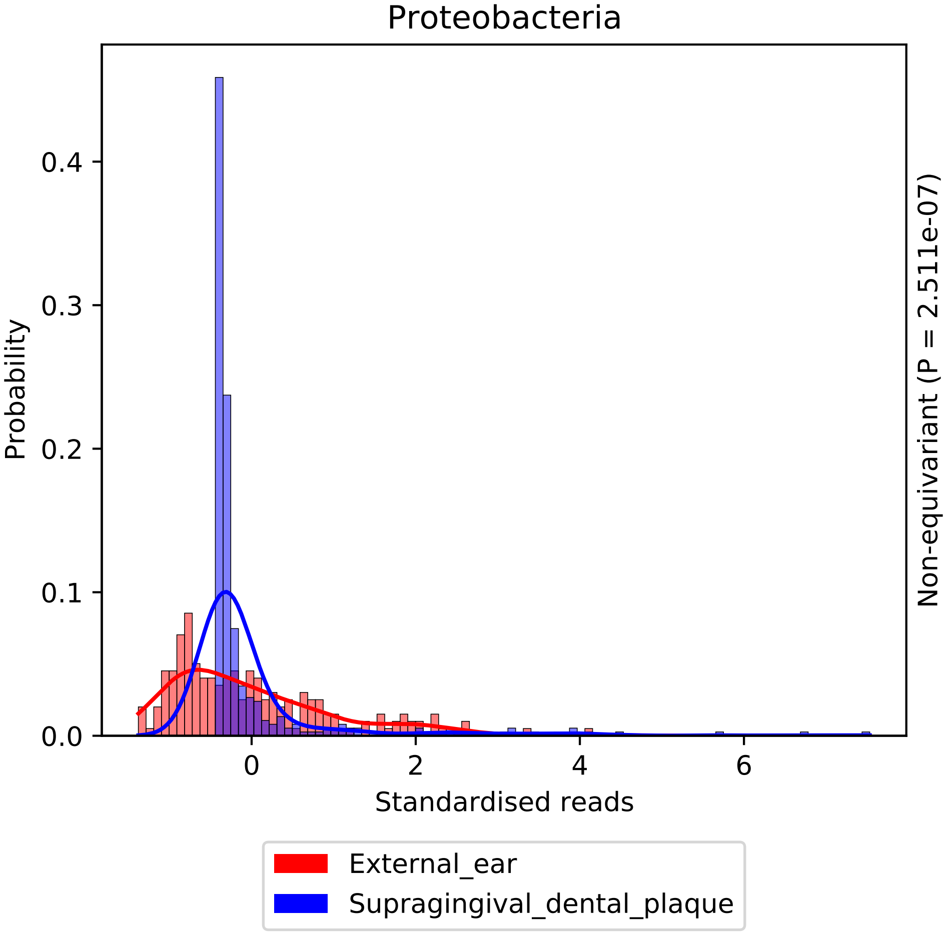

Proteobacteria
0.4
0.3
0.2
0.1
0.0
Probability
0
2
4
6
Standardised reads
Non-equivariant (P = 2.511e-07)
External_ear
Supragingival_dental_plaque

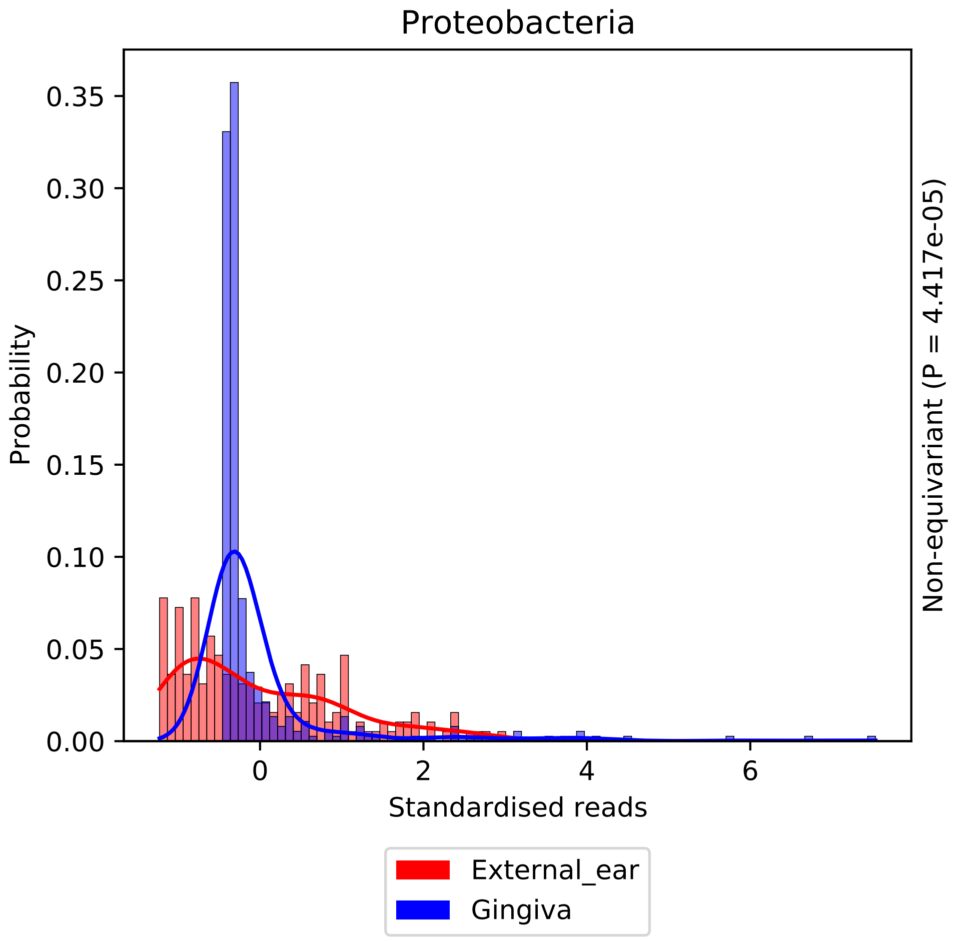

Proteobacteria
Probability
0.35
0.30
0.25
0.20
0.15
0.10
0.05
0.00
0
2
4
6
Standardised reads
Non-equivariant (P = 4.417e-05)
External_ear
Gingiva

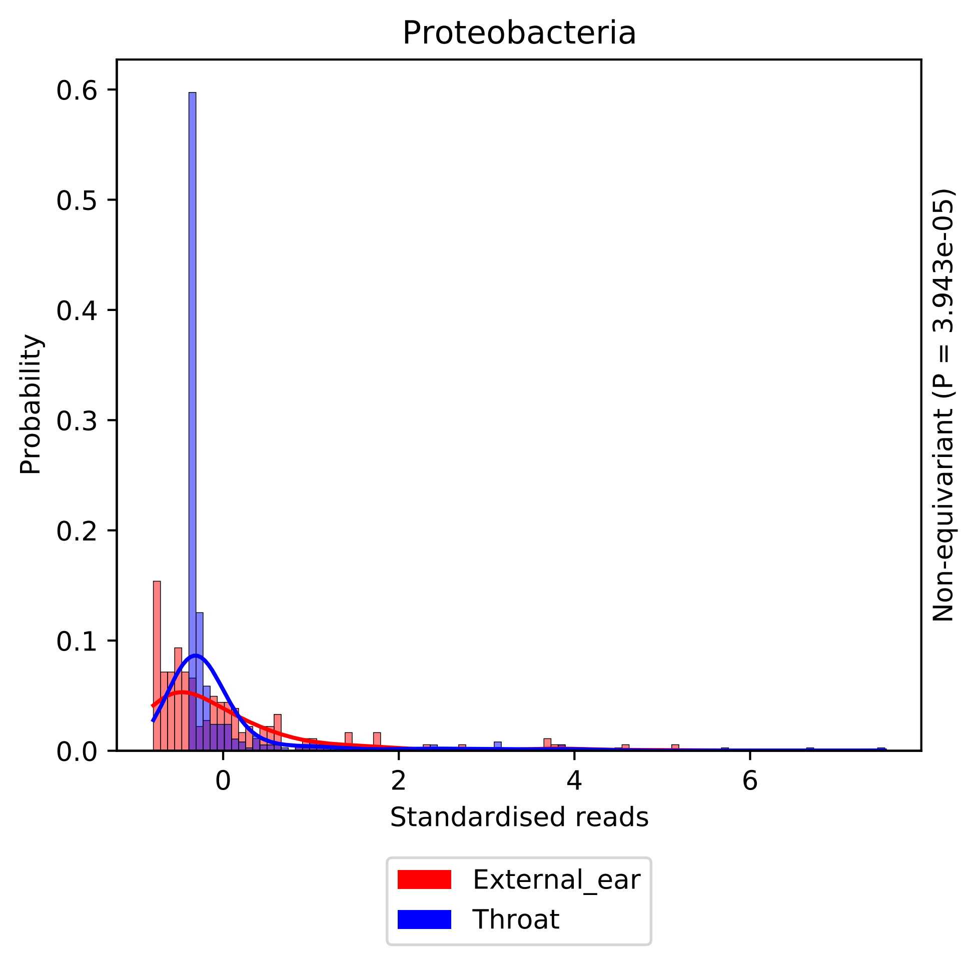
Proteobacteria
Probability
0.6
0.5
0.4
0.3
0.2
0.1
0.0
0
2
4
6
Standardised reads
Non-equivariant (P = 3.943e-05)
External_ear
Throat

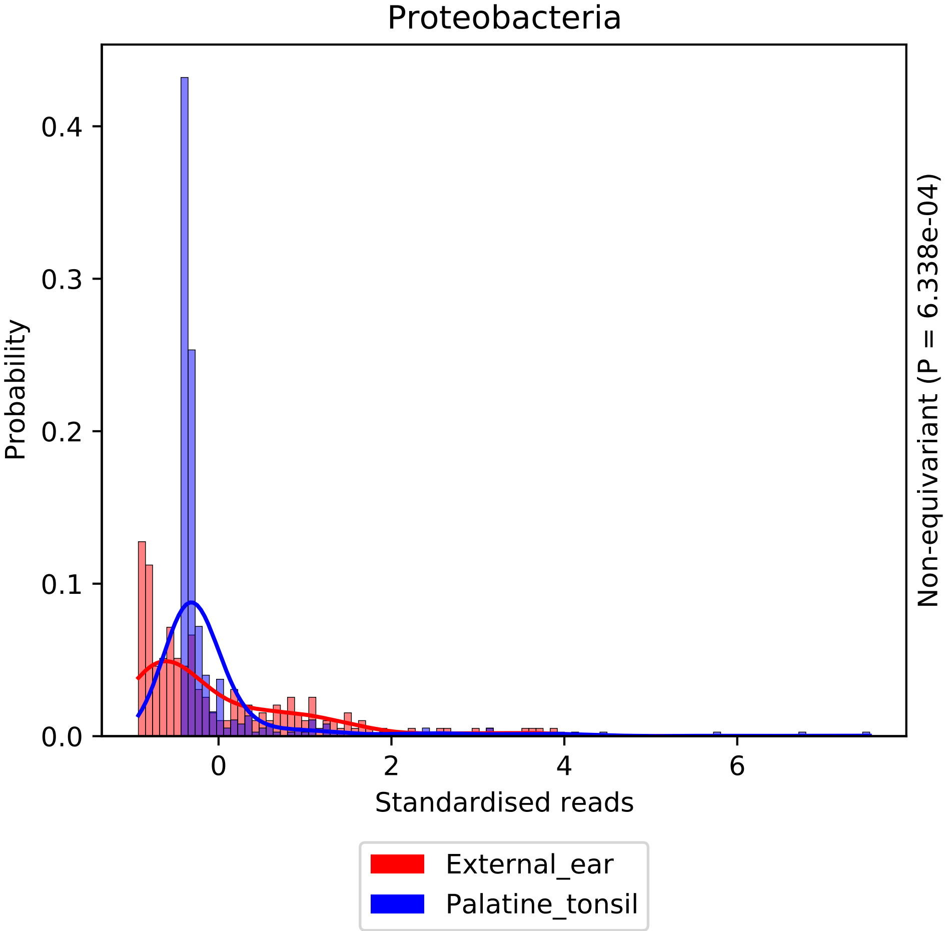
Proteobacteria
Probability
0.0
0.1
0.2
0.3
0.4
0
2
4
6
Standardised reads
Non-equivariant (P = 6.338e-04)
External_ear
Palatine_tonsil

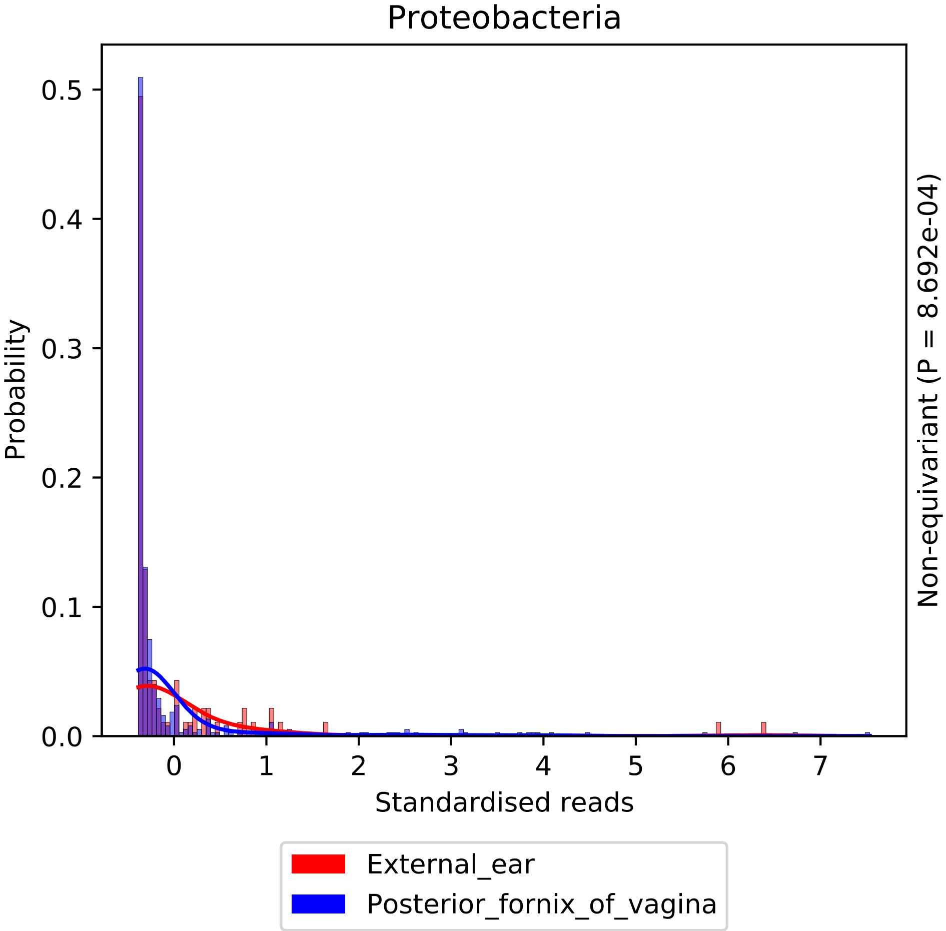

Proteobacteria
Probability
0.5
0.4
0.3
0.2
0.1
0.0
0
1
2
3
4
5
6
7
Standardised reads
Non-equivariant (P = 8.692e-04)
External_ear
Posterior_fornix_of_vagina

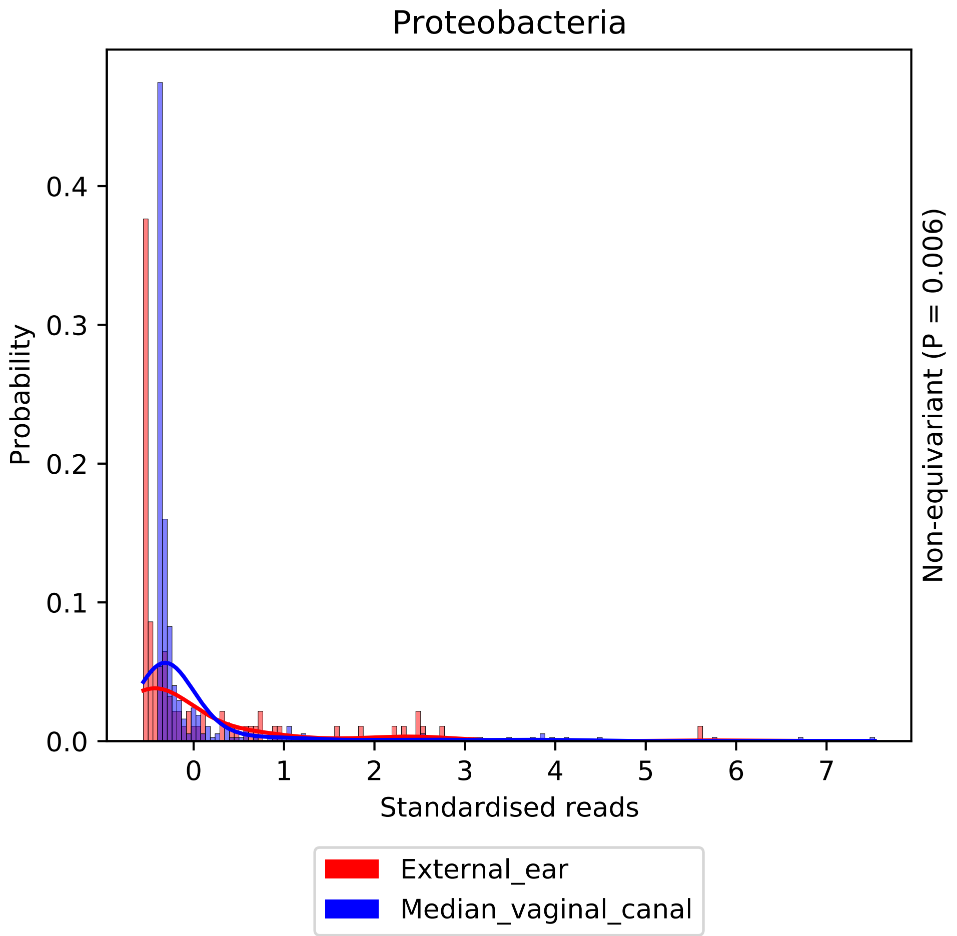

Proteobacteria
Probability
0.0
0.1
0.2
0.3
0.4
0
1
2
3
4
5
6
7
Standardised reads
Non-equivariant (P = 0.006)
External_ear
Median_vaginal_canal

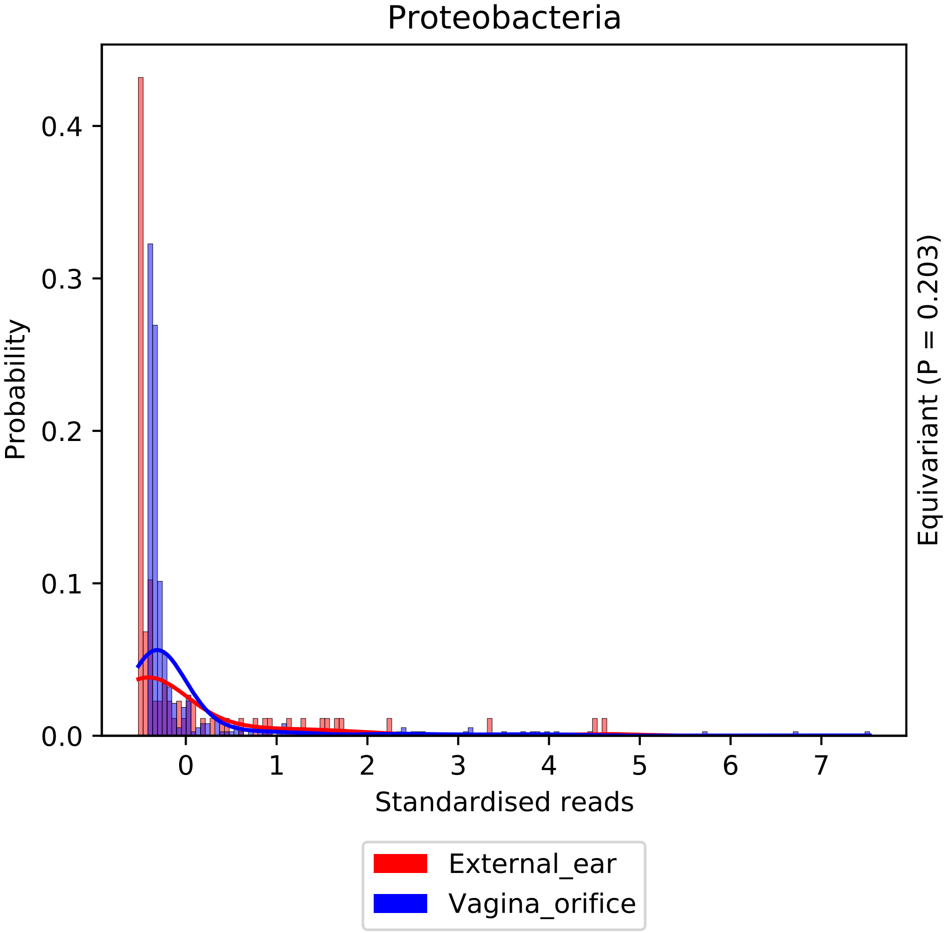

Proteobacteria
Probability
0.0
0.1
0.2
0.3
0.4
0
1
2
3
4
5
6
7
Standardised reads
Equivariant (P = 0.203)
External_ear
Vagina_orifice

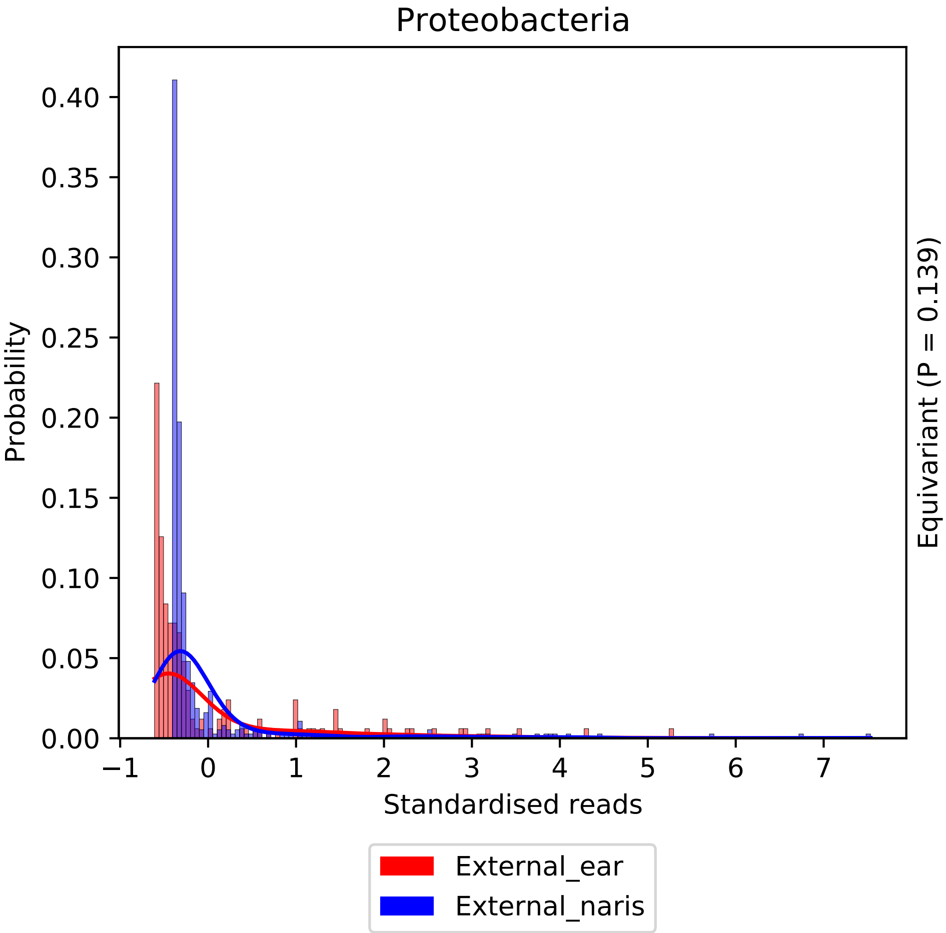

Proteobacteria
Probability
Standardised reads
Equivariant (P = 0.139)
External_ear
External_naris
0.00
0.05
0.10
0.15
0.20
0.25
0.30
0.35
0.40
−1
0
1
2
3
4
5
6
7

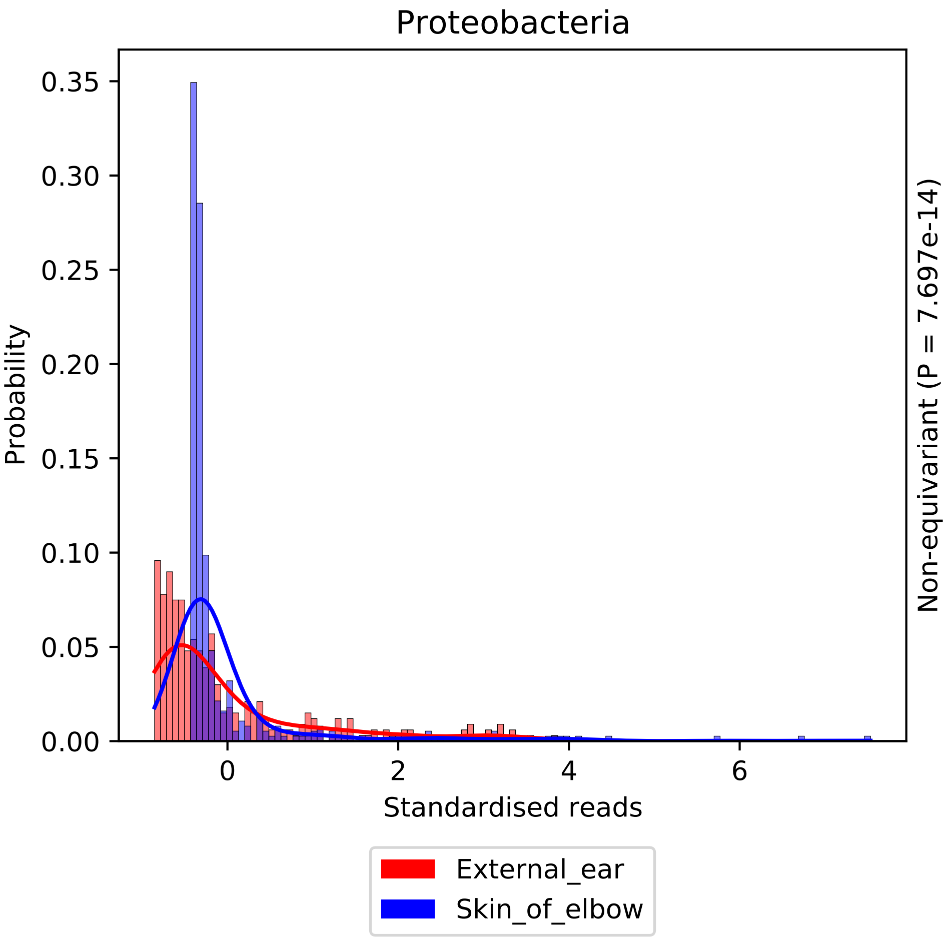

Proteobacteria
Probability
0.35
0.30
0.25
0.20
0.15
0.10
0.05
0.00
0
2
4
6
Standardised reads
Non-equivariant (P = 7.697e-14)
External_ear
Skin_of_elbow

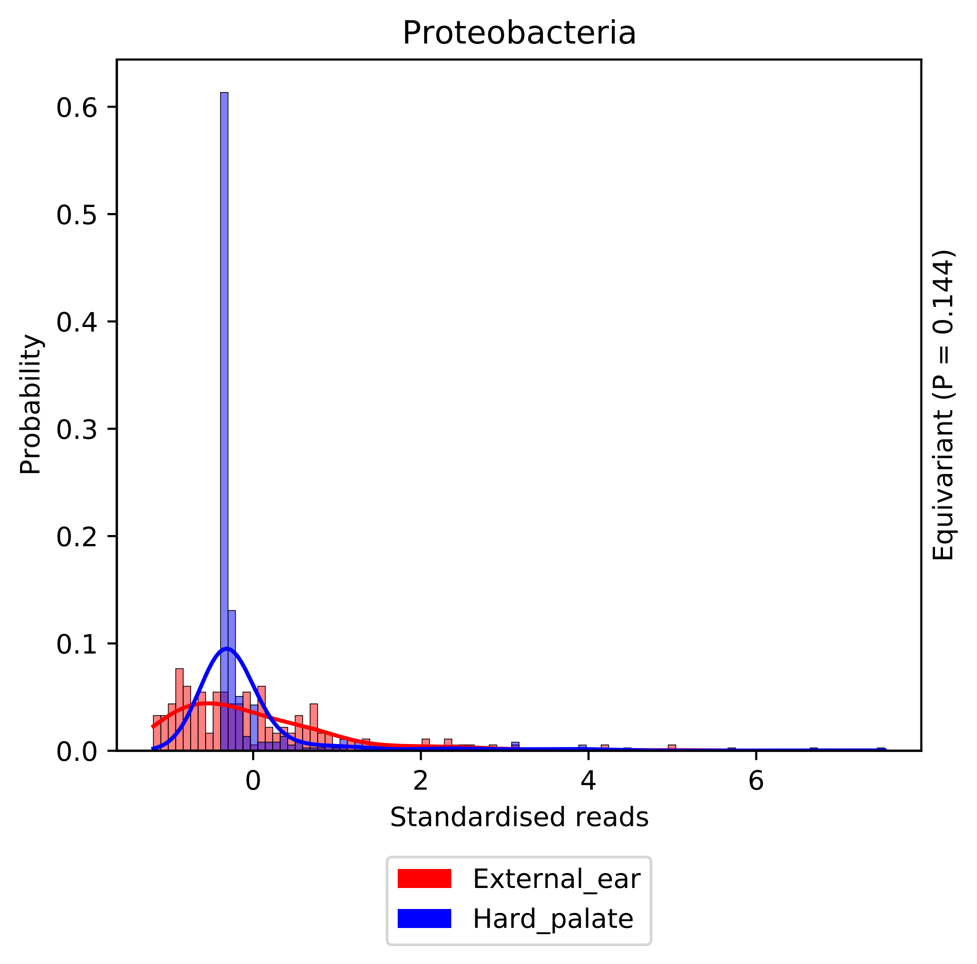

Proteobacteria
Probability
0.6
0.5
0.4
0.3
0.2
0.1
0.0
0
2
4
6
Standardised reads
Equivariant (P = 0.144)
External_ear
Hard_palate

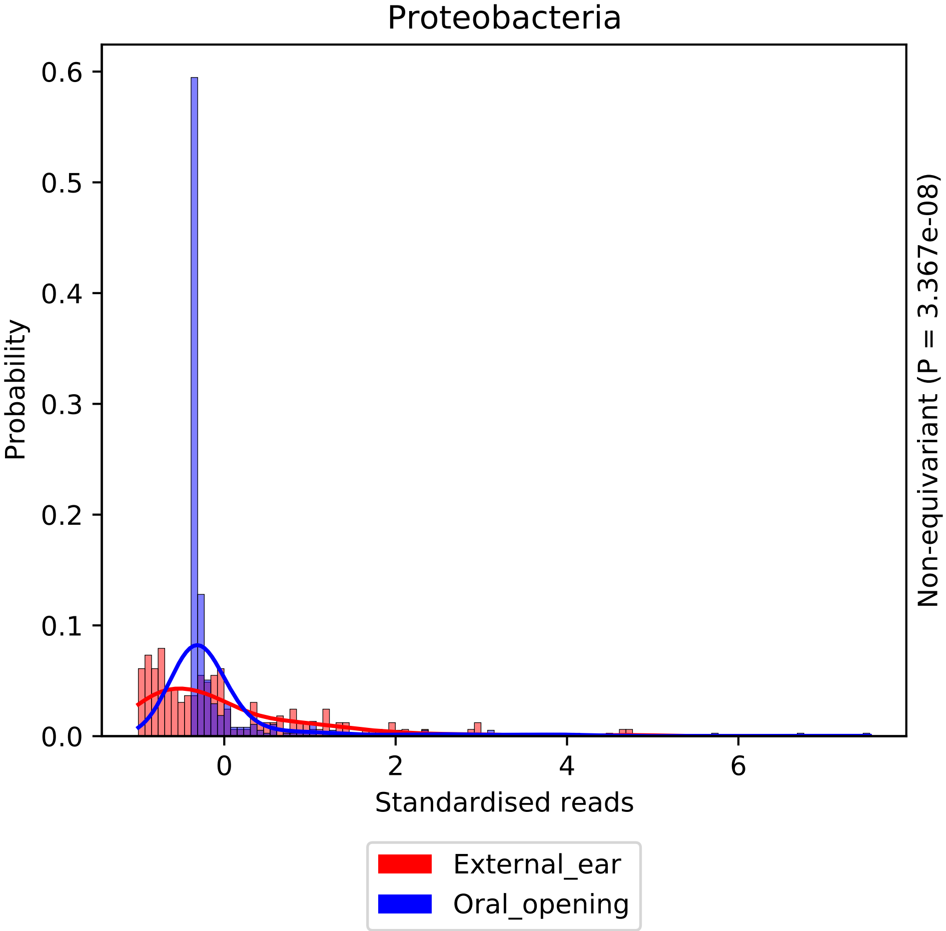

Proteobacteria
Probability
0.6
0.5
0.4
0.3
0.2
0.1
0.0
0
2
4
6
Standardised reads
Non-equivariant (P = 3.367e-08)
External_ear
Oral_opening

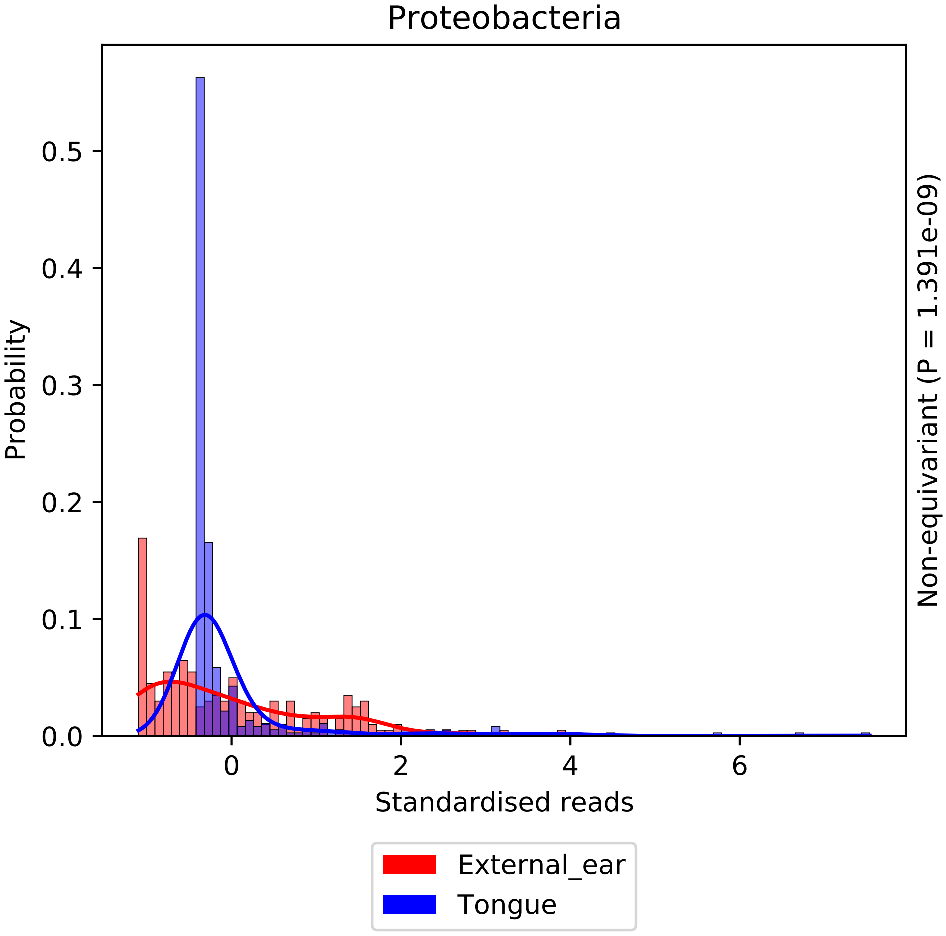

Proteobacteria
Probability
Standardised reads
Non-equivariant (P = 1.391e-09)
External_ear
Tongue
0.0
0.1
0.2
0.3
0.4
0.5
0
2
4
6

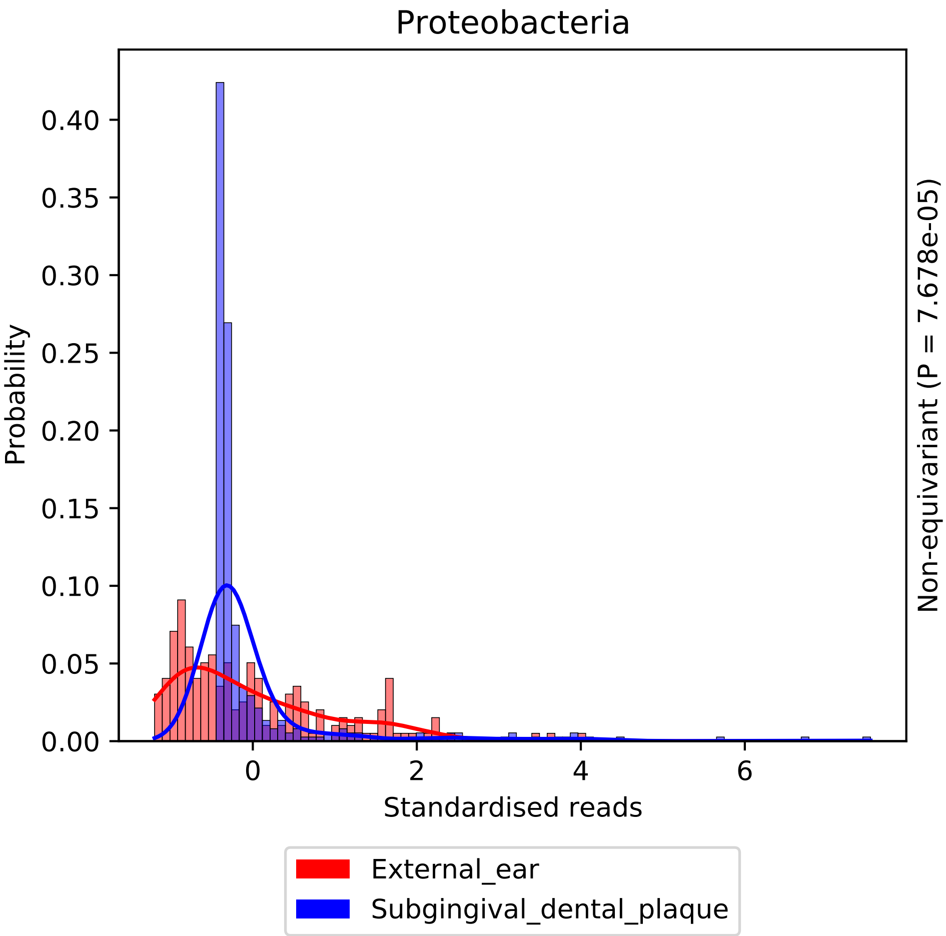

Proteobacteria
Probability
0.00
0.05
0.10
0.15
0.20
0.25
0.30
0.35
0.40
0
2
4
6
Standardised reads
Non-equivariant (P = 7.678e-05)
External_ear
Subgingival_dental_plaque

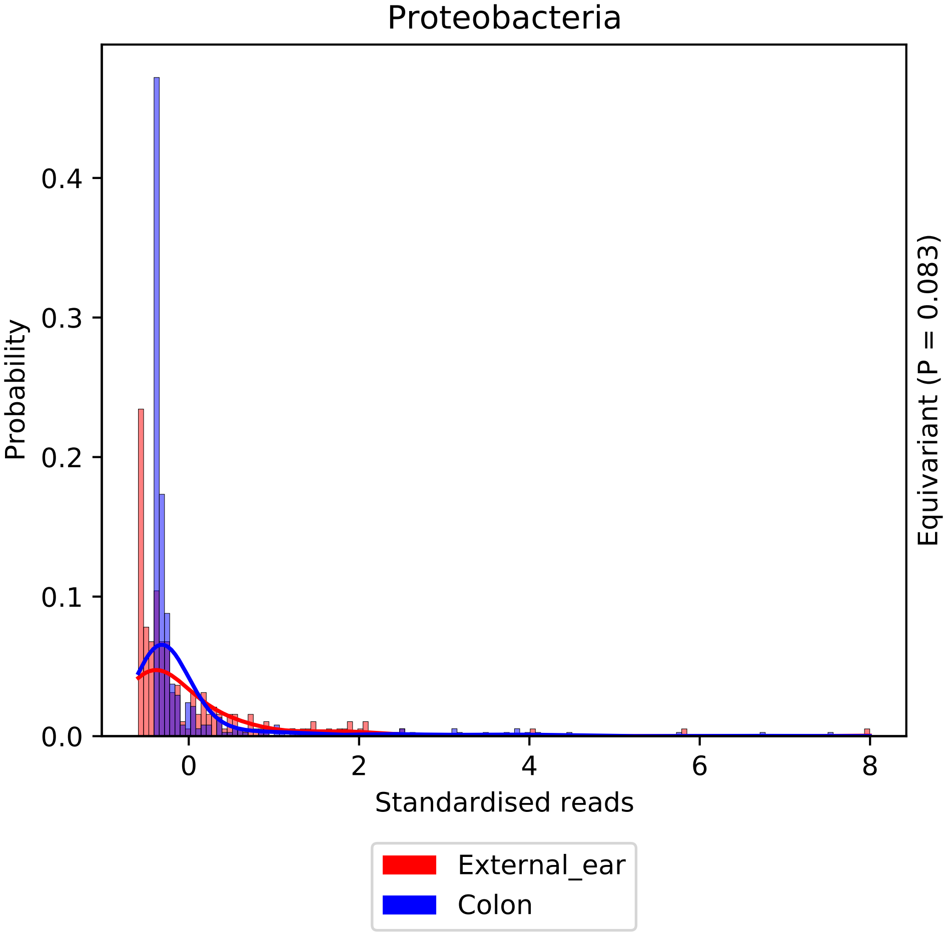
Proteobacteria
Probability
0.0
0.1
0.2
0.3
0.4
0
2
4
6
8
Standardised reads
Equivariant (P = 0.083)
External_ear
Colon

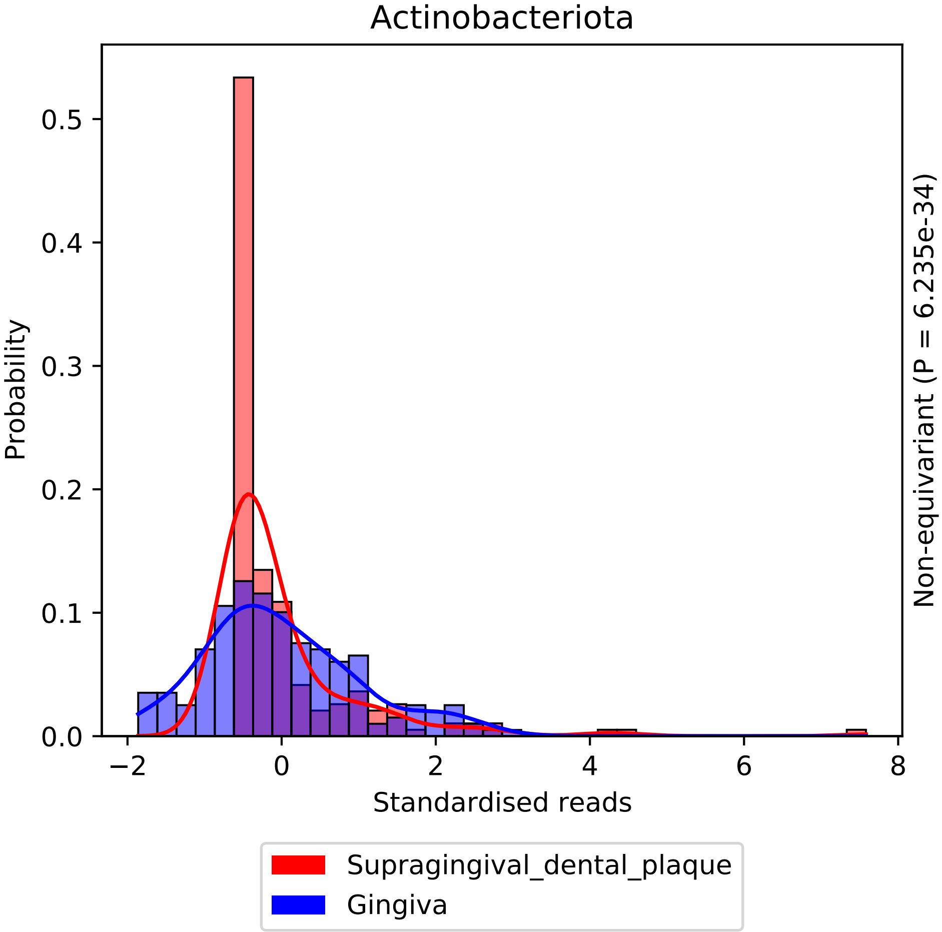
Actinobacteriota
Probability
0.0
0.1
0.2
0.3
0.4
0.5
−2
0
2
4
6
8
Standardised reads
Non-equivariant (P = 6.235e-34)
Supragingival_dental_plaque
Gingiva

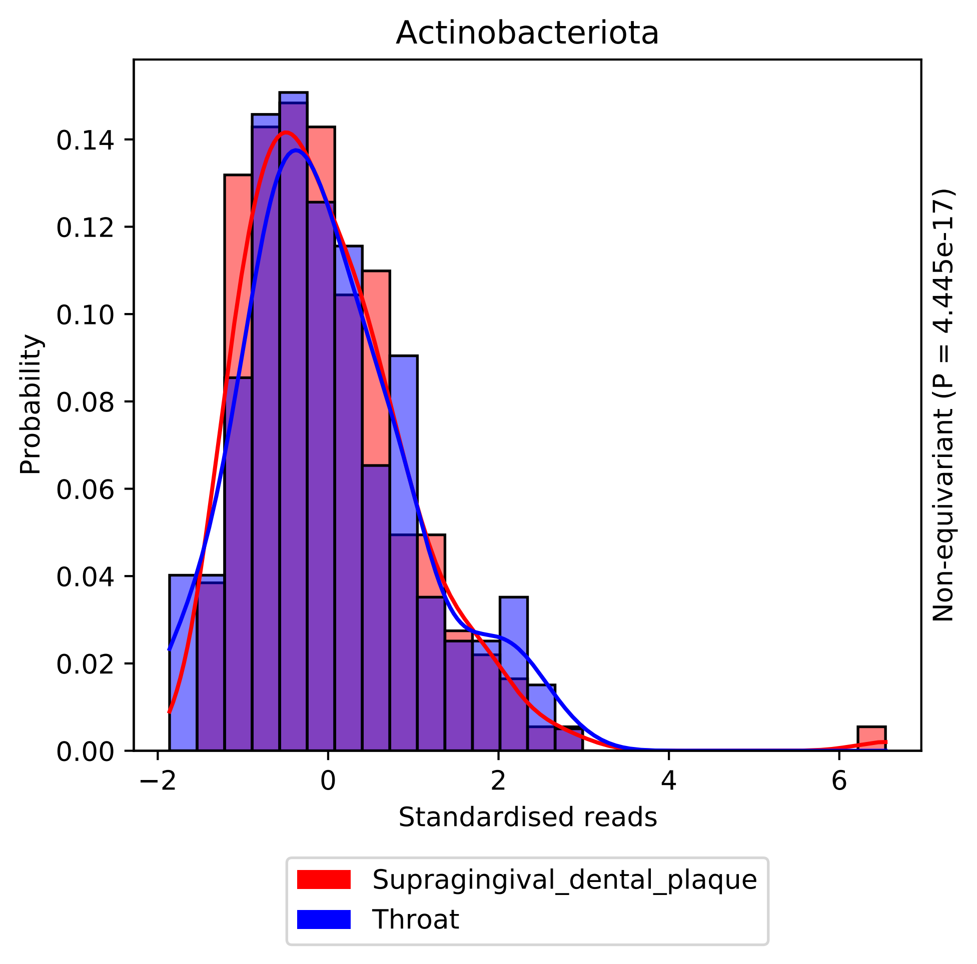

Actinobacteriota
Probability
0.14
0.12
0.10
0.08
0.06
0.04
0.02
0.00
−2
0
2
4
6
Standardised reads
Non-equivariant (P = 4.445e-17)
Supragingival_dental_plaque
Throat

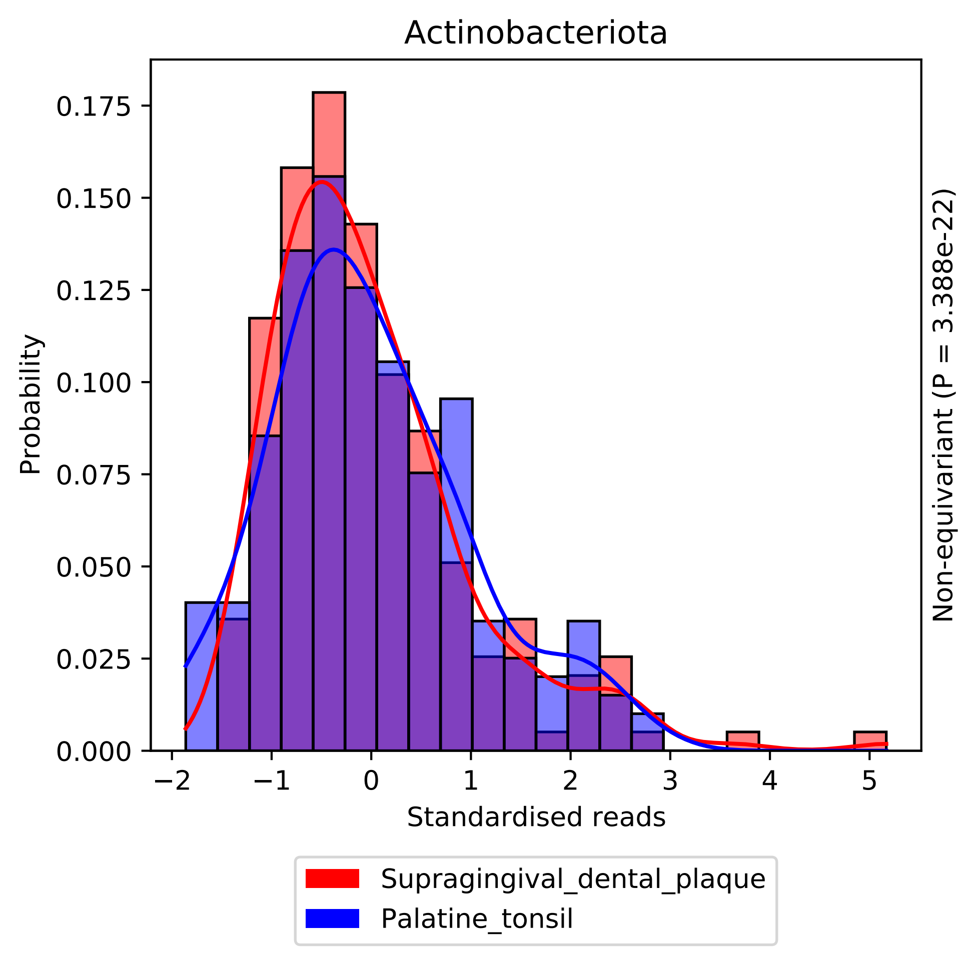
Actinobacteriota
Probability
0.000
0.025
0.050
0.075
0.100
0.125
0.150
0.175
−2
−1
0
1
2
3
4
5
Standardised reads
Non-equivariant (P = 3.388e-22)
Supragingival_dental_plaque
Palatine_tonsil

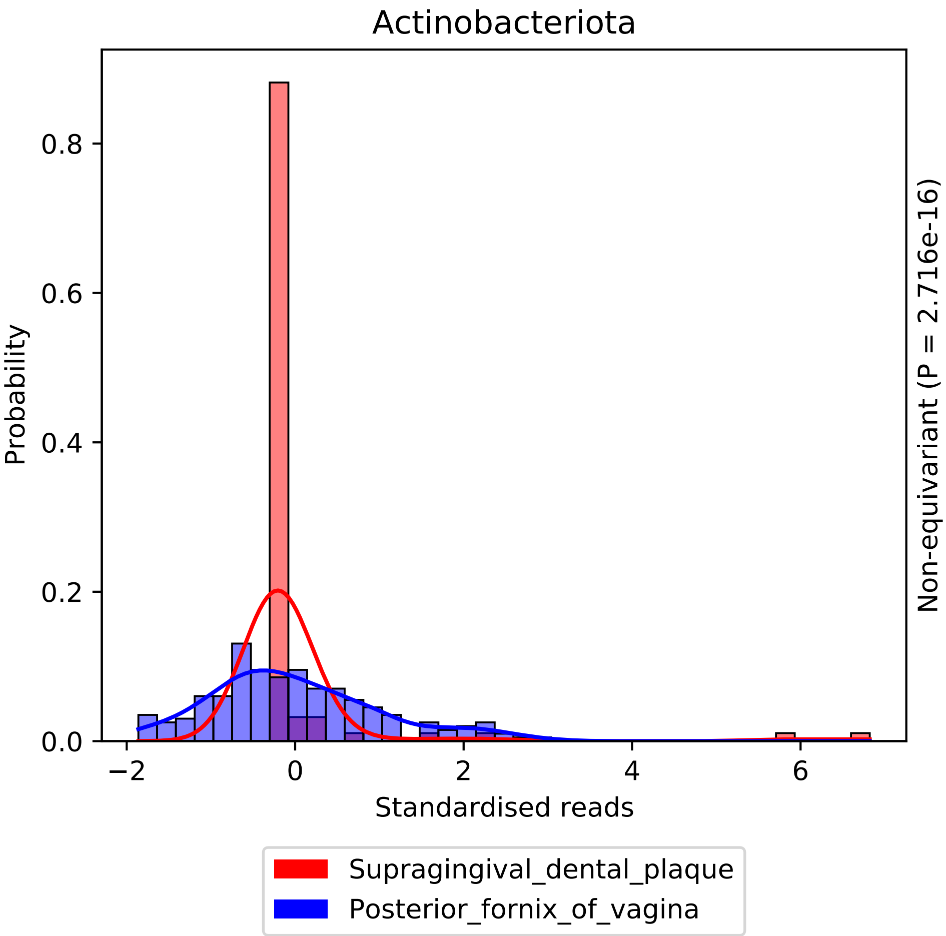

Actinobacteriota
Probability
0.0
0.2
0.4
0.6
0.8
−2
0
2
4
6
Standardised reads
Non-equivariant (P = 2.716e-16)
Supragingival_dental_plaque
Posterior_fornix_of_vagina

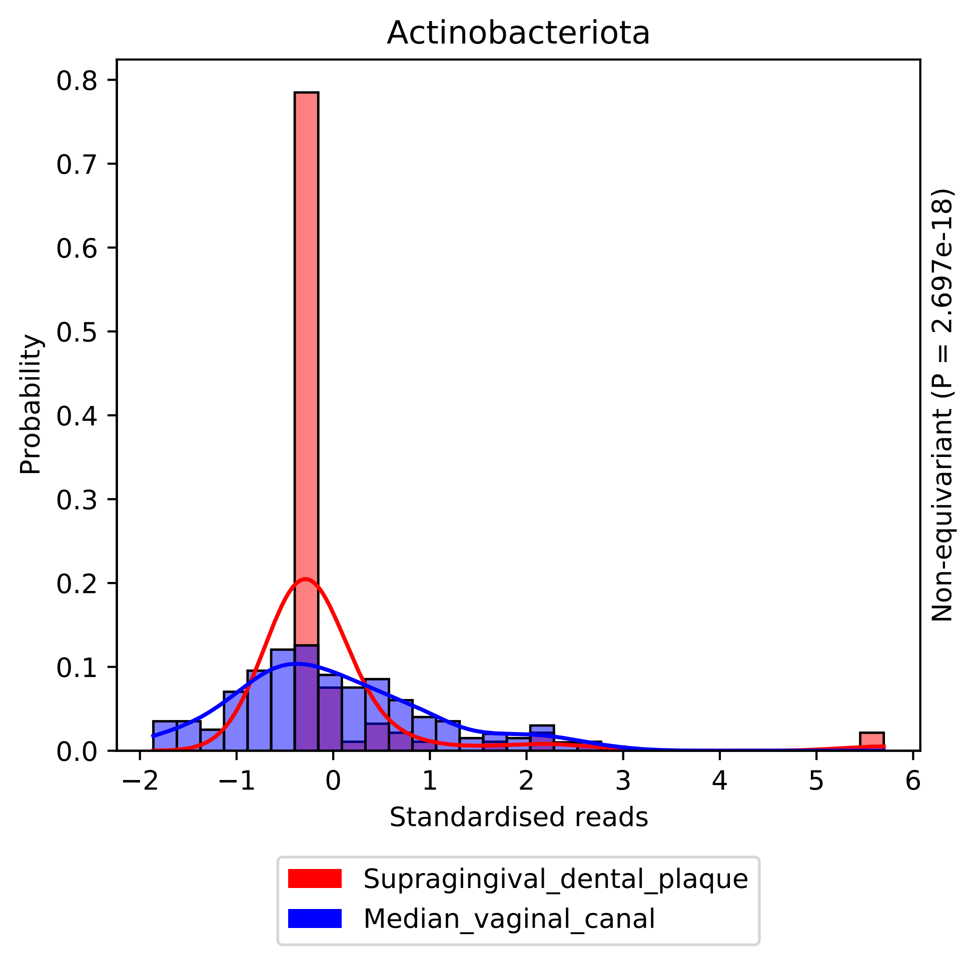

Actinobacteriota
Probability
Standardised reads
Non-equivariant (P = 2.697e-18)
0.0
0.1
0.2
0.3
0.4
0.5
0.6
0.7
0.8
−2
−1
0
1
2
3
4
5
6
Supragingival_dental_plaque
Median_vaginal_canal

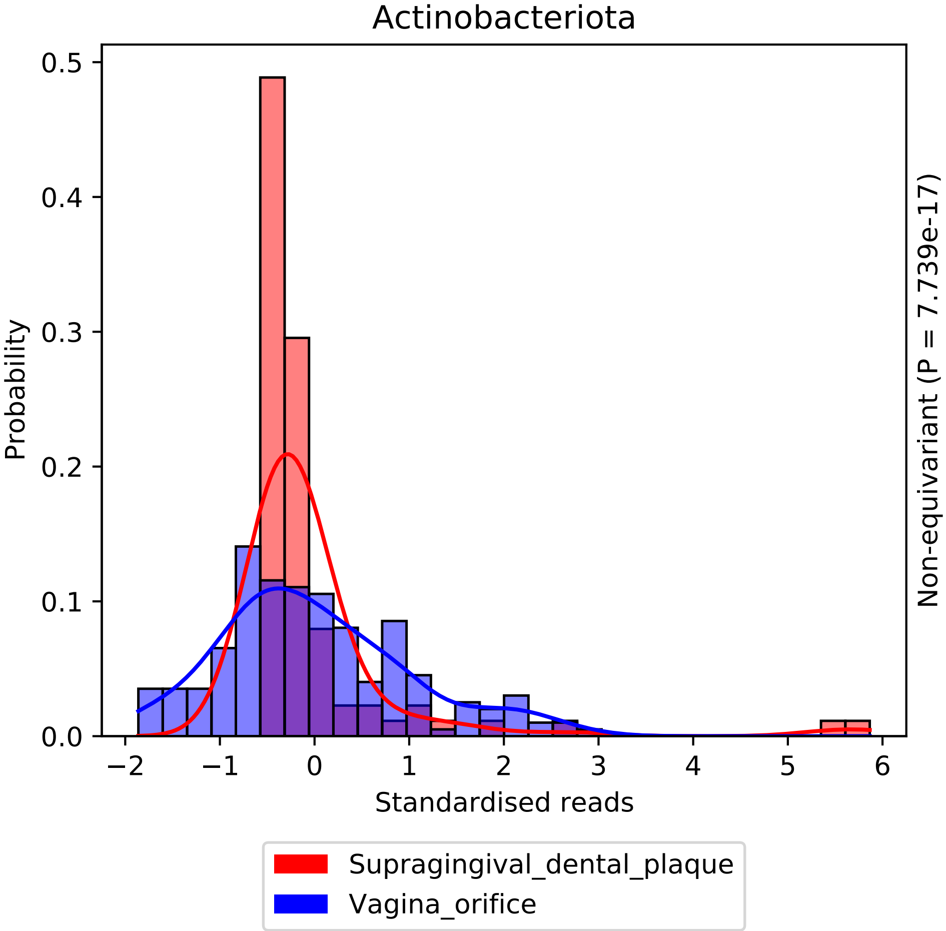
Actinobacteriota
Probability
Standardised reads
Non-equivariant (P = 7.739e-17)
Supragingival_dental_plaque
Vagina_orifice
0.0
0.1
0.2
0.3
0.4
0.5
−2
−1
0
1
2
3
4
5
6

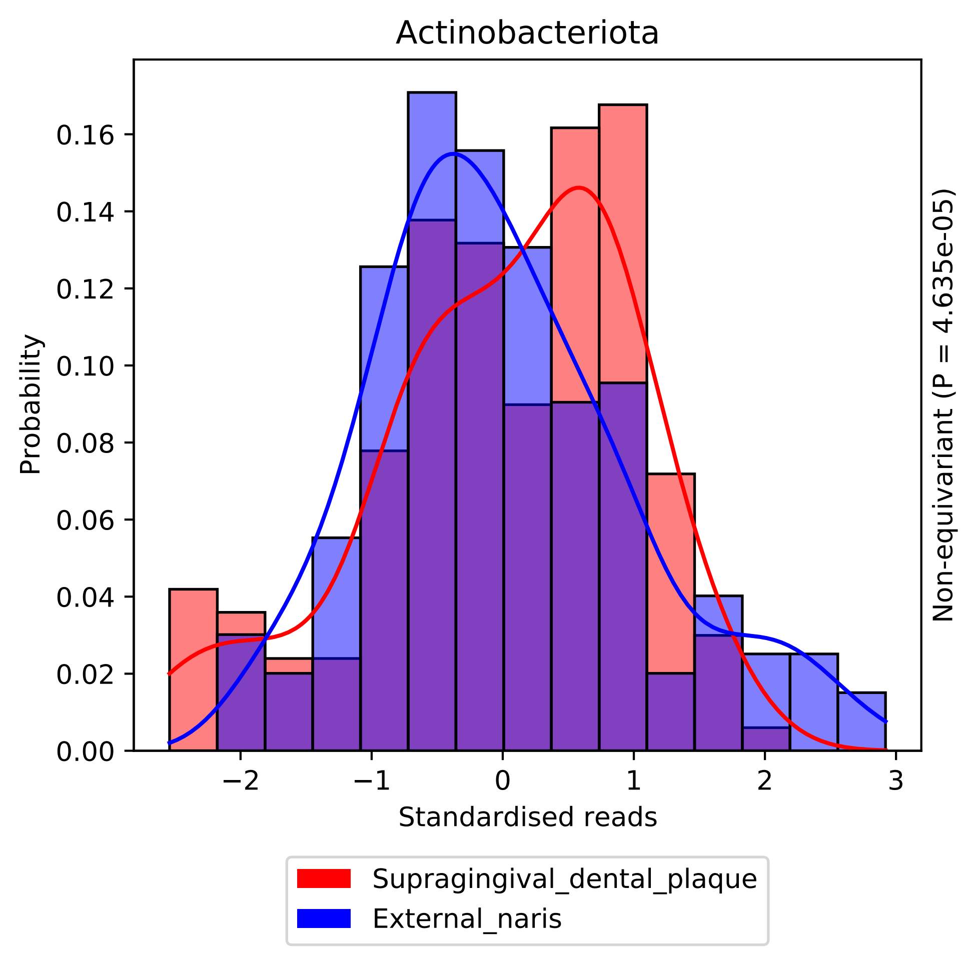

Actinobacteriota
Probability
0.00
0.02
0.04
0.06
0.08
0.10
0.12
0.14
0.16
−2
−1
0
1
2
3
Standardised reads
Non-equivariant (P = 4.635e-05)
Supragingival_dental_plaque
External_naris

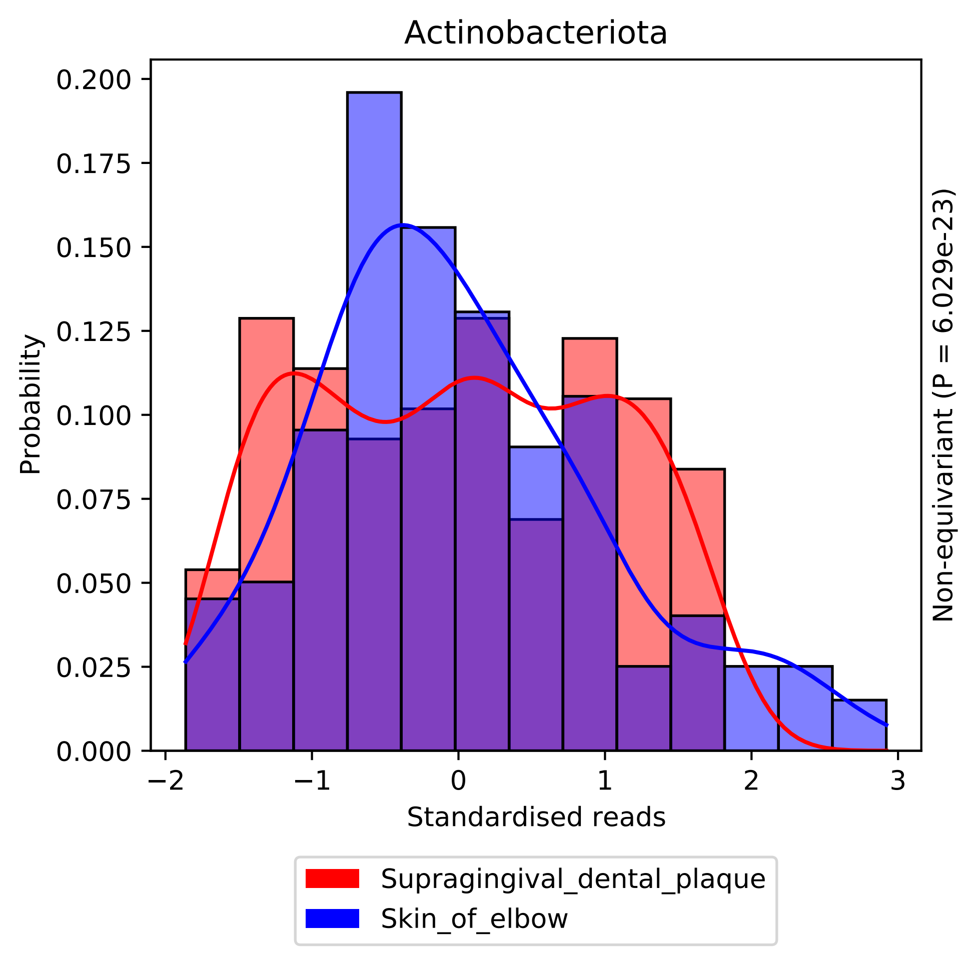

Actinobacteriota
Probability
0.200
0.175
0.150
0.125
0.100
0.075
0.050
0.025
0.000
−2
−1
0
1
2
3
Standardised reads
Non-equivariant (P = 6.029e-23)
Supragingival_dental_plaque
Skin_of_elbow

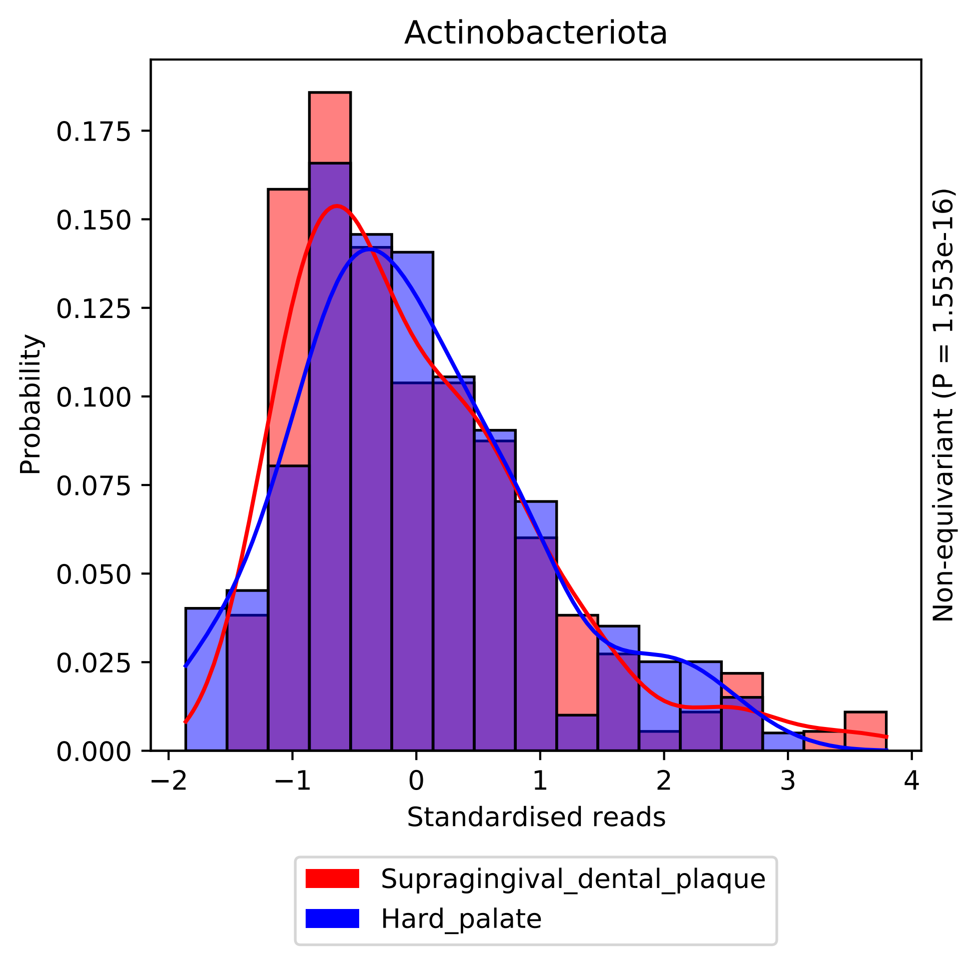

Actinobacteriota
Probability
0.000
0.025
0.050
0.075
0.100
0.125
0.150
0.175
−2
−1
0
1
2
3
4
Standardised reads
Non-equivariant (P = 1.553e-16)
Supragingival_dental_plaque
Hard_palate

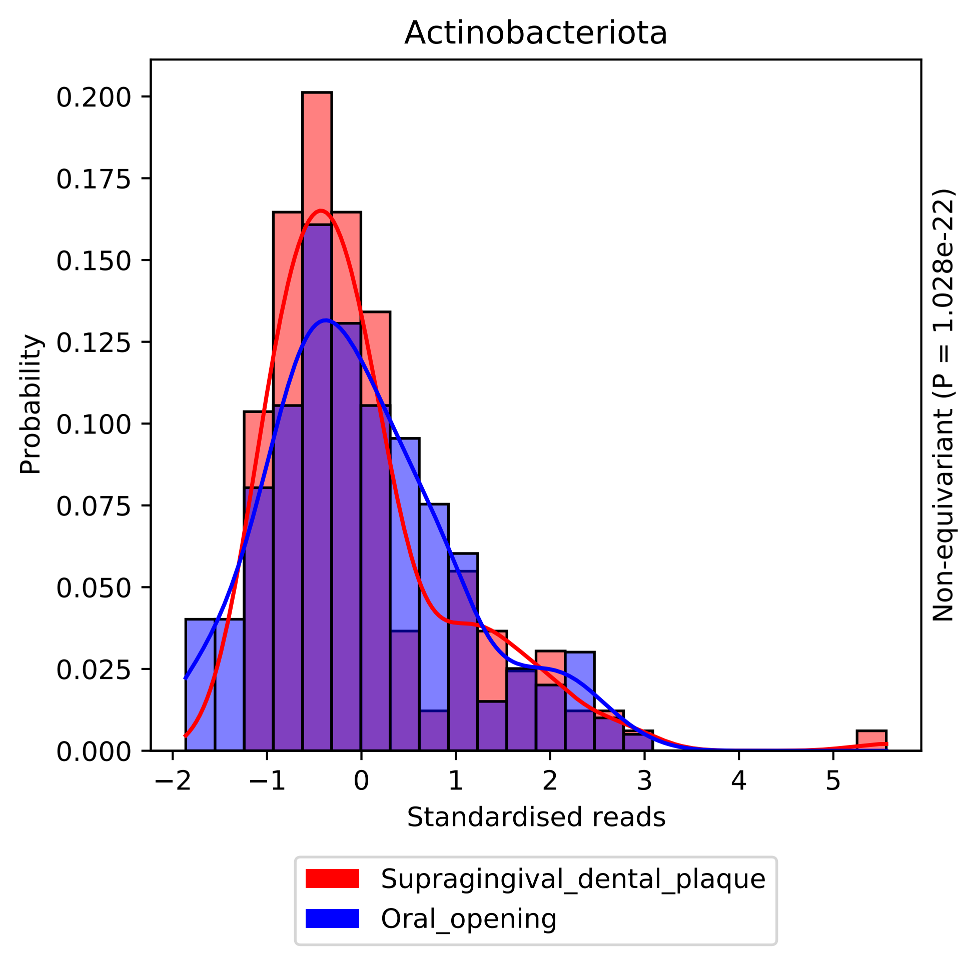
Actinobacteriota
Probability
0.200
0.175
0.150
0.125
0.100
0.075
0.050
0.025
0.000
−2
−1
0
1
2
3
4
5
Standardised reads
Non-equivariant (P = 1.028e-22)
Supragingival_dental_plaque
Oral_opening

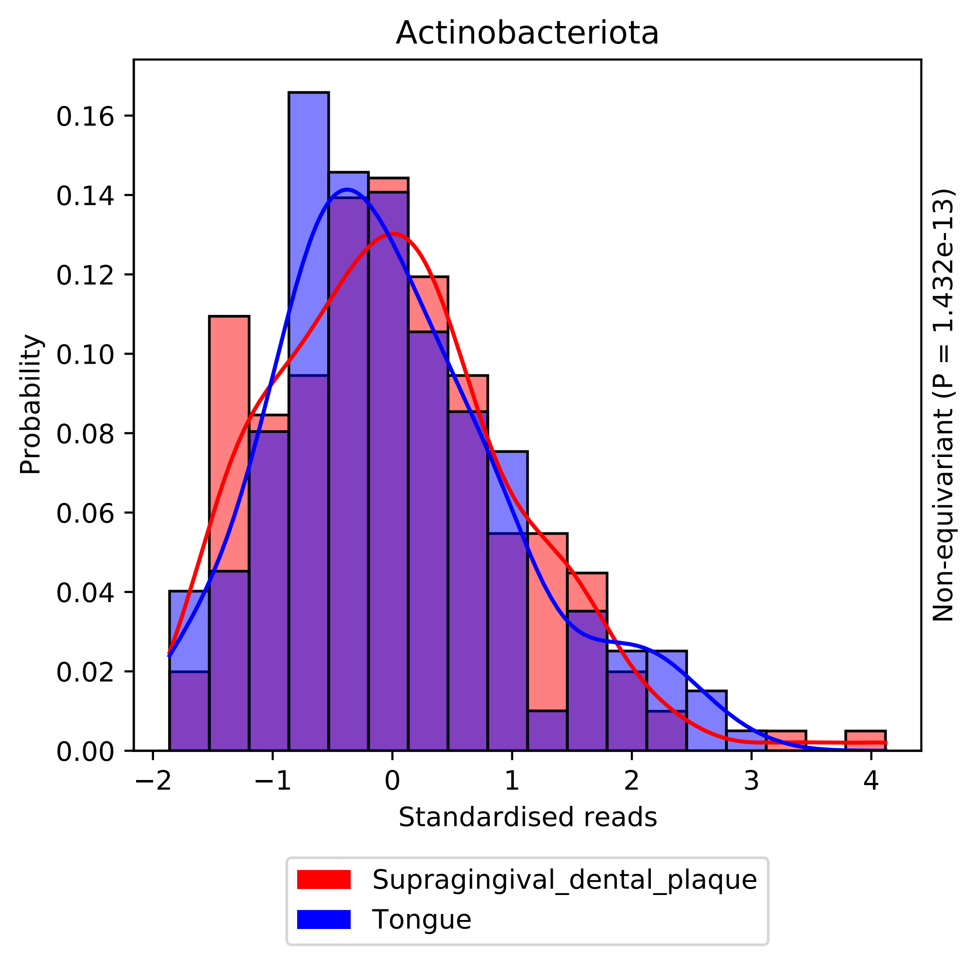

Actinobacteriota
Probability
Standardised reads
0.00
0.02
0.04
0.06
0.08
0.10
0.12
0.14
0.16
−2
−1
0
1
2
3
4
Non-equivariant (P = 1.432e-13)
Supragingival_dental_plaque
Tongue

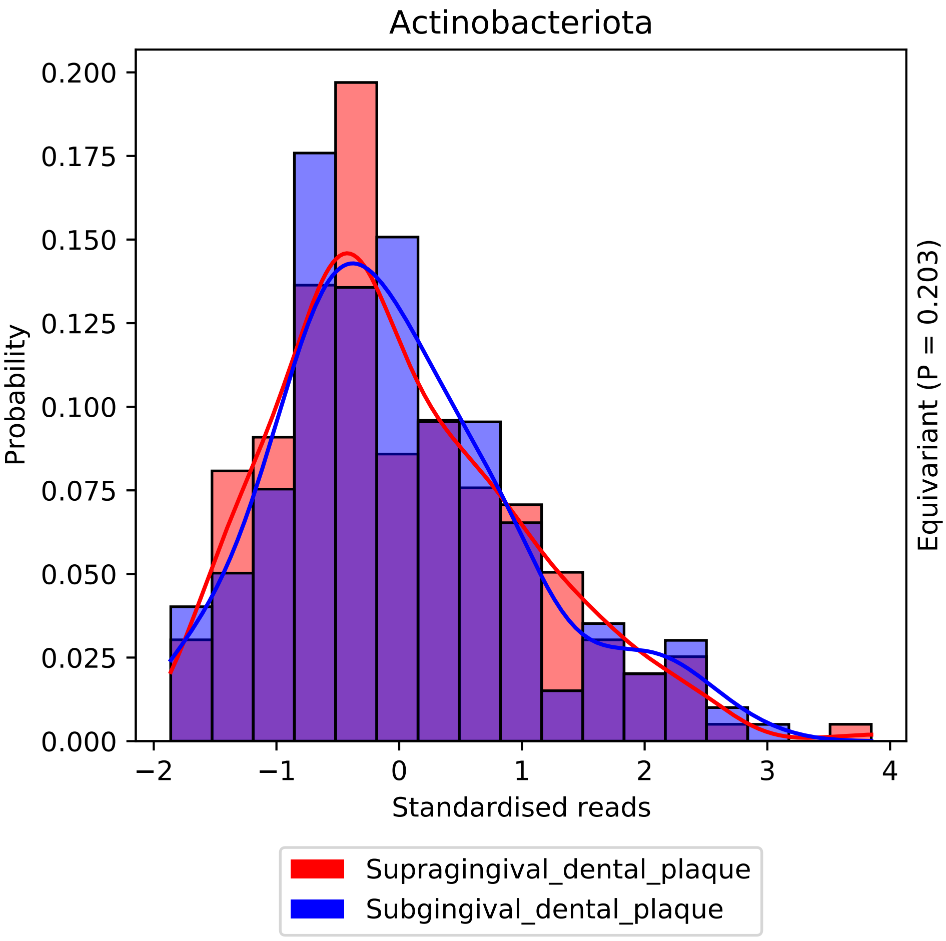
Actinobacteriota
Probability
0.200
0.175
0.150
0.125
0.100
0.075
0.050
0.025
0.000
−2
−1
0
1
2
3
4
Standardised reads
Equivariant (P = 0.203)
Supragingival_dental_plaque
Subgingival_dental_plaque

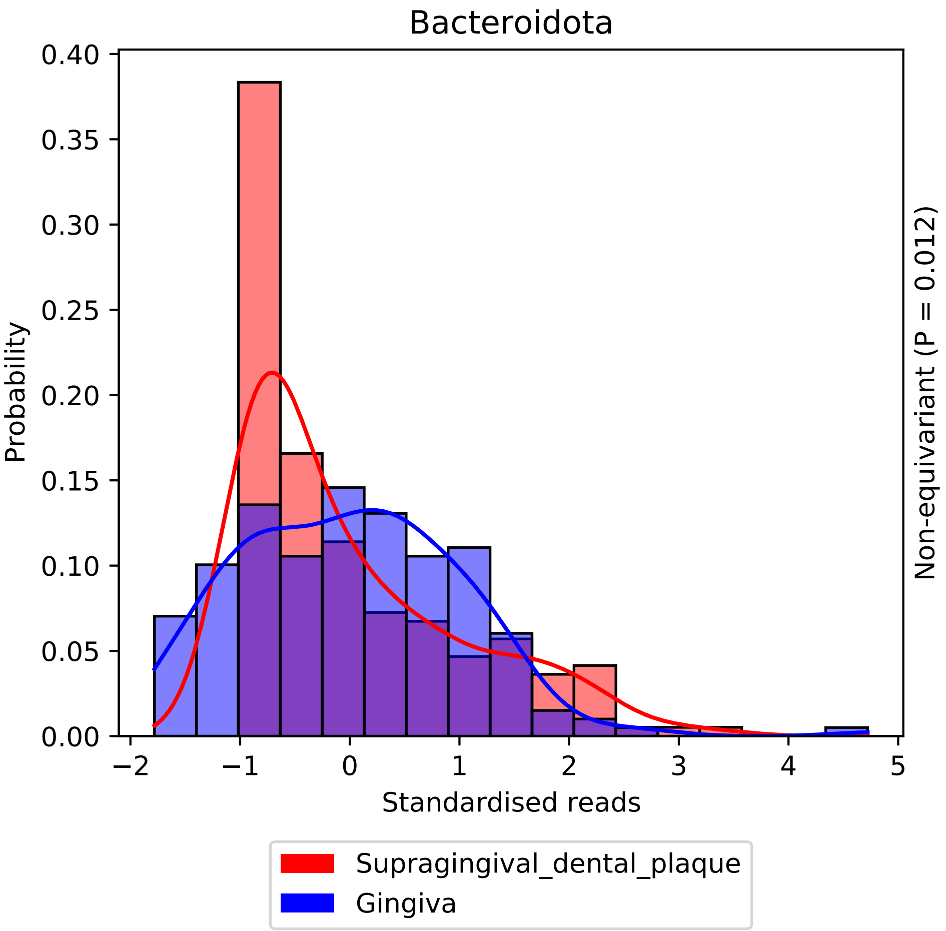
Bacteroidota
Probability
0.40
0.35
0.30
0.25
0.20
0.15
0.10
0.05
0.00
−2
−1
0
1
2
3
4
5
Standardised reads
Non-equivariant (P = 0.012)
Supragingival_dental_plaque
Gingiva

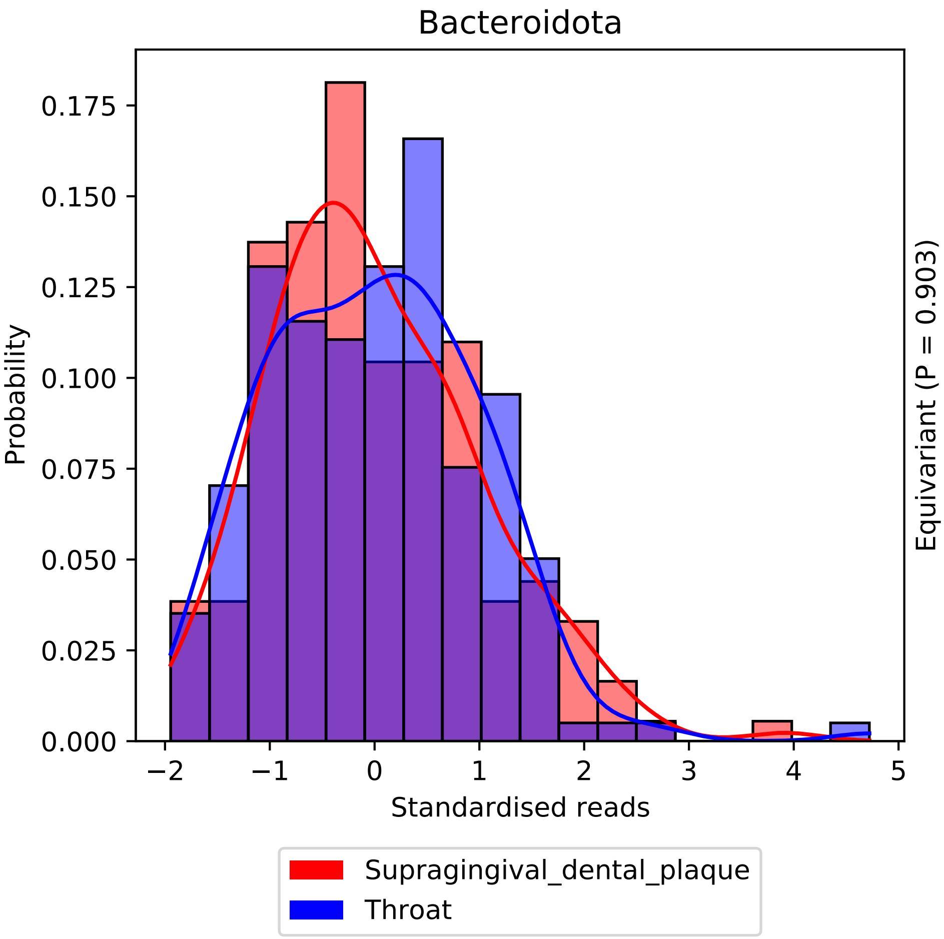
Bacteroidota
Probability
0.000
0.025
0.050
0.075
0.100
0.125
0.150
0.175
−2
−1
0
1
2
3
4
5
Standardised reads
Equivariant (P = 0.903)
Supragingival_dental_plaque
Throat

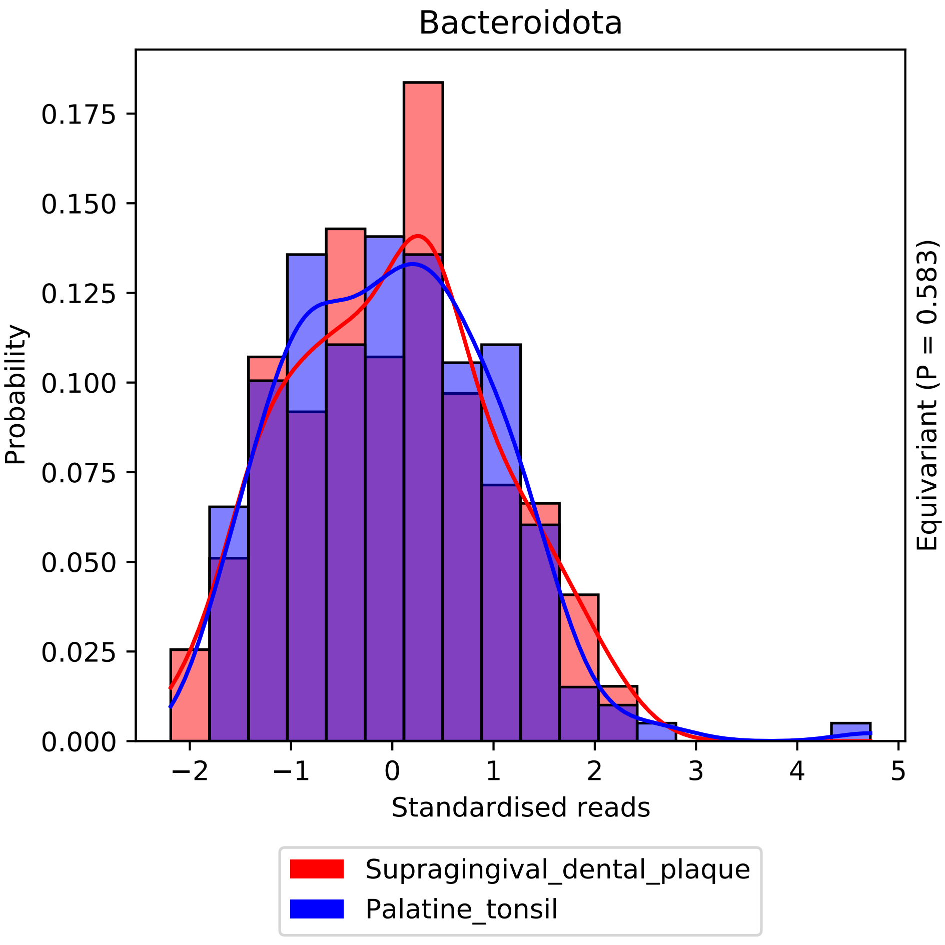
Bacteroidota
Probability
Standardised reads
Equivariant (P = 0.583)
0.000
0.025
0.050
0.075
0.100
0.125
0.150
0.175
−2
−1
0
1
2
3
4
5
Supragingival_dental_plaque
Palatine_tonsil

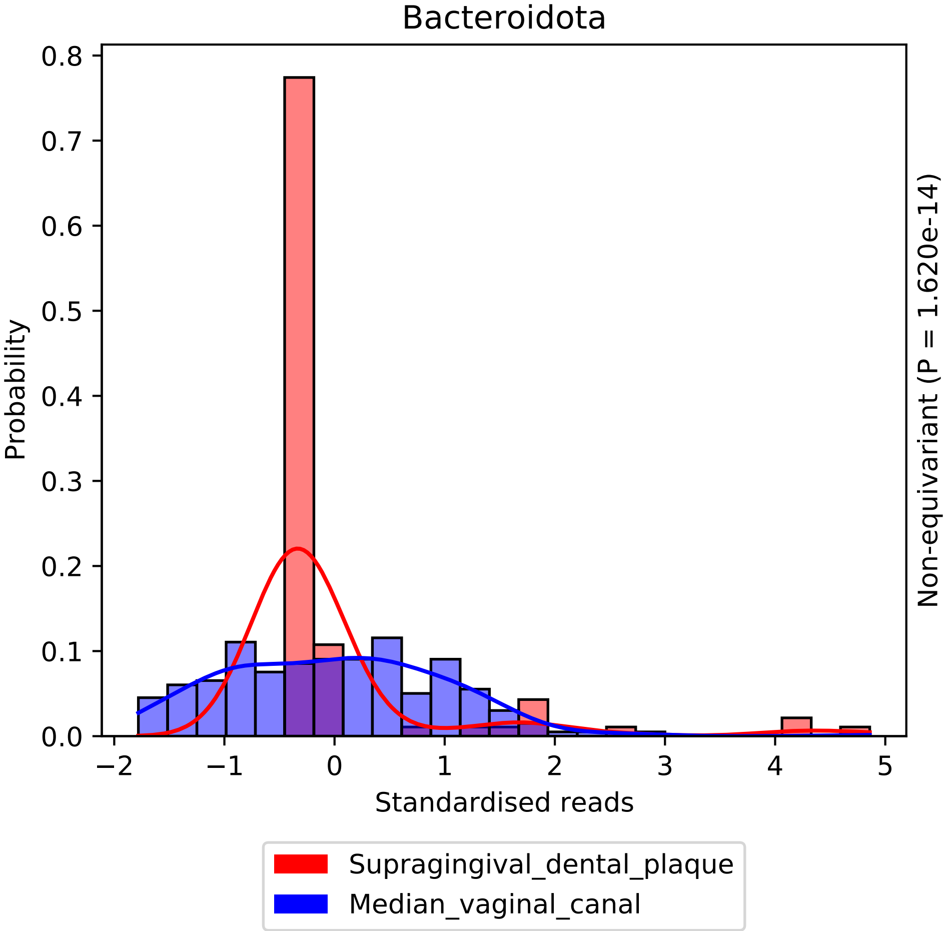

Bacteroidota
Probability
0.0
0.1
0.2
0.3
0.4
0.5
0.6
0.7
0.8
−2
−1
0
1
2
3
4
5
Standardised reads
Non-equivariant (P = 1.620e-14)
Supragingival_dental_plaque
Median_vaginal_canal

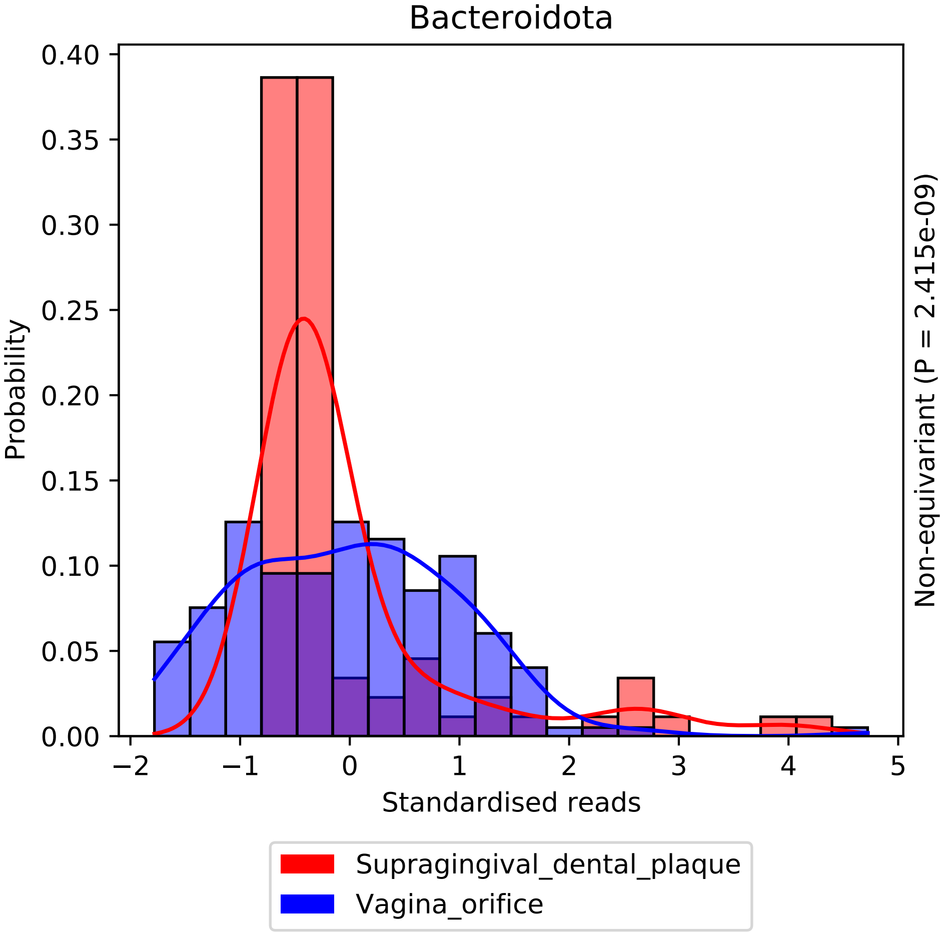

Bacteroidota
0.40
0.35
0.30
0.25
0.20
0.15
0.10
0.05
0.00
Probability
−2
−1
0
1
2
3
4
5
Standardised reads
Non-equivariant (P = 2.415e-09)
Supragingival_dental_plaque
Vagina_orifice

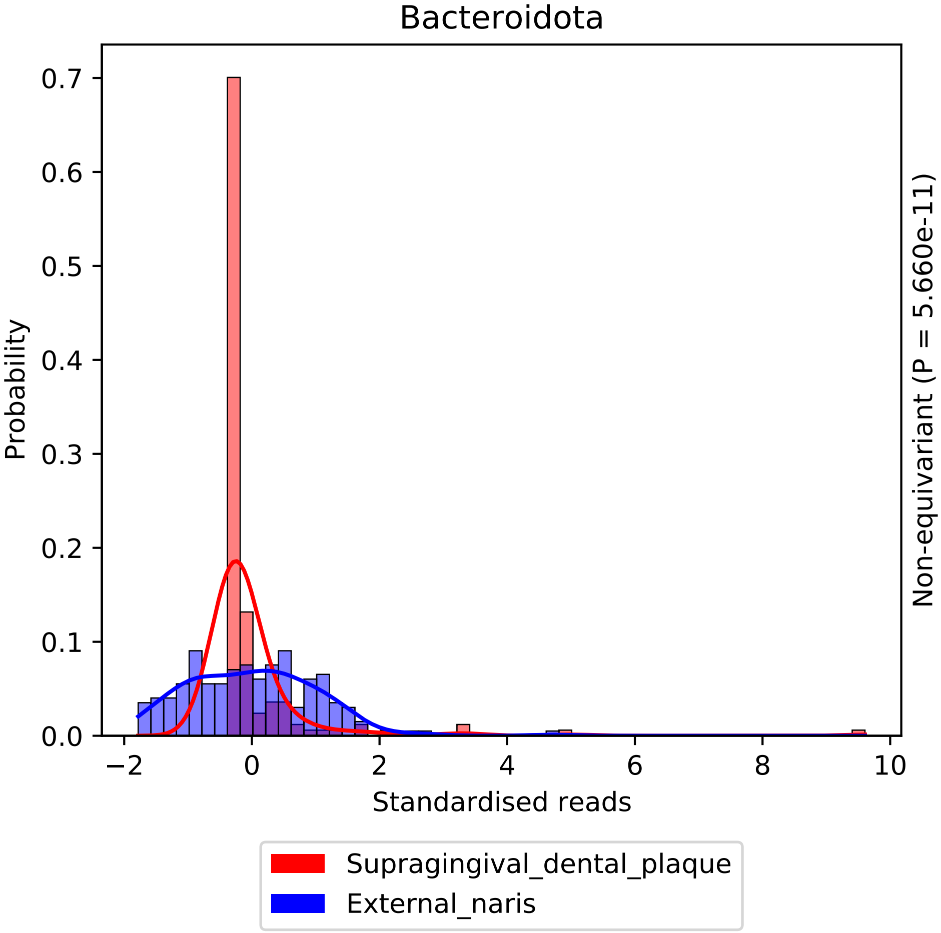

Bacteroidota
Probability
Standardised reads
Non-equivariant (P = 5.660e-11)
Supragingival_dental_plaque
External_naris
0.0
0.1
0.2
0.3
0.4
0.5
0.6
0.7
−2
0
2
4
6
8
10

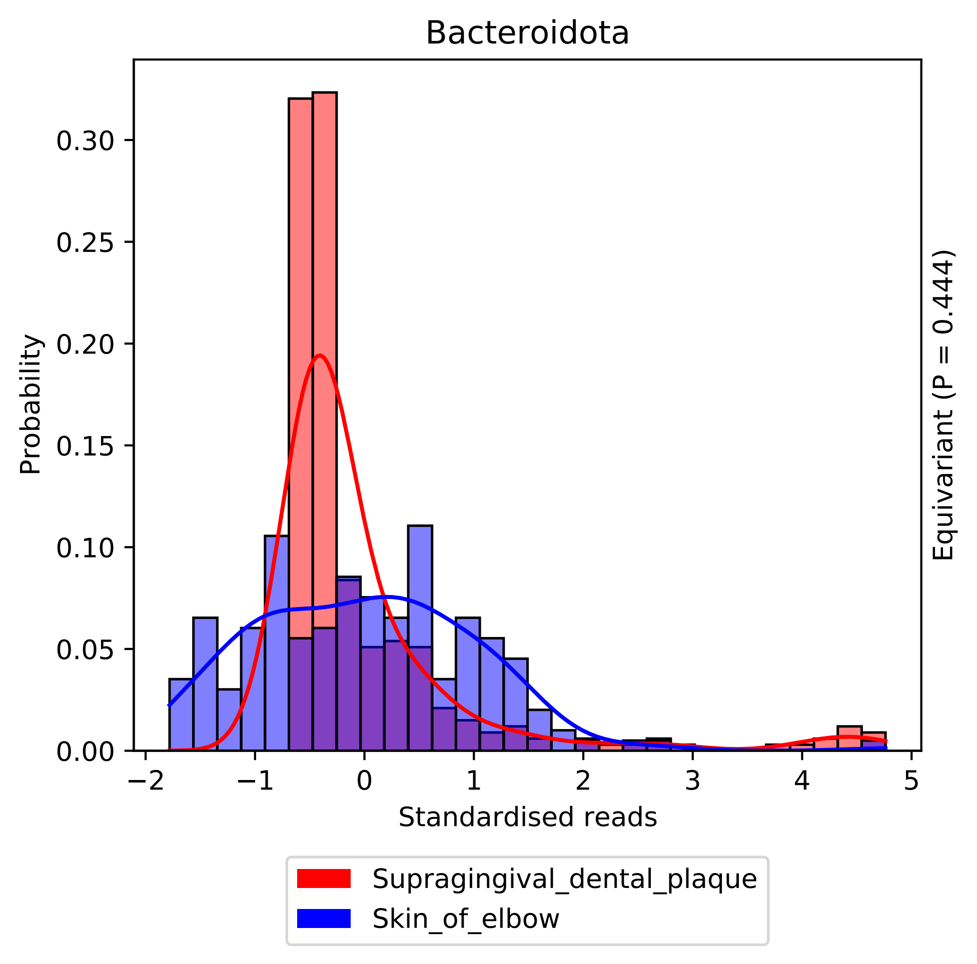

Bacteroidota
Probability
0.30
0.25
0.20
0.15
0.10
0.05
0.00
−2
−1
0
1
2
3
4
5
Standardised reads
Equivariant (P = 0.444)
Supragingival_dental_plaque
Skin_of_elbow

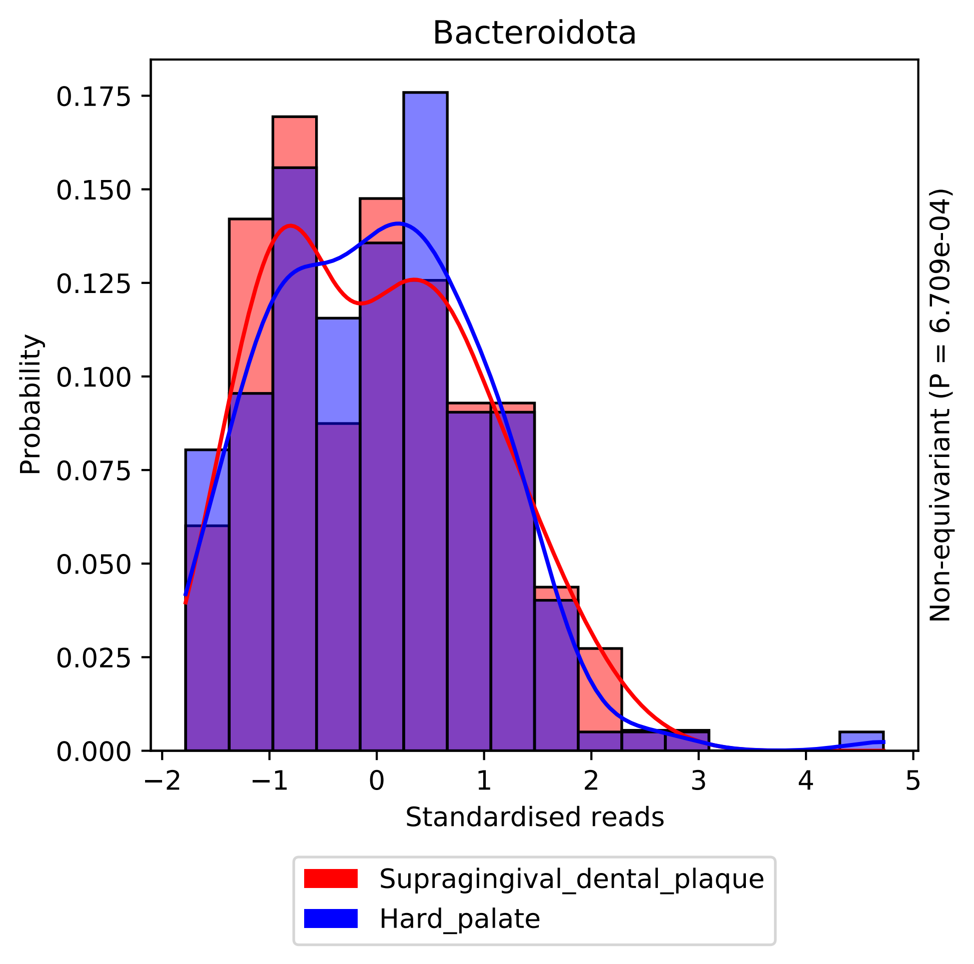

Bacteroidota
Probability
0.000
0.025
0.050
0.075
0.100
0.125
0.150
0.175
−2
−1
0
1
2
3
4
5
Standardised reads
Non-equivariant (P = 6.709e-04)
Supragingival_dental_plaque
Hard_palate

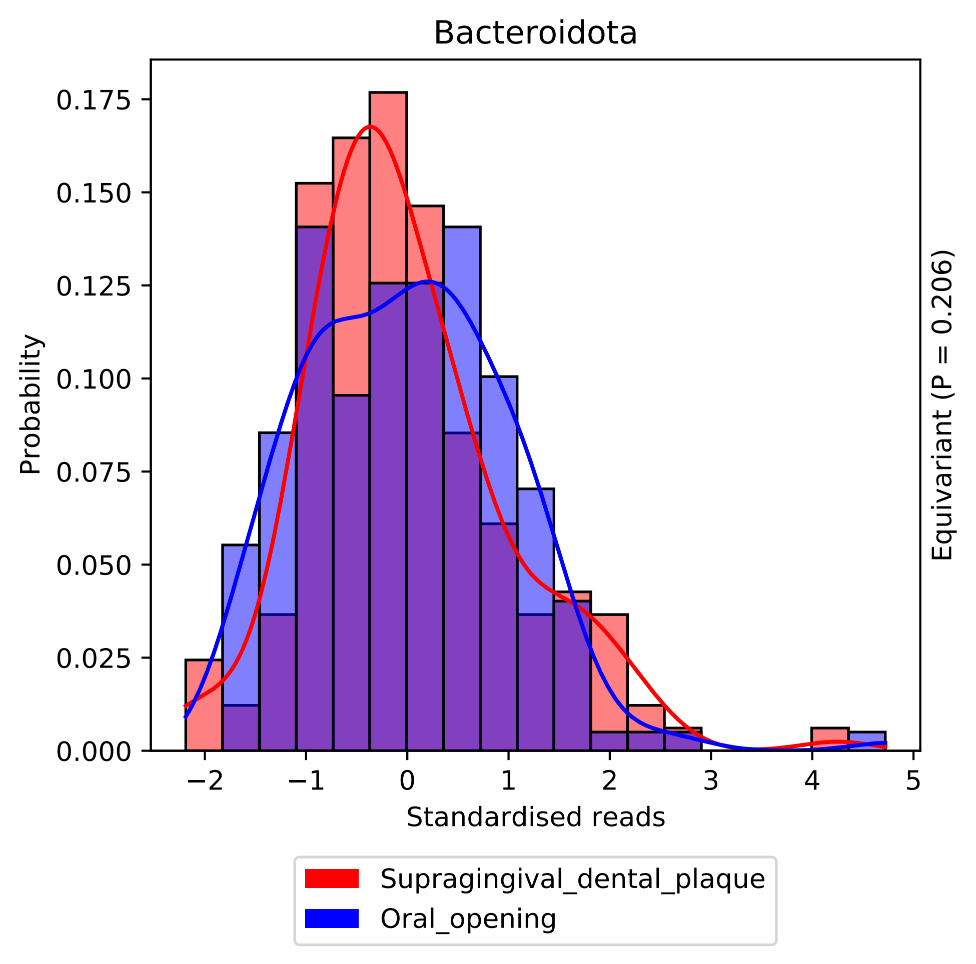

Bacteroidota
Probability
Standardised reads
Equivariant (P = 0.206)
0.000
0.025
0.050
0.075
0.100
0.125
0.150
0.175
−2
−1
0
1
2
3
4
5
Supragingival_dental_plaque
Oral_opening

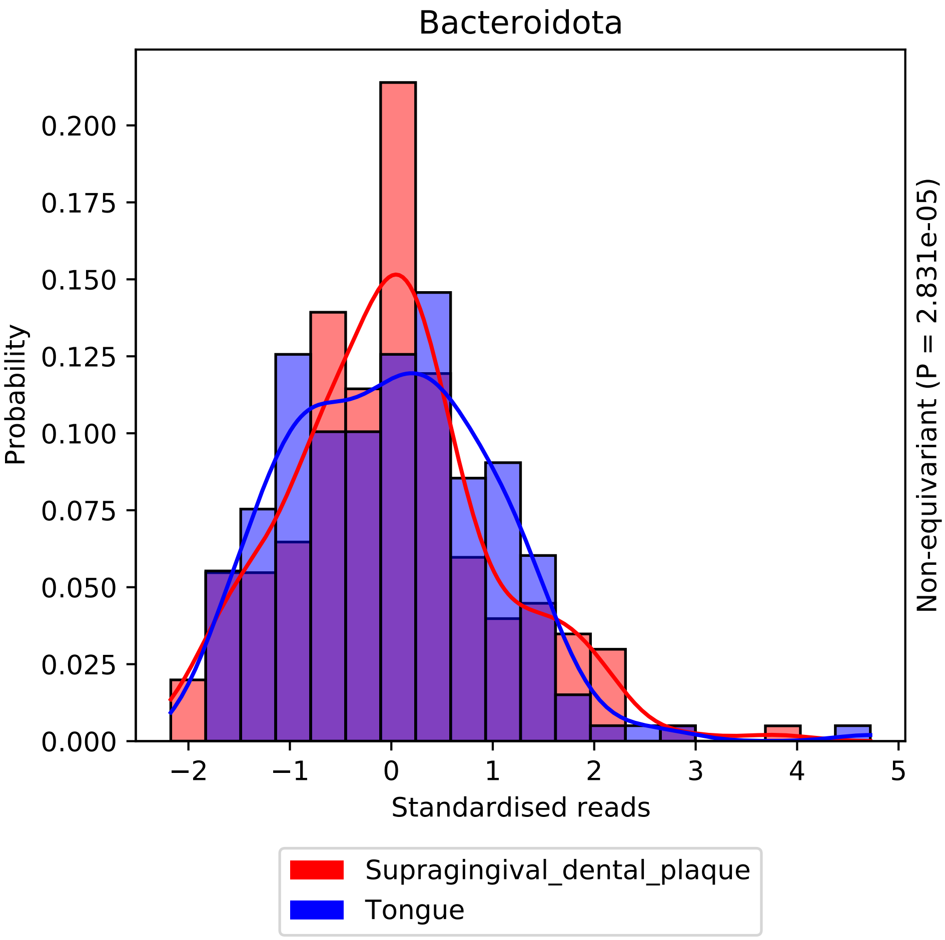

Bacteroidota
Probability
0.200
0.175
0.150
0.125
0.100
0.075
0.050
0.025
0.000
−2
−1
0
1
2
3
4
5
Standardised reads
Non-equivariant (P = 2.831e-05)
Supragingival_dental_plaque
Tongue

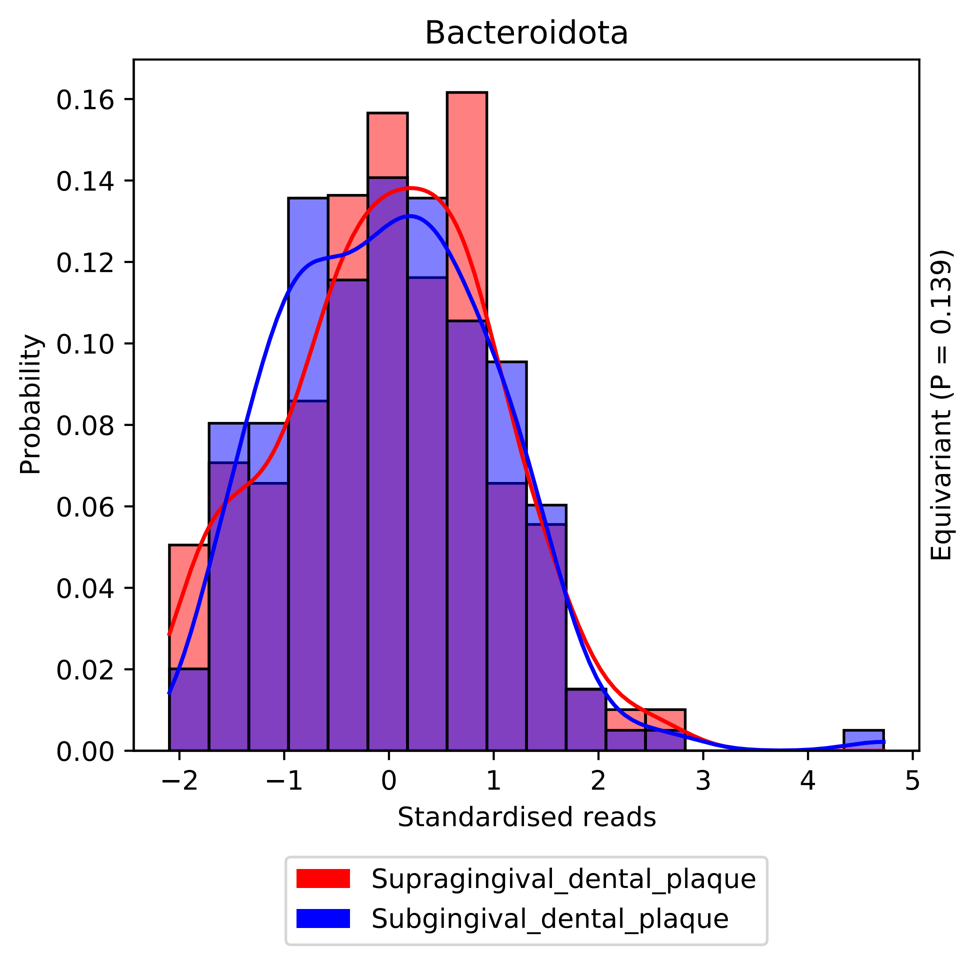

Bacteroidota
Probability
0.16
0.14
0.12
0.10
0.08
0.06
0.04
0.02
0.00
−2
−1
0
1
2
3
4
5
Standardised reads
Equivariant (P = 0.139)
Supragingival_dental_plaque
Subgingival_dental_plaque

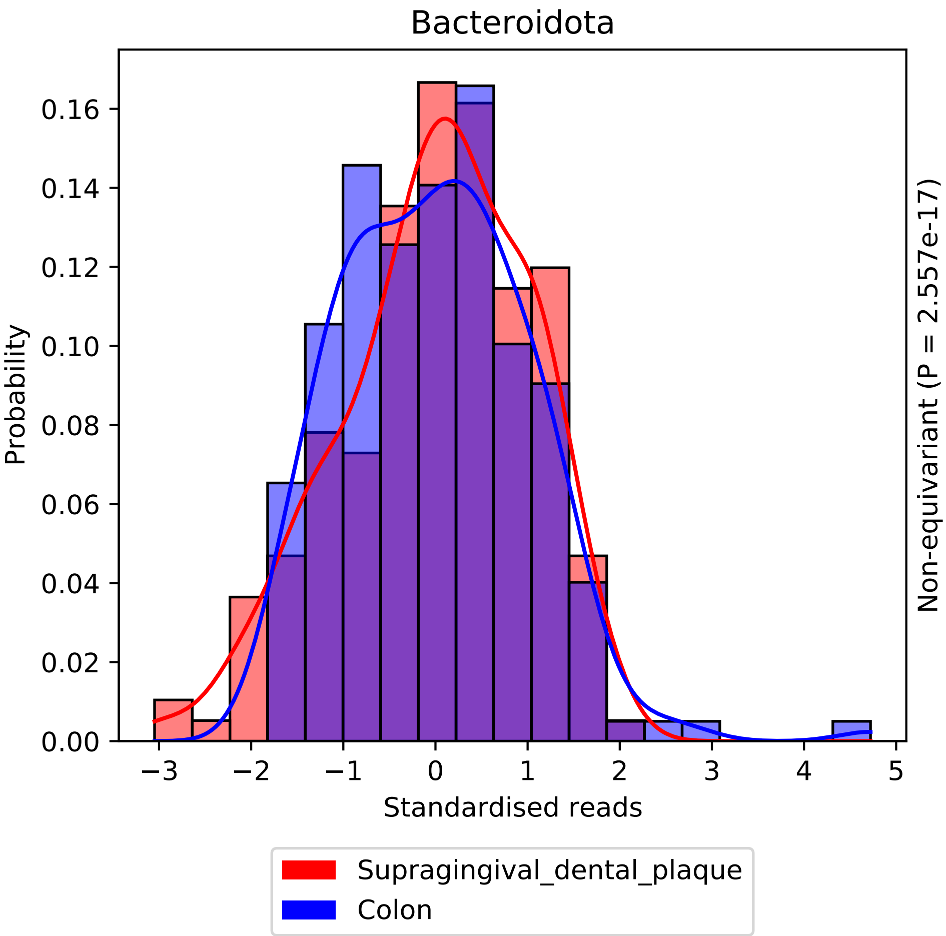

Bacteroidota
Probability
0.00
0.02
0.04
0.06
0.08
0.10
0.12
0.14
0.16
−3
−2
−1
0
1
2
3
4
5
Standardised reads
Non-equivariant (P = 2.557e-17)
Supragingival_dental_plaque
Colon

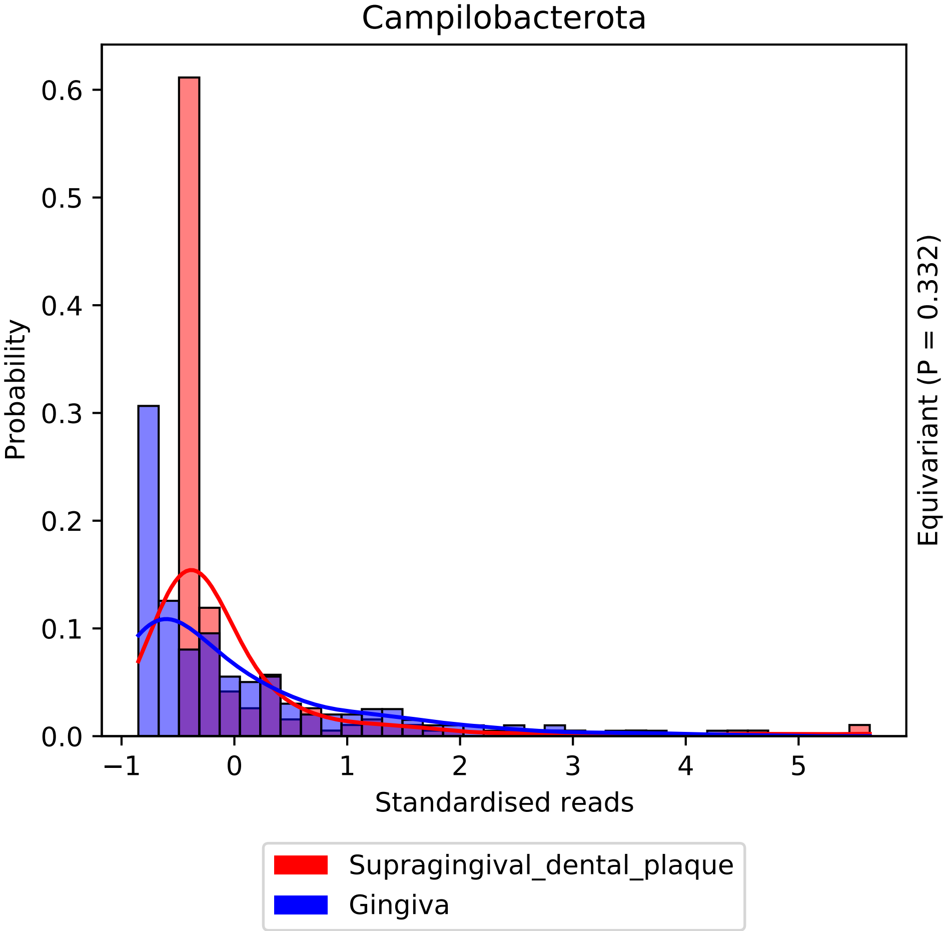

Campilobacterota
Probability
0.0
0.1
0.2
0.3
0.4
0.5
0.6
−1
0
1
2
3
4
5
Standardised reads
Equivariant (P = 0.332)
Supragingival_dental_plaque
Gingiva

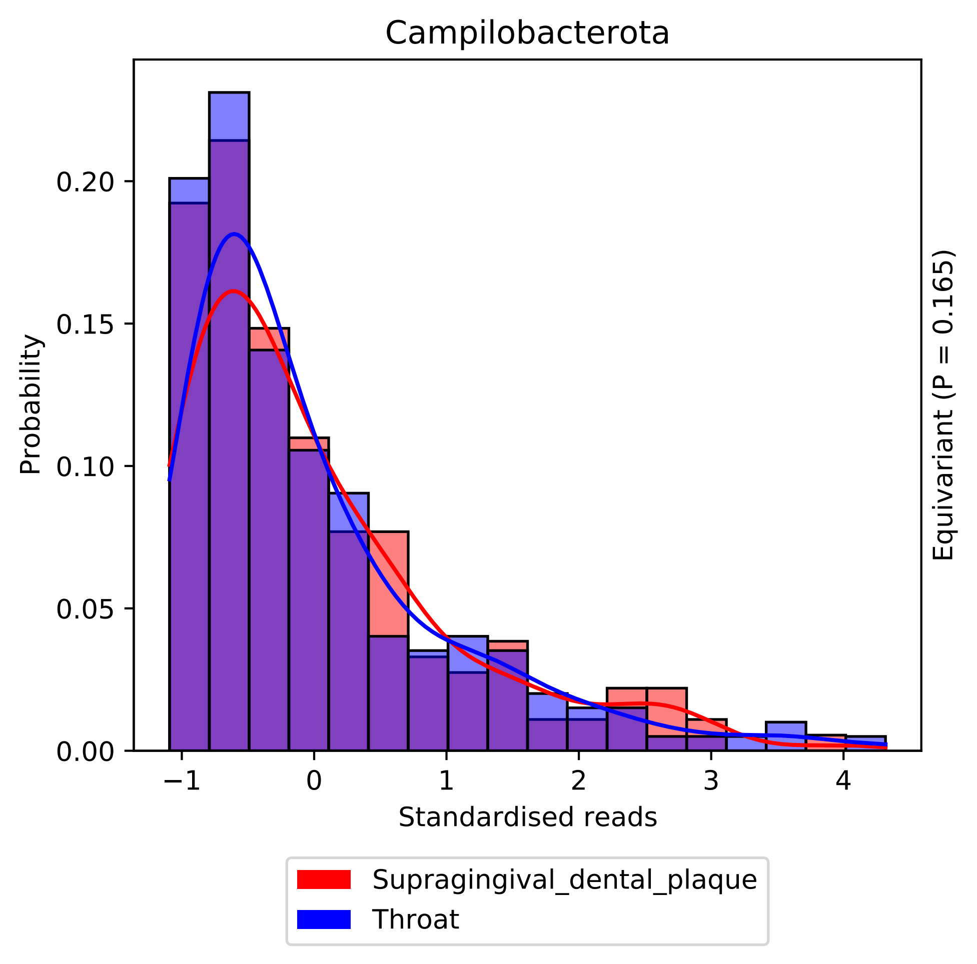
Campilobacterota
Probability
0.00
0.05
0.10
0.15
0.20
−1
0
1
2
3
4
Standardised reads
Equivariant (P = 0.165)
Supragingival_dental_plaque
Throat

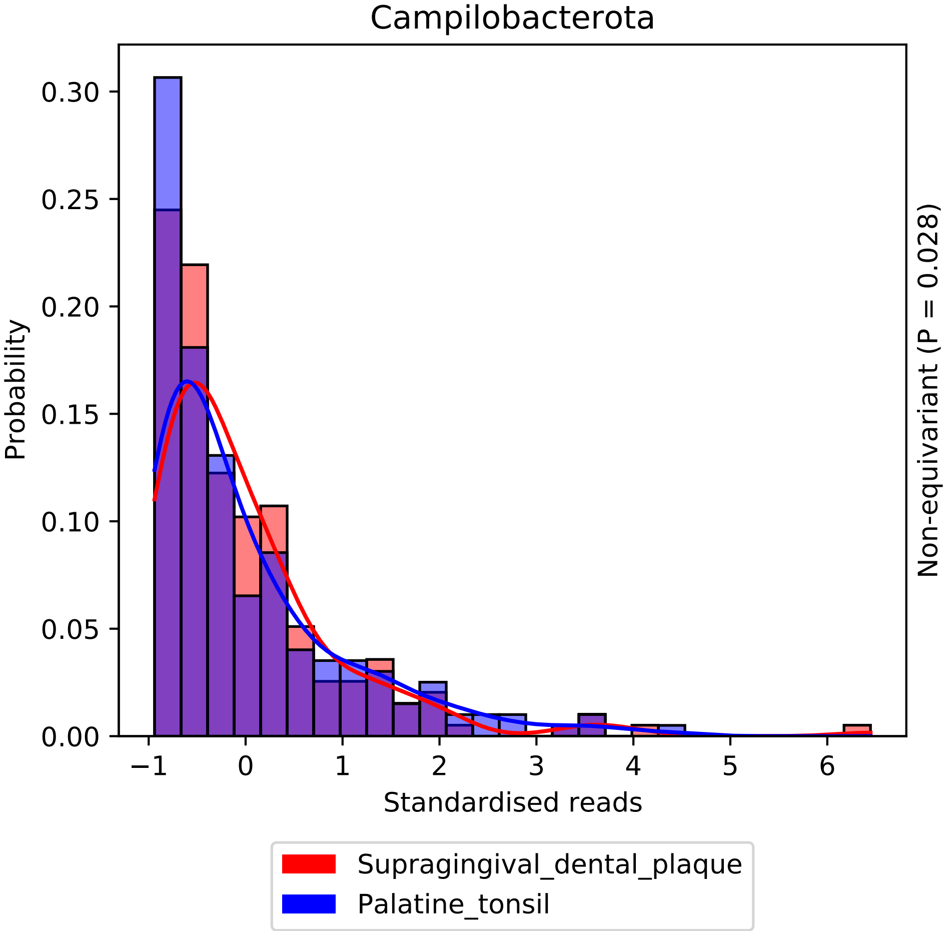

Campilobacterota
Probability
0.00
0.05
0.10
0.15
0.20
0.25
0.30
−1
0
1
2
3
4
5
6
Standardised reads
Non-equivariant (P = 0.028)
Supragingival_dental_plaque
Palatine_tonsil

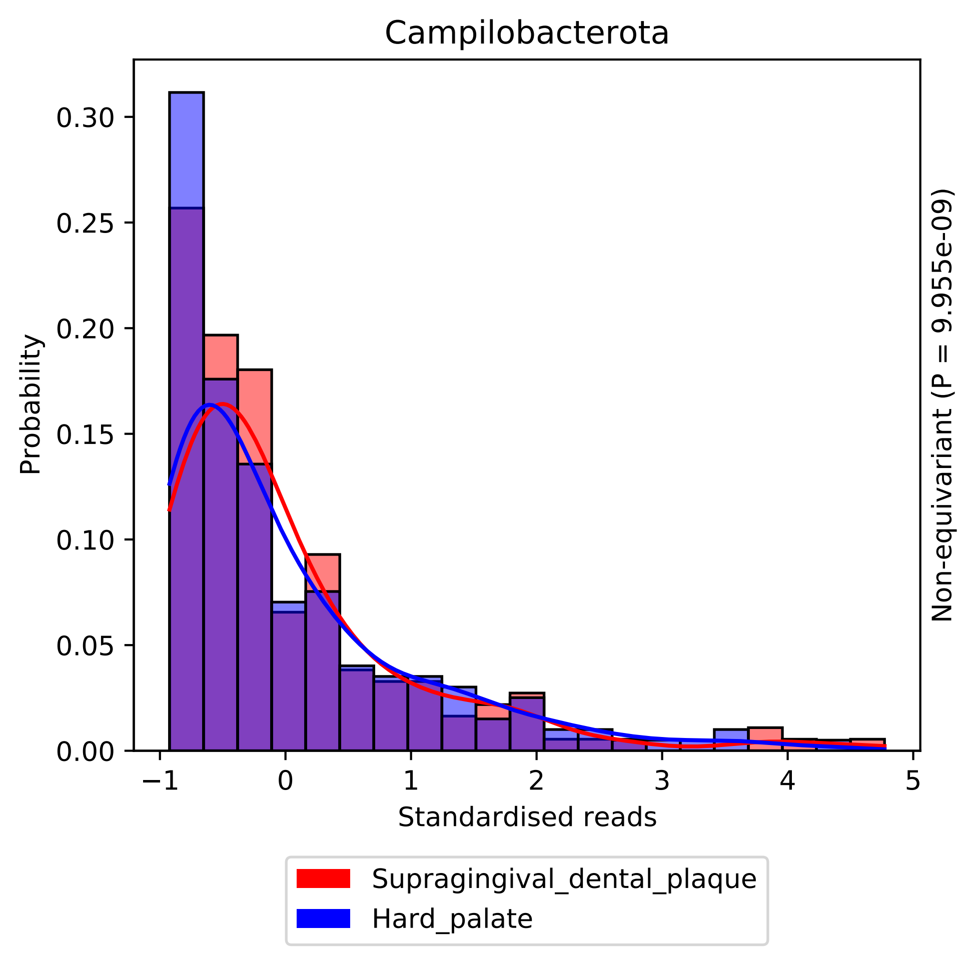
Campilobacterota
Probability
0.00
0.05
0.10
0.15
0.20
0.25
0.30
−1
0
1
2
3
4
5
Standardised reads
Non-equivariant (P = 9.955e-09)
Supragingival_dental_plaque
Hard_palate

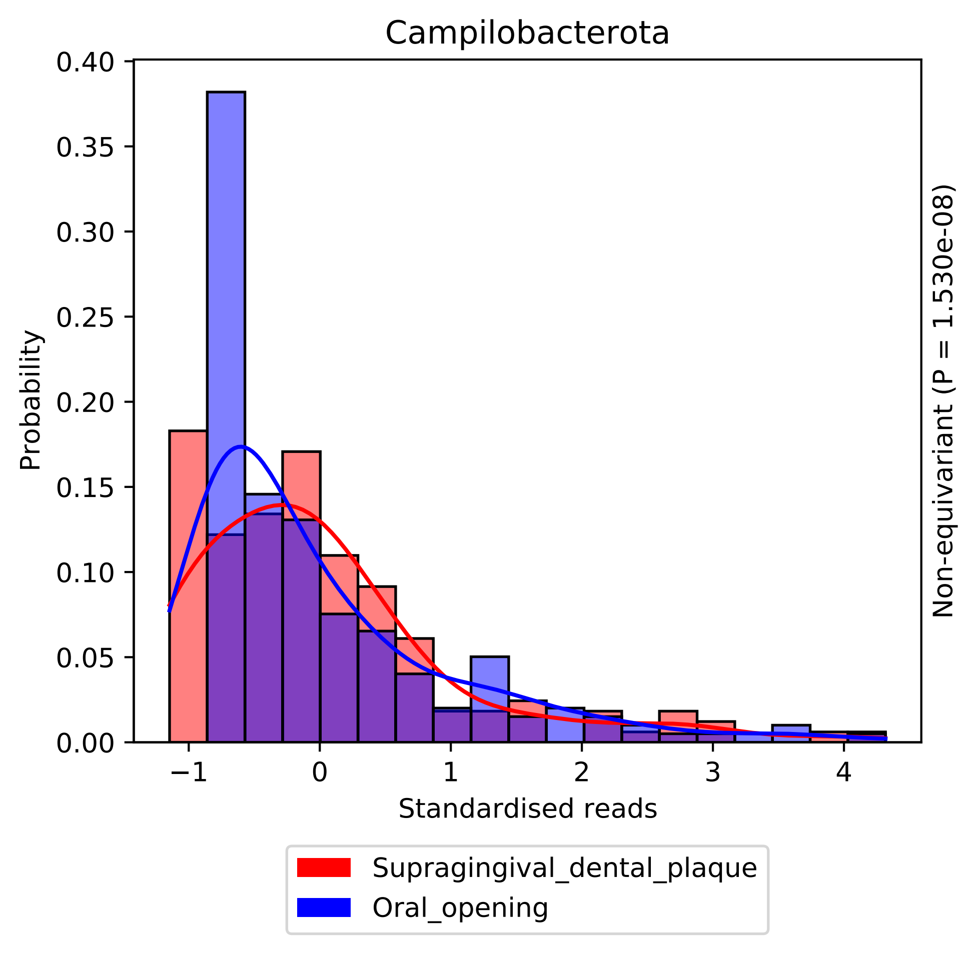

Campilobacterota
Probability
0.00
0.05
0.10
0.15
0.20
0.25
0.30
0.35
0.40
−1
0
1
2
3
4
Standardised reads
Non-equivariant (P = 1.530e-08)
Supragingival_dental_plaque
Oral_opening

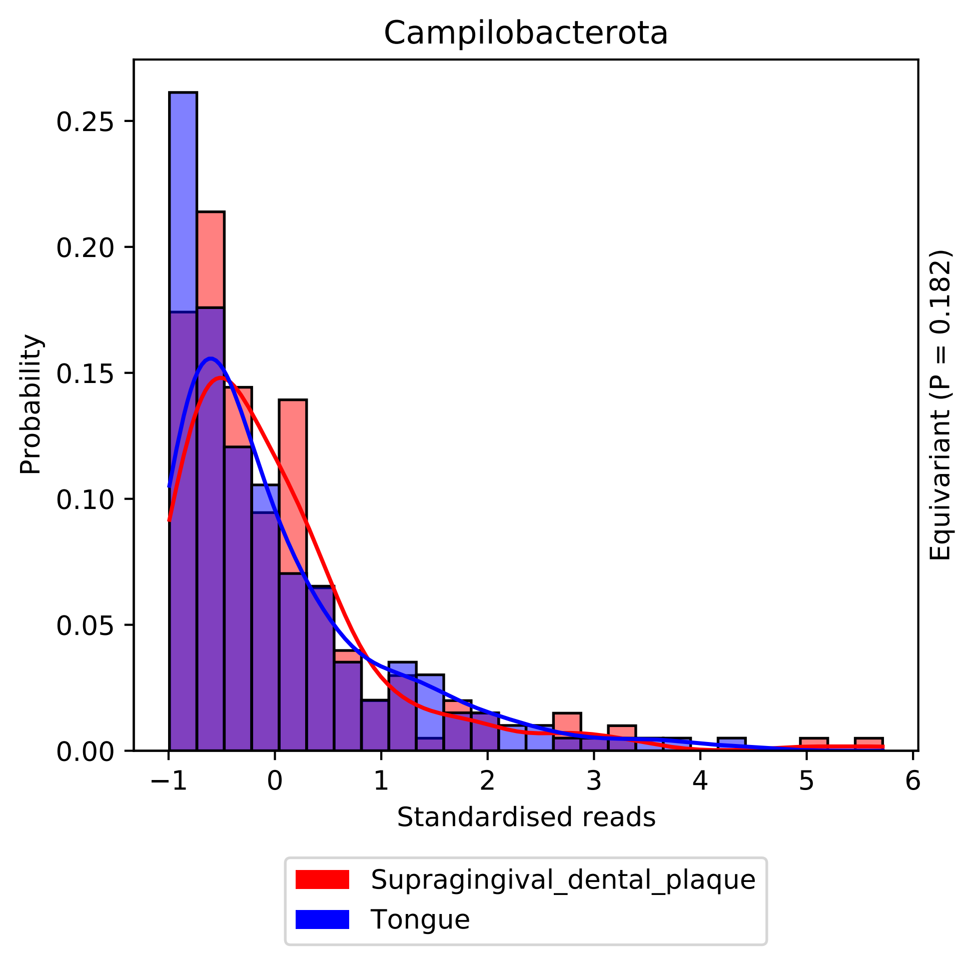
Campilobacterota
Probability
Standardised reads
Equivariant (P = 0.182)
0.00
0.05
0.10
0.15
0.20
0.25
−1
0
1
2
3
4
5
6
Supragingival_dental_plaque
Tongue

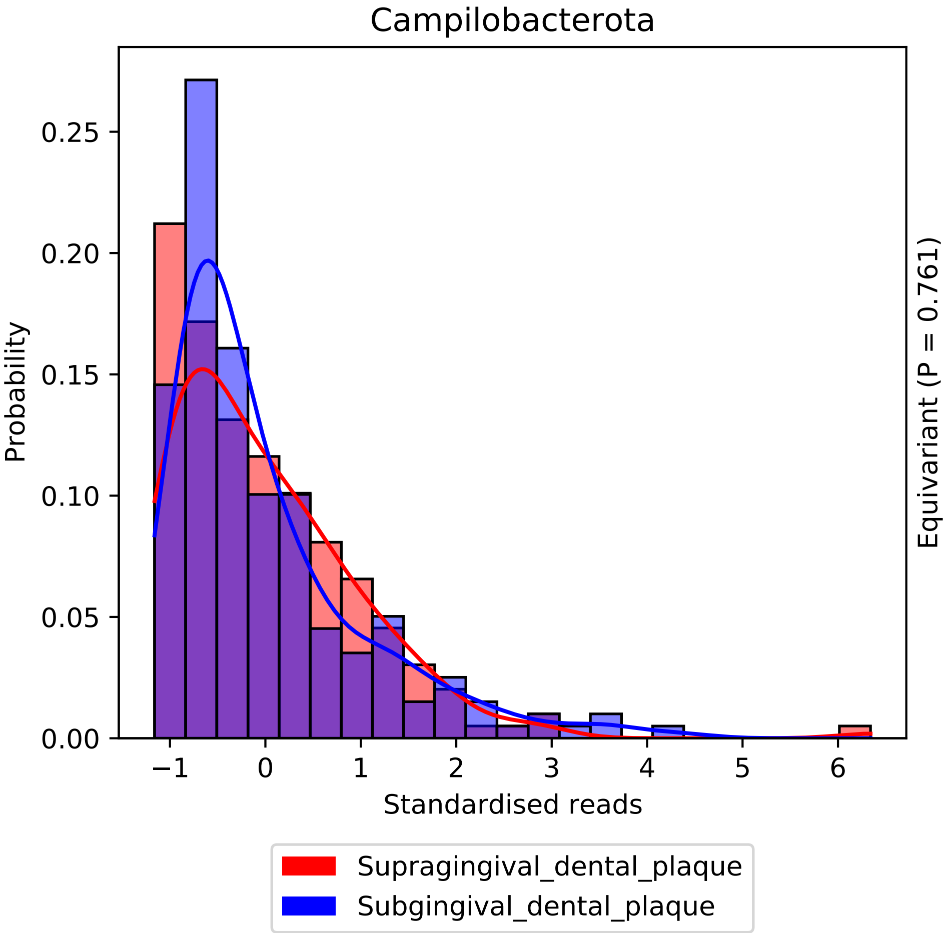

Campilobacterota
Probability
0.00
0.05
0.10
0.15
0.20
0.25
−1
0
1
2
3
4
5
6
Standardised reads
Equivariant (P = 0.761)
Supragingival_dental_plaque
Subgingival_dental_plaque

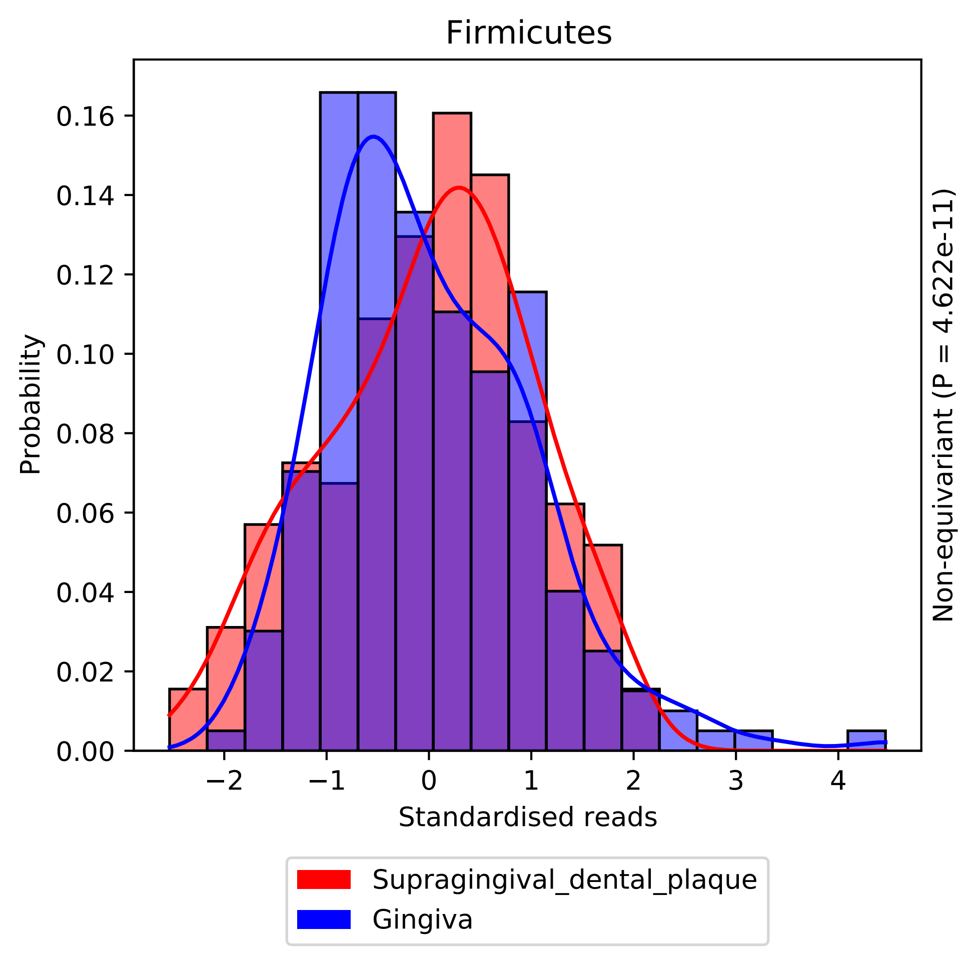

Firmicutes
Probability
0.00
0.02
0.04
0.06
0.08
0.10
0.12
0.14
0.16
−2
−1
0
1
2
3
4
Standardised reads
Non-equivariant (P = 4.622e-11)
Supragingival_dental_plaque
Gingiva

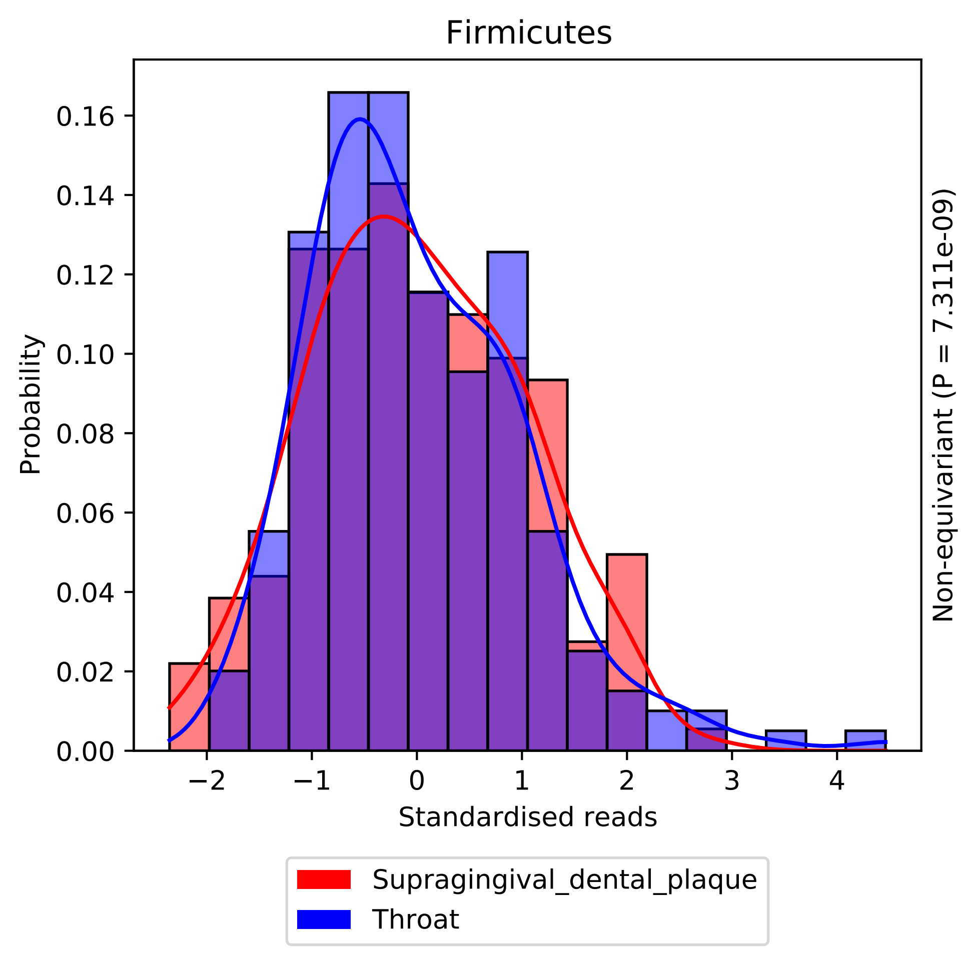

Firmicutes
Probability
0.00
0.02
0.04
0.06
0.08
0.10
0.12
0.14
0.16
−2
−1
0
1
2
3
4
Standardised reads
Non-equivariant (P = 7.311e-09)
Supragingival_dental_plaque
Throat

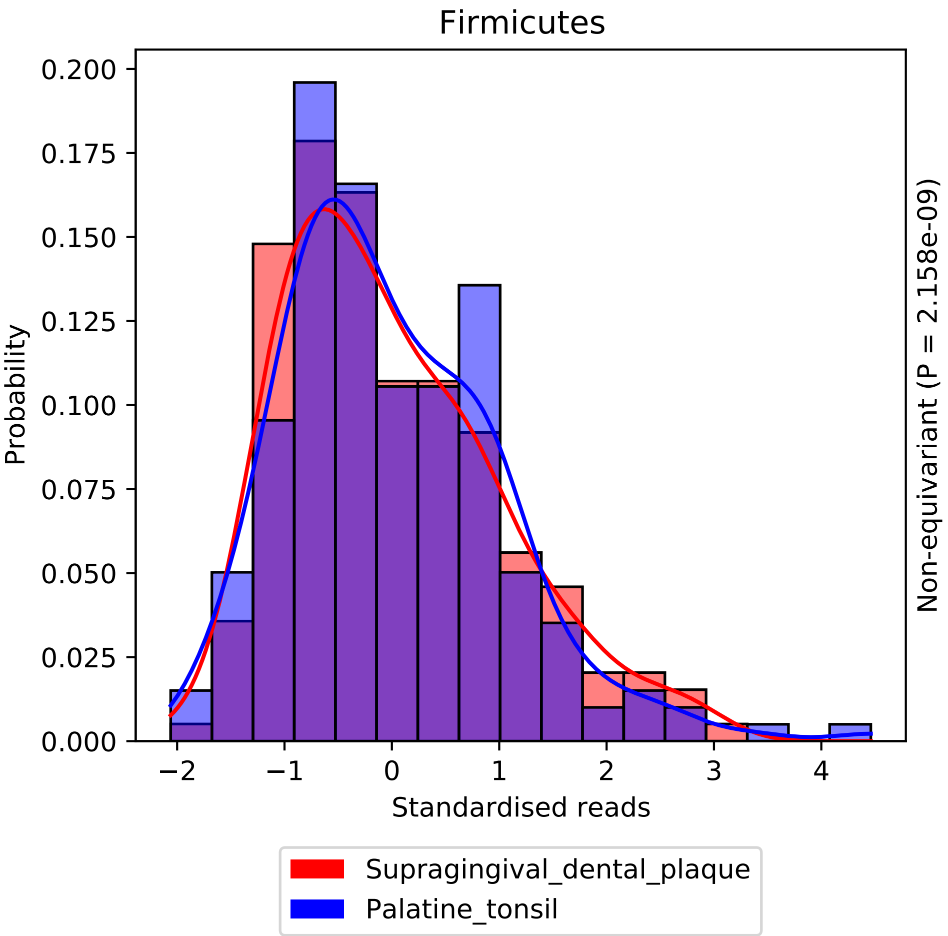

Firmicutes
Probability
Standardised reads
Non-equivariant (P = 2.158e-09)
Supragingival_dental_plaque
Palatine_tonsil
0.000
0.025
0.050
0.075
0.100
0.125
0.150
0.175
0.200
−2
−1
0
1
2
3
4

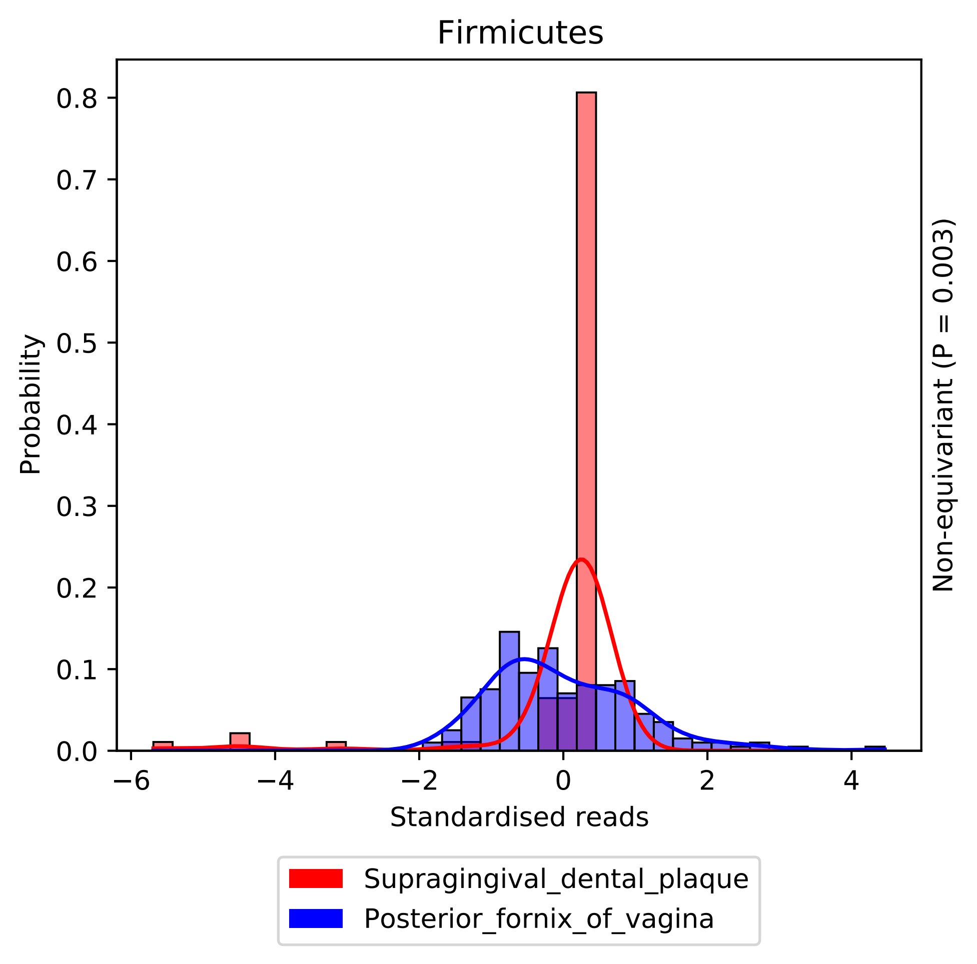

Firmicutes
Probability
Standardised reads
Non-equivariant (P = 0.003)
0.0
0.1
0.2
0.3
0.4
0.5
0.6
0.7
0.8
−6
−4
−2
0
2
4
Supragingival_dental_plaque
Posterior_fornix_of_vagina

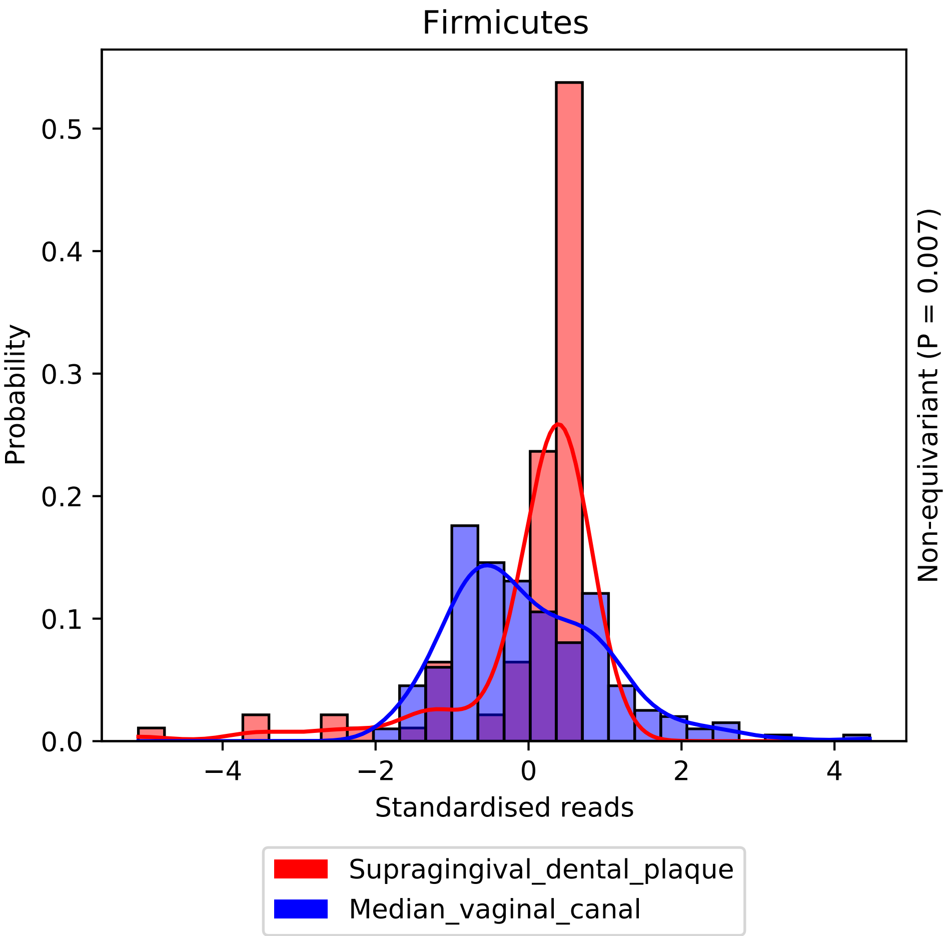
Firmicutes
Probability
Standardised reads
Non-equivariant (P = 0.007)
Supragingival_dental_plaque
Median_vaginal_canal
0.0
0.1
0.2
0.3
0.4
0.5
−4
−2
0
2
4

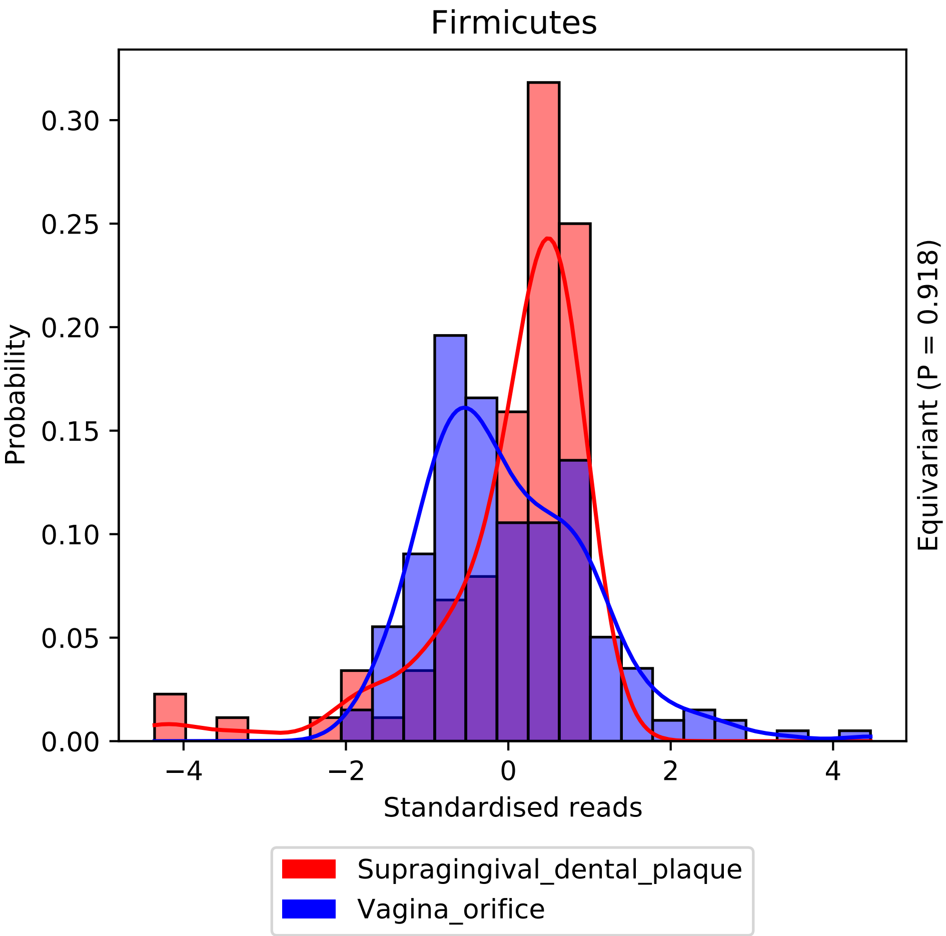
Firmicutes
Probability
Standardised reads
Equivariant (P = 0.918)
0.00
0.05
0.10
0.15
0.20
0.25
0.30
−4
−2
0
2
4
Supragingival_dental_plaque
Vagina_orifice

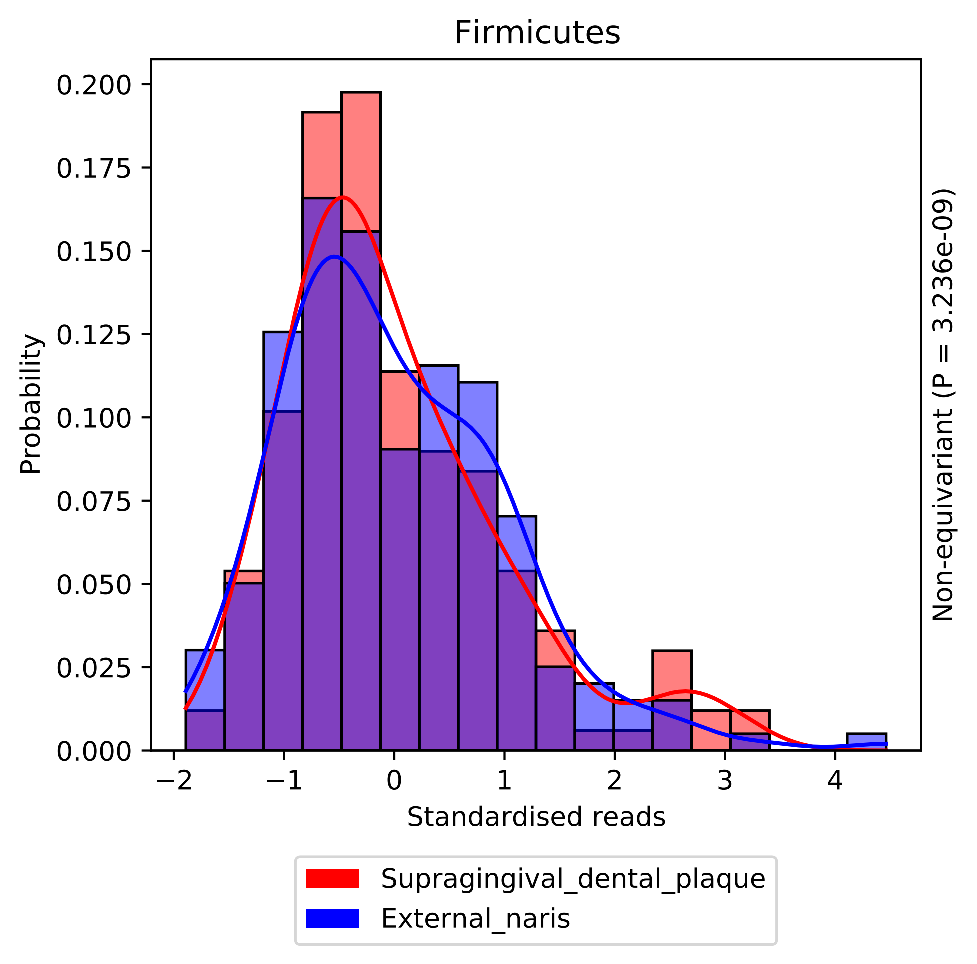

Firmicutes
Probability
0.200
0.175
0.150
0.125
0.100
0.075
0.050
0.025
0.000
−2
−1
0
1
2
3
4
Standardised reads
Non-equivariant (P = 3.236e-09)
Supragingival_dental_plaque
External_naris

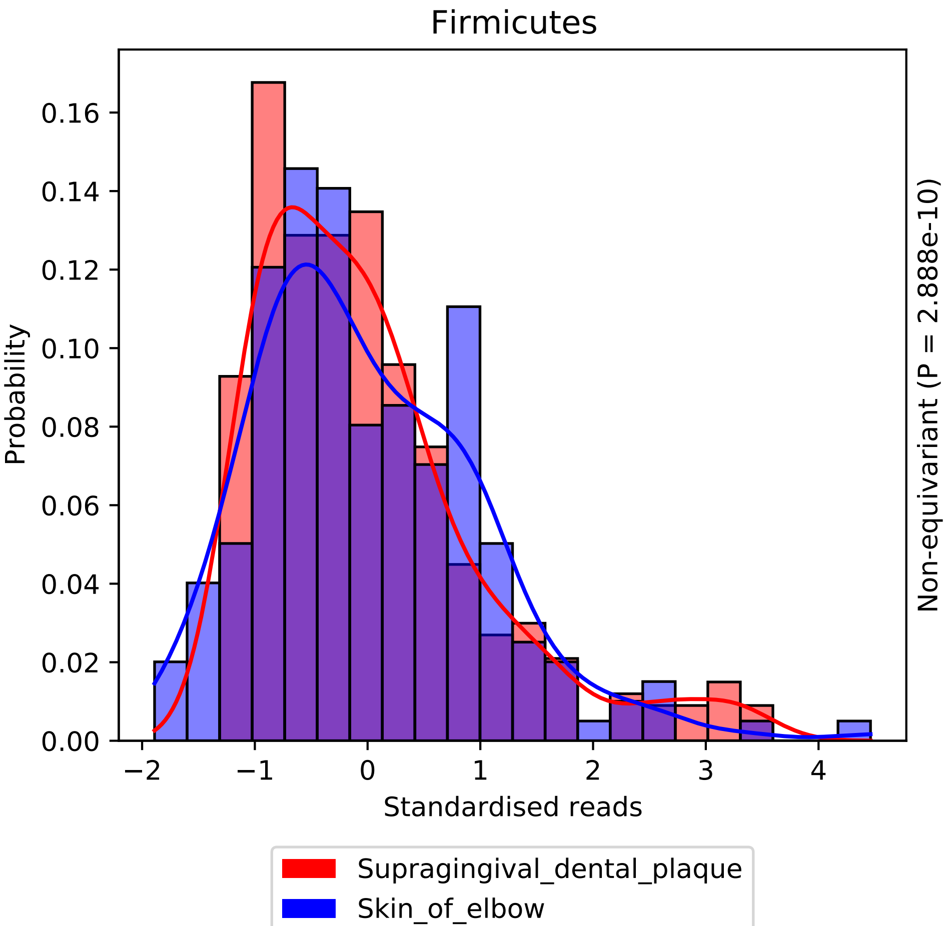

Firmicutes
Probability
0.00
0.02
0.04
0.06
0.08
0.10
0.12
0.14
0.16
−2
−1
0
1
2
3
4
Standardised reads
Non-equivariant (P = 2.888e-10)
Supragingival_dental_plaque
Skin_of_elbow

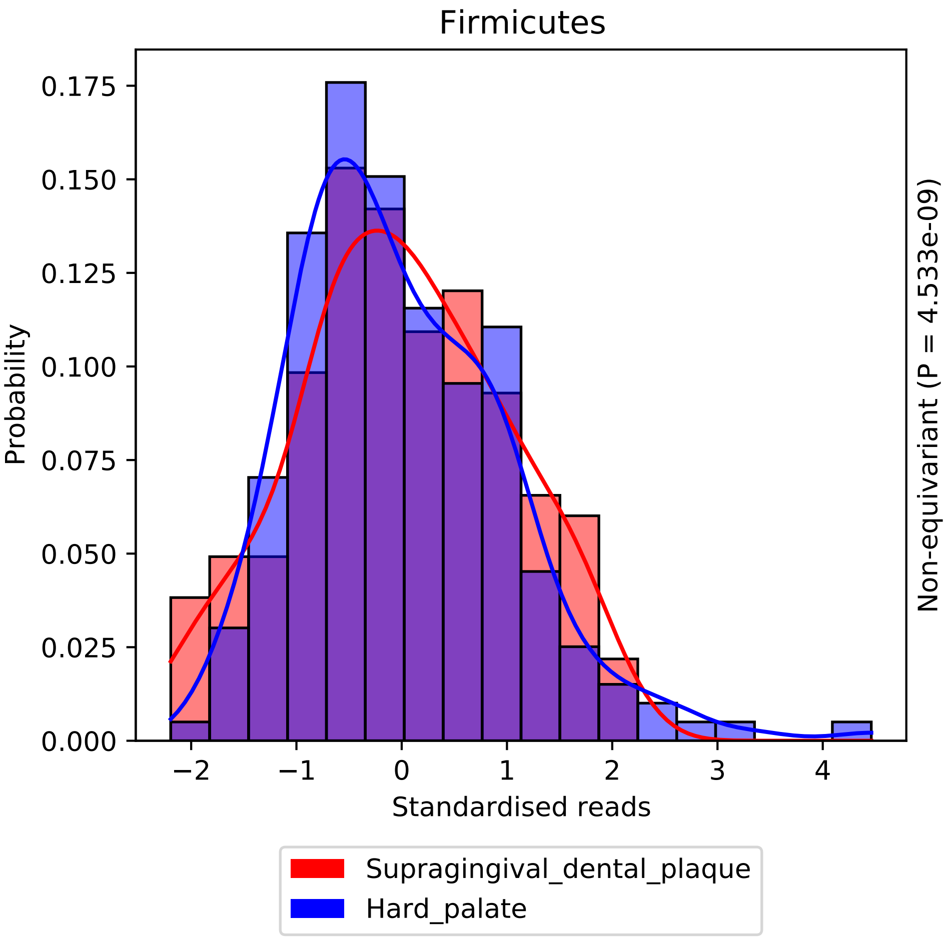

Firmicutes
0.175
0.150
0.125
0.100
0.075
0.050
0.025
0.000
Probability
−2
−1
0
1
2
3
4
Standardised reads
Non-equivariant (P = 4.533e-09)
Supragingival_dental_plaque
Hard_palate

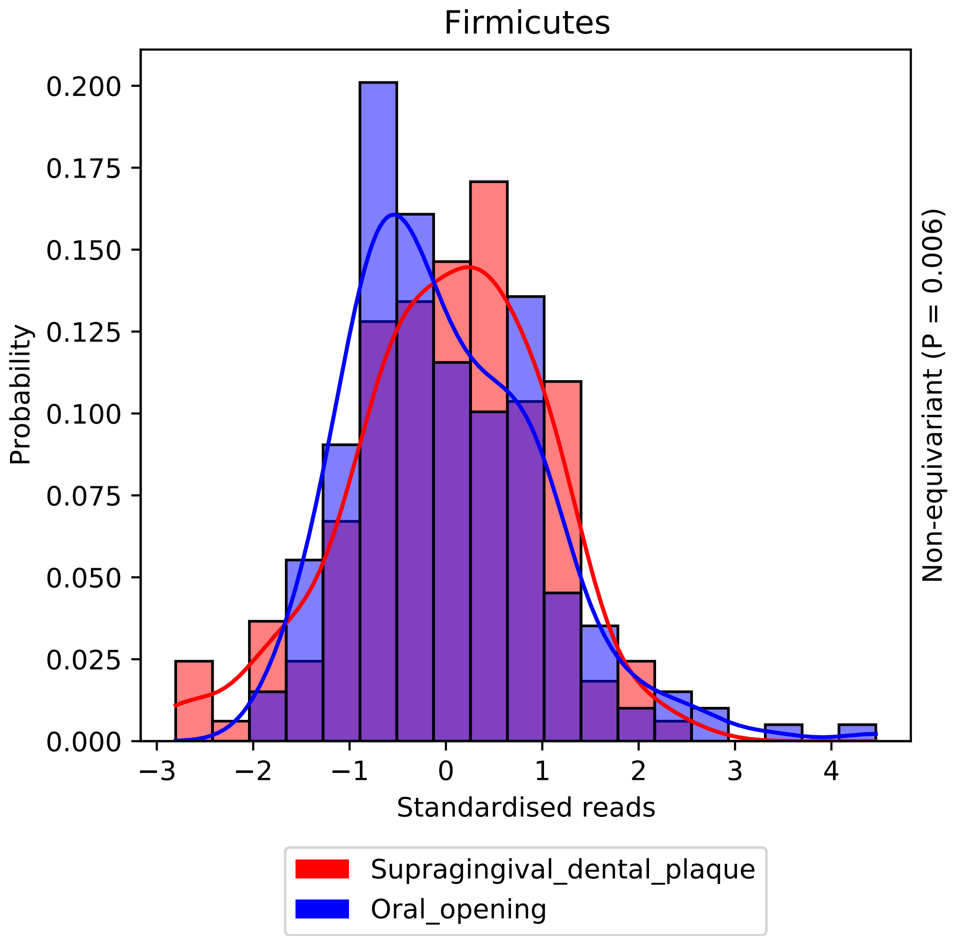

Firmicutes
Probability
0.200
0.175
0.150
0.125
0.100
0.075
0.050
0.025
0.000
−3
−2
−1
0
1
2
3
4
Standardised reads
Non-equivariant (P = 0.006)
Supragingival_dental_plaque
Oral_opening

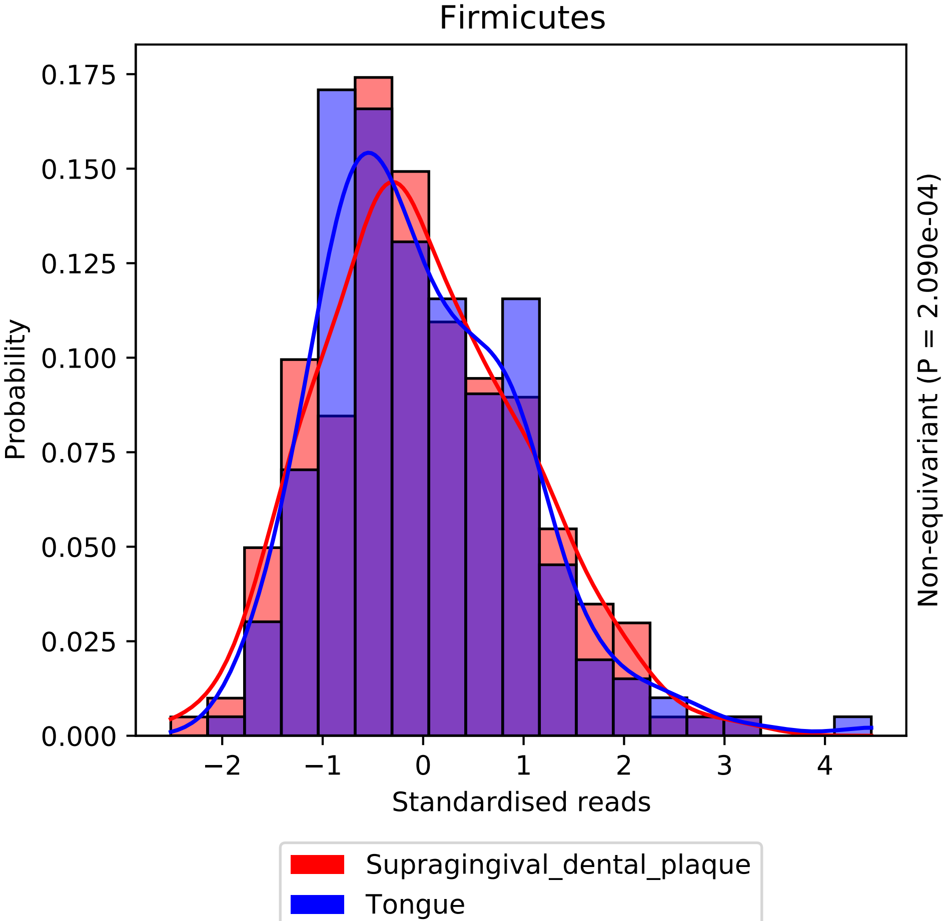
Firmicutes
Probability
0.000
0.025
0.050
0.075
0.100
0.125
0.150
0.175
−2
−1
0
1
2
3
4
Standardised reads
Non-equivariant (P = 2.090e-04)
Supragingival_dental_plaque
Tongue

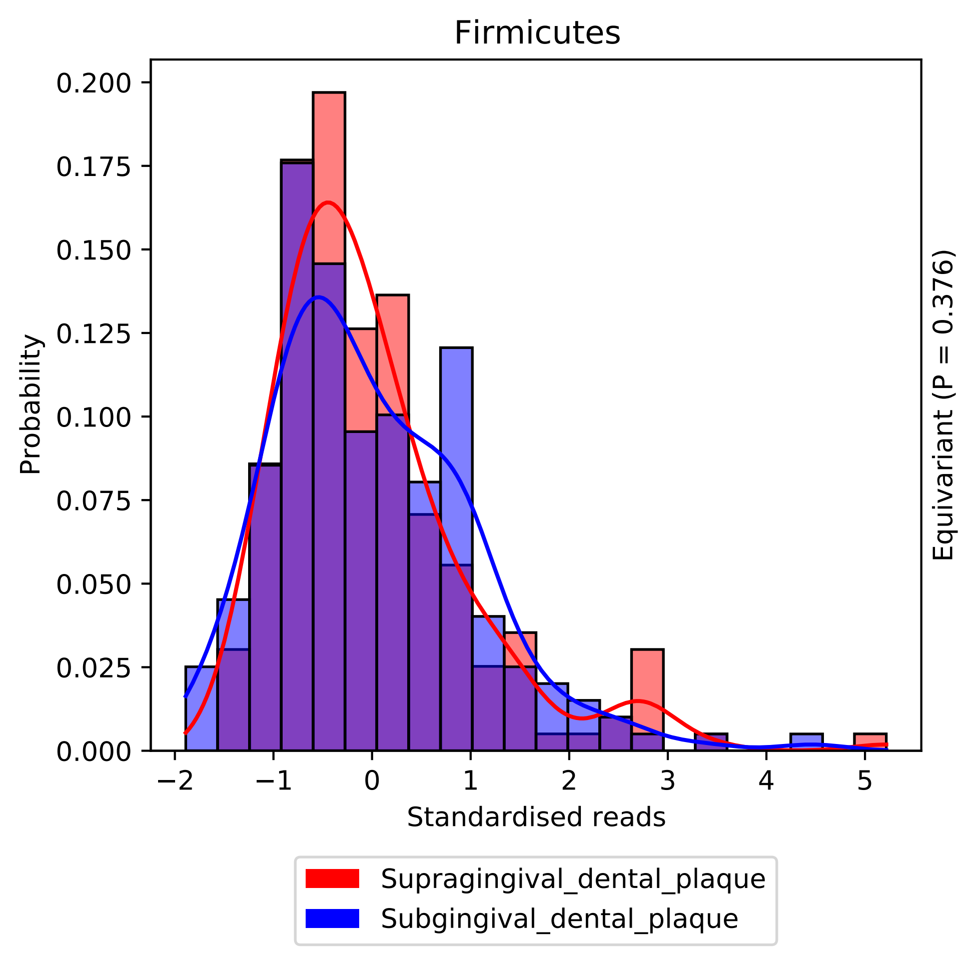
Firmicutes
Probability
Standardised reads
0.000
0.025
0.050
0.075
0.100
0.125
0.150
0.175
0.200
−2
−1
0
1
2
3
4
5
Equivariant (P = 0.376)
Supragingival_dental_plaque
Subgingival_dental_plaque

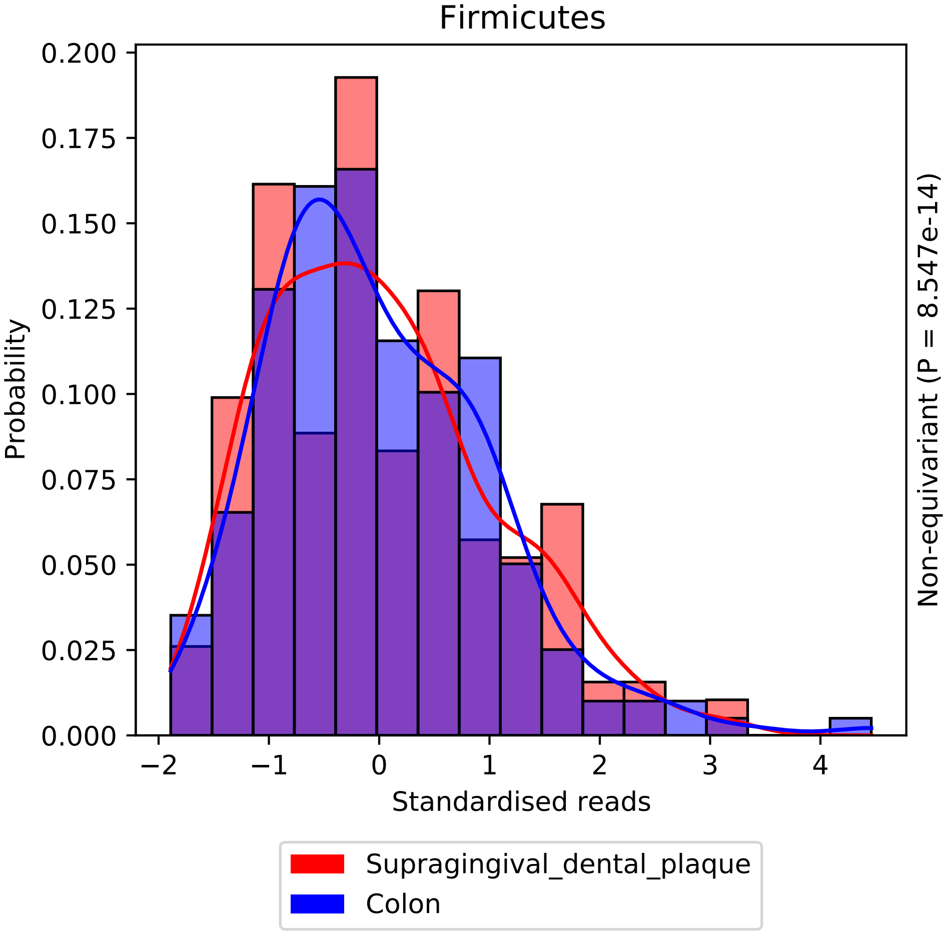

Firmicutes
Probability
0.200
0.175
0.150
0.125
0.100
0.075
0.050
0.025
0.000
−2
−1
0
1
2
3
4
Standardised reads
Non-equivariant (P = 8.547e-14)
Supragingival_dental_plaque
Colon

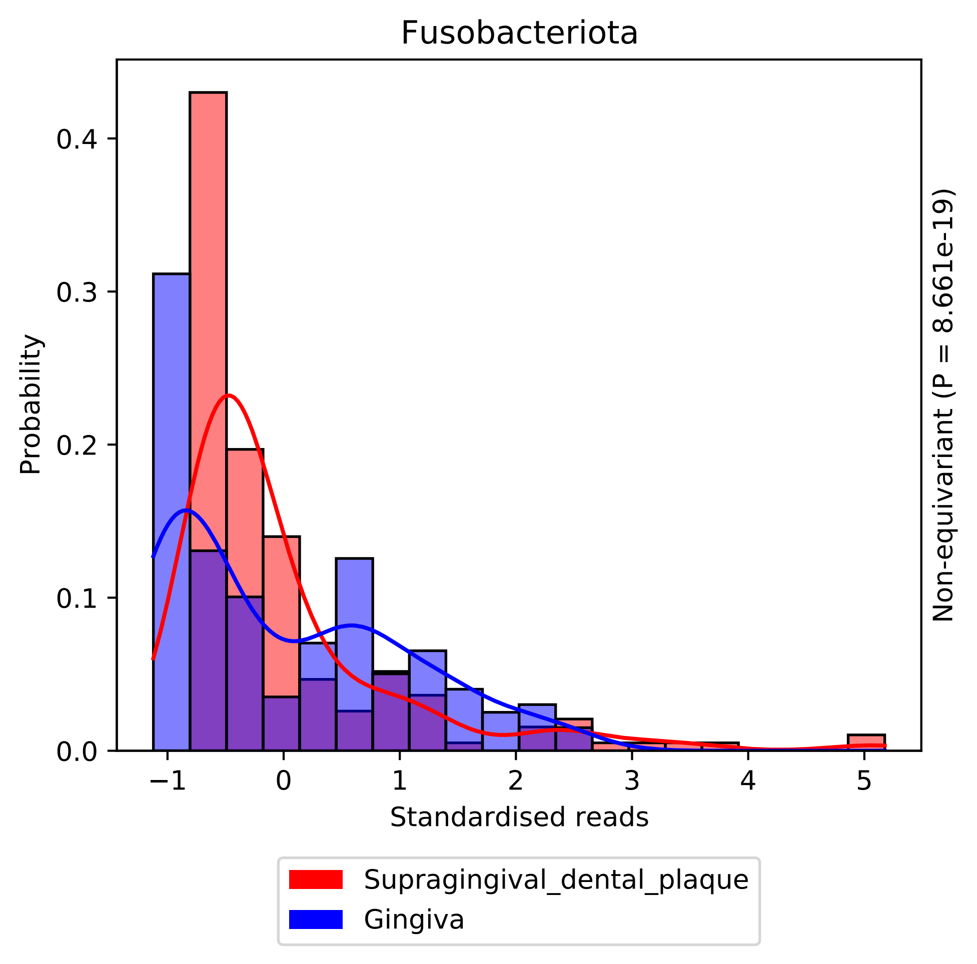

Fusobacteriota
Probability
0.0
0.1
0.2
0.3
0.4
−1
0
1
2
3
4
5
Standardised reads
Non-equivariant (P = 8.661e-19)
Supragingival_dental_plaque
Gingiva

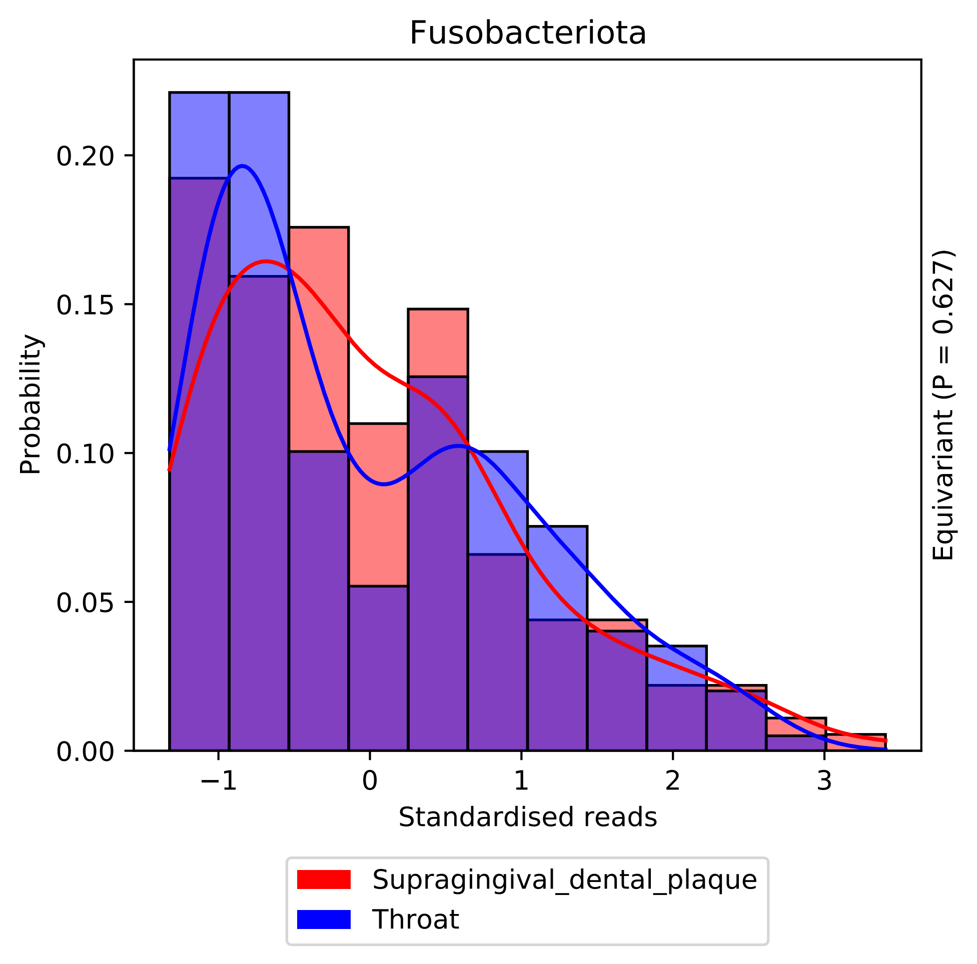

Fusobacteriota
Probability
0.00
0.05
0.10
0.15
0.20
−1
0
1
2
3
Standardised reads
Equivariant (P = 0.627)
Supragingival_dental_plaque
Throat

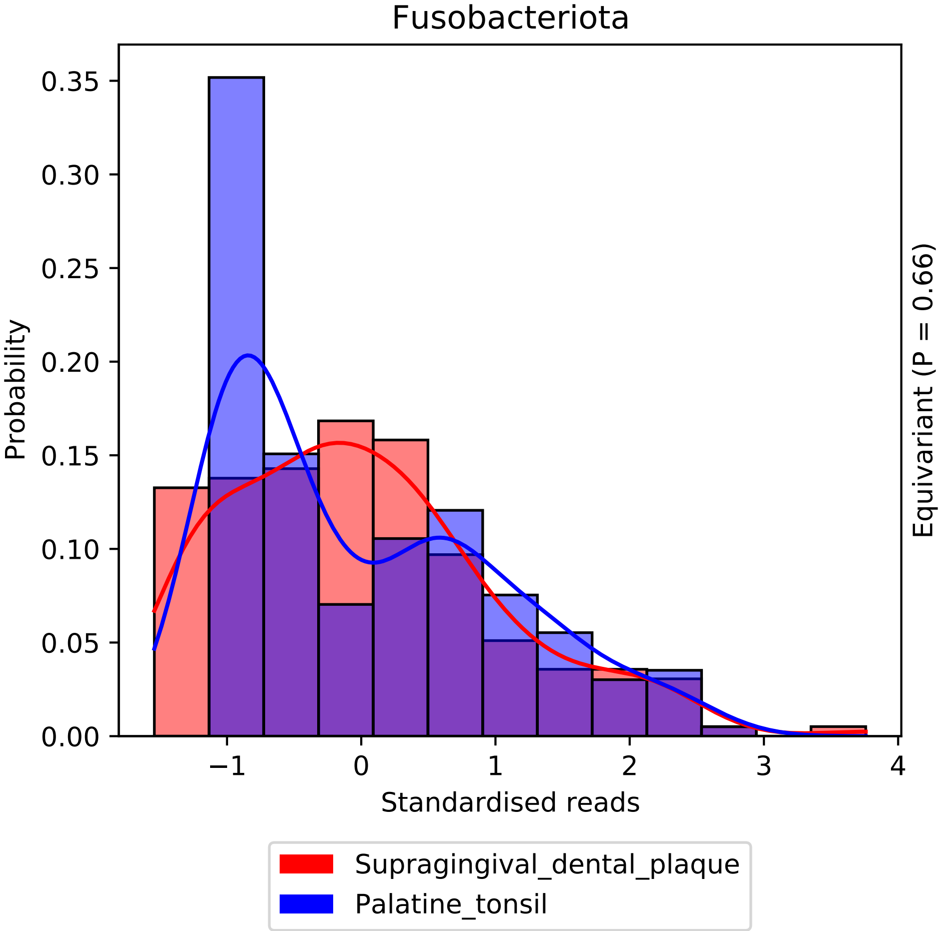
Fusobacteriota
Probability
Standardised reads
Equivariant (P = 0.66)
0.00
0.05
0.10
0.15
0.20
0.25
0.30
0.35
−1
0
1
2
3
4
Supragingival_dental_plaque
Palatine_tonsil

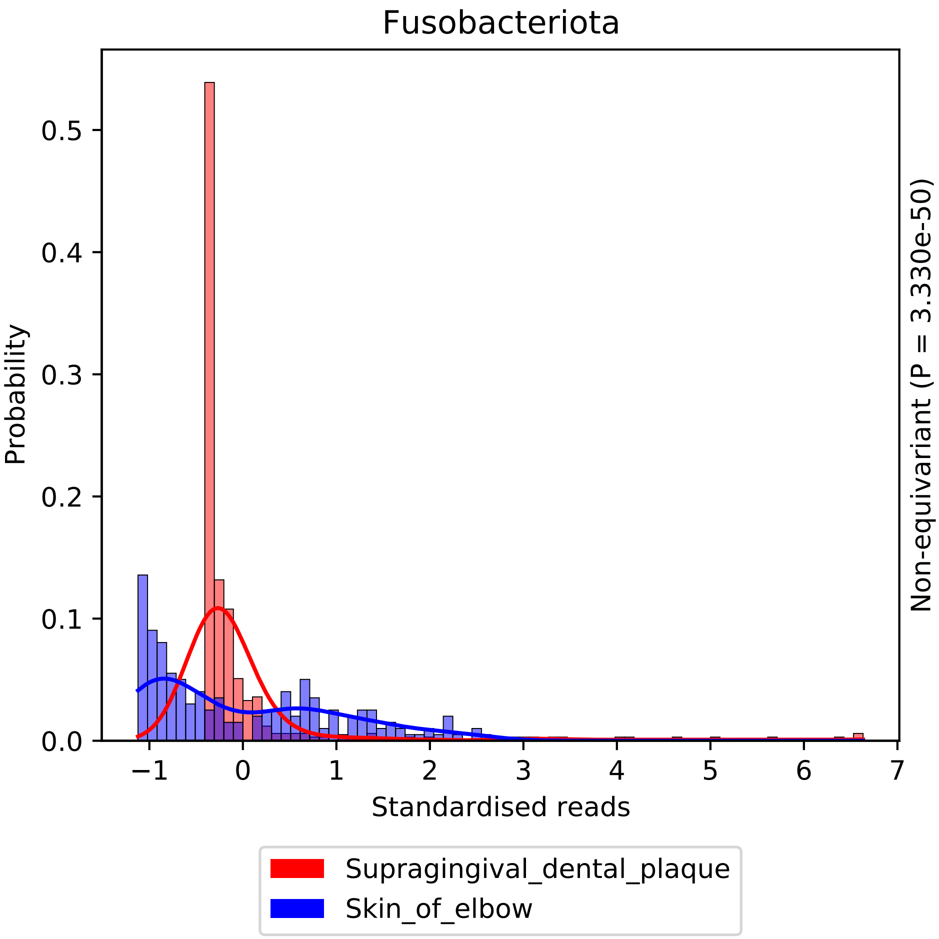

Fusobacteriota
Probability
Standardised reads
Non-equivariant (P = 3.330e-50)
0.0
0.1
0.2
0.3
0.4
0.5
−1
0
1
2
3
4
5
6
7
Supragingival_dental_plaque
Skin_of_elbow

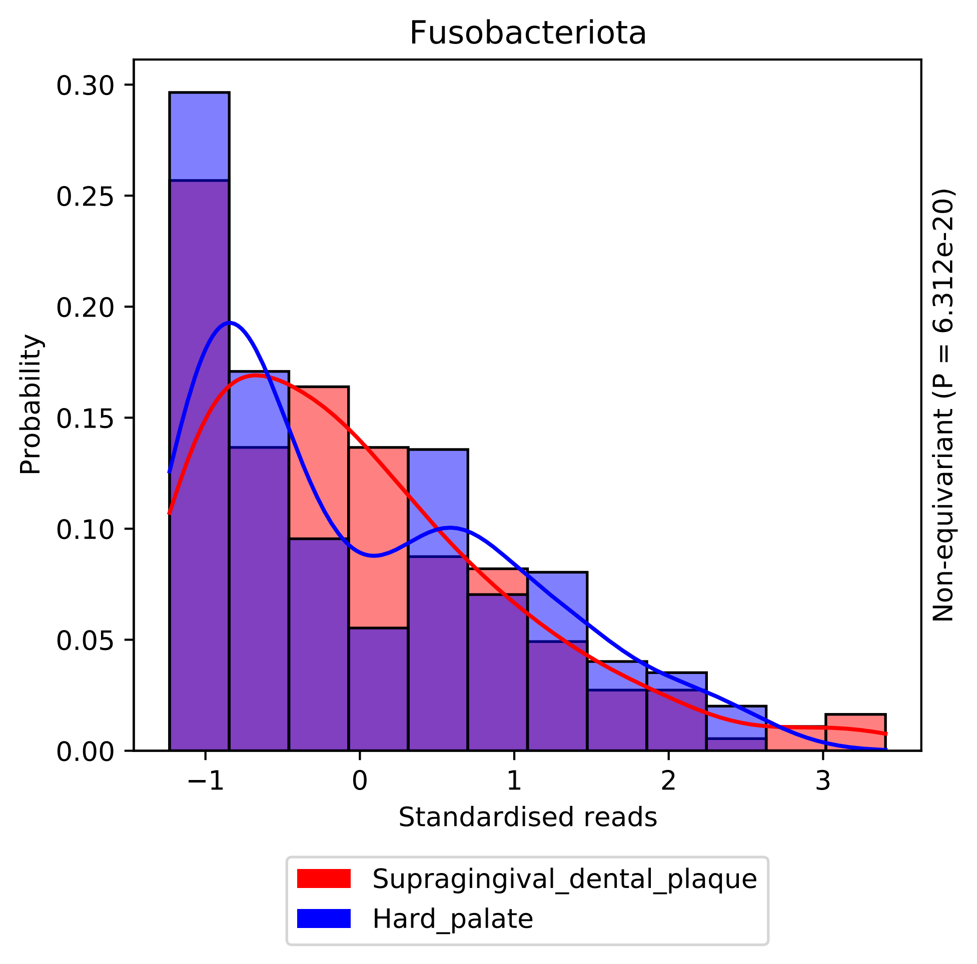

Fusobacteriota
Probability
0.30
0.25
0.20
0.15
0.10
0.05
0.00
−1
0
1
2
3
Standardised reads
Non-equivariant (P = 6.312e-20)
Supragingival_dental_plaque
Hard_palate

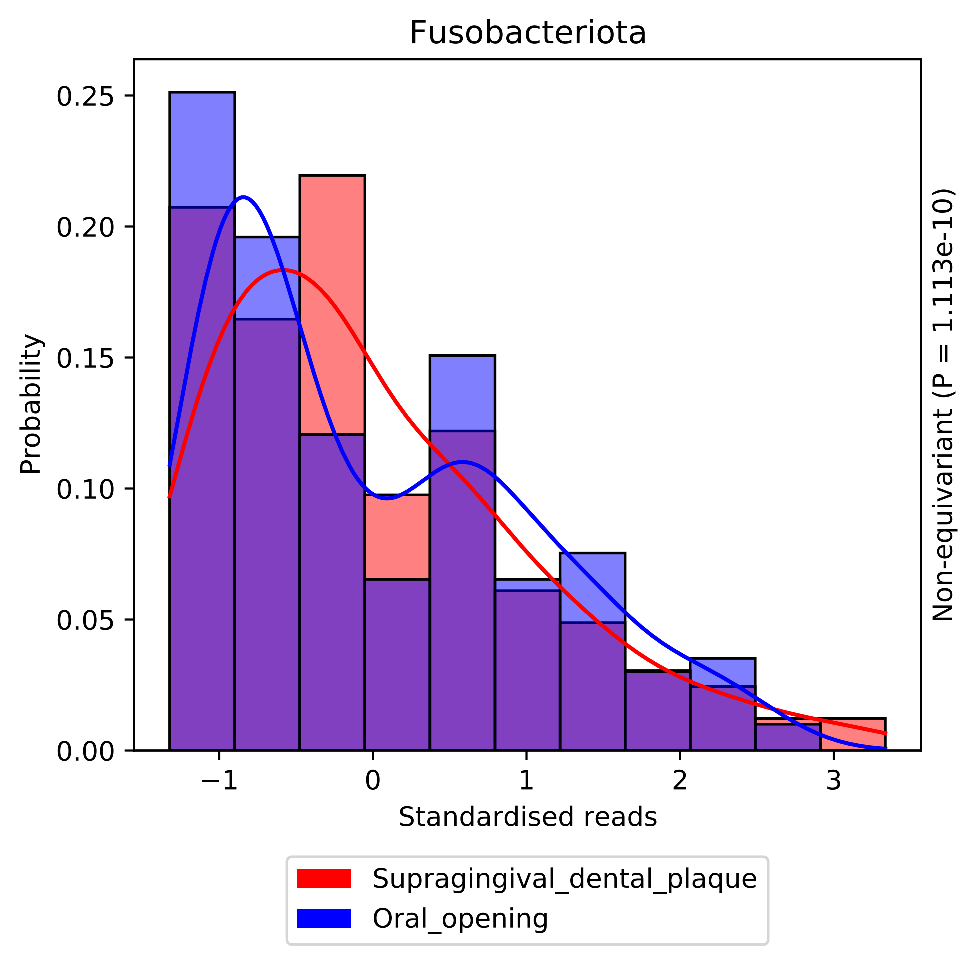
Fusobacteriota
Probability
Standardised reads
Non-equivariant (P = 1.113e-10)
0.00
0.05
0.10
0.15
0.20
0.25
−1
0
1
2
3
Supragingival_dental_plaque
Oral_opening

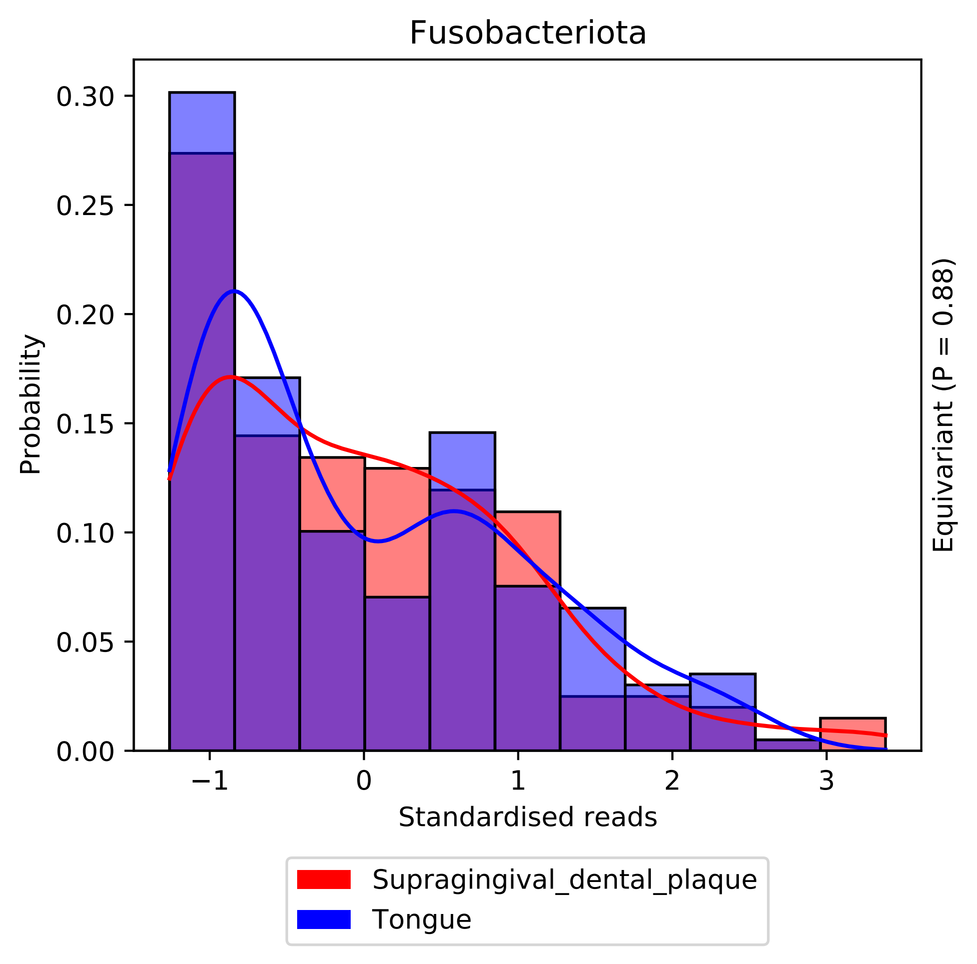
Fusobacteriota
Probability
0.30
0.25
0.20
0.15
0.10
0.05
0.00
−1
0
1
2
3
Standardised reads
Equivariant (P = 0.88)
Supragingival_dental_plaque
Tongue

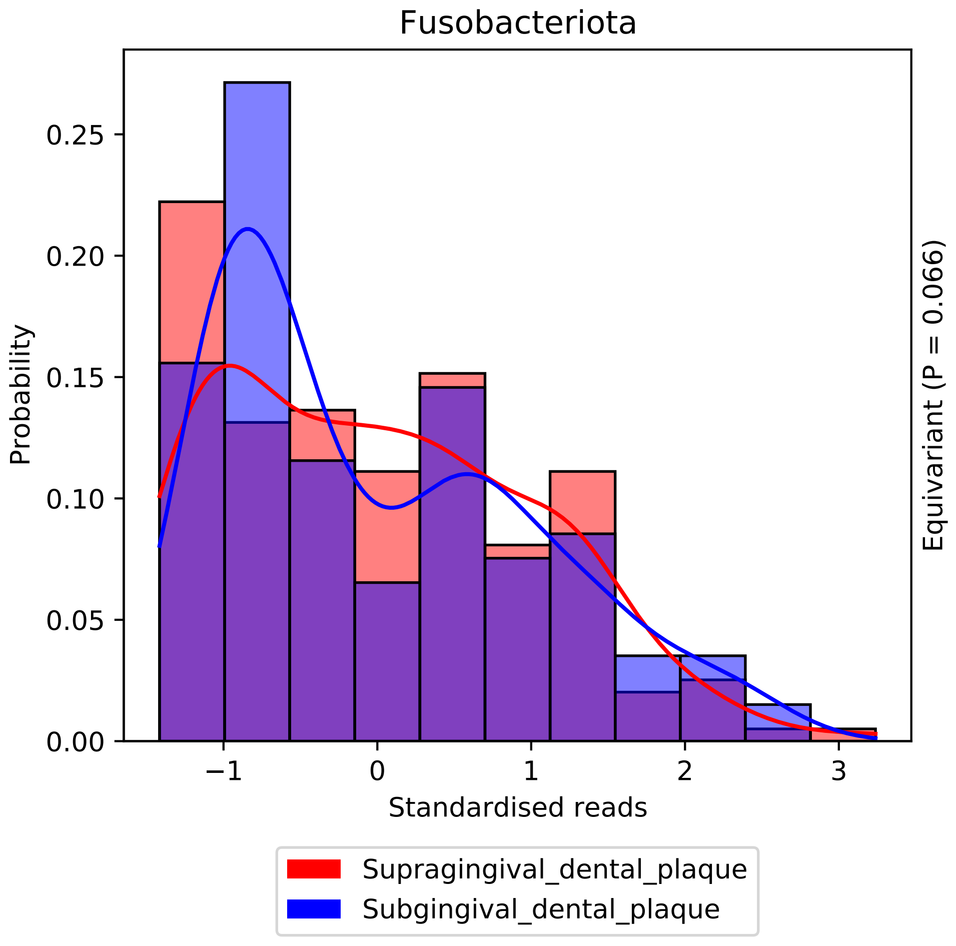
Fusobacteriota
Probability
0.00
0.05
0.10
0.15
0.20
0.25
−1
0
1
2
3
Standardised reads
Equivariant (P = 0.066)
Supragingival_dental_plaque
Subgingival_dental_plaque

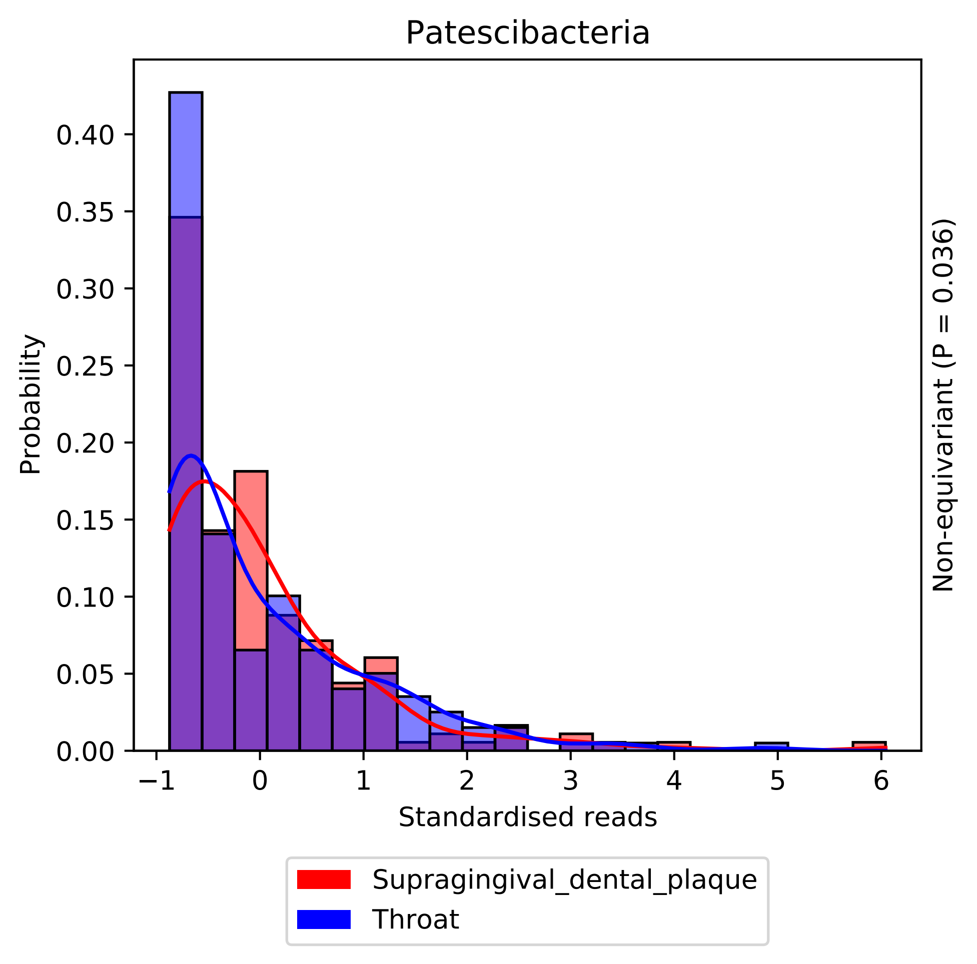
Patescibacteria
Probability
0.00
0.05
0.10
0.15
0.20
0.25
0.30
0.35
0.40
−1
0
1
2
3
4
5
6
Standardised reads
Non-equivariant (P = 0.036)
Supragingival_dental_plaque
Throat

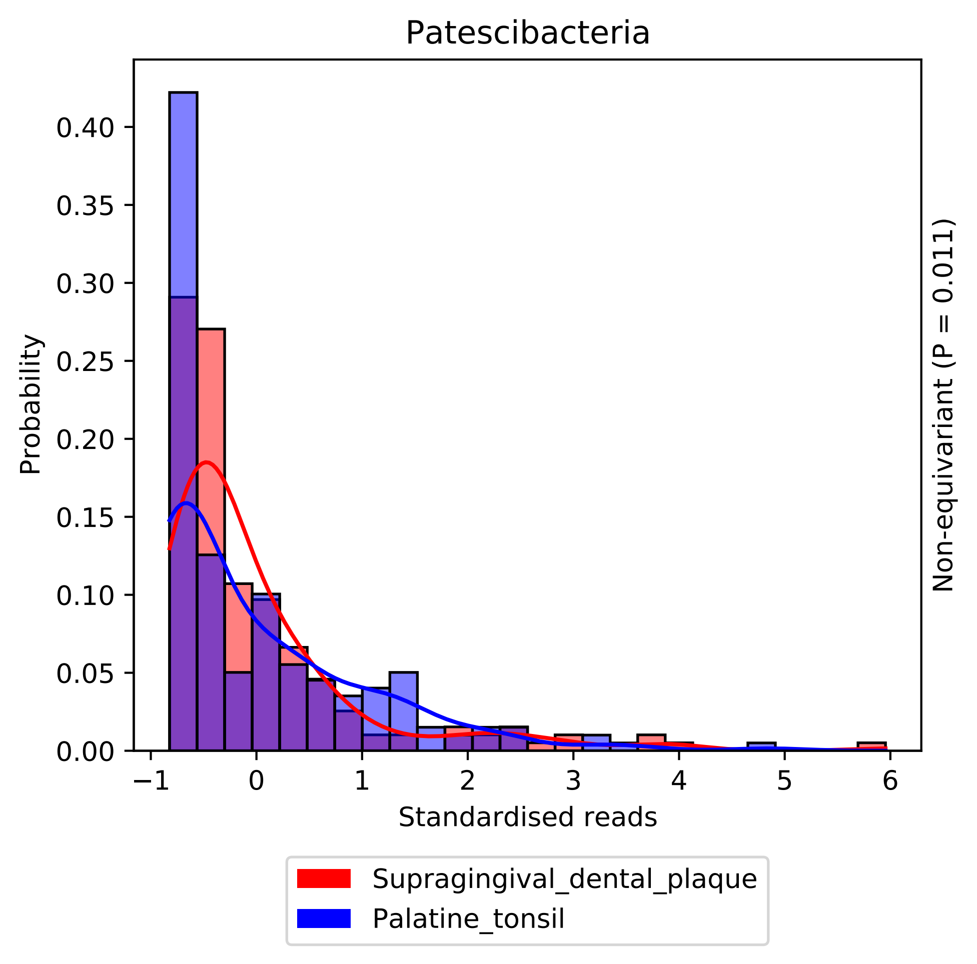

Patescibacteria
Probability
0.00
0.05
0.10
0.15
0.20
0.25
0.30
0.35
0.40
−1
0
1
2
3
4
5
6
Standardised reads
Non-equivariant (P = 0.011)
Supragingival_dental_plaque
Palatine_tonsil

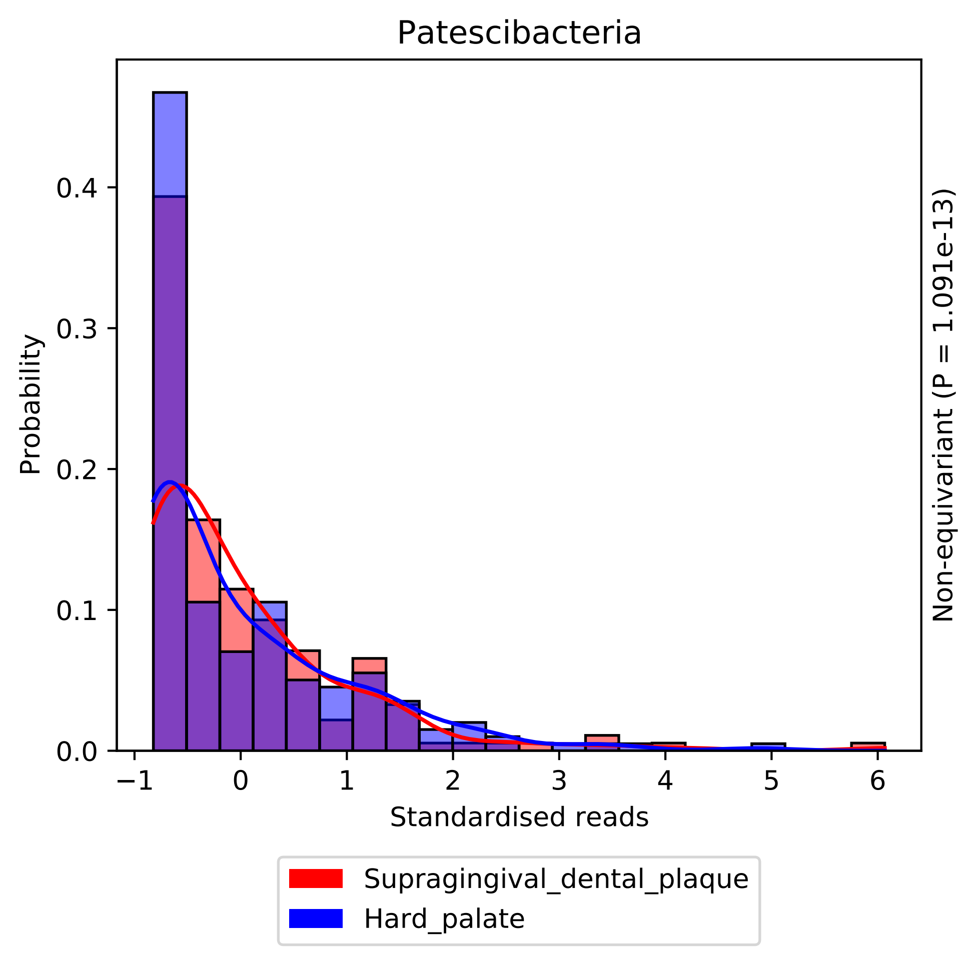

Patescibacteria
Probability
0.0
0.1
0.2
0.3
0.4
−1
0
1
2
3
4
5
6
Standardised reads
Non-equivariant (P = 1.091e-13)
Supragingival_dental_plaque
Hard_palate

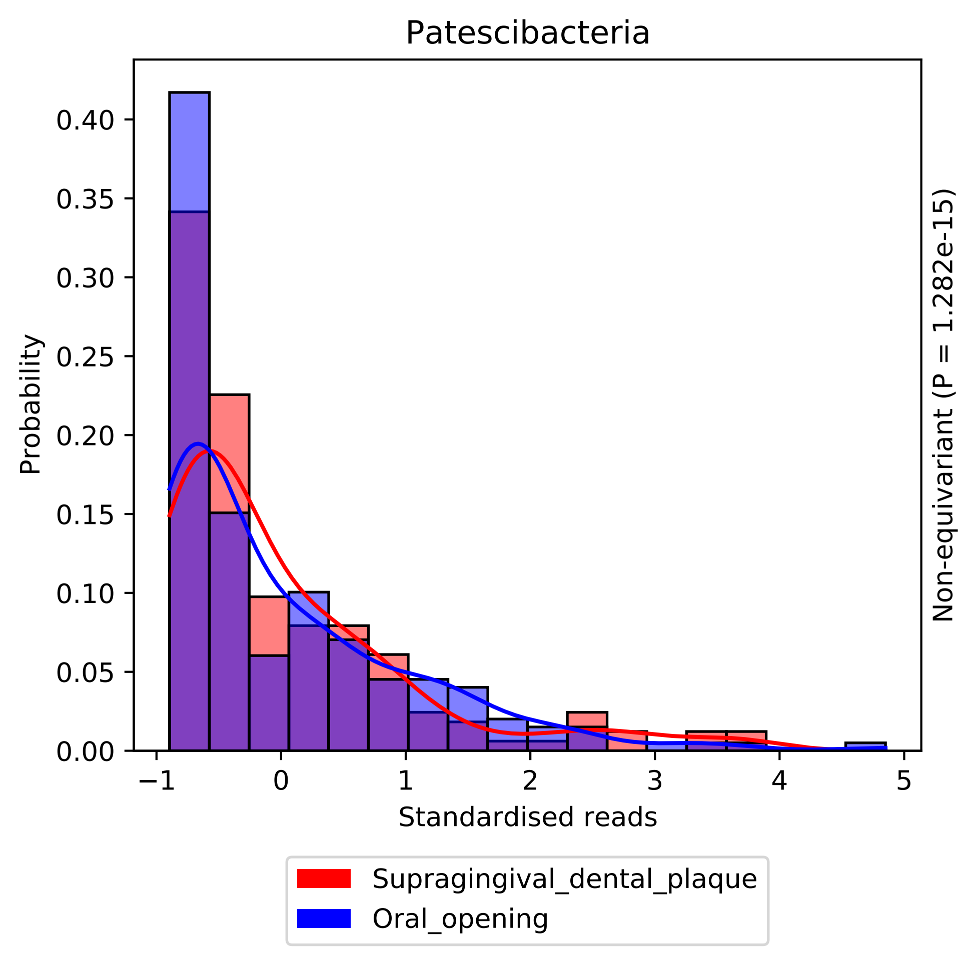

Patescibacteria
Probability
0.00
0.05
0.10
0.15
0.20
0.25
0.30
0.35
0.40
−1
0
1
2
3
4
5
Standardised reads
Non-equivariant (P = 1.282e-15)
Supragingival_dental_plaque
Oral_opening

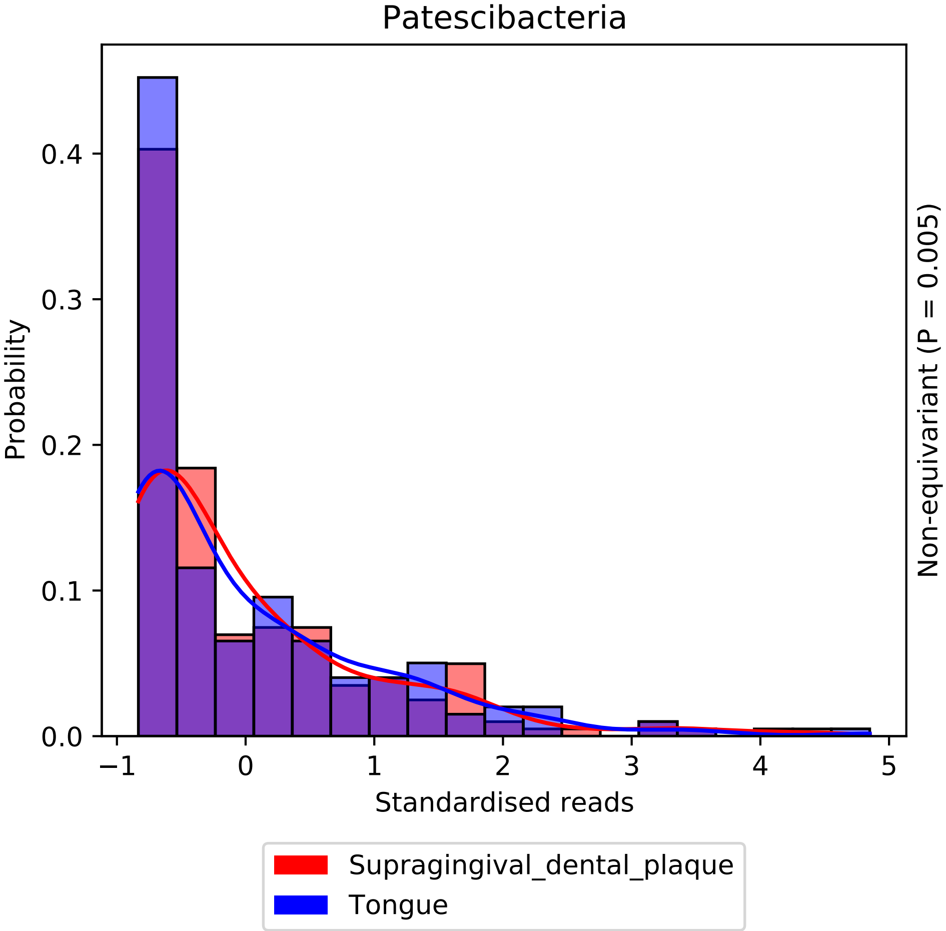

Patescibacteria
Probability
0.0
0.1
0.2
0.3
0.4
−1
0
1
2
3
4
5
Standardised reads
Non-equivariant (P = 0.005)
Supragingival_dental_plaque
Tongue

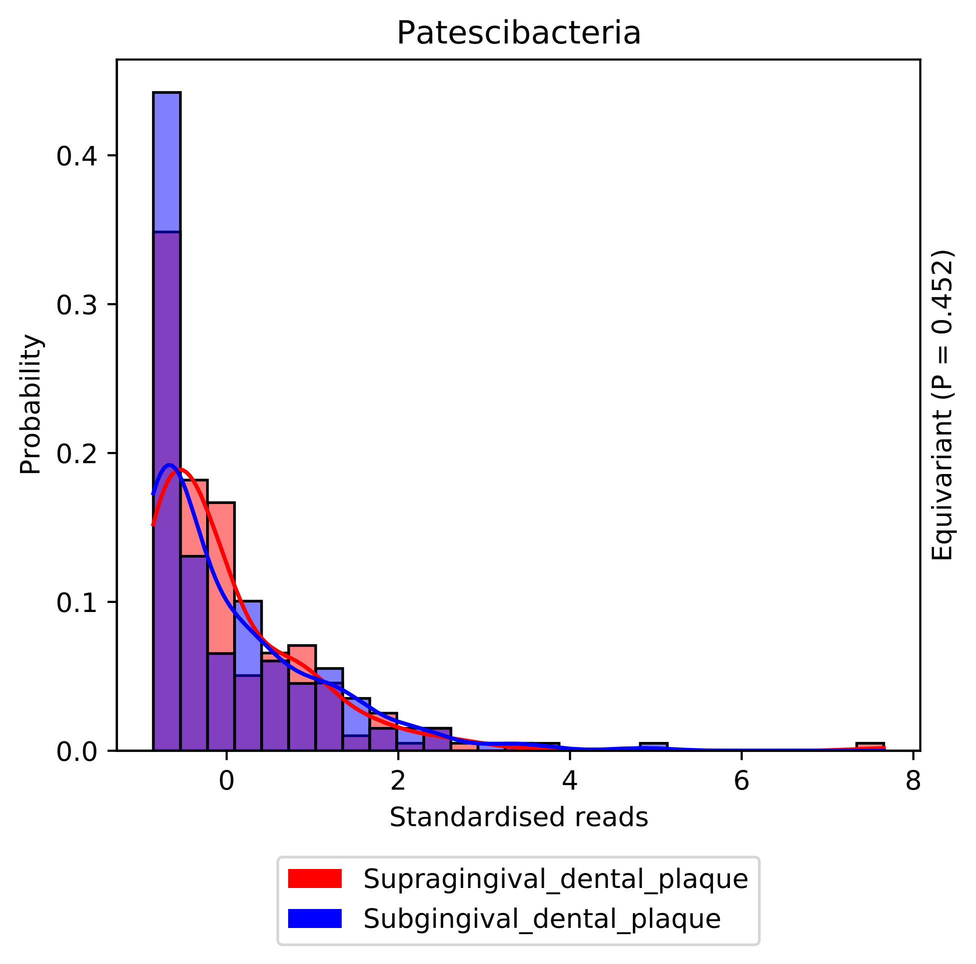
Patescibacteria
Probability
Standardised reads
Equivariant (P = 0.452)
Supragingival_dental_plaque
Subgingival_dental_plaque
0.0
0.1
0.2
0.3
0.4
0
2
4
6
8

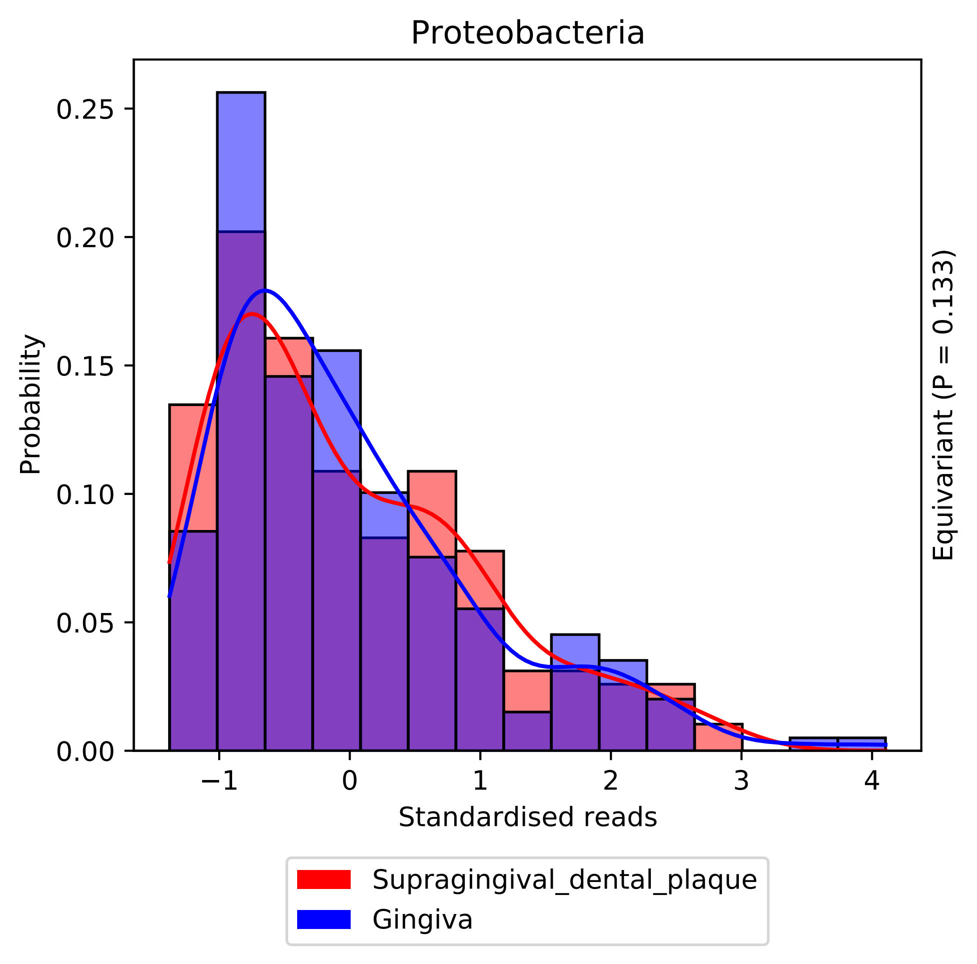

Proteobacteria
Probability
Standardised reads
Equivariant (P = 0.133)
0.00
0.05
0.10
0.15
0.20
0.25
−1
0
1
2
3
4
Supragingival_dental_plaque
Gingiva

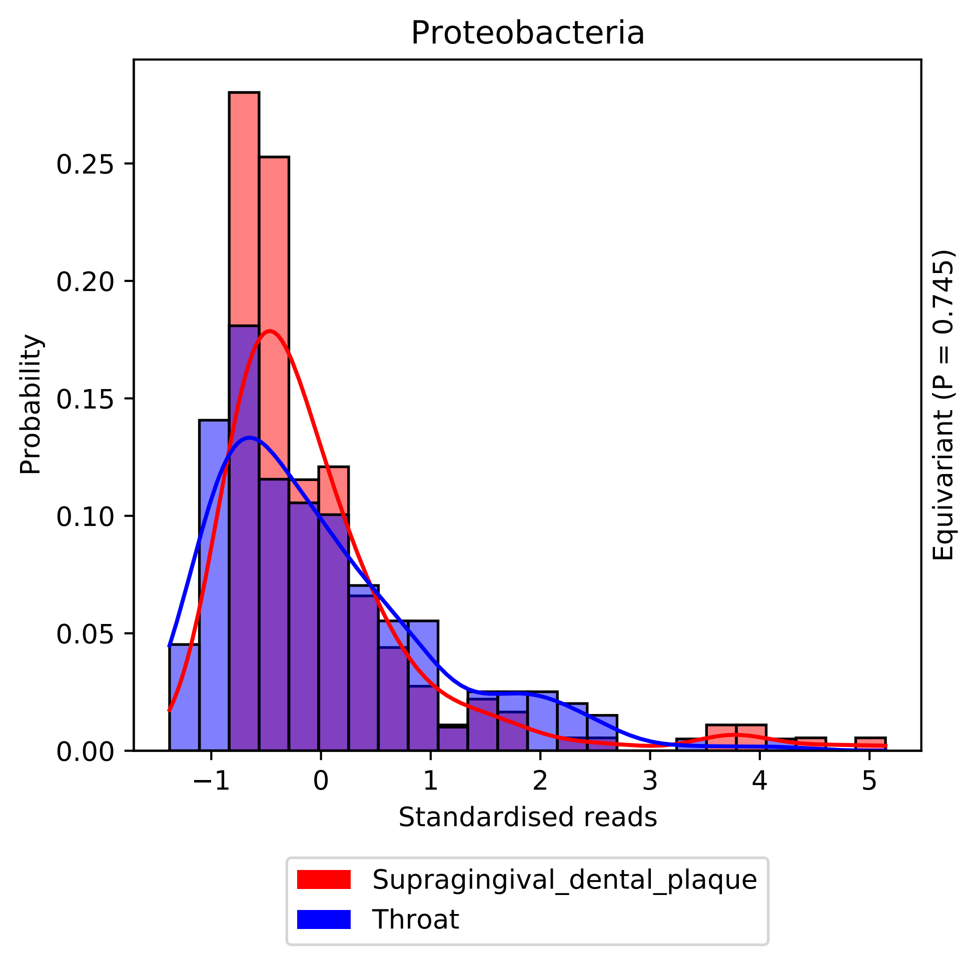

Proteobacteria
Probability
0.00
0.05
0.10
0.15
0.20
0.25
−1
0
1
2
3
4
5
Standardised reads
Equivariant (P = 0.745)
Supragingival_dental_plaque
Throat

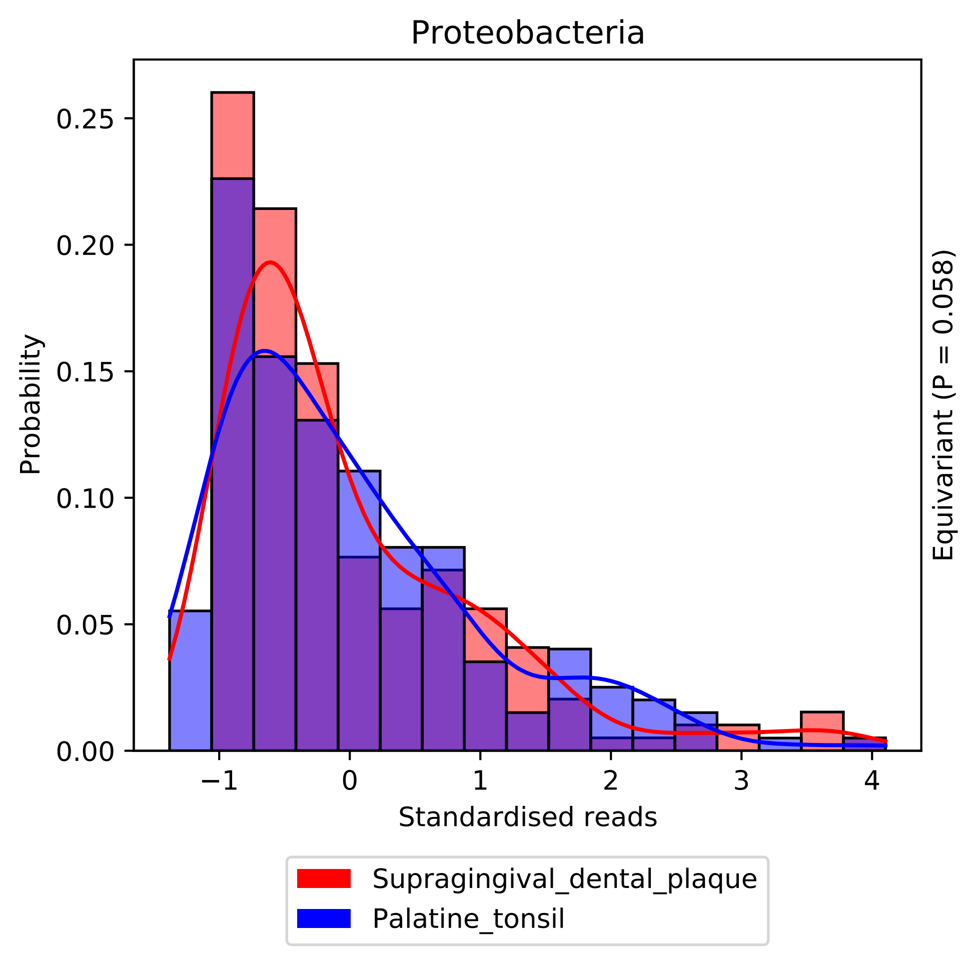
Proteobacteria
Probability
0.00
0.05
0.10
0.15
0.20
0.25
−1
0
1
2
3
4
Standardised reads
Equivariant (P = 0.058)
Supragingival_dental_plaque
Palatine_tonsil

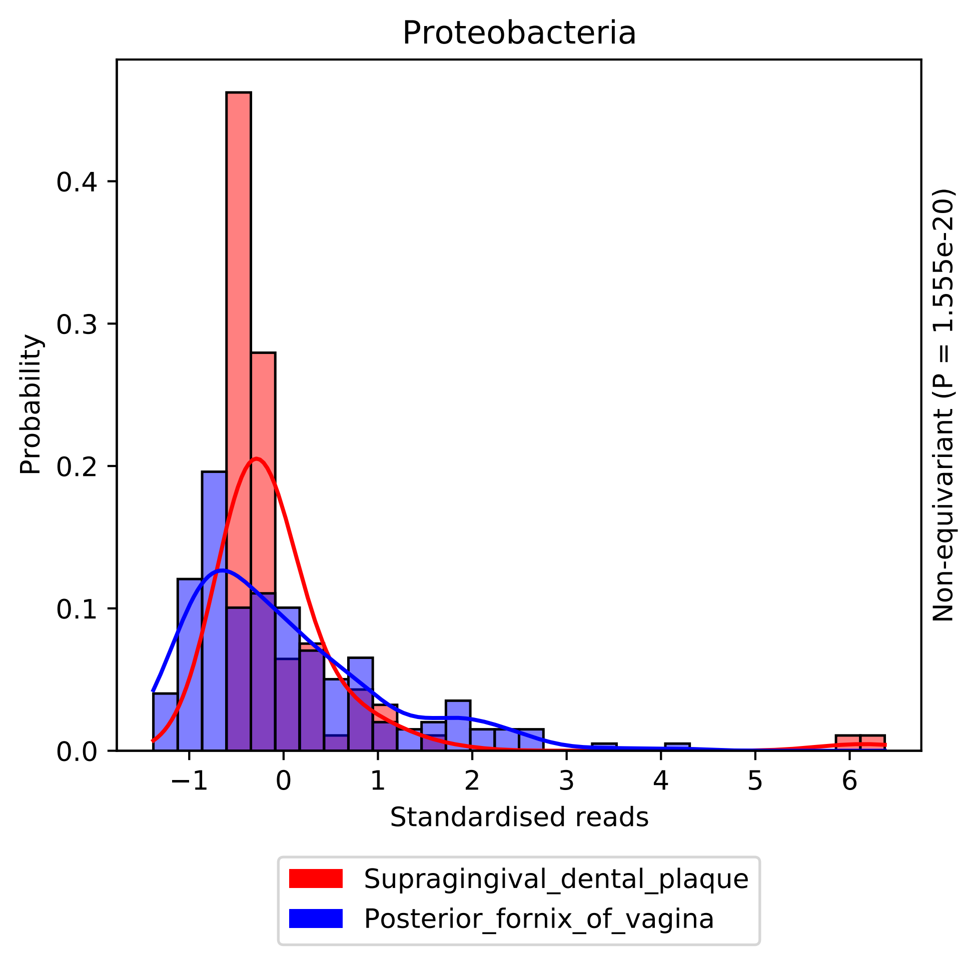

Proteobacteria
Probability
0.0
0.1
0.2
0.3
0.4
−1
0
1
2
3
4
5
6
Standardised reads
Non-equivariant (P = 1.555e-20)
Supragingival_dental_plaque
Posterior_fornix_of_vagina

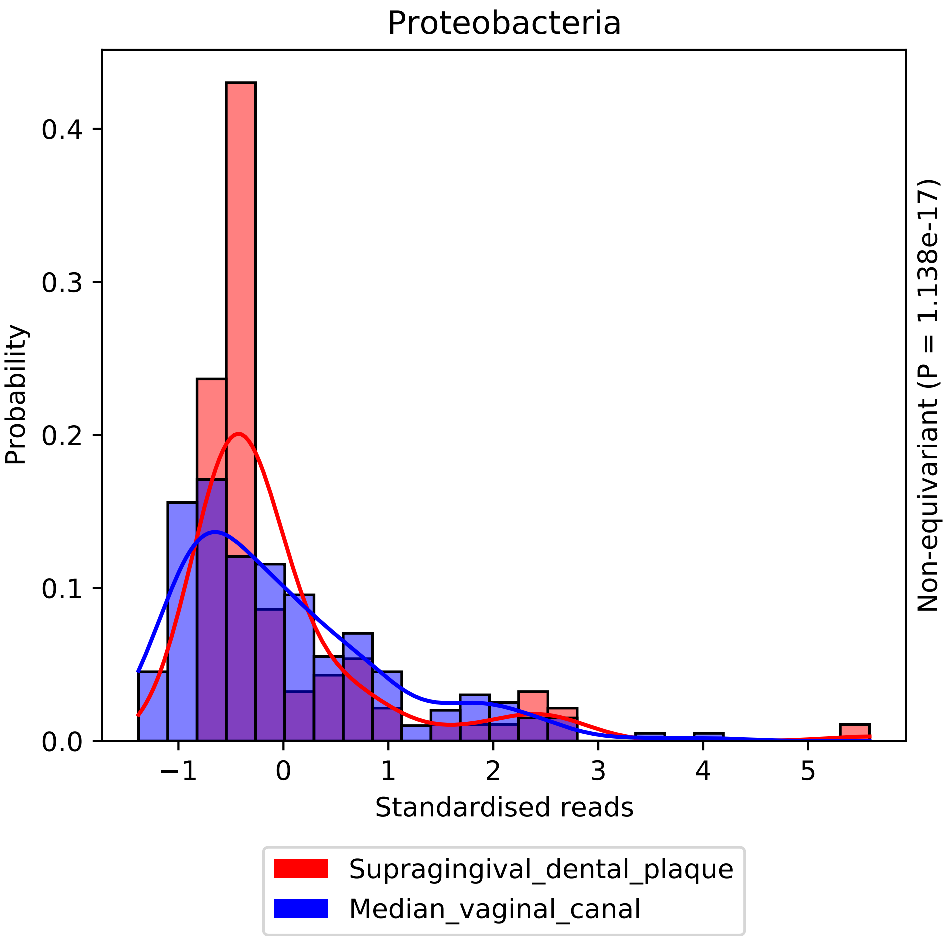

Proteobacteria
Probability
0.0
0.1
0.2
0.3
0.4
−1
0
1
2
3
4
5
Standardised reads
Non-equivariant (P = 1.138e-17)
Supragingival_dental_plaque
Median_vaginal_canal

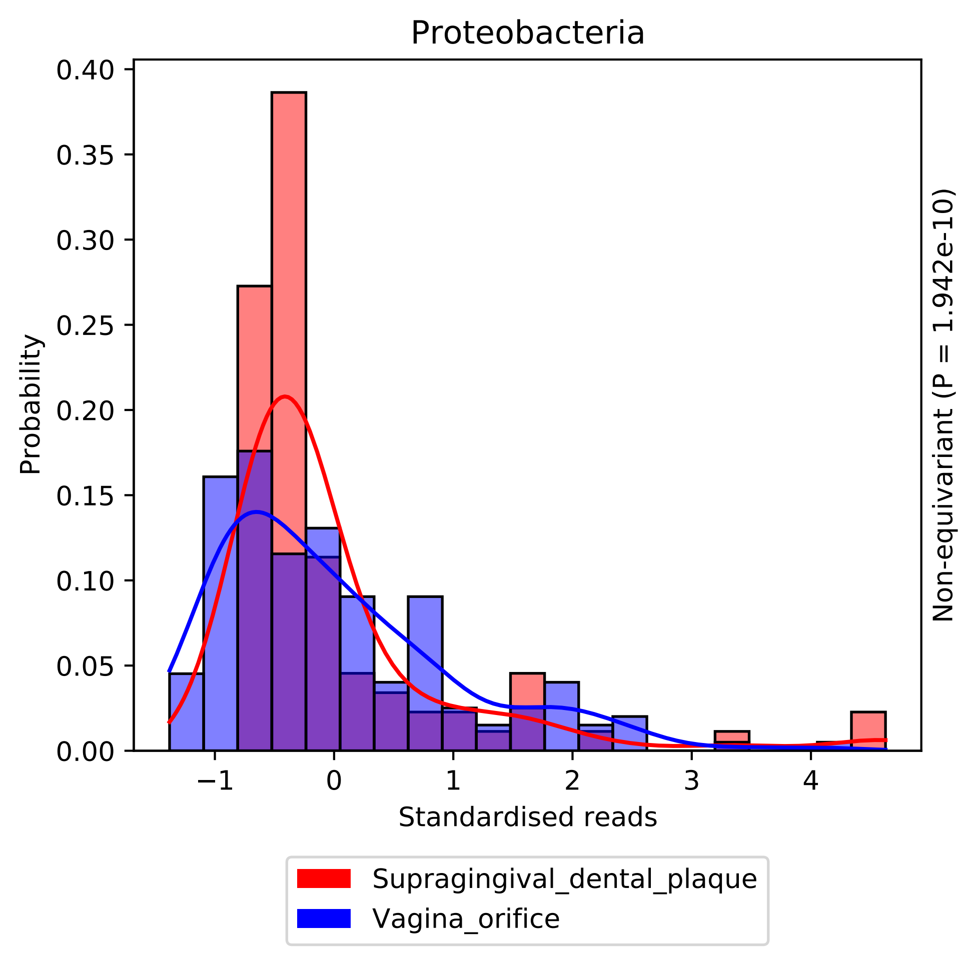

Proteobacteria
Probability
0.40
0.35
0.30
0.25
0.20
0.15
0.10
0.05
0.00
−1
0
1
2
3
4
Standardised reads
Non-equivariant (P = 1.942e-10)
Supragingival_dental_plaque
Vagina_orifice

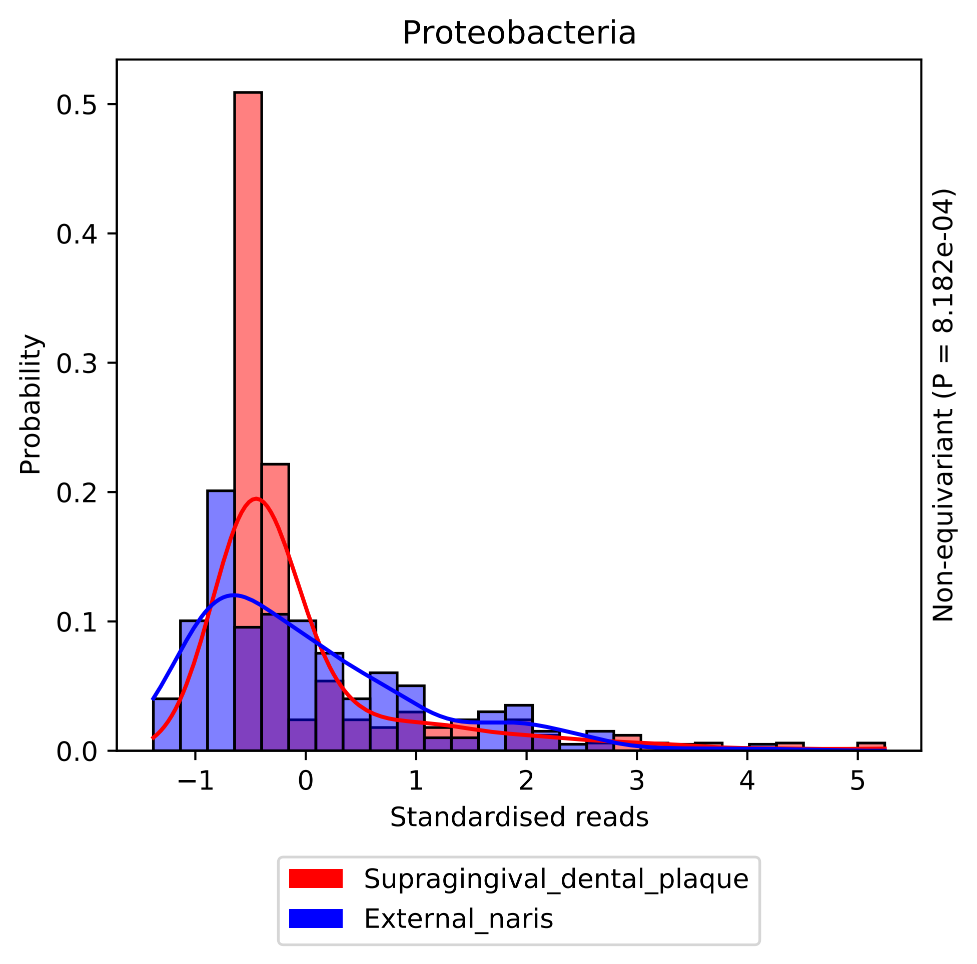

Proteobacteria
Probability
0.0
0.1
0.2
0.3
0.4
0.5
−1
0
1
2
3
4
5
Standardised reads
Non-equivariant (P = 8.182e-04)
Supragingival_dental_plaque
External_naris

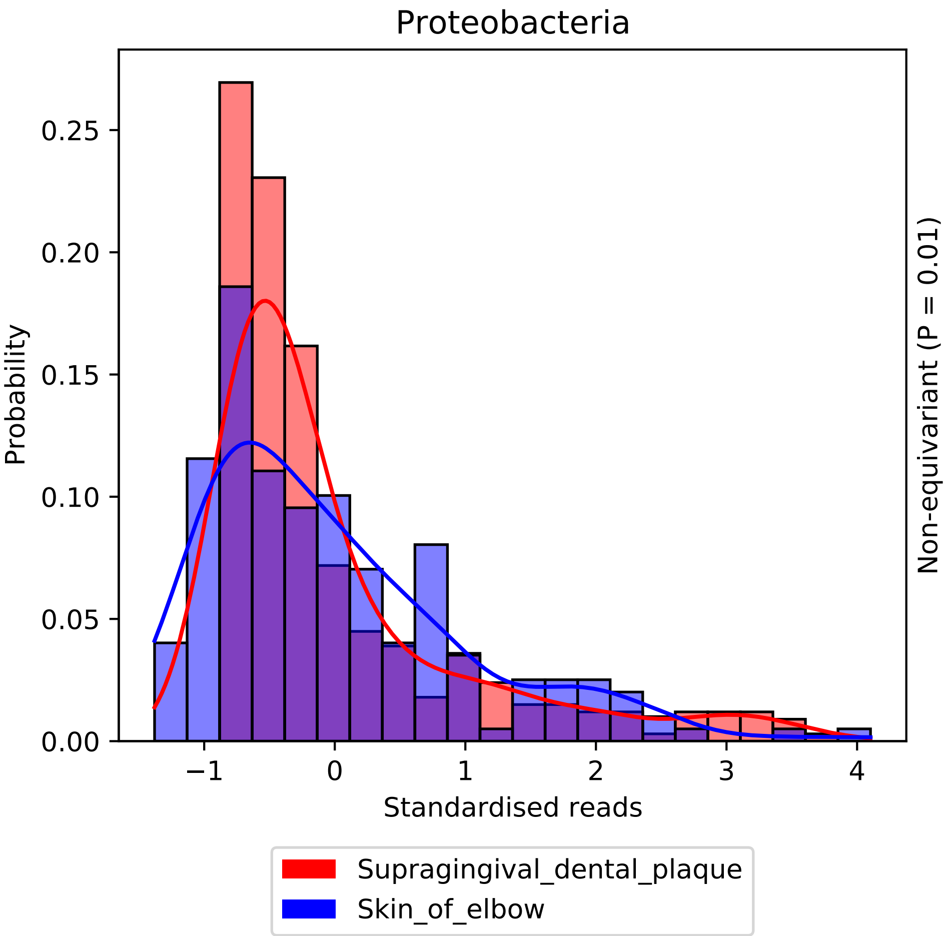
Proteobacteria
Probability
0.00
0.05
0.10
0.15
0.20
0.25
−1
0
1
2
3
4
Standardised reads
Non-equivariant (P = 0.01)
Supragingival_dental_plaque
Skin_of_elbow

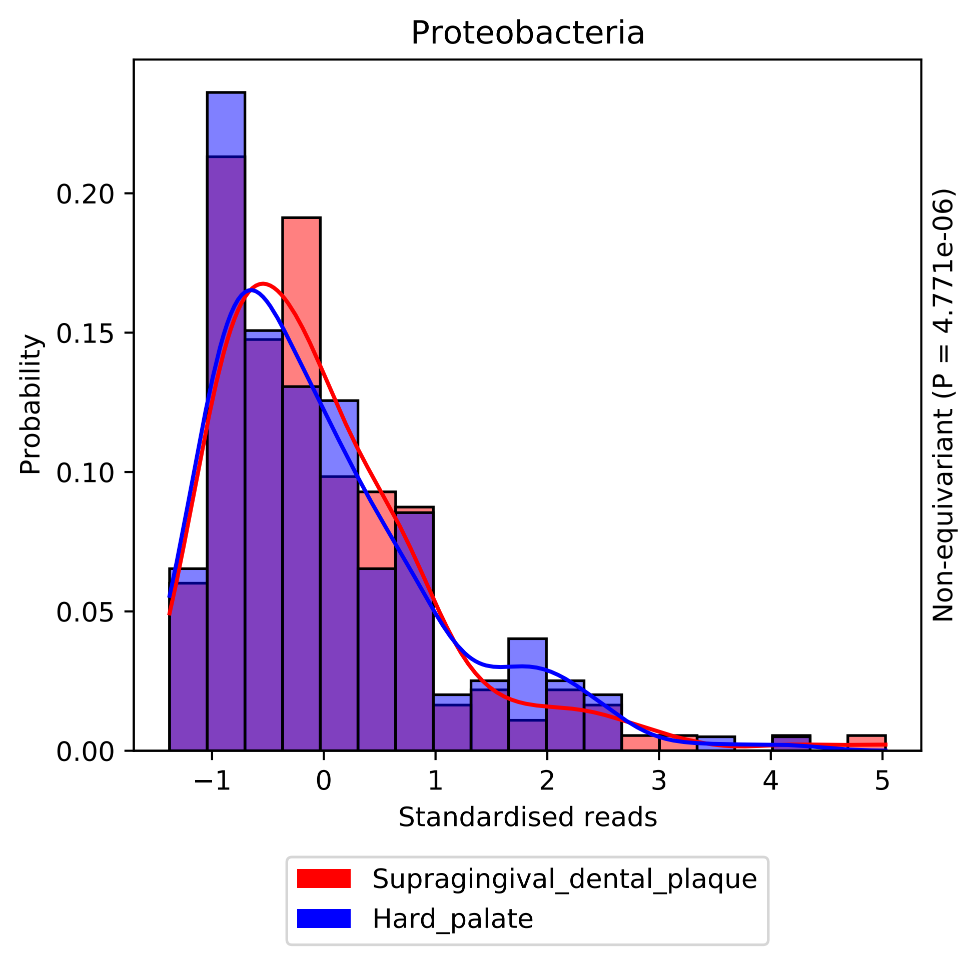

Proteobacteria
Probability
0.00
0.05
0.10
0.15
0.20
−1
0
1
2
3
4
5
Standardised reads
Non-equivariant (P = 4.771e-06)
Supragingival_dental_plaque
Hard_palate

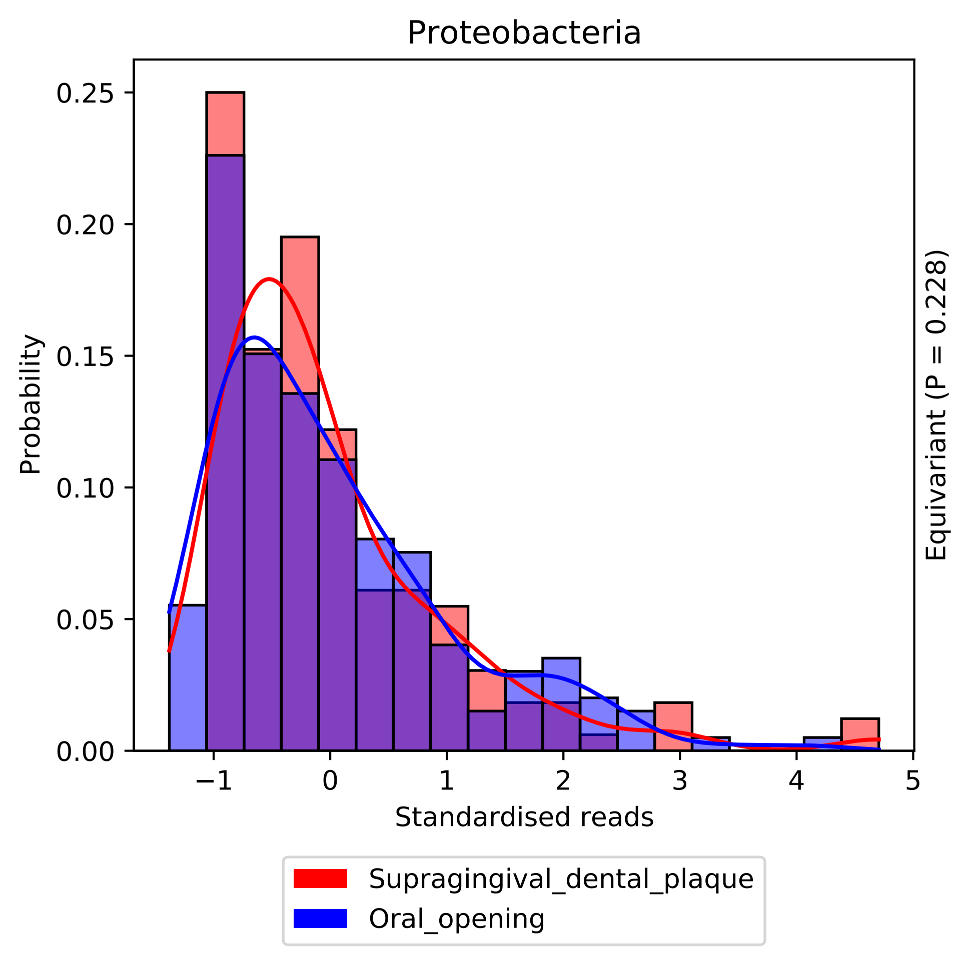

Proteobacteria
Probability
0.25
0.20
0.15
0.10
0.05
0.00
−1
0
1
2
3
4
5
Standardised reads
Equivariant (P = 0.228)
Supragingival_dental_plaque
Oral_opening

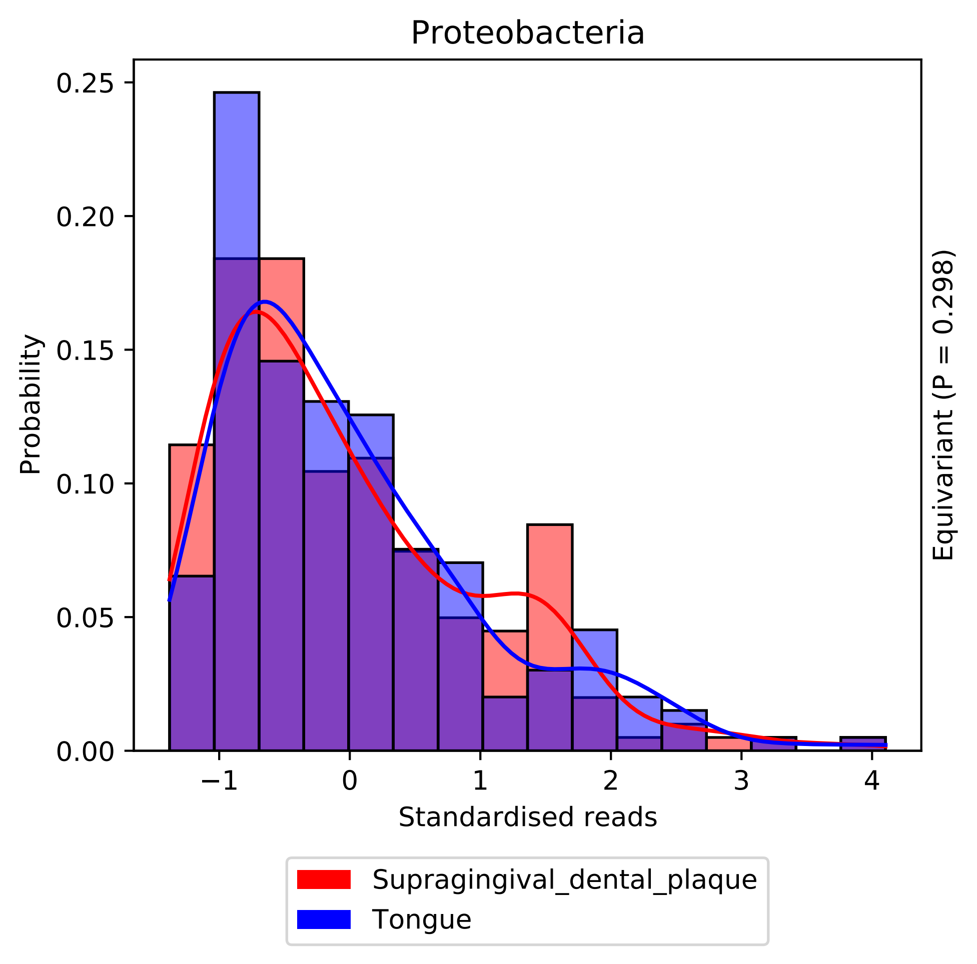

Proteobacteria
Probability
0.25
0.20
0.15
0.10
0.05
0.00
−1
0
1
2
3
4
Standardised reads
Equivariant (P = 0.298)
Supragingival_dental_plaque
Tongue

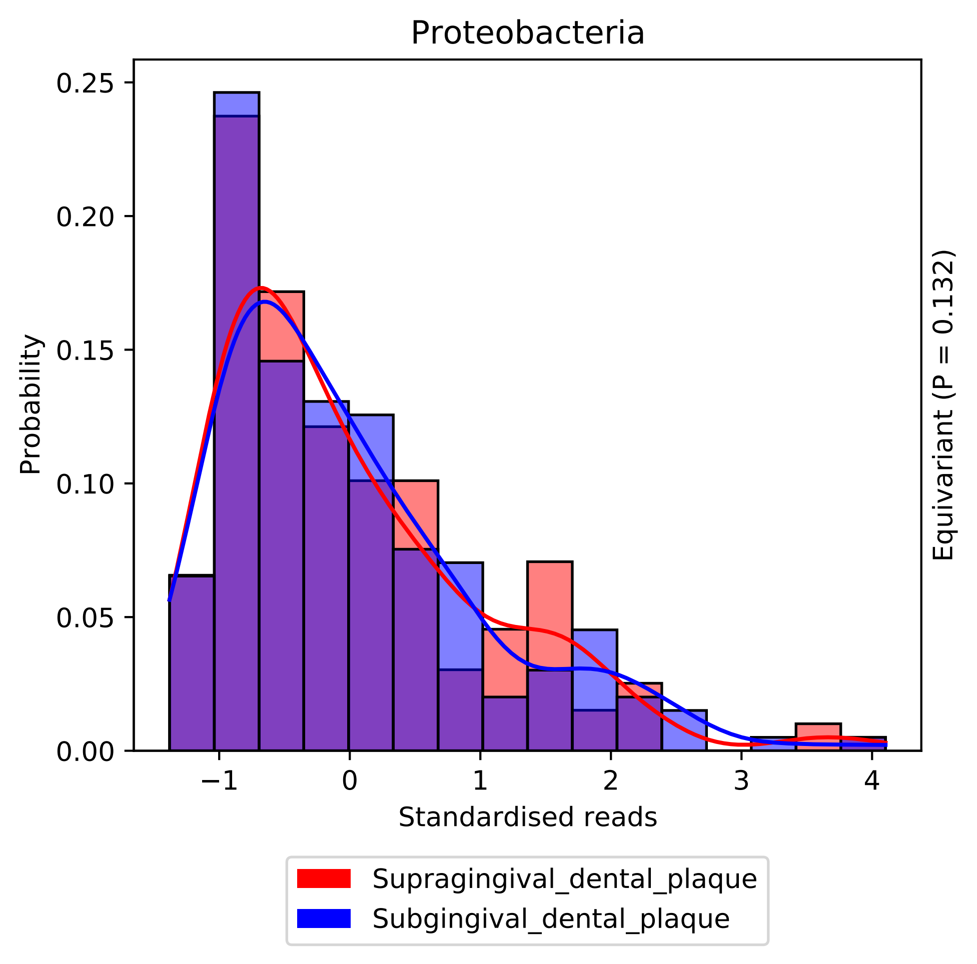
Proteobacteria
Probability
0.25
0.20
0.15
0.10
0.05
0.00
−1
0
1
2
3
4
Standardised reads
Equivariant (P = 0.132)
Supragingival_dental_plaque
Subgingival_dental_plaque

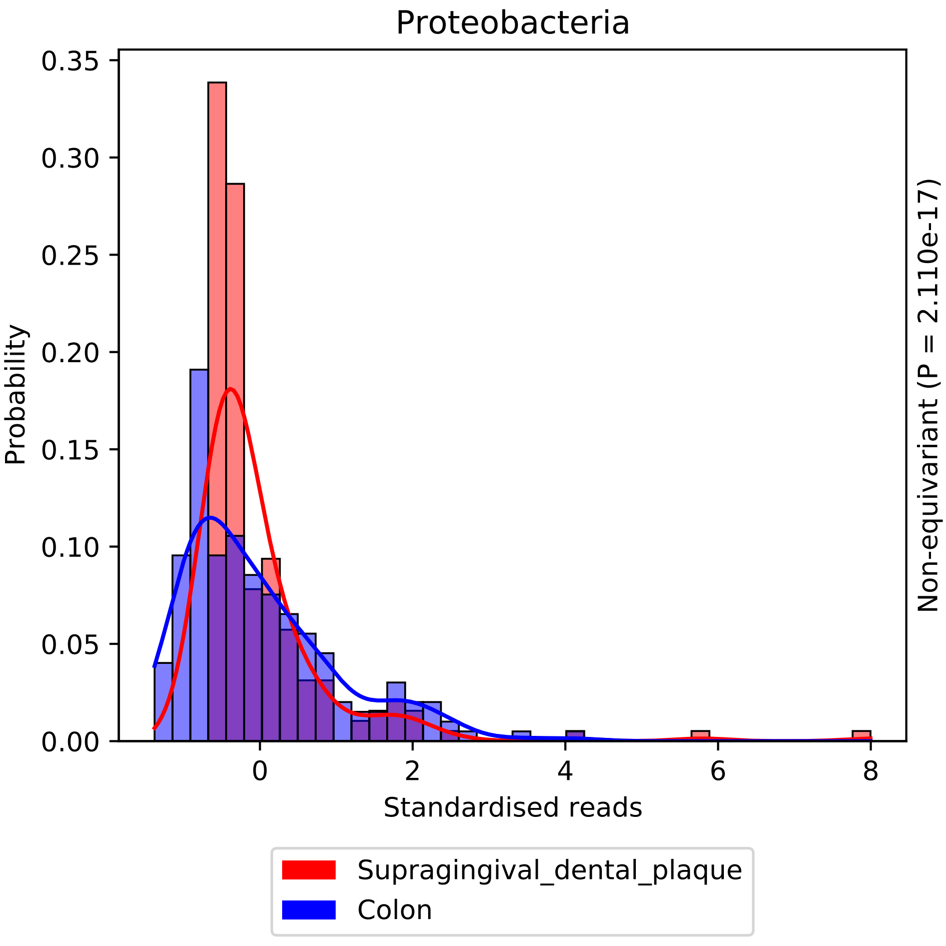
Proteobacteria
Probability
0.35
0.30
0.25
0.20
0.15
0.10
0.05
0.00
0
2
4
6
8
Standardised reads
Non-equivariant (P = 2.110e-17)
Supragingival_dental_plaque
Colon

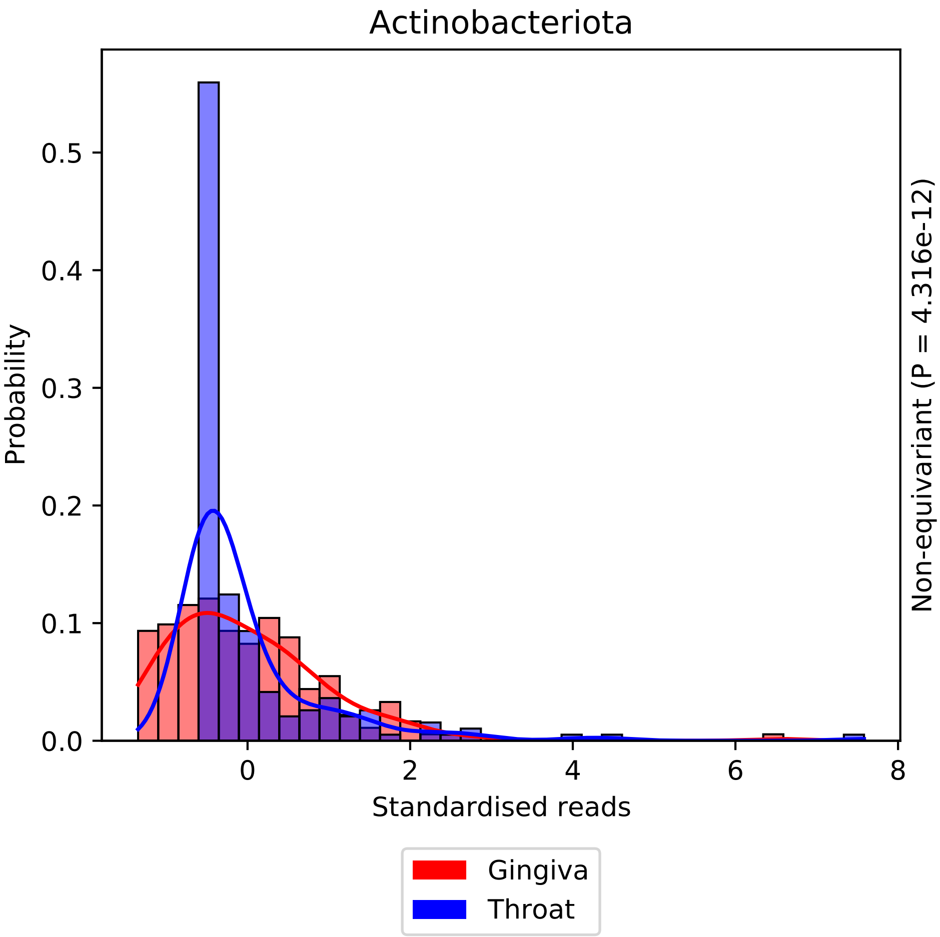

Actinobacteriota
0.0
0.1
0.2
0.3
0.4
0.5
Probability
0
2
4
6
8
Standardised reads
Non-equivariant (P = 4.316e-12)
Gingiva
Throat

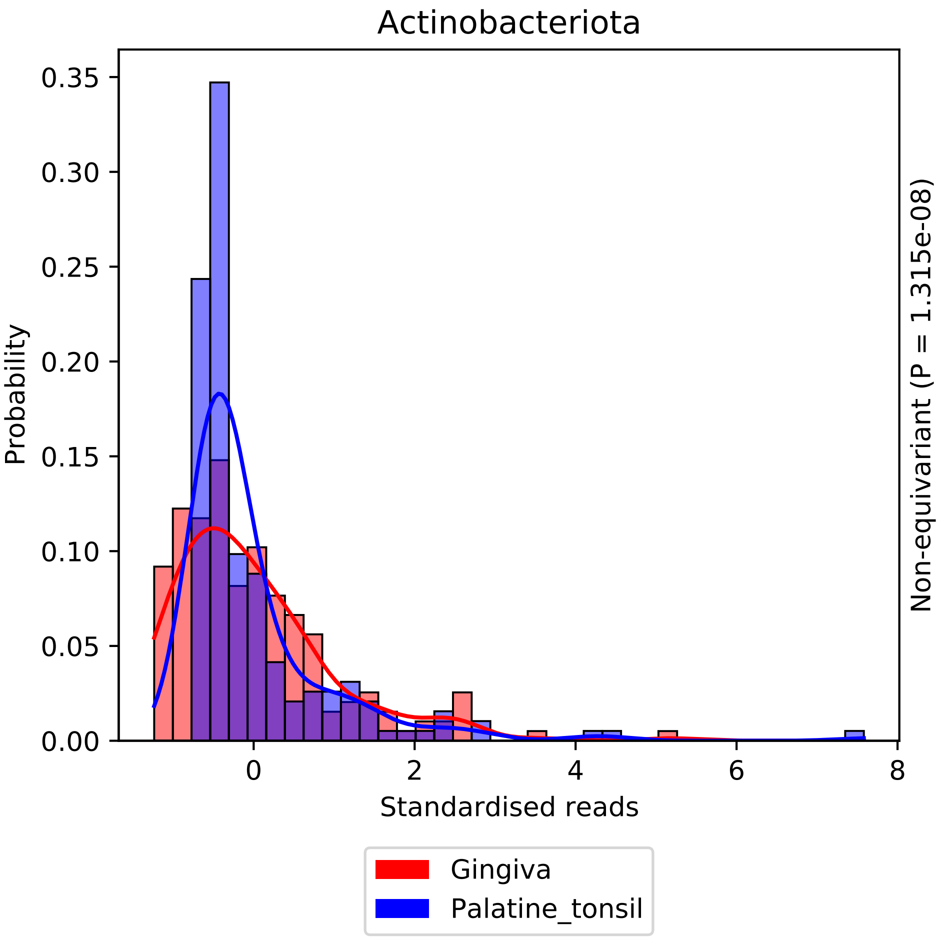

Actinobacteriota
Probability
0.35
0.30
0.25
0.20
0.15
0.10
0.05
0.00
0
2
4
6
8
Standardised reads
Non-equivariant (P = 1.315e-08)
Gingiva
Palatine_tonsil

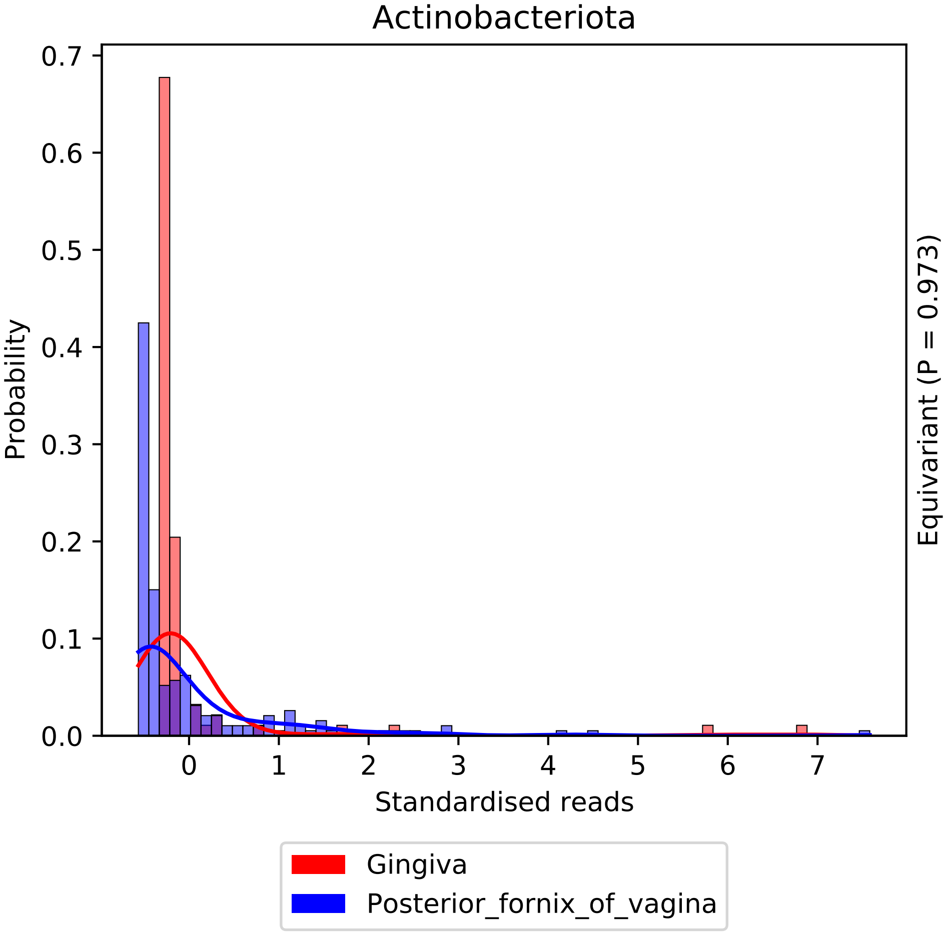

Actinobacteriota
Probability
0.7
0.6
0.5
0.4
0.3
0.2
0.1
0.0
0
1
2
3
4
5
6
7
Standardised reads
Equivariant (P = 0.973)
Gingiva
Posterior_fornix_of_vagina

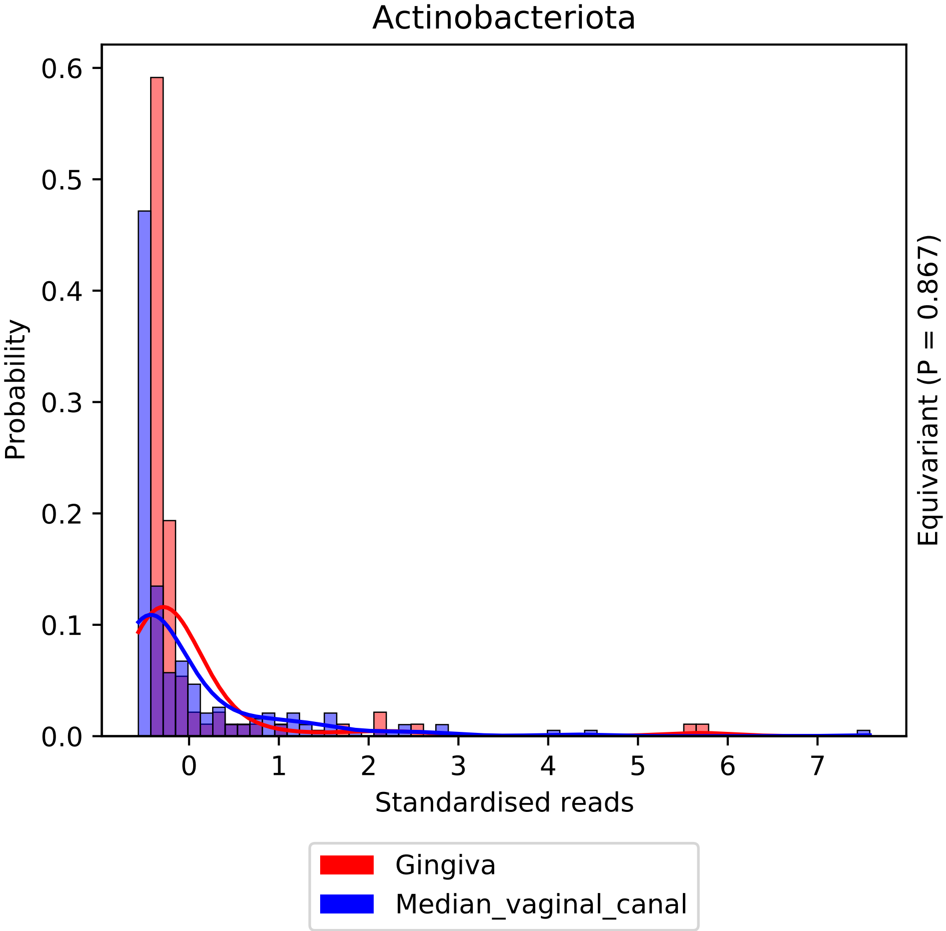

Actinobacteriota
Probability
Standardised reads
Equivariant (P = 0.867)
0.0
0.1
0.2
0.3
0.4
0.5
0.6
0
1
2
3
4
5
6
7
Gingiva
Median_vaginal_canal

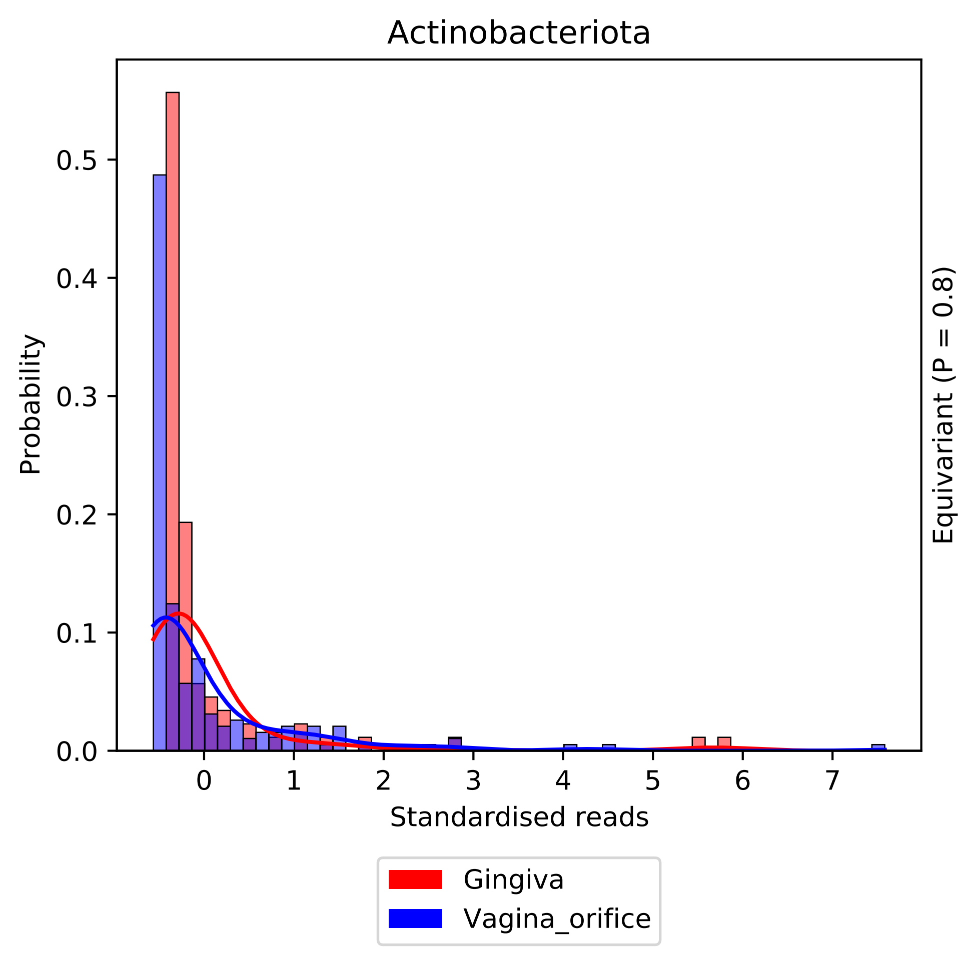
Actinobacteriota
Probability
Standardised reads
Equivariant (P = 0.8)
0.0
0.1
0.2
0.3
0.4
0.5
0
1
2
3
4
5
6
7
Gingiva
Vagina_orifice

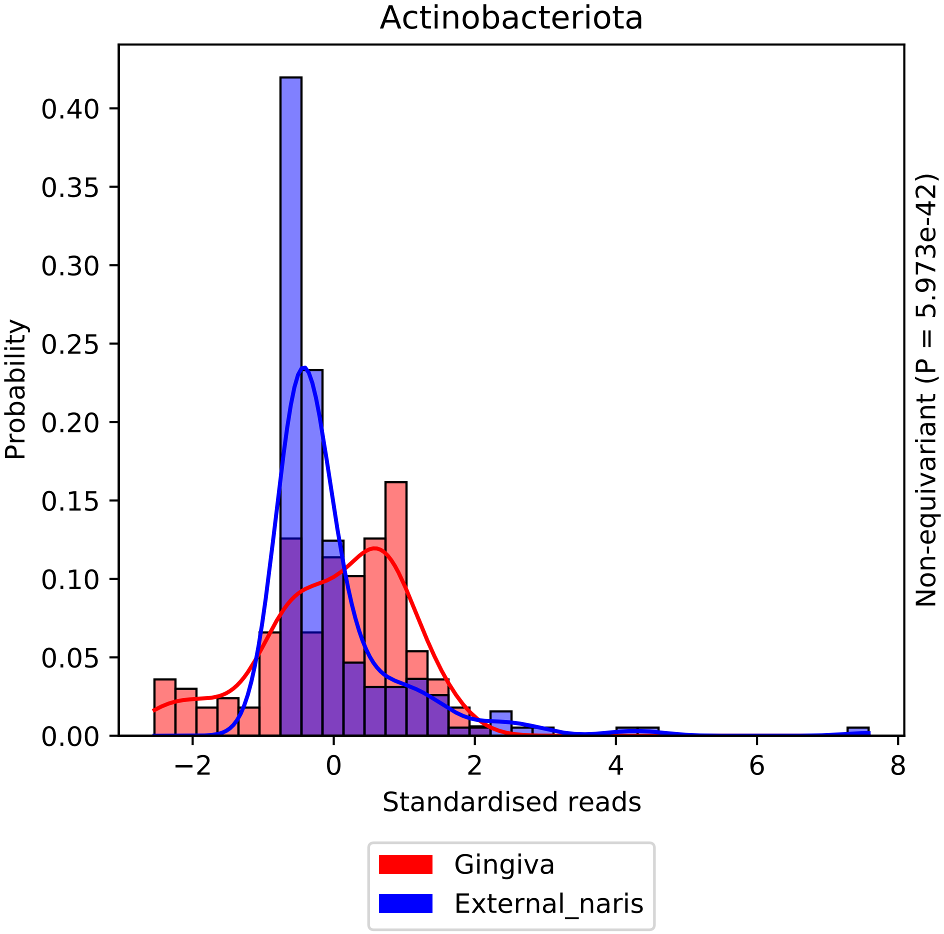

Actinobacteriota
Probability
0.40
0.35
0.30
0.25
0.20
0.15
0.10
0.05
0.00
−2
0
2
4
6
8
Standardised reads
Non-equivariant (P = 5.973e-42)
Gingiva
External_naris

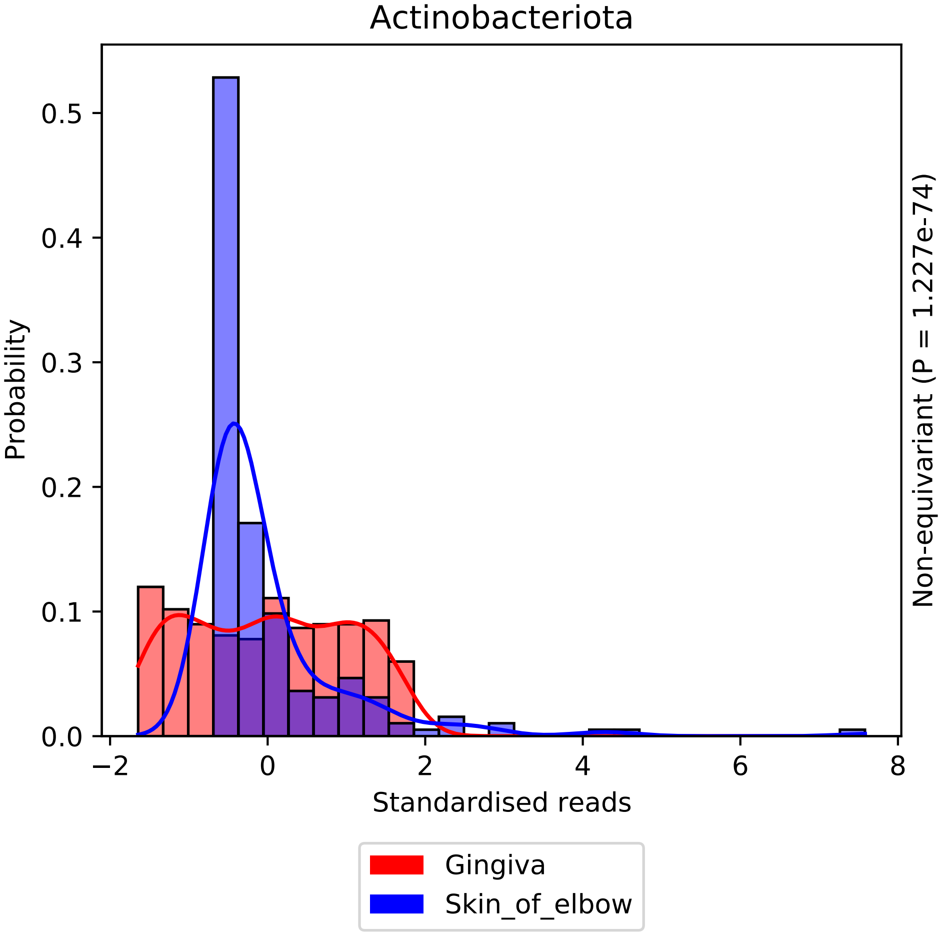

Actinobacteriota
Probability
Standardised reads
Non-equivariant (P = 1.227e-74)
0.0
0.1
0.2
0.3
0.4
0.5
−2
0
2
4
6
8
Gingiva
Skin_of_elbow

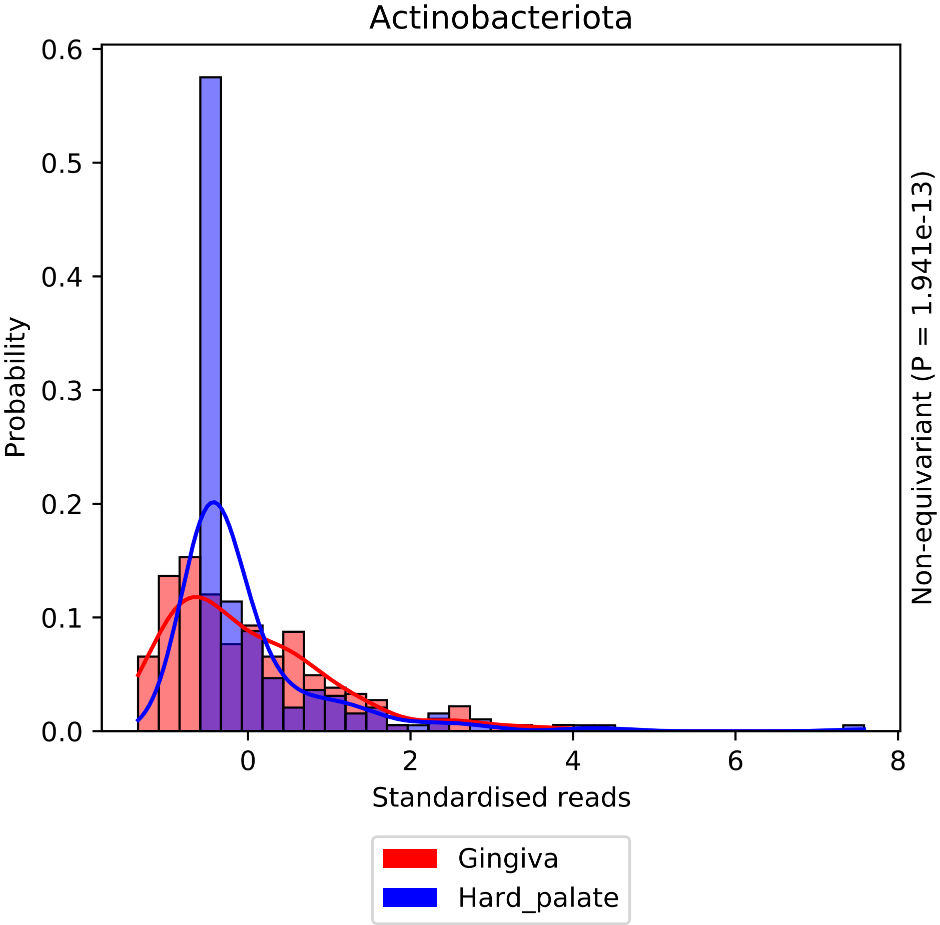

Actinobacteriota
Probability
0.0
0.1
0.2
0.3
0.4
0.5
0.6
0
2
4
6
8
Standardised reads
Non-equivariant (P = 1.941e-13)
Gingiva
Hard_palate

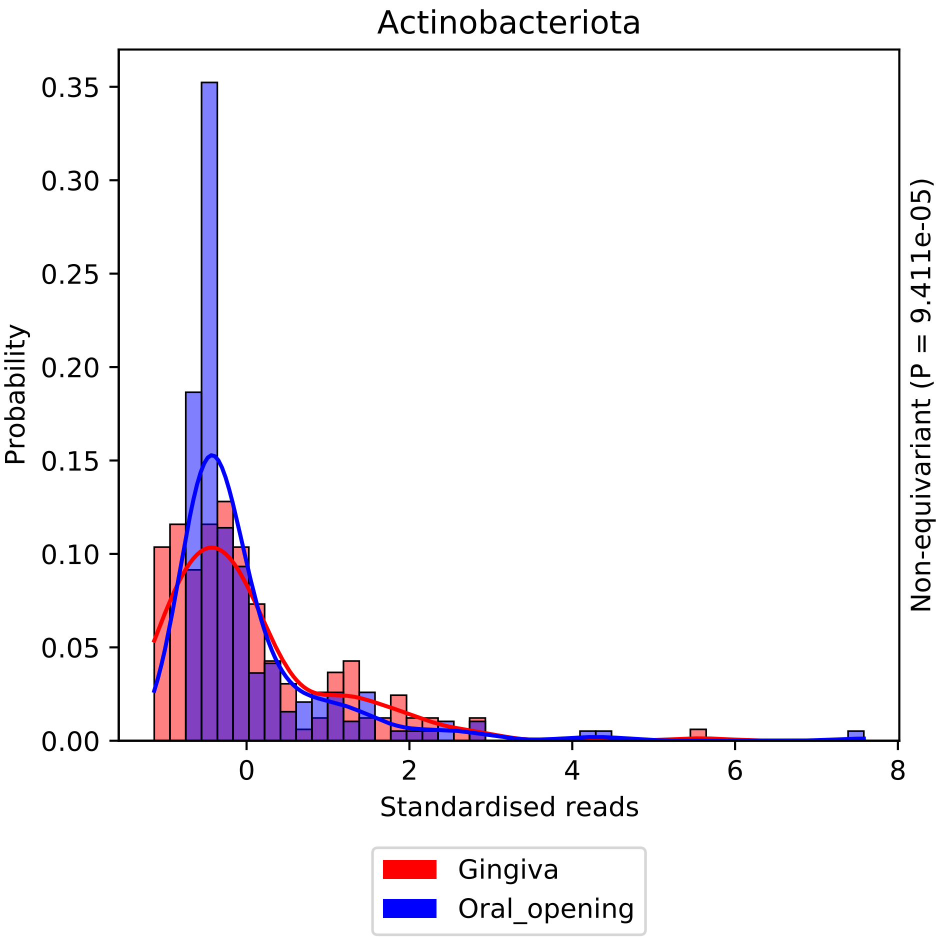

Actinobacteriota
Probability
0.35
0.30
0.25
0.20
0.15
0.10
0.05
0.00
0
2
4
6
8
Standardised reads
Non-equivariant (P = 9.411e-05)
Gingiva
Oral_opening

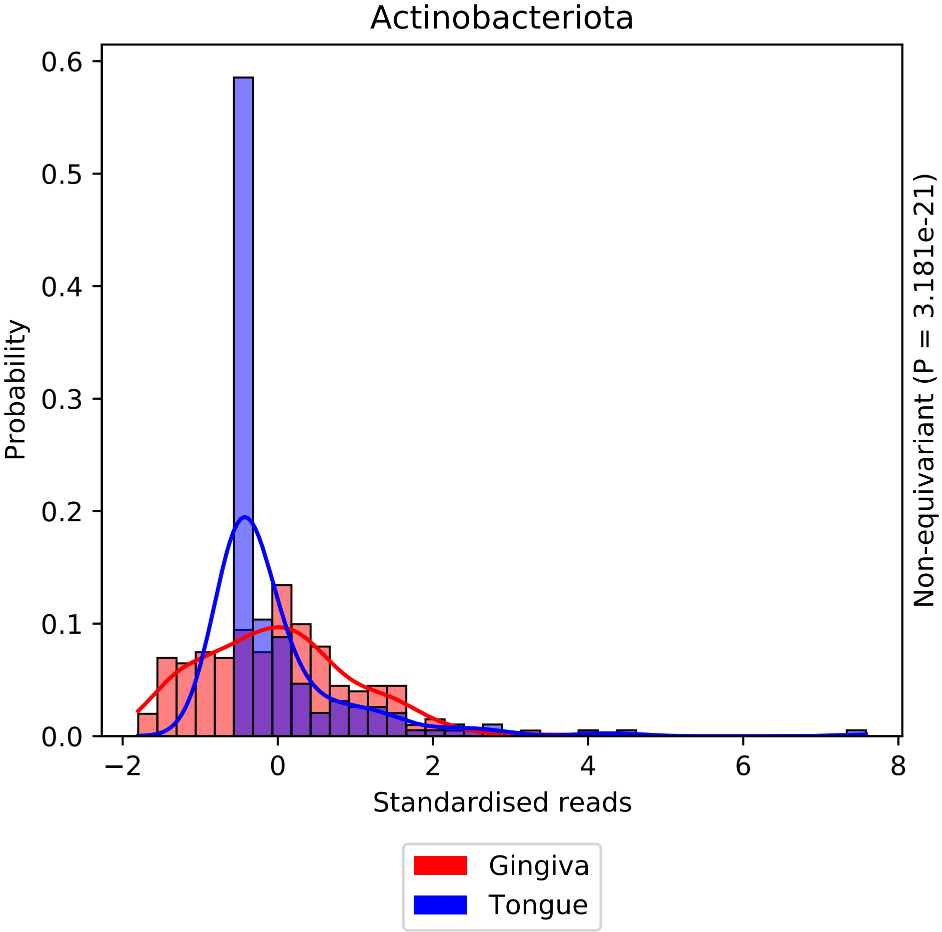

Actinobacteriota
Probability
0.6
0.5
0.4
0.3
0.2
0.1
0.0
−2
0
2
4
6
8
Standardised reads
Non-equivariant (P = 3.181e-21)
Gingiva
Tongue

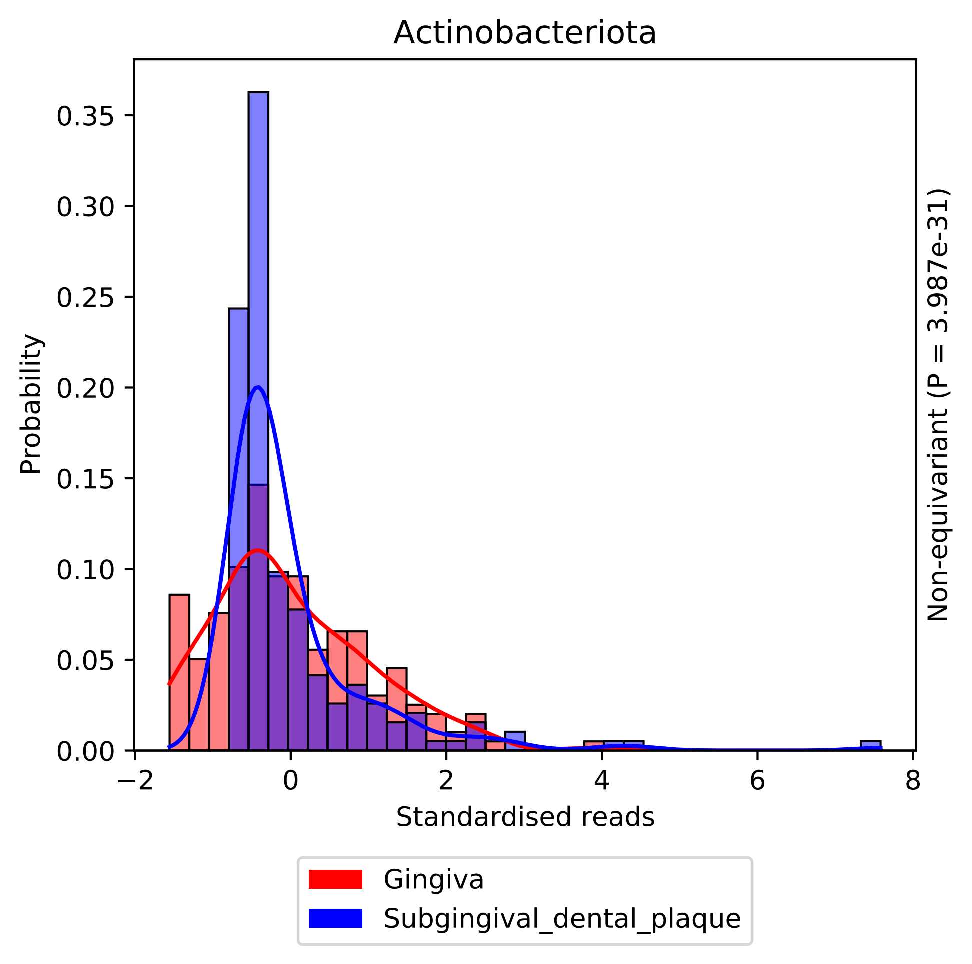

Actinobacteriota
0.35
0.30
0.25
0.20
0.15
0.10
0.05
0.00
Probability
−2
0
2
4
6
8
Standardised reads
Non-equivariant (P = 3.987e-31)
Gingiva
Subgingival_dental_plaque

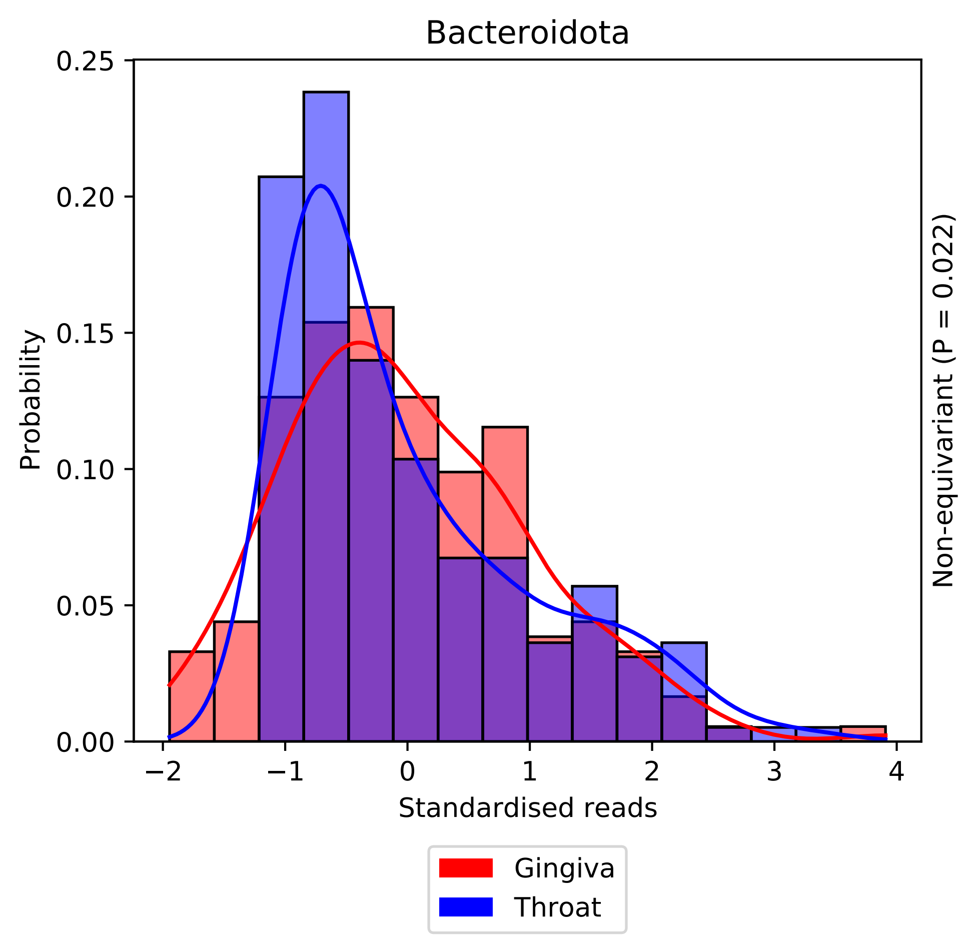

Bacteroidota
0.25
0.20
0.15
0.10
0.05
0.00
Probability
−2
−1
0
1
2
3
4
Standardised reads
Non-equivariant (P = 0.022)
Gingiva
Throat

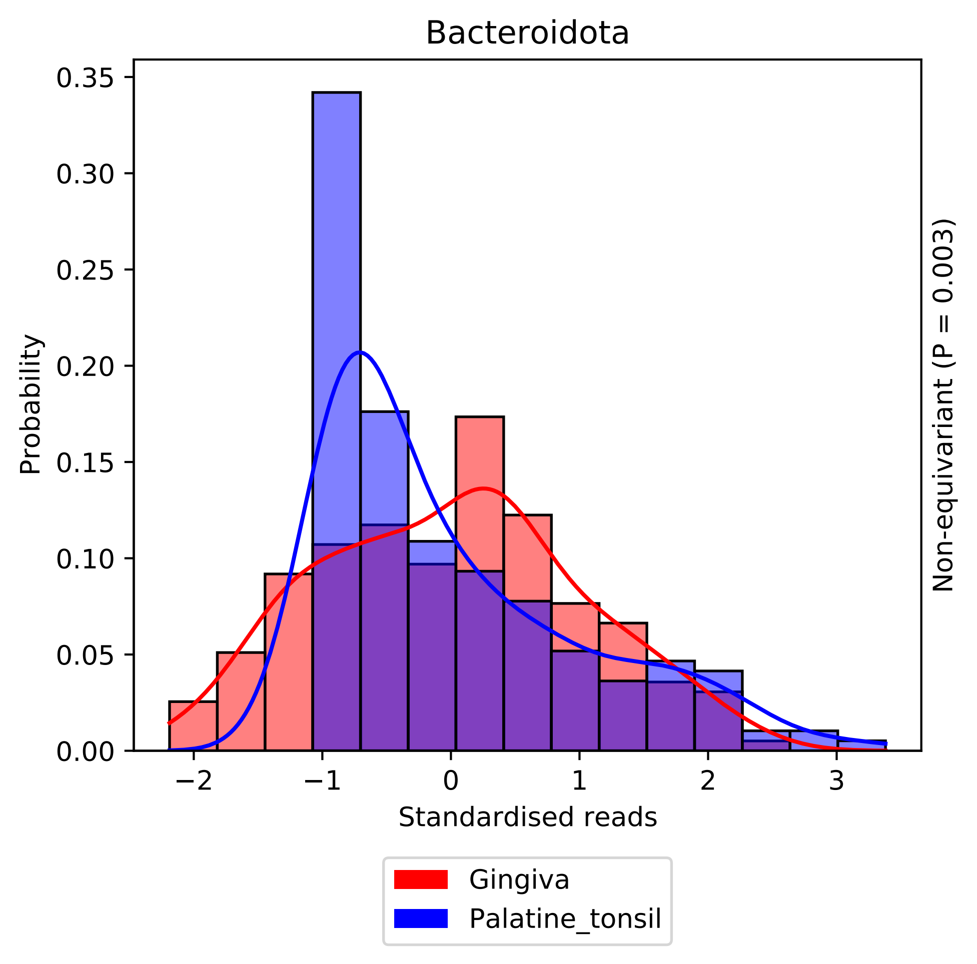

Bacteroidota
Probability
0.35
0.30
0.25
0.20
0.15
0.10
0.05
0.00
−2
−1
0
1
2
3
Standardised reads
Non-equivariant (P = 0.003)
Gingiva
Palatine_tonsil

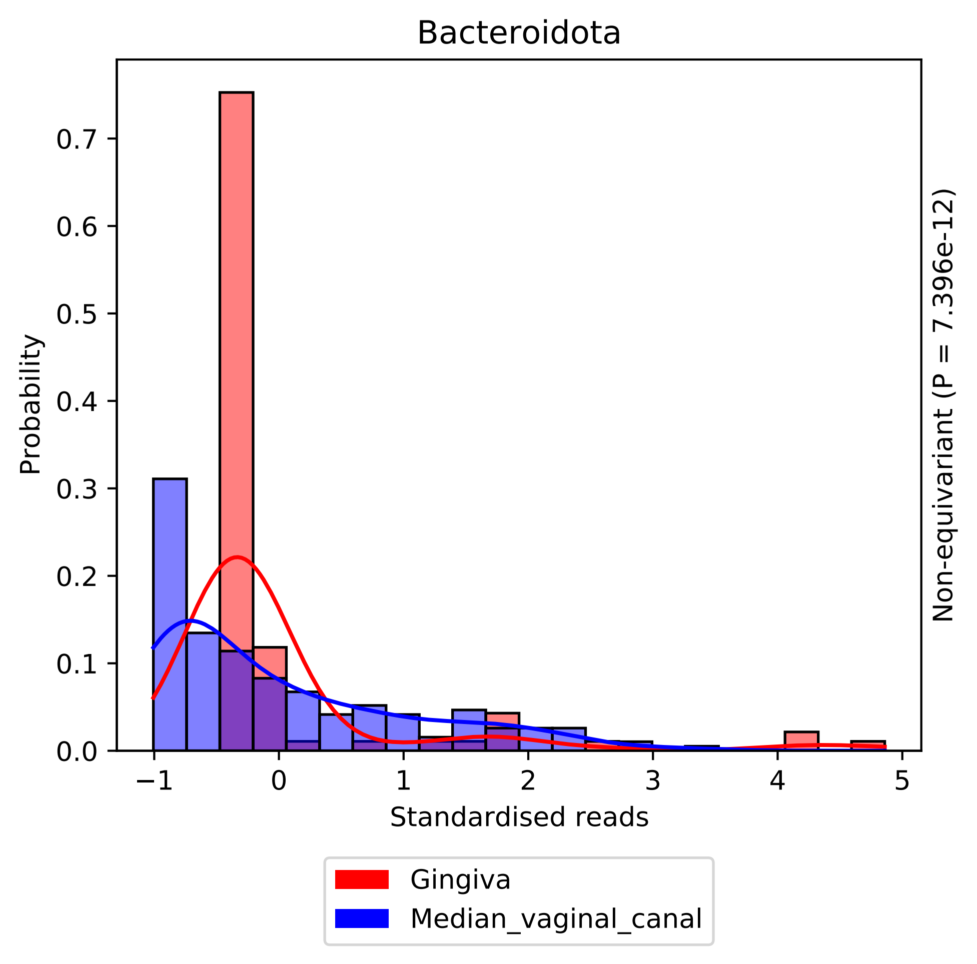
Bacteroidota
Probability
Standardised reads
Non-equivariant (P = 7.396e-12)
0.0
0.1
0.2
0.3
0.4
0.5
0.6
0.7
−1
0
1
2
3
4
5
Gingiva
Median_vaginal_canal

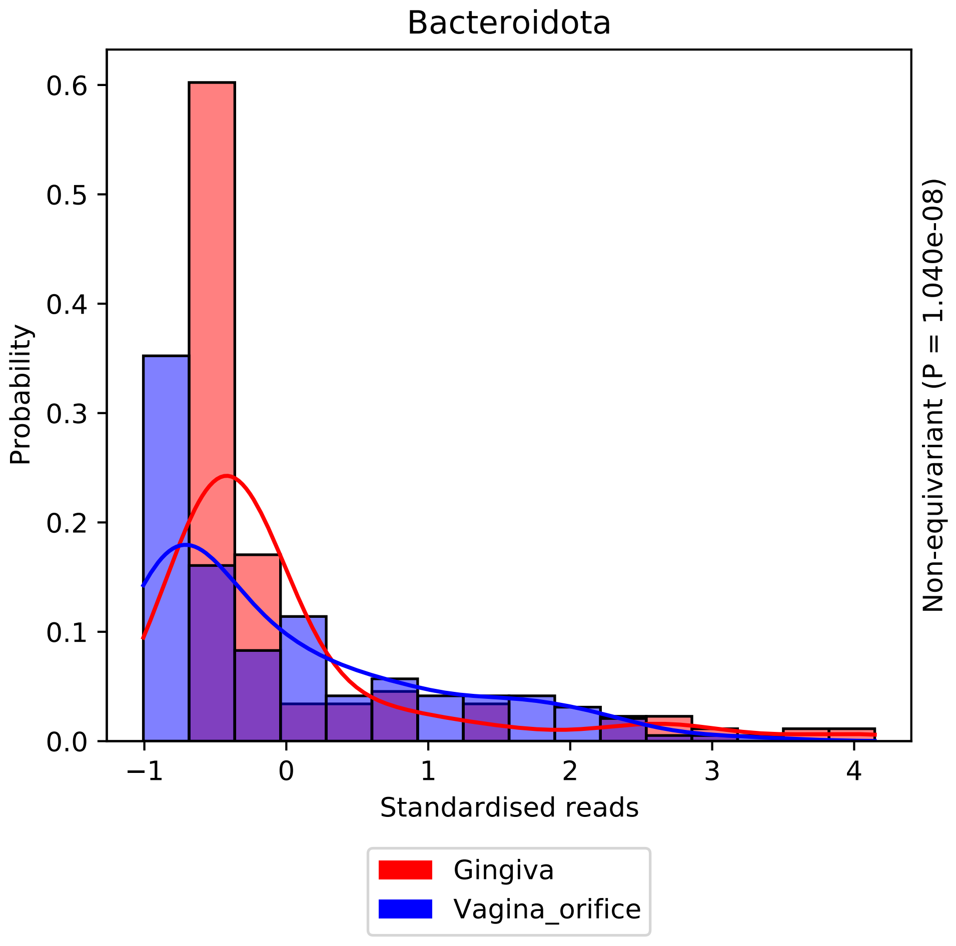

Bacteroidota
Probability
0.0
0.1
0.2
0.3
0.4
0.5
0.6
−1
0
1
2
3
4
Standardised reads
Non-equivariant (P = 1.040e-08)
Gingiva
Vagina_orifice

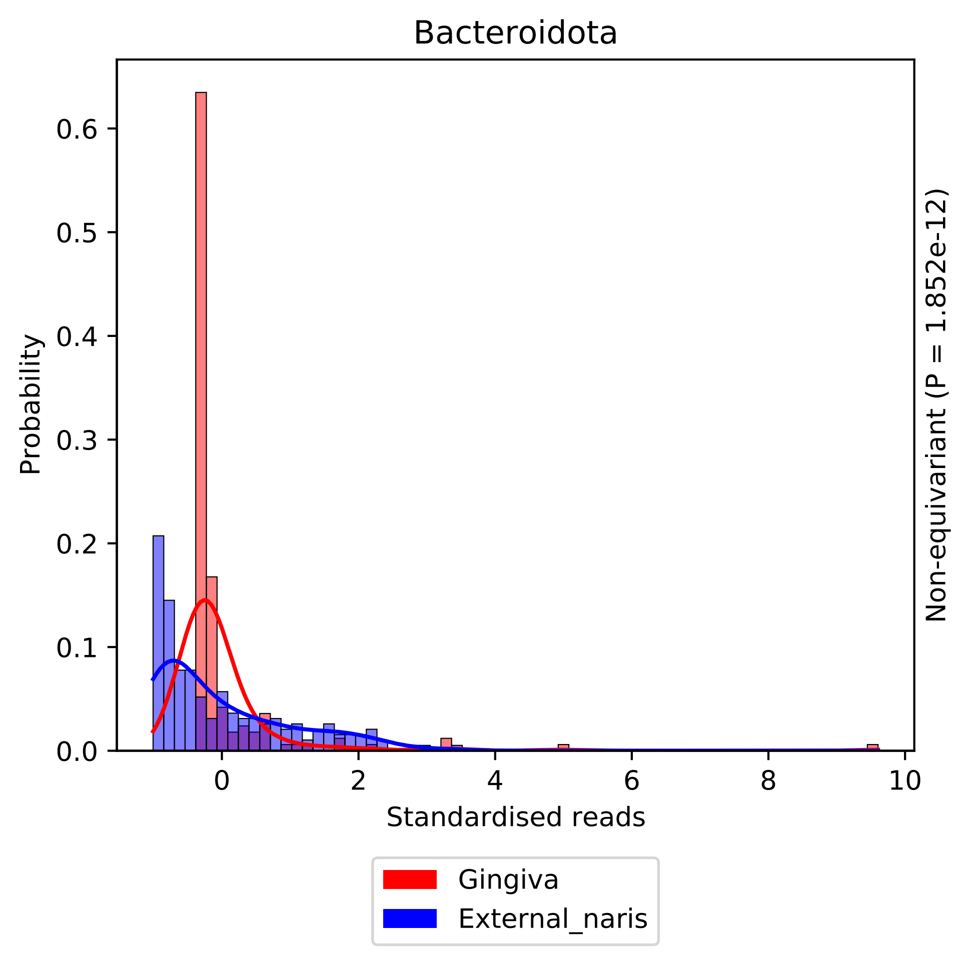

Bacteroidota
Probability
0.0
0.1
0.2
0.3
0.4
0.5
0.6
0
2
4
6
8
10
Standardised reads
Non-equivariant (P = 1.852e-12)
Gingiva
External_naris

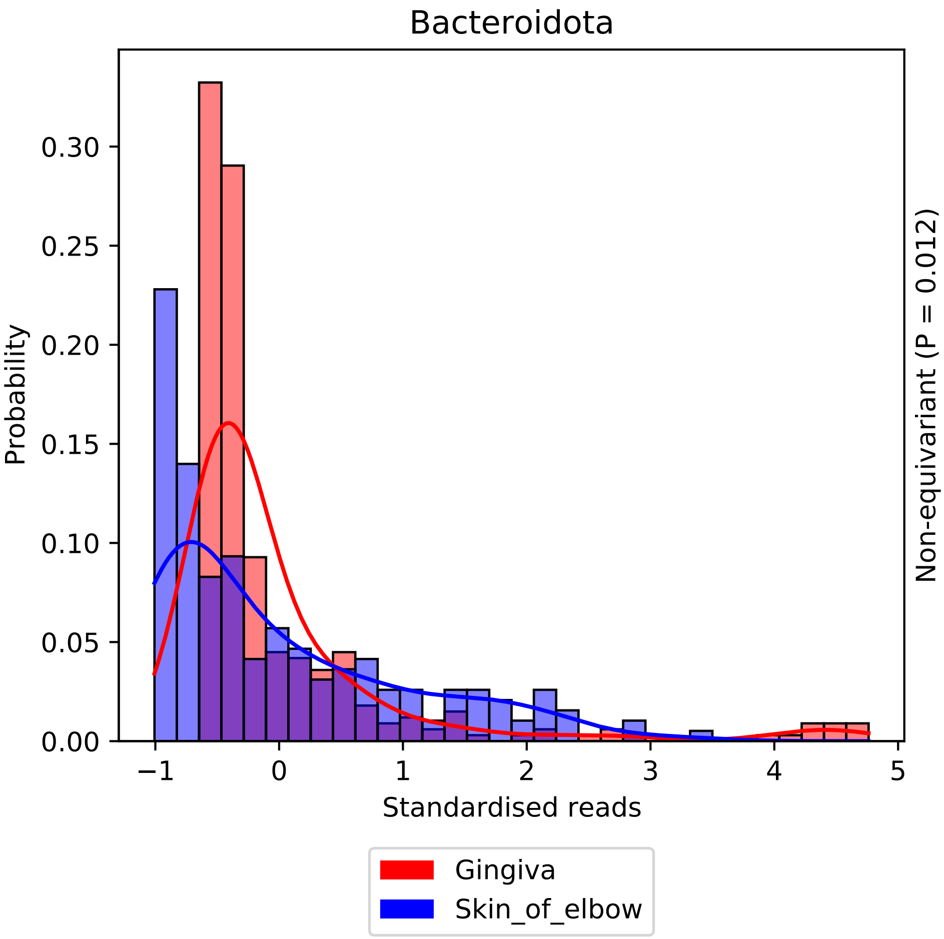

Bacteroidota
Probability
0.30
0.25
0.20
0.15
0.10
0.05
0.00
−1
0
1
2
3
4
5
Standardised reads
Non-equivariant (P = 0.012)
Gingiva
Skin_of_elbow

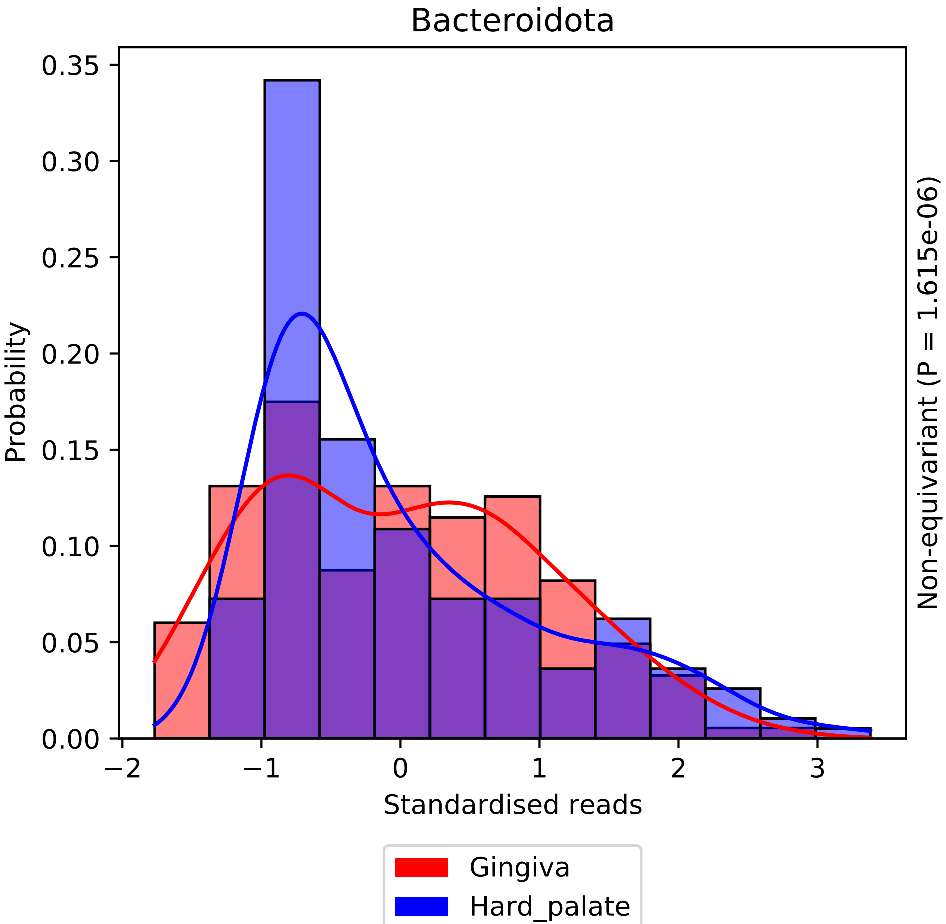
Bacteroidota
Probability
0.35
0.30
0.25
0.20
0.15
0.10
0.05
0.00
−2
−1
0
1
2
3
Standardised reads
Non-equivariant (P = 1.615e-06)
Gingiva
Hard_palate

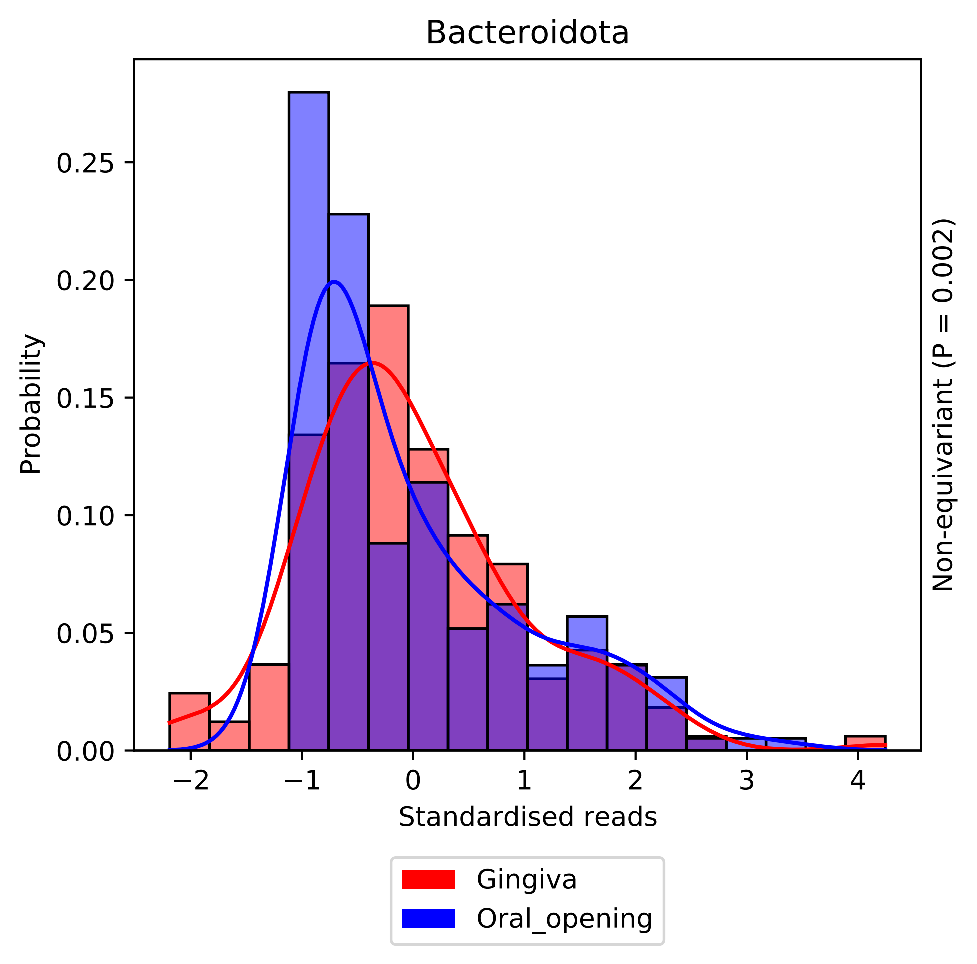

Bacteroidota
Probability
0.00
0.05
0.10
0.15
0.20
0.25
−2
−1
0
1
2
3
4
Standardised reads
Non-equivariant (P = 0.002)
Gingiva
Oral_opening

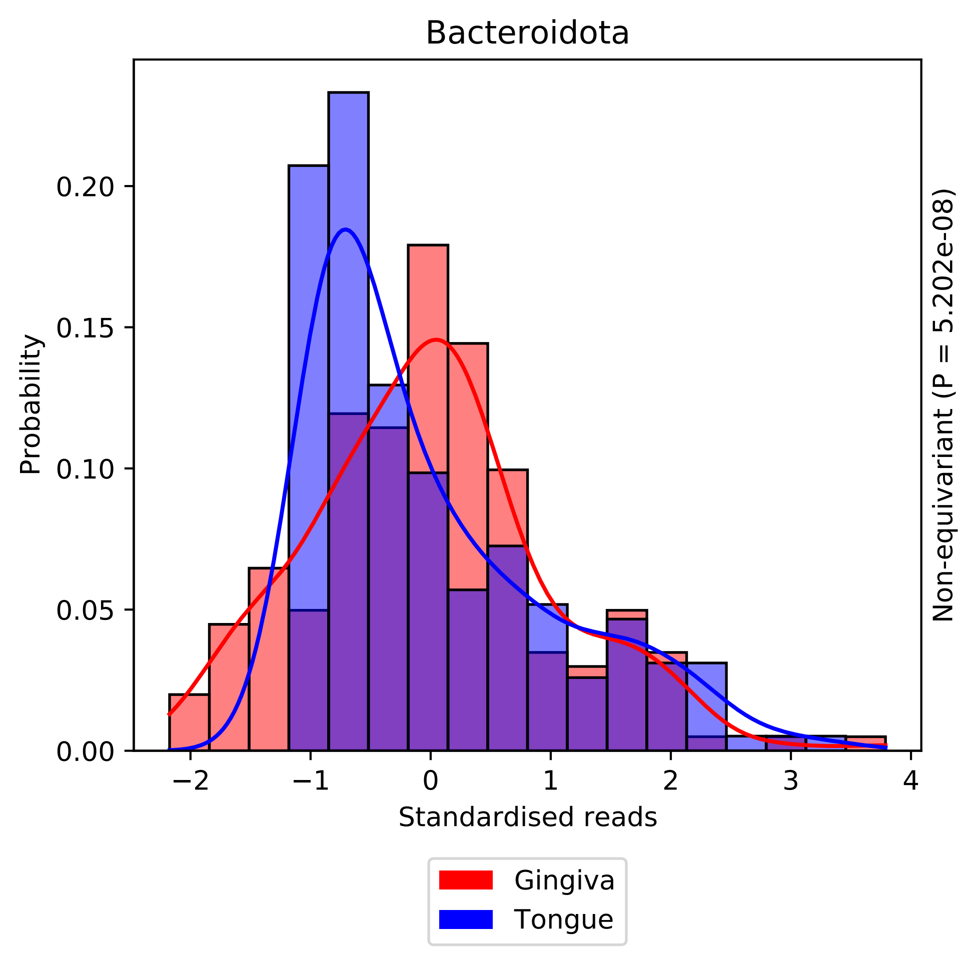

Bacteroidota
Probability
0.00
0.05
0.10
0.15
0.20
−2
−1
0
1
2
3
4
Standardised reads
Non-equivariant (P = 5.202e-08)
Gingiva
Tongue

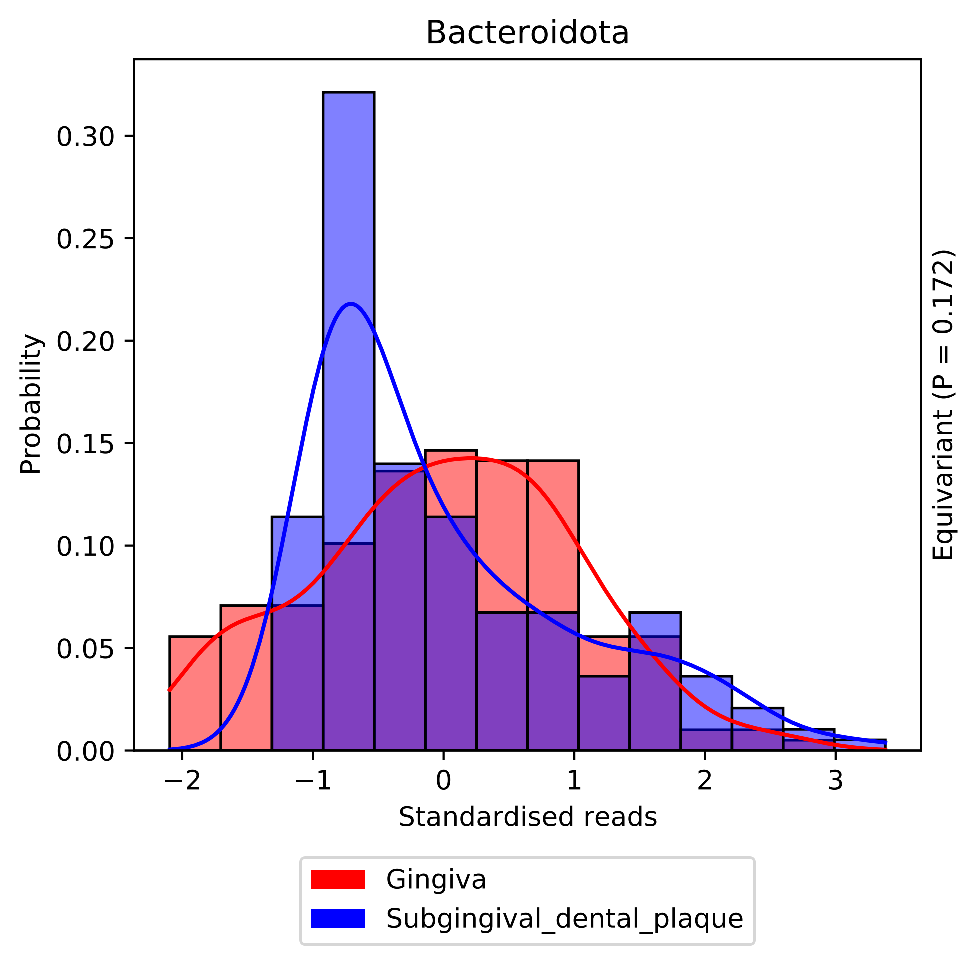

Bacteroidota
Probability
0.00
0.05
0.10
0.15
0.20
0.25
0.30
−2
−1
0
1
2
3
Standardised reads
Equivariant (P = 0.172)
Gingiva
Subgingival_dental_plaque

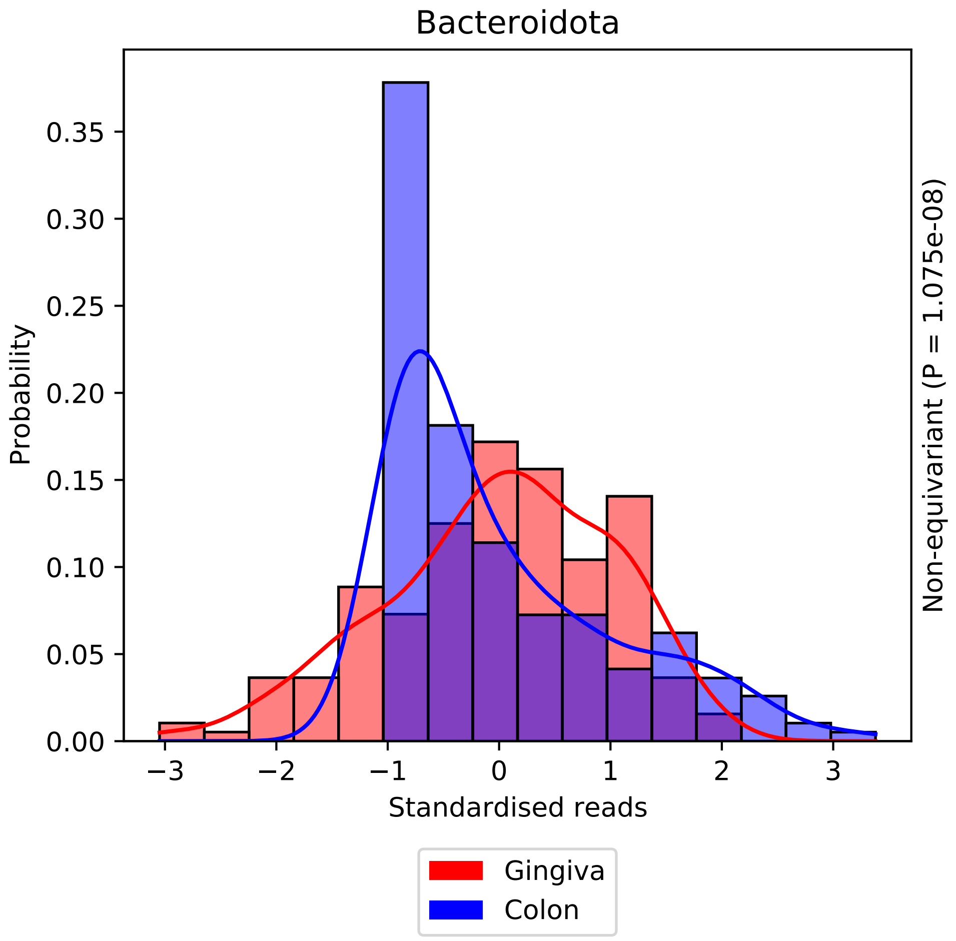

Bacteroidota
0.00
0.05
0.10
0.15
0.20
0.25
0.30
0.35
Probability
−3
−2
−1
0
1
2
3
Standardised reads
Non-equivariant (P = 1.075e-08)
Gingiva
Colon

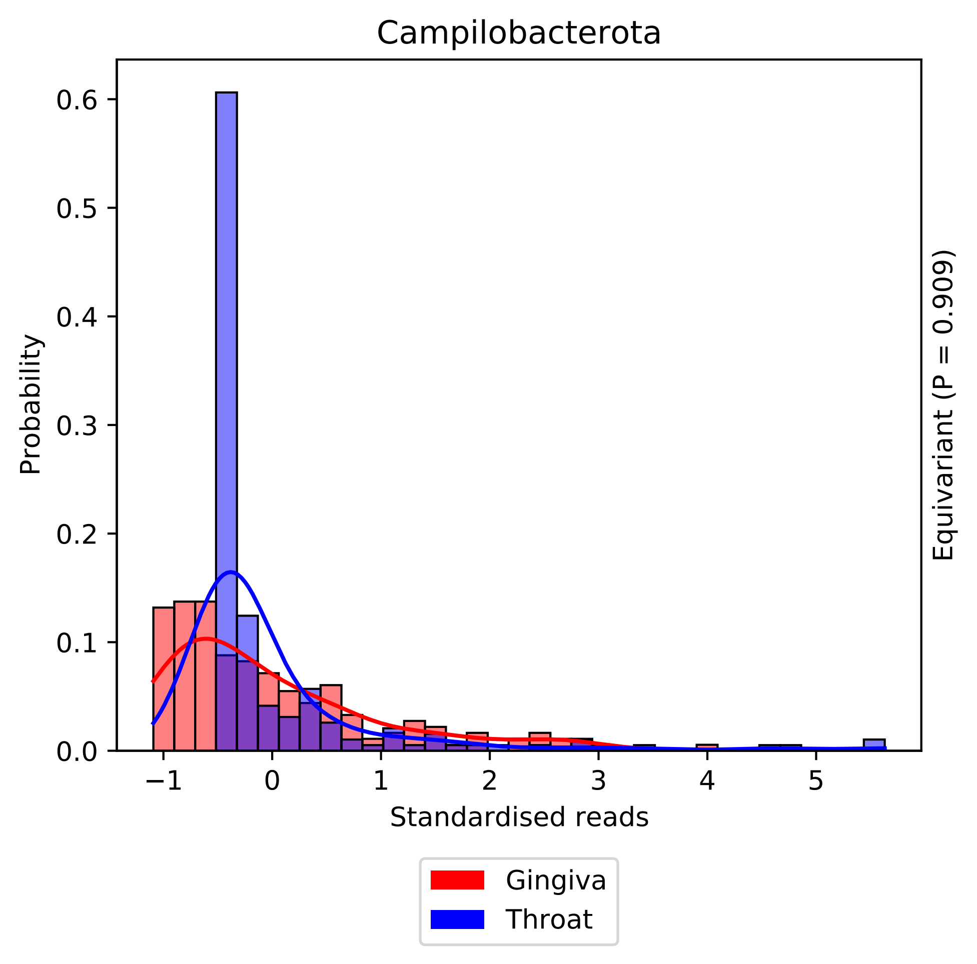
Campilobacterota
Probability
0.0
0.1
0.2
0.3
0.4
0.5
0.6
−1
0
1
2
3
4
5
Standardised reads
Equivariant (P = 0.909)
Gingiva
Throat

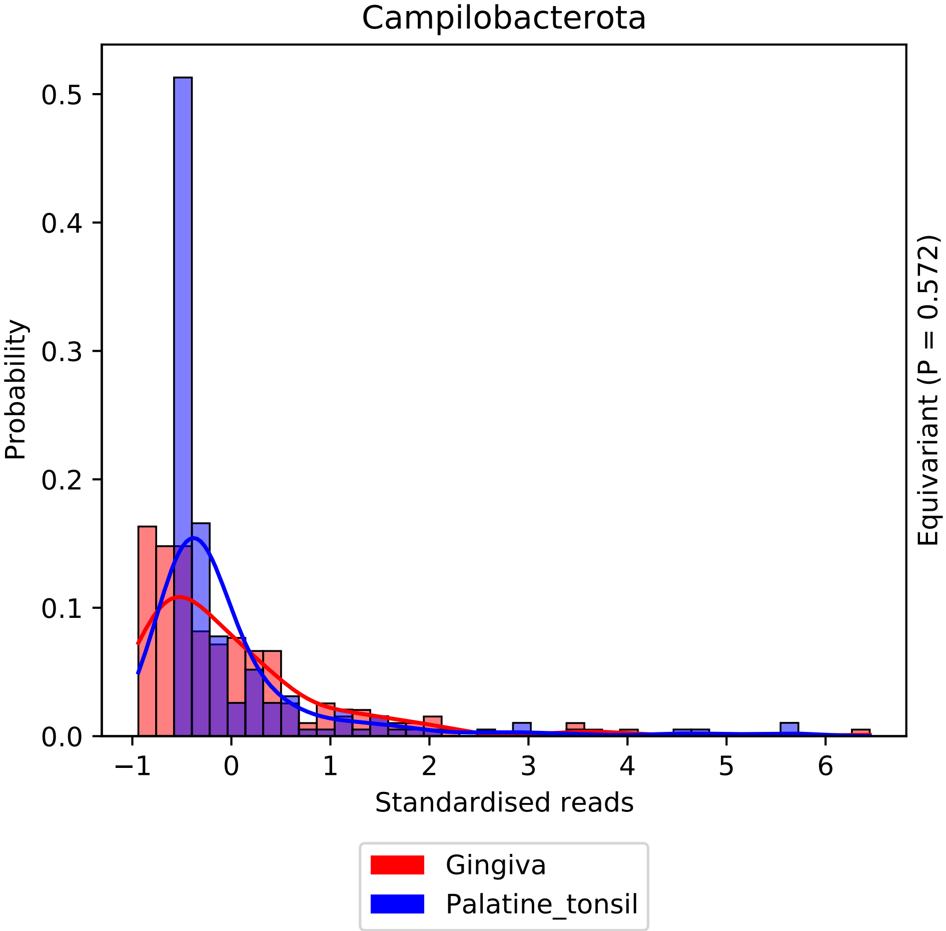
Campilobacterota
Probability
Standardised reads
Equivariant (P = 0.572)
Gingiva
Palatine_tonsil
0.0
0.1
0.2
0.3
0.4
0.5
−1
0
1
2
3
4
5
6

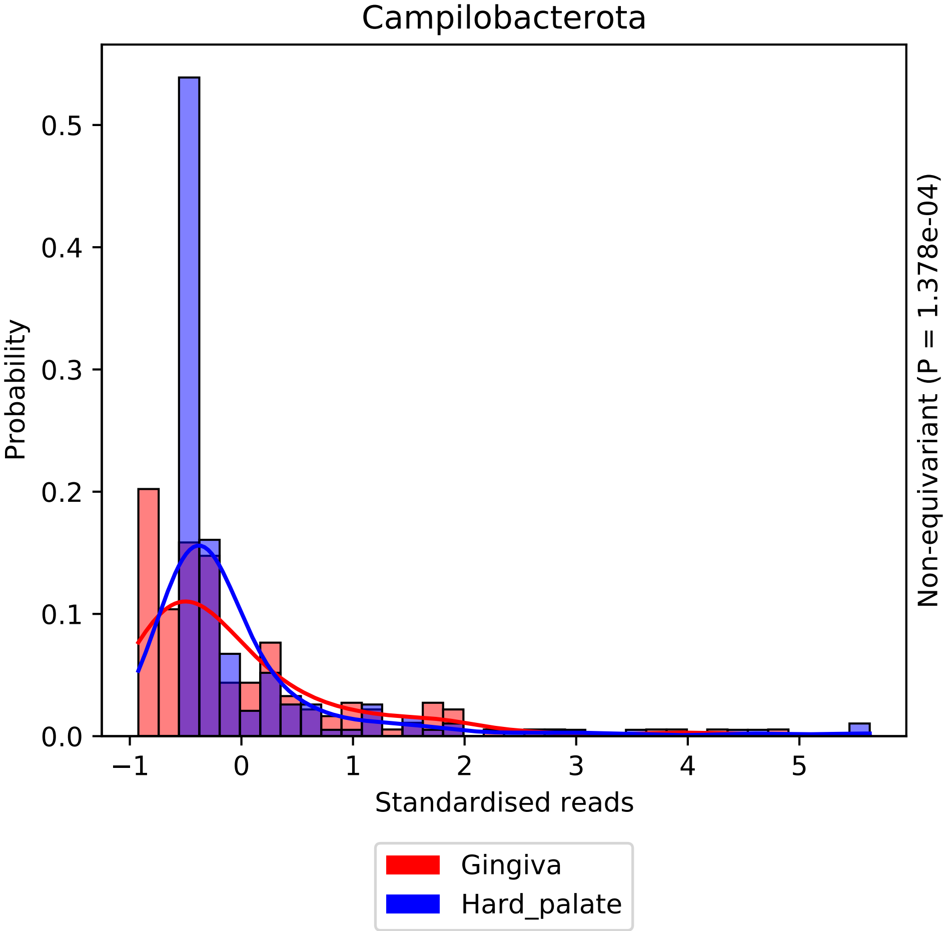

Campilobacterota
Probability
0.0
0.1
0.2
0.3
0.4
0.5
−1
0
1
2
3
4
5
Standardised reads
Non-equivariant (P = 1.378e-04)
Gingiva
Hard_palate

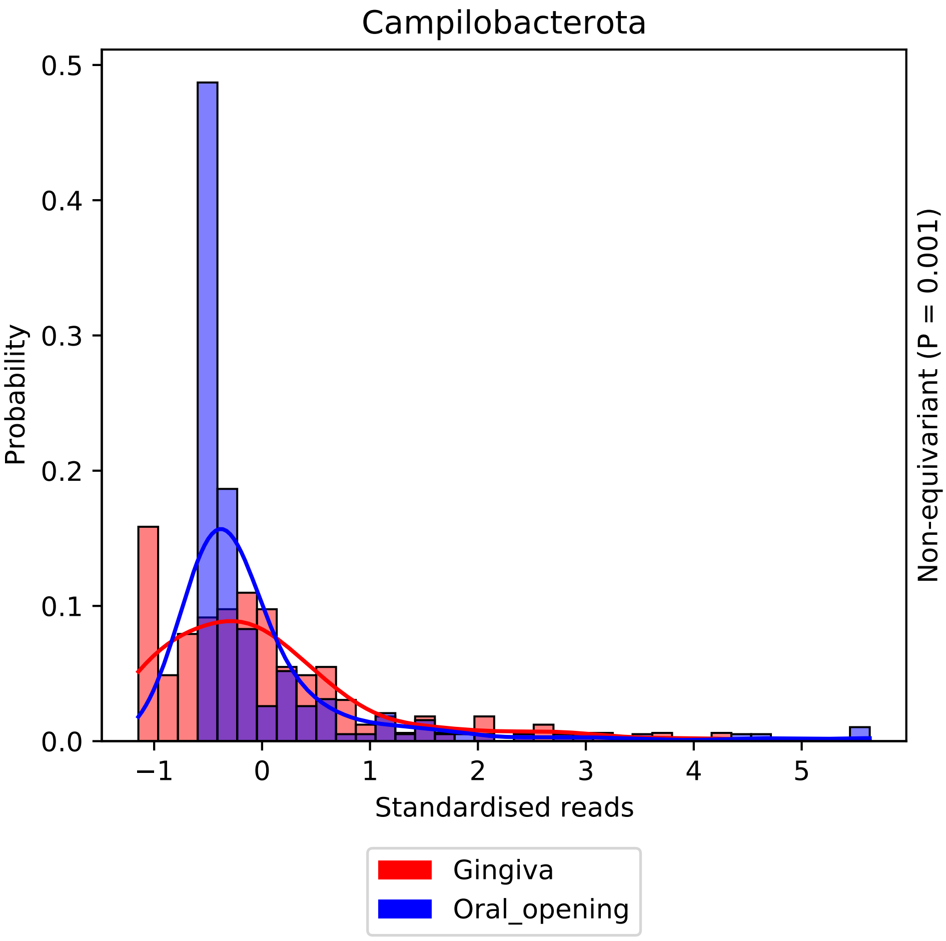

Campilobacterota
Probability
0.5
0.4
0.3
0.2
0.1
0.0
−1
0
1
2
3
4
5
Standardised reads
Non-equivariant (P = 0.001)
Gingiva
Oral_opening

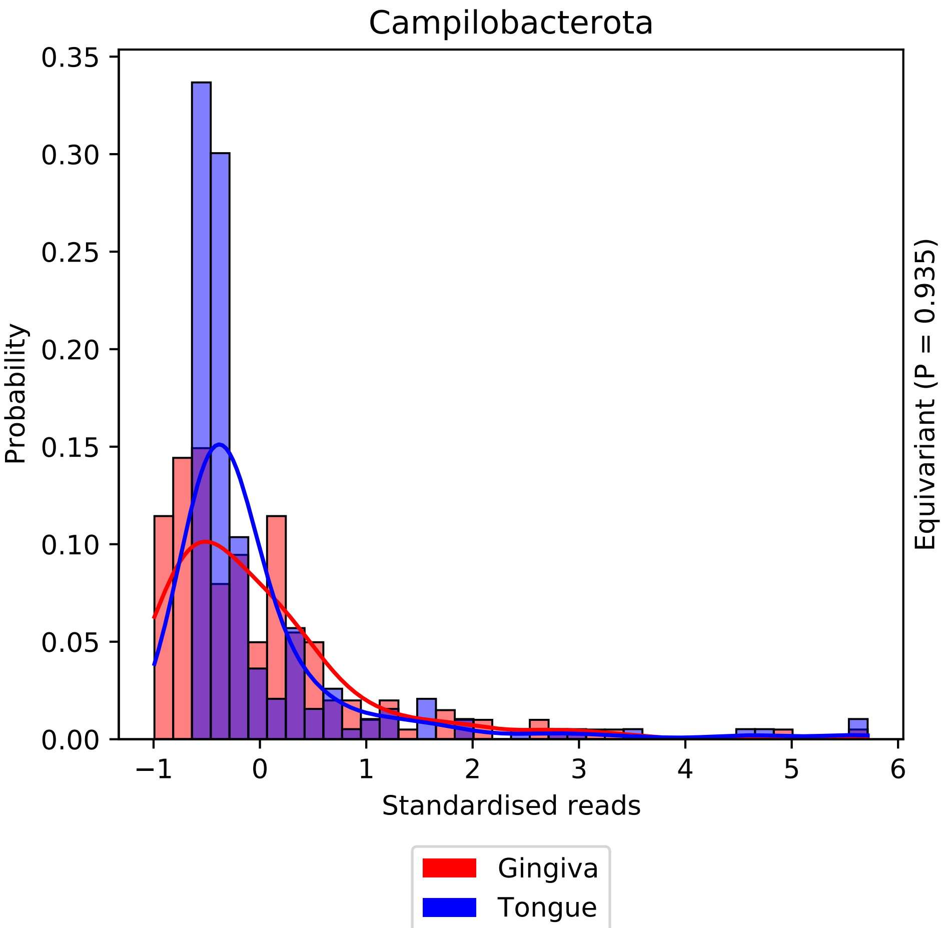
Campilobacterota
Probability
Standardised reads
Equivariant (P = 0.935)
Gingiva
Tongue
0.00
0.05
0.10
0.15
0.20
0.25
0.30
0.35
−1
0
1
2
3
4
5
6

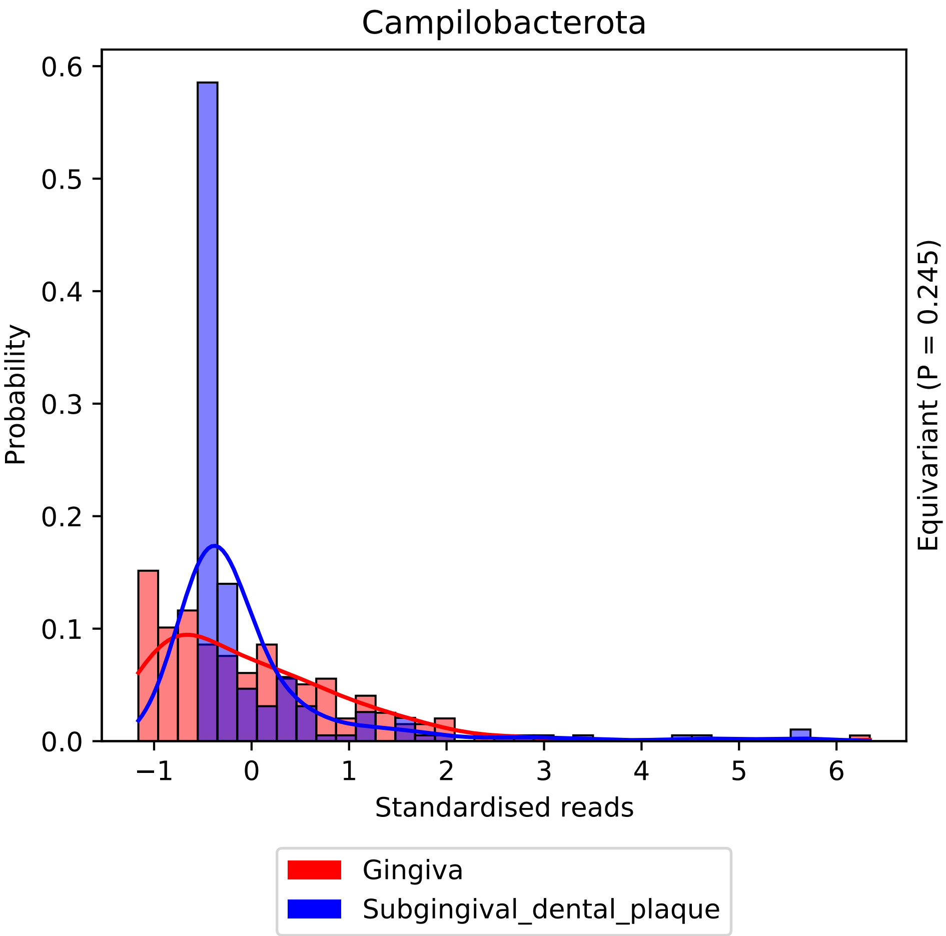
Campilobacterota
Probability
Standardised reads
Equivariant (P = 0.245)
Gingiva
Subgingival_dental_plaque
0.0
0.1
0.2
0.3
0.4
0.5
0.6
−1
0
1
2
3
4
5
6

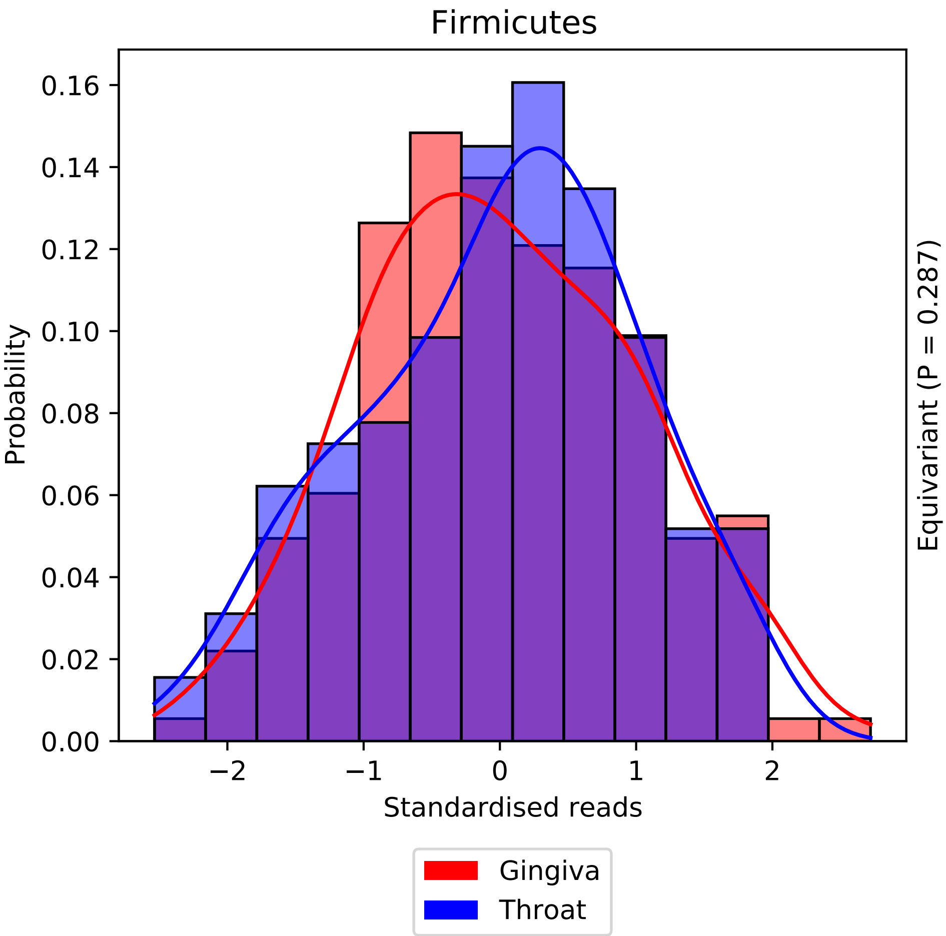
Firmicutes
Probability
0.00
0.02
0.04
0.06
0.08
0.10
0.12
0.14
0.16
−2
−1
0
1
2
Standardised reads
Equivariant (P = 0.287)
Gingiva
Throat

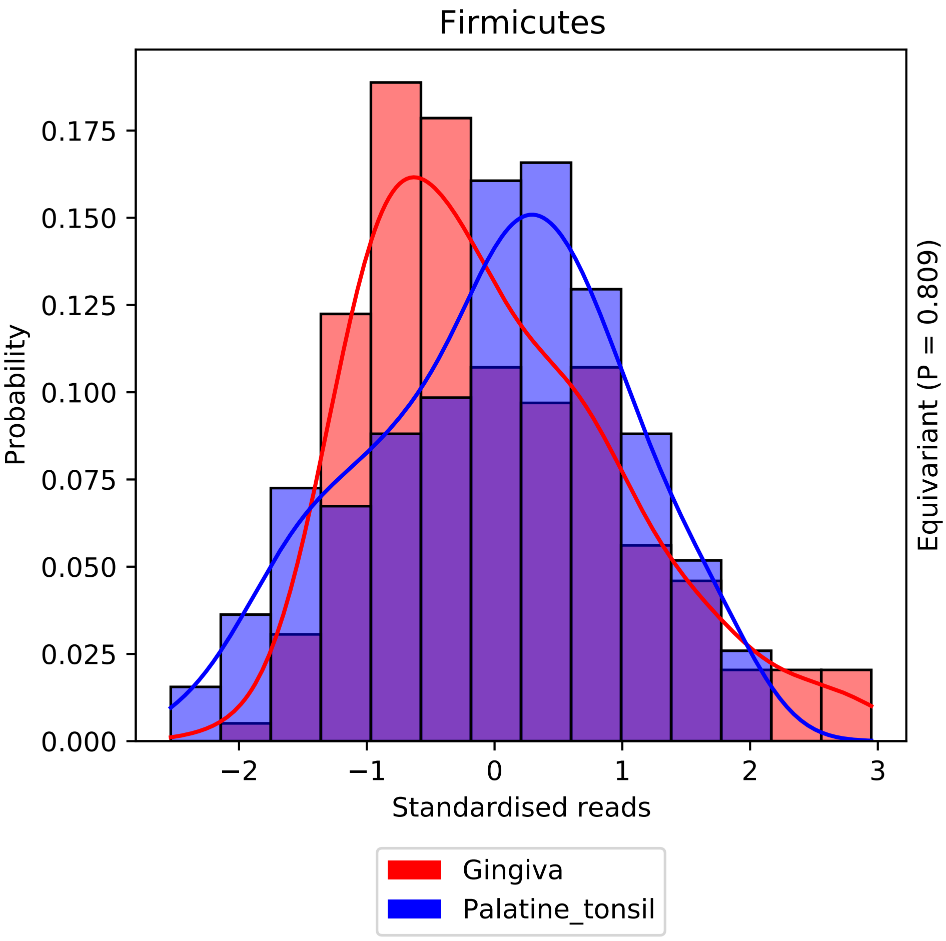

Firmicutes
Probability
0.000
0.025
0.050
0.075
0.100
0.125
0.150
0.175
−2
−1
0
1
2
3
Standardised reads
Equivariant (P = 0.809)
Gingiva
Palatine_tonsil

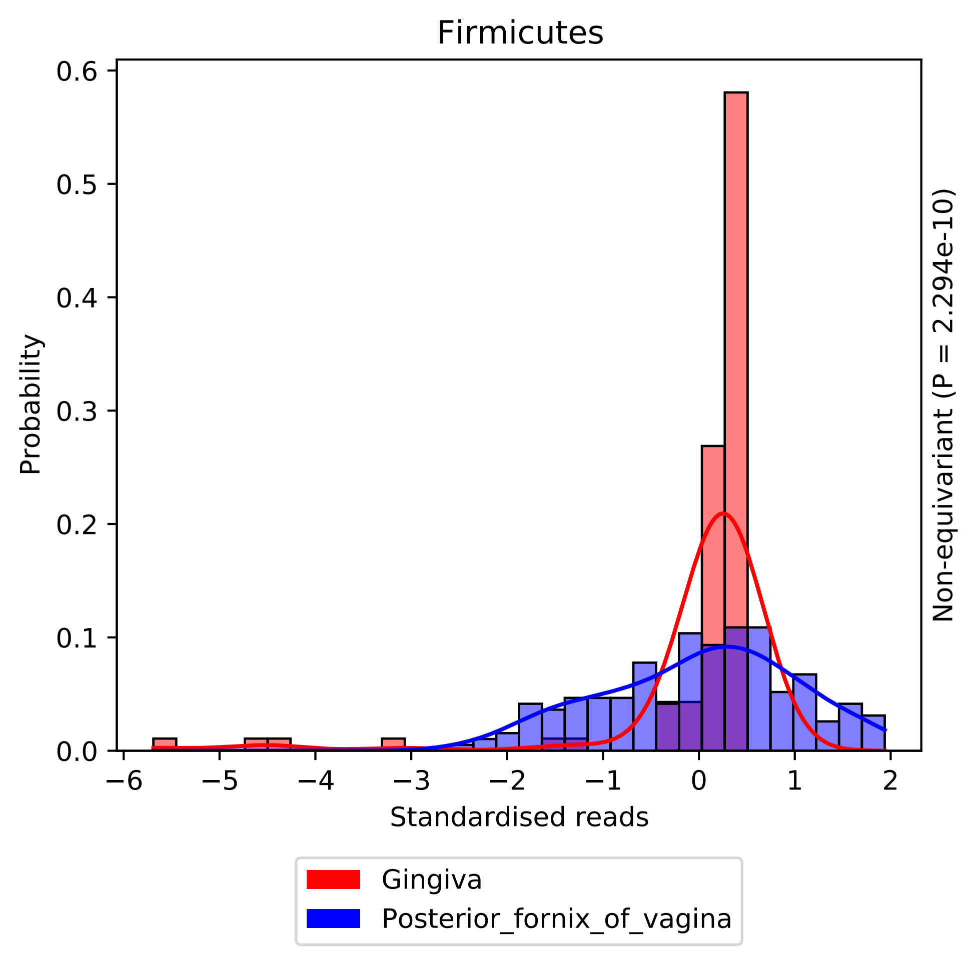

Firmicutes
Probability
0.0
0.1
0.2
0.3
0.4
0.5
0.6
−6
−5
−4
−3
−2
−1
0
1
2
Standardised reads
Non-equivariant (P = 2.294e-10)
Gingiva
Posterior_fornix_of_vagina

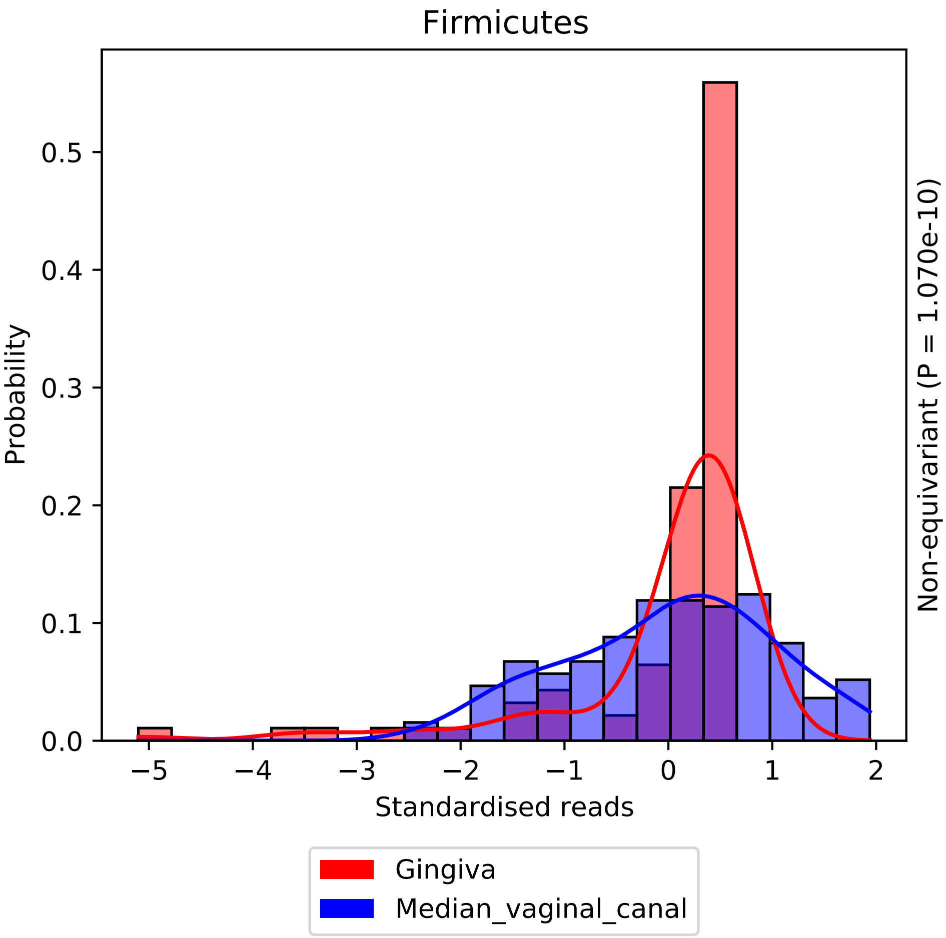

Firmicutes
Probability
0.0
0.1
0.2
0.3
0.4
0.5
−5
−4
−3
−2
−1
0
1
2
Standardised reads
Non-equivariant (P = 1.070e-10)
Gingiva
Median_vaginal_canal

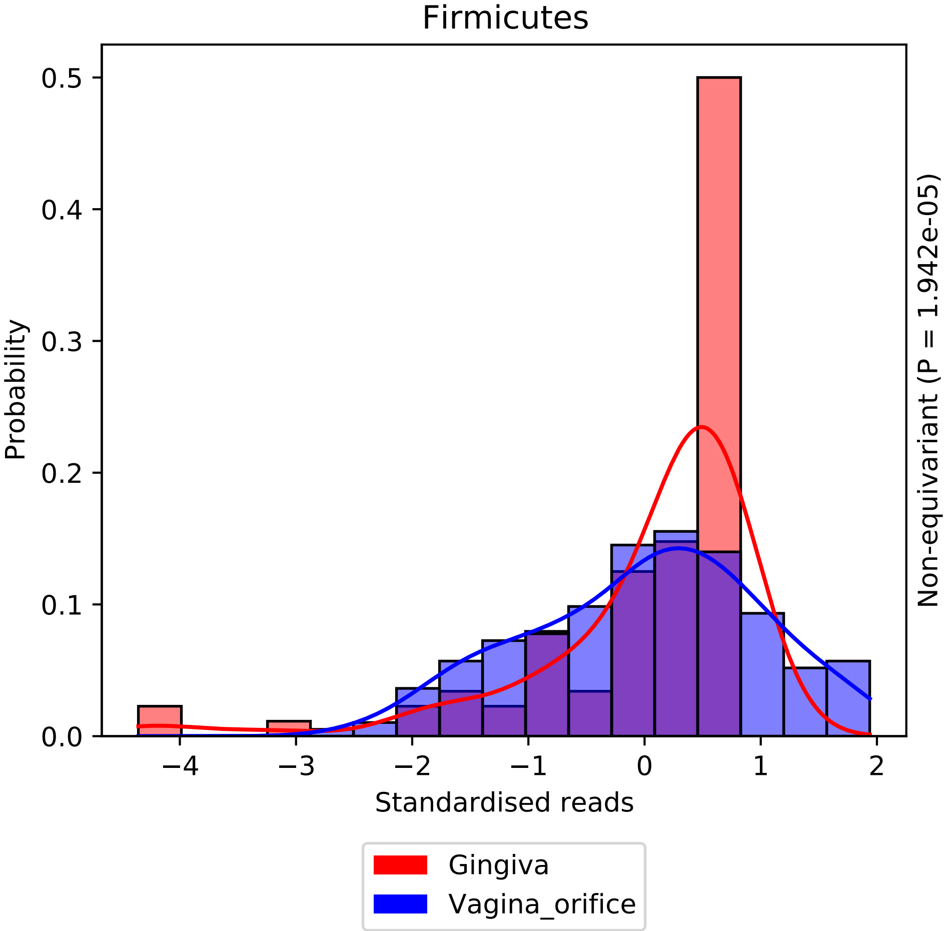

Firmicutes
Probability
0.0
0.1
0.2
0.3
0.4
0.5
−4
−3
−2
−1
0
1
2
Standardised reads
Non-equivariant (P = 1.942e-05)
Gingiva
Vagina_orifice

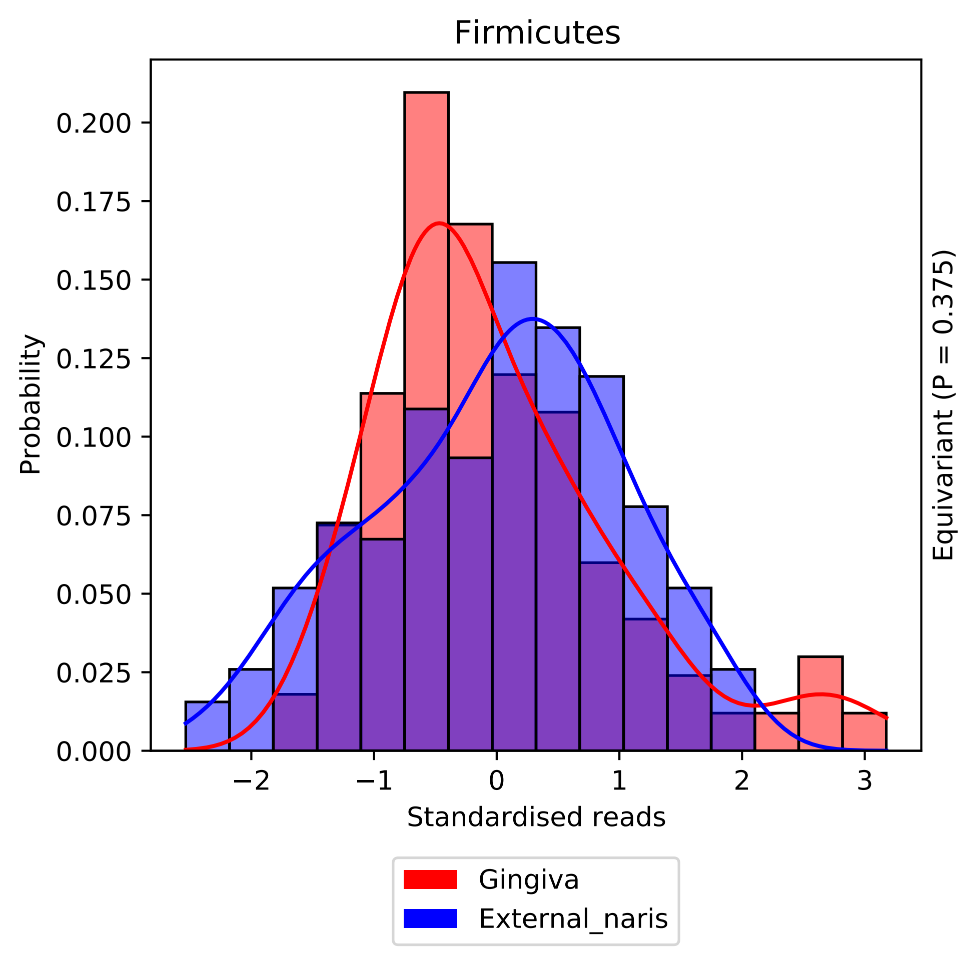
Firmicutes
Probability
0.200
0.175
0.150
0.125
0.100
0.075
0.050
0.025
0.000
−2
−1
0
1
2
3
Standardised reads
Equivariant (P = 0.375)
Gingiva
External_naris

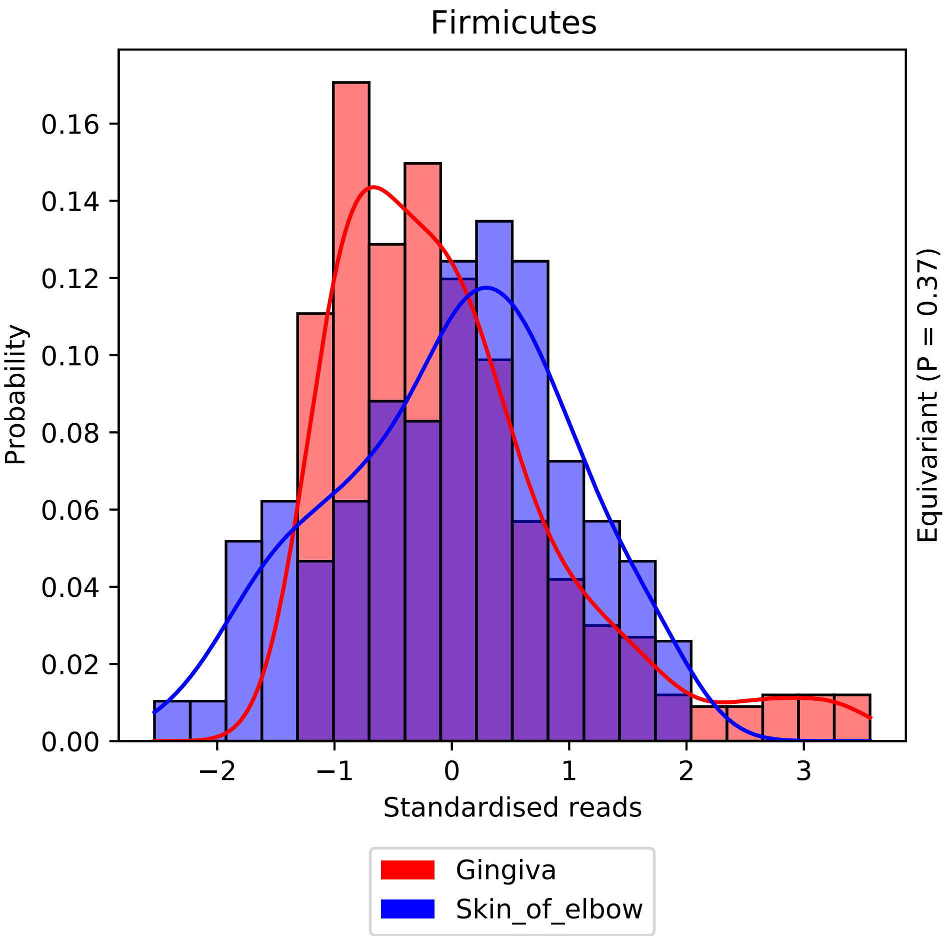
Firmicutes
Probability
0.00
0.02
0.04
0.06
0.08
0.10
0.12
0.14
0.16
−2
−1
0
1
2
3
Standardised reads
Equivariant (P = 0.37)
Gingiva
Skin_of_elbow

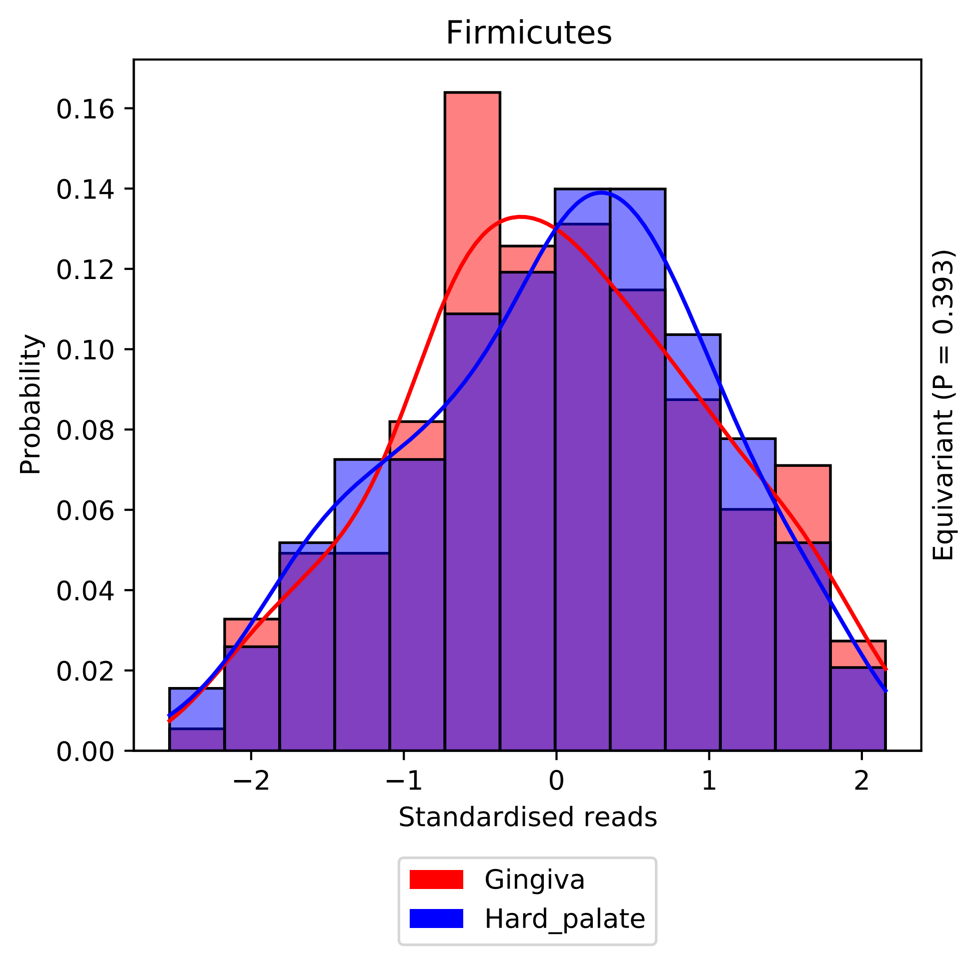

Firmicutes
Probability
Standardised reads
Equivariant (P = 0.393)
Gingiva
Hard_palate
0.00
0.02
0.04
0.06
0.08
0.10
0.12
0.14
0.16
−2
−1
0
1
2

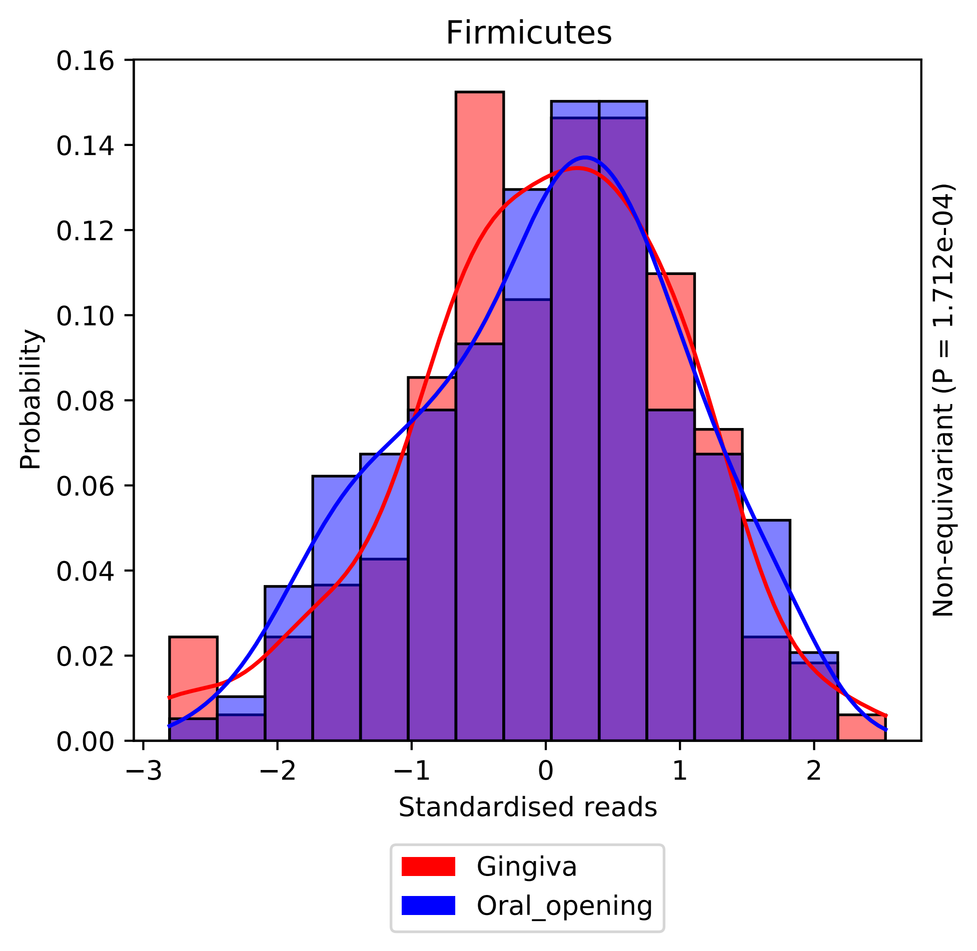
Firmicutes
0.16
0.14
0.12
0.10
0.08
0.06
0.04
0.02
0.00
Probability
−3
−2
−1
0
1
2
Standardised reads
Non-equivariant (P = 1.712e-04)
Gingiva
Oral_opening

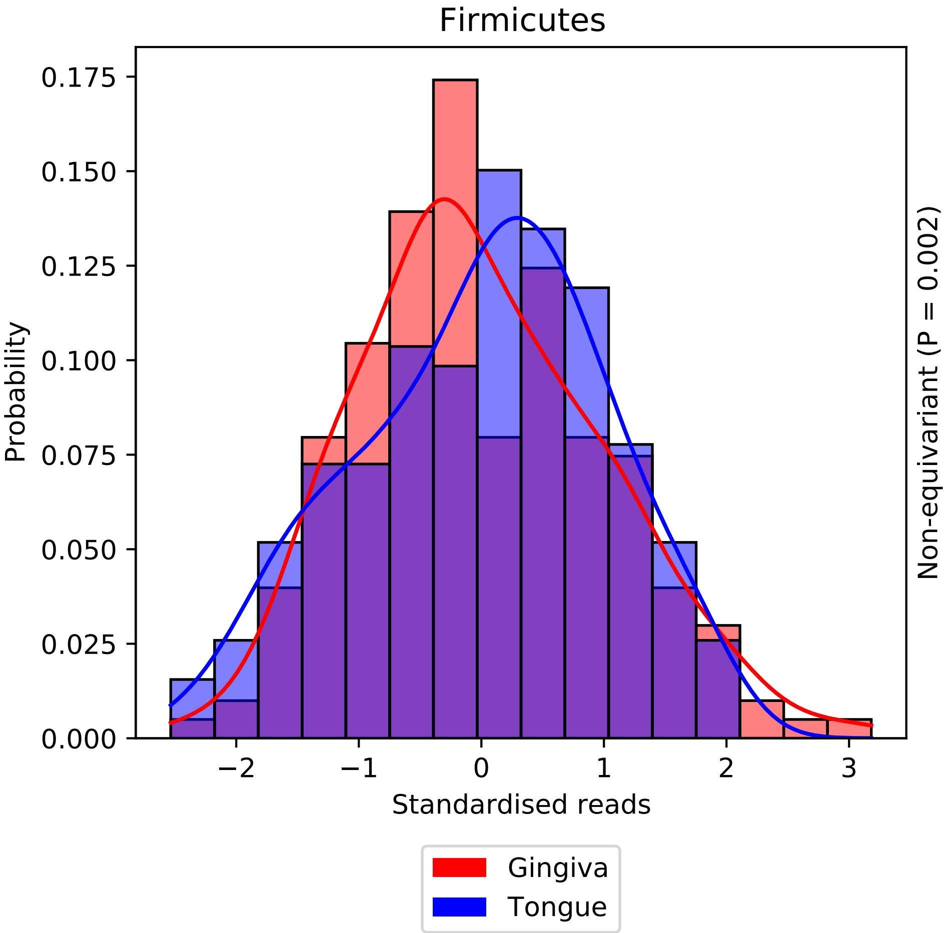

Firmicutes
Probability
0.000
0.025
0.050
0.075
0.100
0.125
0.150
0.175
−2
−1
0
1
2
3
Standardised reads
Non-equivariant (P = 0.002)
Gingiva
Tongue

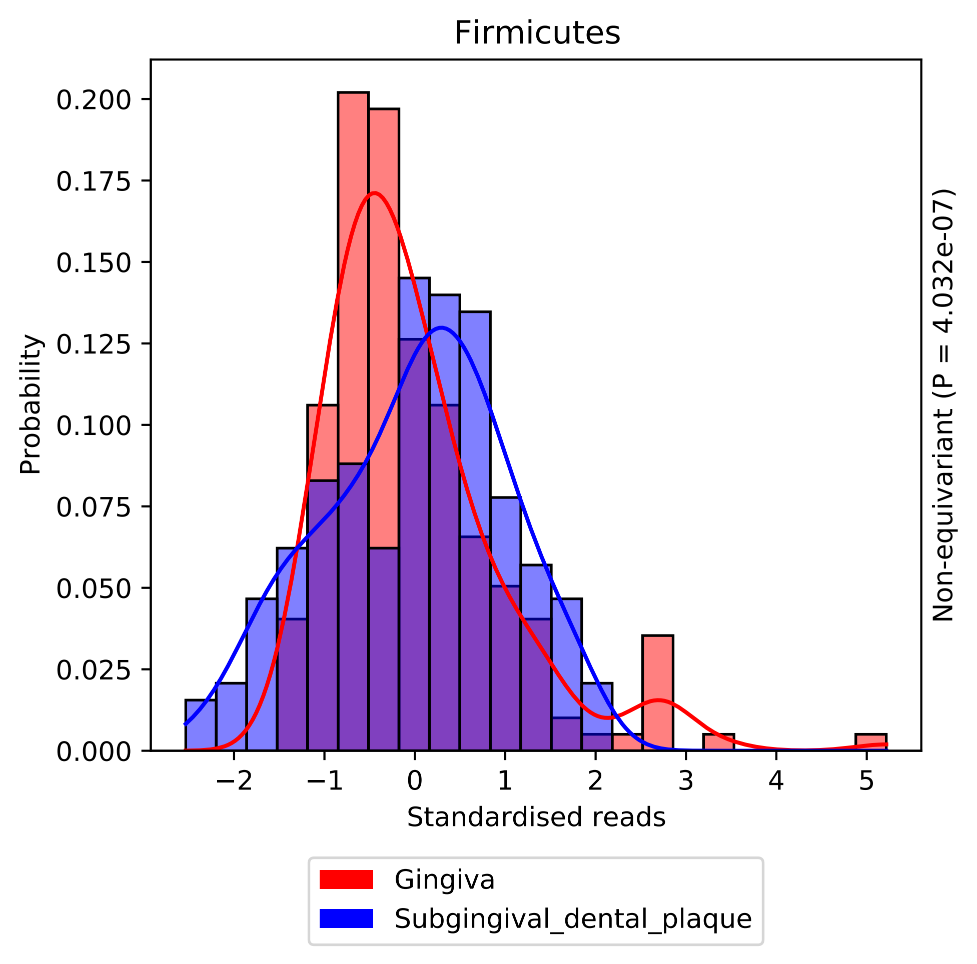

Firmicutes
Probability
Standardised reads
Non-equivariant (P = 4.032e-07)
0.000
0.025
0.050
0.075
0.100
0.125
0.150
0.175
0.200
−2
−1
0
1
2
3
4
5
Gingiva
Subgingival_dental_plaque

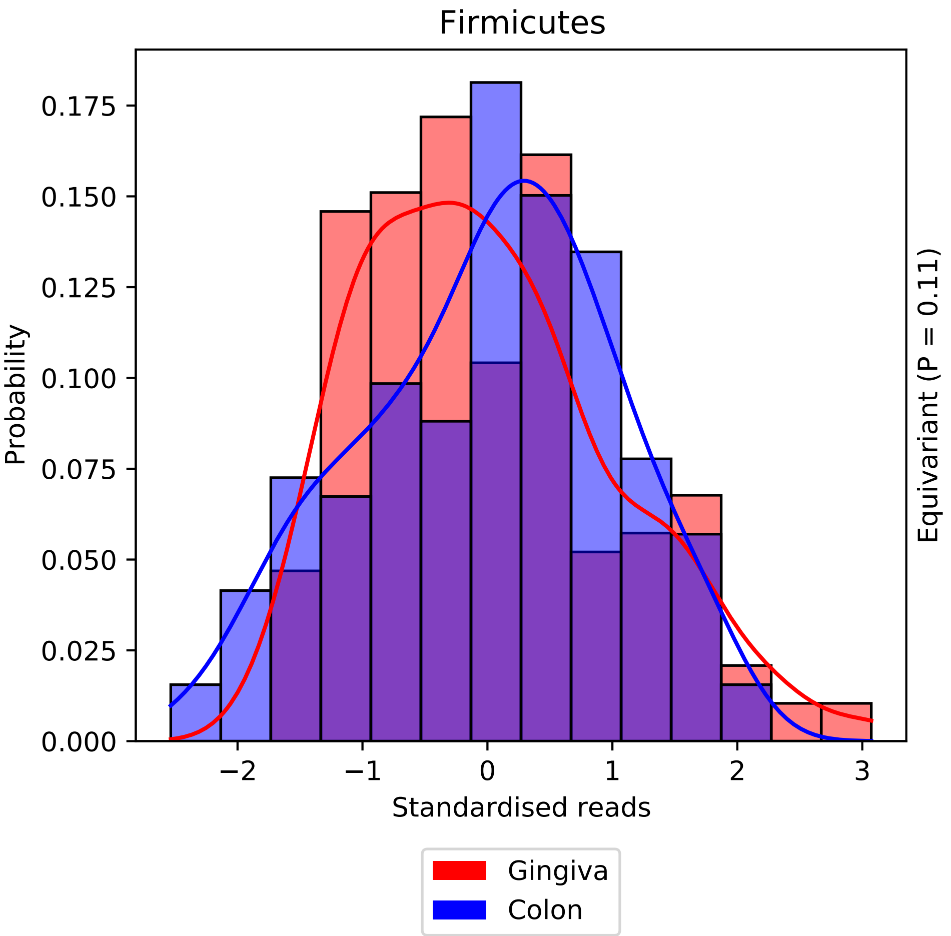

Firmicutes
Probability
0.000
0.025
0.050
0.075
0.100
0.125
0.150
0.175
−2
−1
0
1
2
3
Standardised reads
Equivariant (P = 0.11)
Gingiva
Colon

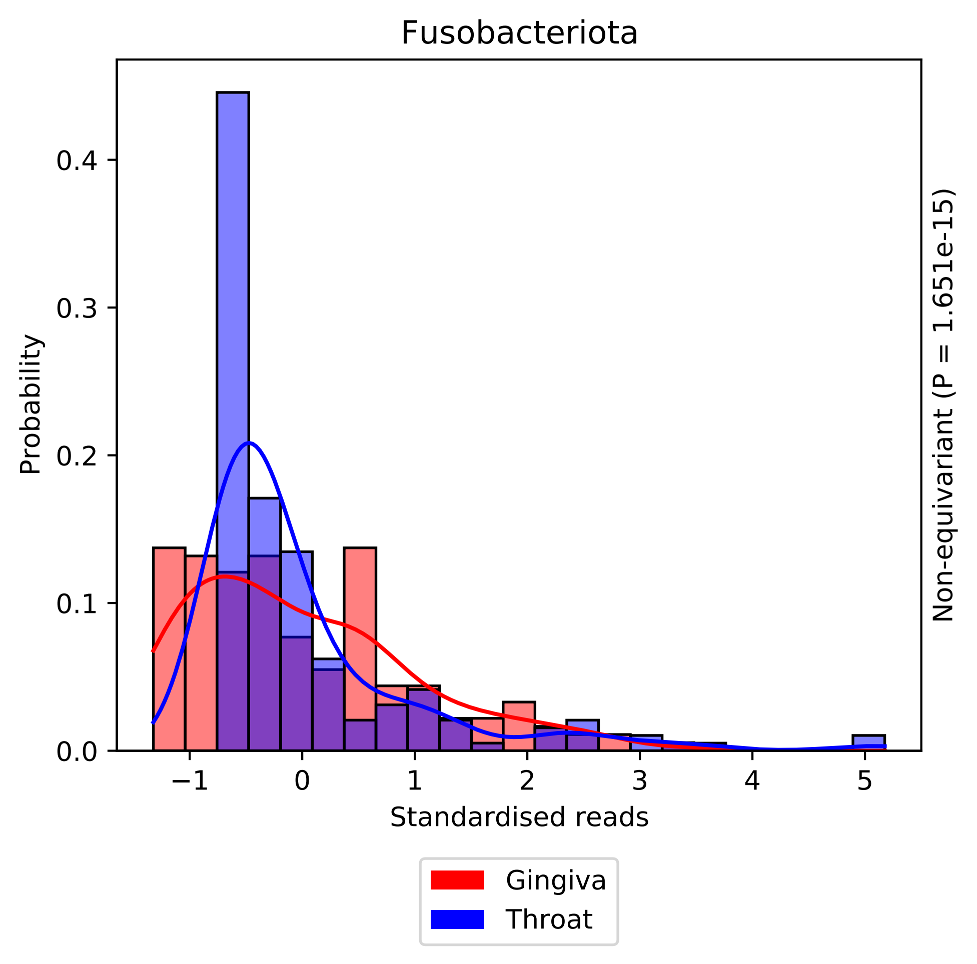

Fusobacteriota
Probability
Standardised reads
Non-equivariant (P = 1.651e-15)
0.0
0.1
0.2
0.3
0.4
−1
0
1
2
3
4
5
Gingiva
Throat

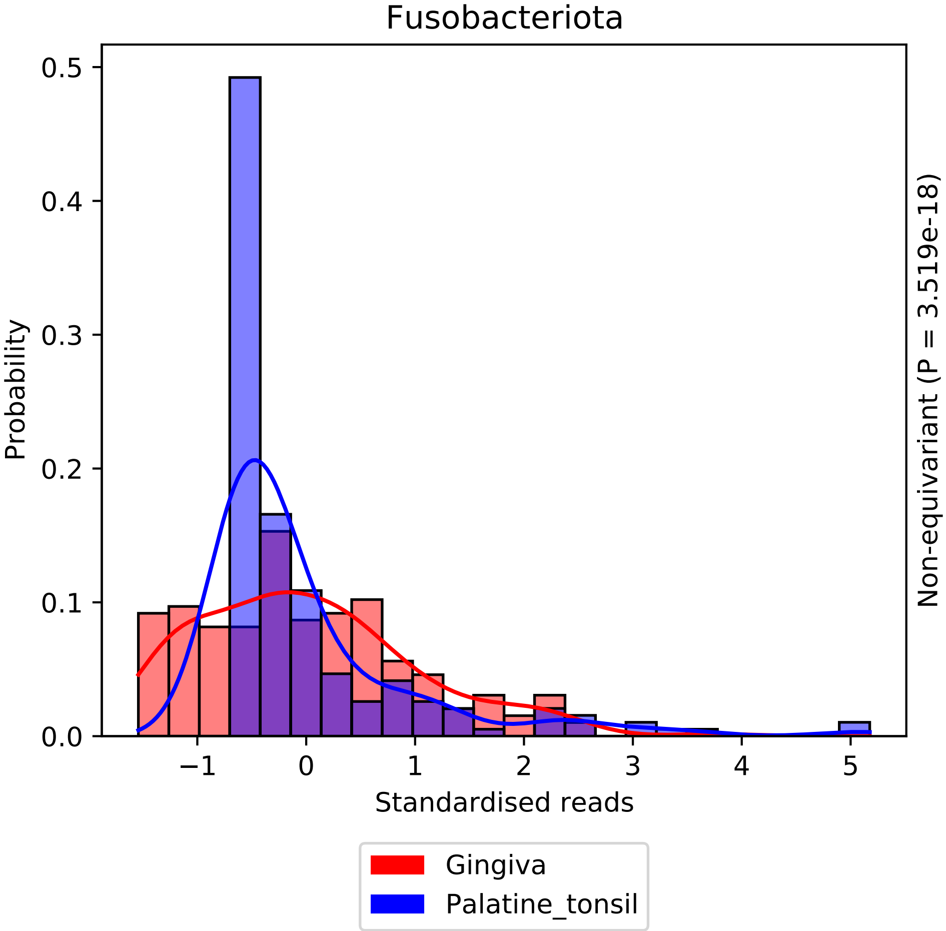

Fusobacteriota
Probability
0.0
0.1
0.2
0.3
0.4
0.5
−1
0
1
2
3
4
5
Standardised reads
Non-equivariant (P = 3.519e-18)
Gingiva
Palatine_tonsil

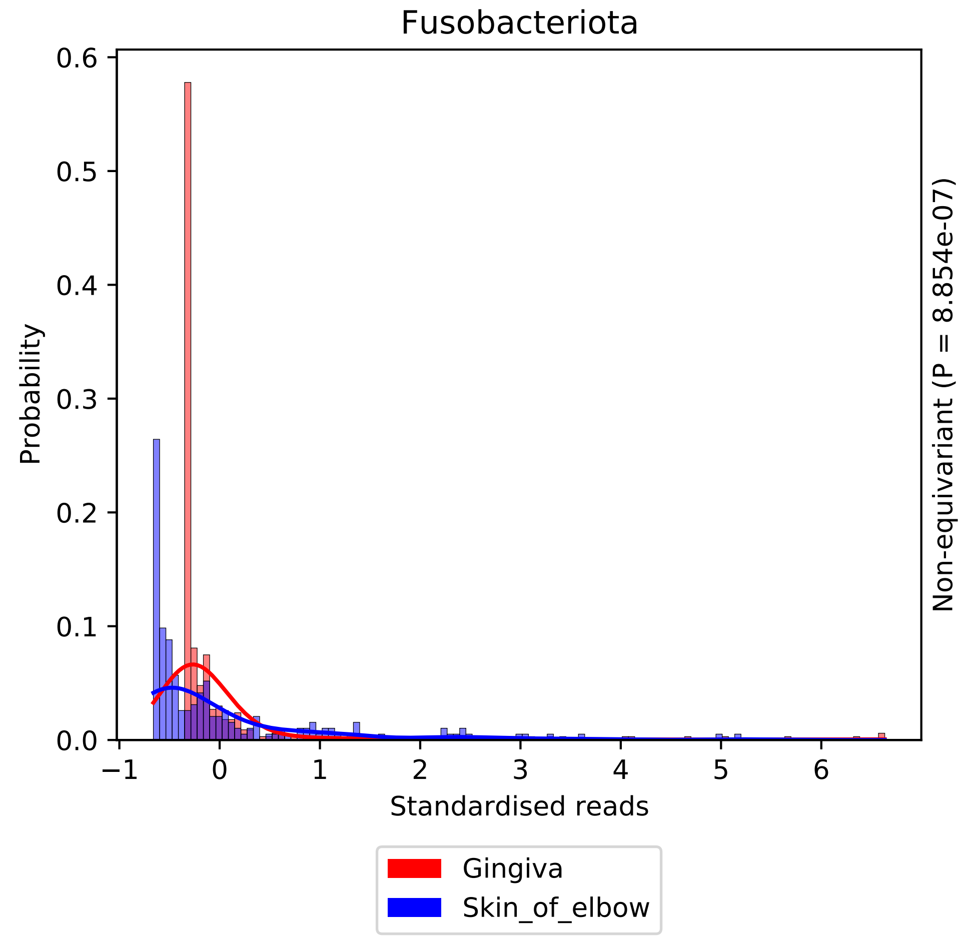

Fusobacteriota
Probability
0.6
0.5
0.4
0.3
0.2
0.1
0.0
−1
0
1
2
3
4
5
6
Standardised reads
Non-equivariant (P = 8.854e-07)
Gingiva
Skin_of_elbow

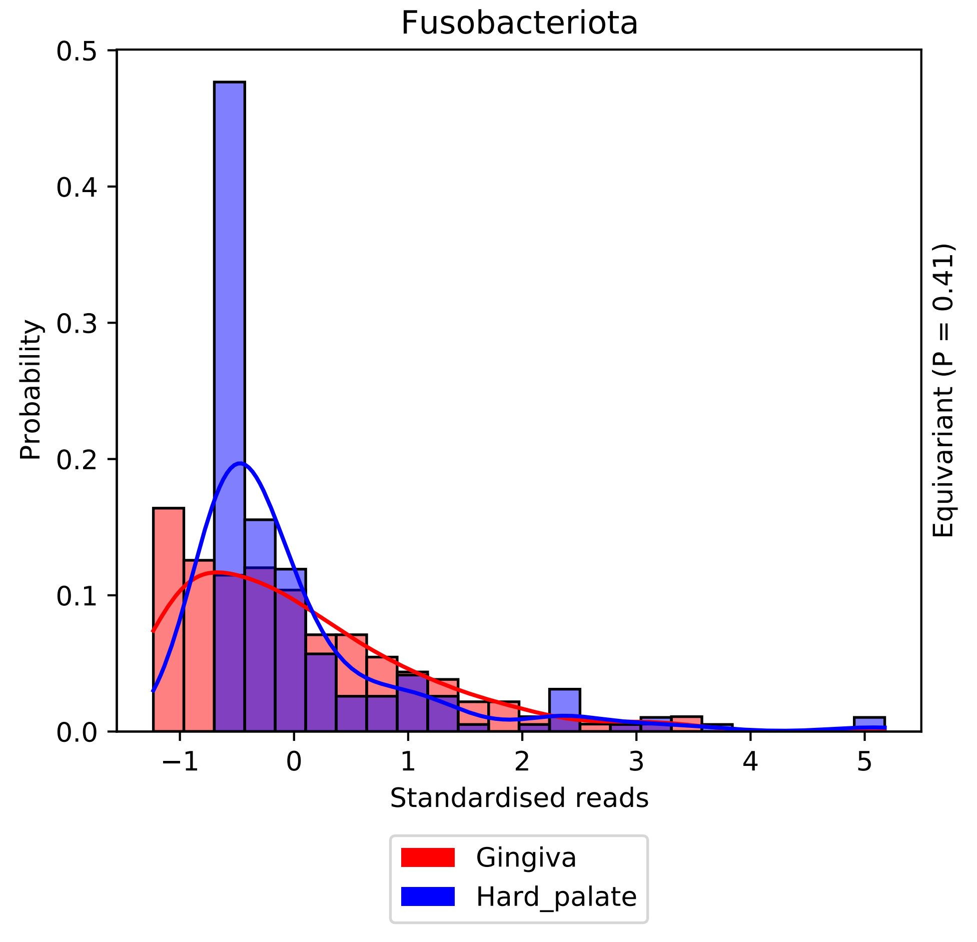

Fusobacteriota
0.5
0.4
0.3
0.2
0.1
0.0
Probability
−1
0
1
2
3
4
5
Standardised reads
Equivariant (P = 0.41)
Gingiva
Hard_palate

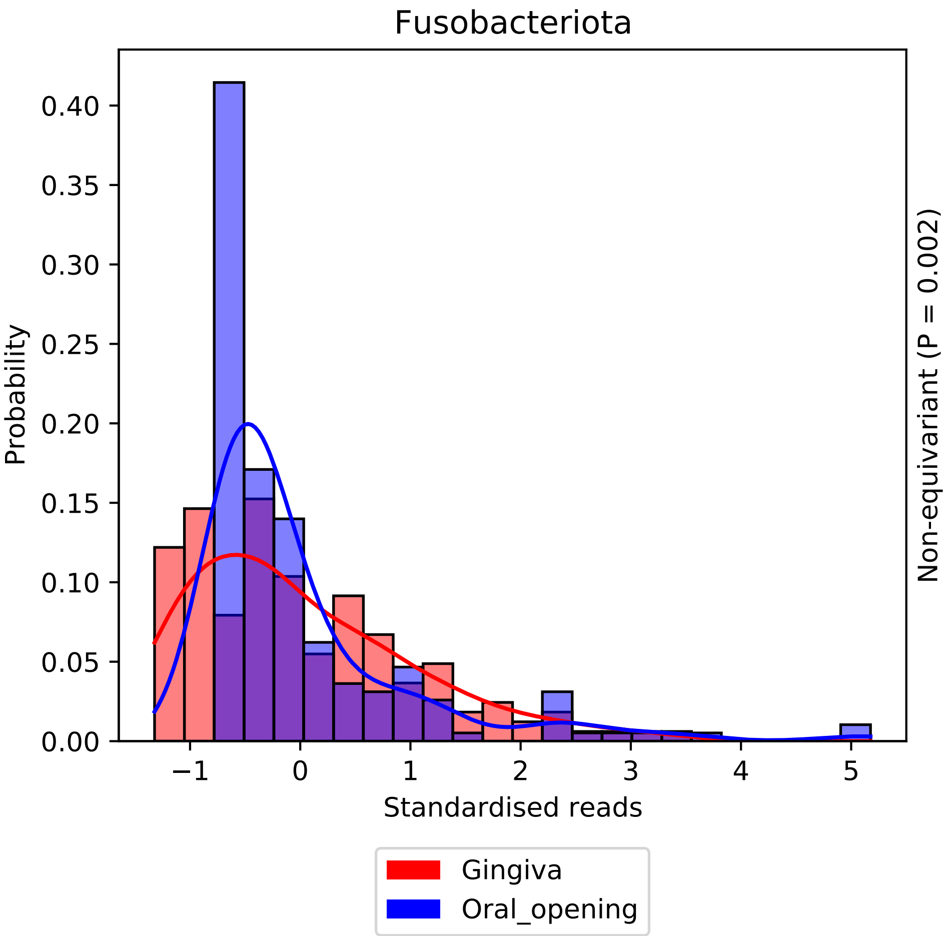

Fusobacteriota
Probability
0.00
0.05
0.10
0.15
0.20
0.25
0.30
0.35
0.40
−1
0
1
2
3
4
5
Standardised reads
Non-equivariant (P = 0.002)
Gingiva
Oral_opening

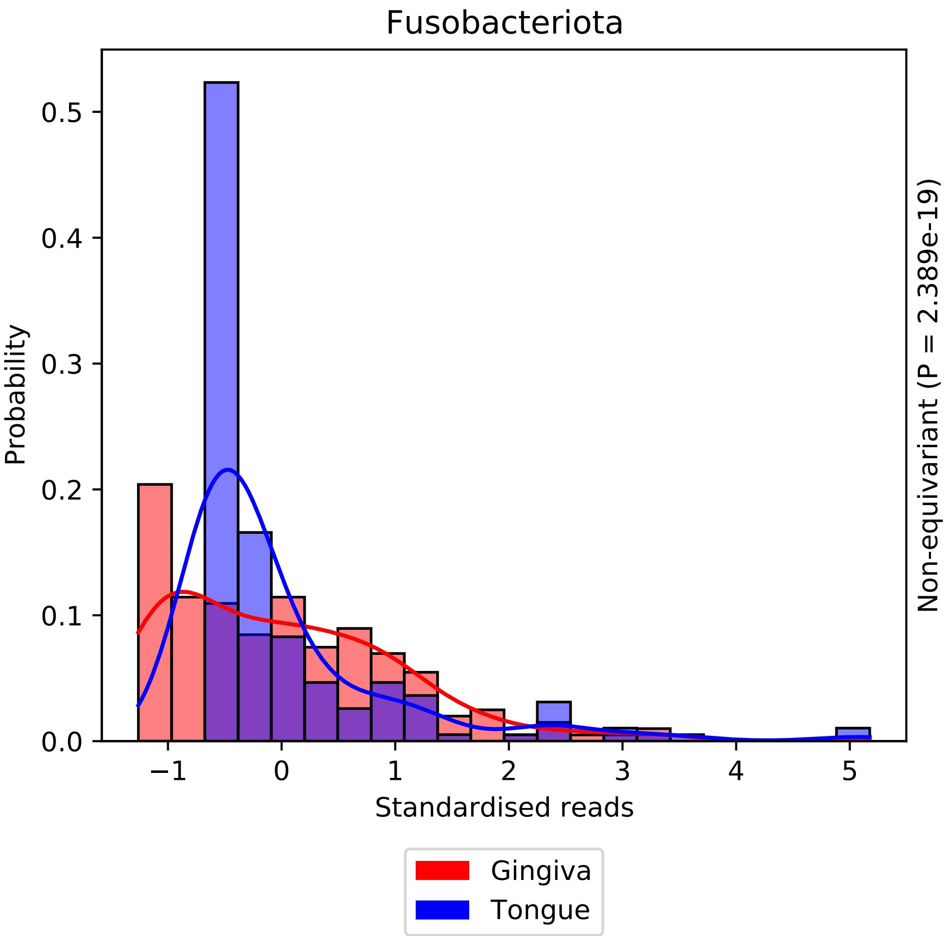

Fusobacteriota
Probability
Standardised reads
Non-equivariant (P = 2.389e-19)
0.0
0.1
0.2
0.3
0.4
0.5
−1
0
1
2
3
4
5
Gingiva
Tongue

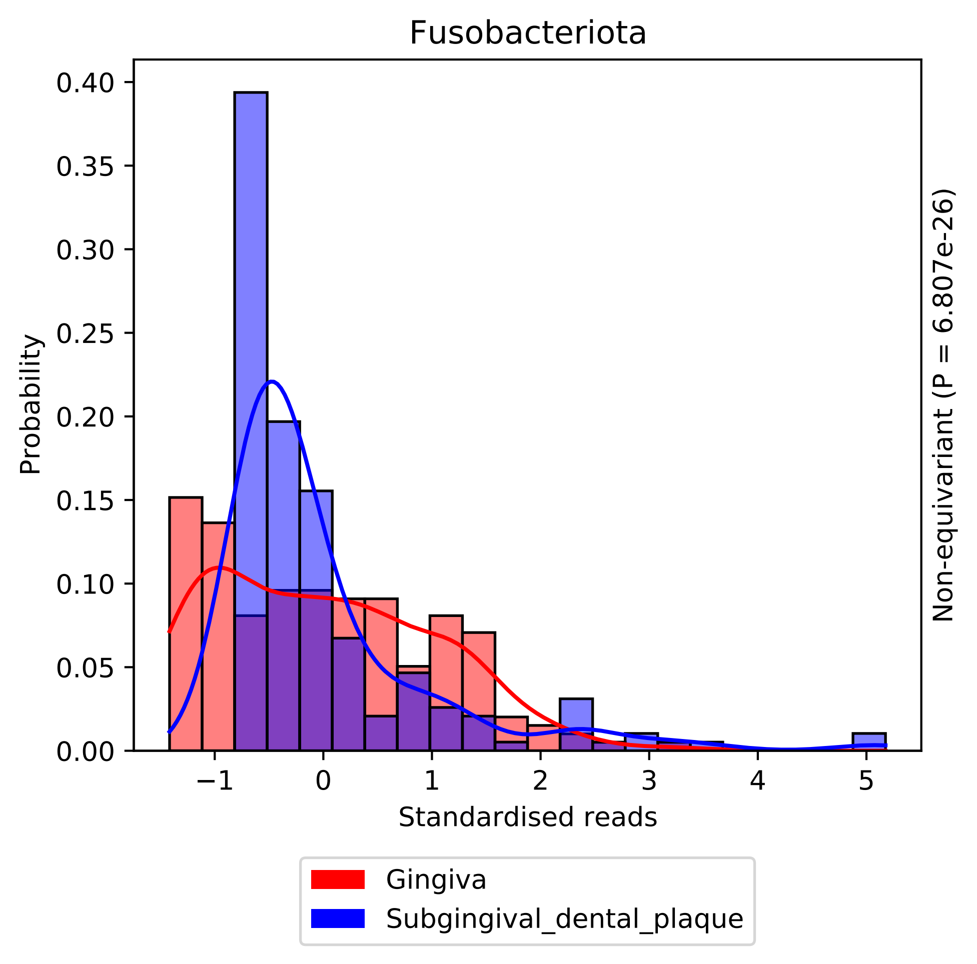

Fusobacteriota
Probability
0.40
0.35
0.30
0.25
0.20
0.15
0.10
0.05
0.00
−1
0
1
2
3
4
5
Standardised reads
Non-equivariant (P = 6.807e-26)
Gingiva
Subgingival_dental_plaque

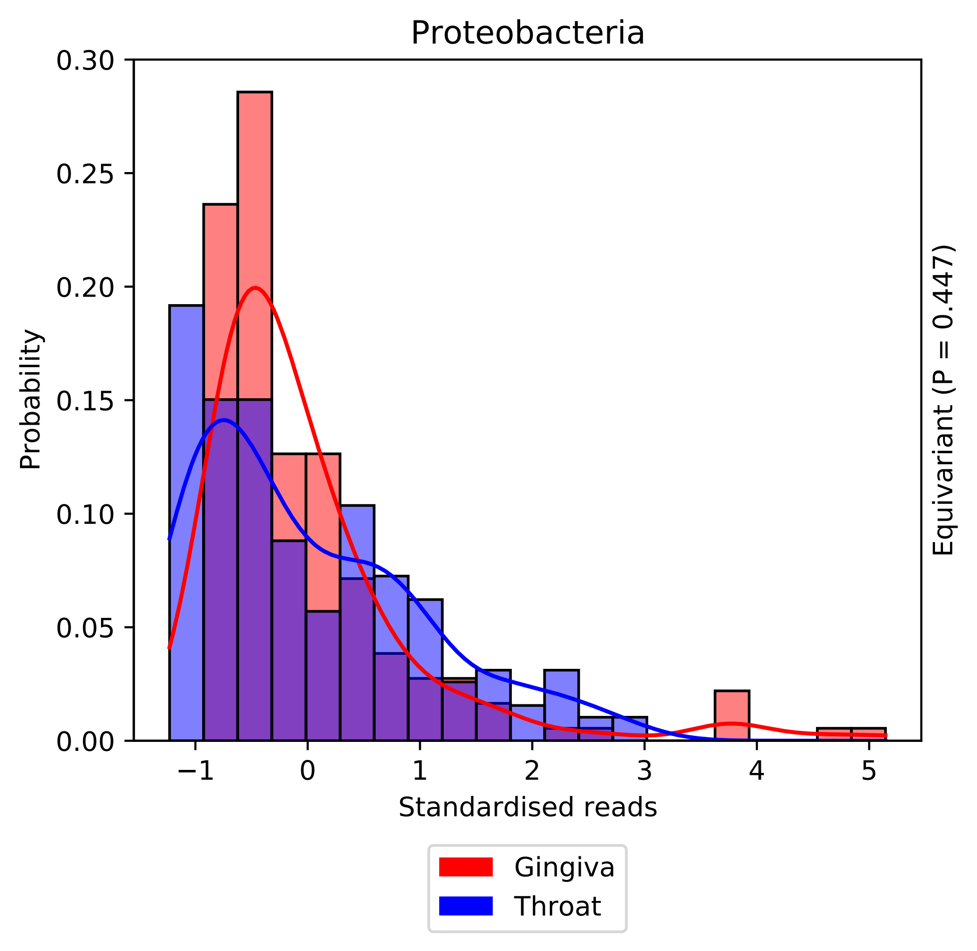

Proteobacteria
Probability
0.30
0.25
0.20
0.15
0.10
0.05
0.00
−1
0
1
2
3
4
5
Standardised reads
Equivariant (P = 0.447)
Gingiva
Throat

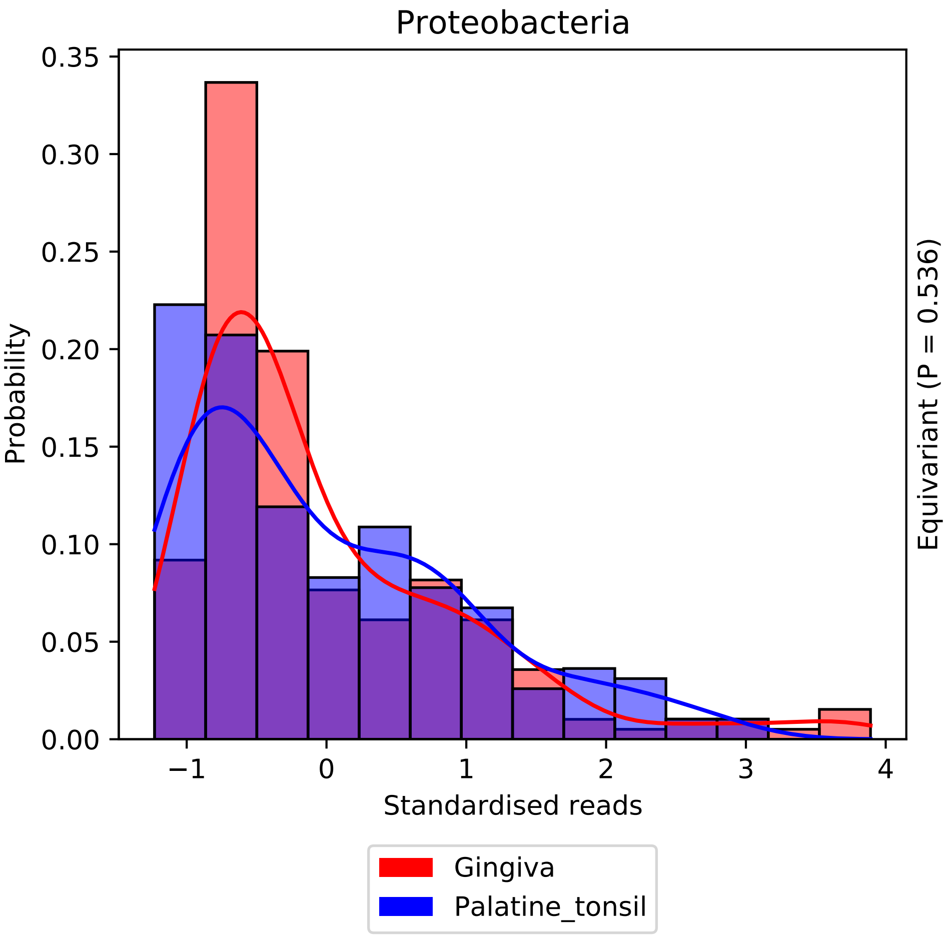
Proteobacteria
0.35
0.30
0.25
0.20
0.15
0.10
0.05
0.00
Probability
−1
0
1
2
3
4
Standardised reads
Equivariant (P = 0.536)
Gingiva
Palatine_tonsil

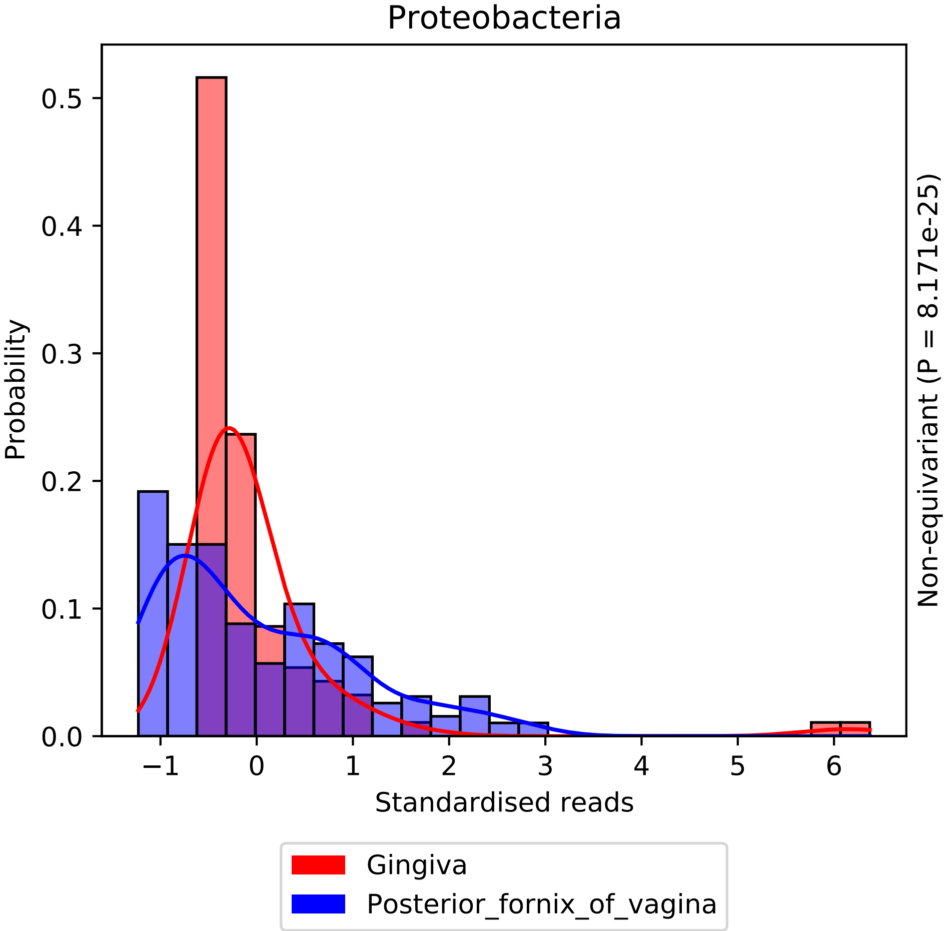

Proteobacteria
Probability
0.0
0.1
0.2
0.3
0.4
0.5
−1
0
1
2
3
4
5
6
Standardised reads
Non-equivariant (P = 8.171e-25)
Gingiva
Posterior_fornix_of_vagina

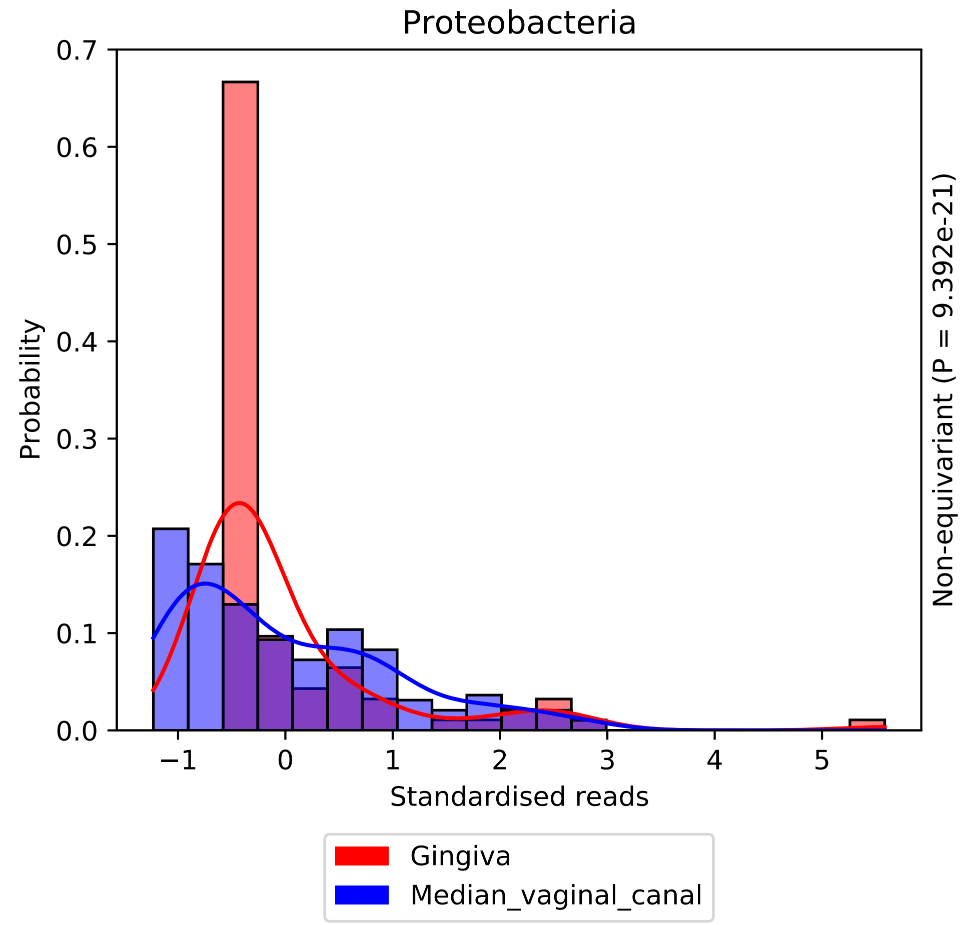
Proteobacteria
Probability
Standardised reads
Non-equivariant (P = 9.392e-21)
0.0
0.1
0.2
0.3
0.4
0.5
0.6
0.7
−1
0
1
2
3
4
5
Gingiva
Median_vaginal_canal

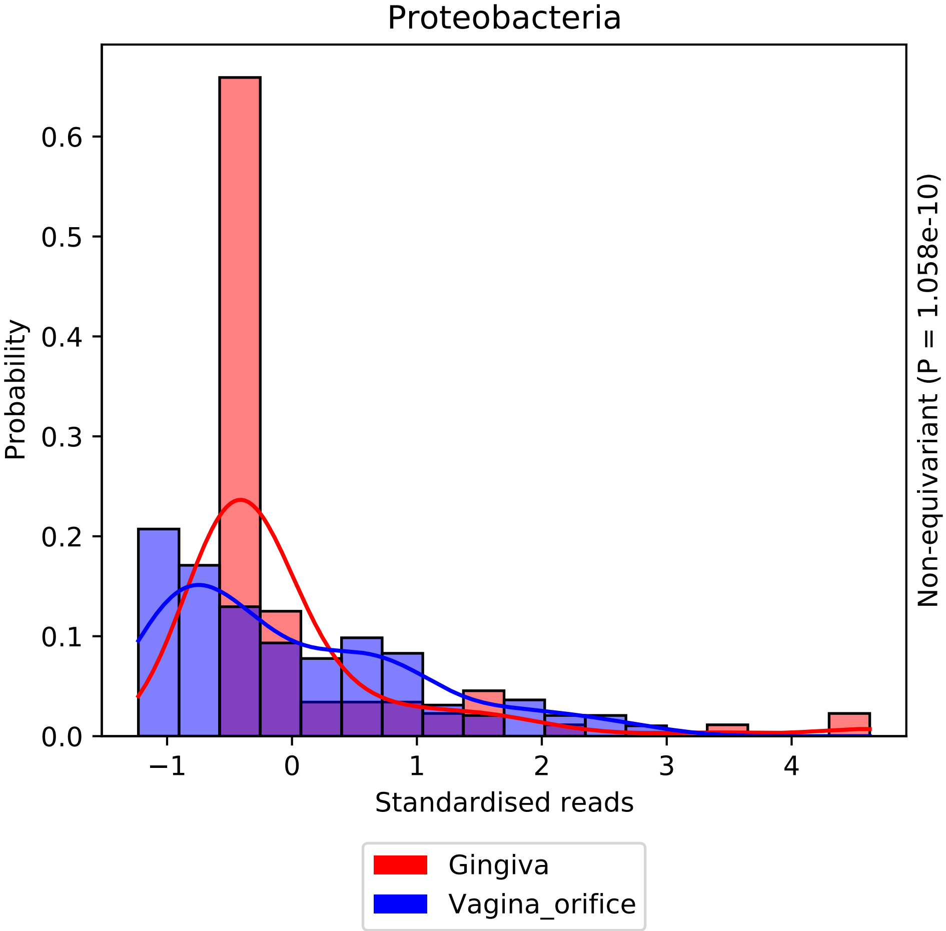
Proteobacteria
Probability
0.0
0.1
0.2
0.3
0.4
0.5
0.6
−1
0
1
2
3
4
Standardised reads
Non-equivariant (P = 1.058e-10)
Gingiva
Vagina_orifice

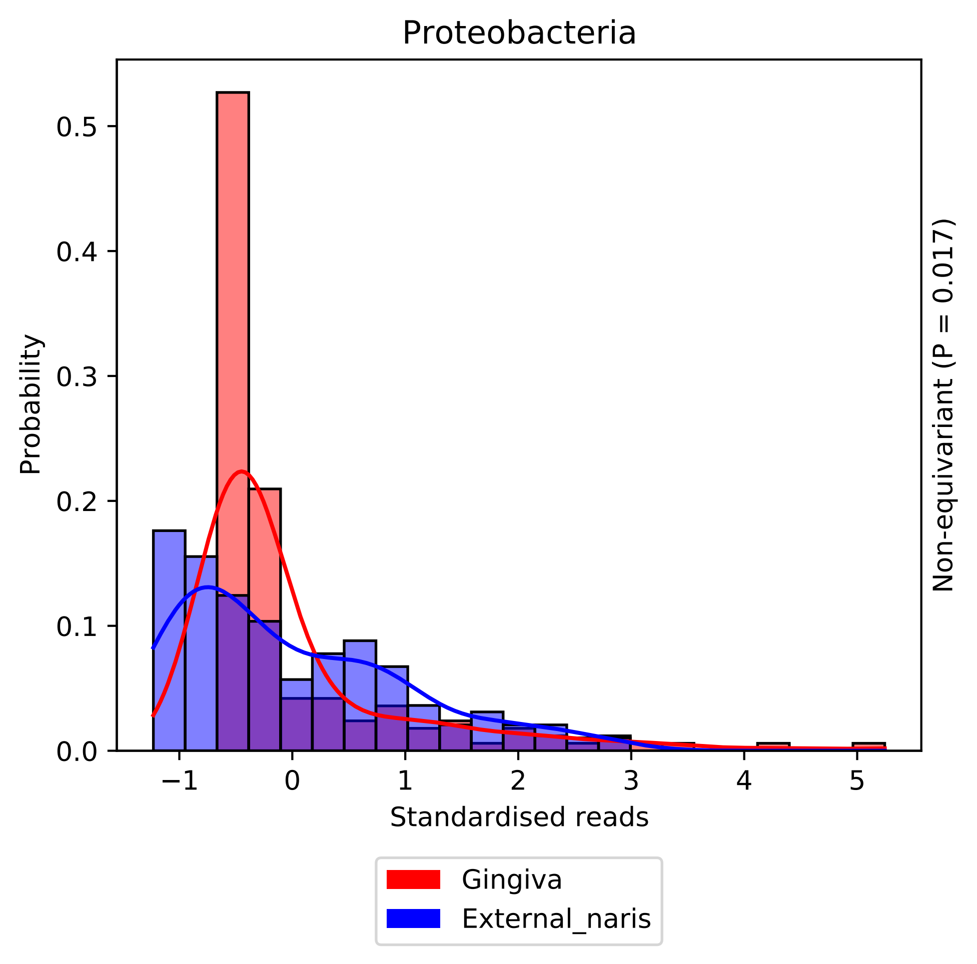

Proteobacteria
Probability
0.0
0.1
0.2
0.3
0.4
0.5
−1
0
1
2
3
4
5
Standardised reads
Non-equivariant (P = 0.017)
Gingiva
External_naris

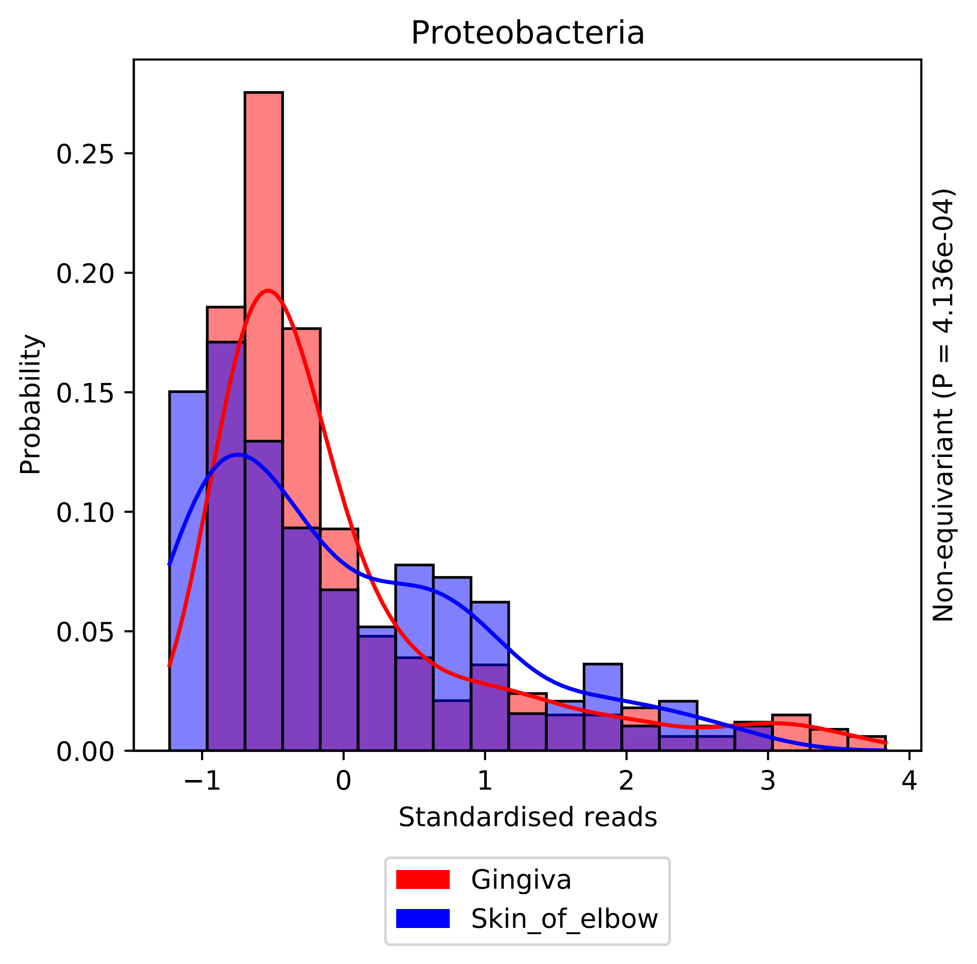

Proteobacteria
Probability
0.00
0.05
0.10
0.15
0.20
0.25
−1
0
1
2
3
4
Standardised reads
Non-equivariant (P = 4.136e-04)
Gingiva
Skin_of_elbow

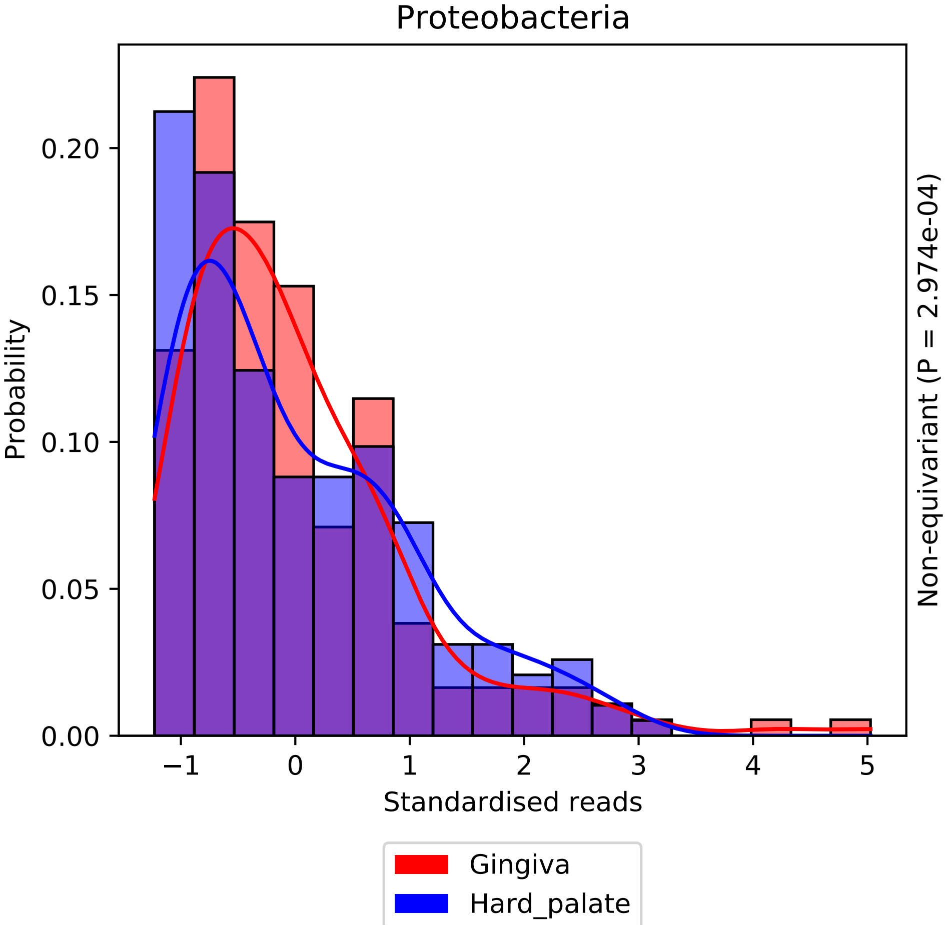

Proteobacteria
Probability
0.00
0.05
0.10
0.15
0.20
−1
0
1
2
3
4
5
Standardised reads
Non-equivariant (P = 2.974e-04)
Gingiva
Hard_palate

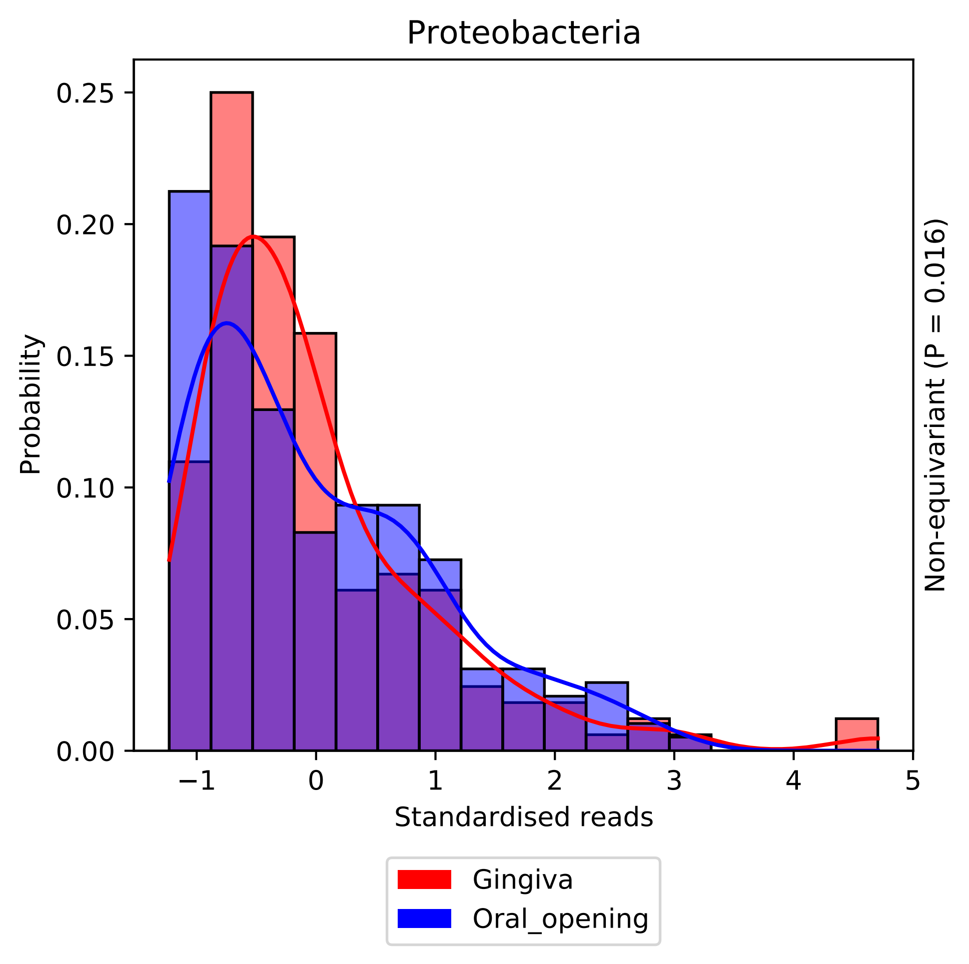

Proteobacteria
Probability
0.25
0.20
0.15
0.10
0.05
0.00
−1
0
1
2
3
4
5
Standardised reads
Non-equivariant (P = 0.016)
Gingiva
Oral_opening

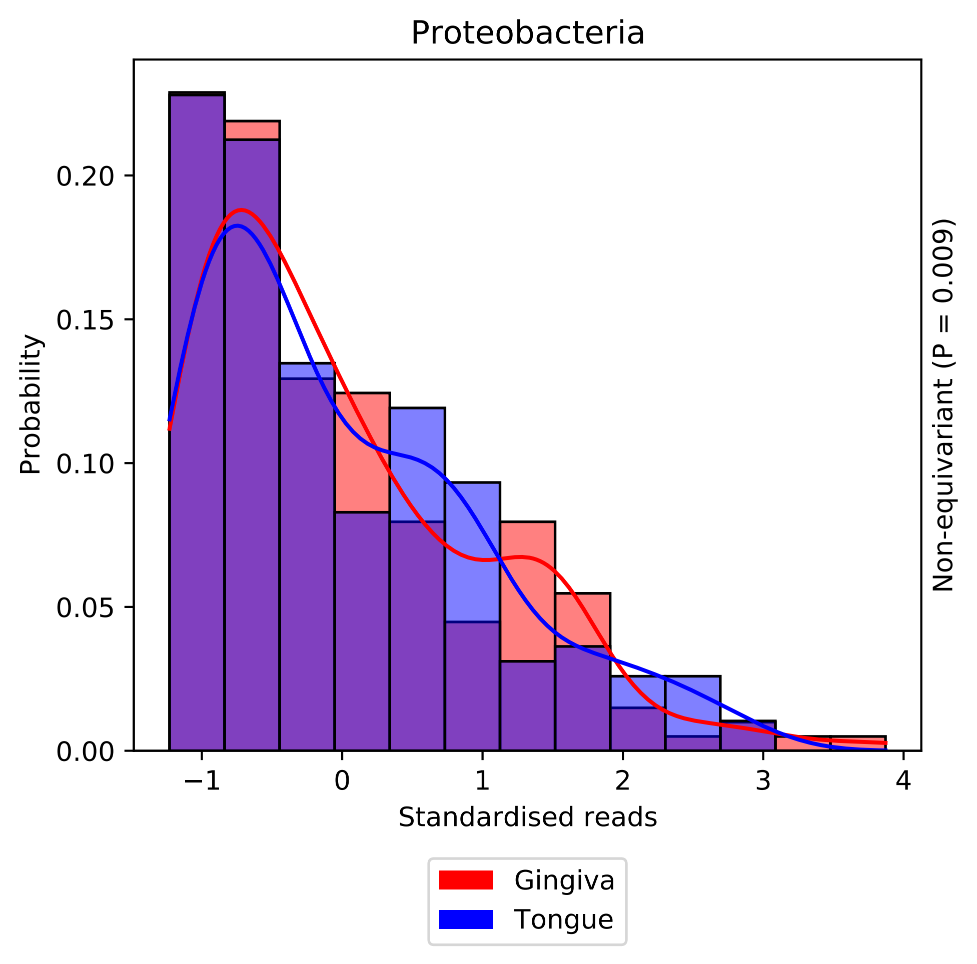

Proteobacteria
Probability
0.00
0.05
0.10
0.15
0.20
−1
0
1
2
3
4
Standardised reads
Non-equivariant (P = 0.009)
Gingiva
Tongue

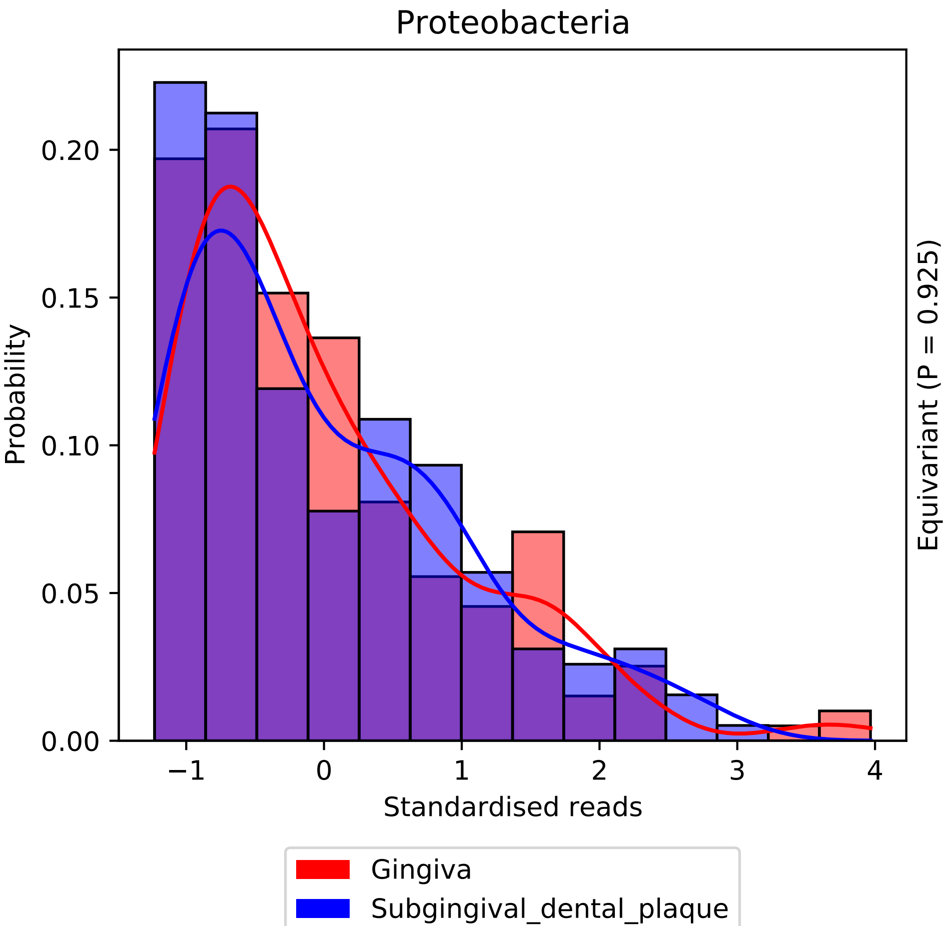

Proteobacteria
Probability
0.00
0.05
0.10
0.15
0.20
−1
0
1
2
3
4
Standardised reads
Equivariant (P = 0.925)
Gingiva
Subgingival_dental_plaque

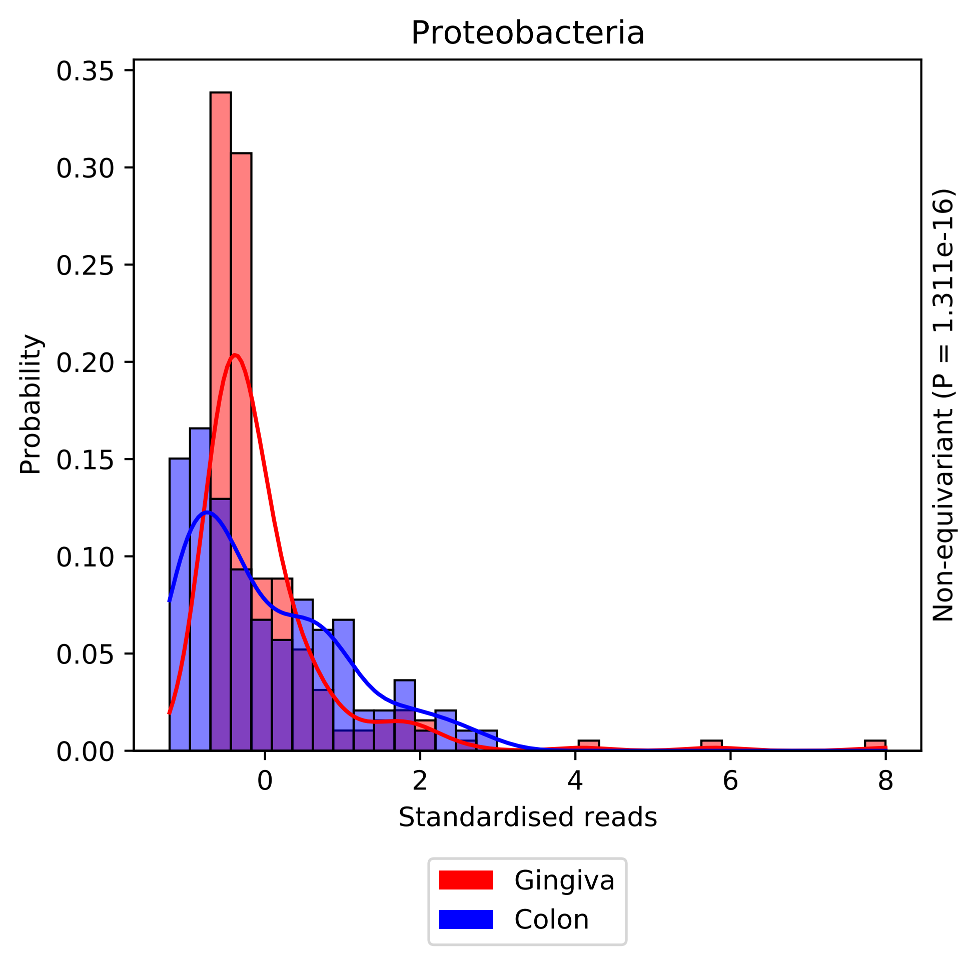
Proteobacteria
Probability
0.35
0.30
0.25
0.20
0.15
0.10
0.05
0.00
0
2
4
6
8
Standardised reads
Non-equivariant (P = 1.311e-16)
Gingiva
Colon

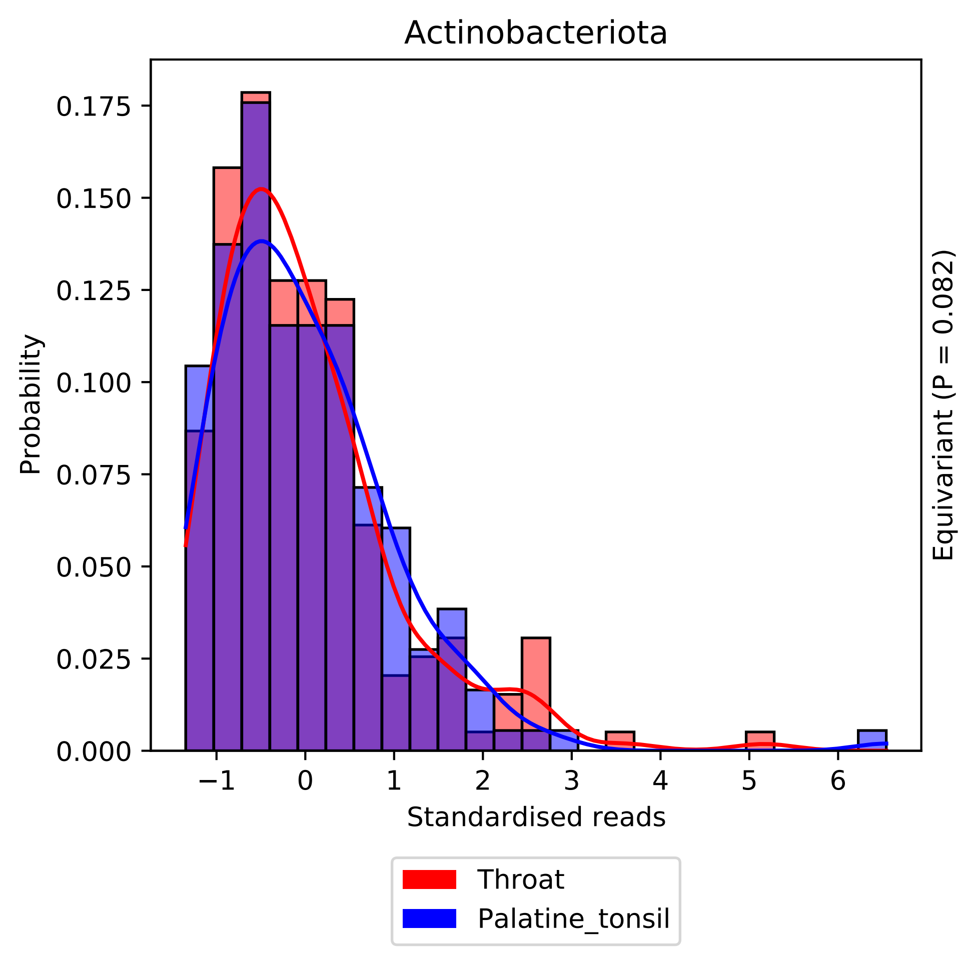

Actinobacteriota
Probability
0.175
0.150
0.125
0.100
0.075
0.050
0.025
0.000
−1
0
1
2
3
4
5
6
Standardised reads
Equivariant (P = 0.082)
Throat
Palatine_tonsil

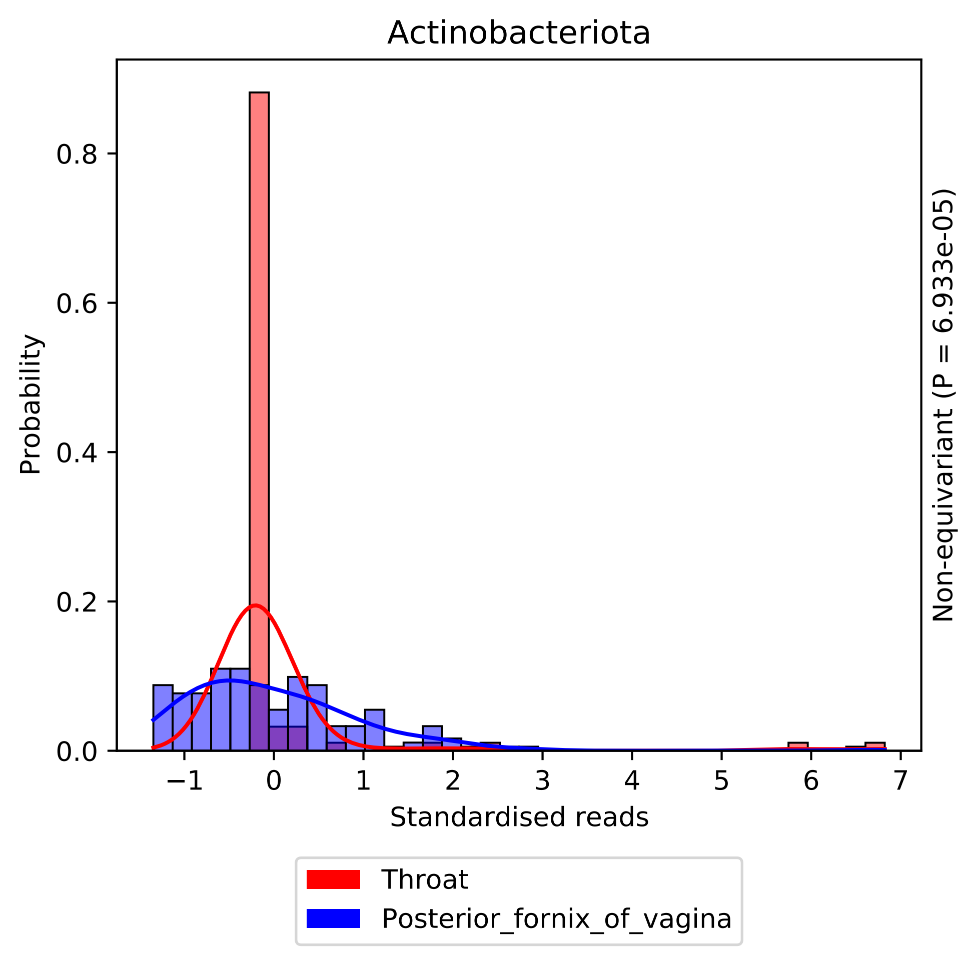

Actinobacteriota
Probability
0.0
0.2
0.4
0.6
0.8
−1
0
1
2
3
4
5
6
7
Standardised reads
Non-equivariant (P = 6.933e-05)
Throat
Posterior_fornix_of_vagina

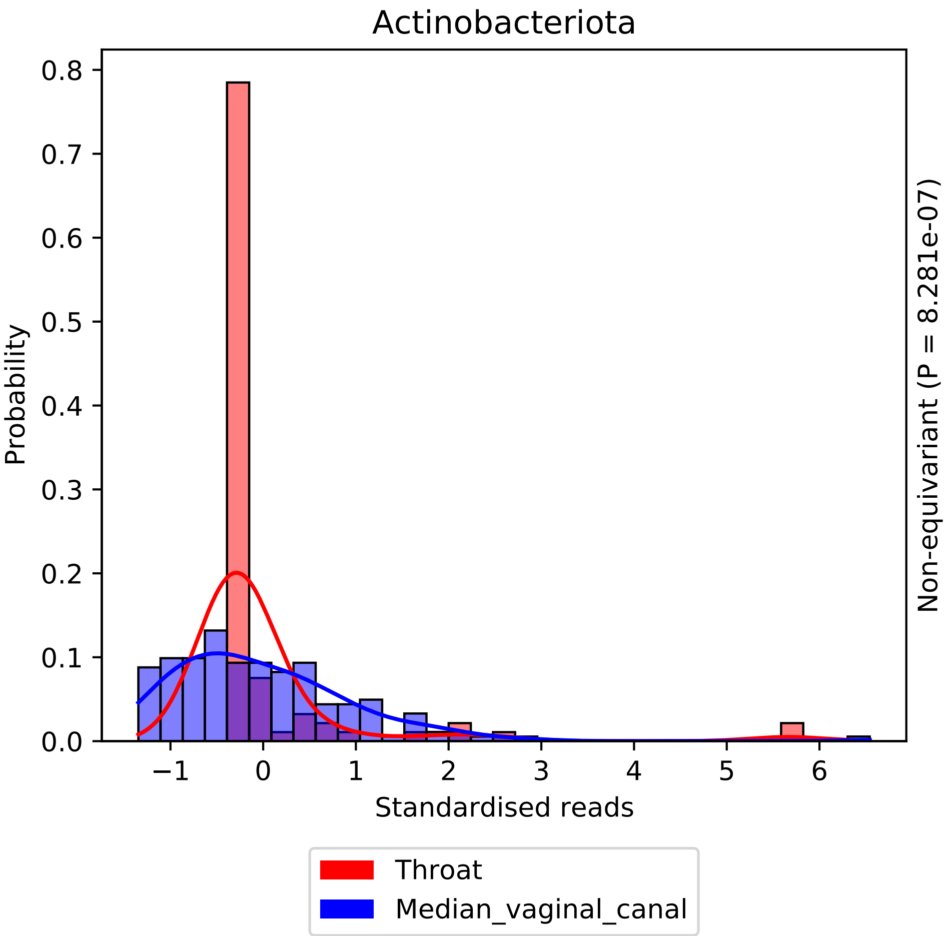

Actinobacteriota
Probability
0.0
0.1
0.2
0.3
0.4
0.5
0.6
0.7
0.8
−1
0
1
2
3
4
5
6
Standardised reads
Non-equivariant (P = 8.281e-07)
Throat
Median_vaginal_canal

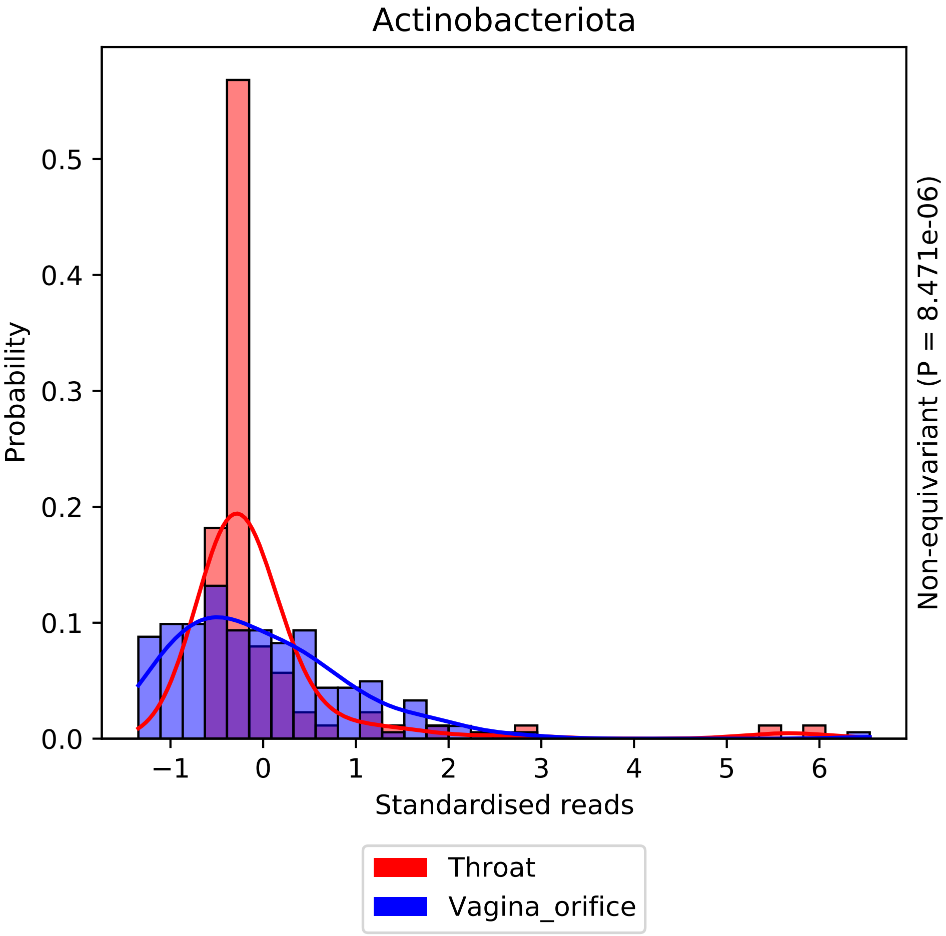

Actinobacteriota
Probability
0.0
0.1
0.2
0.3
0.4
0.5
−1
0
1
2
3
4
5
6
Standardised reads
Non-equivariant (P = 8.471e-06)
Throat
Vagina_orifice

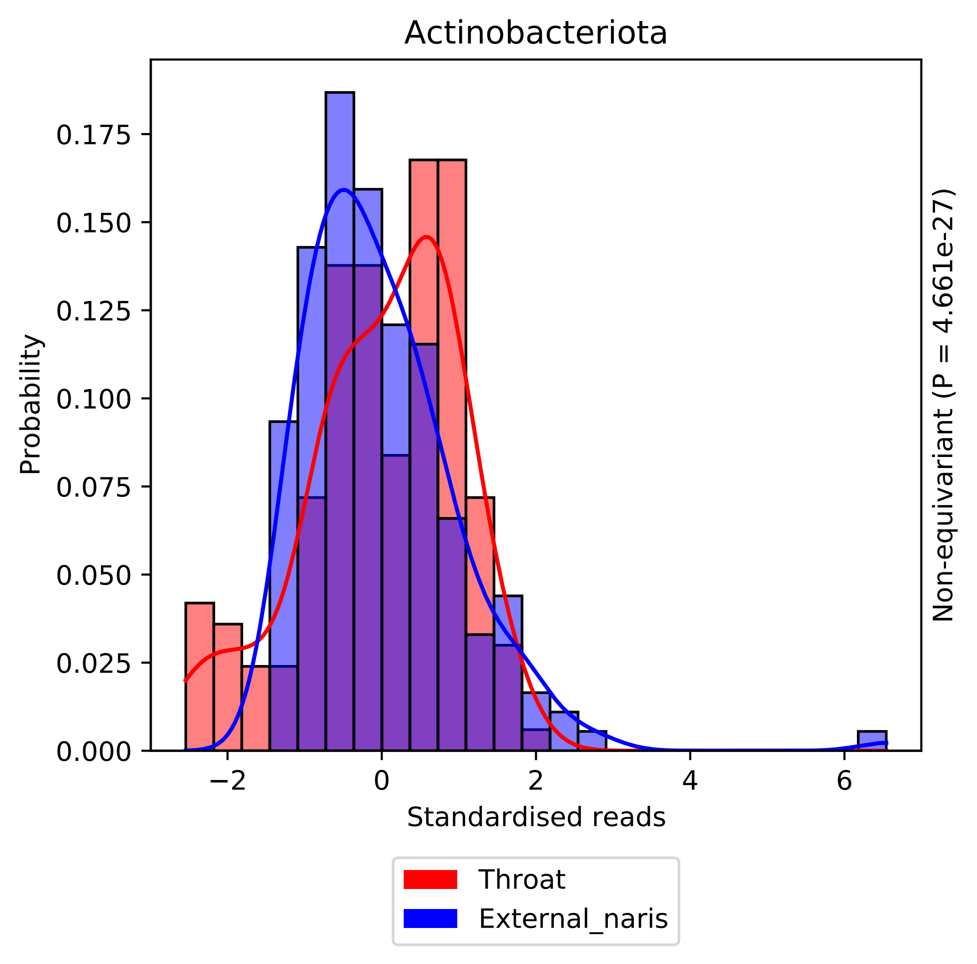

Actinobacteriota
Probability
0.000
0.025
0.050
0.075
0.100
0.125
0.150
0.175
−2
0
2
4
6
Standardised reads
Non-equivariant (P = 4.661e-27)
Throat
External_naris

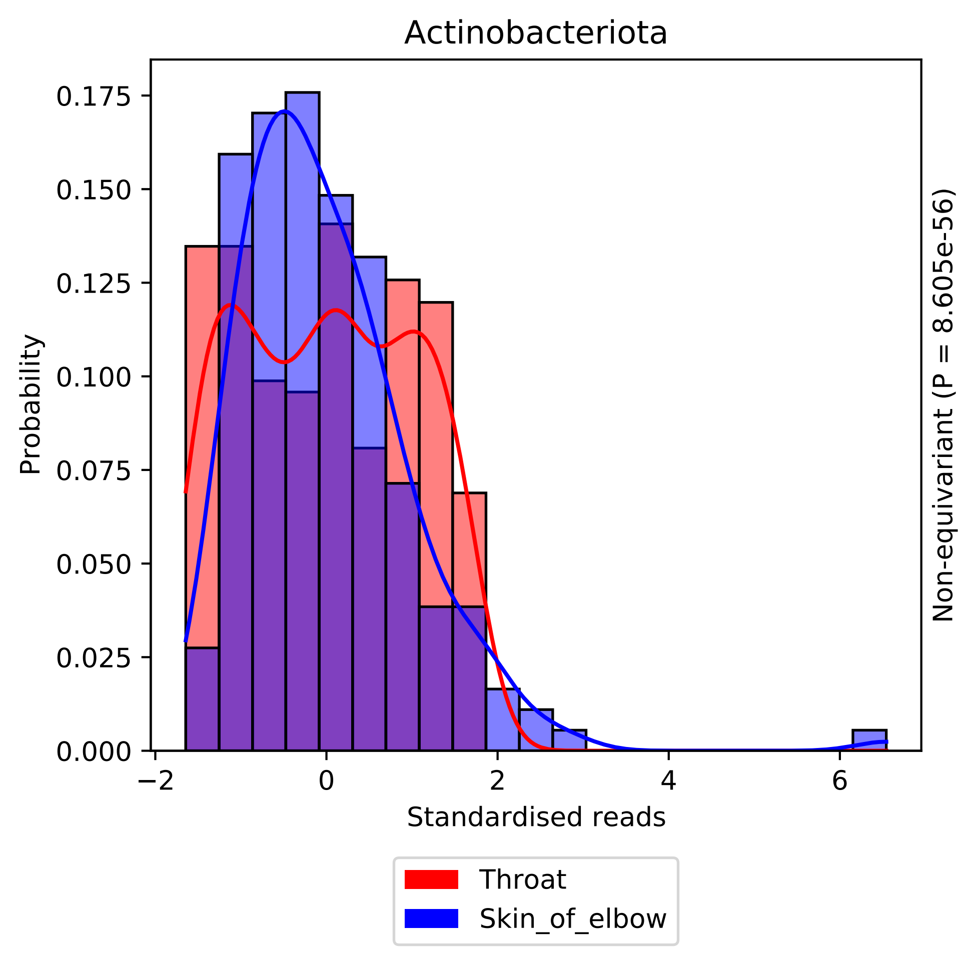

Actinobacteriota
Probability
0.000
0.025
0.050
0.075
0.100
0.125
0.150
0.175
−2
0
2
4
6
Standardised reads
Non-equivariant (P = 8.605e-56)
Throat
Skin_of_elbow

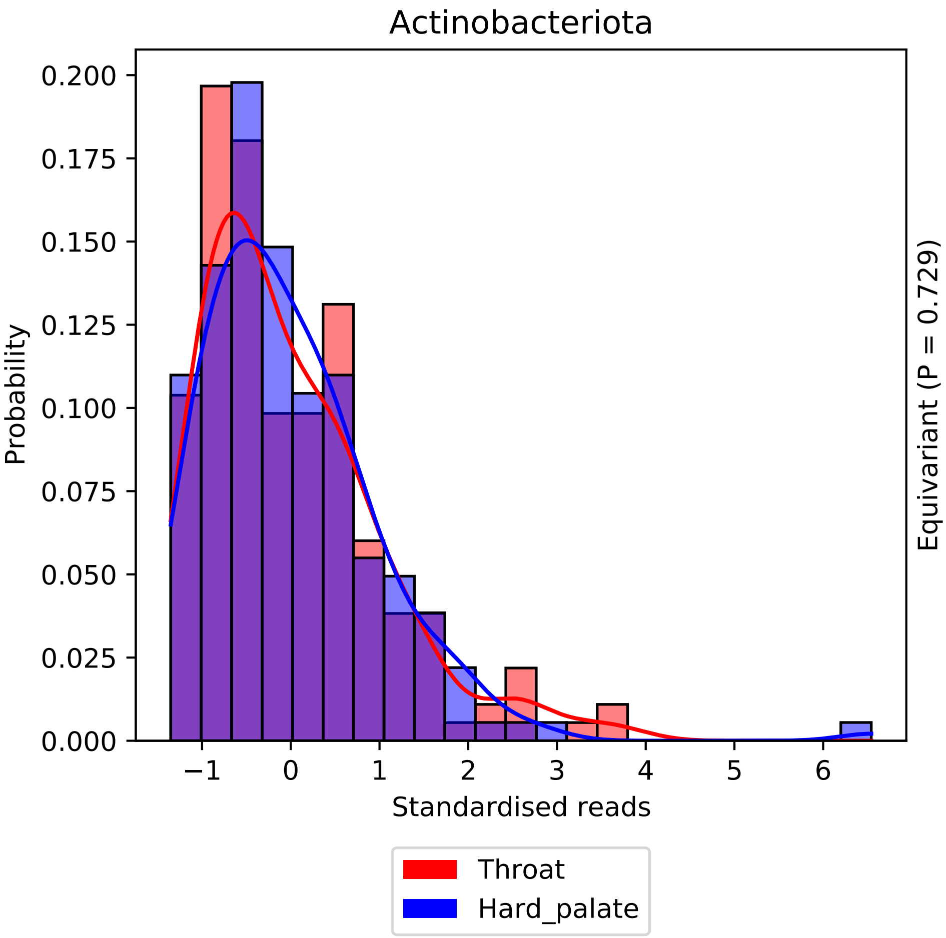

Actinobacteriota
Probability
0.200
0.175
0.150
0.125
0.100
0.075
0.050
0.025
0.000
−1
0
1
2
3
4
5
6
Standardised reads
Equivariant (P = 0.729)
Throat
Hard_palate

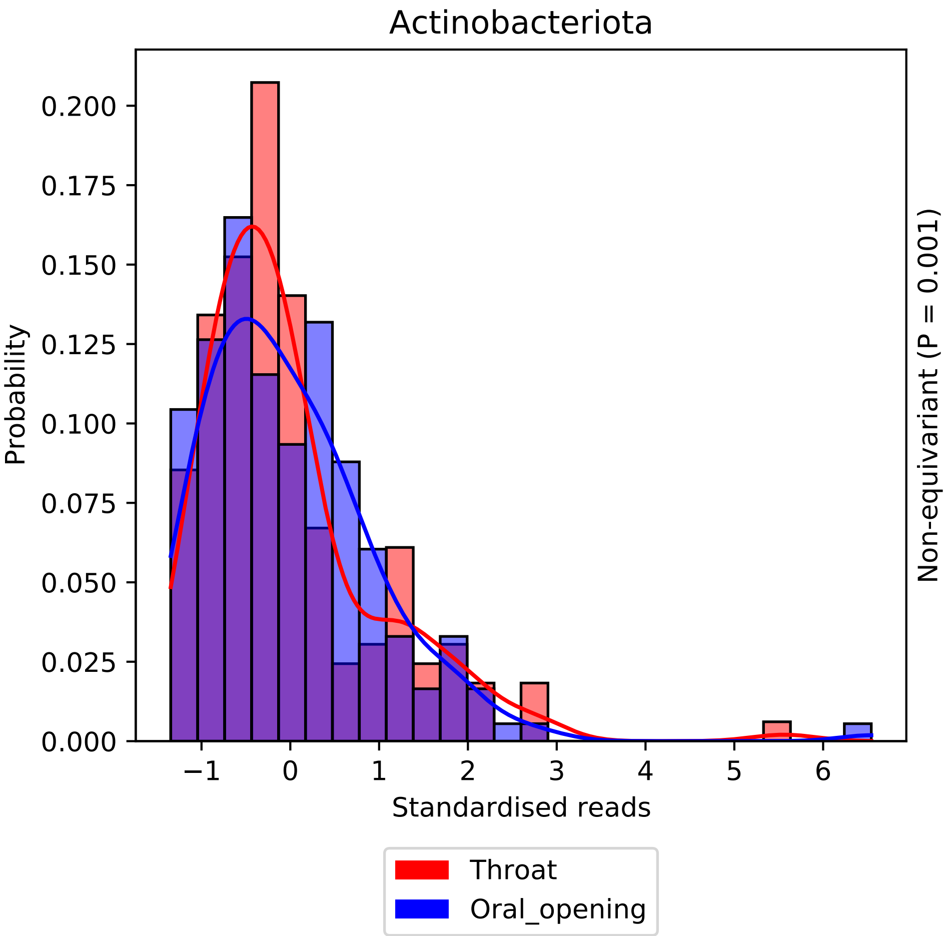

Actinobacteriota
Probability
0.200
0.175
0.150
0.125
0.100
0.075
0.050
0.025
0.000
−1
0
1
2
3
4
5
6
Standardised reads
Non-equivariant (P = 0.001)
Throat
Oral_opening

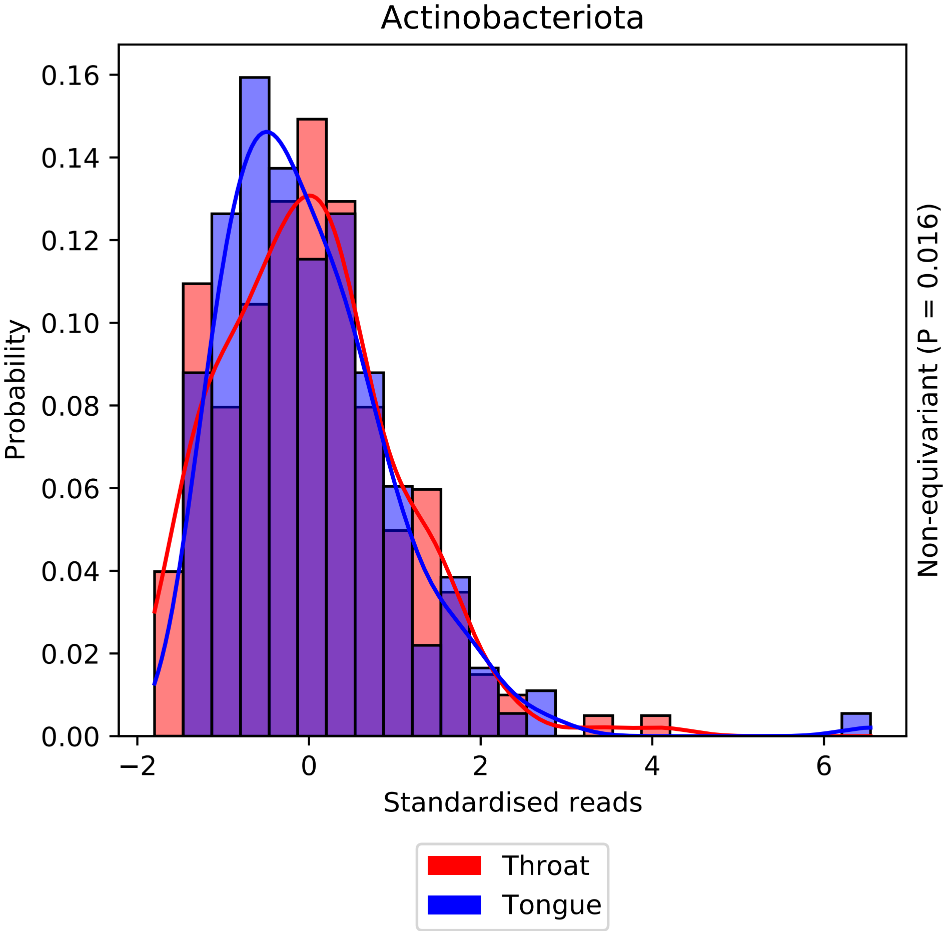

Actinobacteriota
Probability
Standardised reads
Non-equivariant (P = 0.016)
0.00
0.02
0.04
0.06
0.08
0.10
0.12
0.14
0.16
−2
0
2
4
6
Throat
Tongue

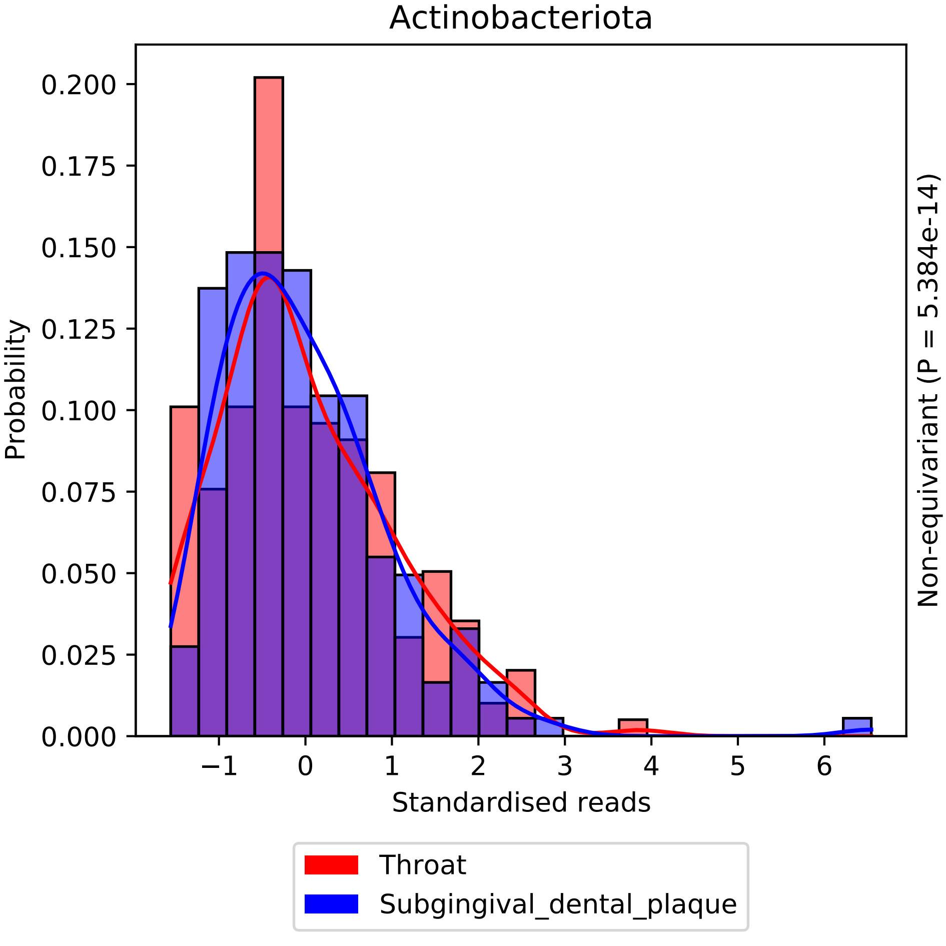

Actinobacteriota
Probability
0.200
0.175
0.150
0.125
0.100
0.075
0.050
0.025
0.000
−1
0
1
2
3
4
5
6
Standardised reads
Non-equivariant (P = 5.384e-14)
Throat
Subgingival_dental_plaque

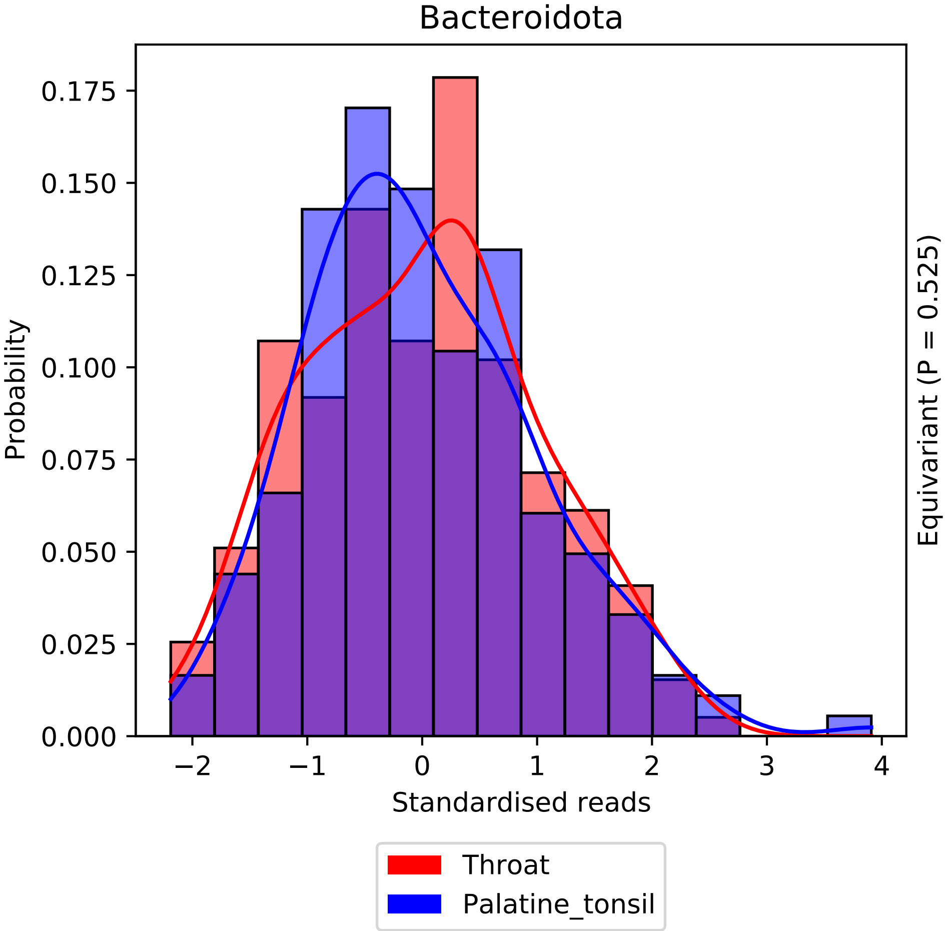

Bacteroidota
Probability
0.000
0.025
0.050
0.075
0.100
0.125
0.150
0.175
−2
−1
0
1
2
3
4
Standardised reads
Equivariant (P = 0.525)
Throat
Palatine_tonsil

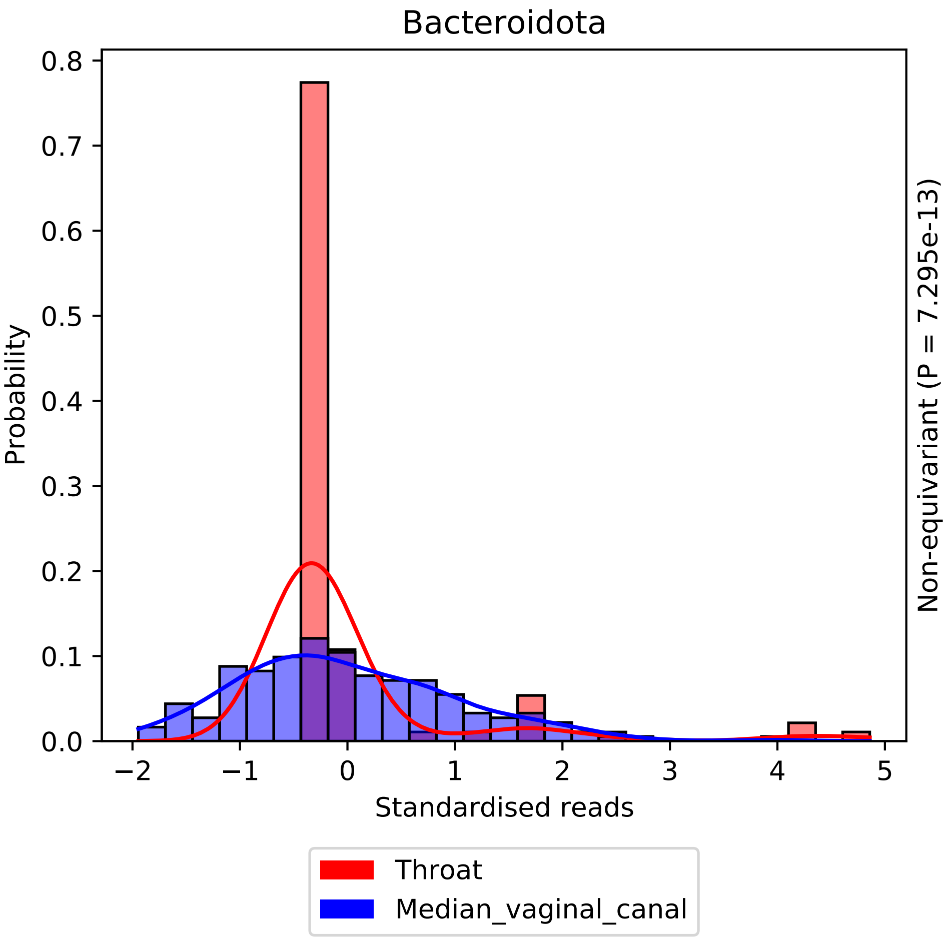
Bacteroidota
Probability
Standardised reads
Non-equivariant (P = 7.295e-13)
Throat
Median_vaginal_canal
0.0
0.1
0.2
0.3
0.4
0.5
0.6
0.7
0.8
−2
−1
0
1
2
3
4
5

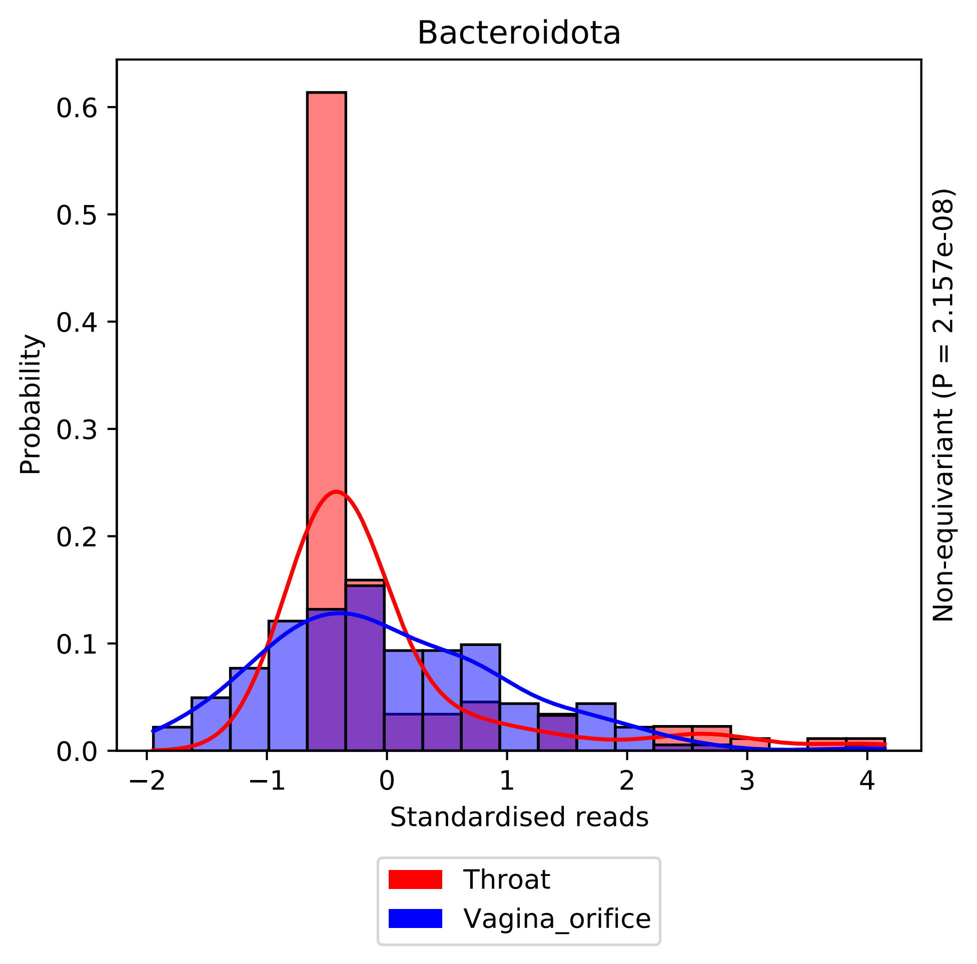
Bacteroidota
Probability
0.0
0.1
0.2
0.3
0.4
0.5
0.6
−2
−1
0
1
2
3
4
Standardised reads
Non-equivariant (P = 2.157e-08)
Throat
Vagina_orifice

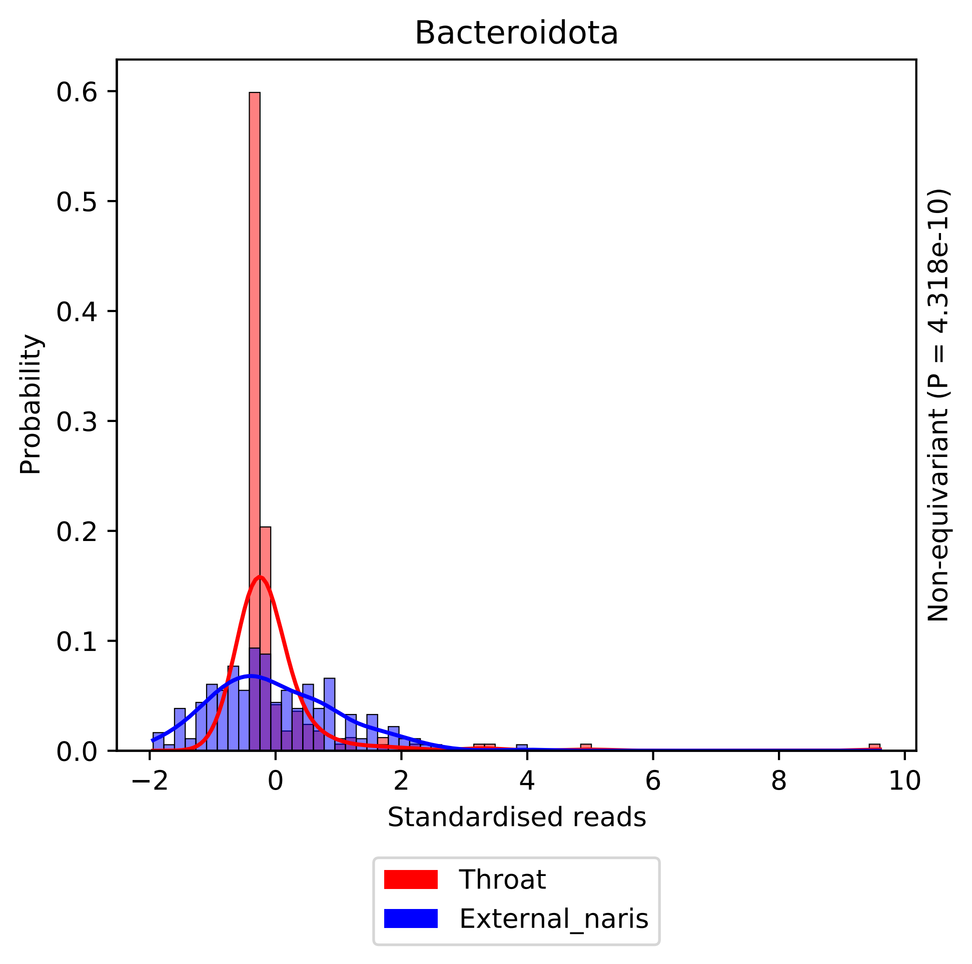

Bacteroidota
Probability
0.0
0.1
0.2
0.3
0.4
0.5
0.6
−2
0
2
4
6
8
10
Standardised reads
Non-equivariant (P = 4.318e-10)
Throat
External_naris

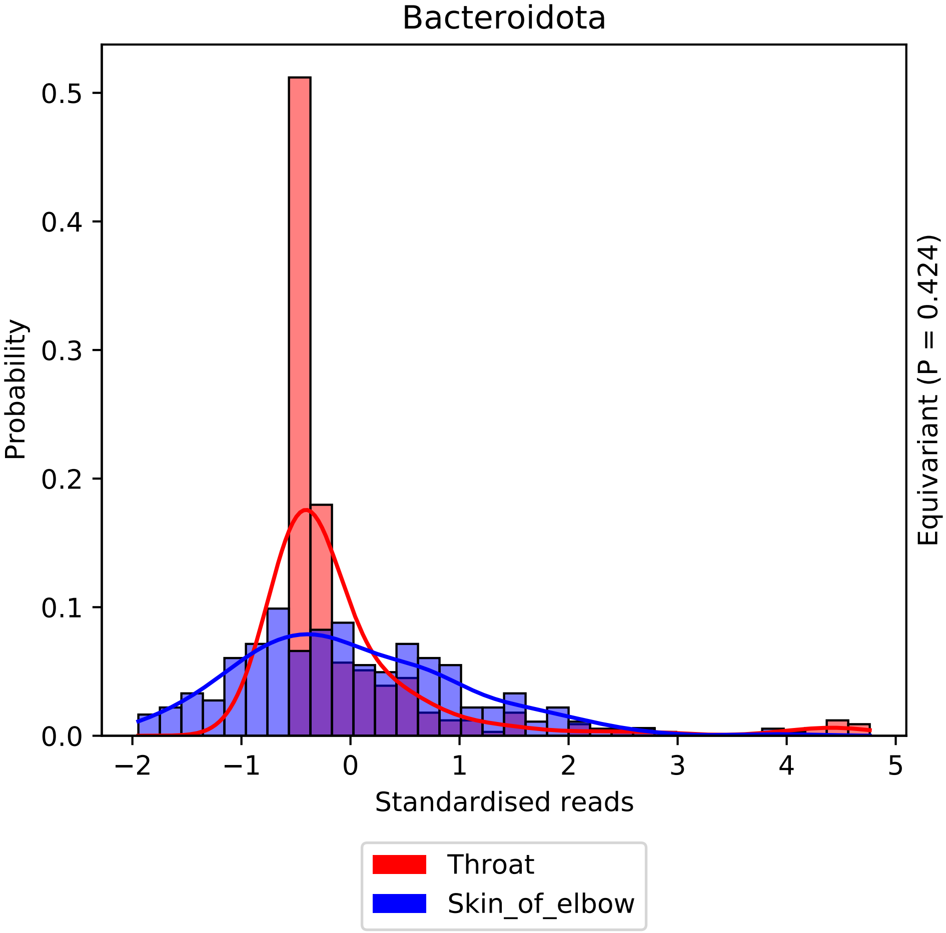

Bacteroidota
Probability
0.0
0.1
0.2
0.3
0.4
0.5
−2
−1
0
1
2
3
4
5
Standardised reads
Equivariant (P = 0.424)
Throat
Skin_of_elbow

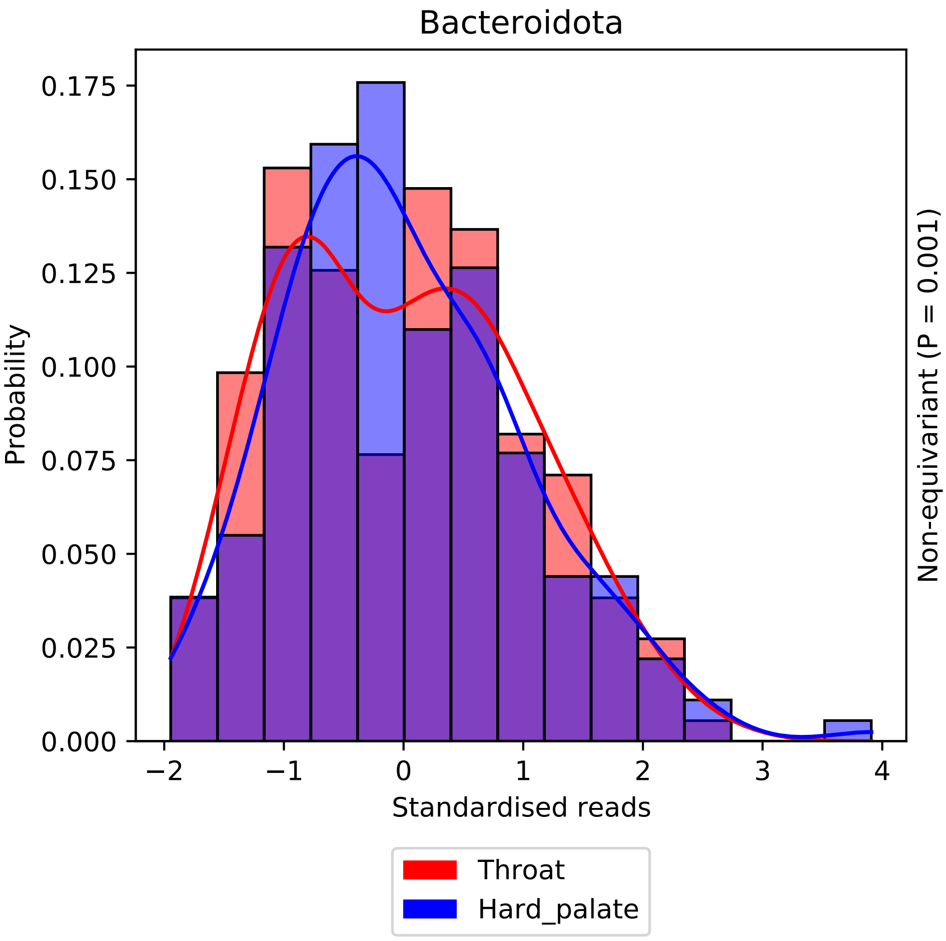

Bacteroidota
Probability
0.000
0.025
0.050
0.075
0.100
0.125
0.150
0.175
−2
−1
0
1
2
3
4
Standardised reads
Non-equivariant (P = 0.001)
Throat
Hard_palate

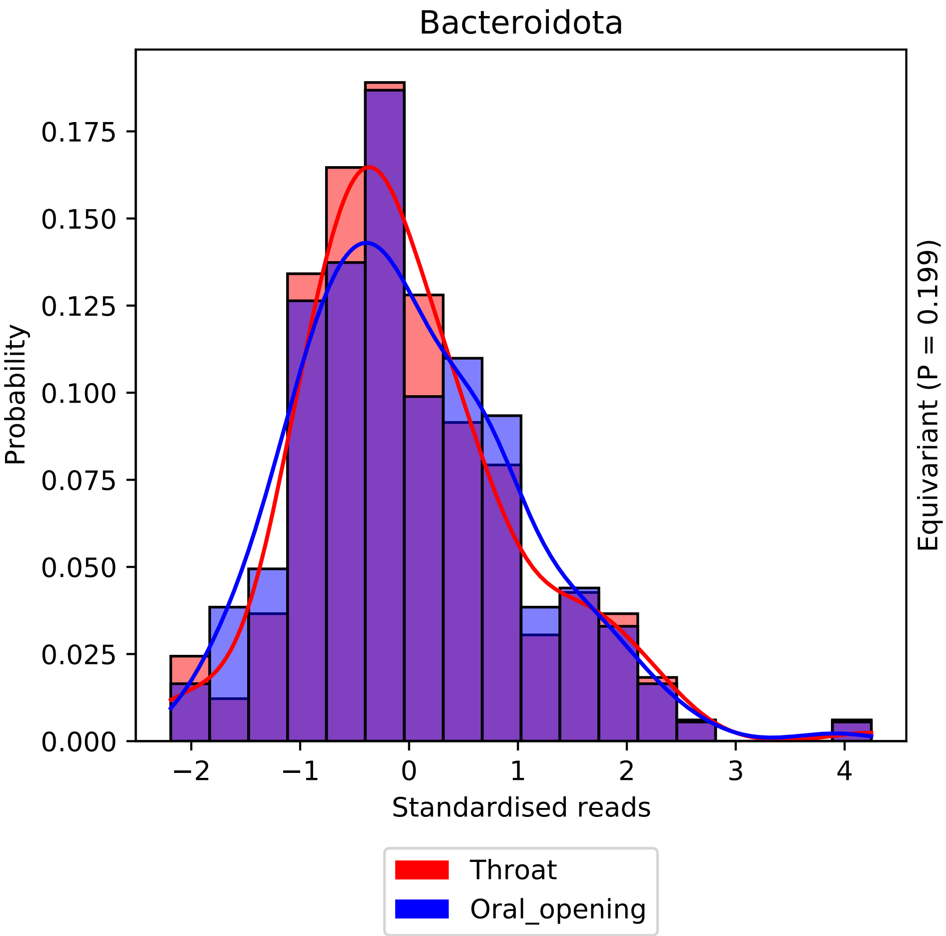
Bacteroidota
Probability
0.000
0.025
0.050
0.075
0.100
0.125
0.150
0.175
−2
−1
0
1
2
3
4
Standardised reads
Equivariant (P = 0.199)
Throat
Oral_opening

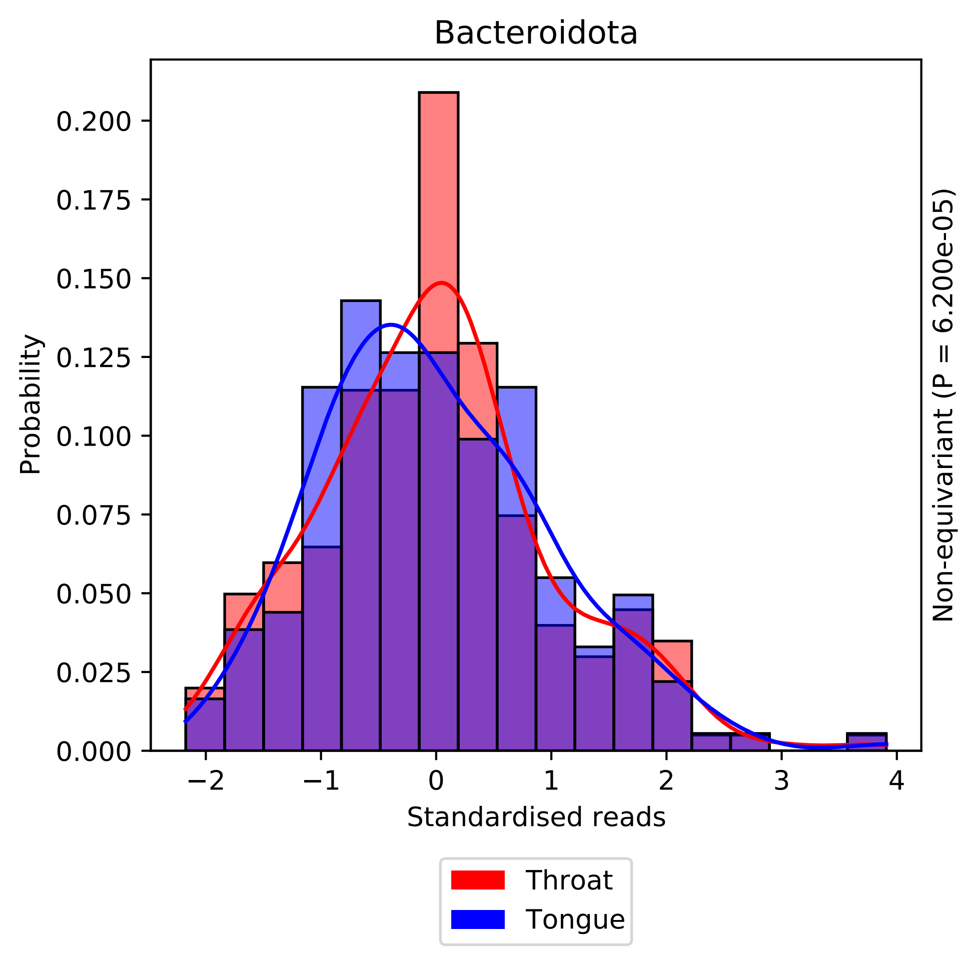

Bacteroidota
Probability
0.000
0.025
0.050
0.075
0.100
0.125
0.150
0.175
0.200
−2
−1
0
1
2
3
4
Standardised reads
Non-equivariant (P = 6.200e-05)
Throat
Tongue

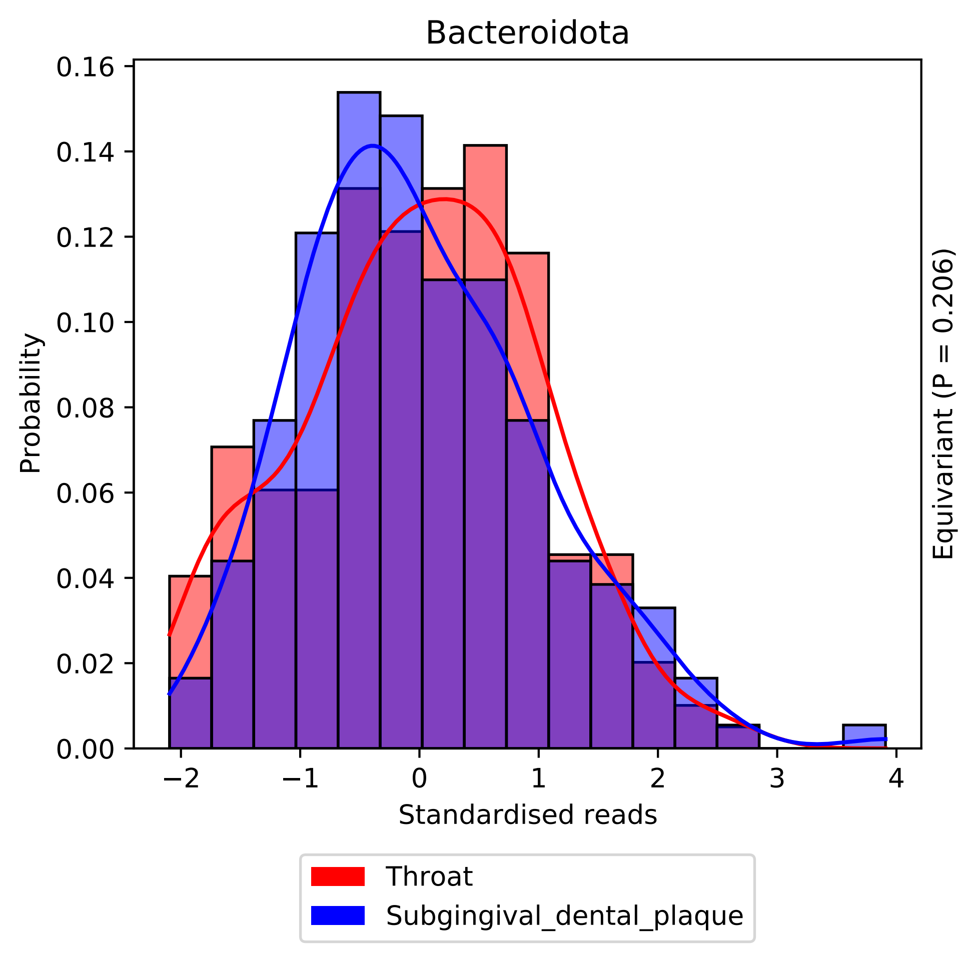
Bacteroidota
Probability
0.16
0.14
0.12
0.10
0.08
0.06
0.04
0.02
0.00
−2
−1
0
1
2
3
4
Standardised reads
Equivariant (P = 0.206)
Throat
Subgingival_dental_plaque

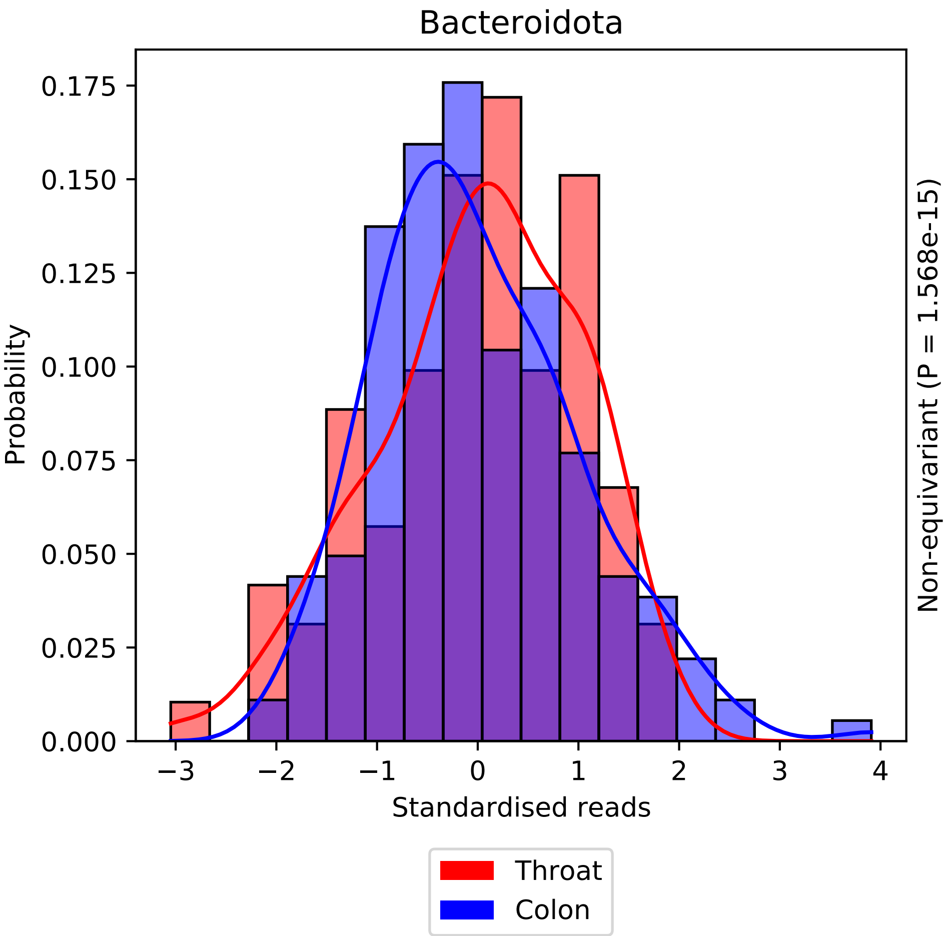

Bacteroidota
Probability
0.000
0.025
0.050
0.075
0.100
0.125
0.150
0.175
−3
−2
−1
0
1
2
3
4
Standardised reads
Non-equivariant (P = 1.568e-15)
Throat
Colon

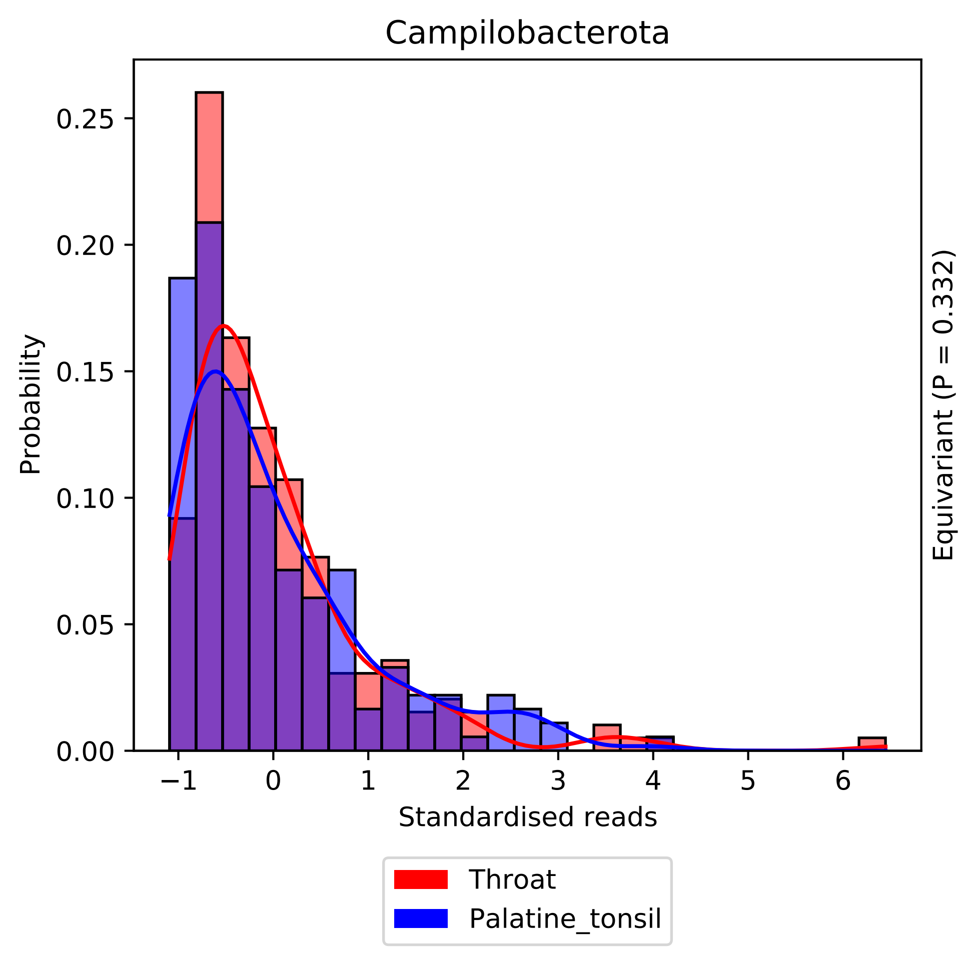

Campilobacterota
Probability
Standardised reads
Equivariant (P = 0.332)
0.00
0.05
0.10
0.15
0.20
0.25
−1
0
1
2
3
4
5
6
Throat
Palatine_tonsil

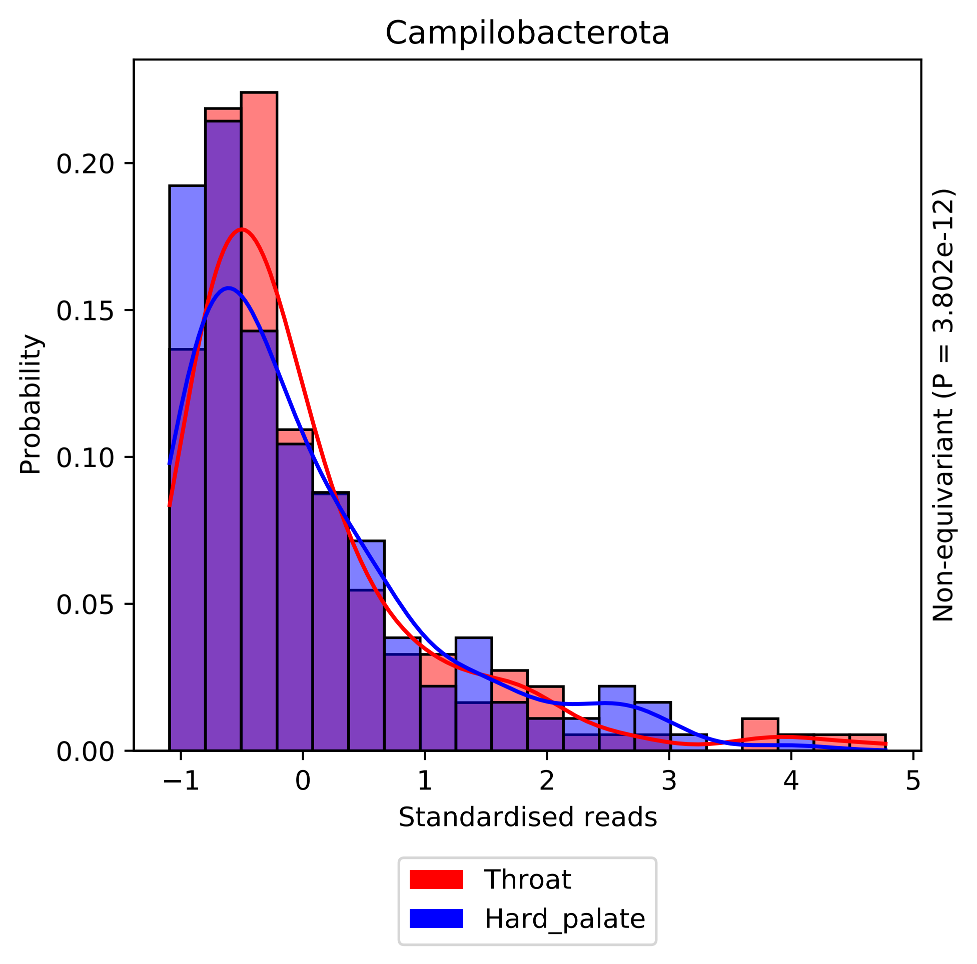

Campilobacterota
Probability
0.00
0.05
0.10
0.15
0.20
−1
0
1
2
3
4
5
Standardised reads
Non-equivariant (P = 3.802e-12)
Throat
Hard_palate

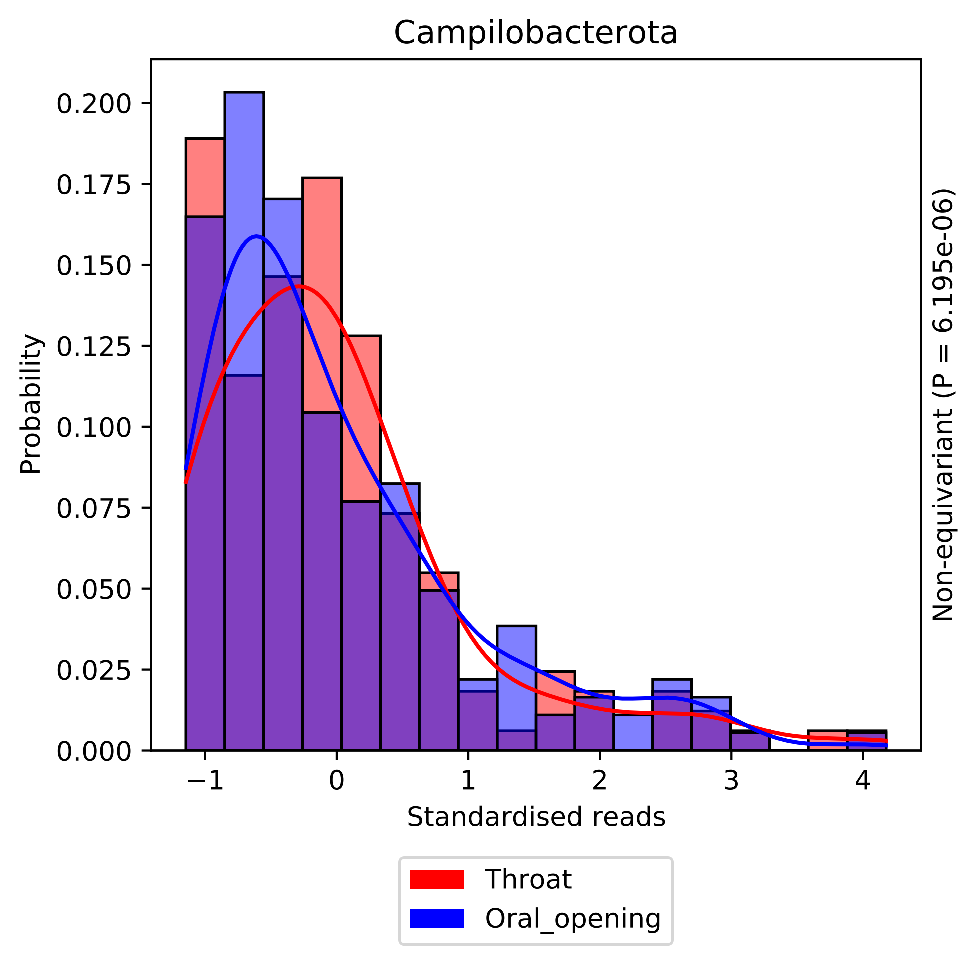

Campilobacterota
Probability
0.000
0.025
0.050
0.075
0.100
0.125
0.150
0.175
0.200
−1
0
1
2
3
4
Standardised reads
Non-equivariant (P = 6.195e-06)
Throat
Oral_opening

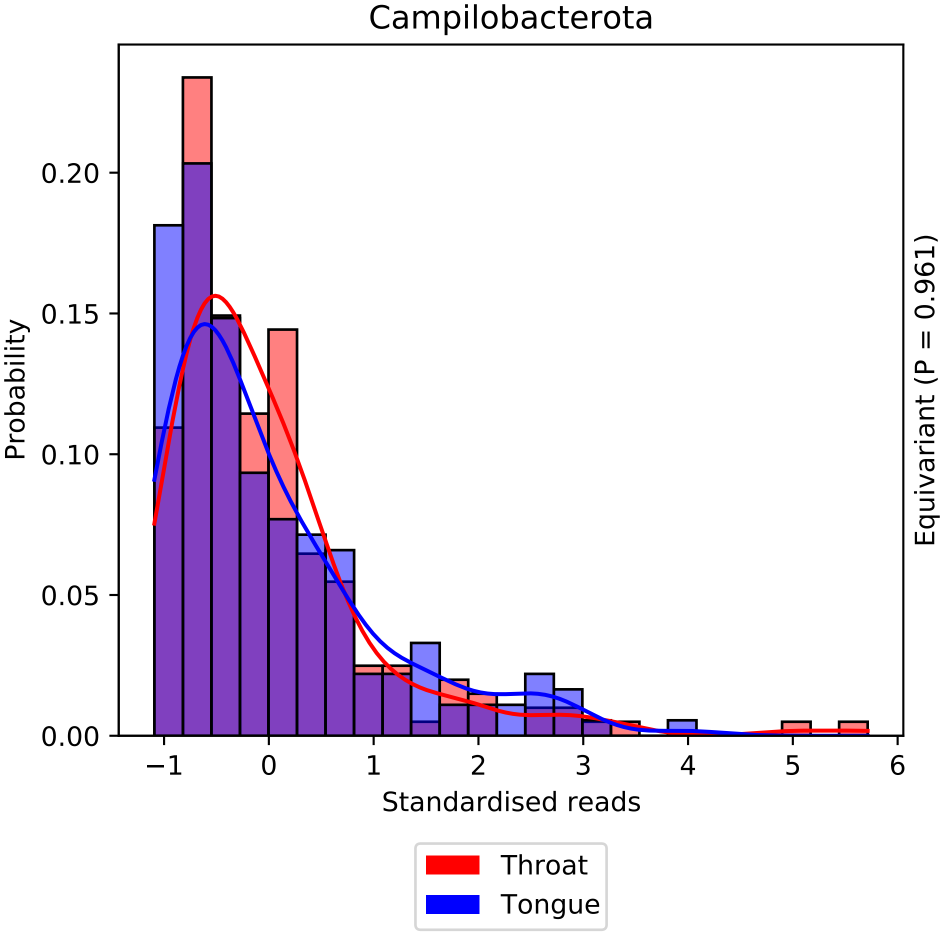

Campilobacterota
Probability
0.00
0.05
0.10
0.15
0.20
−1
0
1
2
3
4
5
6
Standardised reads
Equivariant (P = 0.961)
Throat
Tongue

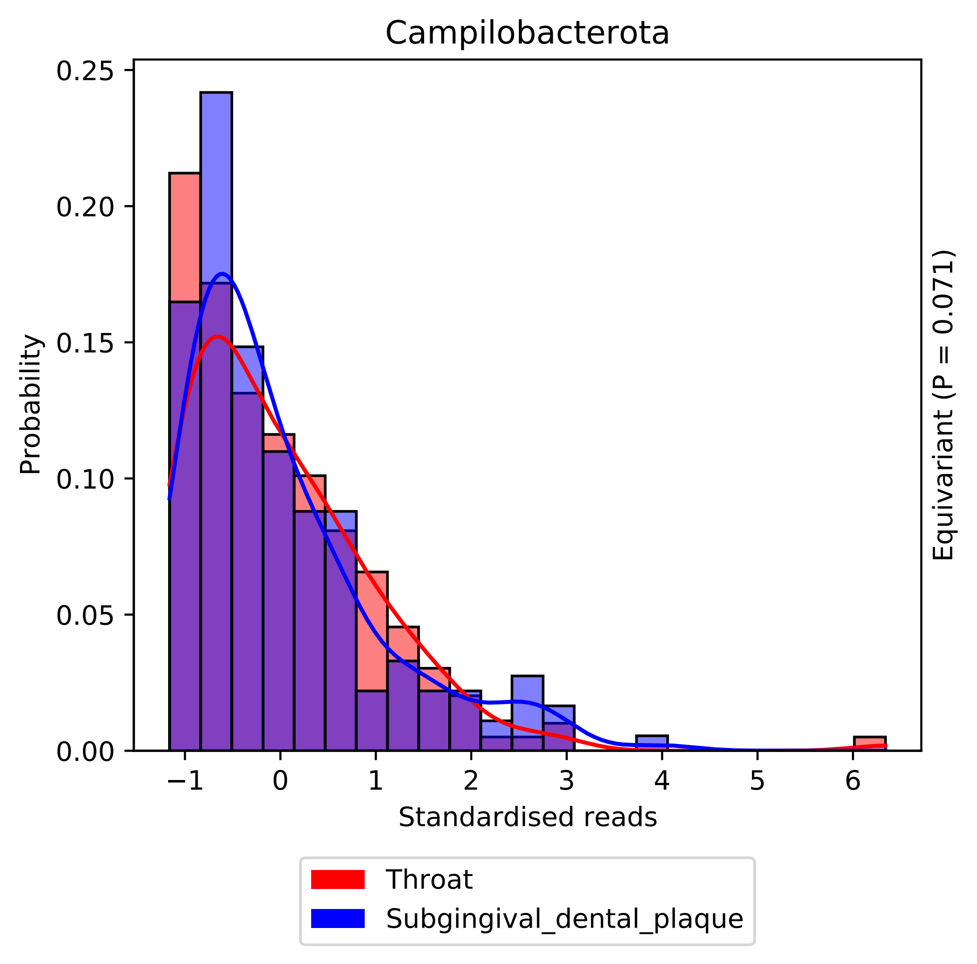

Campilobacterota
Probability
Standardised reads
Equivariant (P = 0.071)
0.00
0.05
0.10
0.15
0.20
0.25
−1
0
1
2
3
4
5
6
Throat
Subgingival_dental_plaque

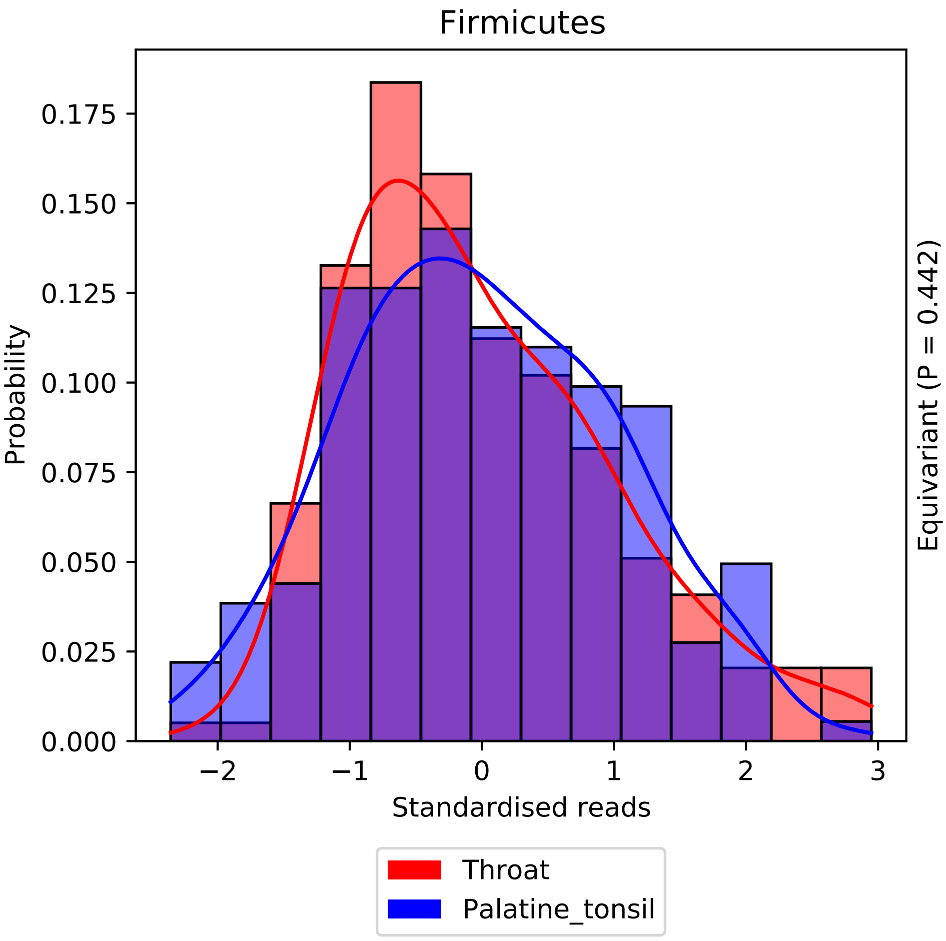

Firmicutes
Probability
0.000
0.025
0.050
0.075
0.100
0.125
0.150
0.175
−2
−1
0
1
2
3
Standardised reads
Equivariant (P = 0.442)
Throat
Palatine_tonsil

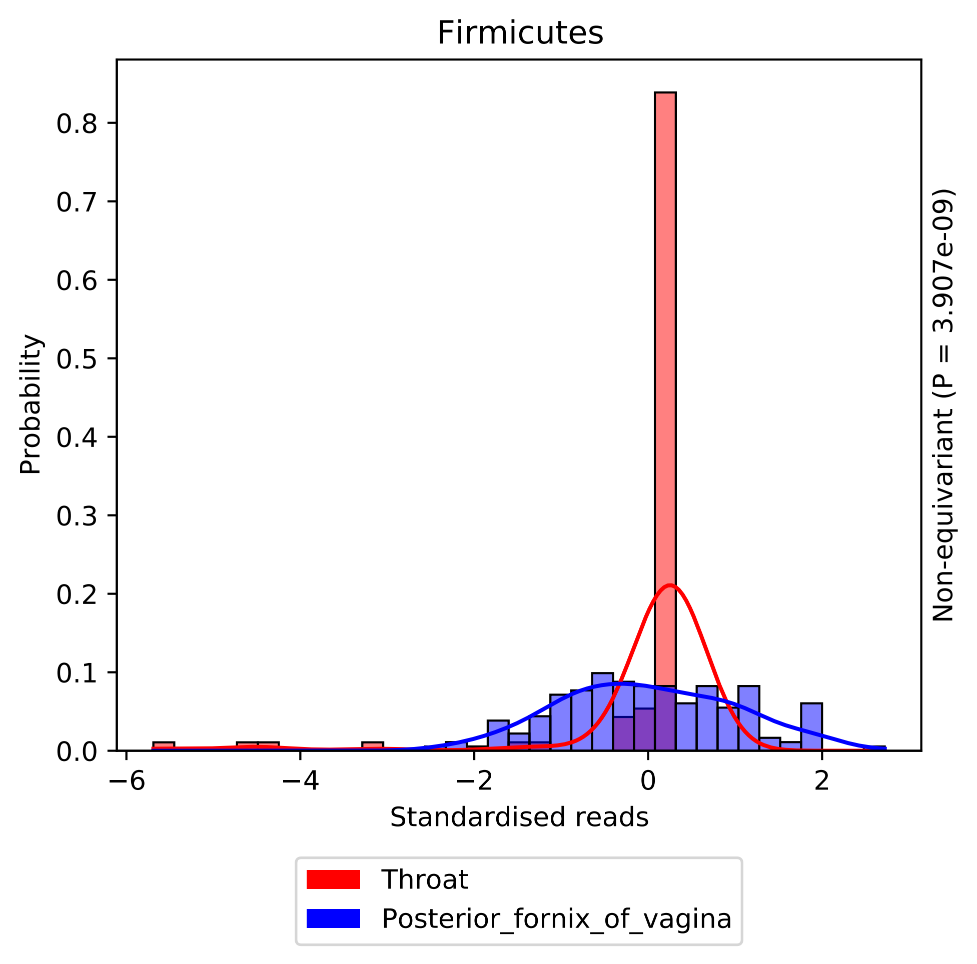

Firmicutes
Probability
0.0
0.1
0.2
0.3
0.4
0.5
0.6
0.7
0.8
−6
−4
−2
0
2
Standardised reads
Non-equivariant (P = 3.907e-09)
Throat
Posterior_fornix_of_vagina

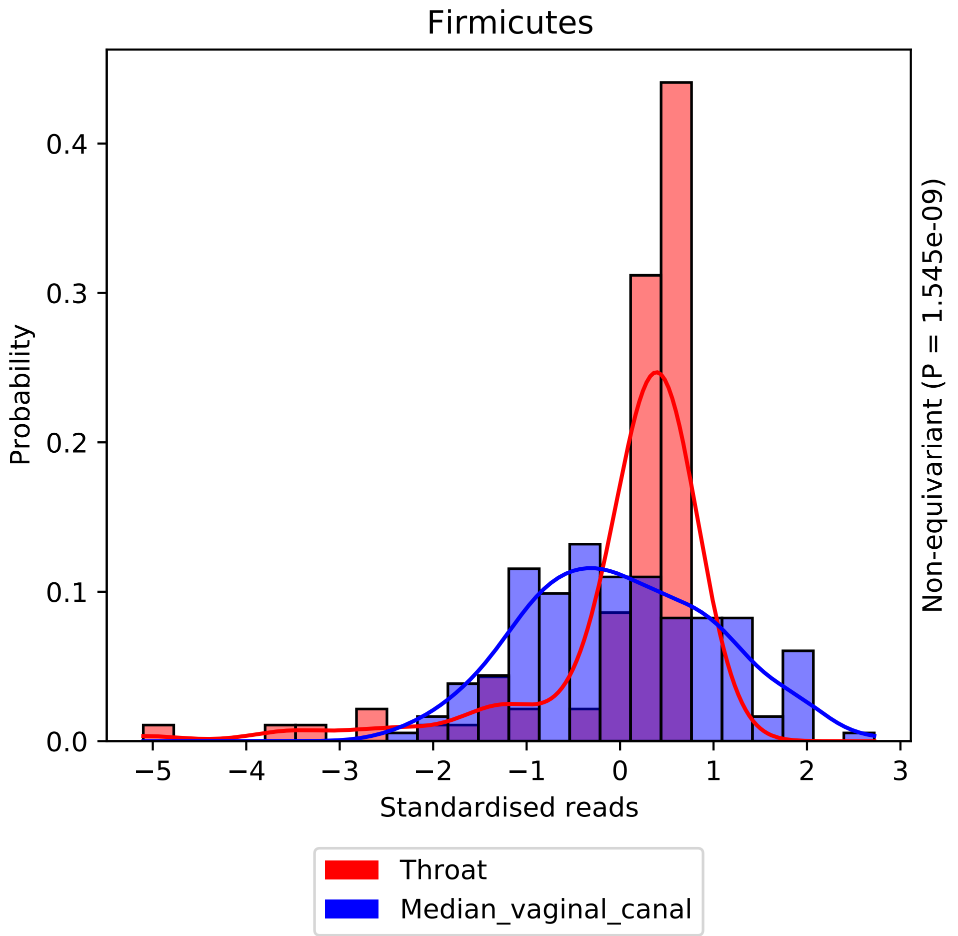

Firmicutes
Probability
Standardised reads
Non-equivariant (P = 1.545e-09)
0.0
0.1
0.2
0.3
0.4
−5
−4
−3
−2
−1
0
1
2
3
Throat
Median_vaginal_canal

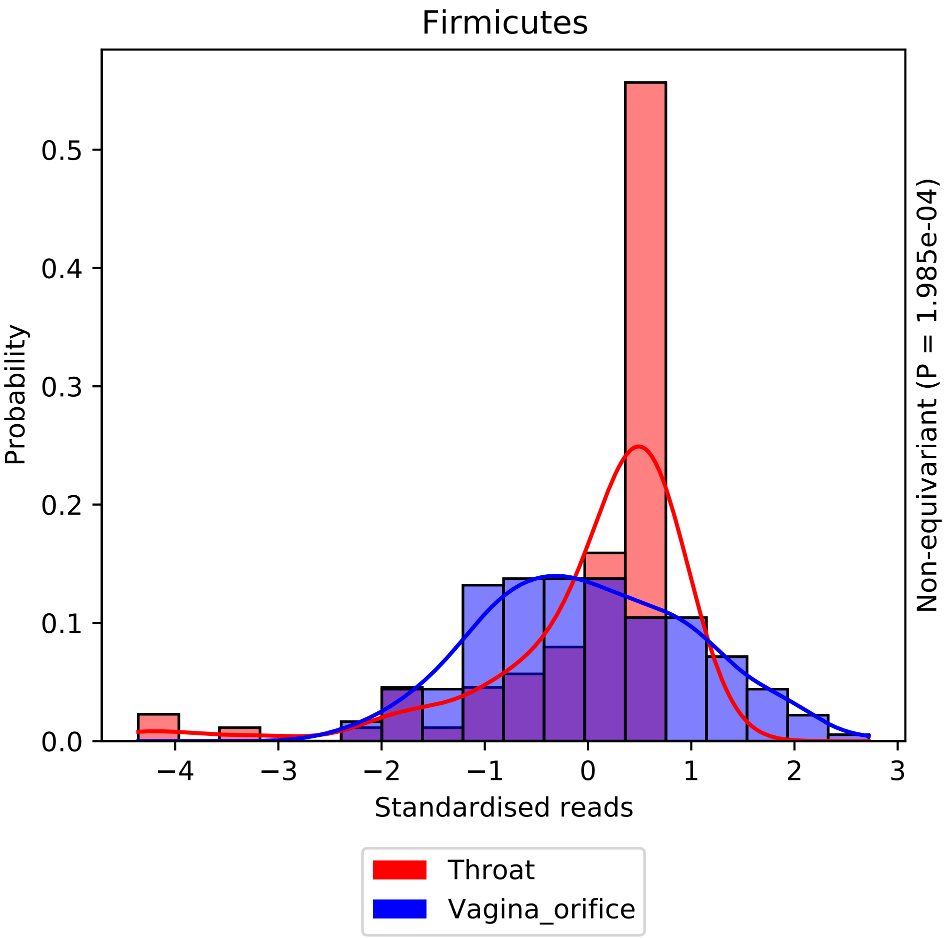
Firmicutes
Probability
0.0
0.1
0.2
0.3
0.4
0.5
−4
−3
−2
−1
0
1
2
3
Standardised reads
Non-equivariant (P = 1.985e-04)
Throat
Vagina_orifice

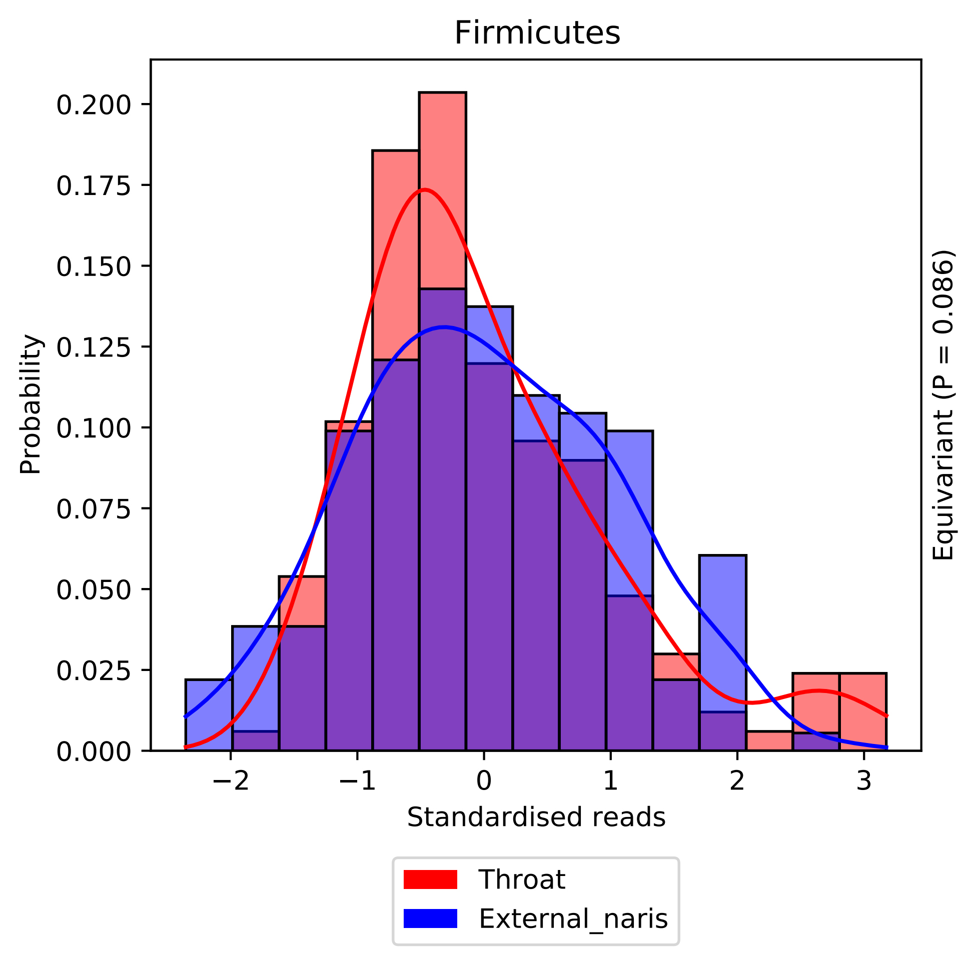

Firmicutes
Probability
0.000
0.025
0.050
0.075
0.100
0.125
0.150
0.175
0.200
−2
−1
0
1
2
3
Standardised reads
Equivariant (P = 0.086)
Throat
External_naris

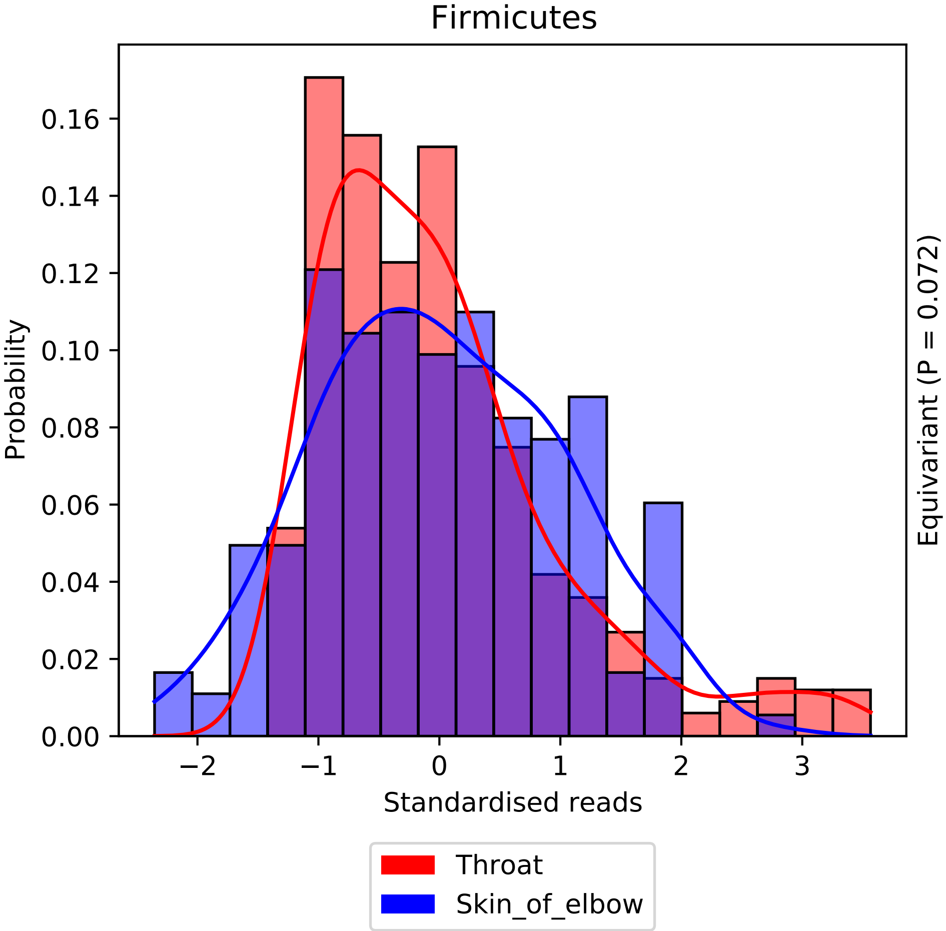
Firmicutes
Probability
0.00
0.02
0.04
0.06
0.08
0.10
0.12
0.14
0.16
−2
−1
0
1
2
3
Standardised reads
Equivariant (P = 0.072)
Throat
Skin_of_elbow

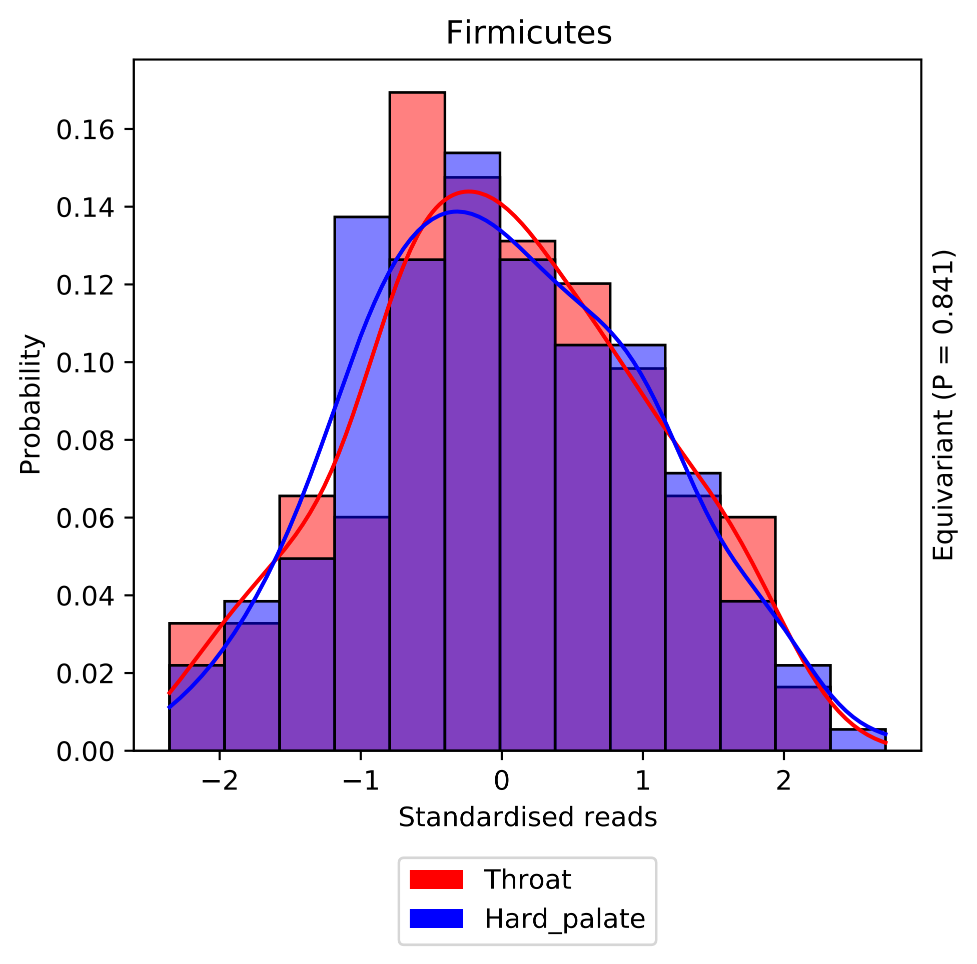

Firmicutes
Probability
0.00
0.02
0.04
0.06
0.08
0.10
0.12
0.14
0.16
−2
−1
0
1
2
Standardised reads
Equivariant (P = 0.841)
Throat
Hard_palate

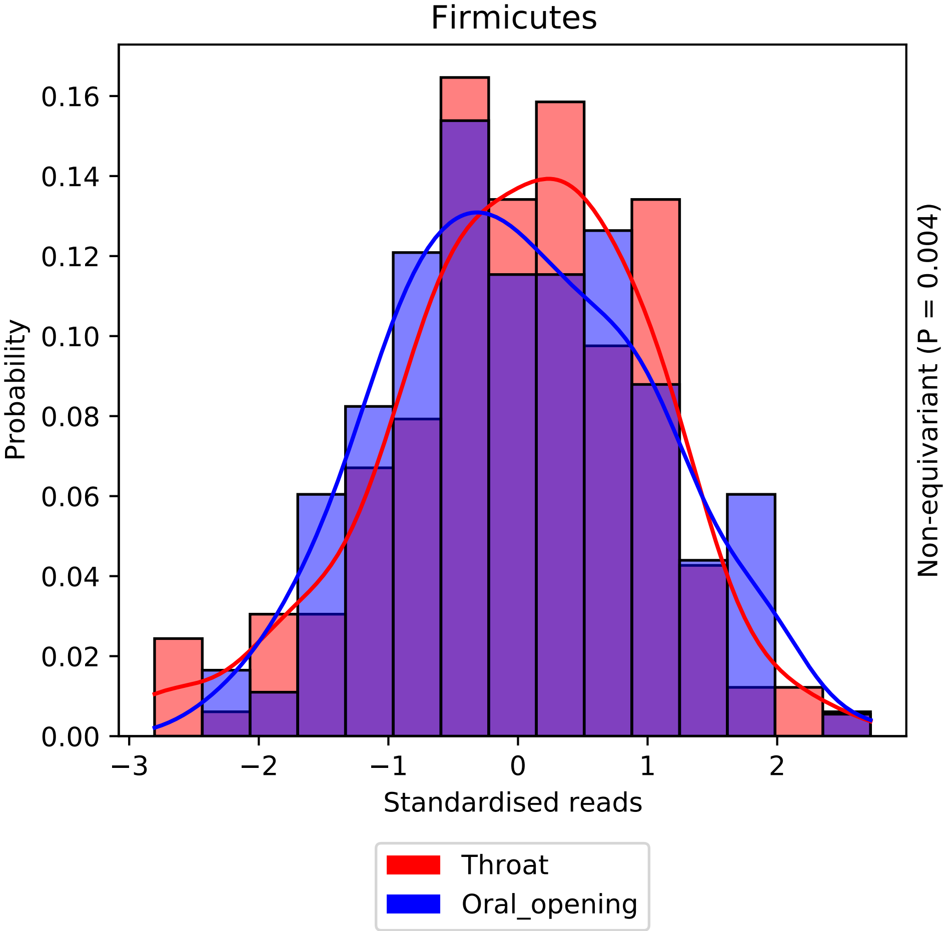
Firmicutes
Probability
0.00
0.02
0.04
0.06
0.08
0.10
0.12
0.14
0.16
−3
−2
−1
0
1
2
Standardised reads
Non-equivariant (P = 0.004)
Throat
Oral_opening

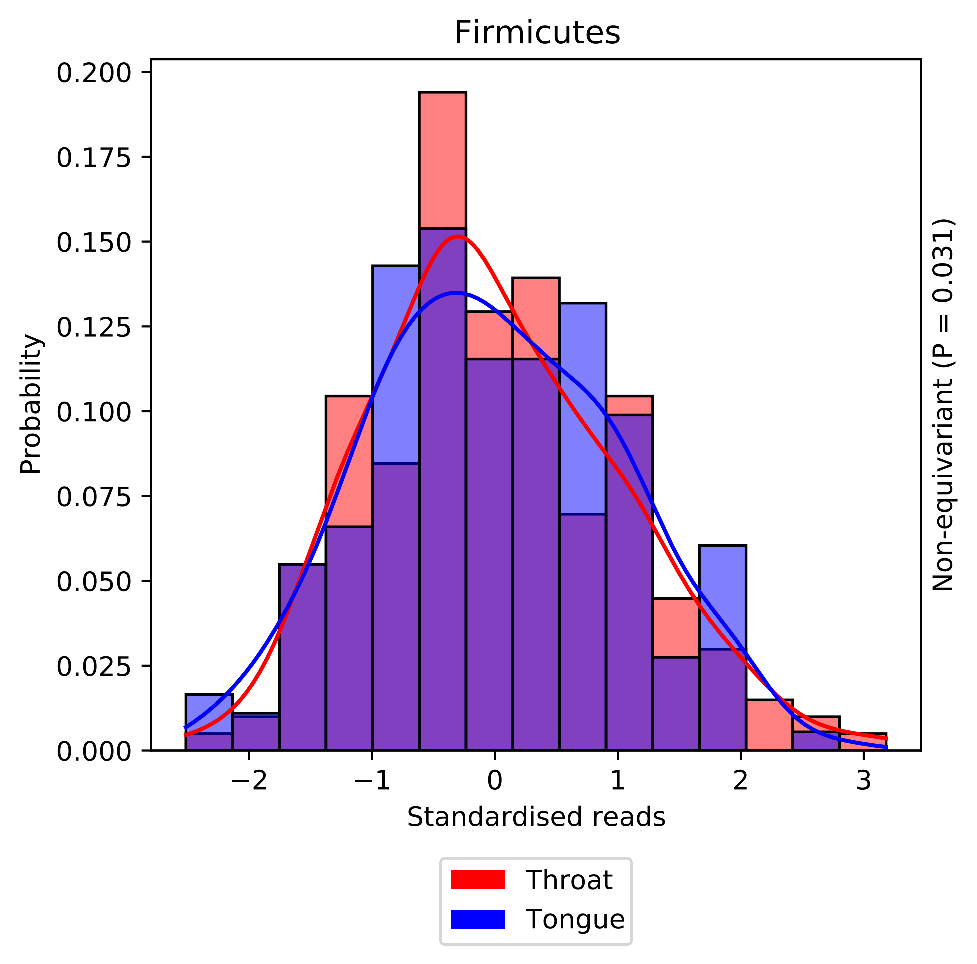

Firmicutes
0.200
0.175
0.150
0.125
0.100
0.075
0.050
0.025
0.000
Probability
−2
−1
0
1
2
3
Standardised reads
Non-equivariant (P = 0.031)
Throat
Tongue

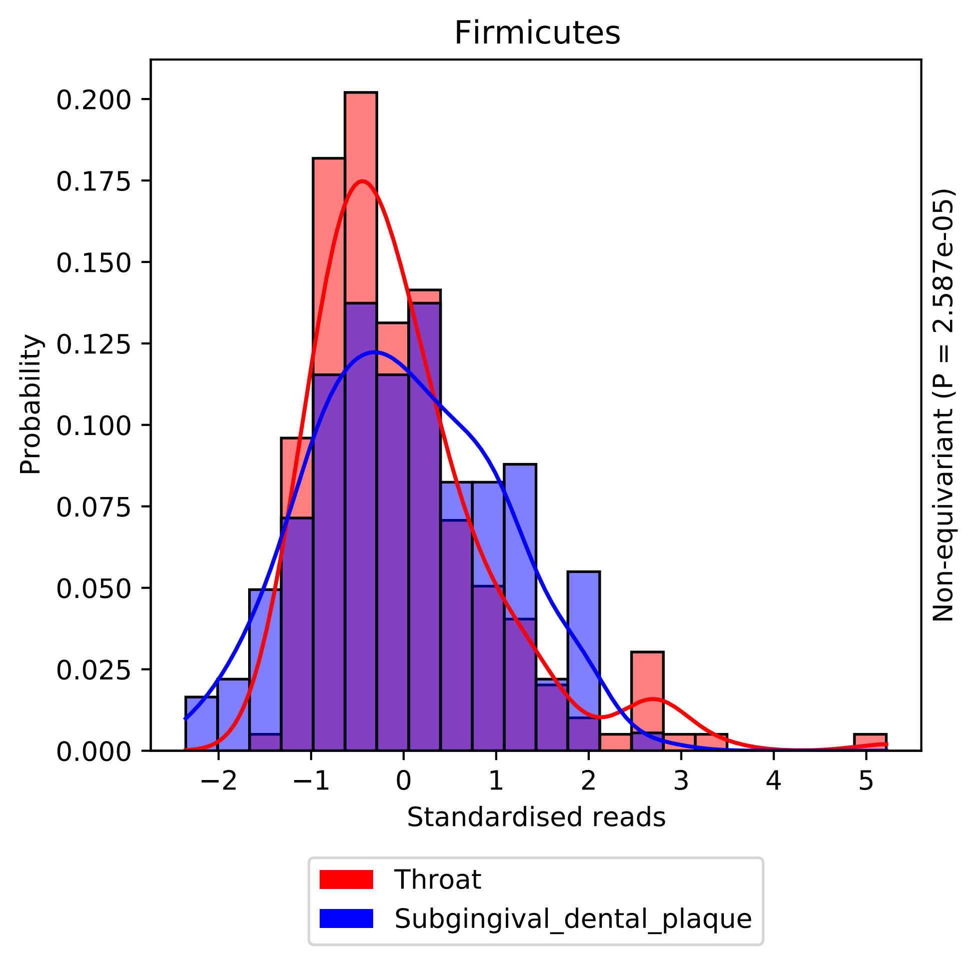

Firmicutes
Probability
0.200
0.175
0.150
0.125
0.100
0.075
0.050
0.025
0.000
−2
−1
0
1
2
3
4
5
Standardised reads
Non-equivariant (P = 2.587e-05)
Throat
Subgingival_dental_plaque

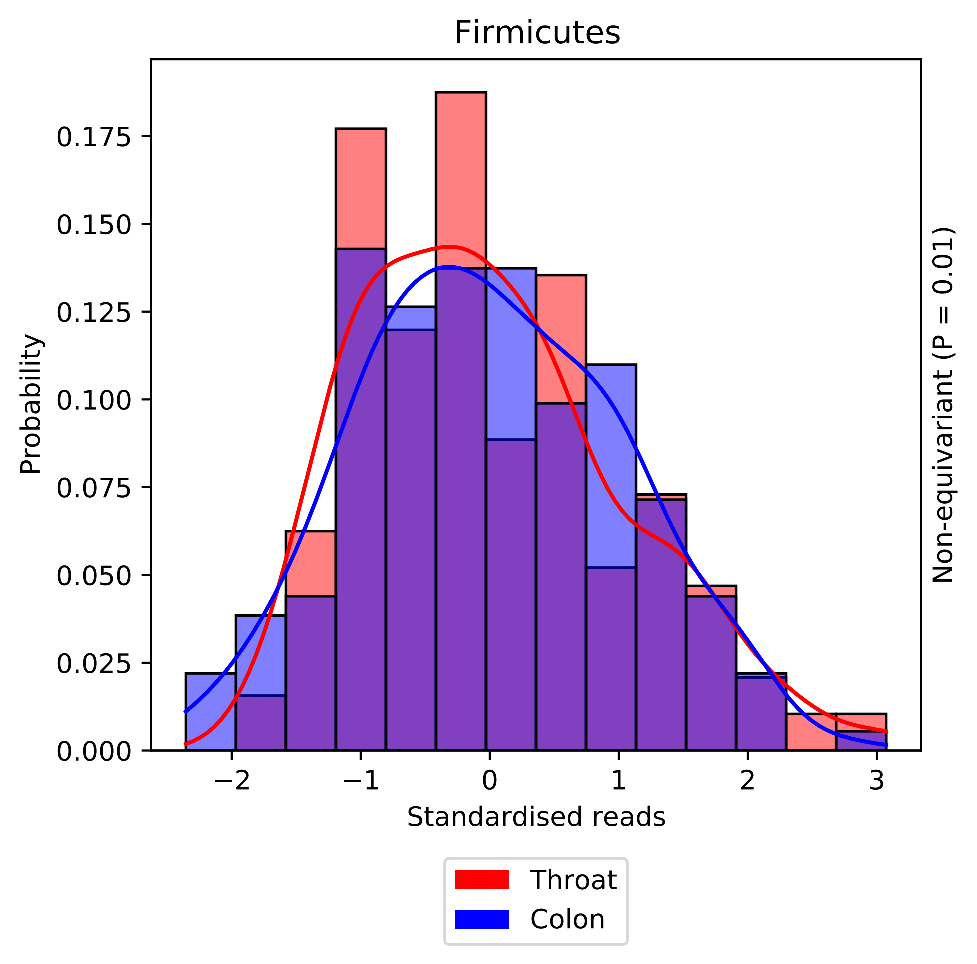

Firmicutes
Probability
0.000
0.025
0.050
0.075
0.100
0.125
0.150
0.175
−2
−1
0
1
2
3
Standardised reads
Non-equivariant (P = 0.01)
Throat
Colon

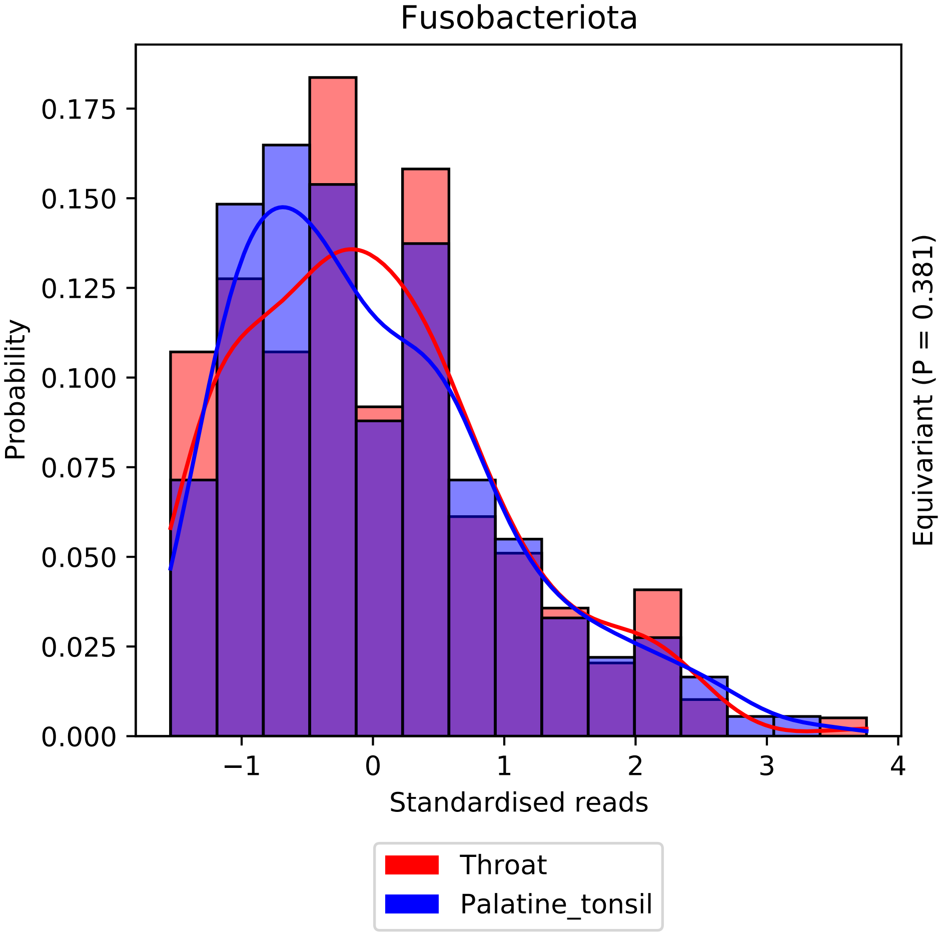
Fusobacteriota
Probability
0.000
0.025
0.050
0.075
0.100
0.125
0.150
0.175
−1
0
1
2
3
4
Standardised reads
Equivariant (P = 0.381)
Throat
Palatine_tonsil

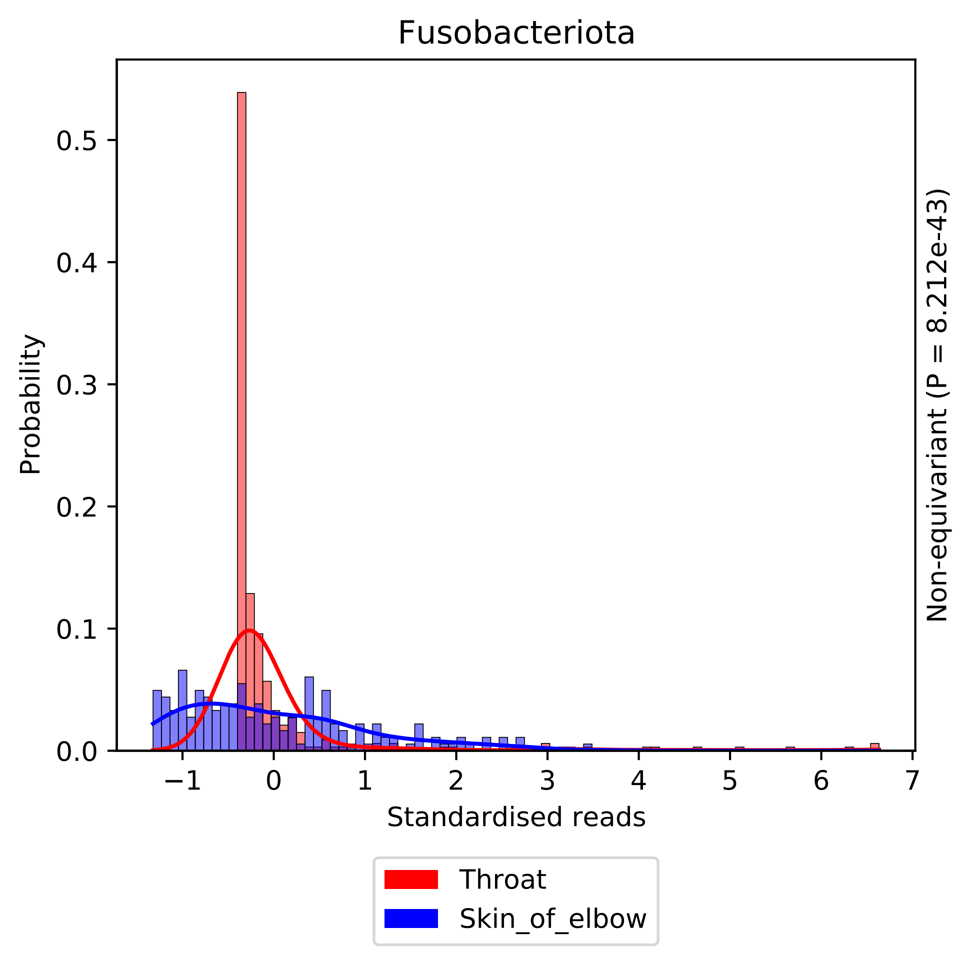

Fusobacteriota
Probability
Standardised reads
Non-equivariant (P = 8.212e-43)
0.0
0.1
0.2
0.3
0.4
0.5
−1
0
1
2
3
4
5
6
7
Throat
Skin_of_elbow

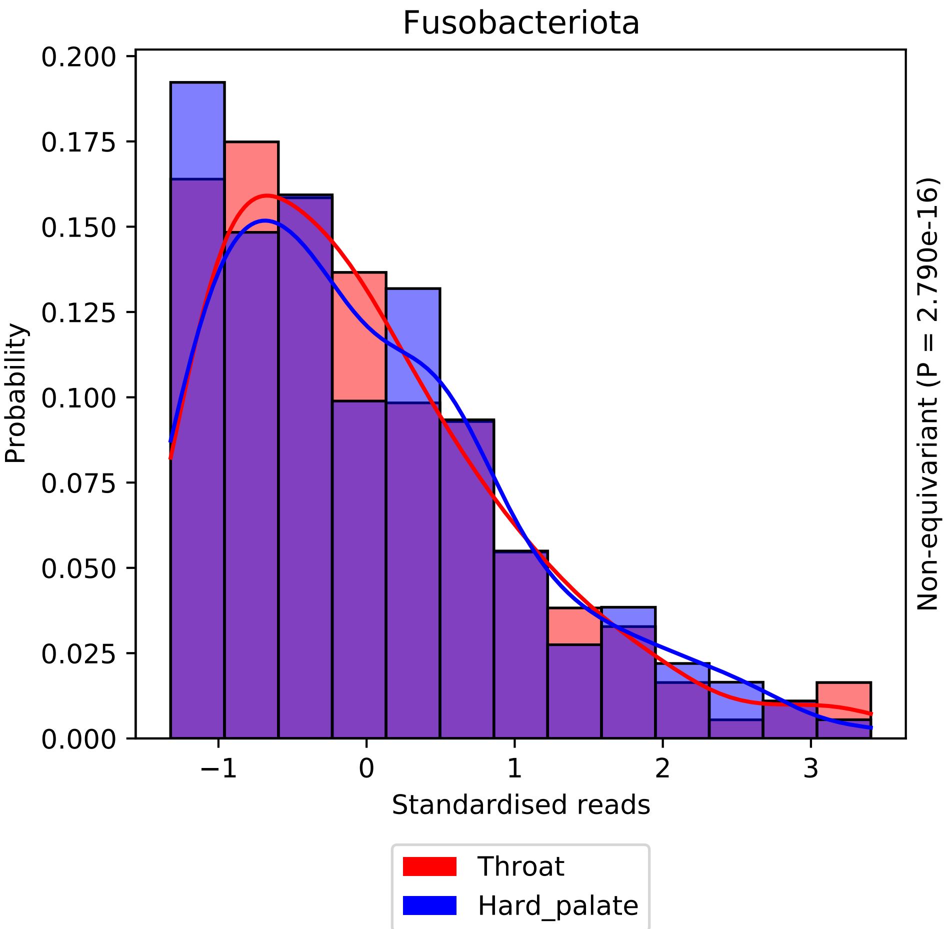
Fusobacteriota
Probability
0.200
0.175
0.150
0.125
0.100
0.075
0.050
0.025
0.000
−1
0
1
2
3
Standardised reads
Non-equivariant (P = 2.790e-16)
Throat
Hard_palate

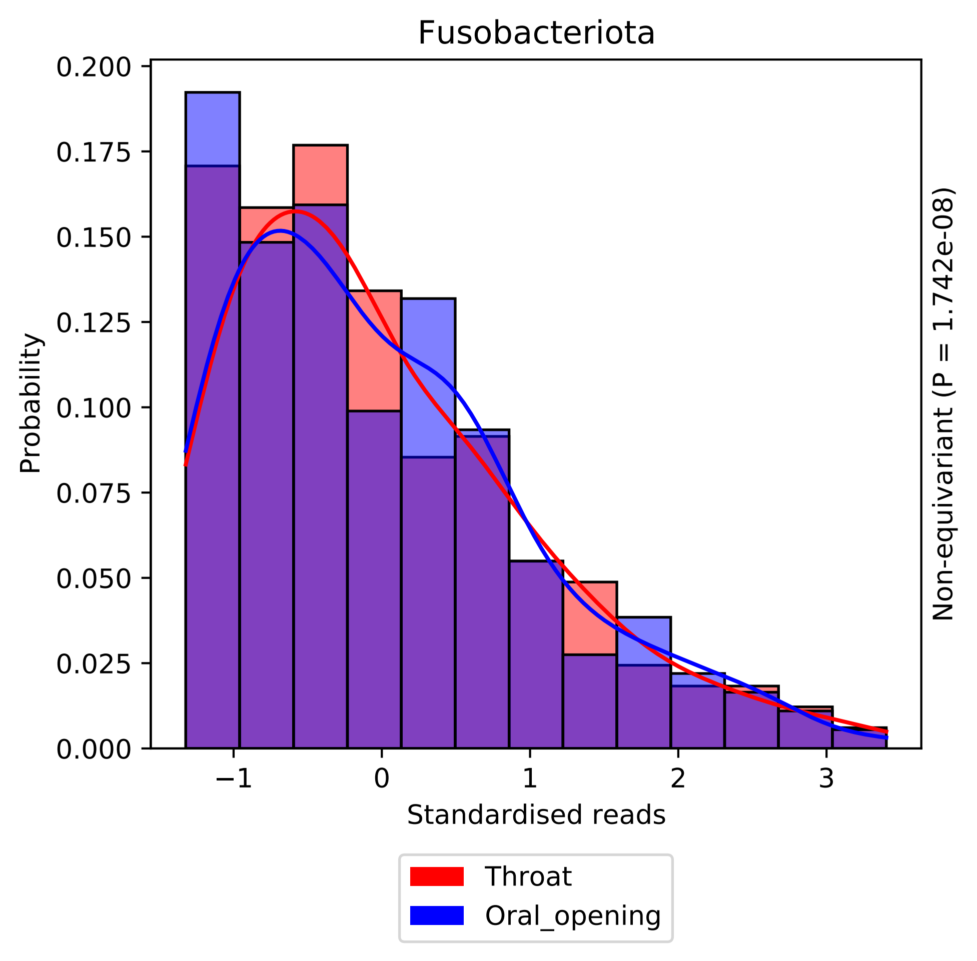
Fusobacteriota
0.200
0.175
0.150
0.125
0.100
0.075
0.050
0.025
0.000
Probability
−1
0
1
2
3
Standardised reads
Non-equivariant (P = 1.742e-08)
Throat
Oral_opening

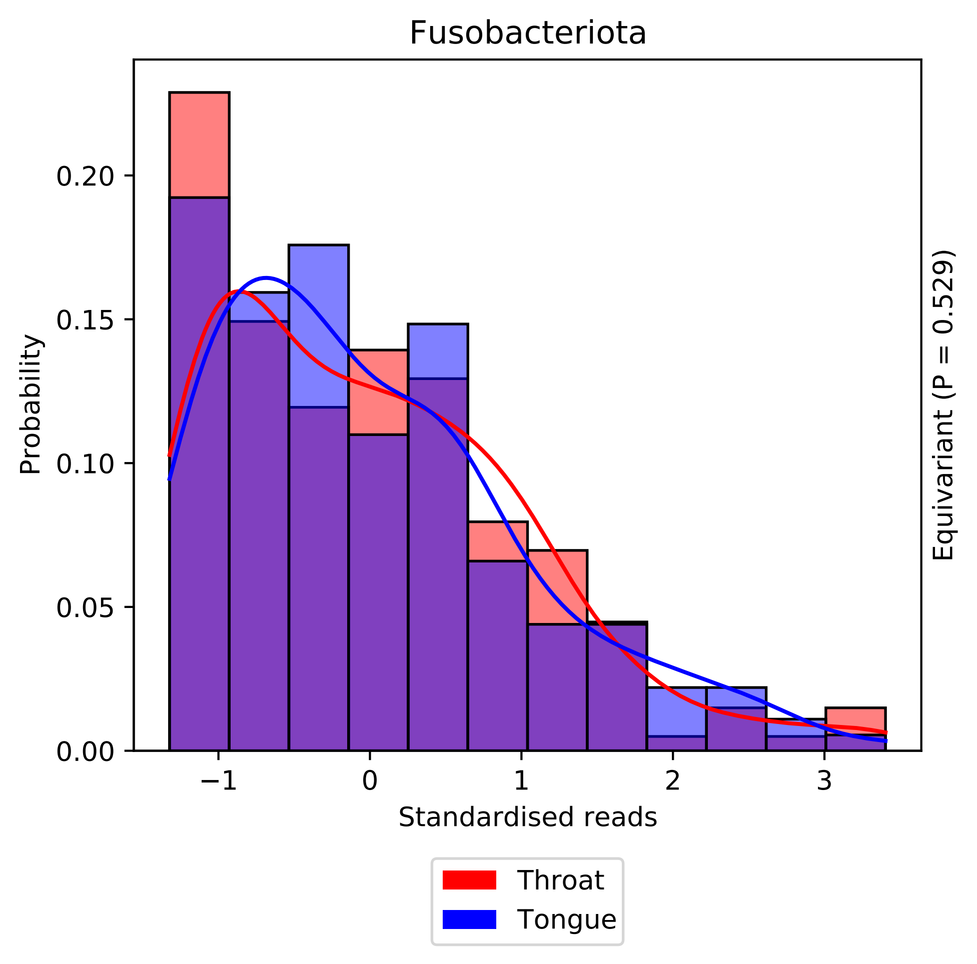

Fusobacteriota
Probability
0.20
0.15
0.10
0.05
0.00
−1
0
1
2
3
Standardised reads
Equivariant (P = 0.529)
Throat
Tongue

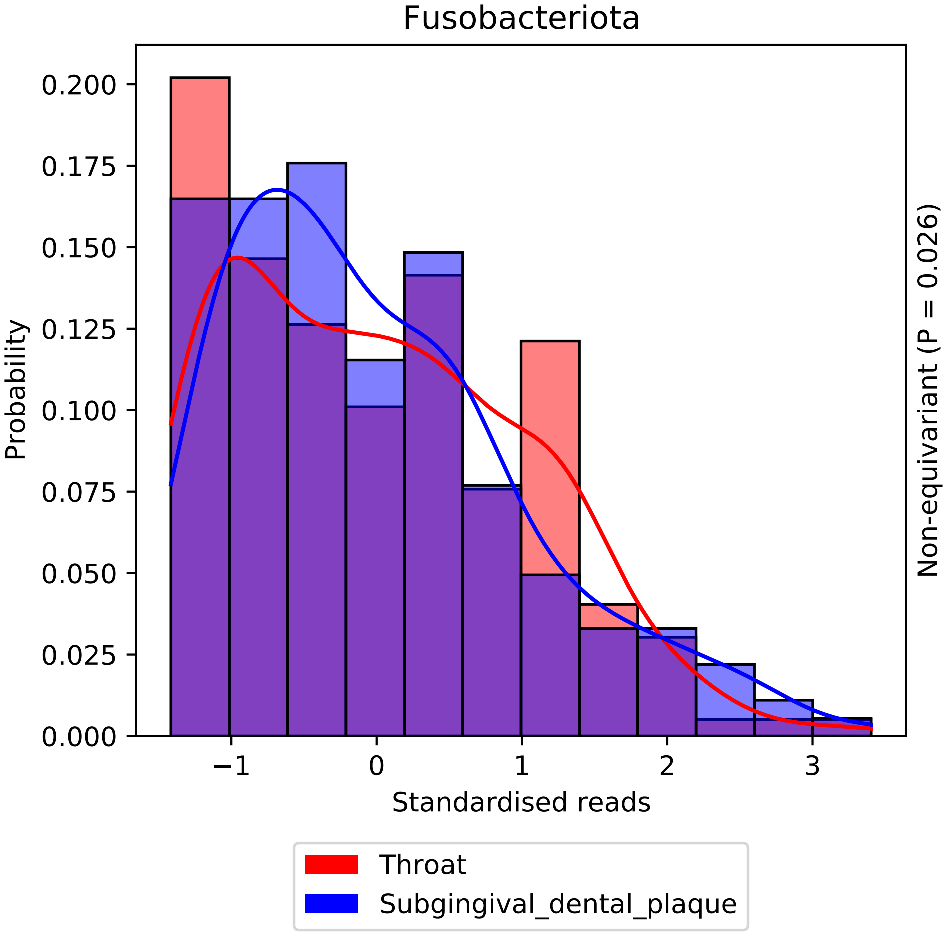

Fusobacteriota
Probability
0.200
0.175
0.150
0.125
0.100
0.075
0.050
0.025
0.000
−1
0
1
2
3
Standardised reads
Non-equivariant (P = 0.026)
Throat
Subgingival_dental_plaque

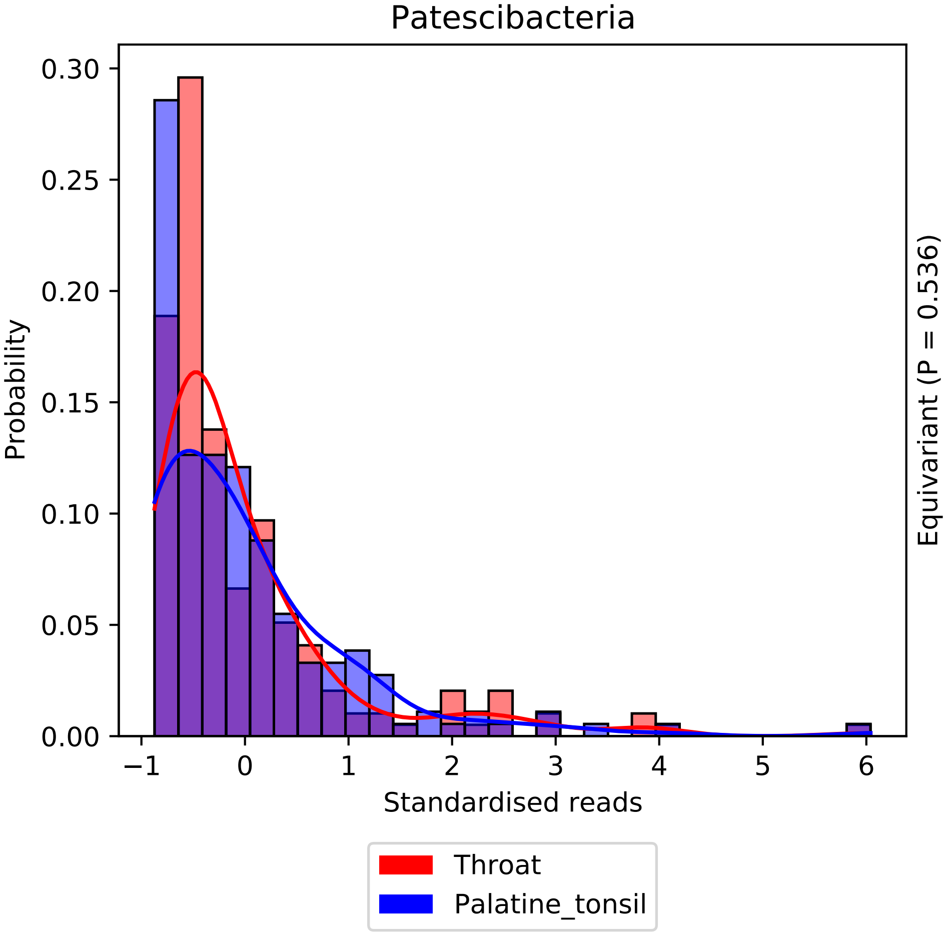
Patescibacteria
Probability
0.30
0.25
0.20
0.15
0.10
0.05
0.00
−1
0
1
2
3
4
5
6
Standardised reads
Equivariant (P = 0.536)
Throat
Palatine_tonsil

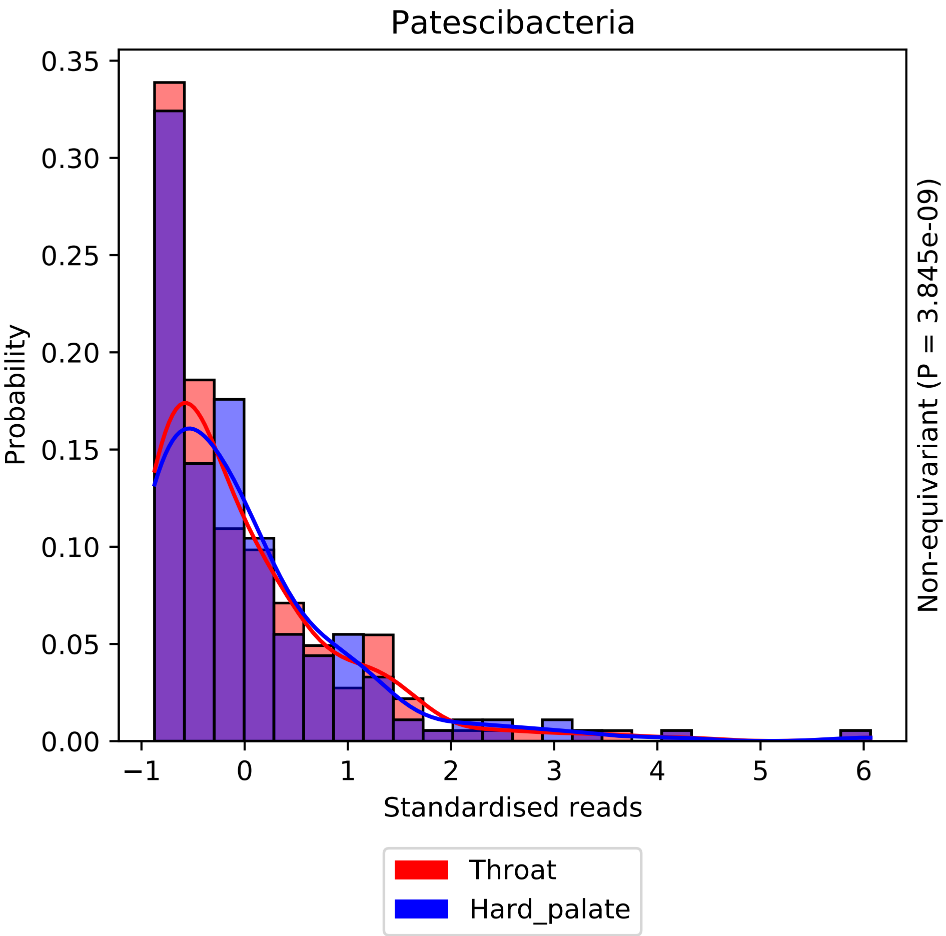
Patescibacteria
Probability
Standardised reads
Non-equivariant (P = 3.845e-09)
Throat
Hard_palate
0.00
0.05
0.10
0.15
0.20
0.25
0.30
0.35
−1
0
1
2
3
4
5
6

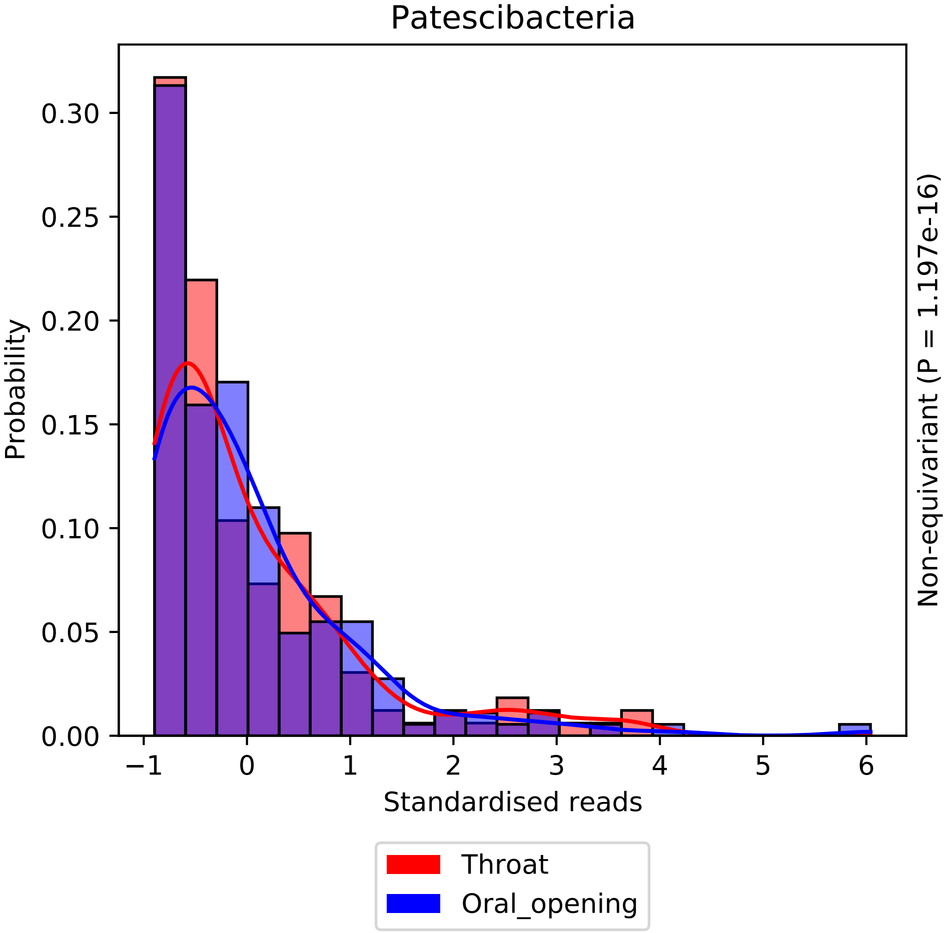
Patescibacteria
Probability
0.00
0.05
0.10
0.15
0.20
0.25
0.30
−1
0
1
2
3
4
5
6
Standardised reads
Non-equivariant (P = 1.197e-16)
Throat
Oral_opening

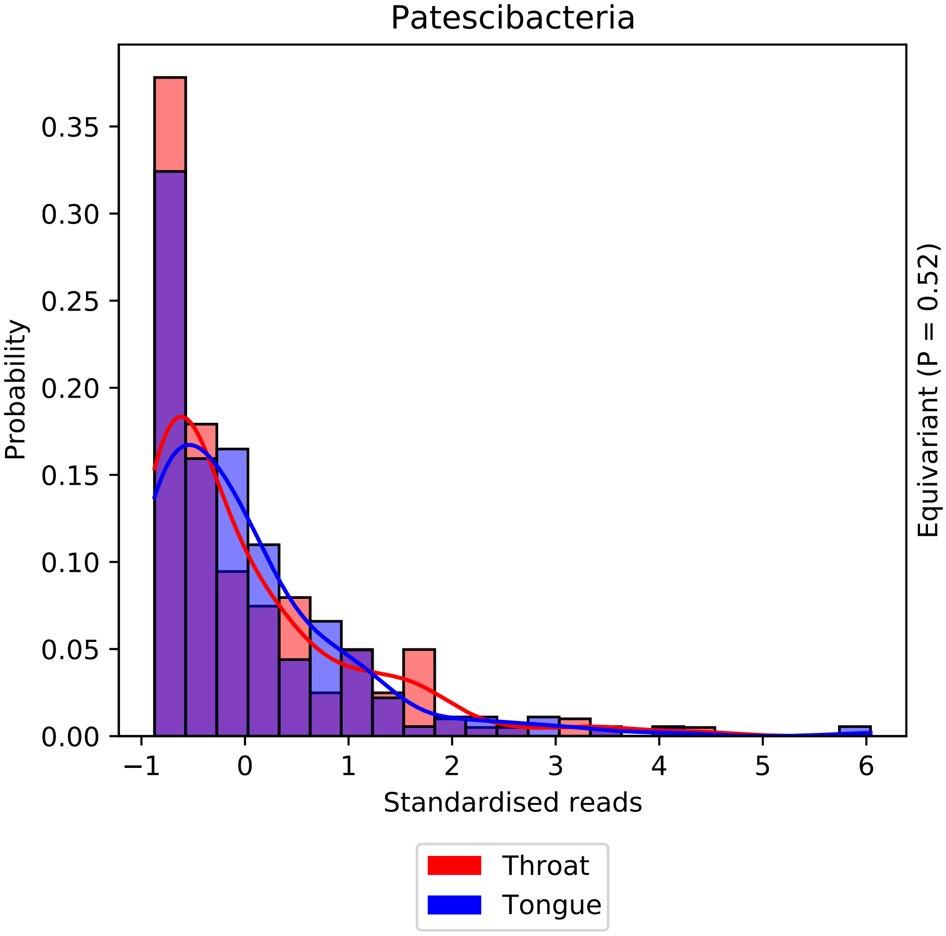

Patescibacteria
Probability
0.00
0.05
0.10
0.15
0.20
0.25
0.30
0.35
−1
0
1
2
3
4
5
6
Standardised reads
Equivariant (P = 0.52)
Throat
Tongue

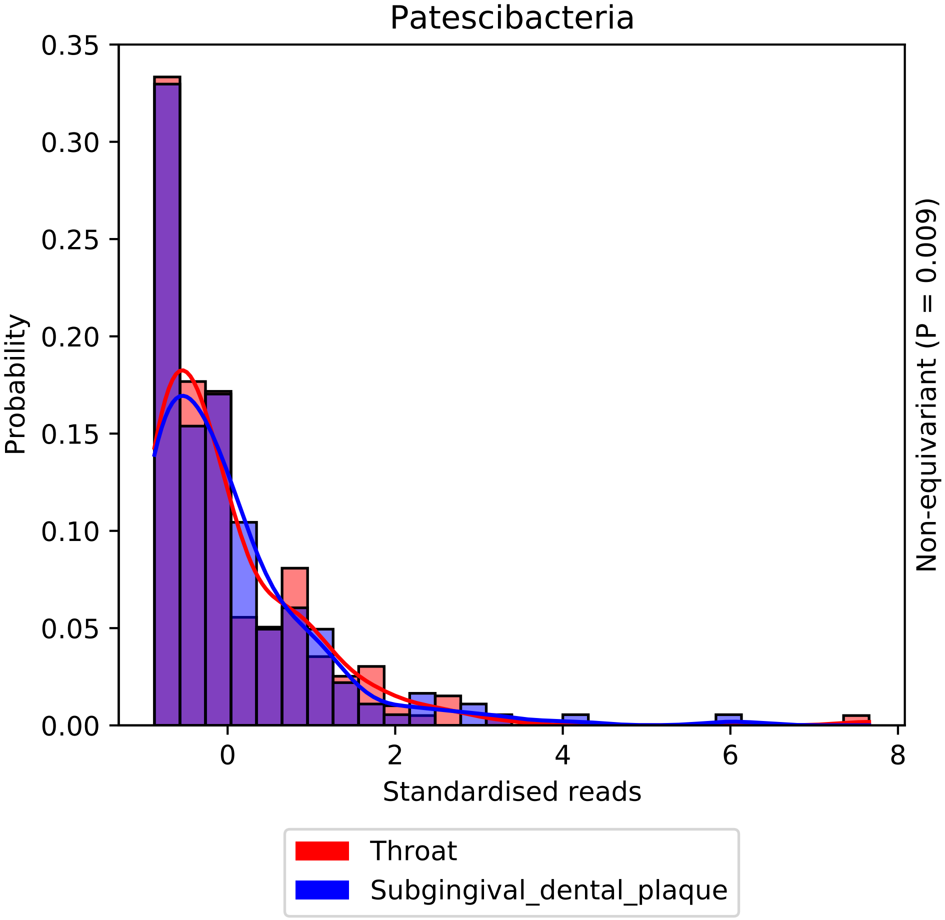
Patescibacteria
Probability
Standardised reads
Non-equivariant (P = 0.009)
0.35
0.30
0.25
0.20
0.15
0.10
0.05
0.00
0
2
4
6
8
Throat
Subgingival_dental_plaque

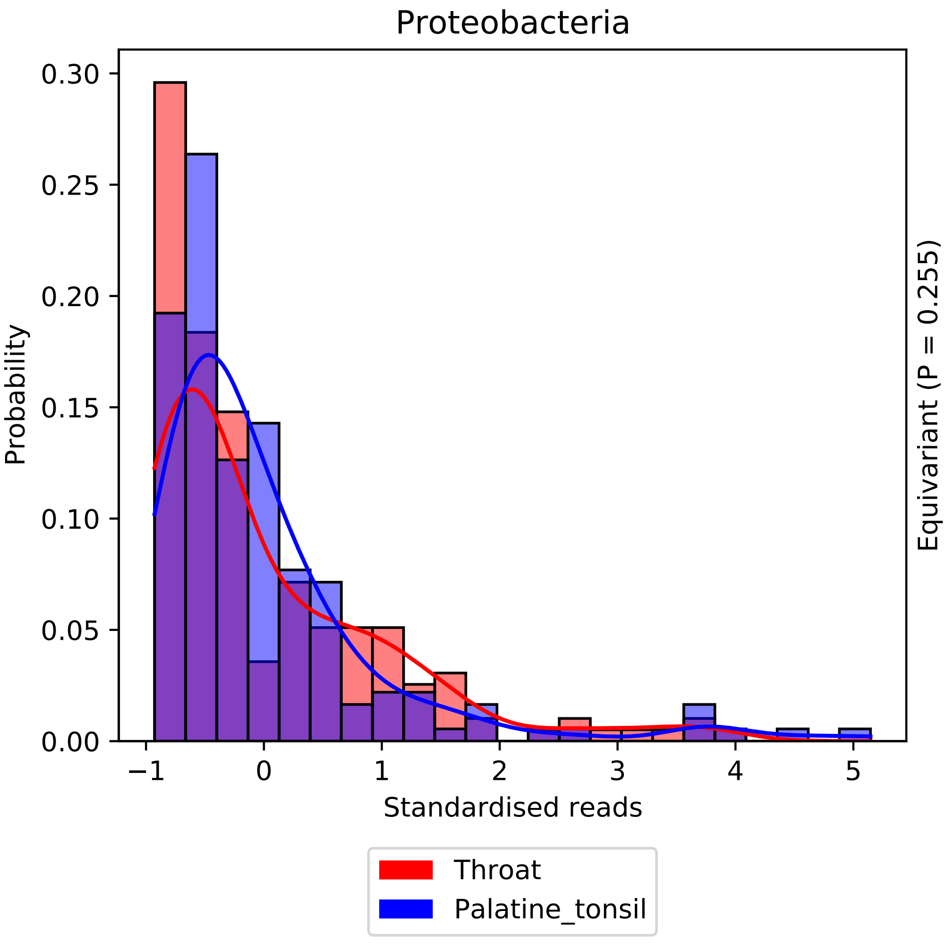
Proteobacteria
0.30
0.25
0.20
0.15
0.10
0.05
0.00
Probability
−1
0
1
2
3
4
5
Standardised reads
Equivariant (P = 0.255)
Throat
Palatine_tonsil

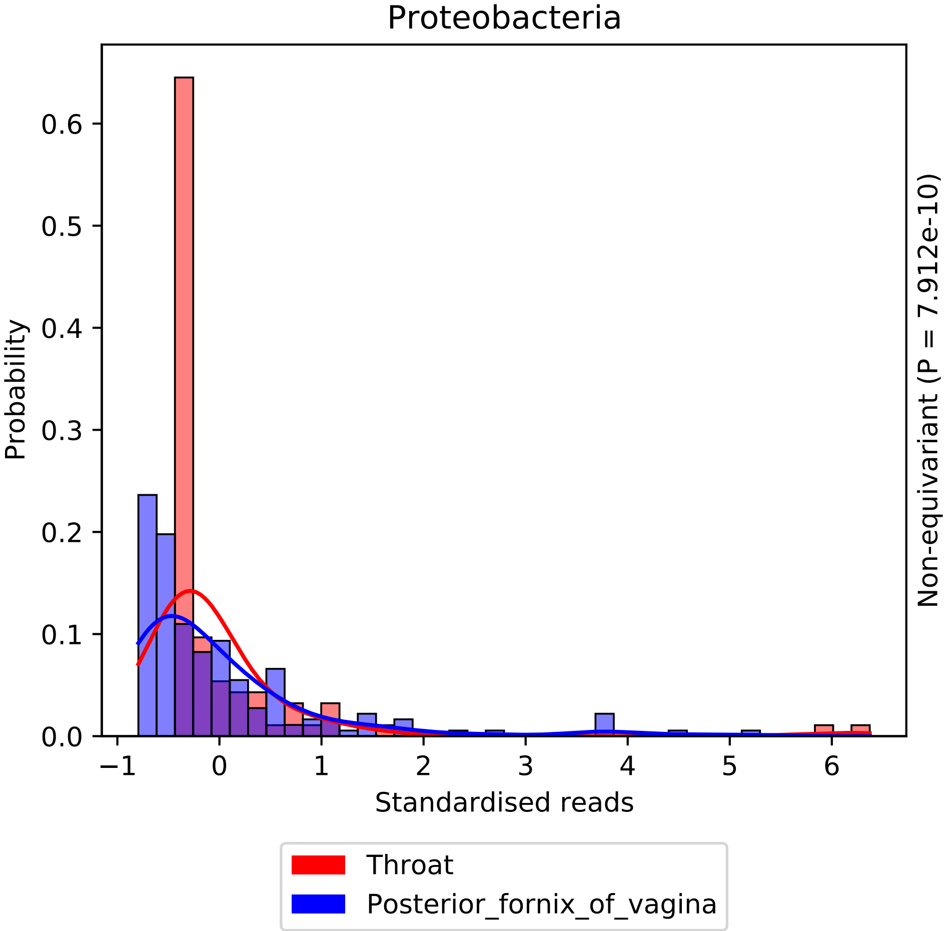

Proteobacteria
Probability
0.0
0.1
0.2
0.3
0.4
0.5
0.6
−1
0
1
2
3
4
5
6
Standardised reads
Non-equivariant (P = 7.912e-10)
Throat
Posterior_fornix_of_vagina

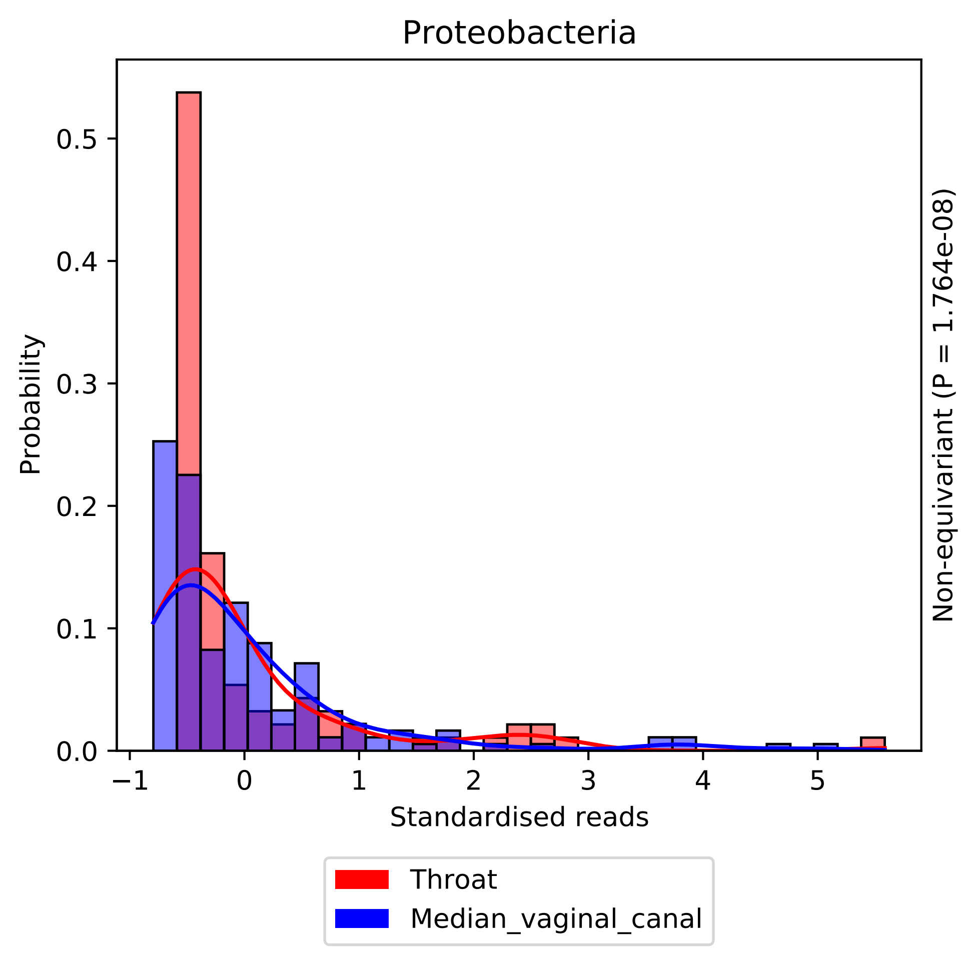
Proteobacteria
Probability
0.0
0.1
0.2
0.3
0.4
0.5
−1
0
1
2
3
4
5
Standardised reads
Non-equivariant (P = 1.764e-08)
Throat
Median_vaginal_canal

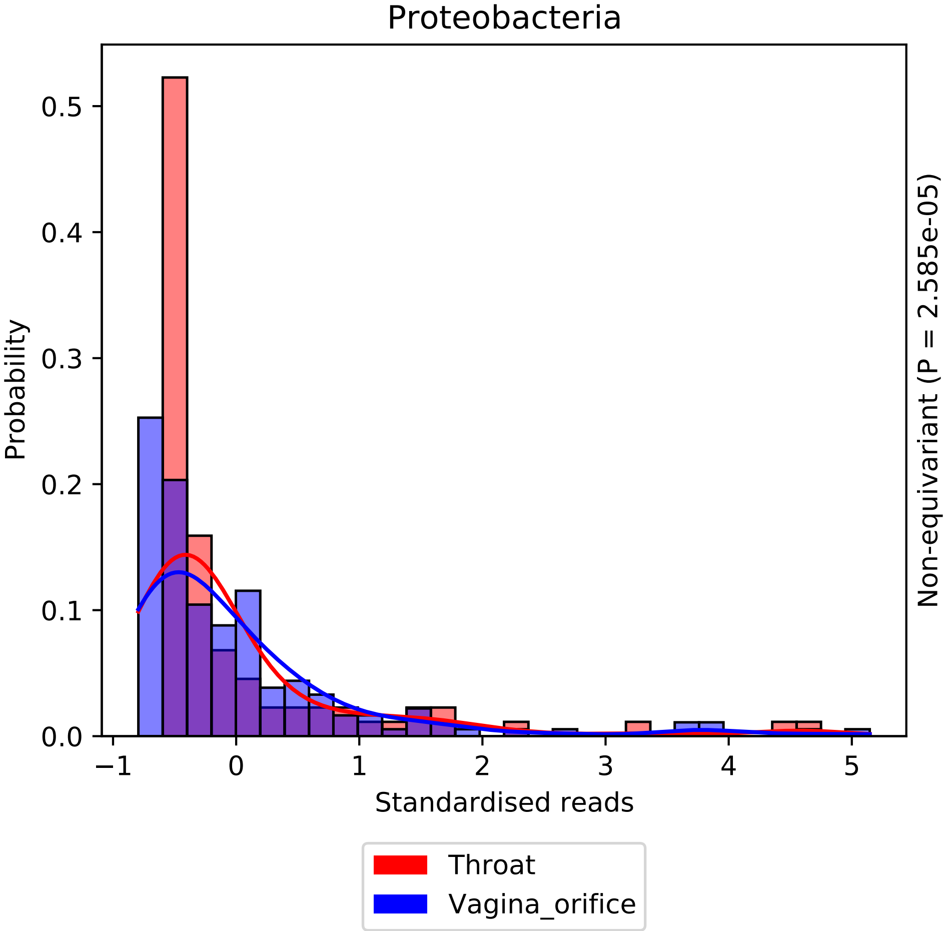

Proteobacteria
Probability
Standardised reads
Non-equivariant (P = 2.585e-05)
0.0
0.1
0.2
0.3
0.4
0.5
−1
0
1
2
3
4
5
Throat
Vagina_orifice

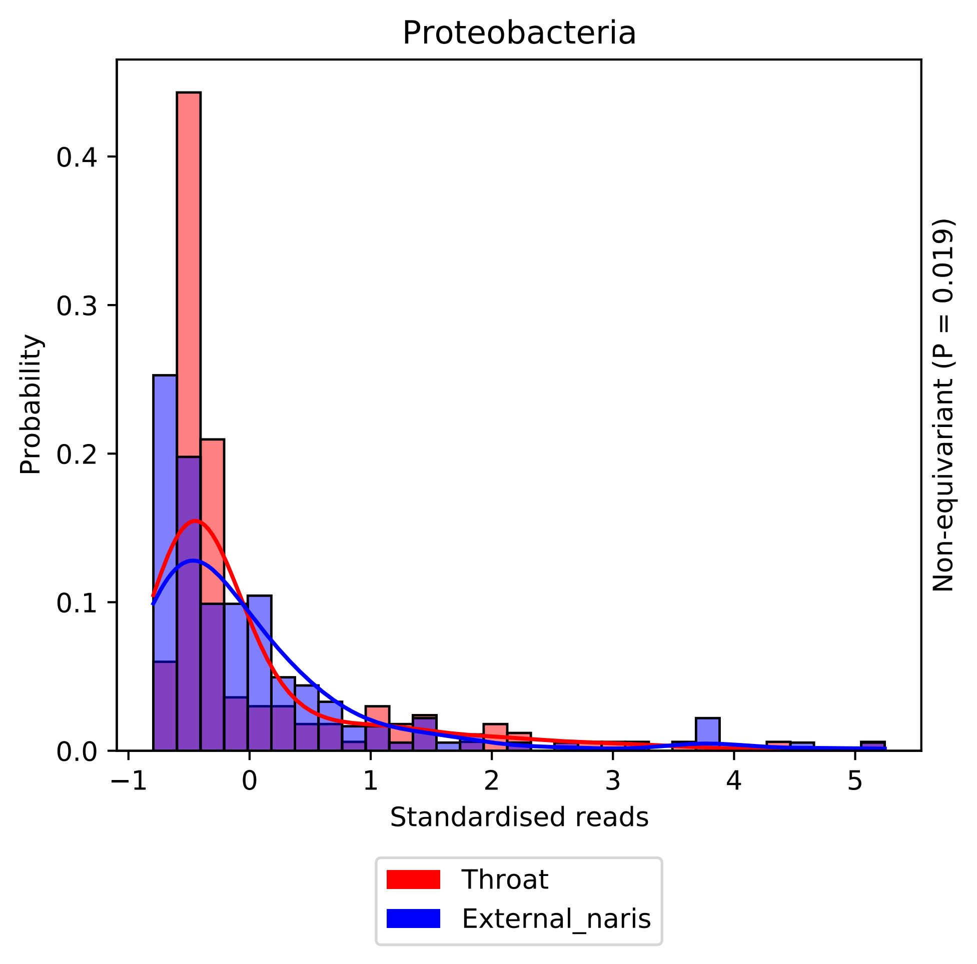

Proteobacteria
Probability
0.0
0.1
0.2
0.3
0.4
−1
0
1
2
3
4
5
Standardised reads
Non-equivariant (P = 0.019)
Throat
External_naris

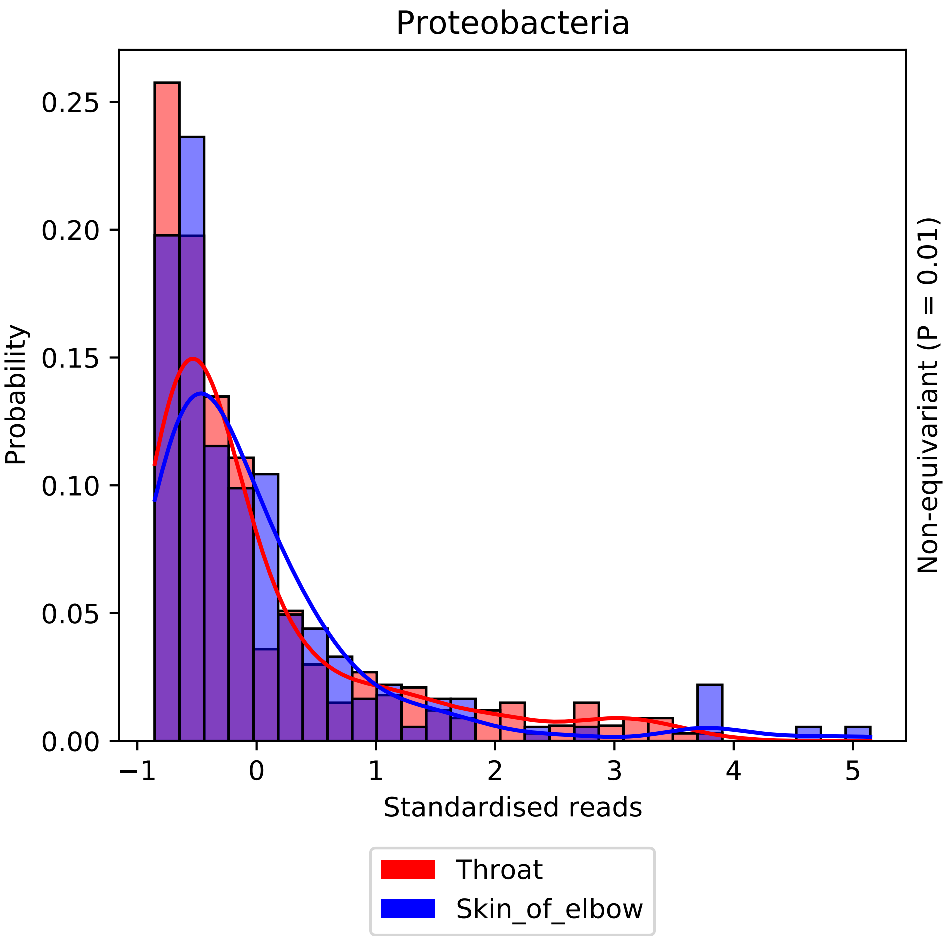

Proteobacteria
Probability
Standardised reads
Non-equivariant (P = 0.01)
0.00
0.05
0.10
0.15
0.20
0.25
−1
0
1
2
3
4
5
Throat
Skin_of_elbow

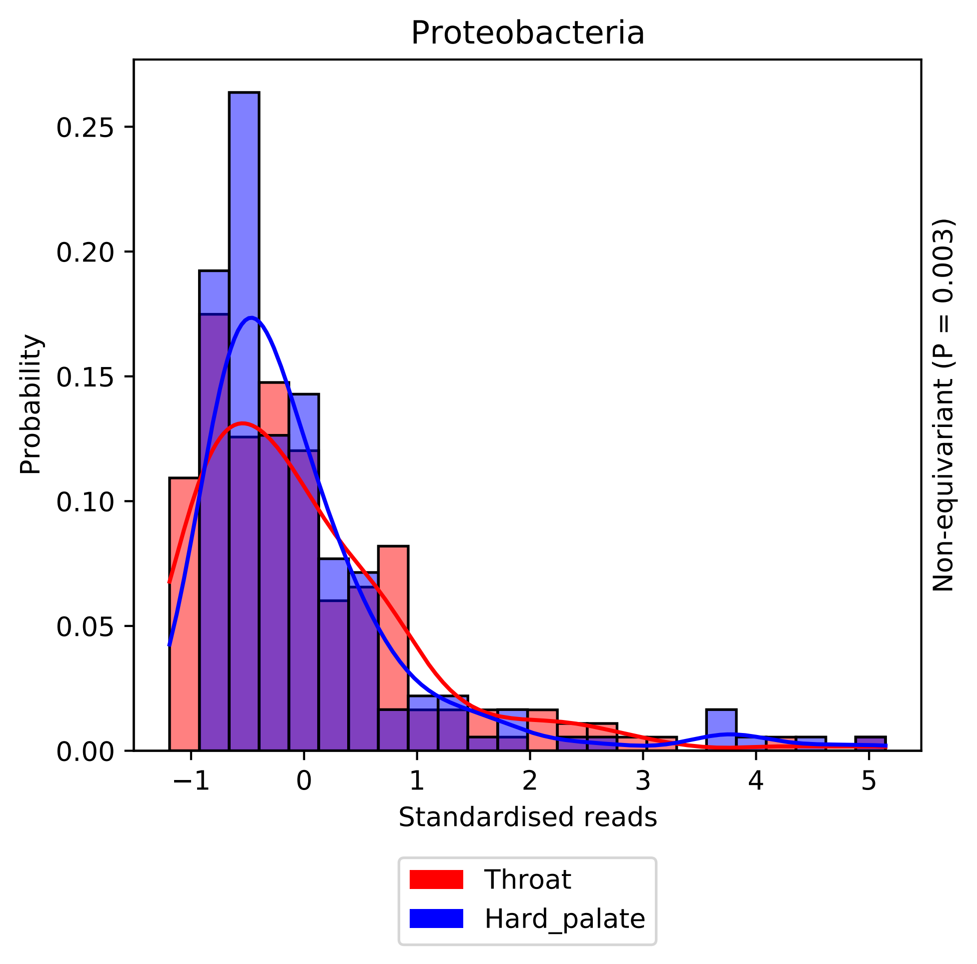

Proteobacteria
Probability
0.00
0.05
0.10
0.15
0.20
0.25
−1
0
1
2
3
4
5
Standardised reads
Non-equivariant (P = 0.003)
Throat
Hard_palate

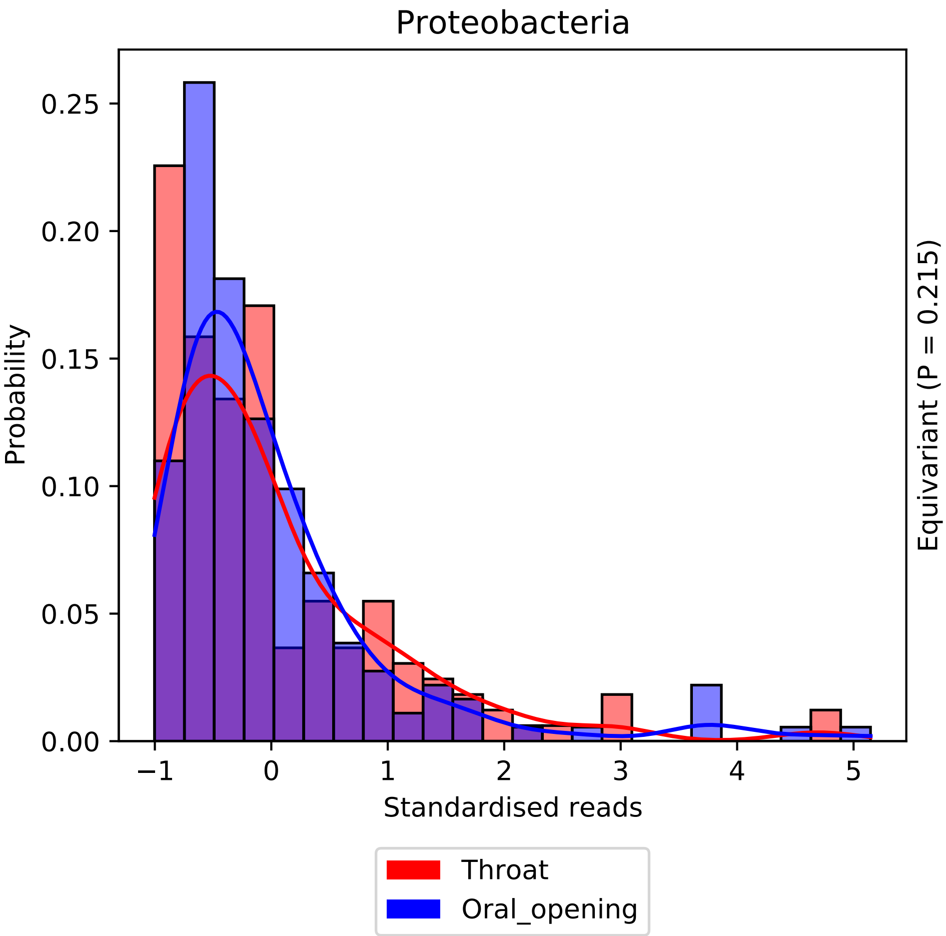
Proteobacteria
Probability
Standardised reads
Equivariant (P = 0.215)
0.00
0.05
0.10
0.15
0.20
0.25
−1
0
1
2
3
4
5
Throat
Oral_opening

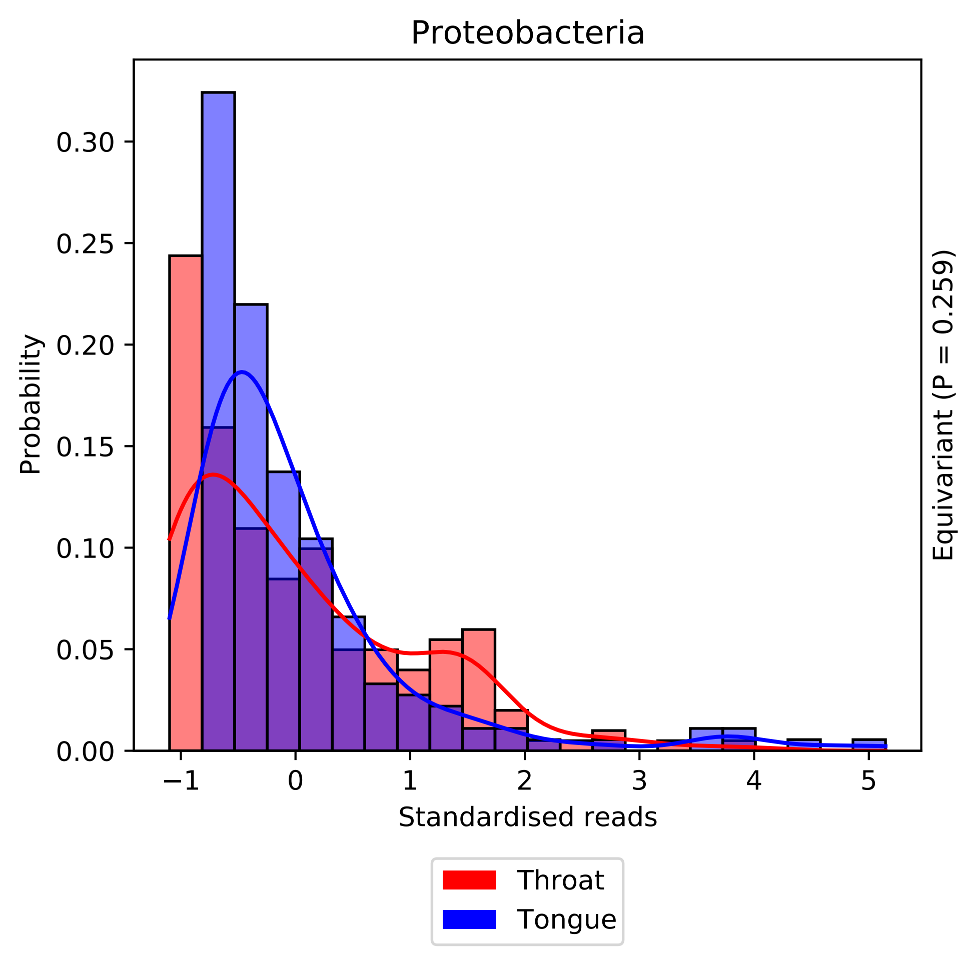

Proteobacteria
Probability
0.00
0.05
0.10
0.15
0.20
0.25
0.30
−1
0
1
2
3
4
5
Standardised reads
Equivariant (P = 0.259)
Throat
Tongue

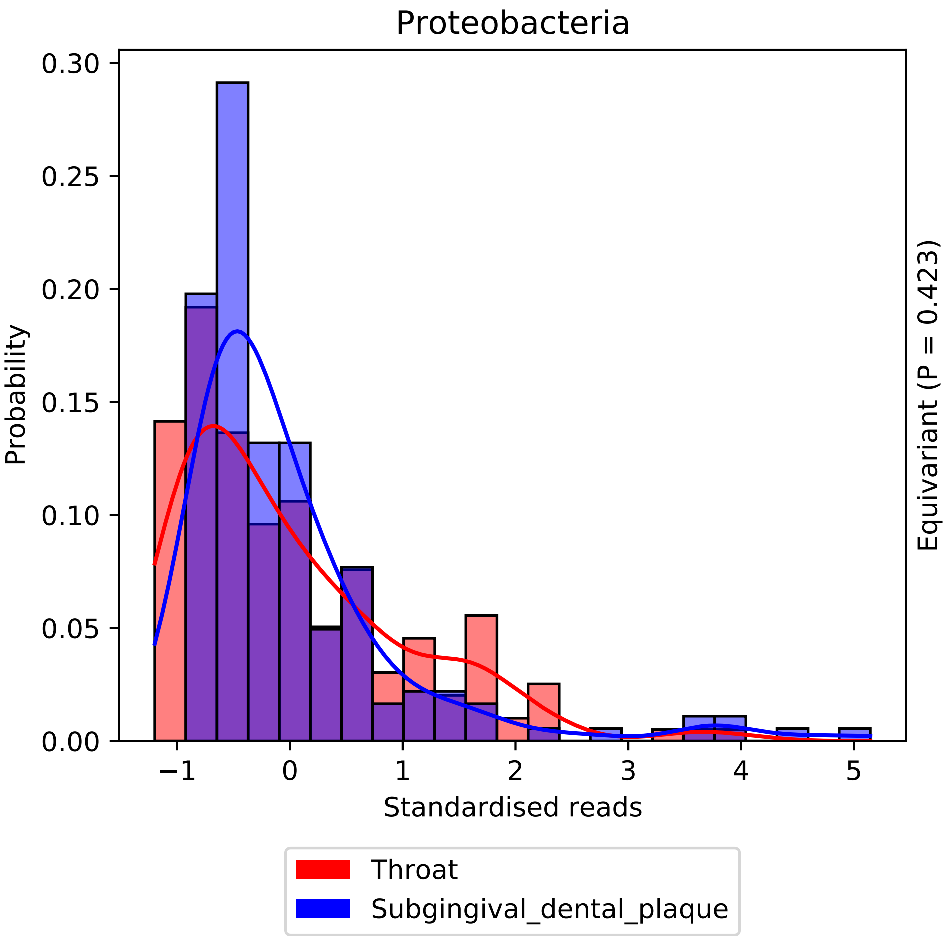

Proteobacteria
Probability
0.30
0.25
0.20
0.15
0.10
0.05
0.00
−1
0
1
2
3
4
5
Standardised reads
Equivariant (P = 0.423)
Throat
Subgingival_dental_plaque

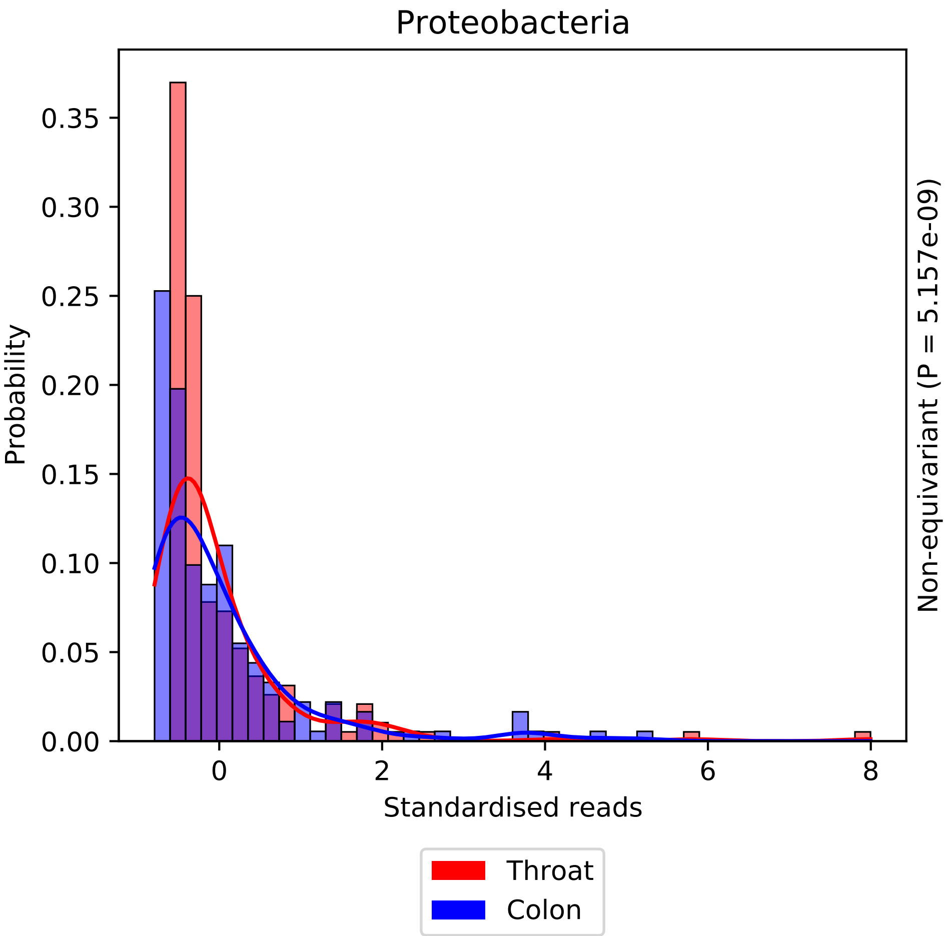

Proteobacteria
Probability
0.00
0.05
0.10
0.15
0.20
0.25
0.30
0.35
0
2
4
6
8
Standardised reads
Non-equivariant (P = 5.157e-09)
Throat
Colon

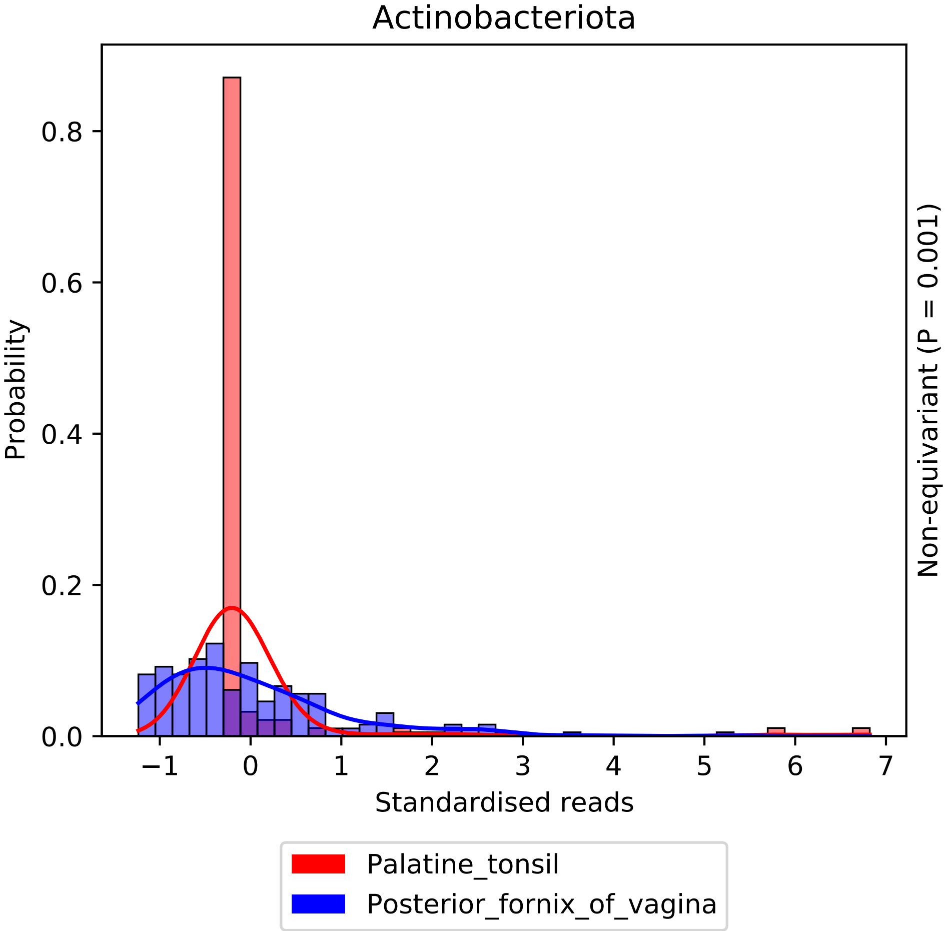

Actinobacteriota
Probability
0.0
0.2
0.4
0.6
0.8
−1
0
1
2
3
4
5
6
7
Standardised reads
Non-equivariant (P = 0.001)
Palatine_tonsil
Posterior_fornix_of_vagina

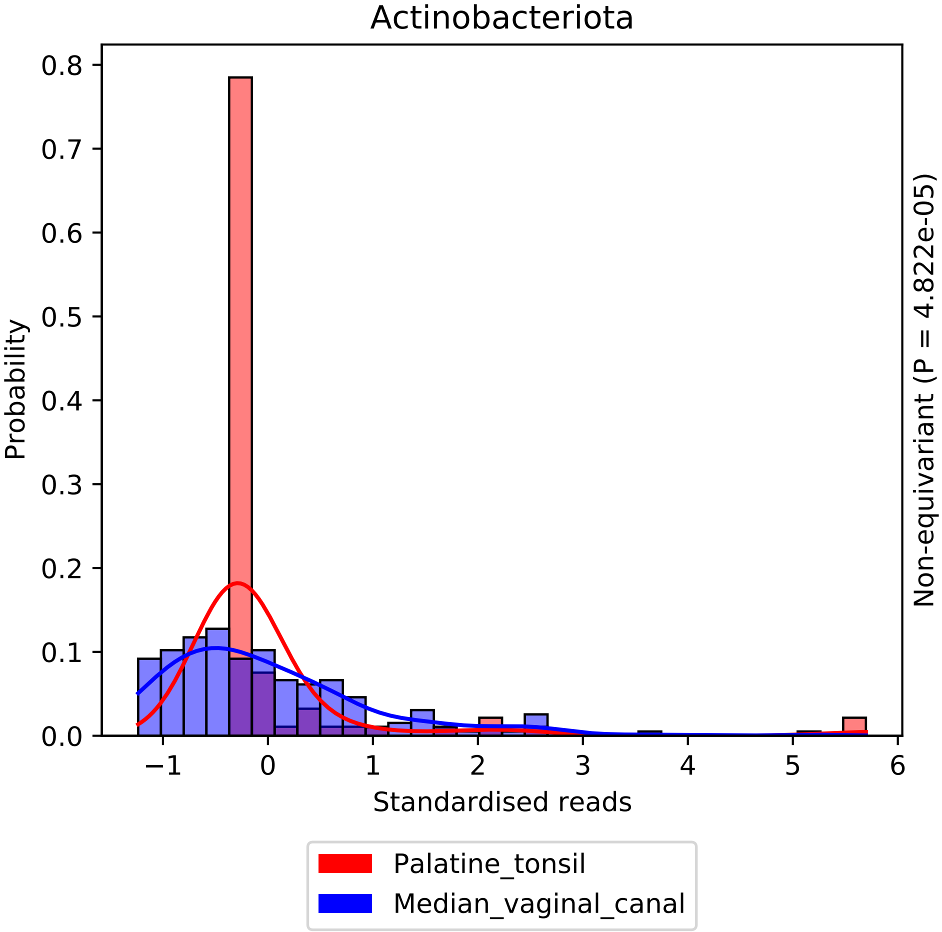

Actinobacteriota
Probability
Standardised reads
Non-equivariant (P = 4.822e-05)
0.0
0.1
0.2
0.3
0.4
0.5
0.6
0.7
0.8
−1
0
1
2
3
4
5
6
Palatine_tonsil
Median_vaginal_canal

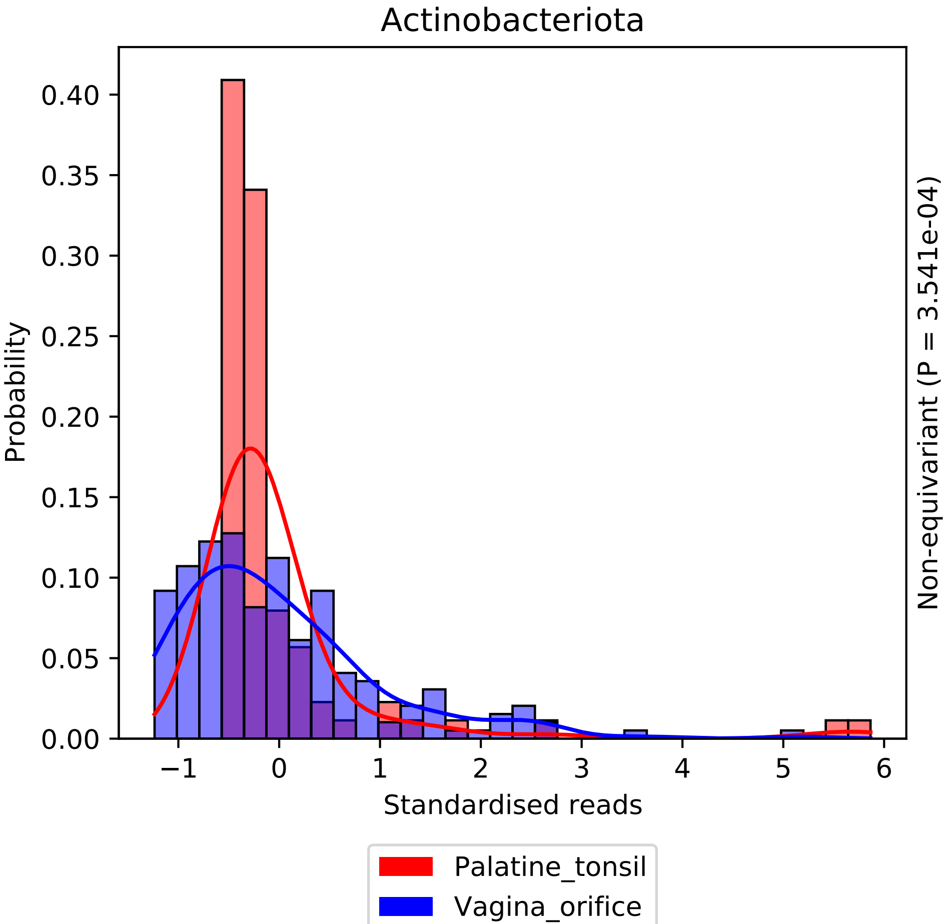

Actinobacteriota
Probability
0.00
0.05
0.10
0.15
0.20
0.25
0.30
0.35
0.40
−1
0
1
2
3
4
5
6
Standardised reads
Non-equivariant (P = 3.541e-04)
Palatine_tonsil
Vagina_orifice

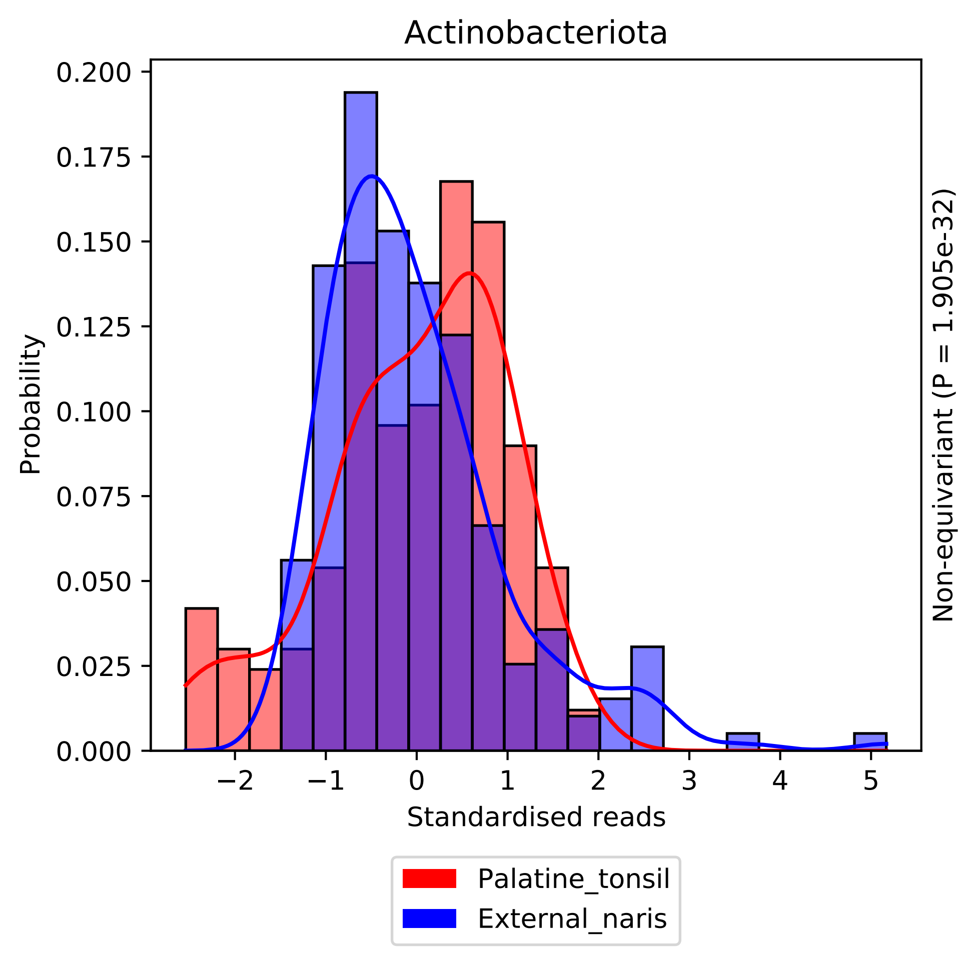
Actinobacteriota
Probability
0.200
0.175
0.150
0.125
0.100
0.075
0.050
0.025
0.000
−2
−1
0
1
2
3
4
5
Standardised reads
Non-equivariant (P = 1.905e-32)
Palatine_tonsil
External_naris

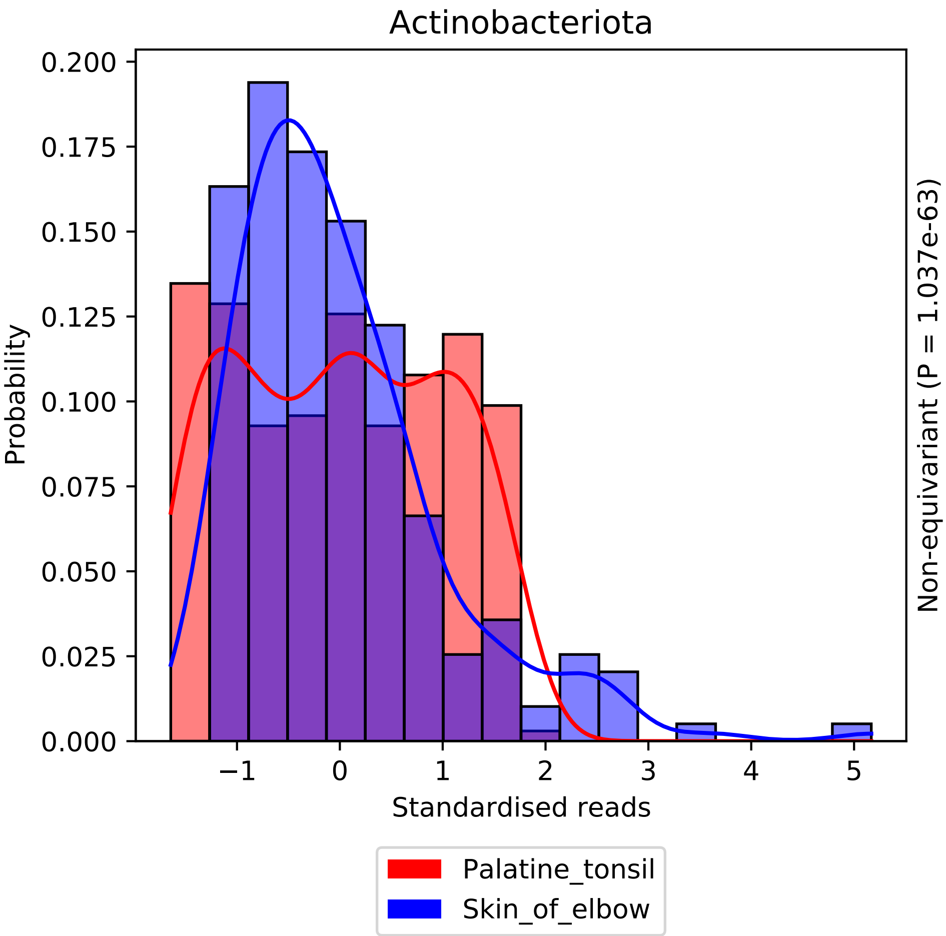
Actinobacteriota
Probability
Standardised reads
Non-equivariant (P = 1.037e-63)
0.000
0.025
0.050
0.075
0.100
0.125
0.150
0.175
0.200
−1
0
1
2
3
4
5
Palatine_tonsil
Skin_of_elbow

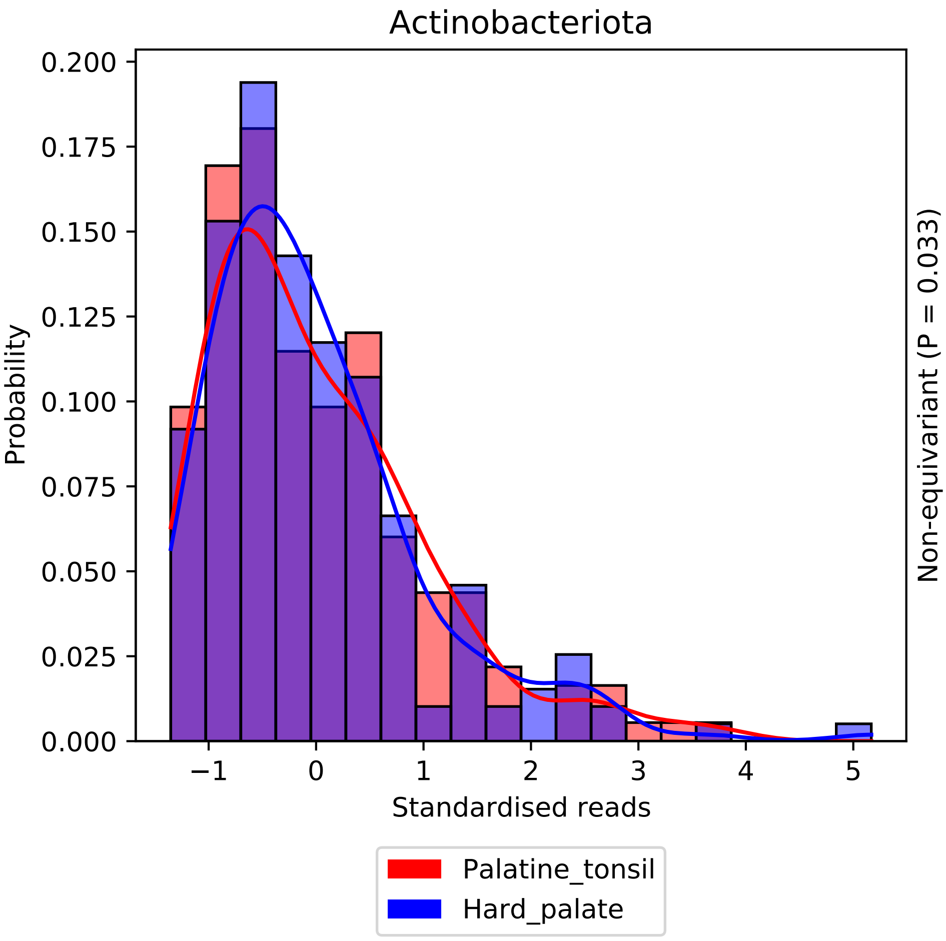
Actinobacteriota
Probability
0.000
0.025
0.050
0.075
0.100
0.125
0.150
0.175
0.200
−1
0
1
2
3
4
5
Standardised reads
Non-equivariant (P = 0.033)
Palatine_tonsil
Hard_palate

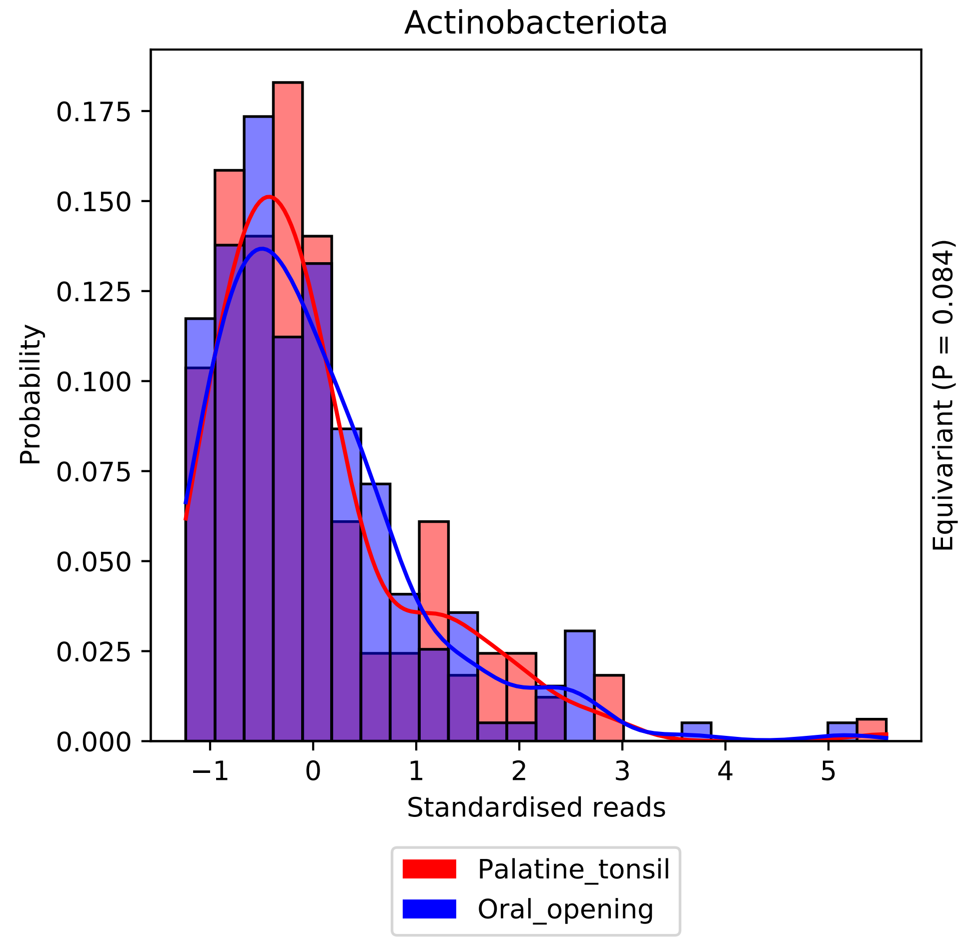
Actinobacteriota
Probability
0.000
0.025
0.050
0.075
0.100
0.125
0.150
0.175
−1
0
1
2
3
4
5
Standardised reads
Equivariant (P = 0.084)
Palatine_tonsil
Oral_opening

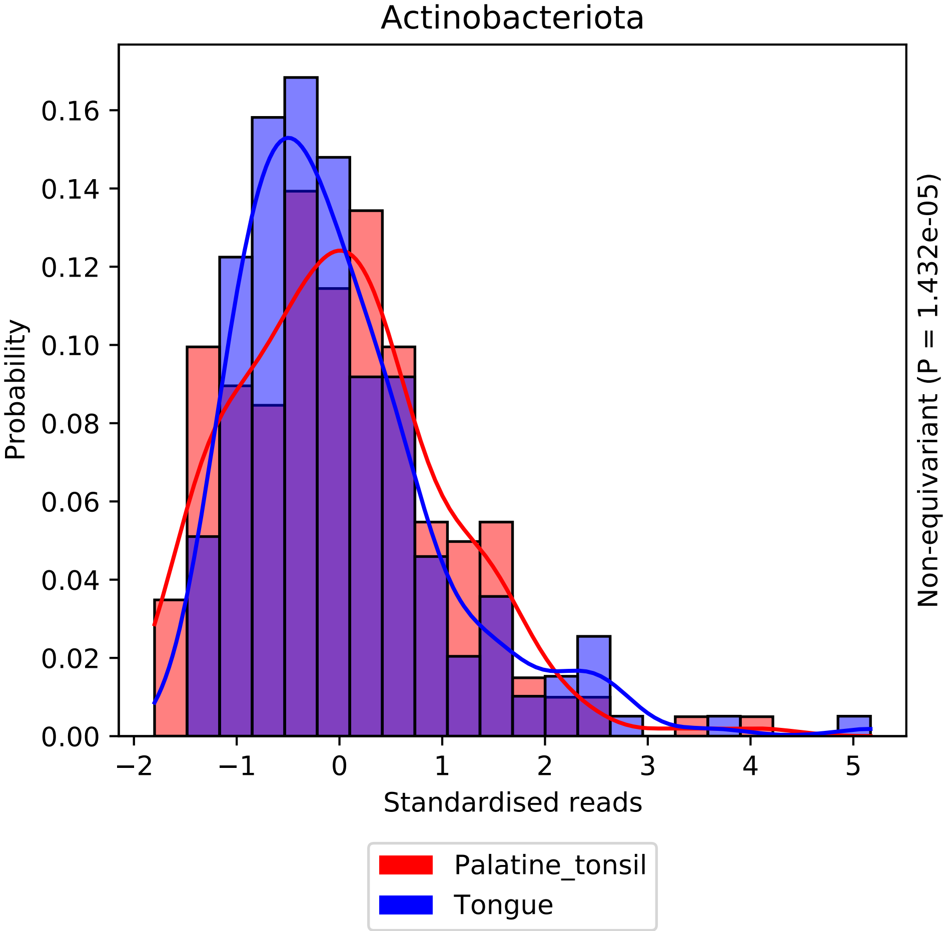
Actinobacteriota
Probability
Standardised reads
Non-equivariant (P = 1.432e-05)
0.00
0.02
0.04
0.06
0.08
0.10
0.12
0.14
0.16
−2
−1
0
1
2
3
4
5
Palatine_tonsil
Tongue

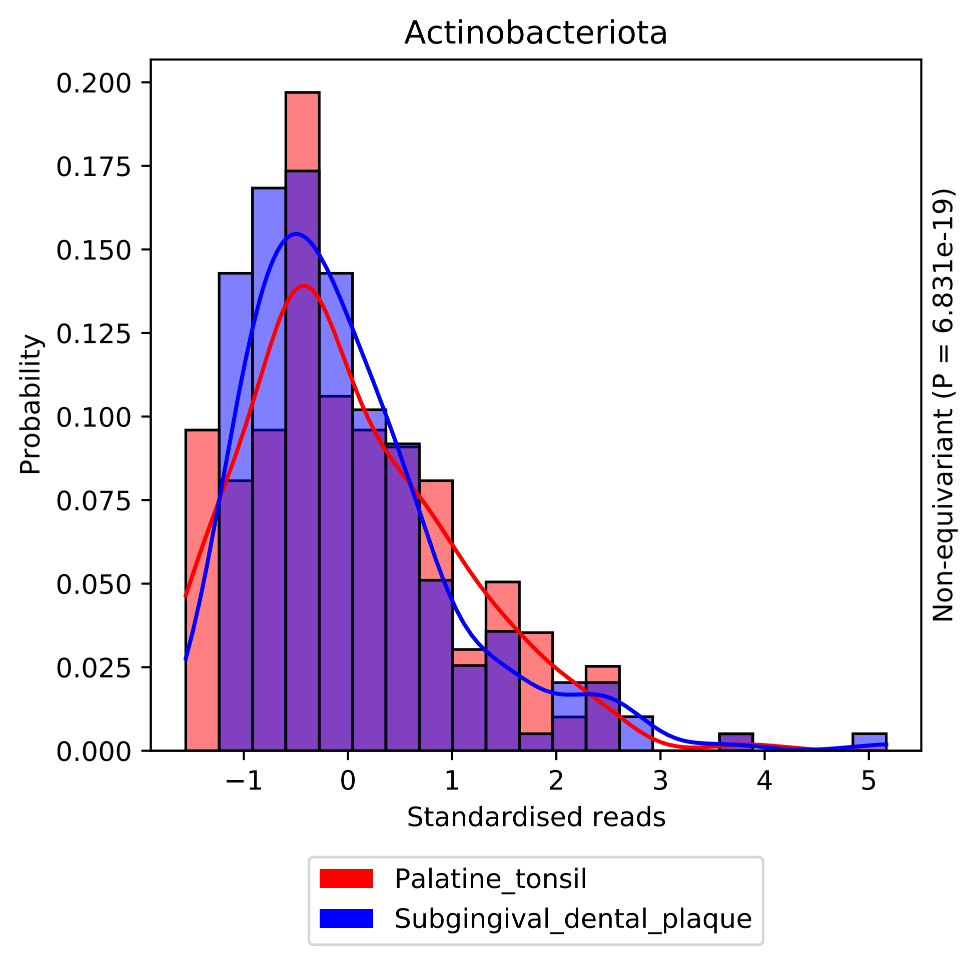
Actinobacteriota
Probability
0.000
0.025
0.050
0.075
0.100
0.125
0.150
0.175
0.200
−1
0
1
2
3
4
5
Standardised reads
Non-equivariant (P = 6.831e-19)
Palatine_tonsil
Subgingival_dental_plaque

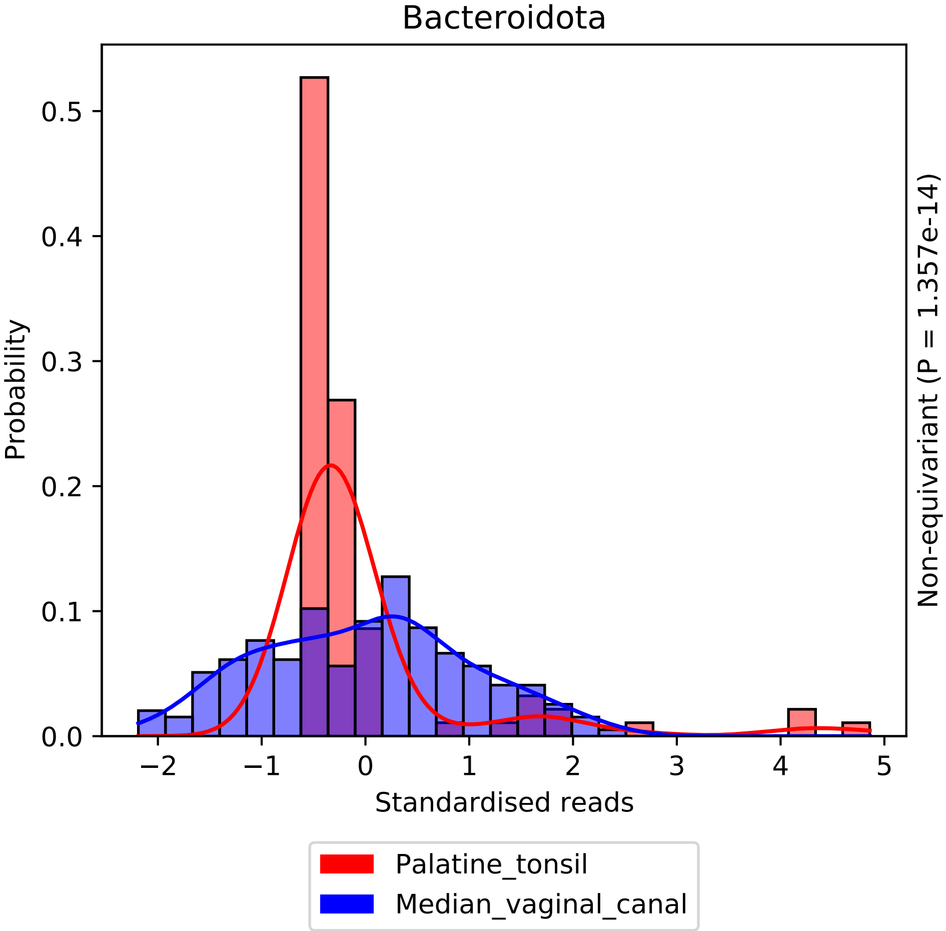

Bacteroidota
Probability
0.0
0.1
0.2
0.3
0.4
0.5
−2
−1
0
1
2
3
4
5
Standardised reads
Non-equivariant (P = 1.357e-14)
Palatine_tonsil
Median_vaginal_canal

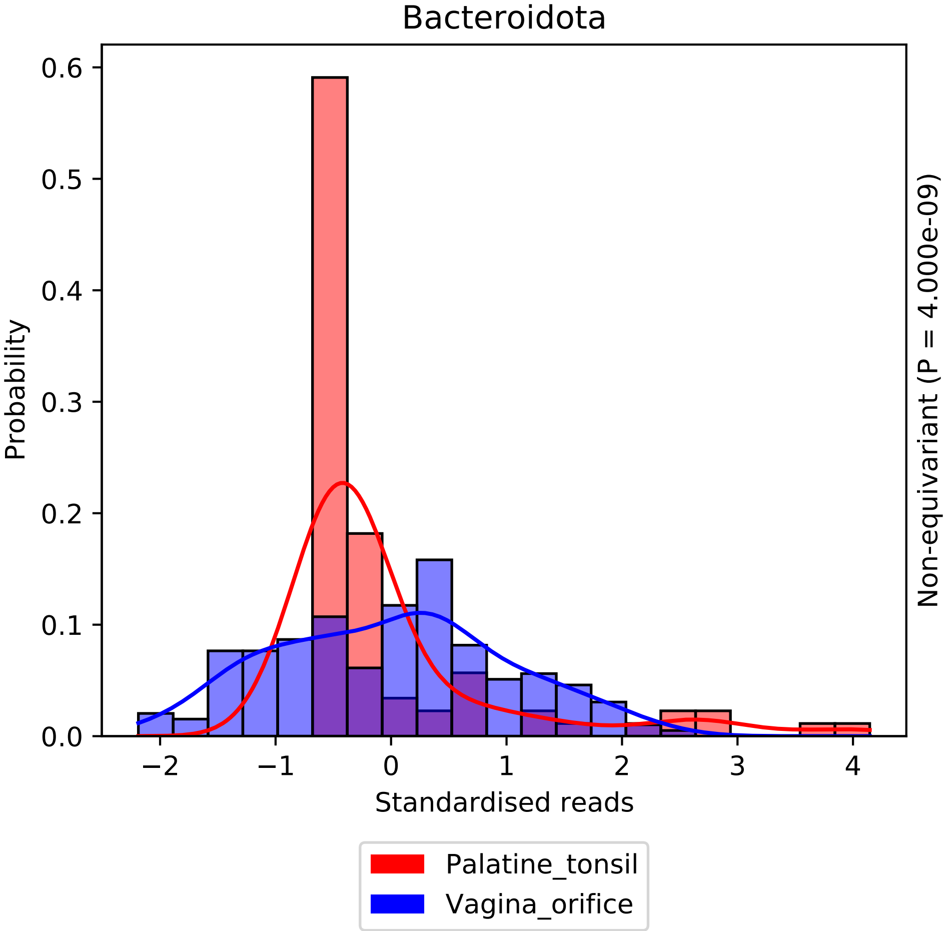
Bacteroidota
Probability
Standardised reads
Non-equivariant (P = 4.000e-09)
0.0
0.1
0.2
0.3
0.4
0.5
0.6
−2
−1
0
1
2
3
4
Palatine_tonsil
Vagina_orifice

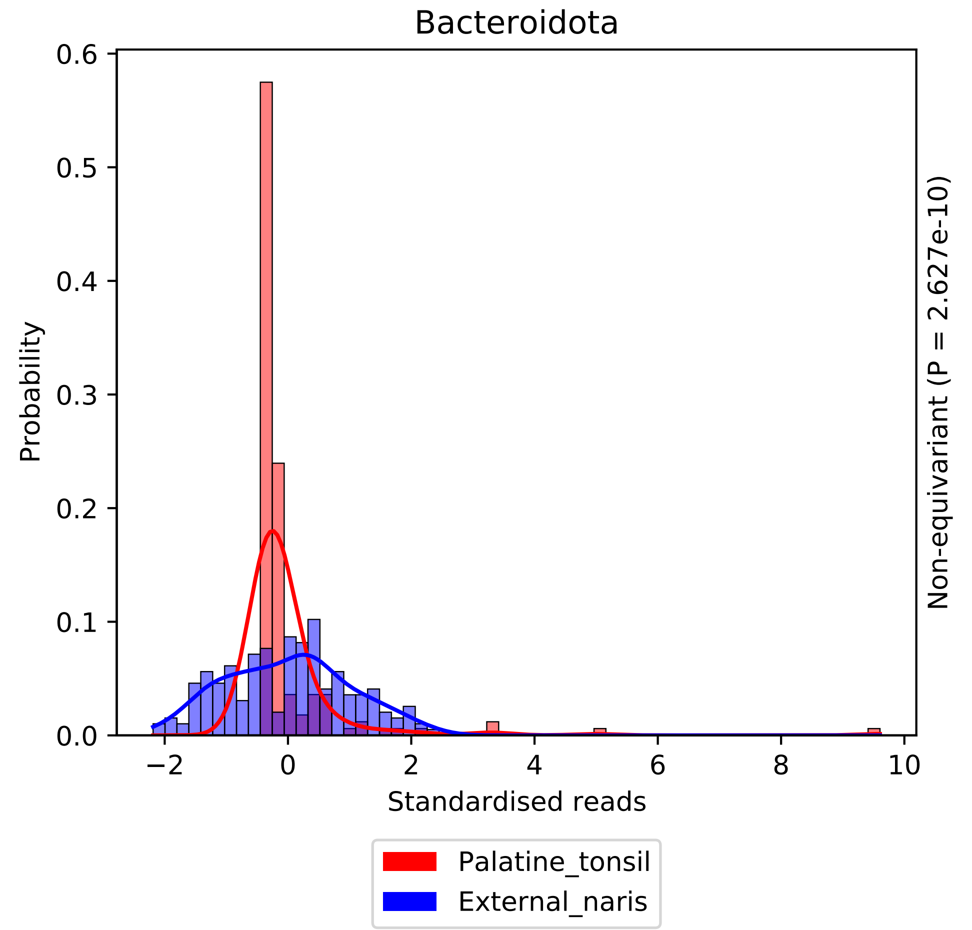

Bacteroidota
Probability
Standardised reads
Non-equivariant (P = 2.627e-10)
Palatine_tonsil
External_naris
0.0
0.1
0.2
0.3
0.4
0.5
0.6
−2
0
2
4
6
8
10

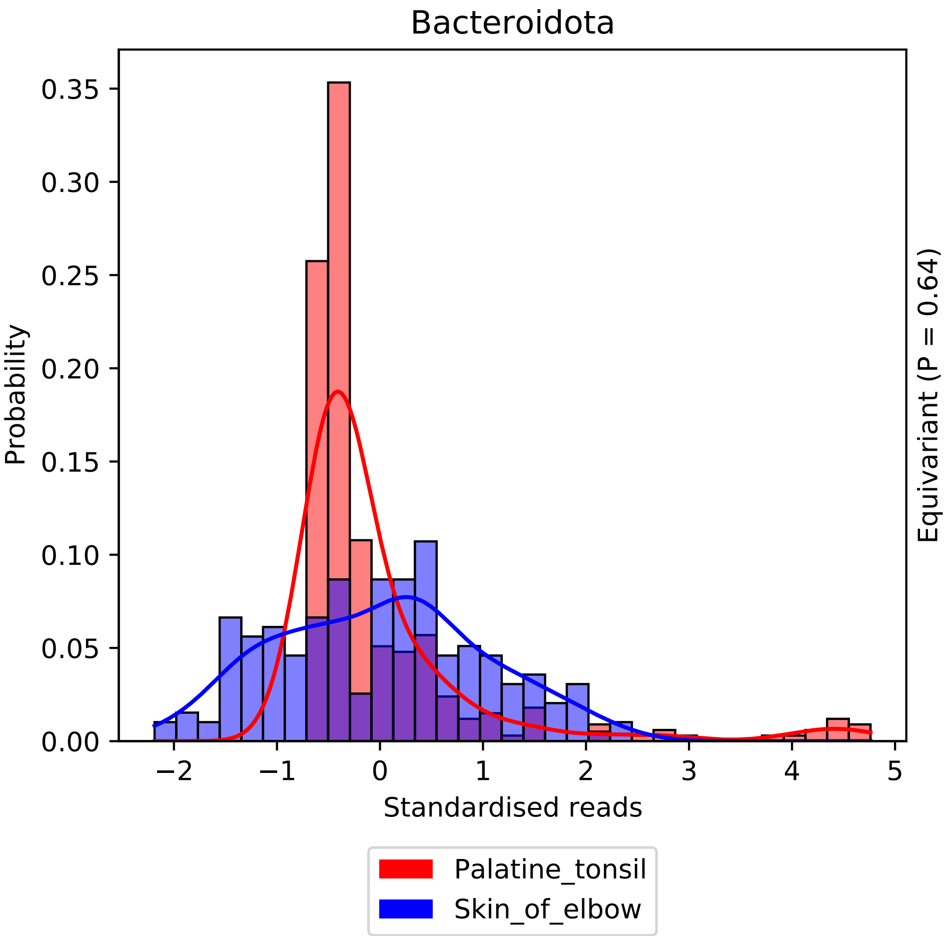

Bacteroidota
Probability
0.35
0.30
0.25
0.20
0.15
0.10
0.05
0.00
−2
−1
0
1
2
3
4
5
Standardised reads
Equivariant (P = 0.64)
Palatine_tonsil
Skin_of_elbow

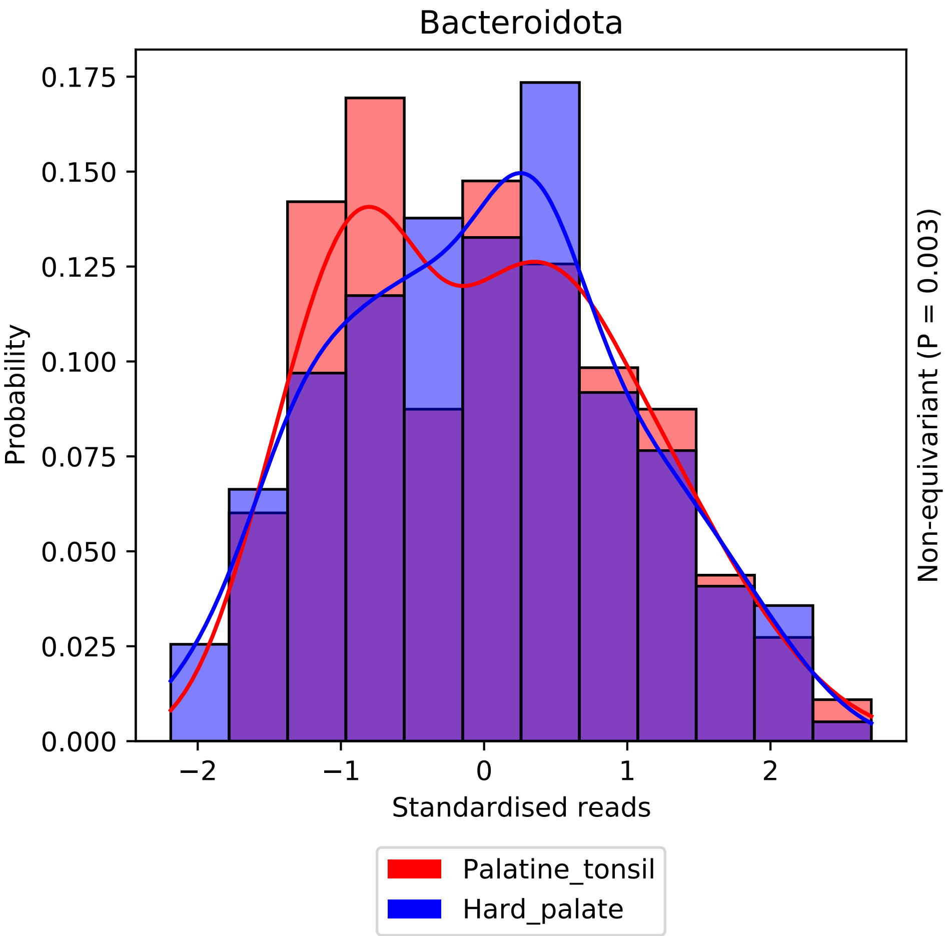

Bacteroidota
Probability
0.175
0.150
0.125
0.100
0.075
0.050
0.025
0.000
−2
−1
0
1
2
Standardised reads
Non-equivariant (P = 0.003)
Palatine_tonsil
Hard_palate

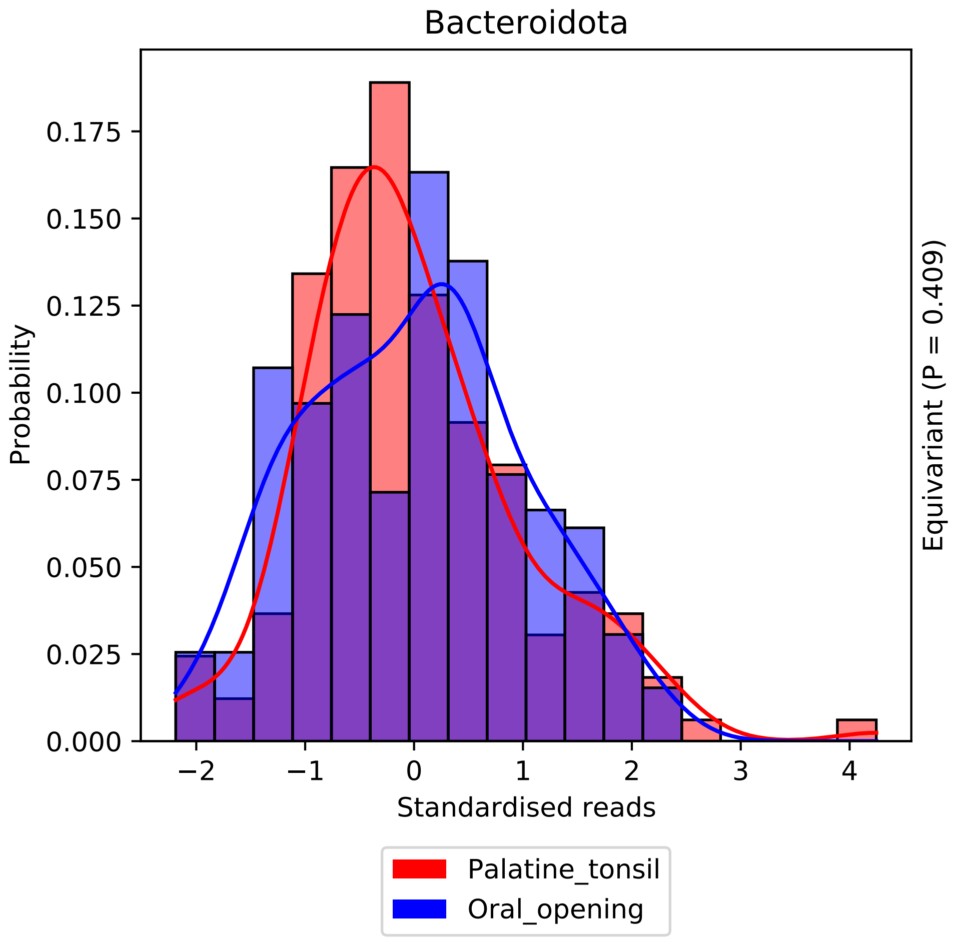

Bacteroidota
Probability
0.000
0.025
0.050
0.075
0.100
0.125
0.150
0.175
−2
−1
0
1
2
3
4
Standardised reads
Equivariant (P = 0.409)
Palatine_tonsil
Oral_opening

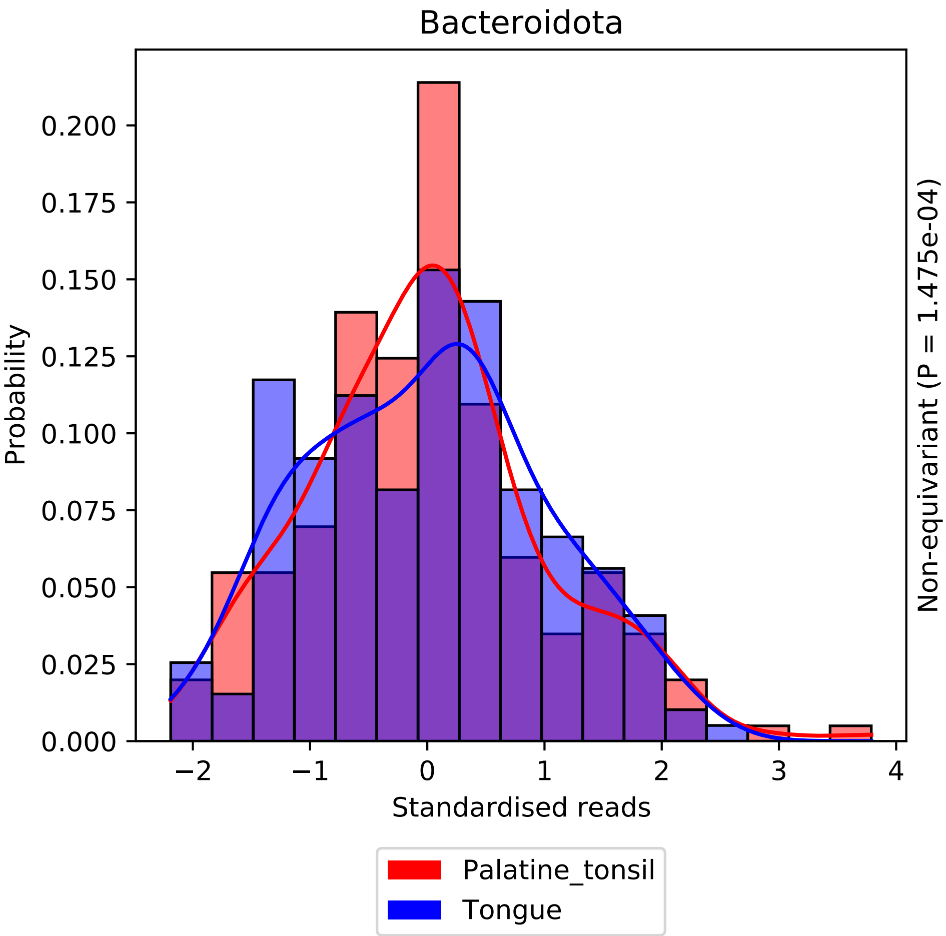

Bacteroidota
Probability
0.000
0.025
0.050
0.075
0.100
0.125
0.150
0.175
0.200
−2
−1
0
1
2
3
4
Standardised reads
Non-equivariant (P = 1.475e-04)
Palatine_tonsil
Tongue

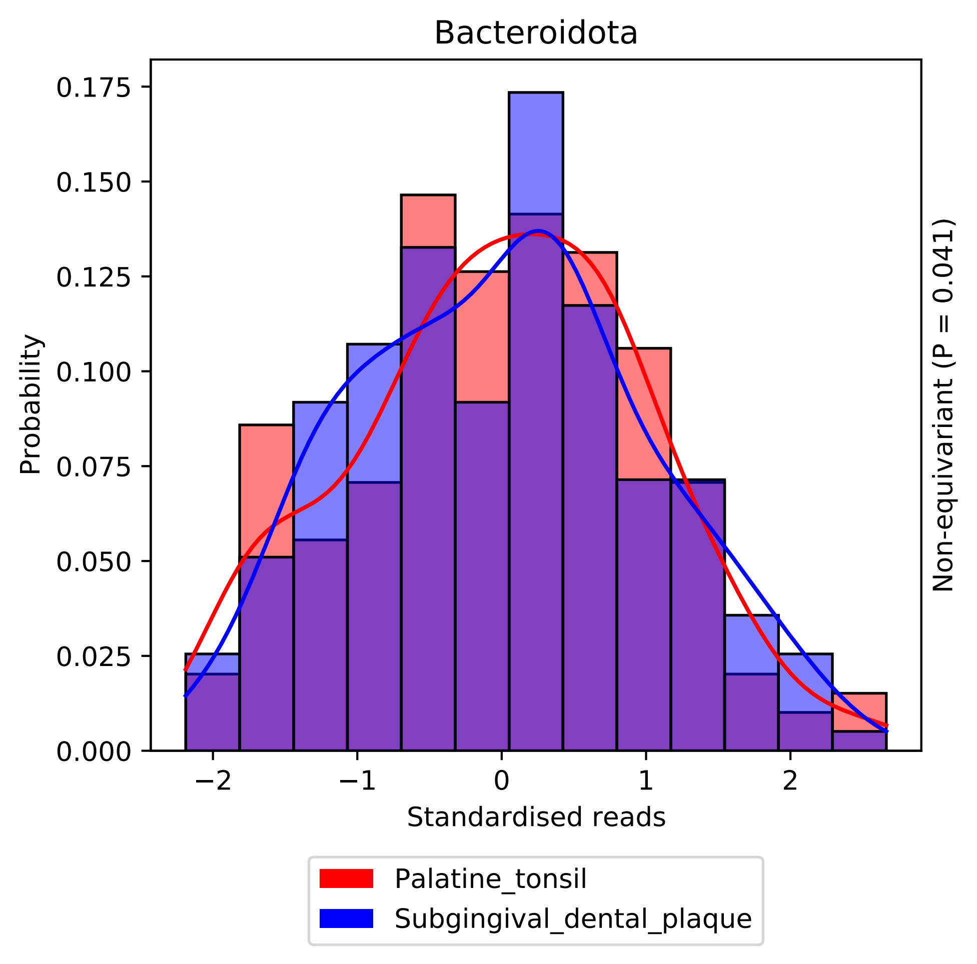
Bacteroidota
Probability
0.000
0.025
0.050
0.075
0.100
0.125
0.150
0.175
−2
−1
0
1
2
Standardised reads
Non-equivariant (P = 0.041)
Palatine_tonsil
Subgingival_dental_plaque

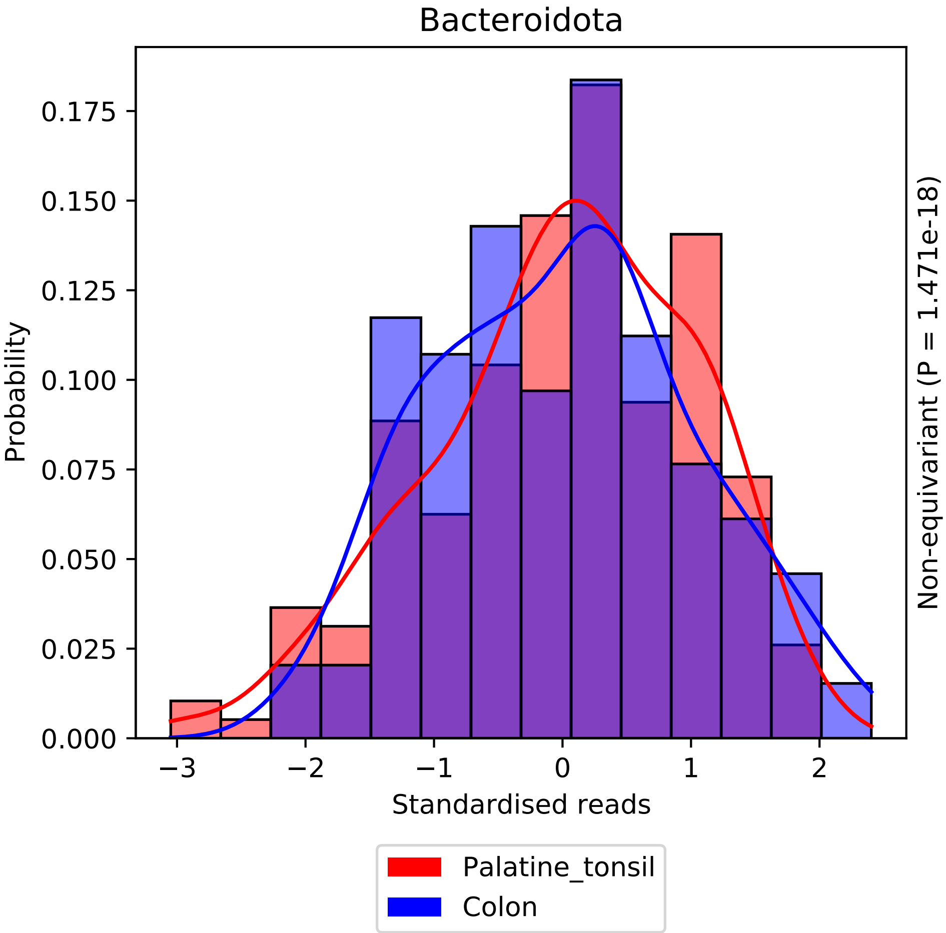

Bacteroidota
Probability
0.000
0.025
0.050
0.075
0.100
0.125
0.150
0.175
−3
−2
−1
0
1
2
Standardised reads
Non-equivariant (P = 1.471e-18)
Palatine_tonsil
Colon

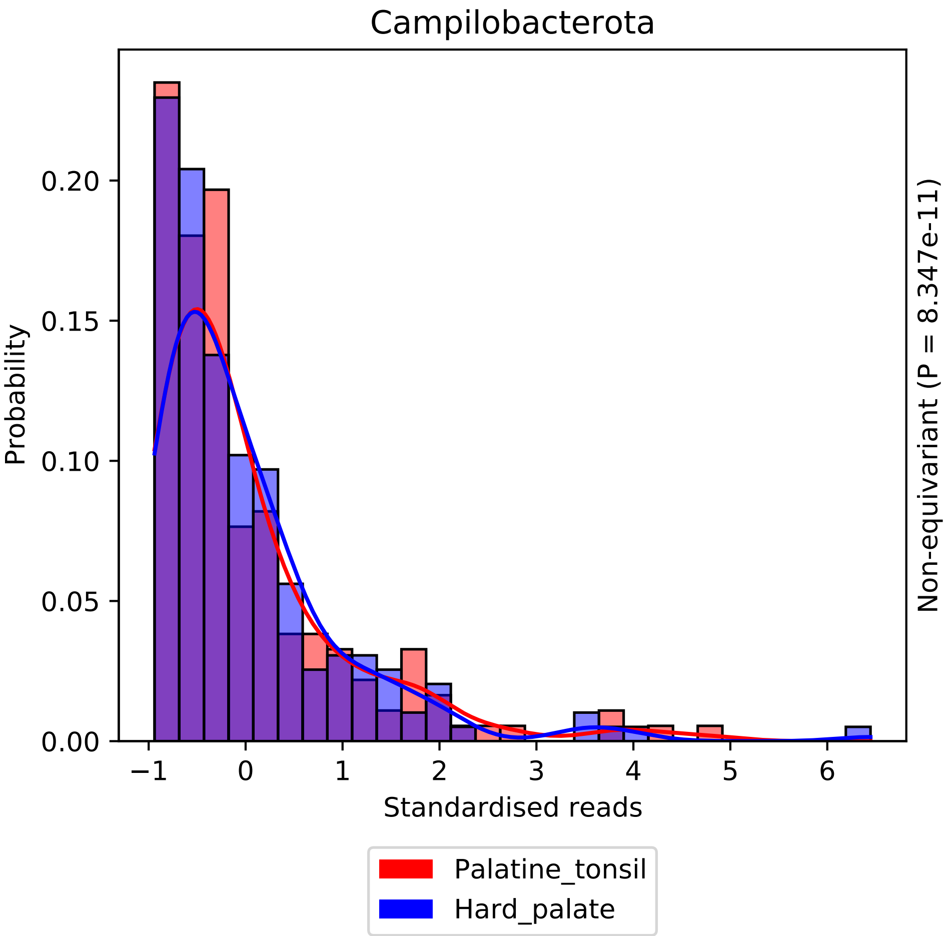
Campilobacterota
Probability
Standardised reads
Non-equivariant (P = 8.347e-11)
0.00
0.05
0.10
0.15
0.20
−1
0
1
2
3
4
5
6
Palatine_tonsil
Hard_palate

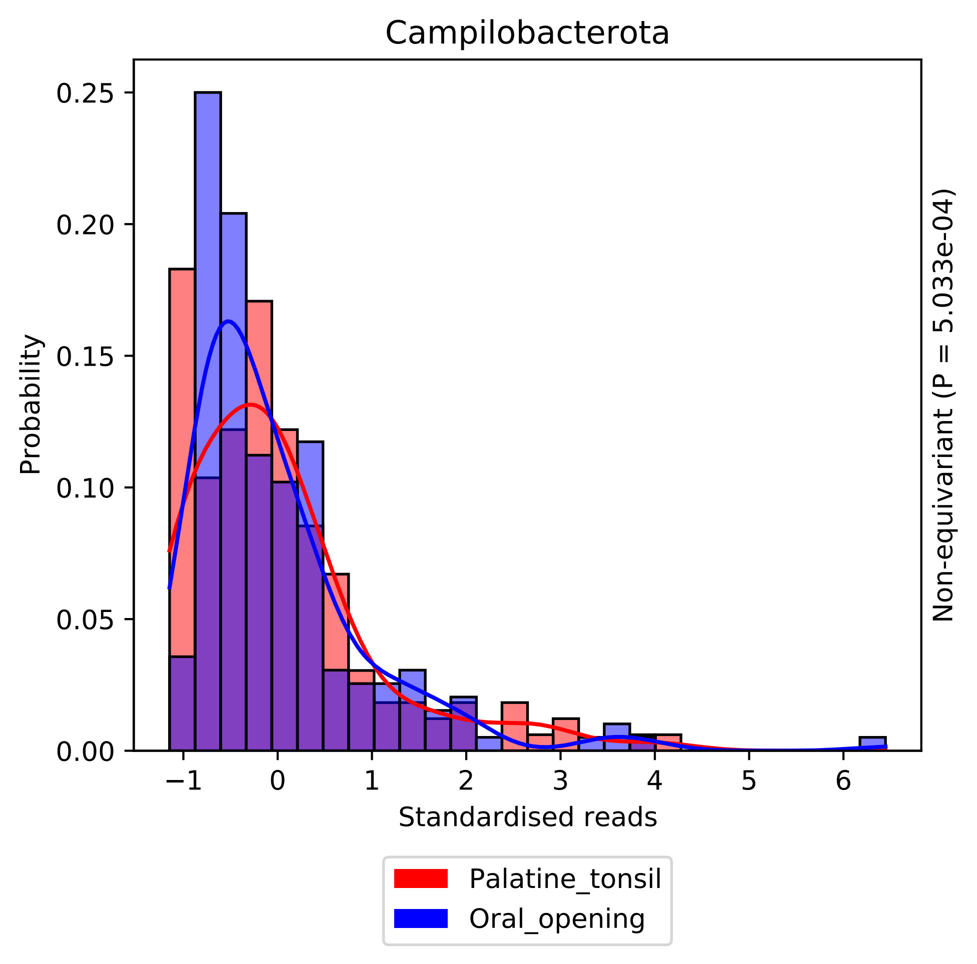

Campilobacterota
Probability
0.00
0.05
0.10
0.15
0.20
0.25
−1
0
1
2
3
4
5
6
Standardised reads
Non-equivariant (P = 5.033e-04)
Palatine_tonsil
Oral_opening

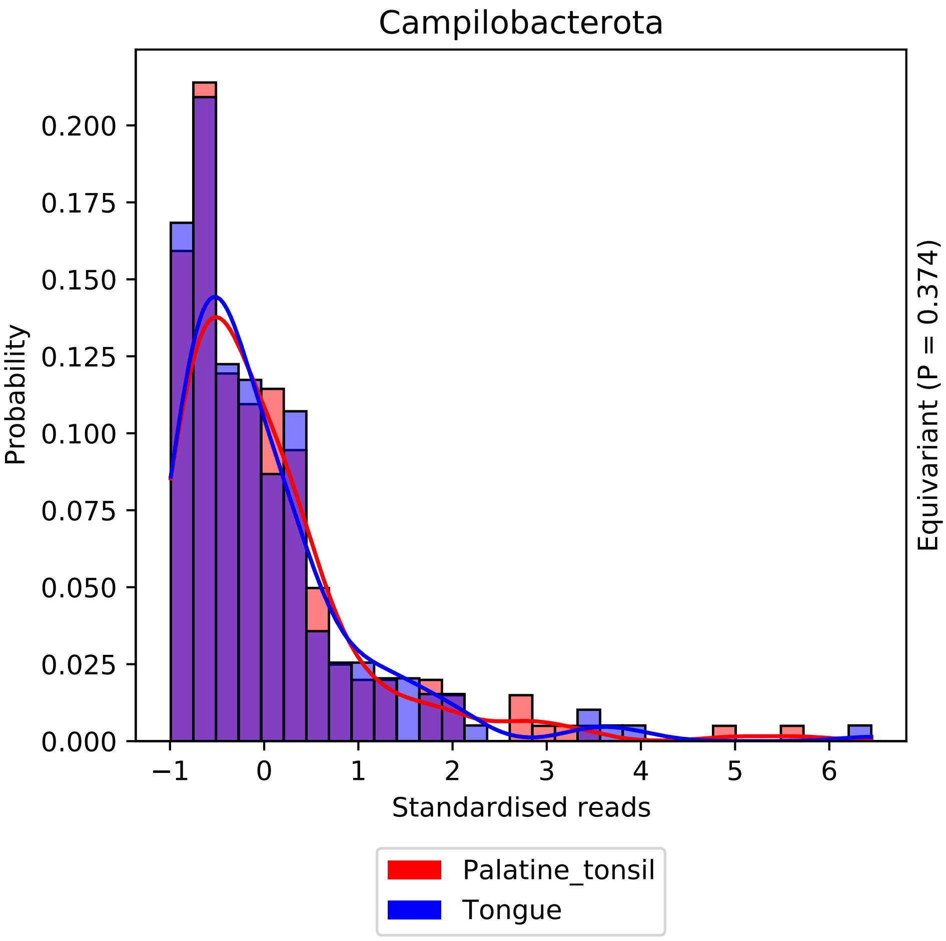

Campilobacterota
Probability
0.000
0.025
0.050
0.075
0.100
0.125
0.150
0.175
0.200
−1
0
1
2
3
4
5
6
Standardised reads
Equivariant (P = 0.374)
Palatine_tonsil
Tongue

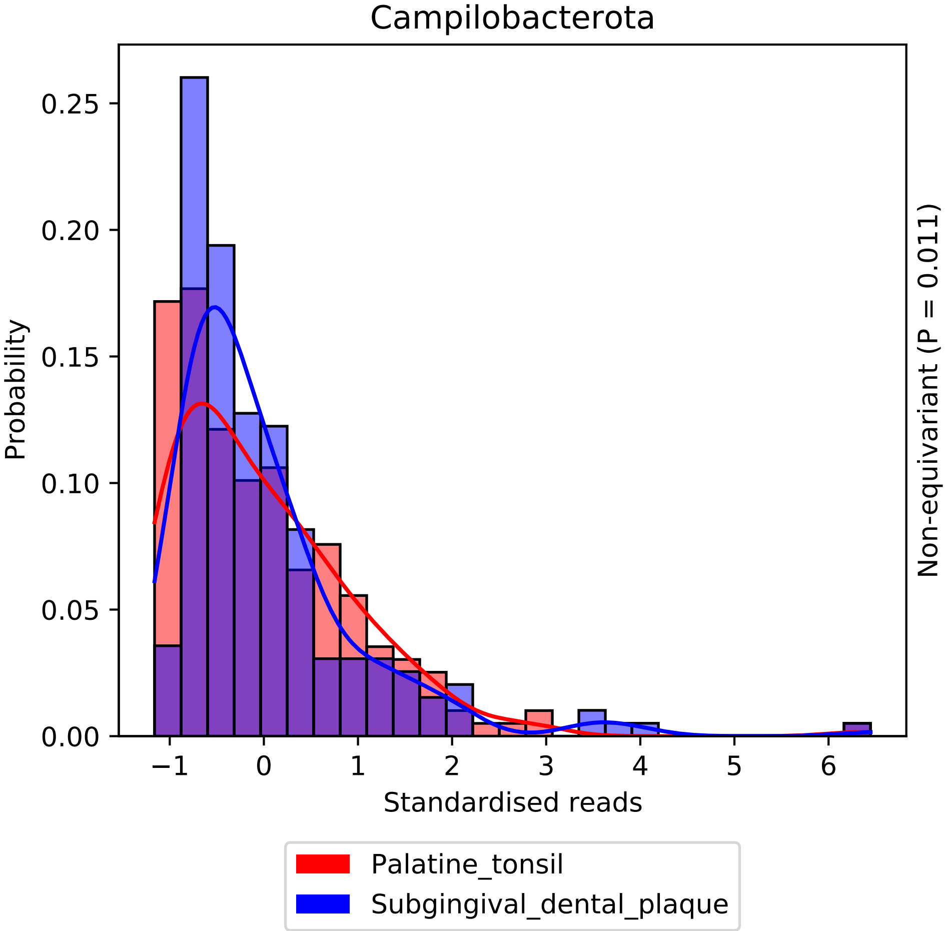

Campilobacterota
Probability
0.00
0.05
0.10
0.15
0.20
0.25
−1
0
1
2
3
4
5
6
Standardised reads
Non-equivariant (P = 0.011)
Palatine_tonsil
Subgingival_dental_plaque

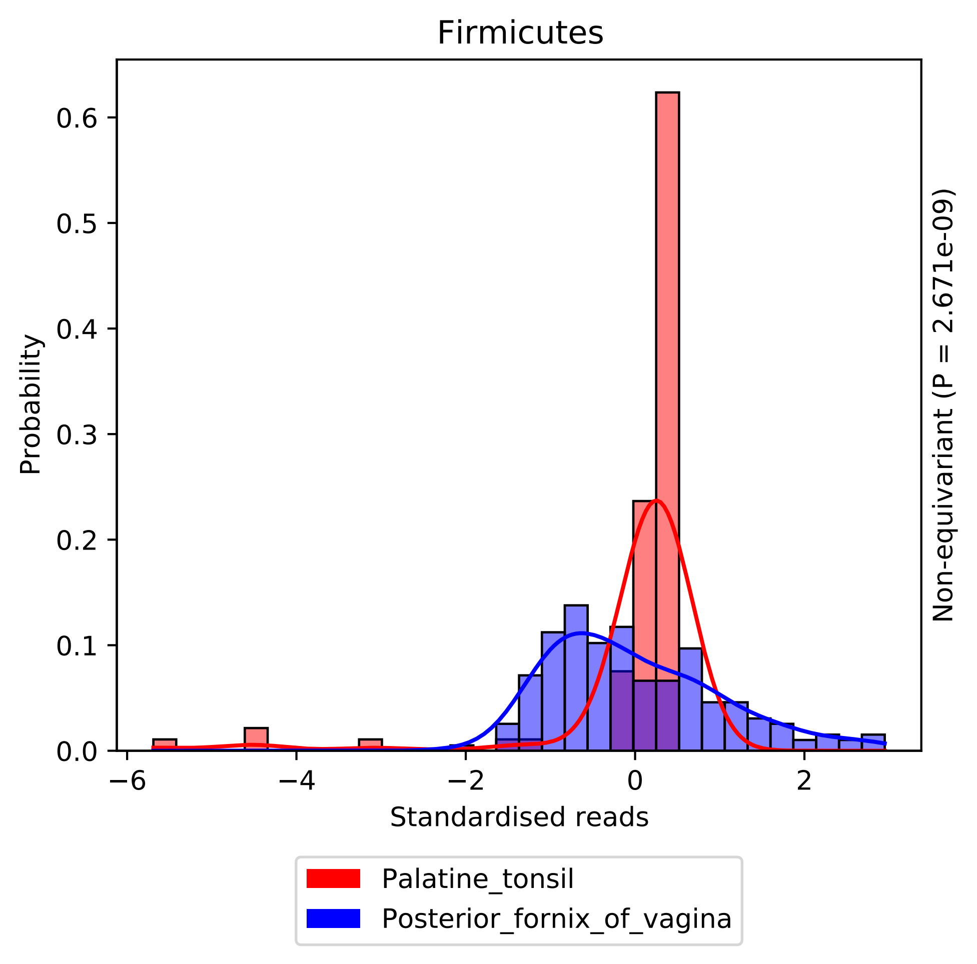

Firmicutes
Probability
0.0
0.1
0.2
0.3
0.4
0.5
0.6
−6
−4
−2
0
2
Standardised reads
Non-equivariant (P = 2.671e-09)
Palatine_tonsil
Posterior_fornix_of_vagina

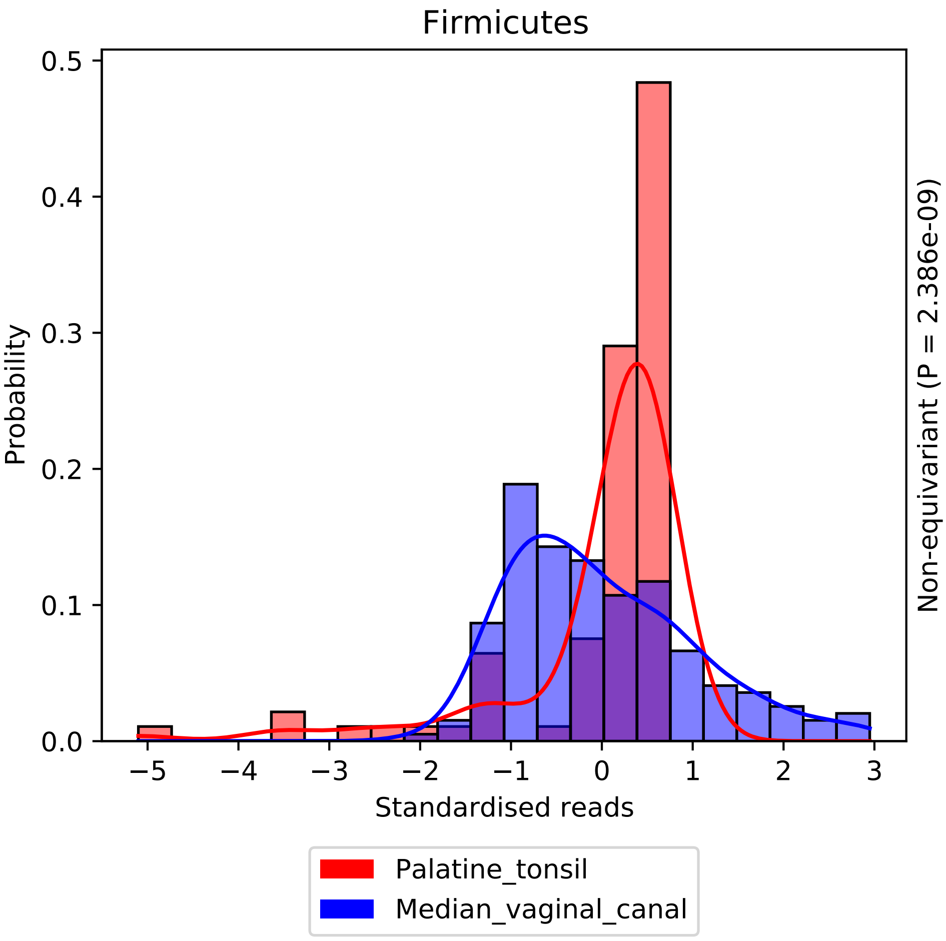
Firmicutes
Probability
Standardised reads
Non-equivariant (P = 2.386e-09)
Palatine_tonsil
Median_vaginal_canal
0.0
0.1
0.2
0.3
0.4
0.5
−5
−4
−3
−2
−1
0
1
2
3

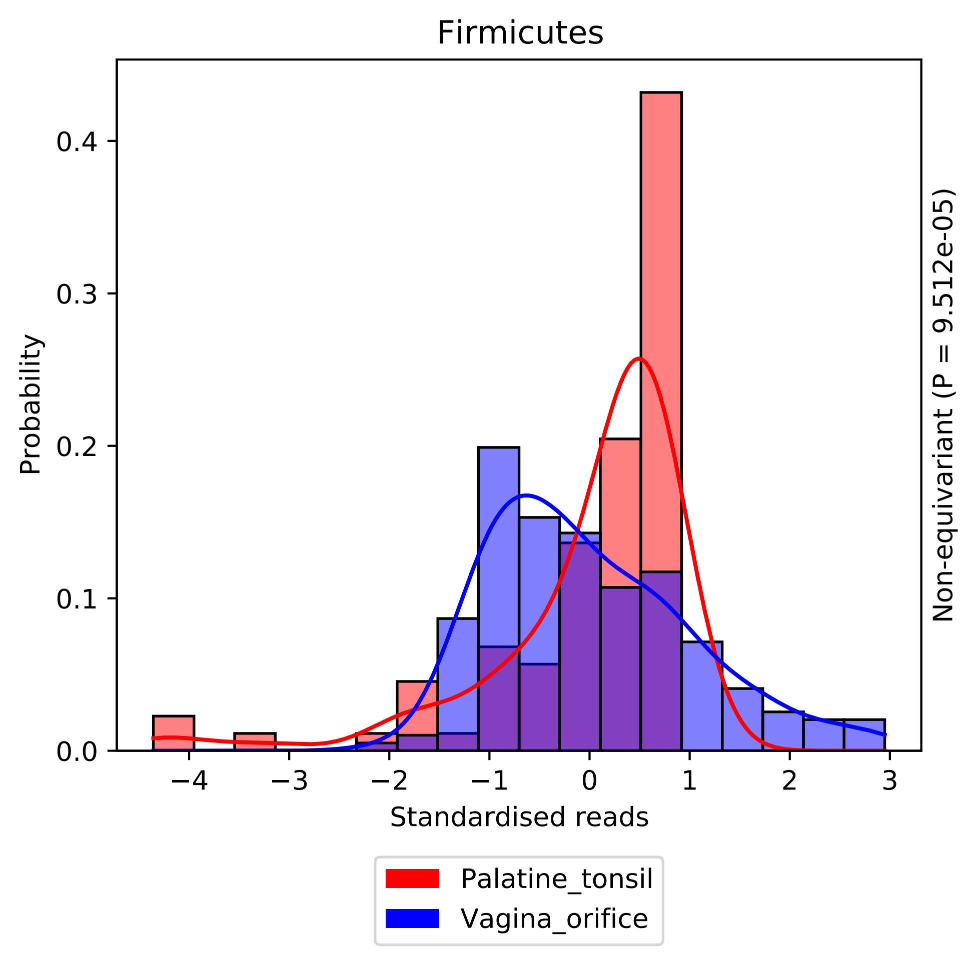
Firmicutes
Probability
0.0
0.1
0.2
0.3
0.4
−4
−3
−2
−1
0
1
2
3
Standardised reads
Non-equivariant (P = 9.512e-05)
Palatine_tonsil
Vagina_orifice

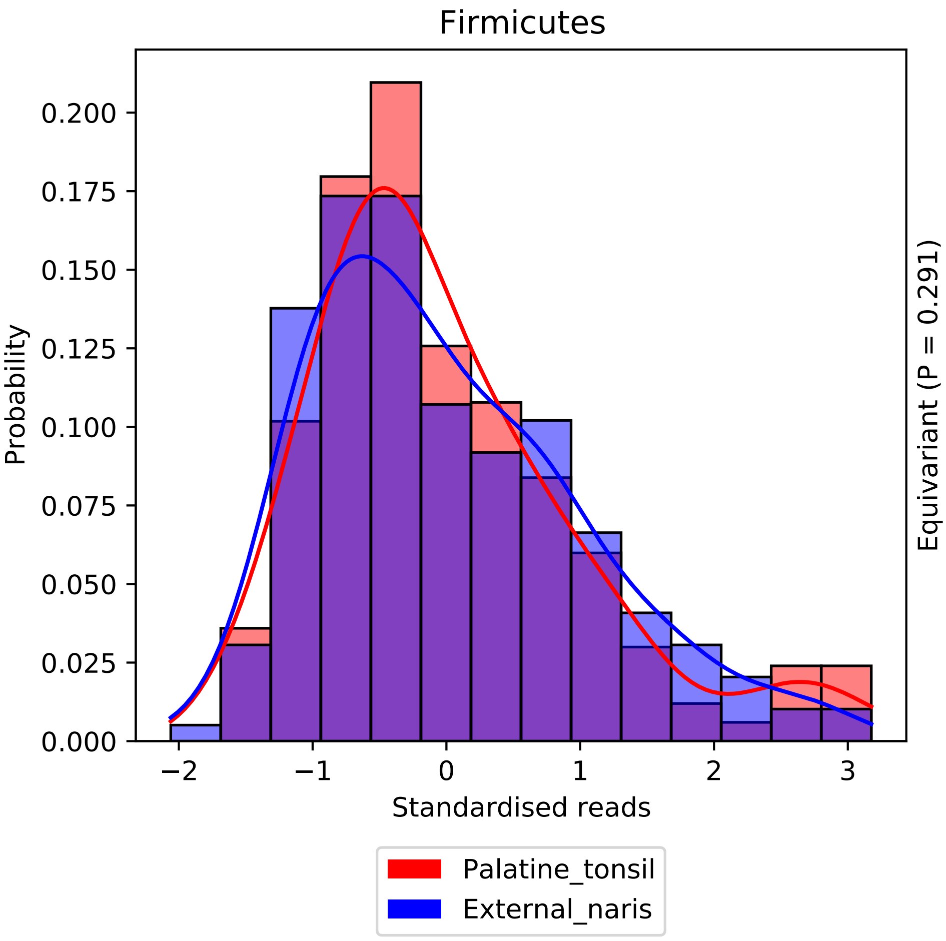

Firmicutes
Probability
0.000
0.025
0.050
0.075
0.100
0.125
0.150
0.175
0.200
−2
−1
0
1
2
3
Standardised reads
Equivariant (P = 0.291)
Palatine_tonsil
External_naris

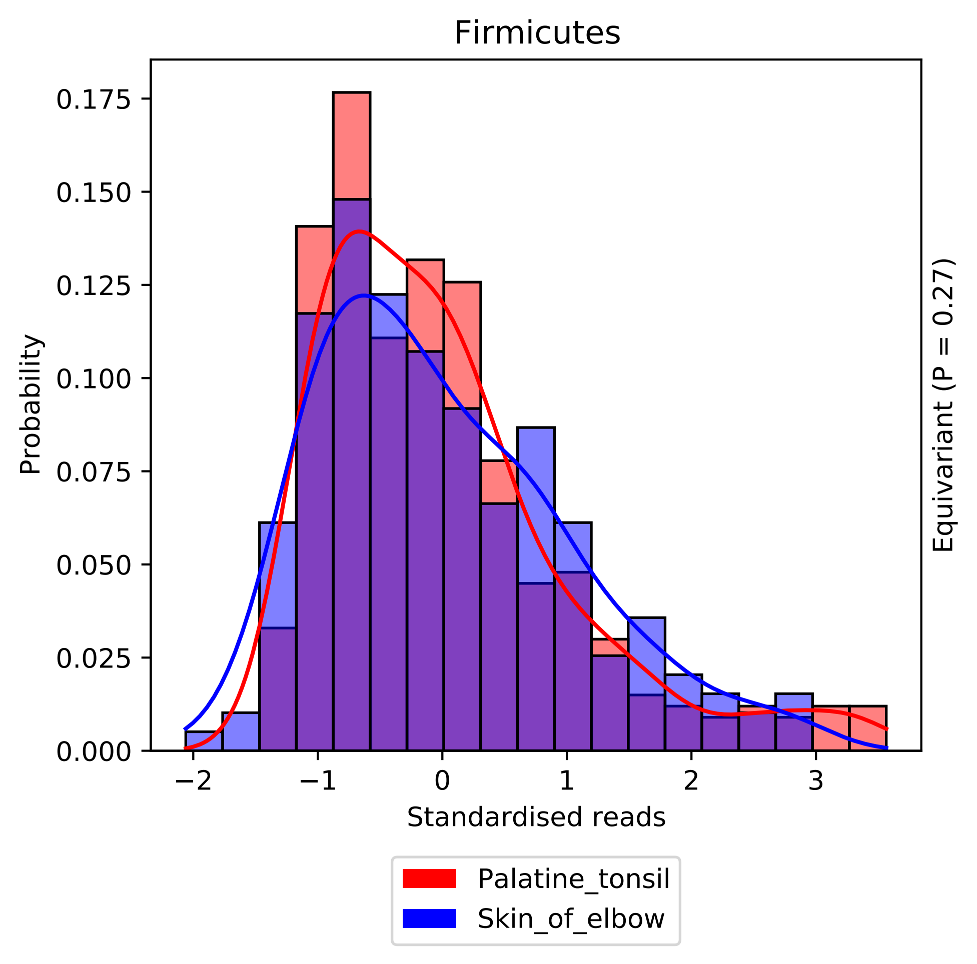
Firmicutes
Probability
0.175
0.150
0.125
0.100
0.075
0.050
0.025
0.000
−2
−1
0
1
2
3
Standardised reads
Equivariant (P = 0.27)
Palatine_tonsil
Skin_of_elbow

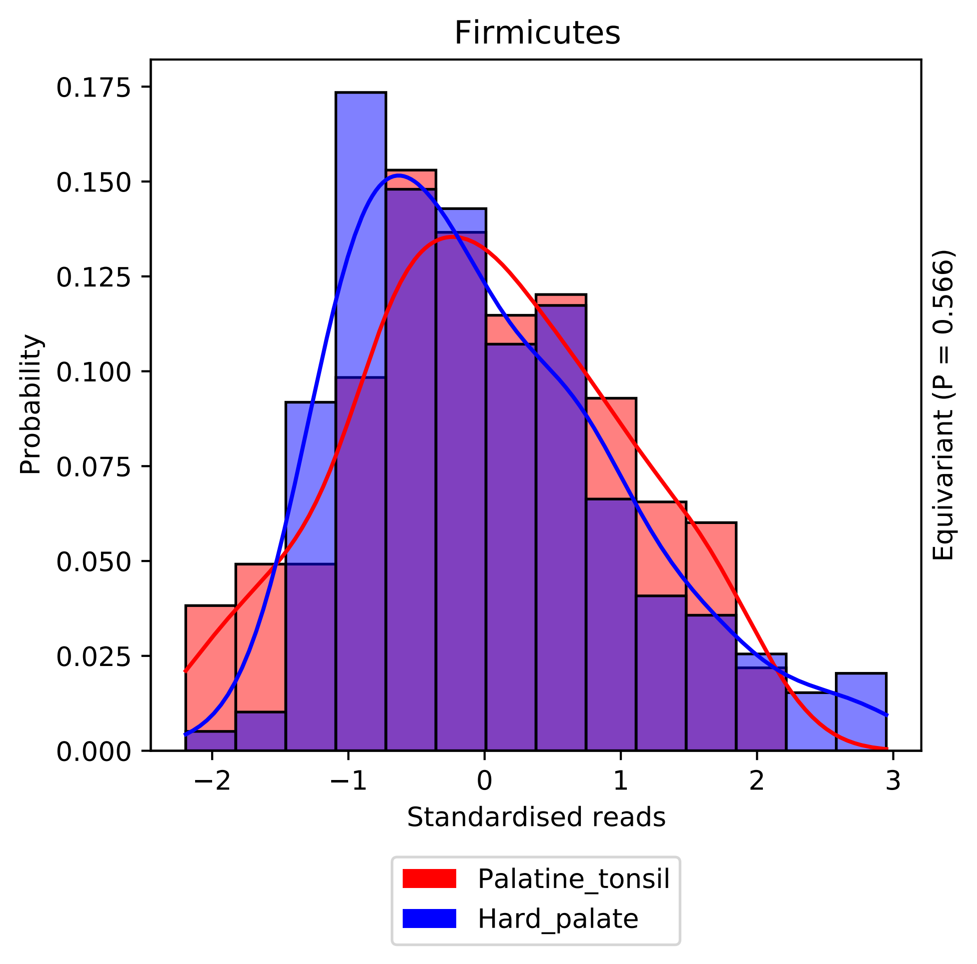

Firmicutes
0.175
0.150
0.125
0.100
0.075
0.050
0.025
0.000
Probability
−2
−1
0
1
2
3
Standardised reads
Equivariant (P = 0.566)
Palatine_tonsil
Hard_palate

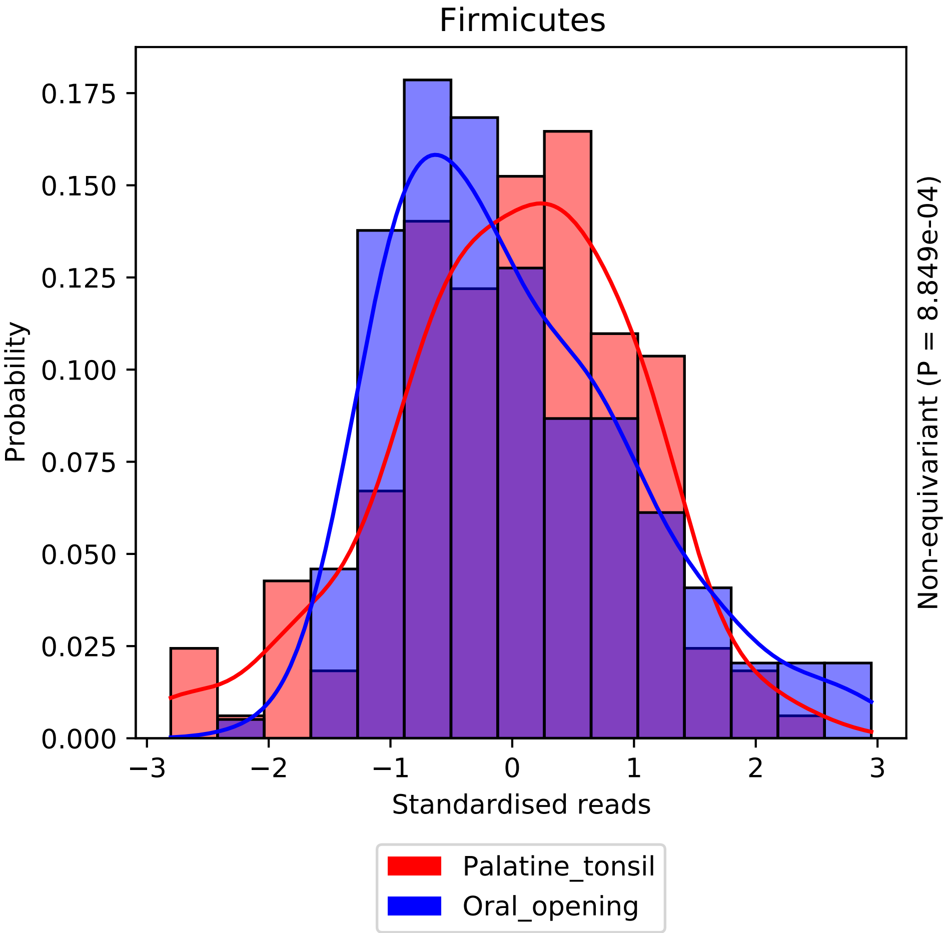
Firmicutes
Probability
0.000
0.025
0.050
0.075
0.100
0.125
0.150
0.175
−3
−2
−1
0
1
2
3
Standardised reads
Non-equivariant (P = 8.849e-04)
Palatine_tonsil
Oral_opening

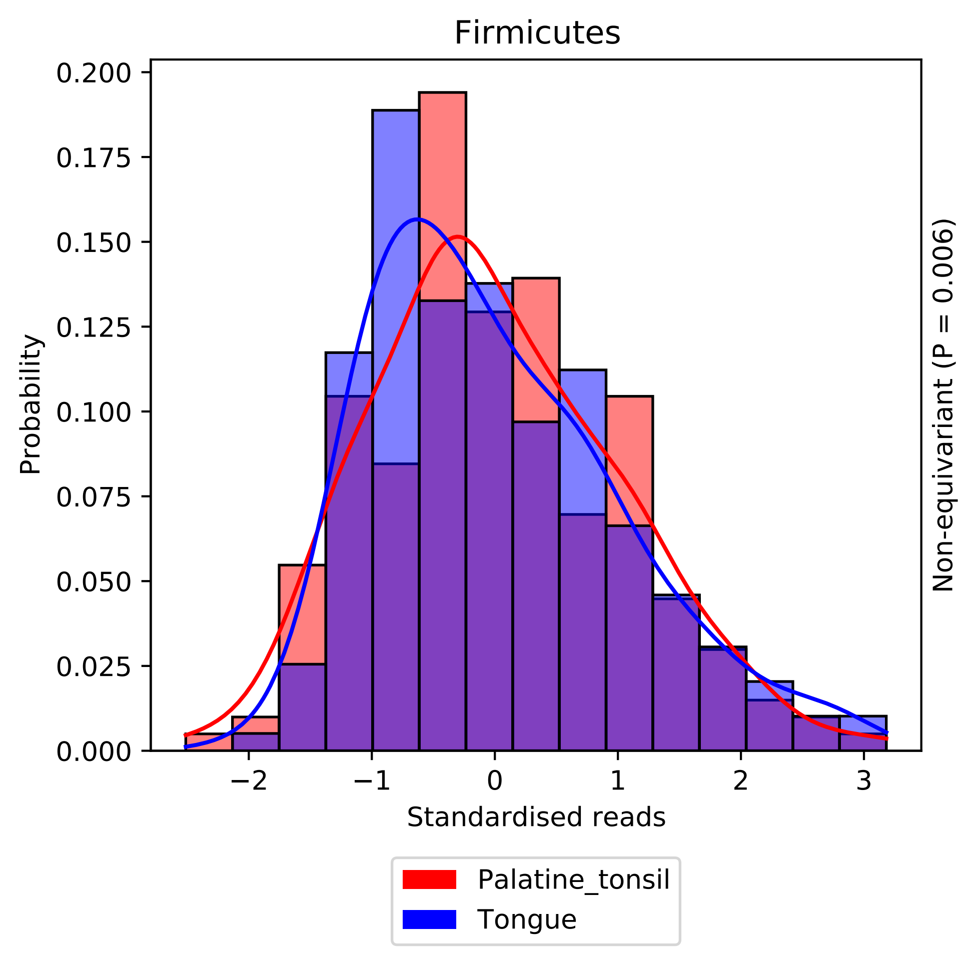

Firmicutes
Probability
0.200
0.175
0.150
0.125
0.100
0.075
0.050
0.025
0.000
−2
−1
0
1
2
3
Standardised reads
Non-equivariant (P = 0.006)
Palatine_tonsil
Tongue

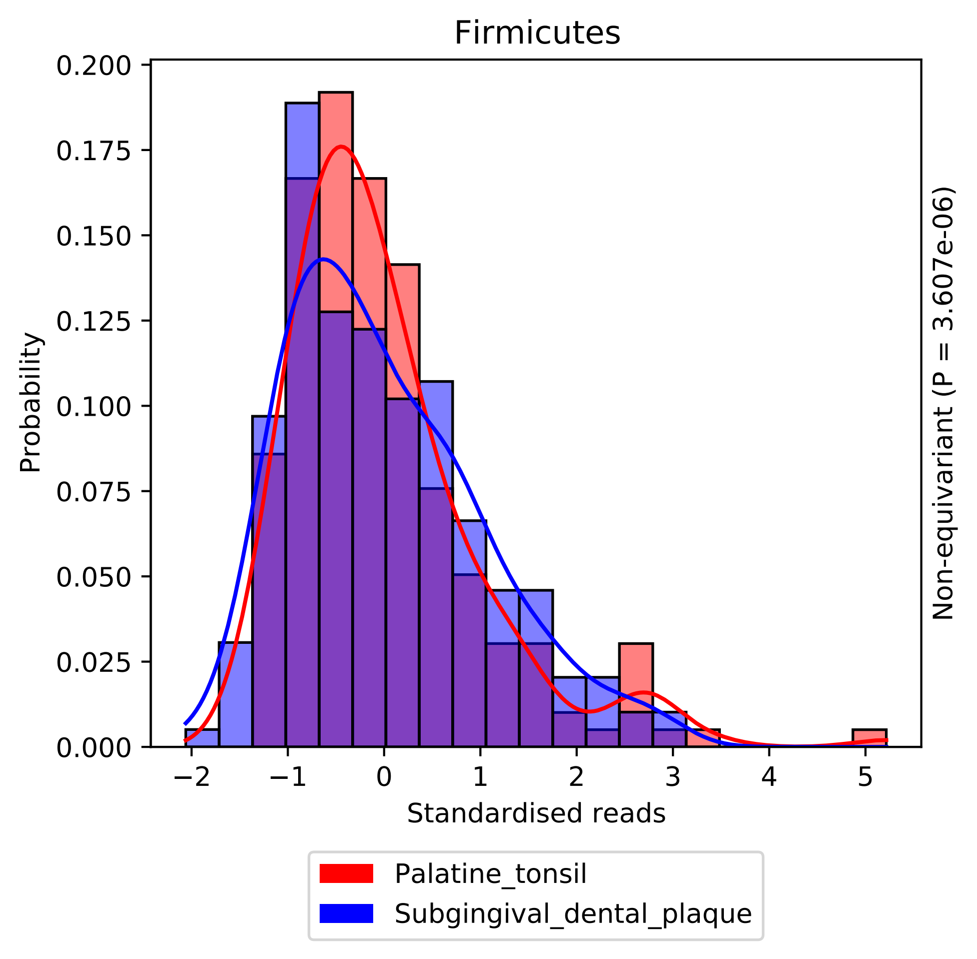

Firmicutes
Probability
0.200
0.175
0.150
0.125
0.100
0.075
0.050
0.025
0.000
−2
−1
0
1
2
3
4
5
Standardised reads
Non-equivariant (P = 3.607e-06)
Palatine_tonsil
Subgingival_dental_plaque

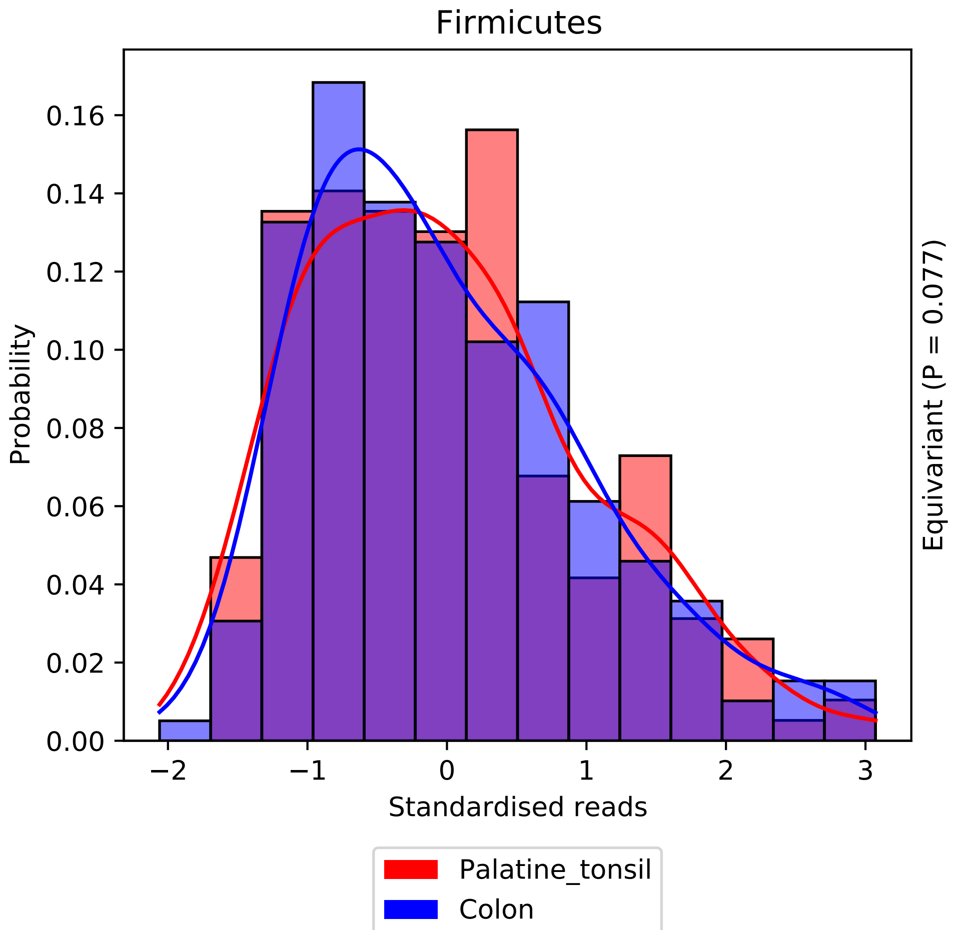

Firmicutes
Probability
0.00
0.02
0.04
0.06
0.08
0.10
0.12
0.14
0.16
−2
−1
0
1
2
3
Standardised reads
Equivariant (P = 0.077)
Palatine_tonsil
Colon

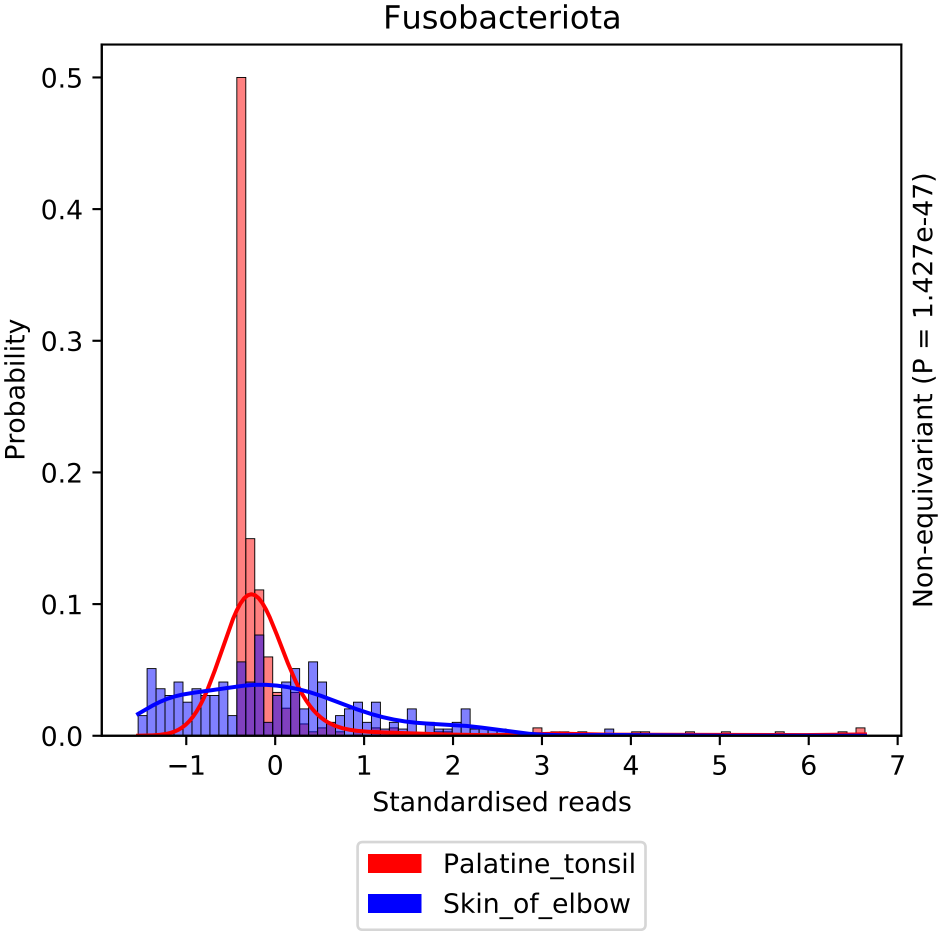
Fusobacteriota
Probability
Standardised reads
Non-equivariant (P = 1.427e-47)
Palatine_tonsil
Skin_of_elbow
0.0
0.1
0.2
0.3
0.4
0.5
−1
0
1
2
3
4
5
6
7

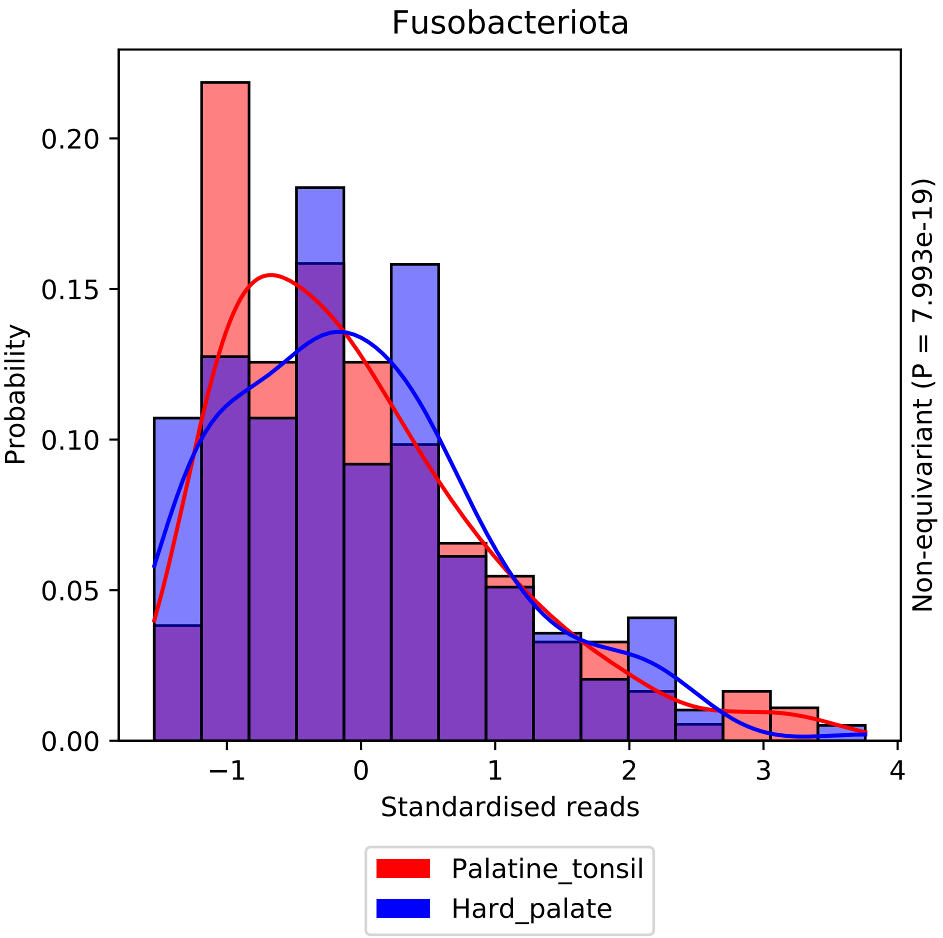

Fusobacteriota
Probability
0.00
0.05
0.10
0.15
0.20
−1
0
1
2
3
4
Standardised reads
Non-equivariant (P = 7.993e-19)
Palatine_tonsil
Hard_palate

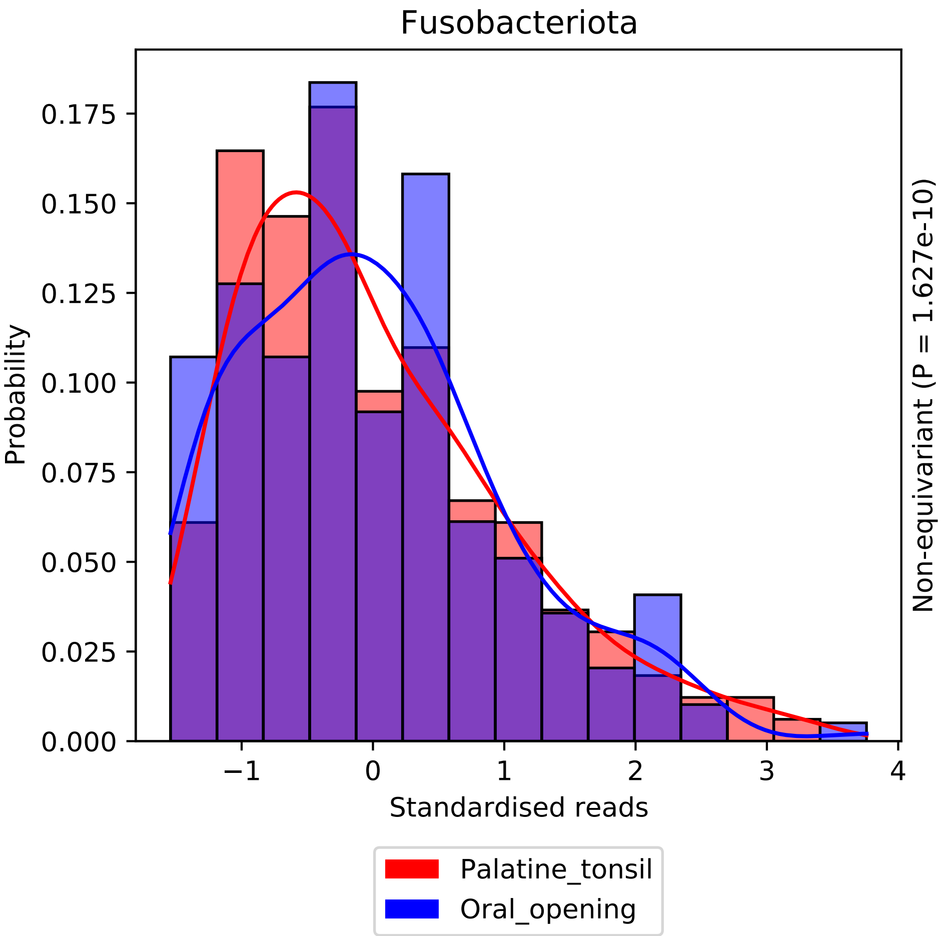

Fusobacteriota
Probability
0.000
0.025
0.050
0.075
0.100
0.125
0.150
0.175
−1
0
1
2
3
4
Standardised reads
Non-equivariant (P = 1.627e-10)
Palatine_tonsil
Oral_opening

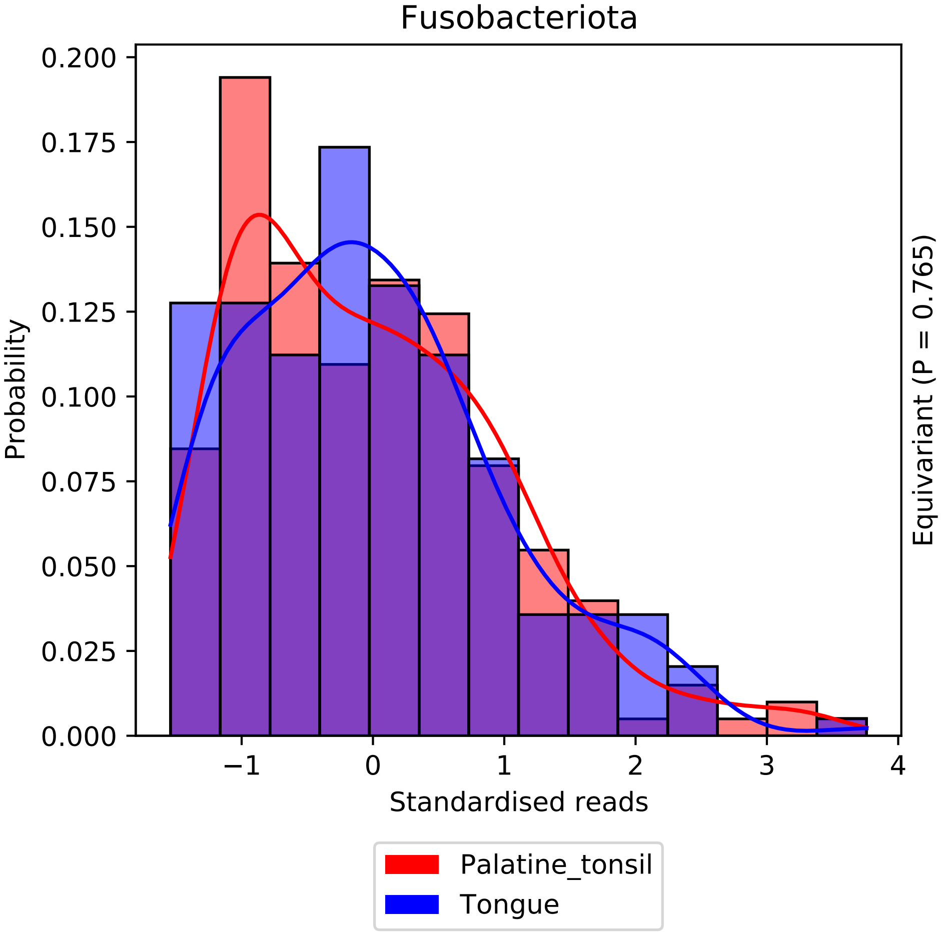
Fusobacteriota
Probability
Standardised reads
Equivariant (P = 0.765)
Palatine_tonsil
Tongue
0.000
0.025
0.050
0.075
0.100
0.125
0.150
0.175
0.200
−1
0
1
2
3
4

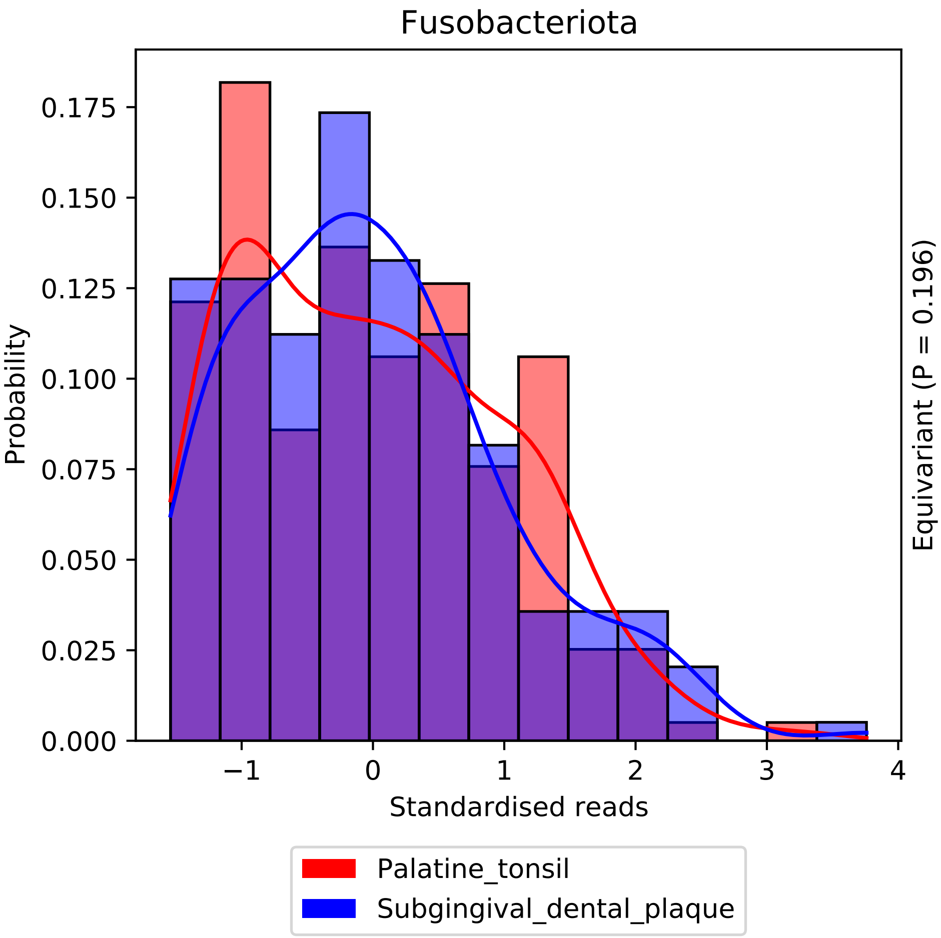
Fusobacteriota
Probability
Standardised reads
Equivariant (P = 0.196)
0.000
0.025
0.050
0.075
0.100
0.125
0.150
0.175
−1
0
1
2
3
4
Palatine_tonsil
Subgingival_dental_plaque

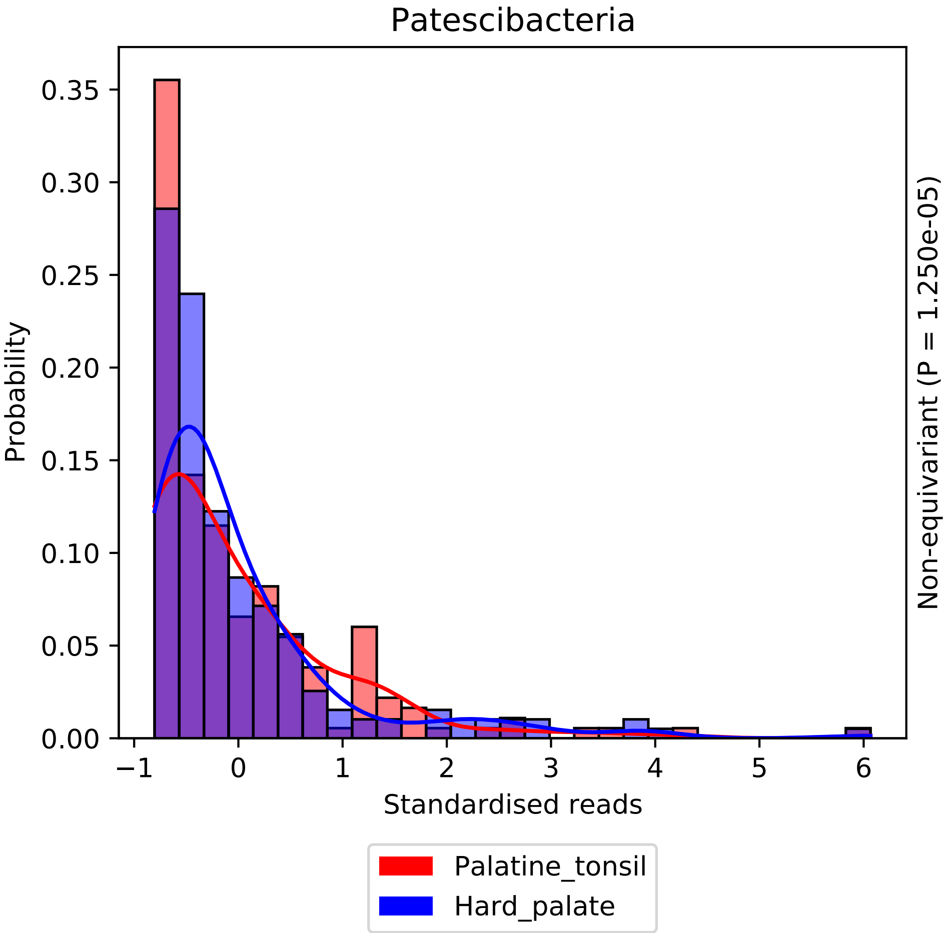

Patescibacteria
Probability
Standardised reads
Non-equivariant (P = 1.250e-05)
Palatine_tonsil
Hard_palate
0.00
0.05
0.10
0.15
0.20
0.25
0.30
0.35
−1
0
1
2
3
4
5
6

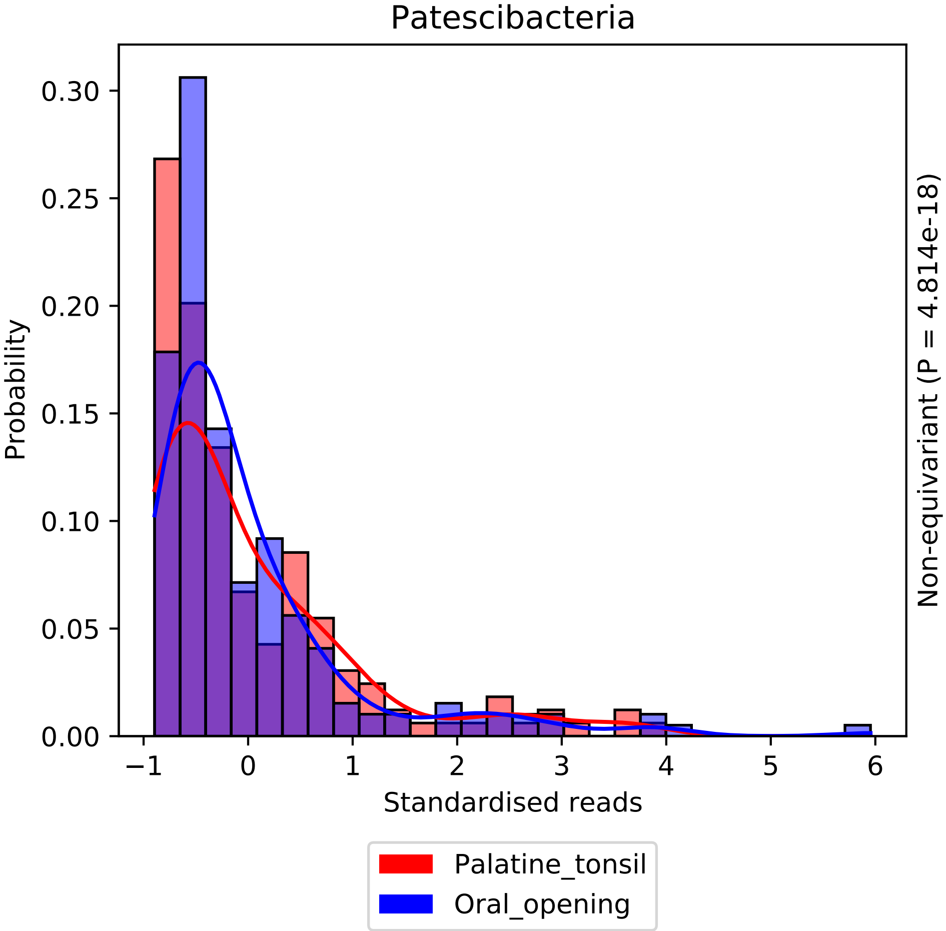

Patescibacteria
Probability
Standardised reads
Non-equivariant (P = 4.814e-18)
0.00
0.05
0.10
0.15
0.20
0.25
0.30
−1
0
1
2
3
4
5
6
Palatine_tonsil
Oral_opening

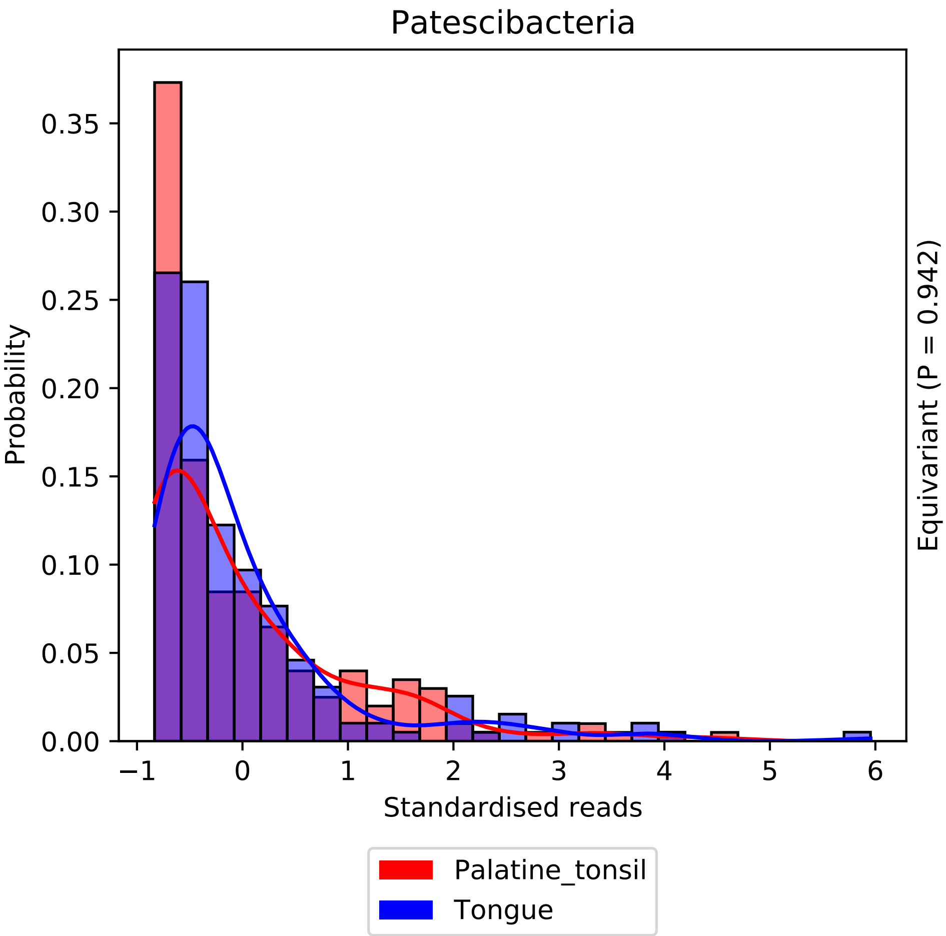
Patescibacteria
Probability
0.00
0.05
0.10
0.15
0.20
0.25
0.30
0.35
−1
0
1
2
3
4
5
6
Standardised reads
Equivariant (P = 0.942)
Palatine_tonsil
Tongue

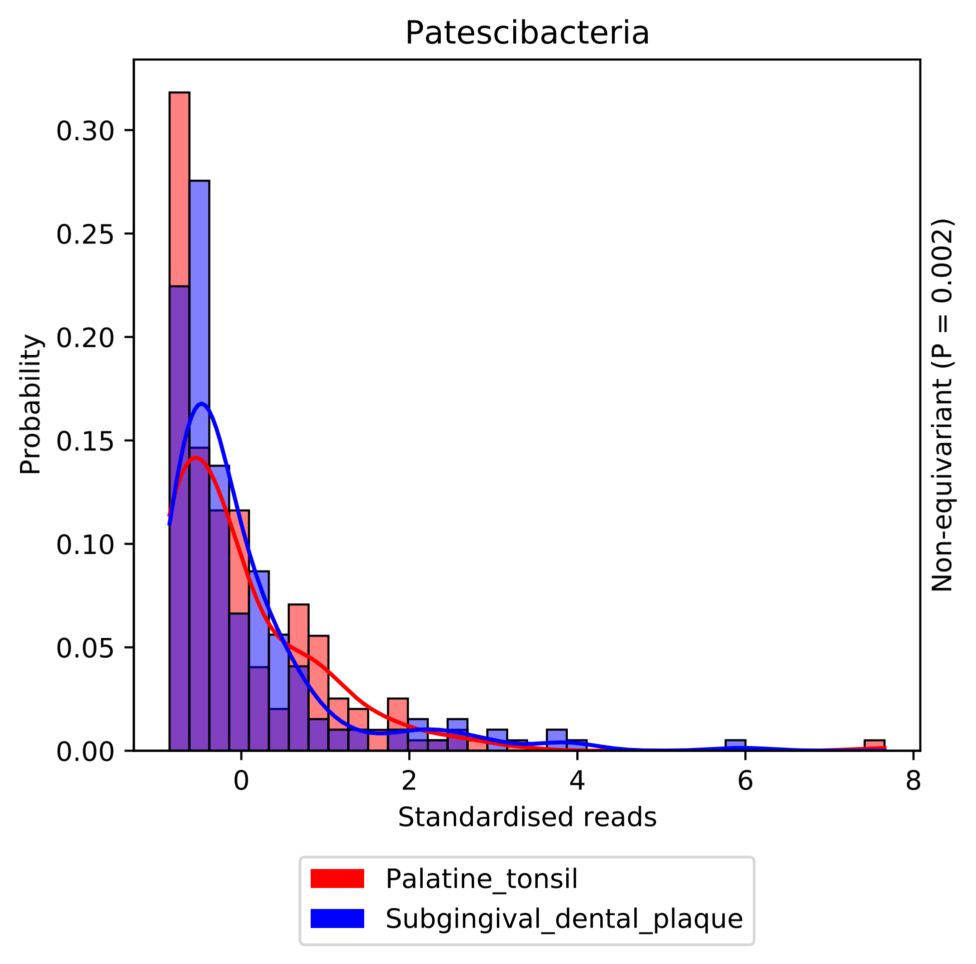
Patescibacteria
Probability
0.00
0.05
0.10
0.15
0.20
0.25
0.30
0
2
4
6
8
Standardised reads
Non-equivariant (P = 0.002)
Palatine_tonsil
Subgingival_dental_plaque

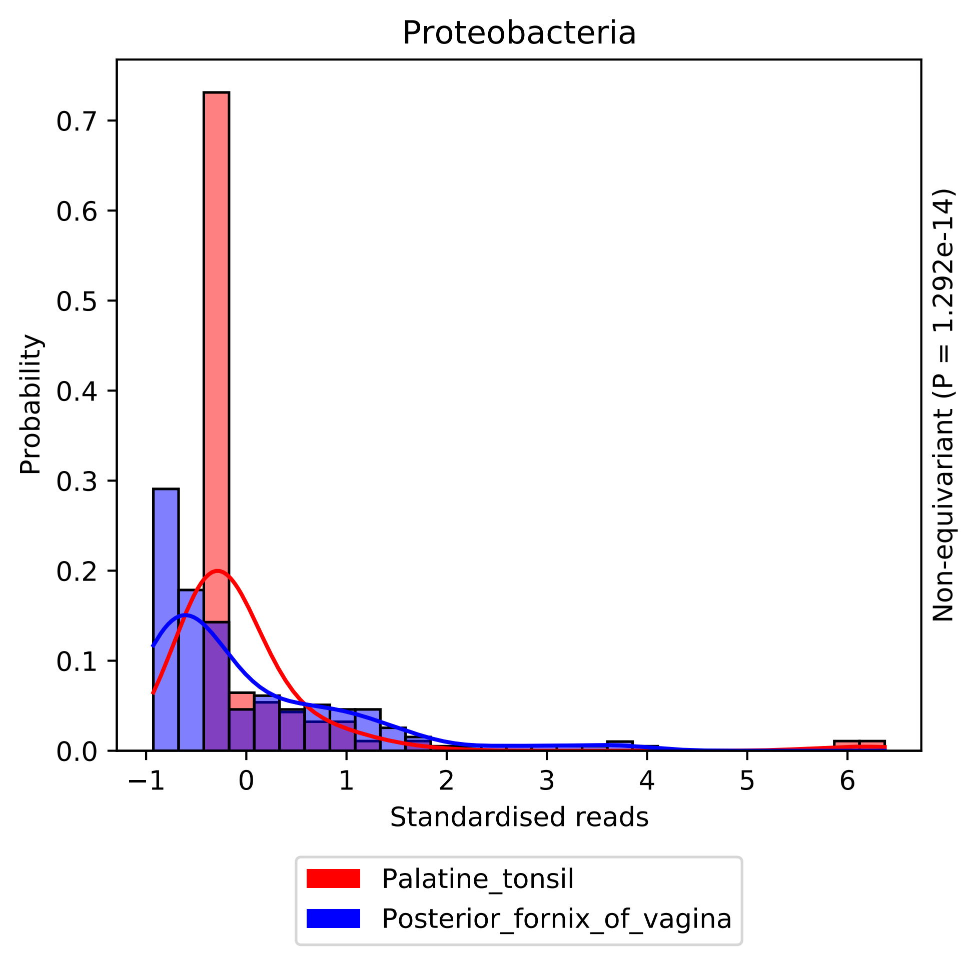

Proteobacteria
Probability
0.0
0.1
0.2
0.3
0.4
0.5
0.6
0.7
−1
0
1
2
3
4
5
6
Standardised reads
Non-equivariant (P = 1.292e-14)
Palatine_tonsil
Posterior_fornix_of_vagina

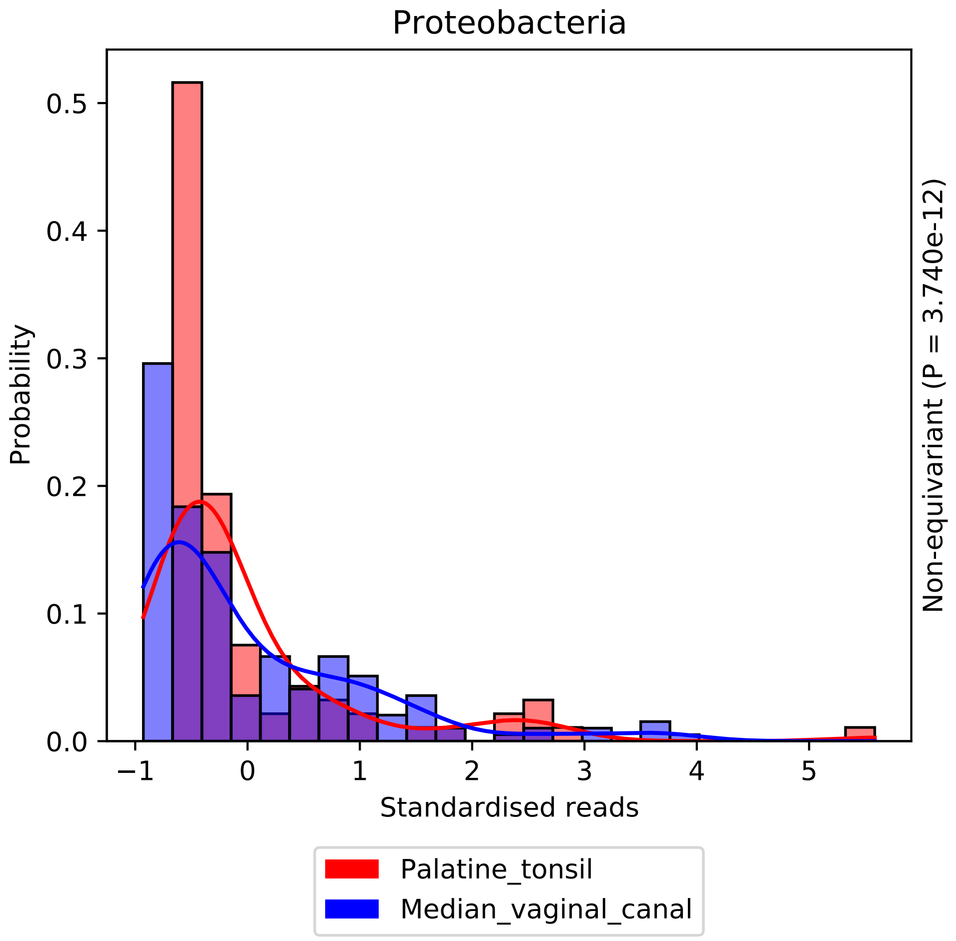

Proteobacteria
Probability
0.0
0.1
0.2
0.3
0.4
0.5
−1
0
1
2
3
4
5
Standardised reads
Non-equivariant (P = 3.740e-12)
Palatine_tonsil
Median_vaginal_canal

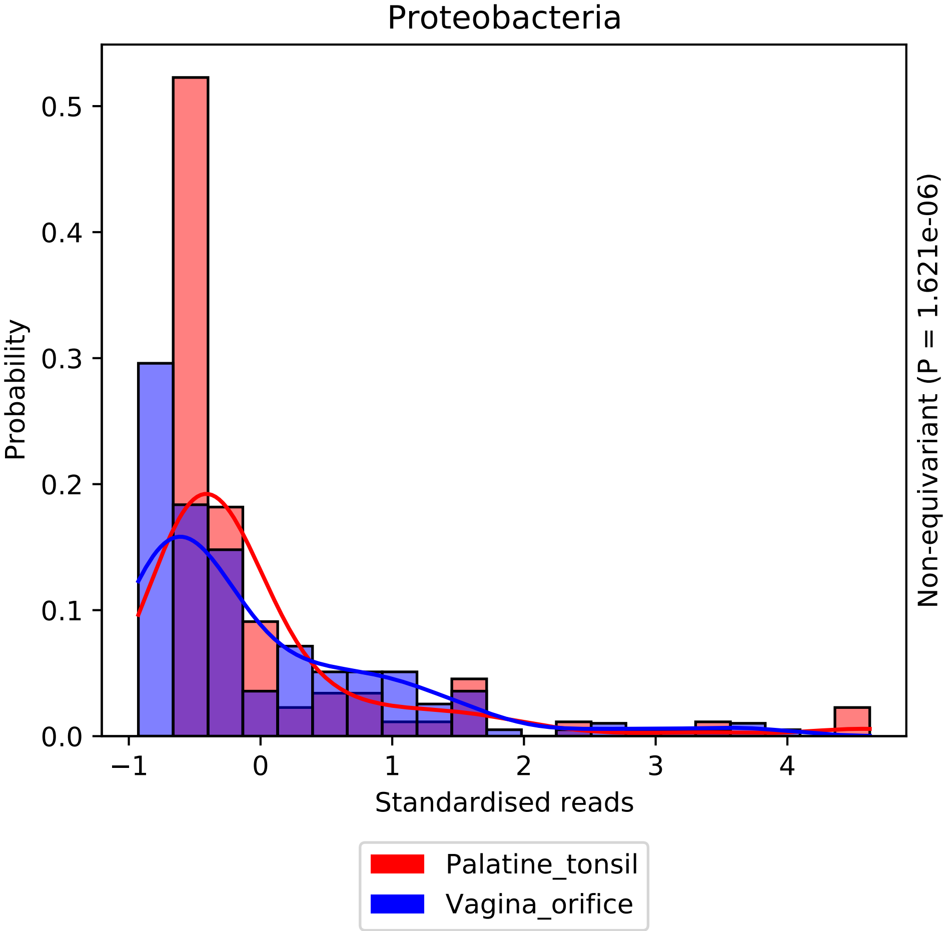

Proteobacteria
Probability
0.0
0.1
0.2
0.3
0.4
0.5
−1
0
1
2
3
4
Standardised reads
Non-equivariant (P = 1.621e-06)
Palatine_tonsil
Vagina_orifice

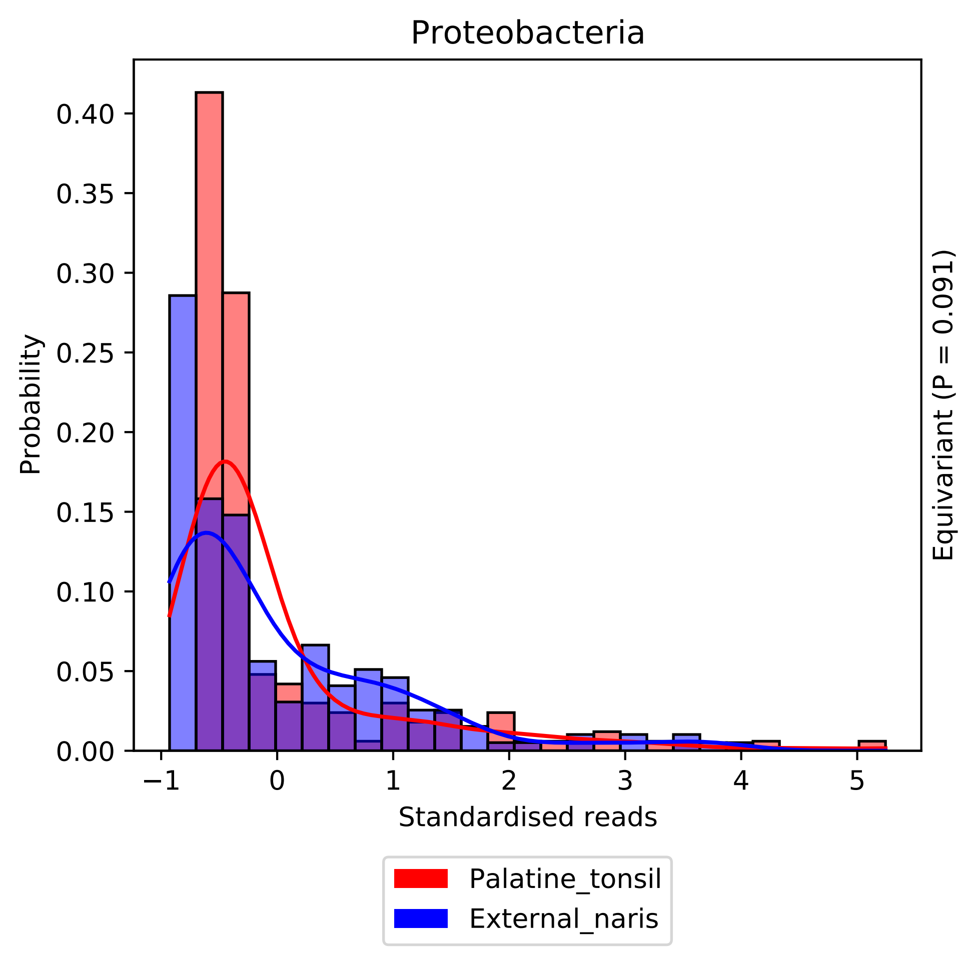

Proteobacteria
Probability
0.00
0.05
0.10
0.15
0.20
0.25
0.30
0.35
0.40
−1
0
1
2
3
4
5
Standardised reads
Equivariant (P = 0.091)
Palatine_tonsil
External_naris

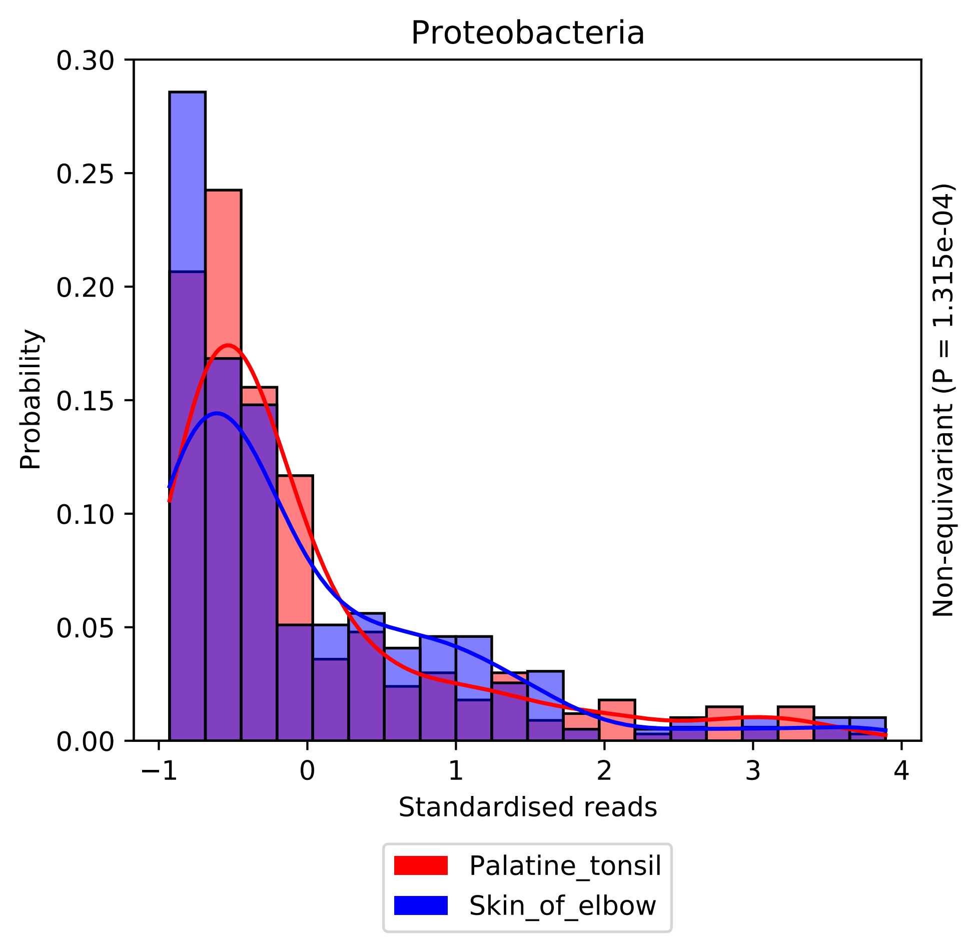

Proteobacteria
Probability
0.30
0.25
0.20
0.15
0.10
0.05
0.00
−1
0
1
2
3
4
Standardised reads
Non-equivariant (P = 1.315e-04)
Palatine_tonsil
Skin_of_elbow

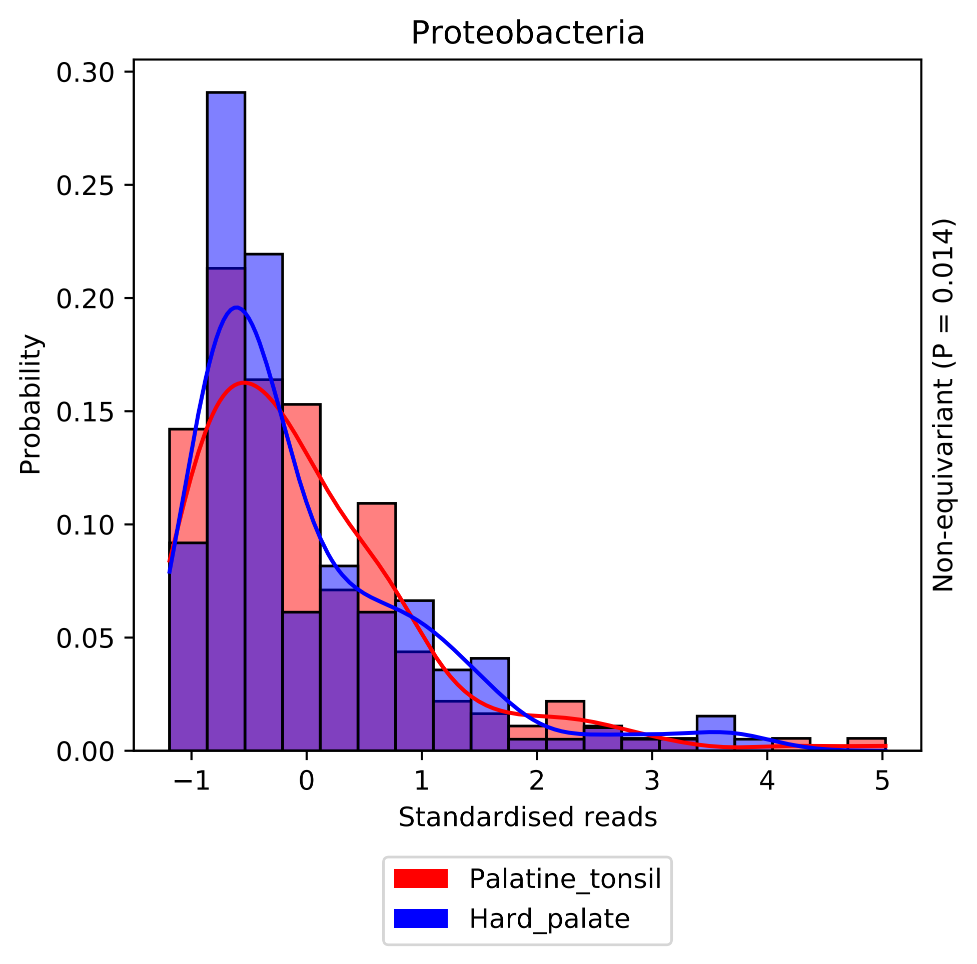
Proteobacteria
0.30
0.25
0.20
0.15
0.10
0.05
0.00
Probability
−1
0
1
2
3
4
5
Standardised reads
Non-equivariant (P = 0.014)
Palatine_tonsil
Hard_palate

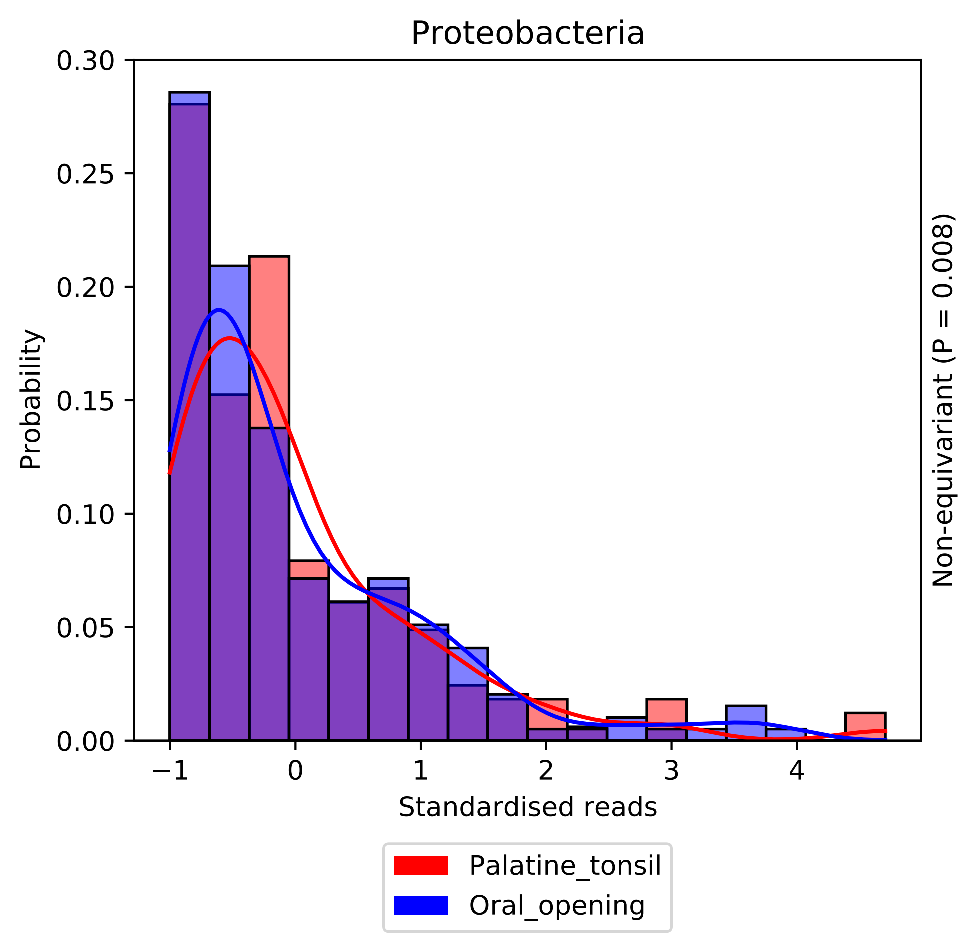
Proteobacteria
0.30
0.25
0.20
0.15
0.10
0.05
0.00
Probability
−1
0
1
2
3
4
Standardised reads
Non-equivariant (P = 0.008)
Palatine_tonsil
Oral_opening

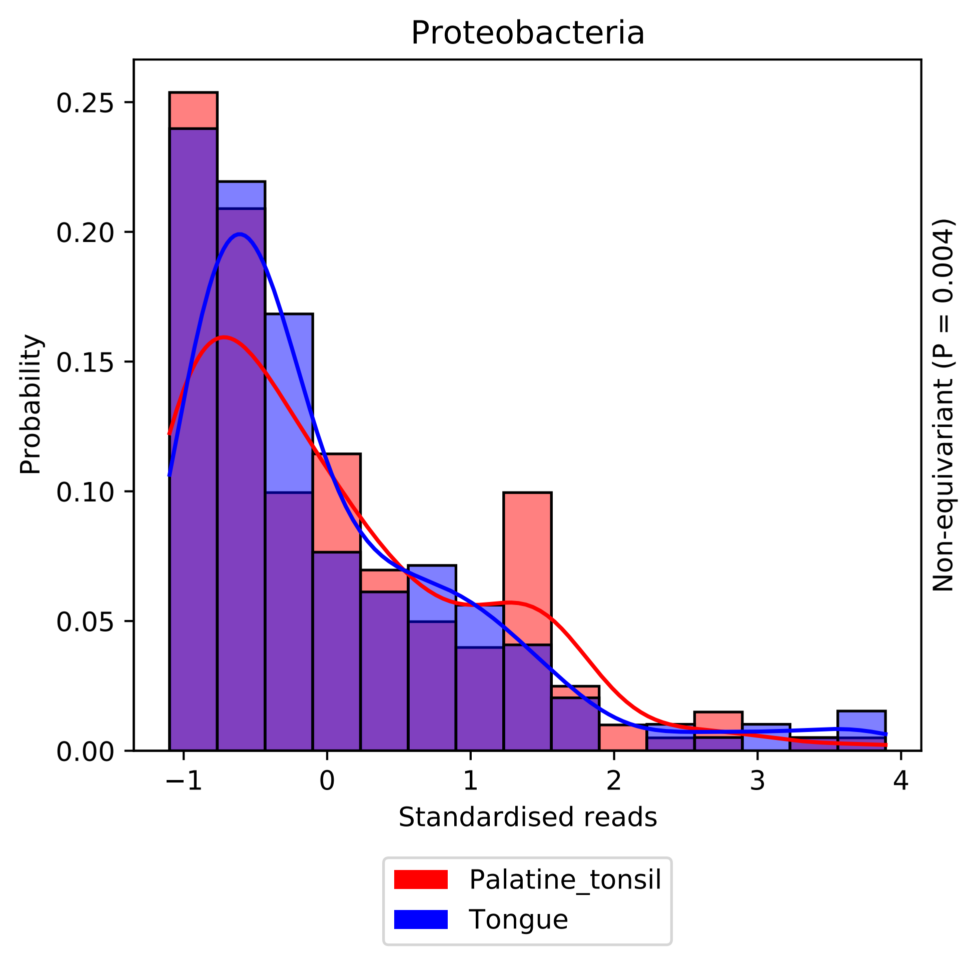

Proteobacteria
Probability
0.25
0.20
0.15
0.10
0.05
0.00
−1
0
1
2
3
4
Standardised reads
Non-equivariant (P = 0.004)
Palatine_tonsil
Tongue

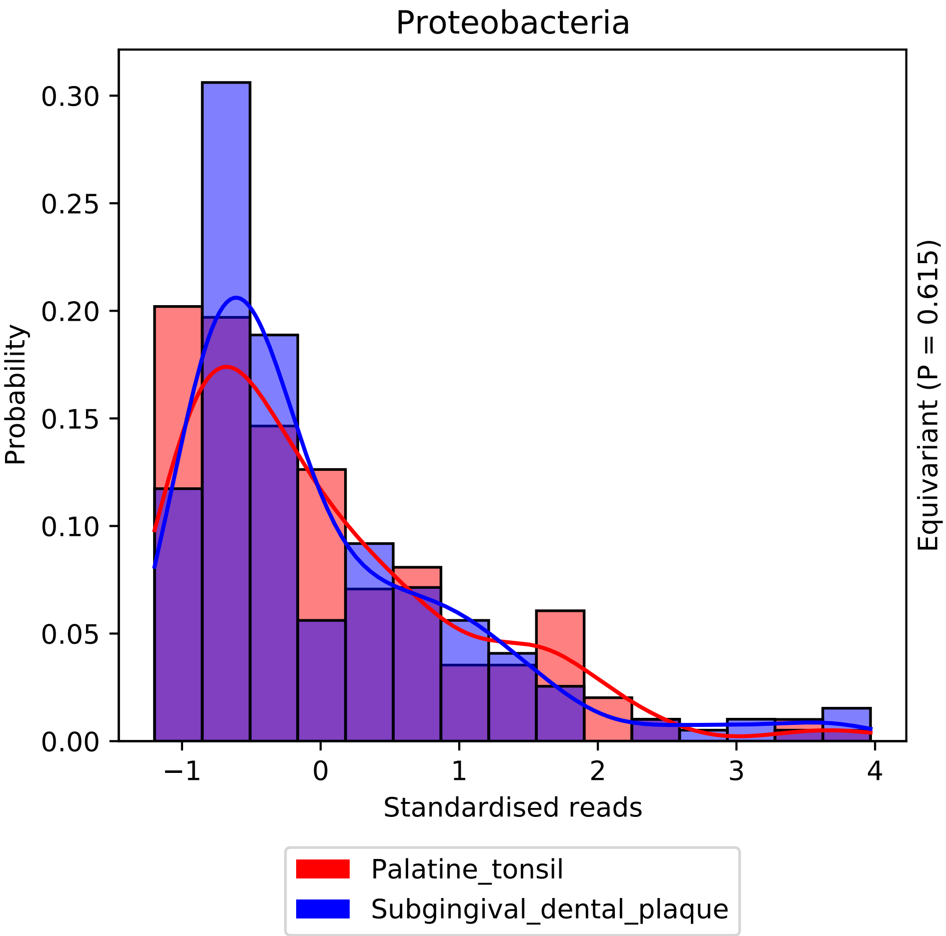

Proteobacteria
Probability
0.00
0.05
0.10
0.15
0.20
0.25
0.30
−1
0
1
2
3
4
Standardised reads
Equivariant (P = 0.615)
Palatine_tonsil
Subgingival_dental_plaque

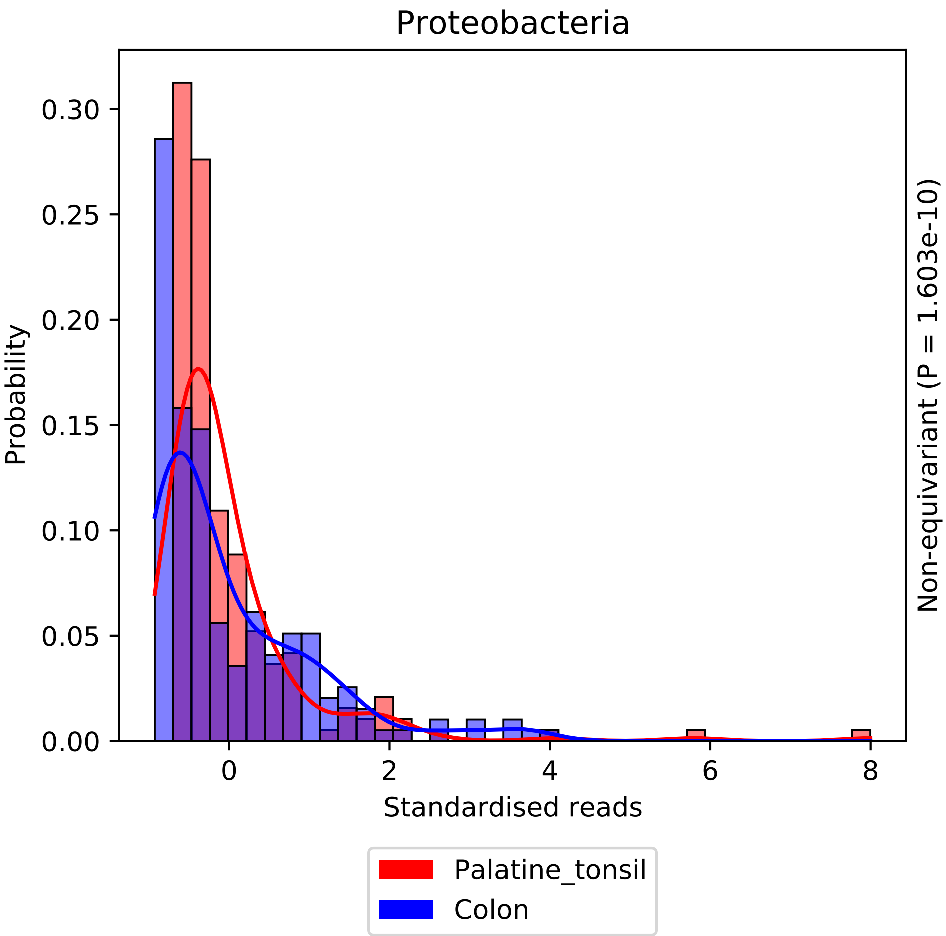

Proteobacteria
Probability
0.00
0.05
0.10
0.15
0.20
0.25
0.30
0
2
4
6
8
Standardised reads
Non-equivariant (P = 1.603e-10)
Palatine_tonsil
Colon

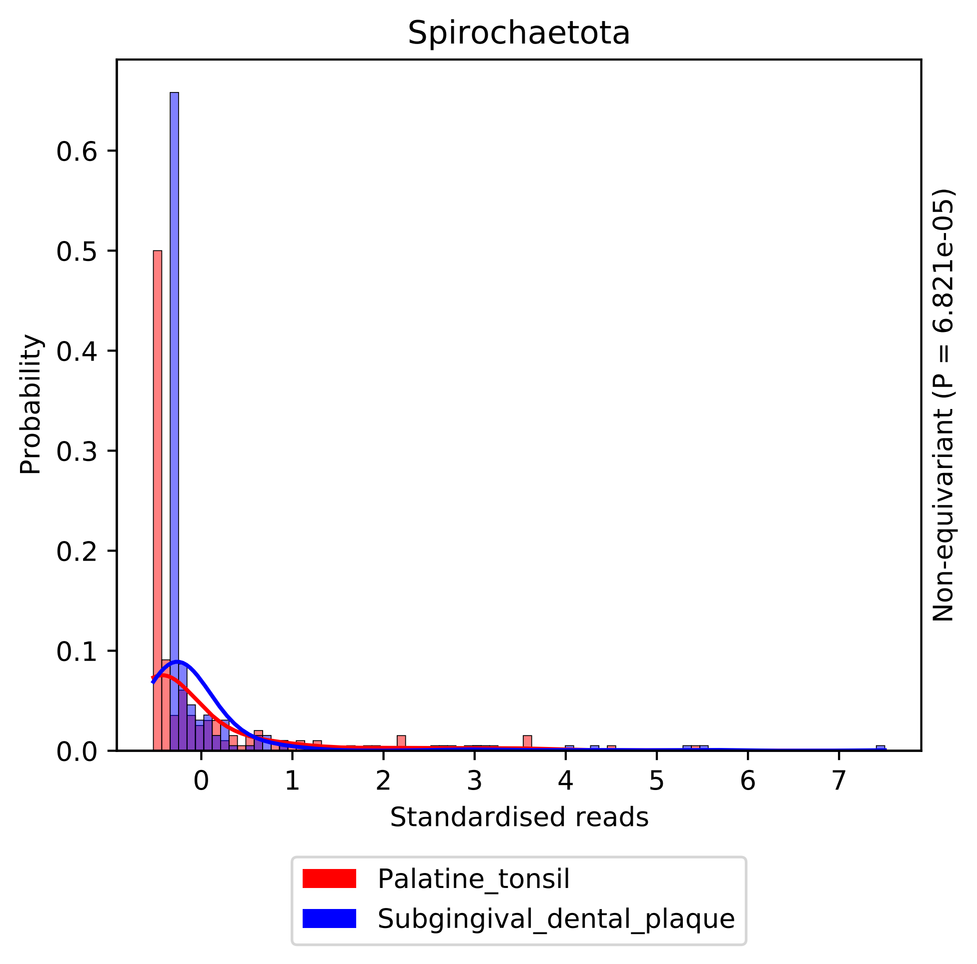

Spirochaetota
Probability
0.0
0.1
0.2
0.3
0.4
0.5
0.6
0
1
2
3
4
5
6
7
Standardised reads
Non-equivariant (P = 6.821e-05)
Palatine_tonsil
Subgingival_dental_plaque

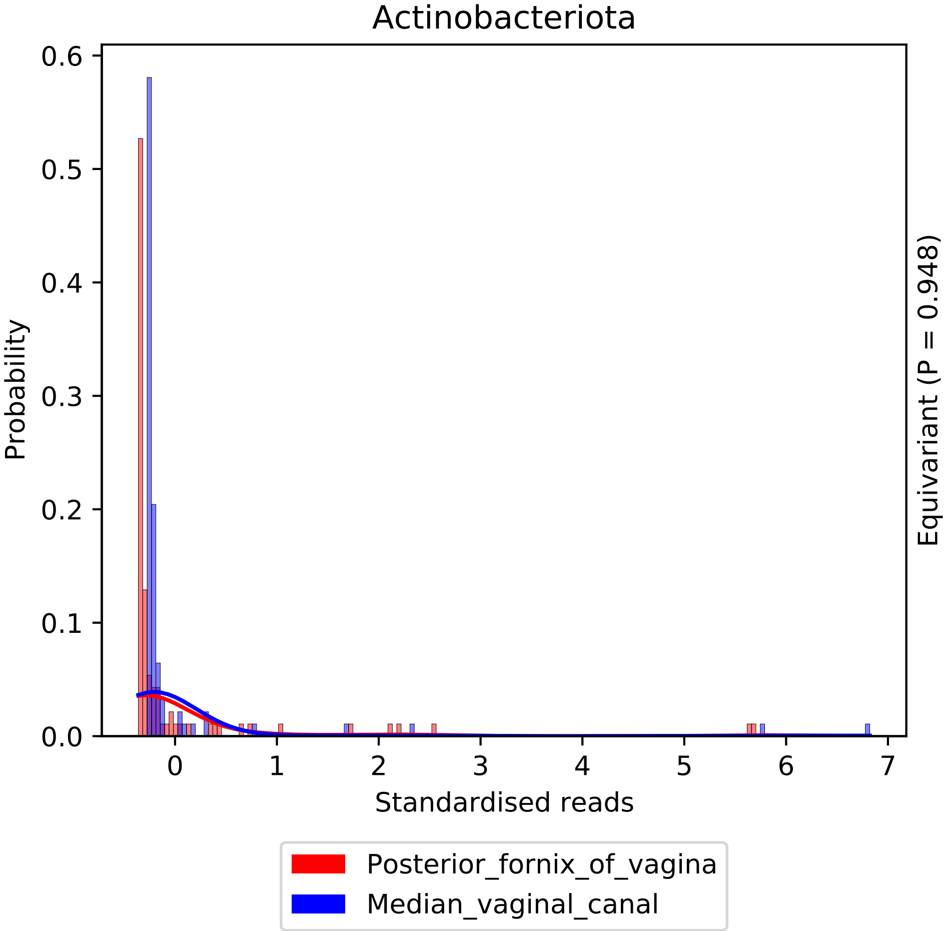

Actinobacteriota
Probability
Standardised reads
Equivariant (P = 0.948)
0.0
0.1
0.2
0.3
0.4
0.5
0.6
0
1
2
3
4
5
6
7
Posterior_fornix_of_vagina
Median_vaginal_canal

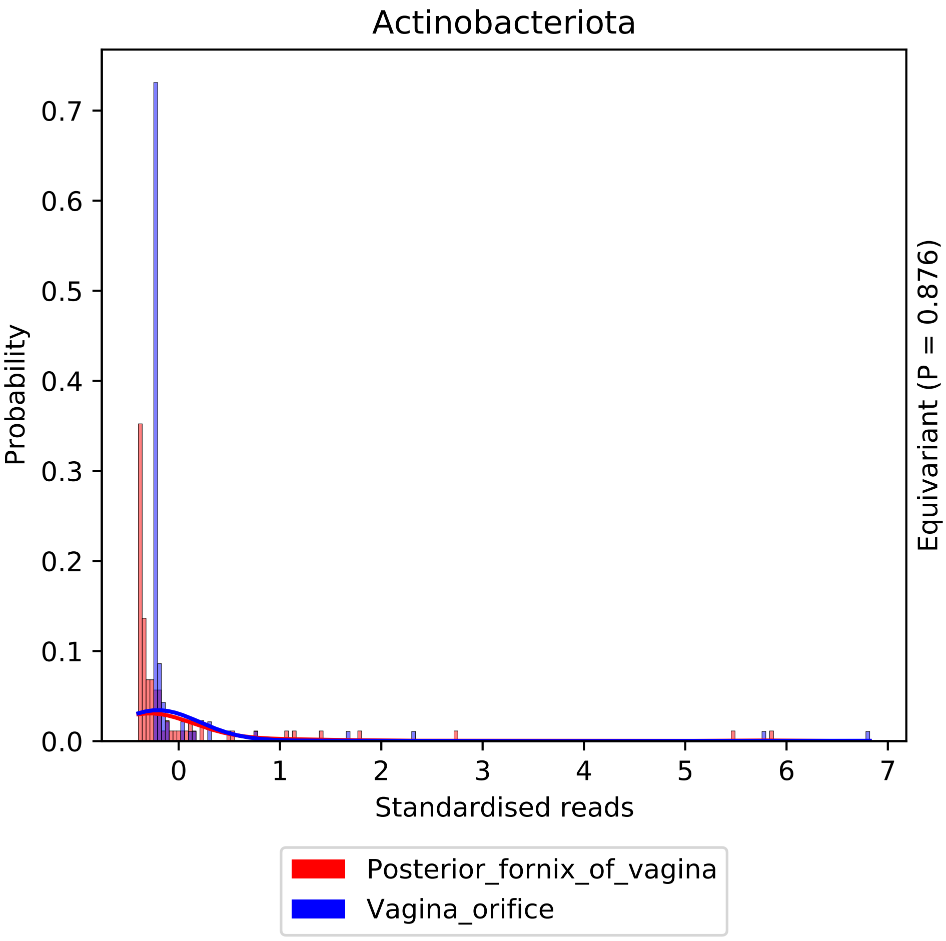

Actinobacteriota
Probability
0.0
0.1
0.2
0.3
0.4
0.5
0.6
0.7
0
1
2
3
4
5
6
7
Standardised reads
Equivariant (P = 0.876)
Posterior_fornix_of_vagina
Vagina_orifice

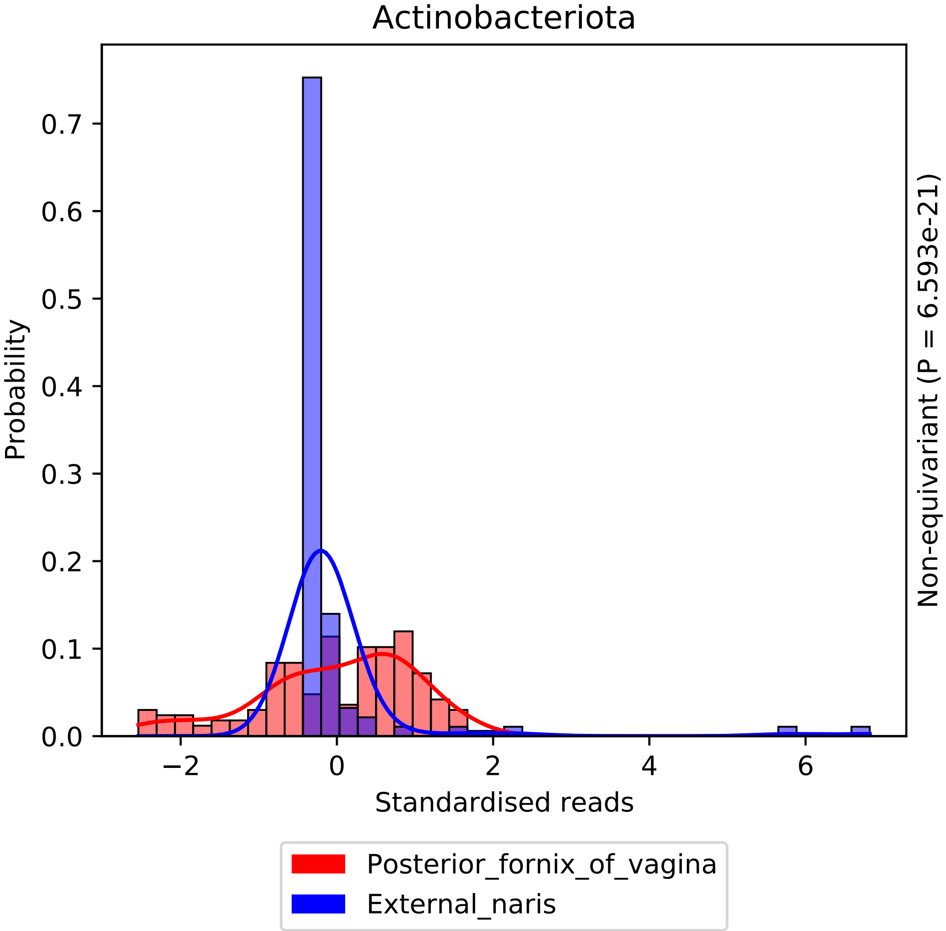

Actinobacteriota
Probability
0.0
0.1
0.2
0.3
0.4
0.5
0.6
0.7
−2
0
2
4
6
Standardised reads
Non-equivariant (P = 6.593e-21)
Posterior_fornix_of_vagina
External_naris

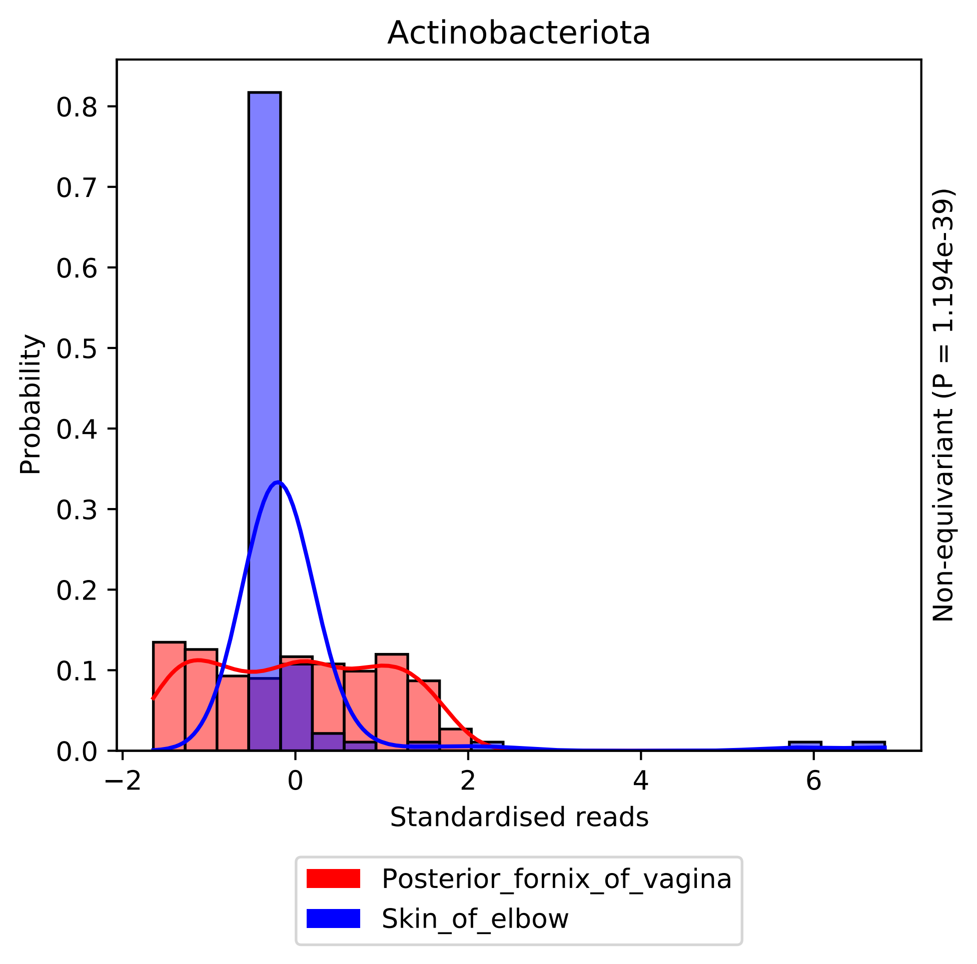

Actinobacteriota
Probability
0.0
0.1
0.2
0.3
0.4
0.5
0.6
0.7
0.8
−2
0
2
4
6
Standardised reads
Non-equivariant (P = 1.194e-39)
Posterior_fornix_of_vagina
Skin_of_elbow

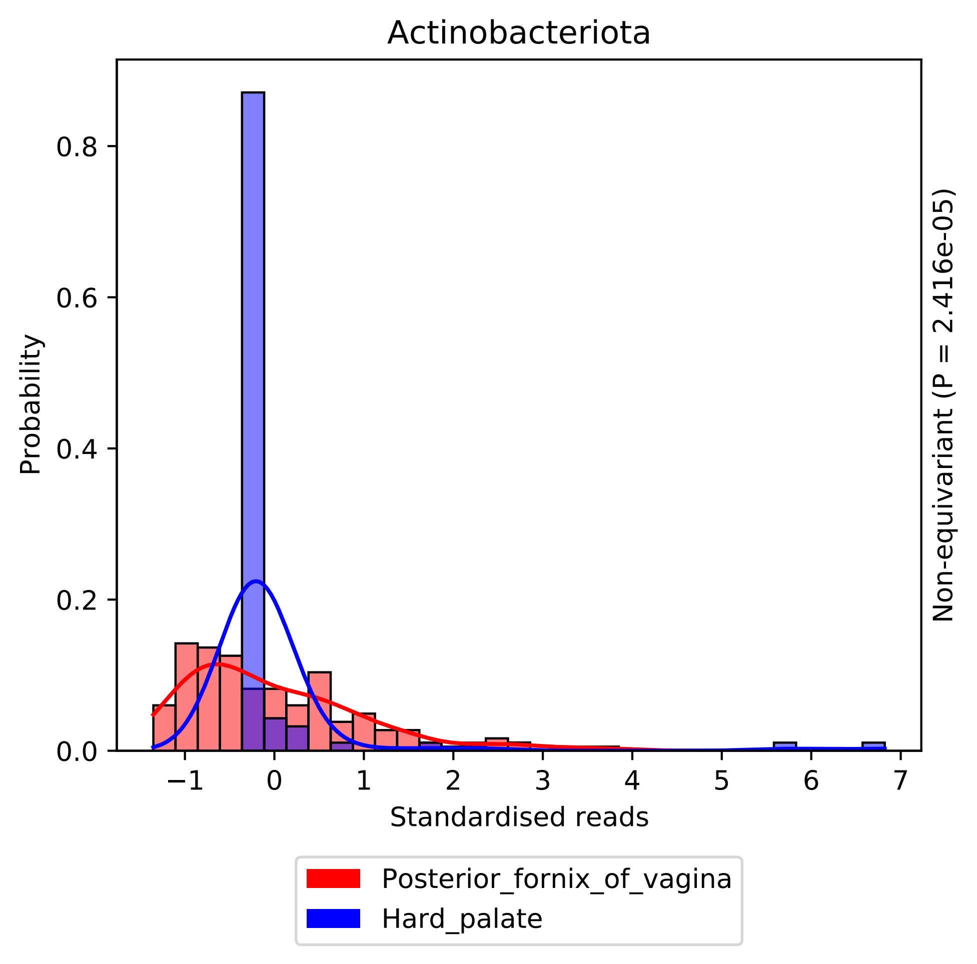
Actinobacteriota
Probability
0.0
0.2
0.4
0.6
0.8
−1
0
1
2
3
4
5
6
7
Standardised reads
Non-equivariant (P = 2.416e-05)
Posterior_fornix_of_vagina
Hard_palate

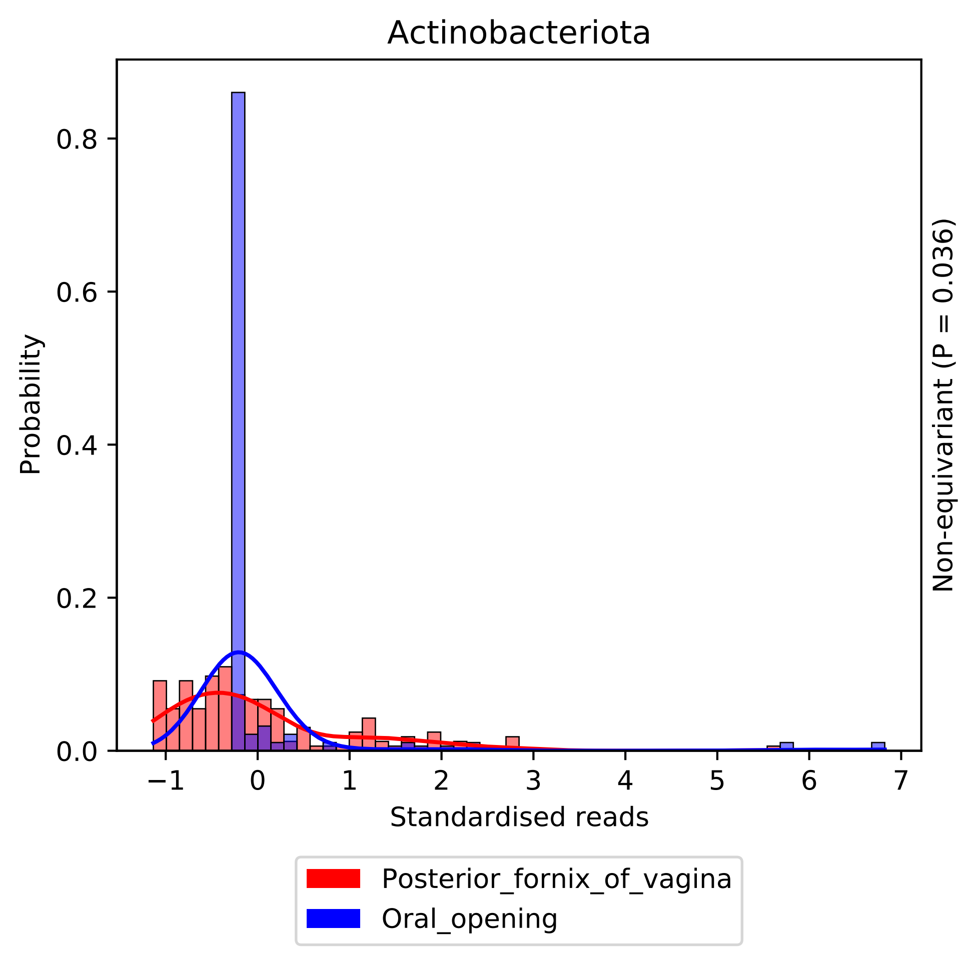
Actinobacteriota
Probability
0.0
0.2
0.4
0.6
0.8
−1
0
1
2
3
4
5
6
7
Standardised reads
Non-equivariant (P = 0.036)
Posterior_fornix_of_vagina
Oral_opening

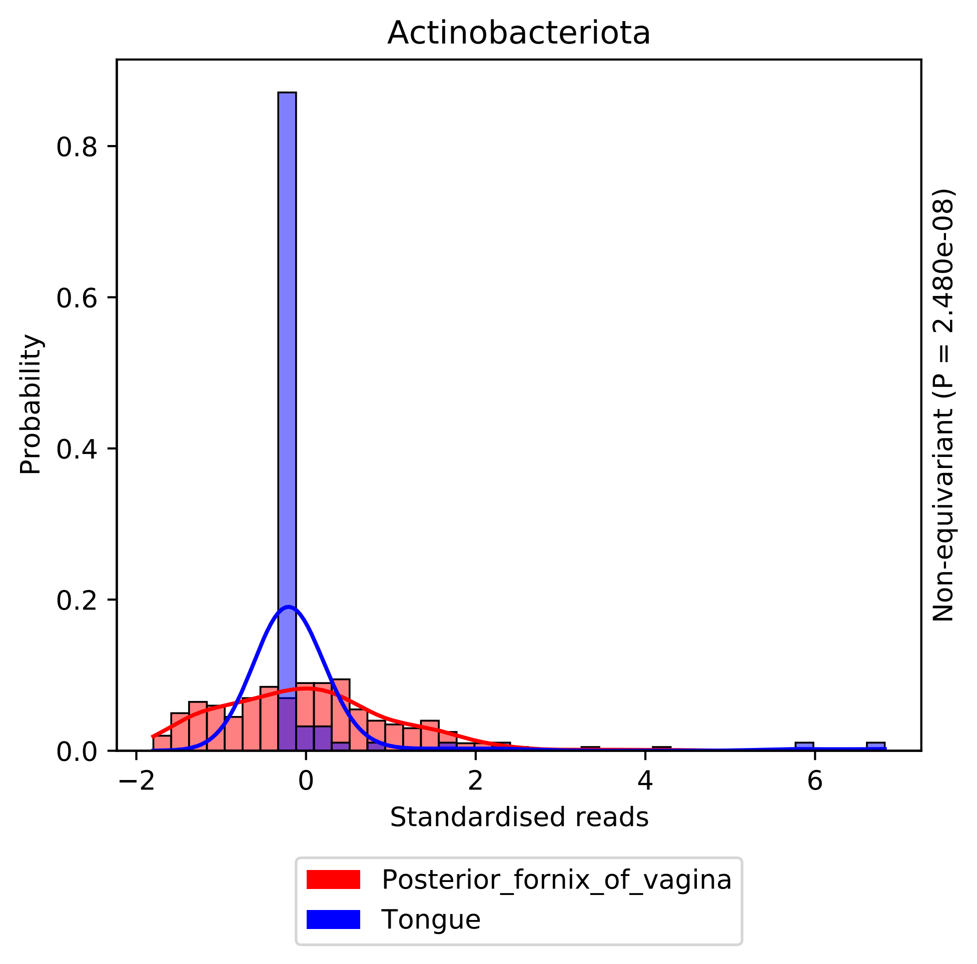
Actinobacteriota
Probability
0.0
0.2
0.4
0.6
0.8
−2
0
2
4
6
Standardised reads
Non-equivariant (P = 2.480e-08)
Posterior_fornix_of_vagina
Tongue

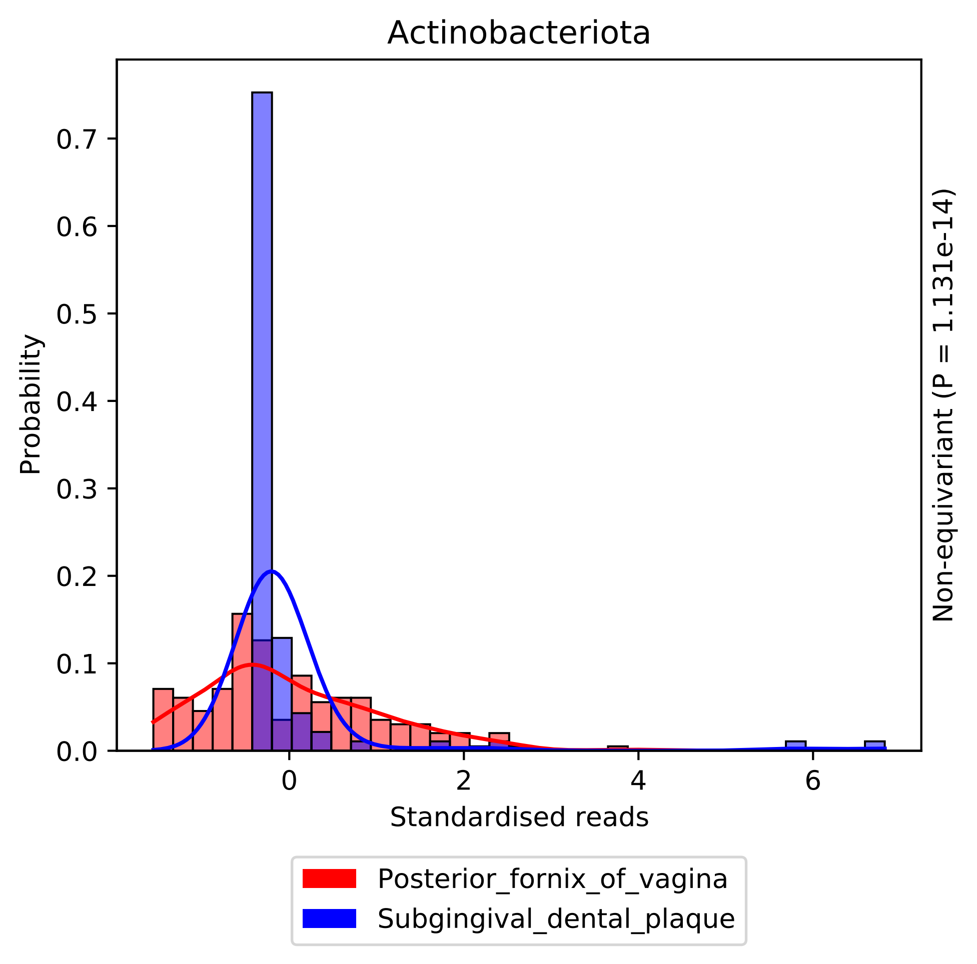

Actinobacteriota
Probability
0.0
0.1
0.2
0.3
0.4
0.5
0.6
0.7
0
2
4
6
Standardised reads
Non-equivariant (P = 1.131e-14)
Posterior_fornix_of_vagina
Subgingival_dental_plaque

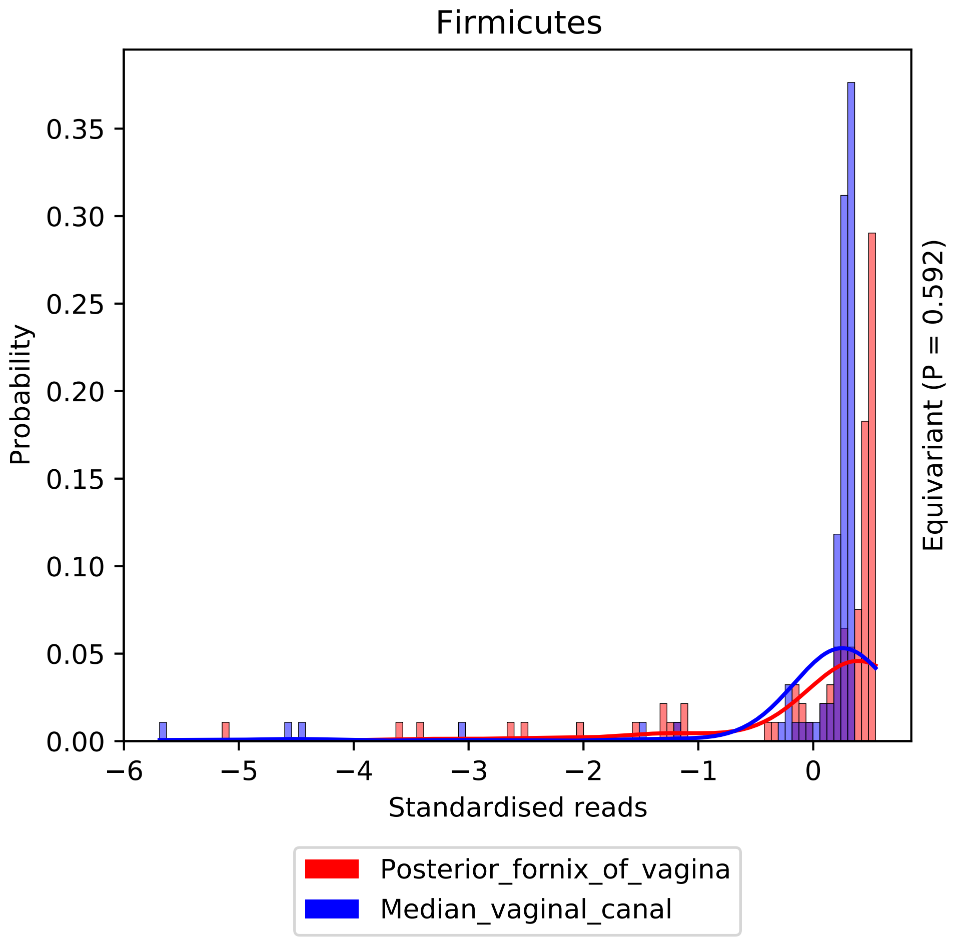

Firmicutes
0.35
0.30
0.25
0.20
0.15
0.10
0.05
0.00
Probability
−6
−5
−4
−3
−2
−1
0
Standardised reads
Equivariant (P = 0.592)
Posterior_fornix_of_vagina
Median_vaginal_canal

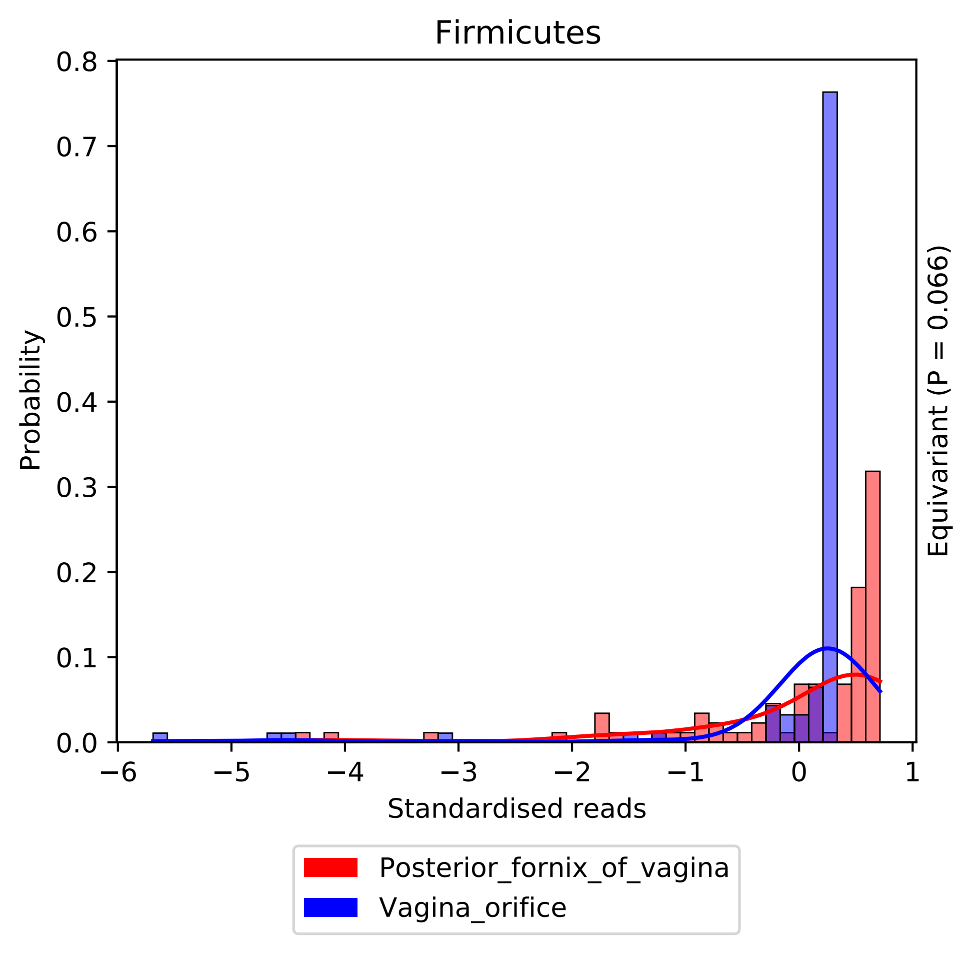

Firmicutes
Probability
0.8
0.7
0.6
0.5
0.4
0.3
0.2
0.1
0.0
−6
−5
−4
−3
−2
−1
0
1
Standardised reads
Equivariant (P = 0.066)
Posterior_fornix_of_vagina
Vagina_orifice

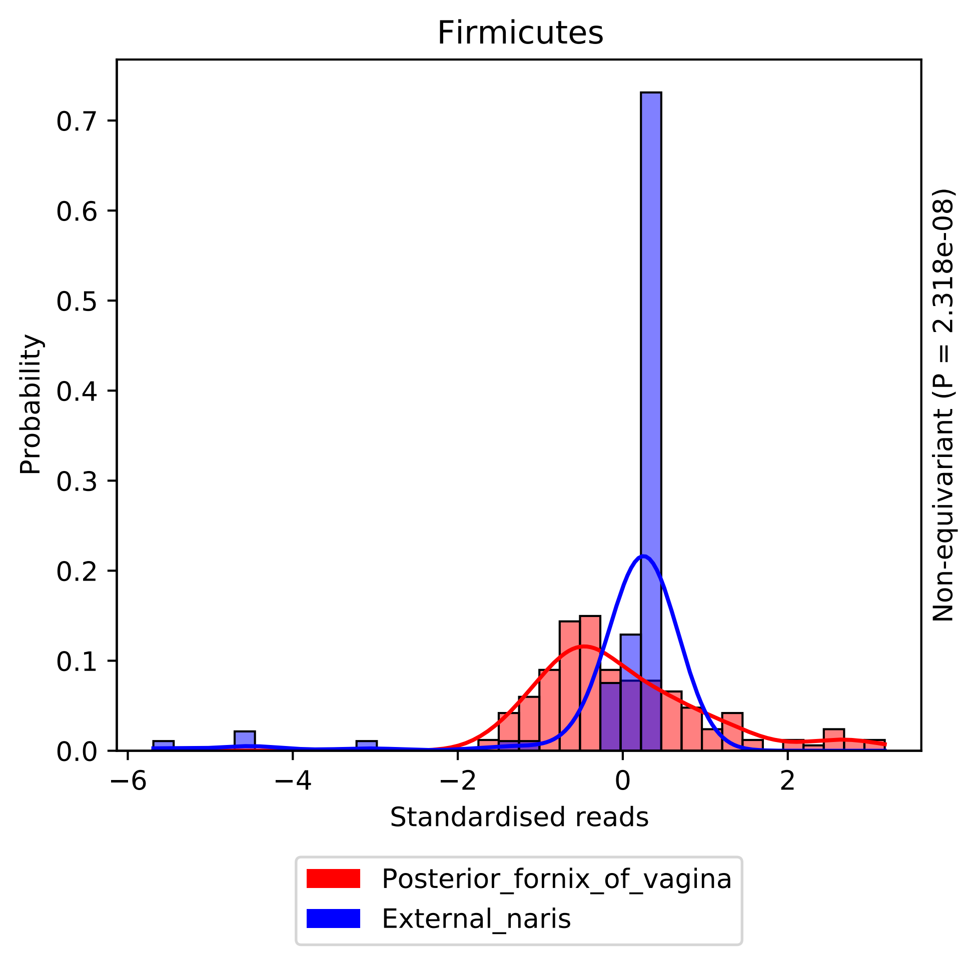
Firmicutes
Probability
0.0
0.1
0.2
0.3
0.4
0.5
0.6
0.7
−6
−4
−2
0
2
Standardised reads
Non-equivariant (P = 2.318e-08)
Posterior_fornix_of_vagina
External_naris

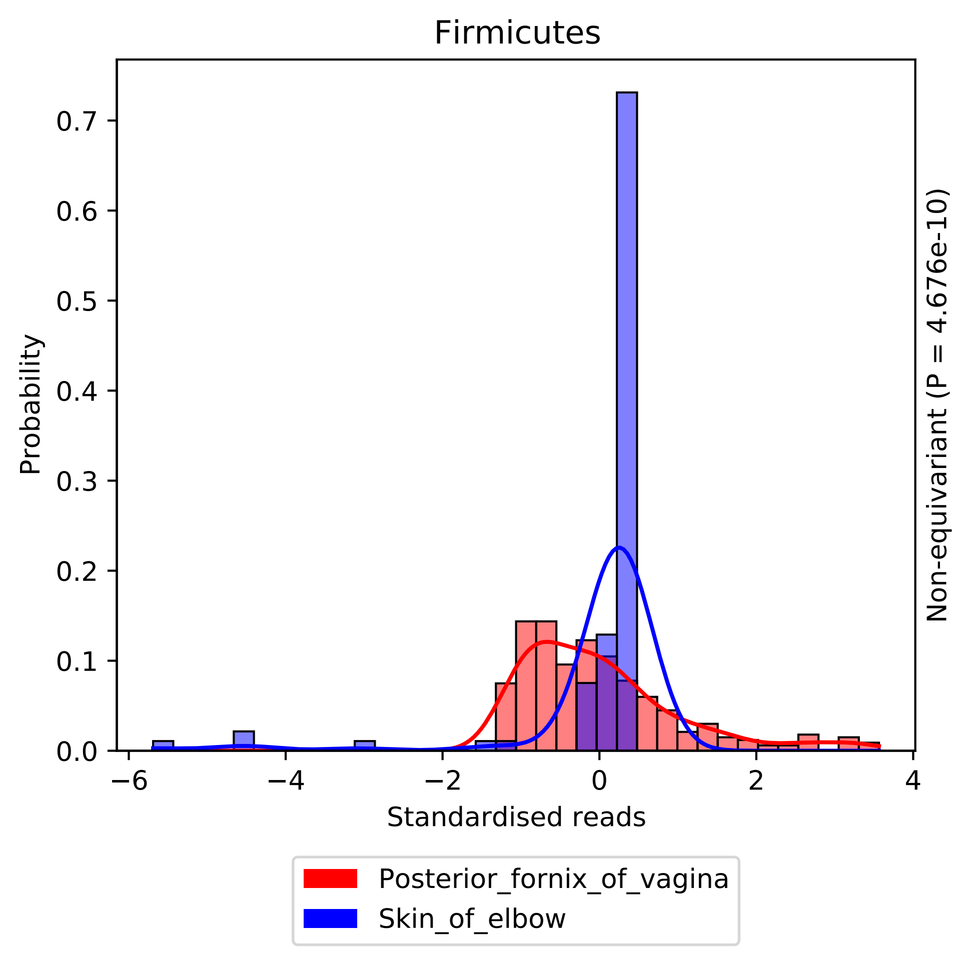

Firmicutes
Probability
Standardised reads
Non-equivariant (P = 4.676e-10)
0.0
0.1
0.2
0.3
0.4
0.5
0.6
0.7
−6
−4
−2
0
2
4
Posterior_fornix_of_vagina
Skin_of_elbow

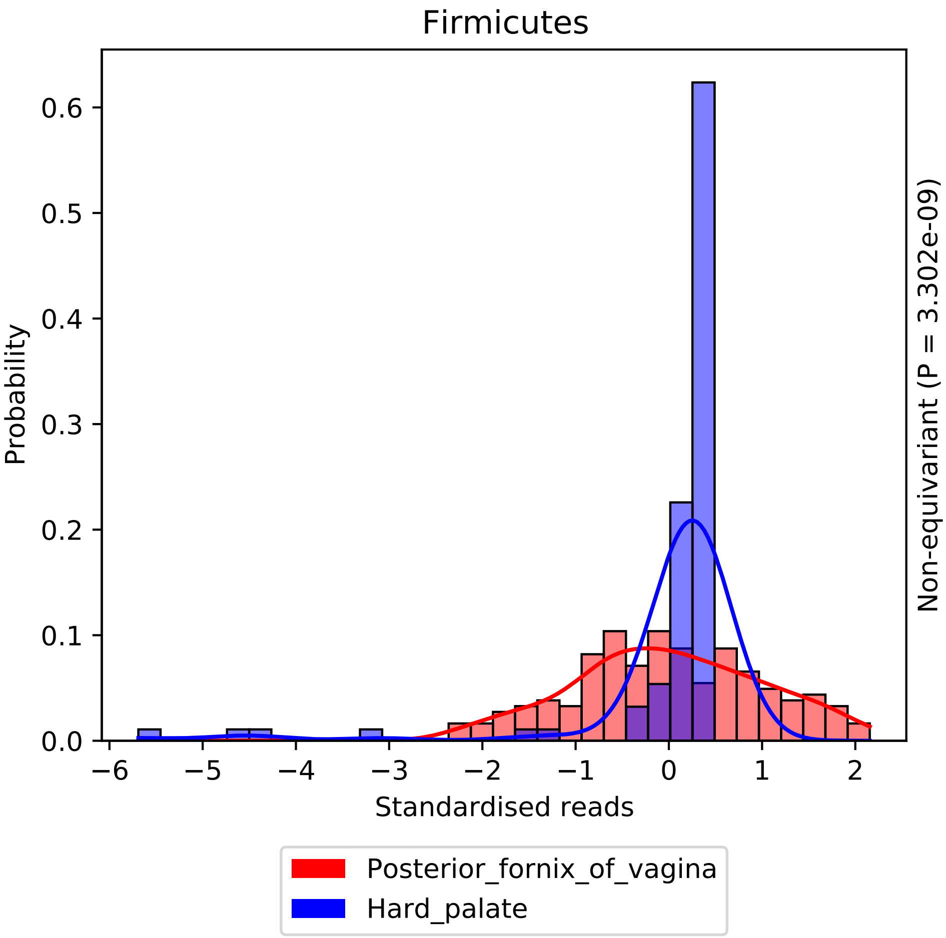

Firmicutes
Probability
0.0
0.1
0.2
0.3
0.4
0.5
0.6
−6
−5
−4
−3
−2
−1
0
1
2
Standardised reads
Non-equivariant (P = 3.302e-09)
Posterior_fornix_of_vagina
Hard_palate

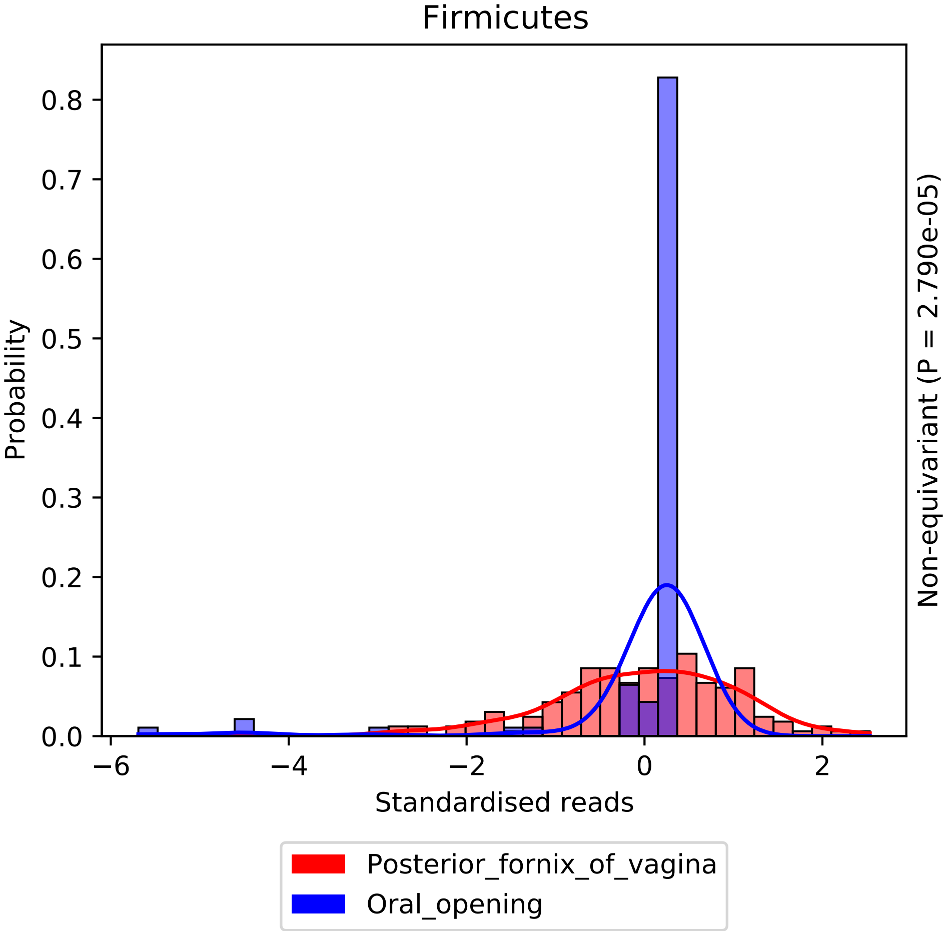
Firmicutes
Probability
0.0
0.1
0.2
0.3
0.4
0.5
0.6
0.7
0.8
−6
−4
−2
0
2
Standardised reads
Non-equivariant (P = 2.790e-05)
Posterior_fornix_of_vagina
Oral_opening

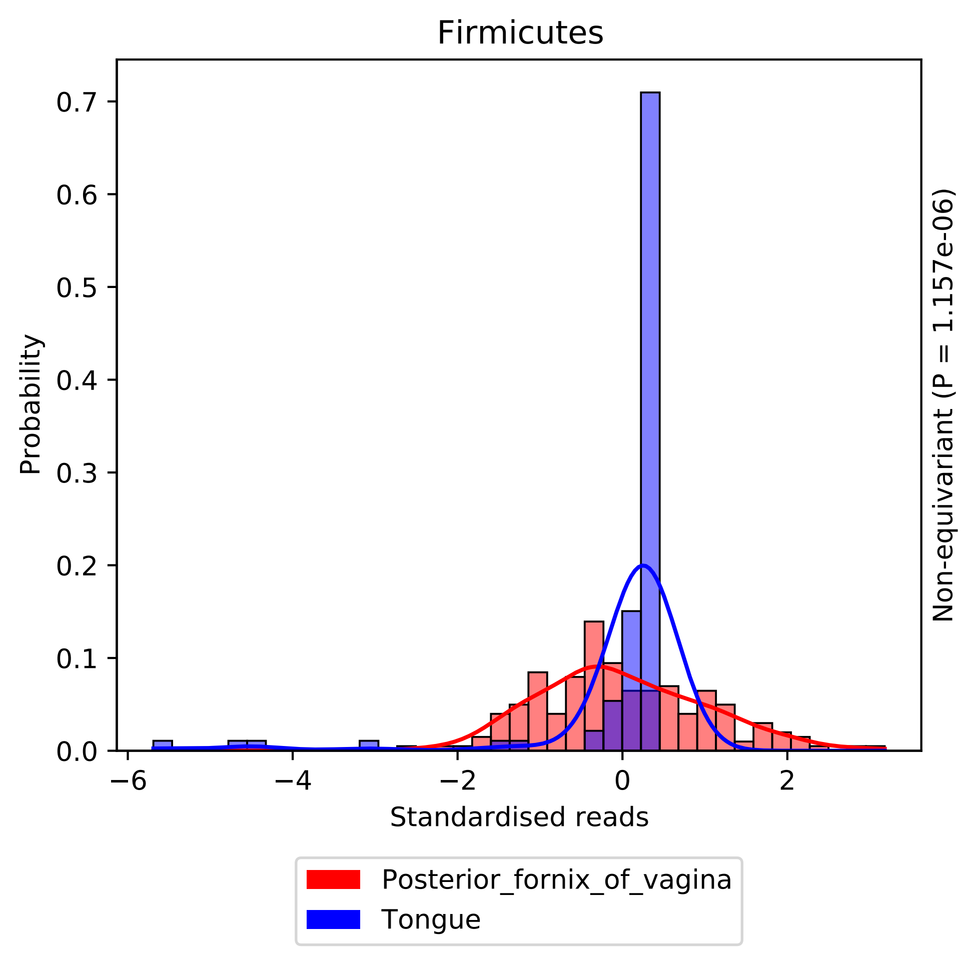

Firmicutes
0.7
0.6
0.5
0.4
0.3
0.2
0.1
0.0
Probability
−6
−4
−2
0
2
Standardised reads
Non-equivariant (P = 1.157e-06)
Posterior_fornix_of_vagina
Tongue

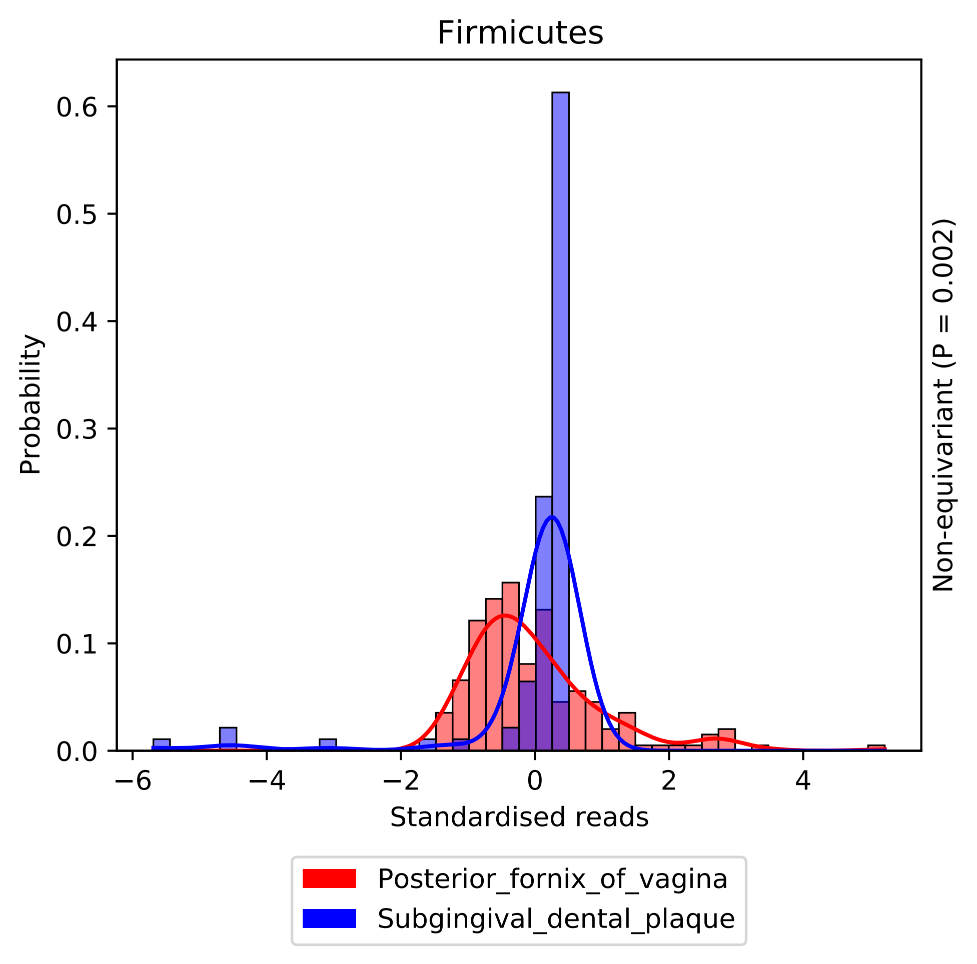

Firmicutes
0.6
0.5
0.4
0.3
0.2
0.1
0.0
Probability
−6
−4
−2
0
2
4
Standardised reads
Non-equivariant (P = 0.002)
Posterior_fornix_of_vagina
Subgingival_dental_plaque

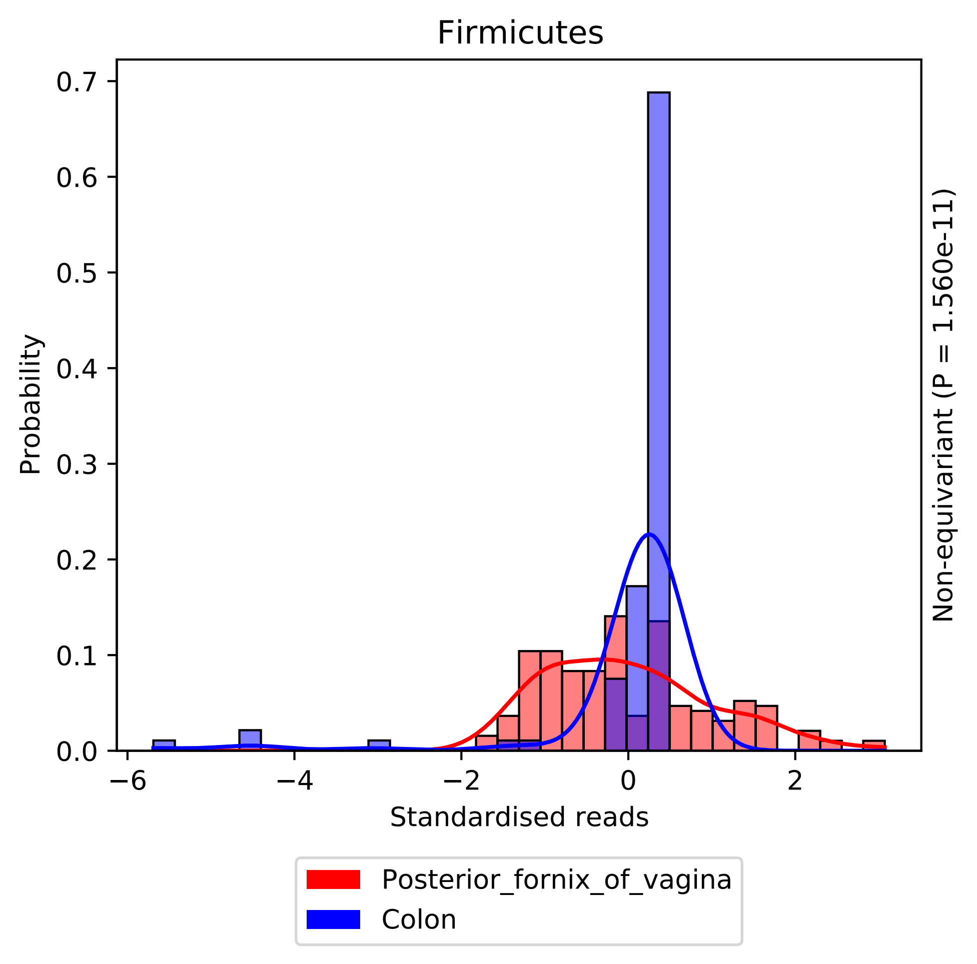

Firmicutes
Probability
Standardised reads
Non-equivariant (P = 1.560e-11)
0.0
0.1
0.2
0.3
0.4
0.5
0.6
0.7
−6
−4
−2
0
2
Posterior_fornix_of_vagina
Colon

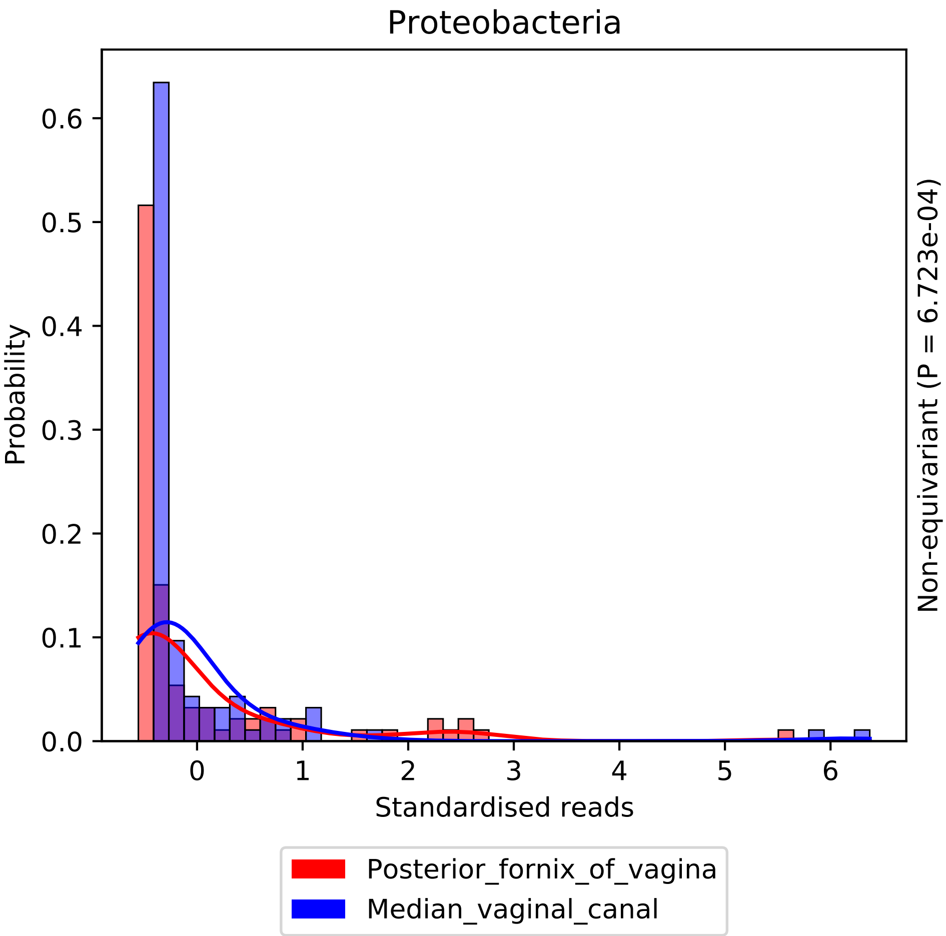
Proteobacteria
Probability
0.0
0.1
0.2
0.3
0.4
0.5
0.6
0
1
2
3
4
5
6
Standardised reads
Non-equivariant (P = 6.723e-04)
Posterior_fornix_of_vagina
Median_vaginal_canal

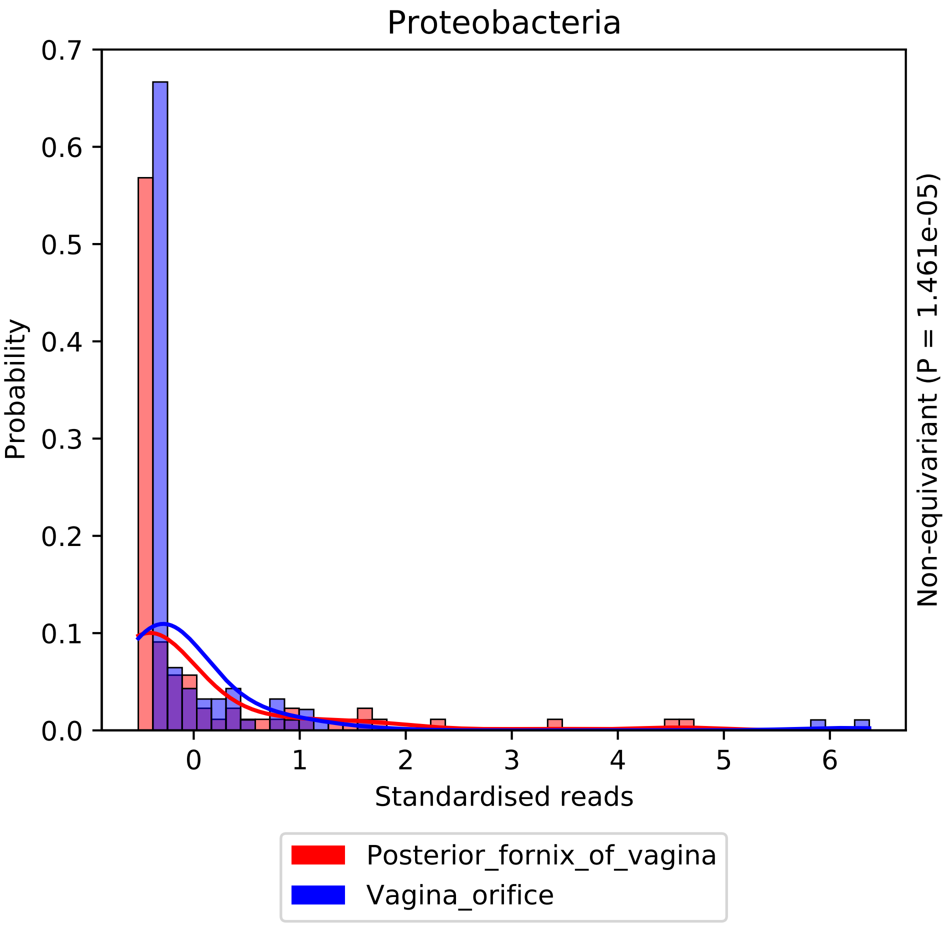
Proteobacteria
Probability
0.0
0.1
0.2
0.3
0.4
0.5
0.6
0.7
0
1
2
3
4
5
6
Standardised reads
Non-equivariant (P = 1.461e-05)
Posterior_fornix_of_vagina
Vagina_orifice

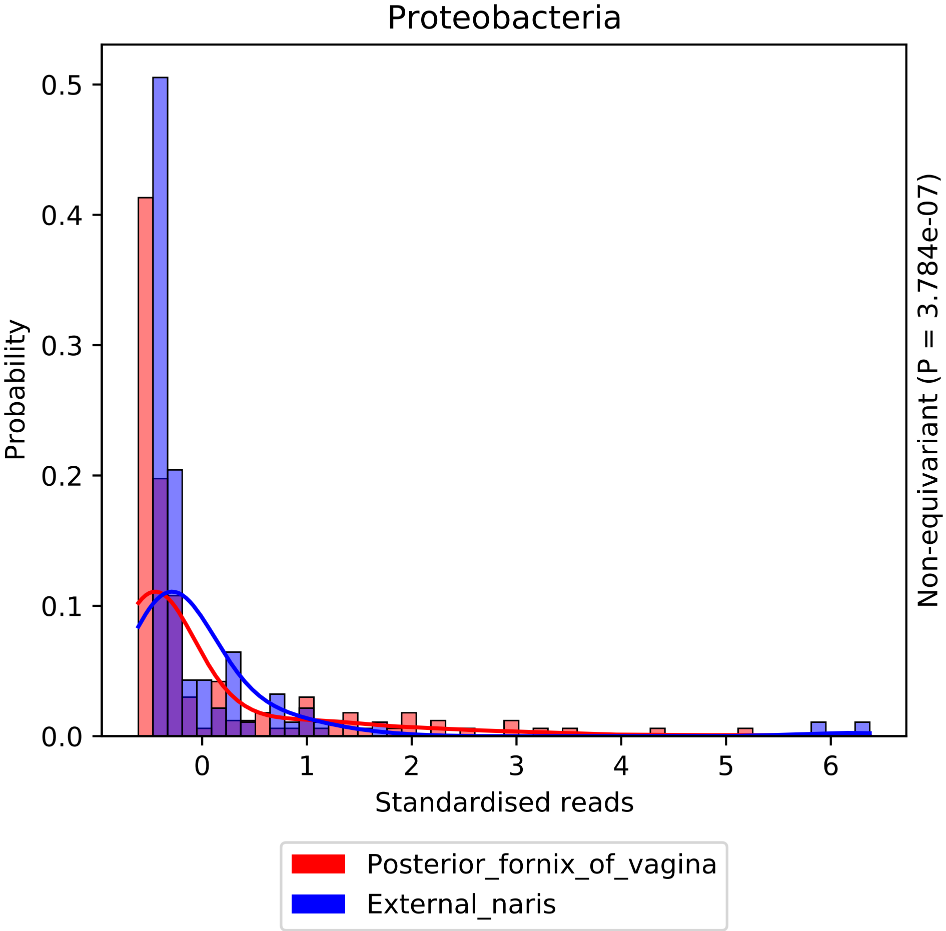

Proteobacteria
Probability
0.0
0.1
0.2
0.3
0.4
0.5
0
1
2
3
4
5
6
Standardised reads
Non-equivariant (P = 3.784e-07)
Posterior_fornix_of_vagina
External_naris

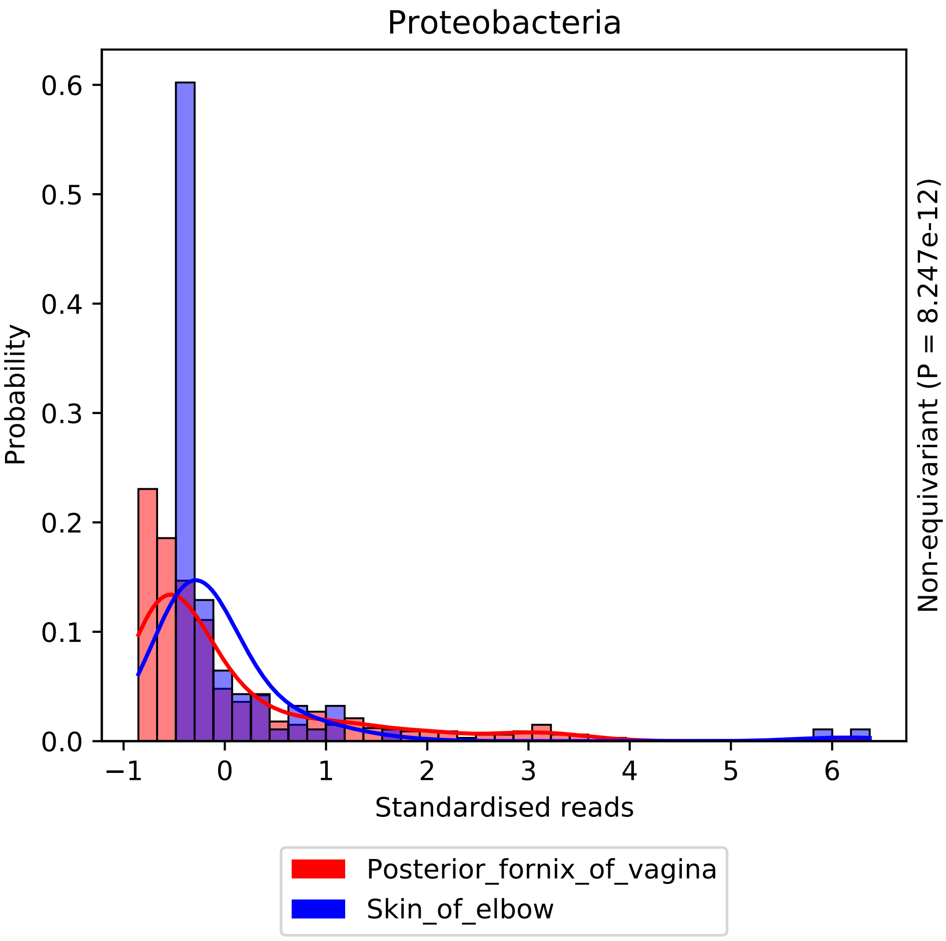

Proteobacteria
Probability
0.0
0.1
0.2
0.3
0.4
0.5
0.6
−1
0
1
2
3
4
5
6
Standardised reads
Non-equivariant (P = 8.247e-12)
Posterior_fornix_of_vagina
Skin_of_elbow

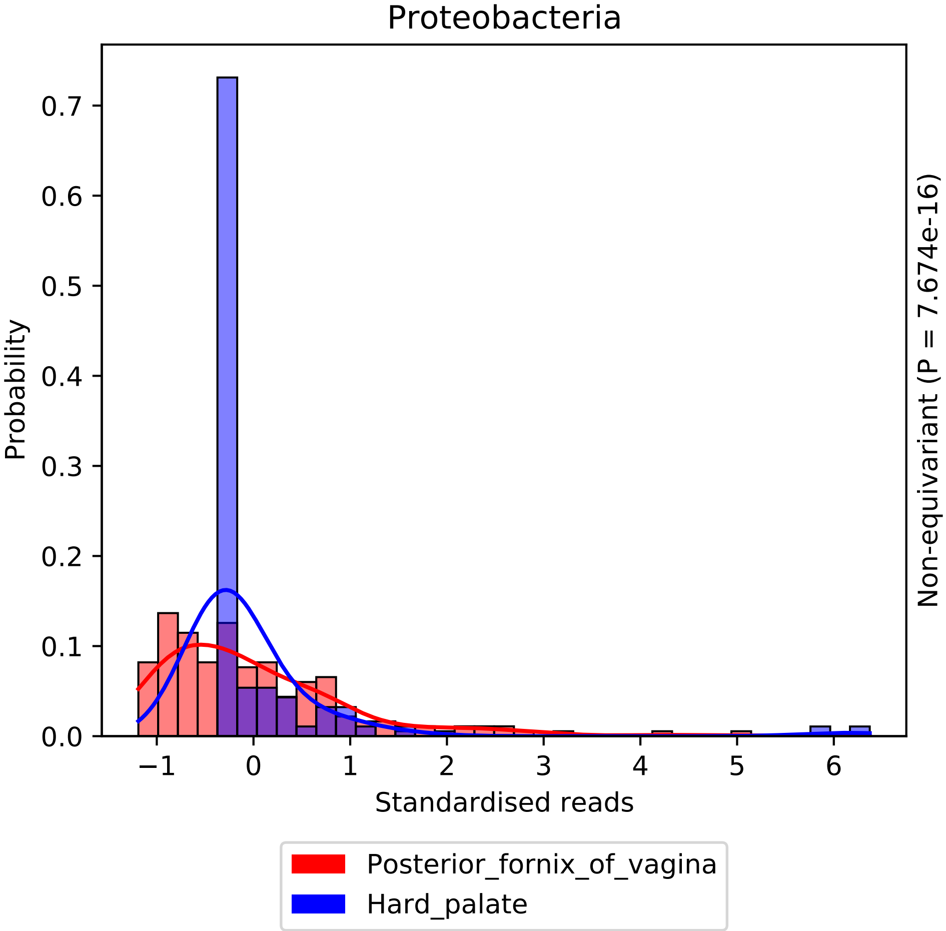
Proteobacteria
Probability
0.0
0.1
0.2
0.3
0.4
0.5
0.6
0.7
−1
0
1
2
3
4
5
6
Standardised reads
Non-equivariant (P = 7.674e-16)
Posterior_fornix_of_vagina
Hard_palate

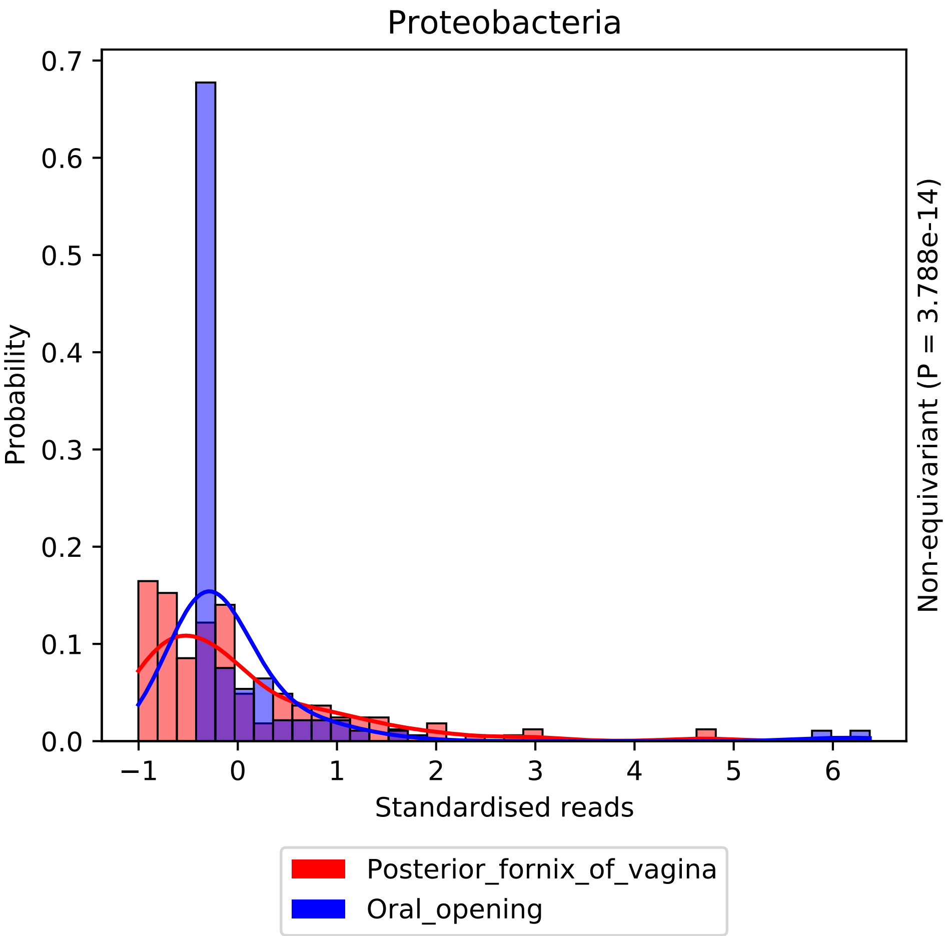

Proteobacteria
Probability
0.0
0.1
0.2
0.3
0.4
0.5
0.6
0.7
−1
0
1
2
3
4
5
6
Standardised reads
Non-equivariant (P = 3.788e-14)
Posterior_fornix_of_vagina
Oral_opening

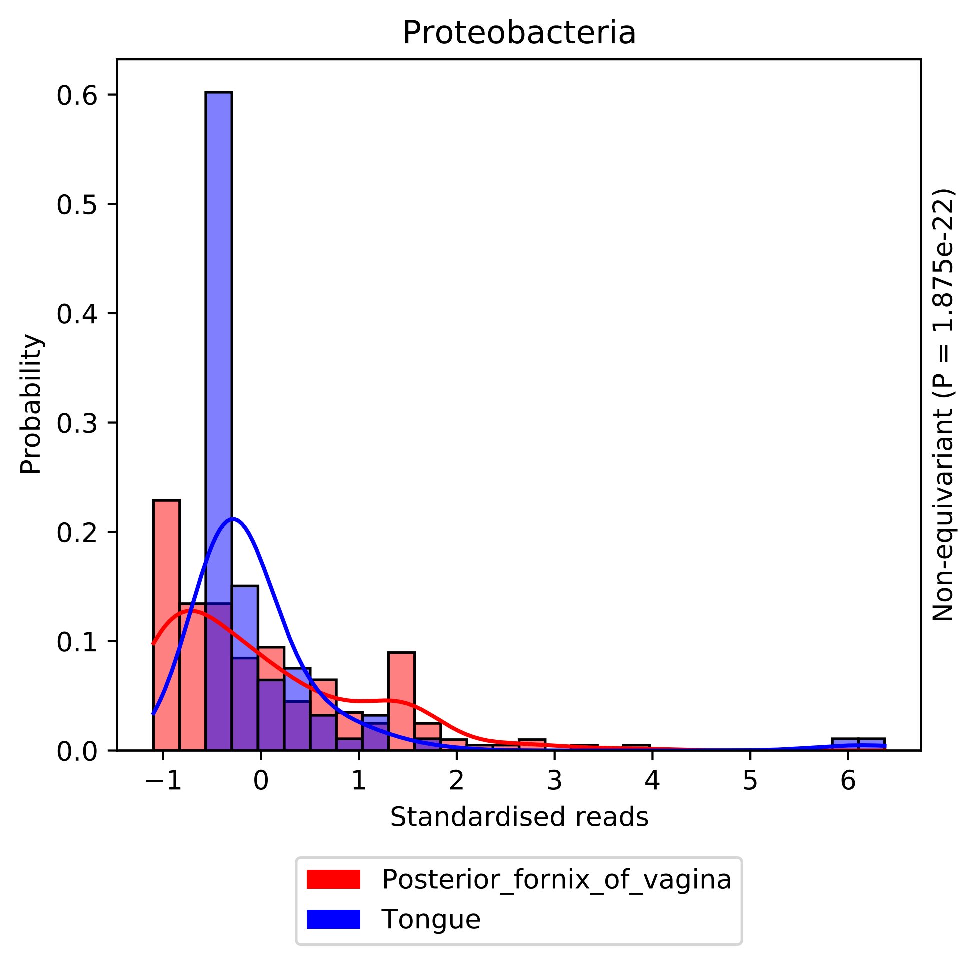

Proteobacteria
Probability
0.0
0.1
0.2
0.3
0.4
0.5
0.6
−1
0
1
2
3
4
5
6
Standardised reads
Non-equivariant (P = 1.875e-22)
Posterior_fornix_of_vagina
Tongue

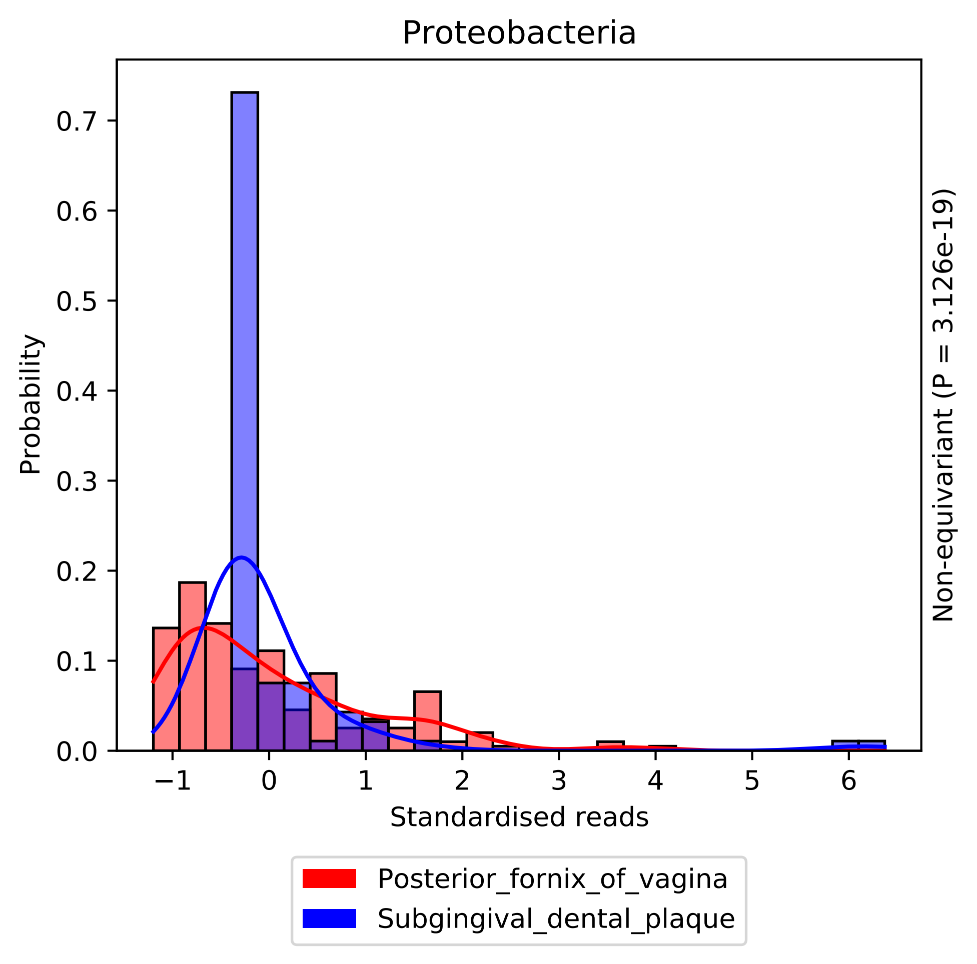

Proteobacteria
Probability
0.0
0.1
0.2
0.3
0.4
0.5
0.6
0.7
−1
0
1
2
3
4
5
6
Standardised reads
Non-equivariant (P = 3.126e-19)
Posterior_fornix_of_vagina
Subgingival_dental_plaque

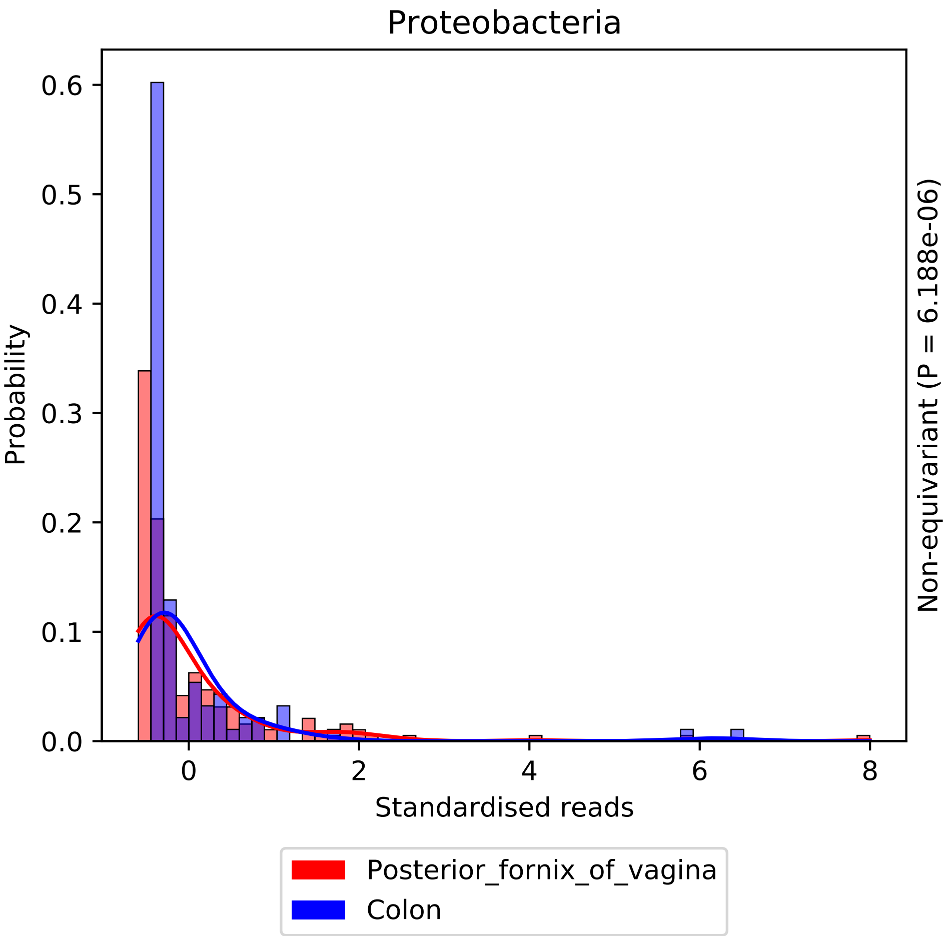

Proteobacteria
Probability
0.0
0.1
0.2
0.3
0.4
0.5
0.6
0
2
4
6
8
Standardised reads
Non-equivariant (P = 6.188e-06)
Posterior_fornix_of_vagina
Colon

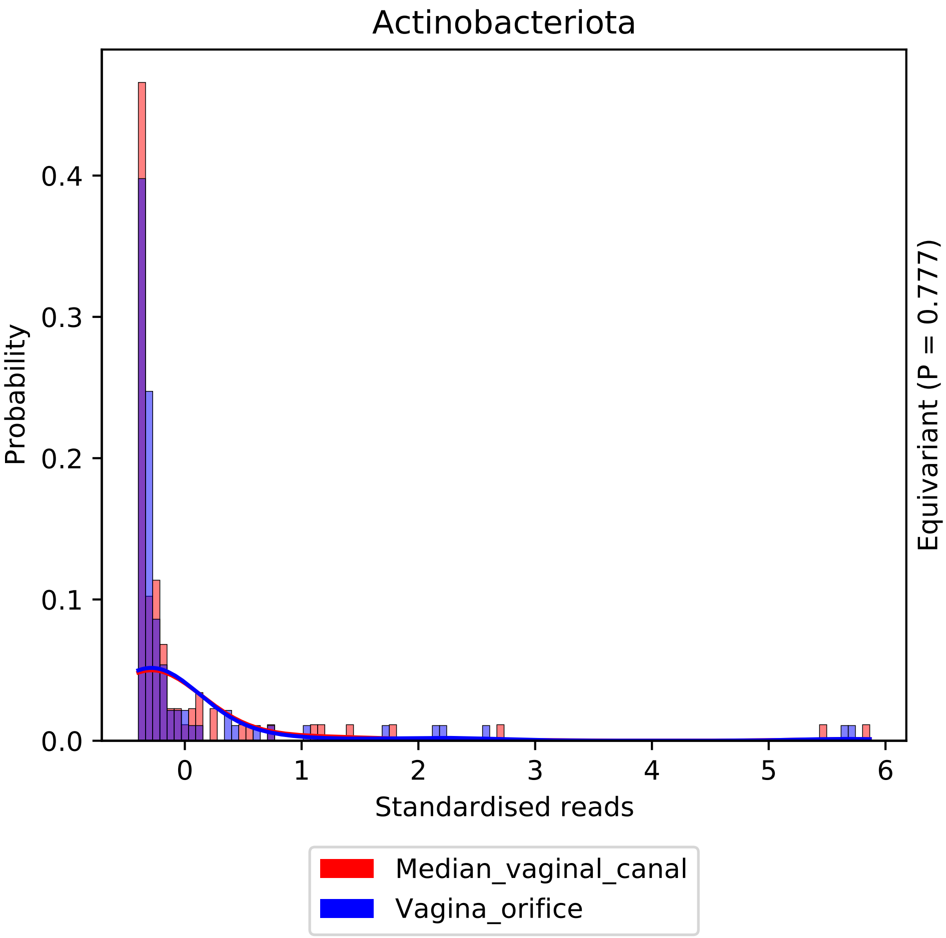
Actinobacteriota
Probability
0.0
0.1
0.2
0.3
0.4
0
1
2
3
4
5
6
Standardised reads
Equivariant (P = 0.777)
Median_vaginal_canal
Vagina_orifice

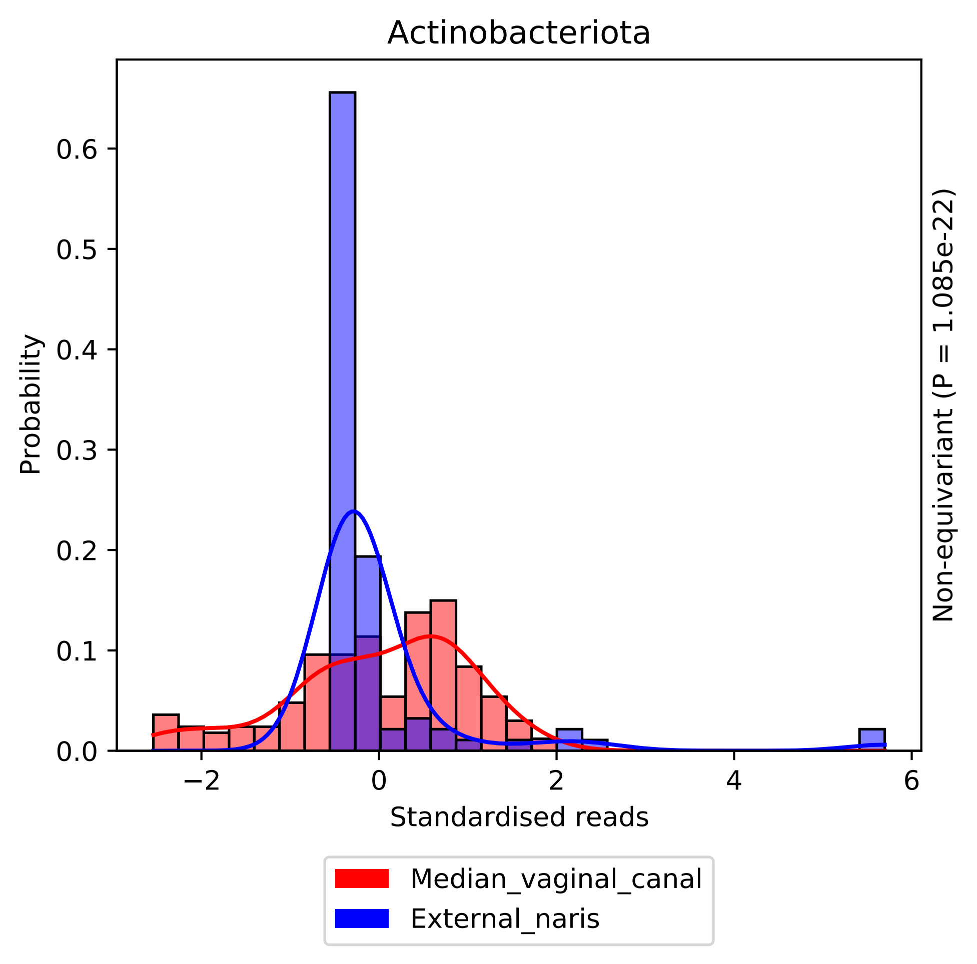
Actinobacteriota
Probability
0.0
0.1
0.2
0.3
0.4
0.5
0.6
−2
0
2
4
6
Standardised reads
Non-equivariant (P = 1.085e-22)
Median_vaginal_canal
External_naris

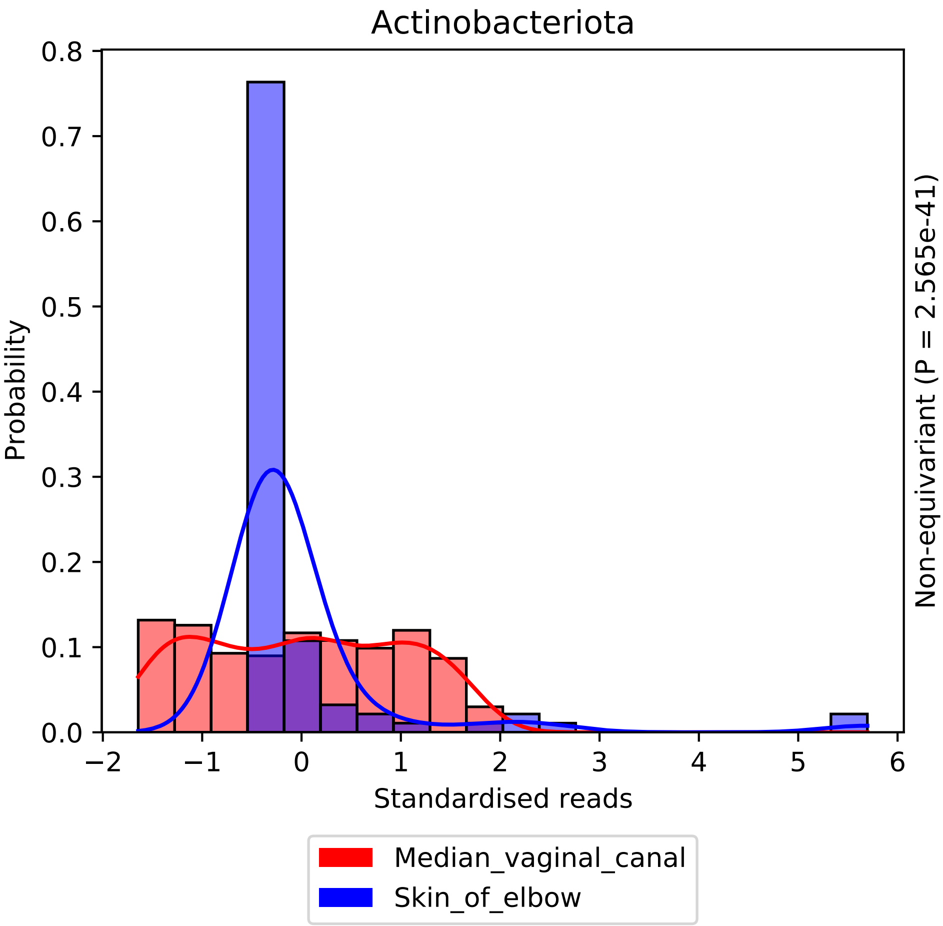

Actinobacteriota
0.8
0.7
0.6
0.5
0.4
0.3
0.2
0.1
0.0
Probability
−2
−1
0
1
2
3
4
5
6
Standardised reads
Non-equivariant (P = 2.565e-41)
Median_vaginal_canal
Skin_of_elbow

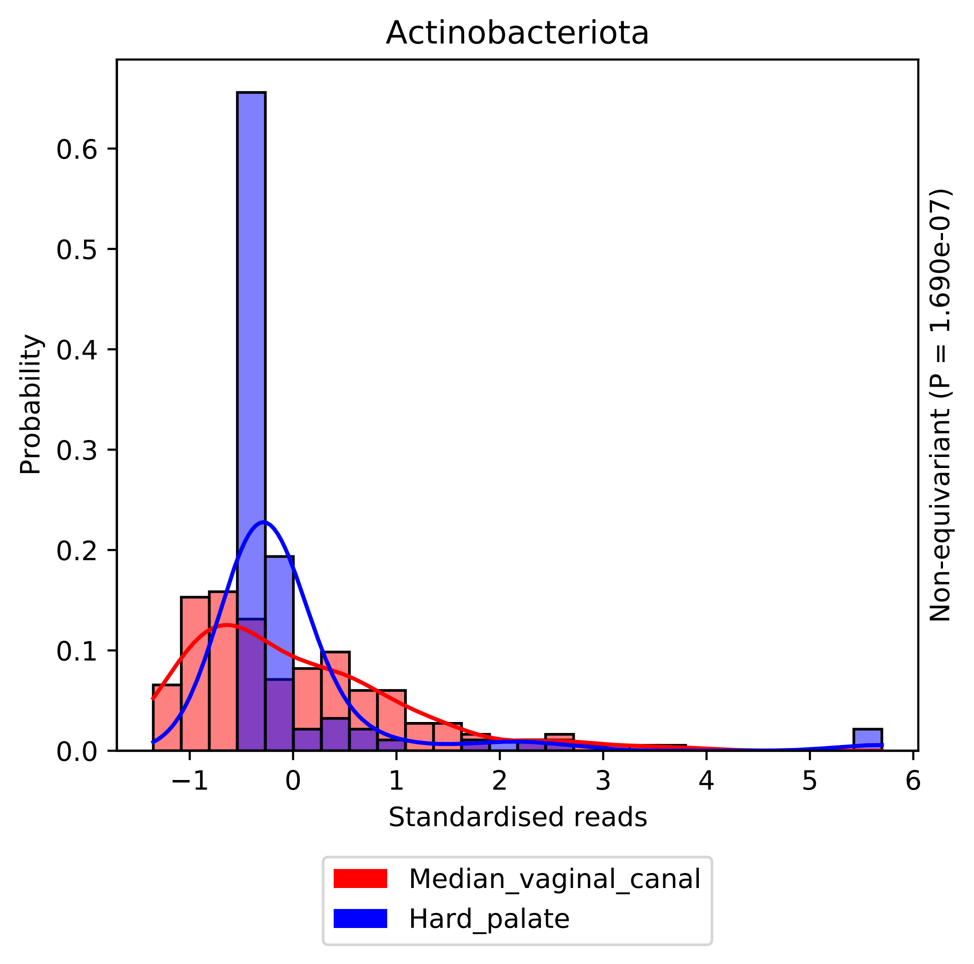

Actinobacteriota
Probability
0.0
0.1
0.2
0.3
0.4
0.5
0.6
−1
0
1
2
3
4
5
6
Standardised reads
Non-equivariant (P = 1.690e-07)
Median_vaginal_canal
Hard_palate

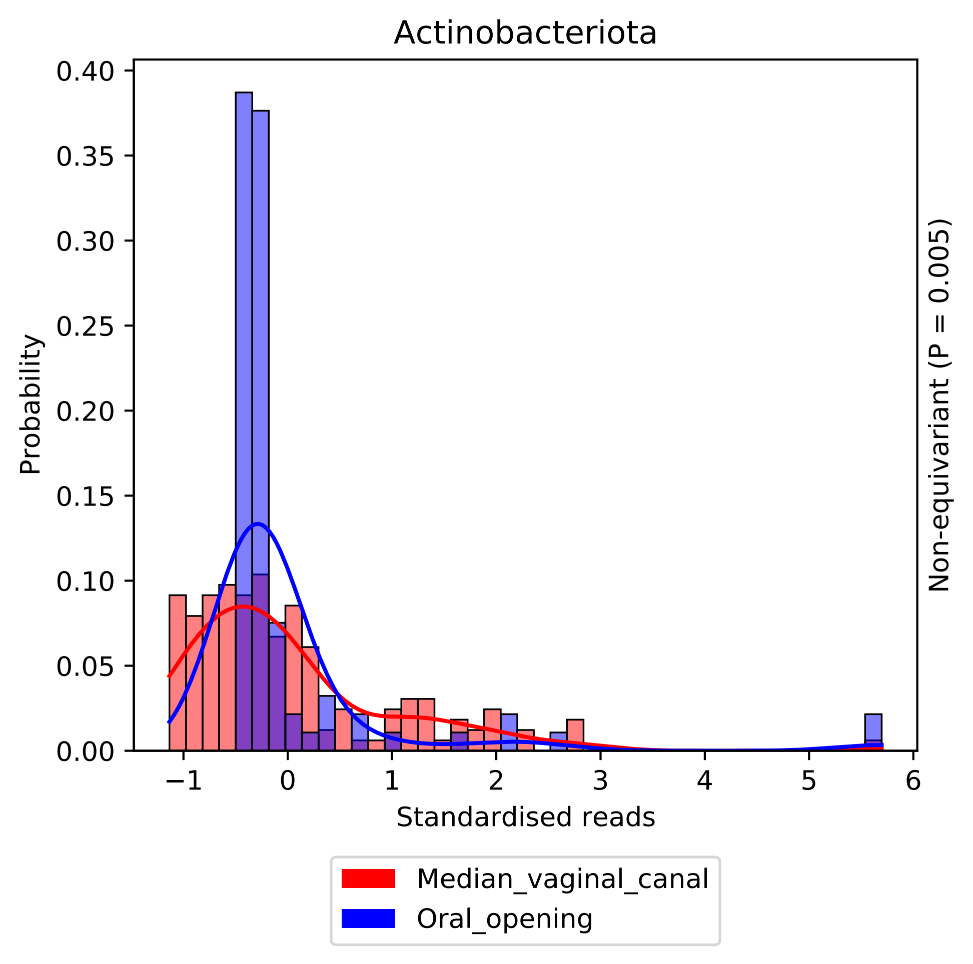

Actinobacteriota
Probability
0.40
0.35
0.30
0.25
0.20
0.15
0.10
0.05
0.00
−1
0
1
2
3
4
5
6
Standardised reads
Non-equivariant (P = 0.005)
Median_vaginal_canal
Oral_opening

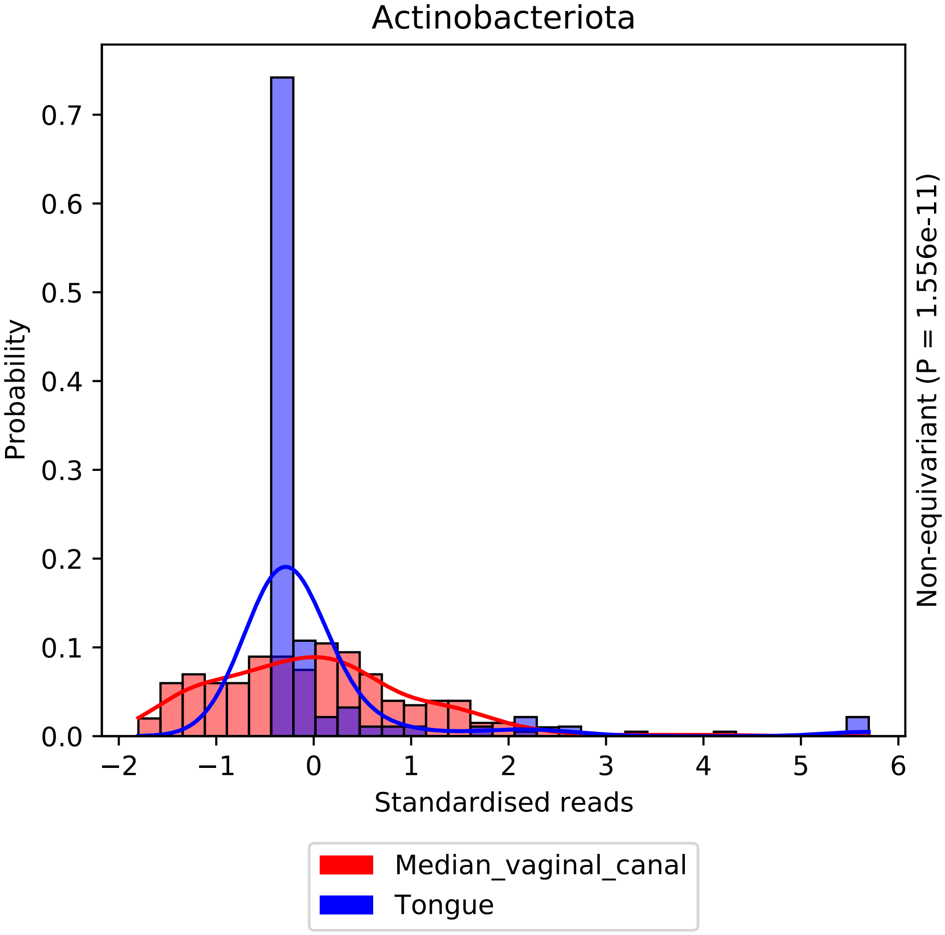

Actinobacteriota
Probability
0.0
0.1
0.2
0.3
0.4
0.5
0.6
0.7
−2
−1
0
1
2
3
4
5
6
Standardised reads
Non-equivariant (P = 1.556e-11)
Median_vaginal_canal
Tongue

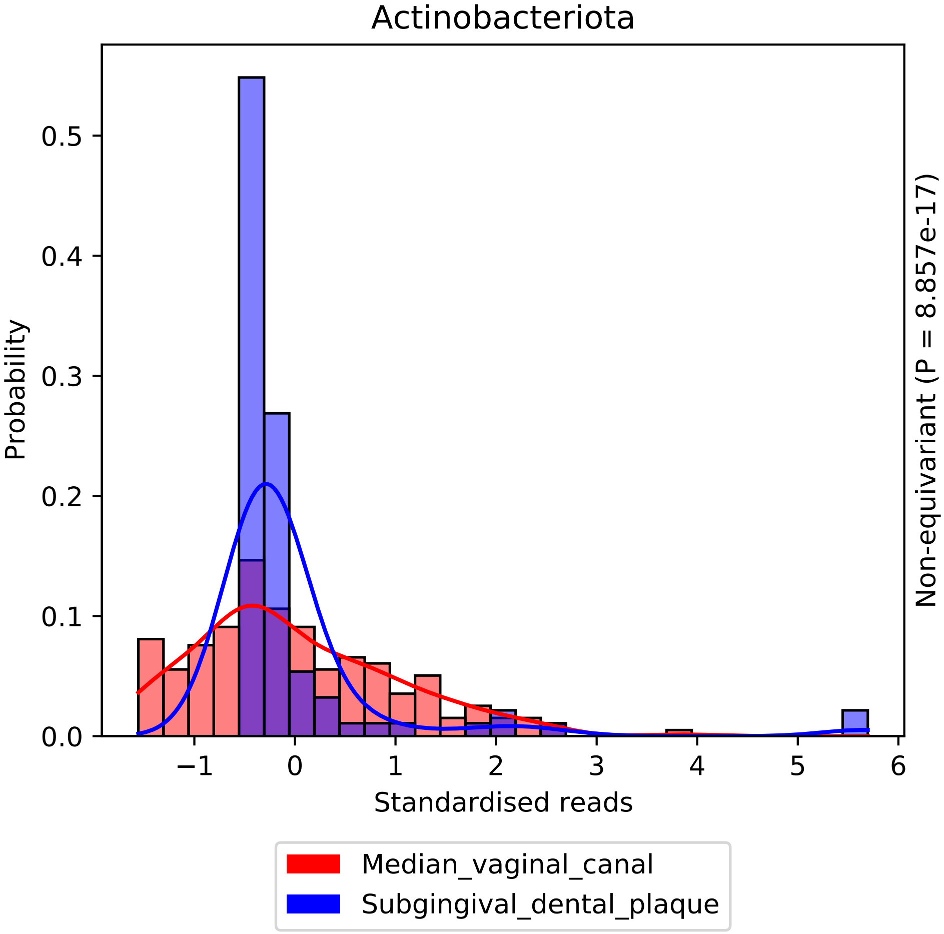

Actinobacteriota
Probability
0.0
0.1
0.2
0.3
0.4
0.5
−1
0
1
2
3
4
5
6
Standardised reads
Non-equivariant (P = 8.857e-17)
Median_vaginal_canal
Subgingival_dental_plaque

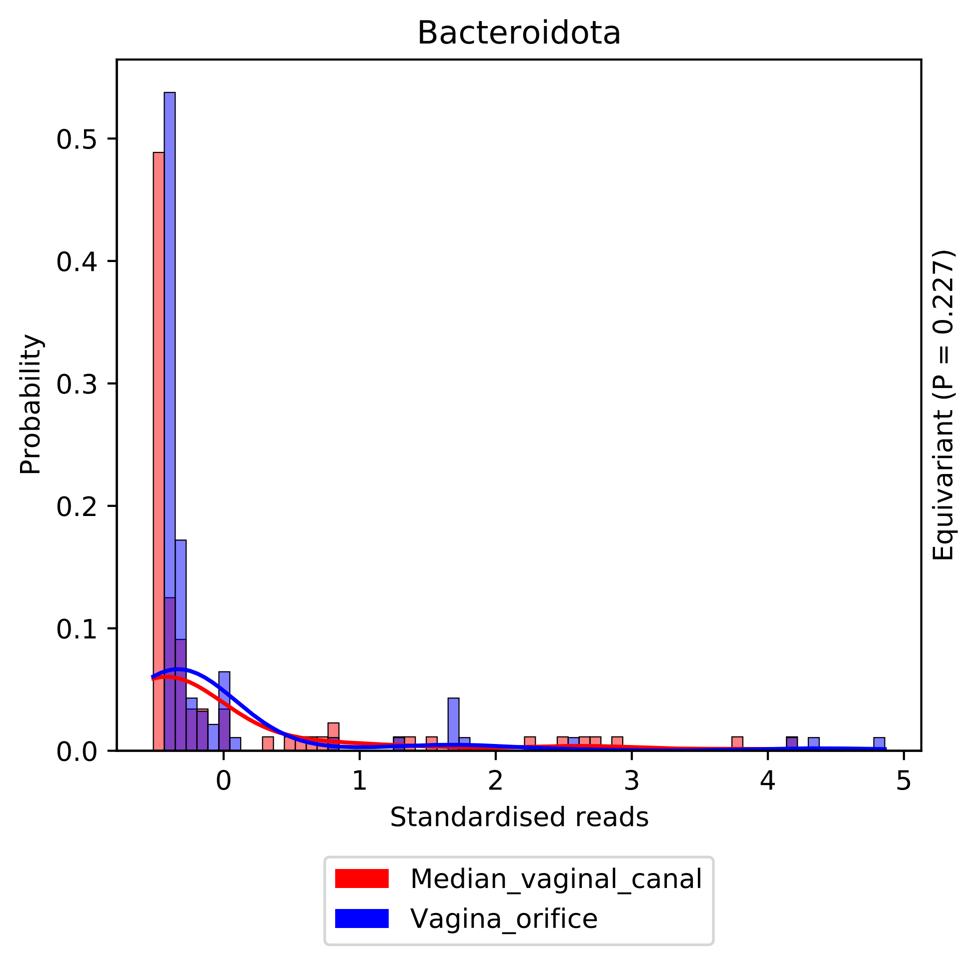
Bacteroidota
Probability
0.0
0.1
0.2
0.3
0.4
0.5
0
1
2
3
4
5
Standardised reads
Equivariant (P = 0.227)
Median_vaginal_canal
Vagina_orifice

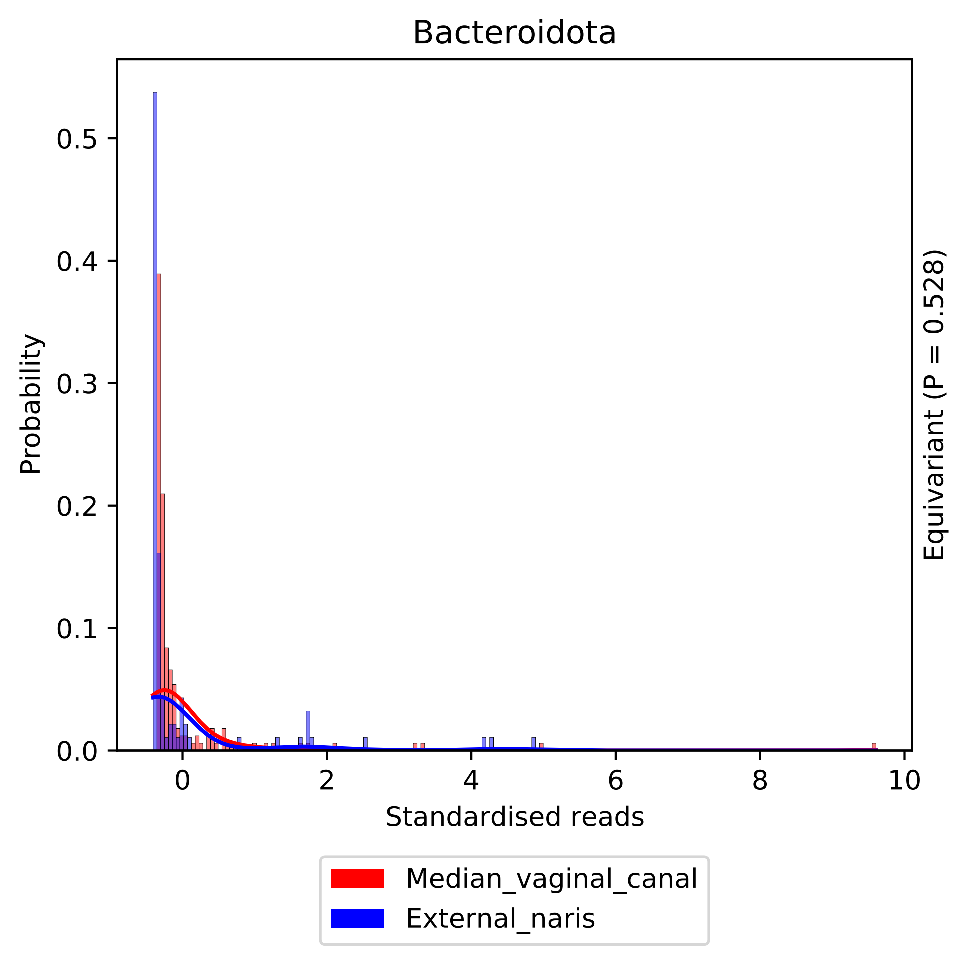

Bacteroidota
Probability
0.0
0.1
0.2
0.3
0.4
0.5
0
2
4
6
8
10
Standardised reads
Equivariant (P = 0.528)
Median_vaginal_canal
External_naris

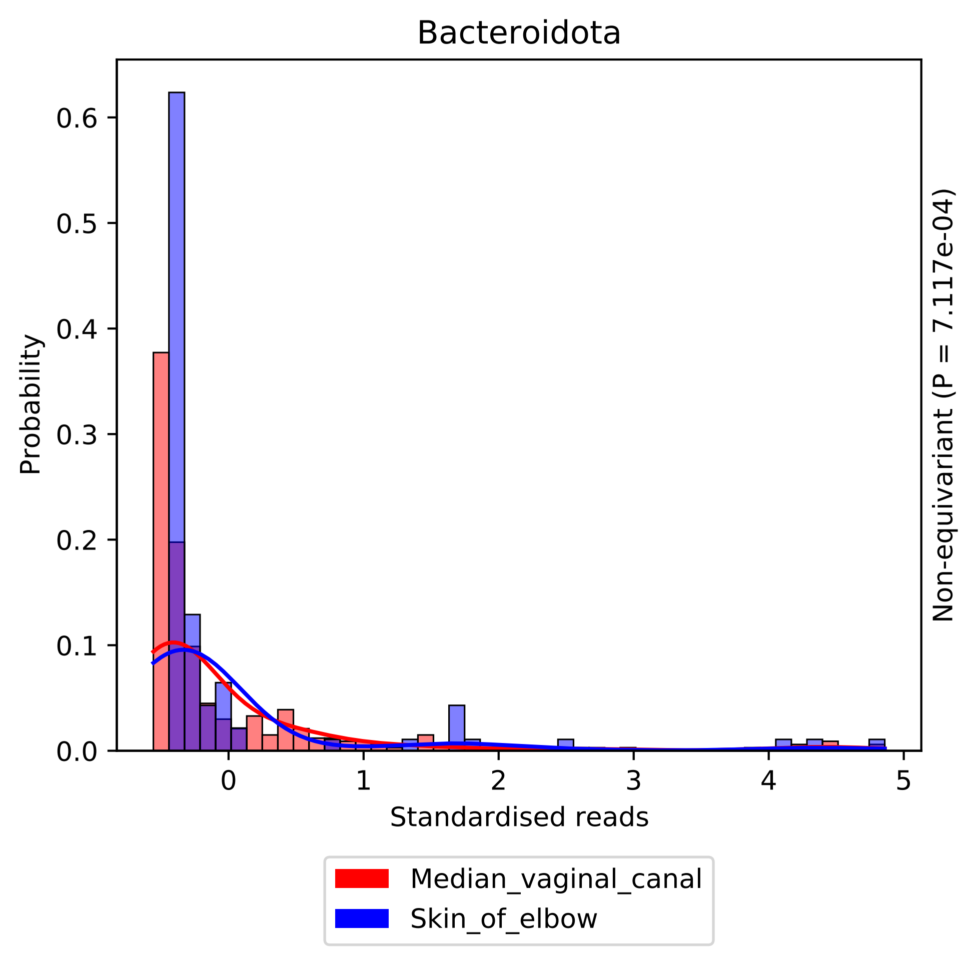
Bacteroidota
Probability
Standardised reads
Non-equivariant (P = 7.117e-04)
Median_vaginal_canal
Skin_of_elbow
0.0
0.1
0.2
0.3
0.4
0.5
0.6
0
1
2
3
4
5

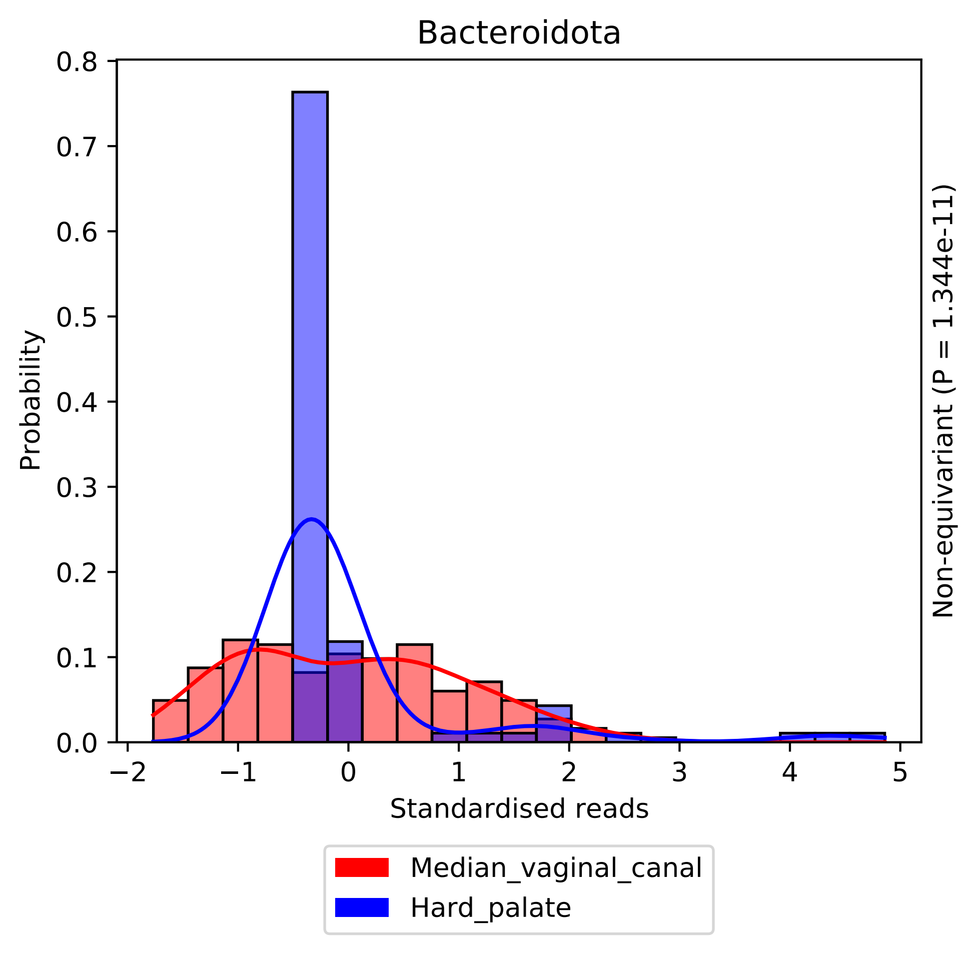

Bacteroidota
Probability
0.0
0.1
0.2
0.3
0.4
0.5
0.6
0.7
0.8
−2
−1
0
1
2
3
4
5
Standardised reads
Non-equivariant (P = 1.344e-11)
Median_vaginal_canal
Hard_palate

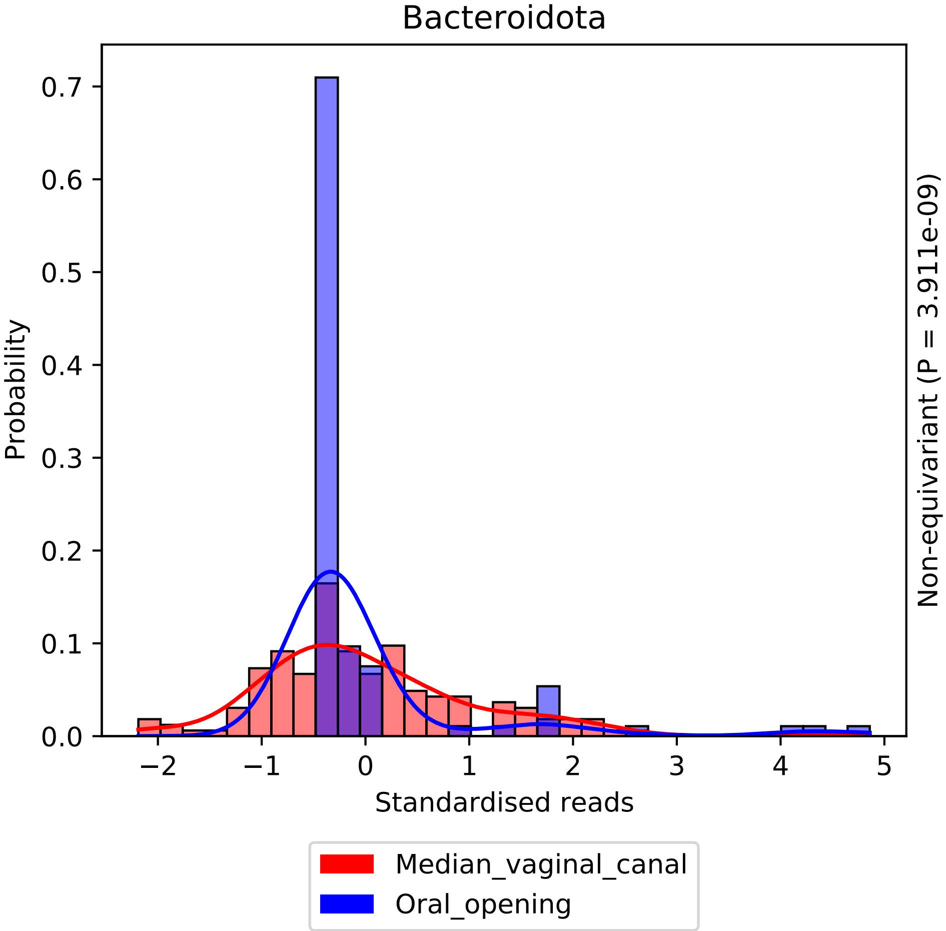
Bacteroidota
Probability
Standardised reads
Non-equivariant (P = 3.911e-09)
0.0
0.1
0.2
0.3
0.4
0.5
0.6
0.7
−2
−1
0
1
2
3
4
5
Median_vaginal_canal
Oral_opening

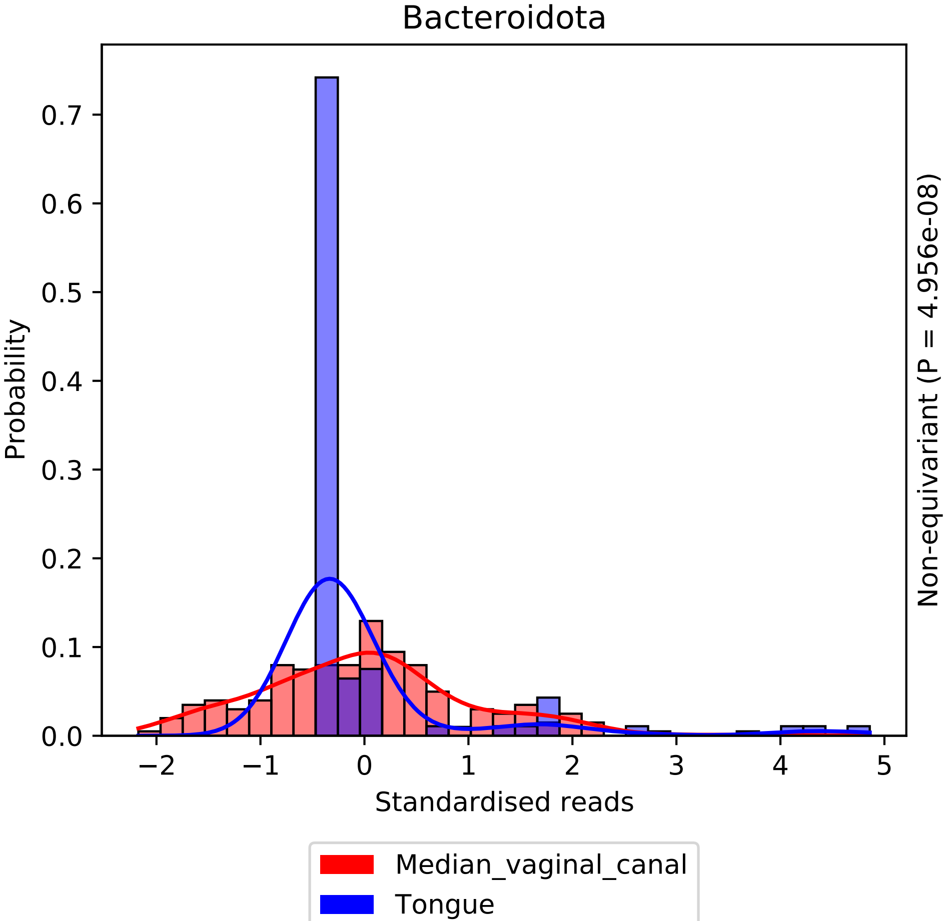

Bacteroidota
Probability
0.0
0.1
0.2
0.3
0.4
0.5
0.6
0.7
−2
−1
0
1
2
3
4
5
Standardised reads
Non-equivariant (P = 4.956e-08)
Median_vaginal_canal
Tongue

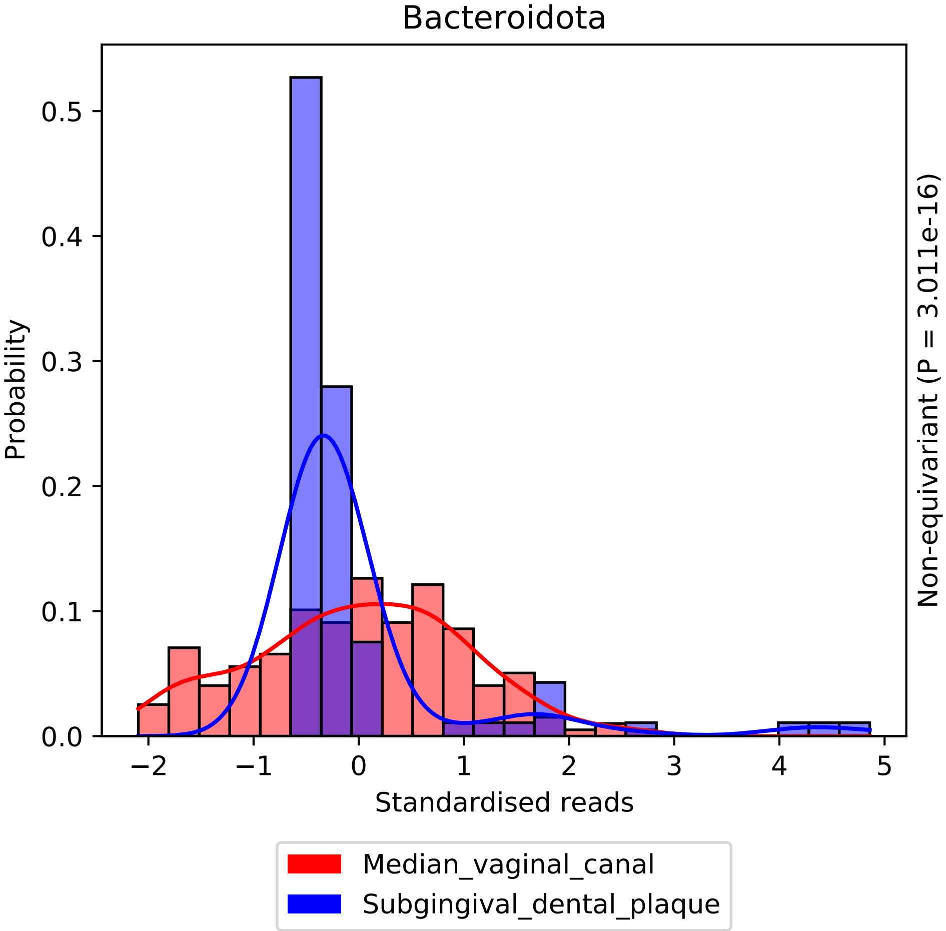

Bacteroidota
Probability
0.0
0.1
0.2
0.3
0.4
0.5
−2
−1
0
1
2
3
4
5
Standardised reads
Non-equivariant (P = 3.011e-16)
Median_vaginal_canal
Subgingival_dental_plaque

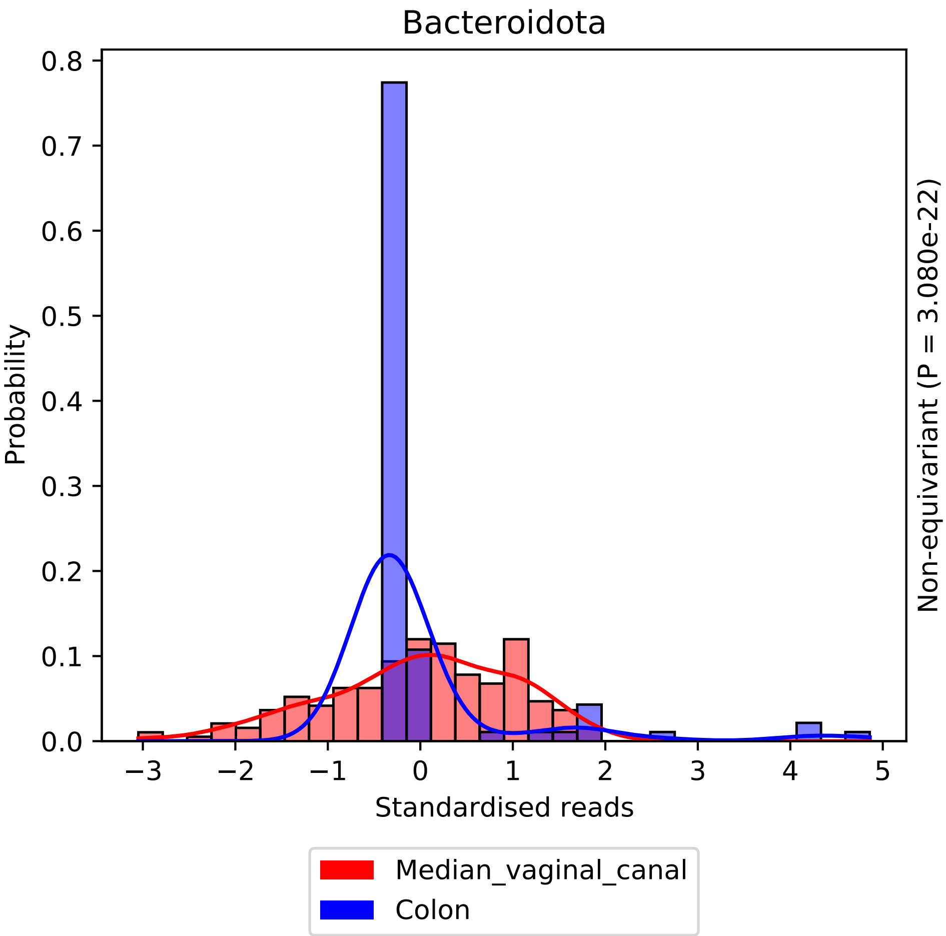

Bacteroidota
Probability
Standardised reads
Non-equivariant (P = 3.080e-22)
Median_vaginal_canal
Colon

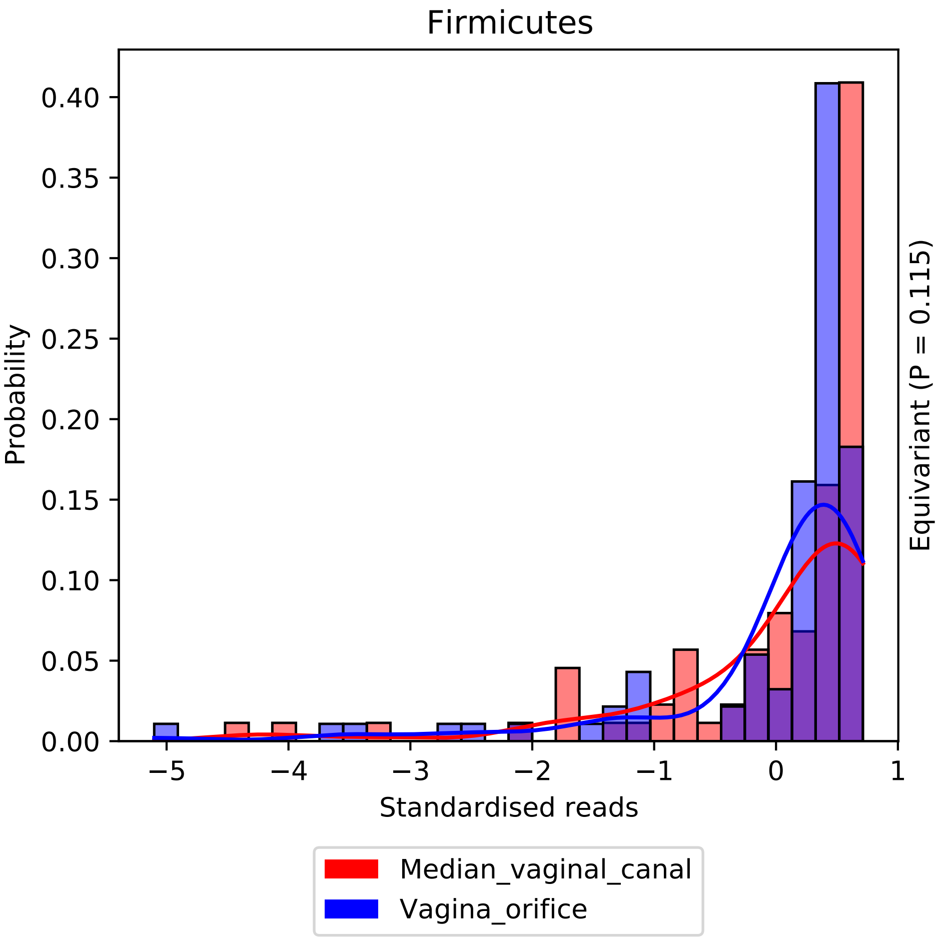

Firmicutes
0.40
0.35
0.30
0.25
0.20
0.15
0.10
0.05
0.00
Probability
−5
−4
−3
−2
−1
0
1
Standardised reads
Equivariant (P = 0.115)
Median_vaginal_canal
Vagina_orifice

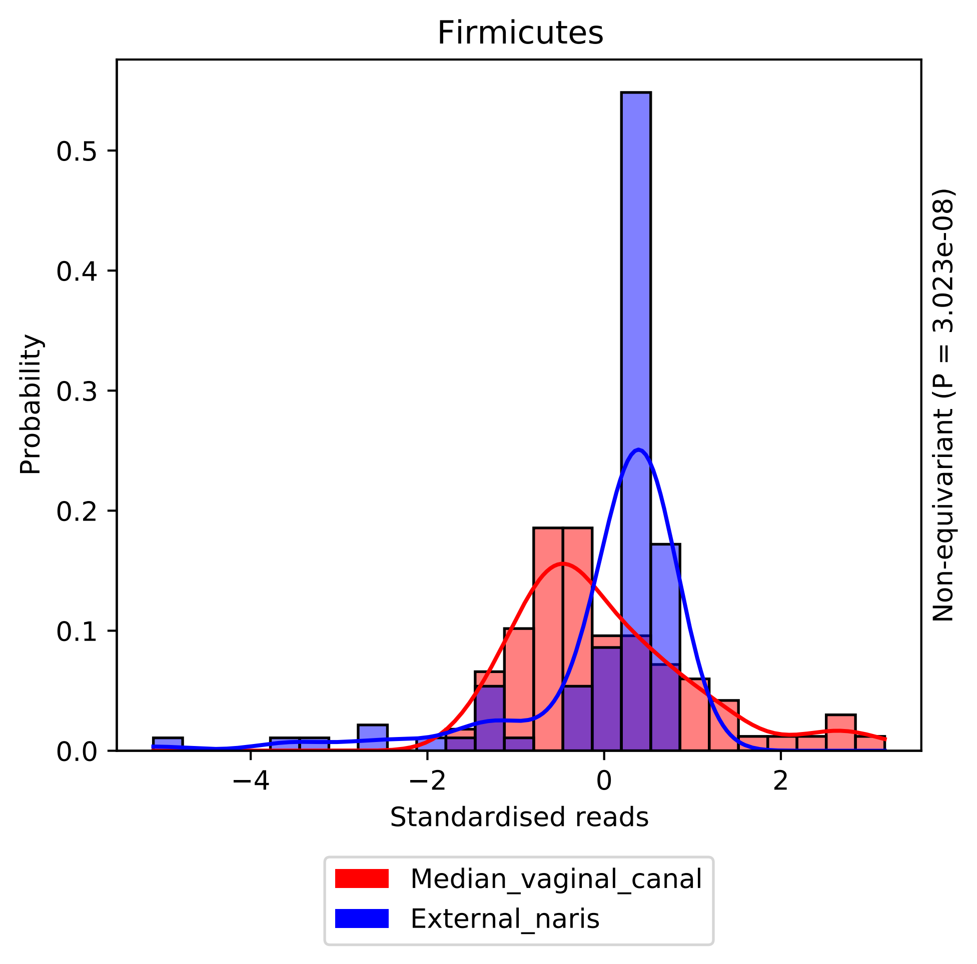

Firmicutes
0.0
0.1
0.2
0.3
0.4
0.5
−4
−2
0
2
Probability
Standardised reads
Non-equivariant (P = 3.023e-08)
Median_vaginal_canal
External_naris

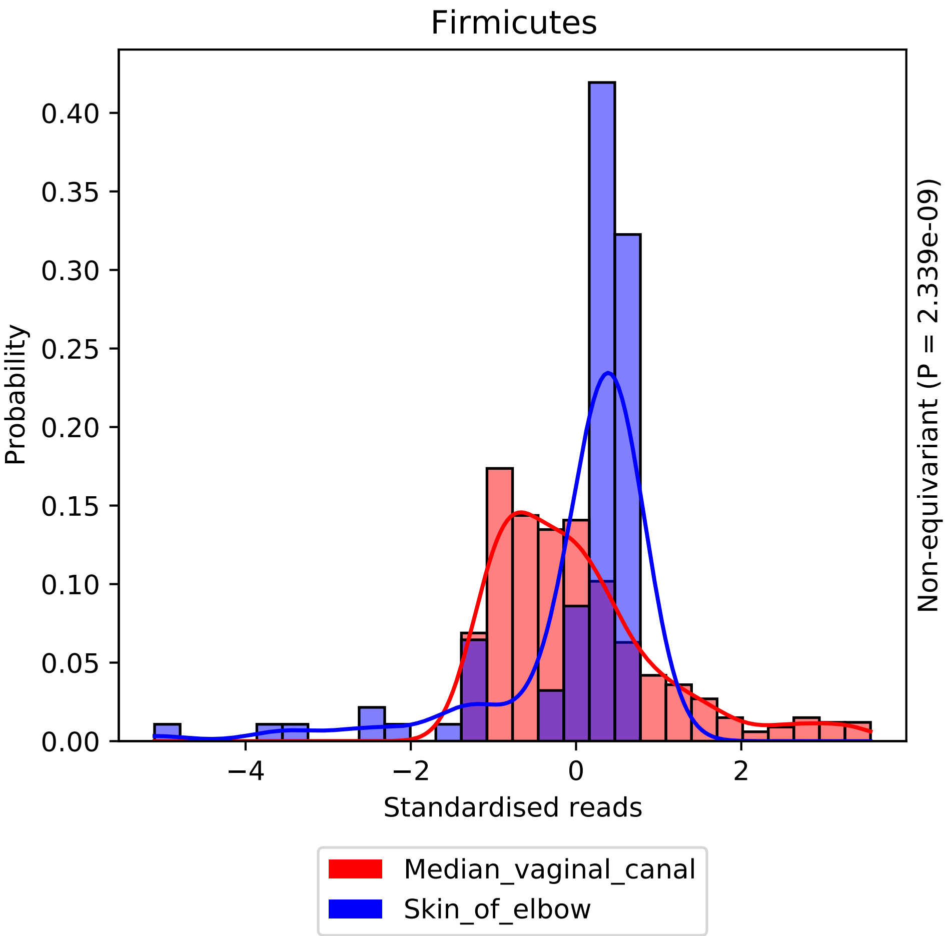

Firmicutes
Probability
0.00
0.05
0.10
0.15
0.20
0.25
0.30
0.35
0.40
−4
−2
0
2
Standardised reads
Non-equivariant (P = 2.339e-09)
Median_vaginal_canal
Skin_of_elbow

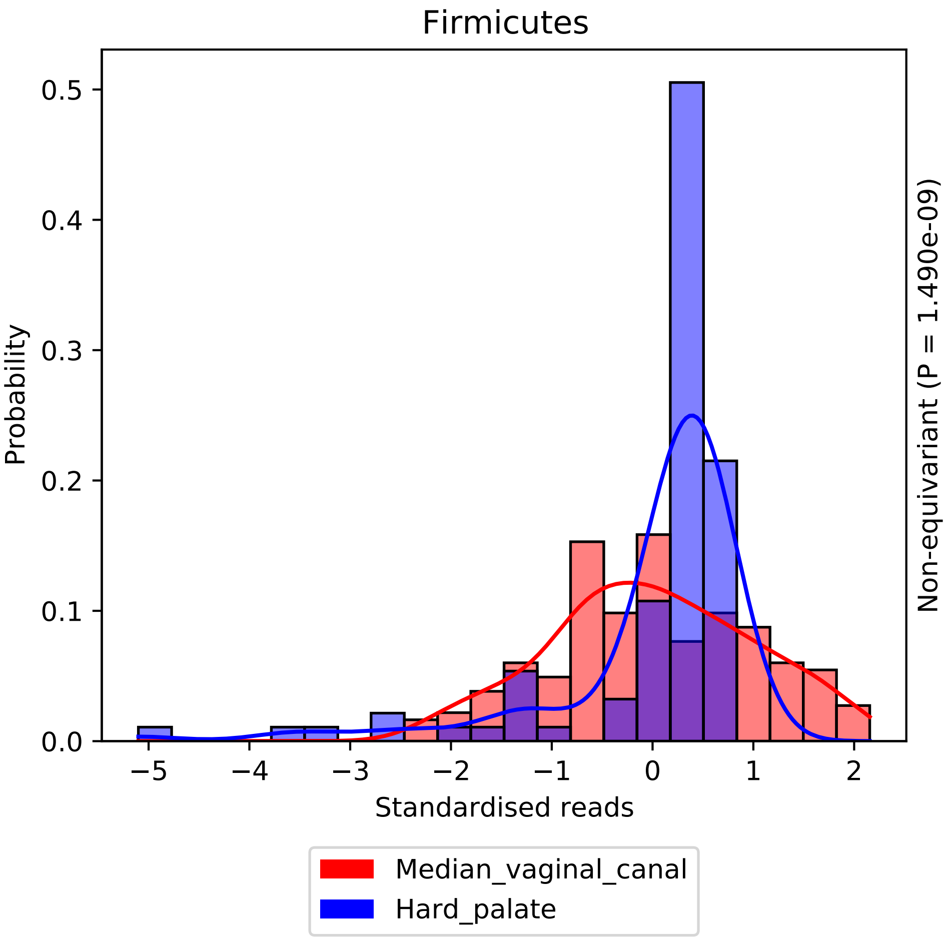
Firmicutes
Probability
Standardised reads
Non-equivariant (P = 1.490e-09)
Median_vaginal_canal
Hard_palate
0.0
0.1
0.2
0.3
0.4
0.5
−5
−4
−3
−2
−1
0
1
2

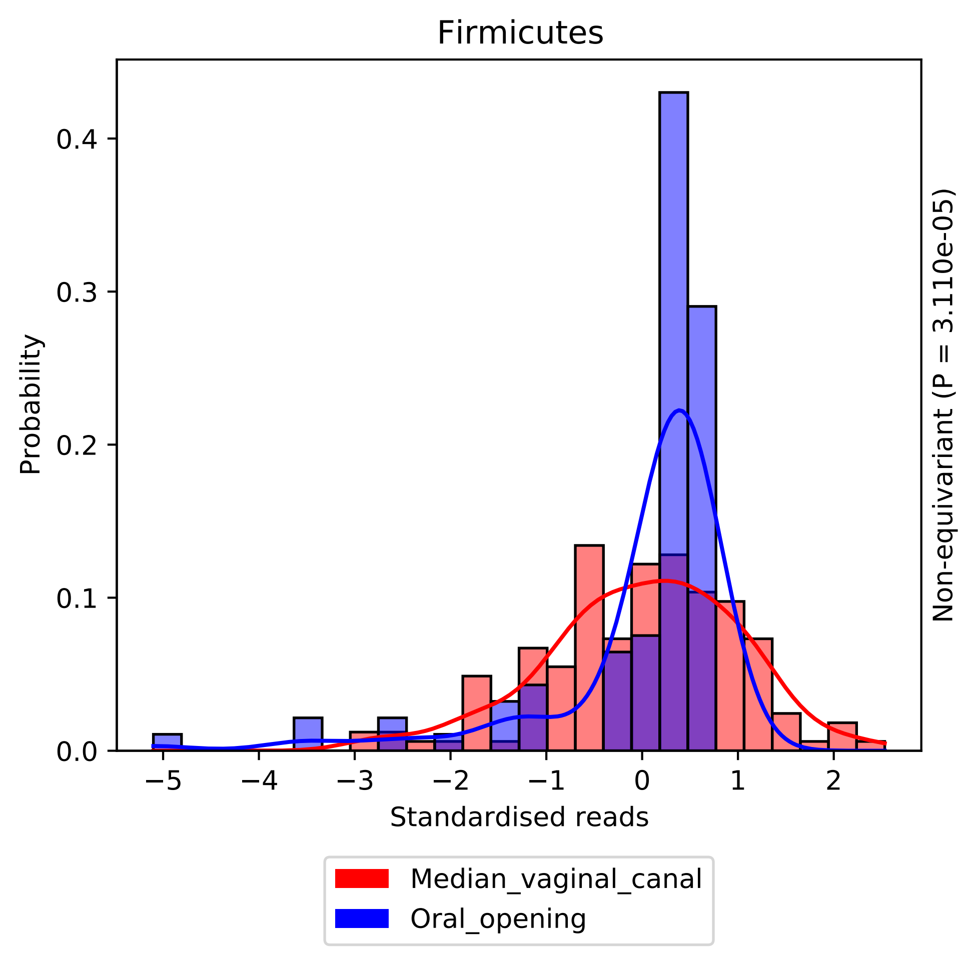
Firmicutes
Probability
Standardised reads
Non-equivariant (P = 3.110e-05)
0.0
0.1
0.2
0.3
0.4
−5
−4
−3
−2
−1
0
1
2
Median_vaginal_canal
Oral_opening

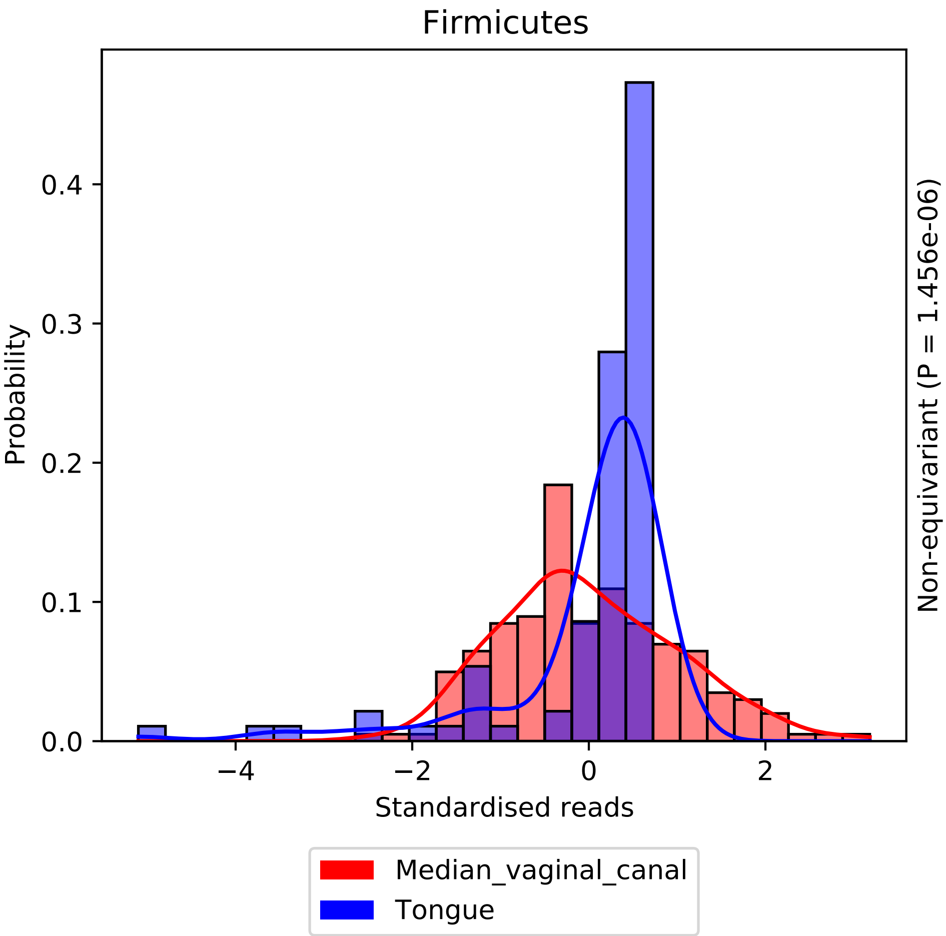

Firmicutes
Probability
0.0
0.1
0.2
0.3
0.4
−4
−2
0
2
Standardised reads
Non-equivariant (P = 1.456e-06)
Median_vaginal_canal
Tongue

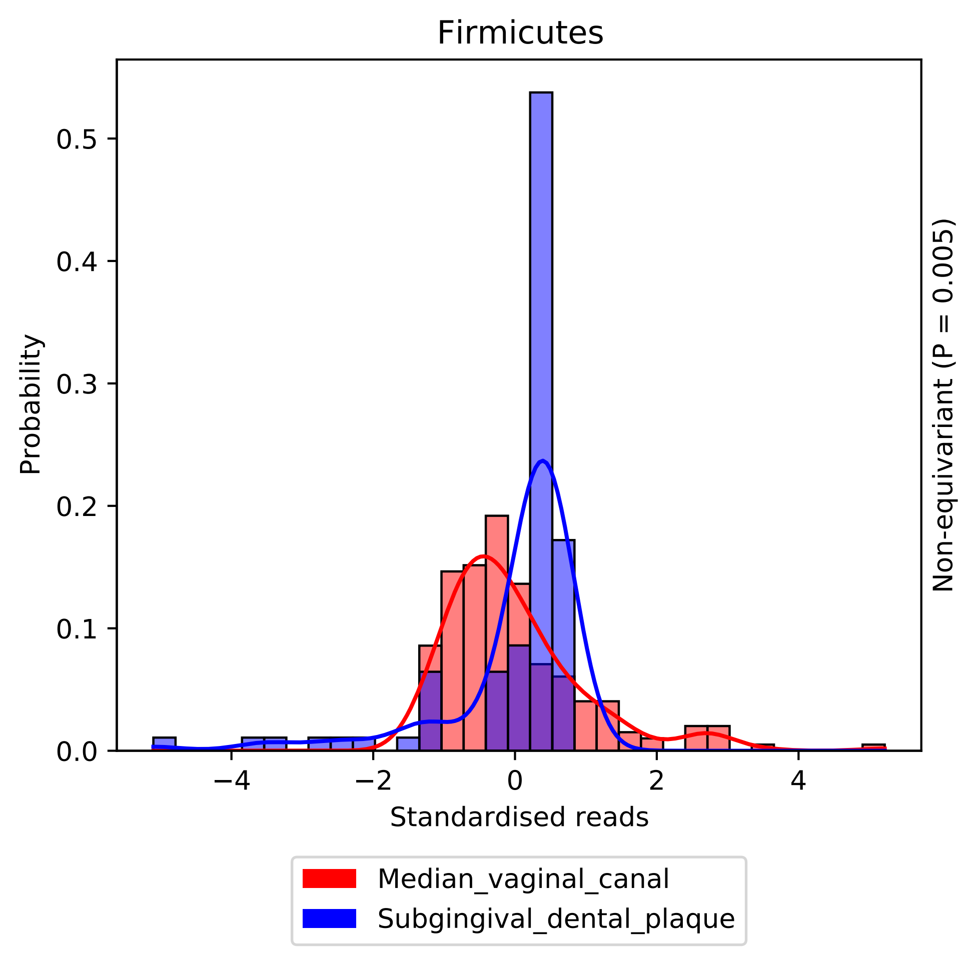

Firmicutes
Probability
0.0
0.1
0.2
0.3
0.4
0.5
−4
−2
0
2
4
Standardised reads
Non-equivariant (P = 0.005)
Median_vaginal_canal
Subgingival_dental_plaque

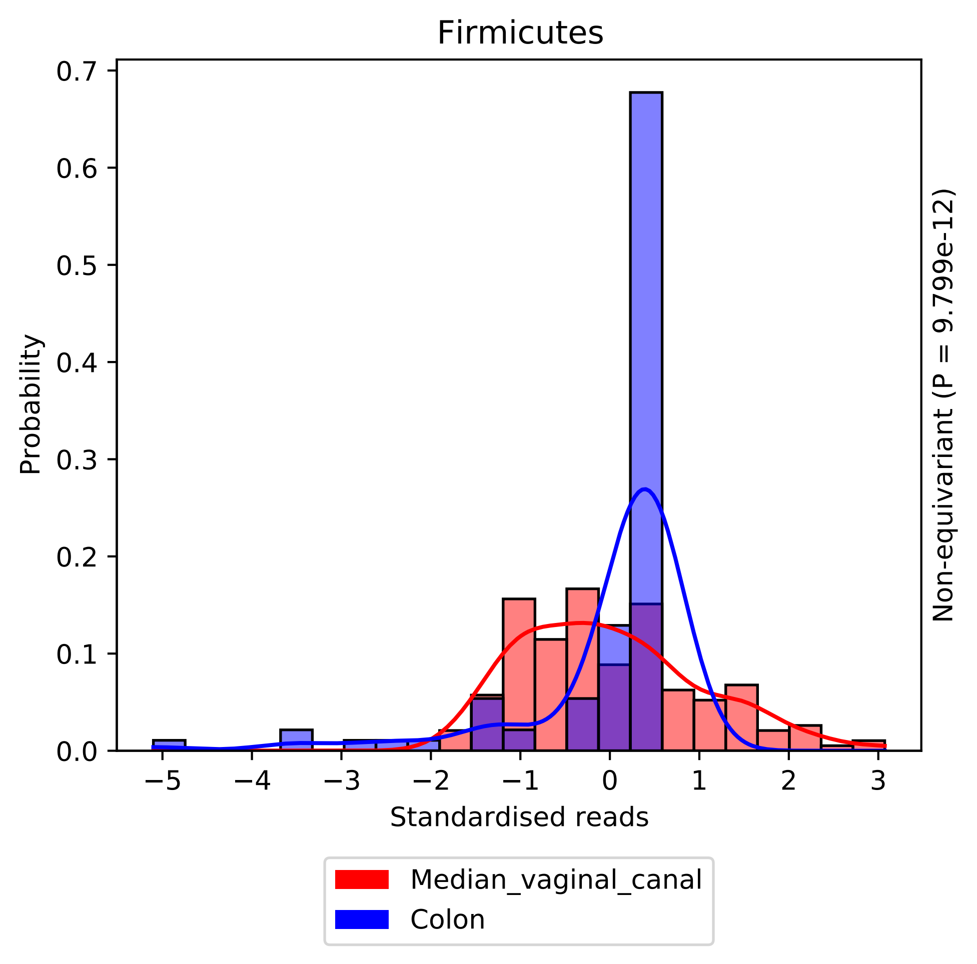

Firmicutes
Probability
0.0
0.1
0.2
0.3
0.4
0.5
0.6
0.7
−5
−4
−3
−2
−1
0
1
2
3
Standardised reads
Non-equivariant (P = 9.799e-12)
Median_vaginal_canal
Colon

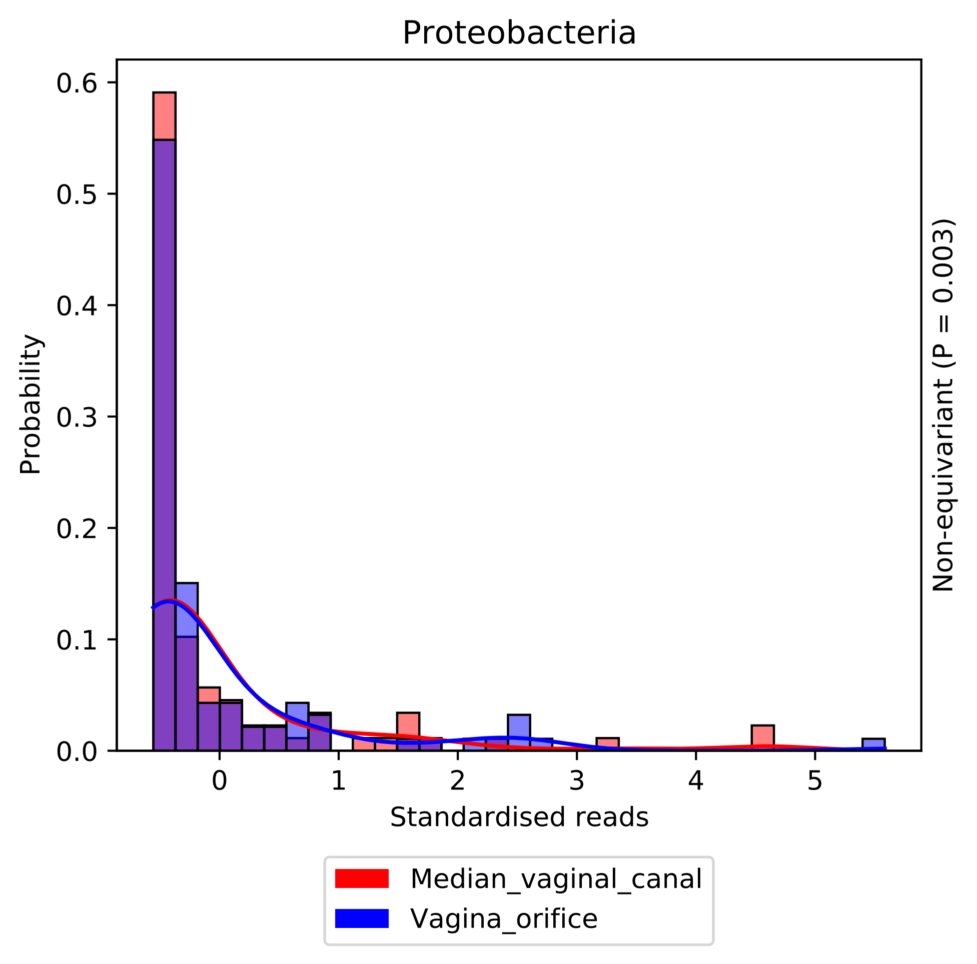

Proteobacteria
Probability
Standardised reads
Non-equivariant (P = 0.003)
0.0
0.1
0.2
0.3
0.4
0.5
0.6
0
1
2
3
4
5
Median_vaginal_canal
Vagina_orifice

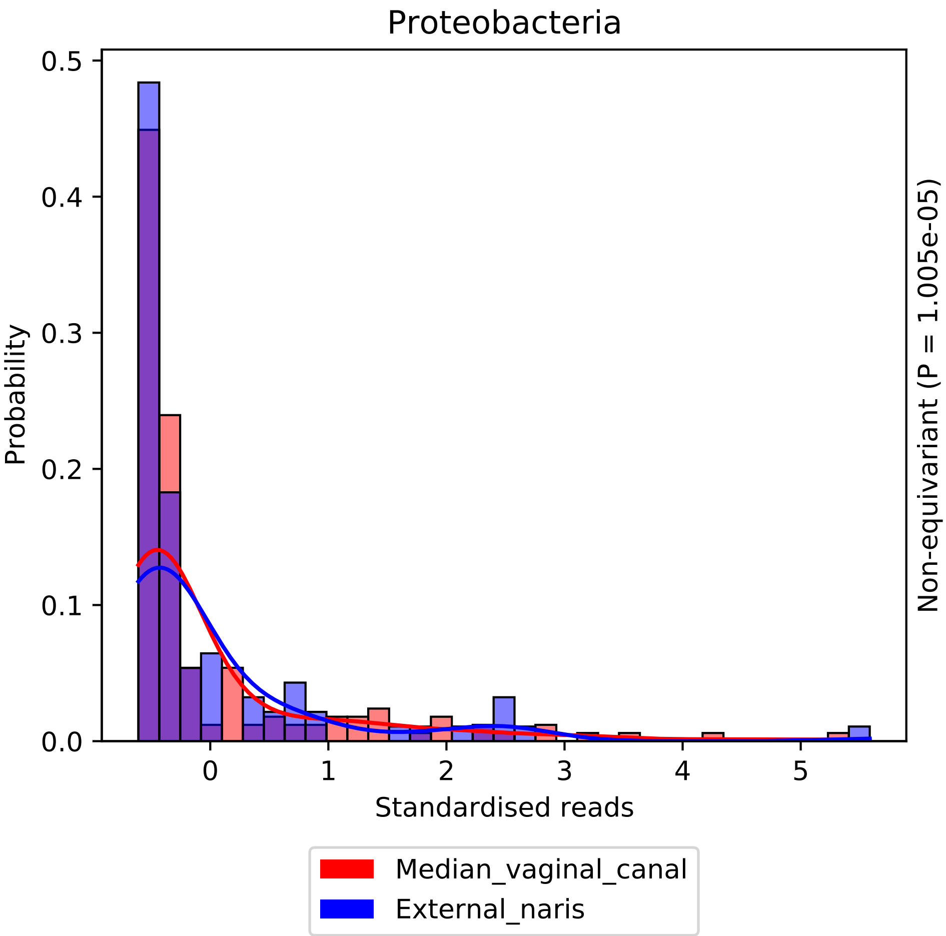
Proteobacteria
Probability
Standardised reads
Non-equivariant (P = 1.005e-05)
0.0
0.1
0.2
0.3
0.4
0.5
0
1
2
3
4
5
Median_vaginal_canal
External_naris

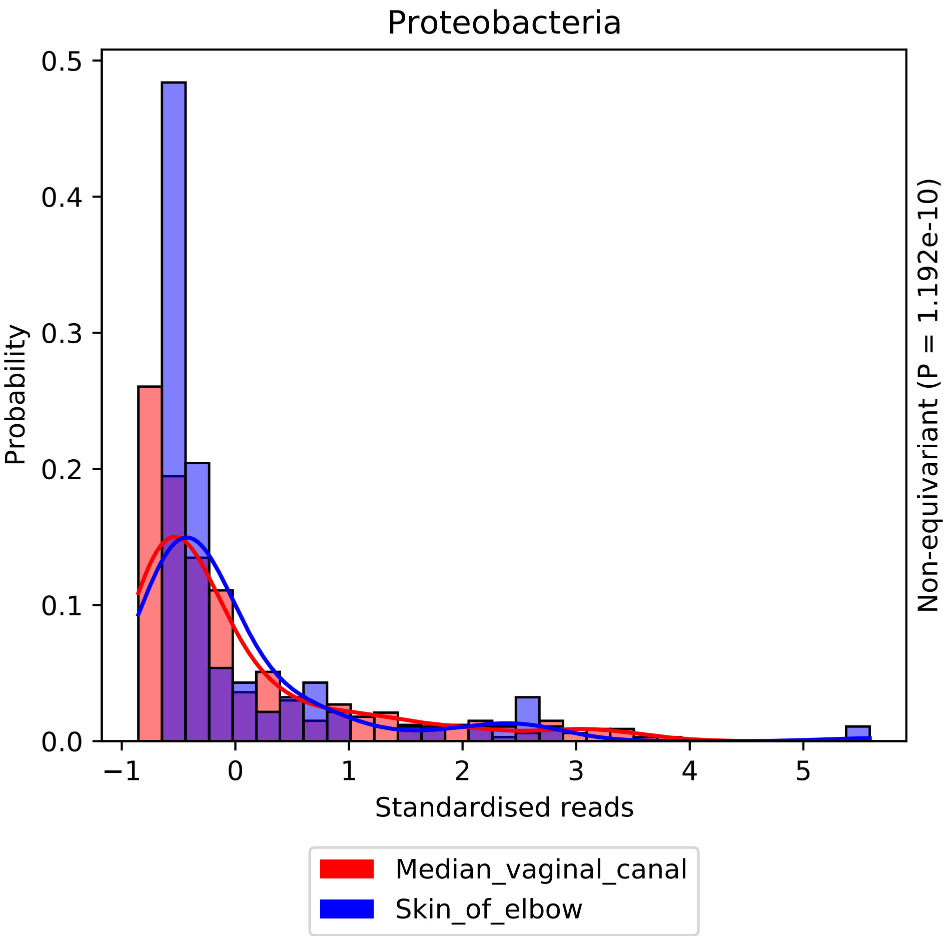

Proteobacteria
Probability
0.0
0.1
0.2
0.3
0.4
0.5
−1
0
1
2
3
4
5
Standardised reads
Non-equivariant (P = 1.192e-10)
Median_vaginal_canal
Skin_of_elbow

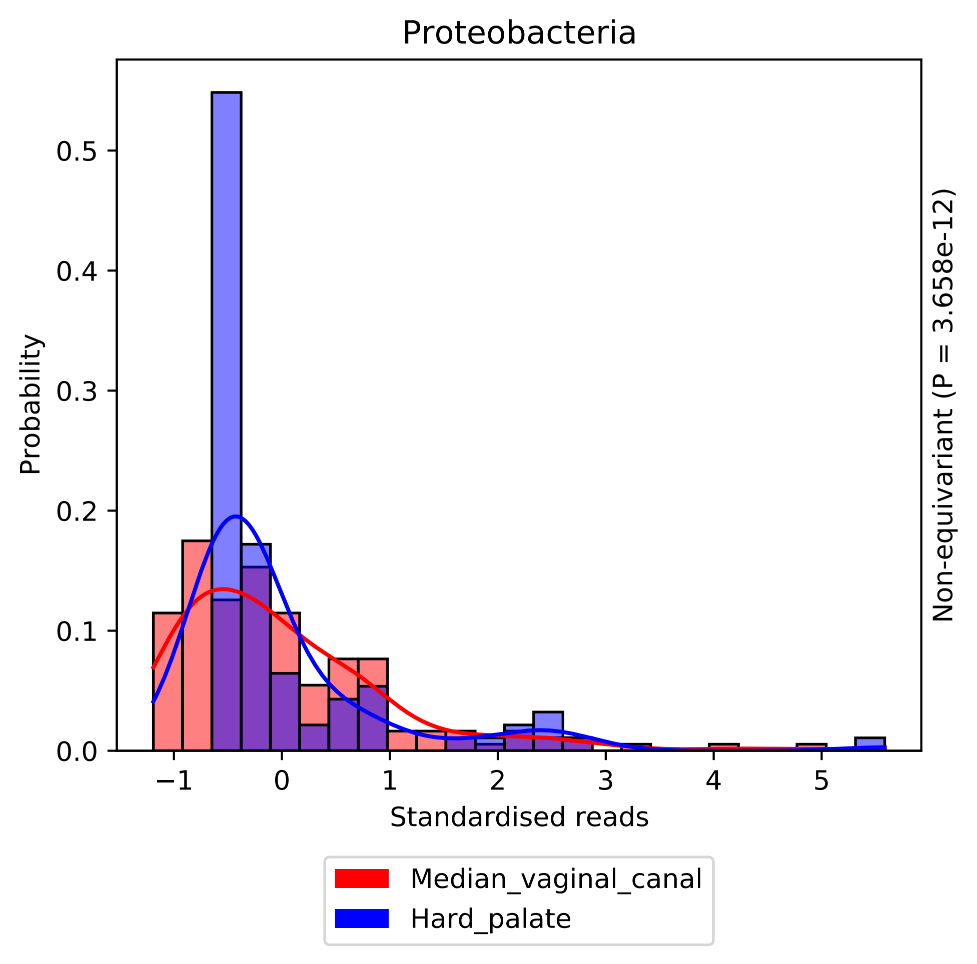

Proteobacteria
Probability
0.0
0.1
0.2
0.3
0.4
0.5
−1
0
1
2
3
4
5
Standardised reads
Non-equivariant (P = 3.658e-12)
Median_vaginal_canal
Hard_palate

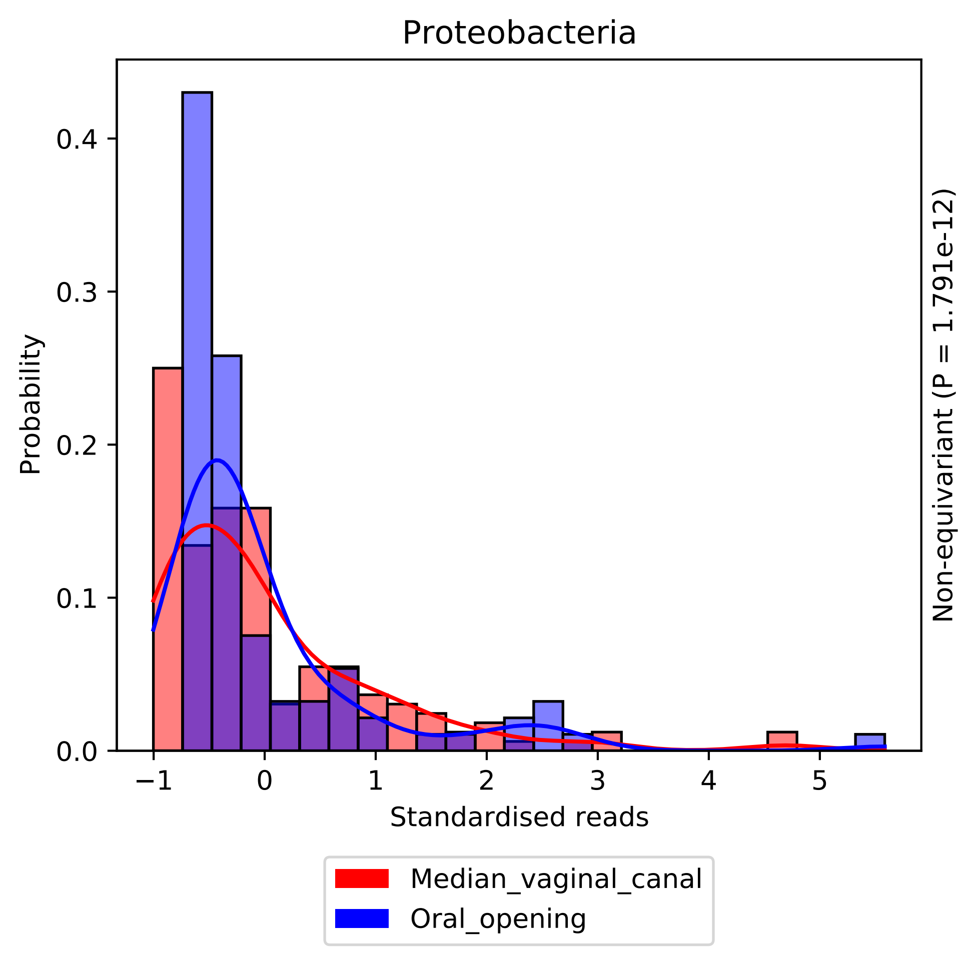

Proteobacteria
Probability
0.0
0.1
0.2
0.3
0.4
−1
0
1
2
3
4
5
Standardised reads
Non-equivariant (P = 1.791e-12)
Median_vaginal_canal
Oral_opening

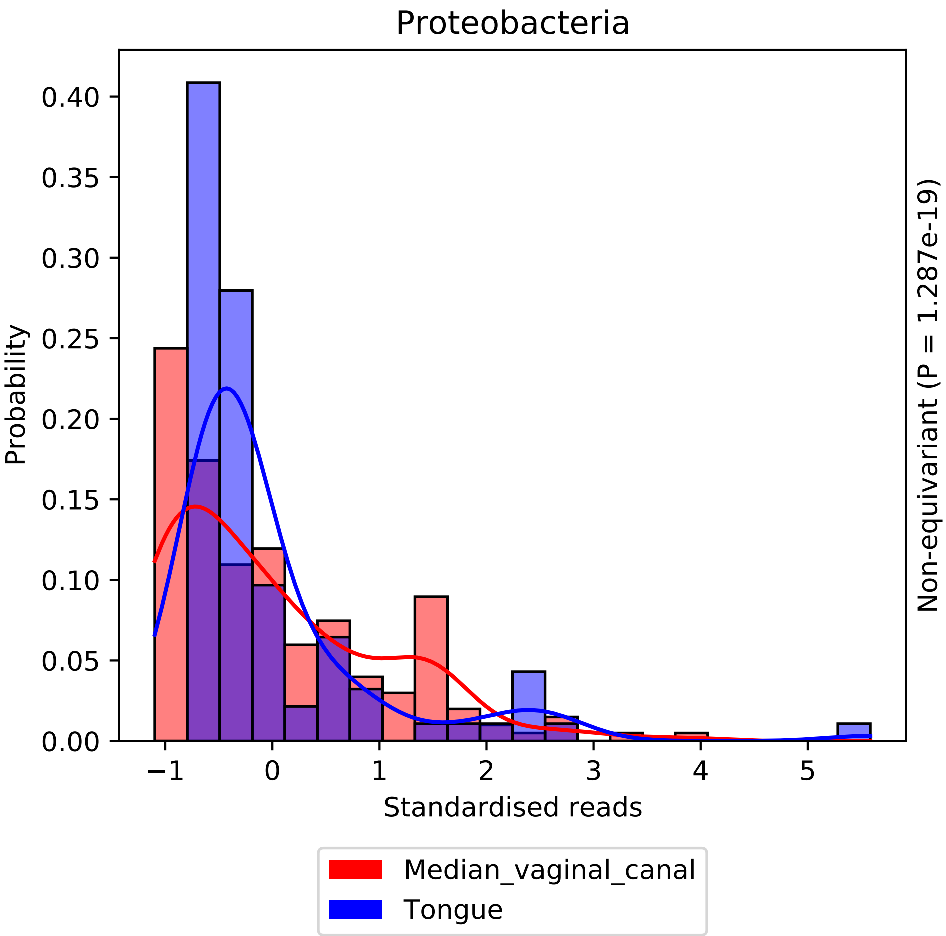

Proteobacteria
Probability
0.00
0.05
0.10
0.15
0.20
0.25
0.30
0.35
0.40
−1
0
1
2
3
4
5
Standardised reads
Non-equivariant (P = 1.287e-19)
Median_vaginal_canal
Tongue

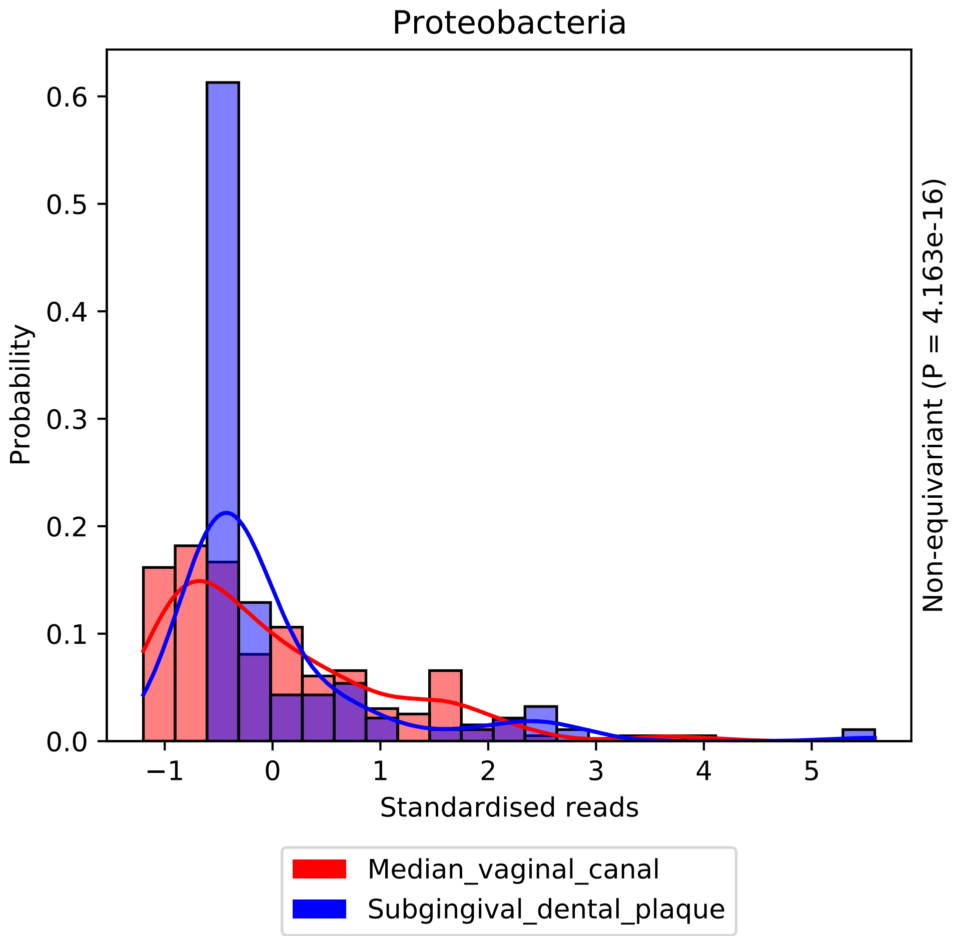

Proteobacteria
Probability
0.0
0.1
0.2
0.3
0.4
0.5
0.6
−1
0
1
2
3
4
5
Standardised reads
Non-equivariant (P = 4.163e-16)
Median_vaginal_canal
Subgingival_dental_plaque

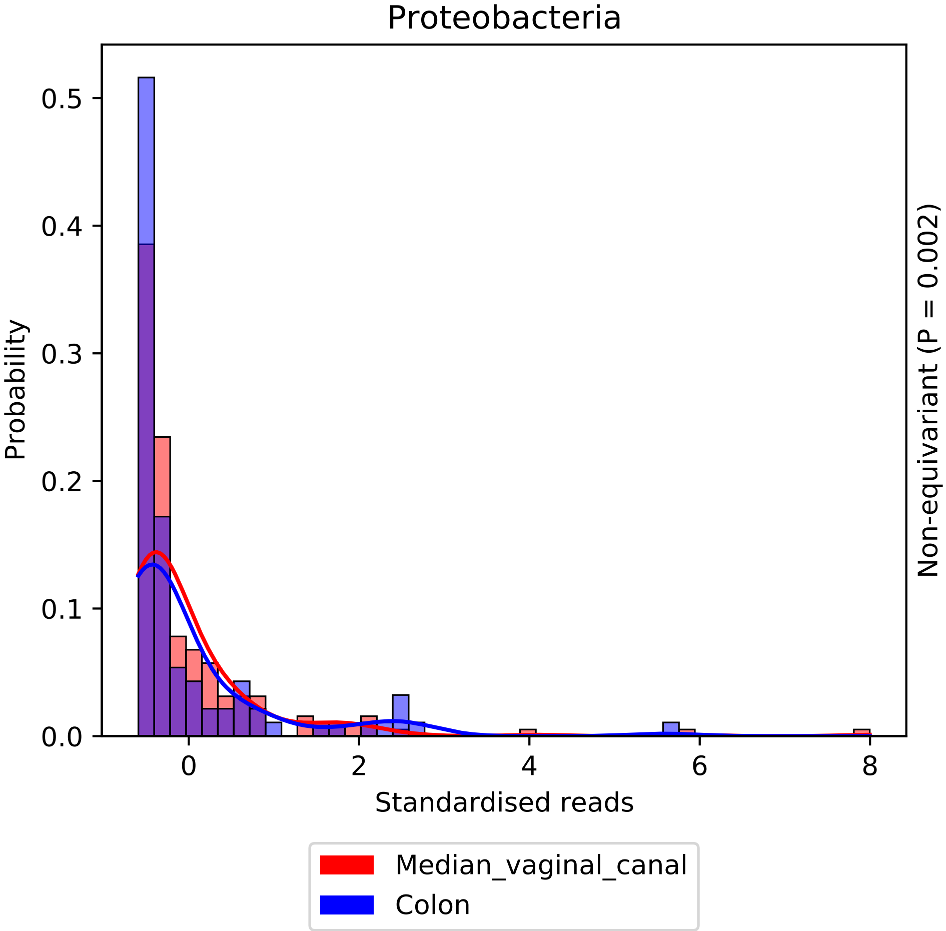

Proteobacteria
Probability
0.0
0.1
0.2
0.3
0.4
0.5
0
2
4
6
8
Standardised reads
Non-equivariant (P = 0.002)
Median_vaginal_canal
Colon

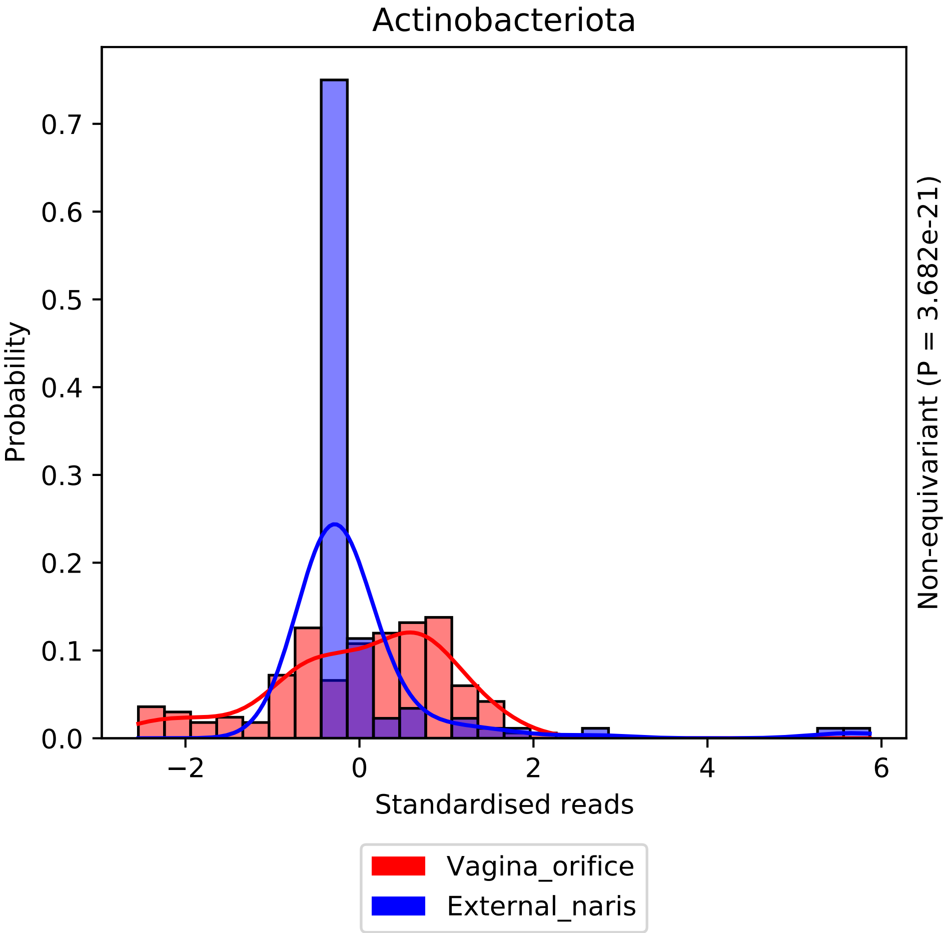

Actinobacteriota
Probability
Standardised reads
Non-equivariant (P = 3.682e-21)
0.0
0.1
0.2
0.3
0.4
0.5
0.6
0.7
−2
0
2
4
6
Vagina_orifice
External_naris

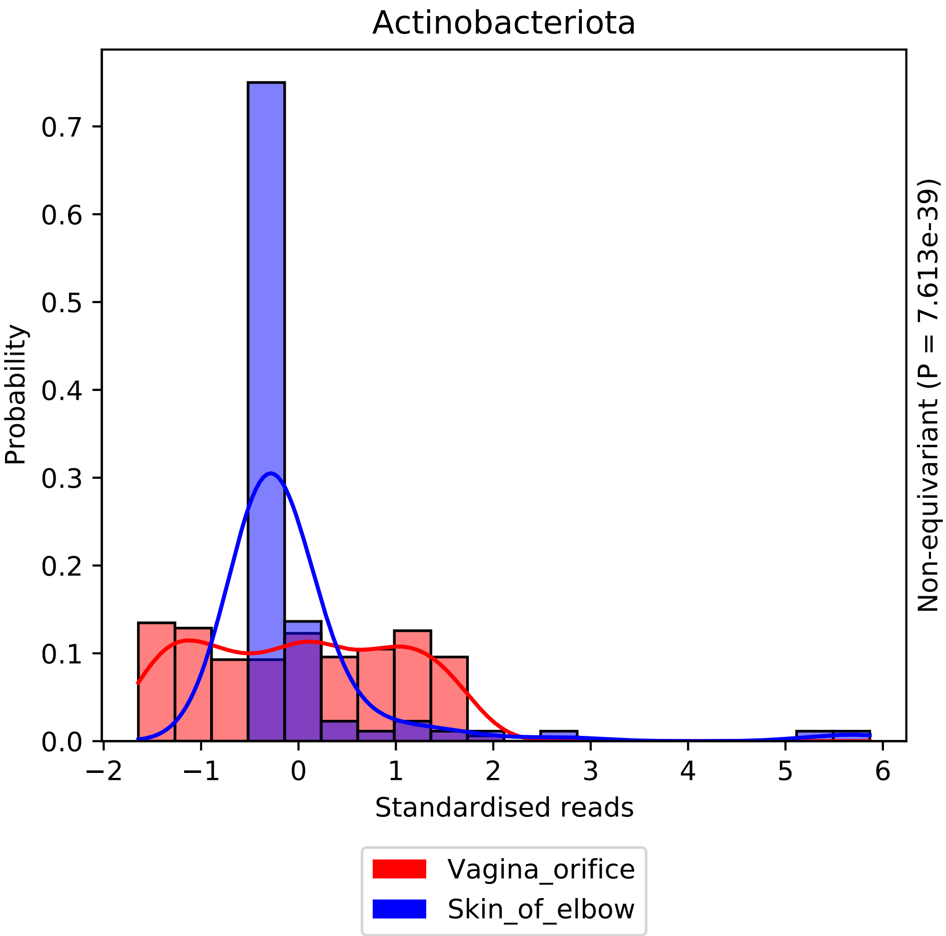

Actinobacteriota
Probability
0.0
0.1
0.2
0.3
0.4
0.5
0.6
0.7
−2
−1
0
1
2
3
4
5
6
Standardised reads
Non-equivariant (P = 7.613e-39)
Vagina_orifice
Skin_of_elbow

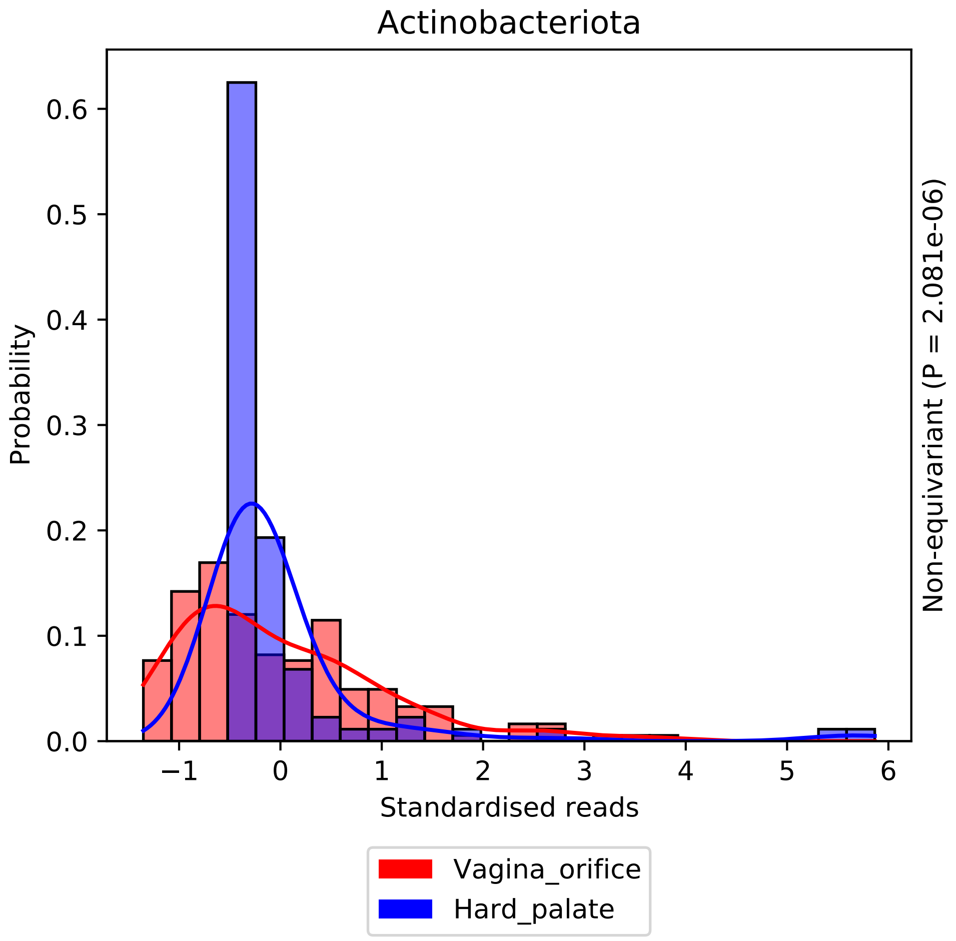

Actinobacteriota
Probability
0.0
0.1
0.2
0.3
0.4
0.5
0.6
−1
0
1
2
3
4
5
6
Standardised reads
Non-equivariant (P = 2.081e-06)
Vagina_orifice
Hard_palate

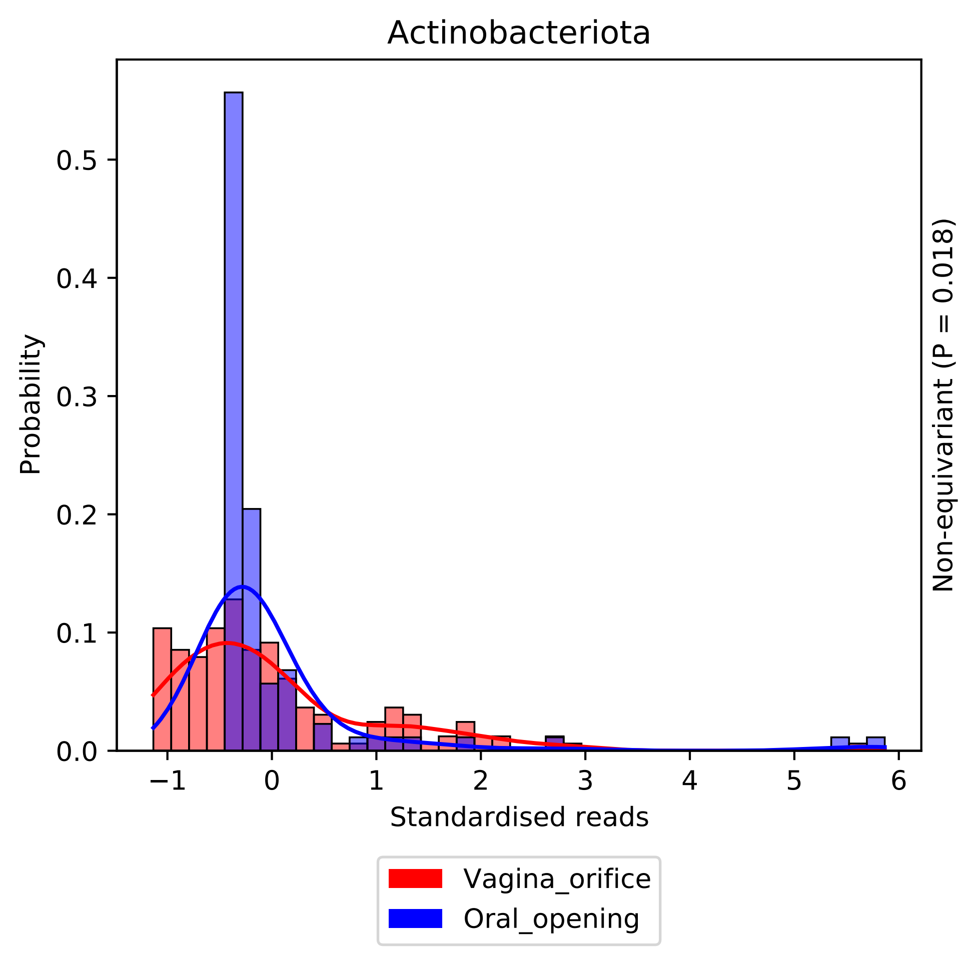
Actinobacteriota
Probability
0.0
0.1
0.2
0.3
0.4
0.5
−1
0
1
2
3
4
5
6
Standardised reads
Non-equivariant (P = 0.018)
Vagina_orifice
Oral_opening

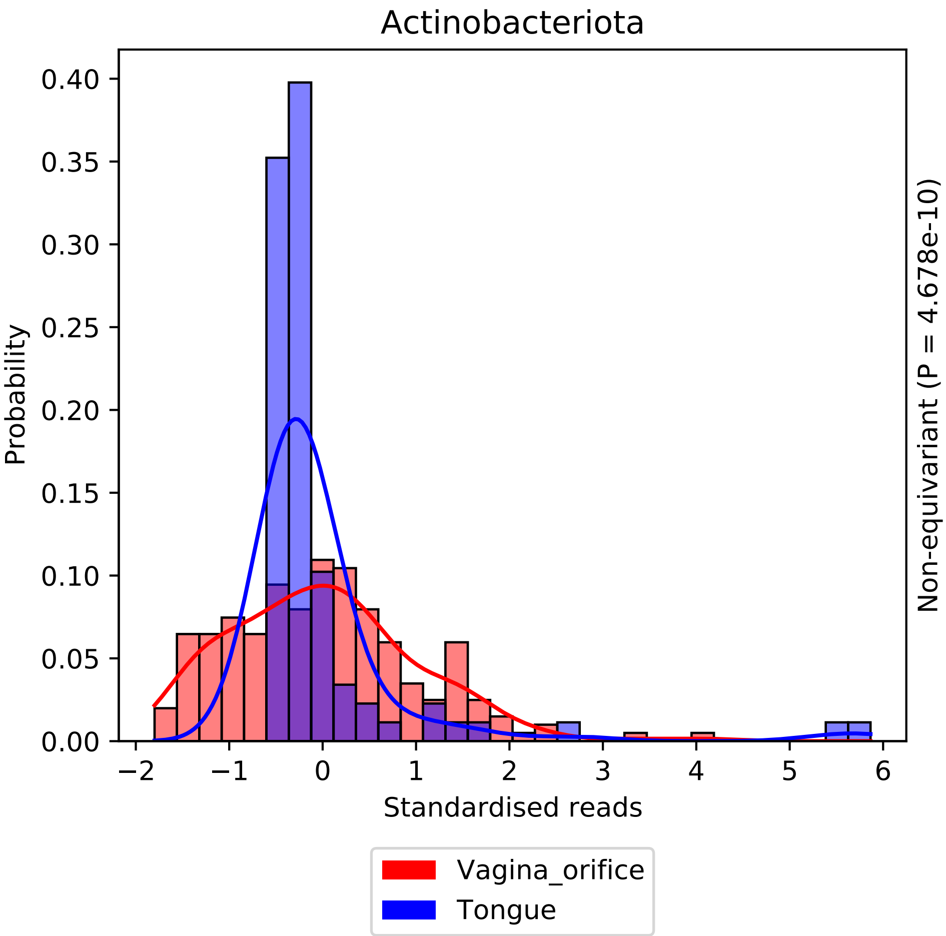

Actinobacteriota
Probability
0.40
0.35
0.30
0.25
0.20
0.15
0.10
0.05
0.00
−2
−1
0
1
2
3
4
5
6
Standardised reads
Non-equivariant (P = 4.678e-10)
Vagina_orifice
Tongue

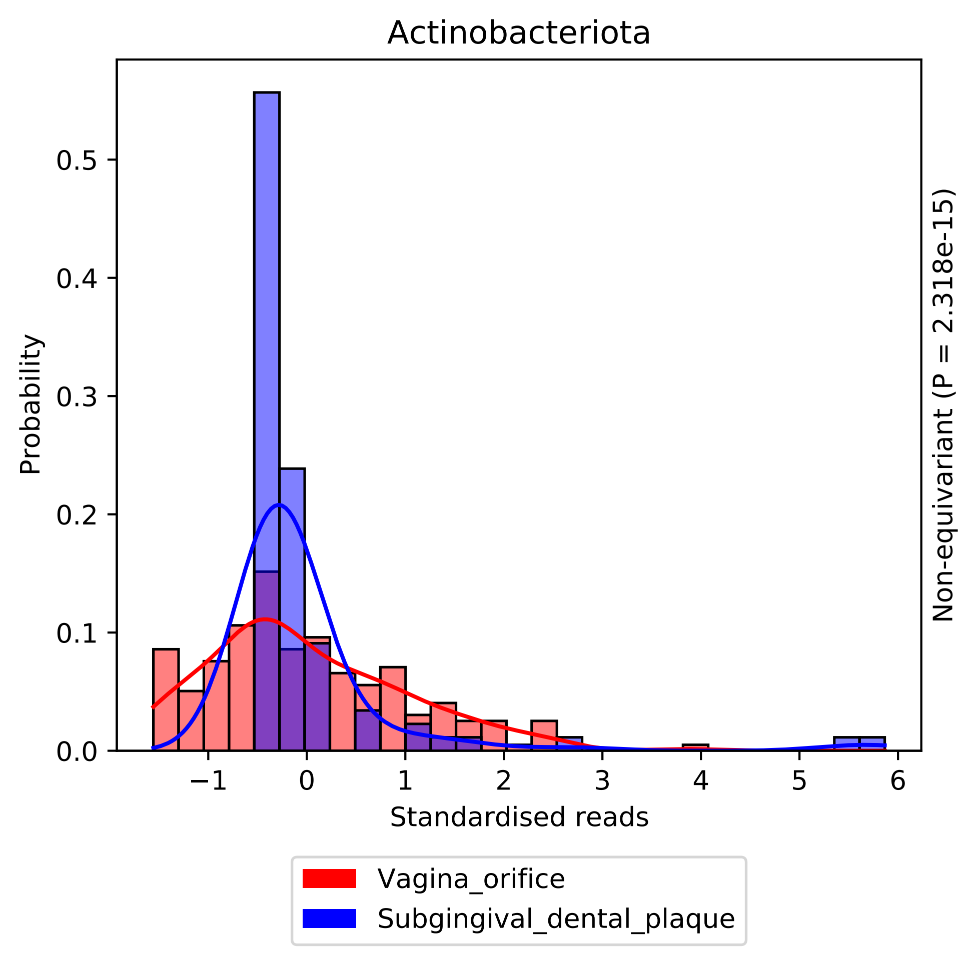

Actinobacteriota
Probability
Standardised reads
Non-equivariant (P = 2.318e-15)
0.0
0.1
0.2
0.3
0.4
0.5
−1
0
1
2
3
4
5
6
Vagina_orifice
Subgingival_dental_plaque

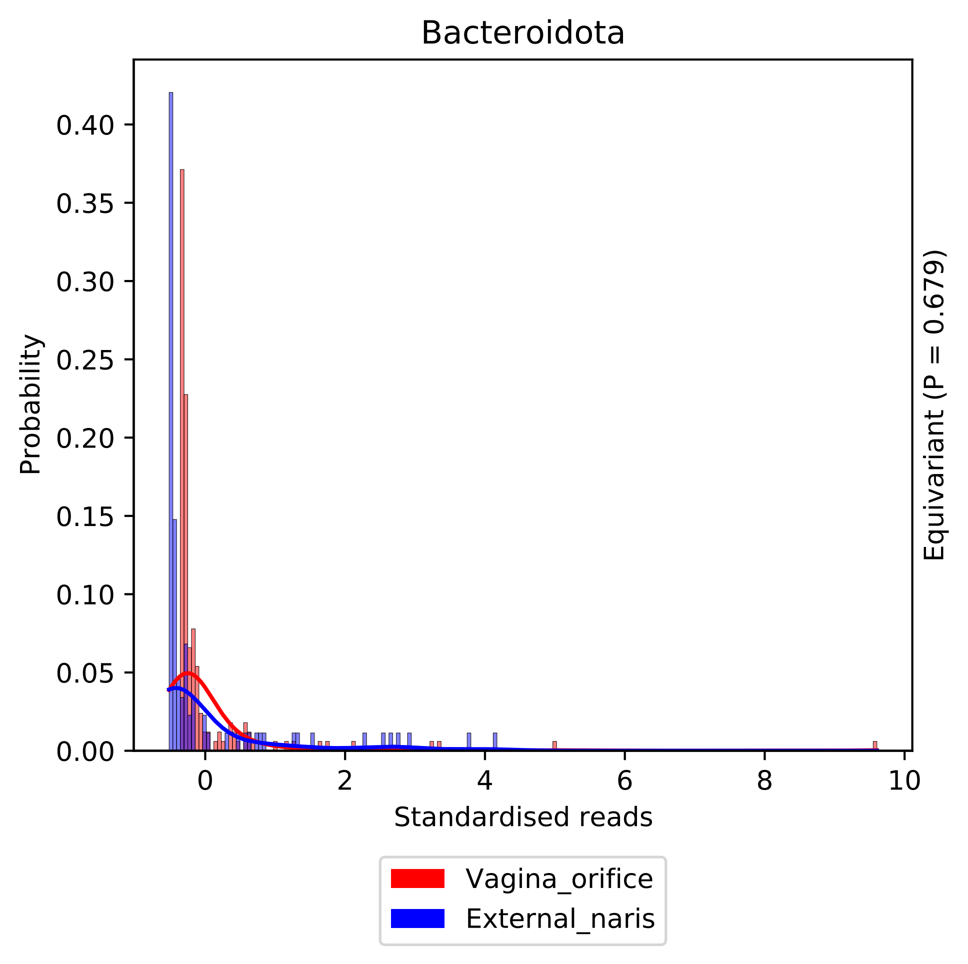

Bacteroidota
Probability
0.40
0.35
0.30
0.25
0.20
0.15
0.10
0.05
0.00
0
2
4
6
8
10
Standardised reads
Equivariant (P = 0.679)
Vagina_orifice
External_naris

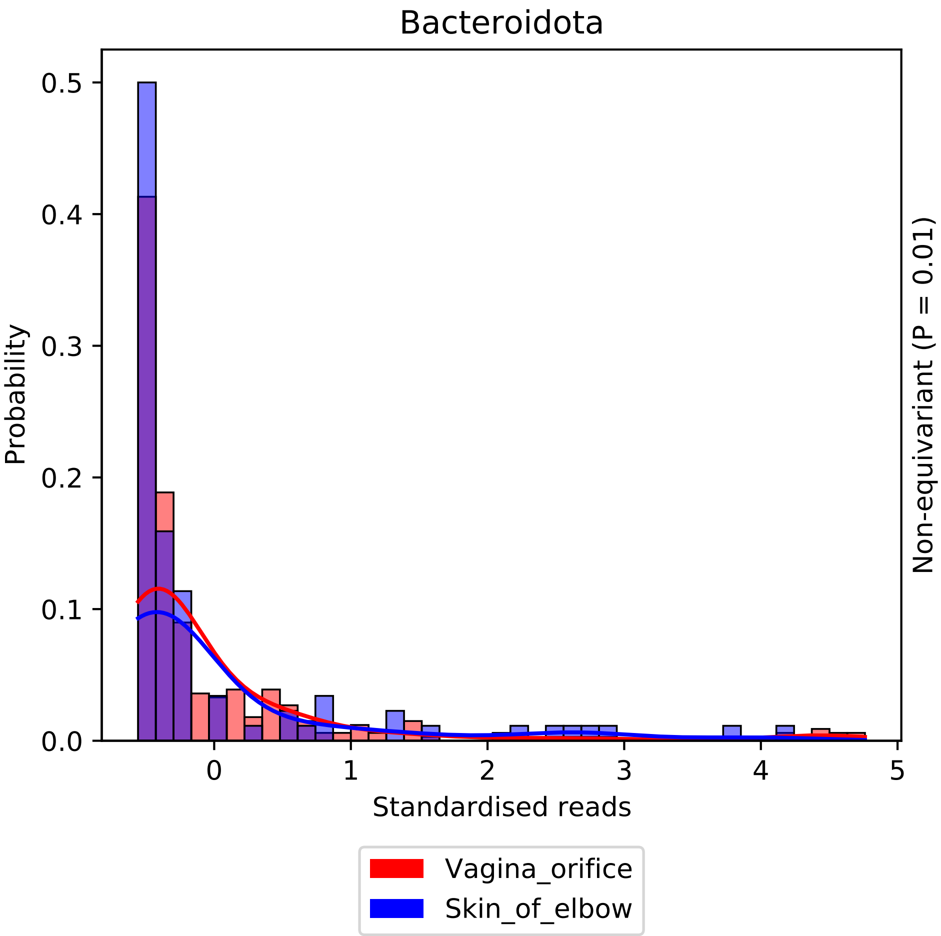

Bacteroidota
Probability
0.5
0.4
0.3
0.2
0.1
0.0
0
1
2
3
4
5
Standardised reads
Non-equivariant (P = 0.01)
Vagina_orifice
Skin_of_elbow

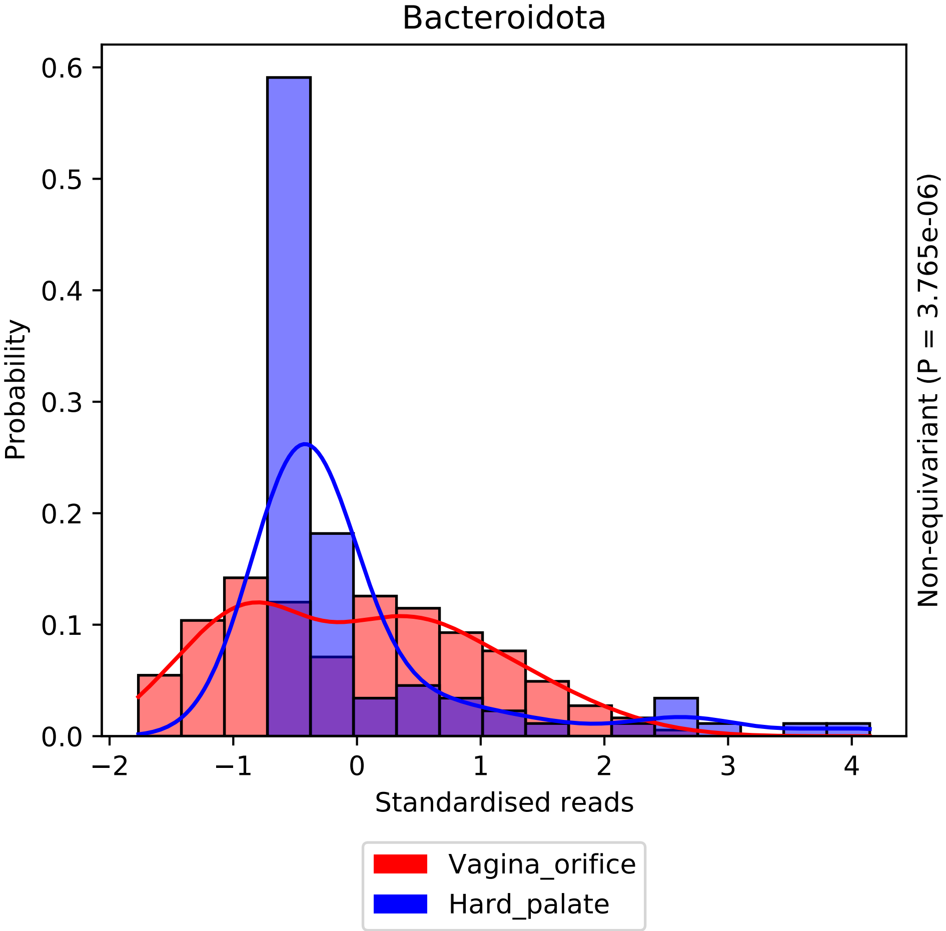
Bacteroidota
Probability
0.0
0.1
0.2
0.3
0.4
0.5
0.6
−2
−1
0
1
2
3
4
Standardised reads
Non-equivariant (P = 3.765e-06)
Vagina_orifice
Hard_palate

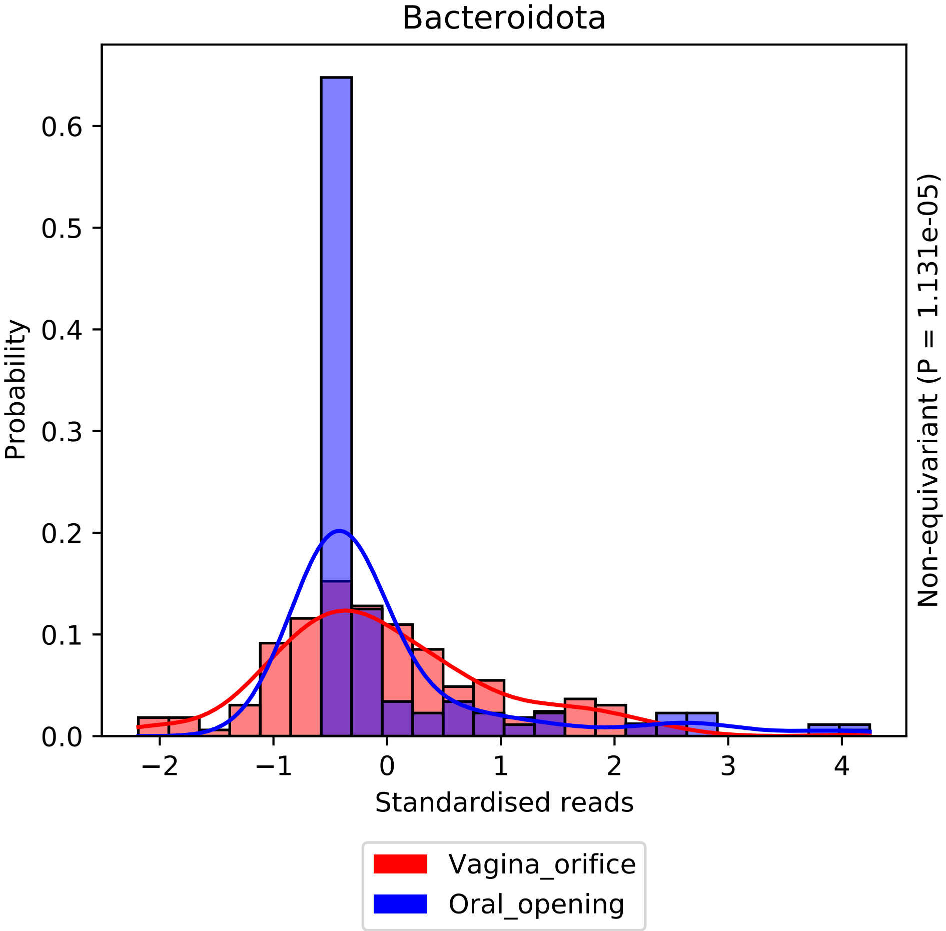
Bacteroidota
Probability
Standardised reads
Non-equivariant (P = 1.131e-05)
Vagina_orifice
Oral_opening
0.0
0.1
0.2
0.3
0.4
0.5
0.6
−2
−1
0
1
2
3
4

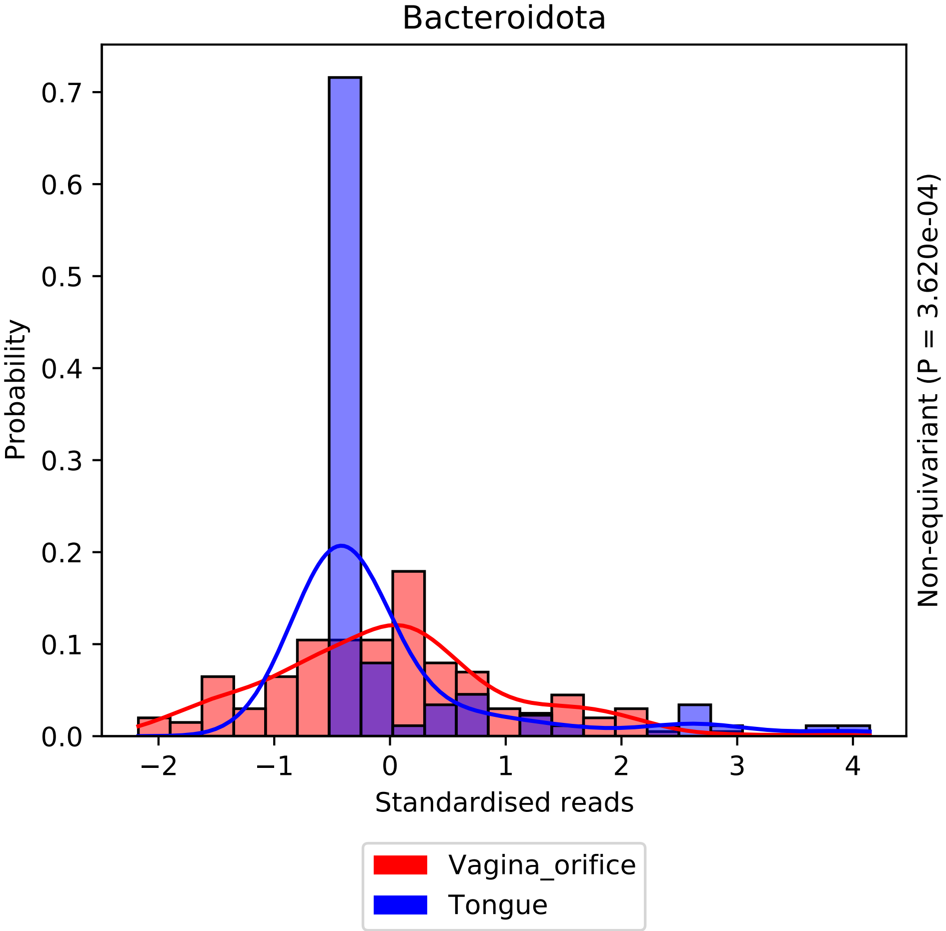

Bacteroidota
Probability
Standardised reads
Non-equivariant (P = 3.620e-04)
0.0
0.1
0.2
0.3
0.4
0.5
0.6
0.7
−2
−1
0
1
2
3
4
Vagina_orifice
Tongue

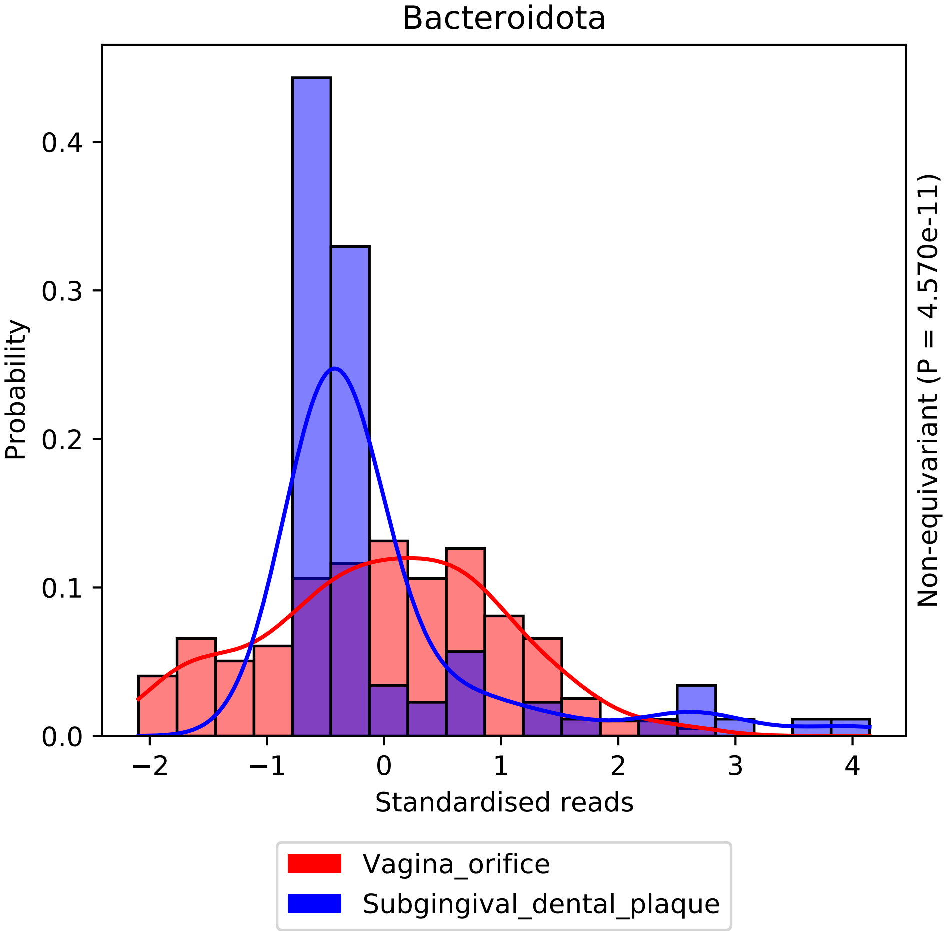
Bacteroidota
Probability
0.0
0.1
0.2
0.3
0.4
−2
−1
0
1
2
3
4
Standardised reads
Non-equivariant (P = 4.570e-11)
Vagina_orifice
Subgingival_dental_plaque

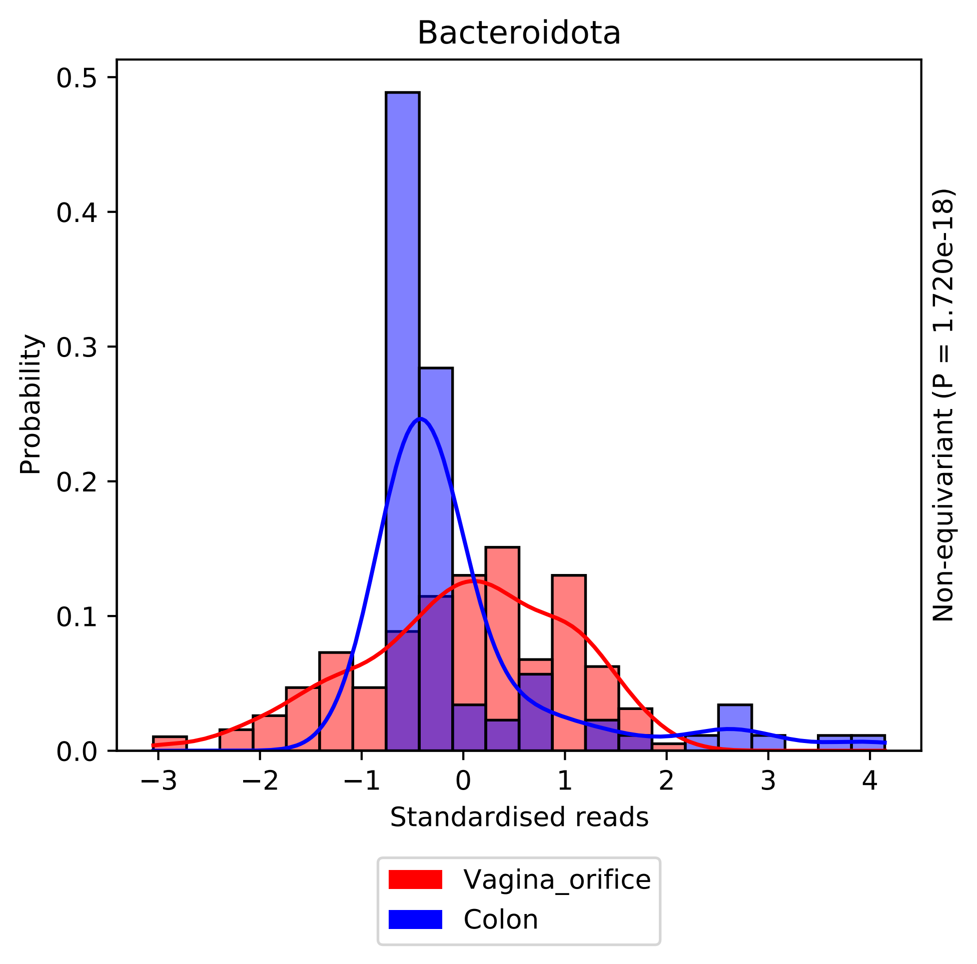
Bacteroidota
Probability
0.0
0.1
0.2
0.3
0.4
0.5
−3
−2
−1
0
1
2
3
4
Standardised reads
Non-equivariant (P = 1.720e-18)
Vagina_orifice
Colon

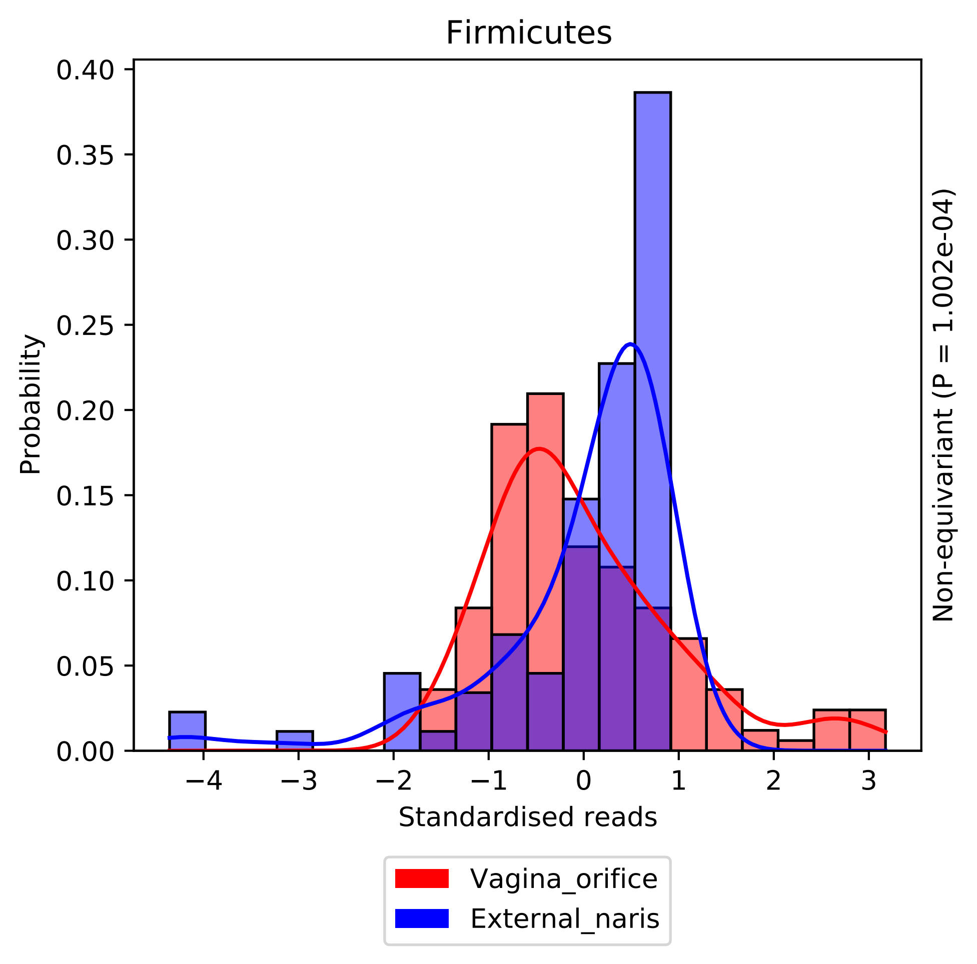

Firmicutes
0.40
0.35
0.30
0.25
0.20
0.15
0.10
0.05
0.00
Probability
−4
−3
−2
−1
0
1
2
3
Standardised reads
Non-equivariant (P = 1.002e-04)
Vagina_orifice
External_naris

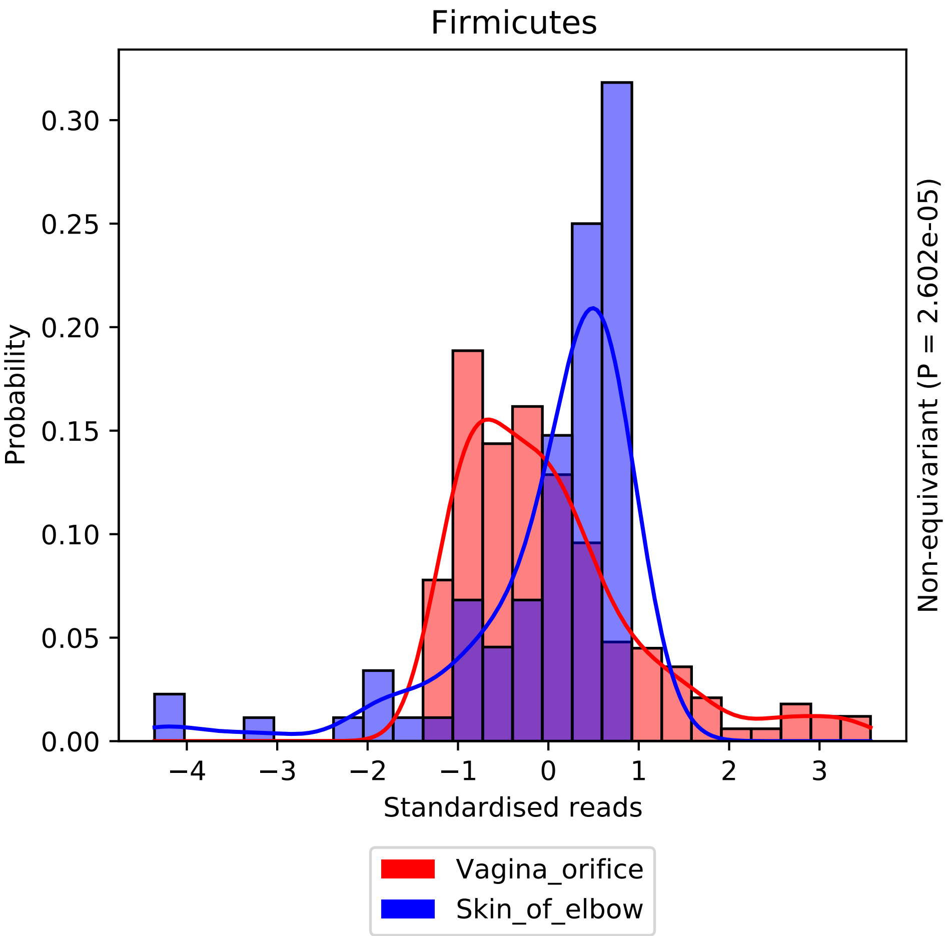

Firmicutes
Probability
0.00
0.05
0.10
0.15
0.20
0.25
0.30
−4
−3
−2
−1
0
1
2
3
Standardised reads
Non-equivariant (P = 2.602e-05)
Vagina_orifice
Skin_of_elbow

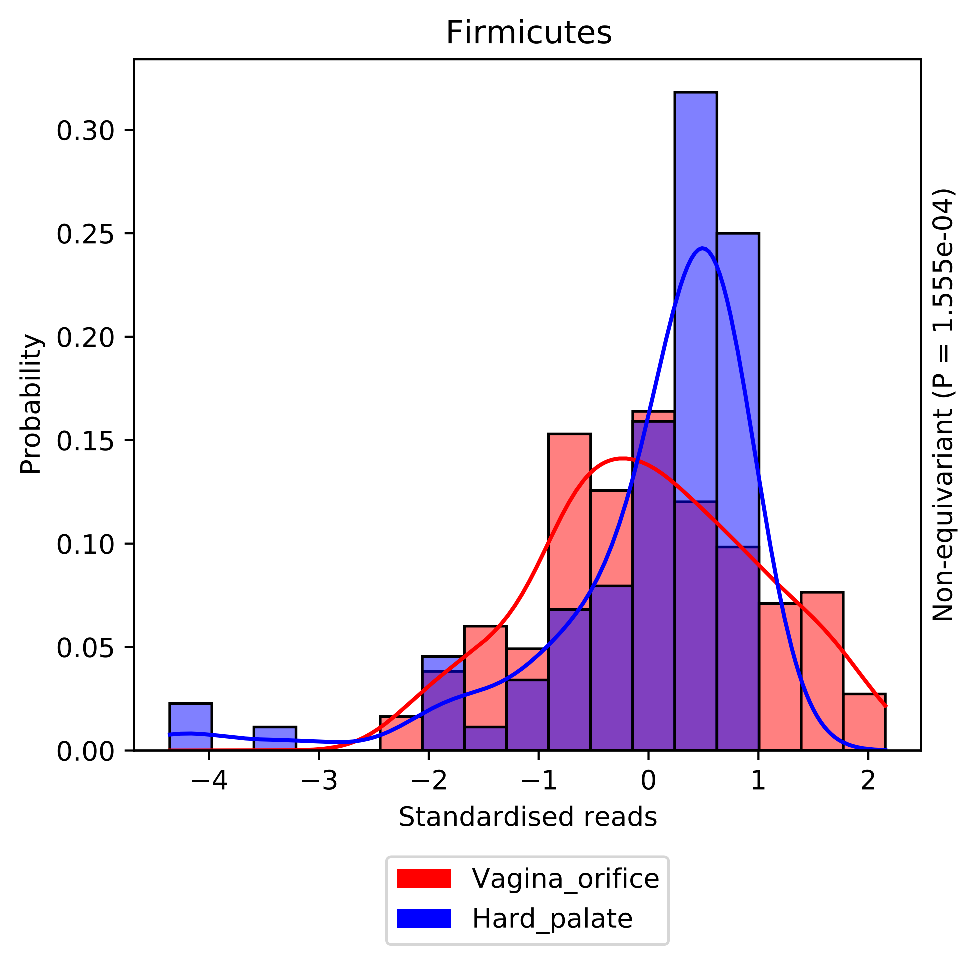

Firmicutes
Probability
0.00
0.05
0.10
0.15
0.20
0.25
0.30
−4
−3
−2
−1
0
1
2
Standardised reads
Non-equivariant (P = 1.555e-04)
Vagina_orifice
Hard_palate

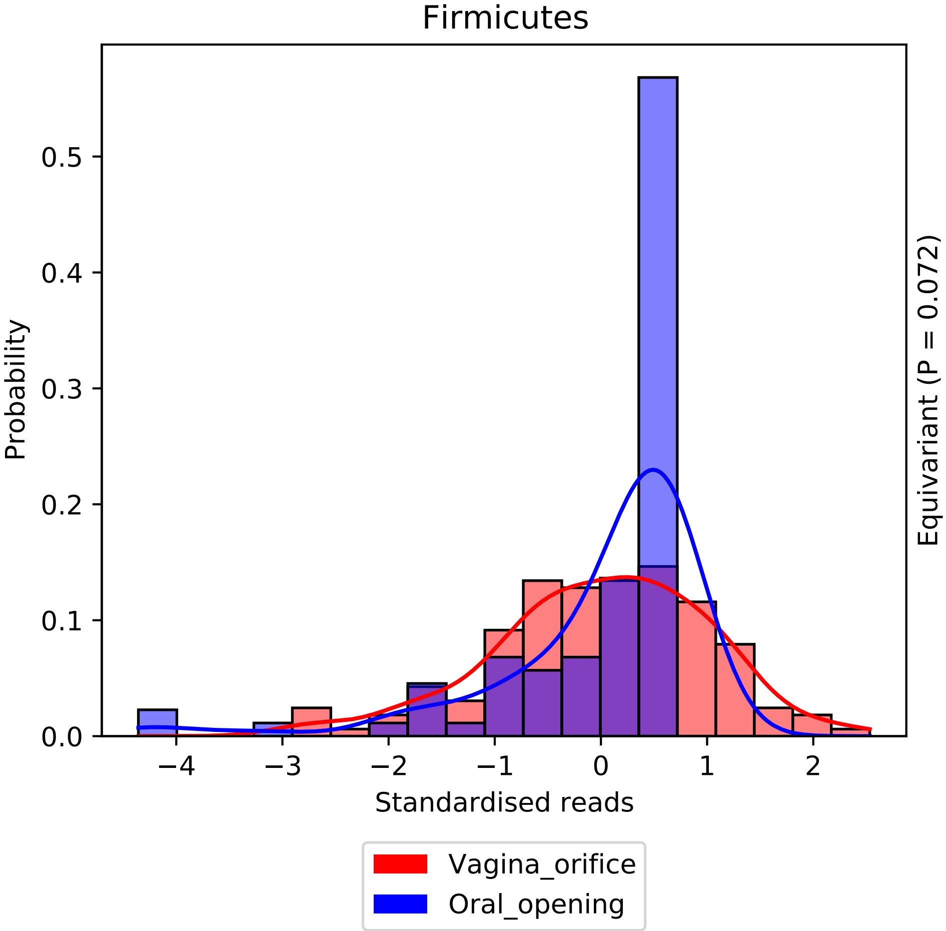
Firmicutes
Probability
Standardised reads
Equivariant (P = 0.072)
0.0
0.1
0.2
0.3
0.4
0.5
−4
−3
−2
−1
0
1
2
Vagina_orifice
Oral_opening

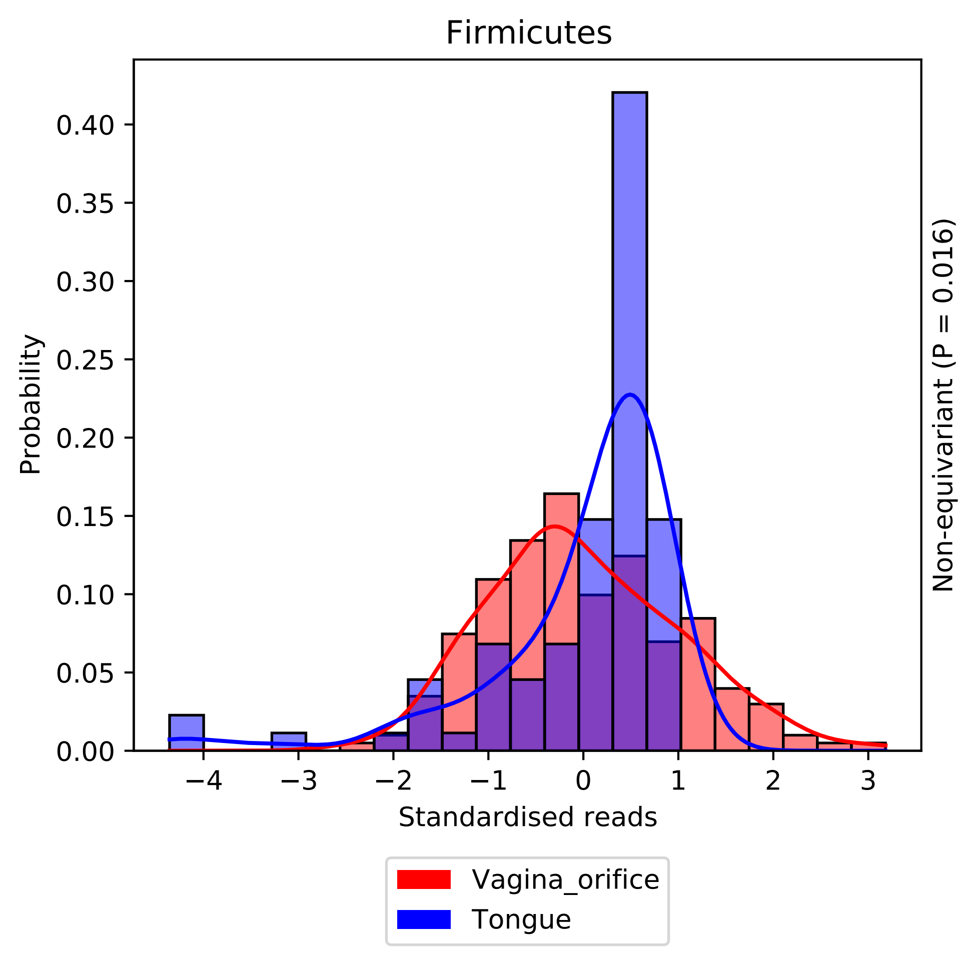
Firmicutes
Probability
0.00
0.05
0.10
0.15
0.20
0.25
0.30
0.35
0.40
−4
−3
−2
−1
0
1
2
3
Standardised reads
Non-equivariant (P = 0.016)
Vagina_orifice
Tongue

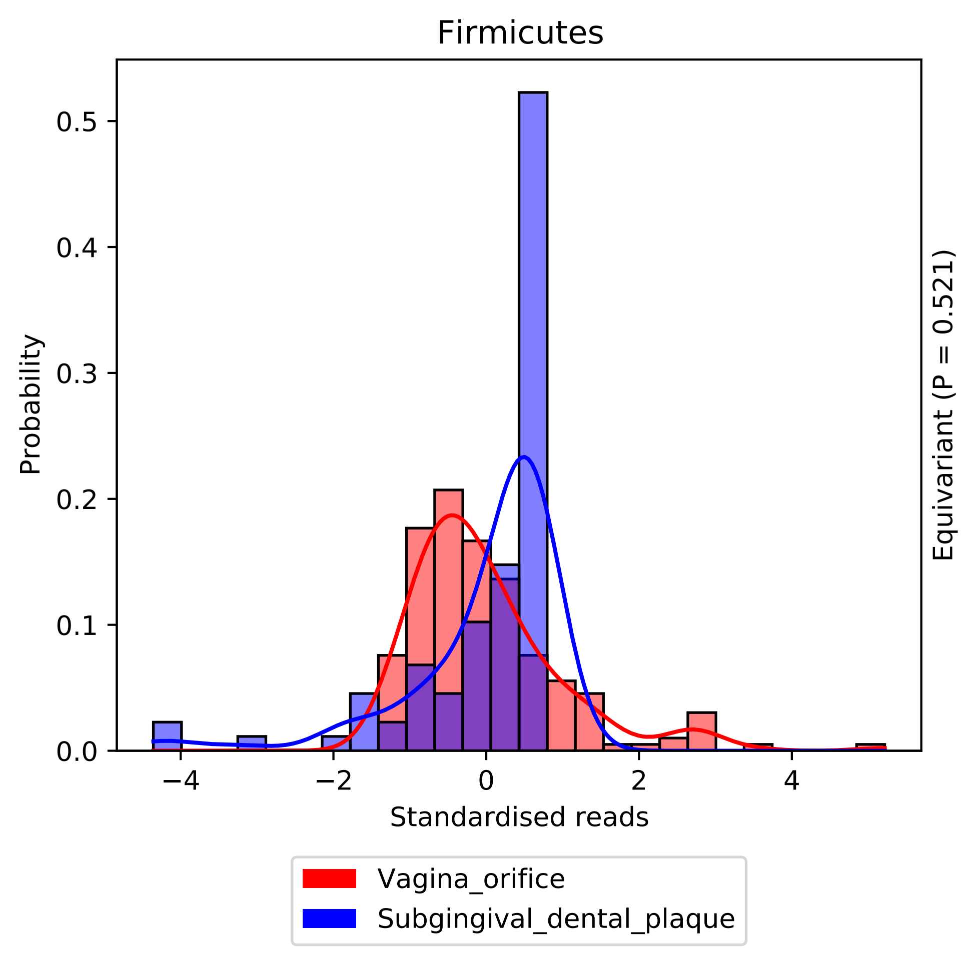

Firmicutes
Probability
Standardised reads
Equivariant (P = 0.521)
0.0
0.1
0.2
0.3
0.4
0.5
−4
−2
0
2
4
Vagina_orifice
Subgingival_dental_plaque

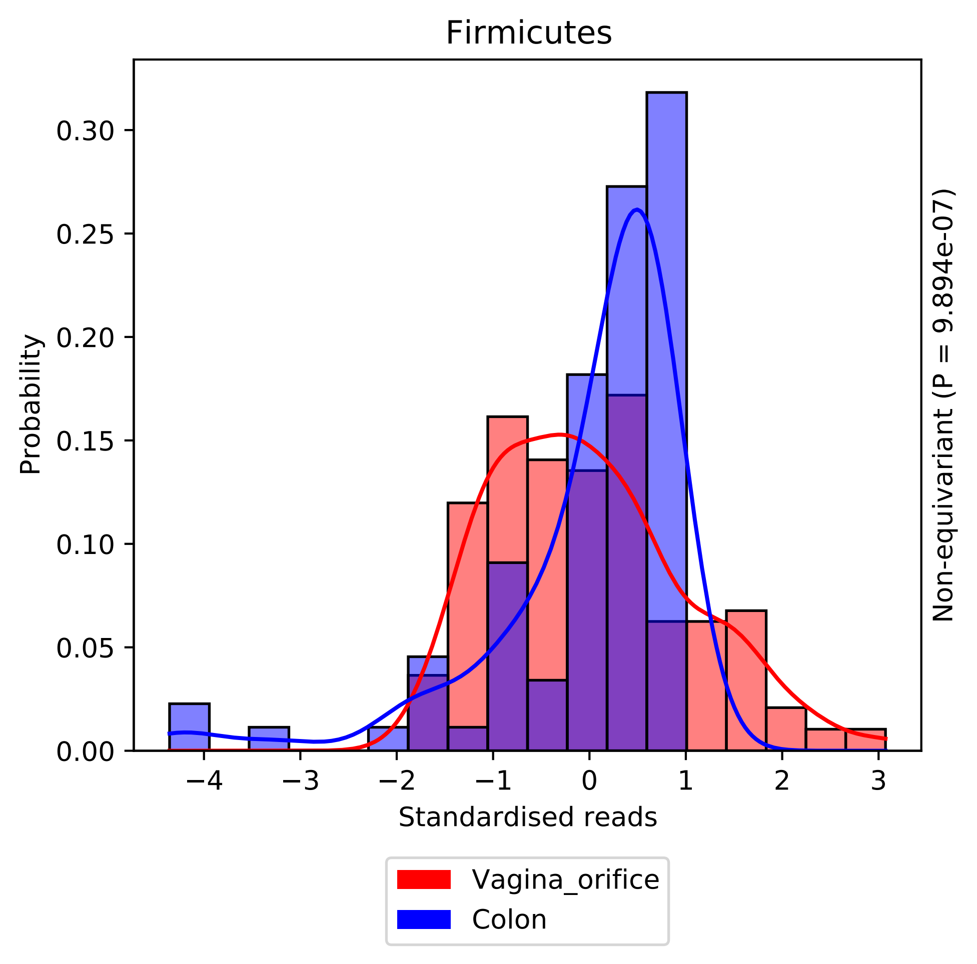

Firmicutes
Probability
0.30
0.25
0.20
0.15
0.10
0.05
0.00
−4
−3
−2
−1
0
1
2
3
Standardised reads
Non-equivariant (P = 9.894e-07)
Vagina_orifice
Colon

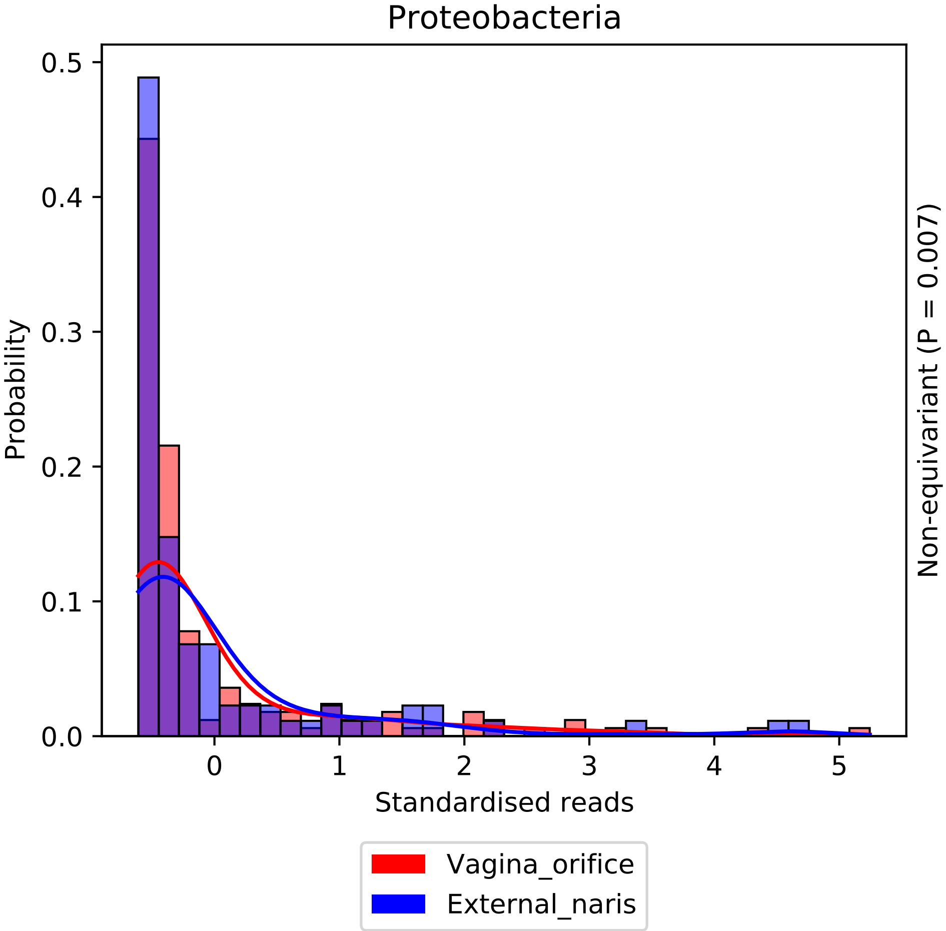

Proteobacteria
Probability
0.0
0.1
0.2
0.3
0.4
0.5
0
1
2
3
4
5
Standardised reads
Non-equivariant (P = 0.007)
Vagina_orifice
External_naris

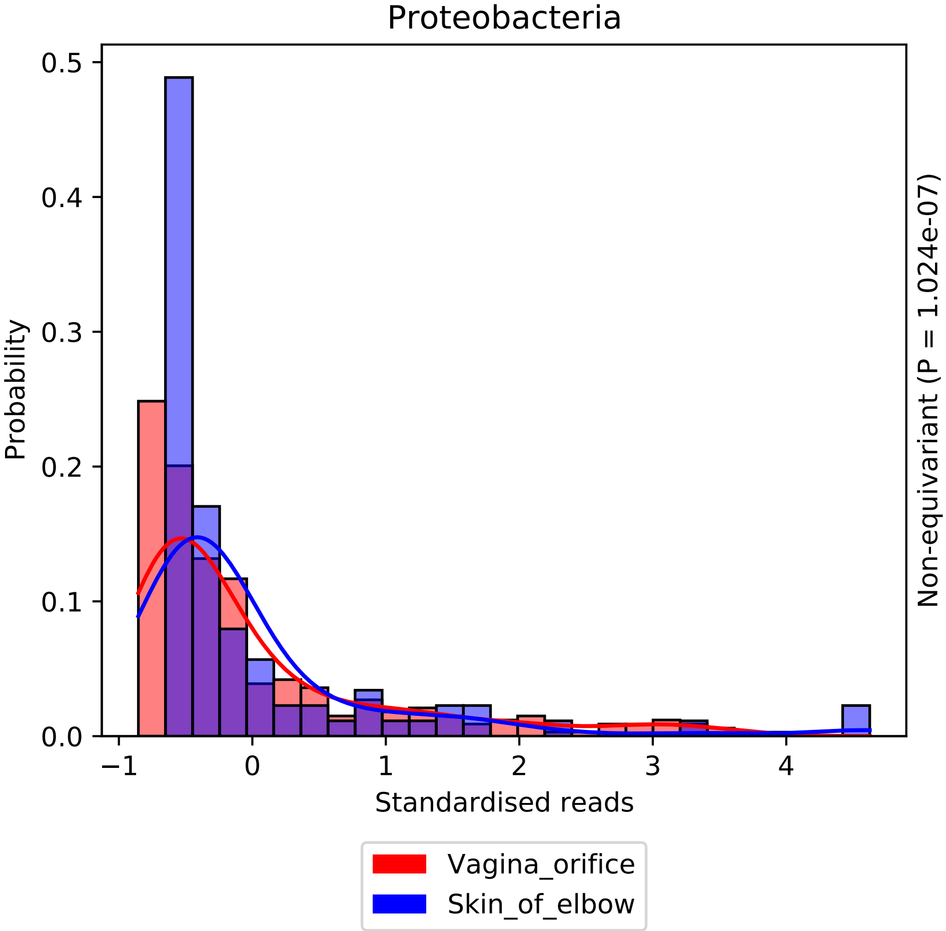

Proteobacteria
Probability
0.0
0.1
0.2
0.3
0.4
0.5
−1
0
1
2
3
4
Standardised reads
Non-equivariant (P = 1.024e-07)
Vagina_orifice
Skin_of_elbow

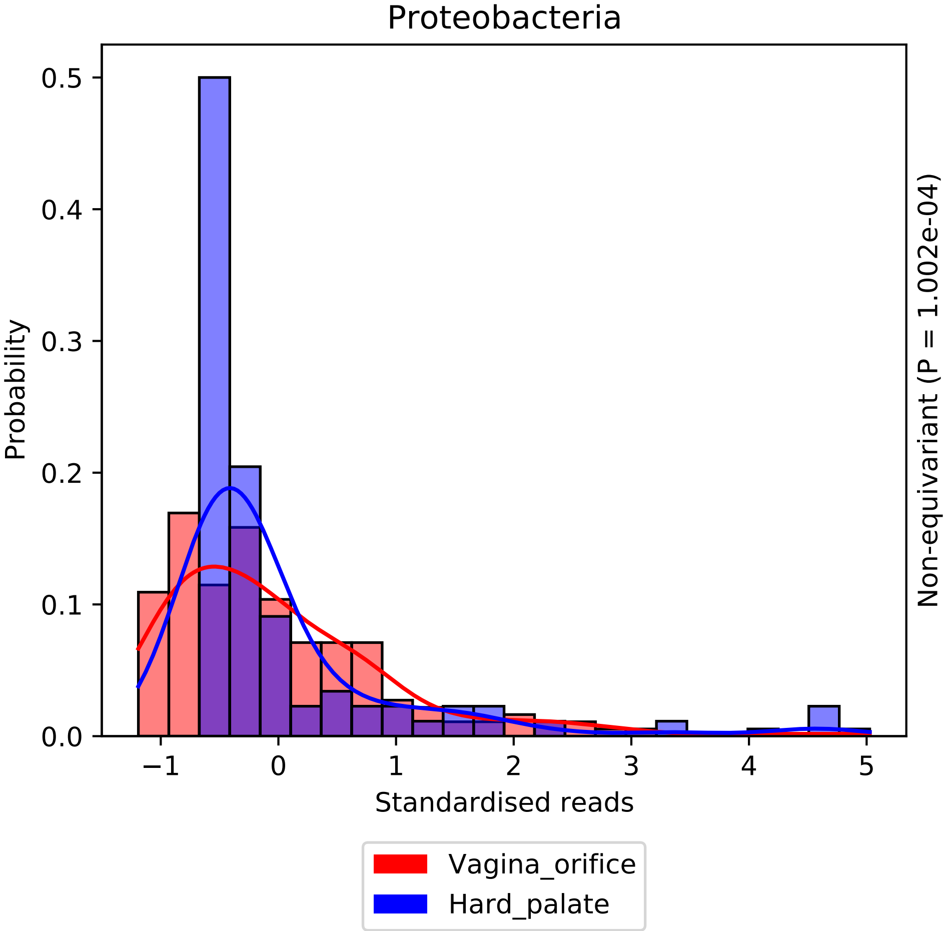

Proteobacteria
Probability
0.0
0.1
0.2
0.3
0.4
0.5
−1
0
1
2
3
4
5
Standardised reads
Non-equivariant (P = 1.002e-04)
Vagina_orifice
Hard_palate

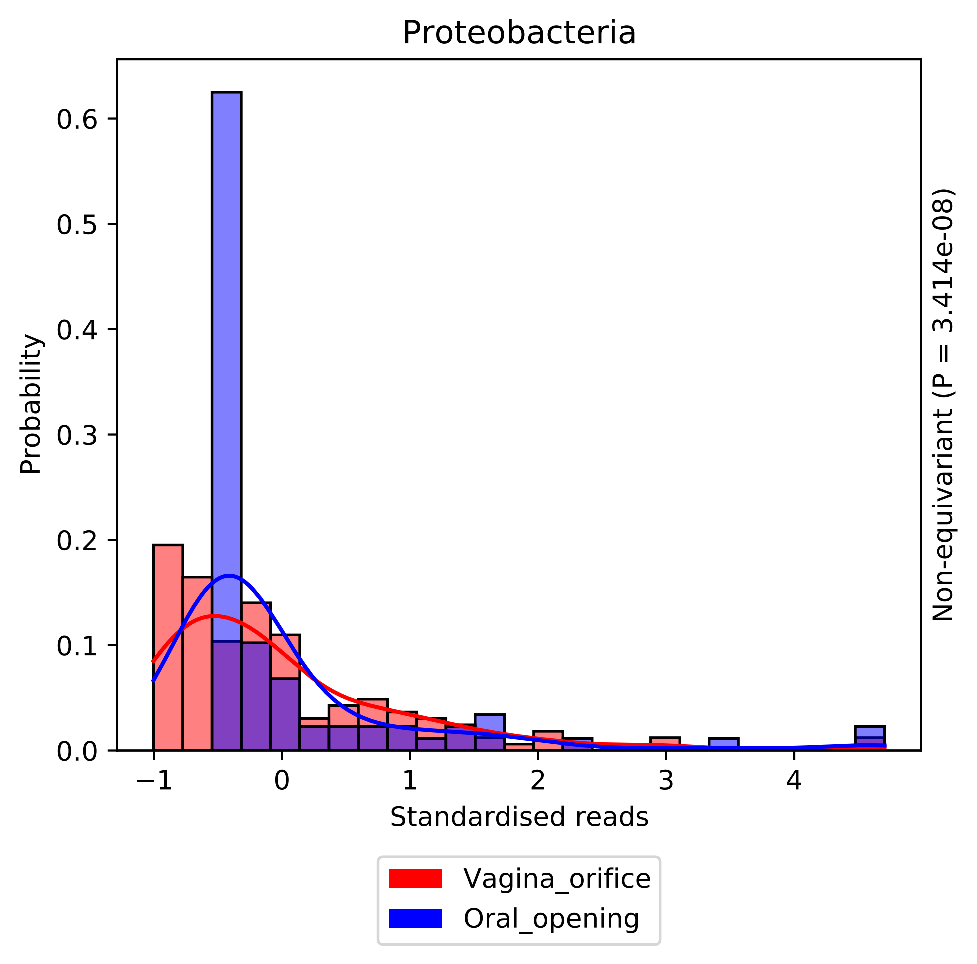

Proteobacteria
Probability
0.0
0.1
0.2
0.3
0.4
0.5
0.6
−1
0
1
2
3
4
Standardised reads
Non-equivariant (P = 3.414e-08)
Vagina_orifice
Oral_opening

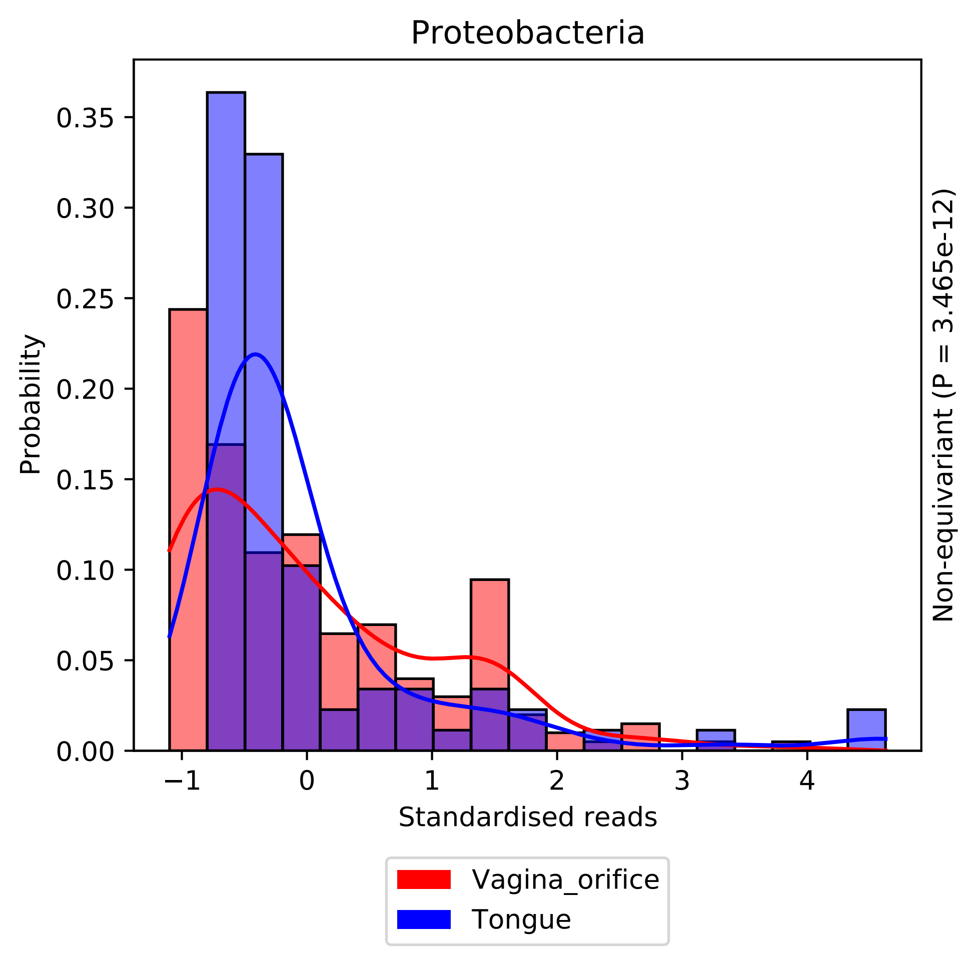
Proteobacteria
Probability
0.00
0.05
0.10
0.15
0.20
0.25
0.30
0.35
−1
0
1
2
3
4
Standardised reads
Non-equivariant (P = 3.465e-12)
Vagina_orifice
Tongue

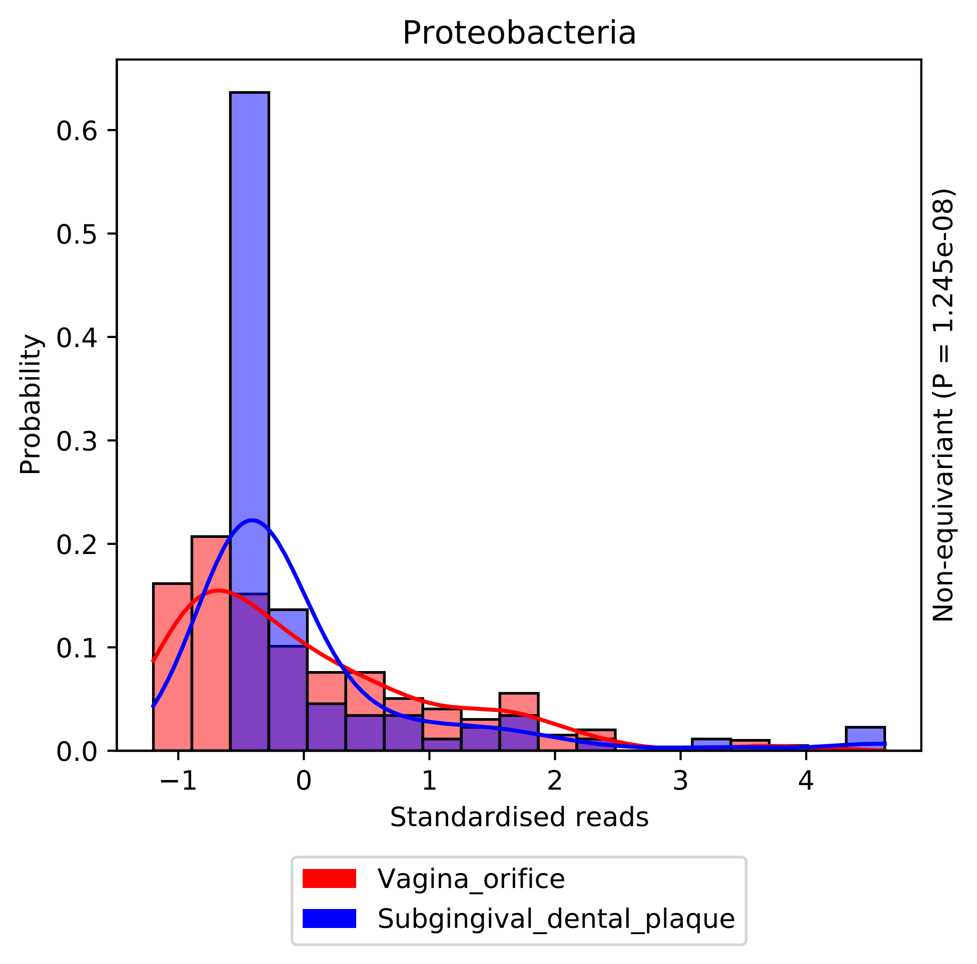

Proteobacteria
Probability
0.0
0.1
0.2
0.3
0.4
0.5
0.6
−1
0
1
2
3
4
Standardised reads
Non-equivariant (P = 1.245e-08)
Vagina_orifice
Subgingival_dental_plaque

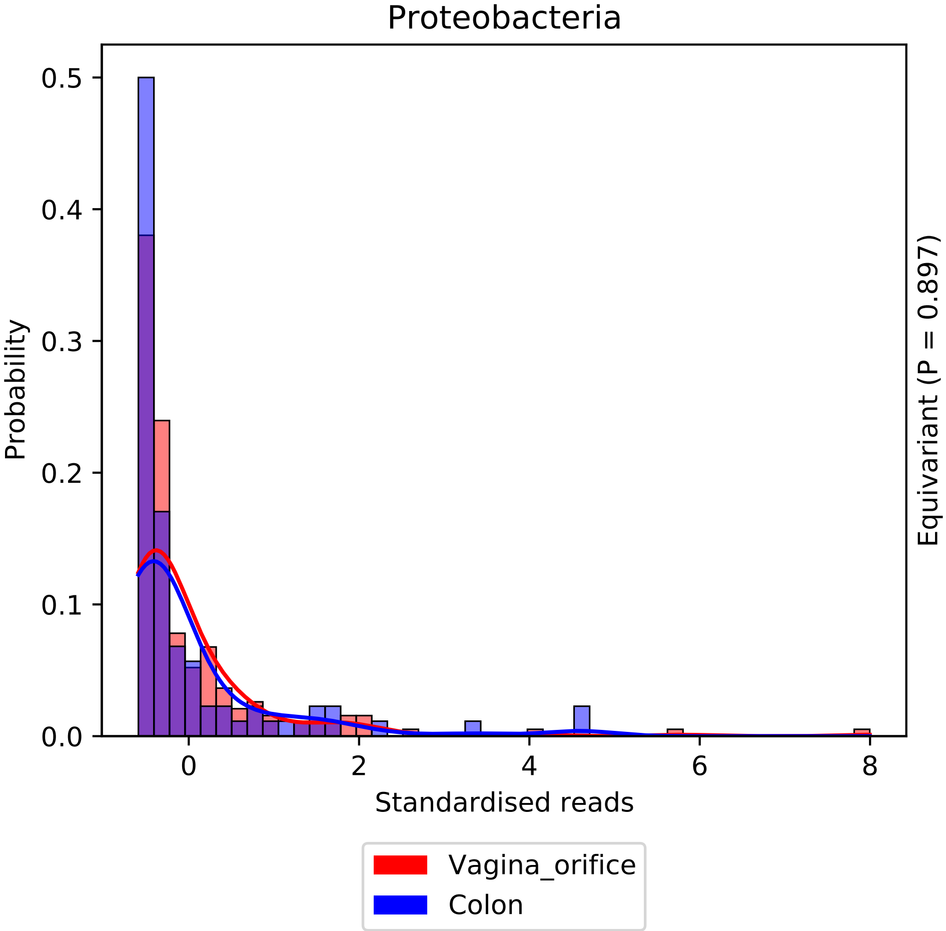
Proteobacteria
Probability
0.5
0.4
0.3
0.2
0.1
0.0
0
2
4
6
8
Standardised reads
Equivariant (P = 0.897)
Vagina_orifice
Colon

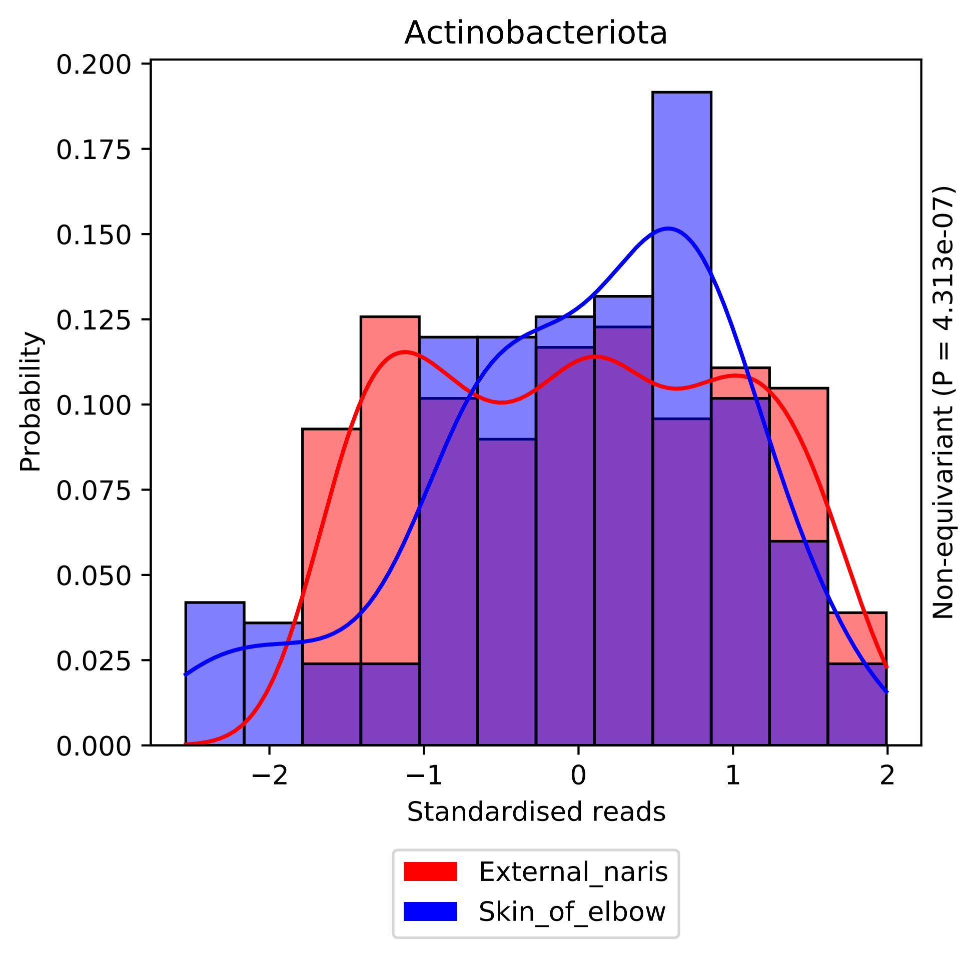

Actinobacteriota
Probability
0.200
0.175
0.150
0.125
0.100
0.075
0.050
0.025
0.000
−2
−1
0
1
2
Standardised reads
Non-equivariant (P = 4.313e-07)
External_naris
Skin_of_elbow

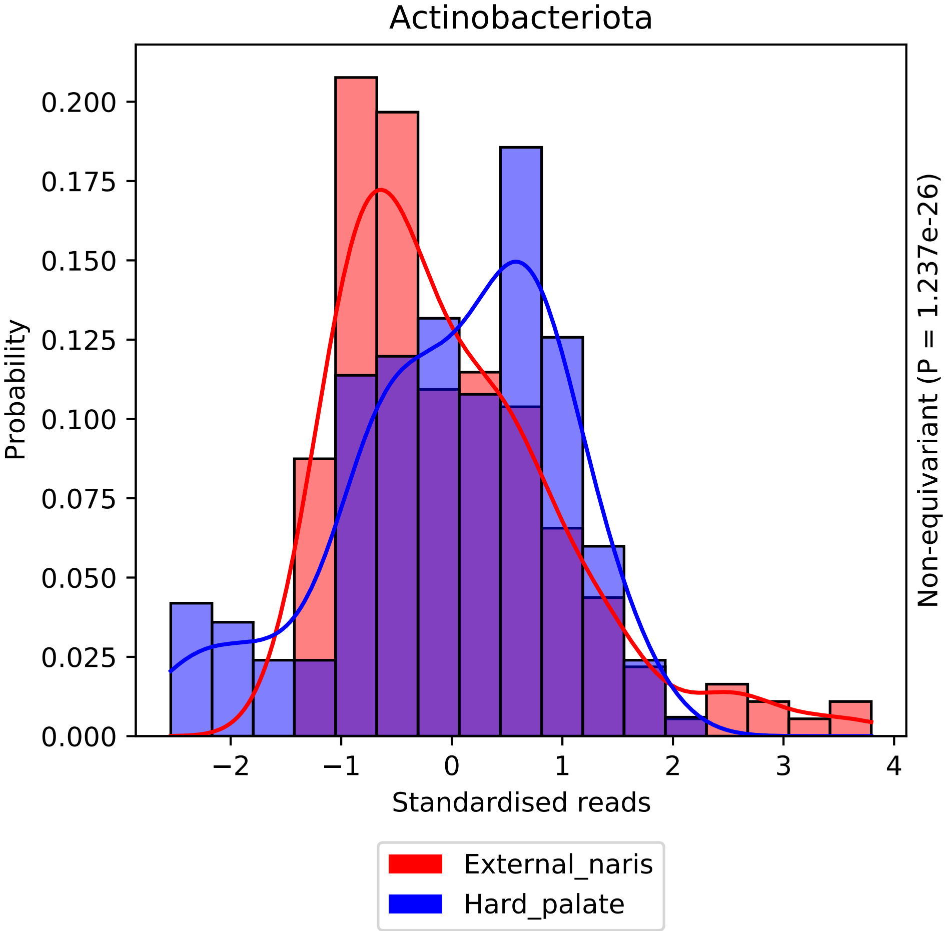

Actinobacteriota
Probability
0.000
0.025
0.050
0.075
0.100
0.125
0.150
0.175
0.200
−2
−1
0
1
2
3
4
Standardised reads
Non-equivariant (P = 1.237e-26)
External_naris
Hard_palate

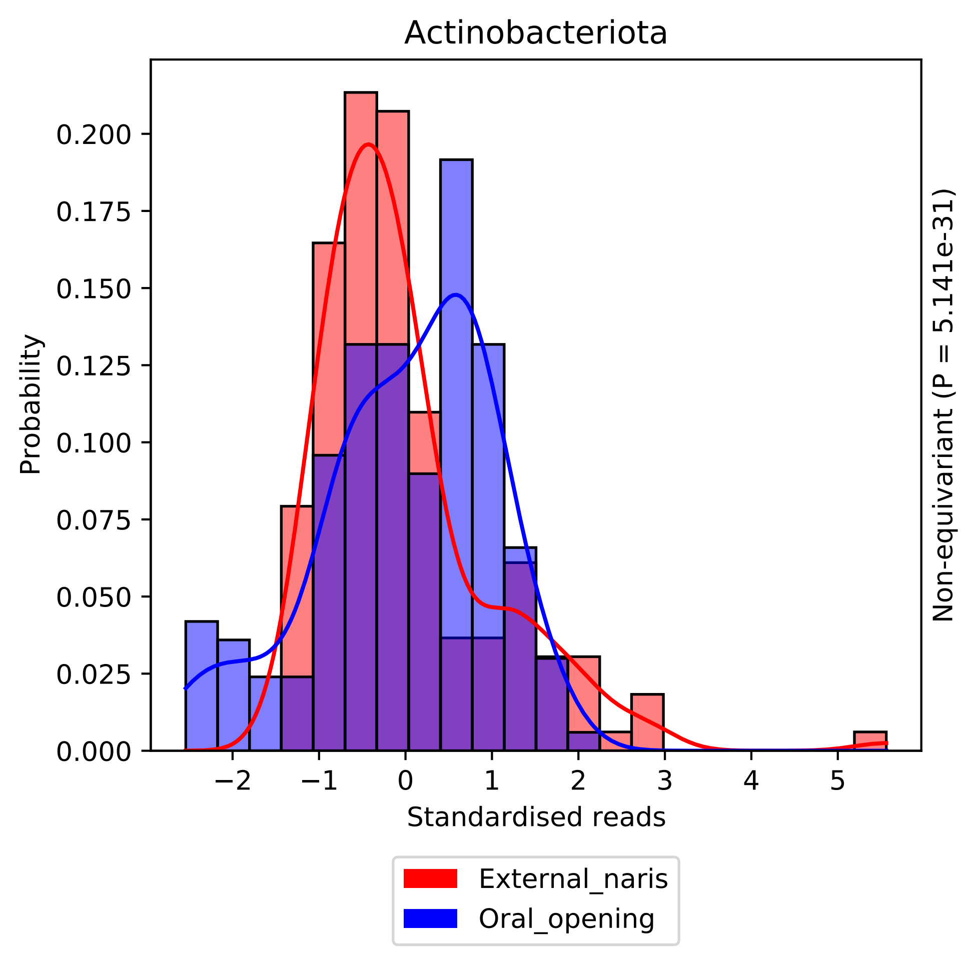

Actinobacteriota
Probability
0.000
0.025
0.050
0.075
0.100
0.125
0.150
0.175
0.200
−2
−1
0
1
2
3
4
5
Standardised reads
Non-equivariant (P = 5.141e-31)
External_naris
Oral_opening

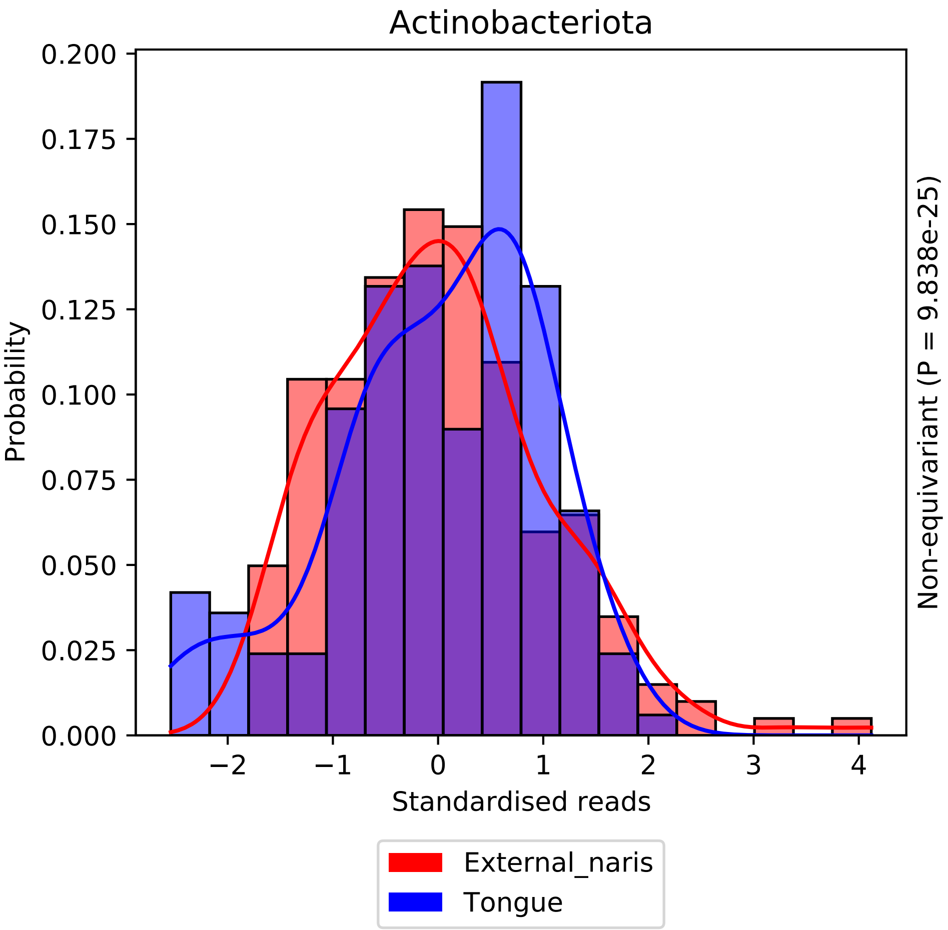

Actinobacteriota
0.200
0.175
0.150
0.125
0.100
0.075
0.050
0.025
0.000
Probability
−2
−1
0
1
2
3
4
Standardised reads
Non-equivariant (P = 9.838e-25)
External_naris
Tongue

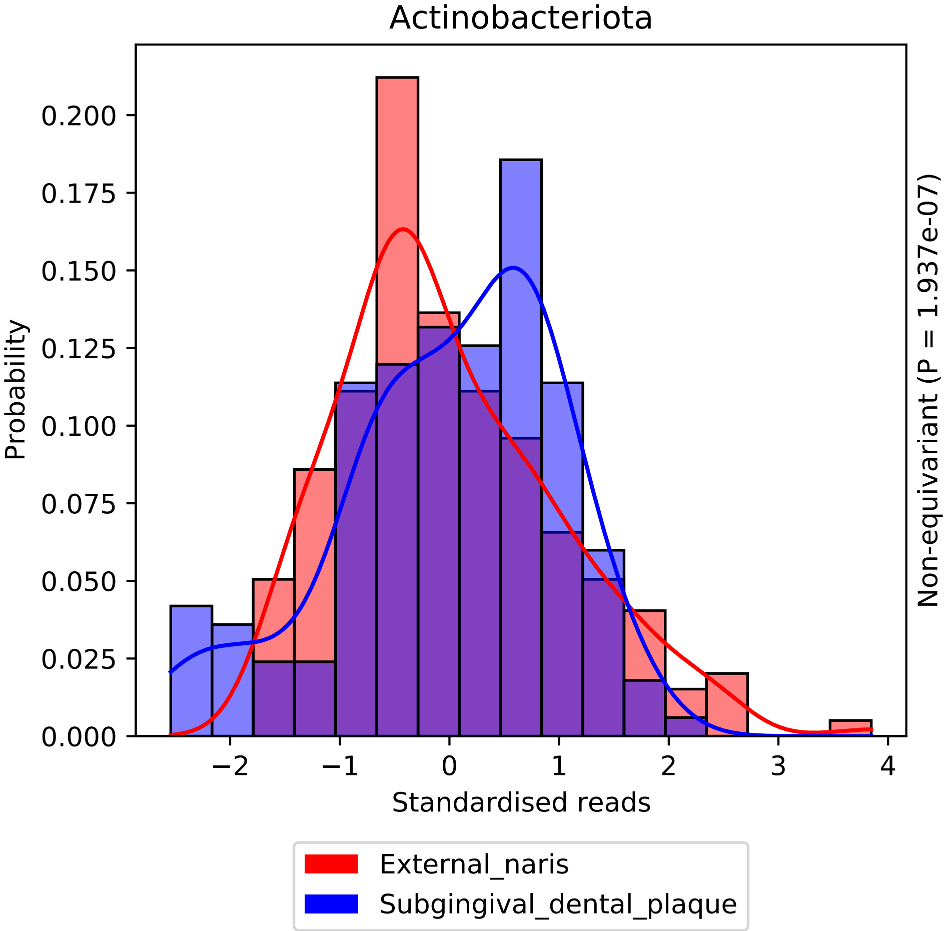
Actinobacteriota
Probability
0.000
0.025
0.050
0.075
0.100
0.125
0.150
0.175
0.200
−2
−1
0
1
2
3
4
Standardised reads
Non-equivariant (P = 1.937e-07)
External_naris
Subgingival_dental_plaque

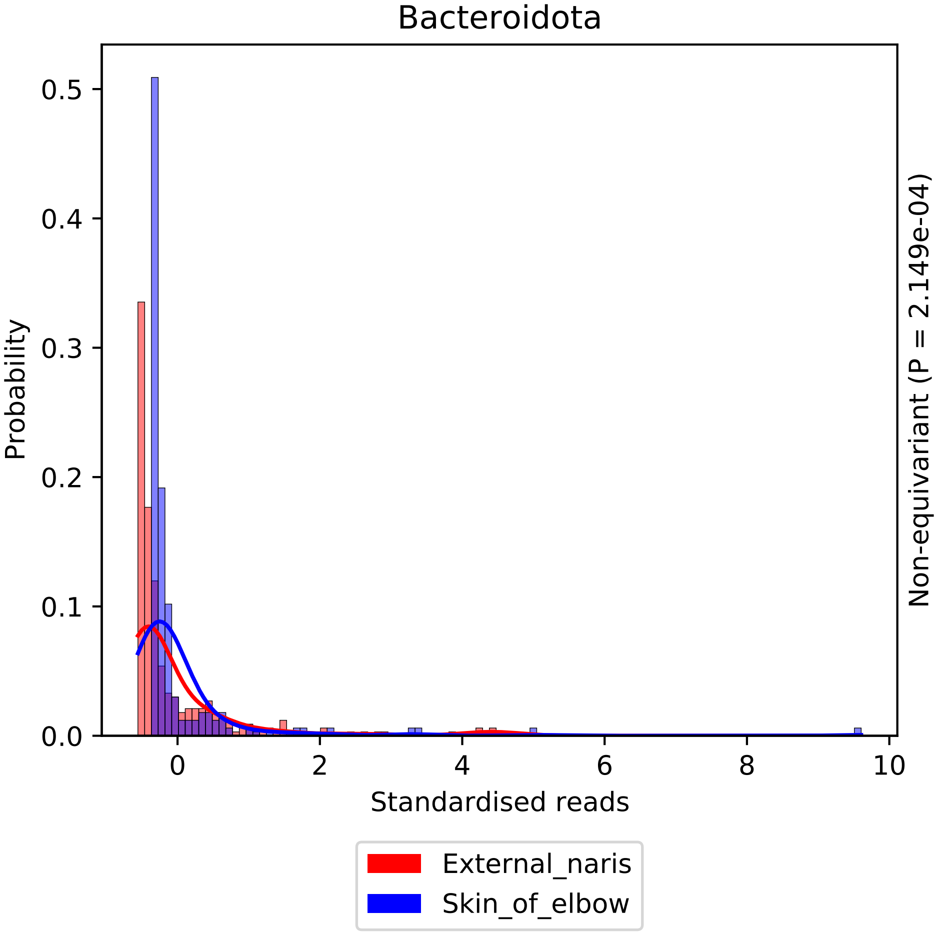

Bacteroidota
Probability
0.0
0.1
0.2
0.3
0.4
0.5
0
2
4
6
8
10
Standardised reads
Non-equivariant (P = 2.149e-04)
External_naris
Skin_of_elbow

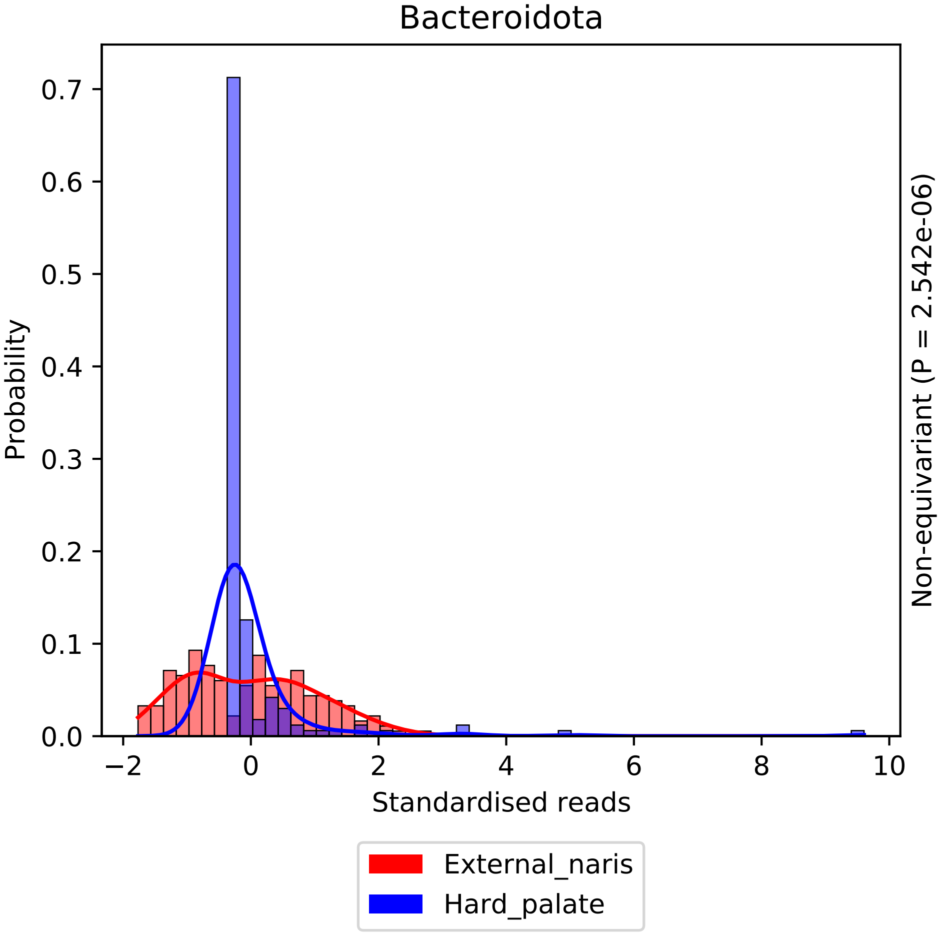

Bacteroidota
Probability
0.0
0.1
0.2
0.3
0.4
0.5
0.6
0.7
−2
0
2
4
6
8
10
Standardised reads
Non-equivariant (P = 2.542e-06)
External_naris
Hard_palate

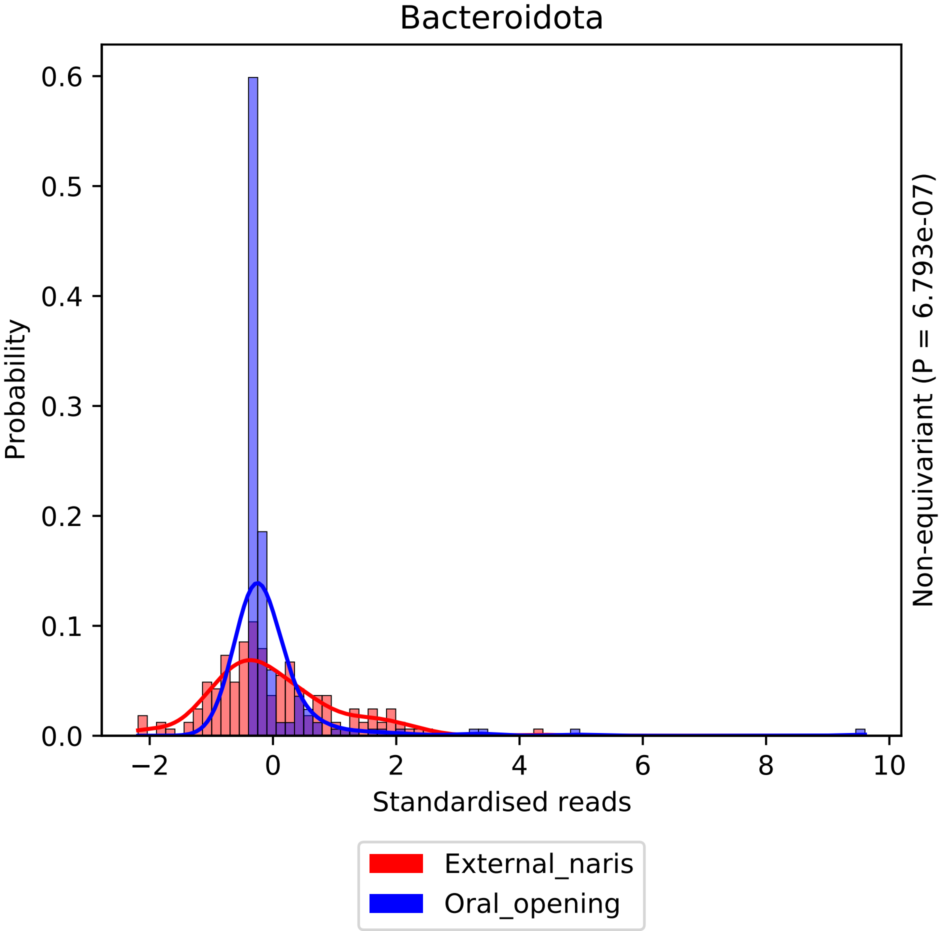

Bacteroidota
Probability
0.0
0.1
0.2
0.3
0.4
0.5
0.6
−2
0
2
4
6
8
10
Standardised reads
Non-equivariant (P = 6.793e-07)
External_naris
Oral_opening

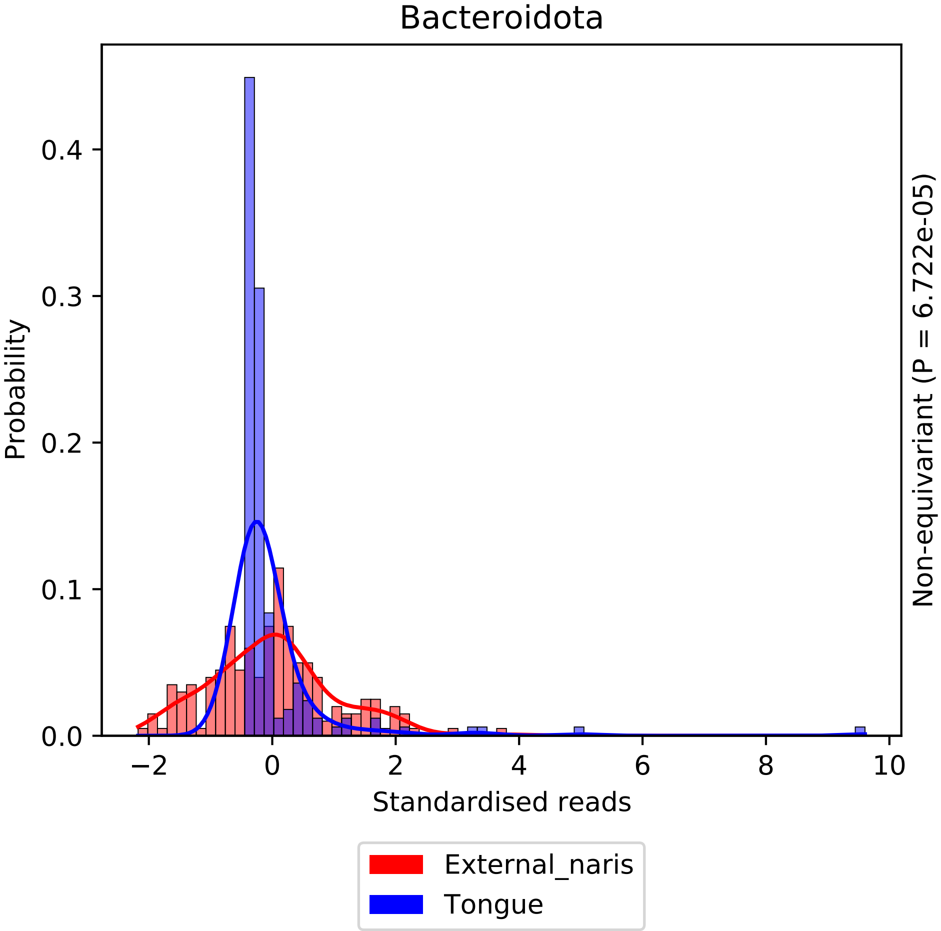

Bacteroidota
Probability
Standardised reads
Non-equivariant (P = 6.722e-05)
0.0
0.1
0.2
0.3
0.4
−2
0
2
4
6
8
10
External_naris
Tongue

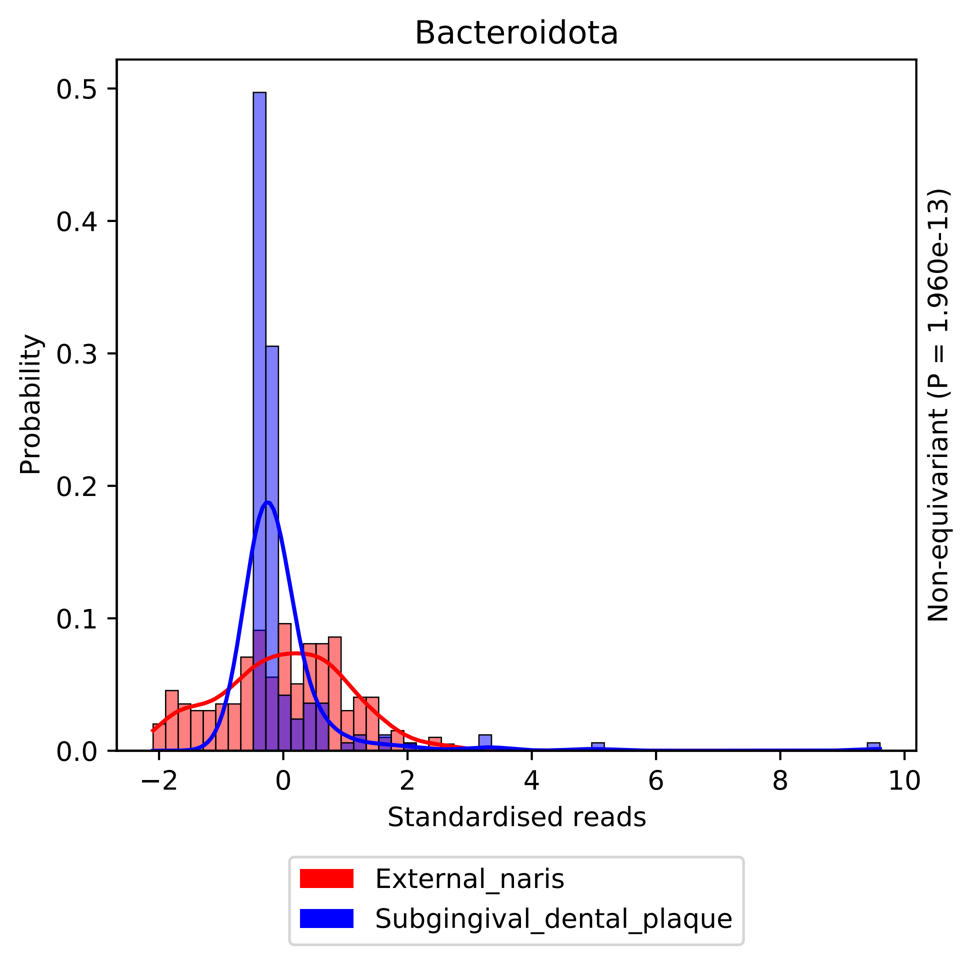

Bacteroidota
Probability
Standardised reads
Non-equivariant (P = 1.960e-13)
0.0
0.1
0.2
0.3
0.4
0.5
−2
0
2
4
6
8
10
External_naris
Subgingival_dental_plaque

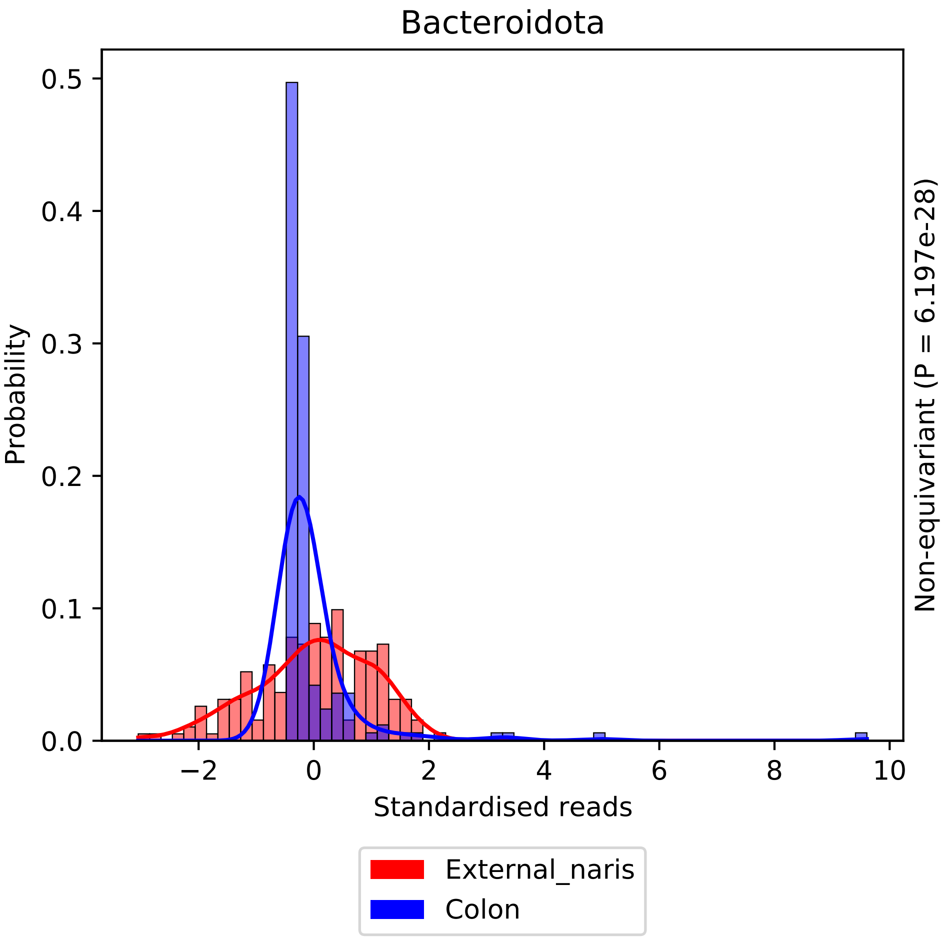

Bacteroidota
Probability
0.0
0.1
0.2
0.3
0.4
0.5
−2
0
2
4
6
8
10
Standardised reads
Non-equivariant (P = 6.197e-28)
External_naris
Colon

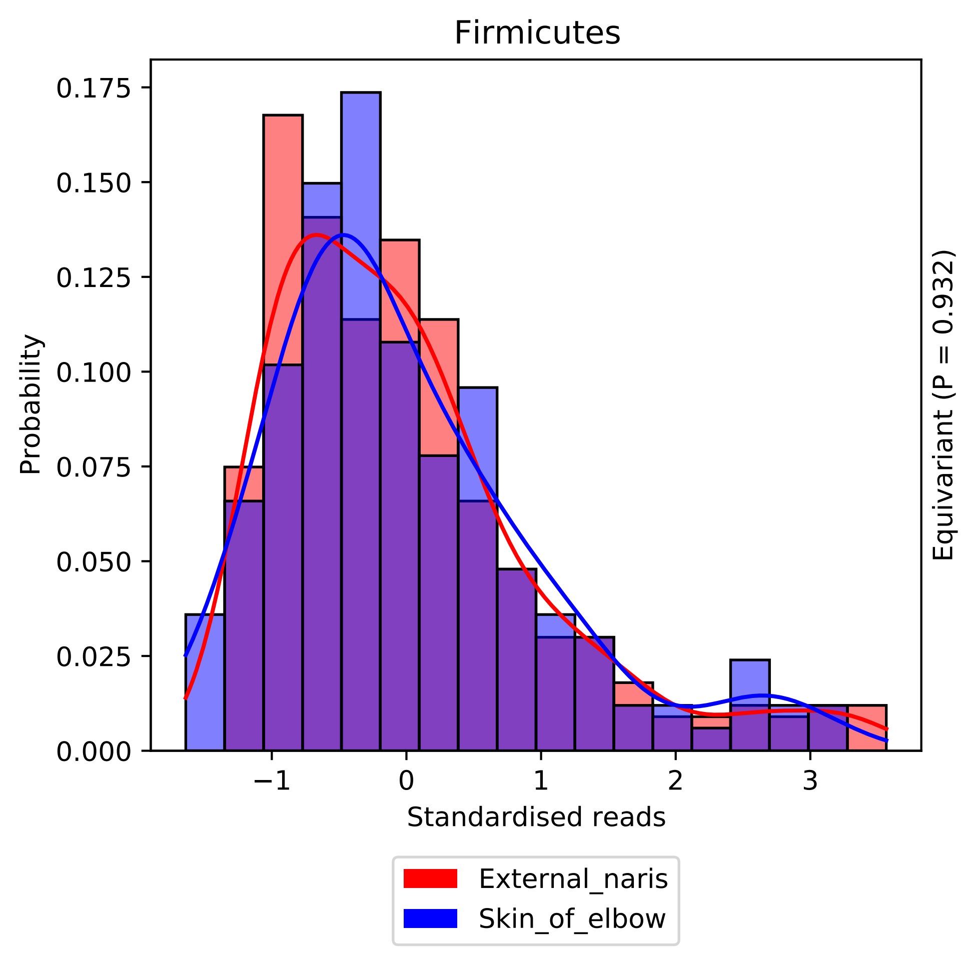
Firmicutes
Probability
0.000
0.025
0.050
0.075
0.100
0.125
0.150
0.175
−1
0
1
2
3
Standardised reads
Equivariant (P = 0.932)
External_naris
Skin_of_elbow

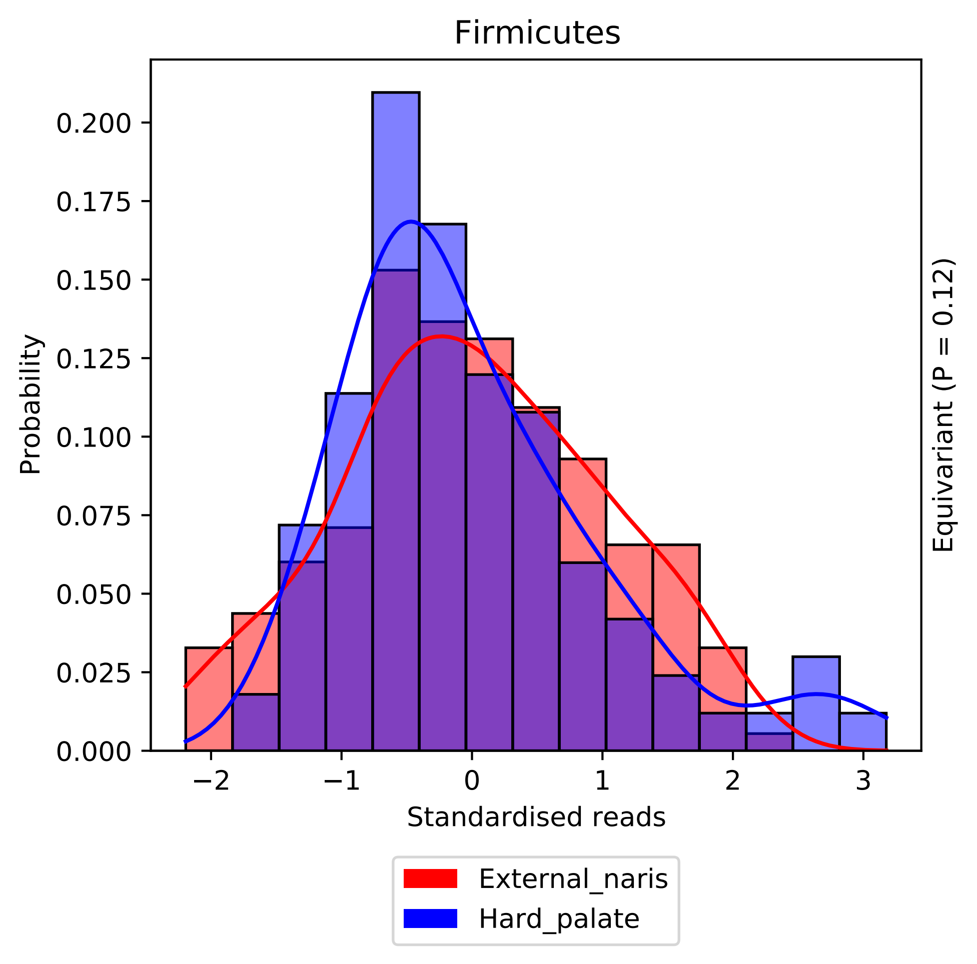

Firmicutes
Probability
0.000
0.025
0.050
0.075
0.100
0.125
0.150
0.175
0.200
−2
−1
0
1
2
3
Standardised reads
Equivariant (P = 0.12)
External_naris
Hard_palate

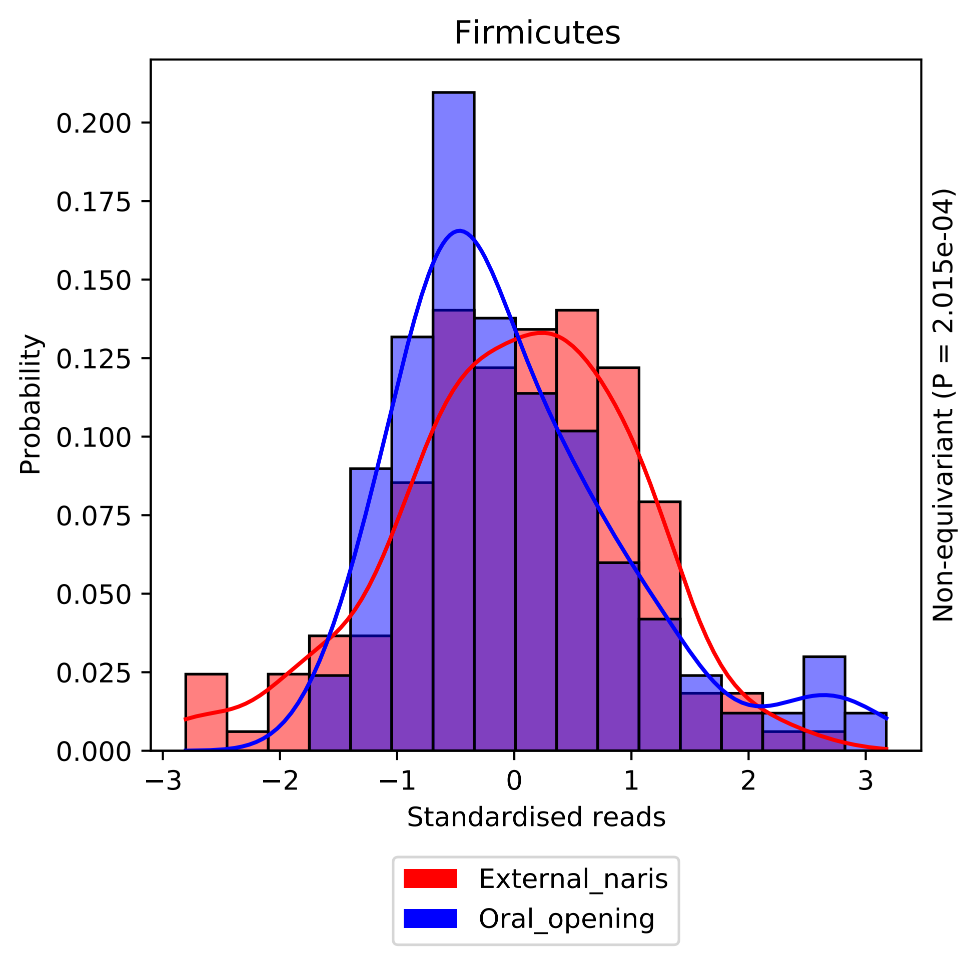
Firmicutes
Probability
0.200
0.175
0.150
0.125
0.100
0.075
0.050
0.025
0.000
−3
−2
−1
0
1
2
3
Standardised reads
Non-equivariant (P = 2.015e-04)
External_naris
Oral_opening

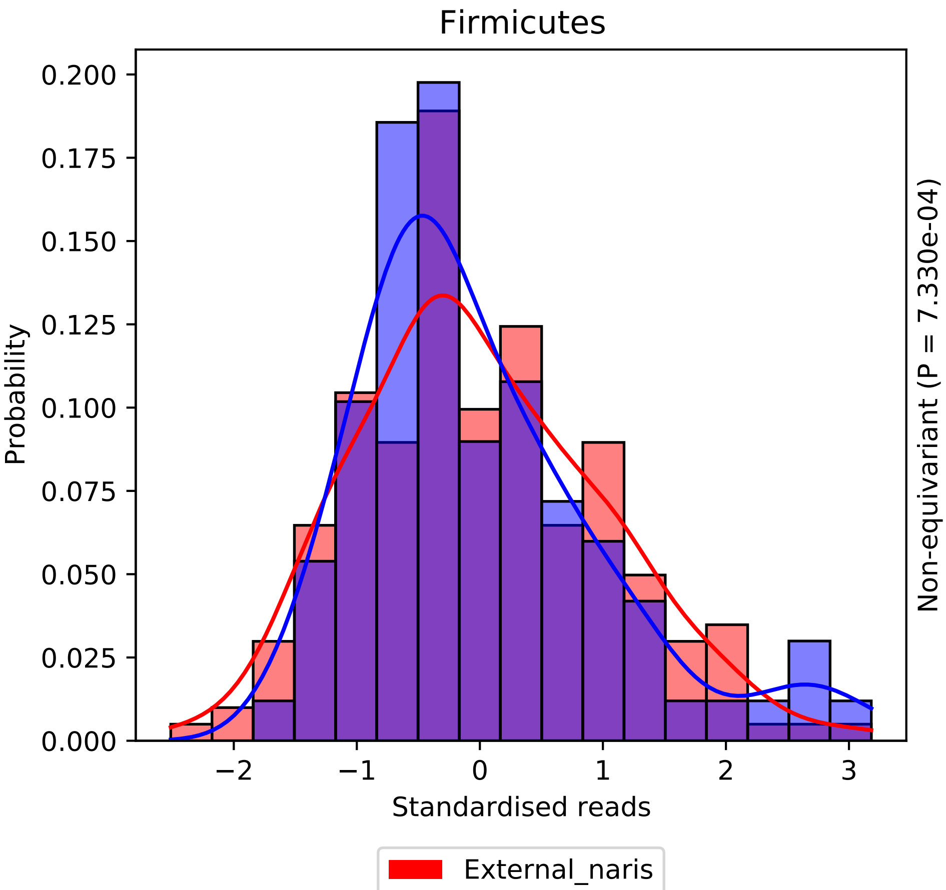

Firmicutes
Probability
0.000
0.025
0.050
0.075
0.100
0.125
0.150
0.175
0.200
−2
−1
0
1
2
3
Standardised reads
Non-equivariant (P = 7.330e-04)
External_naris
Tongue

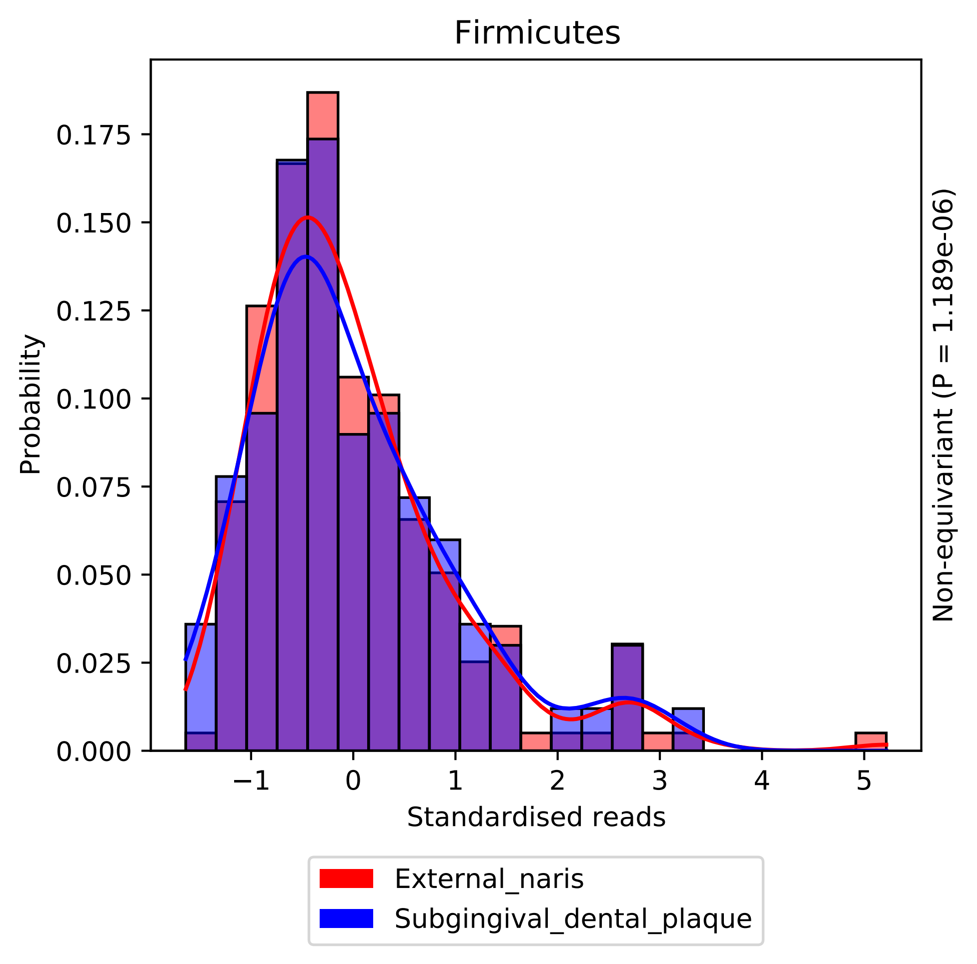
Firmicutes
Probability
Standardised reads
0.000
0.025
0.050
0.075
0.100
0.125
0.150
0.175
−1
0
1
2
3
4
5
Non-equivariant (P = 1.189e-06)
External_naris
Subgingival_dental_plaque

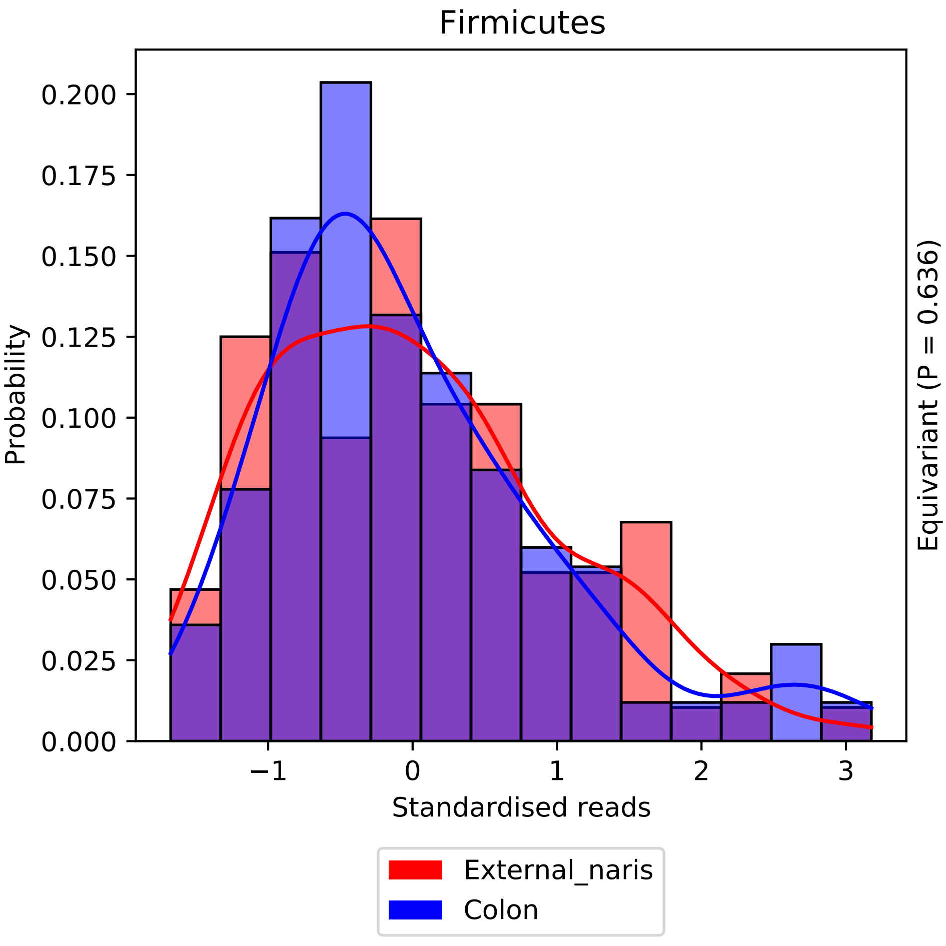
Firmicutes
Probability
Standardised reads
Equivariant (P = 0.636)
0.000
0.025
0.050
0.075
0.100
0.125
0.150
0.175
0.200
−1
0
1
2
3
External_naris
Colon

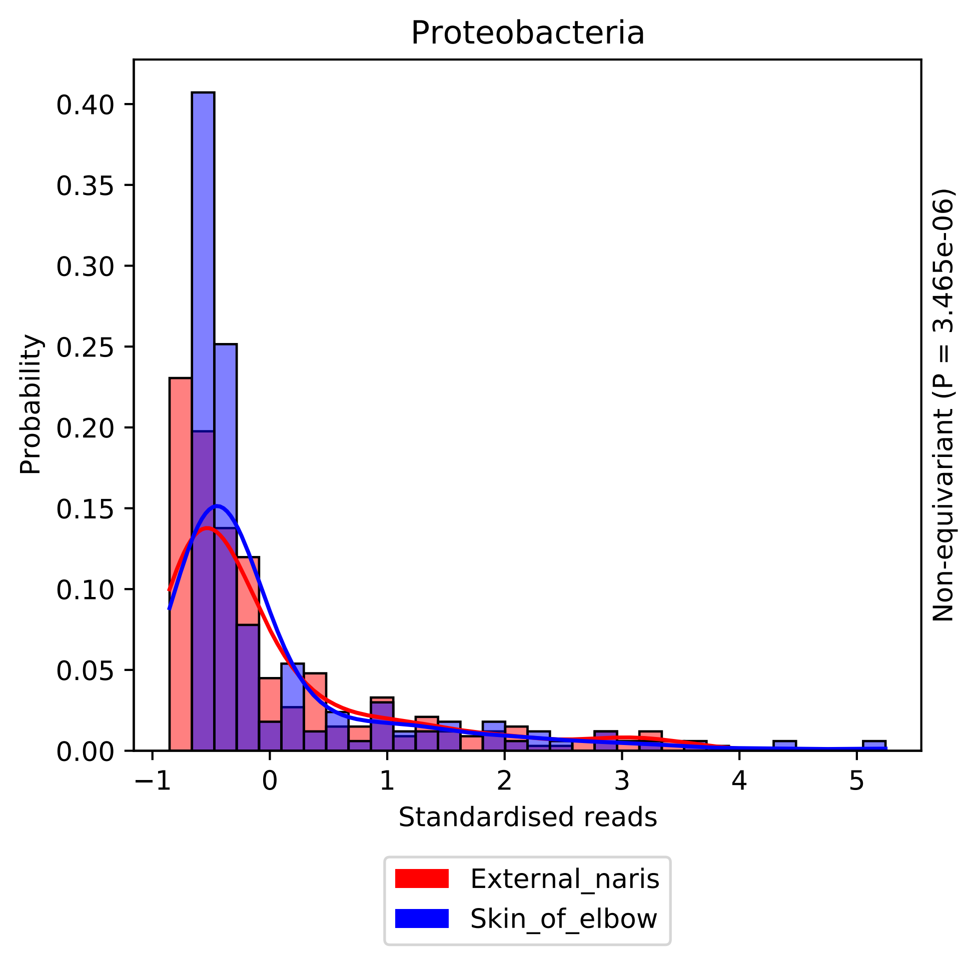

Proteobacteria
Probability
0.00
0.05
0.10
0.15
0.20
0.25
0.30
0.35
0.40
−1
0
1
2
3
4
5
Standardised reads
Non-equivariant (P = 3.465e-06)
External_naris
Skin_of_elbow

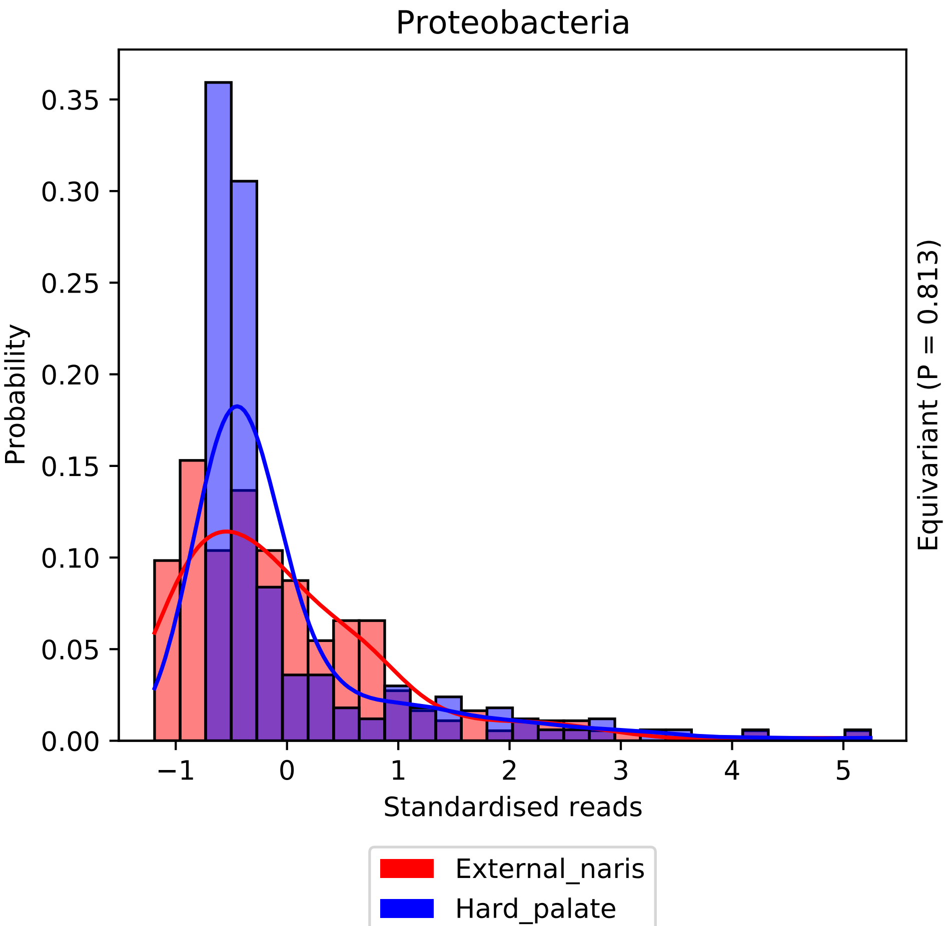
Proteobacteria
Probability
0.35
0.30
0.25
0.20
0.15
0.10
0.05
0.00
−1
0
1
2
3
4
5
Standardised reads
Equivariant (P = 0.813)
External_naris
Hard_palate

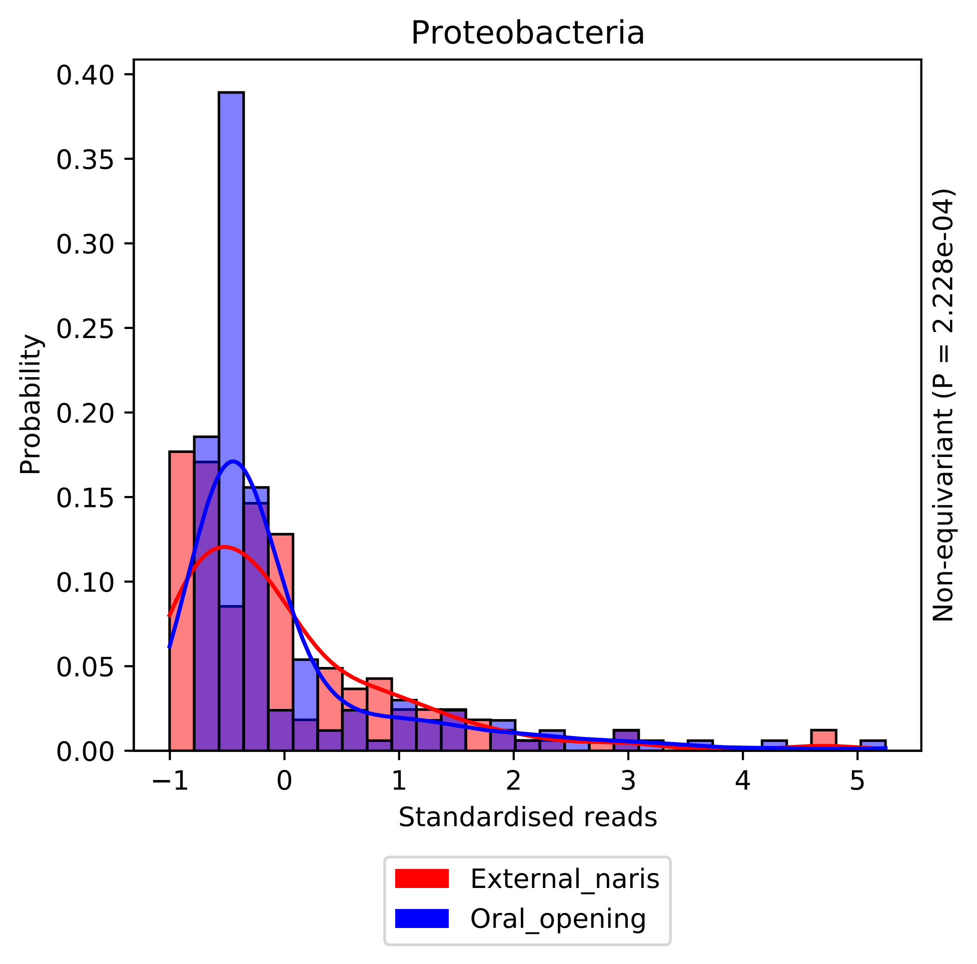
Proteobacteria
Probability
0.40
0.35
0.30
0.25
0.20
0.15
0.10
0.05
0.00
−1
0
1
2
3
4
5
Standardised reads
Non-equivariant (P = 2.228e-04)
External_naris
Oral_opening

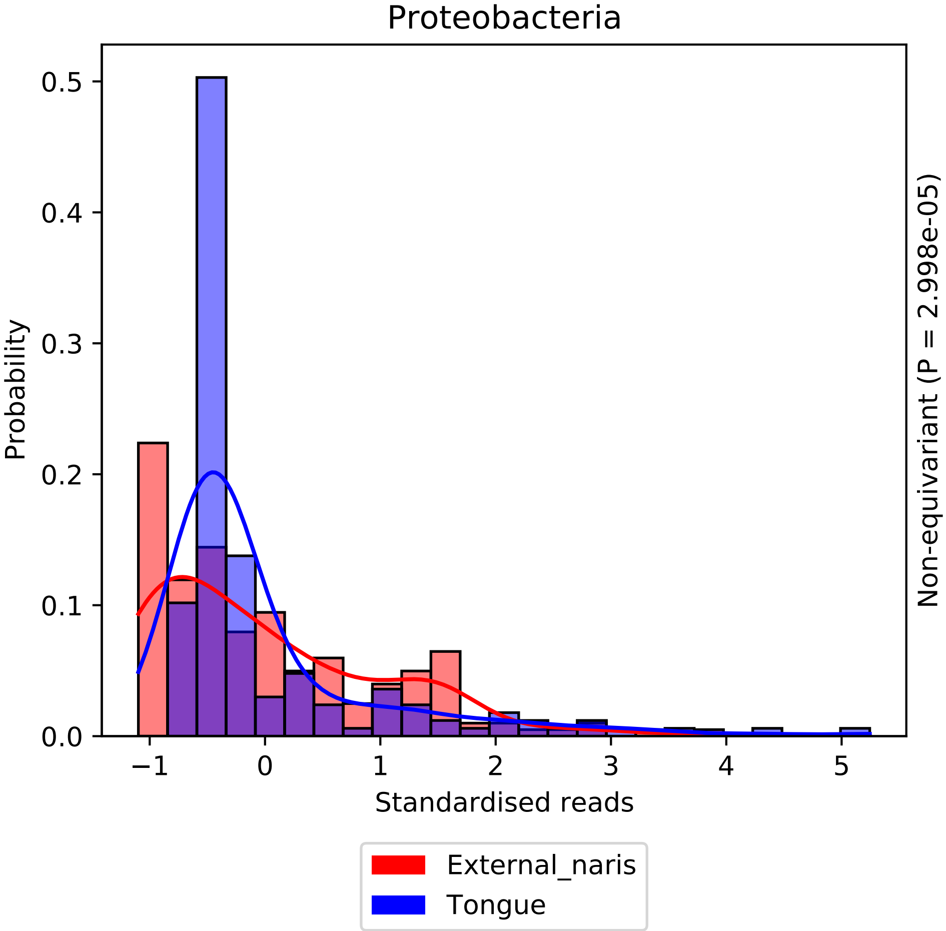
Proteobacteria
Probability
Standardised reads
Non-equivariant (P = 2.998e-05)
External_naris
Tongue
0.0
0.1
0.2
0.3
0.4
0.5
−1
0
1
2
3
4
5

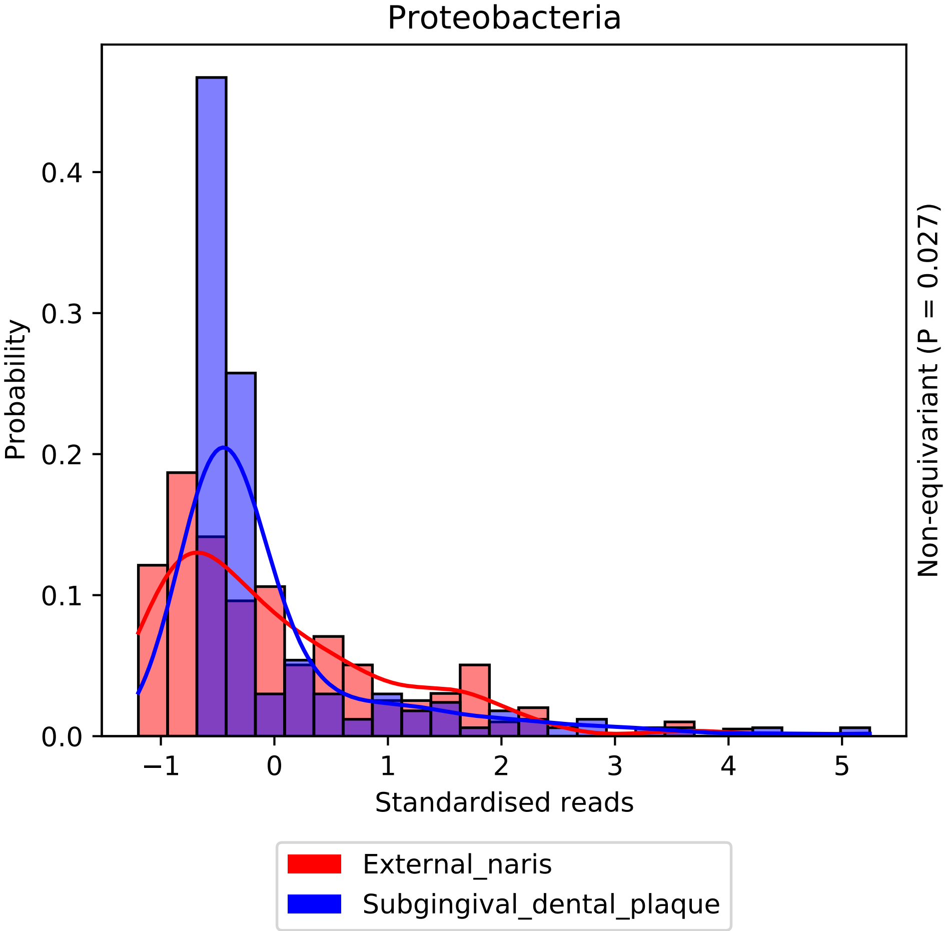
Proteobacteria
Probability
0.0
0.1
0.2
0.3
0.4
−1
0
1
2
3
4
5
Standardised reads
Non-equivariant (P = 0.027)
External_naris
Subgingival_dental_plaque

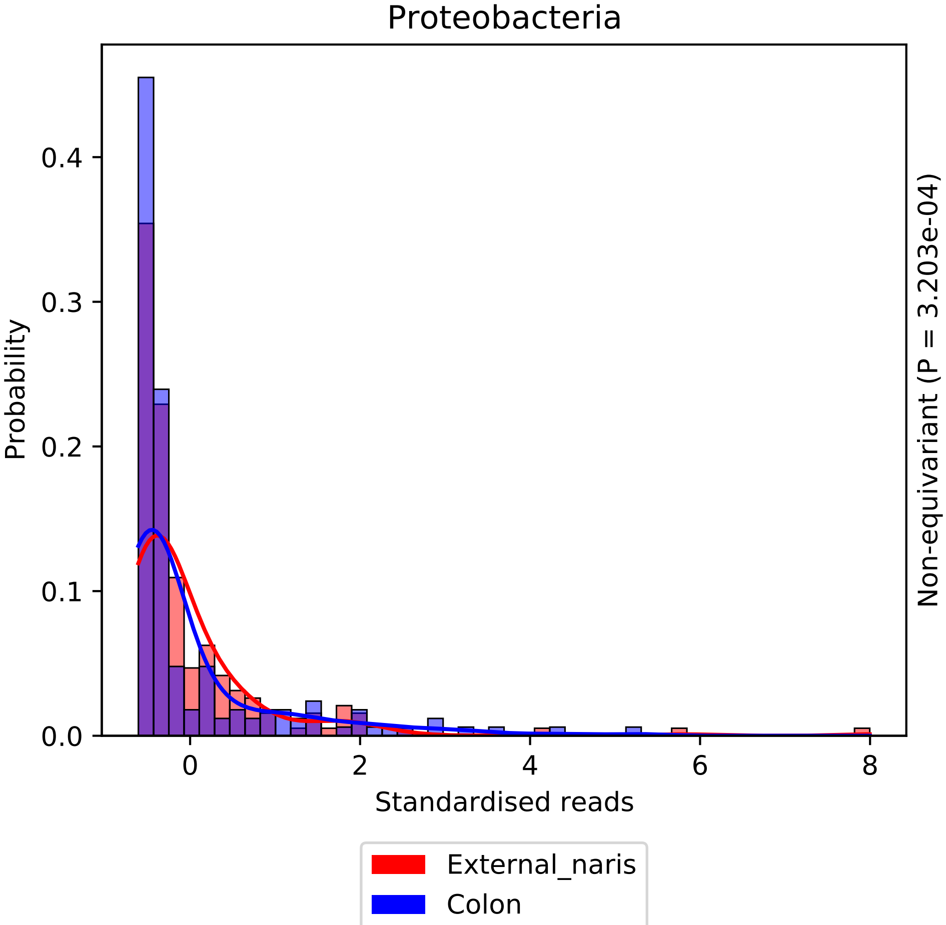

Proteobacteria
Probability
0.0
0.1
0.2
0.3
0.4
0
2
4
6
8
Standardised reads
Non-equivariant (P = 3.203e-04)
External_naris
Colon

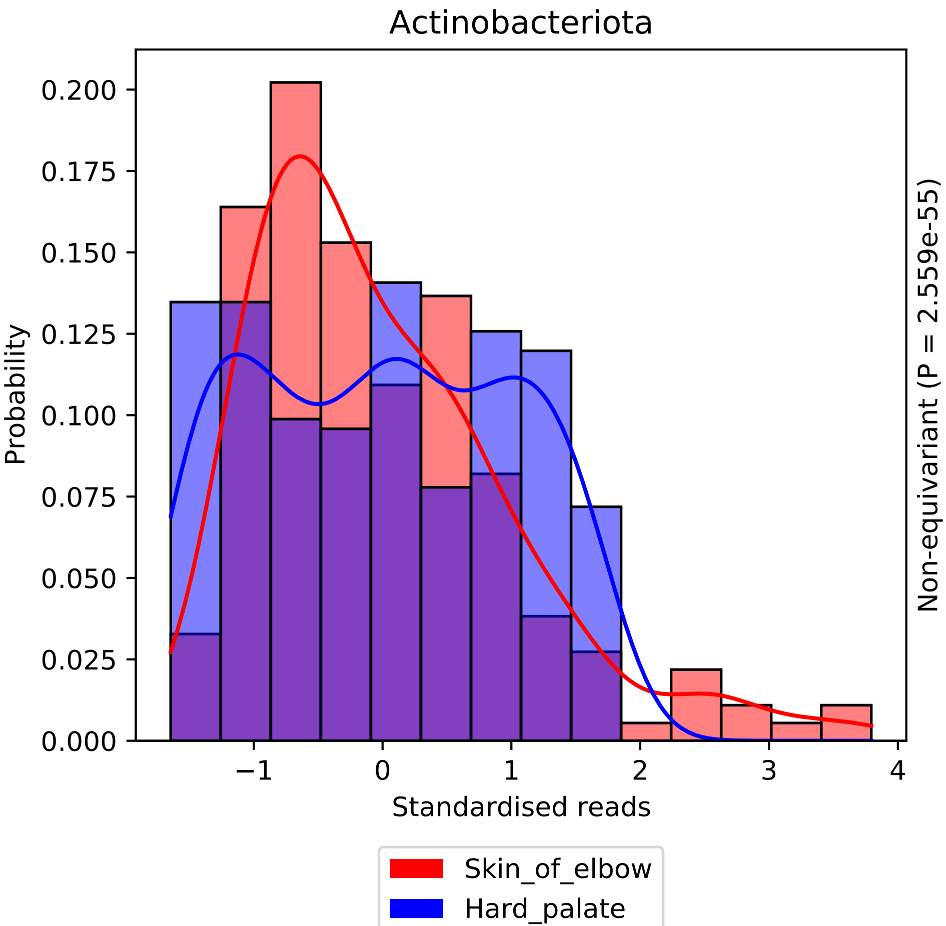

Actinobacteriota
Probability
0.200
0.175
0.150
0.125
0.100
0.075
0.050
0.025
0.000
−1
0
1
2
3
4
Standardised reads
Non-equivariant (P = 2.559e-55)
Skin_of_elbow
Hard_palate

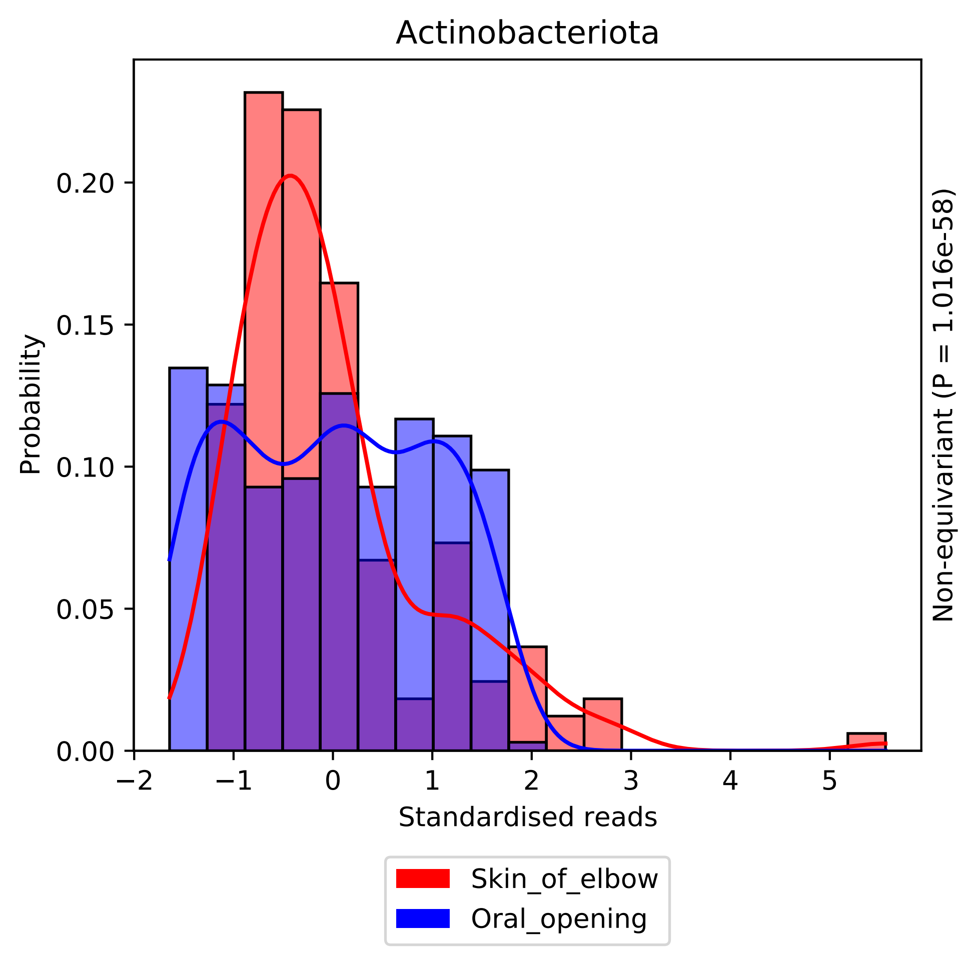
Actinobacteriota
Probability
0.00
0.05
0.10
0.15
0.20
−2
−1
0
1
2
3
4
5
Standardised reads
Non-equivariant (P = 1.016e-58)
Skin_of_elbow
Oral_opening

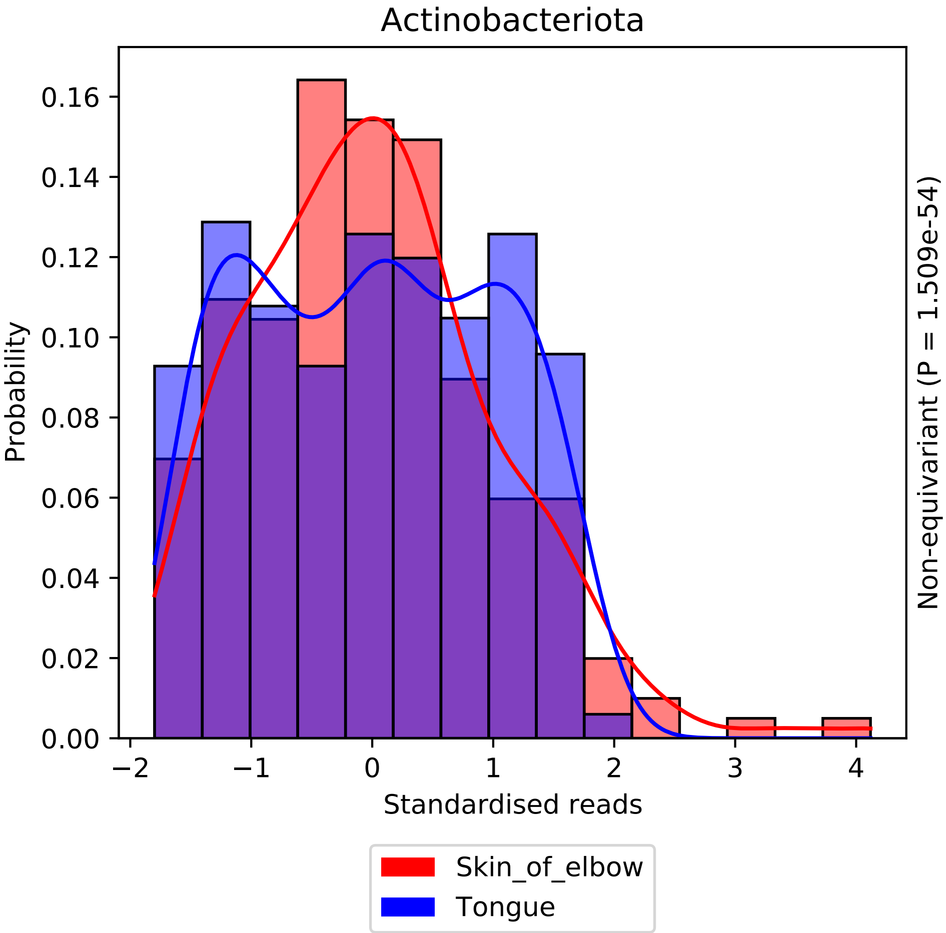

Actinobacteriota
0.16
0.14
0.12
0.10
0.08
0.06
0.04
0.02
0.00
Probability
−2
−1
0
1
2
3
4
Standardised reads
Non-equivariant (P = 1.509e-54)
Skin_of_elbow
Tongue

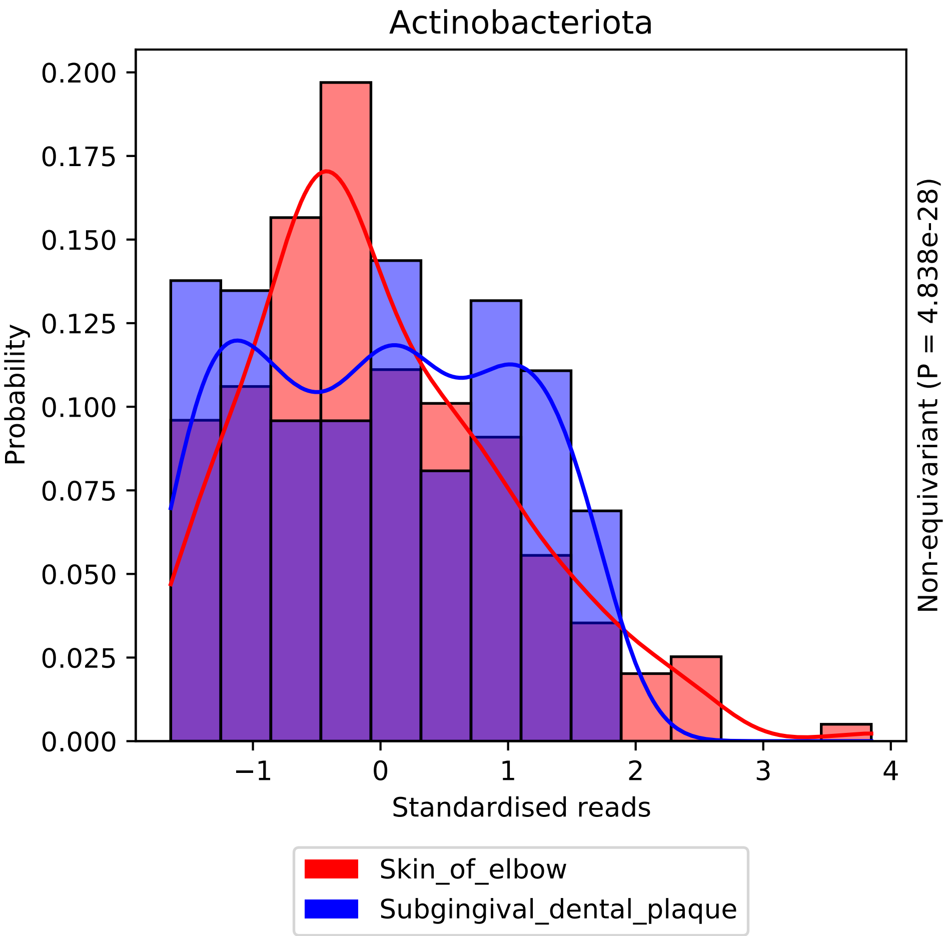

Actinobacteriota
Probability
0.200
0.175
0.150
0.125
0.100
0.075
0.050
0.025
0.000
−1
0
1
2
3
4
Standardised reads
Non-equivariant (P = 4.838e-28)
Skin_of_elbow
Subgingival_dental_plaque

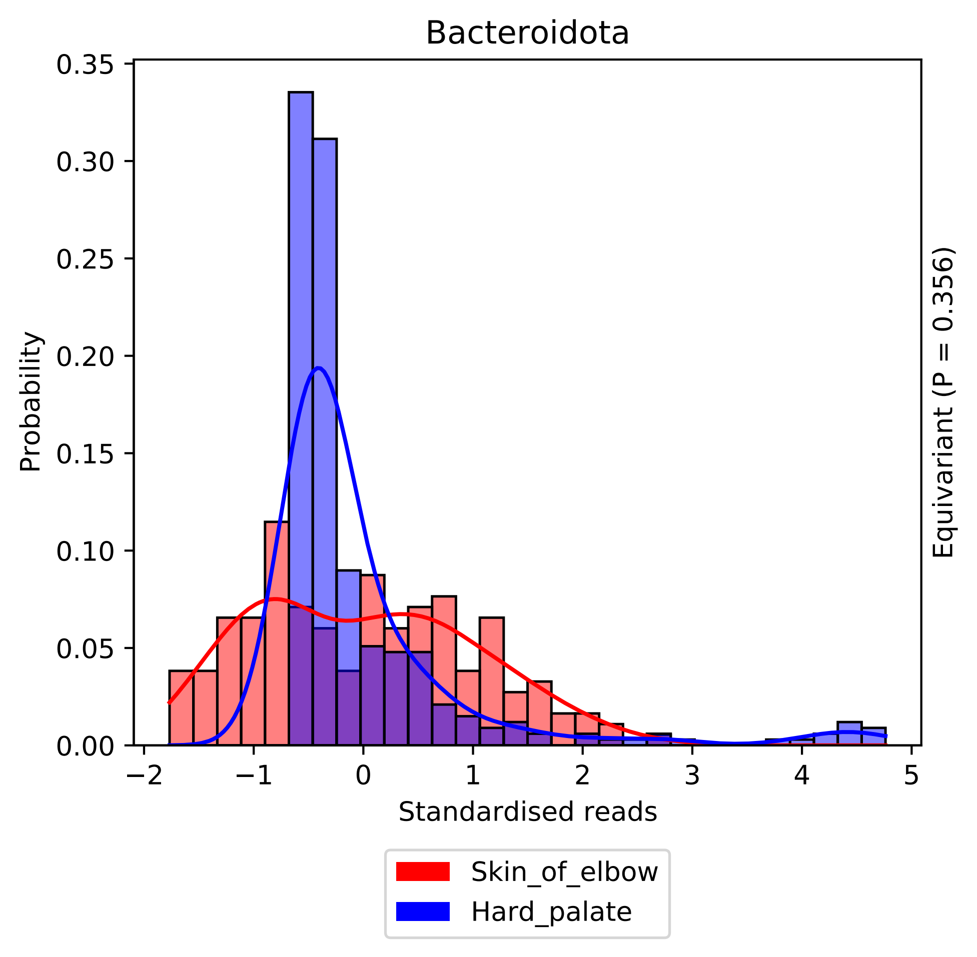

Bacteroidota
Probability
0.00
0.05
0.10
0.15
0.20
0.25
0.30
0.35
−2
−1
0
1
2
3
4
5
Standardised reads
Equivariant (P = 0.356)
Skin_of_elbow
Hard_palate

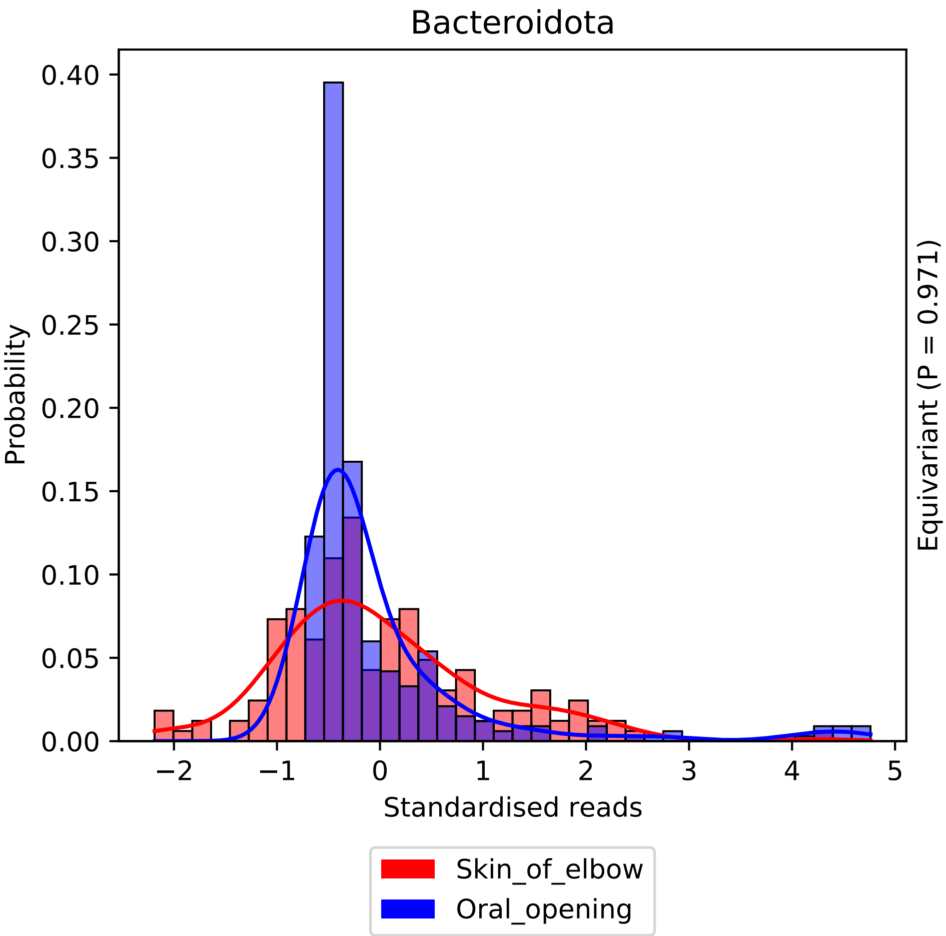

Bacteroidota
Probability
0.00
0.05
0.10
0.15
0.20
0.25
0.30
0.35
0.40
−2
−1
0
1
2
3
4
5
Standardised reads
Equivariant (P = 0.971)
Skin_of_elbow
Oral_opening

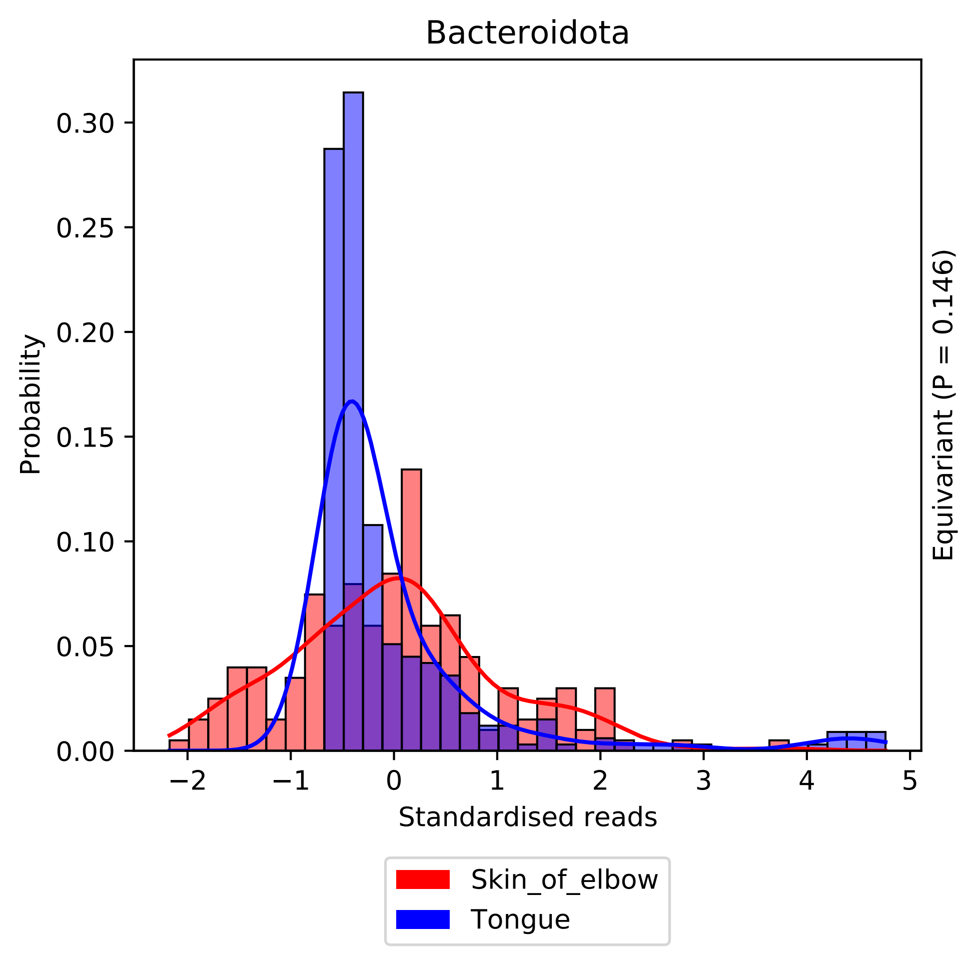

Bacteroidota
Probability
0.00
0.05
0.10
0.15
0.20
0.25
0.30
−2
−1
0
1
2
3
4
5
Standardised reads
Equivariant (P = 0.146)
Skin_of_elbow
Tongue

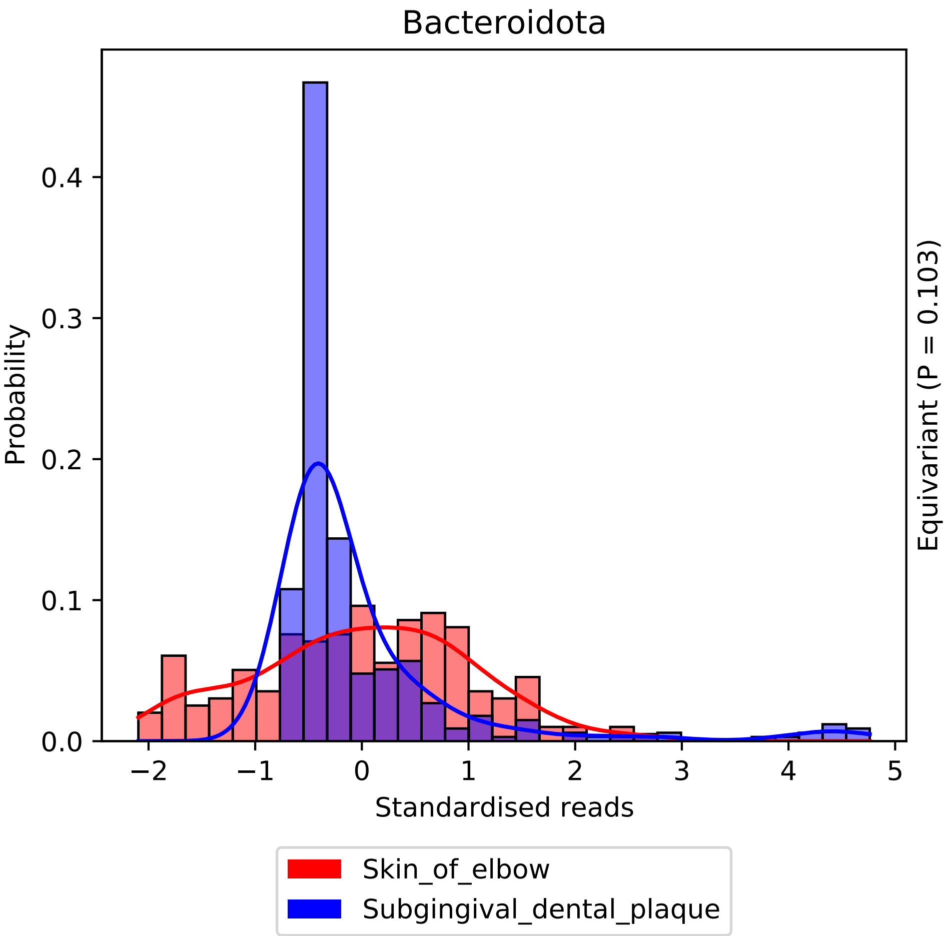

Bacteroidota
Probability
0.0
0.1
0.2
0.3
0.4
−2
−1
0
1
2
3
4
5
Standardised reads
Equivariant (P = 0.103)
Skin_of_elbow
Subgingival_dental_plaque

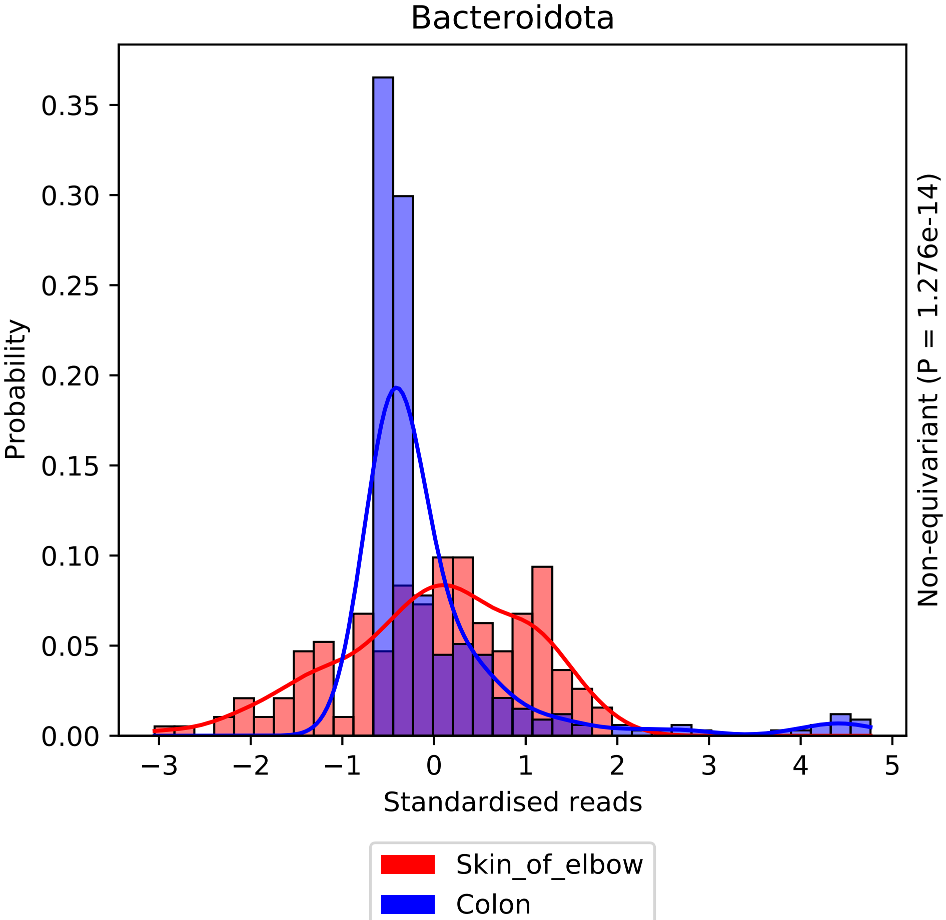

Bacteroidota
Probability
Standardised reads
Non-equivariant (P = 1.276e-14)
0.00
0.05
0.10
0.15
0.20
0.25
0.30
0.35
−3
−2
−1
0
1
2
3
4
5
Skin_of_elbow
Colon

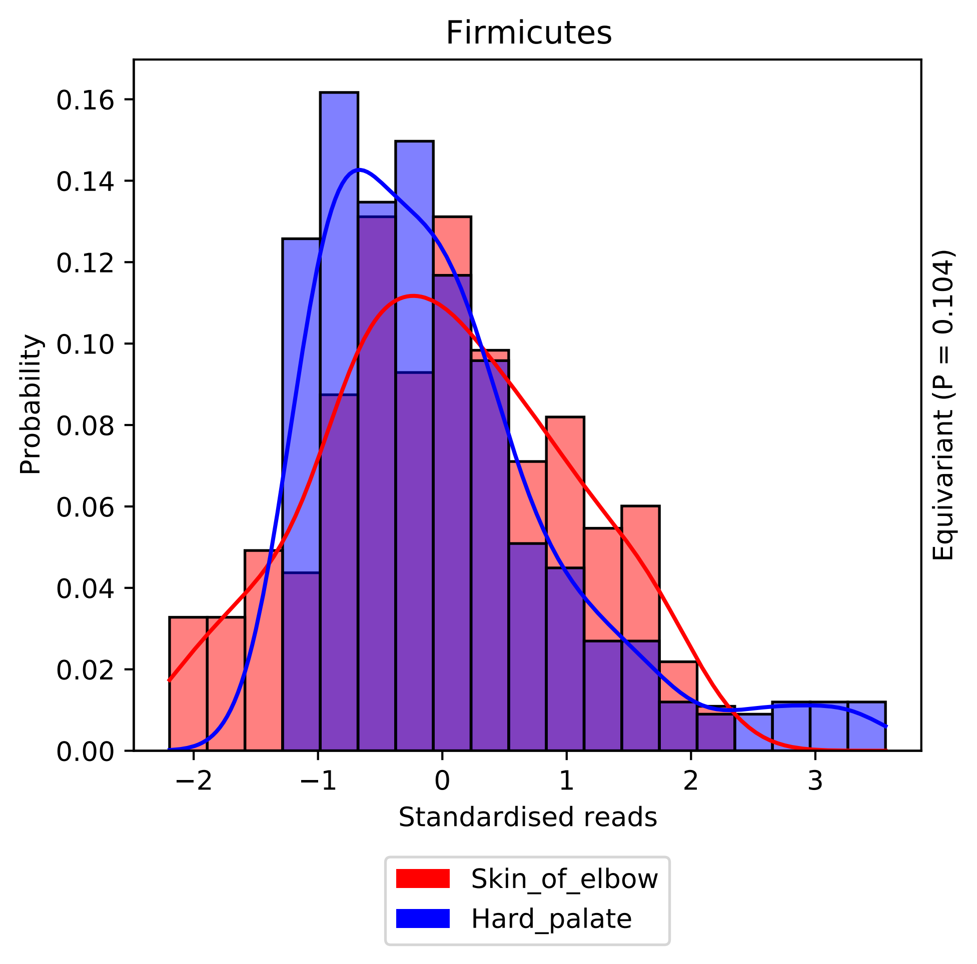

Firmicutes
Probability
0.16
0.14
0.12
0.10
0.08
0.06
0.04
0.02
0.00
−2
−1
0
1
2
3
Standardised reads
Equivariant (P = 0.104)
Skin_of_elbow
Hard_palate

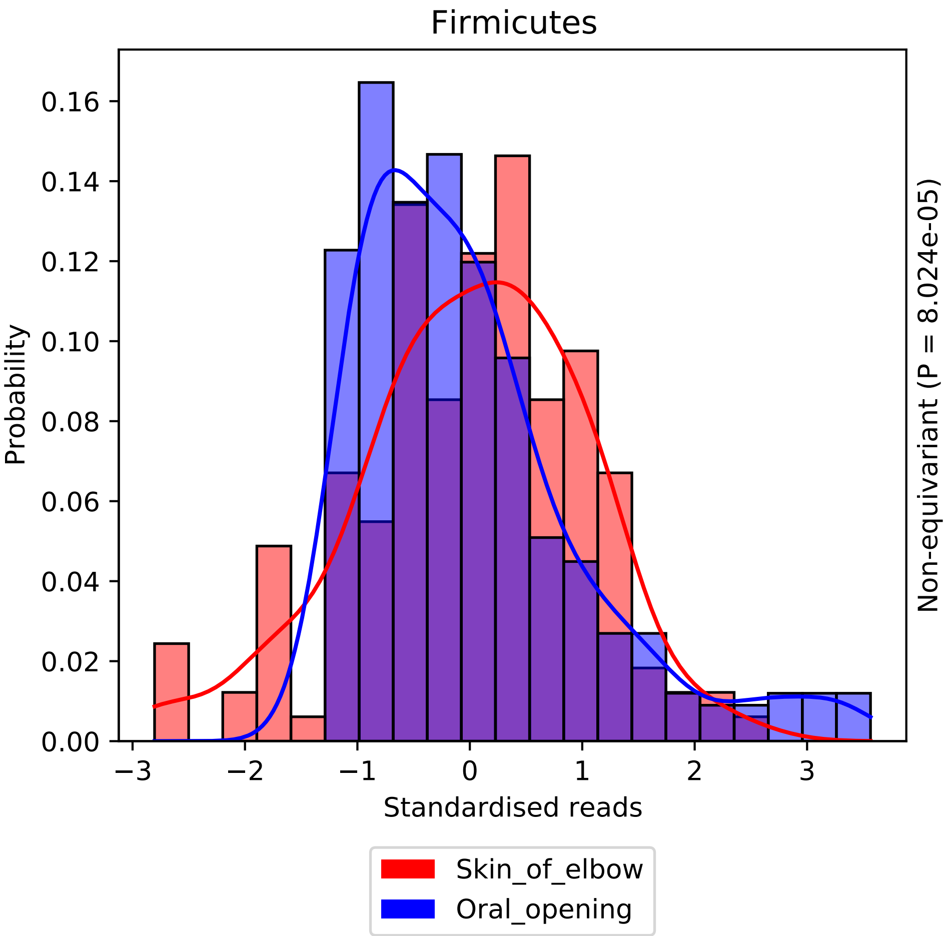

Firmicutes
Probability
0.00
0.02
0.04
0.06
0.08
0.10
0.12
0.14
0.16
−3
−2
−1
0
1
2
3
Standardised reads
Non-equivariant (P = 8.024e-05)
Skin_of_elbow
Oral_opening

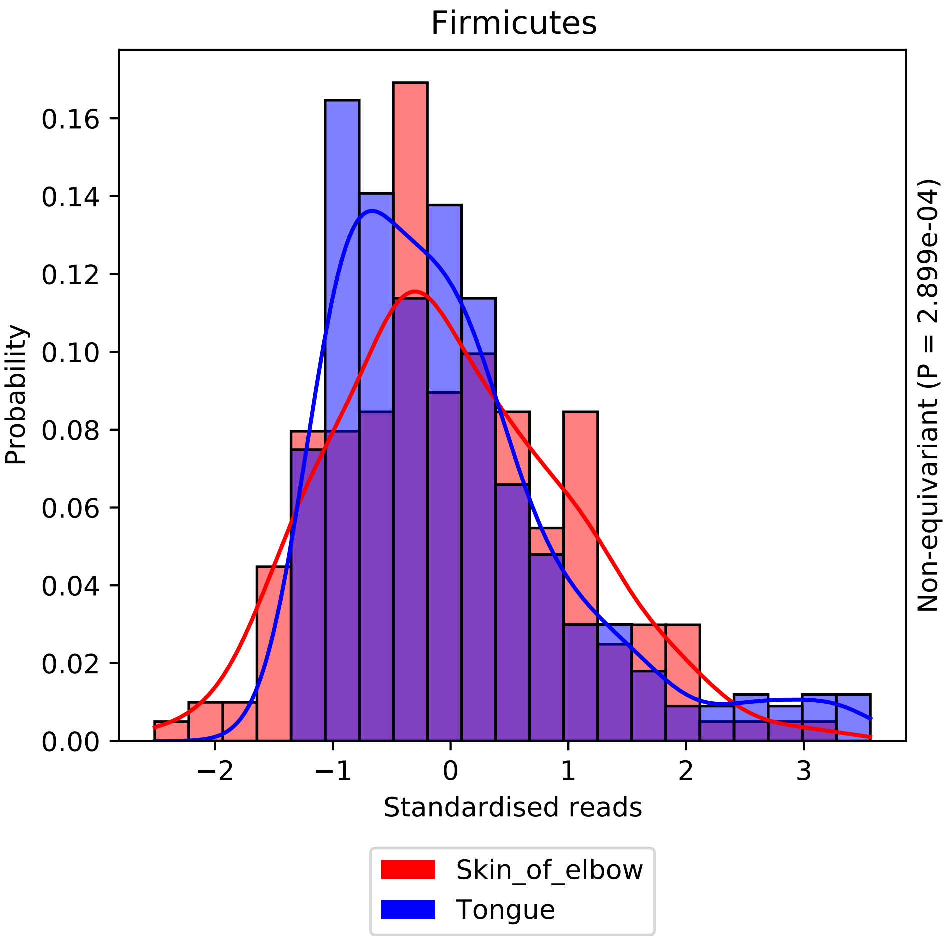

Firmicutes
Probability
0.00
0.02
0.04
0.06
0.08
0.10
0.12
0.14
0.16
−2
−1
0
1
2
3
Standardised reads
Non-equivariant (P = 2.899e-04)
Skin_of_elbow
Tongue

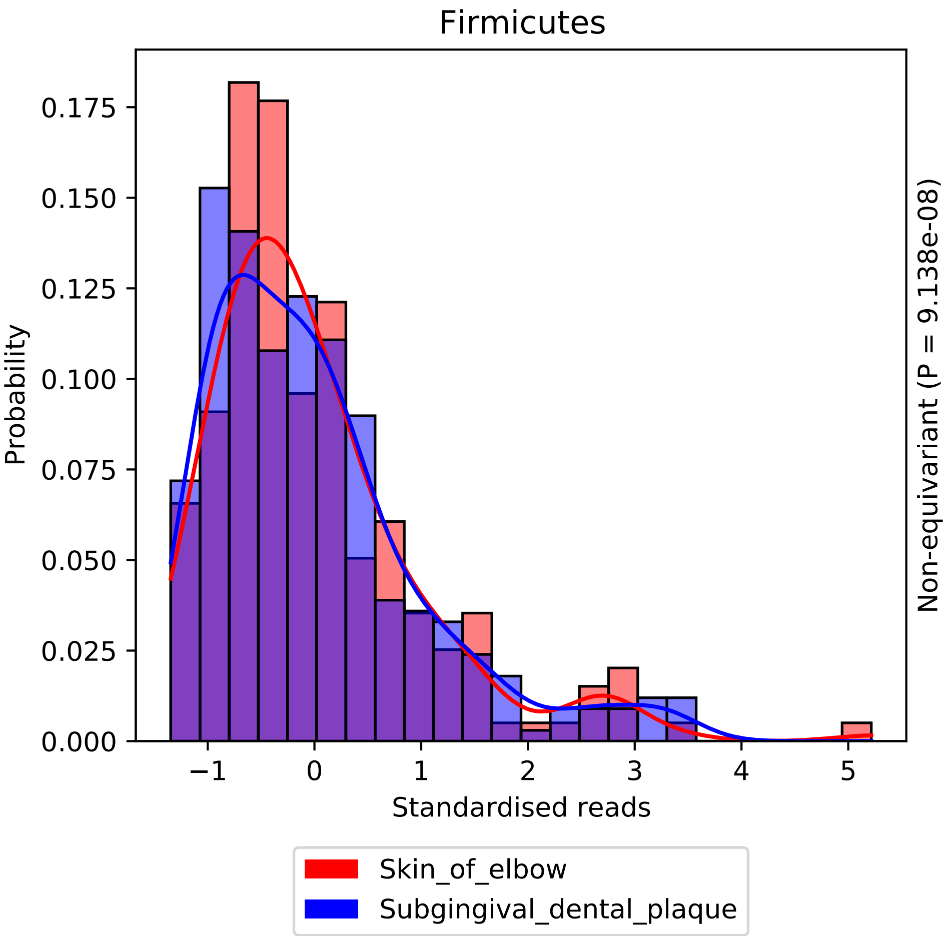
Firmicutes
Probability
0.000
0.025
0.050
0.075
0.100
0.125
0.150
0.175
−1
0
1
2
3
4
5
Standardised reads
Non-equivariant (P = 9.138e-08)
Skin_of_elbow
Subgingival_dental_plaque

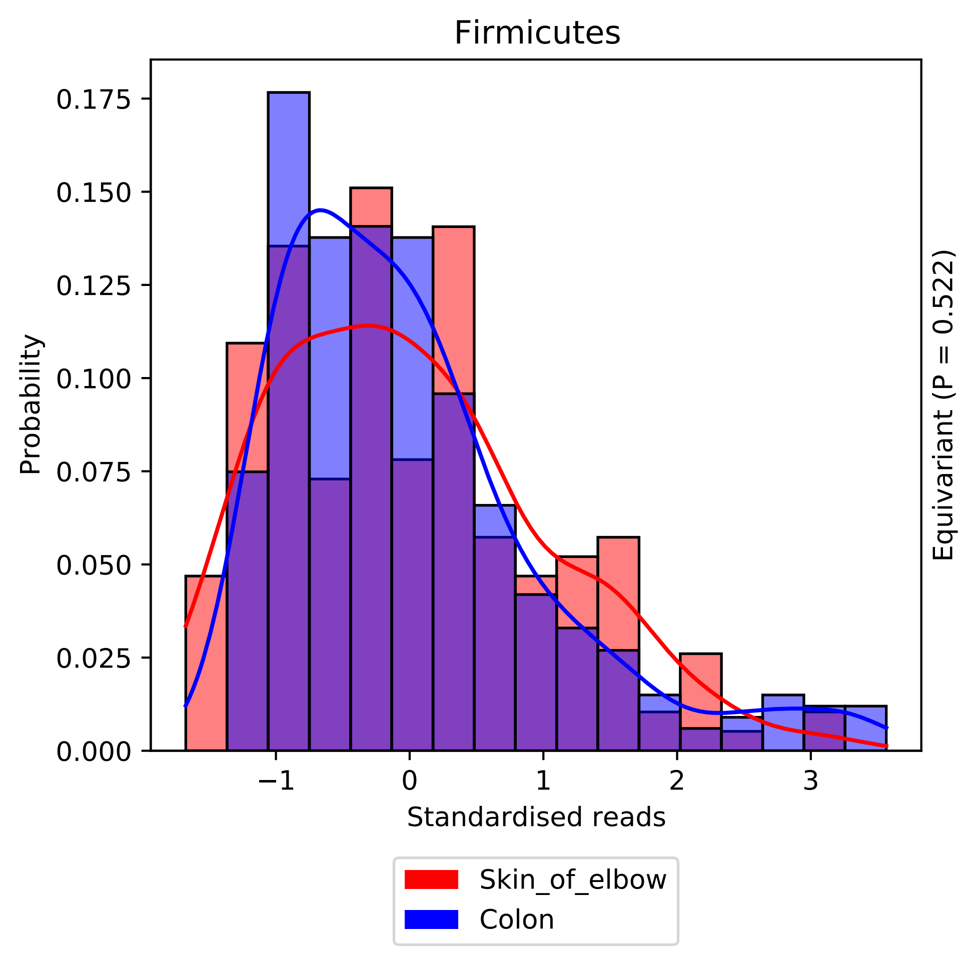

Firmicutes
Probability
0.000
0.025
0.050
0.075
0.100
0.125
0.150
0.175
−1
0
1
2
3
Standardised reads
Equivariant (P = 0.522)
Skin_of_elbow
Colon

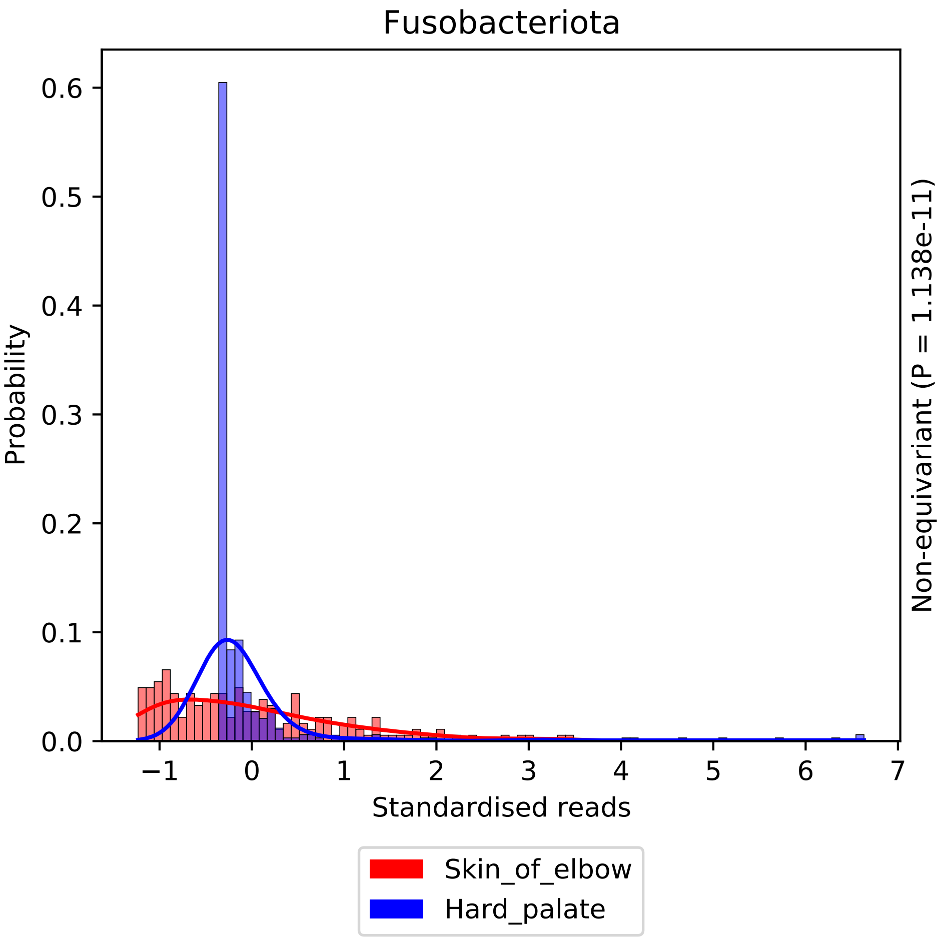

Fusobacteriota
Probability
0.0
0.1
0.2
0.3
0.4
0.5
0.6
−1
0
1
2
3
4
5
6
7
Standardised reads
Non-equivariant (P = 1.138e-11)
Skin_of_elbow
Hard_palate

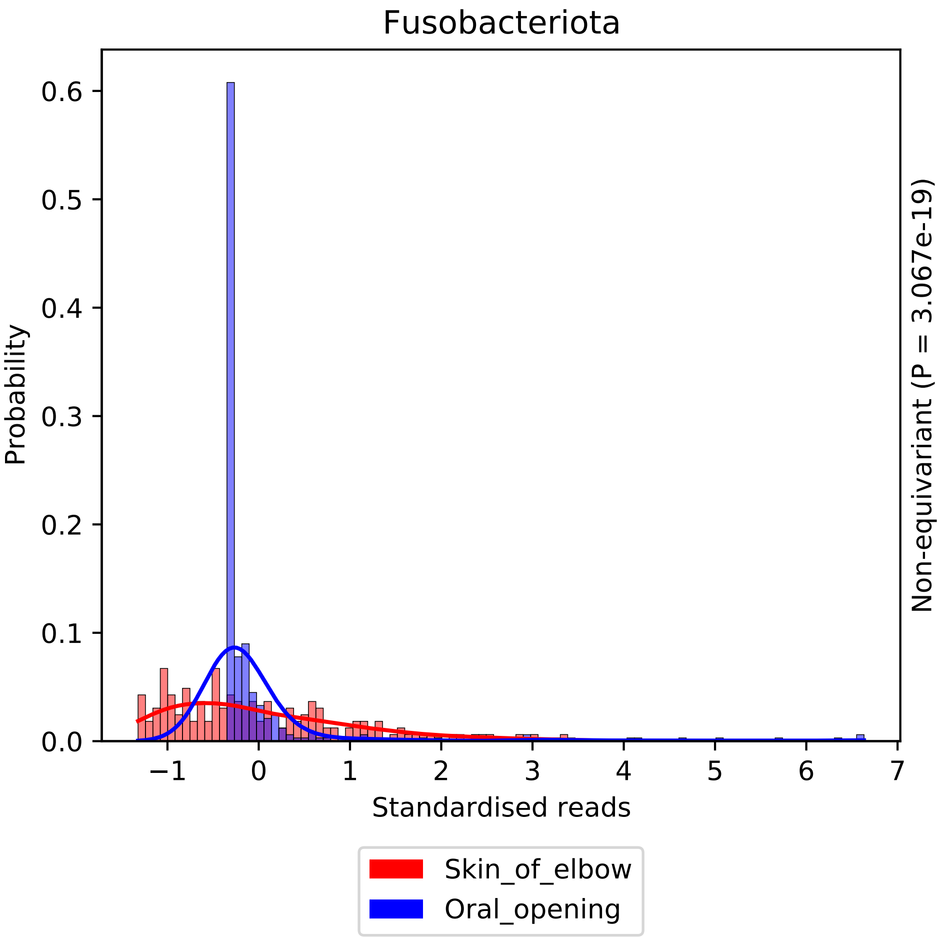

Fusobacteriota
Probability
0.0
0.1
0.2
0.3
0.4
0.5
0.6
−1
0
1
2
3
4
5
6
7
Standardised reads
Non-equivariant (P = 3.067e-19)
Skin_of_elbow
Oral_opening

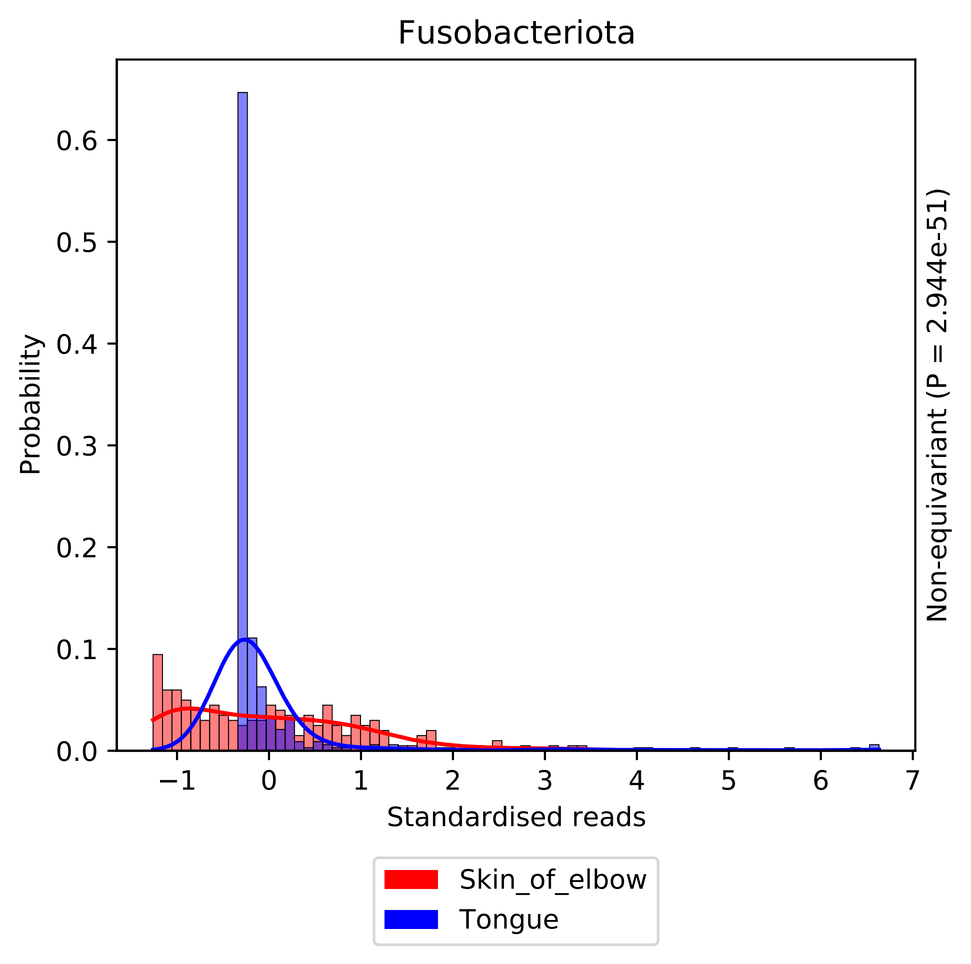

Fusobacteriota
Probability
0.0
0.1
0.2
0.3
0.4
0.5
0.6
−1
0
1
2
3
4
5
6
7
Standardised reads
Non-equivariant (P = 2.944e-51)
Skin_of_elbow
Tongue

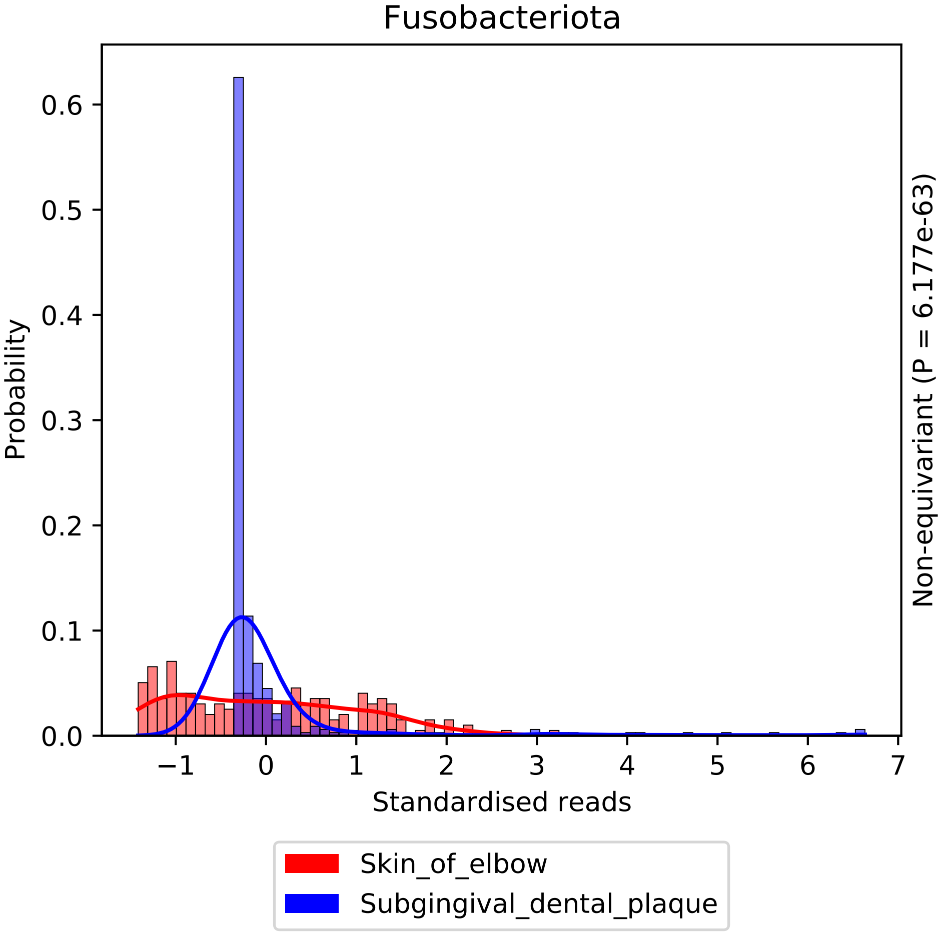

Fusobacteriota
Probability
Standardised reads
Non-equivariant (P = 6.177e-63)
0.0
0.1
0.2
0.3
0.4
0.5
0.6
−1
0
1
2
3
4
5
6
7
Skin_of_elbow
Subgingival_dental_plaque

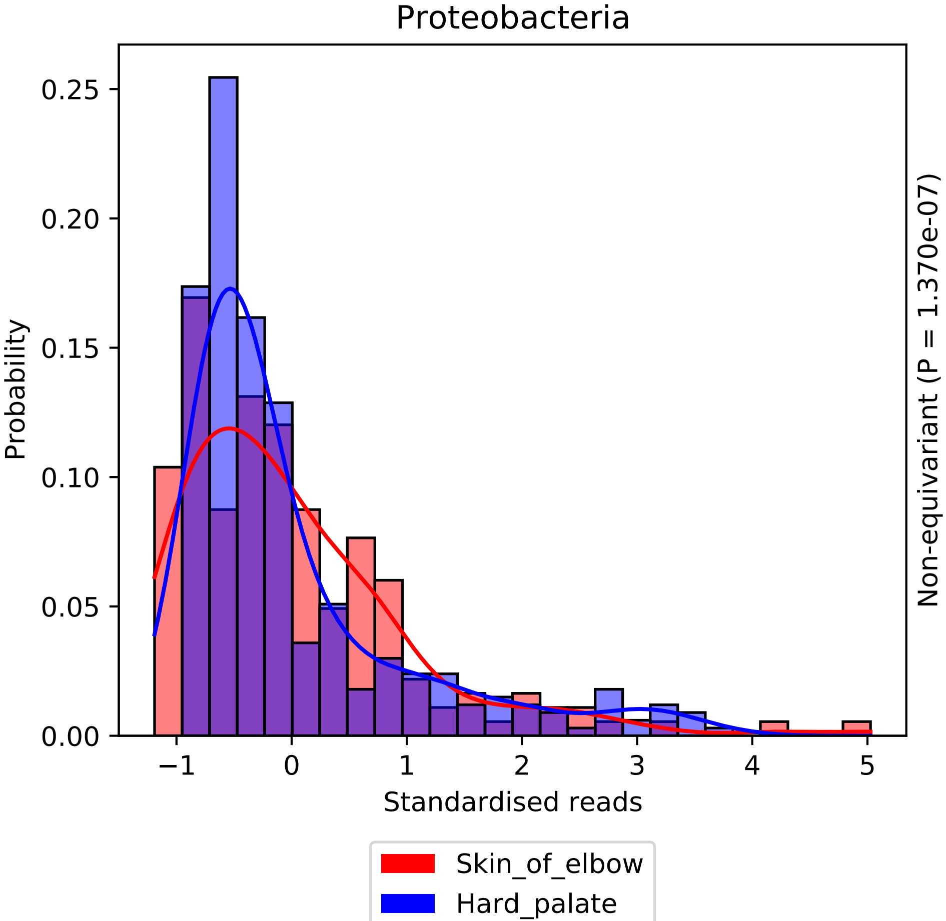

Proteobacteria
Probability
0.25
0.20
0.15
0.10
0.05
0.00
−1
0
1
2
3
4
5
Standardised reads
Non-equivariant (P = 1.370e-07)
Skin_of_elbow
Hard_palate

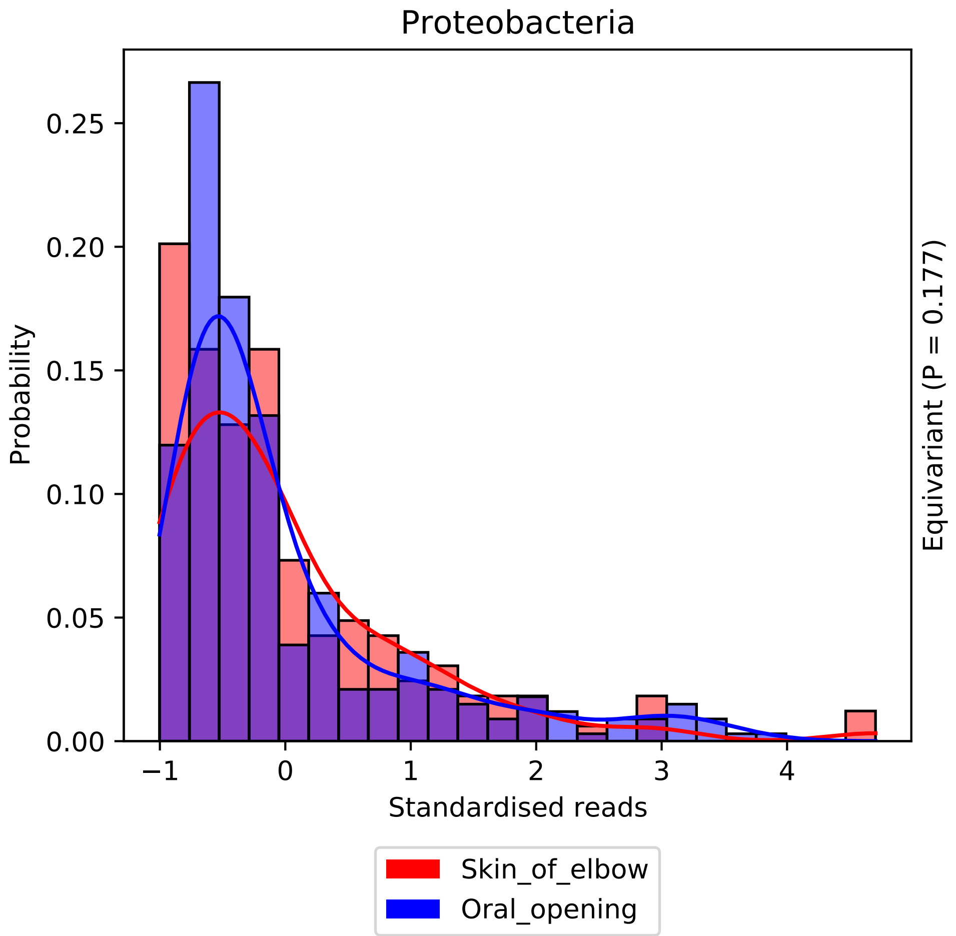
Proteobacteria
Probability
0.25
0.20
0.15
0.10
0.05
0.00
−1
0
1
2
3
4
Standardised reads
Equivariant (P = 0.177)
Skin_of_elbow
Oral_opening

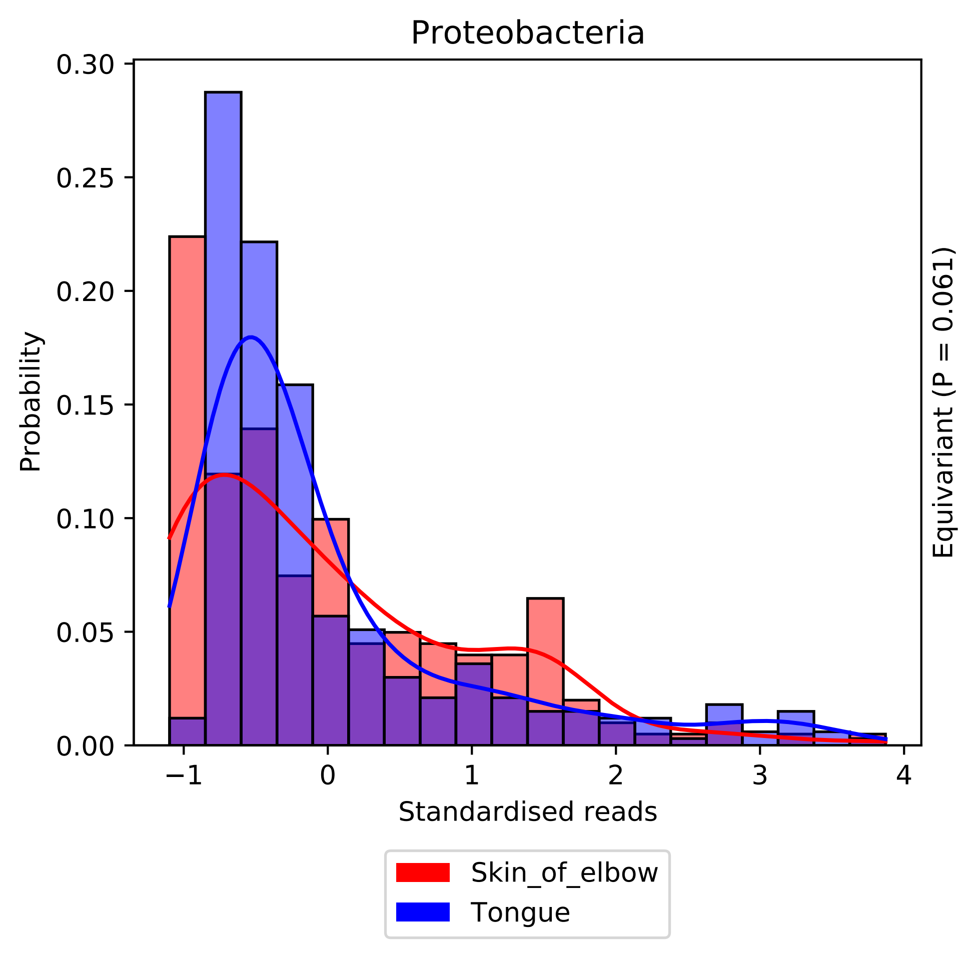

Proteobacteria
0.30
0.25
0.20
0.15
0.10
0.05
0.00
Probability
−1
0
1
2
3
4
Standardised reads
Equivariant (P = 0.061)
Skin_of_elbow
Tongue

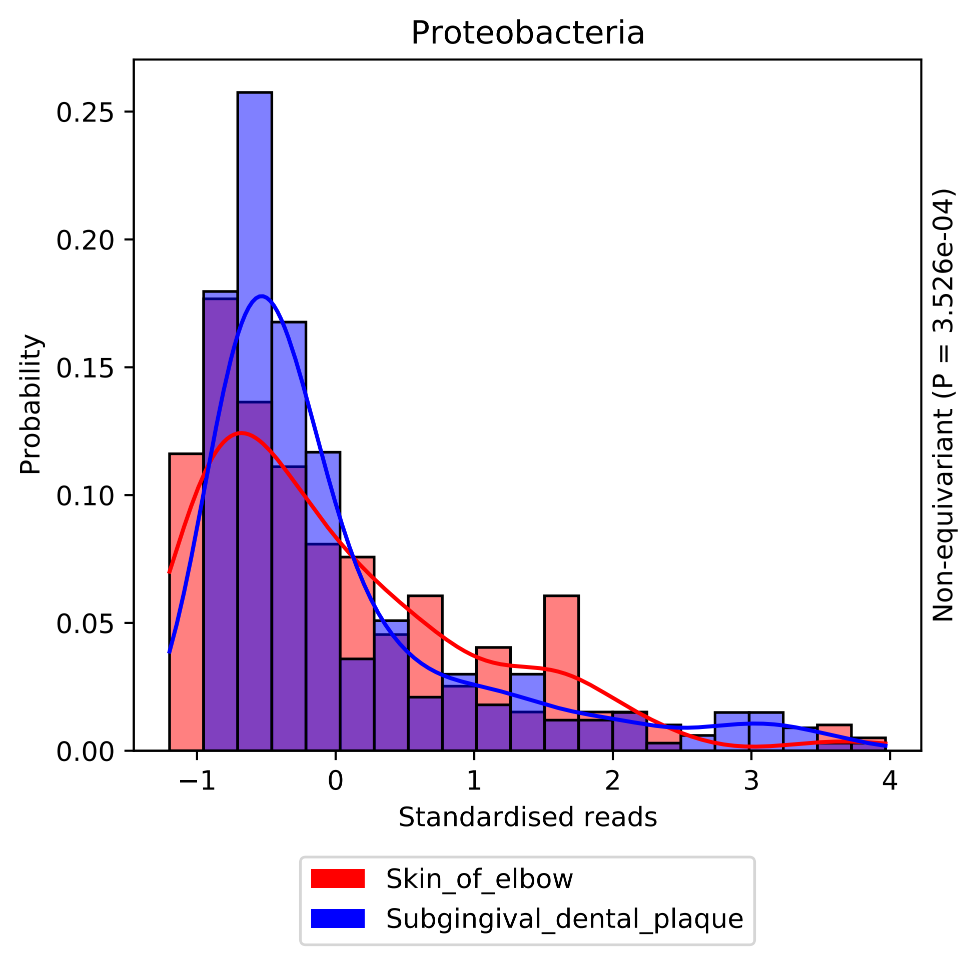
Proteobacteria
Probability
0.00
0.05
0.10
0.15
0.20
0.25
−1
0
1
2
3
4
Standardised reads
Non-equivariant (P = 3.526e-04)
Skin_of_elbow
Subgingival_dental_plaque

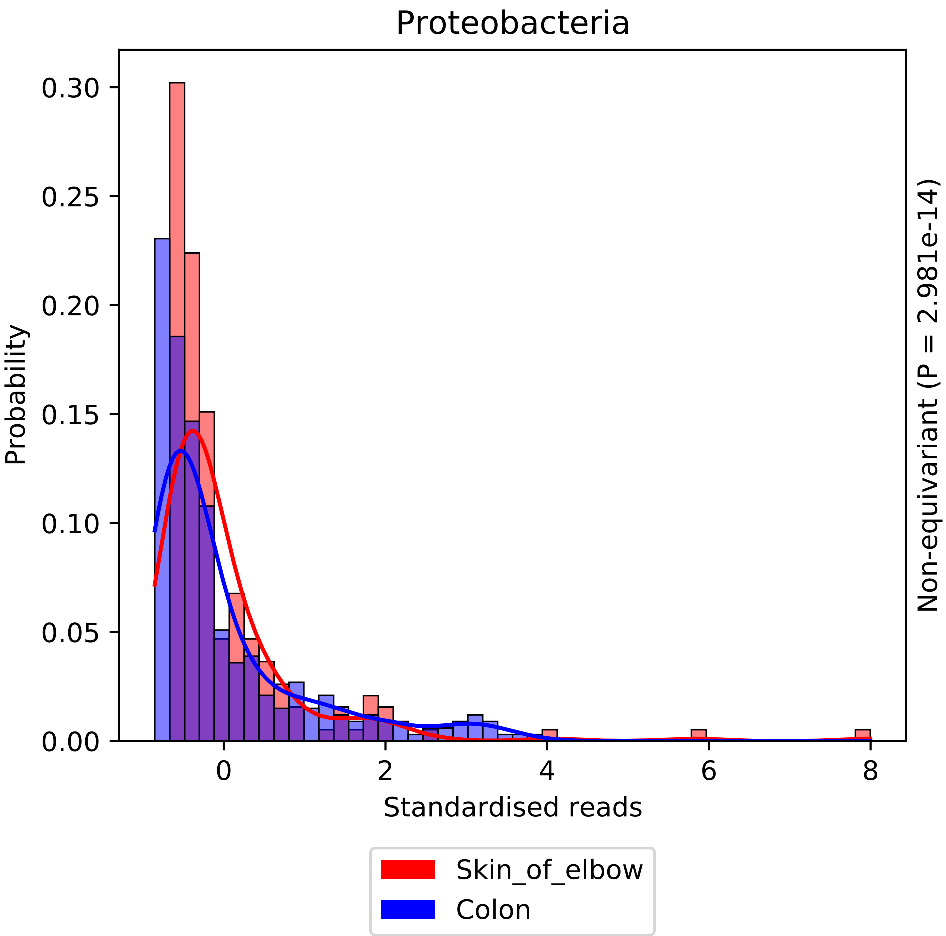
Proteobacteria
Probability
0.00
0.05
0.10
0.15
0.20
0.25
0.30
0
2
4
6
8
Standardised reads
Non-equivariant (P = 2.981e-14)
Skin_of_elbow
Colon

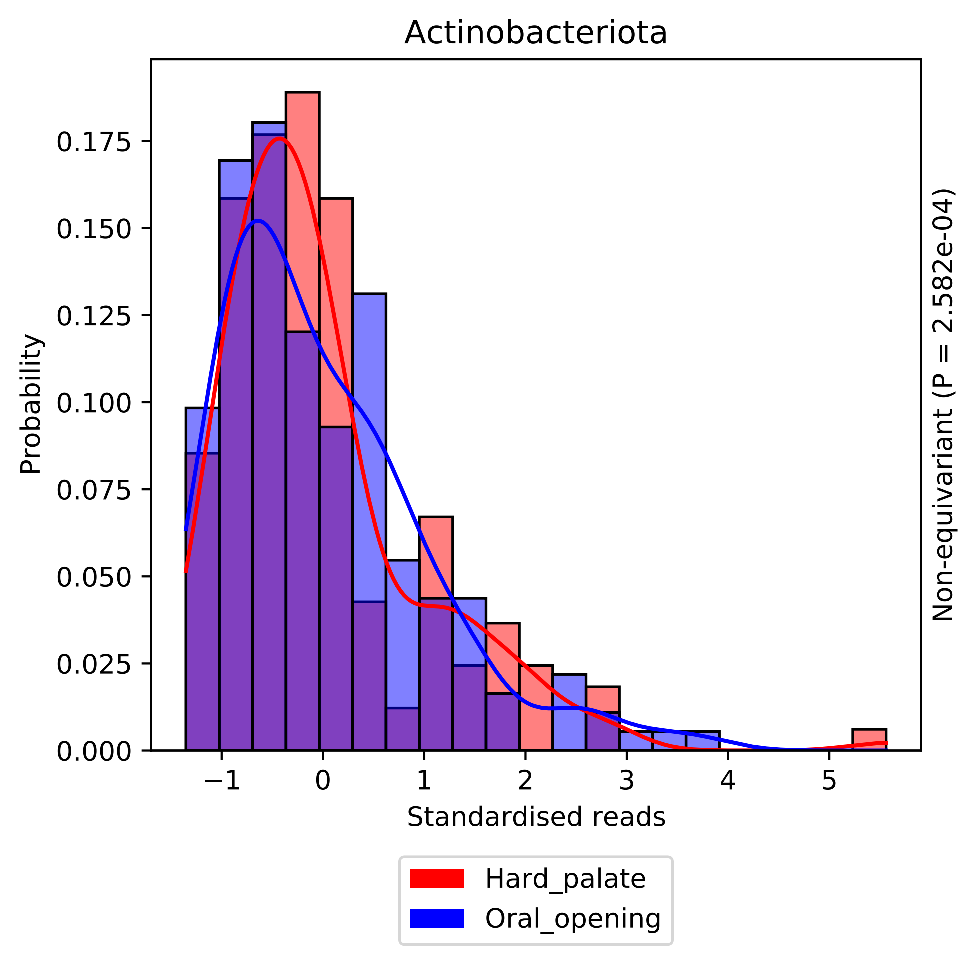

Actinobacteriota
Probability
0.000
0.025
0.050
0.075
0.100
0.125
0.150
0.175
−1
0
1
2
3
4
5
Standardised reads
Non-equivariant (P = 2.582e-04)
Hard_palate
Oral_opening

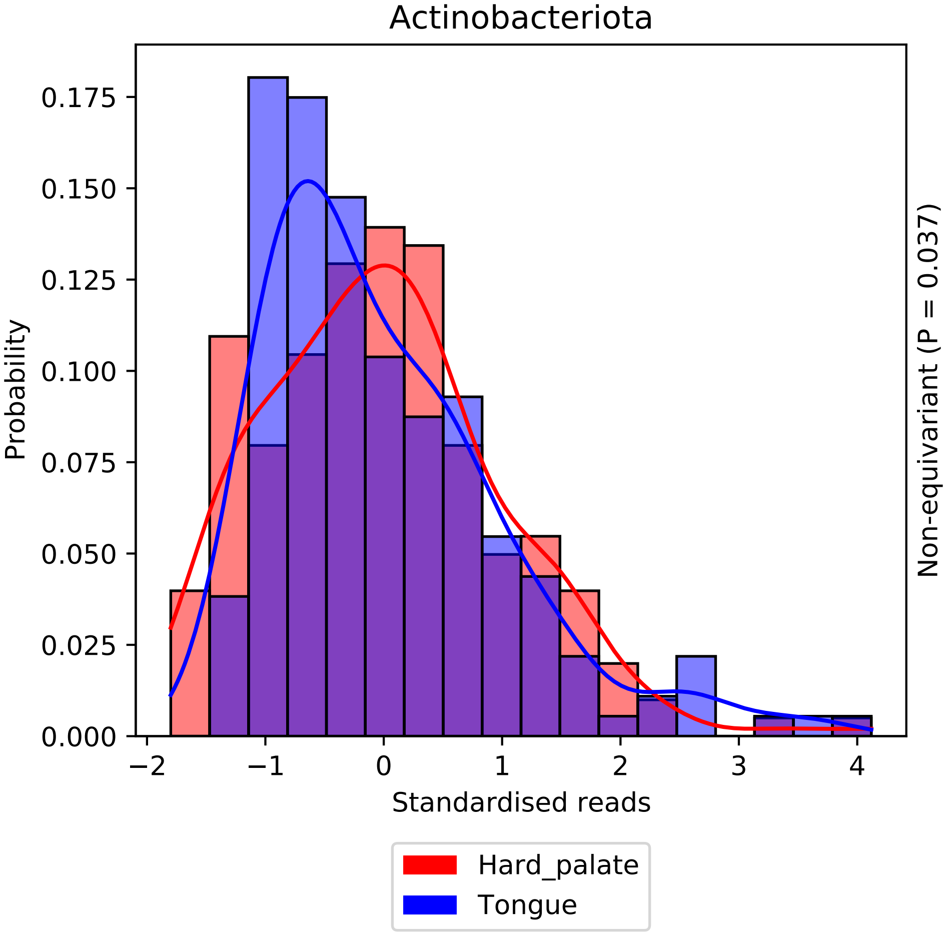

Actinobacteriota
0.175
0.150
0.125
0.100
0.075
0.050
0.025
0.000
Probability
−2
−1
0
1
2
3
4
Standardised reads
Non-equivariant (P = 0.037)
Hard_palate
Tongue

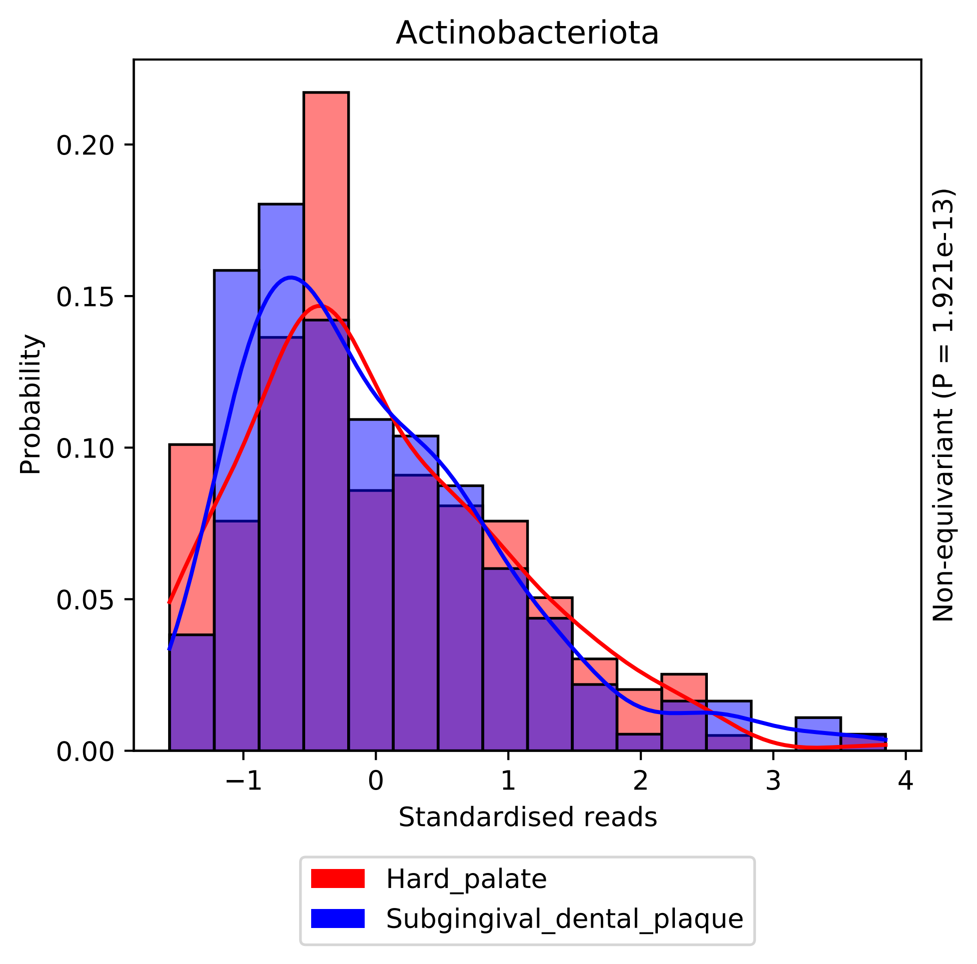

Actinobacteriota
Probability
0.00
0.05
0.10
0.15
0.20
−1
0
1
2
3
4
Standardised reads
Non-equivariant (P = 1.921e-13)
Hard_palate
Subgingival_dental_plaque

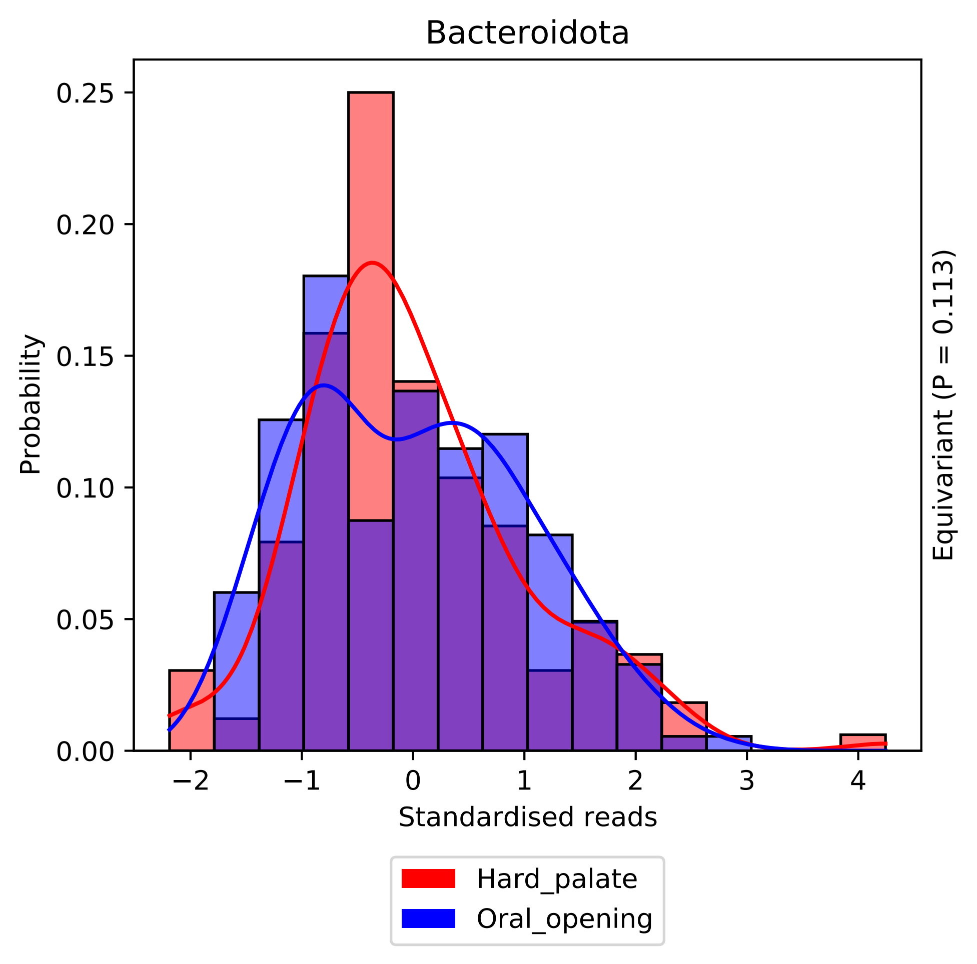

Bacteroidota
Probability
0.25
0.20
0.15
0.10
0.05
0.00
−2
−1
0
1
2
3
4
Standardised reads
Equivariant (P = 0.113)
Hard_palate
Oral_opening

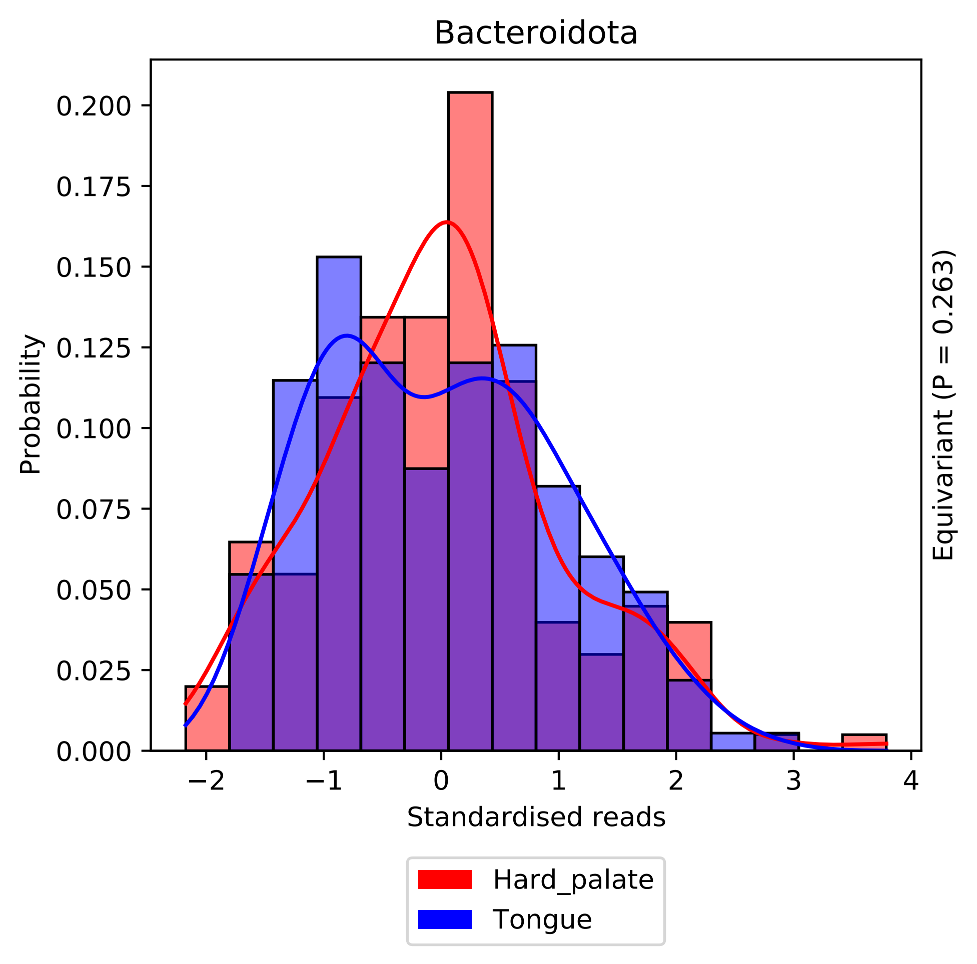

Bacteroidota
Probability
0.000
0.025
0.050
0.075
0.100
0.125
0.150
0.175
0.200
−2
−1
0
1
2
3
4
Standardised reads
Equivariant (P = 0.263)
Hard_palate
Tongue

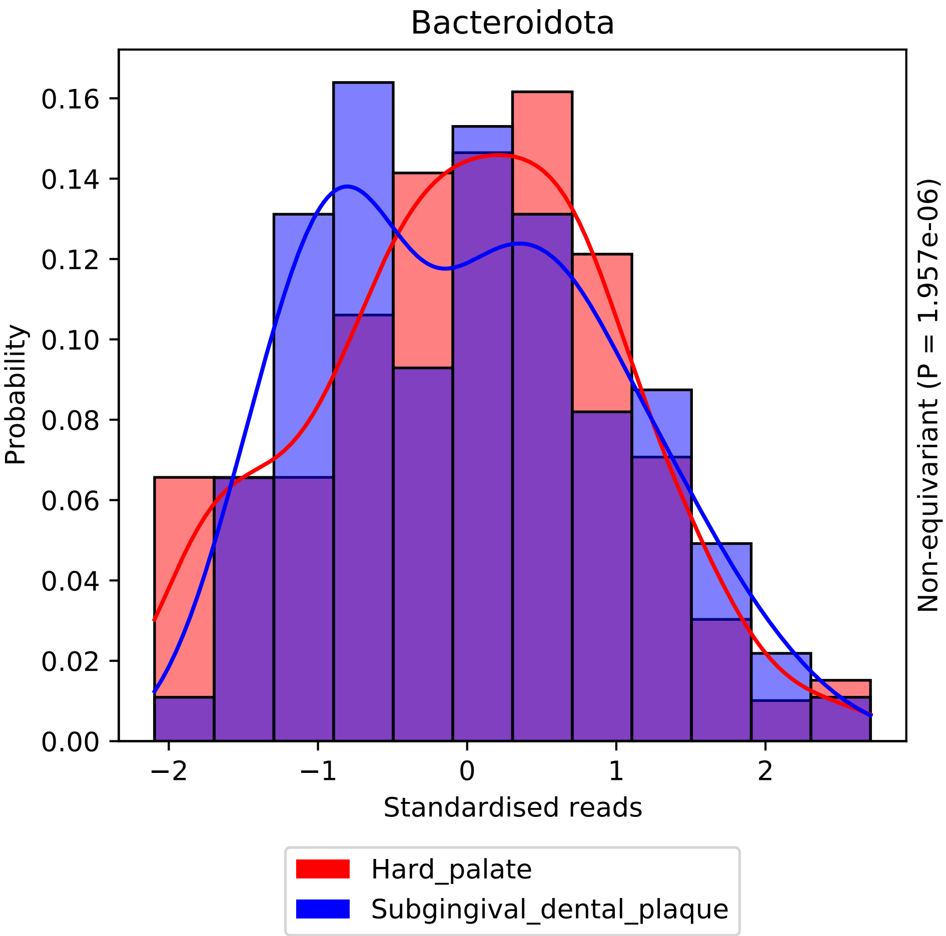

Bacteroidota
Probability
Standardised reads
Non-equivariant (P = 1.957e-06)
0.00
0.02
0.04
0.06
0.08
0.10
0.12
0.14
0.16
−2
−1
0
1
2
Hard_palate
Subgingival_dental_plaque

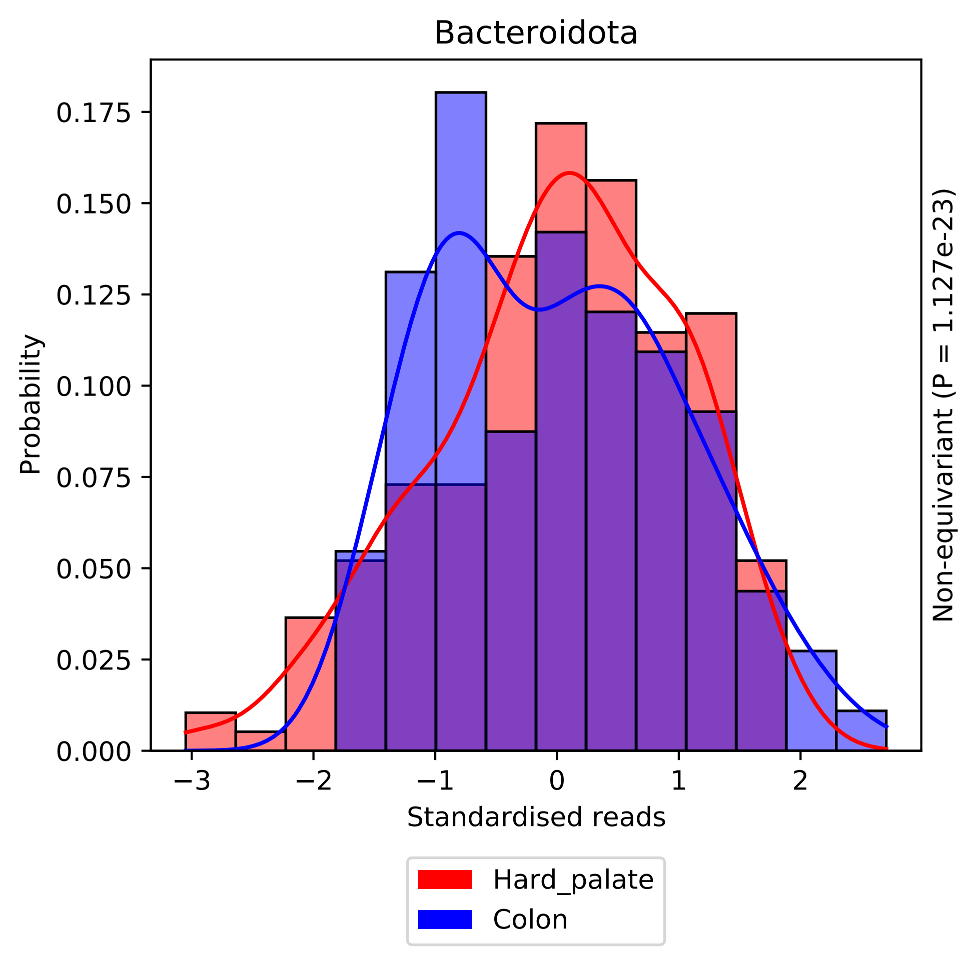

Bacteroidota
Probability
0.000
0.025
0.050
0.075
0.100
0.125
0.150
0.175
−3
−2
−1
0
1
2
Standardised reads
Non-equivariant (P = 1.127e-23)
Hard_palate
Colon

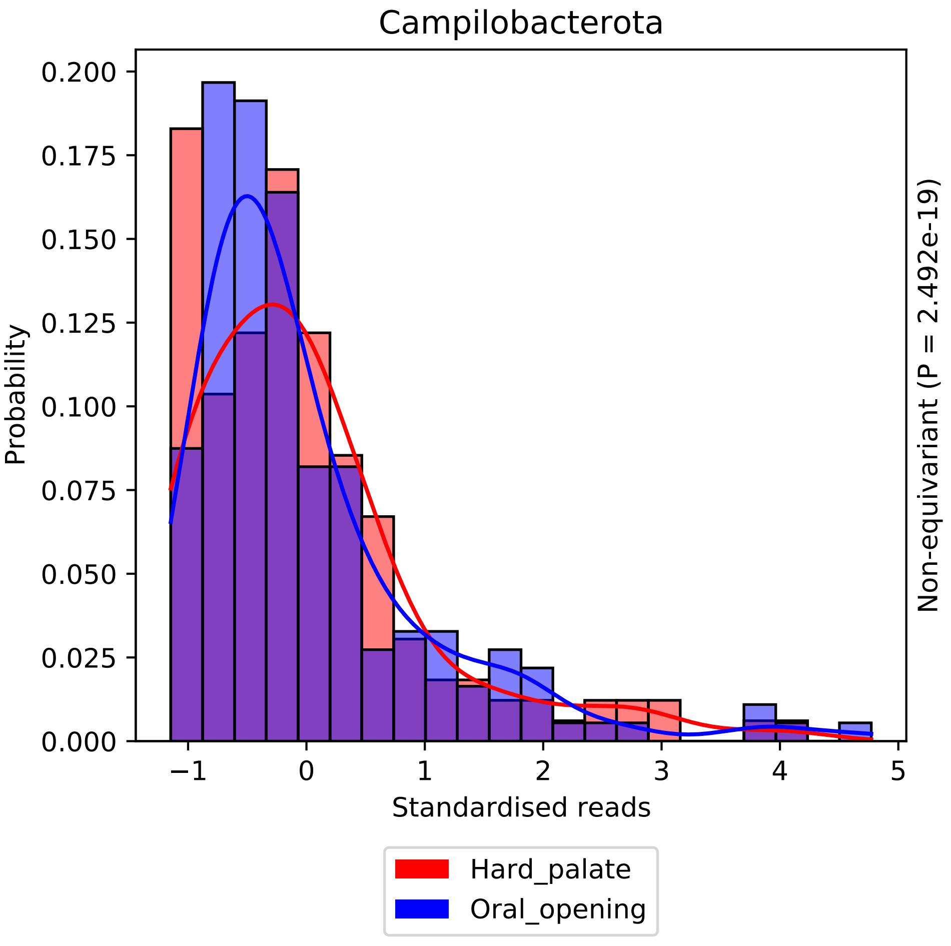

Campilobacterota
Probability
Standardised reads
Non-equivariant (P = 2.492e-19)
0.000
0.025
0.050
0.075
0.100
0.125
0.150
0.175
0.200
−1
0
1
2
3
4
5
Hard_palate
Oral_opening

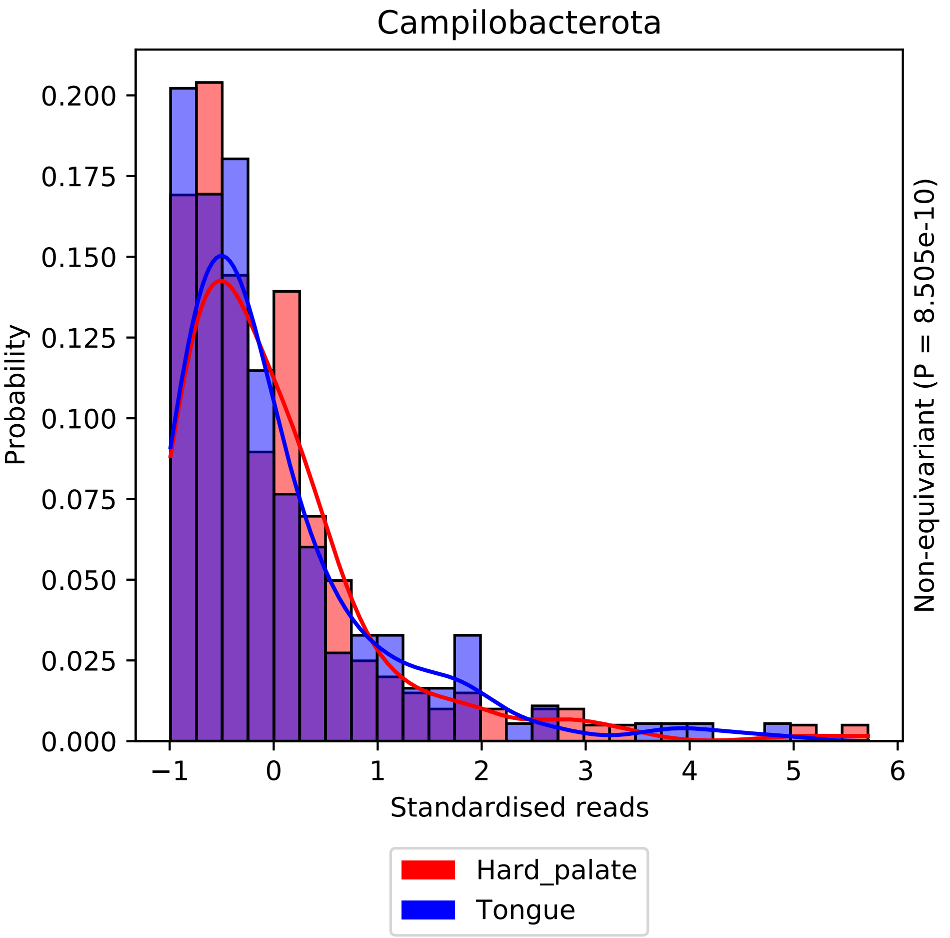

Campilobacterota
Probability
0.000
0.025
0.050
0.075
0.100
0.125
0.150
0.175
0.200
−1
0
1
2
3
4
5
6
Standardised reads
Non-equivariant (P = 8.505e-10)
Hard_palate
Tongue

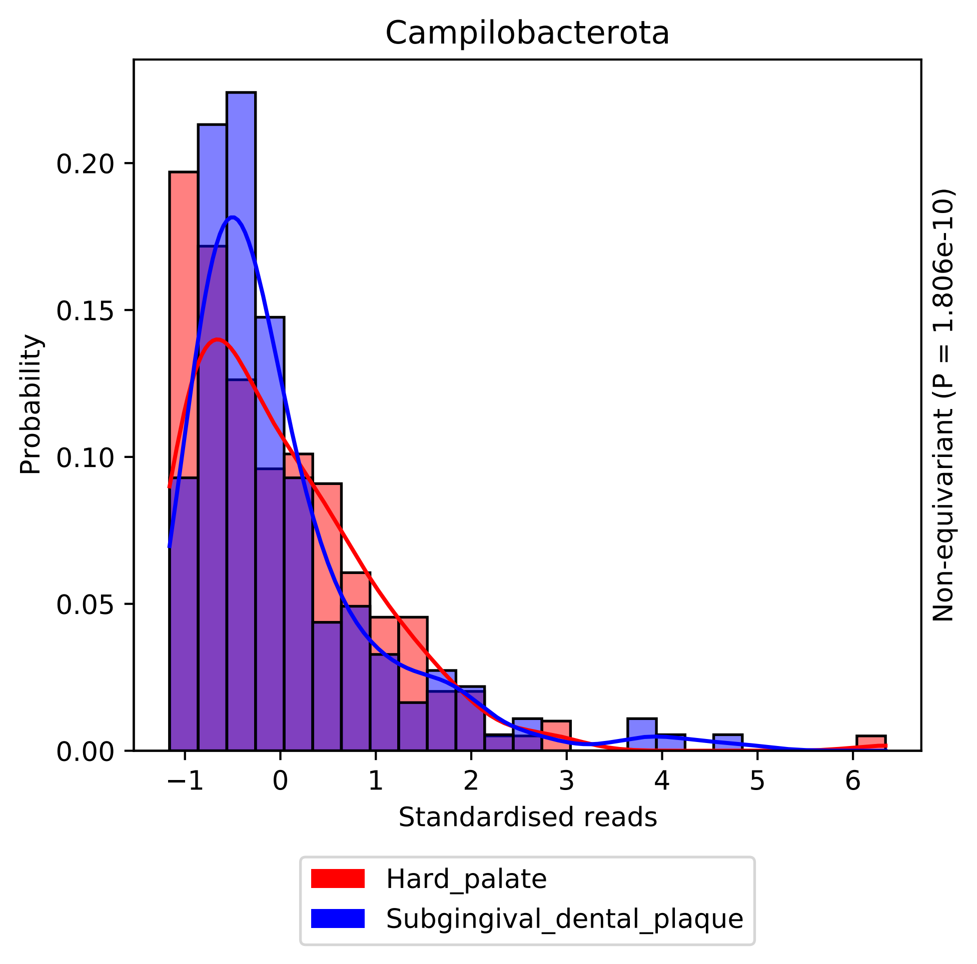

Campilobacterota
Probability
0.00
0.05
0.10
0.15
0.20
−1
0
1
2
3
4
5
6
Standardised reads
Non-equivariant (P = 1.806e-10)
Hard_palate
Subgingival_dental_plaque

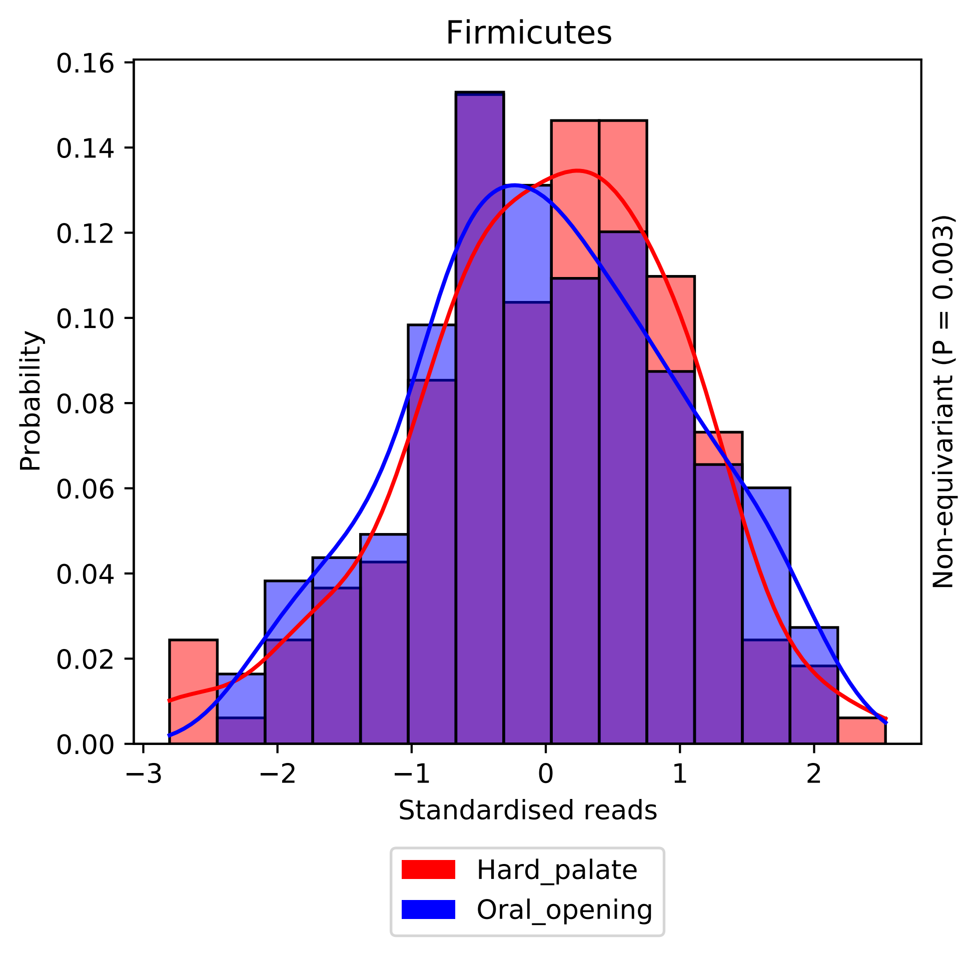
Firmicutes
Probability
Standardised reads
0.00
0.02
0.04
0.06
0.08
0.10
0.12
0.14
0.16
−3
−2
−1
0
1
2
Non-equivariant (P = 0.003)
Hard_palate
Oral_opening

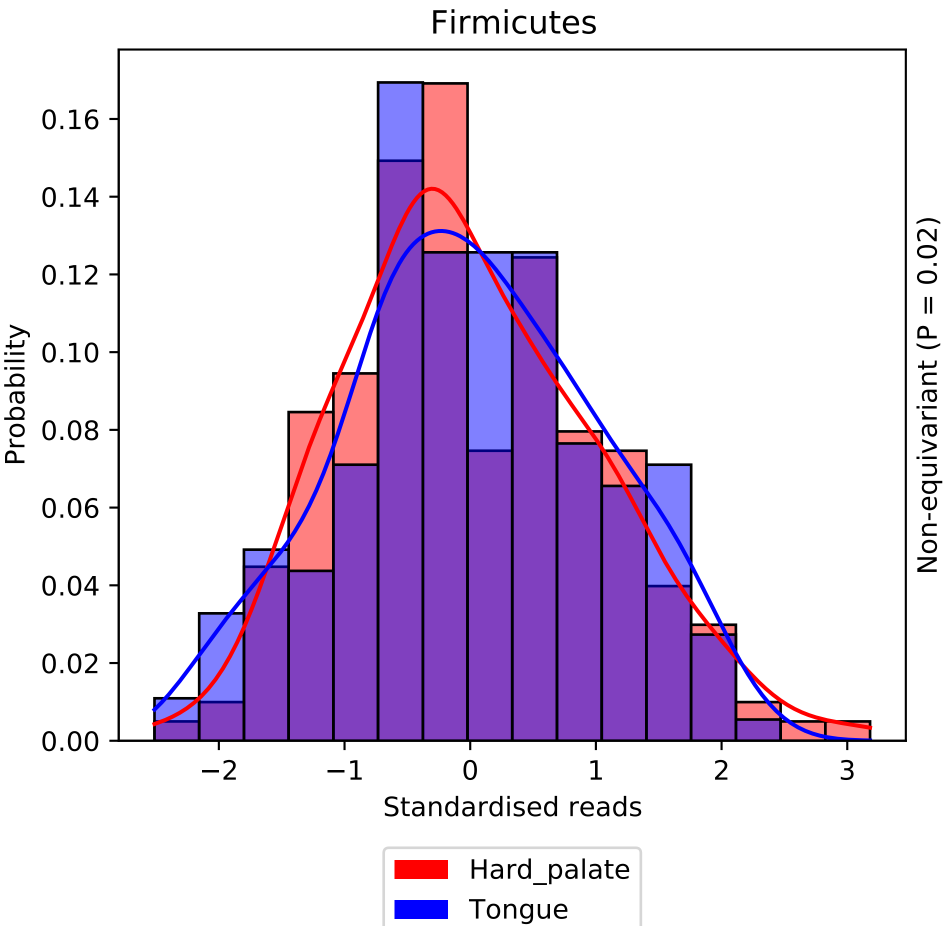

Firmicutes
Probability
0.00
0.02
0.04
0.06
0.08
0.10
0.12
0.14
0.16
−2
−1
0
1
2
3
Standardised reads
Non-equivariant (P = 0.02)
Hard_palate
Tongue

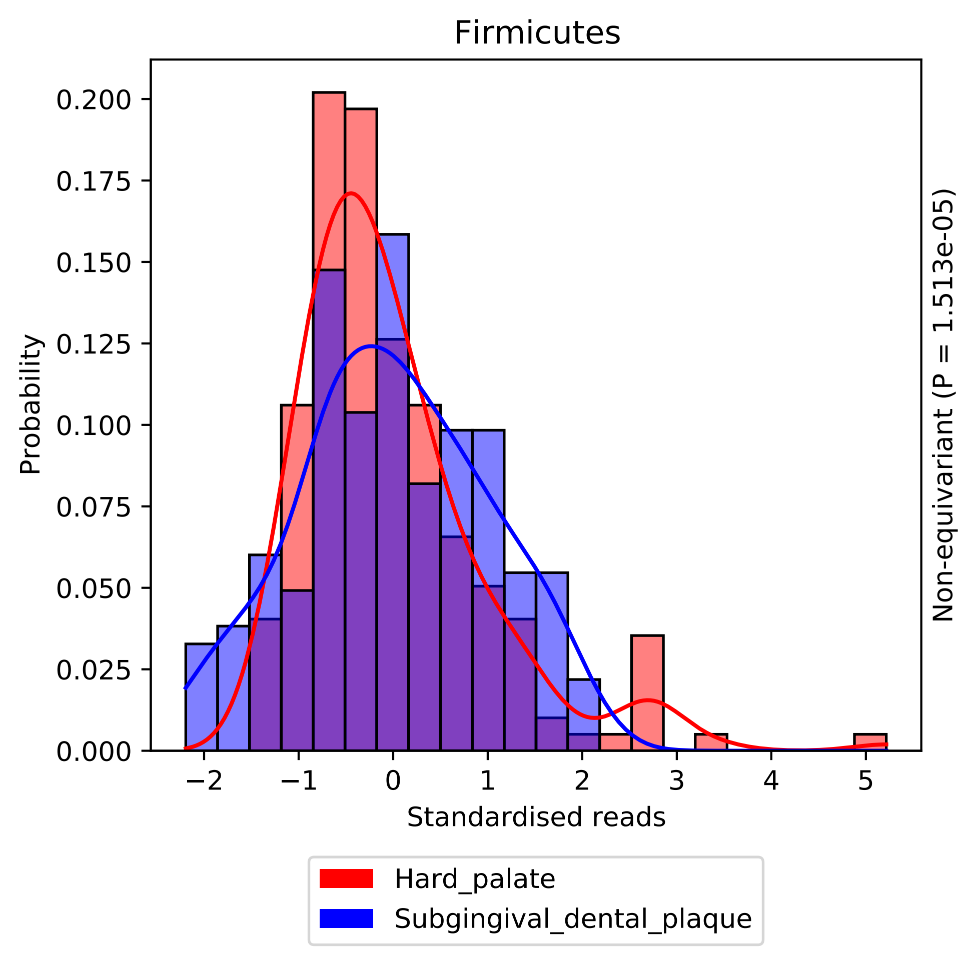

Firmicutes
Probability
Standardised reads
Non-equivariant (P = 1.513e-05)
0.000
0.025
0.050
0.075
0.100
0.125
0.150
0.175
0.200
−2
−1
0
1
2
3
4
5
Hard_palate
Subgingival_dental_plaque

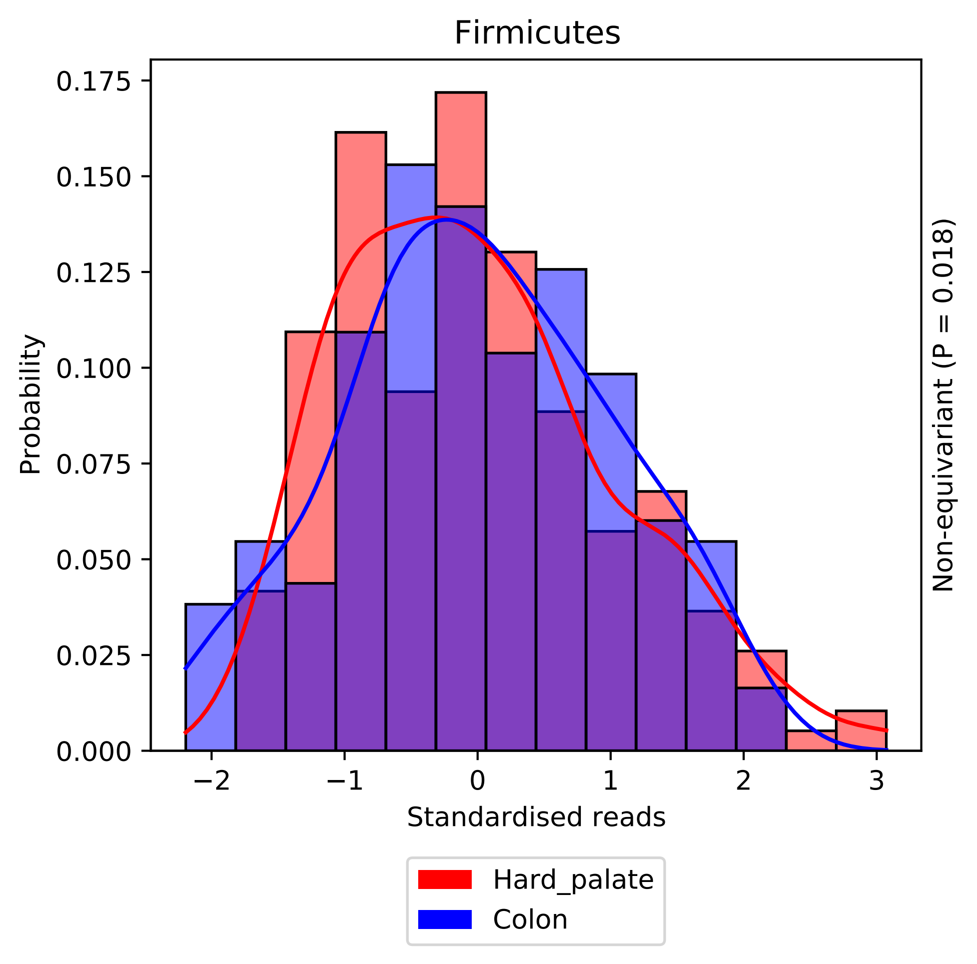

Firmicutes
Probability
0.000
0.025
0.050
0.075
0.100
0.125
0.150
0.175
−2
−1
0
1
2
3
Standardised reads
Non-equivariant (P = 0.018)
Hard_palate
Colon

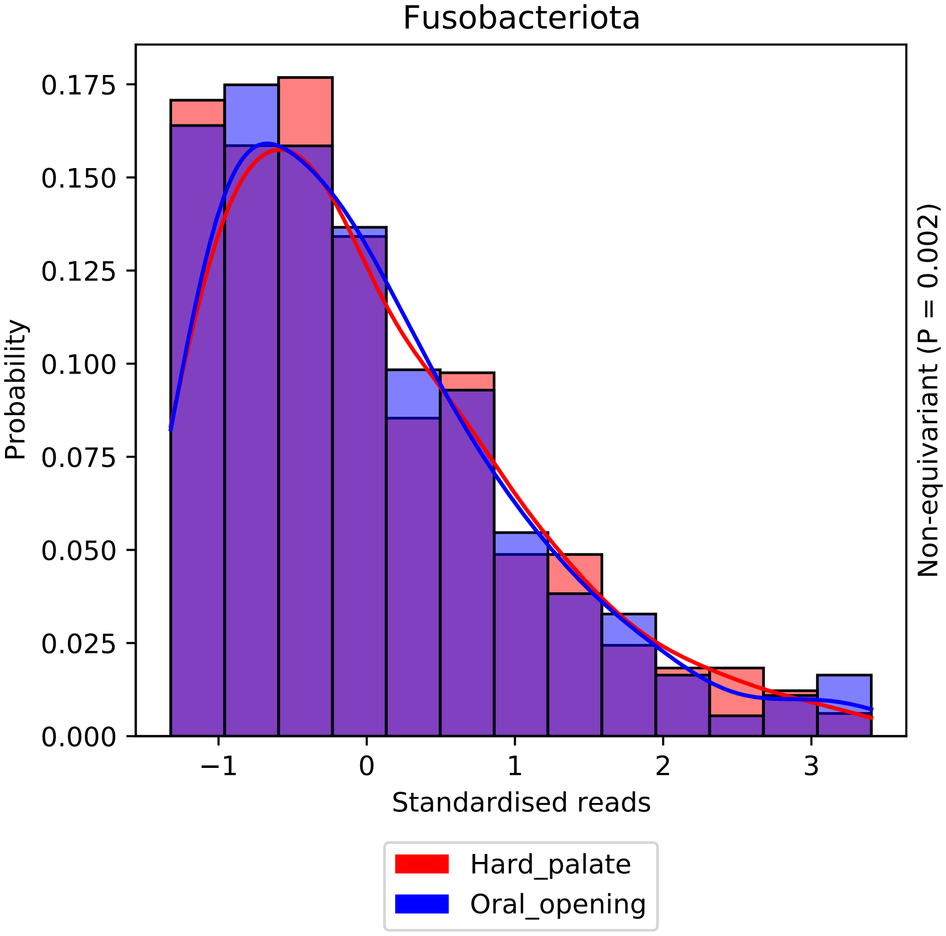
Fusobacteriota
Probability
0.000
0.025
0.050
0.075
0.100
0.125
0.150
0.175
−1
0
1
2
3
Standardised reads
Non-equivariant (P = 0.002)
Hard_palate
Oral_opening

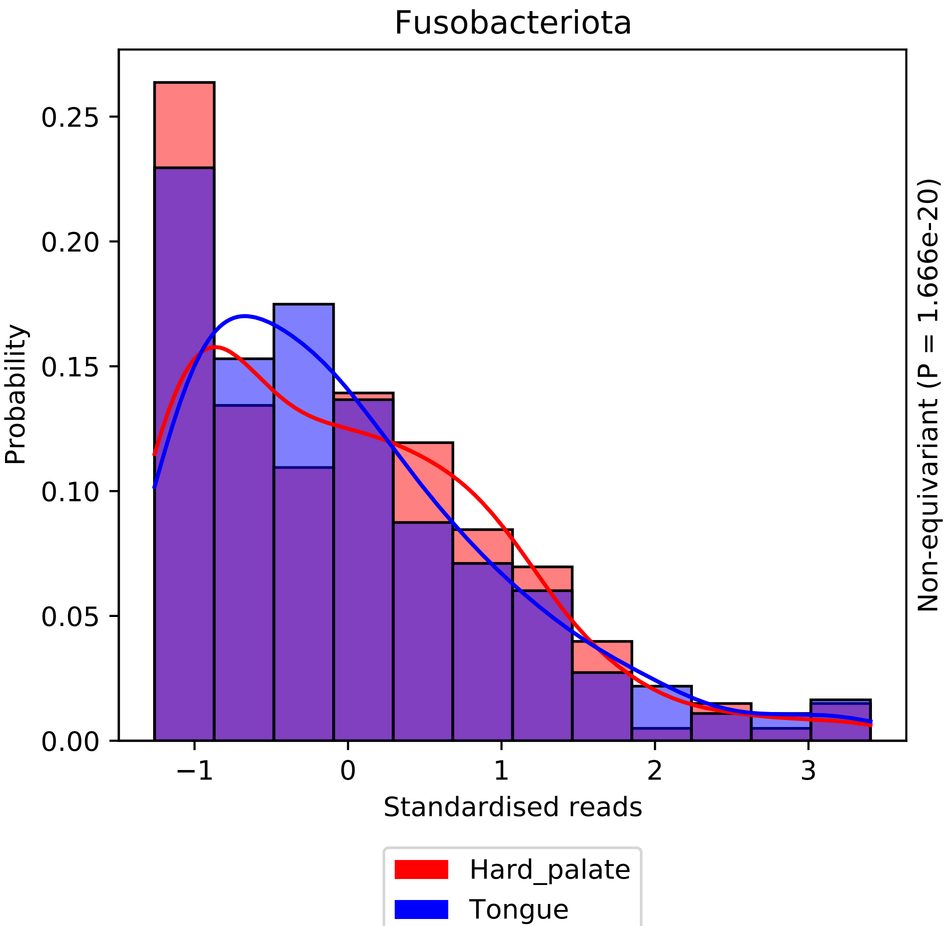

Fusobacteriota
Probability
0.00
0.05
0.10
0.15
0.20
0.25
−1
0
1
2
3
Standardised reads
Non-equivariant (P = 1.666e-20)
Hard_palate
Tongue

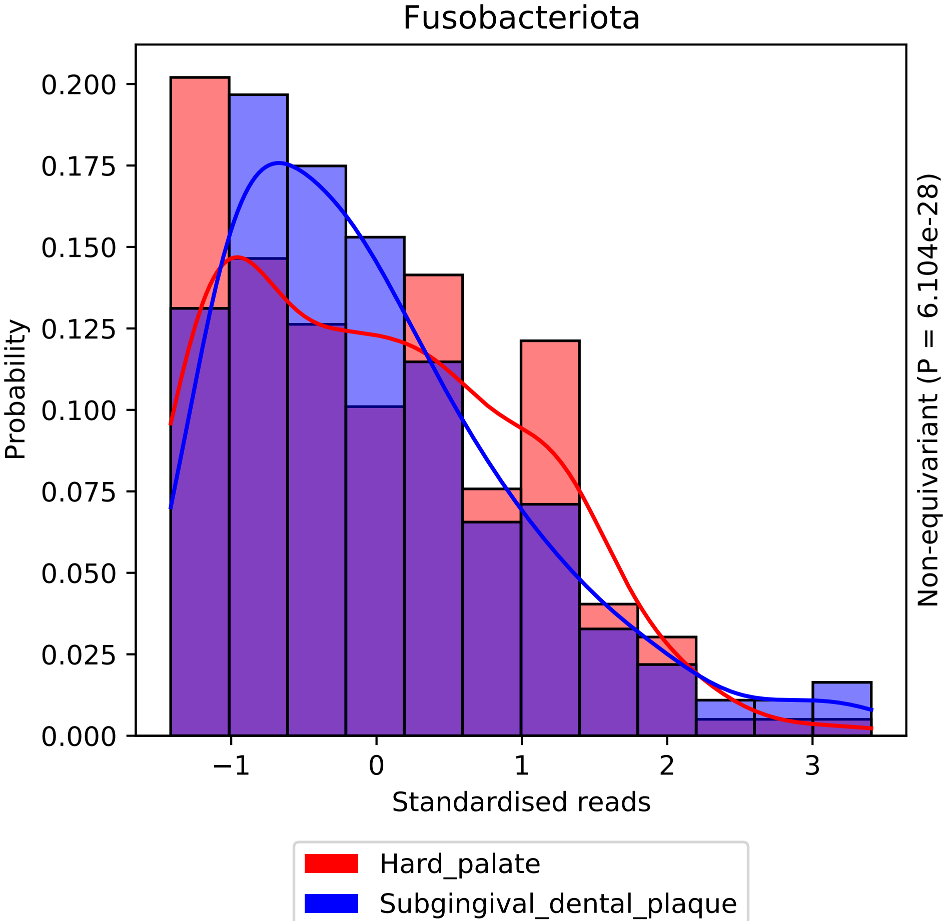
Fusobacteriota
0.200
0.175
0.150
0.125
0.100
0.075
0.050
0.025
0.000
Probability
−1
0
1
2
3
Standardised reads
Non-equivariant (P = 6.104e-28)
Hard_palate
Subgingival_dental_plaque

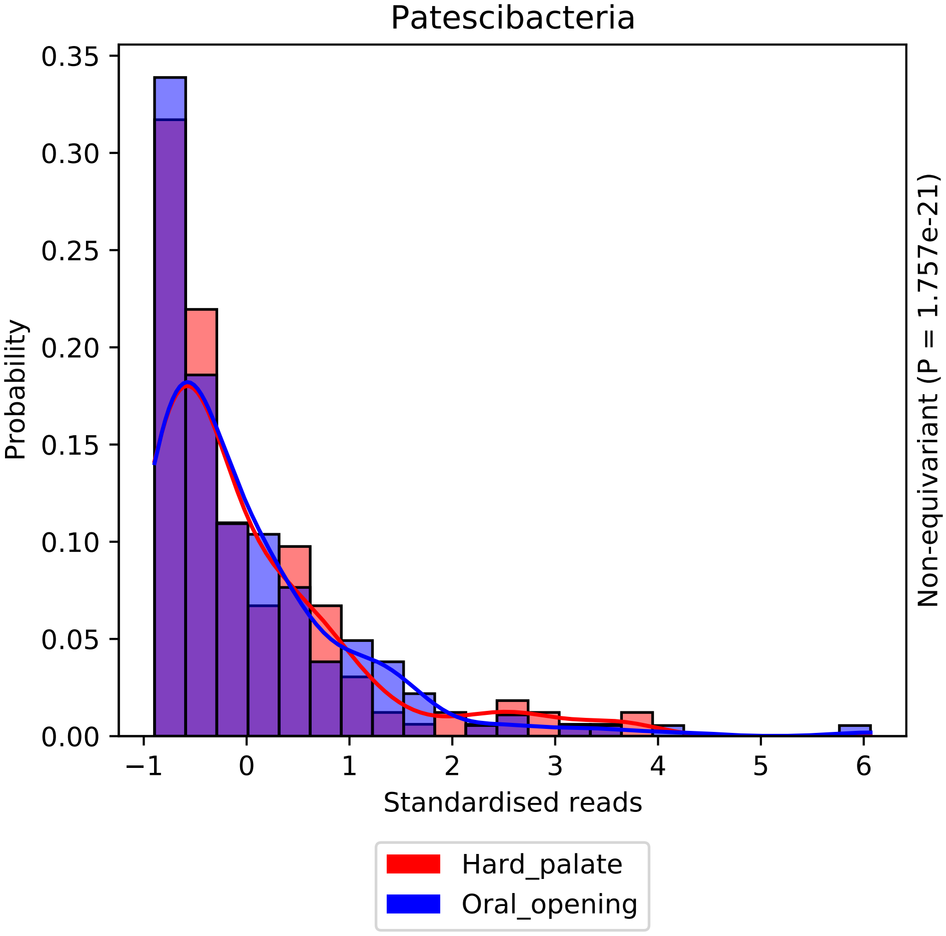
Patescibacteria
Probability
0.35
0.30
0.25
0.20
0.15
0.10
0.05
0.00
−1
0
1
2
3
4
5
6
Standardised reads
Non-equivariant (P = 1.757e-21)
Hard_palate
Oral_opening

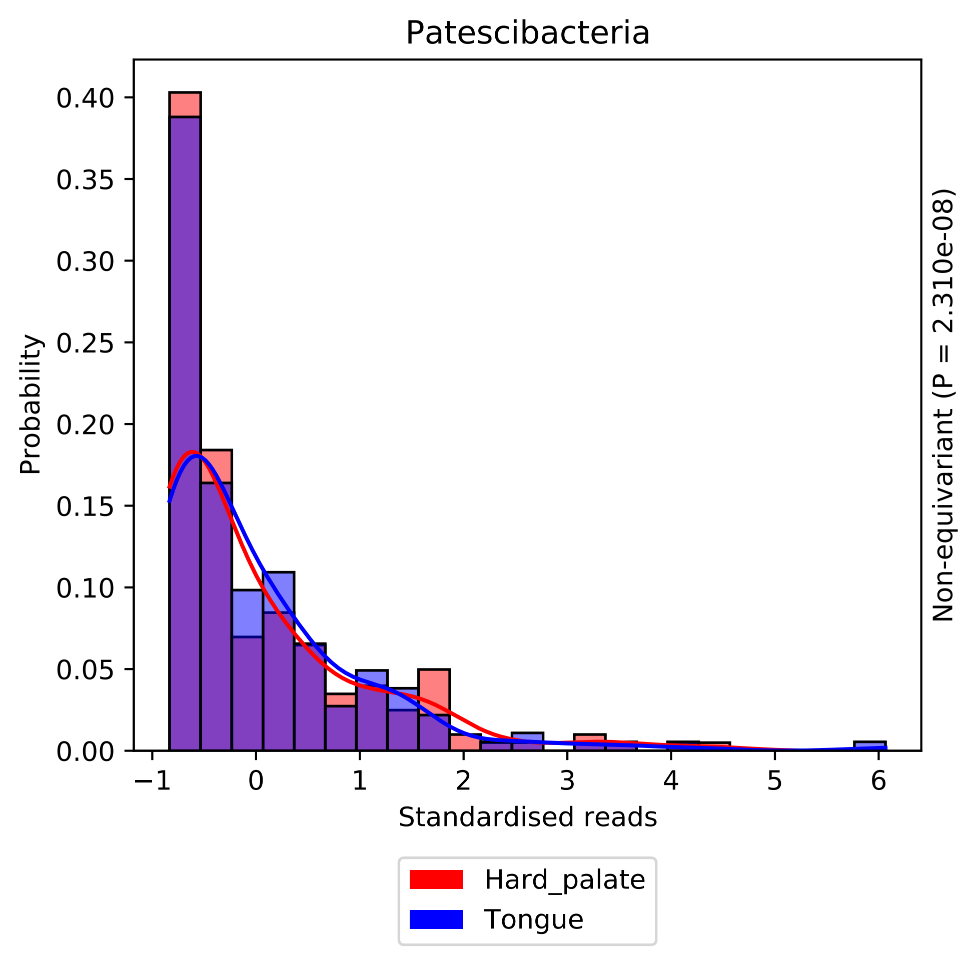

Patescibacteria
Probability
Standardised reads
Non-equivariant (P = 2.310e-08)
0.00
0.05
0.10
0.15
0.20
0.25
0.30
0.35
0.40
−1
0
1
2
3
4
5
6
Hard_palate
Tongue

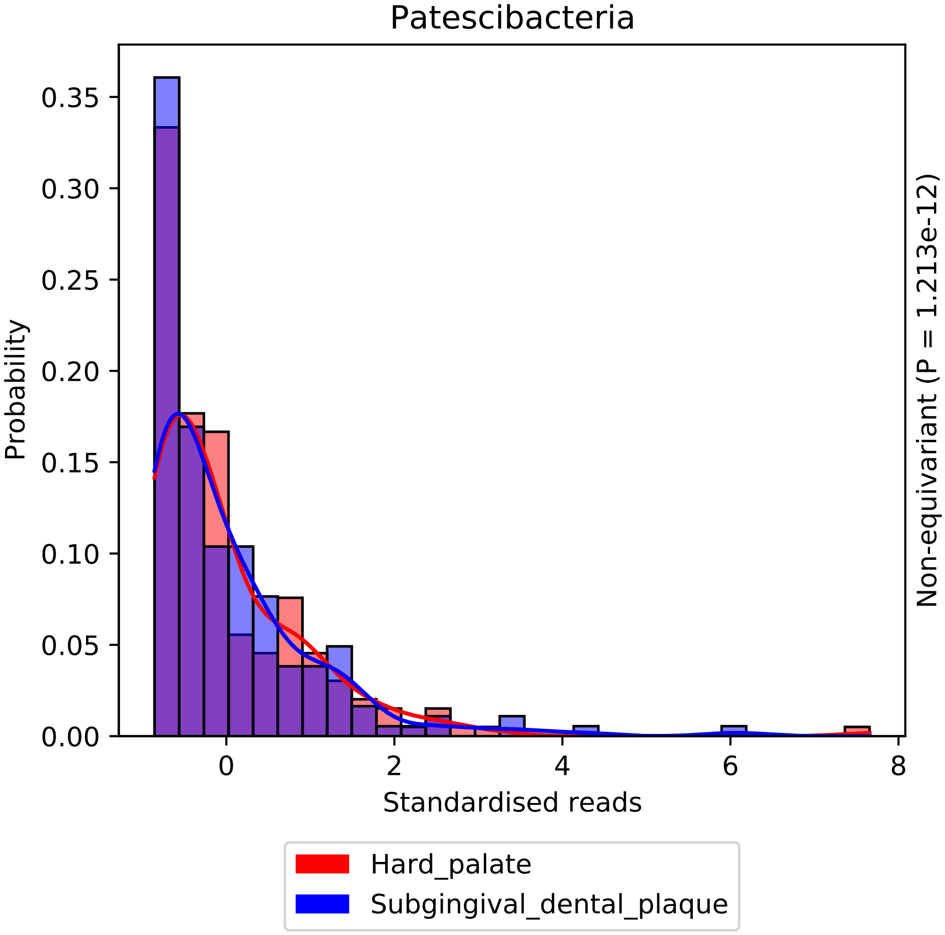
Patescibacteria
Probability
0.00
0.05
0.10
0.15
0.20
0.25
0.30
0.35
0
2
4
6
8
Standardised reads
Non-equivariant (P = 1.213e-12)
Hard_palate
Subgingival_dental_plaque

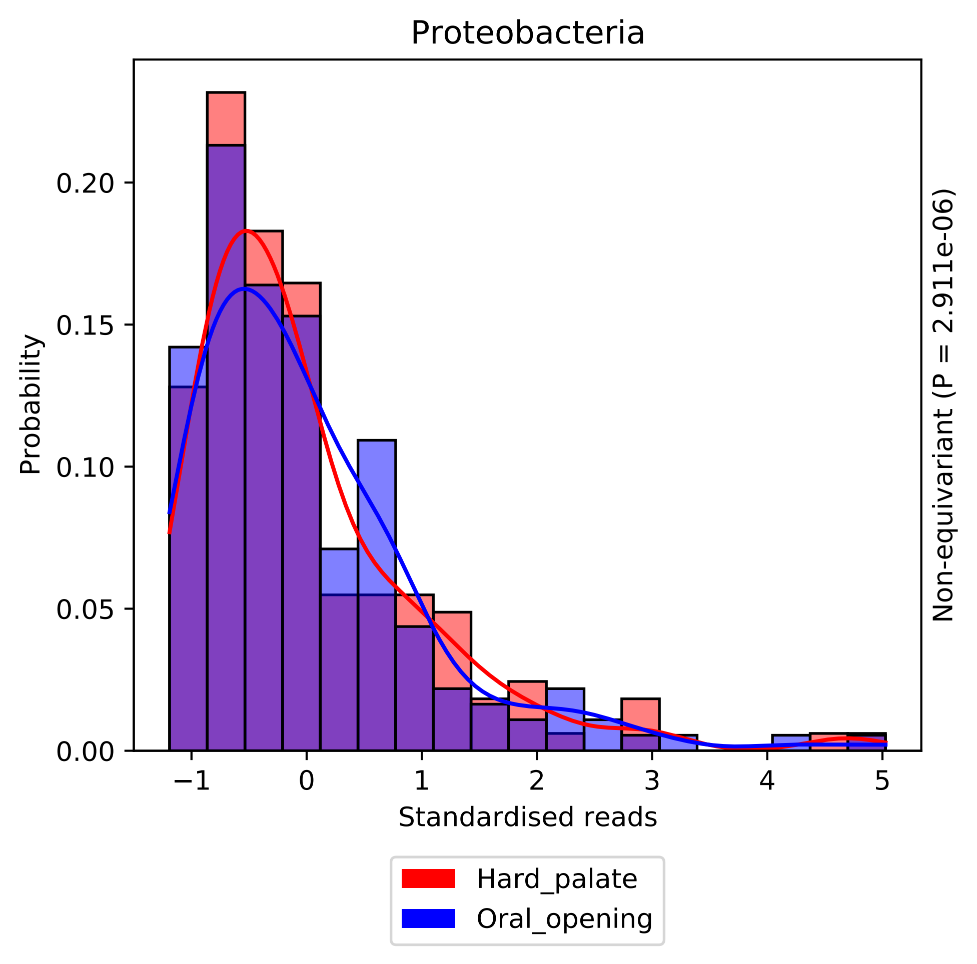

Proteobacteria
Probability
0.00
0.05
0.10
0.15
0.20
−1
0
1
2
3
4
5
Standardised reads
Non-equivariant (P = 2.911e-06)
Hard_palate
Oral_opening

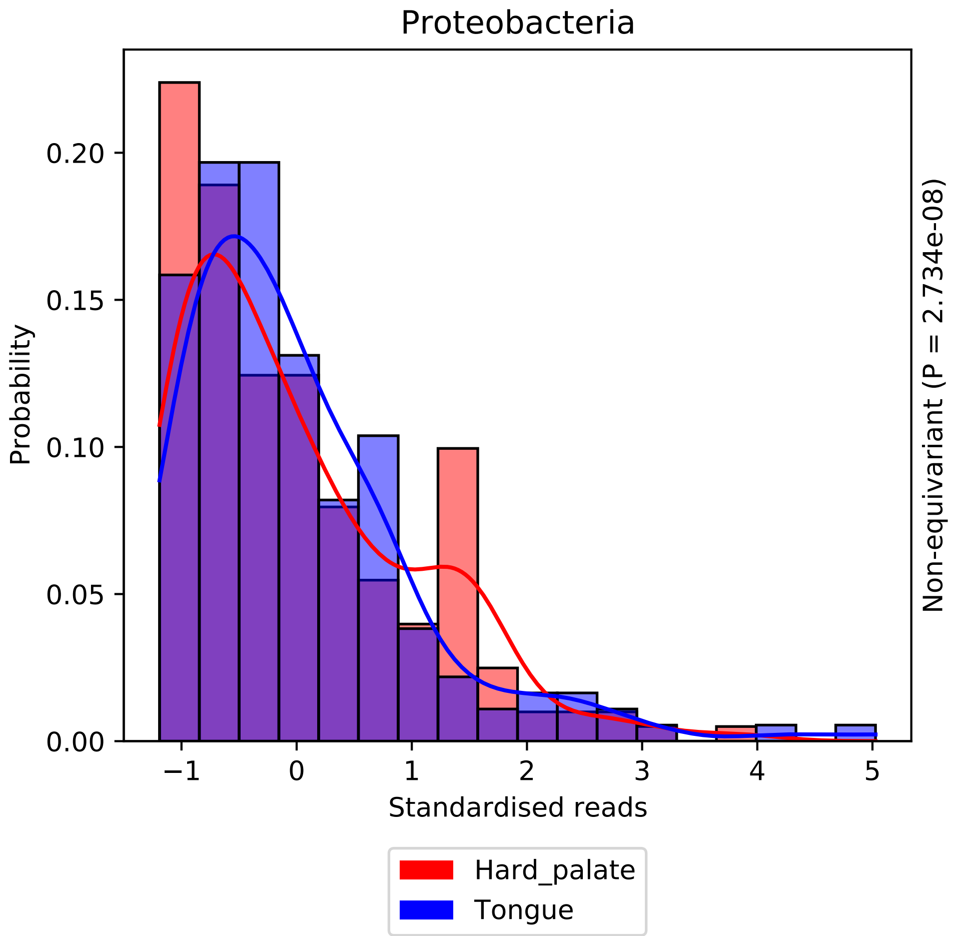

Proteobacteria
Probability
0.00
0.05
0.10
0.15
0.20
−1
0
1
2
3
4
5
Standardised reads
Non-equivariant (P = 2.734e-08)
Hard_palate
Tongue

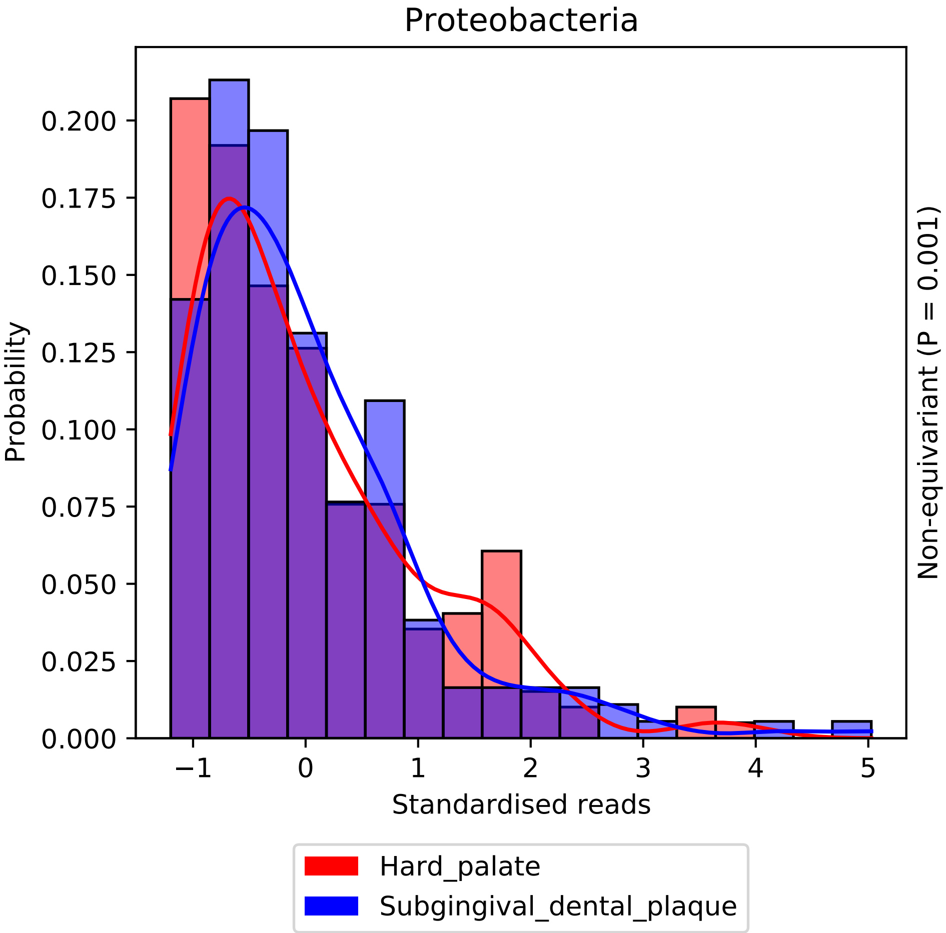
Proteobacteria
Probability
Standardised reads
Non-equivariant (P = 0.001)
0.000
0.025
0.050
0.075
0.100
0.125
0.150
0.175
0.200
−1
0
1
2
3
4
5
Hard_palate
Subgingival_dental_plaque

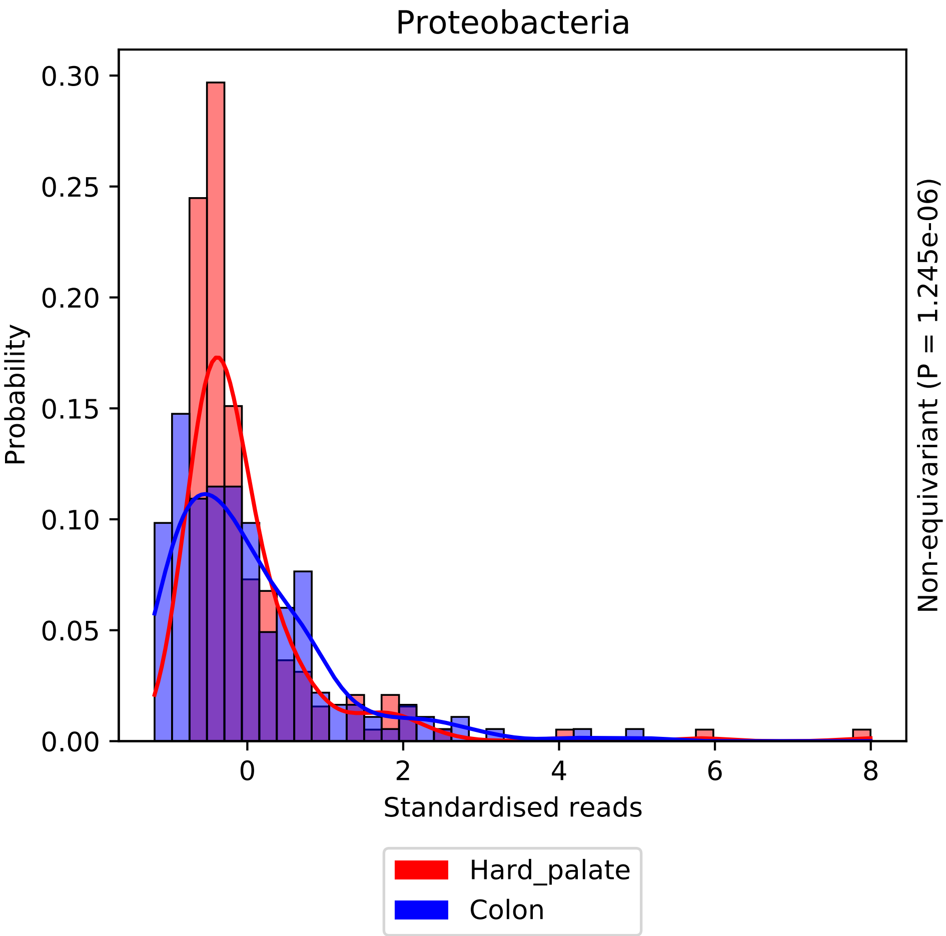
Proteobacteria
0.30
0.25
0.20
0.15
0.10
0.05
0.00
Probability
0
2
4
6
8
Standardised reads
Non-equivariant (P = 1.245e-06)
Hard_palate
Colon

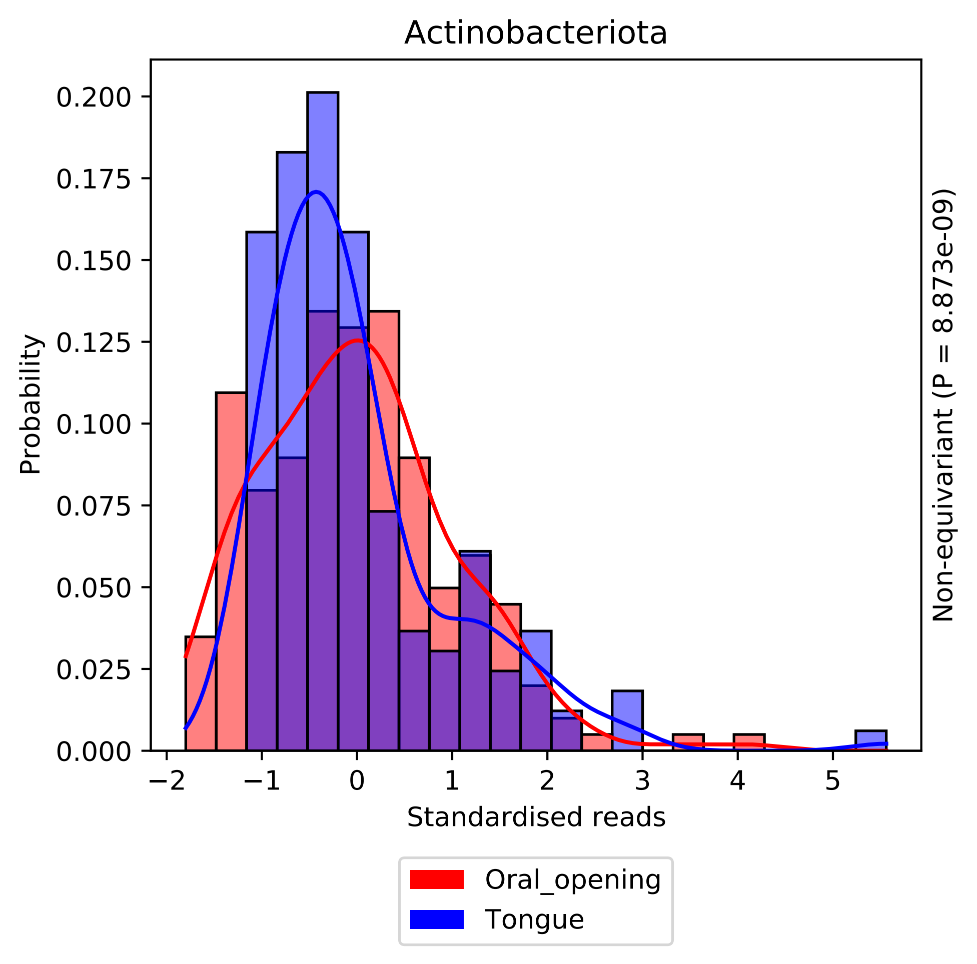

Actinobacteriota
Probability
0.200
0.175
0.150
0.125
0.100
0.075
0.050
0.025
0.000
−2
−1
0
1
2
3
4
5
Standardised reads
Non-equivariant (P = 8.873e-09)
Oral_opening
Tongue

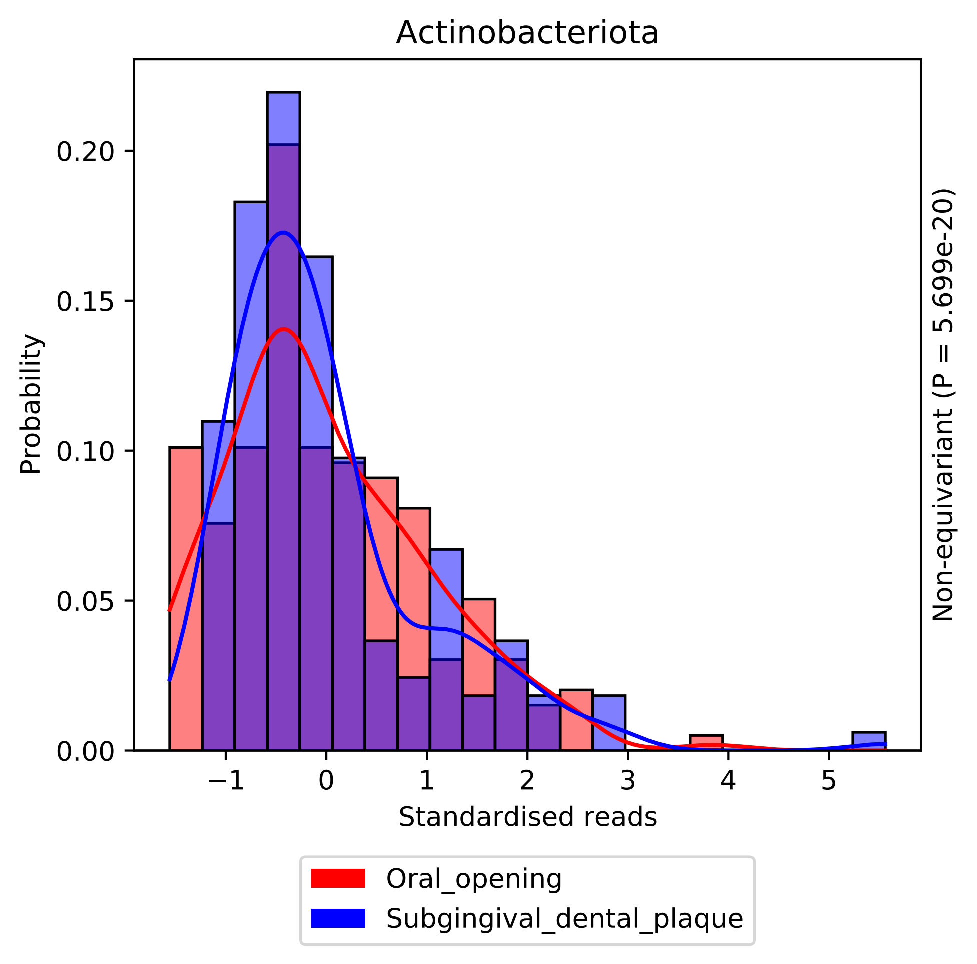

Actinobacteriota
0.00
0.05
0.10
0.15
0.20
Probability
−1
0
1
2
3
4
5
Standardised reads
Non-equivariant (P = 5.699e-20)
Oral_opening
Subgingival_dental_plaque

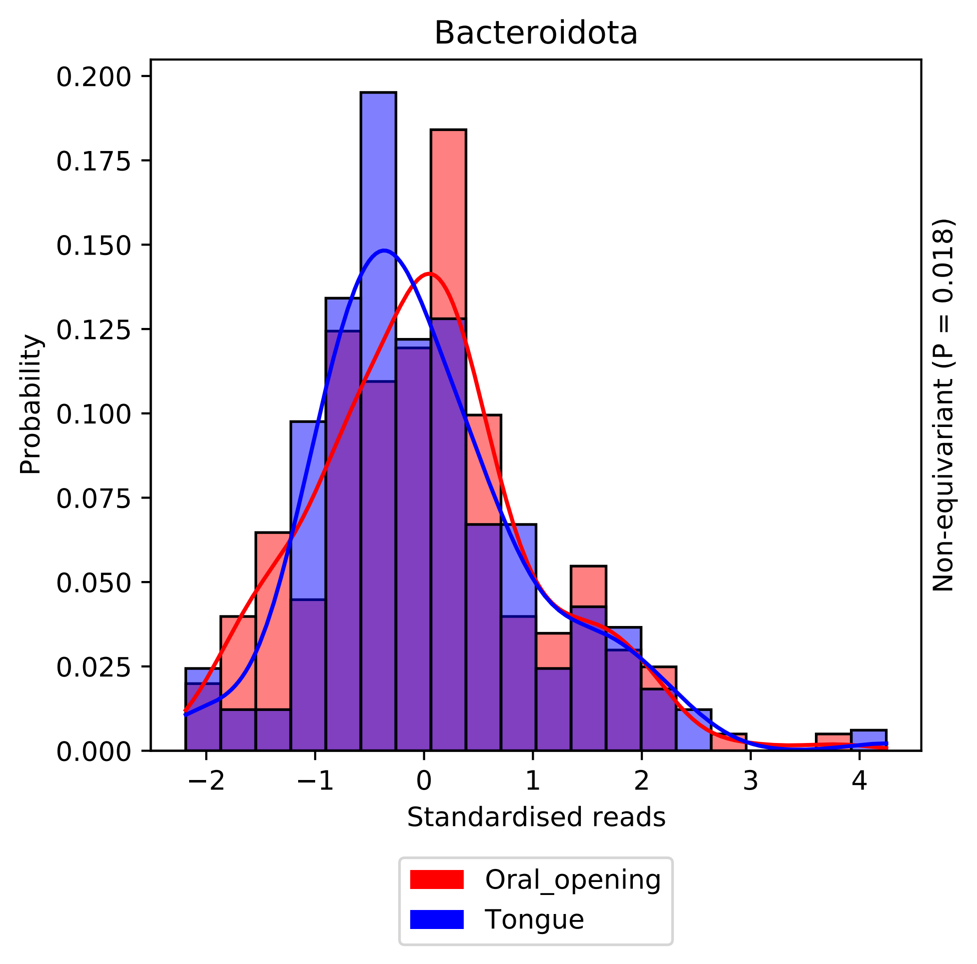

Bacteroidota
Probability
Standardised reads
Non-equivariant (P = 0.018)
Oral_opening
Tongue
0.000
0.025
0.050
0.075
0.100
0.125
0.150
0.175
0.200
−2
−1
0
1
2
3
4

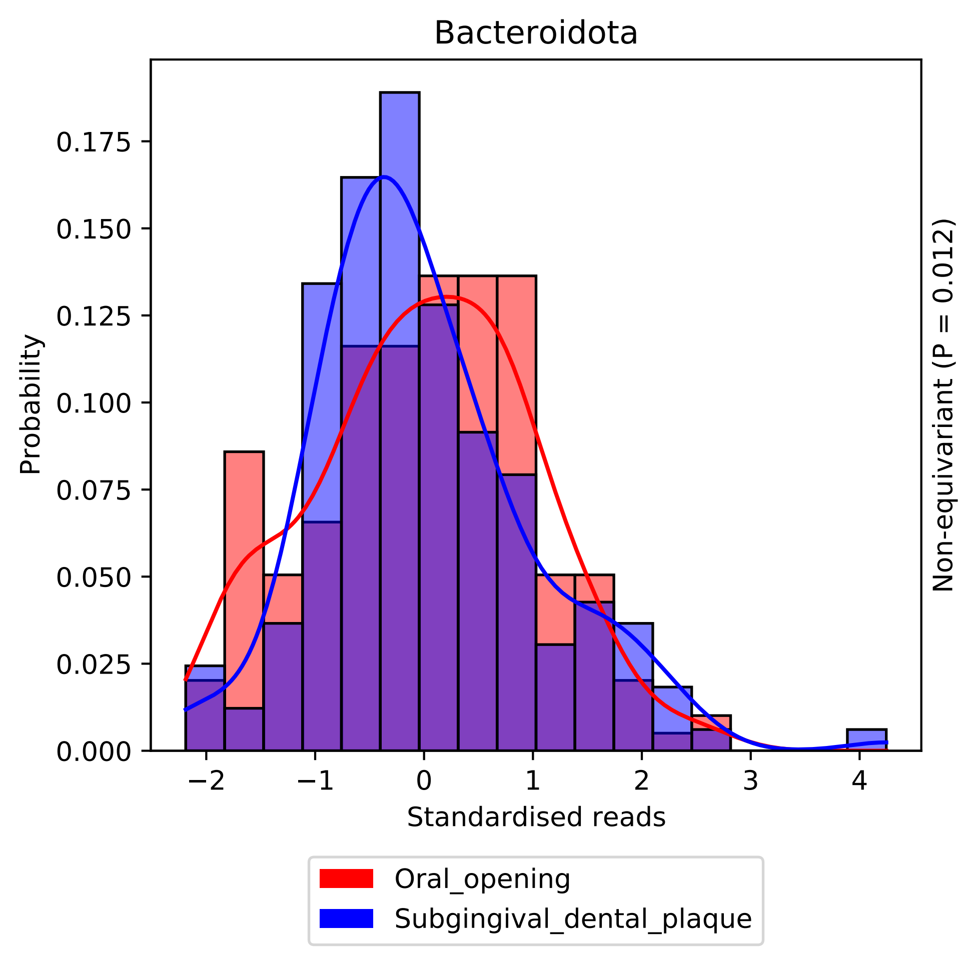
Bacteroidota
Probability
0.000
0.025
0.050
0.075
0.100
0.125
0.150
0.175
−2
−1
0
1
2
3
4
Standardised reads
Non-equivariant (P = 0.012)
Oral_opening
Subgingival_dental_plaque

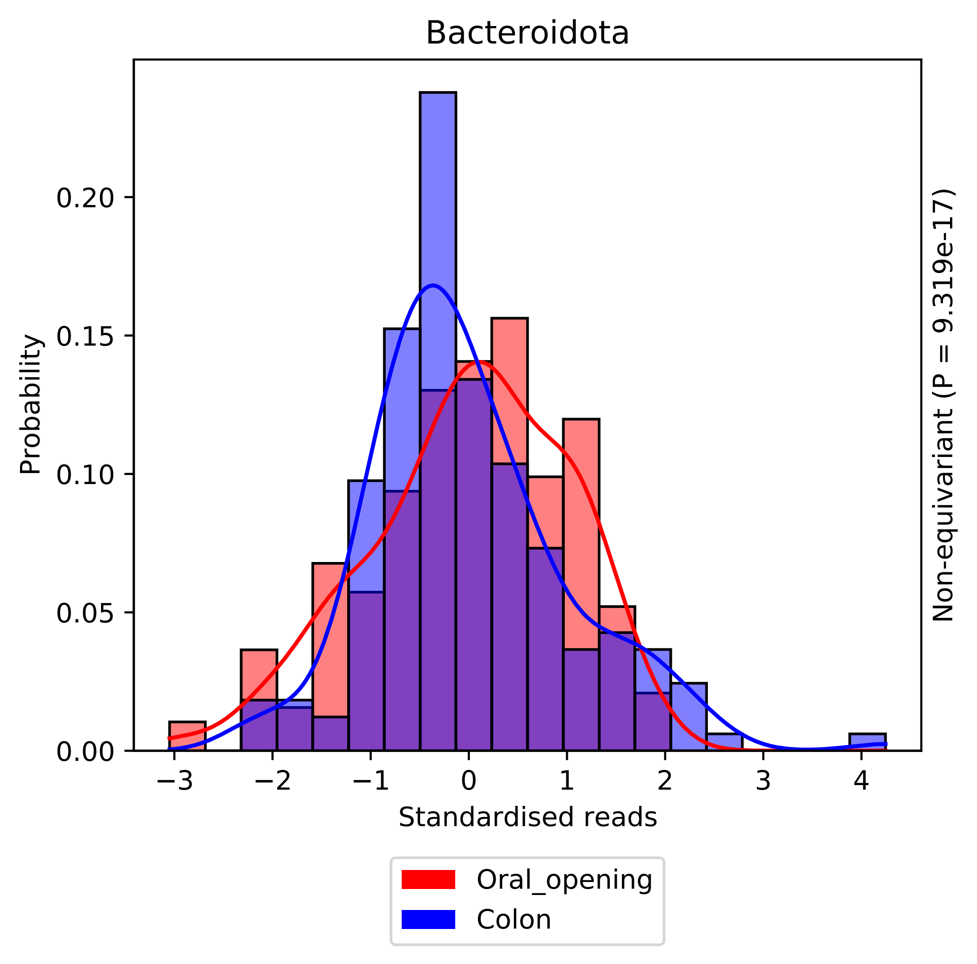

Bacteroidota
Probability
0.00
0.05
0.10
0.15
0.20
−3
−2
−1
0
1
2
3
4
Standardised reads
Non-equivariant (P = 9.319e-17)
Oral_opening
Colon

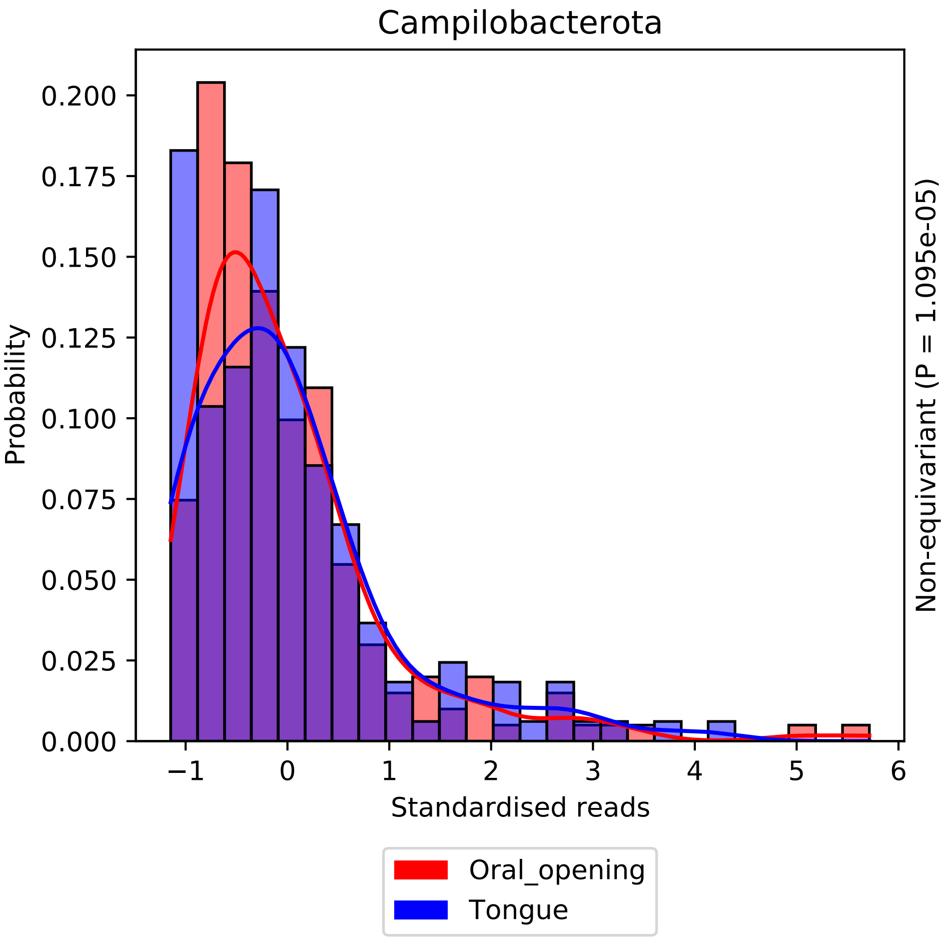
Campilobacterota
Probability
Standardised reads
Non-equivariant (P = 1.095e-05)
Oral_opening
Tongue
0.000
0.025
0.050
0.075
0.100
0.125
0.150
0.175
0.200
−1
0
1
2
3
4
5
6

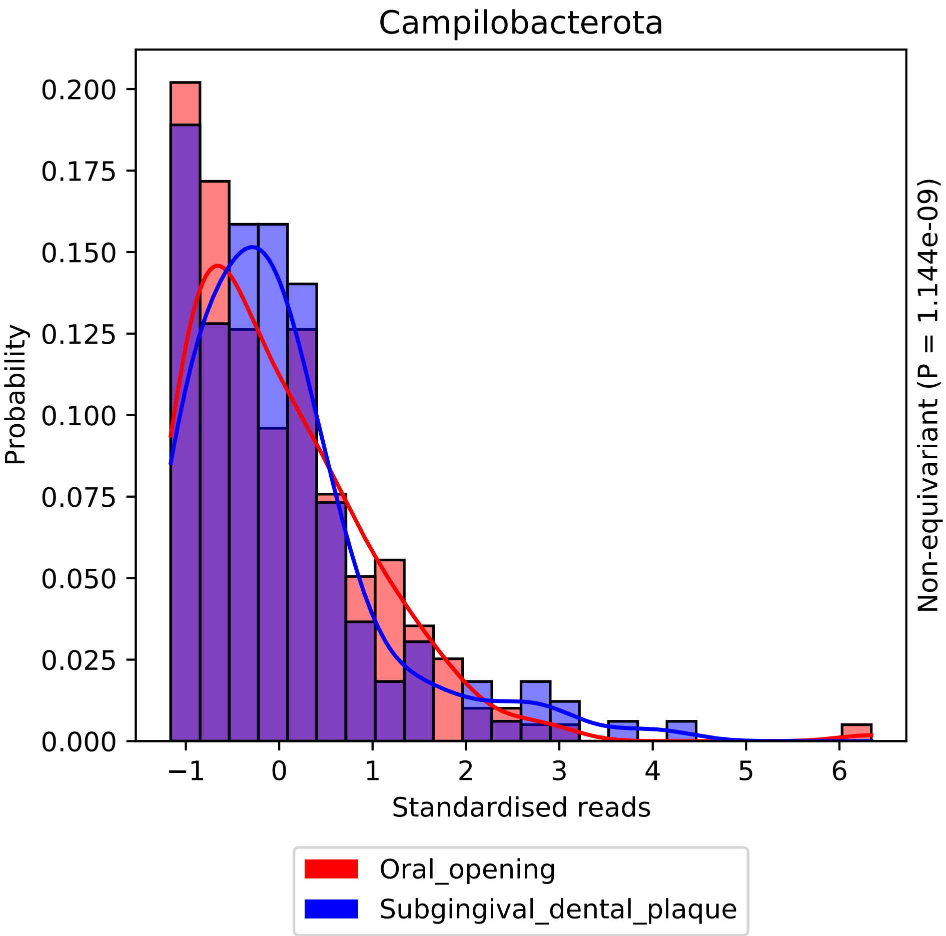
Campilobacterota
Probability
0.200
0.175
0.150
0.125
0.100
0.075
0.050
0.025
0.000
−1
0
1
2
3
4
5
6
Standardised reads
Non-equivariant (P = 1.144e-09)
Oral_opening
Subgingival_dental_plaque

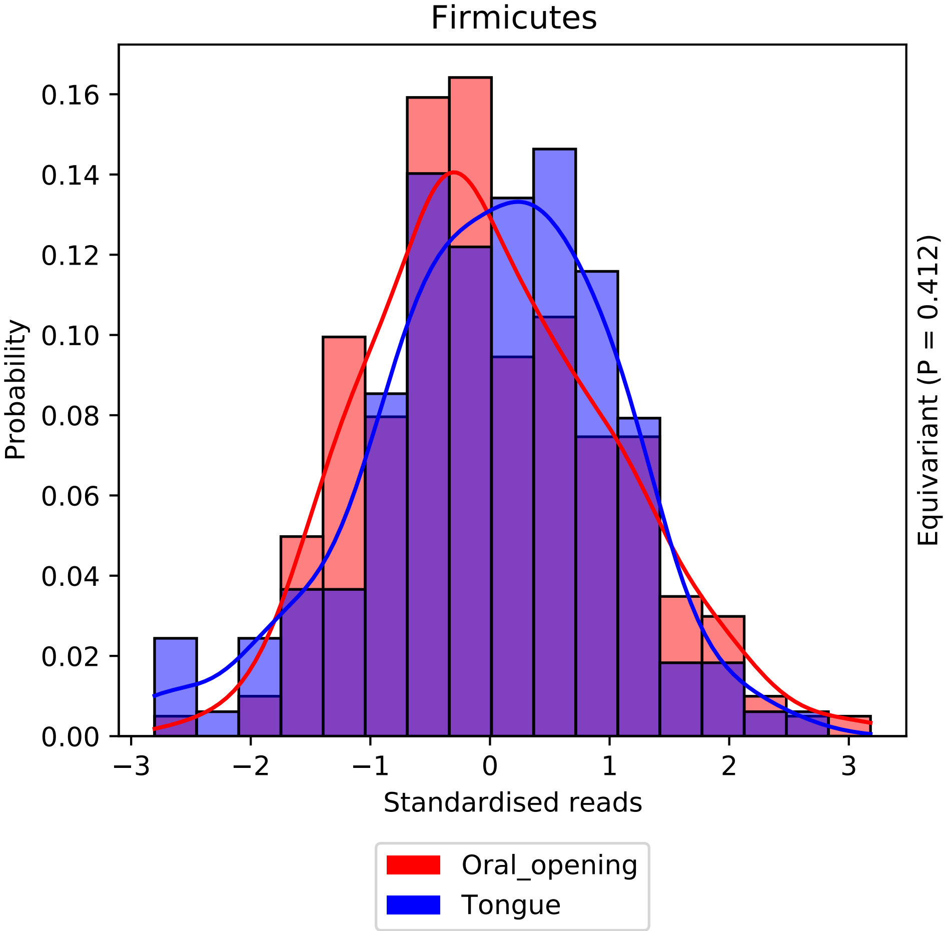

Firmicutes
Probability
0.00
0.02
0.04
0.06
0.08
0.10
0.12
0.14
0.16
−3
−2
−1
0
1
2
3
Standardised reads
Equivariant (P = 0.412)
Oral_opening
Tongue

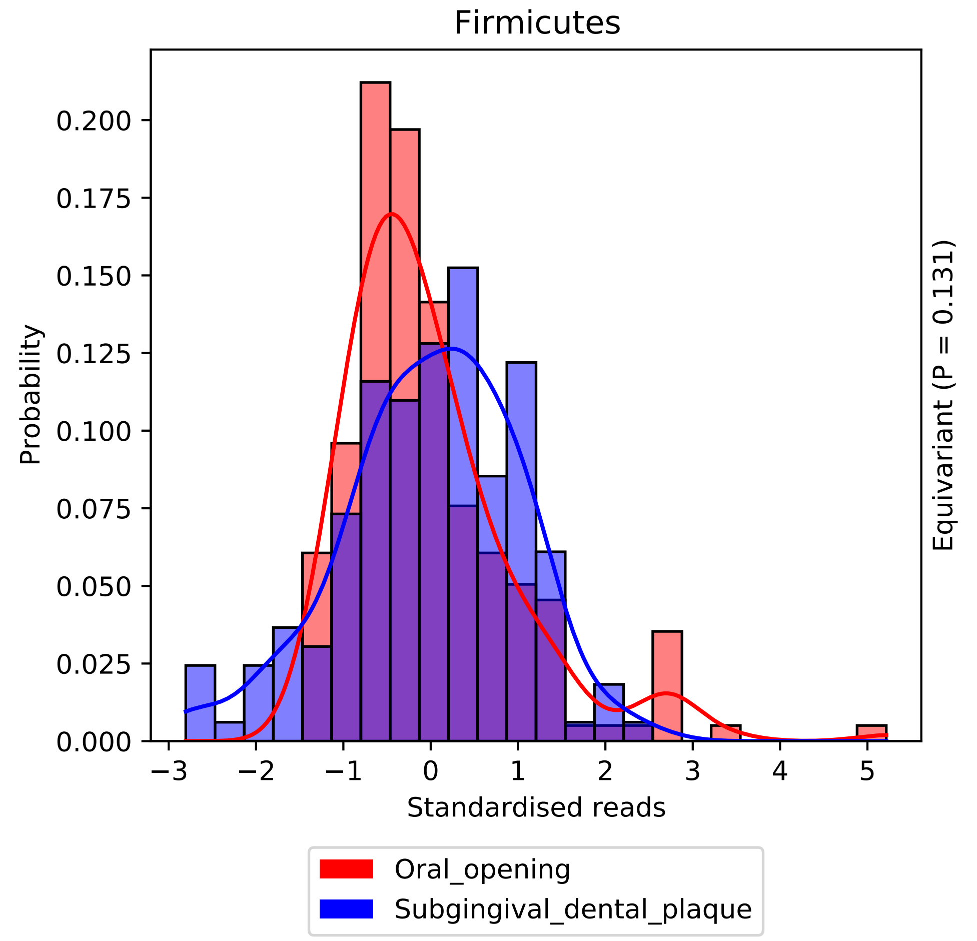

Firmicutes
0.200
0.175
0.150
0.125
0.100
0.075
0.050
0.025
0.000
Probability
−3
−2
−1
0
1
2
3
4
5
Standardised reads
Equivariant (P = 0.131)
Oral_opening
Subgingival_dental_plaque

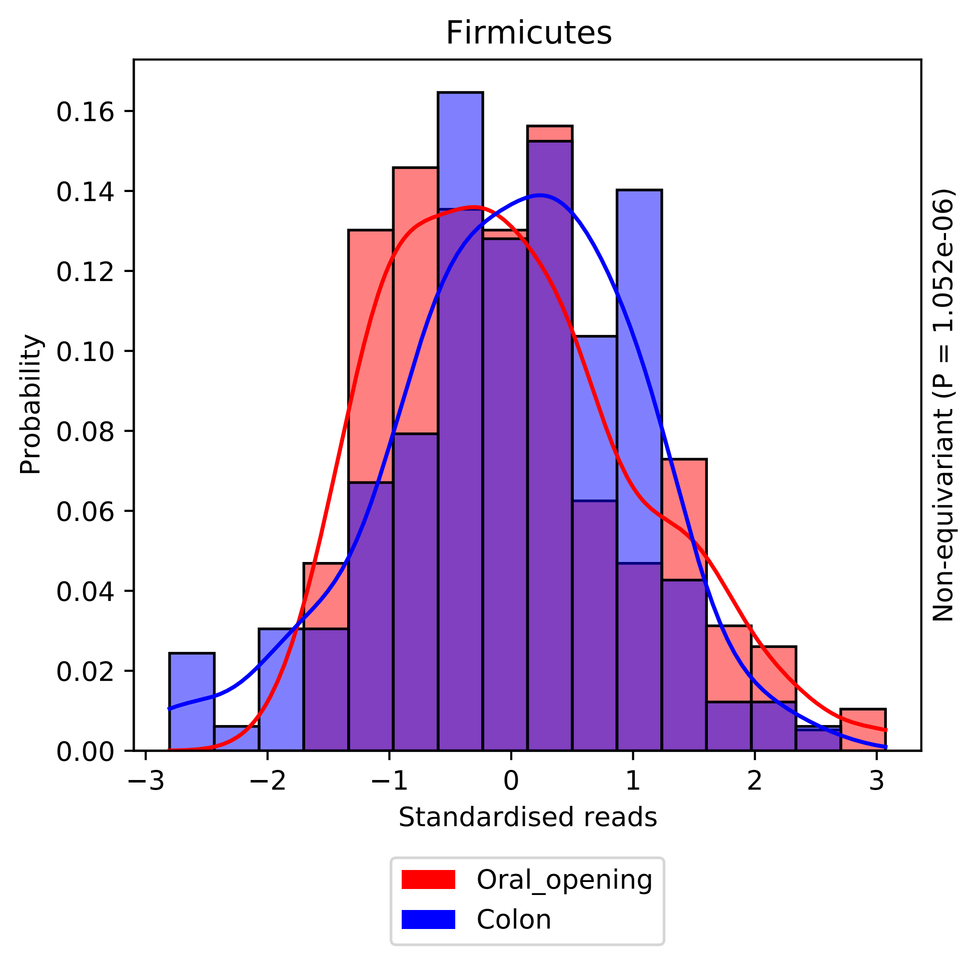

Firmicutes
Probability
Standardised reads
Non-equivariant (P = 1.052e-06)
0.00
0.02
0.04
0.06
0.08
0.10
0.12
0.14
0.16
−3
−2
−1
0
1
2
3
Oral_opening
Colon

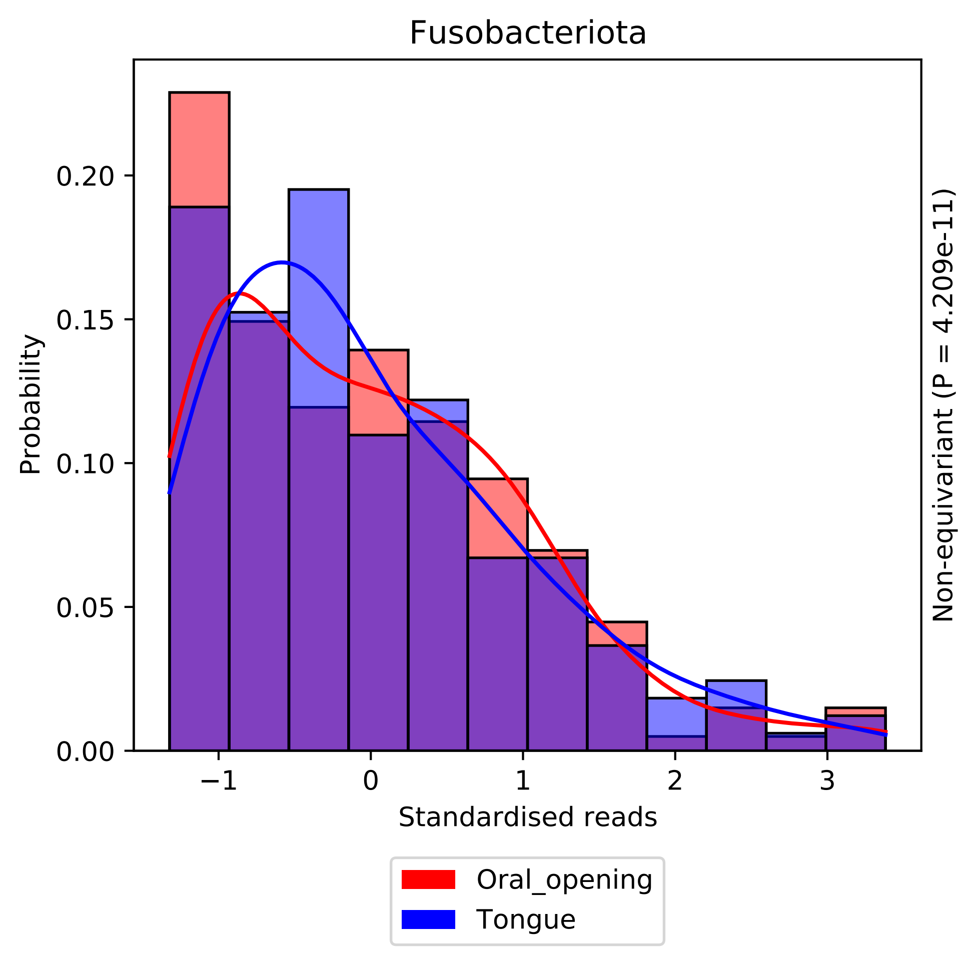

Fusobacteriota
Probability
0.00
0.05
0.10
0.15
0.20
−1
0
1
2
3
Standardised reads
Non-equivariant (P = 4.209e-11)
Oral_opening
Tongue

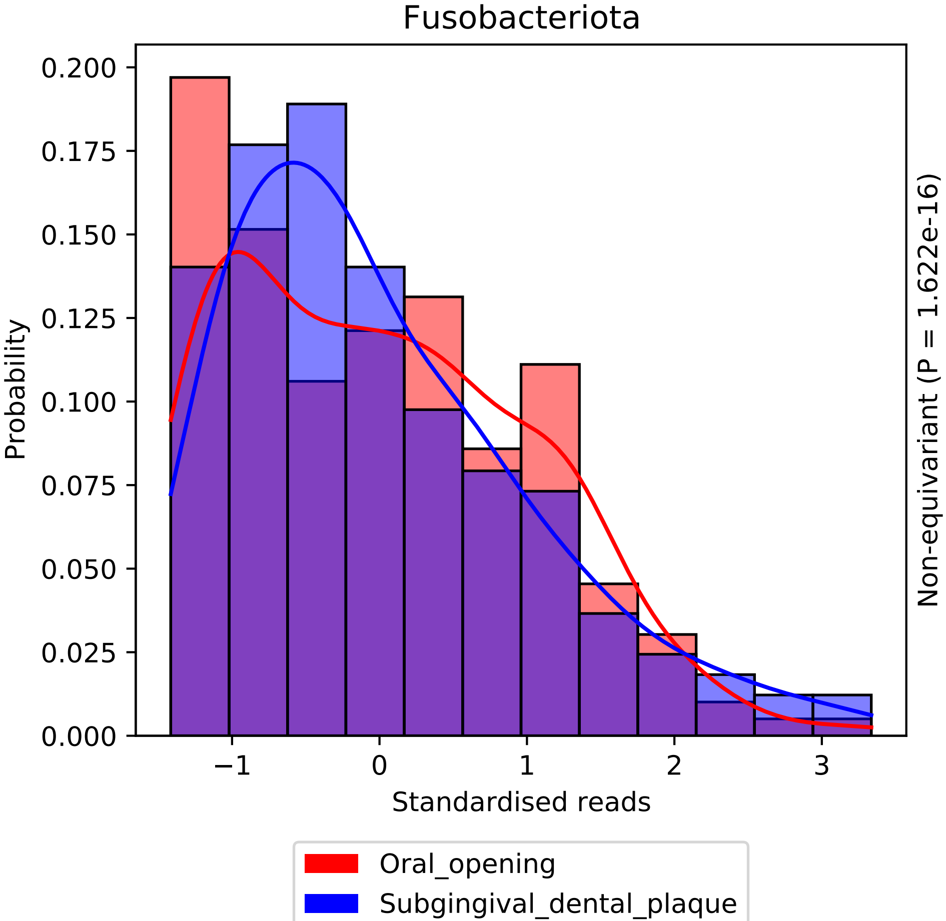
Fusobacteriota
Probability
Standardised reads
Non-equivariant (P = 1.622e-16)
0.000
0.025
0.050
0.075
0.100
0.125
0.150
0.175
0.200
−1
0
1
2
3
Oral_opening
Subgingival_dental_plaque

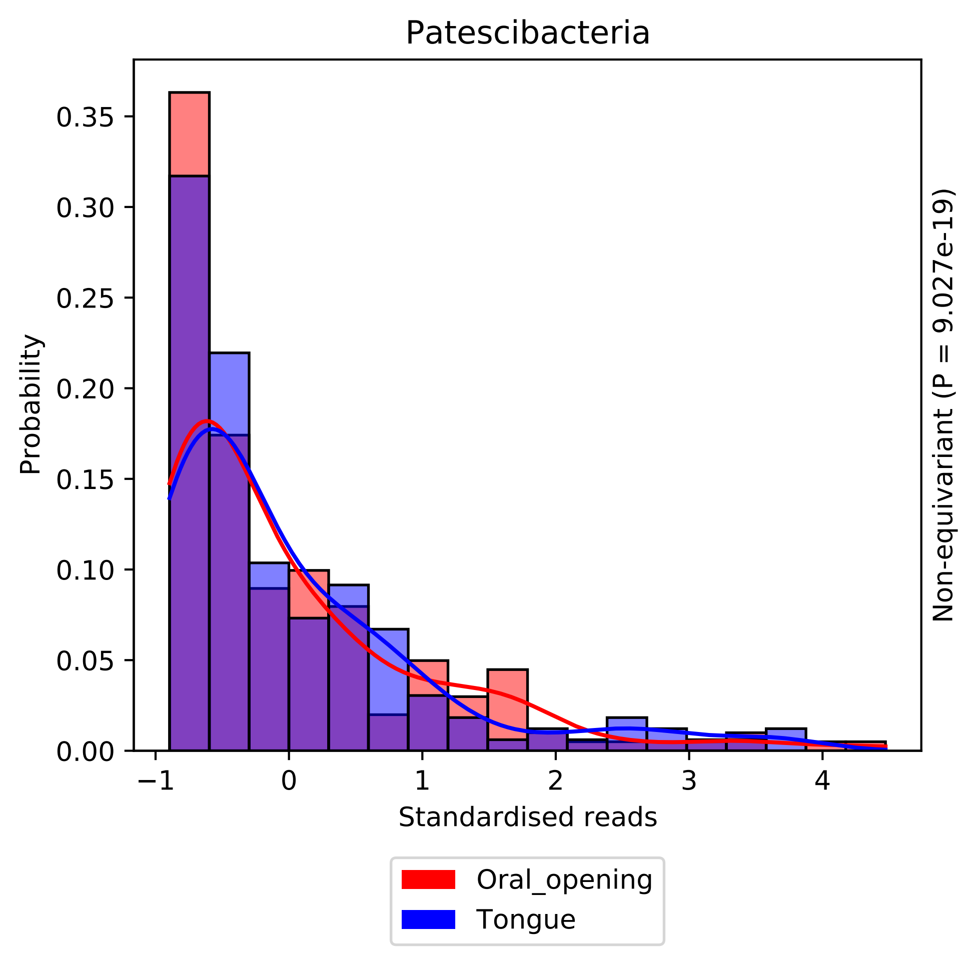

Patescibacteria
Probability
0.00
0.05
0.10
0.15
0.20
0.25
0.30
0.35
−1
0
1
2
3
4
Standardised reads
Non-equivariant (P = 9.027e-19)
Oral_opening
Tongue

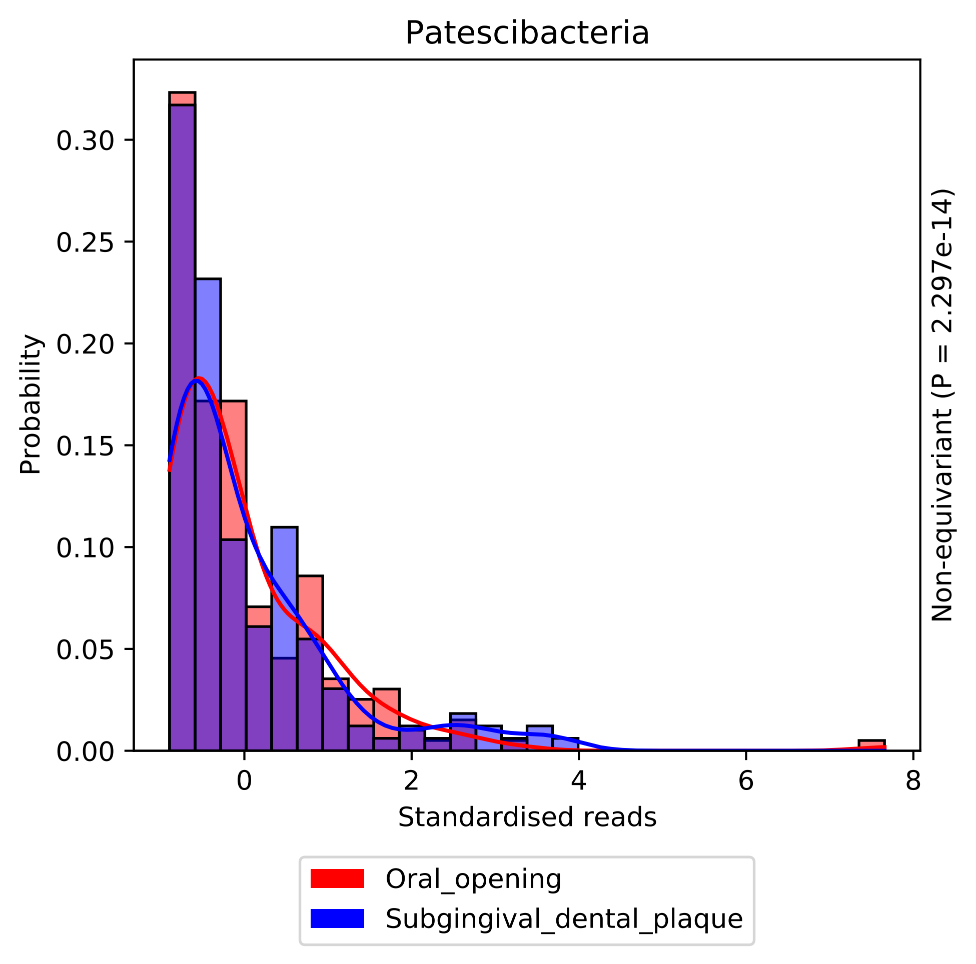

Patescibacteria
Probability
Standardised reads
Non-equivariant (P = 2.297e-14)
0.00
0.05
0.10
0.15
0.20
0.25
0.30
0
2
4
6
8
Oral_opening
Subgingival_dental_plaque

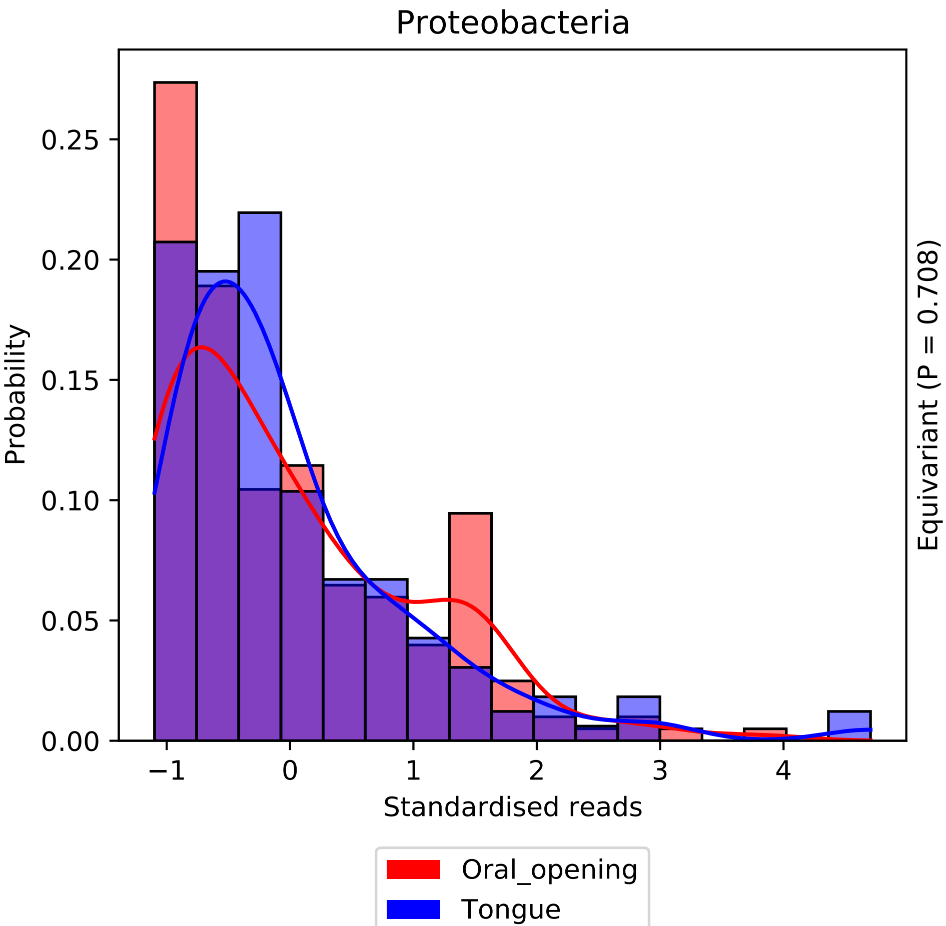

Proteobacteria
Probability
Standardised reads
Equivariant (P = 0.708)
0.00
0.05
0.10
0.15
0.20
0.25
−1
0
1
2
3
4
Oral_opening
Tongue

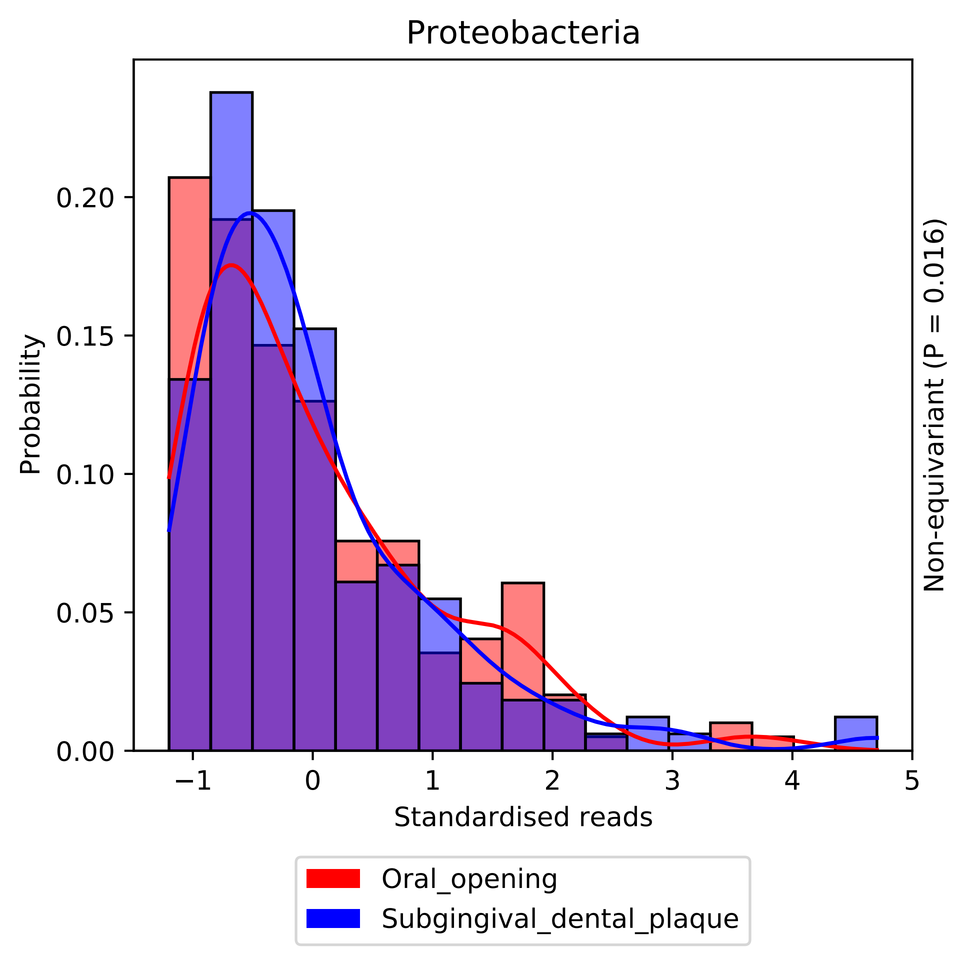

Proteobacteria
Probability
0.00
0.05
0.10
0.15
0.20
−1
0
1
2
3
4
5
Standardised reads
Non-equivariant (P = 0.016)
Oral_opening
Subgingival_dental_plaque

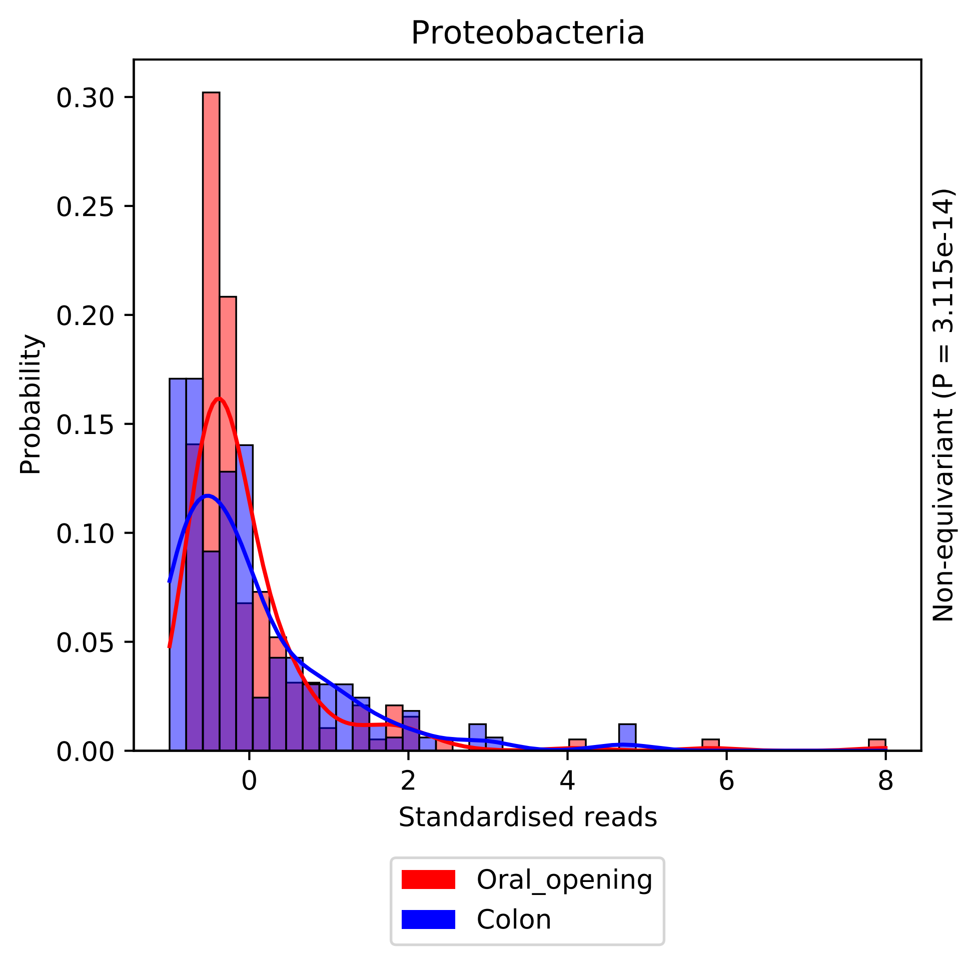
Proteobacteria
Probability
0.30
0.25
0.20
0.15
0.10
0.05
0.00
0
2
4
6
8
Standardised reads
Non-equivariant (P = 3.115e-14)
Oral_opening
Colon

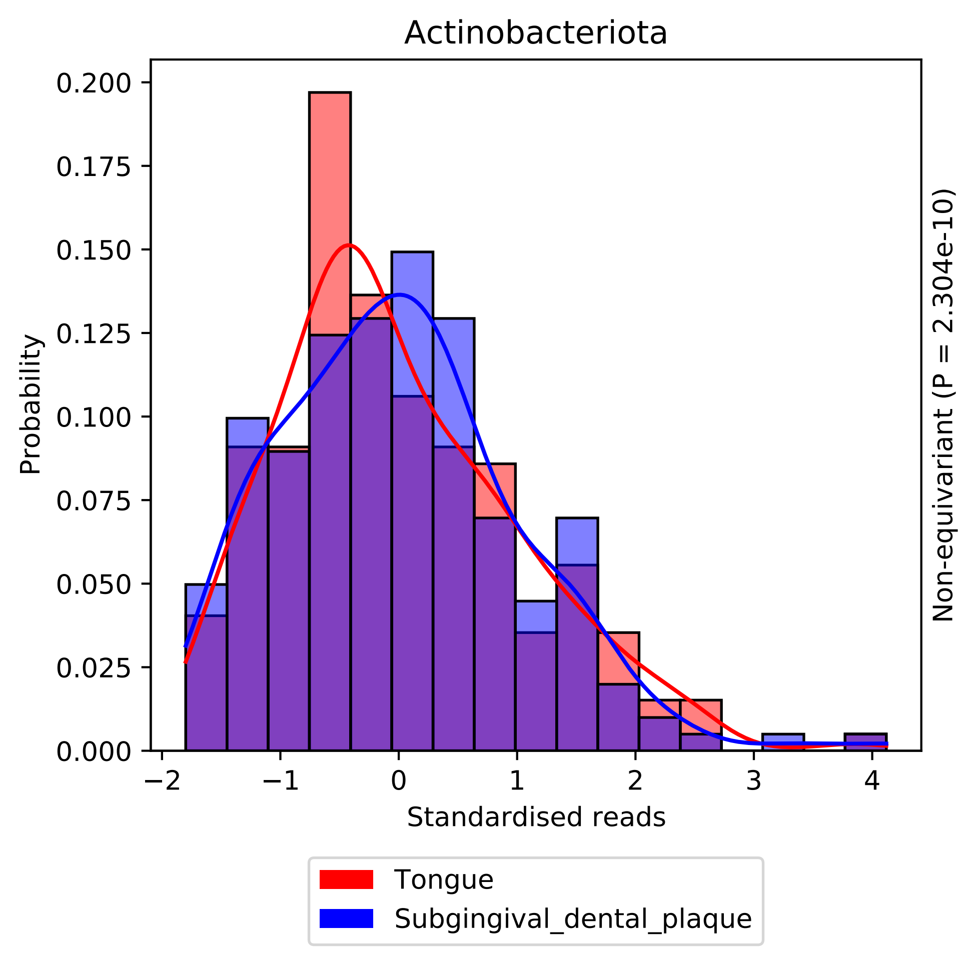

Actinobacteriota
Probability
0.200
0.175
0.150
0.125
0.100
0.075
0.050
0.025
0.000
−2
−1
0
1
2
3
4
Standardised reads
Non-equivariant (P = 2.304e-10)
Tongue
Subgingival_dental_plaque

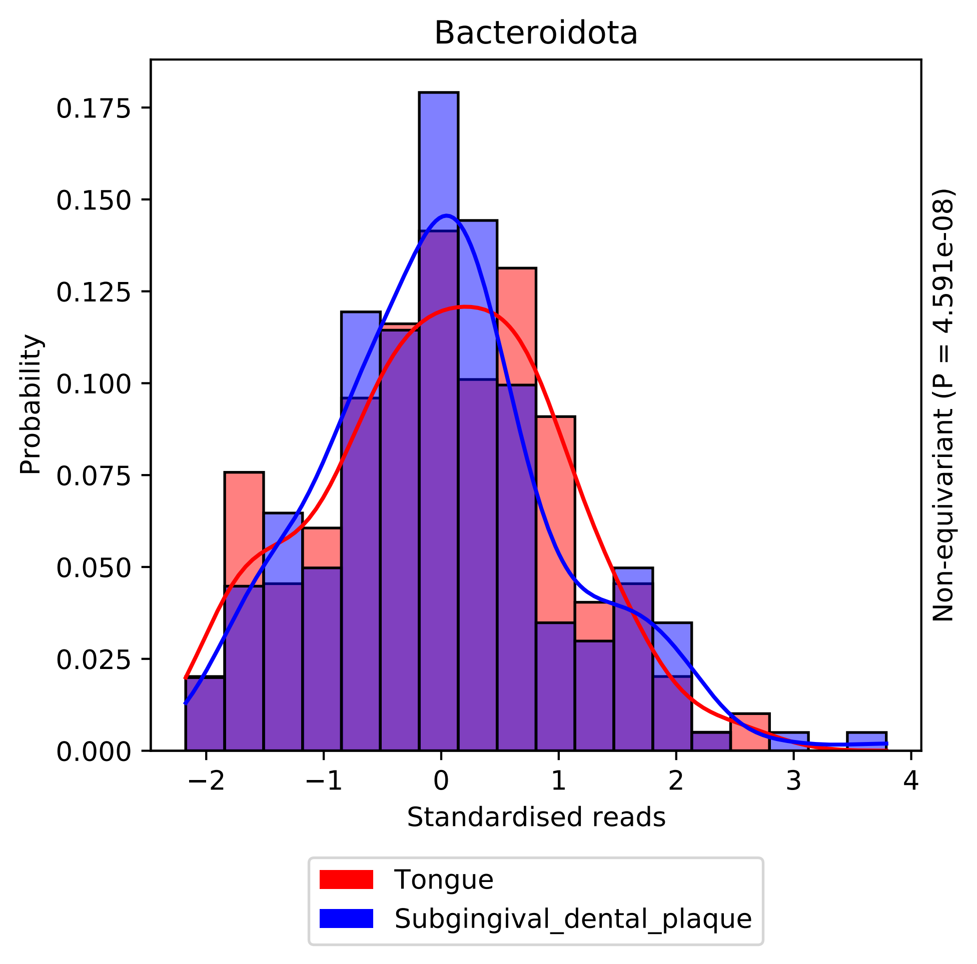

Bacteroidota
Probability
0.000
0.025
0.050
0.075
0.100
0.125
0.150
0.175
−2
−1
0
1
2
3
4
Standardised reads
Non-equivariant (P = 4.591e-08)
Tongue
Subgingival_dental_plaque

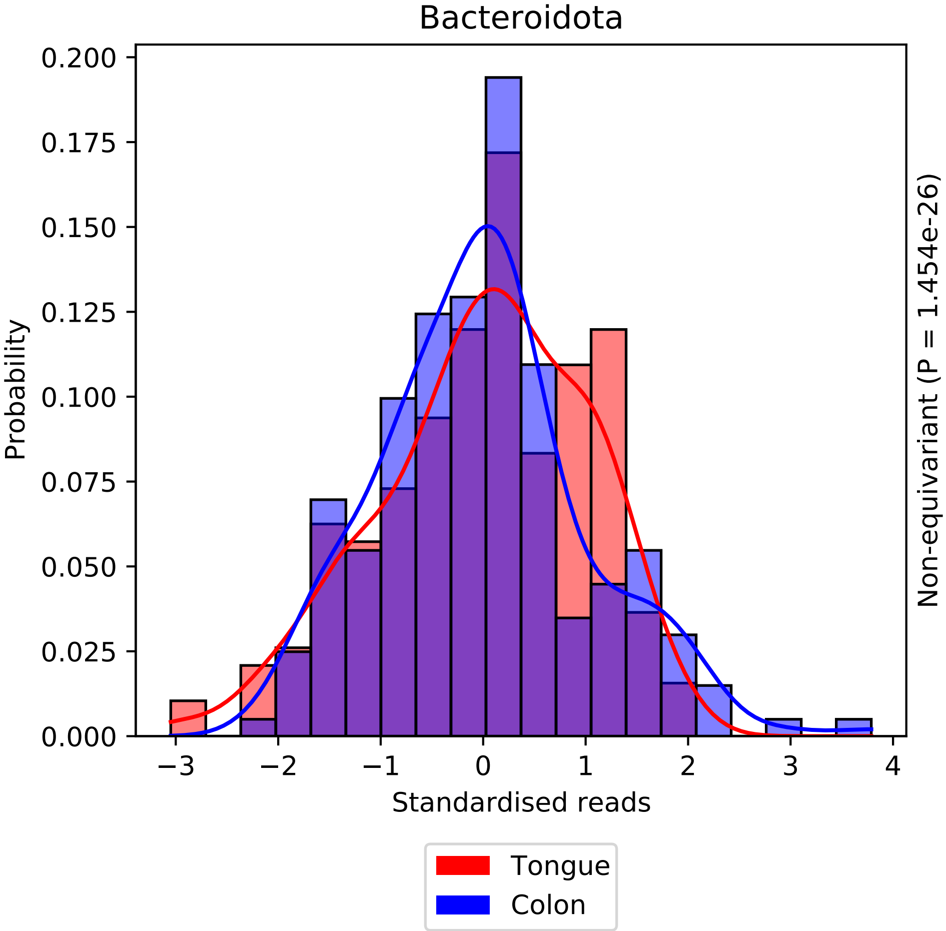
Bacteroidota
Probability
Standardised reads
Non-equivariant (P = 1.454e-26)
Tongue
Colon
0.200
0.175
0.150
0.125
0.100
0.075
0.050
0.025
0.000
−3
−2
−1
0
1
2
3
4

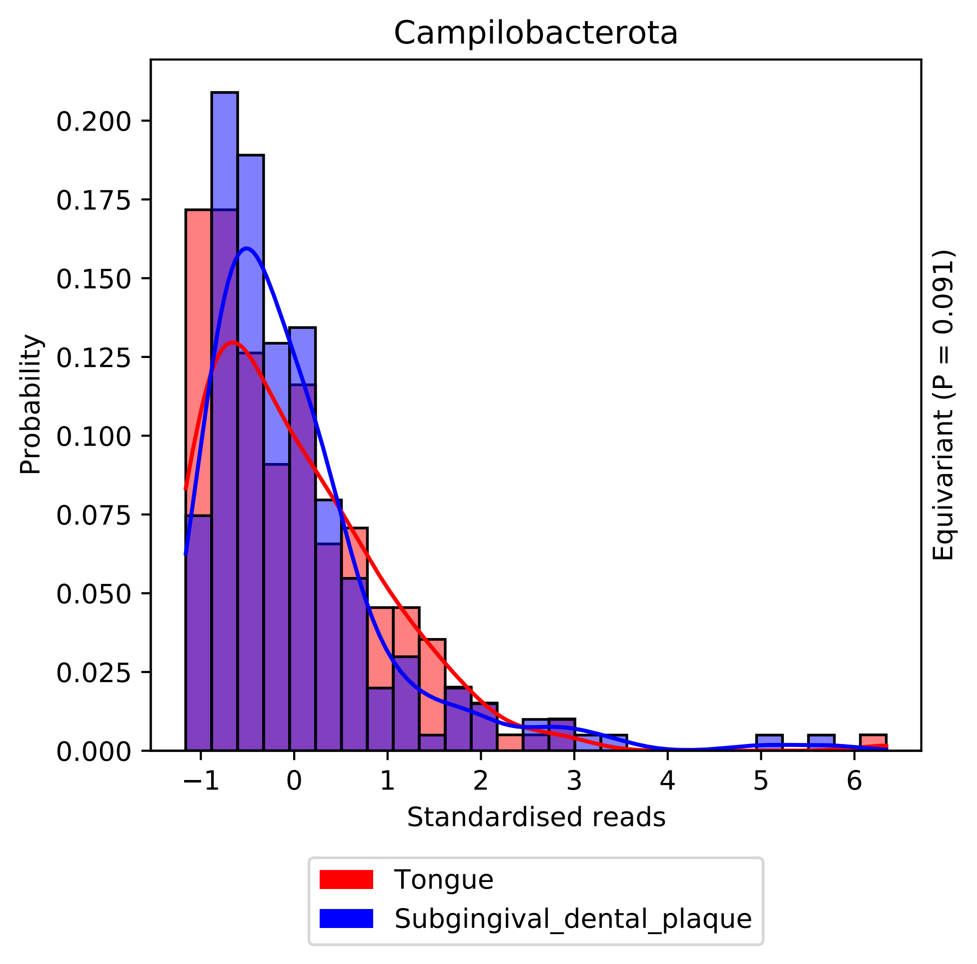
Campilobacterota
Probability
Standardised reads
Equivariant (P = 0.091)
0.000
0.025
0.050
0.075
0.100
0.125
0.150
0.175
0.200
−1
0
1
2
3
4
5
6
Tongue
Subgingival_dental_plaque

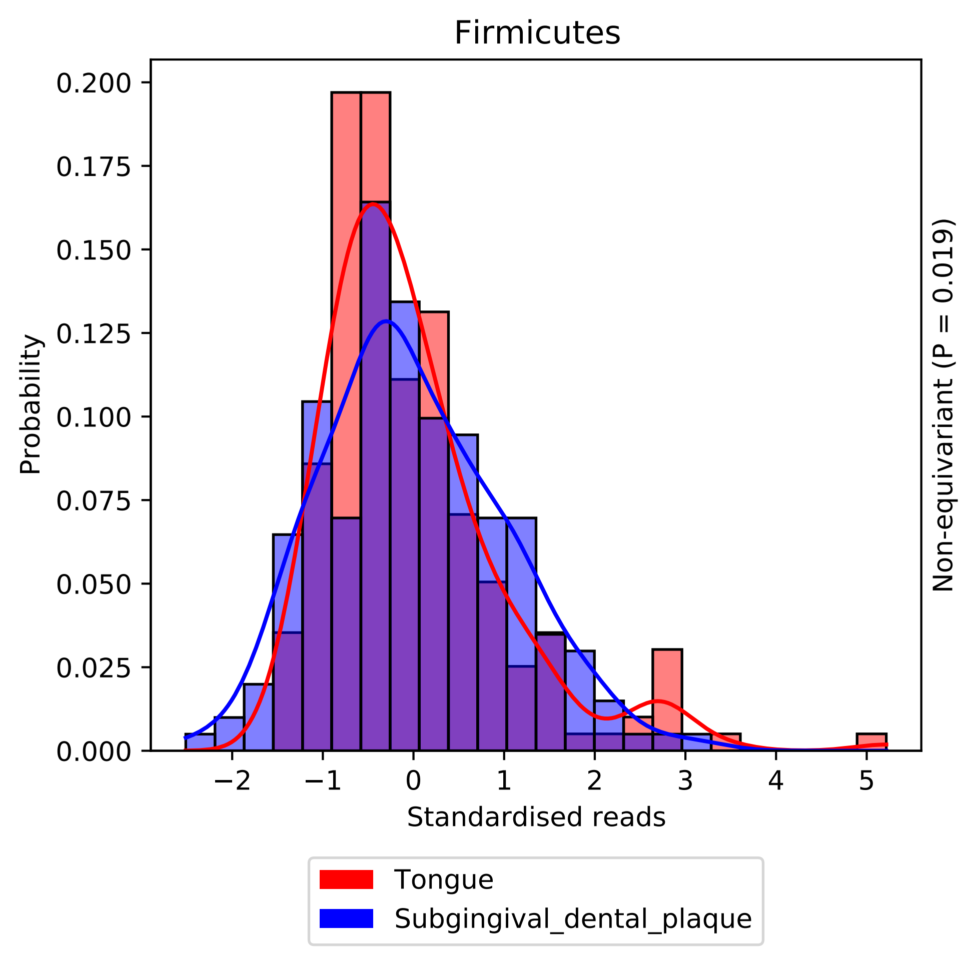

Firmicutes
Probability
0.200
0.175
0.150
0.125
0.100
0.075
0.050
0.025
0.000
−2
−1
0
1
2
3
4
5
Standardised reads
Non-equivariant (P = 0.019)
Tongue
Subgingival_dental_plaque

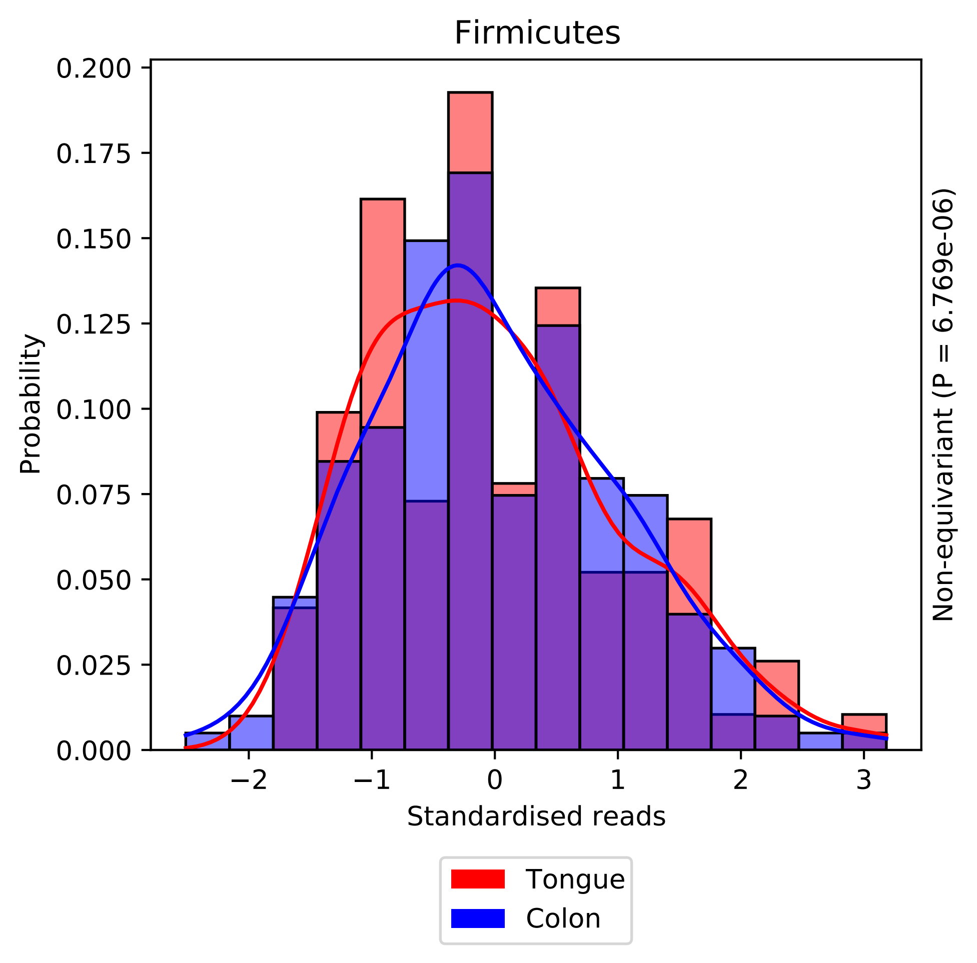
Firmicutes
Probability
0.200
0.175
0.150
0.125
0.100
0.075
0.050
0.025
0.000
−2
−1
0
1
2
3
Standardised reads
Non-equivariant (P = 6.769e-06)
Tongue
Colon

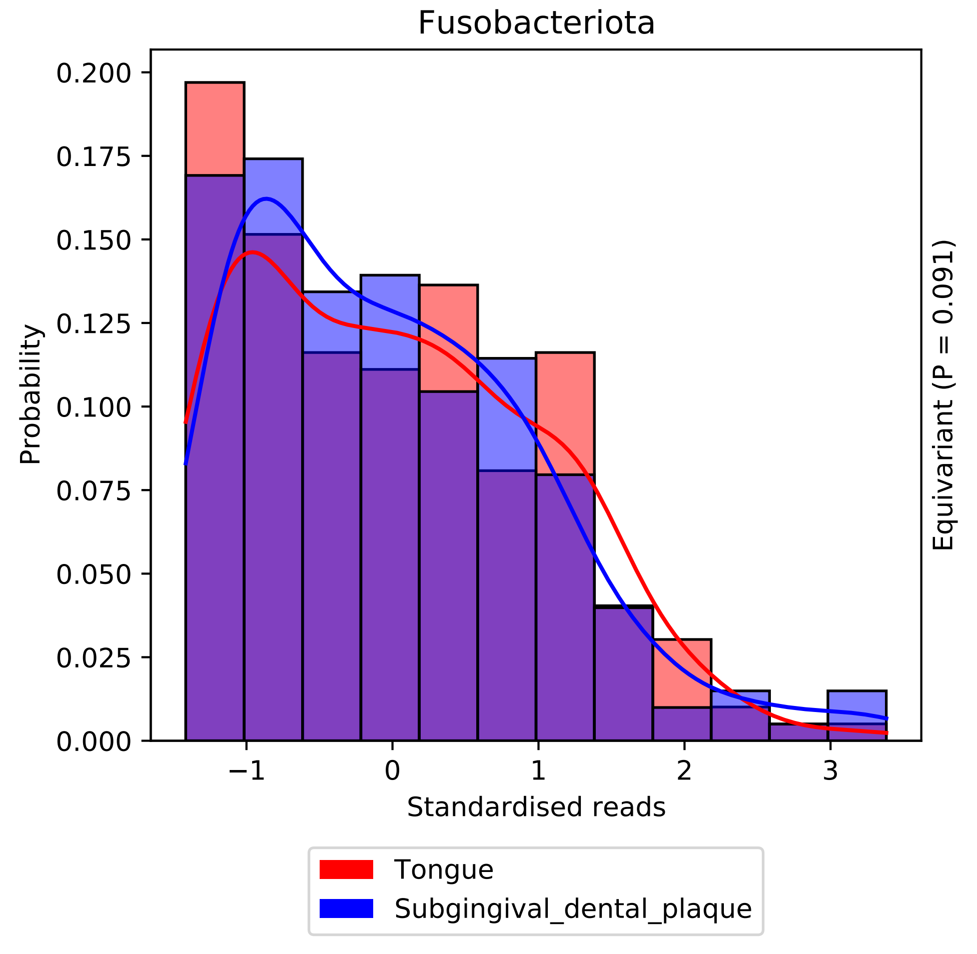

Fusobacteriota
Probability
0.200
0.175
0.150
0.125
0.100
0.075
0.050
0.025
0.000
−1
0
1
2
3
Standardised reads
Equivariant (P = 0.091)
Tongue
Subgingival_dental_plaque

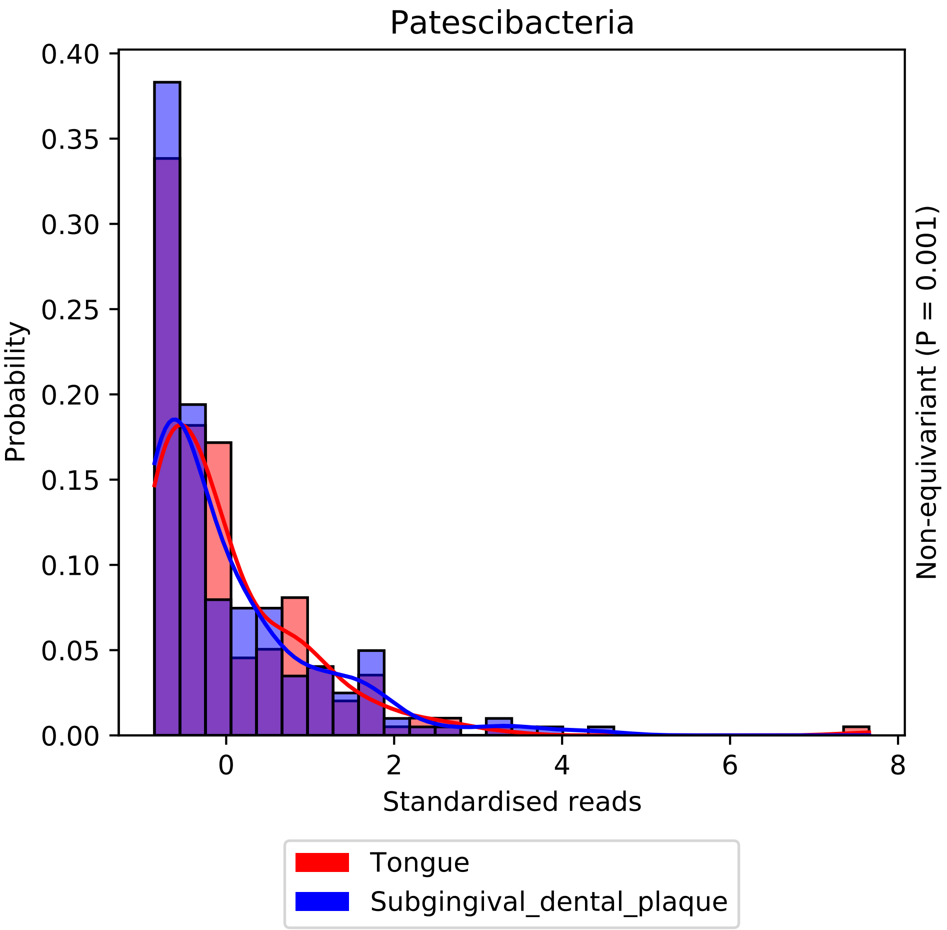

Patescibacteria
Probability
0.00
0.05
0.10
0.15
0.20
0.25
0.30
0.35
0.40
0
2
4
6
8
Standardised reads
Non-equivariant (P = 0.001)
Tongue
Subgingival_dental_plaque

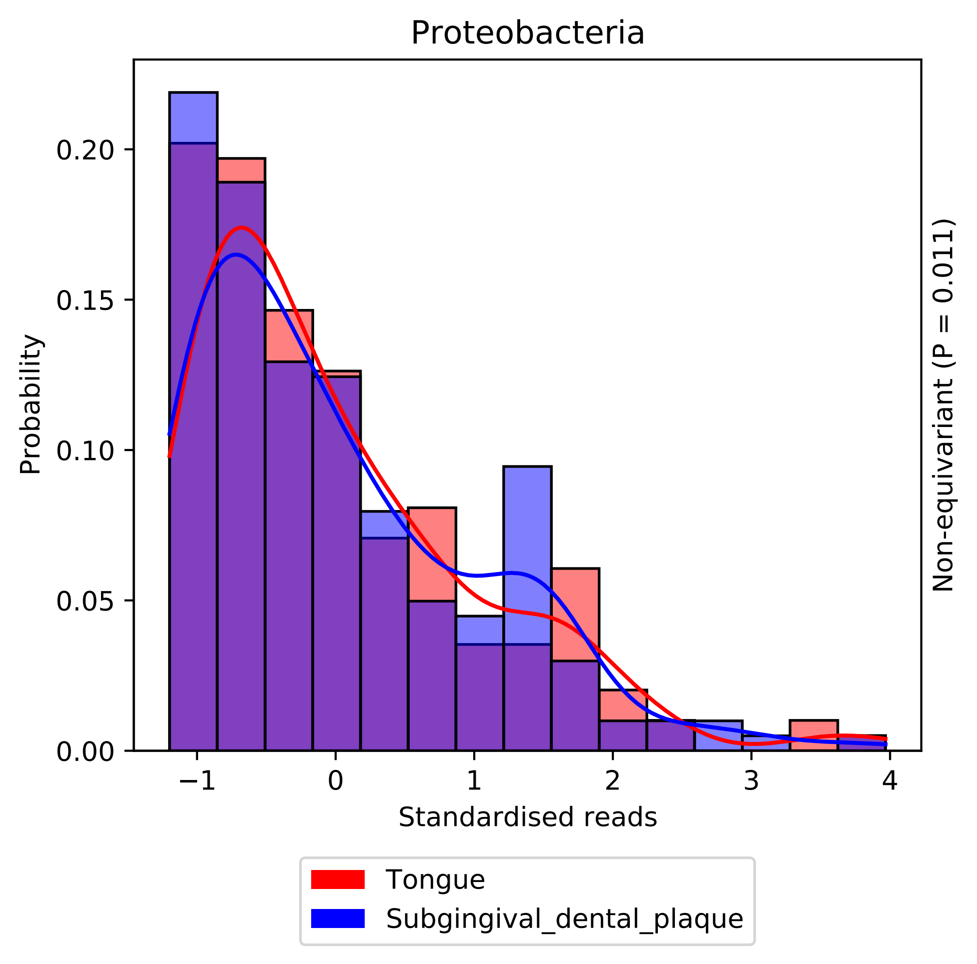

Proteobacteria
Probability
0.00
0.05
0.10
0.15
0.20
−1
0
1
2
3
4
Standardised reads
Non-equivariant (P = 0.011)
Tongue
Subgingival_dental_plaque

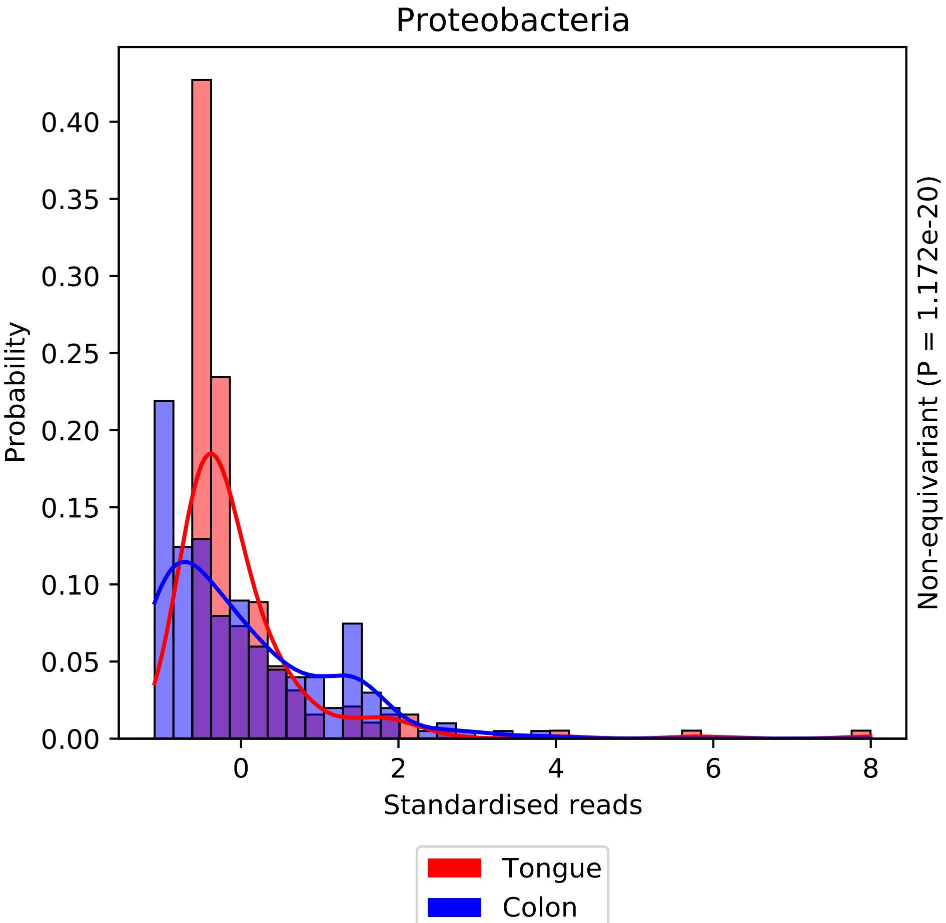
Proteobacteria
Probability
0.00
0.05
0.10
0.15
0.20
0.25
0.30
0.35
0.40
0
2
4
6
8
Standardised reads
Non-equivariant (P = 1.172e-20)
Tongue
Colon

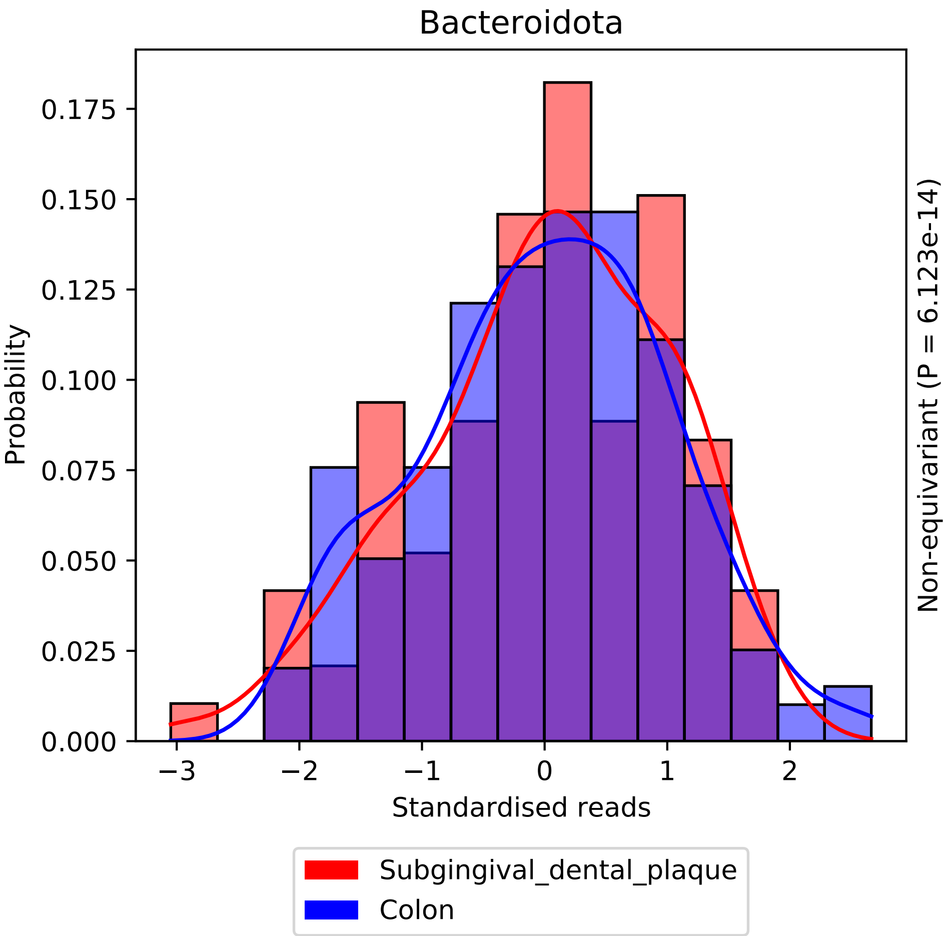

Bacteroidota
0.175
0.150
0.125
0.100
0.075
0.050
0.025
0.000
Probability
−3
−2
−1
0
1
2
Standardised reads
Non-equivariant (P = 6.123e-14)
Subgingival_dental_plaque
Colon

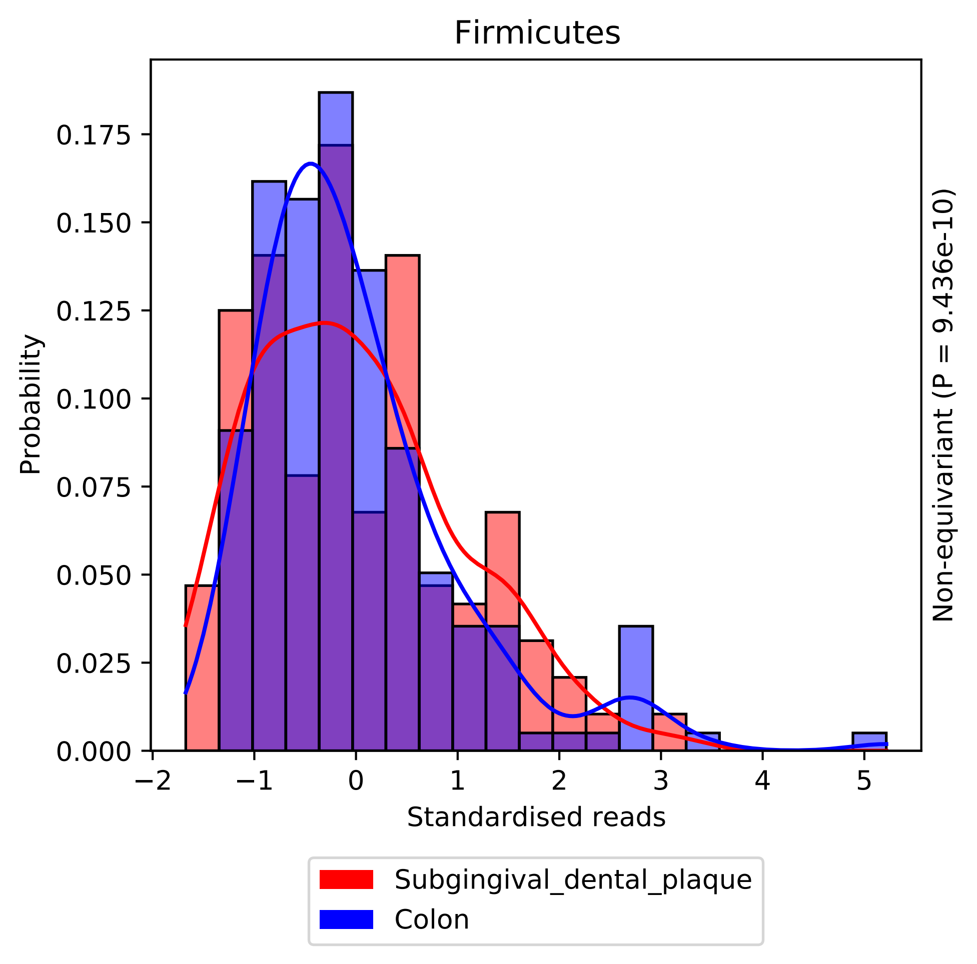
Firmicutes
Probability
Standardised reads
Non-equivariant (P = 9.436e-10)
0.000
0.025
0.050
0.075
0.100
0.125
0.150
0.175
−2
−1
0
1
2
3
4
5
Subgingival_dental_plaque
Colon

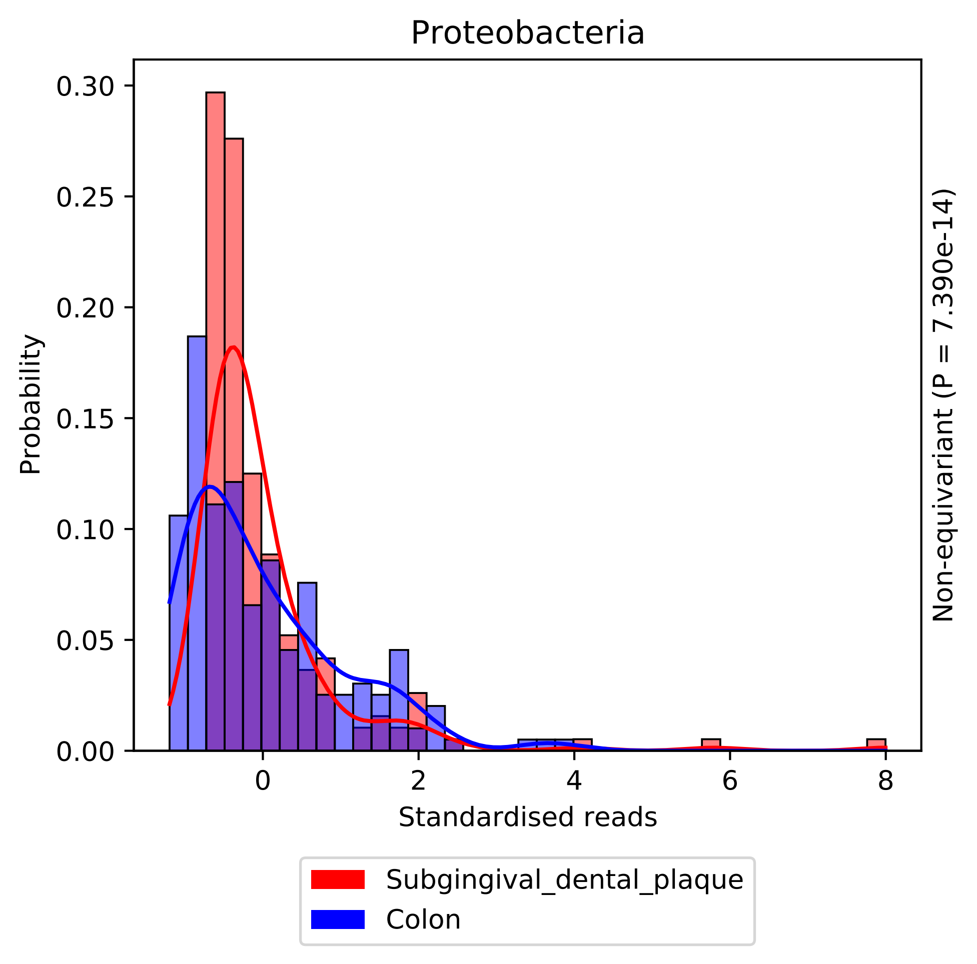

Proteobacteria
Probability
0.00
0.05
0.10
0.15
0.20
0.25
0.30
0
2
4
6
8
Standardised reads
Non-equivariant (P = 7.390e-14)
Subgingival_dental_plaque
Colon
