## Supplementary material for "statSuma: automated selection and performance of statistical comparisons for microbiome studies": SI Figure 2 (Gaussian comparisons)

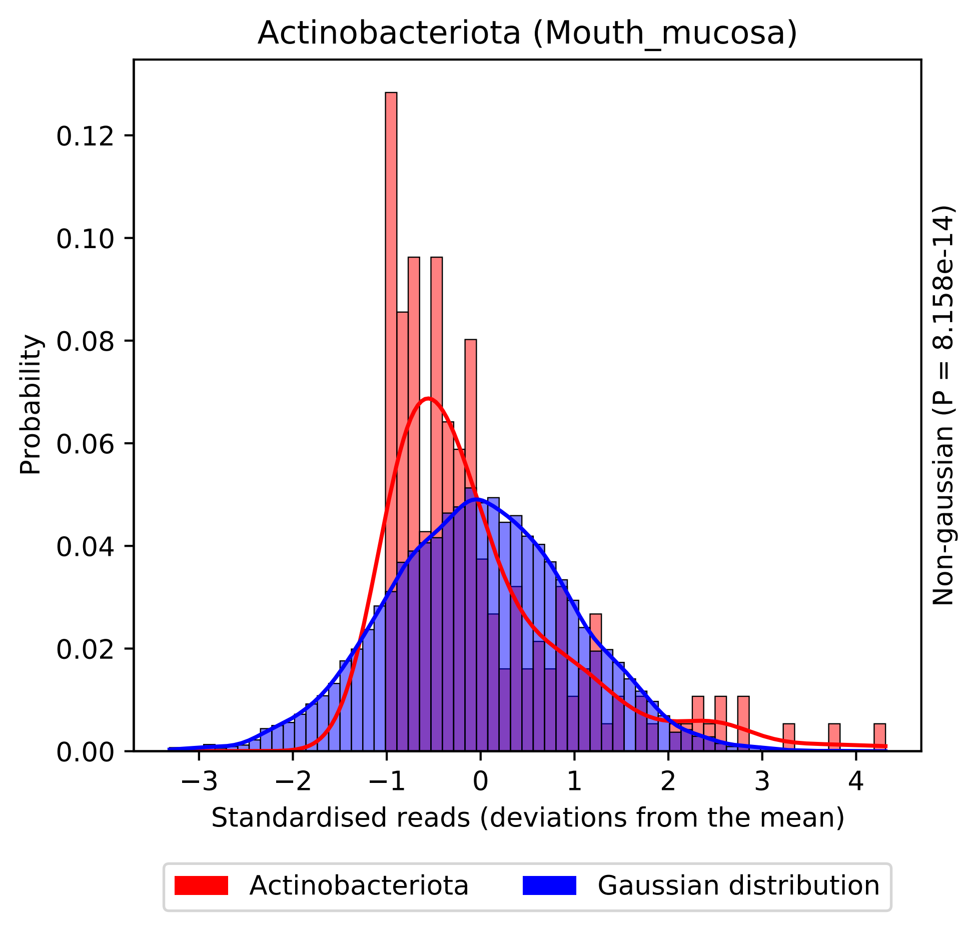

Actinobacteriota (Mouth_mucosa)
Probability
0.12
0.10
0.08
0.06
0.04
0.02
0.00
−3
−2
−1
0
1
2
3
4
Standardised reads (deviations from the mean)
Non-gaussian (P = 8.158e-14)
Actinobacteriota
Gaussian distribution

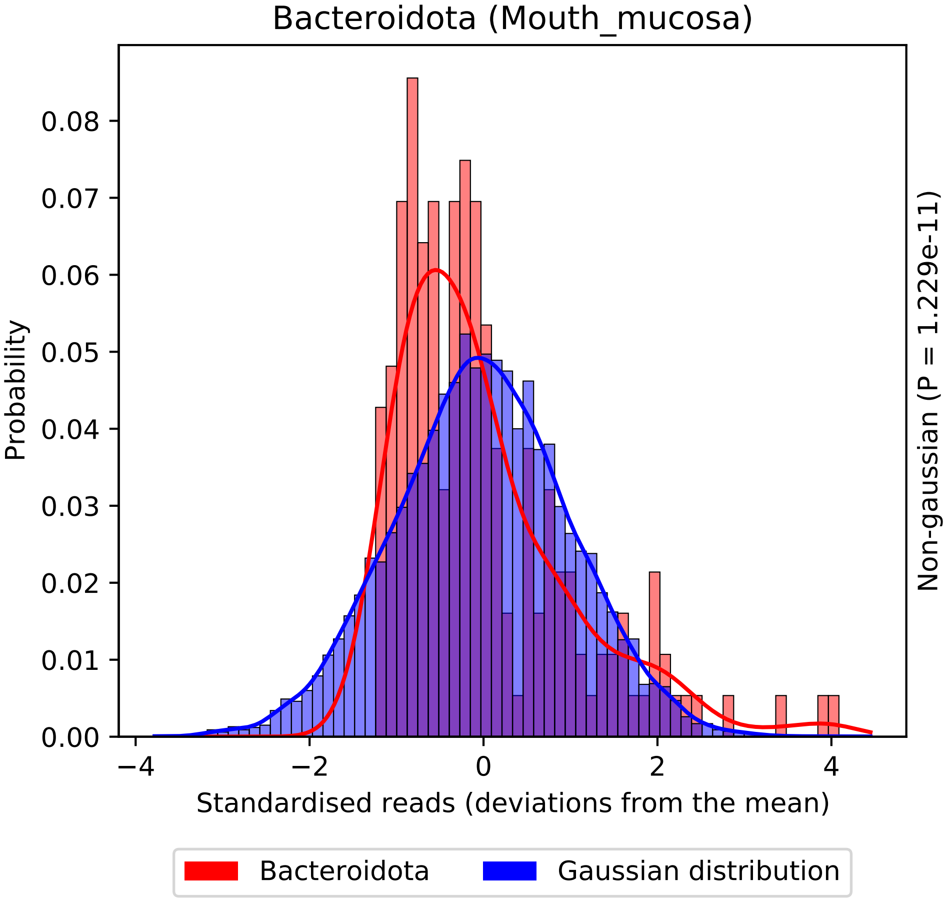

Bacteroidota (Mouth_mucosa)
Probability
0.00
0.01
0.02
0.03
0.04
0.05
0.06
0.07
0.08
−4
−2
0
2
4
Standardised reads (deviations from the mean)
Non-gaussian (P = 1.229e-11)
Bacteroidota
Gaussian distribution

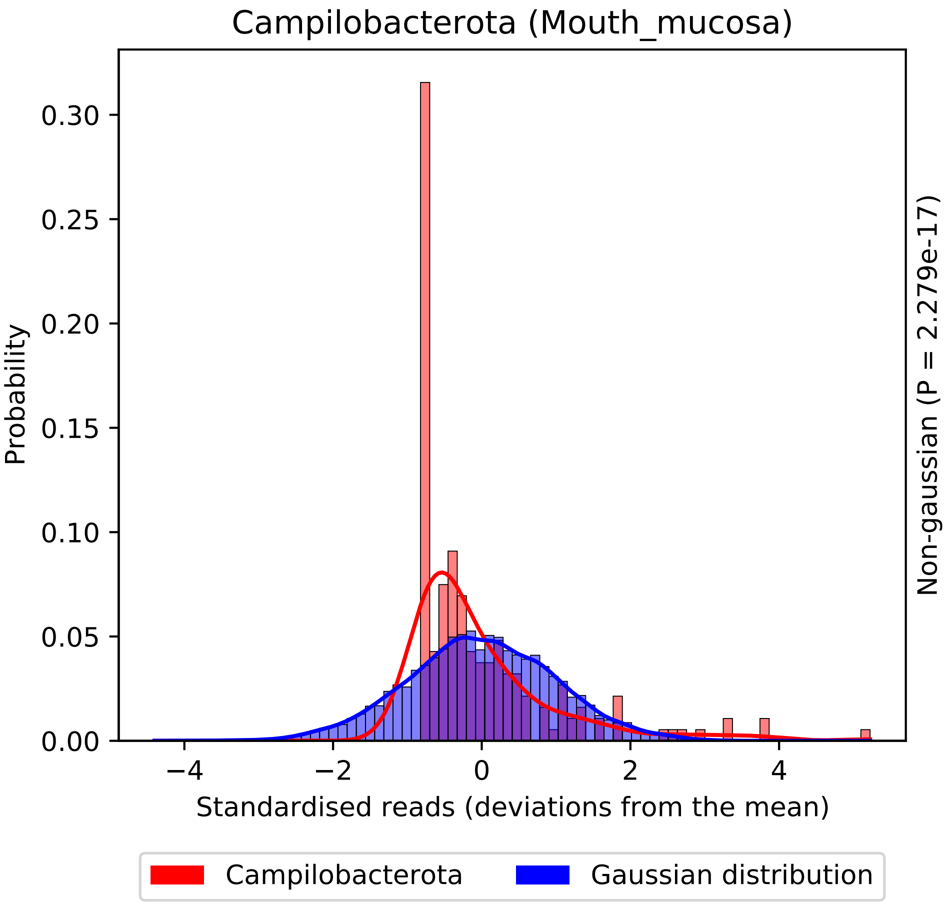

Campilobacterota (Mouth_mucosa)
Probability
0.30
0.25
0.20
0.15
0.10
0.05
0.00
−4
−2
0
2
4
Standardised reads (deviations from the mean)
Non-gaussian (P = 2.279e-17)
Campilobacterota
Gaussian distribution

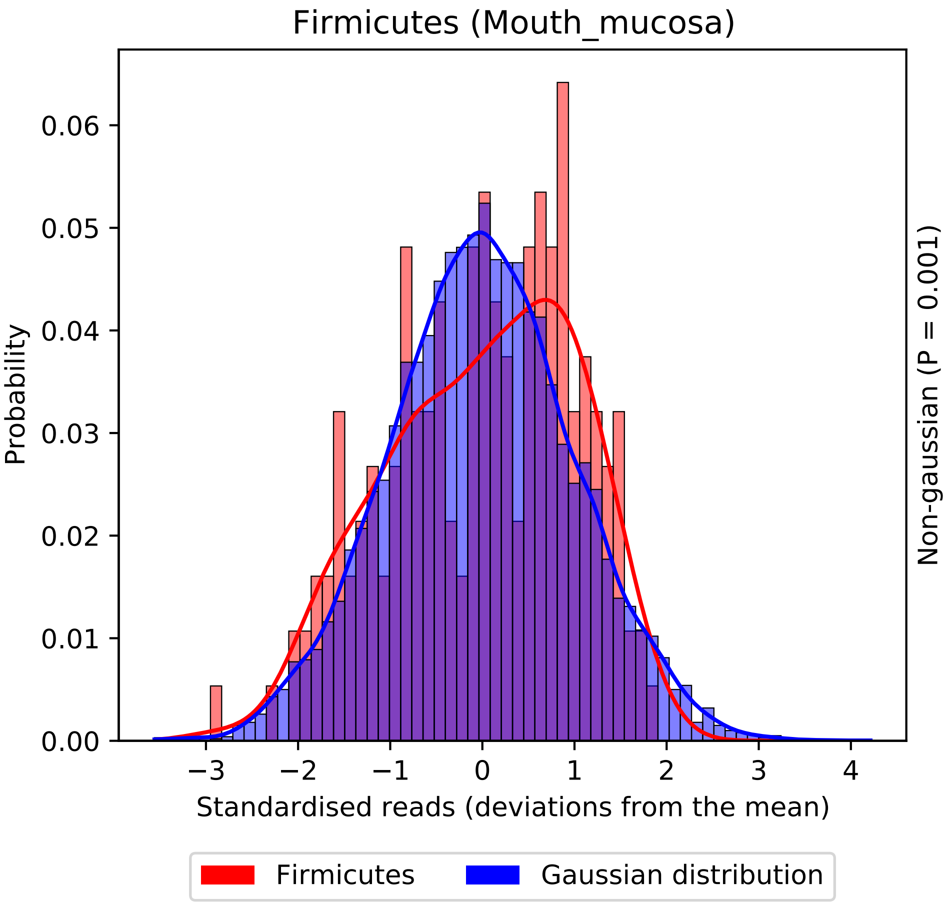

Firmicutes (Mouth_mucosa)
Probability
0.00
0.01
0.02
0.03
0.04
0.05
0.06
−3
−2
−1
0
1
2
3
4
Standardised reads (deviations from the mean)
Non-gaussian (P = 0.001)
Firmicutes
Gaussian distribution

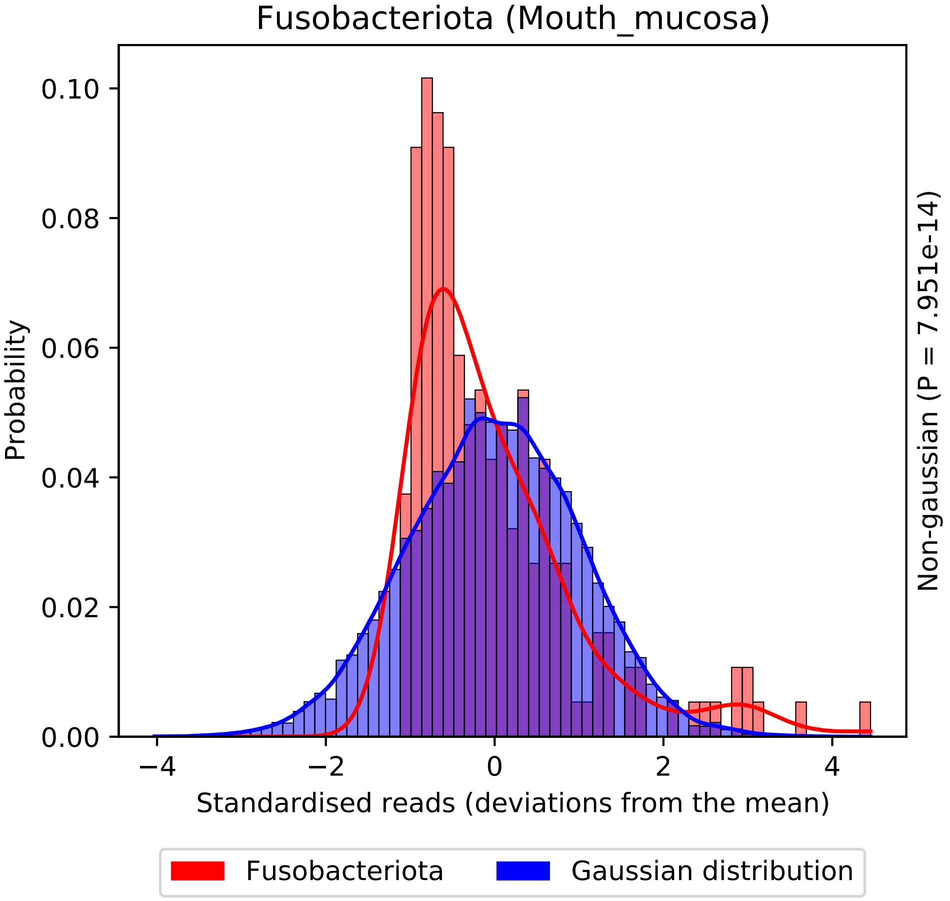
Fusobacteriota (Mouth_mucosa)
Probability
0.10
0.08
0.06
0.04
0.02
0.00
−4
−2
0
2
4
Standardised reads (deviations from the mean)
Non-gaussian (P = 7.951e-14)
Fusobacteriota
Gaussian distribution

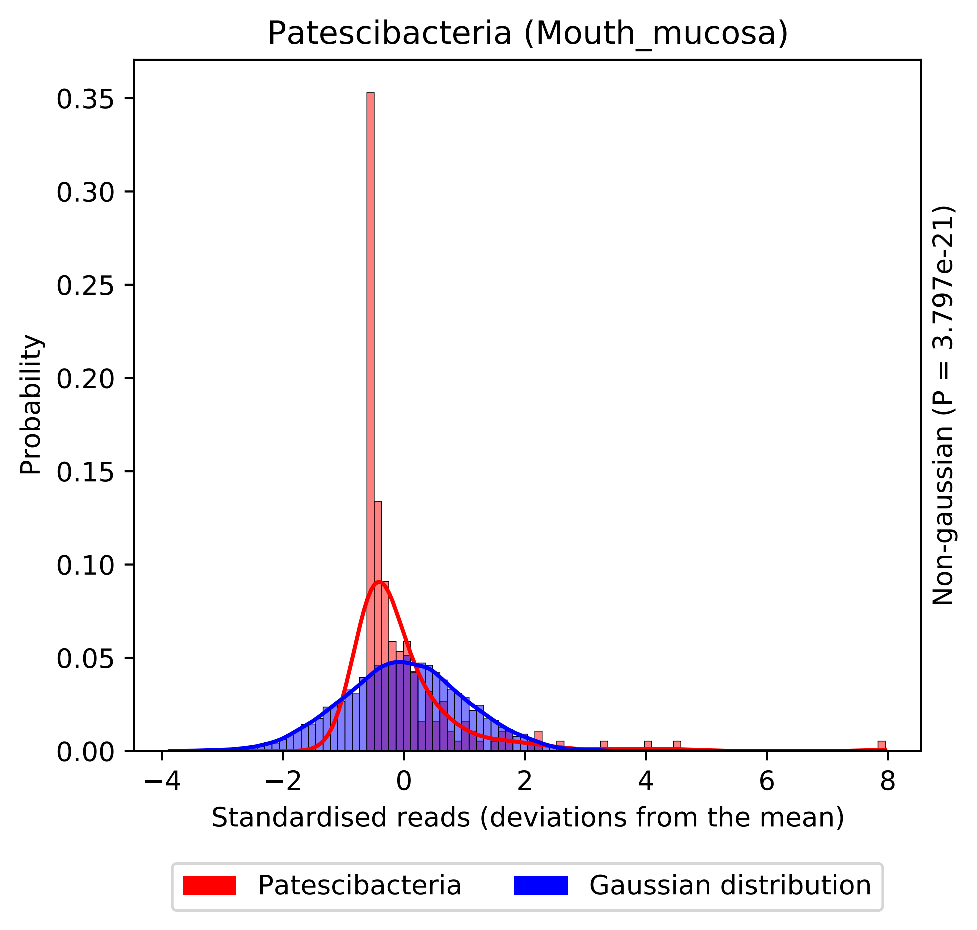
Patescibacteria (Mouth_mucosa)
Probability
0.00
0.05
0.10
0.15
0.20
0.25
0.30
0.35
−4
−2
0
2
4
6
8
Standardised reads (deviations from the mean)
Non-gaussian (P = 3.797e-21)
Patescibacteria
Gaussian distribution

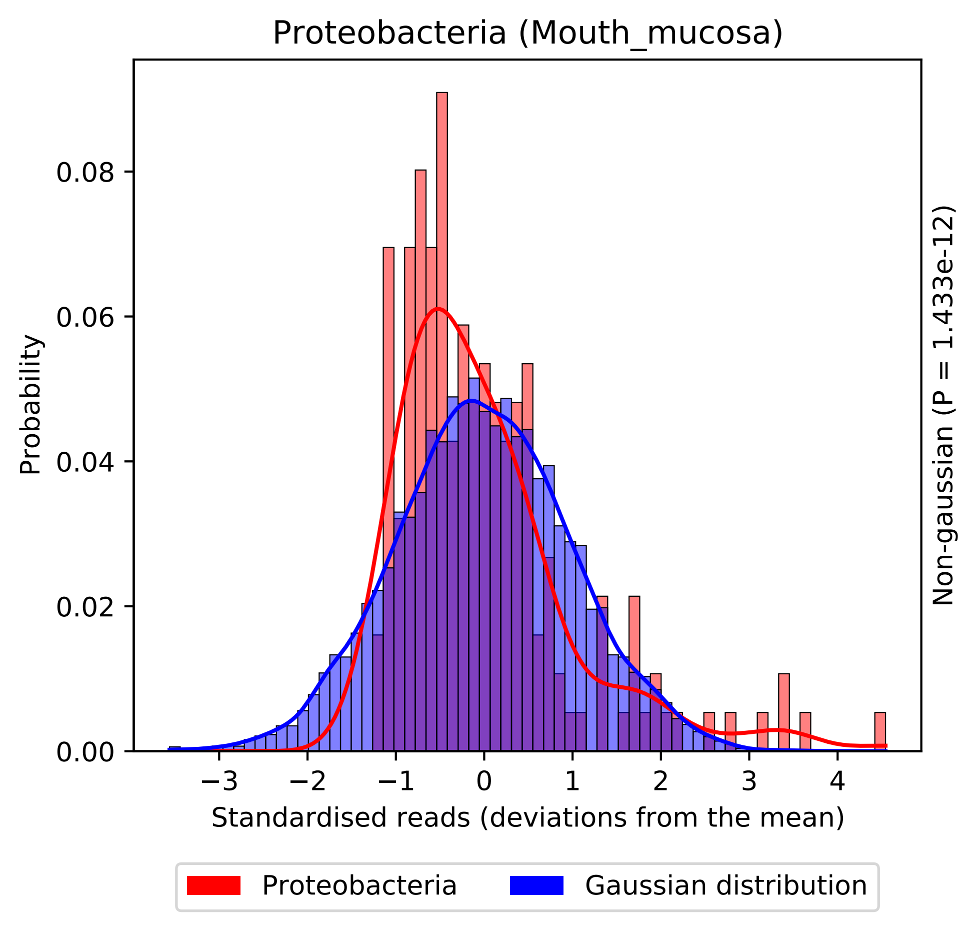

Proteobacteria (Mouth_mucosa)
Probability
0.00
0.02
0.04
0.06
0.08
−3
−2
−1
0
1
2
3
4
Standardised reads (deviations from the mean)
Non-gaussian (P = 1.433e-12)
Proteobacteria
Gaussian distribution

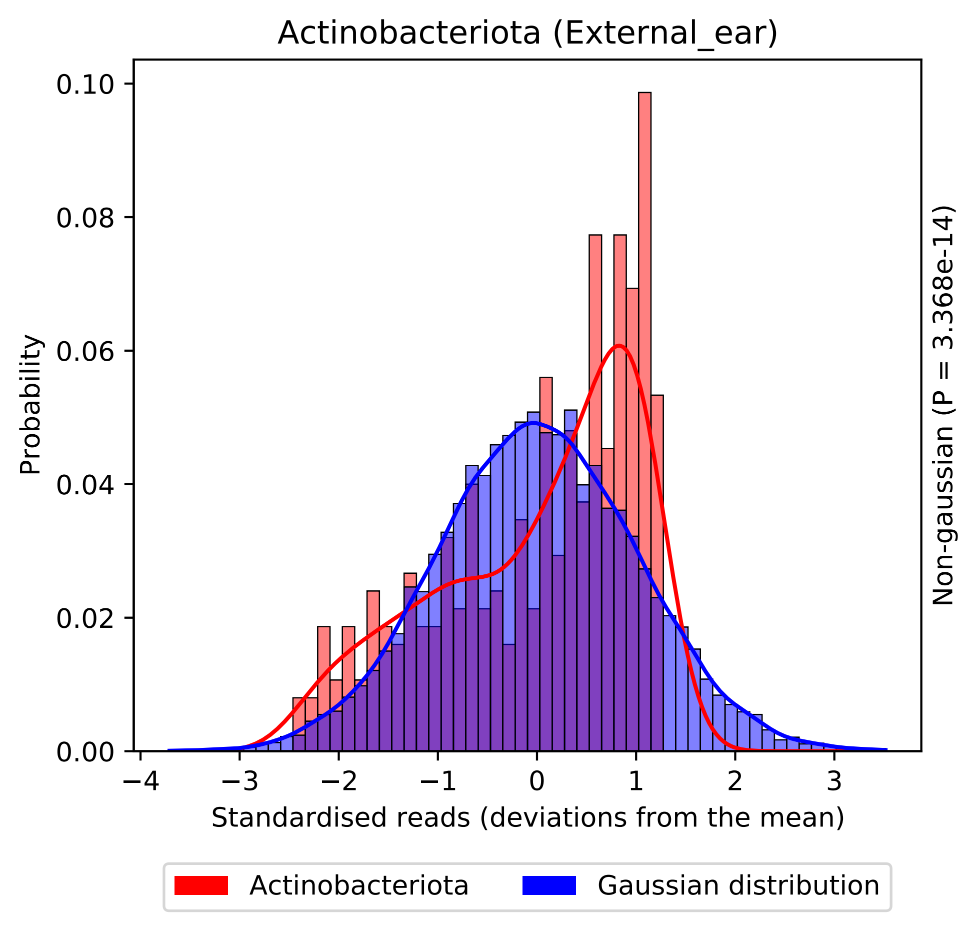

Actinobacteriota (External_ear)
Probability
0.00
0.02
0.04
0.06
0.08
0.10
−4
−3
−2
−1
0
1
2
3
Standardised reads (deviations from the mean)
Non-gaussian (P = 3.368e-14)
Actinobacteriota
Gaussian distribution

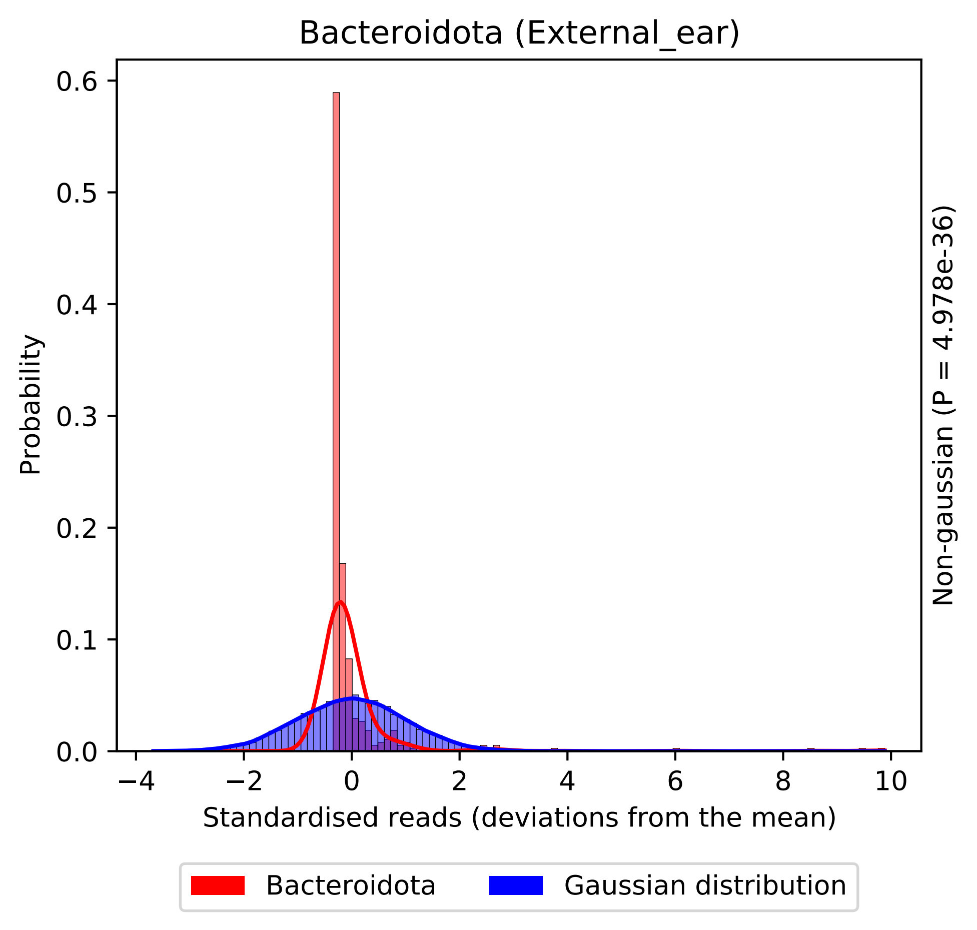

Bacteroidota (External_ear)
Probability
0.0
0.1
0.2
0.3
0.4
0.5
0.6
−4
−2
0
2
4
6
8
10
Standardised reads (deviations from the mean)
Non-gaussian (P = 4.978e-36)
Bacteroidota
Gaussian distribution

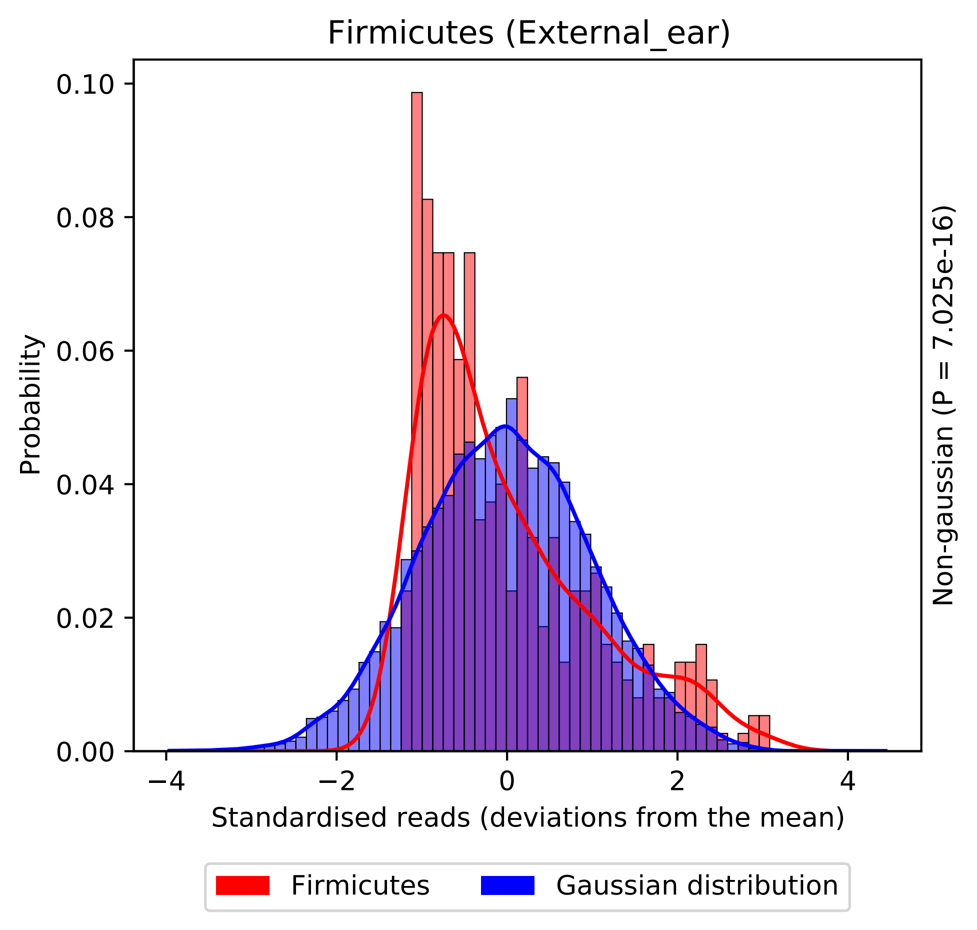
Firmicutes (External_ear)
Probability
0.00
0.02
0.04
0.06
0.08
0.10
−4
−2
0
2
4
Standardised reads (deviations from the mean)
Non-gaussian (P = 7.025e-16)
Firmicutes
Gaussian distribution

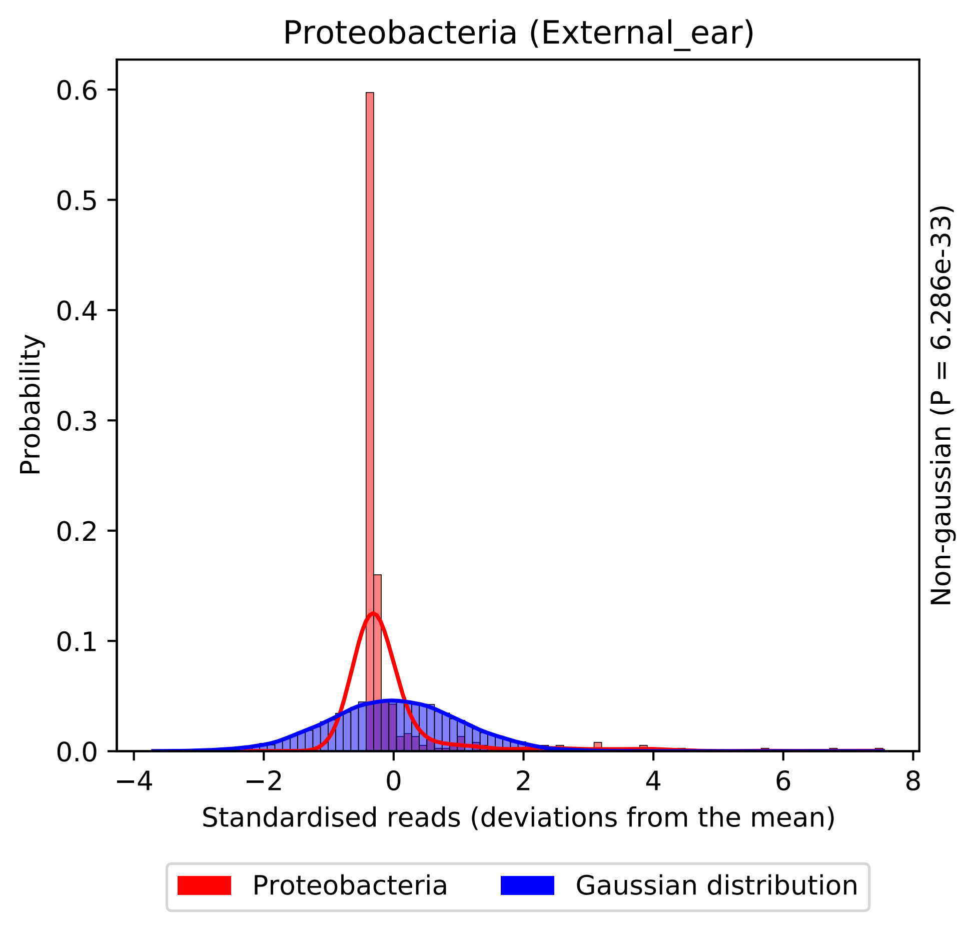
Proteobacteria (External_ear)
Probability
0.0
0.1
0.2
0.3
0.4
0.5
0.6
−4
−2
0
2
4
6
8
Standardised reads (deviations from the mean)
Non-gaussian (P = 6.286e-33)
Proteobacteria
Gaussian distribution

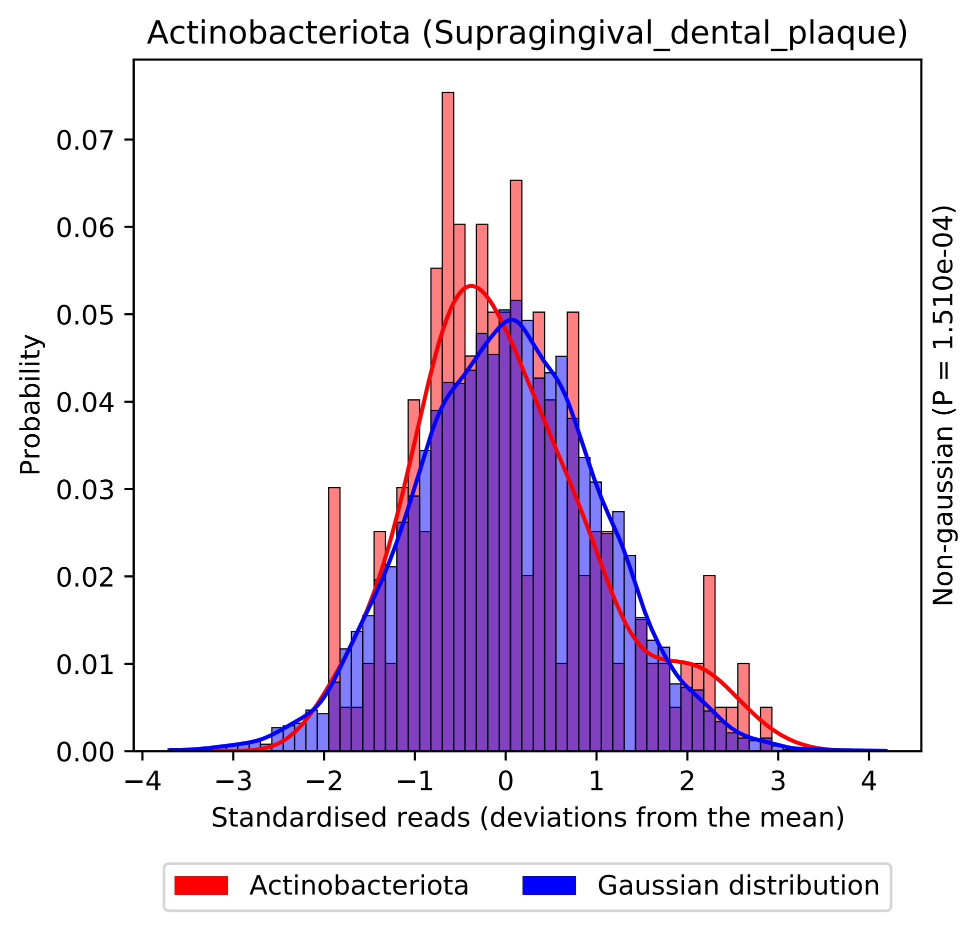

Actinobacteriota (Supragingival_dental_plaque)
Probability
0.00
0.01
0.02
0.03
0.04
0.05
0.06
0.07
−4
−3
−2
−1
0
1
2
3
4
Standardised reads (deviations from the mean)
Non-gaussian (P = 1.510e-04)
Actinobacteriota
Gaussian distribution

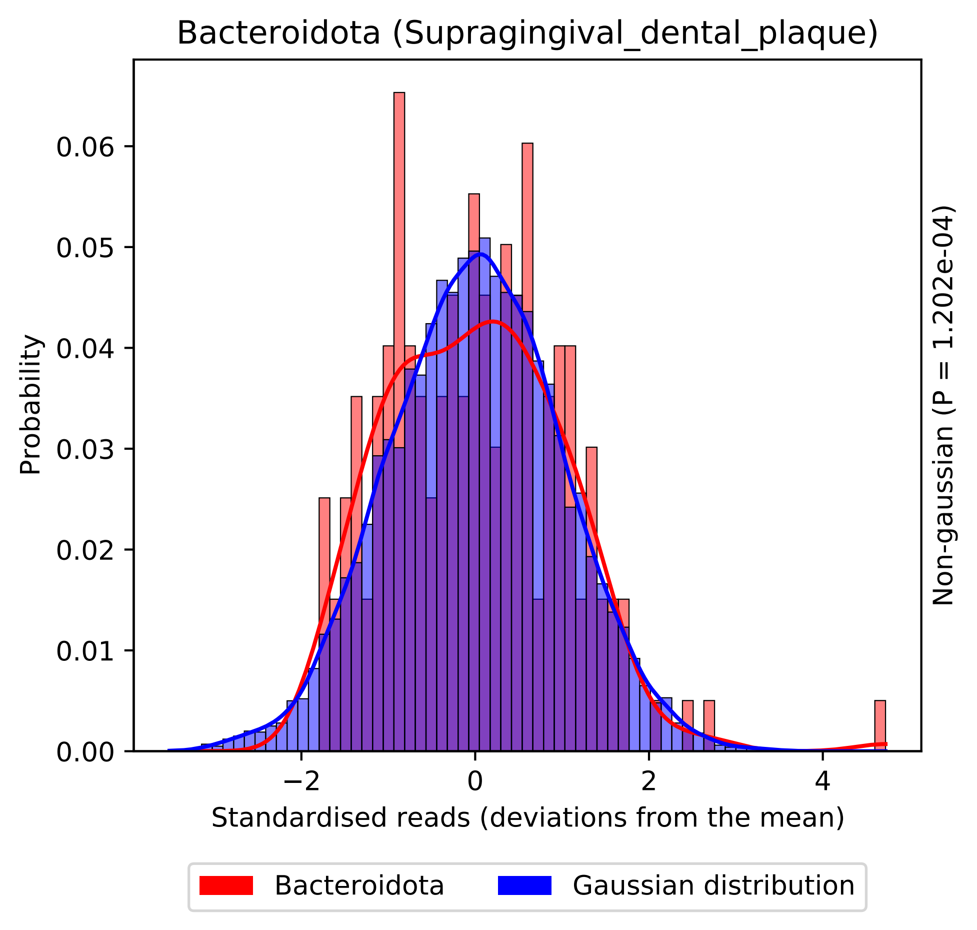

Bacteroidota (Supragingival_dental_plaque)
Probability
0.00
0.01
0.02
0.03
0.04
0.05
0.06
−2
0
2
4
Standardised reads (deviations from the mean)
Non-gaussian (P = 1.202e-04)
Bacteroidota
Gaussian distribution

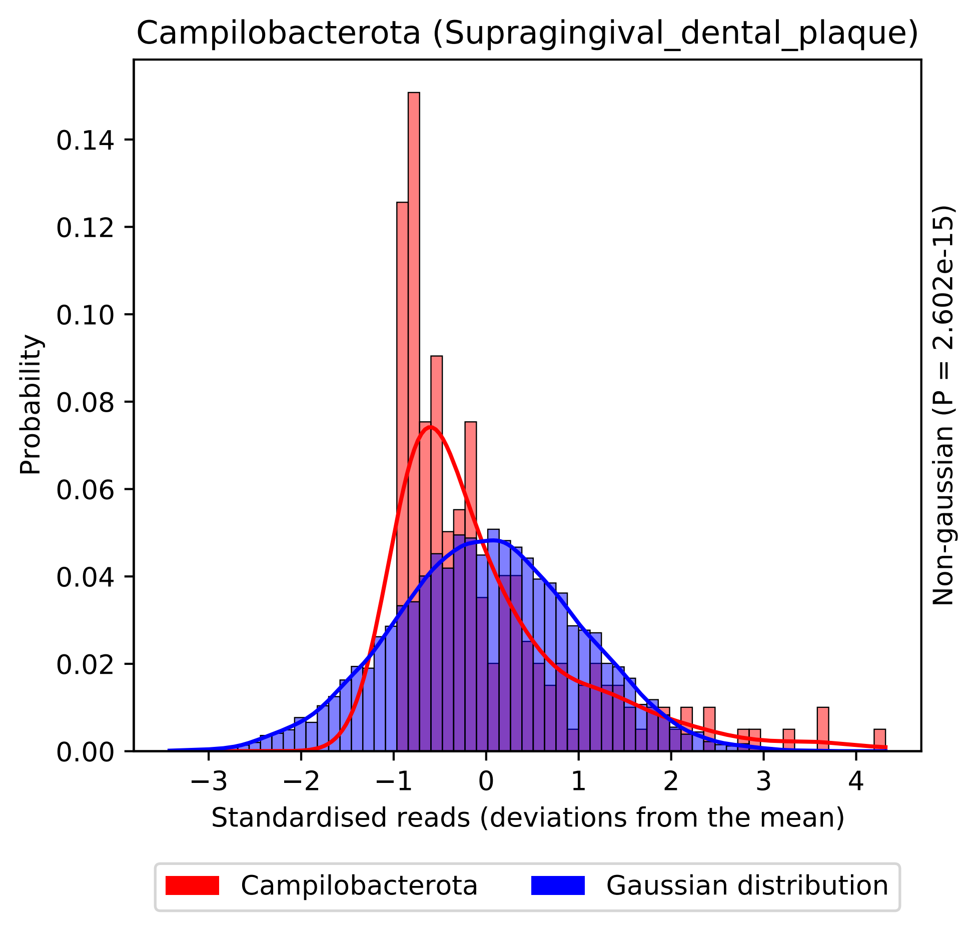
Campilobacterota (Supragingival_dental_plaque)
Probability
0.00
0.02
0.04
0.06
0.08
0.10
0.12
0.14
−3
−2
−1
0
1
2
3
4
Standardised reads (deviations from the mean)
Non-gaussian (P = 2.602e-15)
Campilobacterota
Gaussian distribution

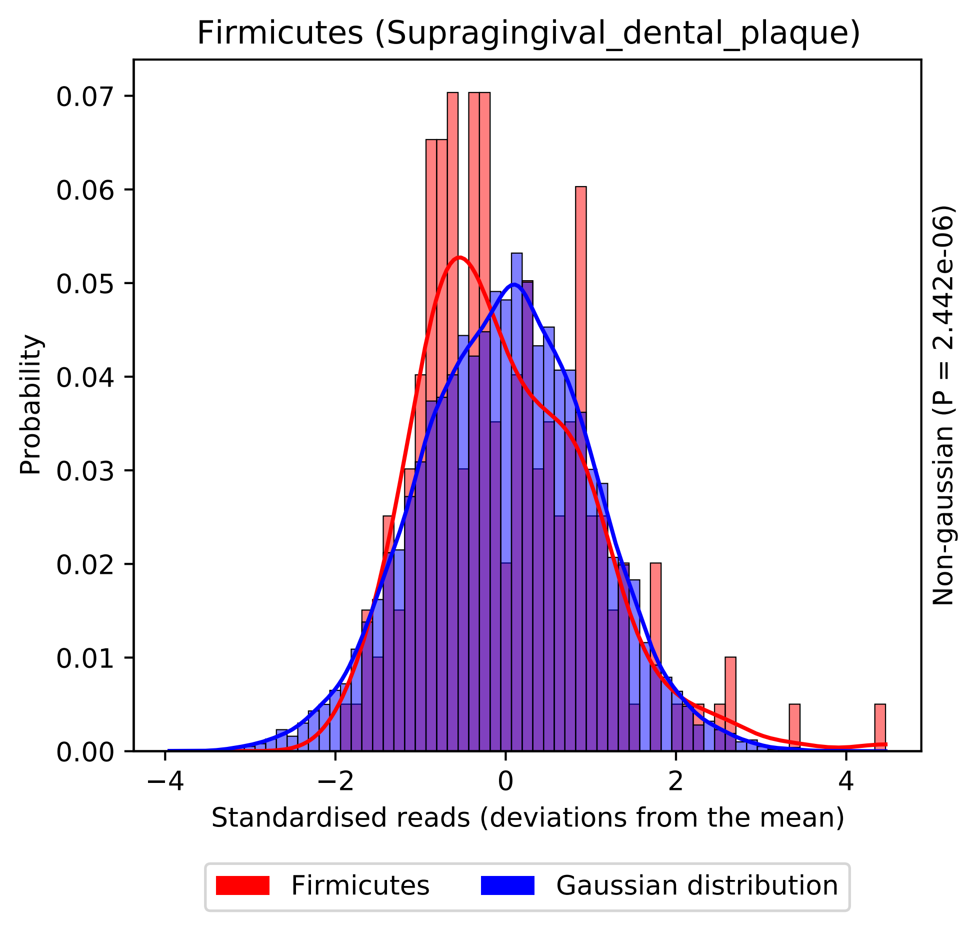

Firmicutes (Supragingival_dental_plaque)
Probability
0.07
0.06
0.05
0.04
0.03
0.02
0.01
0.00
−4
−2
0
2
4
Standardised reads (deviations from the mean)
Non-gaussian (P = 2.442e-06)
Firmicutes
Gaussian distribution

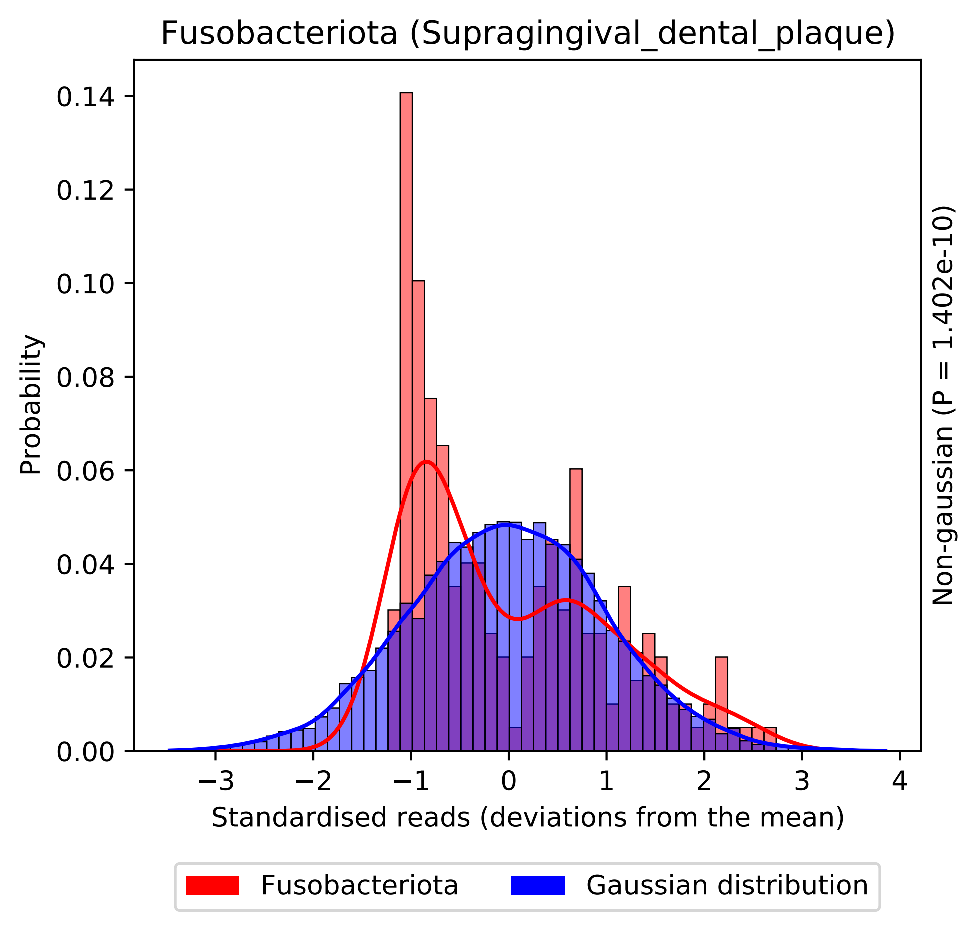

Fusobacteriota (Supragingival_dental_plaque)
Probability
0.00
0.02
0.04
0.06
0.08
0.10
0.12
0.14
−3
−2
−1
0
1
2
3
4
Standardised reads (deviations from the mean)
Non-gaussian (P = 1.402e-10)
Fusobacteriota
Gaussian distribution

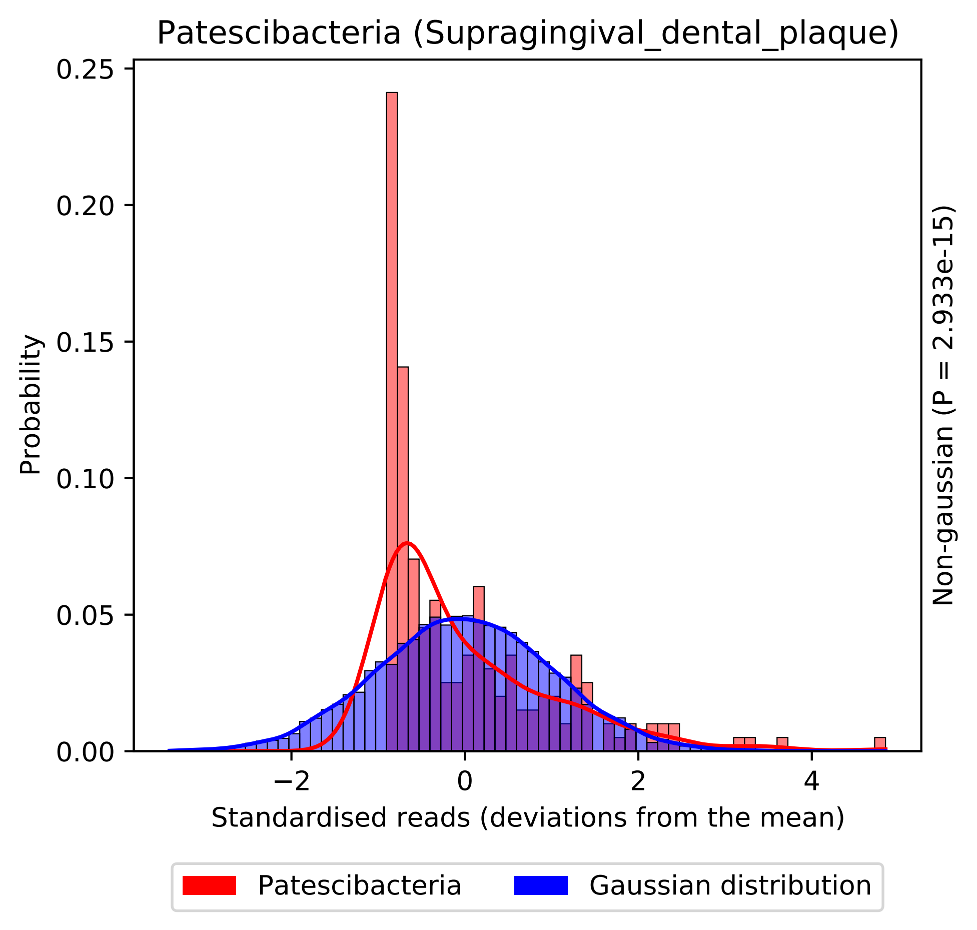

Patescibacteria (Supragingival_dental_plaque)
Probability
0.25
0.20
0.15
0.10
0.05
0.00
−2
0
2
4
Standardised reads (deviations from the mean)
Non-gaussian (P = 2.933e-15)
Patescibacteria
Gaussian distribution

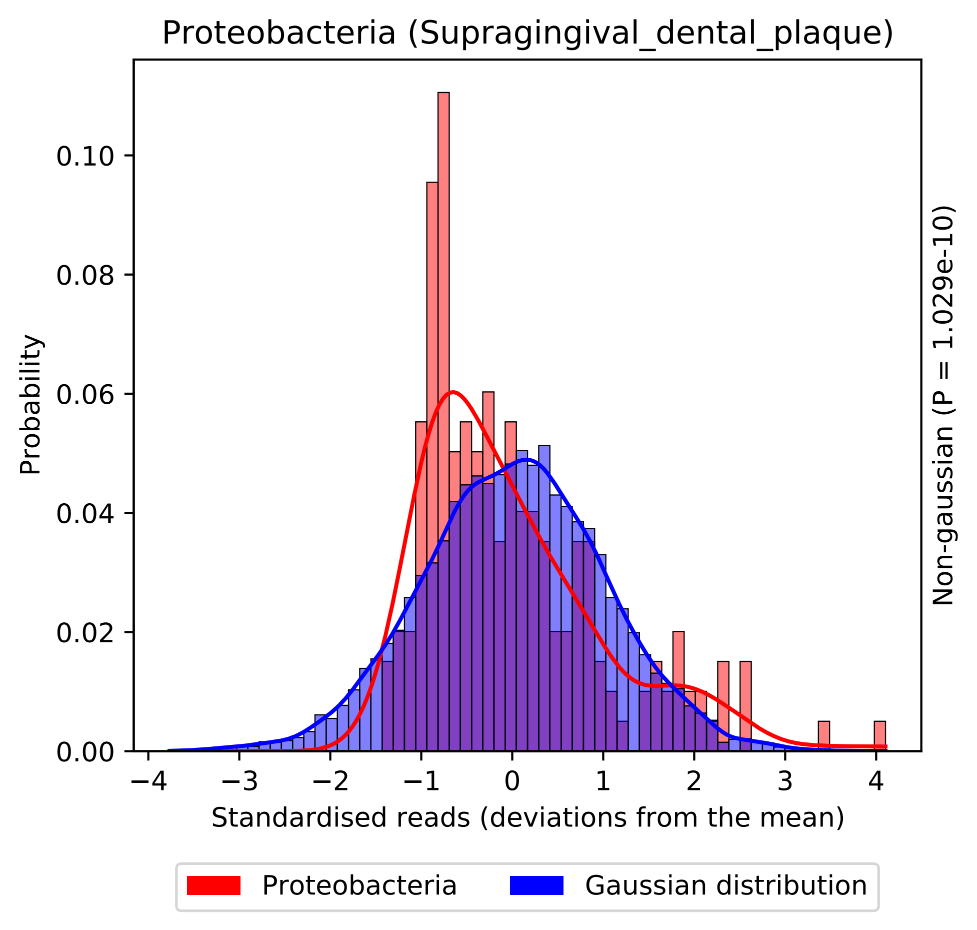

Proteobacteria (Supragingival_dental_plaque)
Probability
0.00
0.02
0.04
0.06
0.08
0.10
−4
−3
−2
−1
0
1
2
3
4
Standardised reads (deviations from the mean)
Non-gaussian (P = 1.029e-10)
Proteobacteria
Gaussian distribution

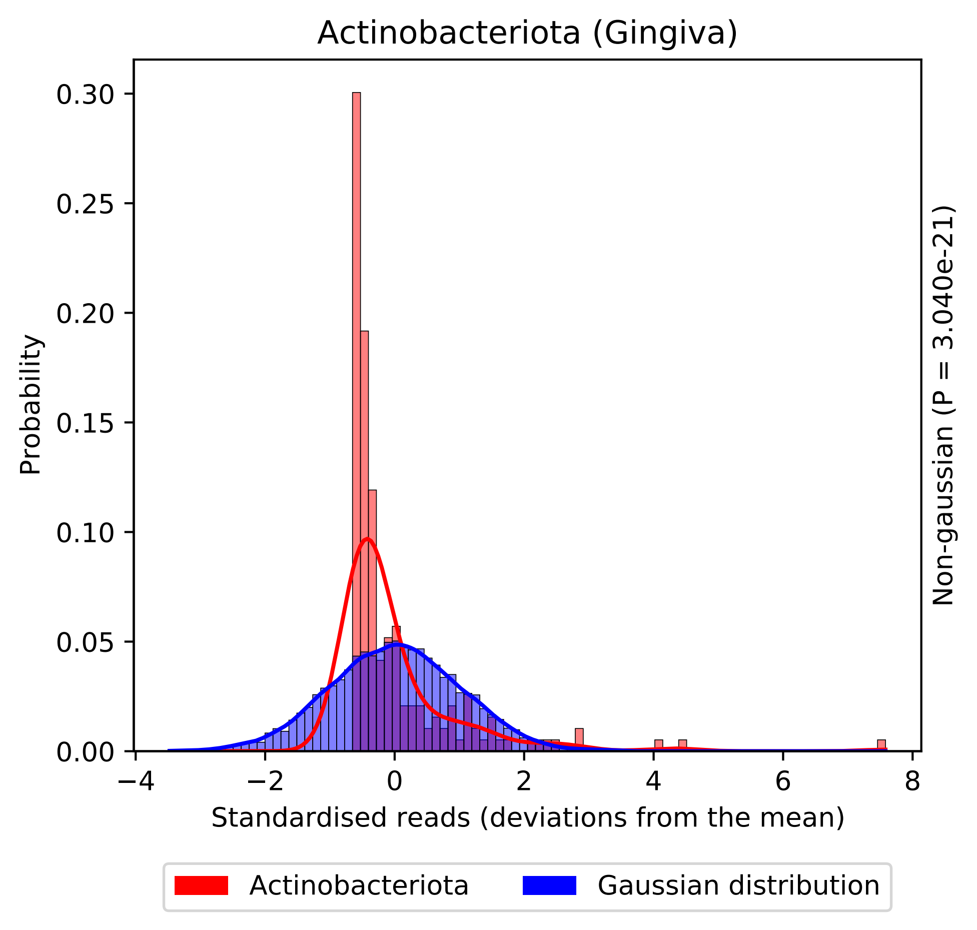
Actinobacteriota (Gingiva)
Probability
0.30
0.25
0.20
0.15
0.10
0.05
0.00
−4
−2
0
2
4
6
8
Standardised reads (deviations from the mean)
Non-gaussian (P = 3.040e-21)
Actinobacteriota
Gaussian distribution

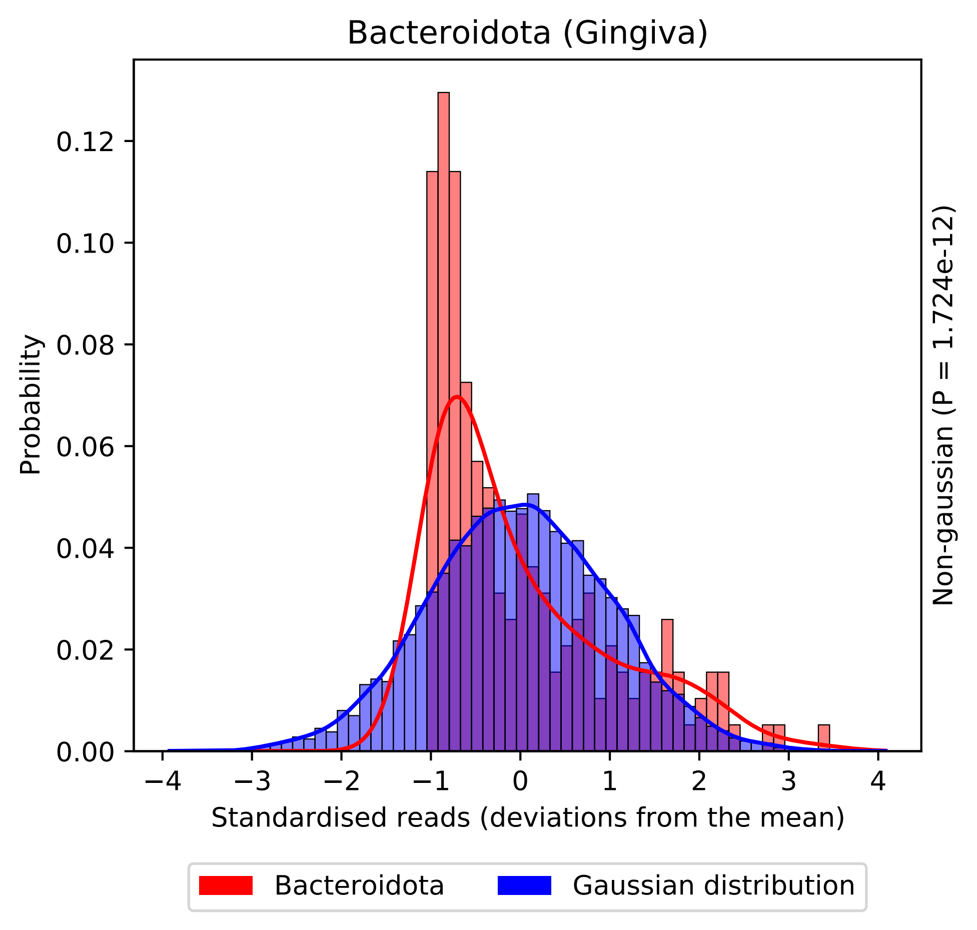
Bacteroidota (Gingiva)
Probability
0.00
0.02
0.04
0.06
0.08
0.10
0.12
−4
−3
−2
−1
0
1
2
3
4
Standardised reads (deviations from the mean)
Non-gaussian (P = 1.724e-12)
Bacteroidota
Gaussian distribution

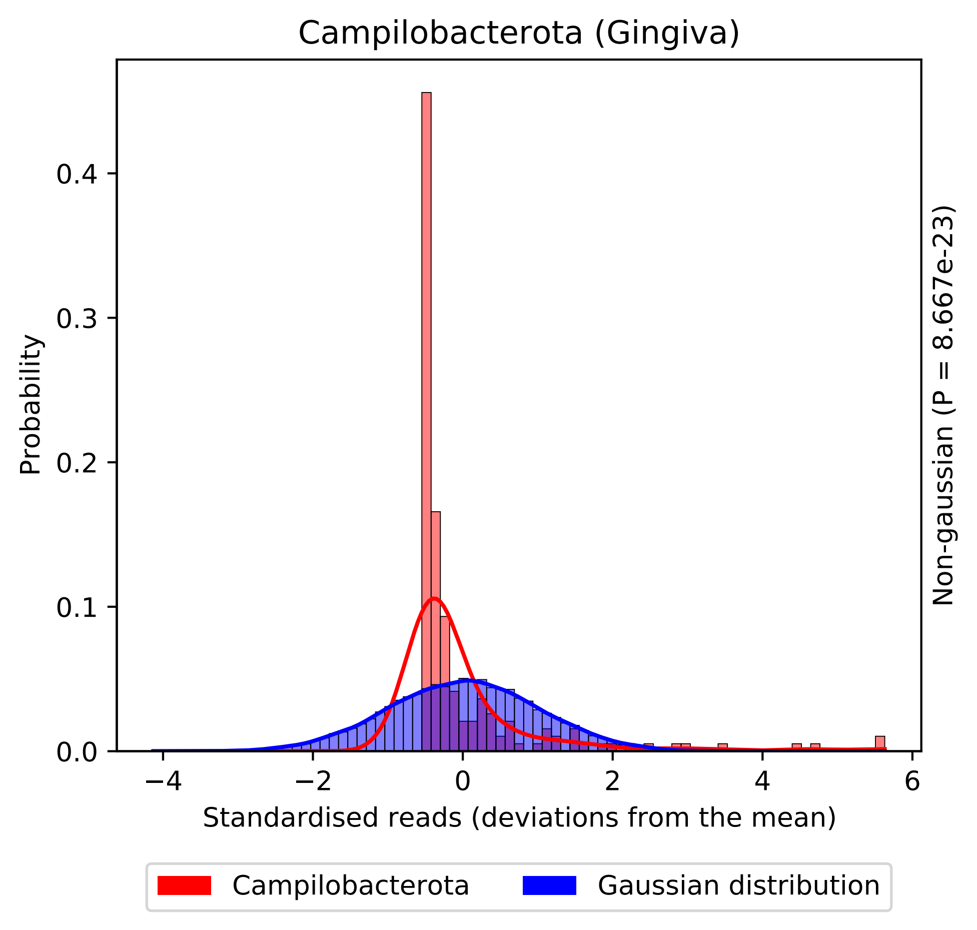
Campilobacterota (Gingiva)
Probability
0.0
0.1
0.2
0.3
0.4
−4
−2
0
2
4
6
Standardised reads (deviations from the mean)
Non-gaussian (P = 8.667e-23)
Campilobacterota
Gaussian distribution

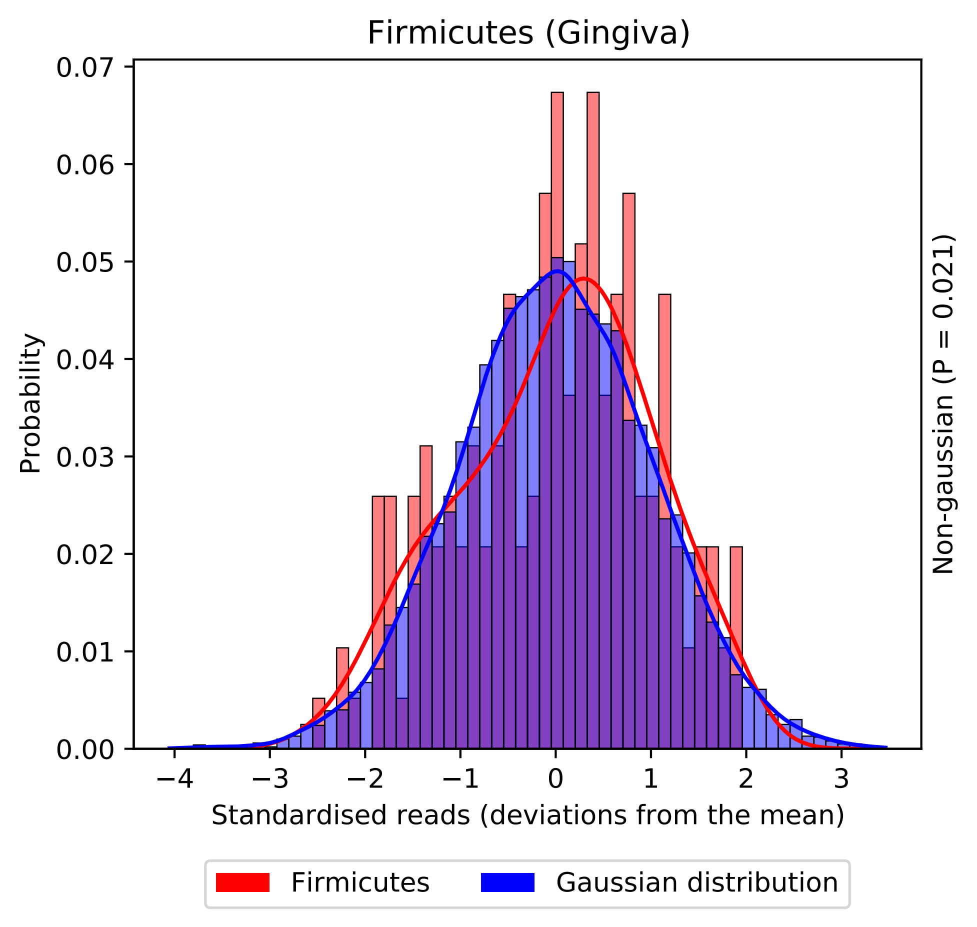

Firmicutes (Gingiva)
0.07
0.06
0.05
0.04
0.03
0.02
0.01
0.00
Probability
−4
−3
−2
−1
0
1
2
3
Standardised reads (deviations from the mean)
Non-gaussian (P = 0.021)
Firmicutes
Gaussian distribution

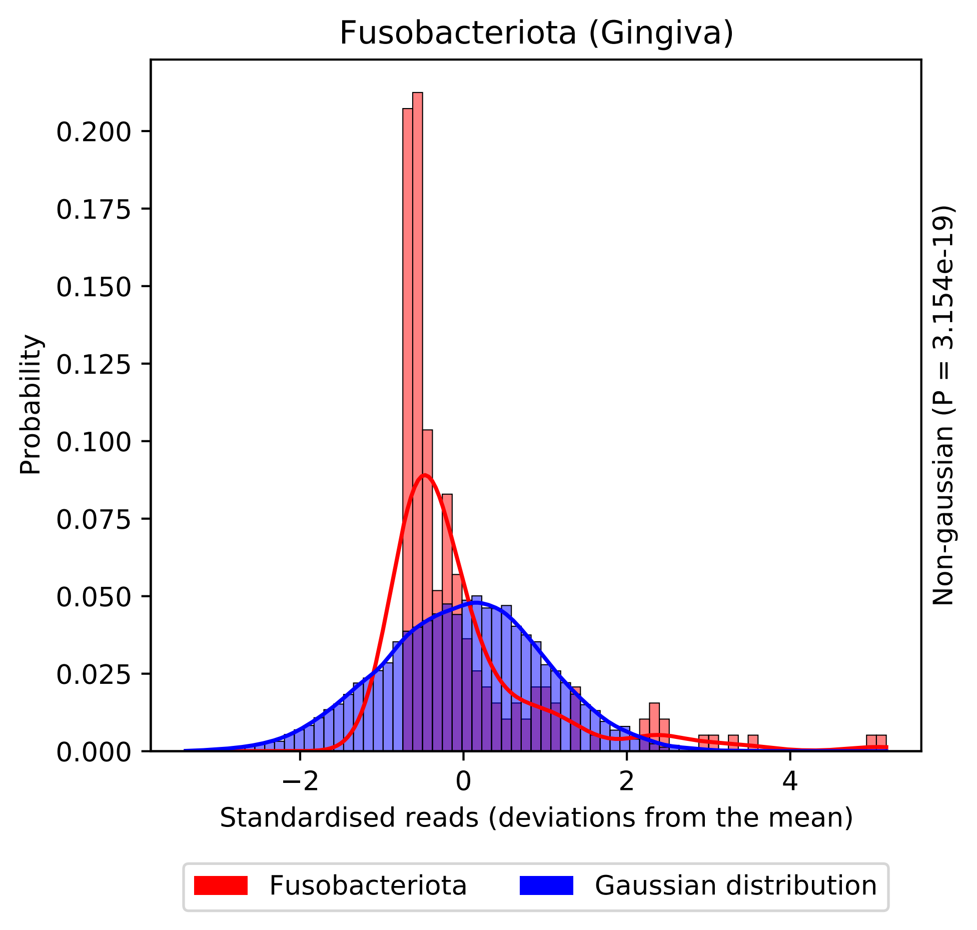
Fusobacteriota (Gingiva)
Probability
0.200
0.175
0.150
0.125
0.100
0.075
0.050
0.025
0.000
−2
0
2
4
Standardised reads (deviations from the mean)
Non-gaussian (P = 3.154e-19)
Fusobacteriota
Gaussian distribution

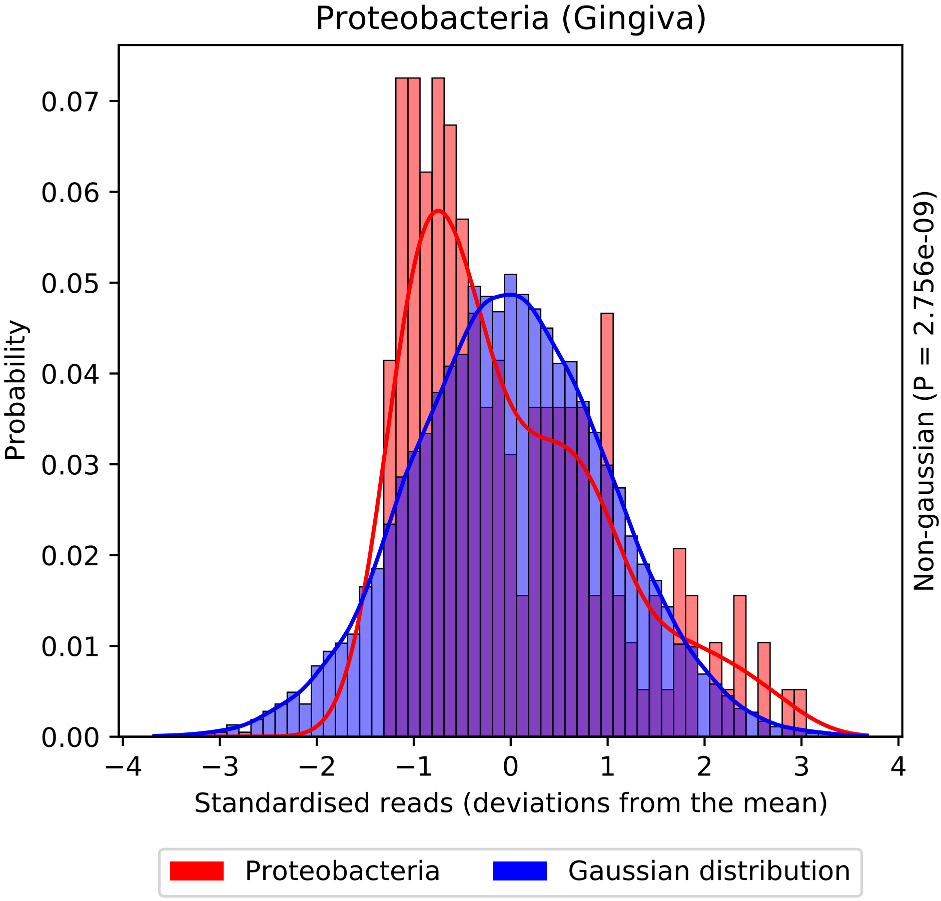
Proteobacteria (Gingiva)
Probability
0.07
0.06
0.05
0.04
0.03
0.02
0.01
0.00
−4
−3
−2
−1
0
1
2
3
4
Standardised reads (deviations from the mean)
Non-gaussian (P = 2.756e-09)
Proteobacteria
Gaussian distribution

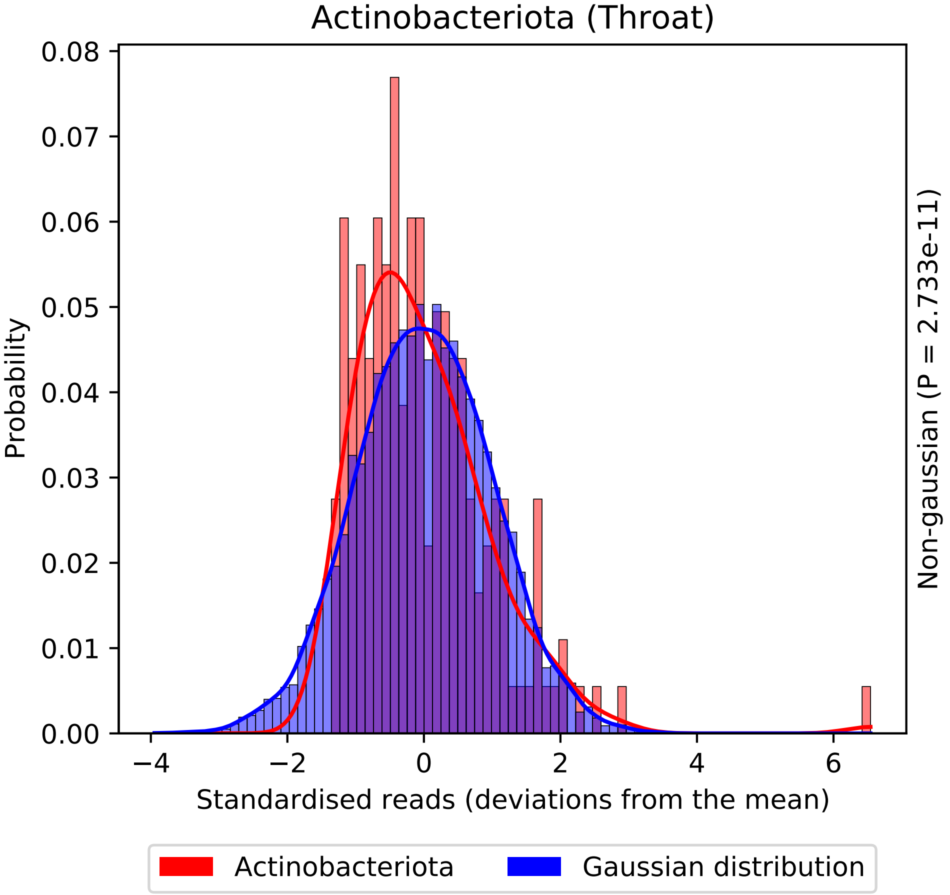

Actinobacteriota (Throat)
Probability
0.08
0.07
0.06
0.05
0.04
0.03
0.02
0.01
0.00
−4
−2
0
2
4
6
Standardised reads (deviations from the mean)
Non-gaussian (P = 2.733e-11)
Actinobacteriota
Gaussian distribution

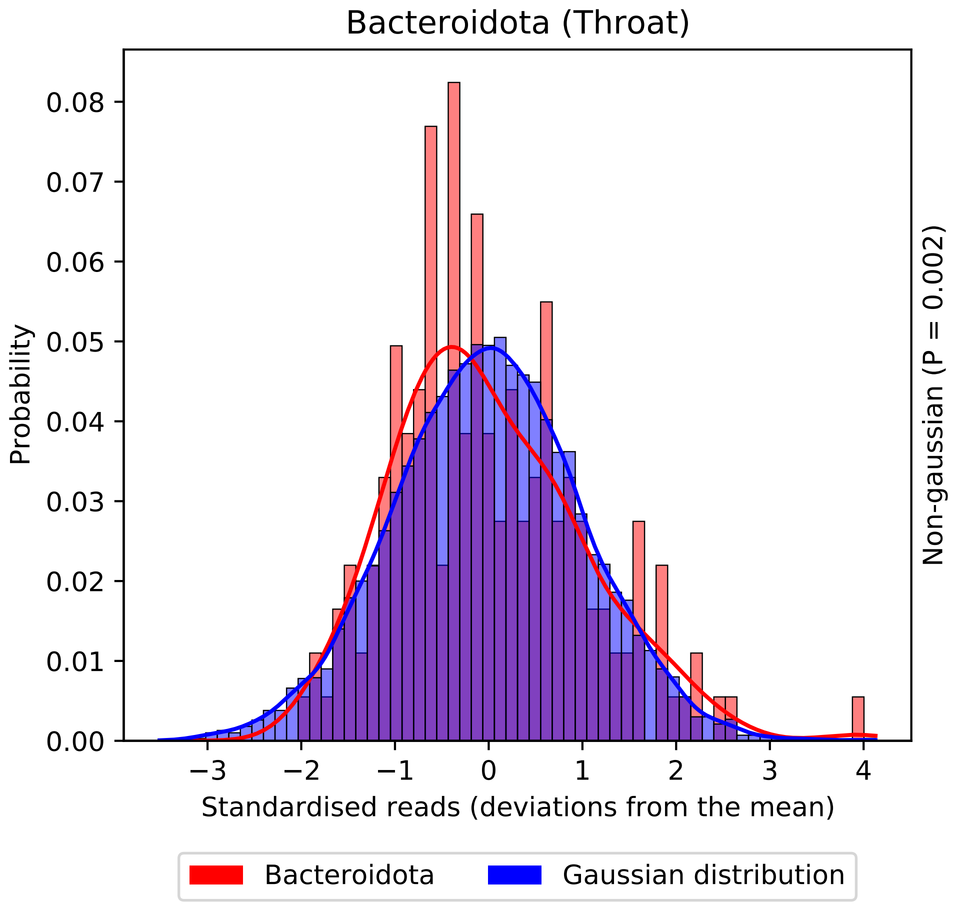

Bacteroidota (Throat)
Probability
0.00
0.01
0.02
0.03
0.04
0.05
0.06
0.07
0.08
−3
−2
−1
0
1
2
3
4
Standardised reads (deviations from the mean)
Non-gaussian (P = 0.002)
Bacteroidota
Gaussian distribution

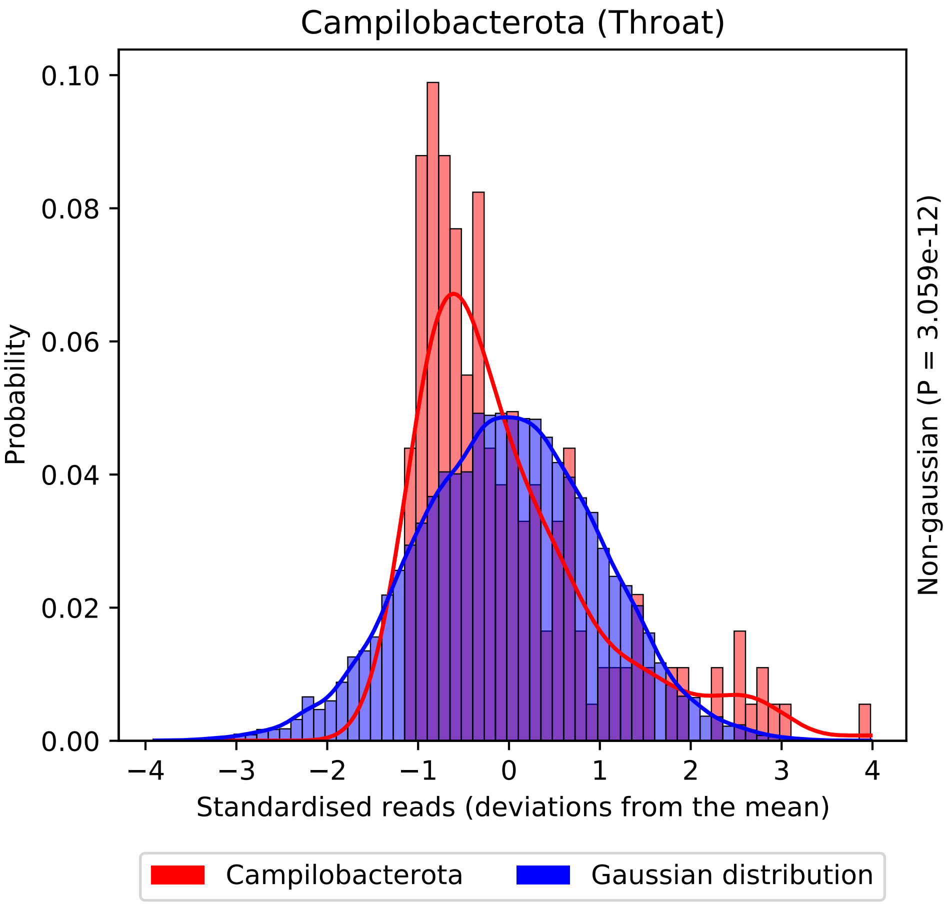
Campilobacterota (Throat)
Probability
0.00
0.02
0.04
0.06
0.08
0.10
−4
−3
−2
−1
0
1
2
3
4
Standardised reads (deviations from the mean)
Non-gaussian (P = 3.059e-12)
Campilobacterota
Gaussian distribution

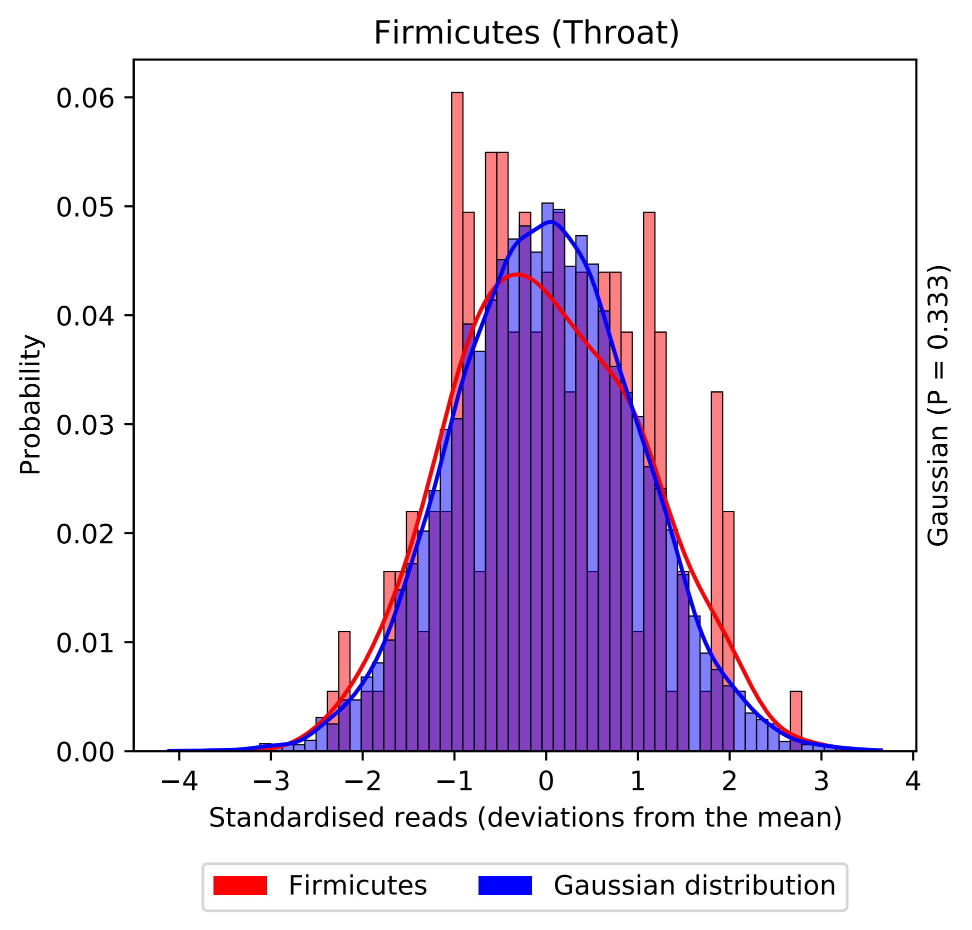

Firmicutes (Throat)
Probability
0.06
0.05
0.04
0.03
0.02
0.01
0.00
−4
−3
−2
−1
0
1
2
3
4
Standardised reads (deviations from the mean)
Gaussian (P = 0.333)
Firmicutes
Gaussian distribution

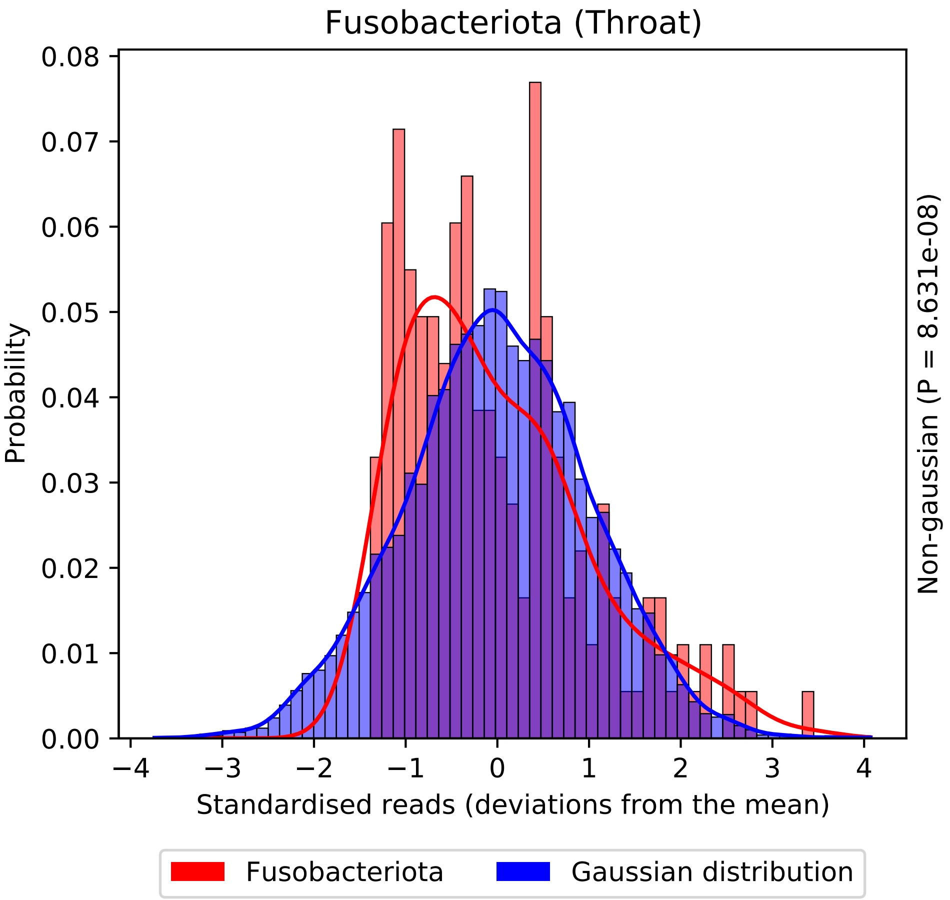
Fusobacteriota (Throat)
Probability
0.08
0.07
0.06
0.05
0.04
0.03
0.02
0.01
0.00
−4
−3
−2
−1
0
1
2
3
4
Standardised reads (deviations from the mean)
Non-gaussian (P = 8.631e-08)
Fusobacteriota
Gaussian distribution

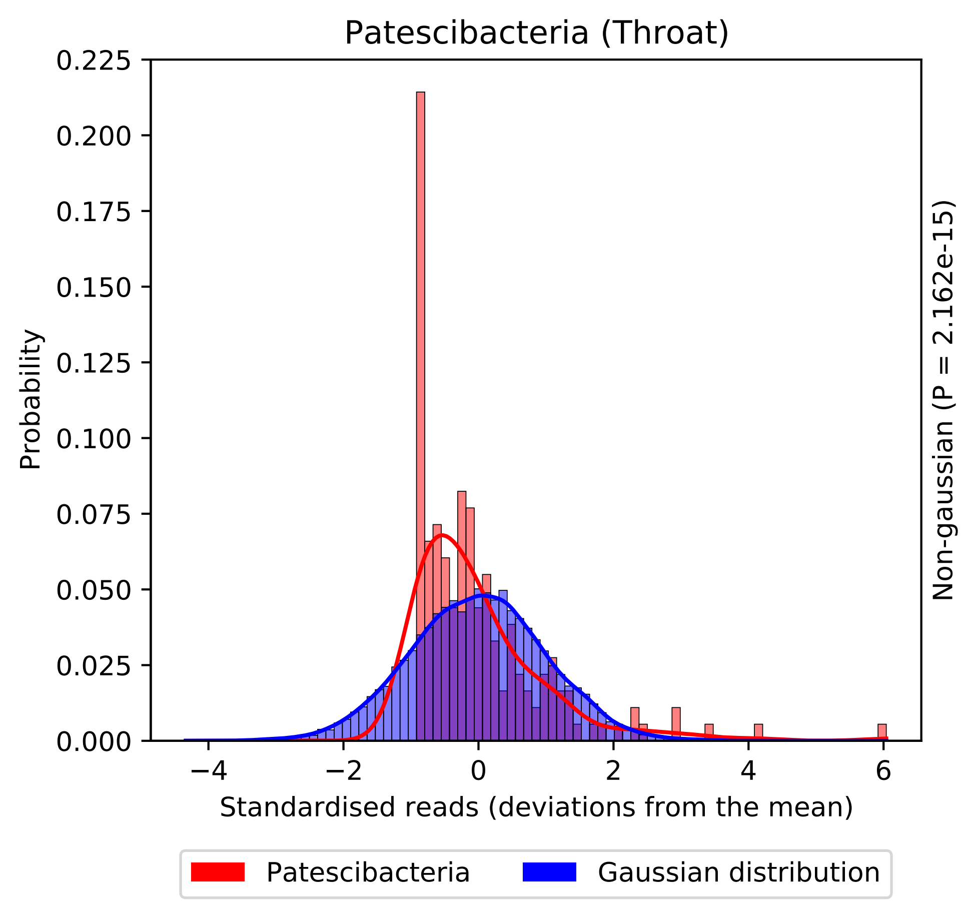
Patescibacteria (Throat)
0.225
0.200
0.175
0.150
0.125
0.100
0.075
0.050
0.025
0.000
Probability
−4
−2
0
2
4
6
Standardised reads (deviations from the mean)
Non-gaussian (P = 2.162e-15)
Patescibacteria
Gaussian distribution

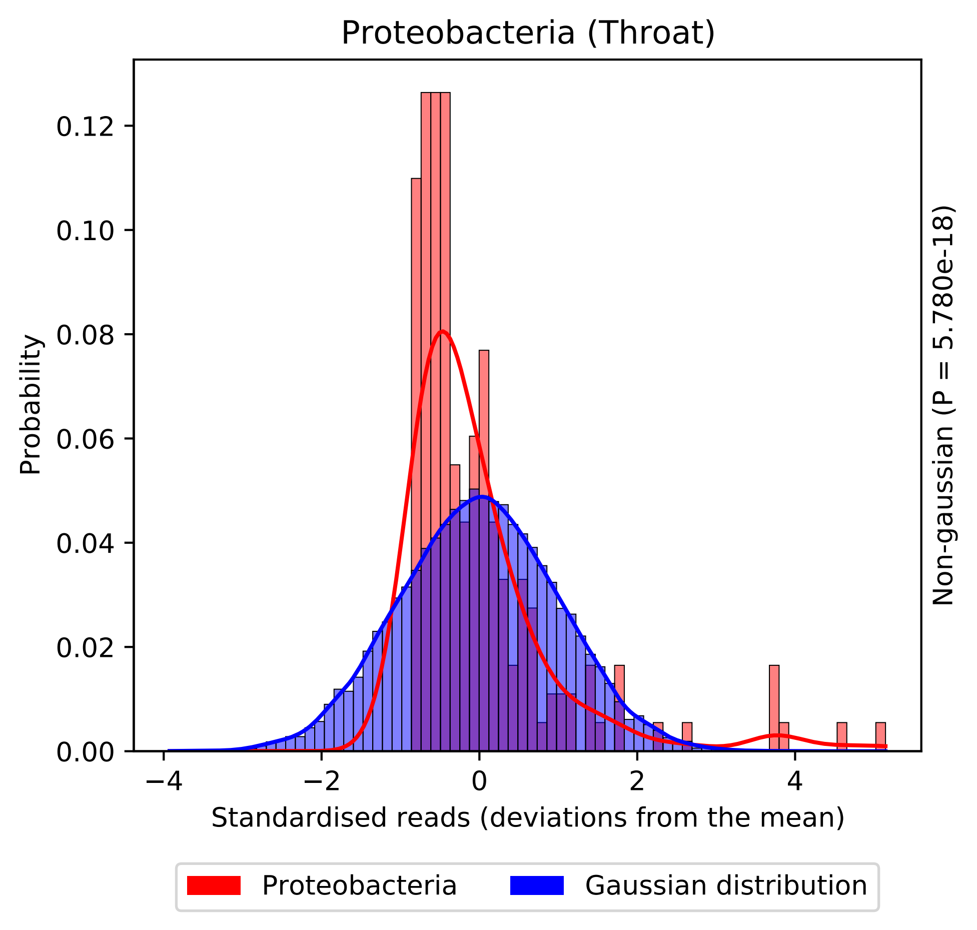

Proteobacteria (Throat)
Probability
0.12
0.10
0.08
0.06
0.04
0.02
0.00
−4
−2
0
2
4
Standardised reads (deviations from the mean)
Non-gaussian (P = 5.780e-18)
Proteobacteria
Gaussian distribution

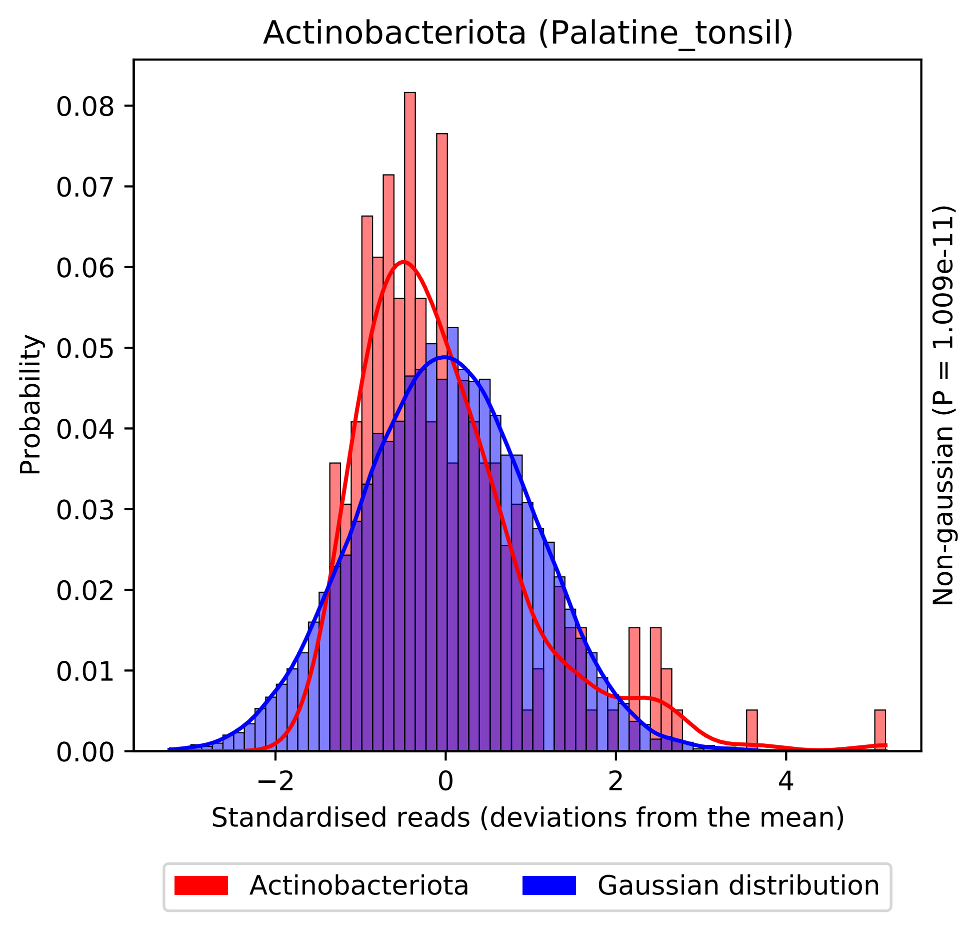

Actinobacteriota (Palatine_tonsil)
Probability
0.08
0.07
0.06
0.05
0.04
0.03
0.02
0.01
0.00
−2
0
2
4
Standardised reads (deviations from the mean)
Non-gaussian (P = 1.009e-11)
Actinobacteriota
Gaussian distribution

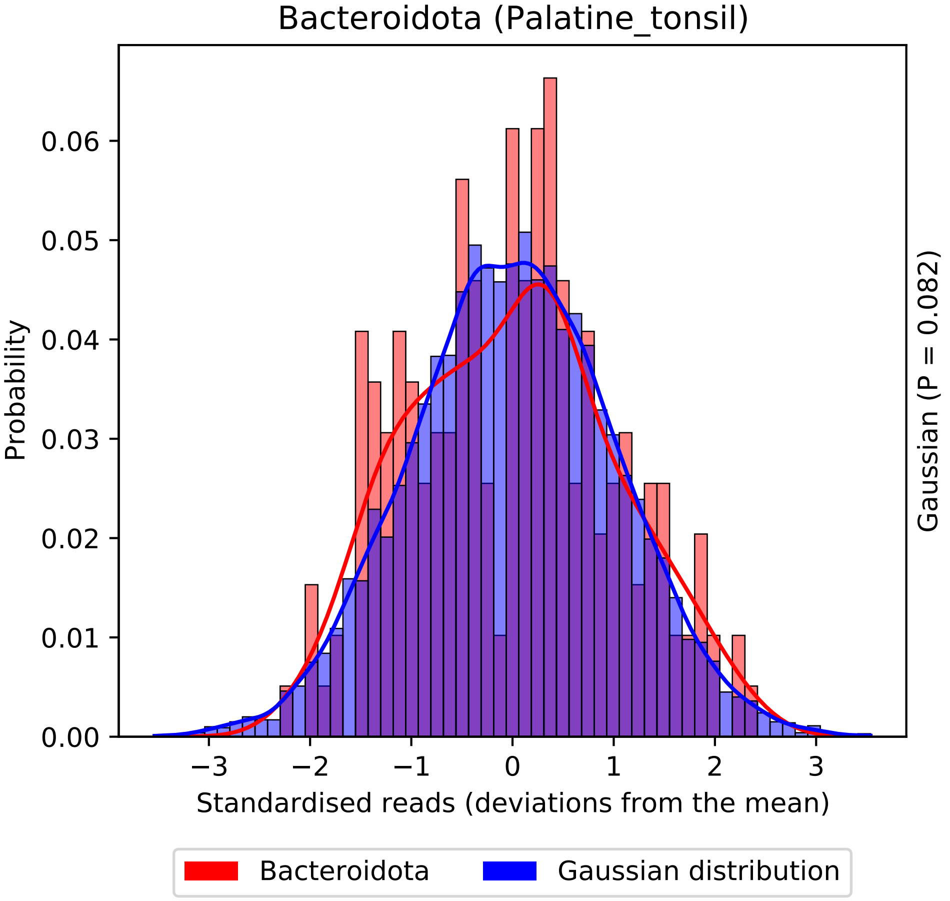

Bacteroidota (Palatine_tonsil)
Probability
0.00
0.01
0.02
0.03
0.04
0.05
0.06
−3
−2
−1
0
1
2
3
Standardised reads (deviations from the mean)
Gaussian (P = 0.082)
Bacteroidota
Gaussian distribution

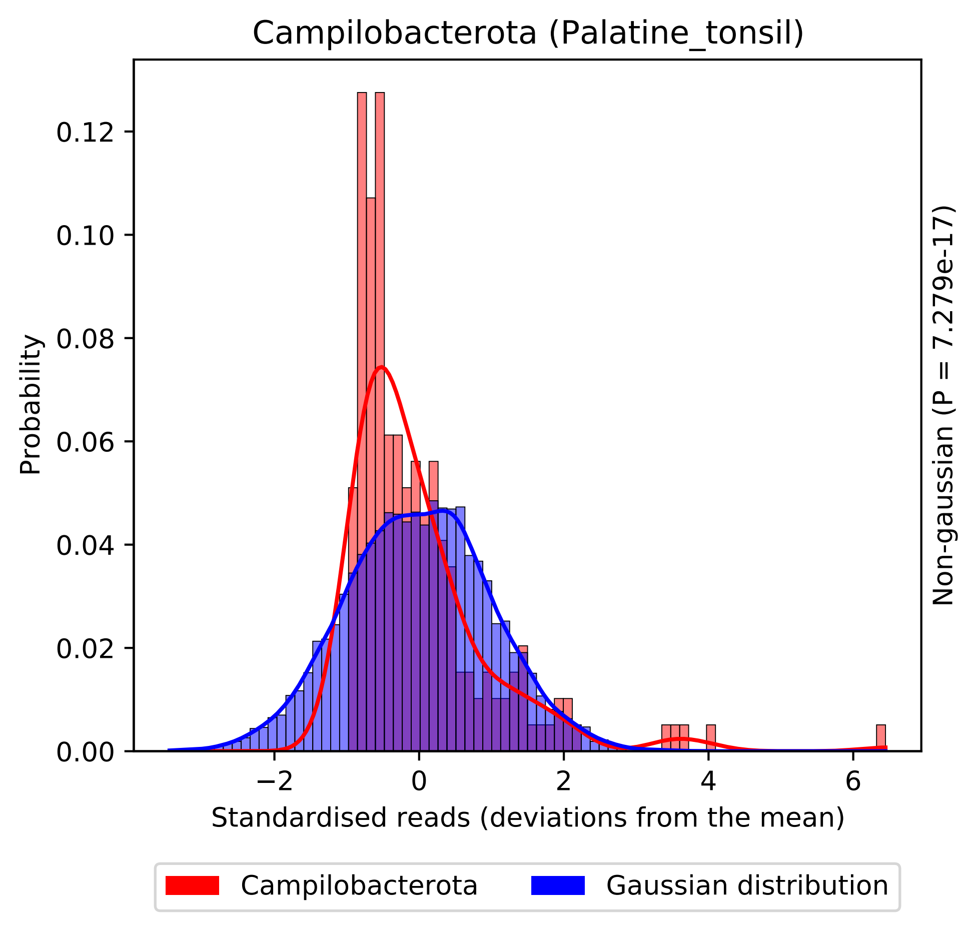

Campilobacterota (Palatine_tonsil)
Probability
0.00
0.02
0.04
0.06
0.08
0.10
0.12
−2
0
2
4
6
Standardised reads (deviations from the mean)
Non-gaussian (P = 7.279e-17)
Campilobacterota
Gaussian distribution

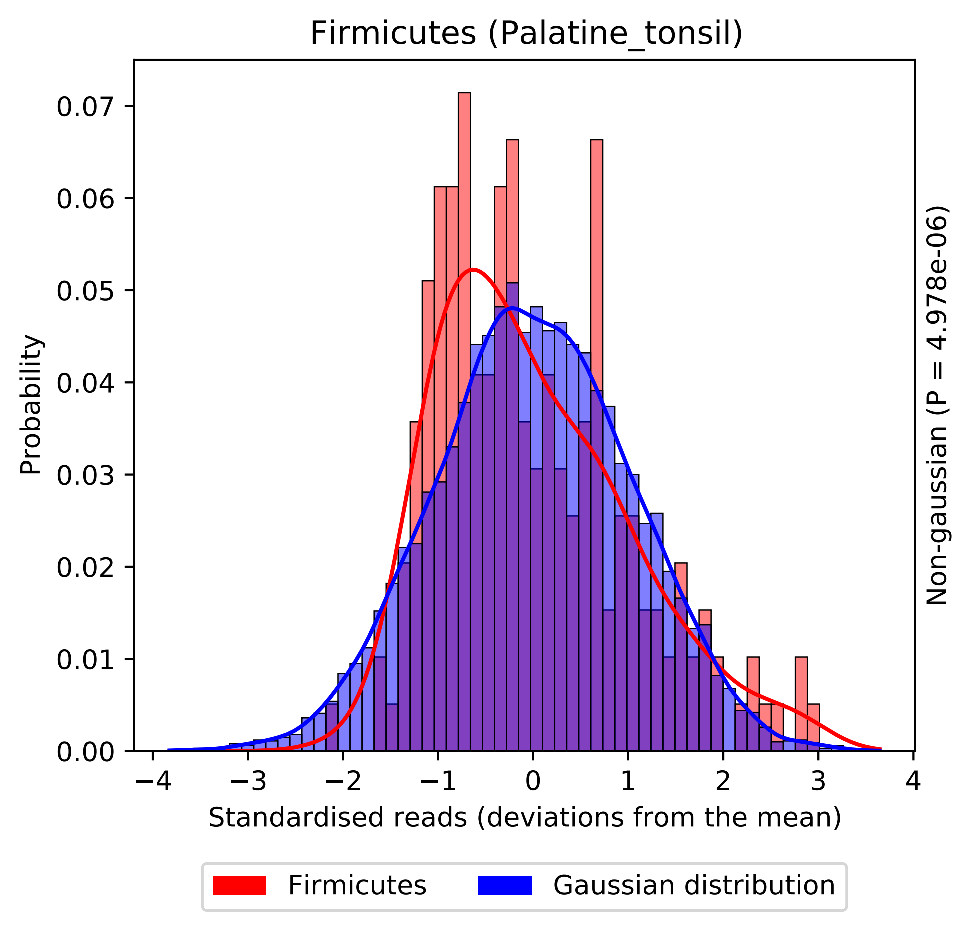

Firmicutes (Palatine_tonsil)
Probability
0.00
0.01
0.02
0.03
0.04
0.05
0.06
0.07
−4
−3
−2
−1
0
1
2
3
4
Standardised reads (deviations from the mean)
Non-gaussian (P = 4.978e-06)
Firmicutes
Gaussian distribution

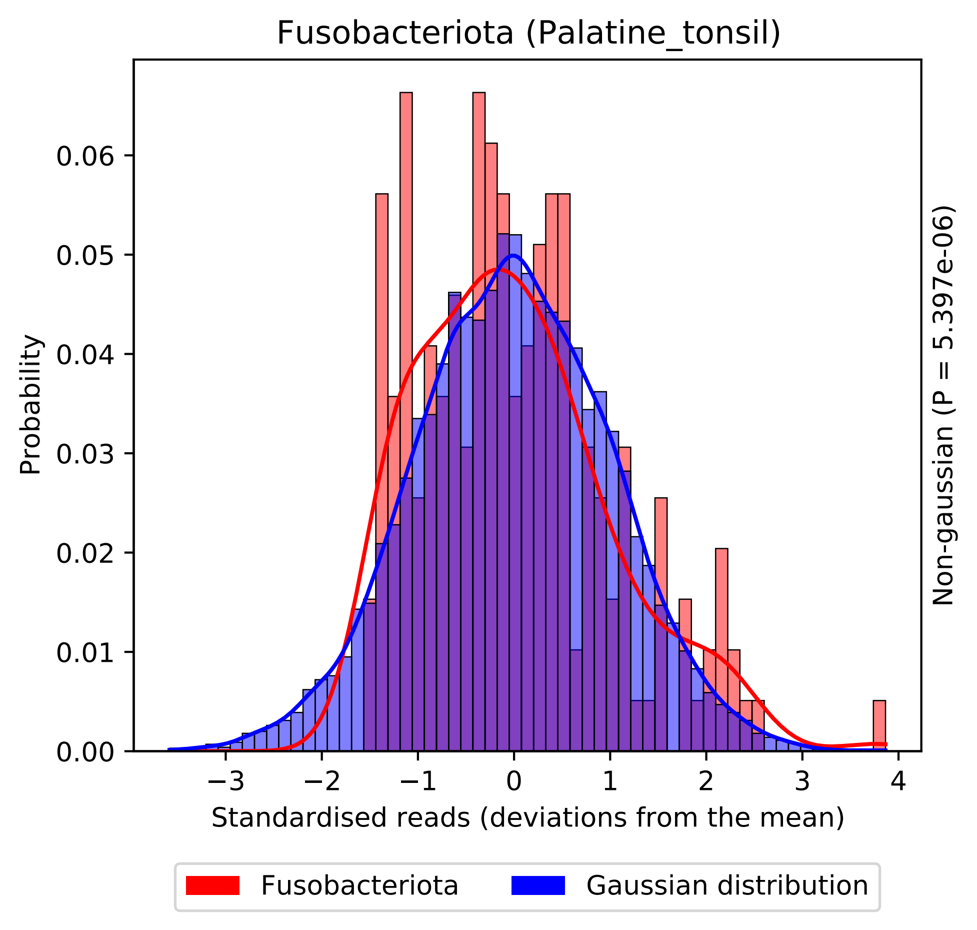
Fusobacteriota (Palatine_tonsil)
Probability
0.00
0.01
0.02
0.03
0.04
0.05
0.06
−3
−2
−1
0
1
2
3
4
Standardised reads (deviations from the mean)
Non-gaussian (P = 5.397e-06)
Fusobacteriota
Gaussian distribution

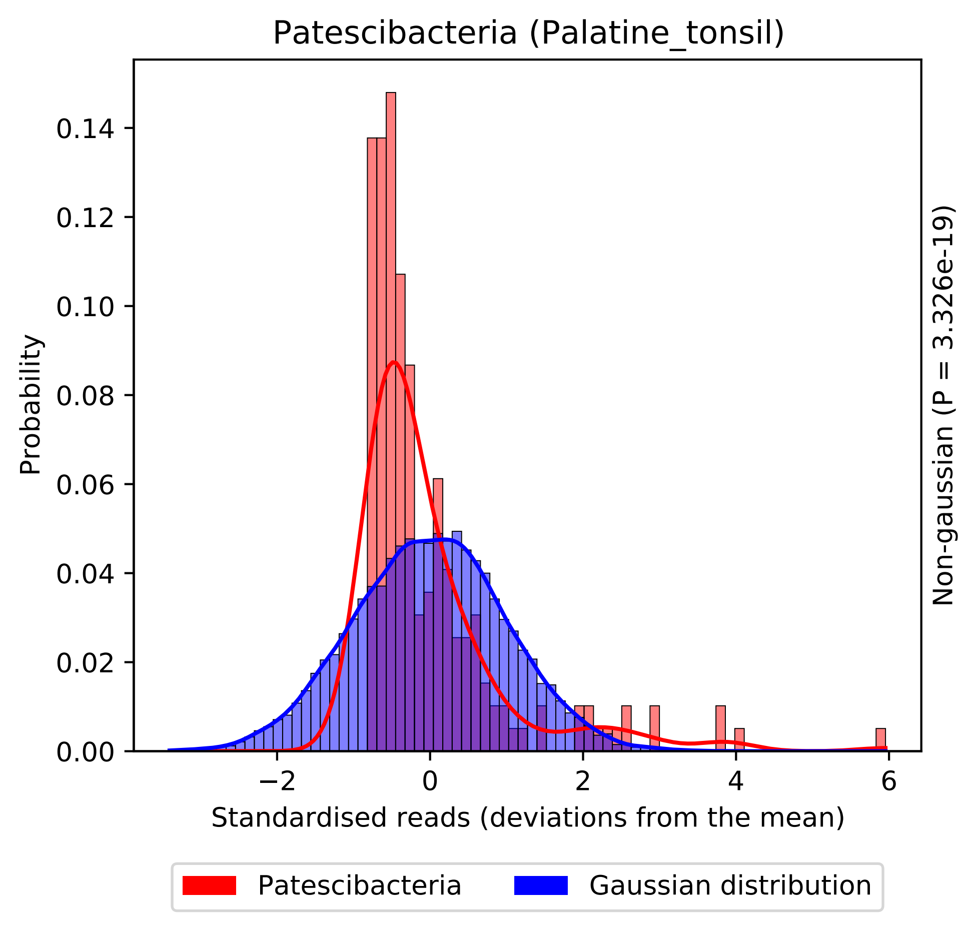

Patescibacteria (Palatine_tonsil)
Probability
0.00
0.02
0.04
0.06
0.08
0.10
0.12
0.14
−2
0
2
4
6
Standardised reads (deviations from the mean)
Non-gaussian (P = 3.326e-19)
Patescibacteria
Gaussian distribution

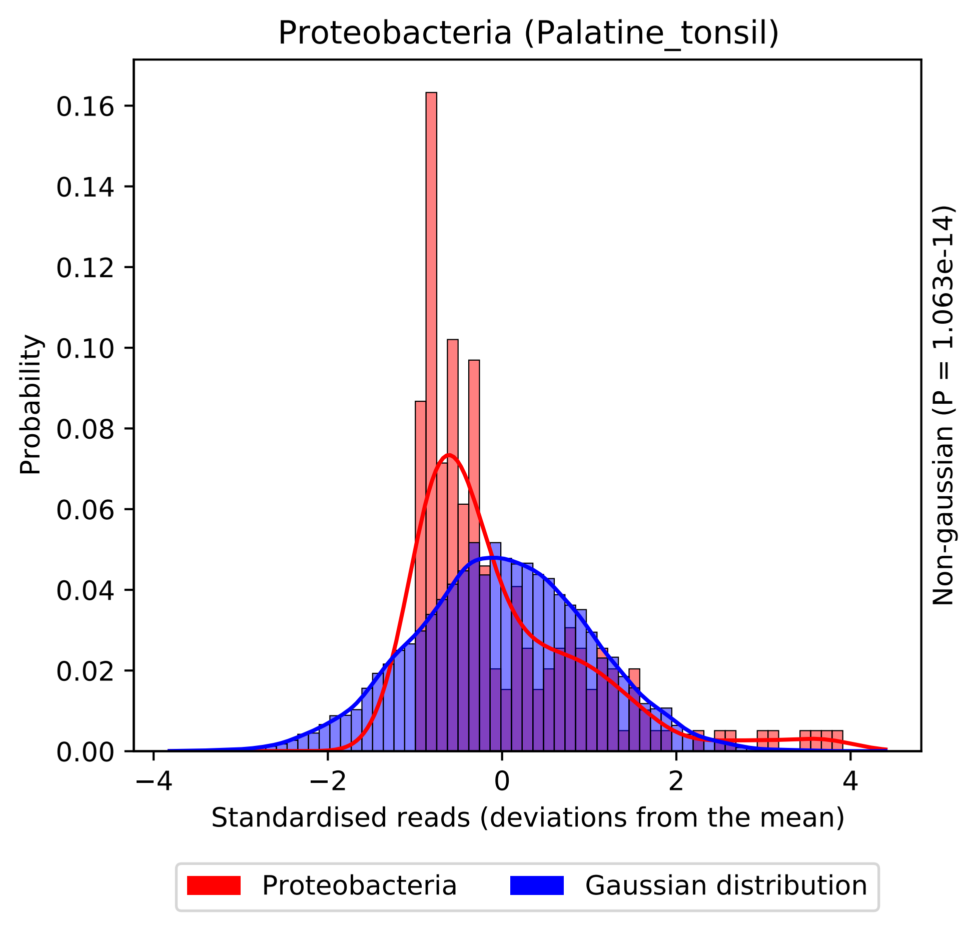
Proteobacteria (Palatine_tonsil)
Probability
0.00
0.02
0.04
0.06
0.08
0.10
0.12
0.14
0.16
−4
−2
0
2
4
Standardised reads (deviations from the mean)
Non-gaussian (P = 1.063e-14)
Proteobacteria
Gaussian distribution

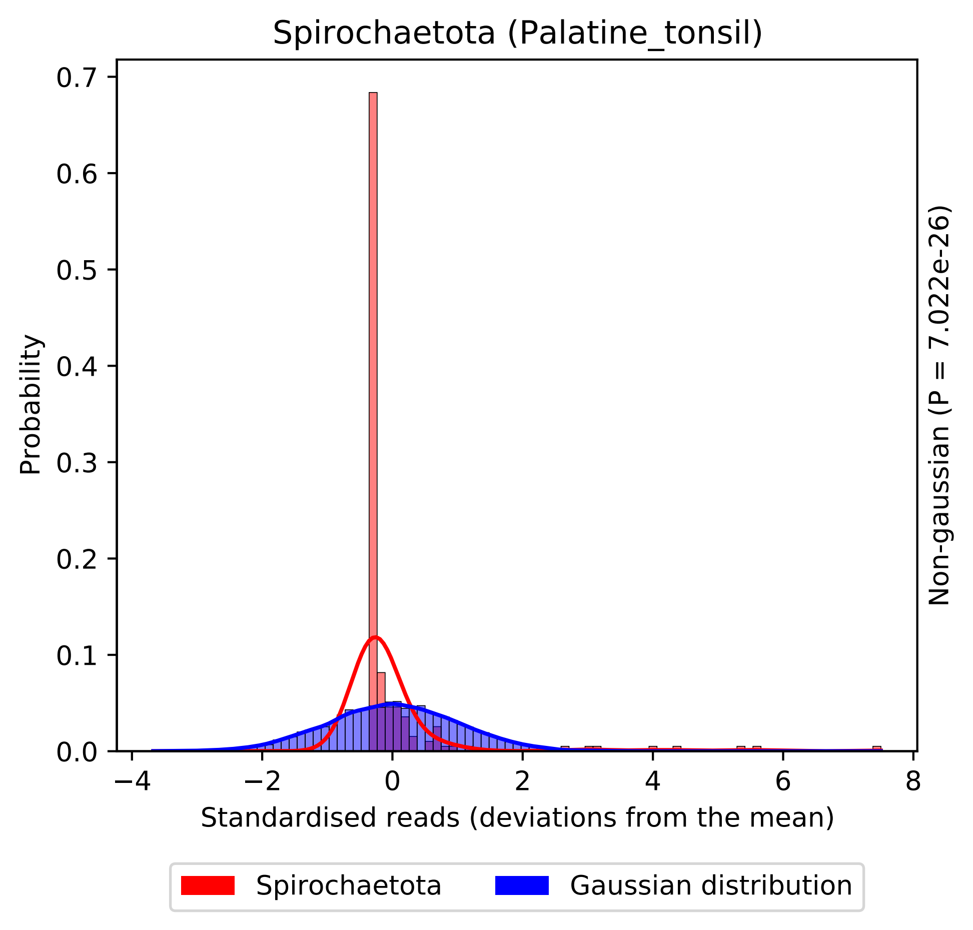

Spirochaetota (Palatine_tonsil)
Probability
0.0
0.1
0.2
0.3
0.4
0.5
0.6
0.7
−4
−2
0
2
4
6
8
Standardised reads (deviations from the mean)
Non-gaussian (P = 7.022e-26)
Spirochaetota
Gaussian distribution

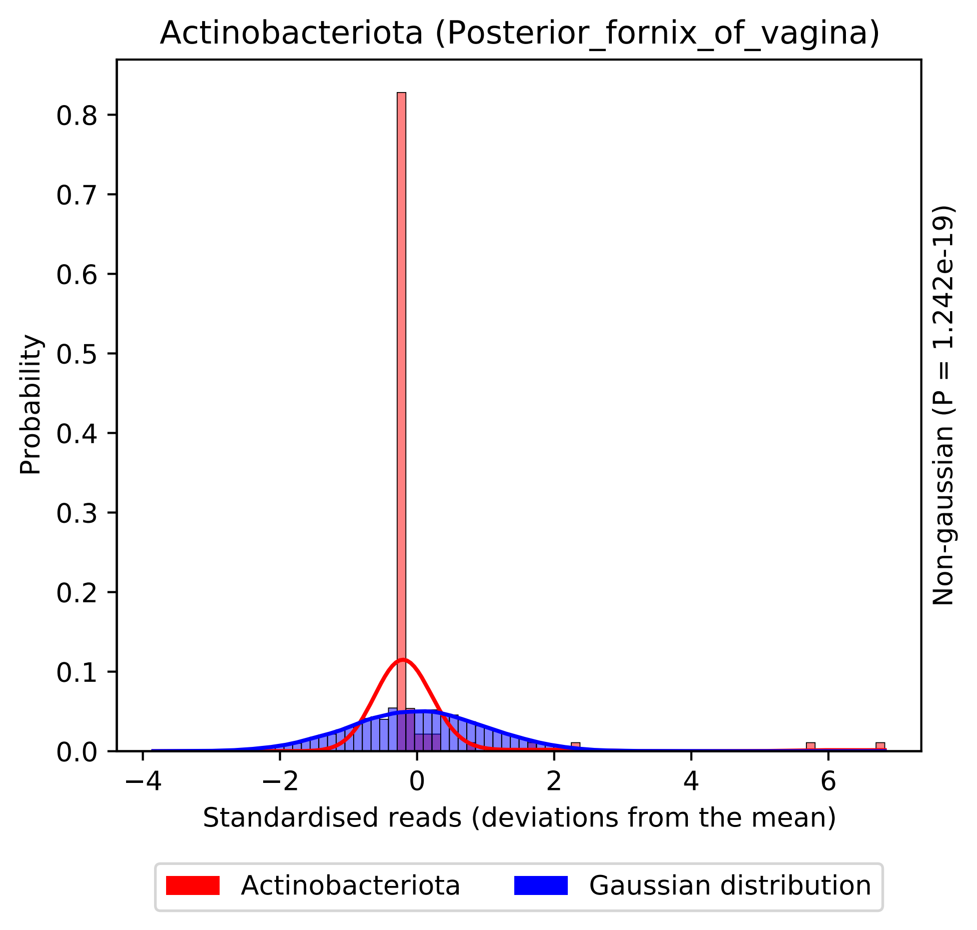
Actinobacteriota (Posterior_fornix_of_vagina)
Probability
0.0
0.1
0.2
0.3
0.4
0.5
0.6
0.7
0.8
−4
−2
0
2
4
6
Standardised reads (deviations from the mean)
Non-gaussian (P = 1.242e-19)
Actinobacteriota
Gaussian distribution

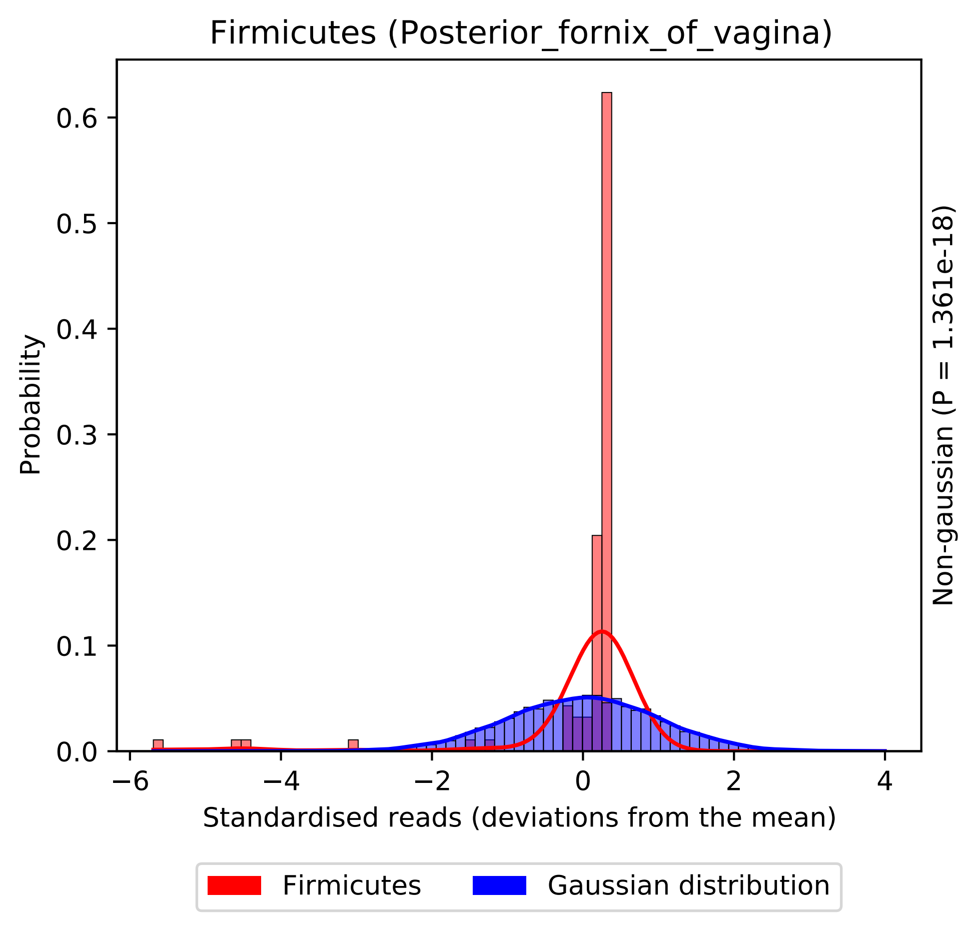
Firmicutes (Posterior_fornix_of_vagina)
Probability
0.0
0.1
0.2
0.3
0.4
0.5
0.6
−6
−4
−2
0
2
4
Standardised reads (deviations from the mean)
Non-gaussian (P = 1.361e-18)
Firmicutes
Gaussian distribution

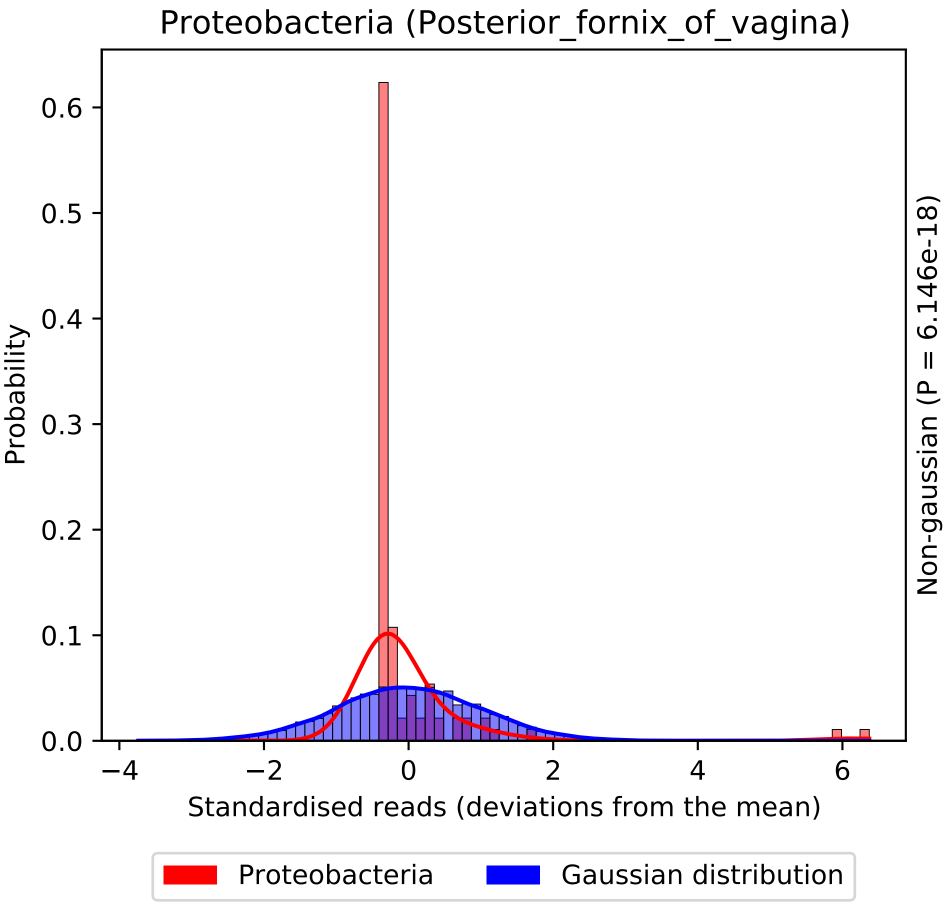
Proteobacteria (Posterior_fornix_of_vagina)
Probability
0.0
0.1
0.2
0.3
0.4
0.5
0.6
−4
−2
0
2
4
6
Standardised reads (deviations from the mean)
Non-gaussian (P = 6.146e-18)
Proteobacteria
Gaussian distribution

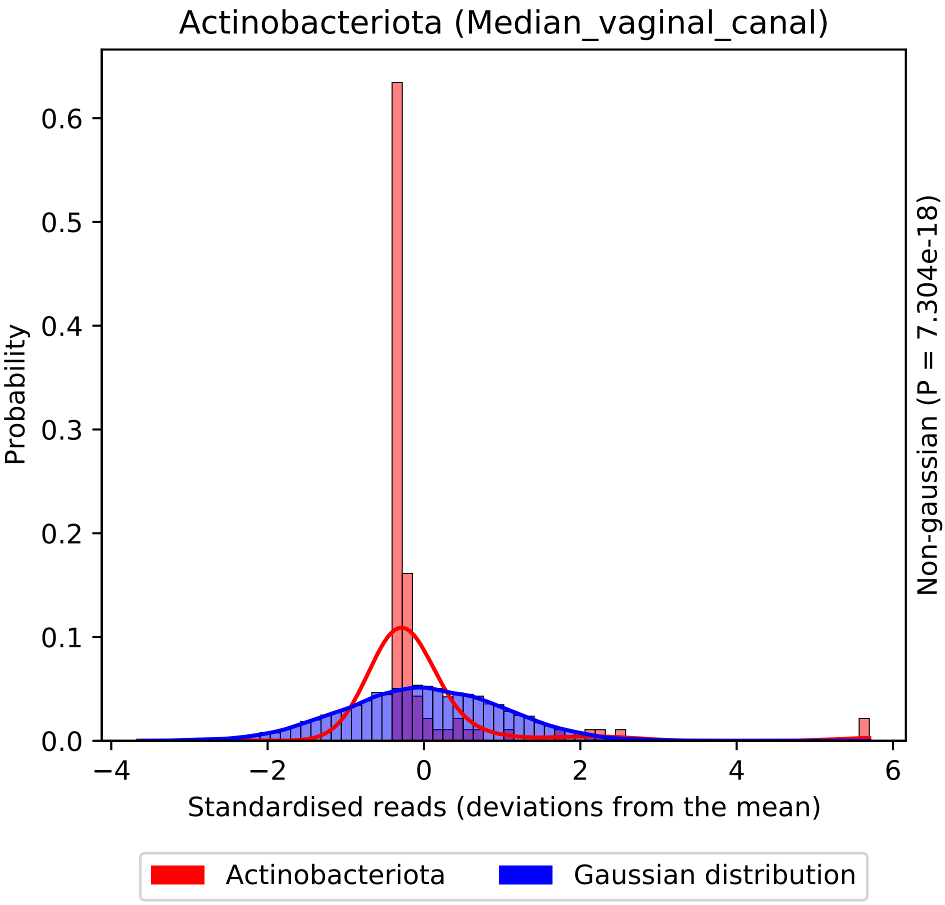

Actinobacteriota (Median_vaginal_canal)
Probability
0.0
0.1
0.2
0.3
0.4
0.5
0.6
−4
−2
0
2
4
6
Standardised reads (deviations from the mean)
Non-gaussian (P = 7.304e-18)
Actinobacteriota
Gaussian distribution

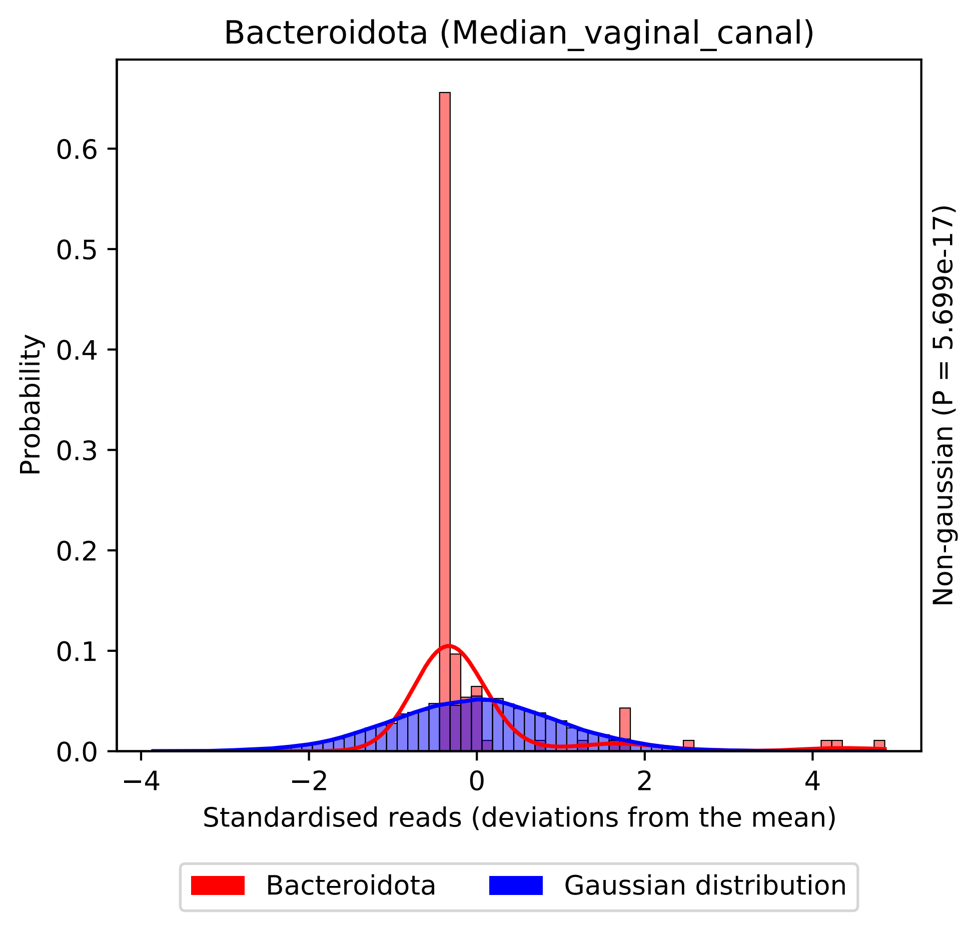

Bacteroidota (Median_vaginal_canal)
Probability
0.0
0.1
0.2
0.3
0.4
0.5
0.6
−4
−2
0
2
4
Standardised reads (deviations from the mean)
Non-gaussian (P = 5.699e-17)
Bacteroidota
Gaussian distribution

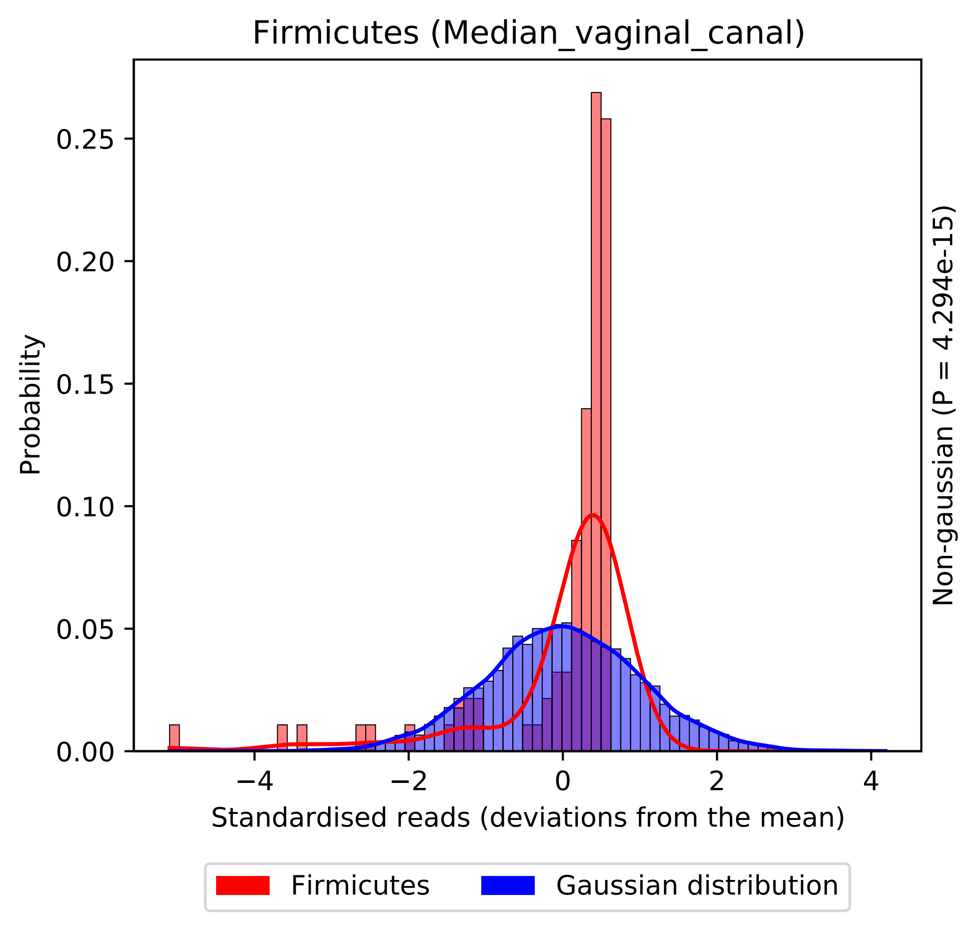
Firmicutes (Median_vaginal_canal)
Probability
0.00
0.05
0.10
0.15
0.20
0.25
−4
−2
0
2
4
Standardised reads (deviations from the mean)
Non-gaussian (P = 4.294e-15)
Firmicutes
Gaussian distribution

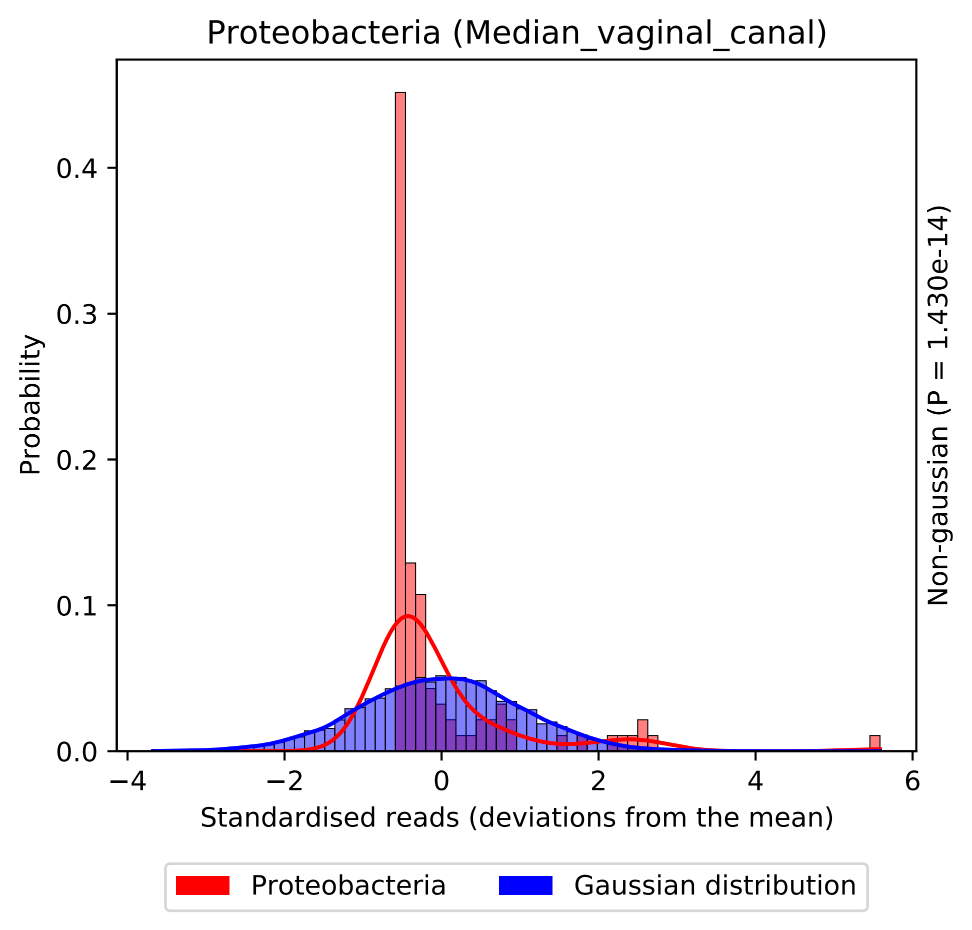

Proteobacteria (Median_vaginal_canal)
Probability
0.0
0.1
0.2
0.3
0.4
−4
−2
0
2
4
6
Standardised reads (deviations from the mean)
Non-gaussian (P = 1.430e-14)
Proteobacteria
Gaussian distribution

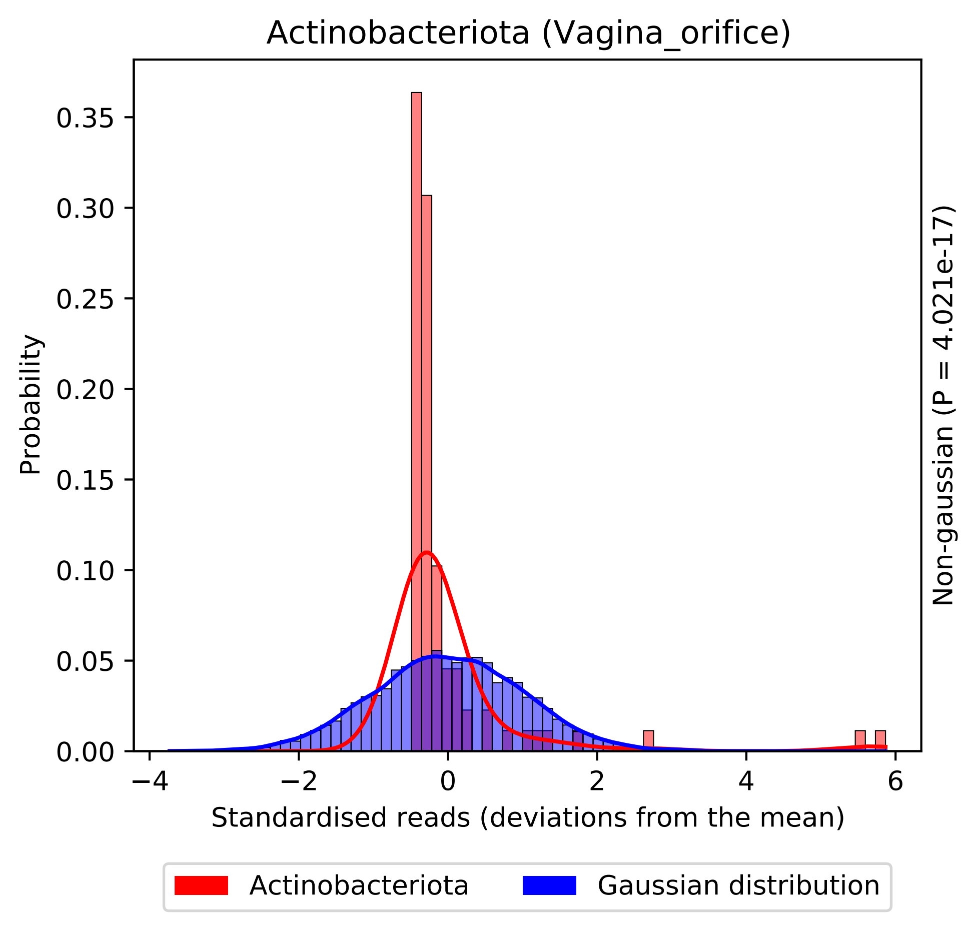

Actinobacteriota (Vagina_orifice)
Probability
0.00
0.05
0.10
0.15
0.20
0.25
0.30
0.35
−4
−2
0
2
4
6
Standardised reads (deviations from the mean)
Non-gaussian (P = 4.021e-17)
Actinobacteriota
Gaussian distribution

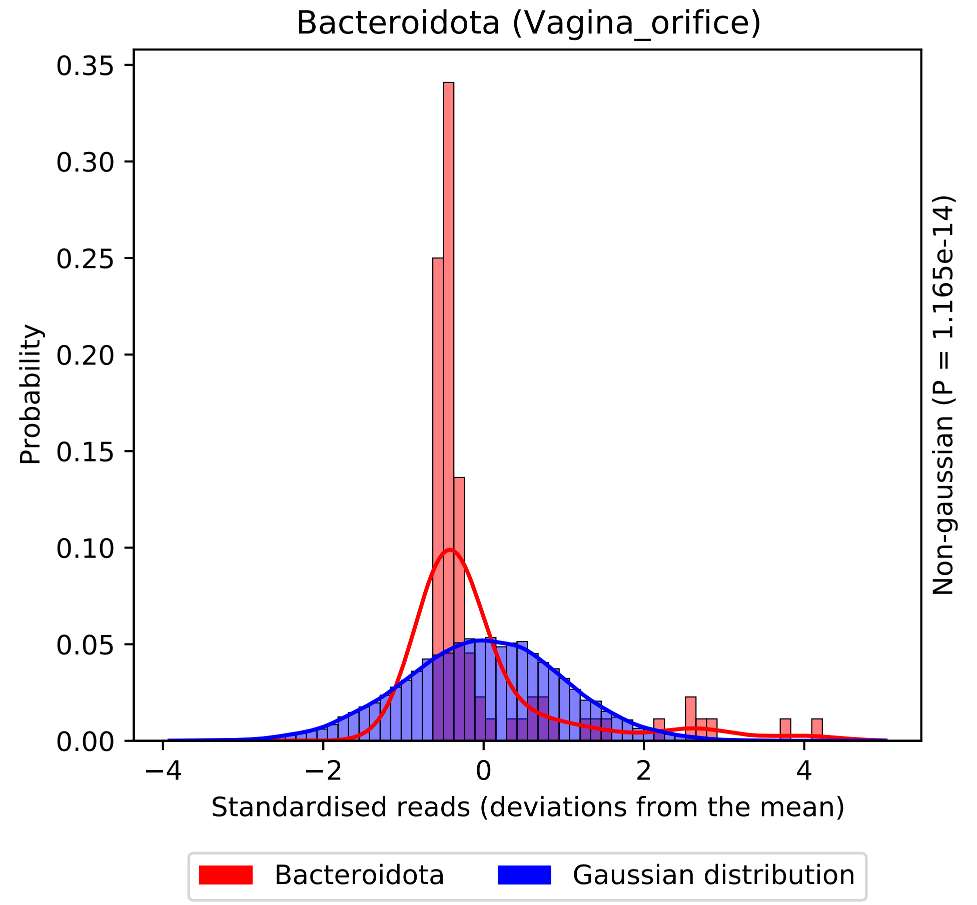
Bacteroidota (Vagina_orifice)
Probability
Standardised reads (deviations from the mean)
Non-gaussian (P = 1.165e-14)
0.00
0.05
0.10
0.15
0.20
0.25
0.30
0.35
−4
−2
0
2
4
Bacteroidota
Gaussian distribution

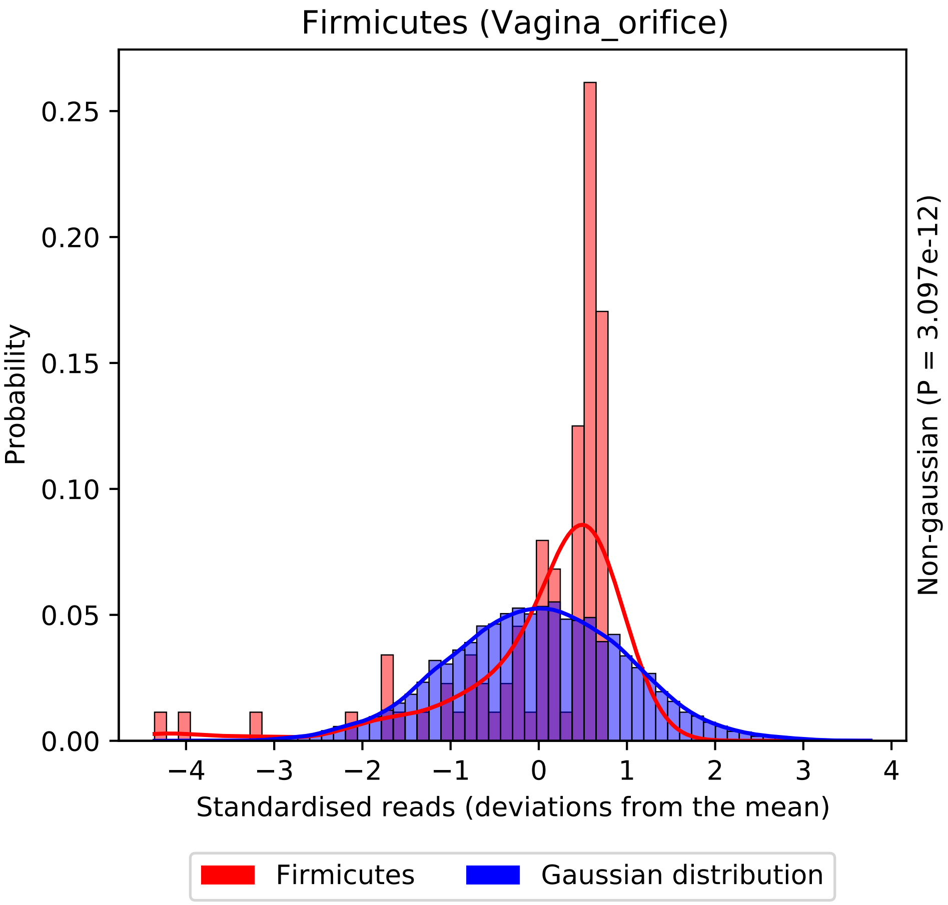
Firmicutes (Vagina_orifice)
Probability
0.00
0.05
0.10
0.15
0.20
0.25
−4
−3
−2
−1
0
1
2
3
4
Standardised reads (deviations from the mean)
Non-gaussian (P = 3.097e-12)
Firmicutes
Gaussian distribution

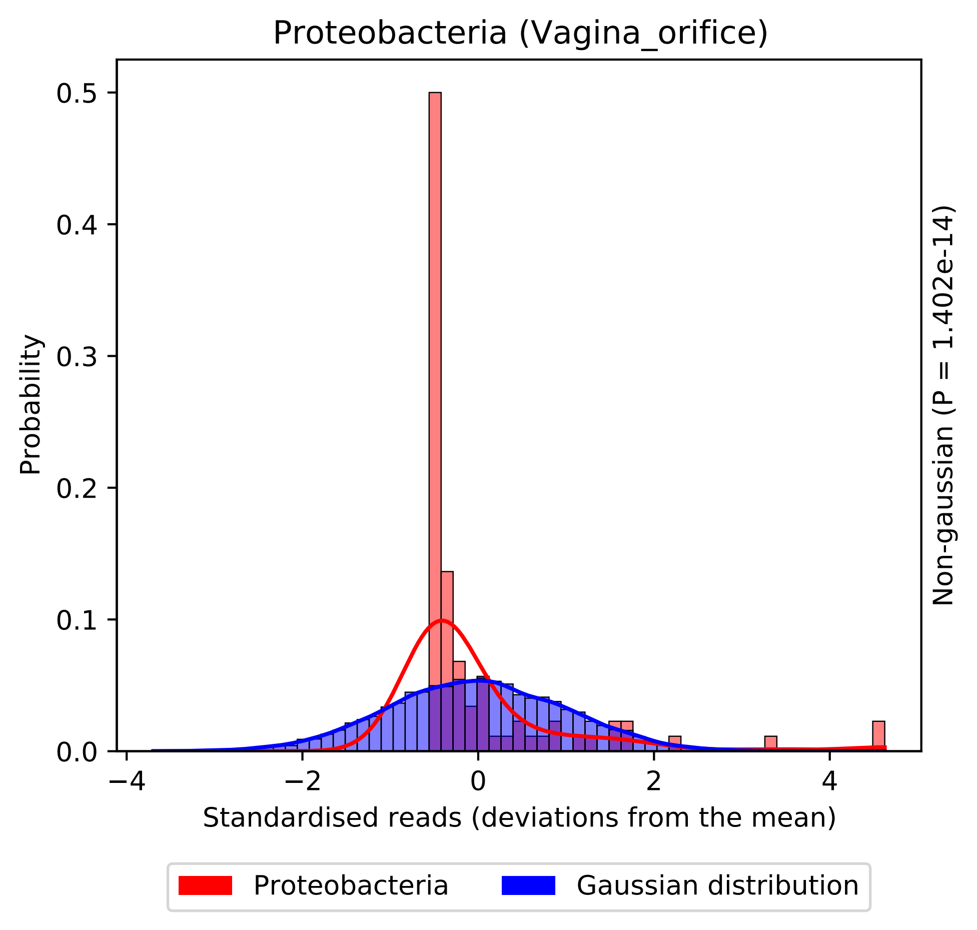
Proteobacteria (Vagina_orifice)
Probability
0.0
0.1
0.2
0.3
0.4
0.5
−4
−2
0
2
4
Standardised reads (deviations from the mean)
Non-gaussian (P = 1.402e-14)
Proteobacteria
Gaussian distribution

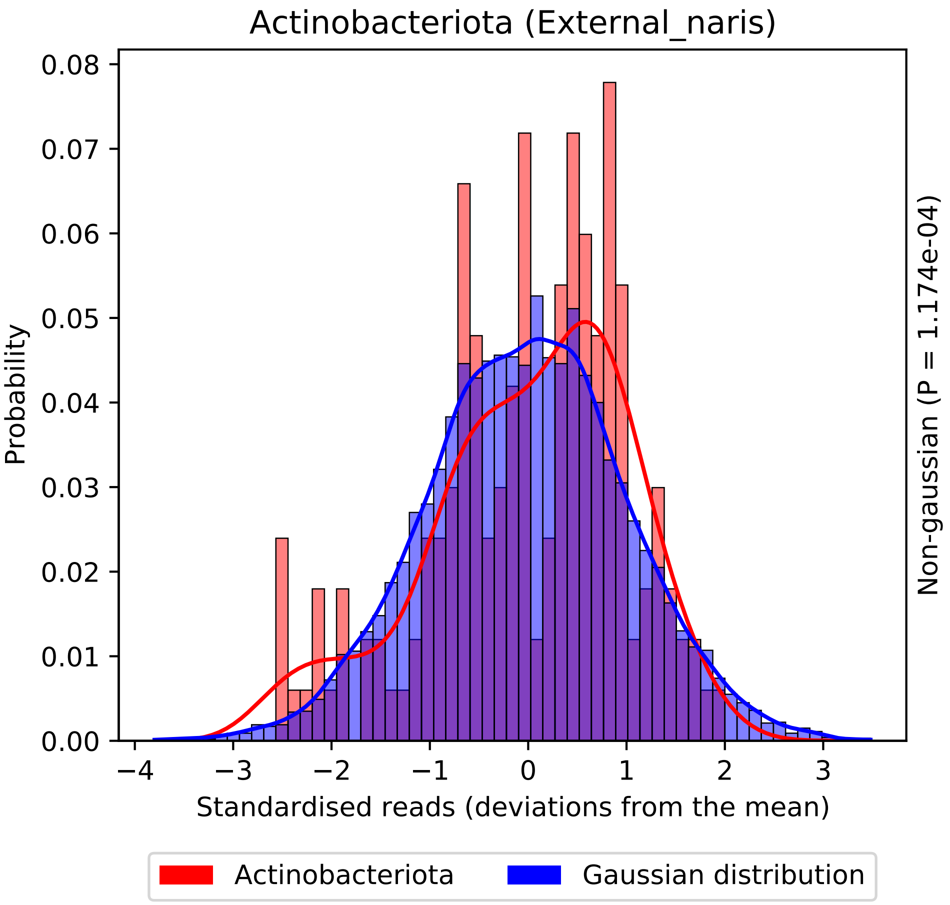

Actinobacteriota (External_naris)
Probability
0.00
0.01
0.02
0.03
0.04
0.05
0.06
0.07
0.08
−4
−3
−2
−1
0
1
2
3
Standardised reads (deviations from the mean)
Non-gaussian (P = 1.174e-04)
Actinobacteriota
Gaussian distribution

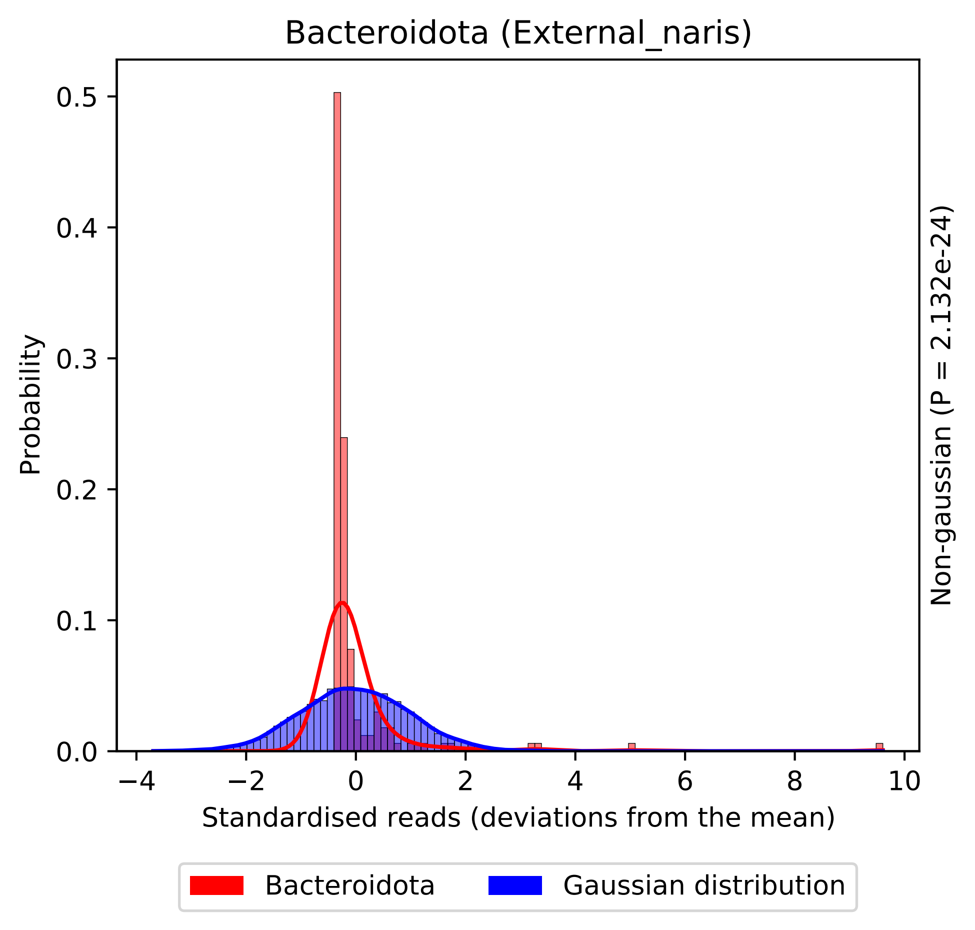
Bacteroidota (External_naris)
Probability
0.0
0.1
0.2
0.3
0.4
0.5
−4
−2
0
2
4
6
8
10
Standardised reads (deviations from the mean)
Non-gaussian (P = 2.132e-24)
Bacteroidota
Gaussian distribution

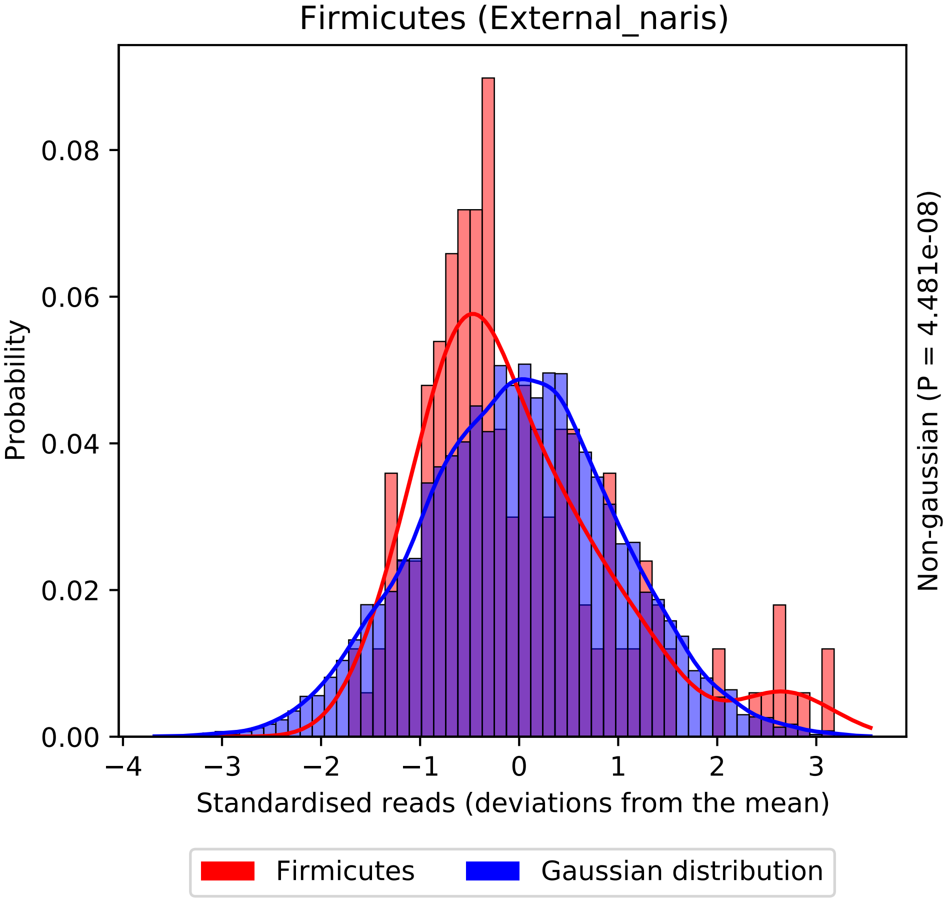

Firmicutes (External_naris)
Probability
0.00
0.02
0.04
0.06
0.08
−4
−3
−2
−1
0
1
2
3
Standardised reads (deviations from the mean)
Non-gaussian (P = 4.481e-08)
Firmicutes
Gaussian distribution

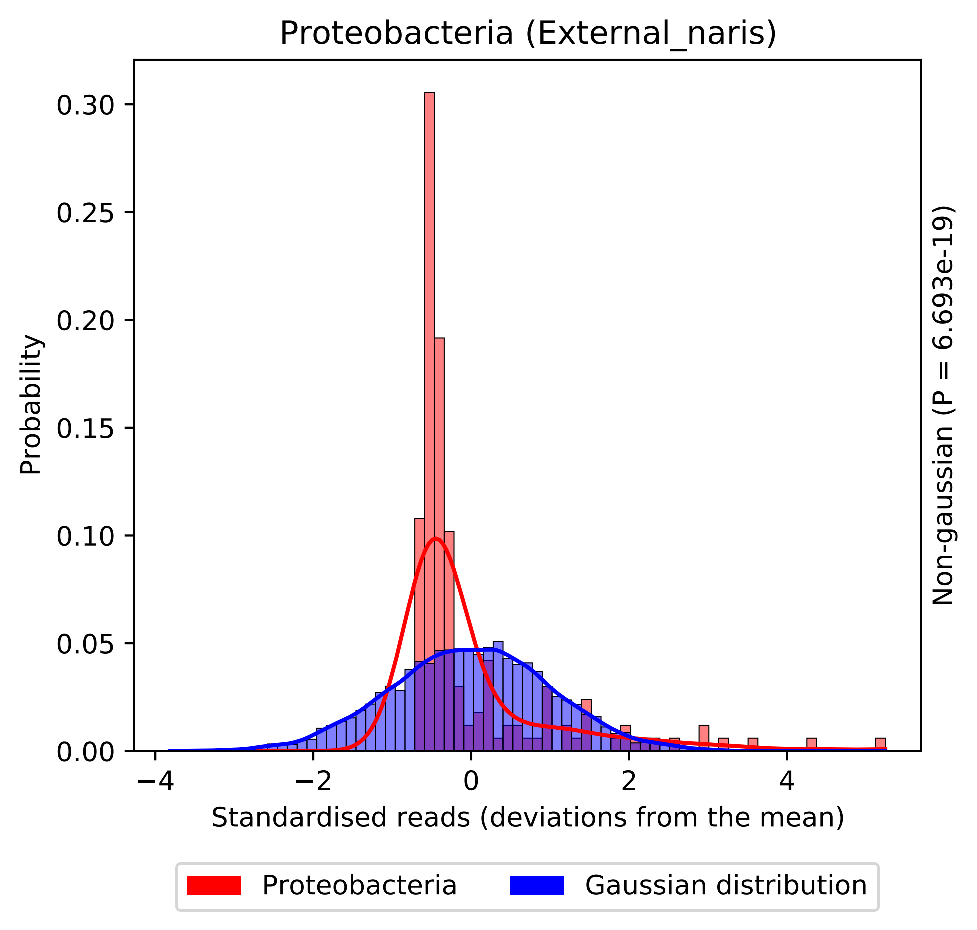

Proteobacteria (External_naris)
0.30
0.25
0.20
0.15
0.10
0.05
0.00
Probability
−4
−2
0
2
4
Standardised reads (deviations from the mean)
Non-gaussian (P = 6.693e-19)
Proteobacteria
Gaussian distribution

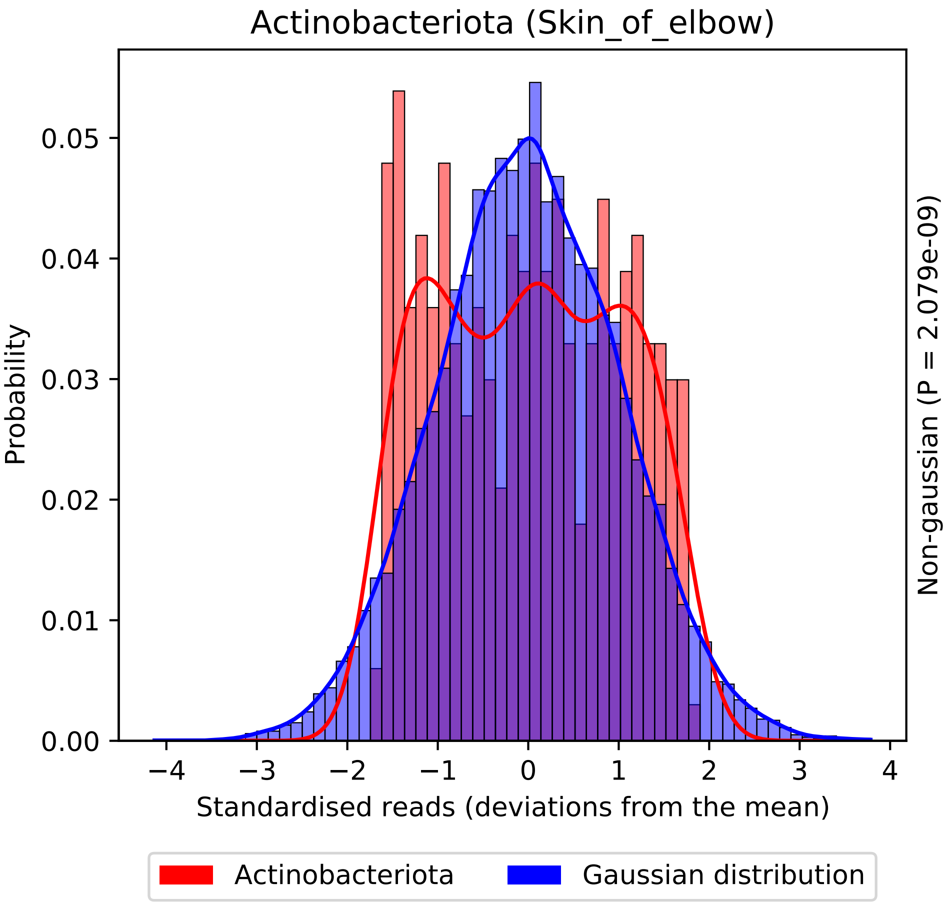

Actinobacteriota (Skin_of_elbow)
Probability
0.00
0.01
0.02
0.03
0.04
0.05
−4
−3
−2
−1
0
1
2
3
4
Standardised reads (deviations from the mean)
Non-gaussian (P = 2.079e-09)
Actinobacteriota
Gaussian distribution

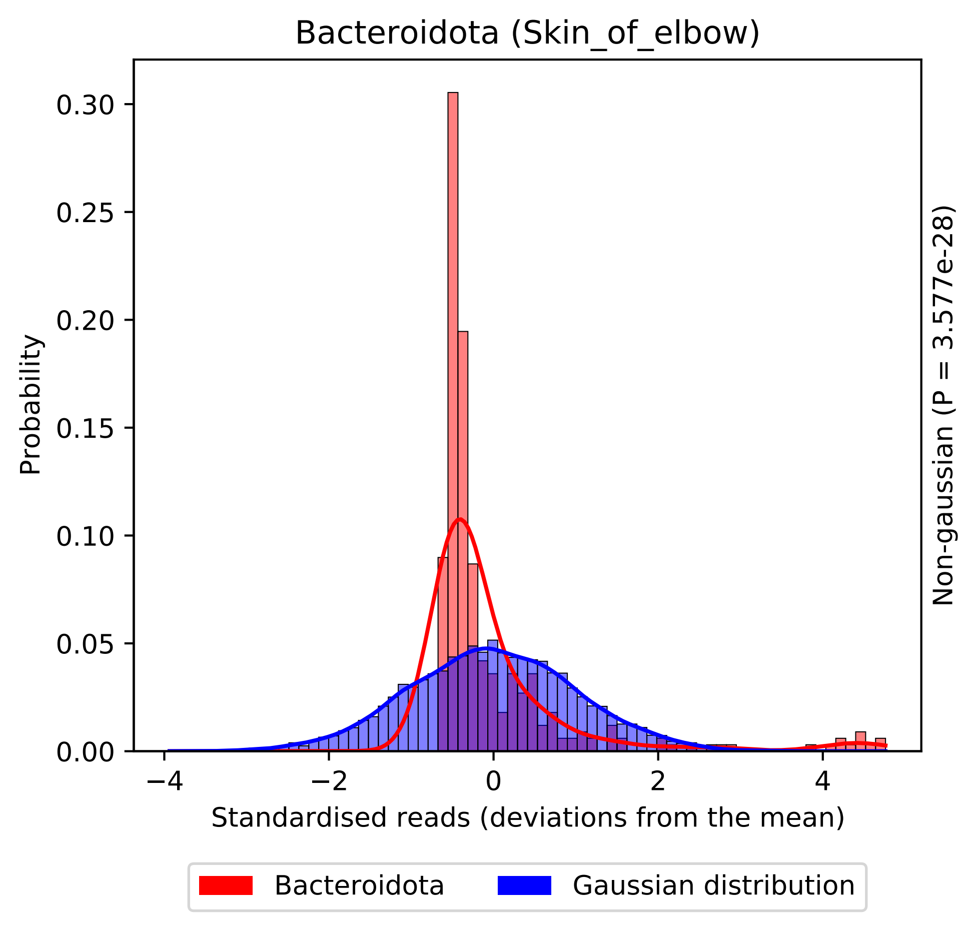

Bacteroidota (Skin_of_elbow)
Probability
0.30
0.25
0.20
0.15
0.10
0.05
0.00
−4
−2
0
2
4
Standardised reads (deviations from the mean)
Non-gaussian (P = 3.577e-28)
Bacteroidota
Gaussian distribution

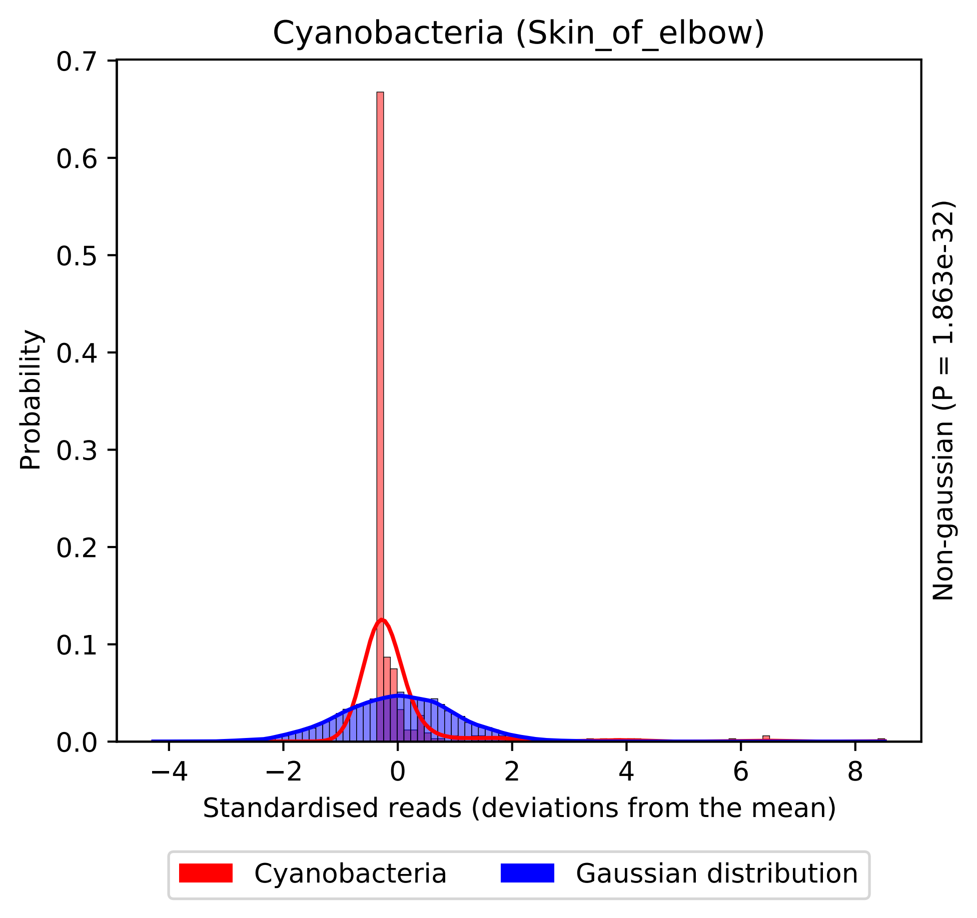

Cyanobacteria (Skin_of_elbow)
Probability
0.0
0.1
0.2
0.3
0.4
0.5
0.6
0.7
−4
−2
0
2
4
6
8
Standardised reads (deviations from the mean)
Non-gaussian (P = 1.863e-32)
Cyanobacteria
Gaussian distribution

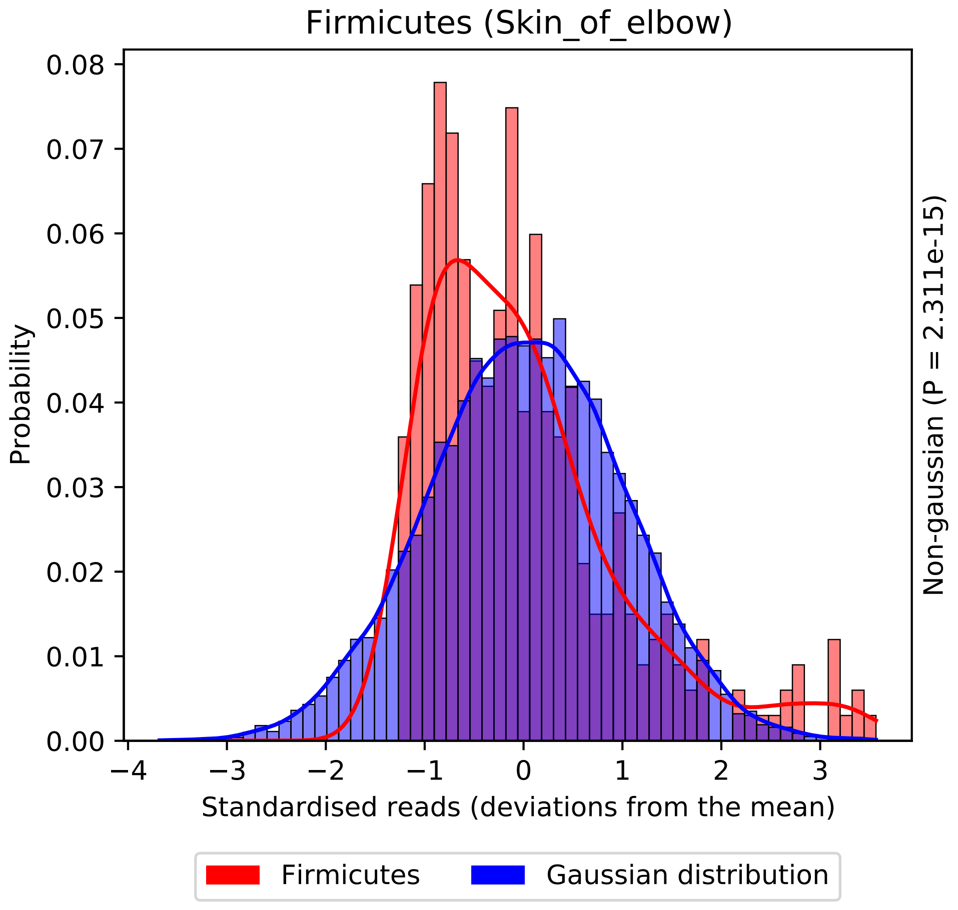
Firmicutes (Skin_of_elbow)
Probability
0.00
0.01
0.02
0.03
0.04
0.05
0.06
0.07
0.08
−4
−3
−2
−1
0
1
2
3
Standardised reads (deviations from the mean)
Non-gaussian (P = 2.311e-15)
Firmicutes
Gaussian distribution

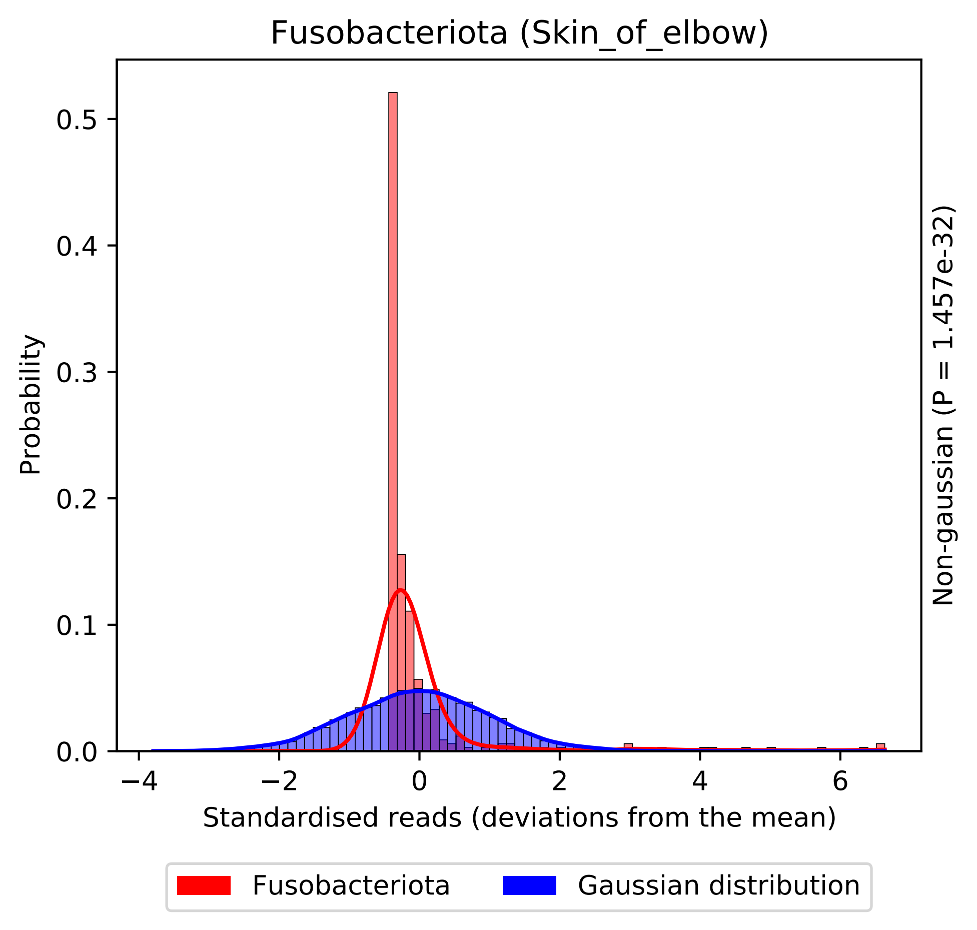
Fusobacteriota (Skin_of_elbow)
Probability
0.0
0.1
0.2
0.3
0.4
0.5
−4
−2
0
2
4
6
Standardised reads (deviations from the mean)
Non-gaussian (P = 1.457e-32)
Fusobacteriota
Gaussian distribution

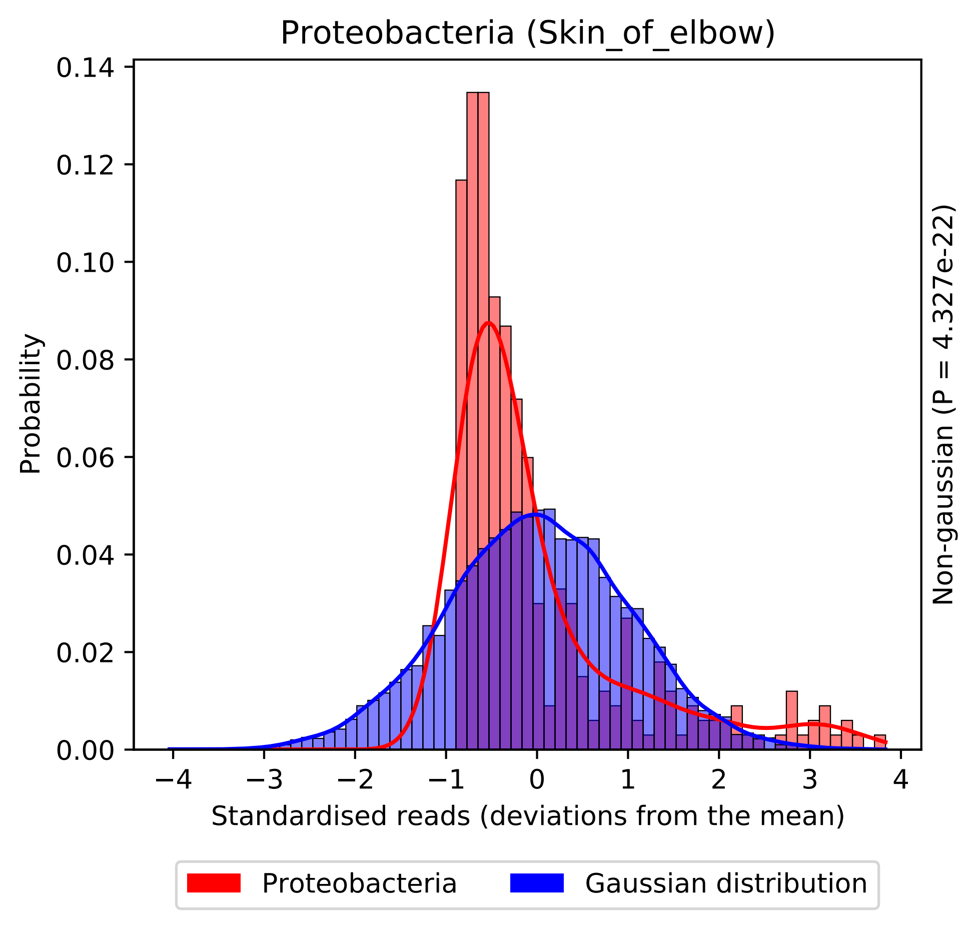

Proteobacteria (Skin_of_elbow)
Probability
0.00
0.02
0.04
0.06
0.08
0.10
0.12
0.14
−4
−3
−2
−1
0
1
2
3
4
Standardised reads (deviations from the mean)
Non-gaussian (P = 4.327e-22)
Proteobacteria
Gaussian distribution

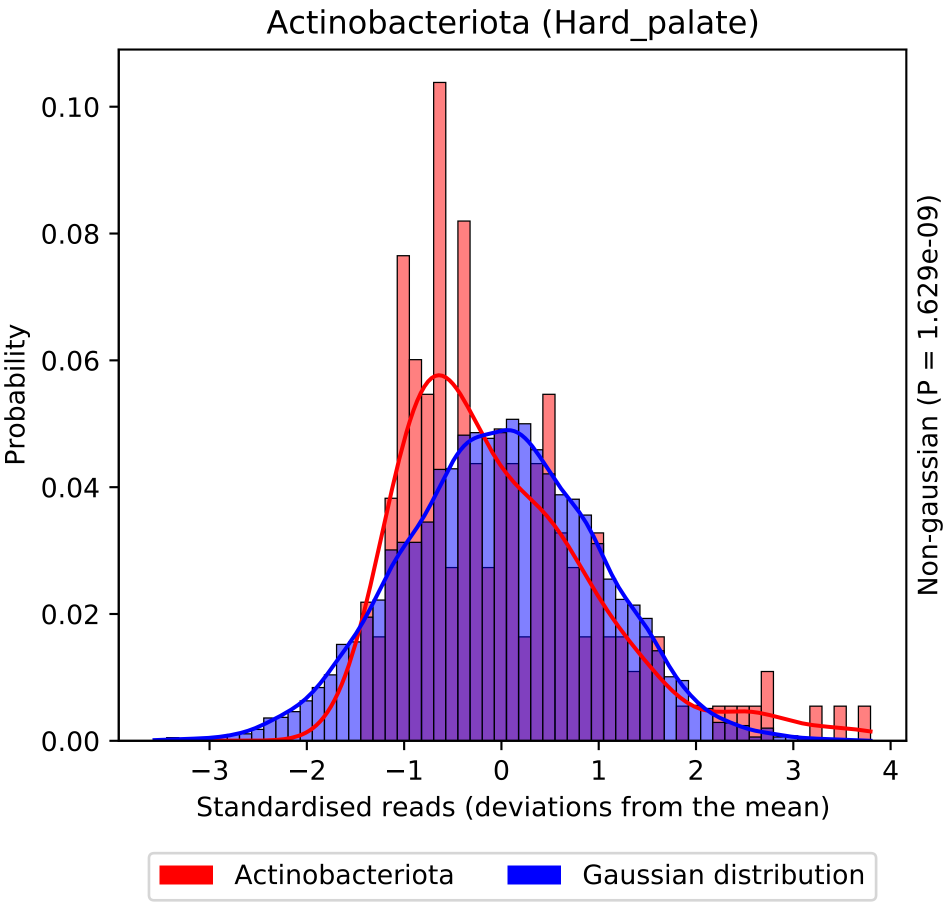
Actinobacteriota (Hard_palate)
Probability
0.00
0.02
0.04
0.06
0.08
0.10
−3
−2
−1
0
1
2
3
4
Standardised reads (deviations from the mean)
Non-gaussian (P = 1.629e-09)
Actinobacteriota
Gaussian distribution

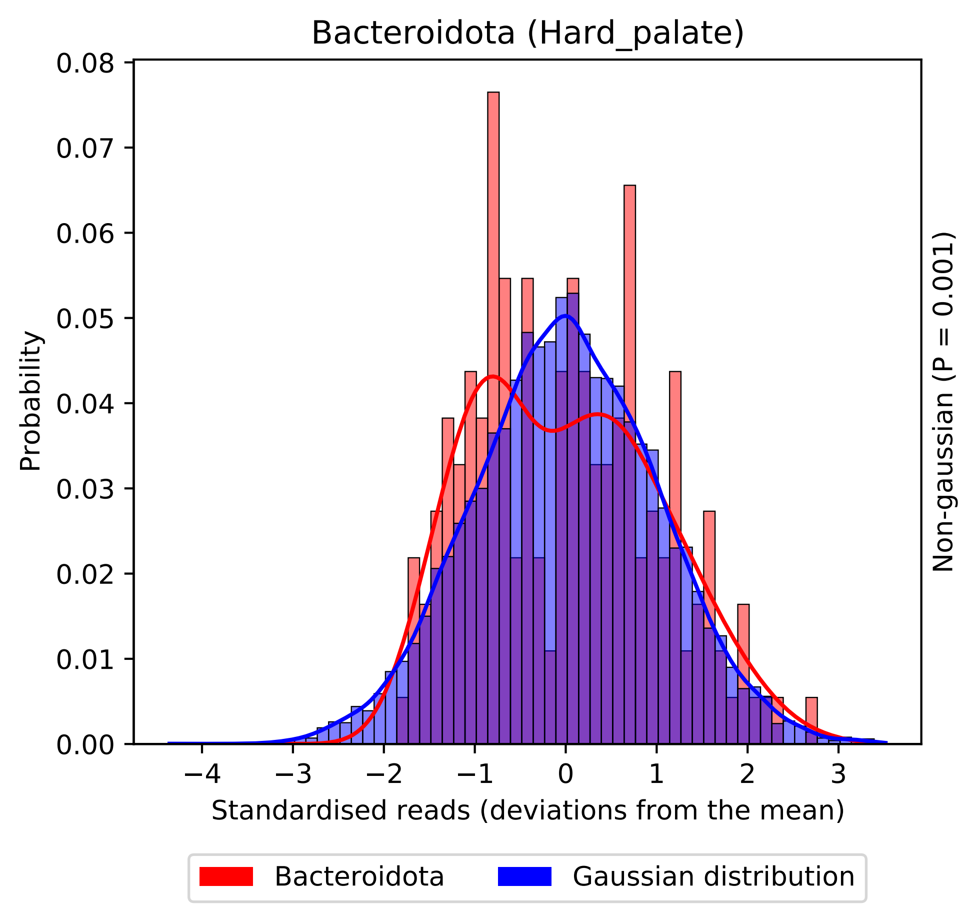

Bacteroidota (Hard_palate)
Probability
0.00
0.01
0.02
0.03
0.04
0.05
0.06
0.07
0.08
−4
−3
−2
−1
0
1
2
3
Standardised reads (deviations from the mean)
Non-gausian (P = 0.001)
Bacteroidota
Gaussian distribution

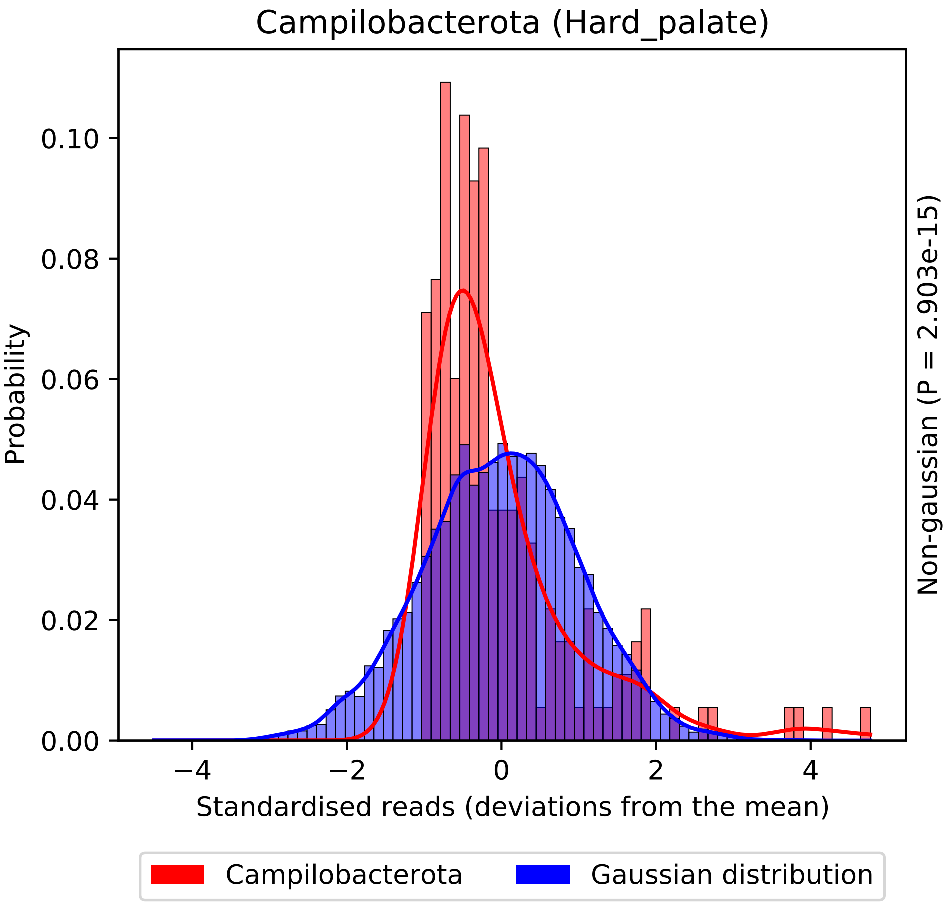
Campilobacterota (Hard_palate)
Probability
0.00
0.02
0.04
0.06
0.08
0.10
−4
−2
0
2
4
Standardised reads (deviations from the mean)
Non-gaussian (P = 2.903e-15)
Campilobacterota
Gaussian distribution

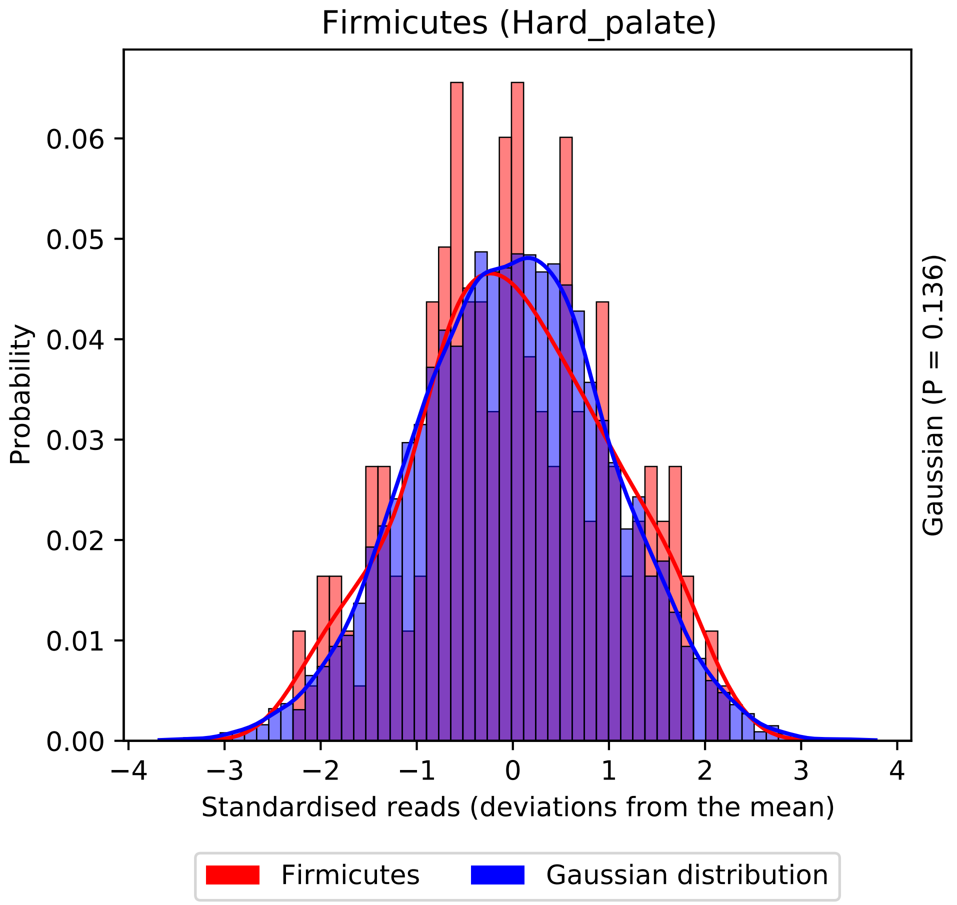
Firmicutes (Hard_palate)
Probability
0.00
0.01
0.02
0.03
0.04
0.05
0.06
−4
−3
−2
−1
0
1
2
3
4
Standardised reads (deviations from the mean)
Gaussian (P = 0.136)
Firmicutes
Gaussian distribution

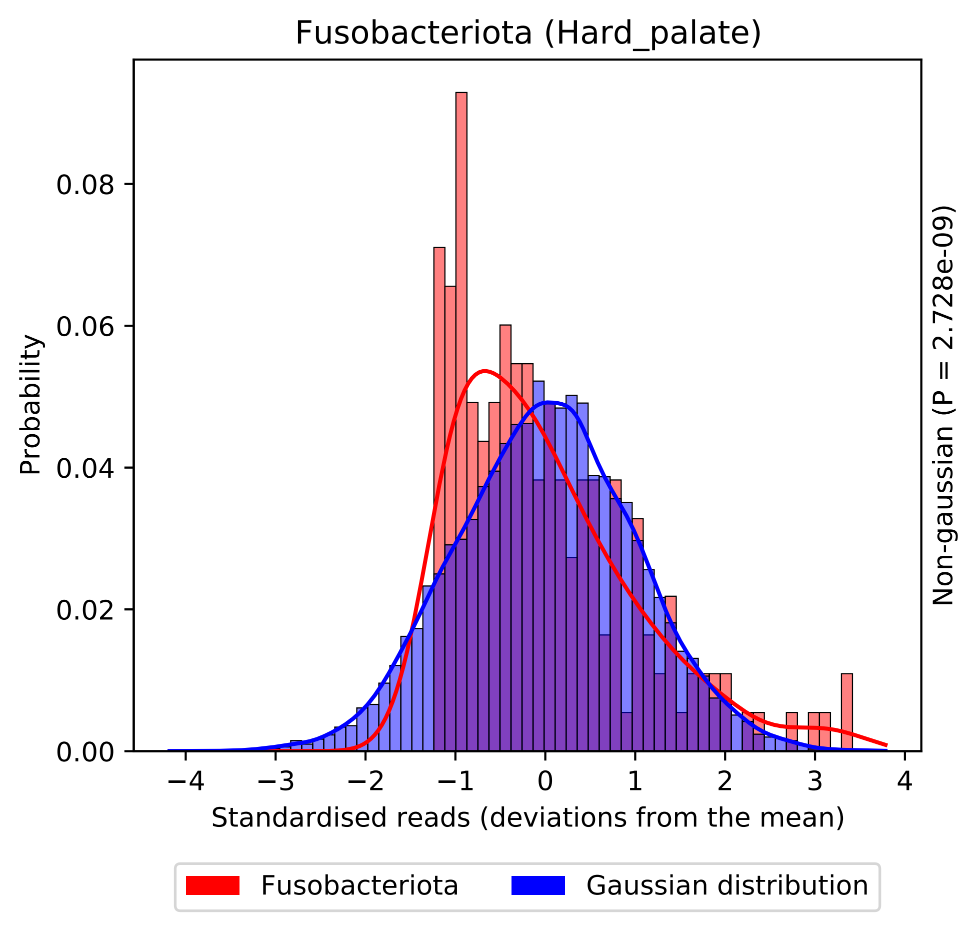
Fusobacteriota (Hard_palate)
Probability
0.00
0.02
0.04
0.06
0.08
−4
−3
−2
−1
0
1
2
3
4
Standardised reads (deviations from the mean)
Non-gaussian (P = 2.728e-09)
Fusobacteriota
Gaussian distribution

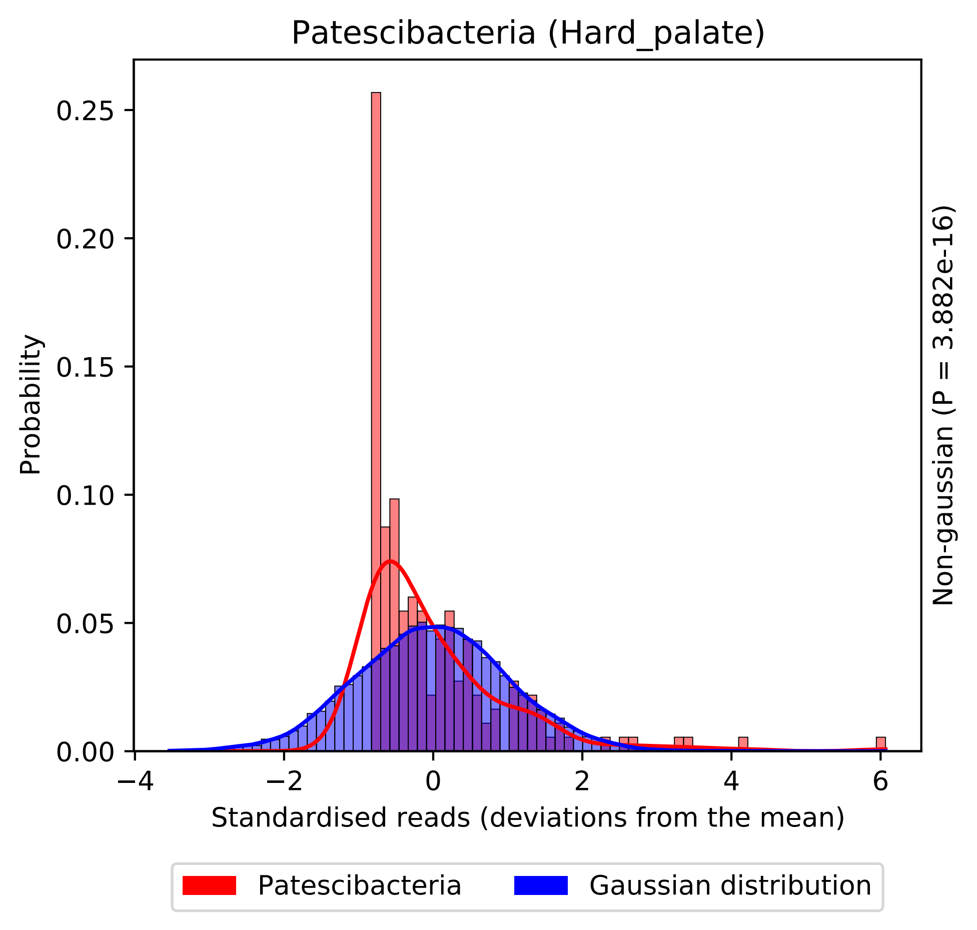
Patescibacteria (Hard_palate)
Probability
0.00
0.05
0.10
0.15
0.20
0.25
−4
−2
0
2
4
6
Standardised reads (deviations from the mean)
Non-gaussian (P = 3.882e-16)
Patescibacteria
Gaussian distribution

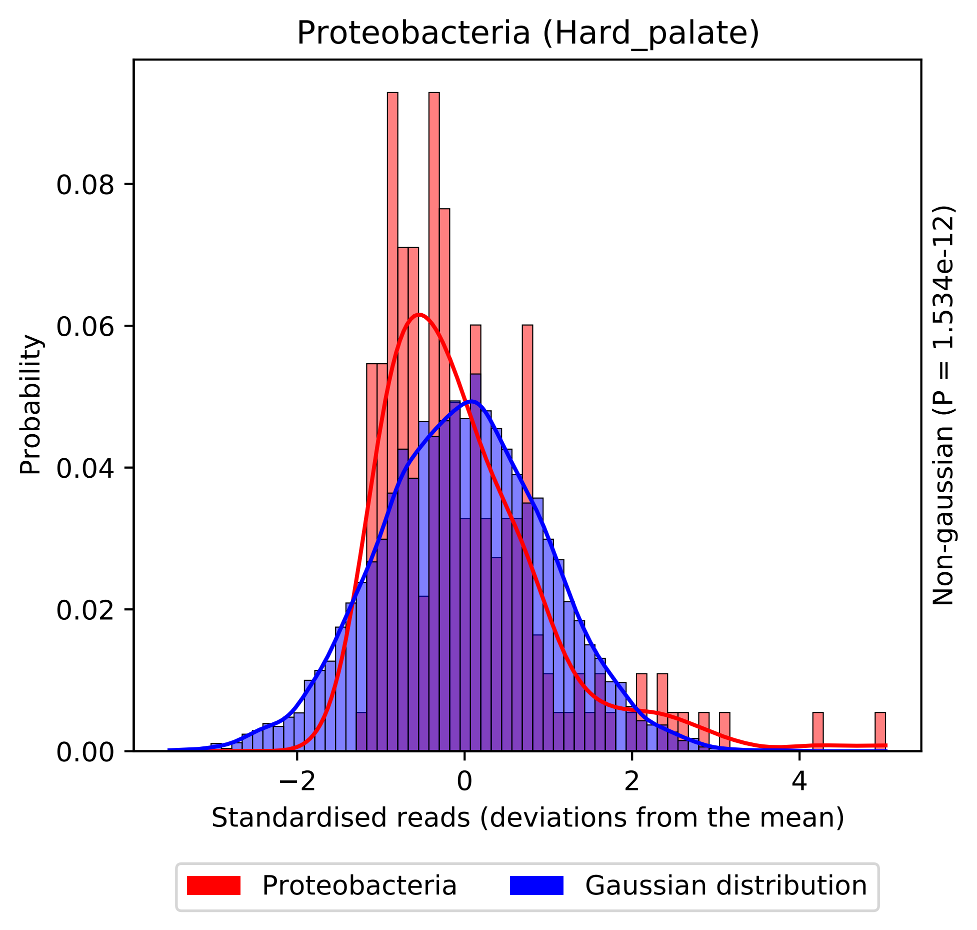

Proteobacteria (Hard_palate)
Probability
0.00
0.02
0.04
0.06
0.08
−2
0
2
4
Standardised reads (deviations from the mean)
Non-gaussian (P = 1.534e-12)
Proteobacteria
Gaussian distribution

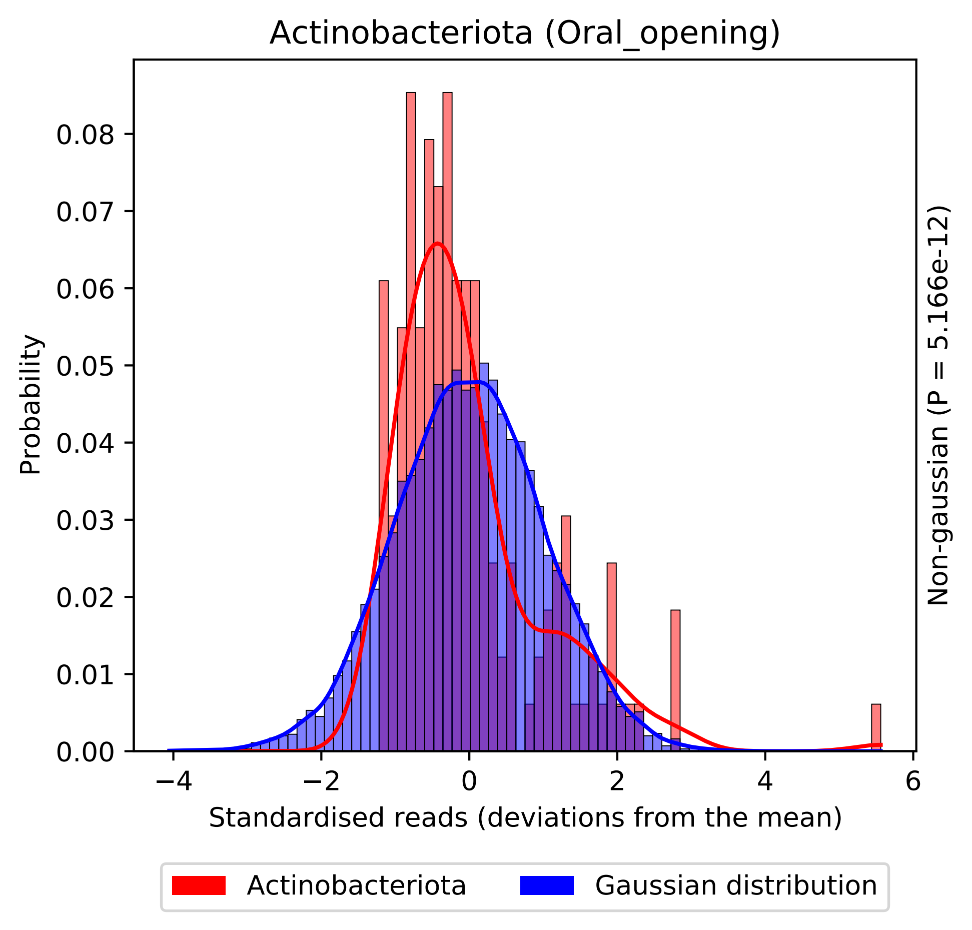
Actinobacteriota (Oral_opening)
Probability
0.00
0.01
0.02
0.03
0.04
0.05
0.06
0.07
0.08
−4
−2
0
2
4
6
Standardised reads (deviations from the mean)
Non-gaussian (P = 5.166e-12)
Actinobacteriota
Gaussian distribution

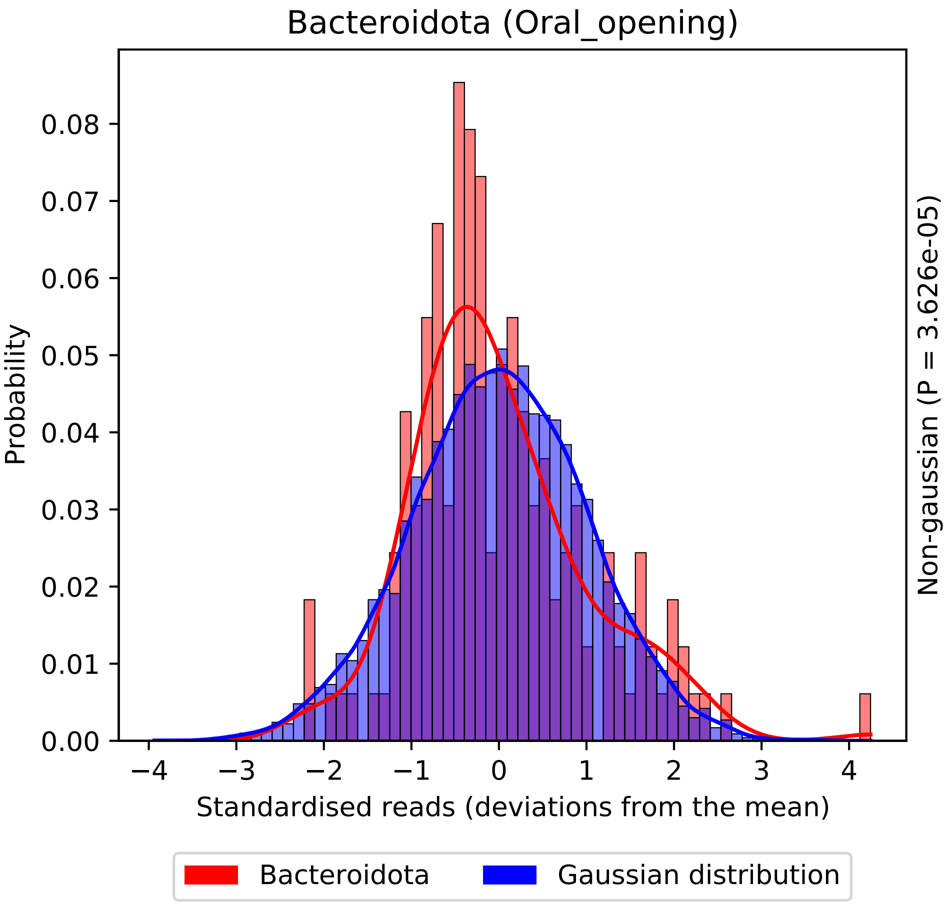
Bacteroidota (Oral_opening)
Probability
0.00
0.01
0.02
0.03
0.04
0.05
0.06
0.07
0.08
−4
−3
−2
−1
0
1
2
3
4
Standardised reads (deviations from the mean)
Non-gaussian (P = 3.626e-05)
Bacteroidota
Gaussian distribution

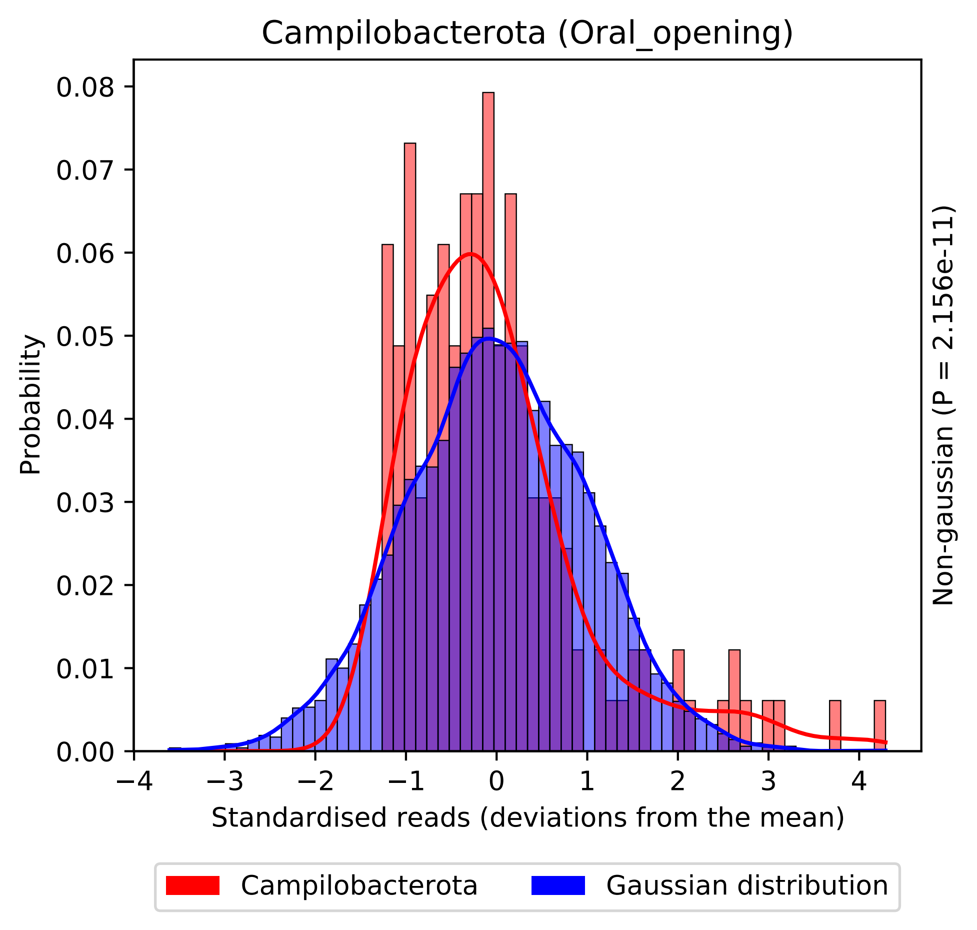

Campilobacterota (Oral_opening)
Probability
Standardised reads (deviations from the mean)
Non-gaussian (P = 2.156e-11)
0.00
0.01
0.02
0.03
0.04
0.05
0.06
0.07
0.08
−4
−3
−2
−1
0
1
2
3
4
Campilobacterota
Gaussian distribution

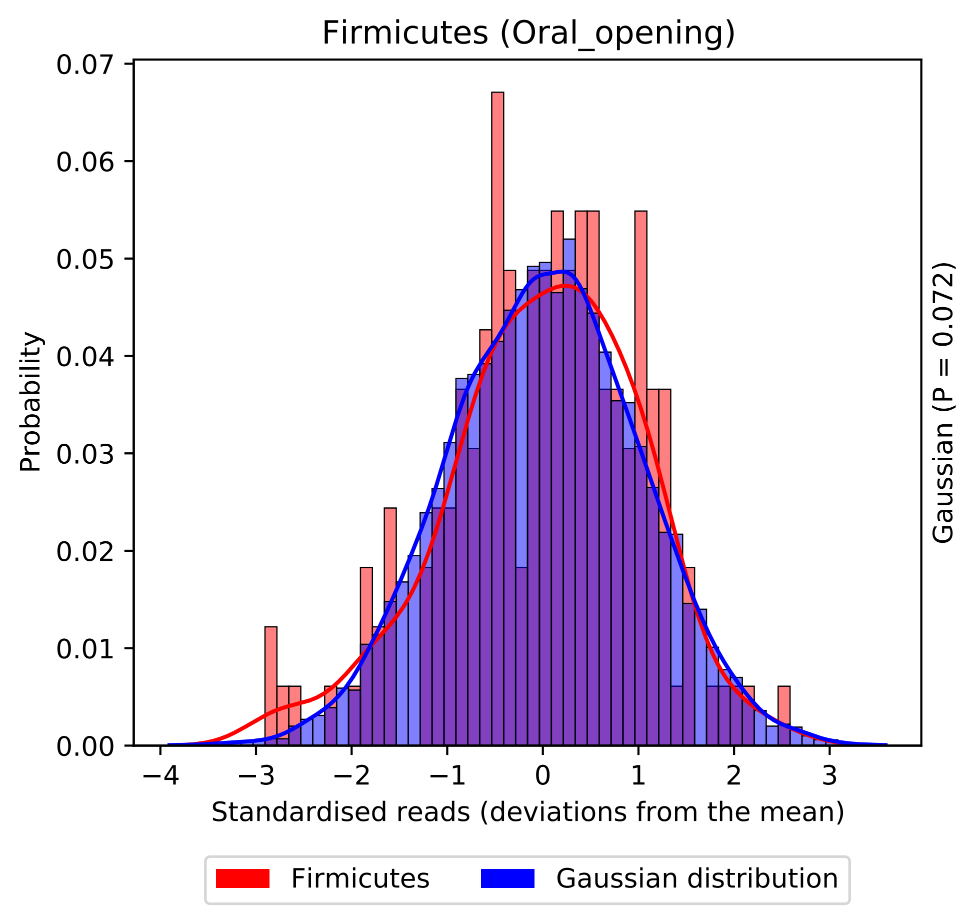
Firmicutes (Oral_opening)
0.07
0.06
0.05
0.04
0.03
0.02
0.01
0.00
Probability
−4
−3
−2
−1
0
1
2
3
Standardised reads (deviations from the mean)
Gaussian (P = 0.072)
Firmicutes
Gaussian distribution

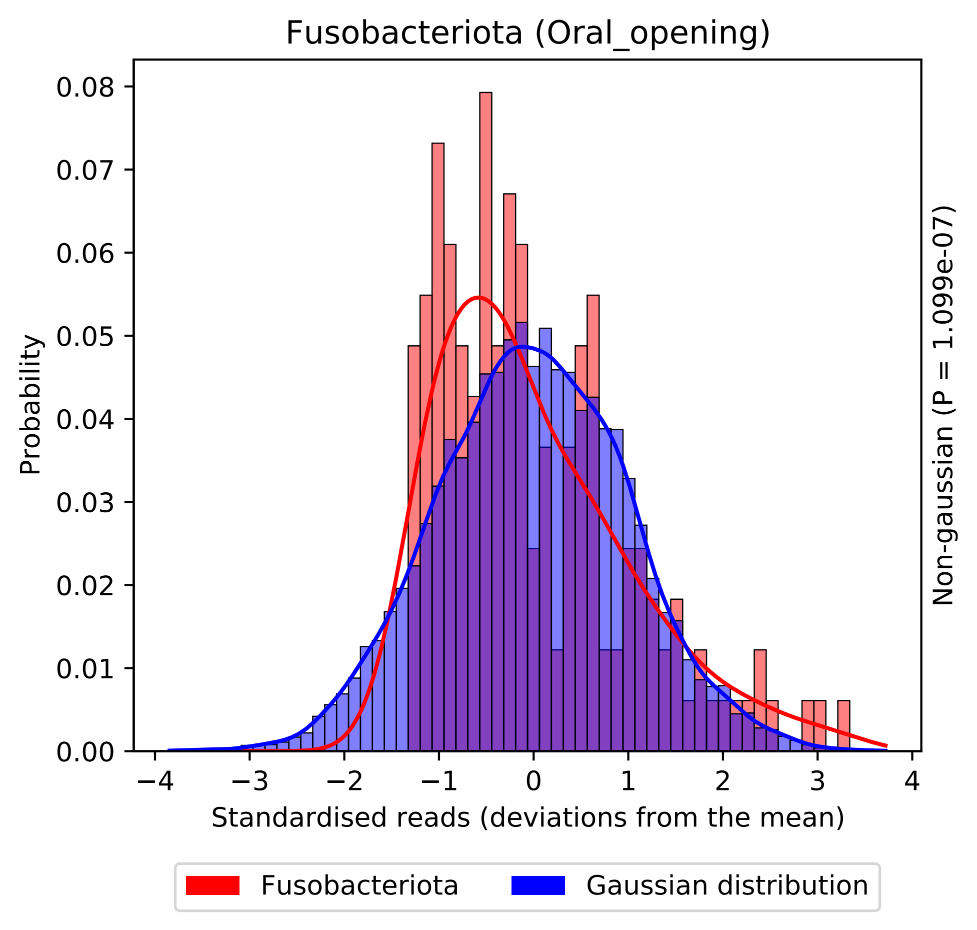
Fusobacteriota (Oral_opening)
Probability
0.00
0.01
0.02
0.03
0.04
0.05
0.06
0.07
0.08
−4
−3
−2
−1
0
1
2
3
4
Standardised reads (deviations from the mean)
Non-gaussian (P = 1.099e-07)
Fusobacteriota
Gaussian distribution

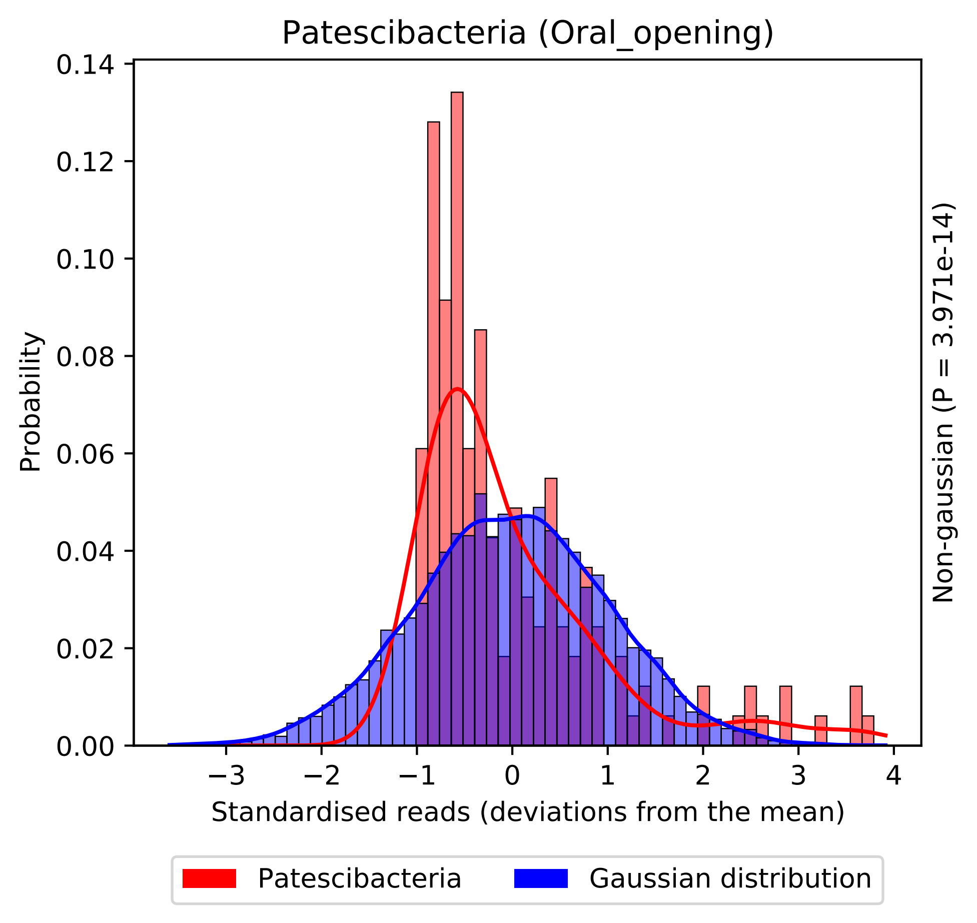

Patescibacteria (Oral_opening)
0.14
0.12
0.10
0.08
0.06
0.04
0.02
0.00
Probability
−3
−2
−1
0
1
2
3
4
Standardised reads (deviations from the mean)
Non-gaussian (P = 3.971e-14)
Patescibacteria
Gaussian distribution

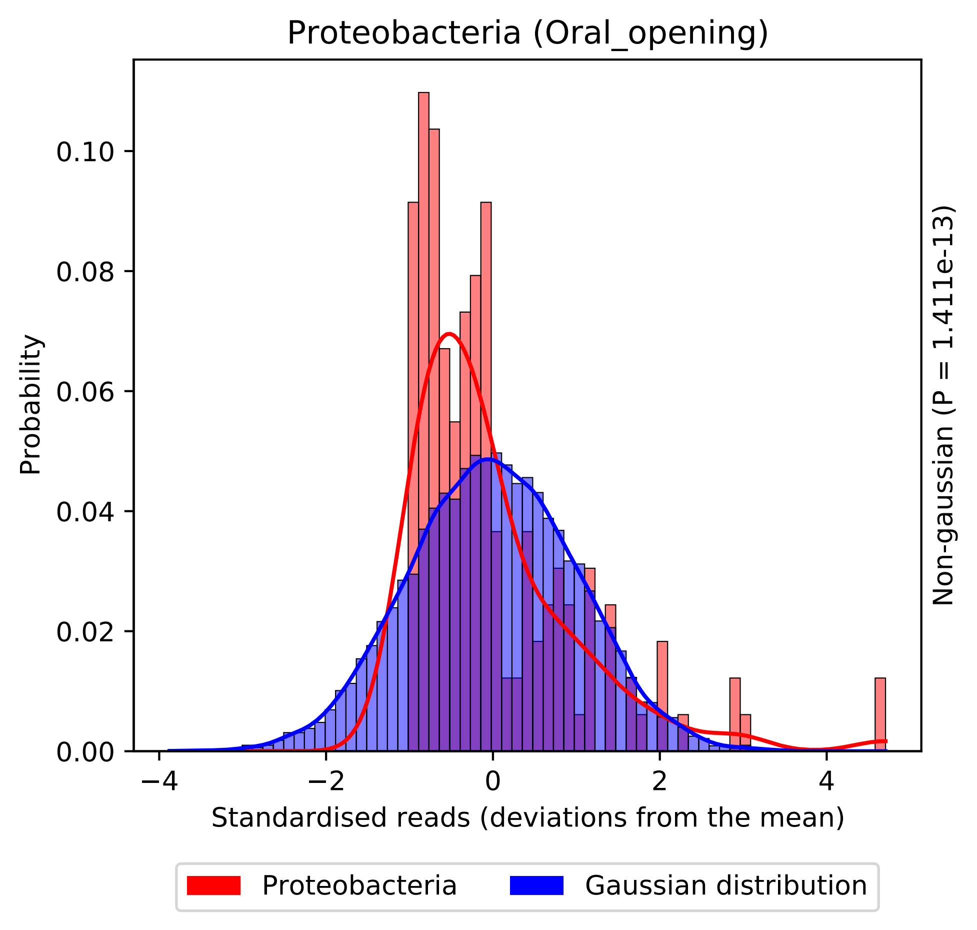
Proteobacteria (Oral_opening)
Probability
0.10
0.08
0.06
0.04
0.02
0.00
−4
−2
0
2
4
Standardised reads (deviations from the mean)
Non-gaussian (P = 1.411e-13)
Proteobacteria
Gaussian distribution

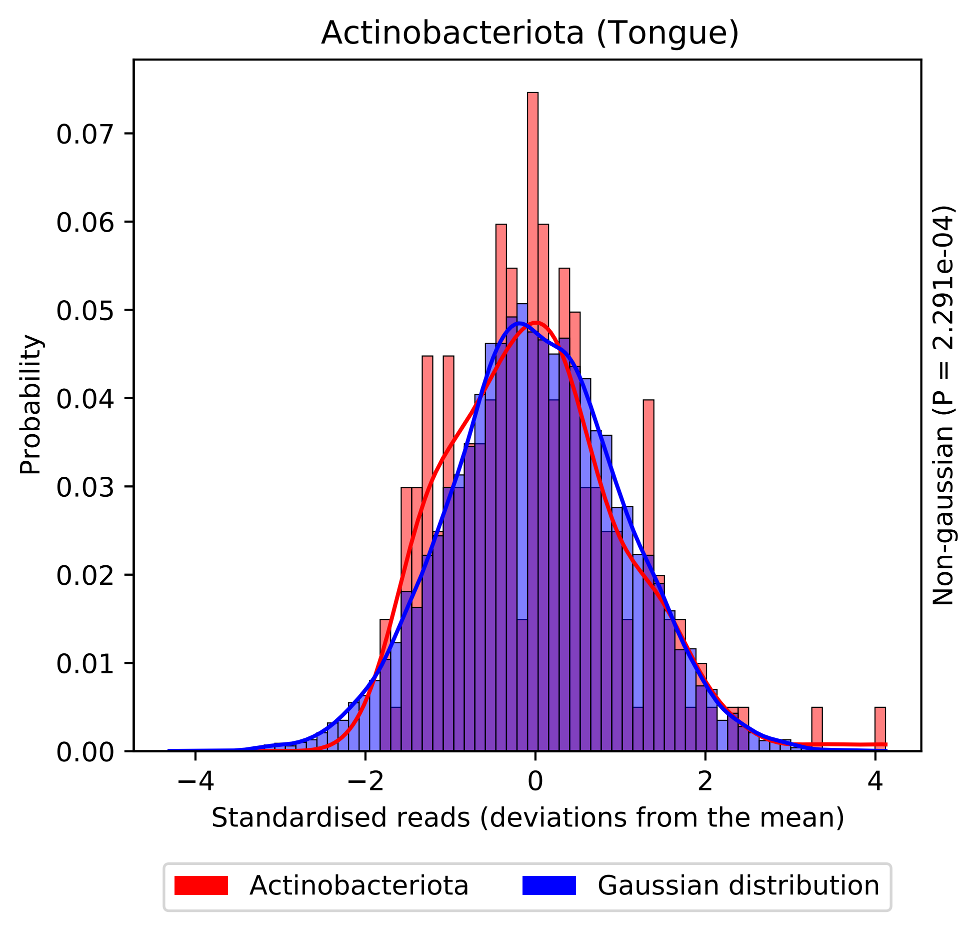

Actinobacteriota (Tongue)
Probability
0.00
0.01
0.02
0.03
0.04
0.05
0.06
0.07
−4
−2
0
2
4
Standardised reads (deviations from the mean)
Non-gaussian (P = 2.291e-04)
Actinobacteriota
Gaussian distribution

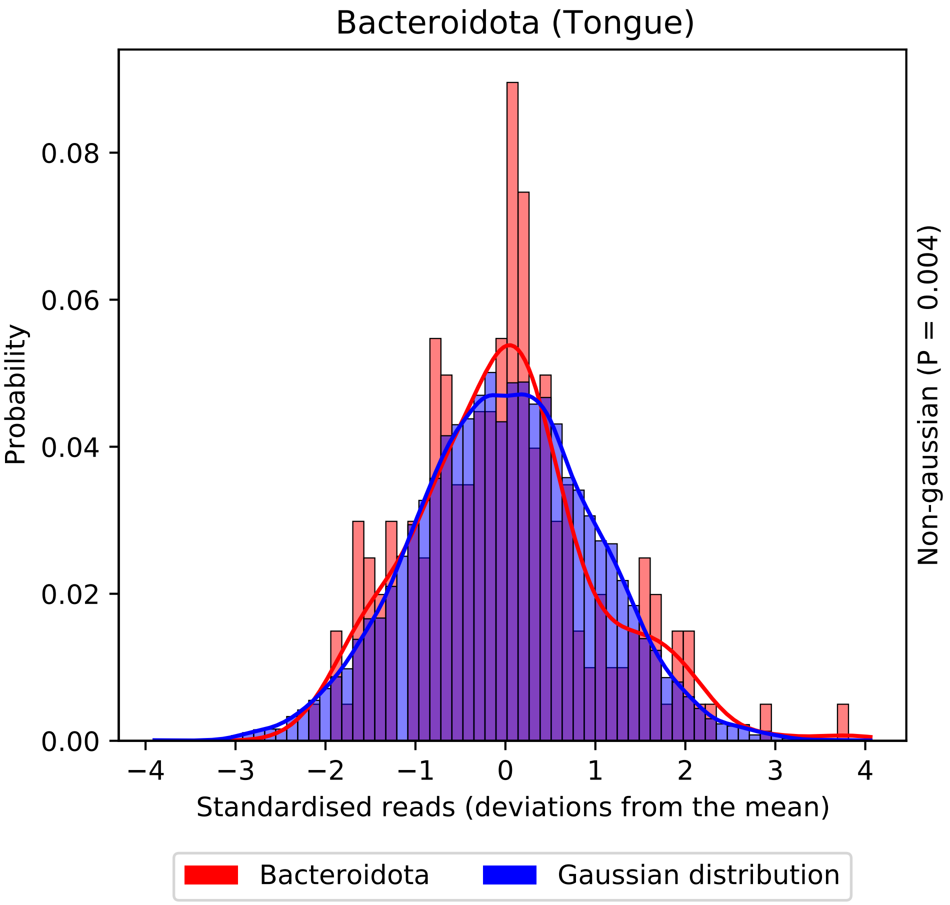
Bacteroidota (Tongue)
Probability
0.00
0.02
0.04
0.06
0.08
−4
−3
−2
−1
0
1
2
3
4
Standardised reads (deviations from the mean)
Non-gaussian (P = 0.004)
Bacteroidota
Gaussian distribution

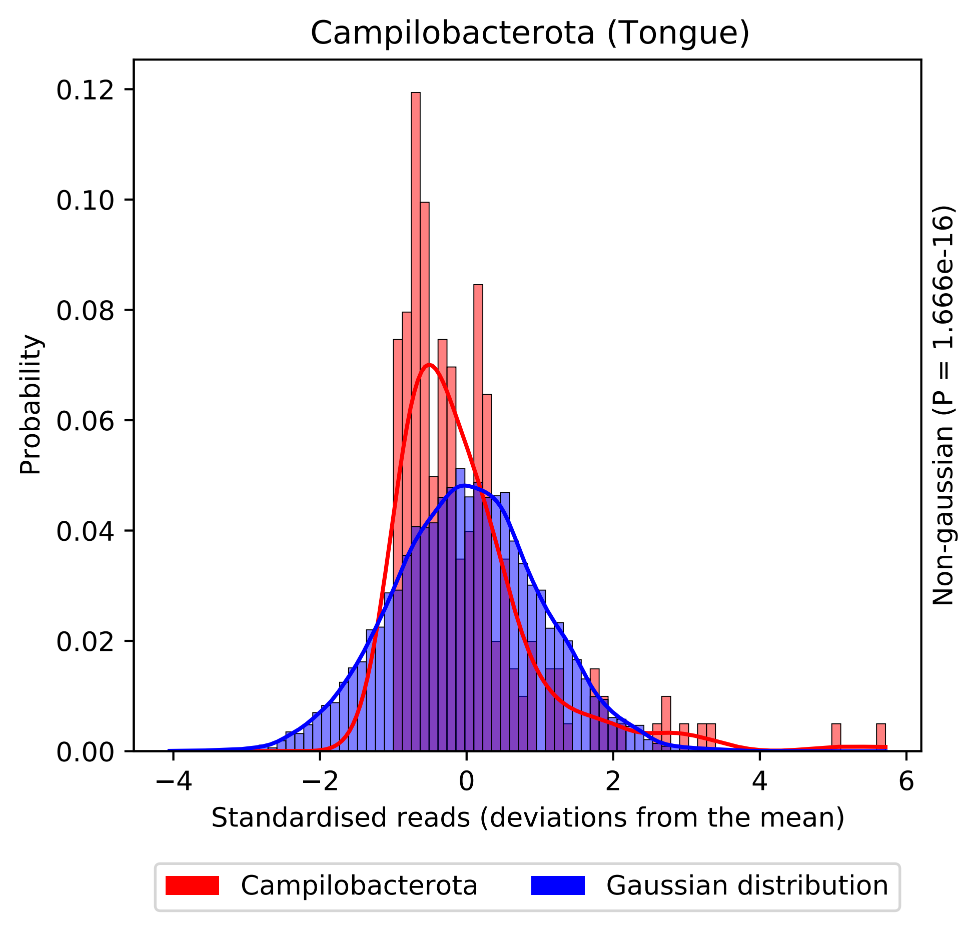

Campilobacterota (Tongue)
Probability
0.12
0.10
0.08
0.06
0.04
0.02
0.00
−4
−2
0
2
4
6
Standardised reads (deviations from the mean)
Non-gaussian (P = 1.666e-16)
Campilobacterota
Gaussian distribution

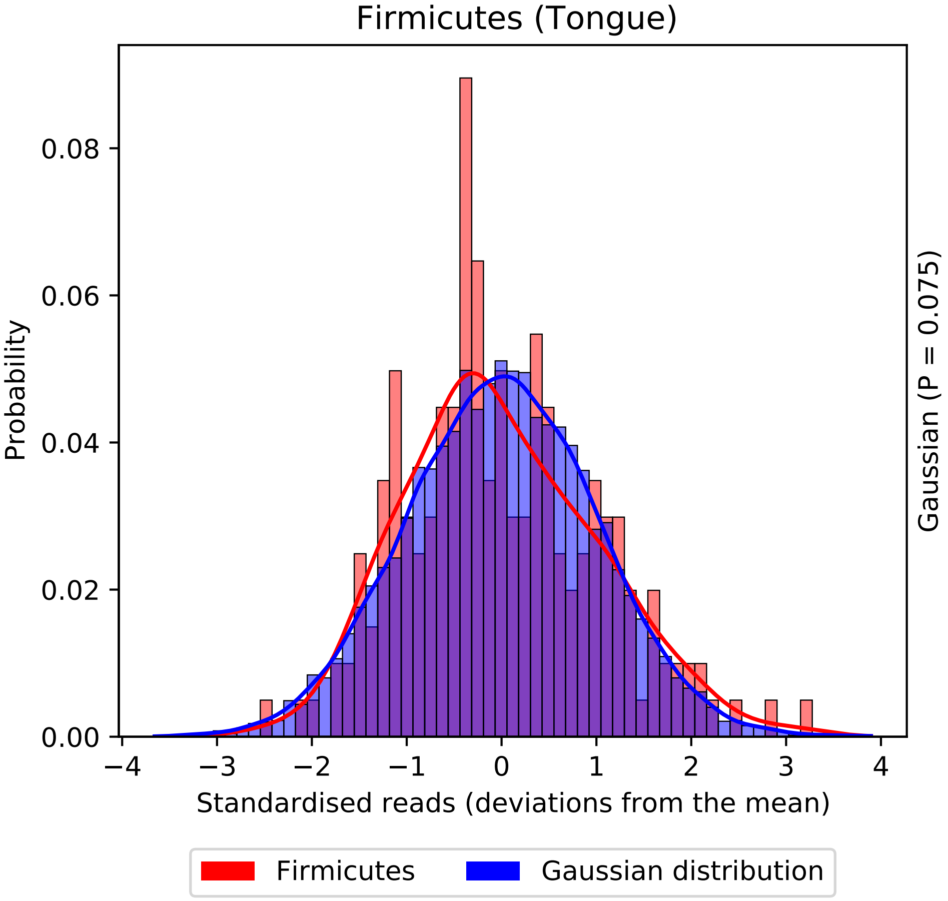
Firmicutes (Tongue)
Probability
0.00
0.02
0.04
0.06
0.08
−4
−3
−2
−1
0
1
2
3
4
Standardised reads (deviations from the mean)
Gaussian (P = 0.075)
Firmicutes
Gaussian distribution

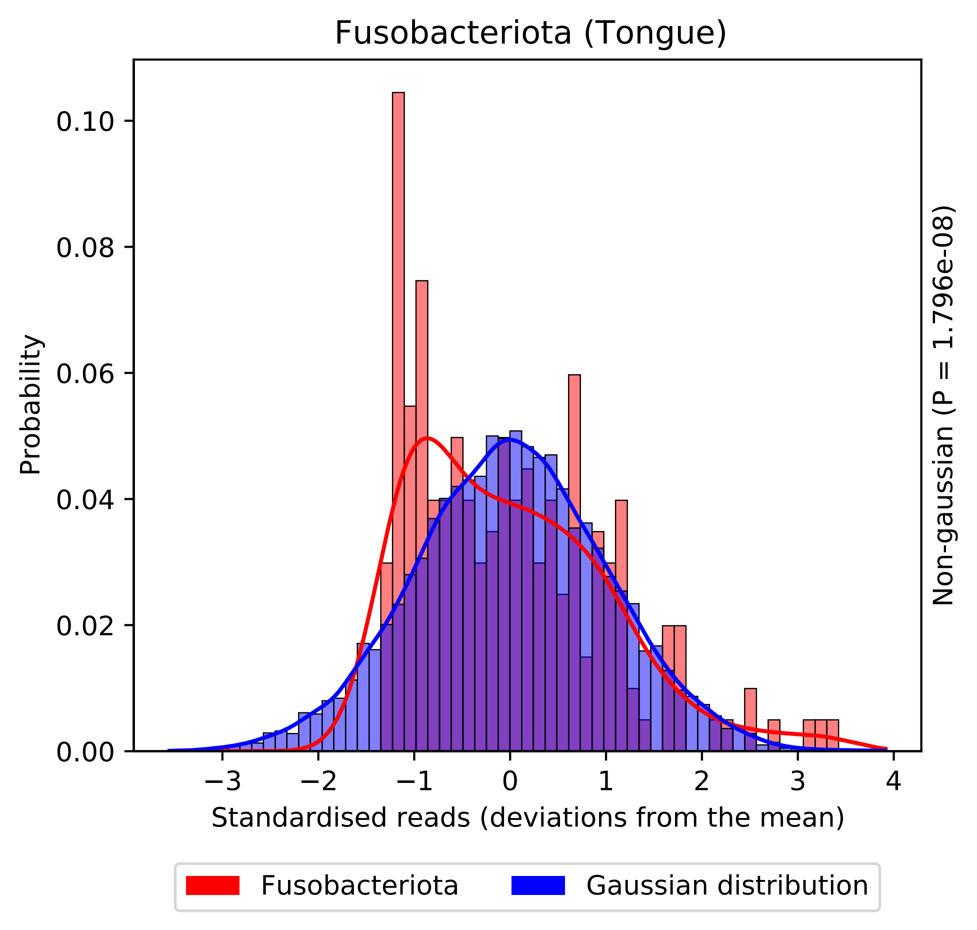
Fusobacteriota (Tongue)
Probability
0.00
0.02
0.04
0.06
0.08
0.10
−3
−2
−1
0
1
2
3
4
Standardised reads (deviations from the mean)
Non-gaussian (P = 1.796e-08)
Fusobacteriota
Gaussian distribution

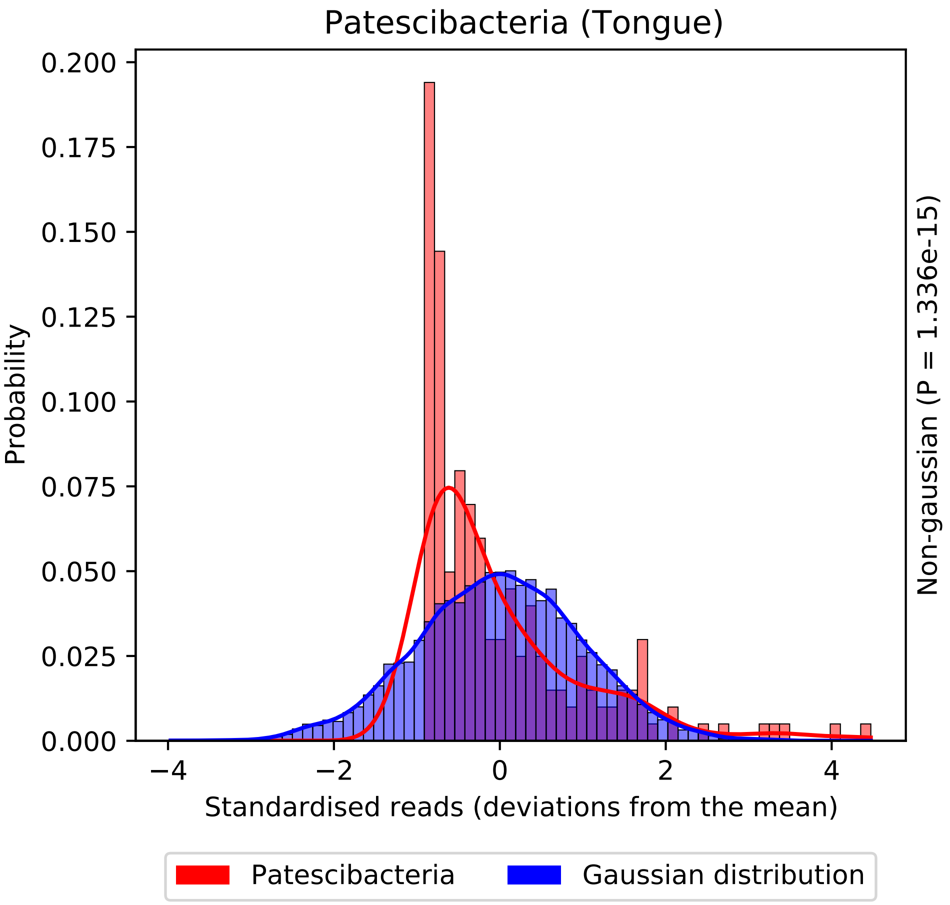
Patescibacteria (Tongue)
Probability
0.200
0.175
0.150
0.125
0.100
0.075
0.050
0.025
0.000
−4
−2
0
2
4
Standardised reads (deviations from the mean)
Non-gaussian (P = 1.336e-15)
Patescibacteria
Gaussian distribution

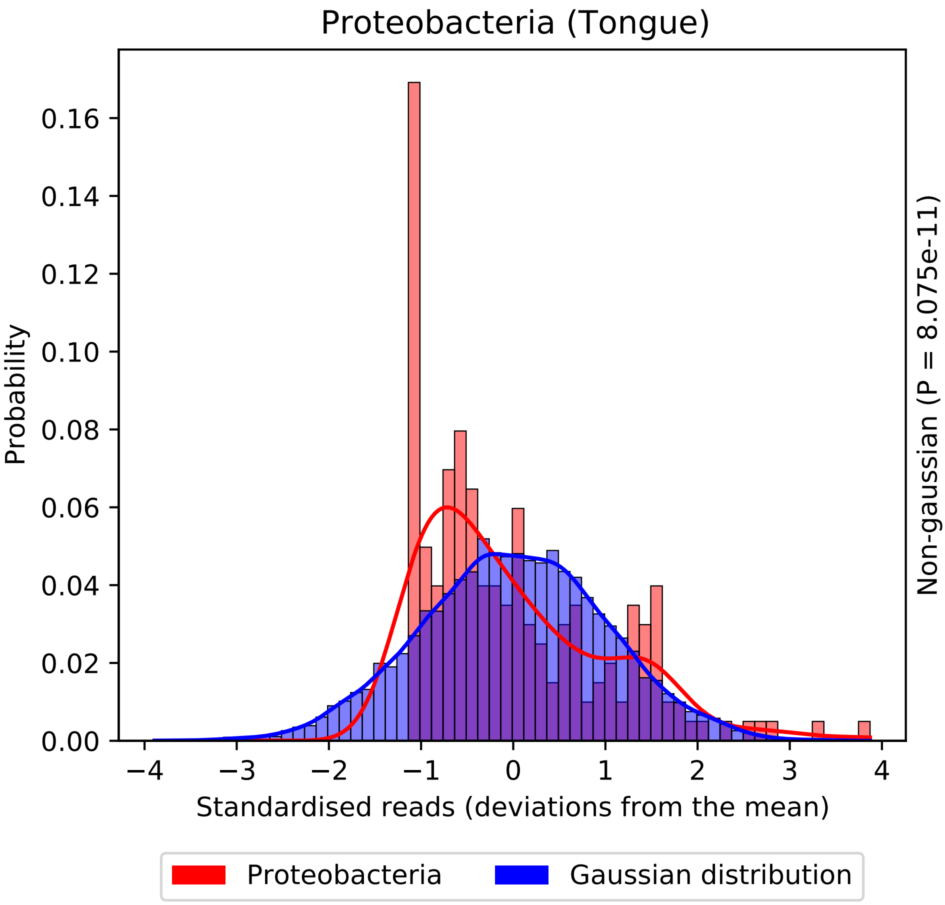

Proteobacteria (Tongue)
Probability
0.00
0.02
0.04
0.06
0.08
0.10
0.12
0.14
0.16
−4
−3
−2
−1
0
1
2
3
4
Standardised reads (deviations from the mean)
Non-gaussian (P = 8.075e-11)
Proteobacteria
Gaussian distribution

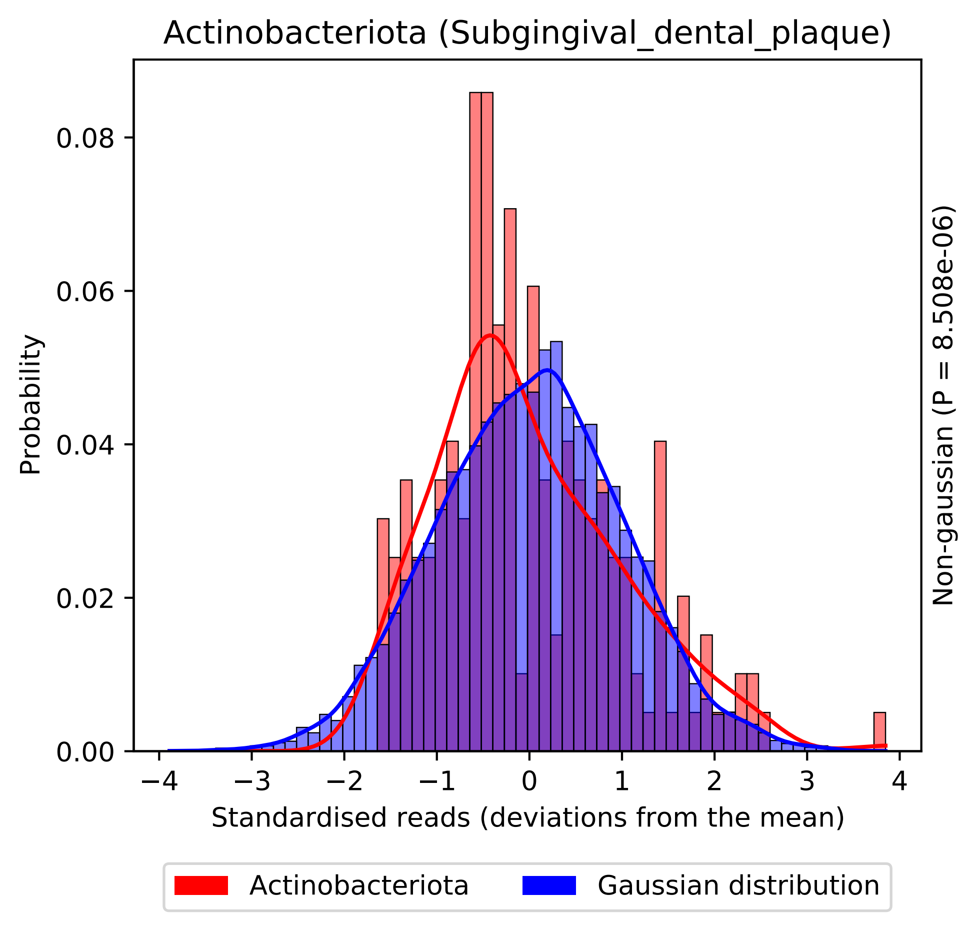
Actinobacteriota (Subgingival_dental_plaque)
Probability
0.00
0.02
0.04
0.06
0.08
−4
−3
−2
−1
0
1
2
3
4
Standardised reads (deviations from the mean)
Non-gaussian (P = 8.508e-06)
Actinobacteriota
Gaussian distribution

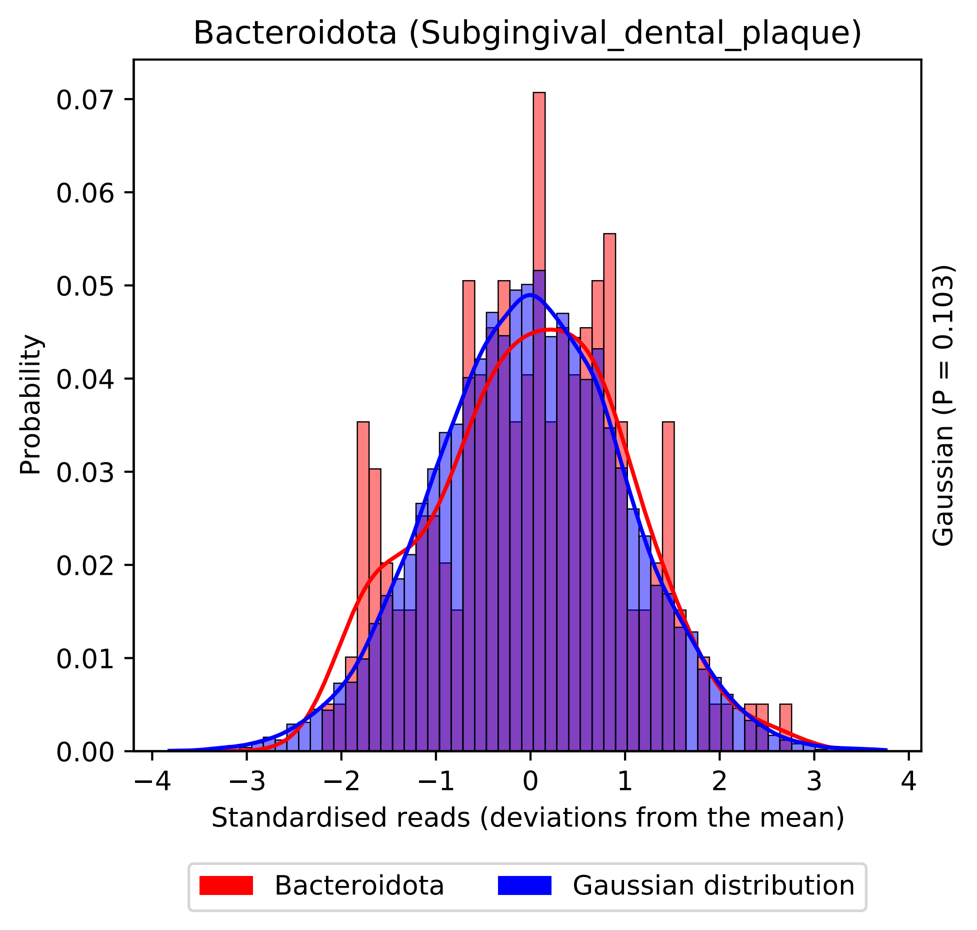
Bacteroidota (Subgingival_dental_plaque)
Probability
0.00
0.01
0.02
0.03
0.04
0.05
0.06
0.07
−4
−3
−2
−1
0
1
2
3
4
Standardised reads (deviations from the mean)
Gaussian (P = 0.103)
Bacteroidota
Gaussian distribution

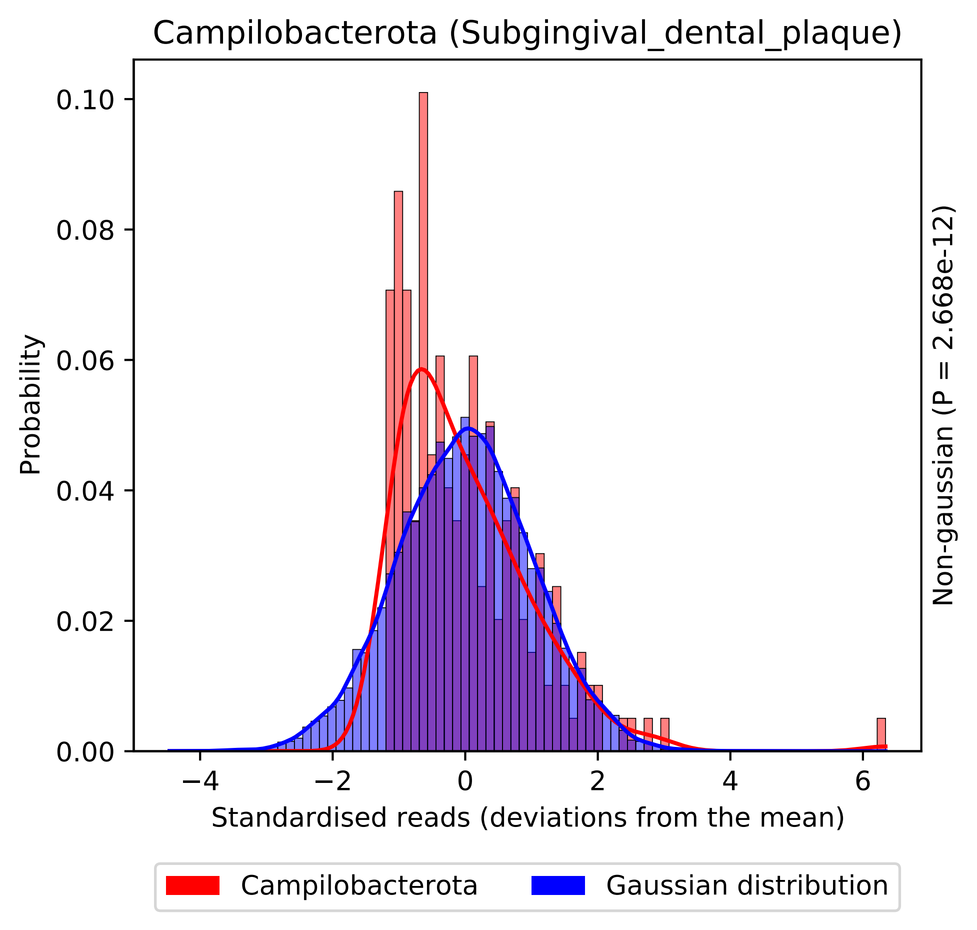

Campilobacterota (Subgingival_dental_plaque)
Probability
0.00
0.02
0.04
0.06
0.08
0.10
−4
−2
0
2
4
6
Standardised reads (deviations from the mean)
Non-gaussian (P = 2.668e-12)
Campilobacterota
Gaussian distribution

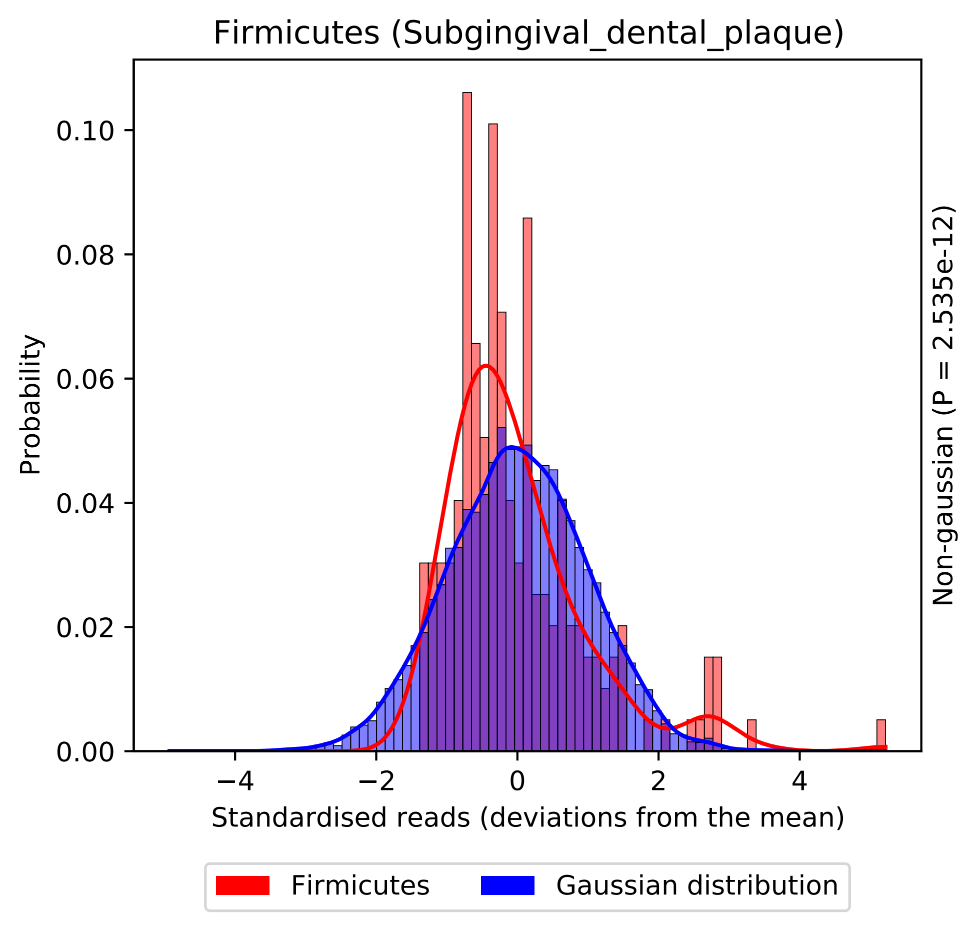
Firmicutes (Subgingival_dental_plaque)
Probability
0.00
0.02
0.04
0.06
0.08
0.10
−4
−2
0
2
4
Standardised reads (deviations from the mean)
Non-gaussian (P = 2.535e-12)
Firmicutes
Gaussian distribution

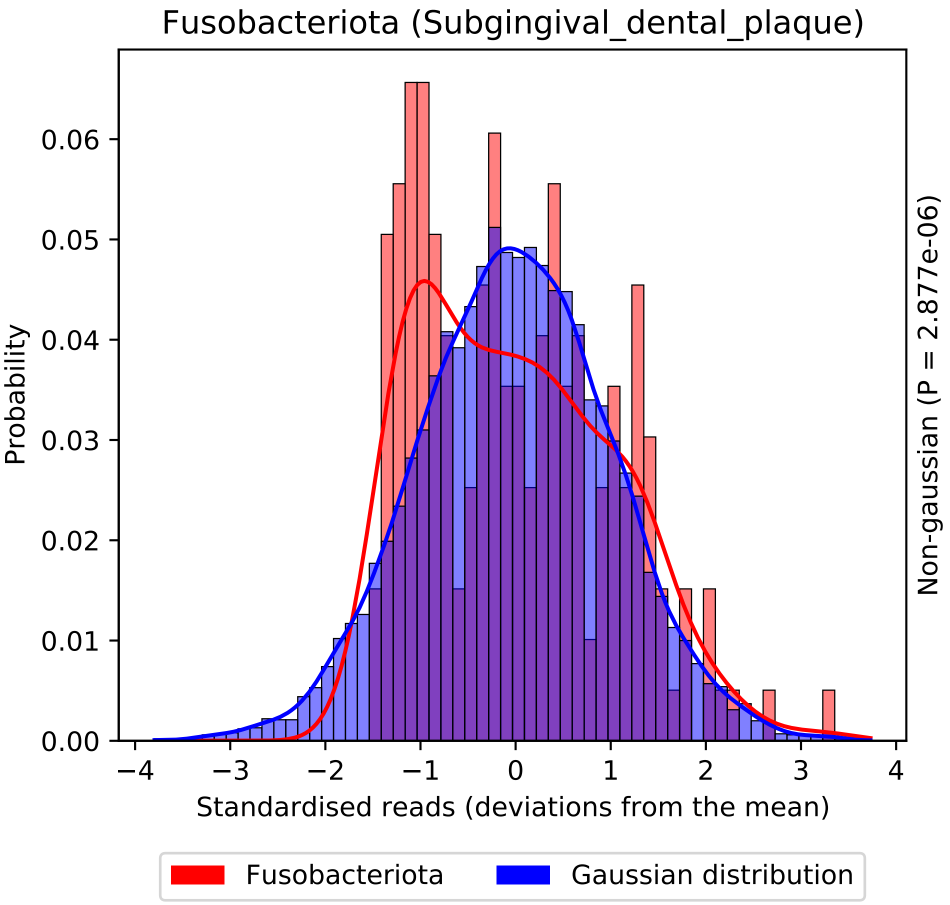
Fusobacteriota (Subgingival_dental_plaque)
Probability
0.00
0.01
0.02
0.03
0.04
0.05
0.06
−4
−3
−2
−1
0
1
2
3
4
Standardised reads (deviations from the mean)
Non-gaussian (P = 2.877e-06)
Fusobacteriota
Gaussian distribution

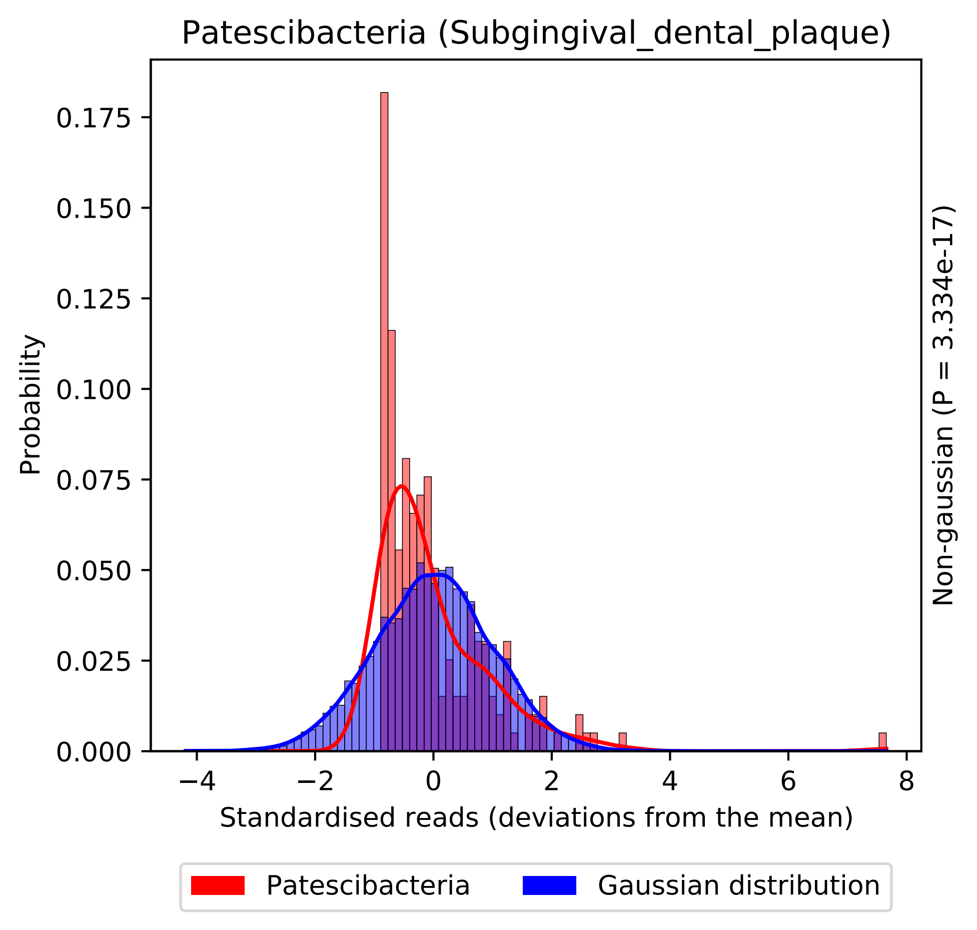

Patescibacteria (Subgingival_dental_plaque)
Probability
0.000
0.025
0.050
0.075
0.100
0.125
0.150
0.175
−4
−2
0
2
4
6
8
Standardised reads (deviations from the mean)
Non-gaussian (P = 3.334e-17)
Patescibacteria
Gaussian distribution

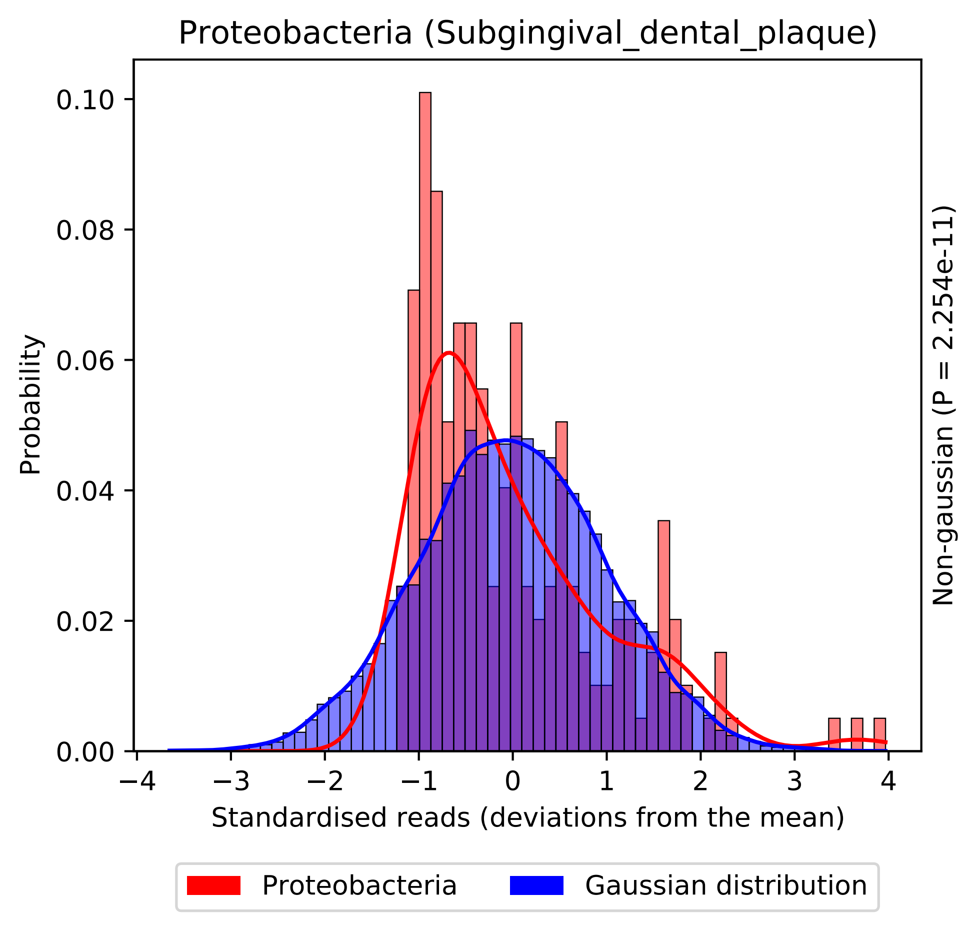

Proteobacteria (Subgingival_dental_plaque)
Probability
0.00
0.02
0.04
0.06
0.08
0.10
−4
−3
−2
−1
0
1
2
3
4
Standardised reads (deviations from the mean)
Non-gaussian (P = 2.254e-11)
Proteobacteria
Gaussian distribution

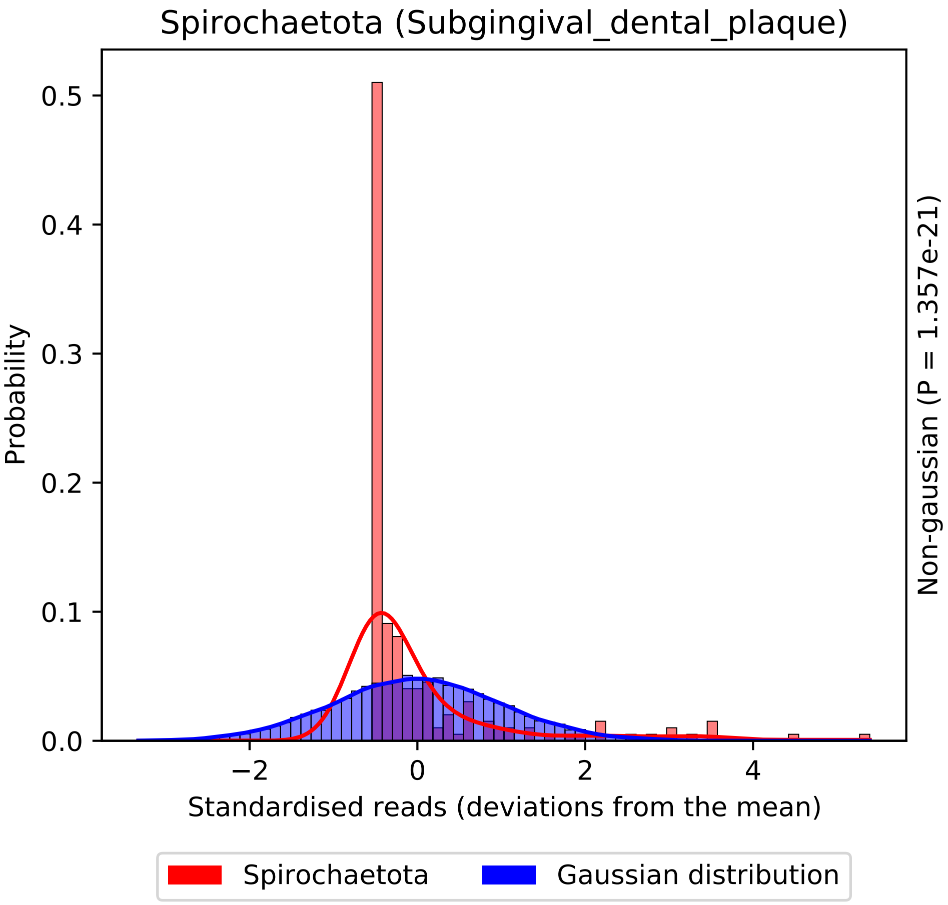
Spirochaetota (Subgingival_dental_plaque)
Probability
0.0
0.1
0.2
0.3
0.4
0.5
−2
0
2
4
Standardised reads (deviations from the mean)
Non-gaussian (P = 1.357e-21)
Spirochaetota
Gaussian distribution

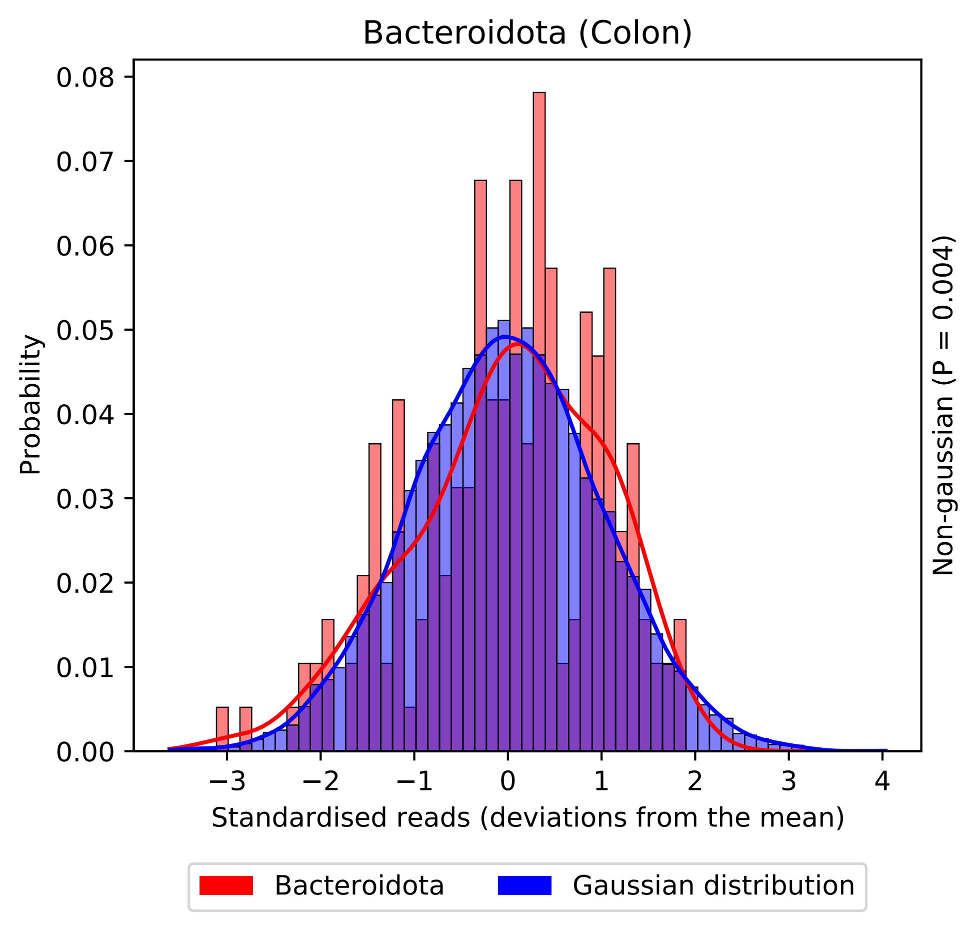
Bacteroidota (Colon)
Probability
0.00
0.01
0.02
0.03
0.04
0.05
0.06
0.07
0.08
−3
−2
−1
0
1
2
3
4
Standardised reads (deviations from the mean)
Non-gaussian (P = 0.004)
Bacteroidota
Gaussian distribution

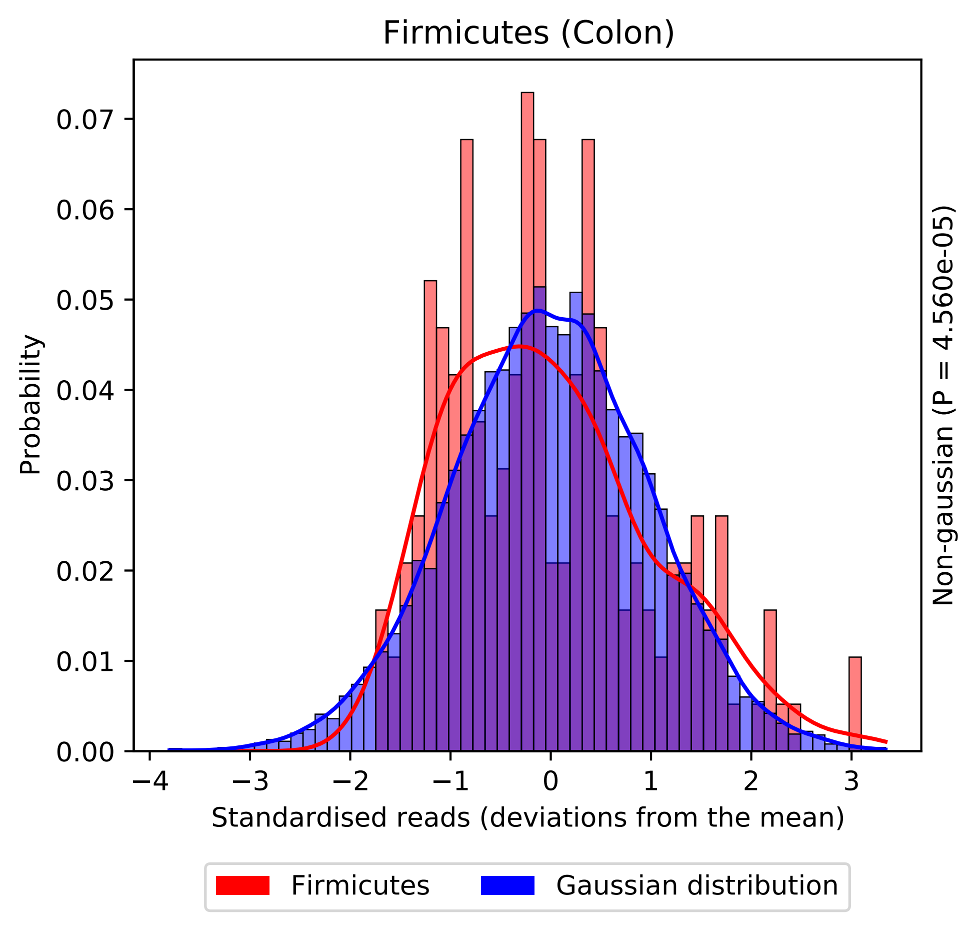

Firmicutes (Colon)
Probability
0.00
0.01
0.02
0.03
0.04
0.05
0.06
0.07
−4
−3
−2
−1
0
1
2
3
Standardised reads (deviations from the mean)
Non-gaussian (P = 4.560e-05)
Firmicutes
Gaussian distribution

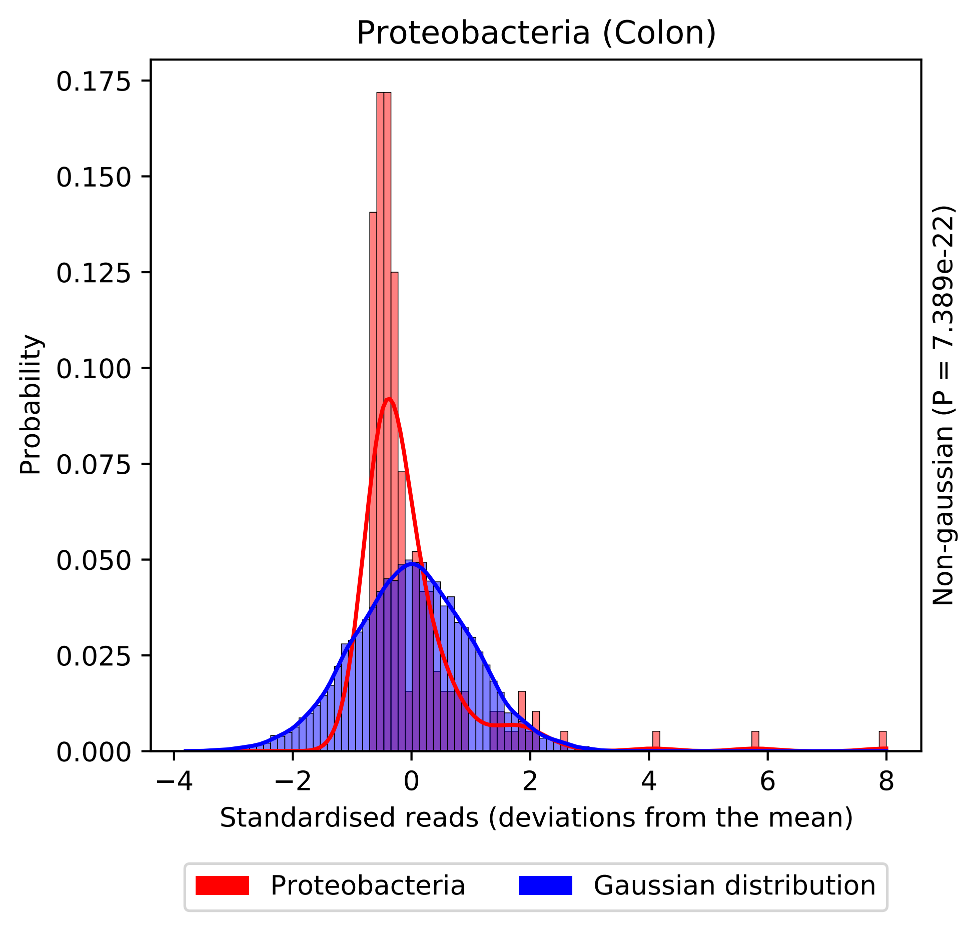
Proteobacteria (Colon)
Probability
0.175
0.150
0.125
0.100
0.075
0.050
0.025
0.000
−4
−2
0
2
4
6
8
Standardised reads (deviations from the mean)
Non-gaussian (P = 7.389e-22)
Proteobacteria
Gaussian distribution

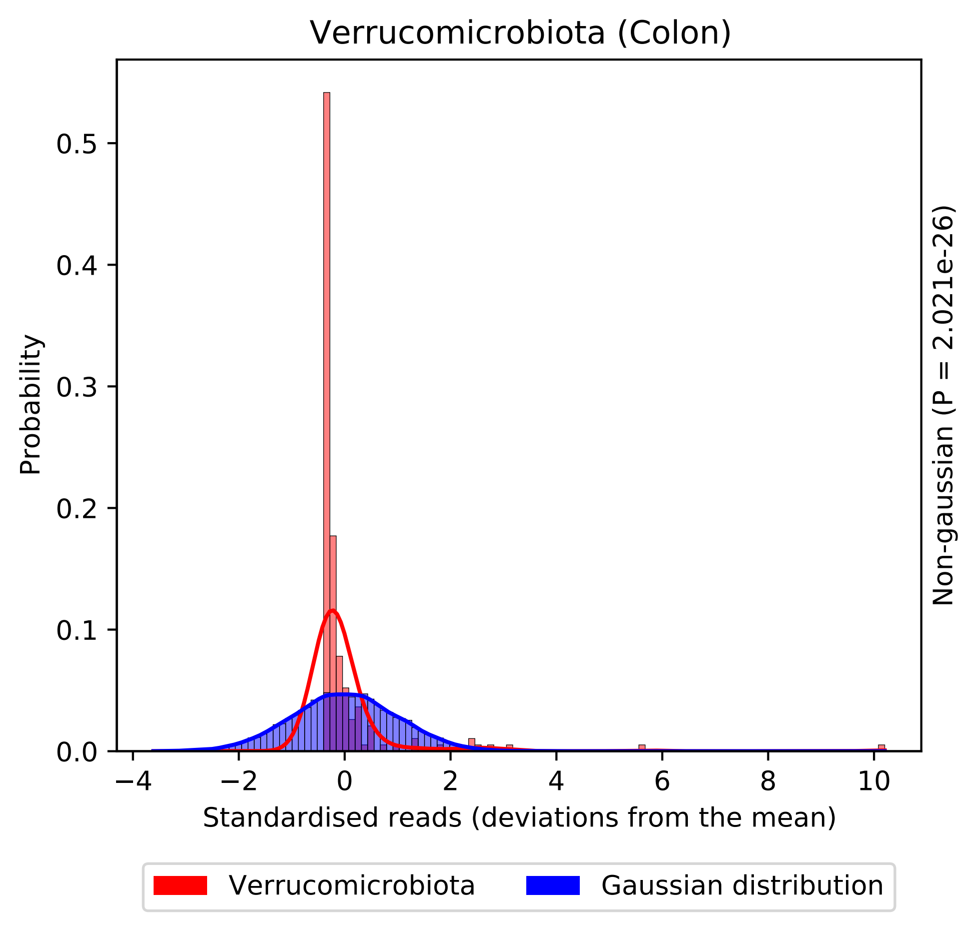
Verrucomicrobiota (Colon)
Probability
0.0
0.1
0.2
0.3
0.4
0.5
−4
−2
0
2
4
6
8
10
Standardised reads (deviations from the mean)
Non-gaussian (P = 2.021e-26)
Verrucomicrobiota
Gaussian distribution
