## Supplementary material for "statSuma: automated selection and performance of statistical comparisons for microbiome studies": SI Figure 3 (QQ plots)

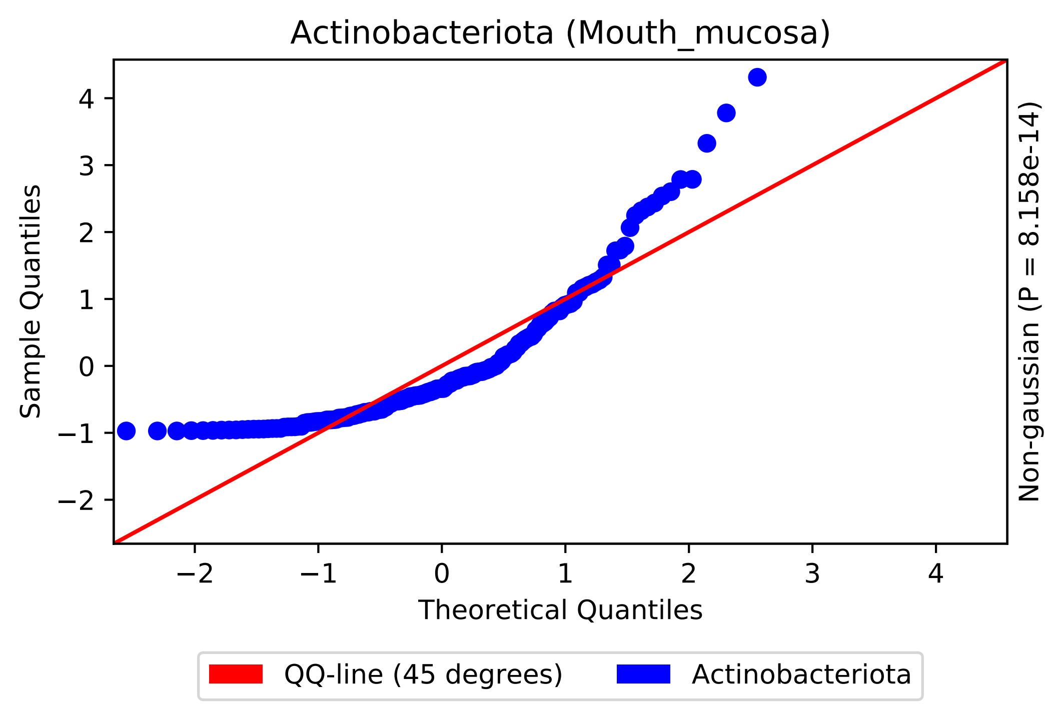

Actinobacteriota (Mouth_mucosa)
Sample Quantiles
Theoretical Quantiles
Non-gaussian (P = 8.158e-14)
−2
−1
0
1
2
3
4
QQ-line (45 degrees)
Actinobacteriota

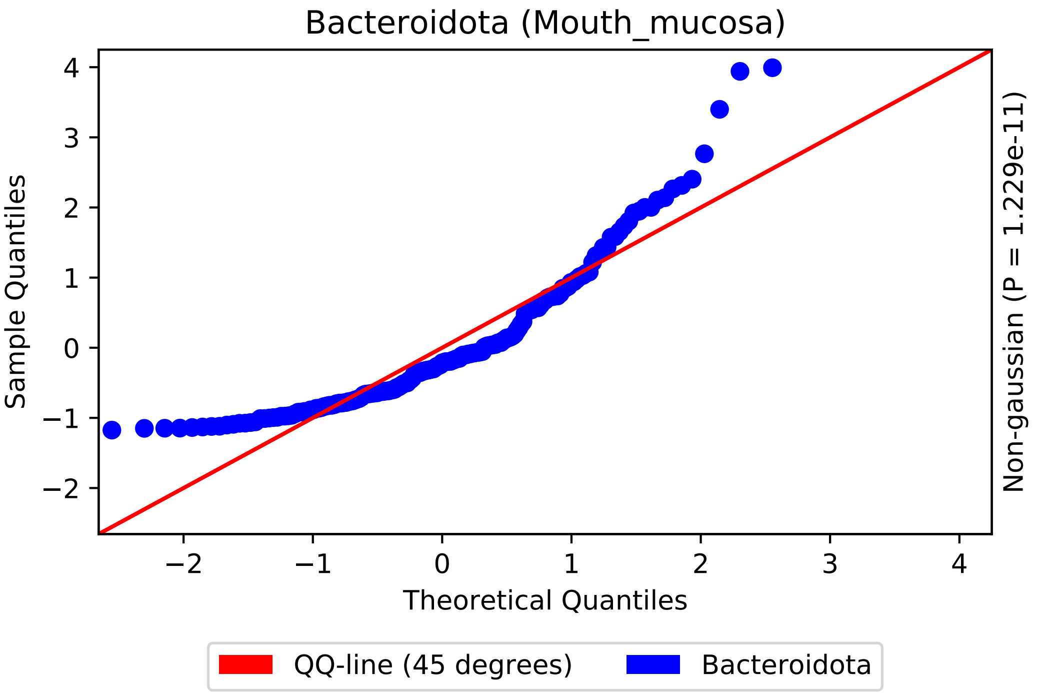
Bacteroidota (Mouth_mucosa)
Sample Quantiles
Theoretical Quantiles
Non-gaussian (P = 1.229e-11)
QQ-line (45 degrees)
Bacteroidota
4
3
2
1
0
−1
−2
−2
−1
0
1
2
3
4

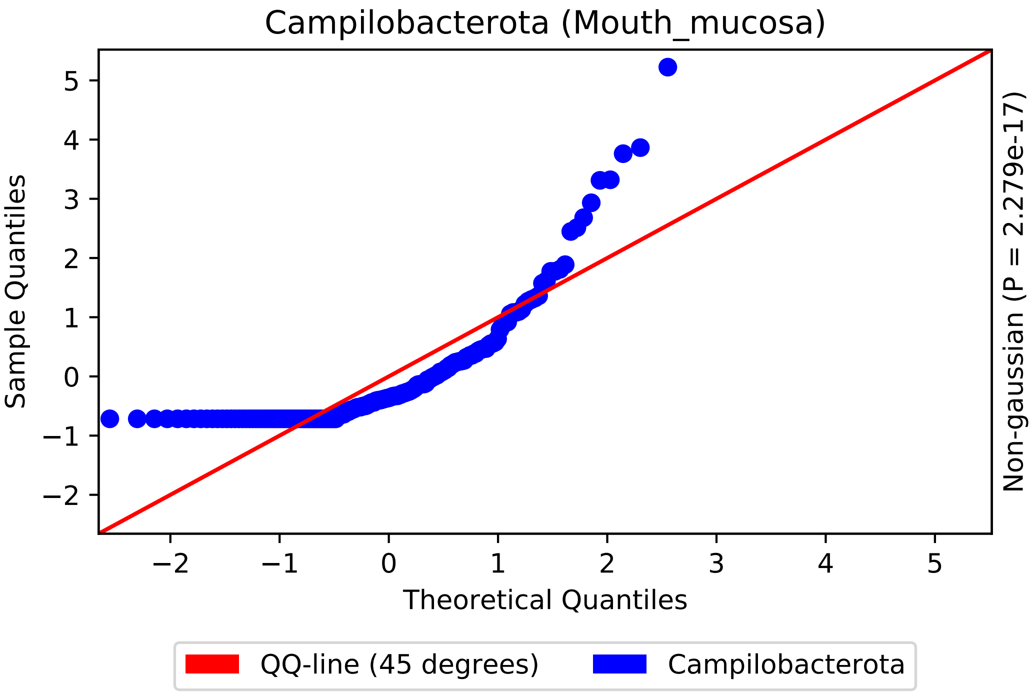

Campilobacterota (Mouth_mucosa)
Sample Quantiles
Theoretical Quantiles
Non-gaussian (P = 2.279e-17)
5
4
3
2
1
0
−1
−2
−2
−1
0
1
2
3
4
5
QQ-line (45 degrees)
Campilobacterota

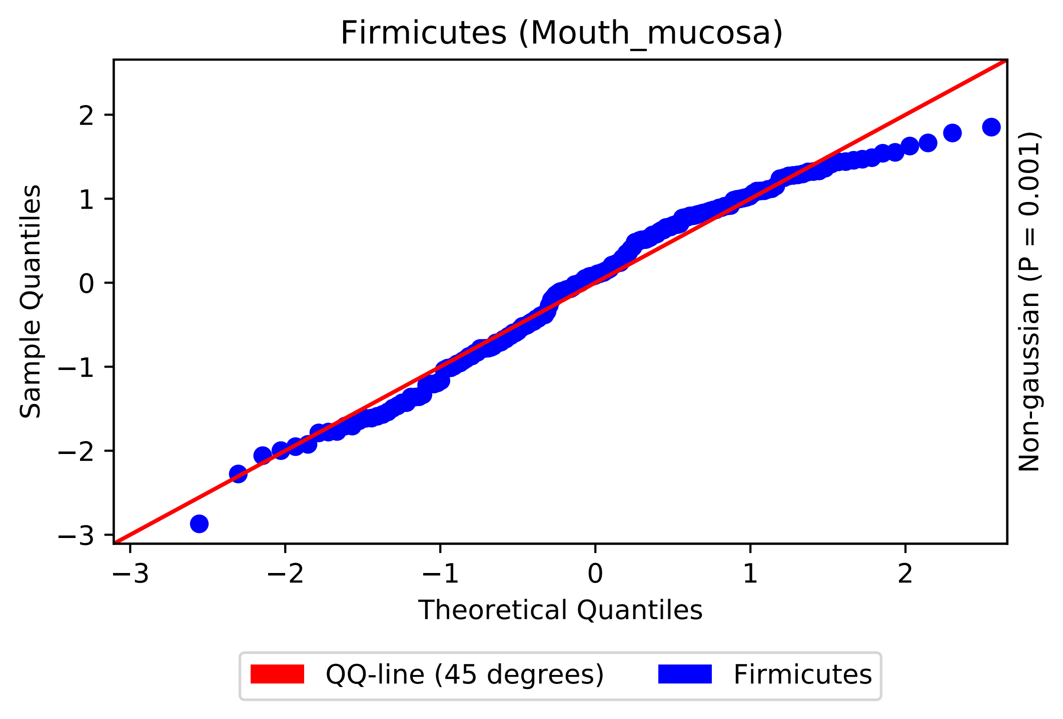

Firmicutes (Mouth_mucosa)
Sample Quantiles
Theoretical Quantiles
Non-gaussian (P = 0.001)
2
1
0
−1
−2
−3
−3
−2
−1
0
1
2
QQ-line (45 degrees)
Firmicutes

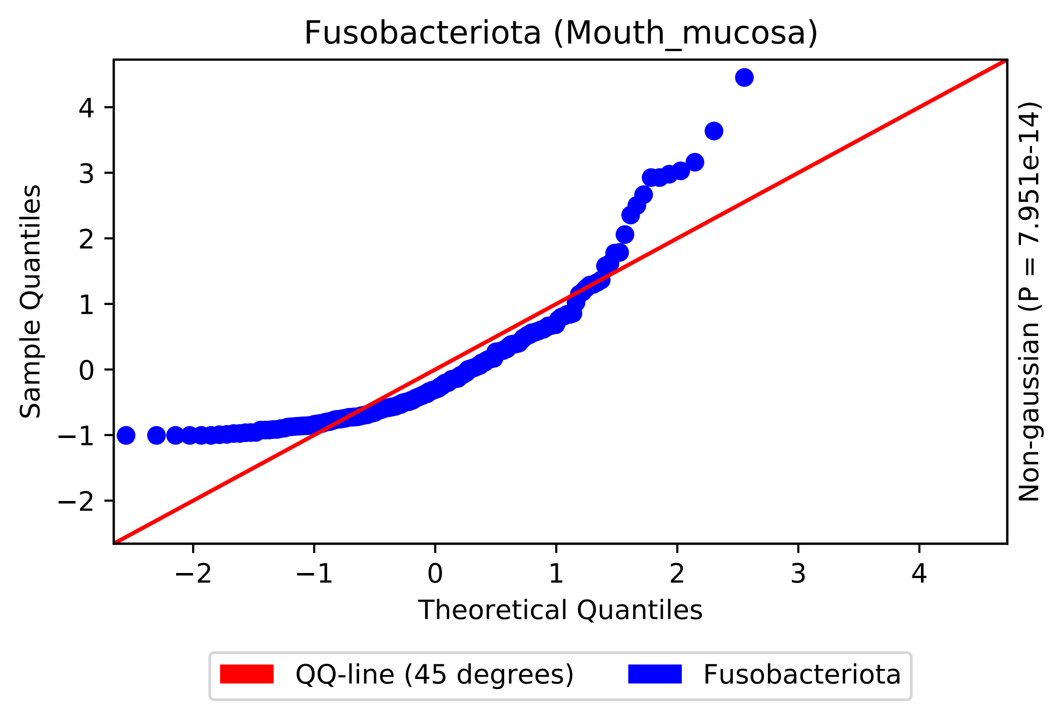
Fusobacteriota (Mouth_mucosa)
Sample Quantiles
Theoretical Quantiles
Non-gaussian (P = 7.951e-14)
−2
−1
0
1
2
3
4
QQ-line (45 degrees)
Fusobacteriota

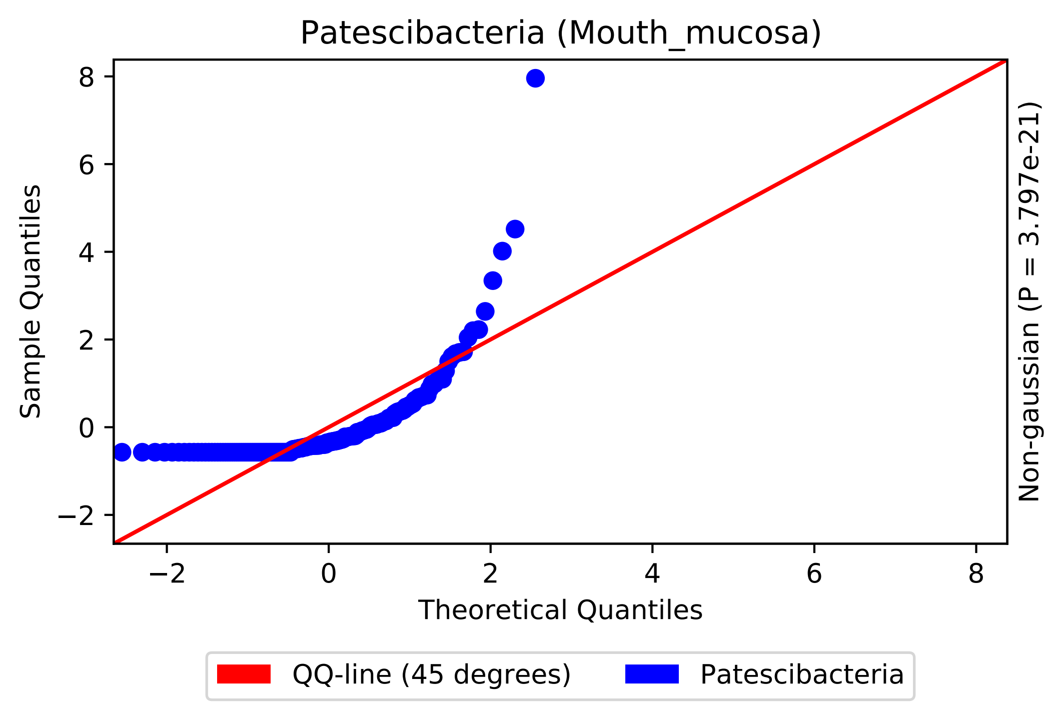
Patescibacteria (Mouth_mucosa)
Sample Quantiles
Theoretical Quantiles
Non-gaussian (P = 3.797e-21)
QQ-line (45 degrees)
Patescibacteria
8
6
4
2
0
−2
−2
0
2
4
6
8

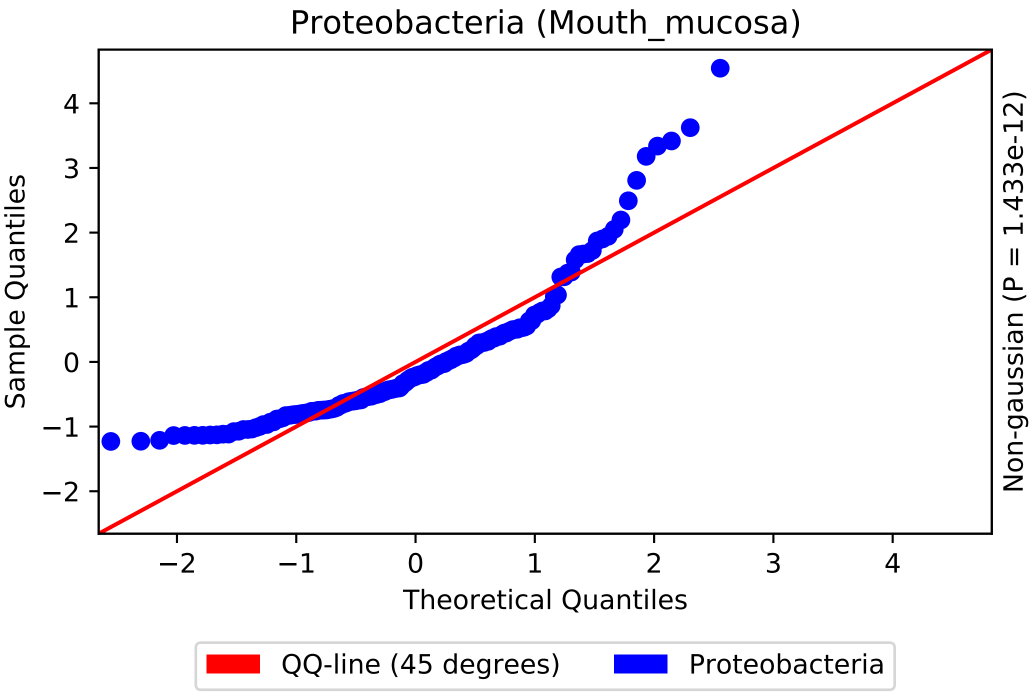

Proteobacteria (Mouth_mucosa)
Sample Quantiles
Theoretical Quantiles
Non-gaussian (P = 1.433e-12)
−2
−1
0
1
2
3
4
QQ-line (45 degrees)
Proteobacteria

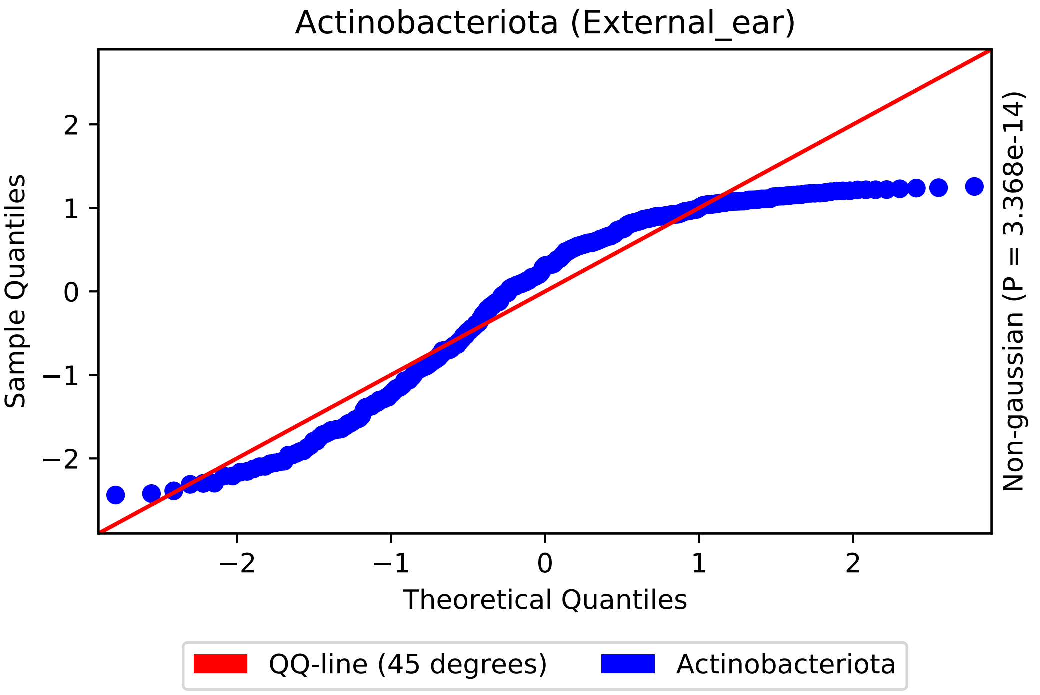

Actinobacteriota (External_ear)
Sample Quantiles
Theoretical Quantiles
Non-gaussian (P = 3.368e-14)
2
1
0
−1
−2
−2
−1
0
1
2
QQ-line (45 degrees)
Actinobacteriota

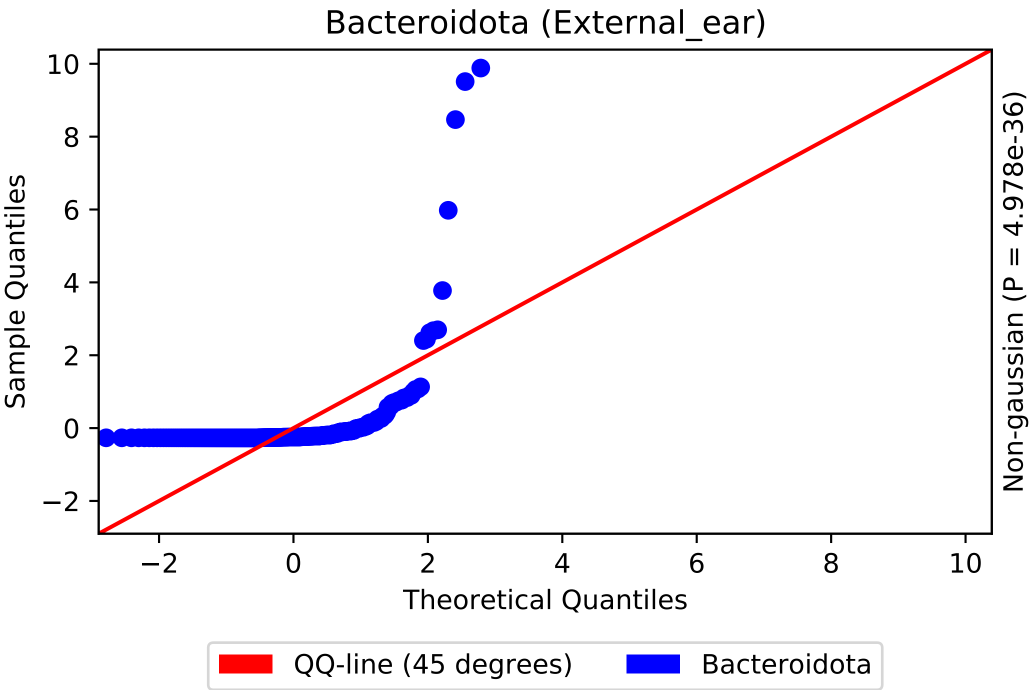
Bacteroidota (External_ear)
Sample Quantiles
Theoretical Quantiles
Non-gaussian (P = 4.978e-36)
QQ-line (45 degrees)
Bacteroidota
10
8
6
4
2
0
−2
−2
0
2
4
6
8
10

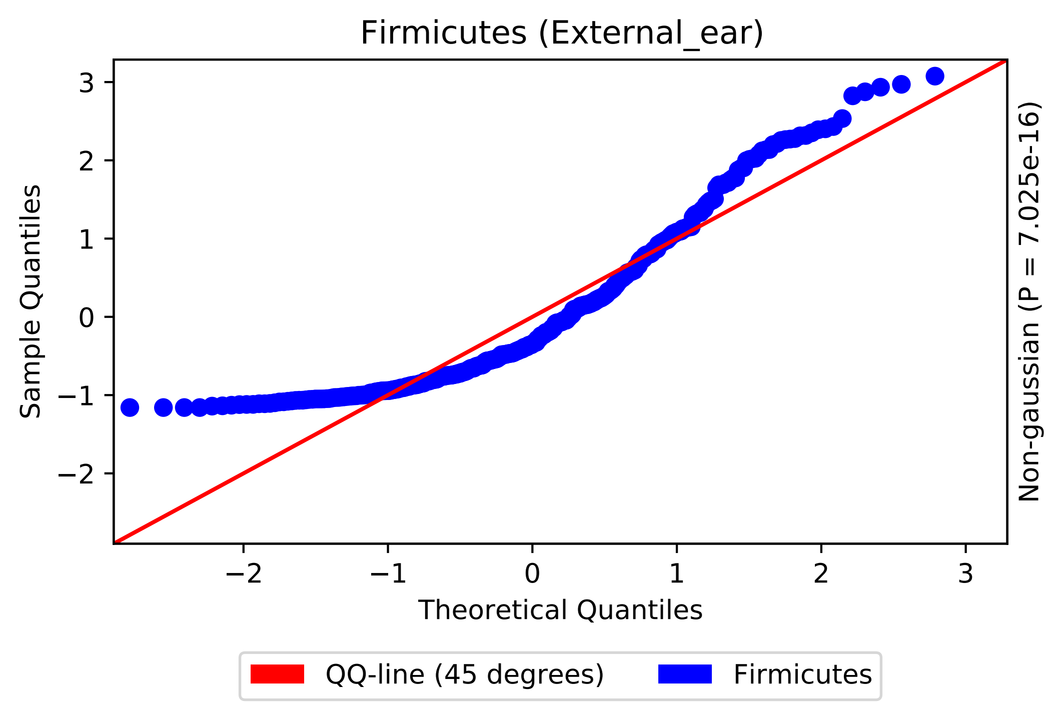
Firmicutes (External_ear)
Sample Quantiles
Theoretical Quantiles
Non-gaussian (P = 7.025e-16)
QQ-line (45 degrees)
Firmicutes
3
2
1
0
−1
−2
−2
−1
0
1
2
3

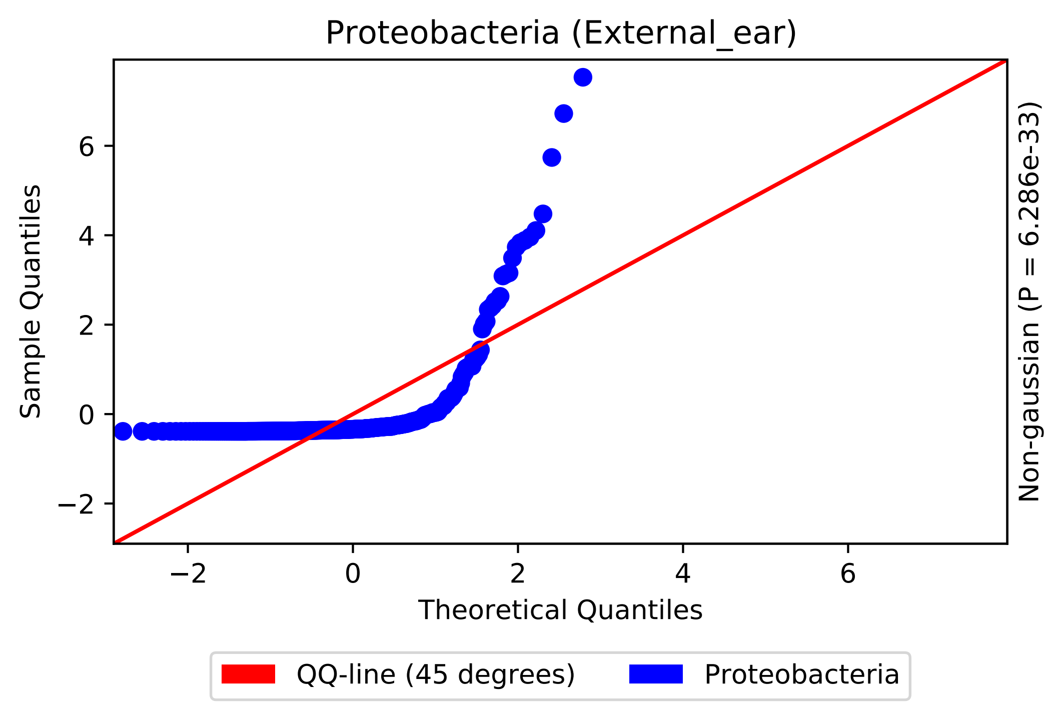
Proteobacteria (External_ear)
Sample Quantiles
Theoretical Quantiles
Non-gaussian (P = 6.286e-33)
−2
0
2
4
6
QQ-line (45 degrees)
Proteobacteria

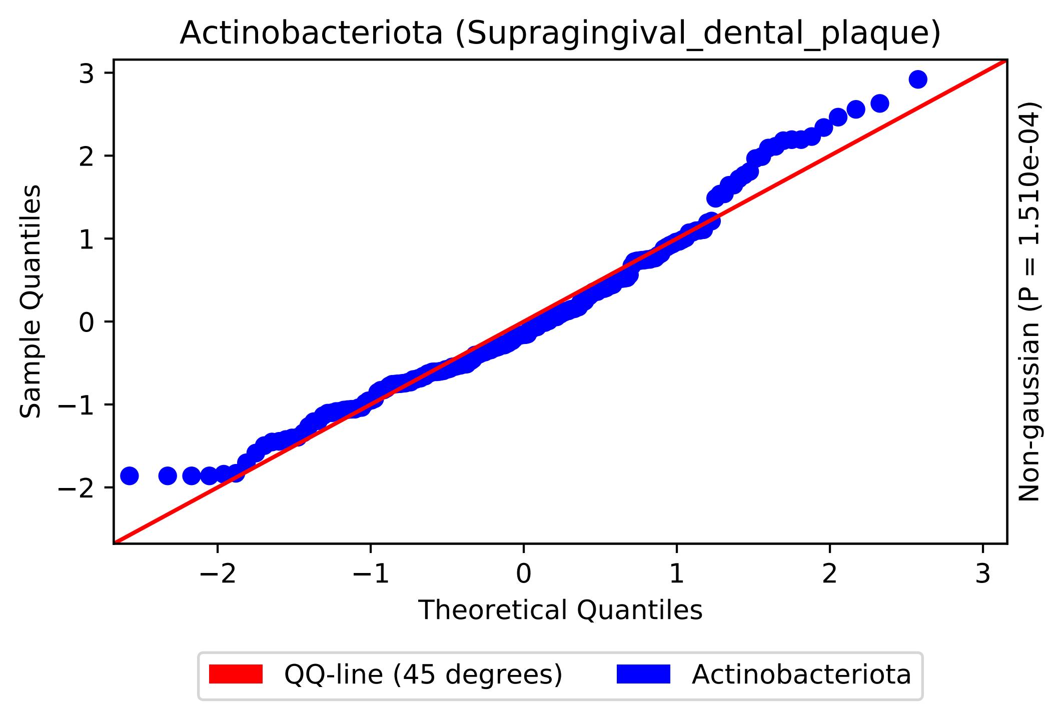

Actinobacteriota (Supragingival_dental_plaque)
Sample Quantiles
Theoretical Quantiles
Non-gaussian (P = 1.510e-04)
QQ-line (45 degrees)
Actinobacteriota
3
2
1
0
−1
−2
−2
−1
0
1
2
3

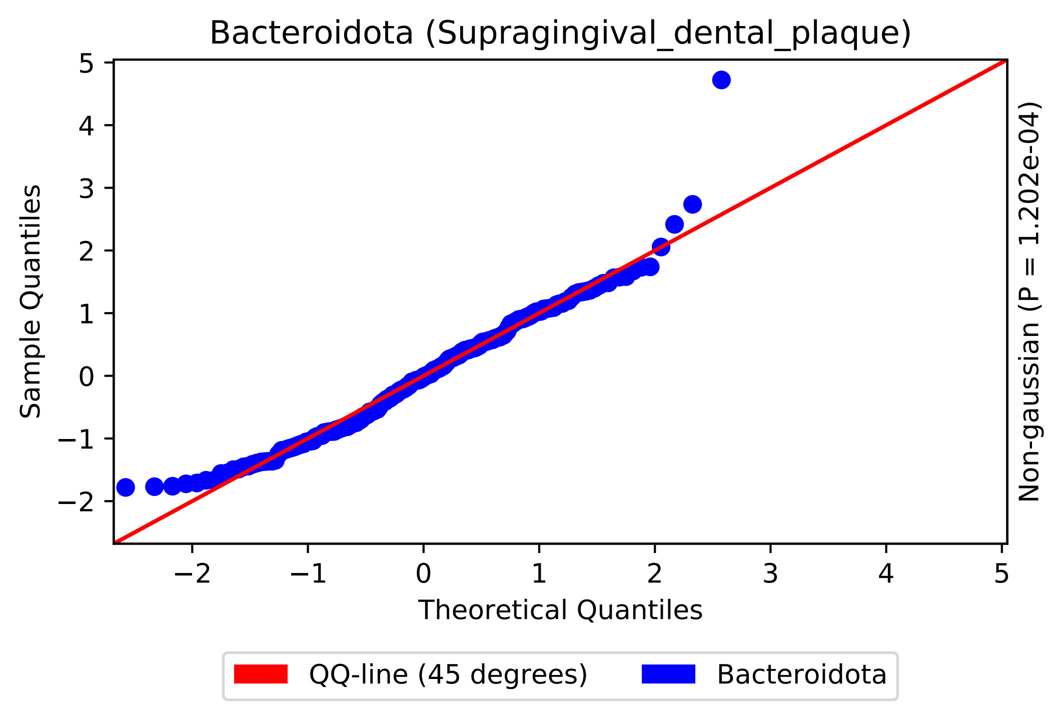
Bacteroidota (Supragingival_dental_plaque)
Sample Quantiles
Theoretical Quantiles
Non-gaussian (P = 1.202e-04)
QQ-line (45 degrees)
Bacteroidota

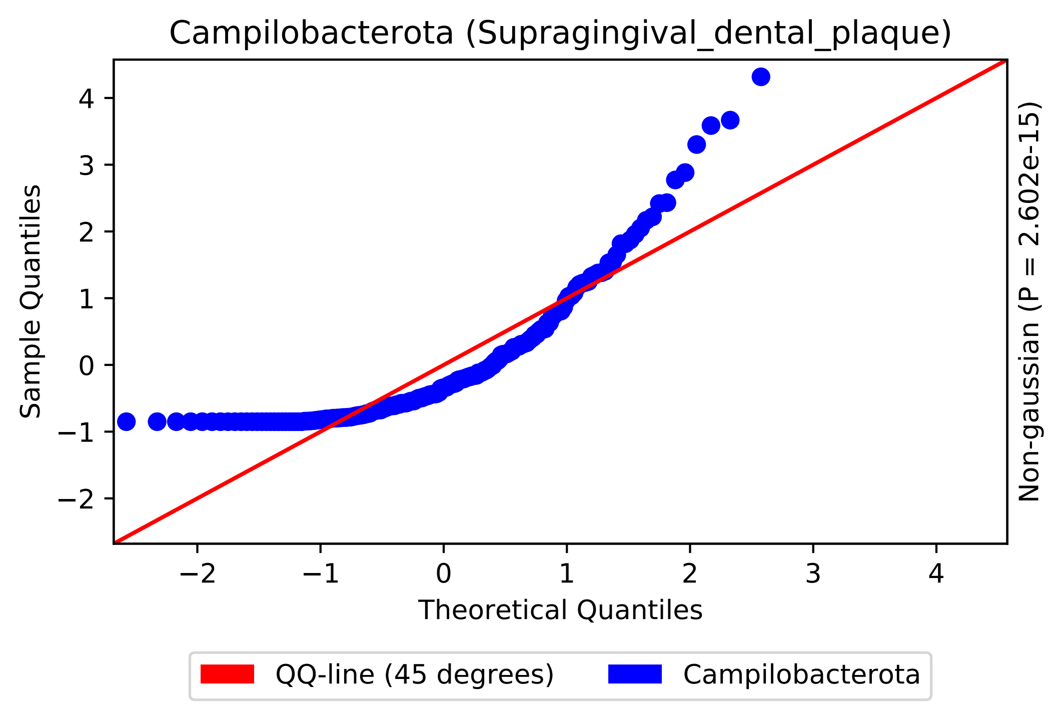

Campilobacterota (Supragingival_dental_plaque)
Sample Quantiles
Theoretical Quantiles
Non-gaussian (P = 2.602e-15)
4
3
2
1
0
−1
−2
−2
−1
0
1
2
3
4
QQ-line (45 degrees)
Campilobacterota

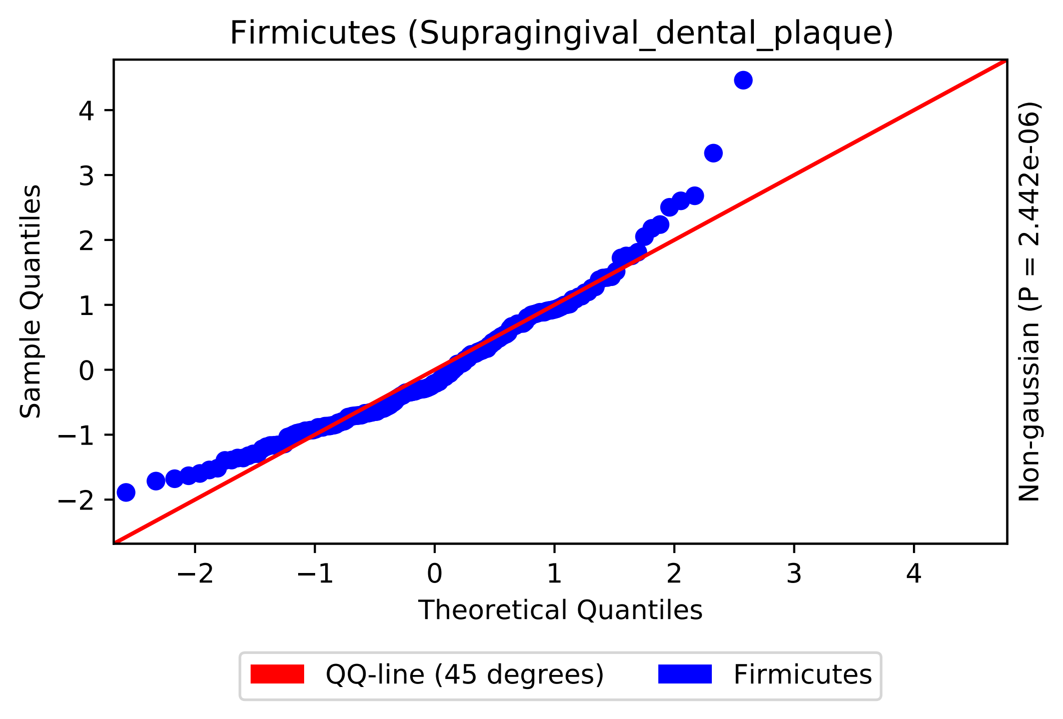
Firmicutes (Supragingival_dental_plaque)
Sample Quantiles
Theoretical Quantiles
Non-gaussian (P = 2.442e-06)
QQ-line (45 degrees)
Firmicutes

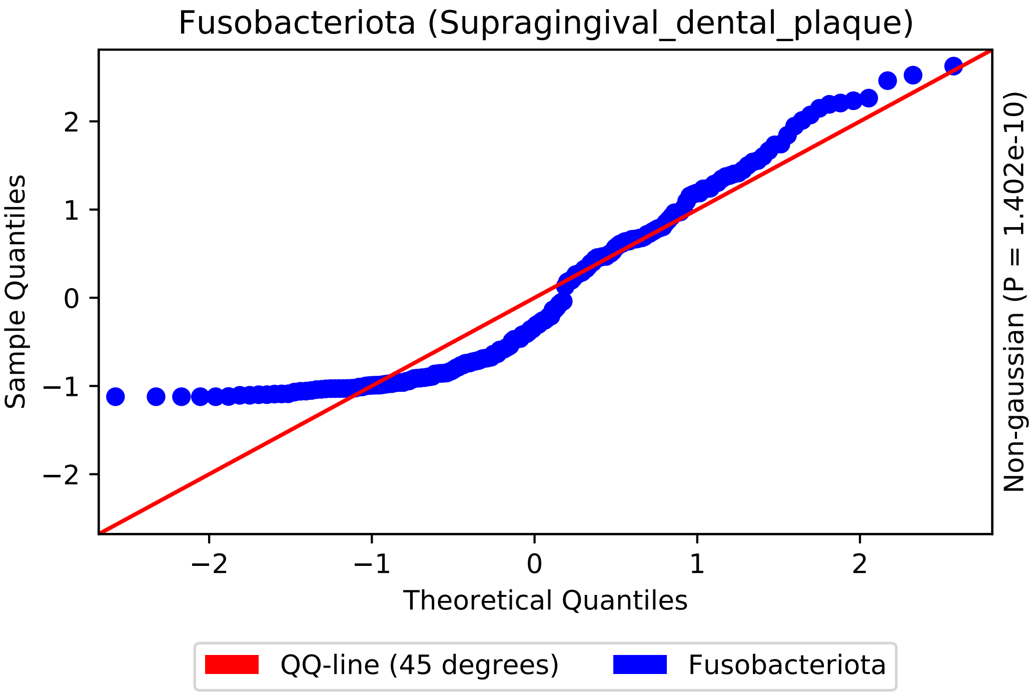
Fusobacteriota (Supragingival_dental_plaque)
Sample Quantiles
Theoretical Quantiles
Non-gaussian (P = 1.402e-10)
QQ-line (45 degrees)
Fusobacteriota
−2
−1
0
1
2

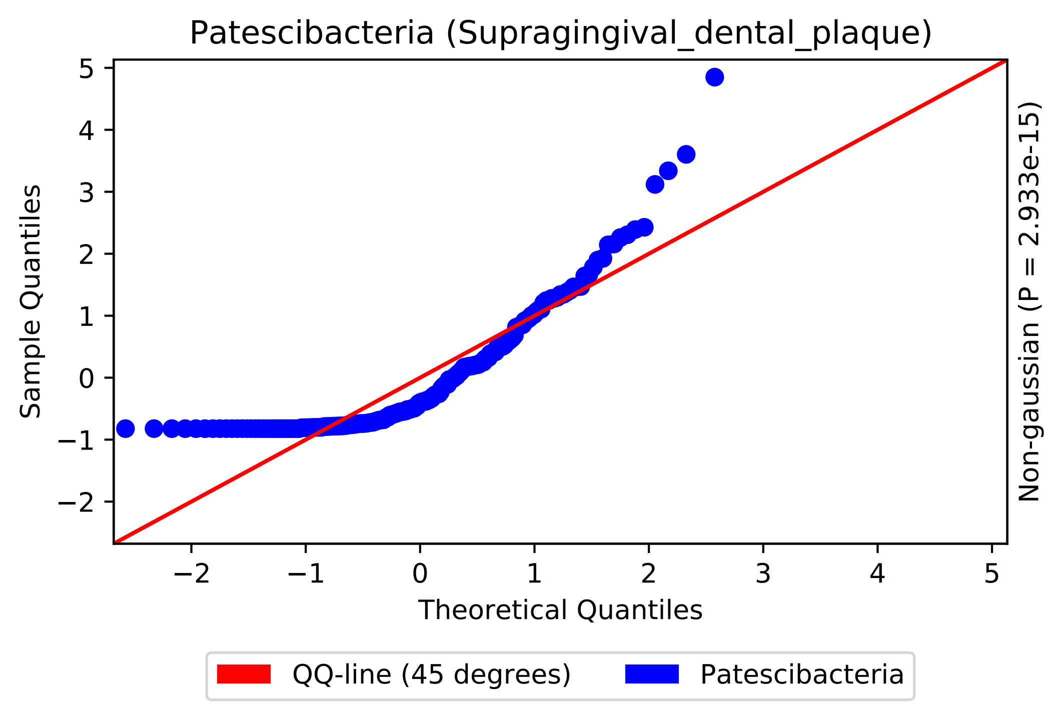
Patescibacteria (Supragingival_dental_plaque)
Sample Quantiles
Theoretical Quantiles
Non-gaussian (P = 2.933e-15)
QQ-line (45 degrees)
Patescibacteria
5
4
3
2
1
0
−1
−2
−2
−1
0
1
2
3
4
5

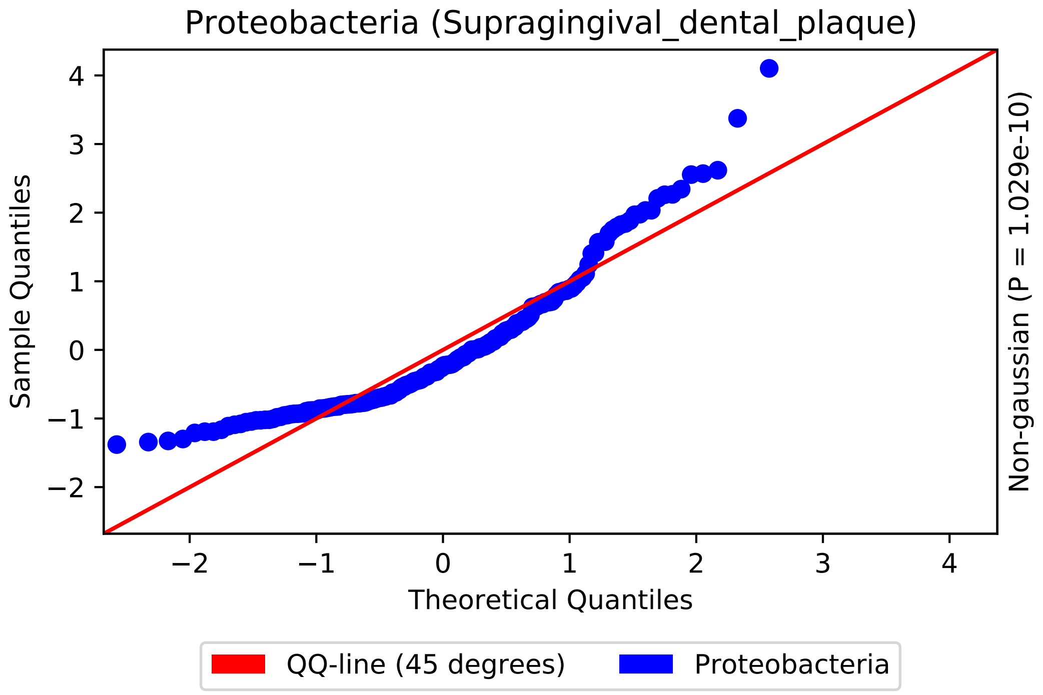
Proteobacteria (Supragingival_dental_plaque)
Sample Quantiles
Theoretical Quantiles
Non-gaussian (P = 1.029e-10)
QQ-line (45 degrees)
Proteobacteria
4
3
2
1
0
−1
−2
−2
−1
0
1
2
3
4

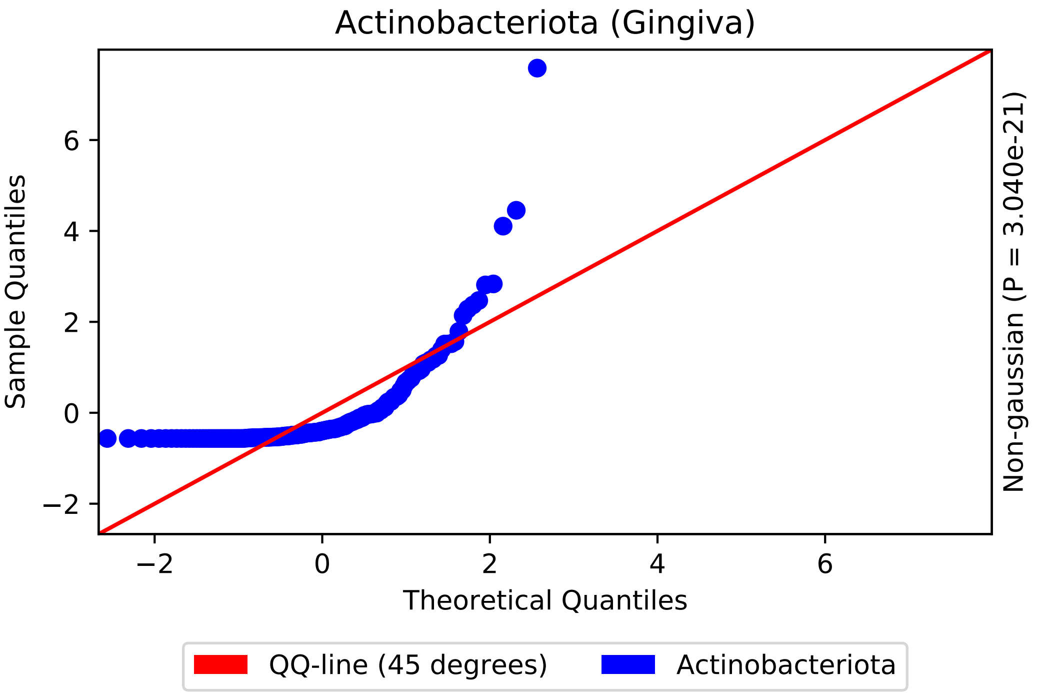
Actinobacteriota (Gingiva)
Sample Quantiles
Theoretical Quantiles
Non-gaussian (P = 3.040e-21)
QQ-line (45 degrees)
Actinobacteriota
−2
0
2
4
6

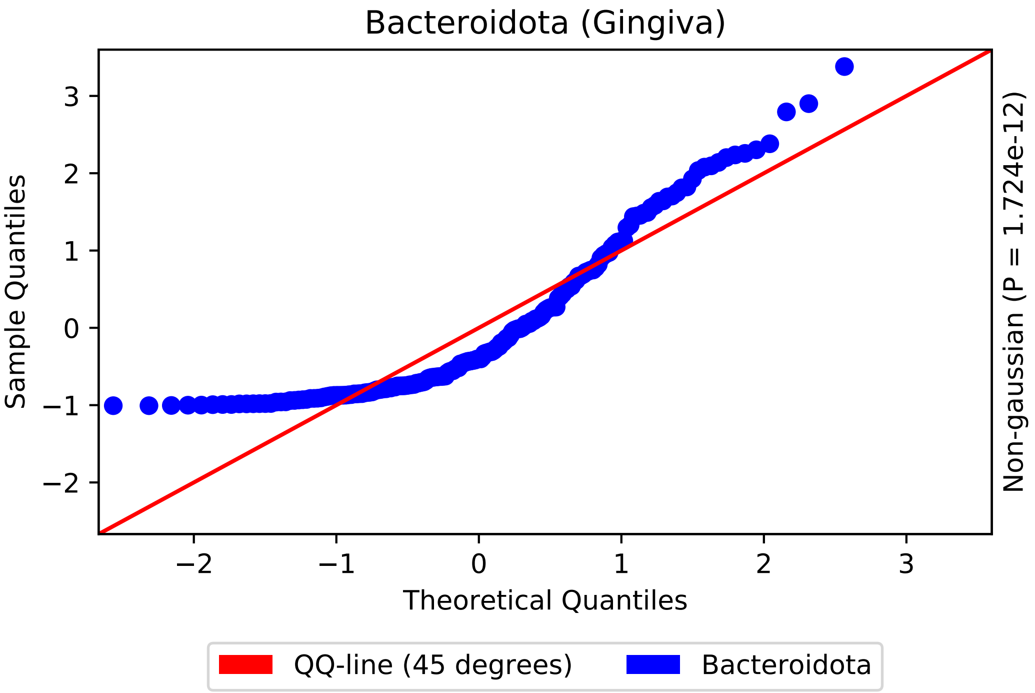

Bacteroidota (Gingiva)
Sample Quantiles
Theoretical Quantiles
Non-gaussian (P = 1.724e-12)
−2
−1
0
1
2
3
QQ-line (45 degrees)
Bacteroidota

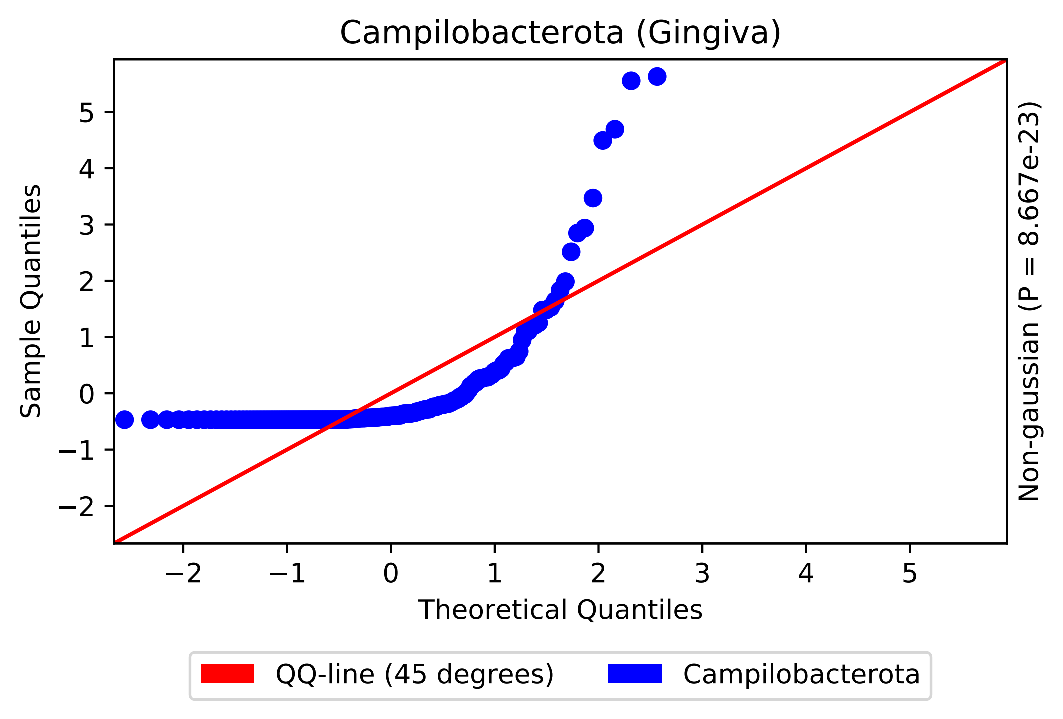

Campilobacterota (Gingiva)
Sample Quantiles
Theoretical Quantiles
Non-gaussian (P = 8.667e-23)
QQ-line (45 degrees)
Campilobacterota

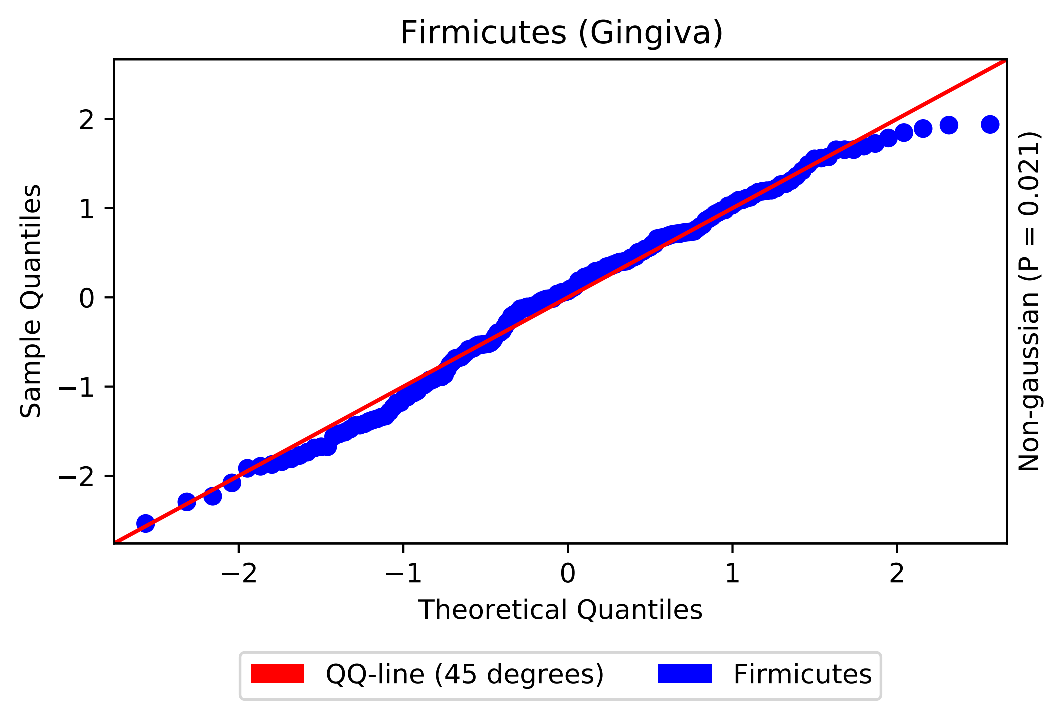

Firmicutes (Gingiva)
Sample Quantiles
Theoretical Quantiles
Non-gaussian (P = 0.021)
QQ-line (45 degrees)
Firmicutes
2
1
0
−1
−2

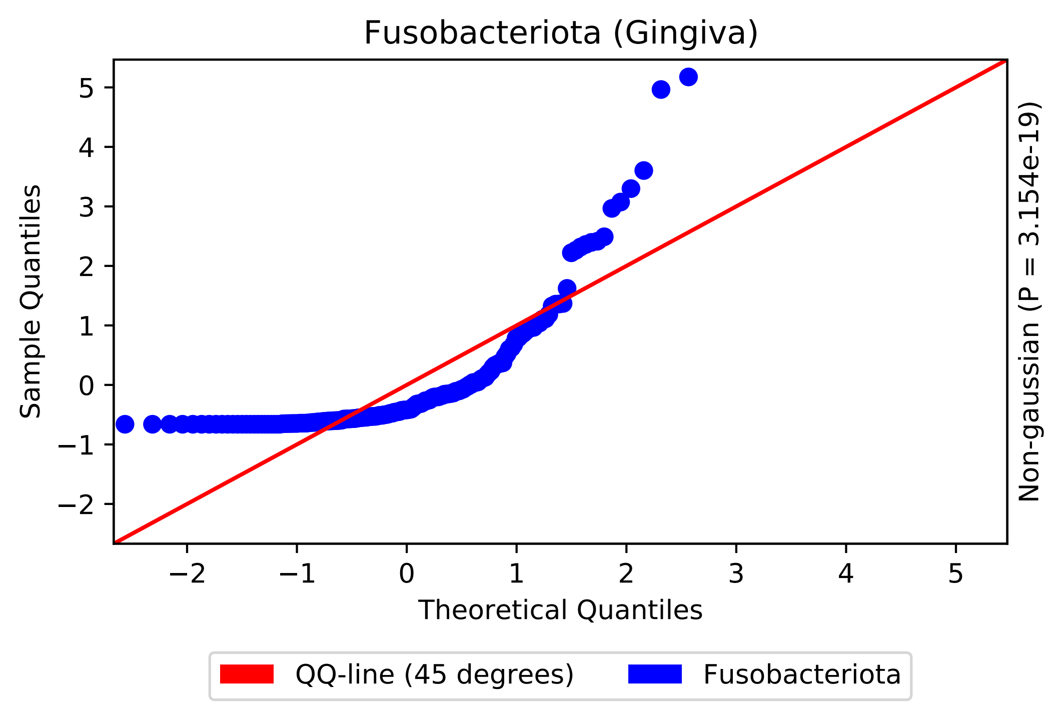

Fusobacteriota (Gingiva)
Sample Quantiles
Theoretical Quantiles
Non-gaussian (P = 3.154e-19)
−2
−1
0
1
2
3
4
5
QQ-line (45 degrees)
Fusobacteriota

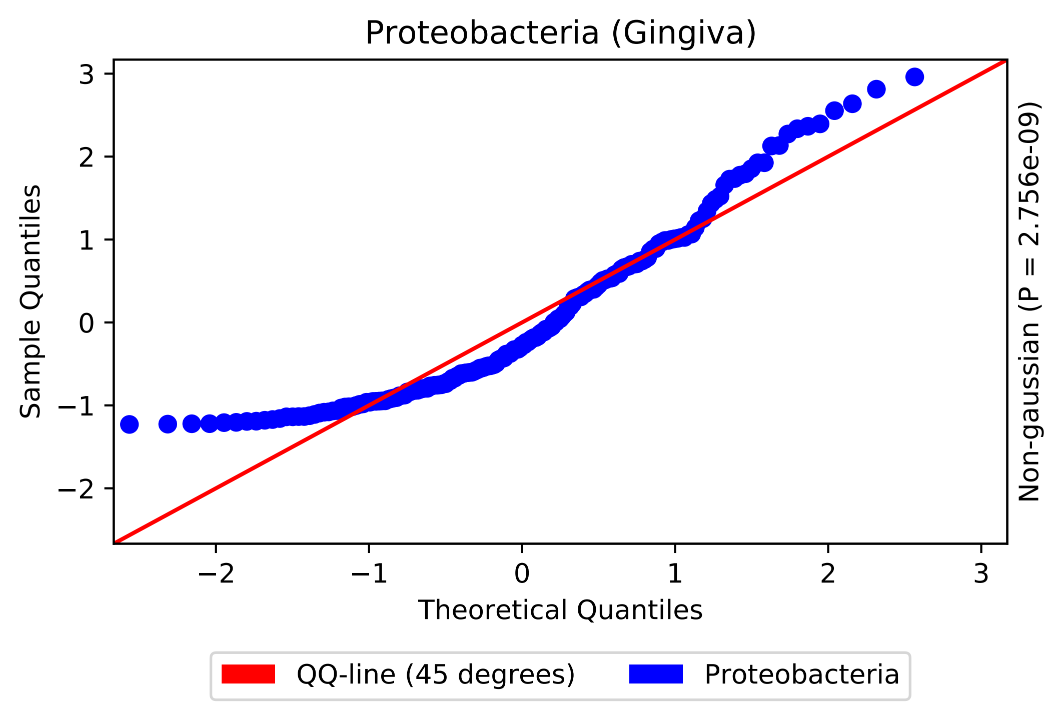
Proteobacteria (Gingiva)
Sample Quantiles
Theoretical Quantiles
Non-gaussian (P = 2.756e-09)
3
2
1
0
−1
−2
−2
−1
0
1
2
3
QQ-line (45 degrees)
Proteobacteria

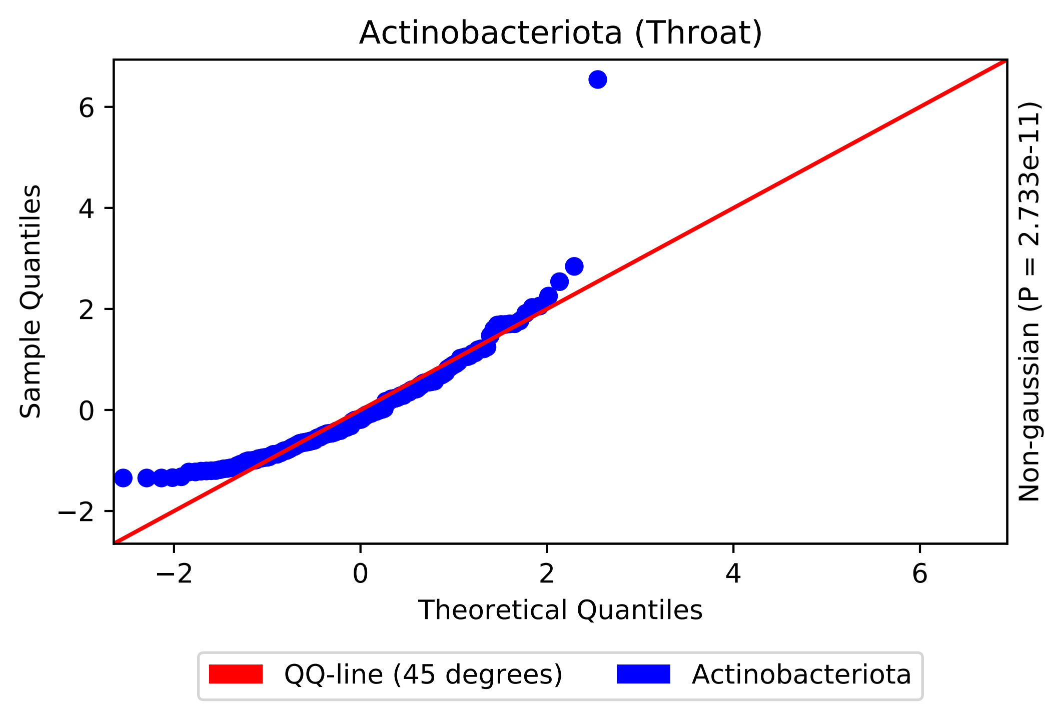
Actinobacteriota (Throat)
Sample Quantiles
Theoretical Quantiles
Non-gaussian (P = 2.733e-11)
−2
0
2
4
6
QQ-line (45 degrees)
Actinobacteriota

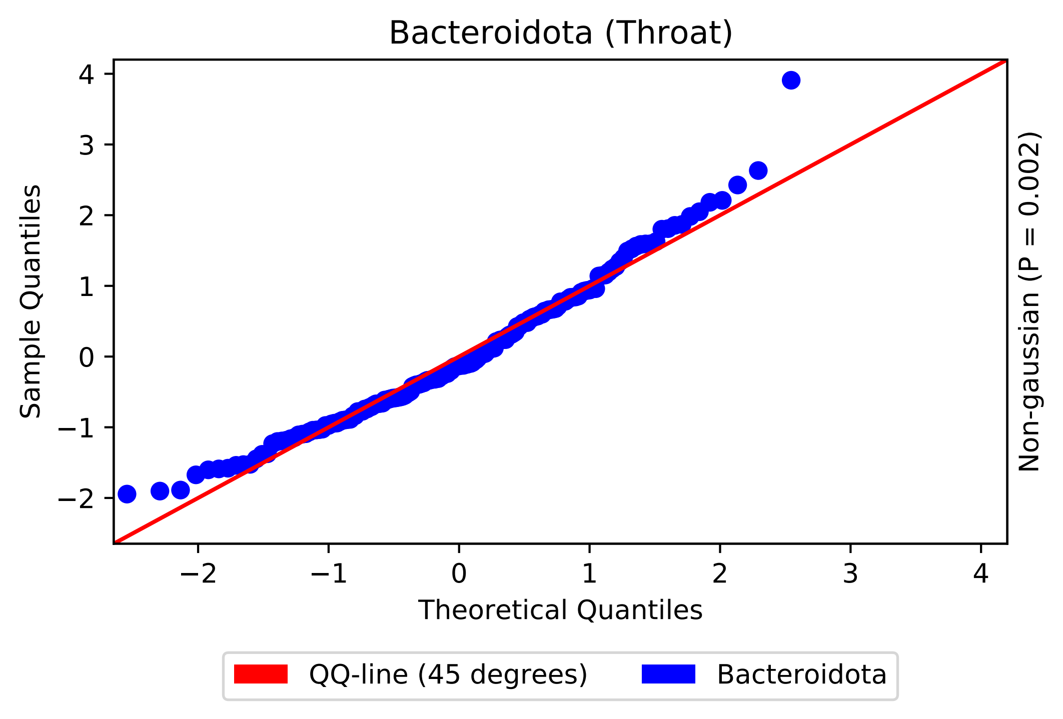
Bacteroidota (Throat)
Sample Quantiles
4
3
2
1
0
−1
−2
−2
−1
0
1
2
3
4
Theoretical Quantiles
Non-gaussian (P = 0.002)
QQ-line (45 degrees)
Bacteroidota

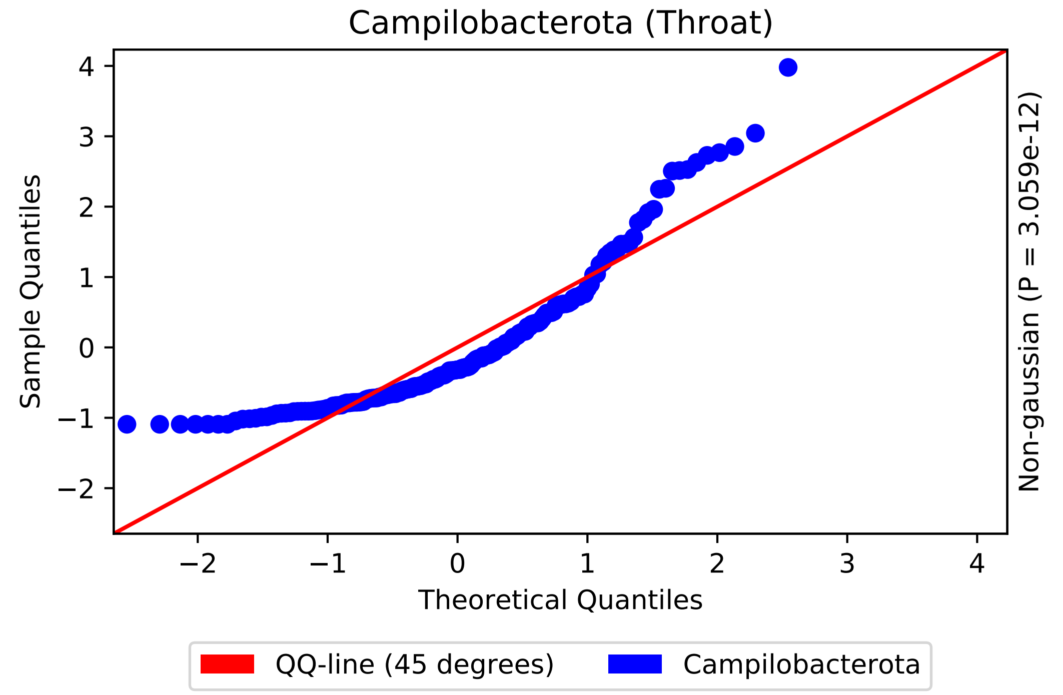
Campilobacterota (Throat)
Sample Quantiles
Theoretical Quantiles
Non-gaussian (P = 3.059e-12)
−2
−1
0
1
2
3
4
QQ-line (45 degrees)
Campilobacterota

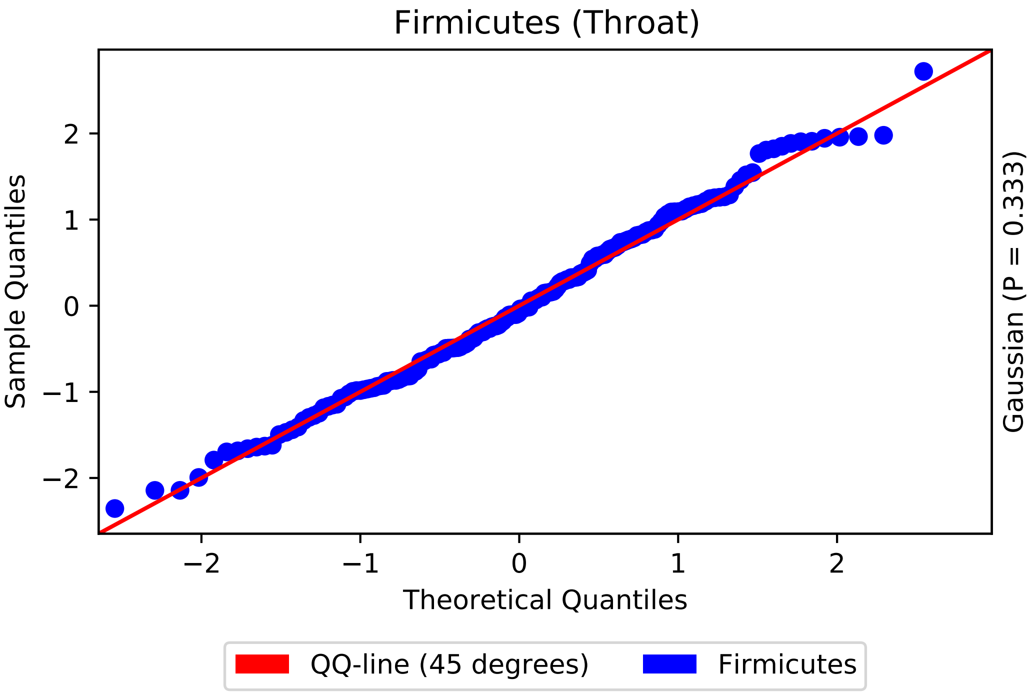

Firmicutes (Throat)
Sample Quantiles
Theoretical Quantiles
Gaussian (P = 0.333)
QQ-line (45 degrees)
Firmicutes
−2
−1
0
1
2

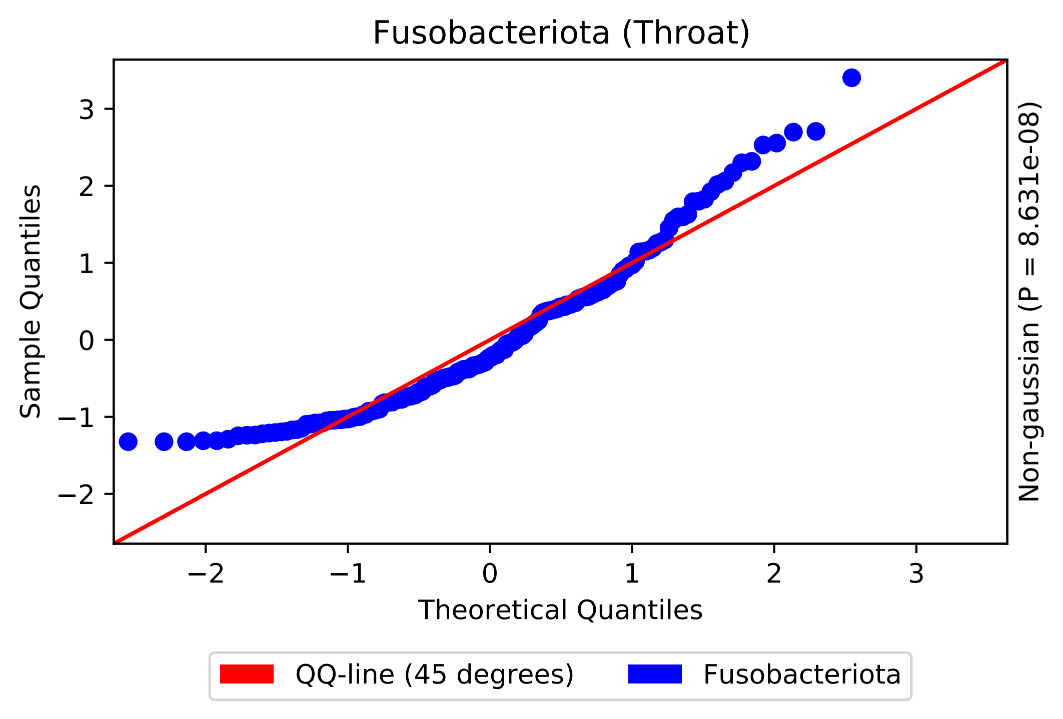
Fusobacteriota (Throat)
Sample Quantiles
Theoretical Quantiles
Non-gaussian (P = 8.631e-08)
3
2
1
0
−1
−2
−2
−1
0
1
2
3
QQ-line (45 degrees)
Fusobacteriota

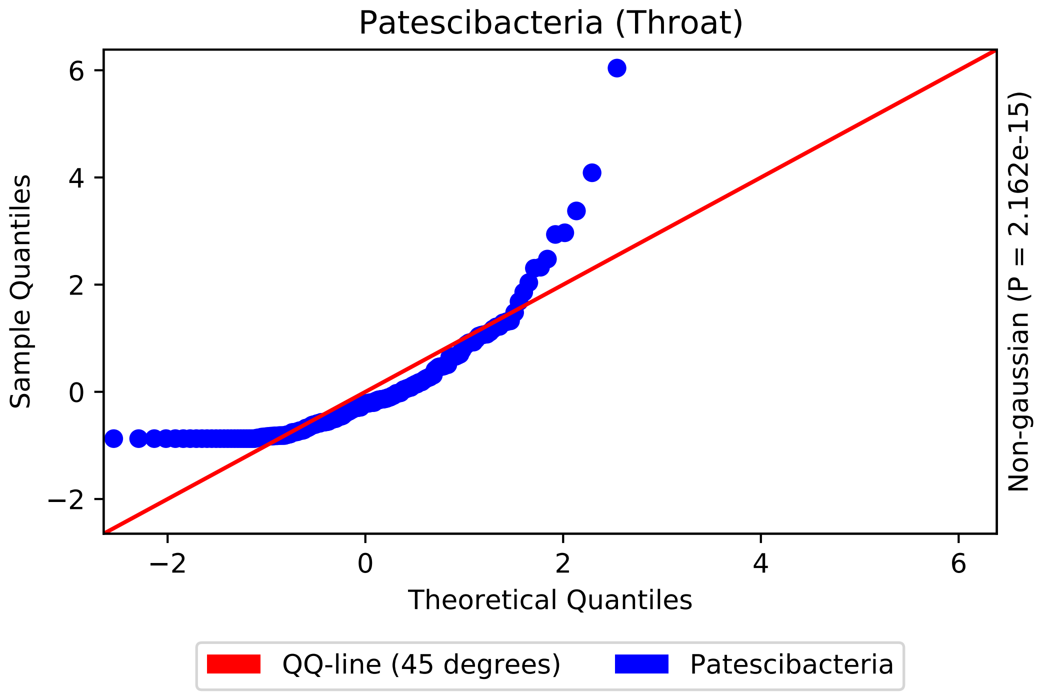
Patescibacteria (Throat)
Sample Quantiles
Theoretical Quantiles
Non-gaussian (P = 2.162e-15)
6
4
2
0
−2
−2
0
2
4
6
QQ-line (45 degrees)
Patescibacteria

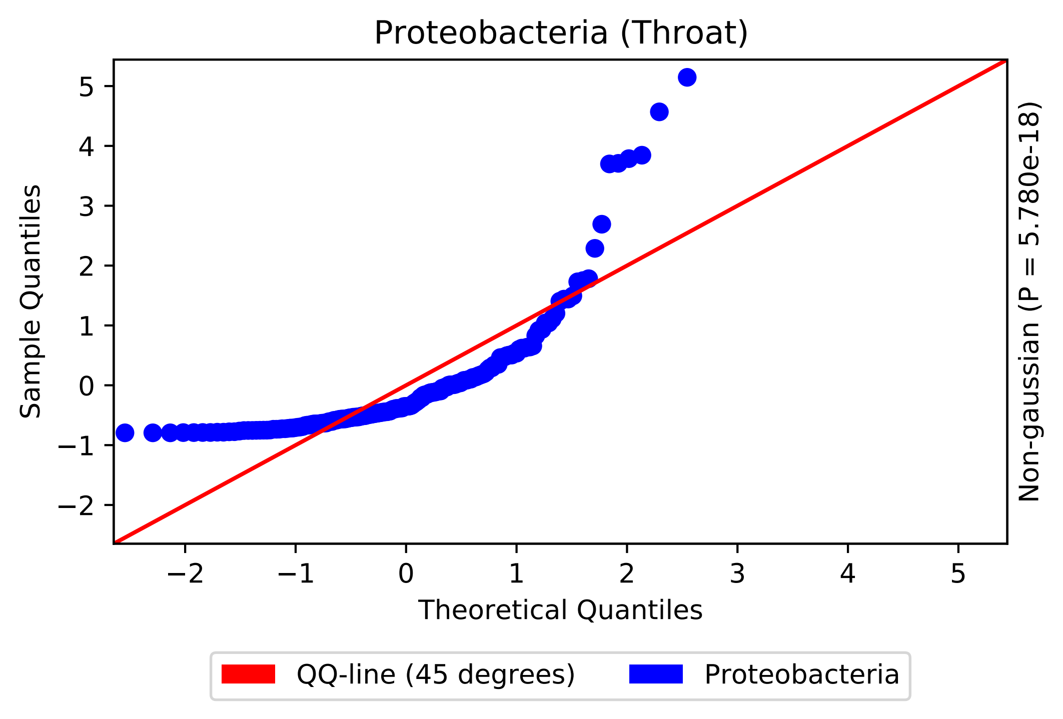

Proteobacteria (Throat)
Sample Quantiles
Theoretical Quantiles
Non-gaussian (P = 5.780e-18)
QQ-line (45 degrees)
Proteobacteria

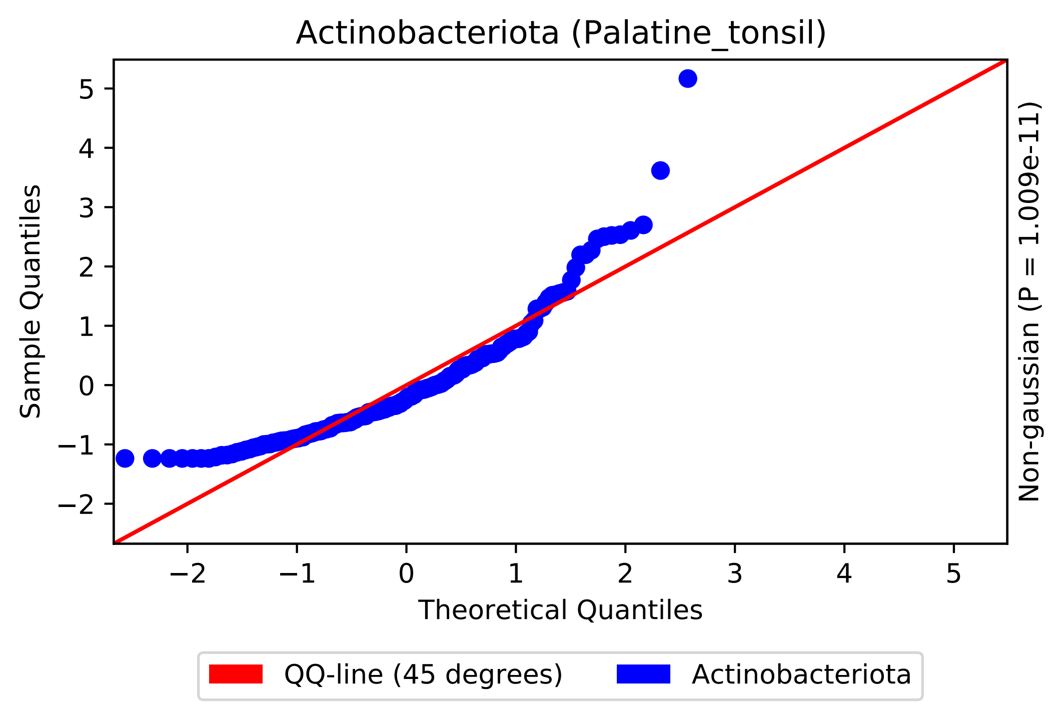

Actinobacteriota (Palatine_tonsil)
Sample Quantiles
Theoretical Quantiles
Non-gaussian (P = 1.009e-11)
QQ-line (45 degrees)
Actinobacteriota

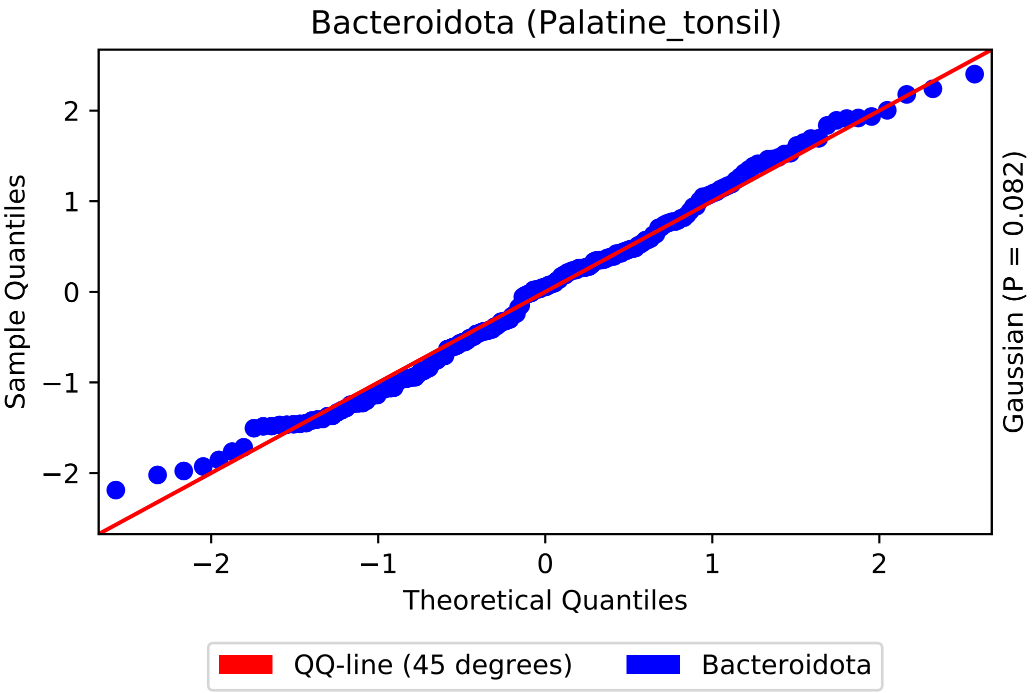
Bacteroidota (Palatine_tonsil)
Sample Quantiles
Theoretical Quantiles
Gaussian (P = 0.082)
2
1
0
−1
−2
−2
−1
0
1
2
QQ-line (45 degrees)
Bacteroidota

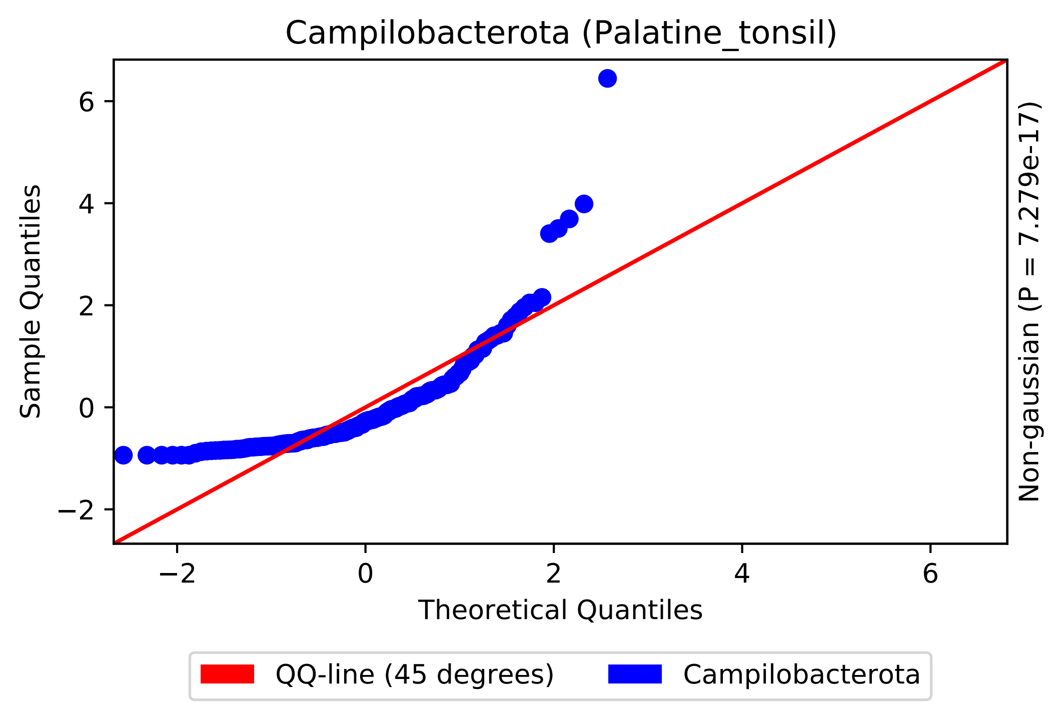
Campilobacterota (Palatine_tonsil)
Sample Quantiles
Theoretical Quantiles
Non-gaussian (P = 7.279e-17)
−2
0
2
4
6
QQ-line (45 degrees)
Campilobacterota

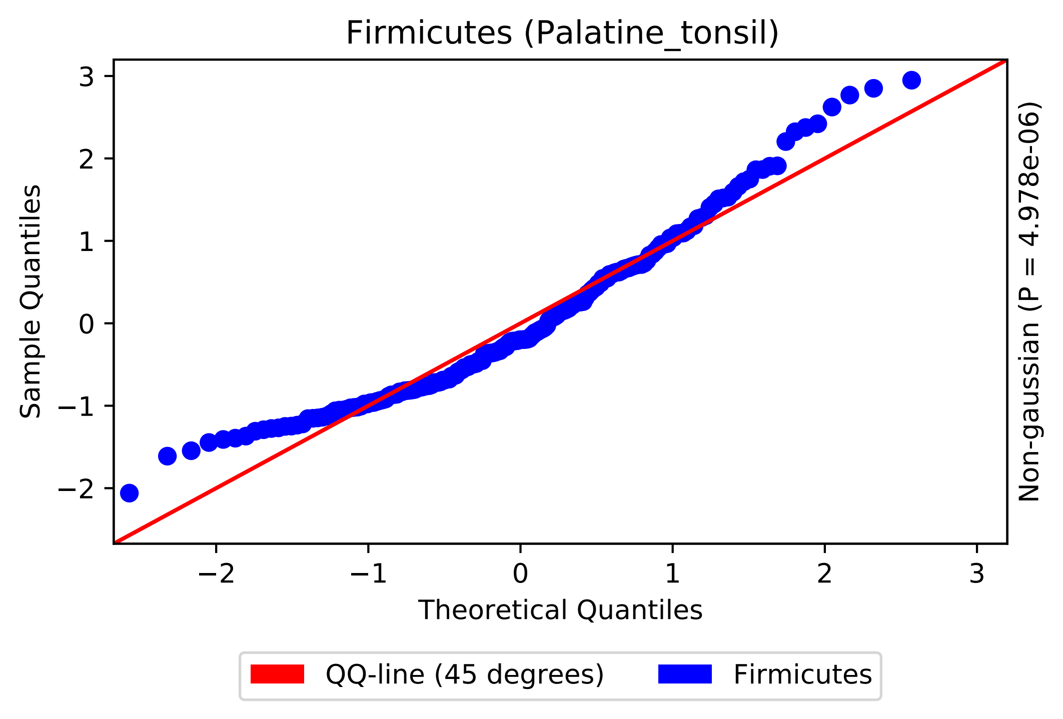
Firmicutes (Palatine_tonsil)
Sample Quantiles
Theoretical Quantiles
Non-gaussian (P = 4.978e-06)
QQ-line (45 degrees)
Firmicutes
3
2
1
0
−1
−2
−2
−1
0
1
2
3

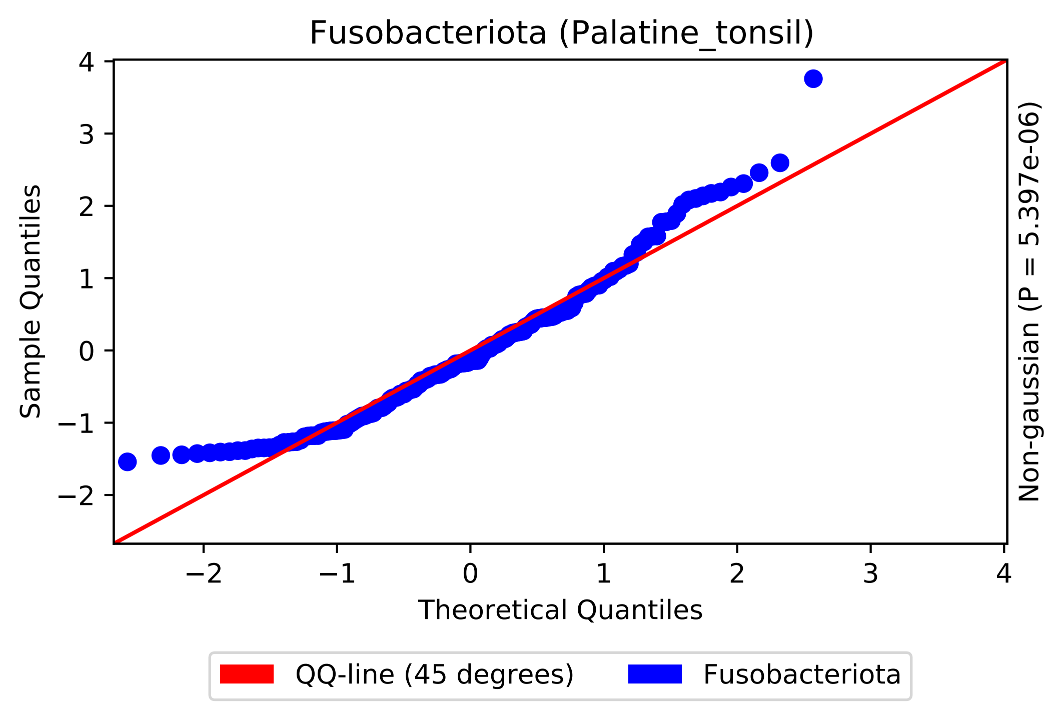

Fusobacteriota (Palatine_tonsil)
Sample Quantiles
Theoretical Quantiles
Non-gaussian (P = 5.397e-06)
QQ-line (45 degrees)
Fusobacteriota
4
3
2
1
0
−1
−2
−2
−1
0
1
2
3
4

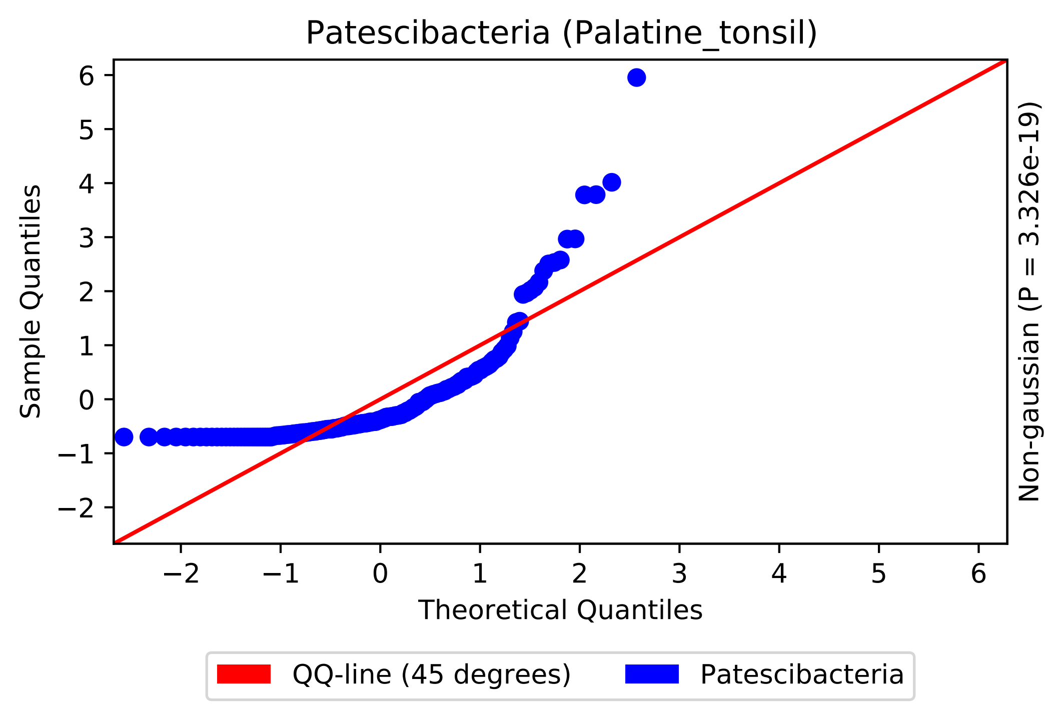

Patescibacteria (Palatine_tonsil)
Sample Quantiles
Theoretical Quantiles
Non-gaussian (P = 3.326e-19)
−2
−1
0
1
2
3
4
5
6
QQ-line (45 degrees)
Patescibacteria

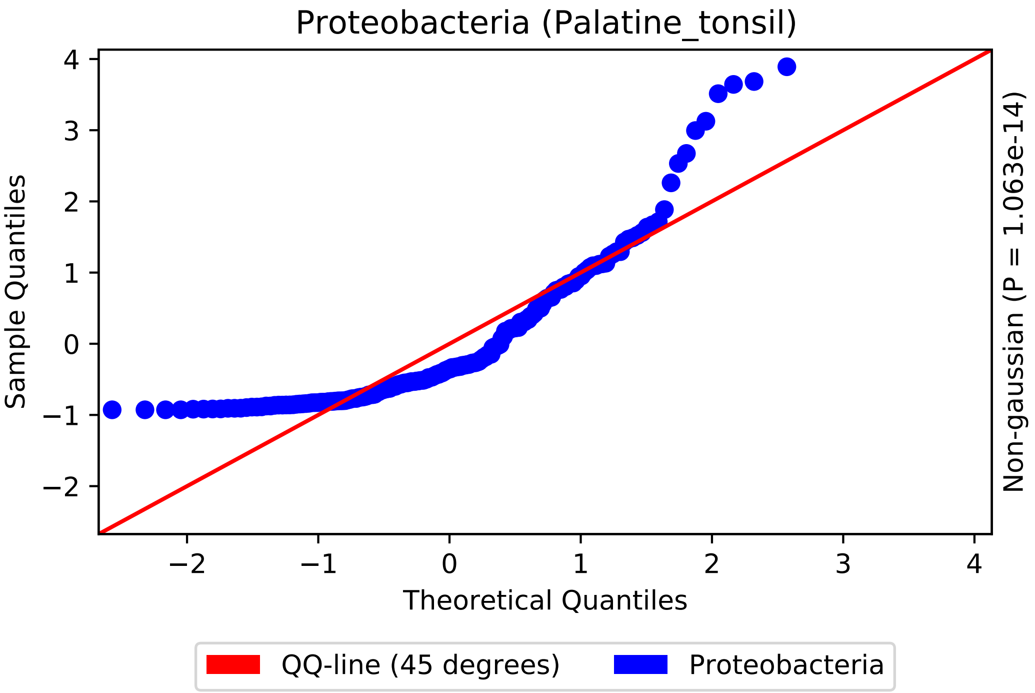

Proteobacteria (Palatine_tonsil)
Sample Quantiles
Theoretical Quantiles
Non-gaussian (P = 1.063e-14)
4
3
2
1
0
−1
−2
−2
−1
0
1
2
3
4
QQ-line (45 degrees)
Proteobacteria

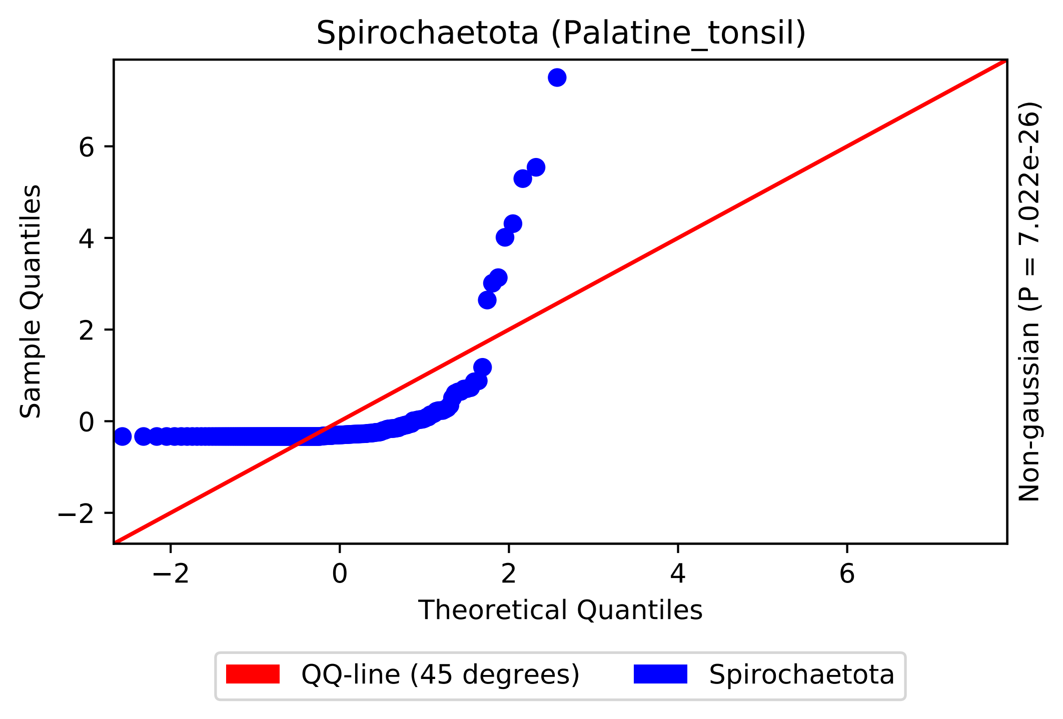

Spirochaetota (Palatine_tonsil)
Sample Quantiles
Theoretical Quantiles
Non-gaussian (P = 7.022e-26)
−2
0
2
4
6
QQ-line (45 degrees)
Spirochaetota

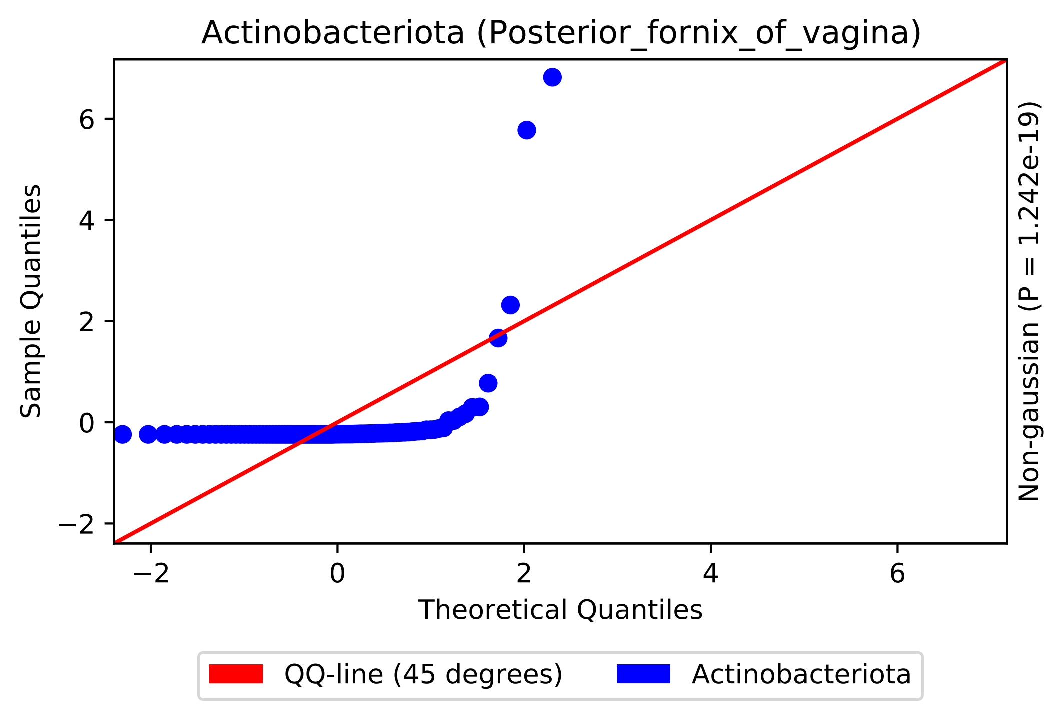

Actinobacteriota (Posterior_fornix_of_vagina)
Sample Quantiles
Theoretical Quantiles
Non-gaussian (P = 1.242e-19)
−2
0
2
4
6
QQ-line (45 degrees)
Actinobacteriota

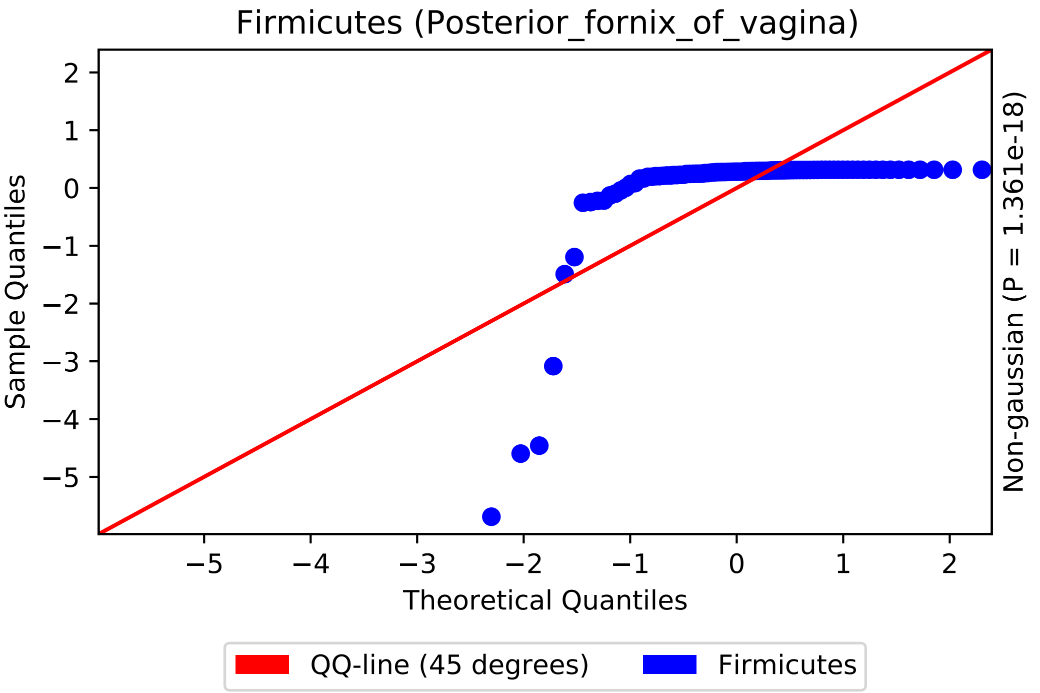
Firmicutes (Posterior_fornix_of_vagina)
Sample Quantiles
Theoretical Quantiles
Non-gaussian (P = 1.361e-18)
QQ-line (45 degrees)
Firmicutes
2
1
0
−1
−2
−3
−4
−5
−5
−4
−3
−2
−1
0
1
2

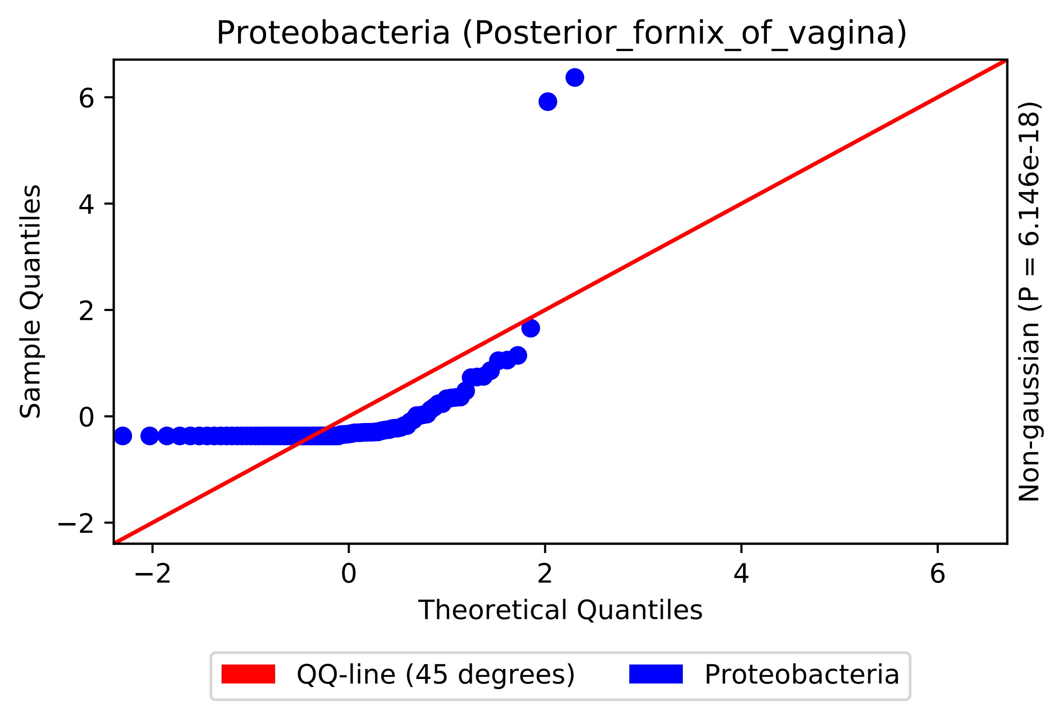

Proteobacteria (Posterior_fornix_of_vagina)
Sample Quantiles
Theoretical Quantiles
Non-gaussian (P = 6.146e-18)
−2
0
2
4
6
QQ-line (45 degrees)
Proteobacteria

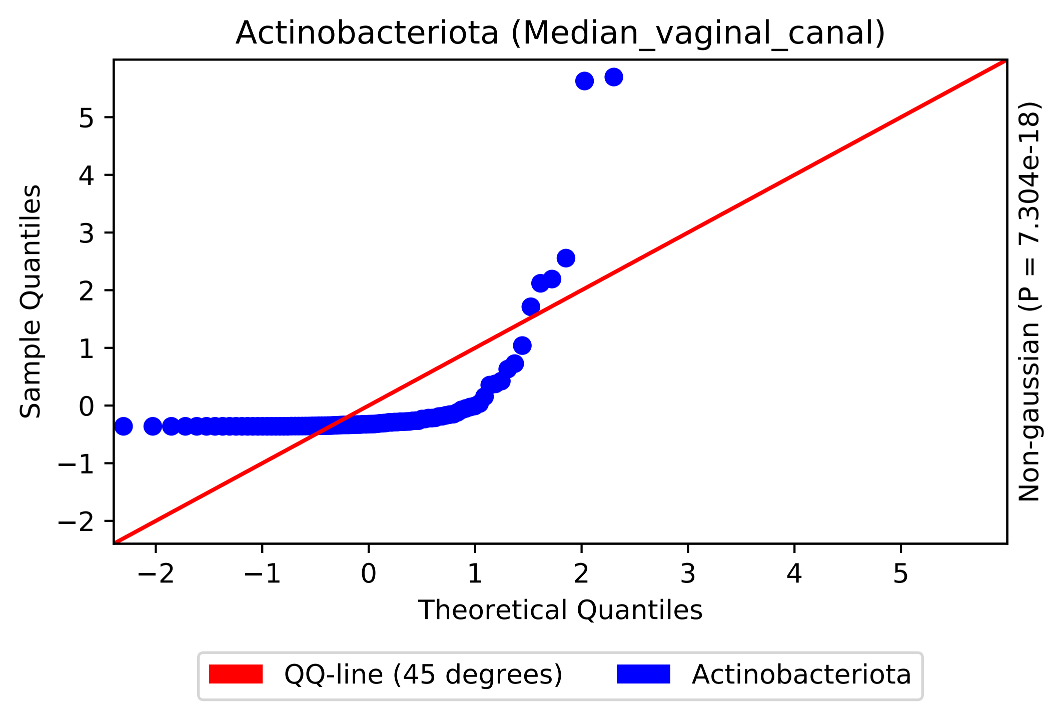
Actinobacteriota (Median_vaginal_canal)
Sample Quantiles
Theoretical Quantiles
Non-gaussian (P = 7.304e-18)
5
4
3
2
1
0
−1
−2
−2
−1
0
1
2
3
4
5
QQ-line (45 degrees)
Actinobacteriota

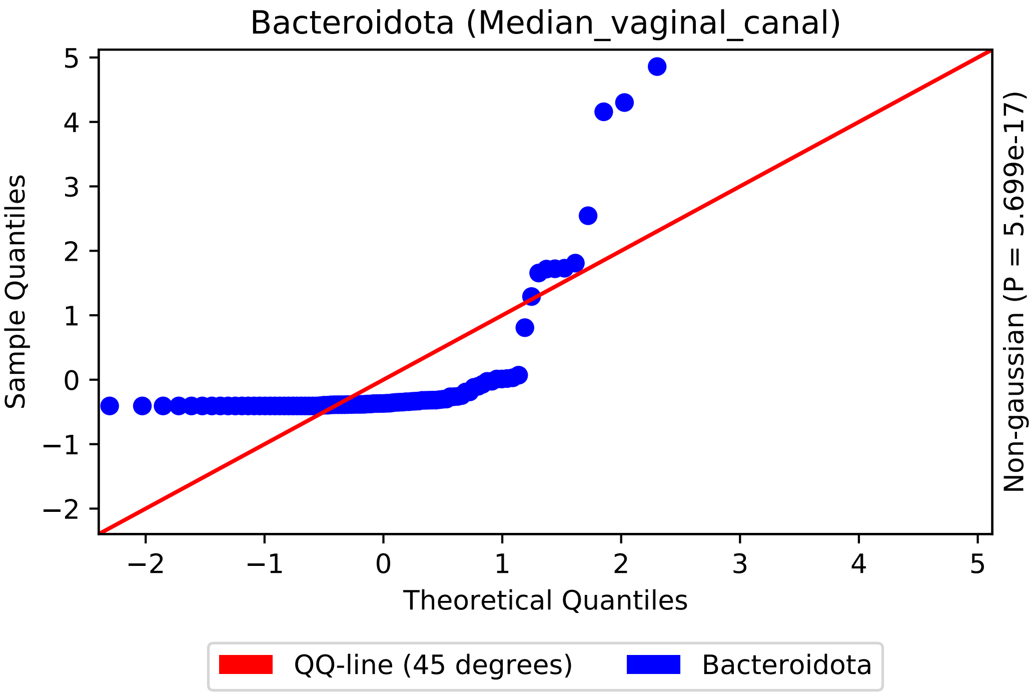

Bacteroidota (Median_vaginal_canal)
Sample Quantiles
Theoretical Quantiles
Non-gaussian (P = 5.699e-17)
−2
−1
0
1
2
3
4
5
QQ-line (45 degrees)
Bacteroidota

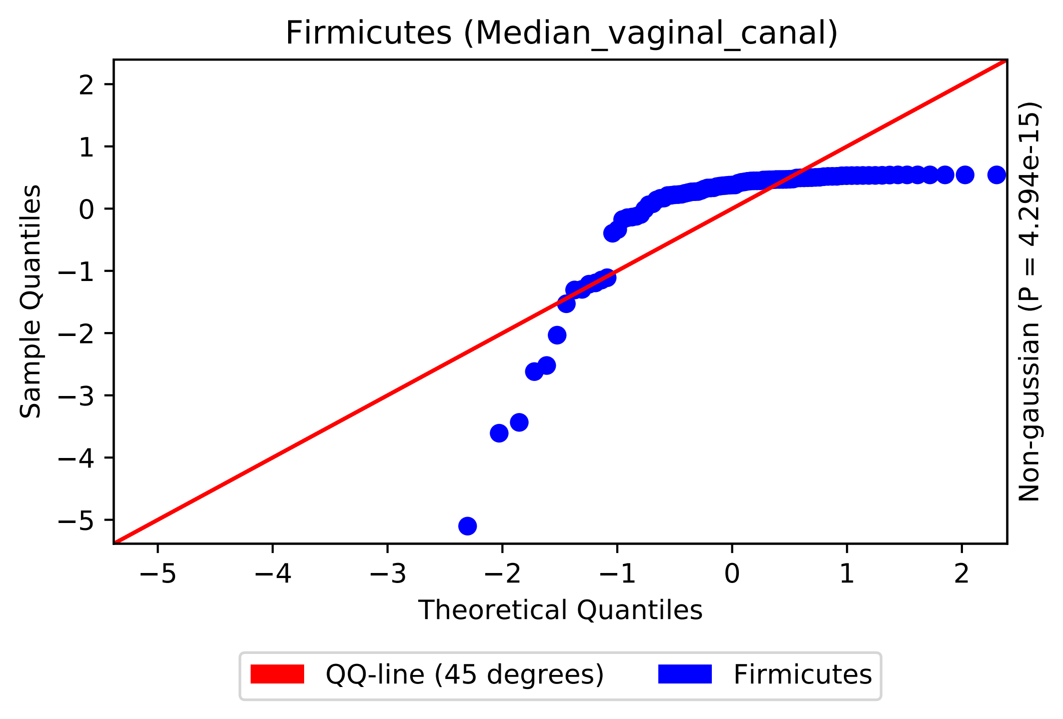

Firmicutes (Median_vaginal_canal)
Sample Quantiles
Theoretical Quantiles
Non-gaussian (P = 4.294e-15)
QQ-line (45 degrees)
Firmicutes

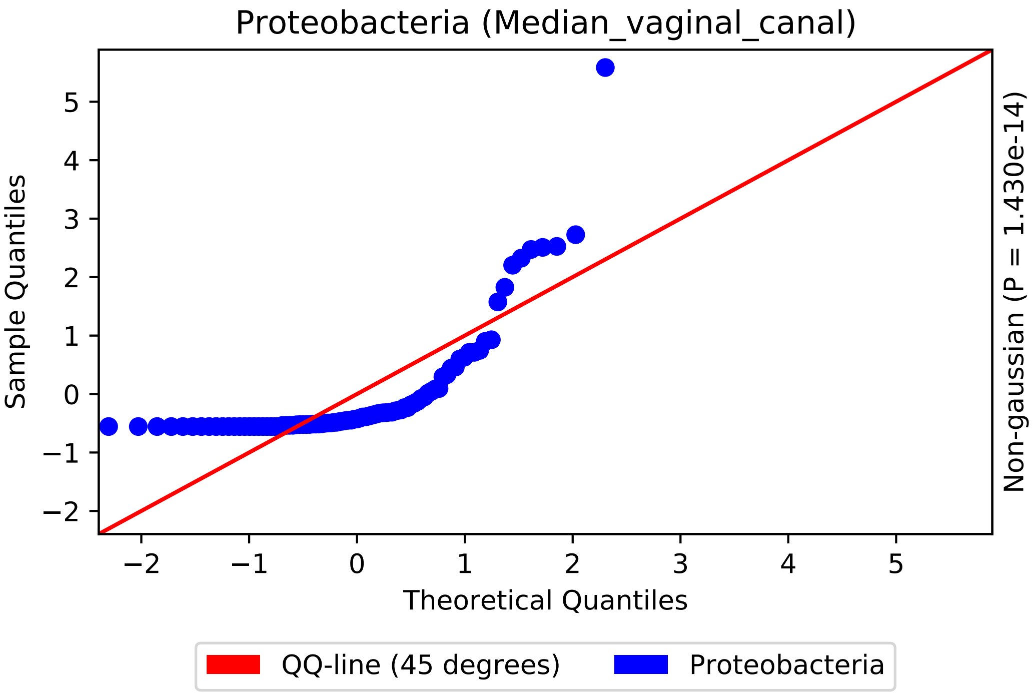

Proteobacteria (Median_vaginal_canal)
Sample Quantiles
Theoretical Quantiles
Non-gaussian (P = 1.430e-14)
5
4
3
2
1
0
−1
−2
−2
−1
0
1
2
3
4
5
QQ-line (45 degrees)
Proteobacteria

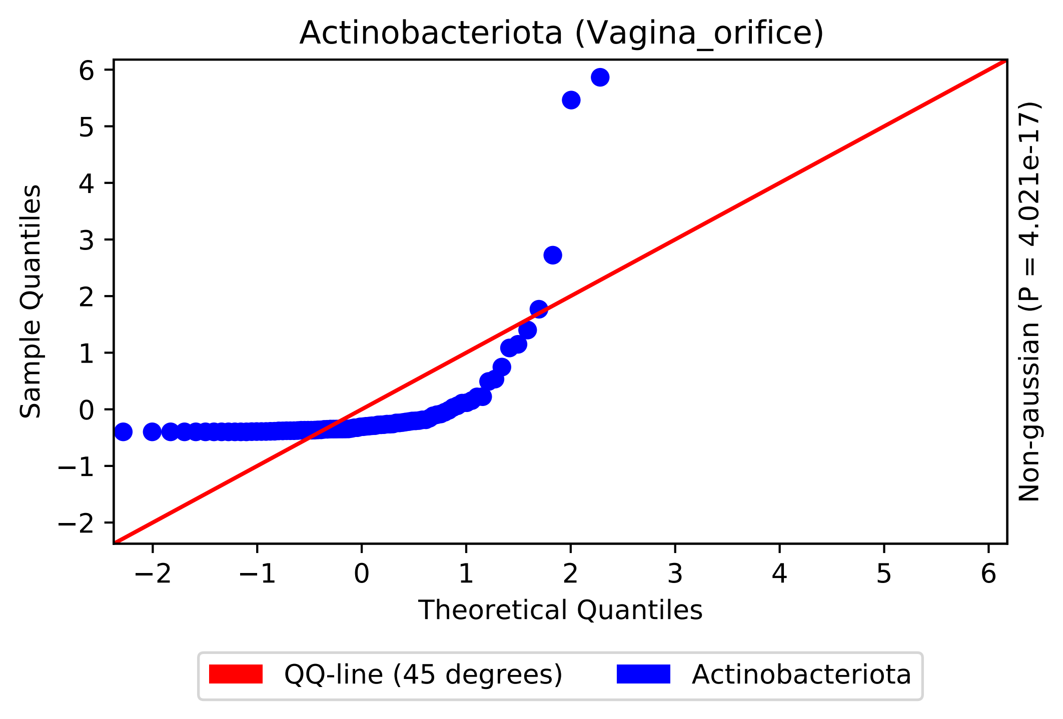

Actinobacteriota (Vagina_orifice)
Sample Quantiles
Theoretical Quantiles
Non-gaussian (P = 4.021e-17)
6
5
4
3
2
1
0
−1
−2
−2
−1
0
1
2
3
4
5
6
QQ-line (45 degrees)
Actinobacteriota

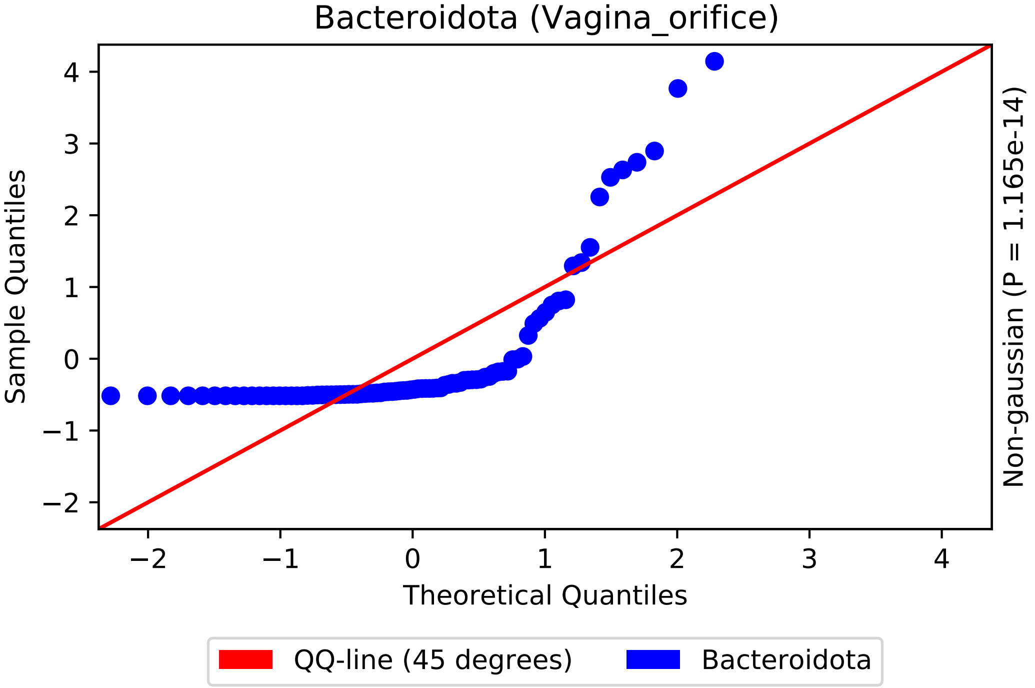

Bacteroidota (Vagina_orifice)
Sample Quantiles
Theoretical Quantiles
Non-gaussian (P = 1.165e-14)
QQ-line (45 degrees)
Bacteroidota

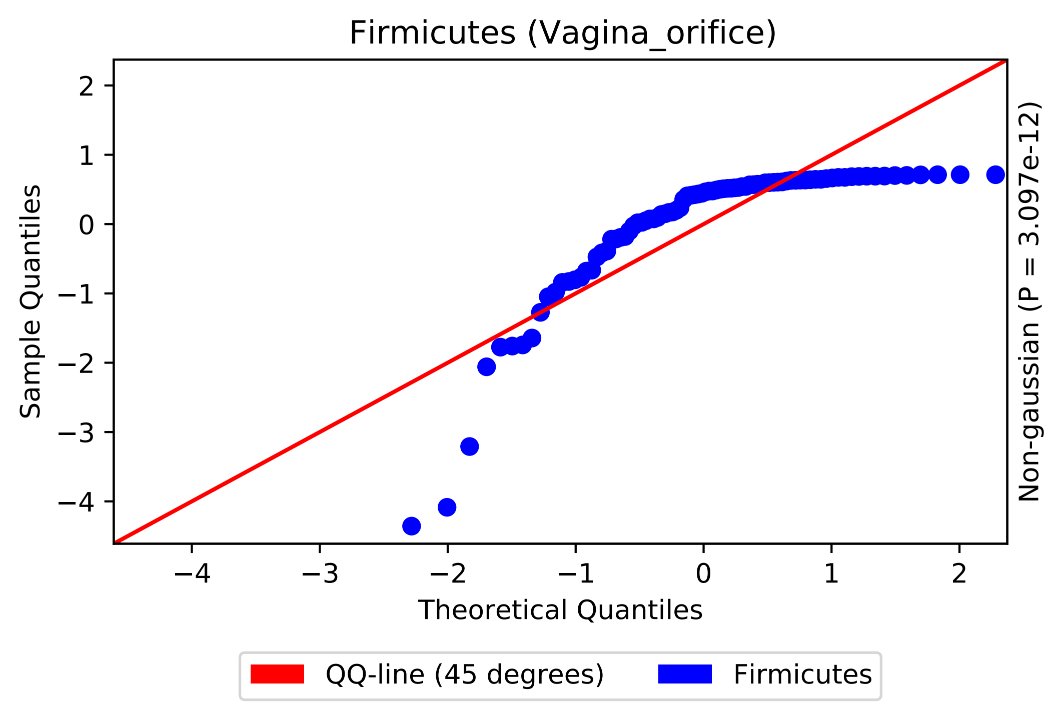

Firmicutes (Vagina_orifice)
Sample Quantiles
Theoretical Quantiles
Non-gaussian (P = 3.097e-12)
2
1
0
−1
−2
−3
−4
−4
−3
−2
−1
0
1
2
QQ-line (45 degrees)
Firmicutes

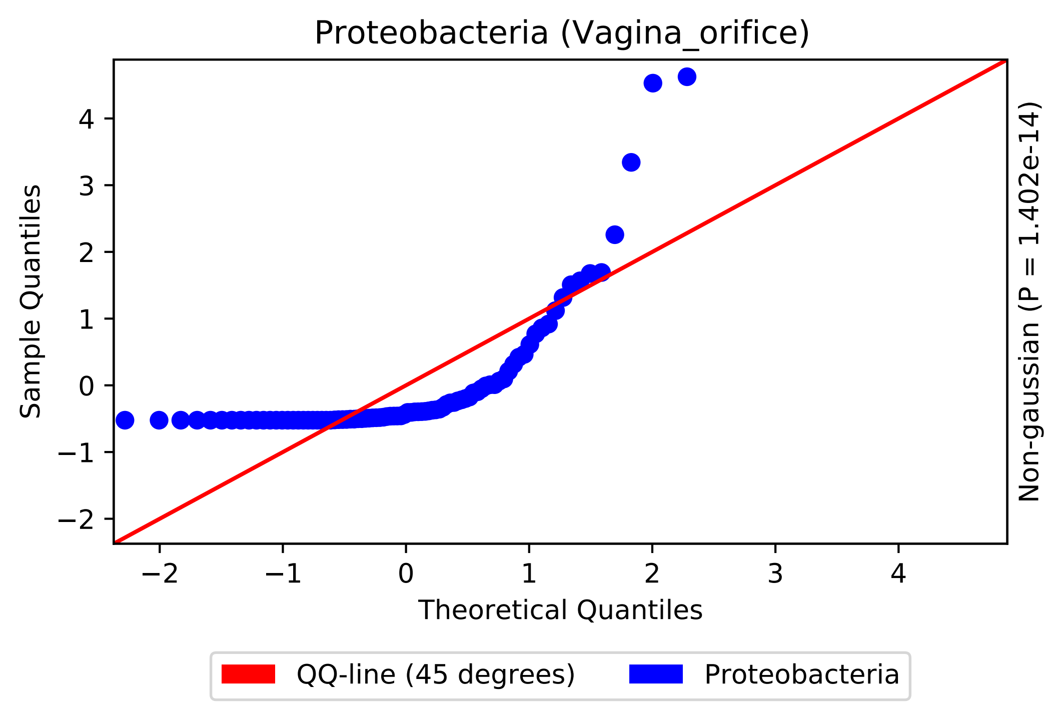
Proteobacteria (Vagina_orifice)
Sample Quantiles
Theoretical Quantiles
Non-gaussian (P = 1.402e-14)
−2
−1
0
1
2
3
4
QQ-line (45 degrees)
Proteobacteria

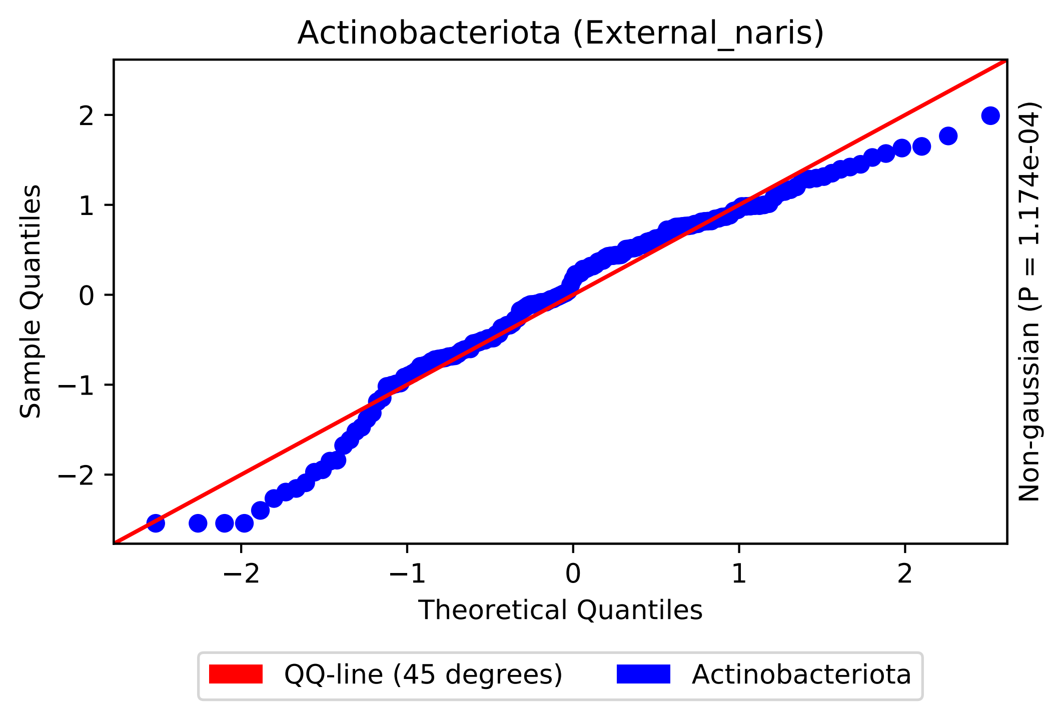

Actinobacteriota (External_naris)
Sample Quantiles
Theoretical Quantiles
Non-gaussian (P = 1.174e-04)
2
1
0
−1
−2
QQ-line (45 degrees)
Actinobacteriota

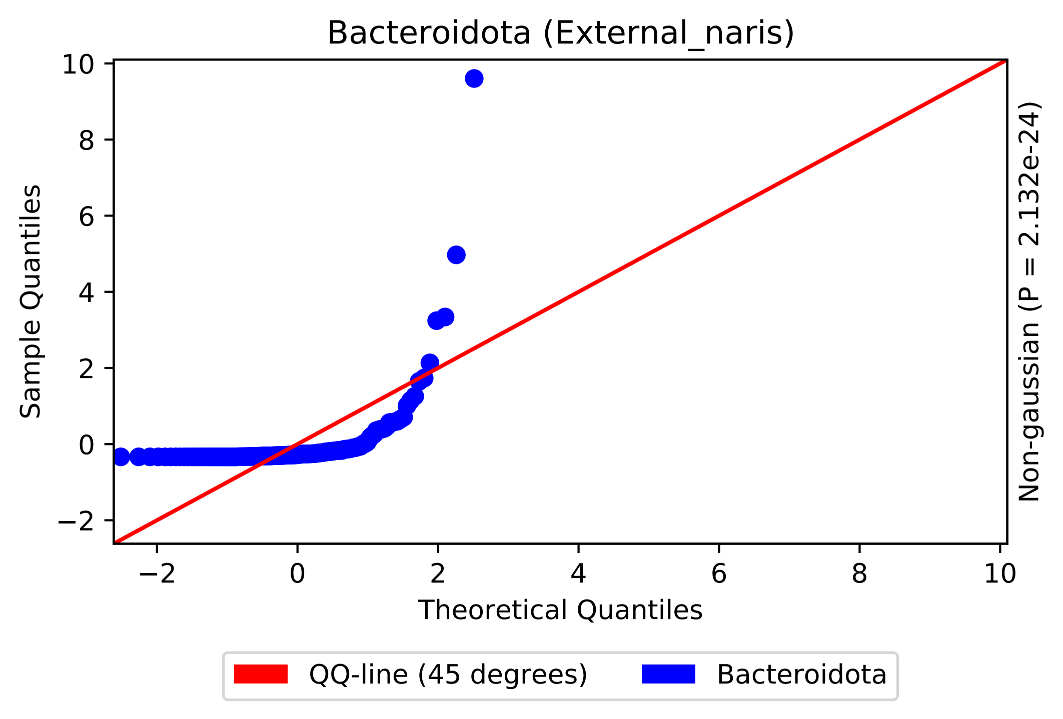
Bacteroidota (External_naris)
Sample Quantiles
Theoretical Quantiles
Non-gaussian (P = 2.132e-24)
−2
0
2
4
6
8
10
QQ-line (45 degrees)
Bacteroidota

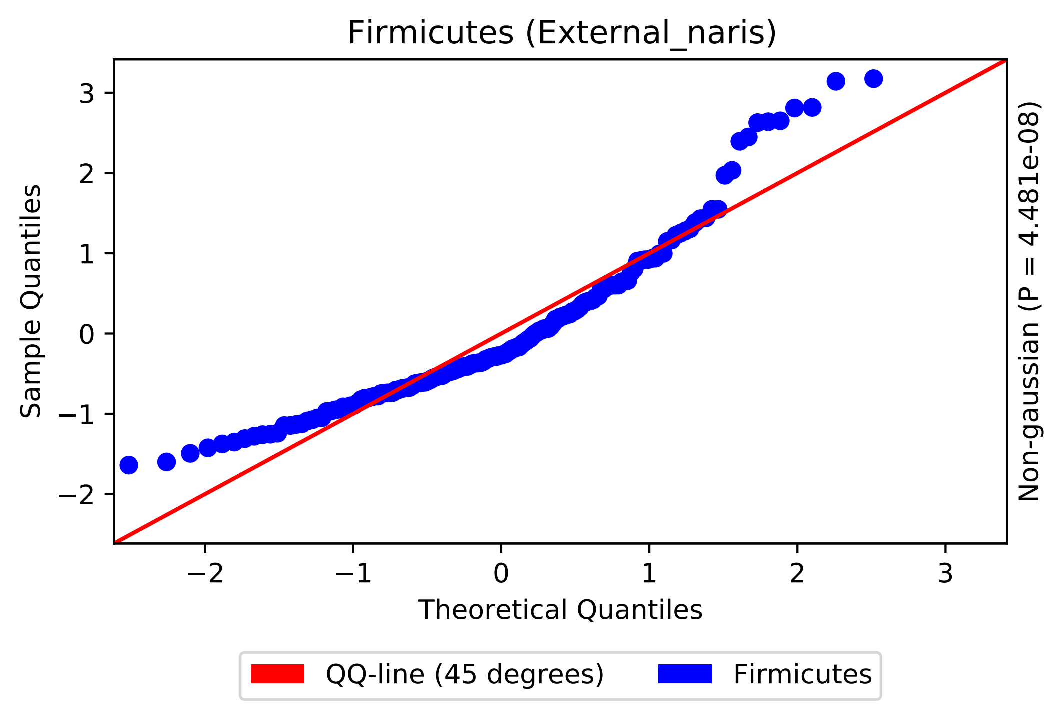
Firmicutes (External_naris)
Sample Quantiles
Theoretical Quantiles
Non-gaussian (P = 4.481e-08)
QQ-line (45 degrees)
Firmicutes
−2
−1
0
1
2
3

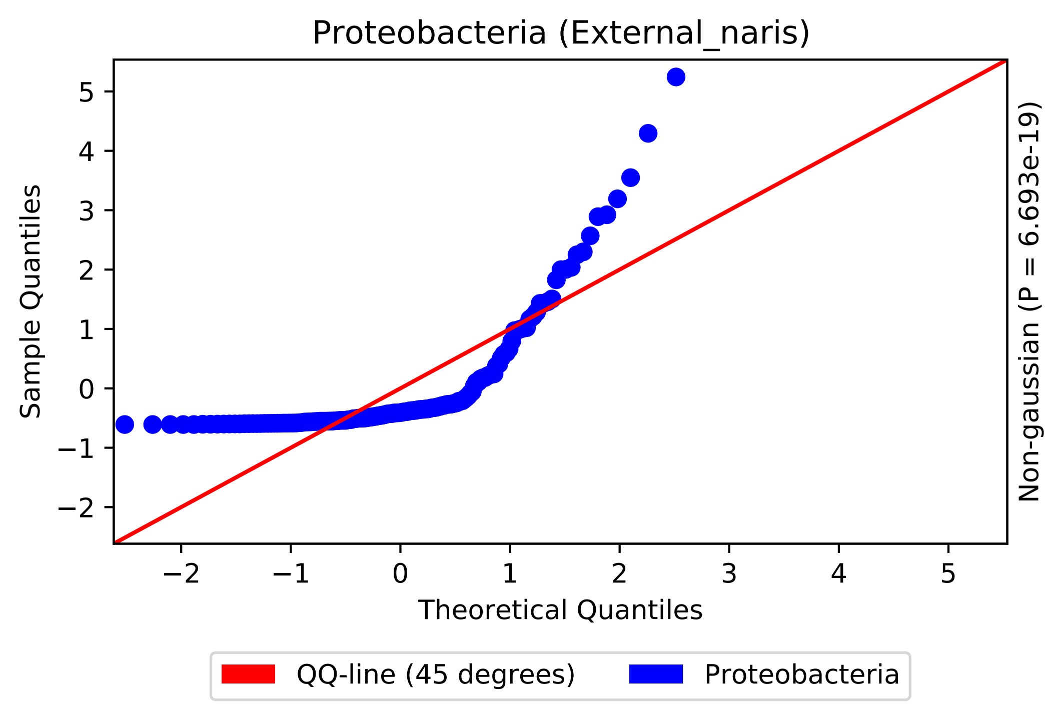

Proteobacteria (External_naris)
Sample Quantiles
Theoretical Quantiles
Non-gaussian (P = 6.693e-19)
5
4
3
2
1
0
−1
−2
−2
−1
0
1
2
3
4
5
QQ-line (45 degrees)
Proteobacteria

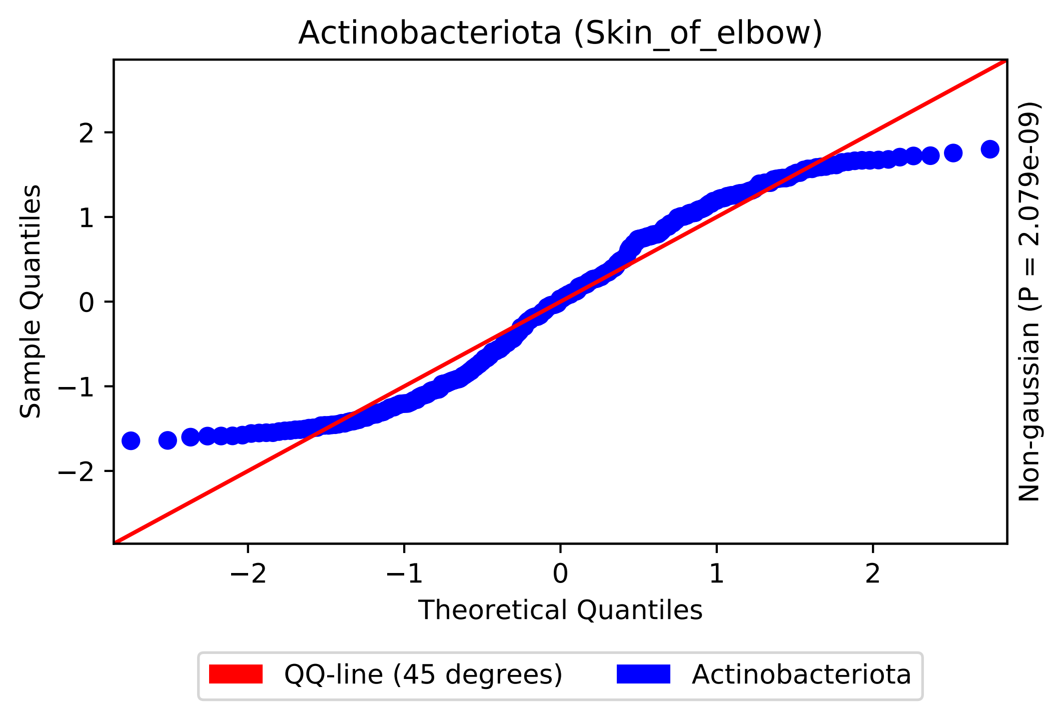

Actinobacteriota (Skin_of_elbow)
Sample Quantiles
Theoretical Quantiles
Non-gaussian (P = 2.079e-09)
2
1
0
−1
−2
−2
−1
0
1
2
QQ-line (45 degrees)
Actinobacteriota

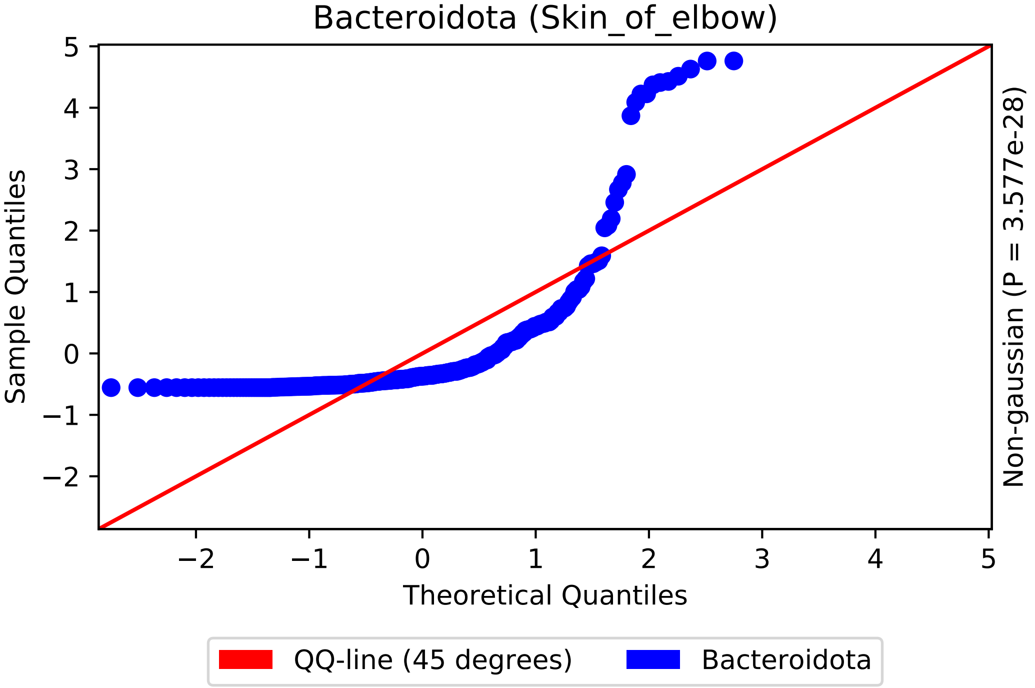

Bacteroidota (Skin_of_elbow)
Sample Quantiles
Theoretical Quantiles
Non-gaussian (P = 3.577e-28)
5
4
3
2
1
0
−1
−2
−2
−1
0
1
2
3
4
5
QQ-line (45 degrees)
Bacteroidota

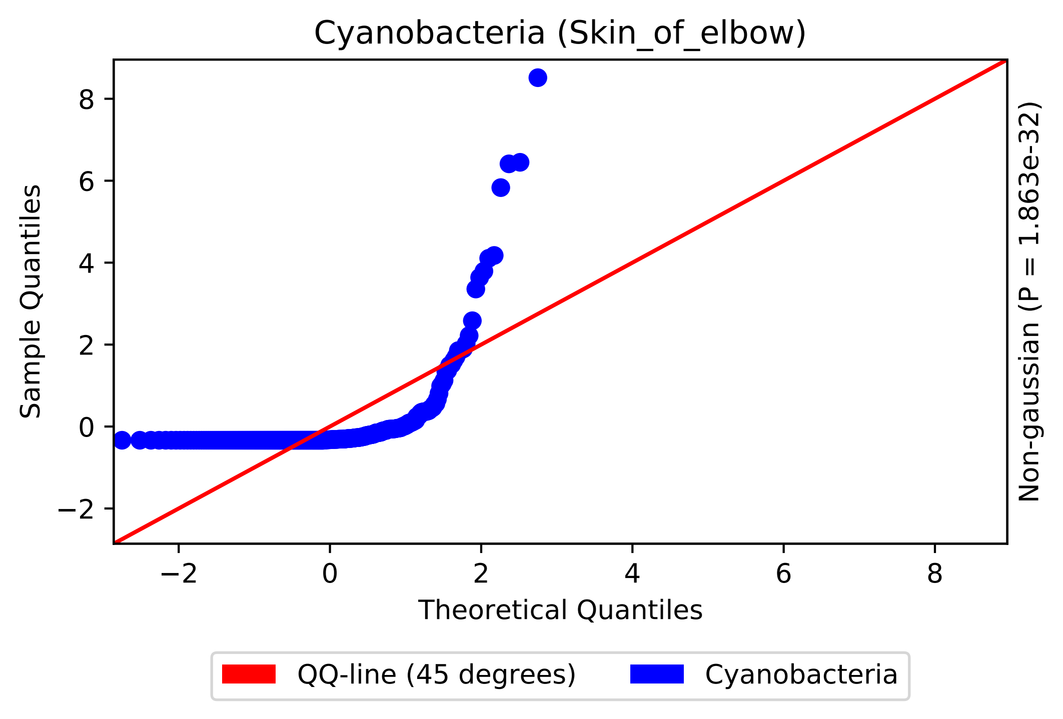

Cyanobacteria (Skin_of_elbow)
Sample Quantiles
Theoretical Quantiles
Non-gaussian (P = 1.863e-32)
−2
0
2
4
6
8
QQ-line (45 degrees)
Cyanobacteria

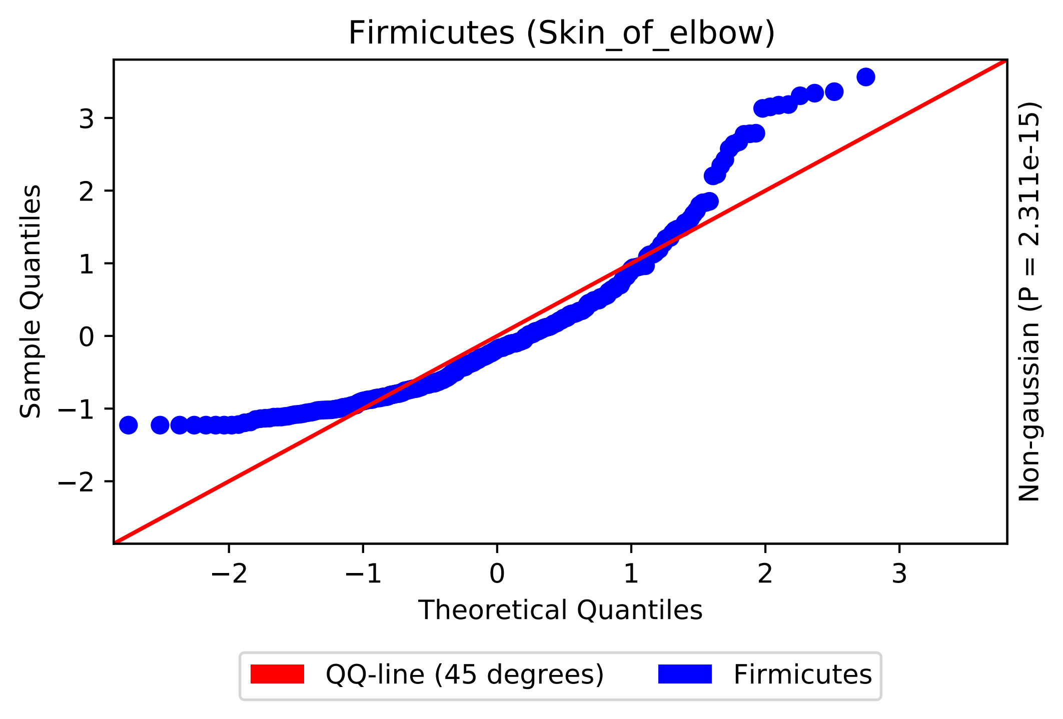
Firmicutes (Skin_of_elbow)
Sample Quantiles
Theoretical Quantiles
Non-gaussian (P = 2.311e-15)
3
2
1
0
−1
−2
−2
−1
0
1
2
3
QQ-line (45 degrees)
Firmicutes

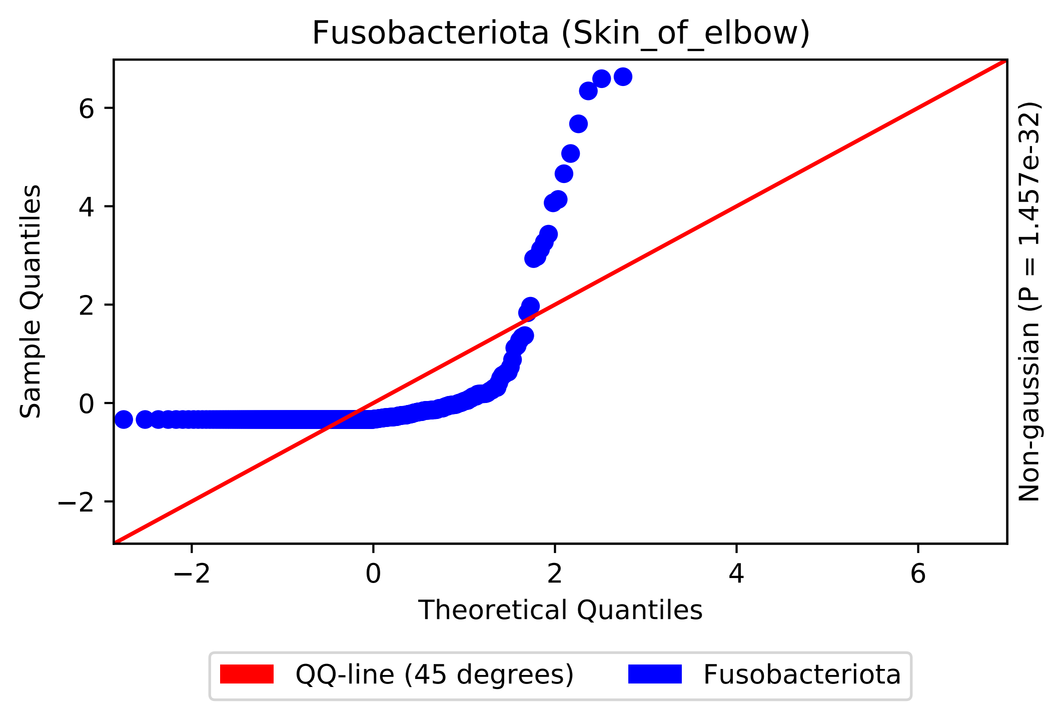

Fusobacteriota (Skin_of_elbow)
Sample Quantiles
Theoretical Quantiles
Non-gaussian (P = 1.457e-32)
−2
0
2
4
6
QQ-line (45 degrees)
Fusobacteriota

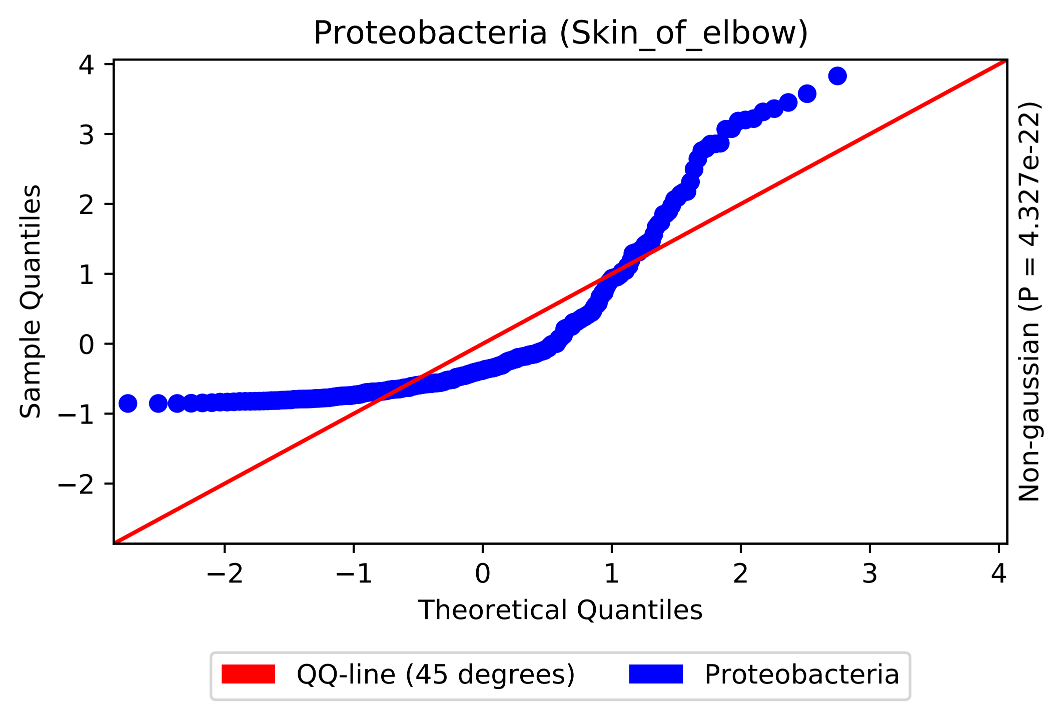
Proteobacteria (Skin_of_elbow)
Sample Quantiles
Theoretical Quantiles
Non-gaussian (P = 4.327e-22)
QQ-line (45 degrees)
Proteobacteria
4
3
2
1
0
−1
−2
−2
−1
0
1
2
3
4

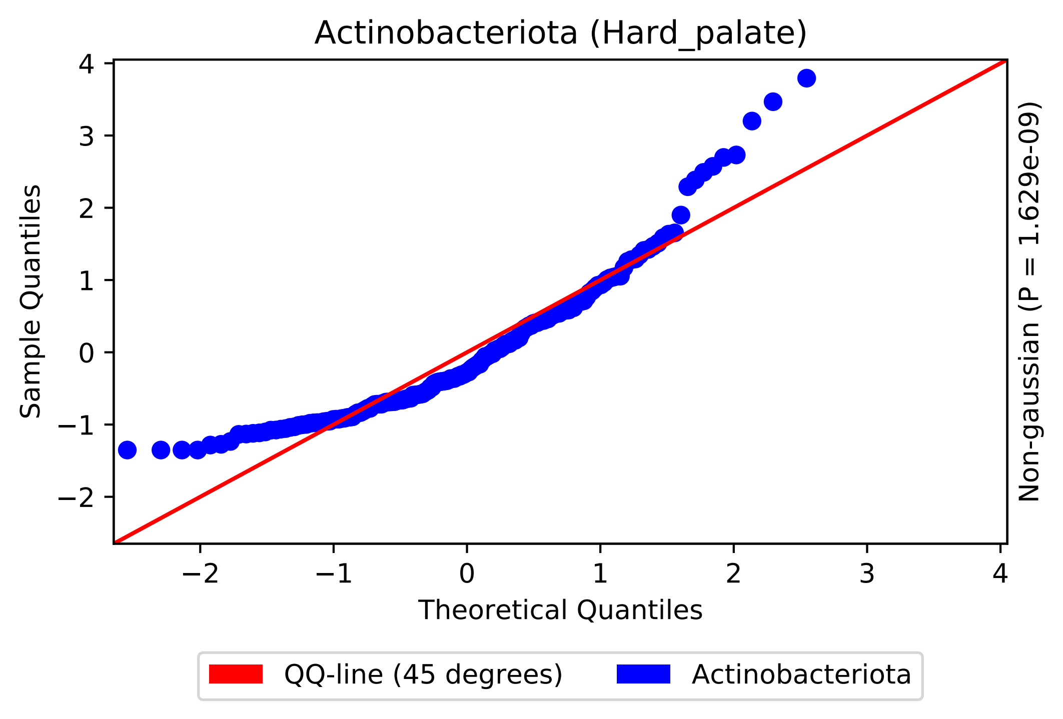

Actinobacteriota (Hard_palate)
Sample Quantiles
Theoretical Quantiles
Non-gaussian (P = 1.629e-09)
4
3
2
1
0
−1
−2
−2
−1
0
1
2
3
4
QQ-line (45 degrees)
Actinobacteriota

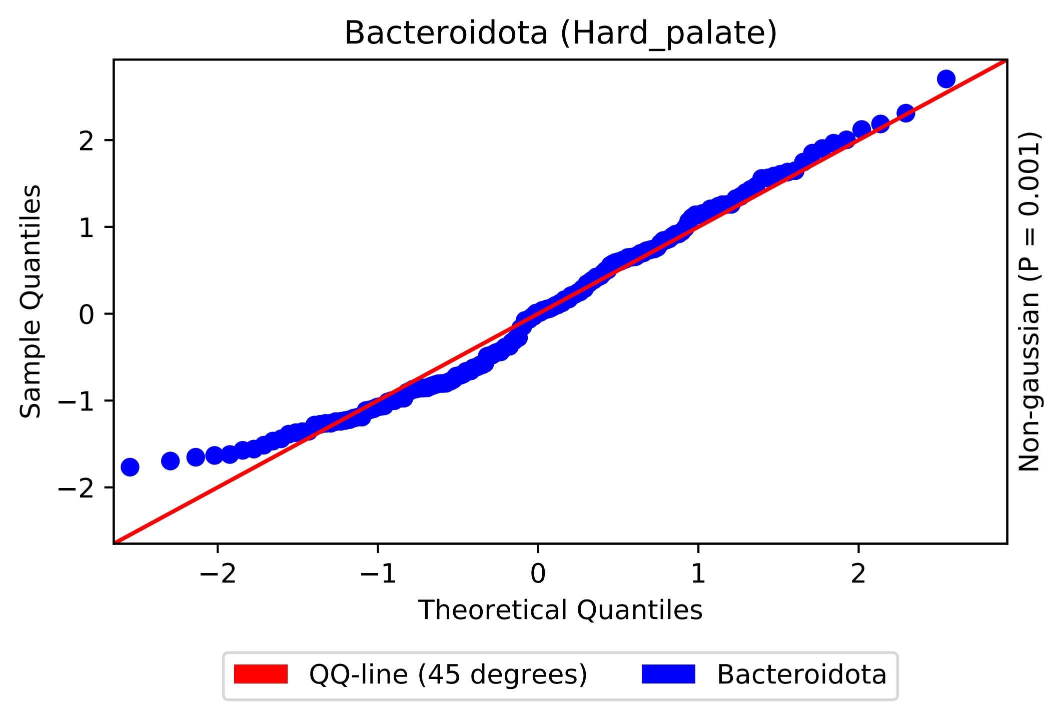
Bacteroidota (Hard_palate)
Sample Quantiles
Theoretical Quantiles
Non-gaussian (P = 0.001)
QQ-line (45 degrees)
Bacteroidota
−2
−1
0
1
2

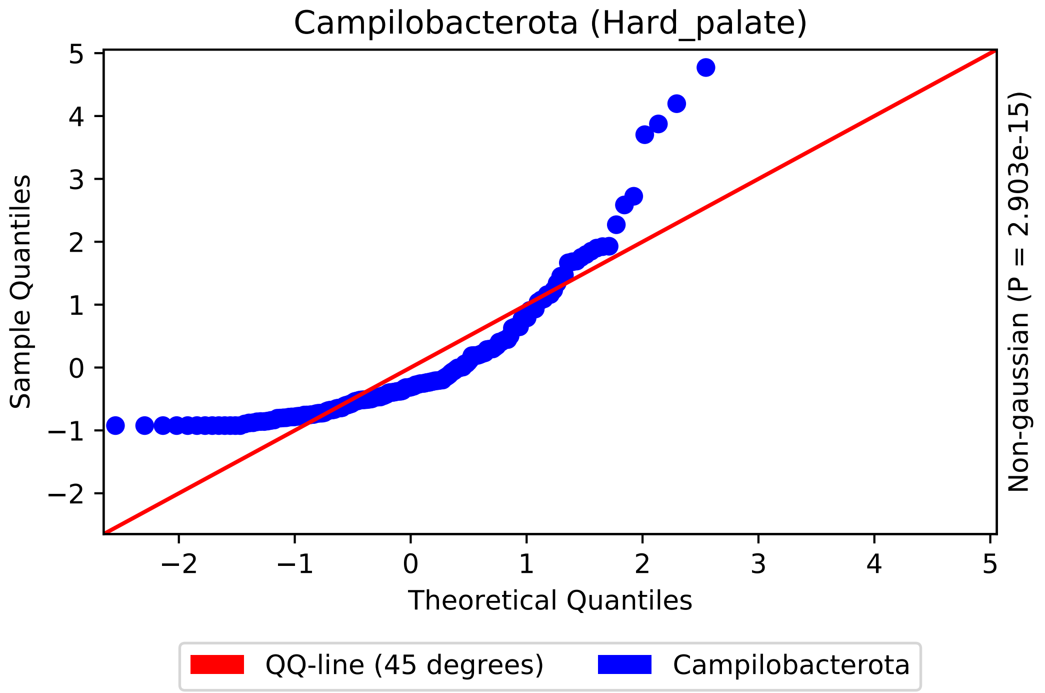

Campilobacterota (Hard_palate)
Sample Quantiles
Theoretical Quantiles
Non-gaussian (P = 2.903e-15)
5
4
3
2
1
0
−1
−2
−2
−1
0
1
2
3
4
5
QQ-line (45 degrees)
Campilobacterota

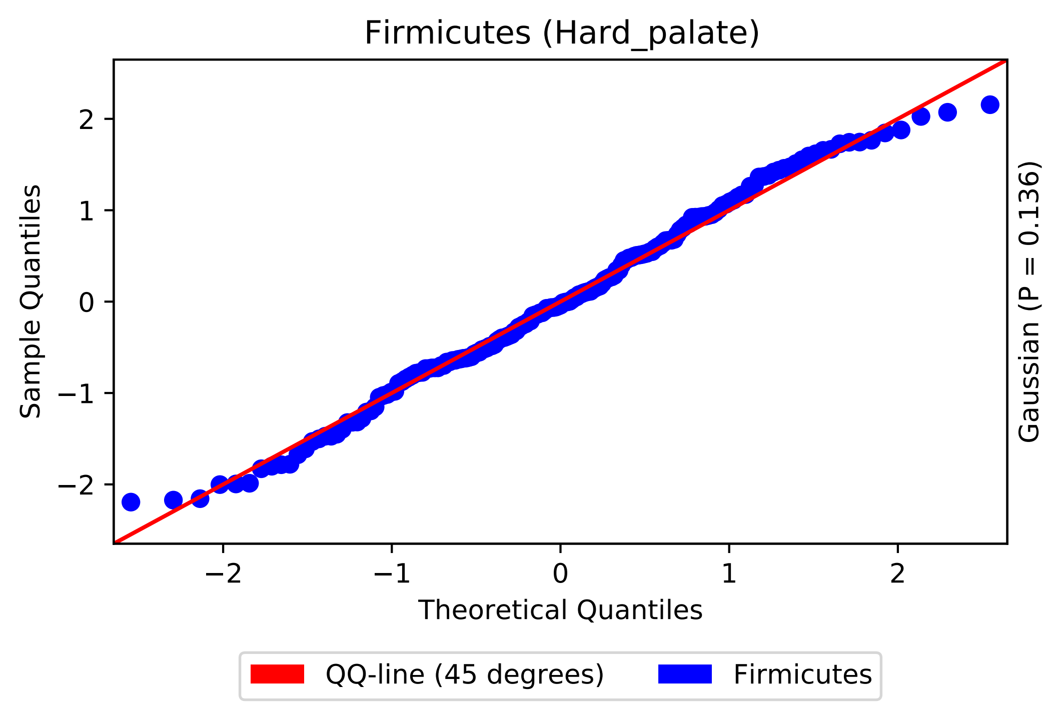

Firmicutes (Hard_palate)
Sample Quantiles
Theoretical Quantiles
Gaussian (P = 0.136)
−2
−1
0
1
2
QQ-line (45 degrees)
Firmicutes

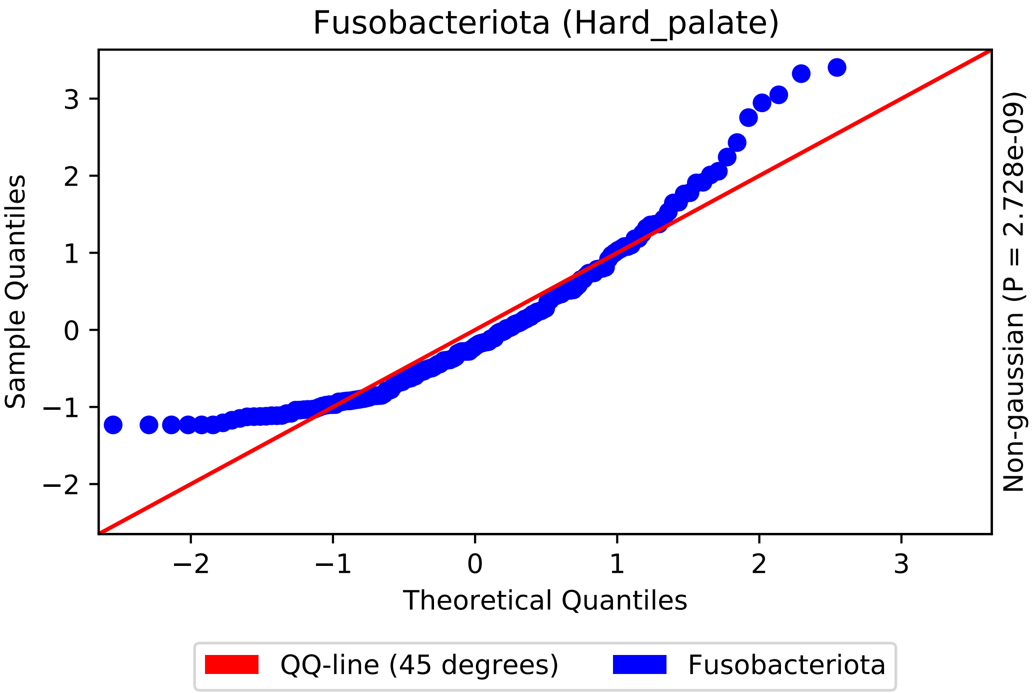

Fusobacteriota (Hard_palate)
Sample Quantiles
Theoretical Quantiles
Non-gaussian (P = 2.728e-09)
−2
−1
0
1
2
3
QQ-line (45 degrees)
Fusobacteriota

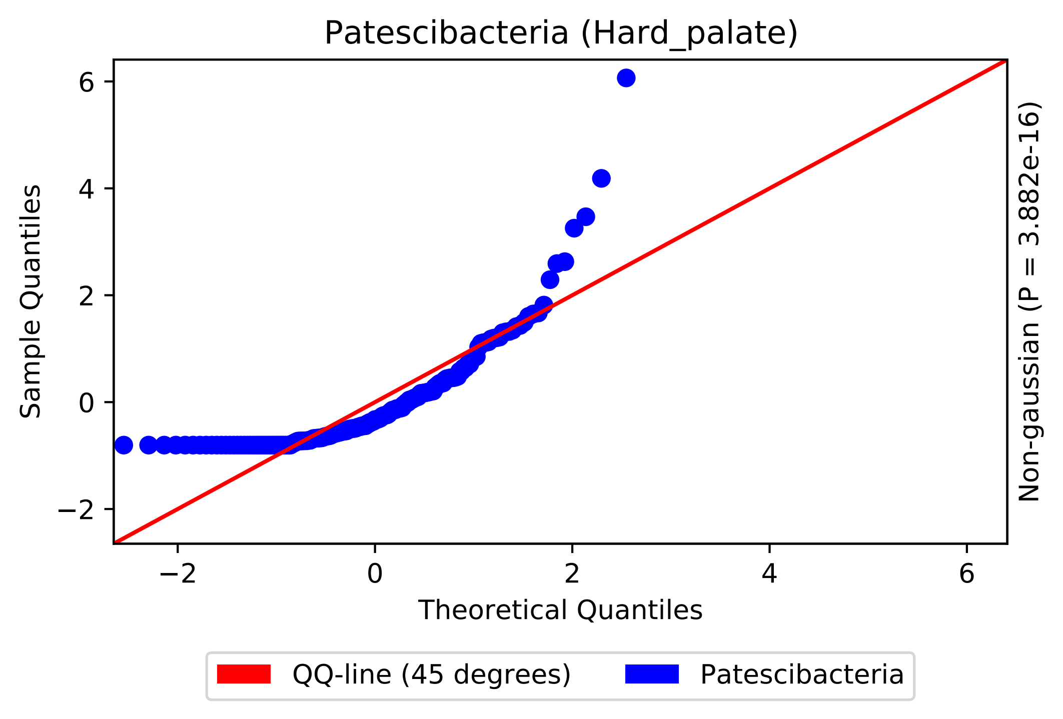

Patescibacteria (Hard_palate)
Sample Quantiles
Theoretical Quantiles
Non-gaussian (P = 3.882e-16)
−2
0
2
4
6
QQ-line (45 degrees)
Patescibacteria

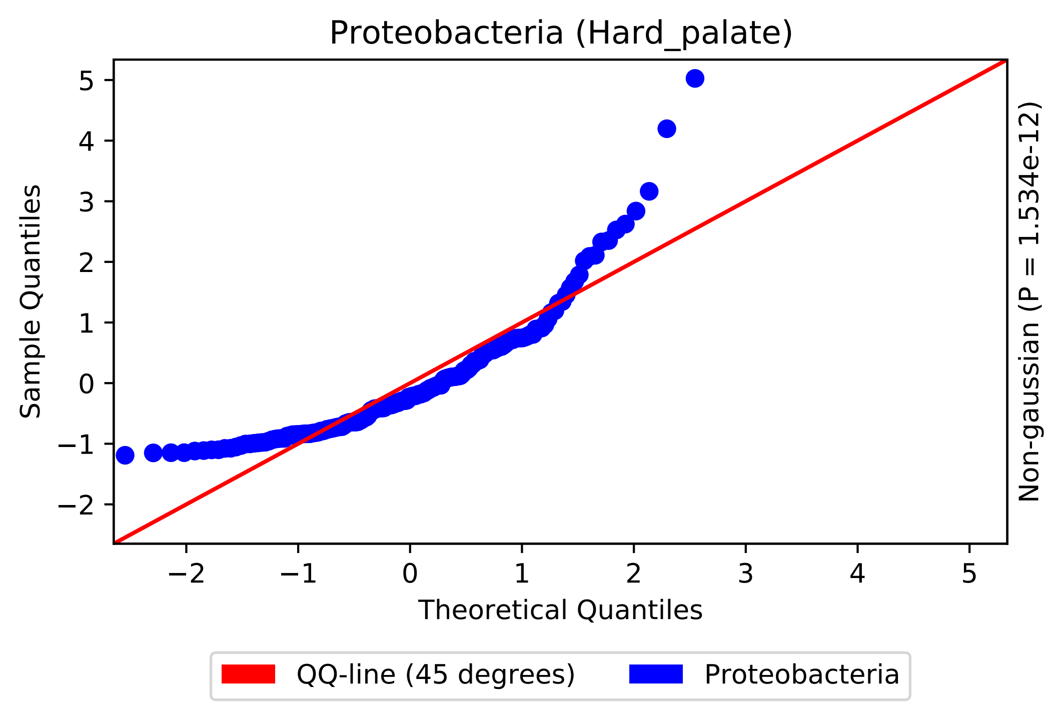

Proteobacteria (Hard_palate)
Sample Quantiles
Theoretical Quantiles
Non-gaussian (P = 1.534e-12)
QQ-line (45 degrees)
Proteobacteria

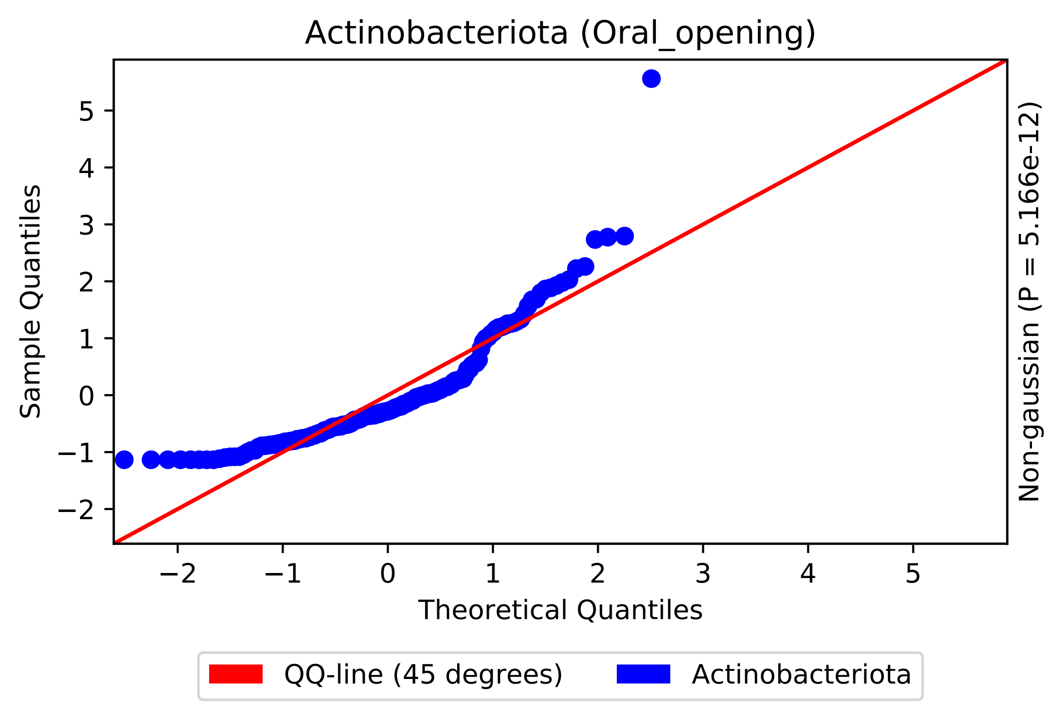

Actinobacteriota (Oral_opening)
Sample Quantiles
Theoretical Quantiles
Non-gaussian (P = 5.166e-12)
−2
−1
0
1
2
3
4
5
QQ-line (45 degrees)
Actinobacteriota

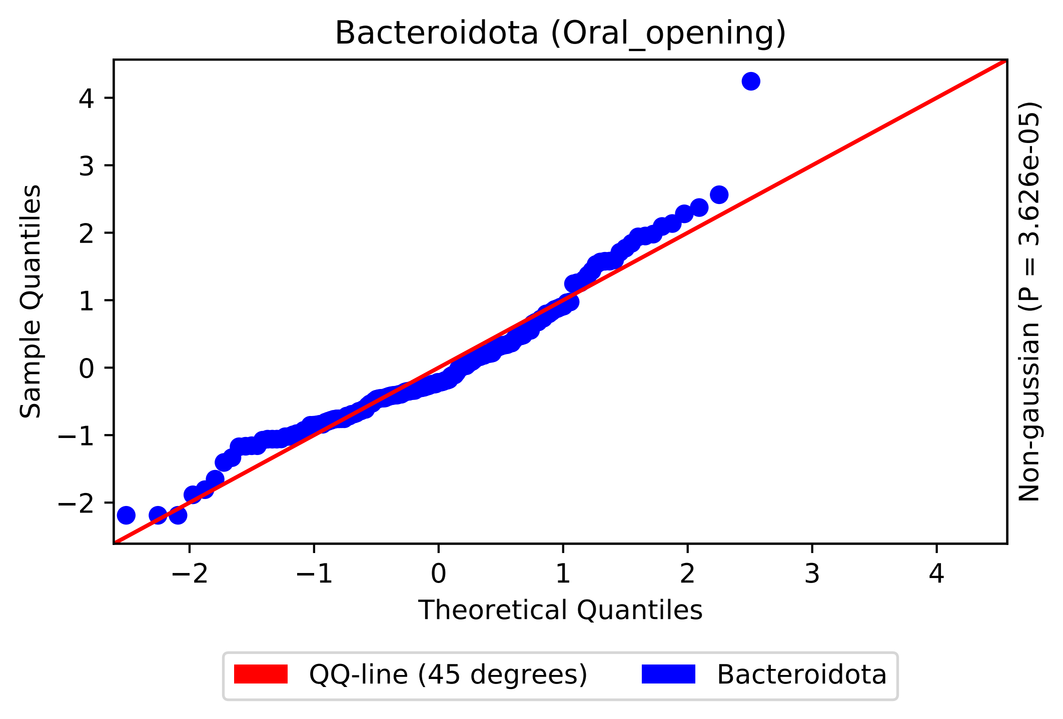
Bacteroidota (Oral_opening)
Sample Quantiles
Theoretical Quantiles
Non-gaussian (P = 3.626e-05)
4
3
2
1
0
−1
−2
−2
−1
0
1
2
3
4
QQ-line (45 degrees)
Bacteroidota

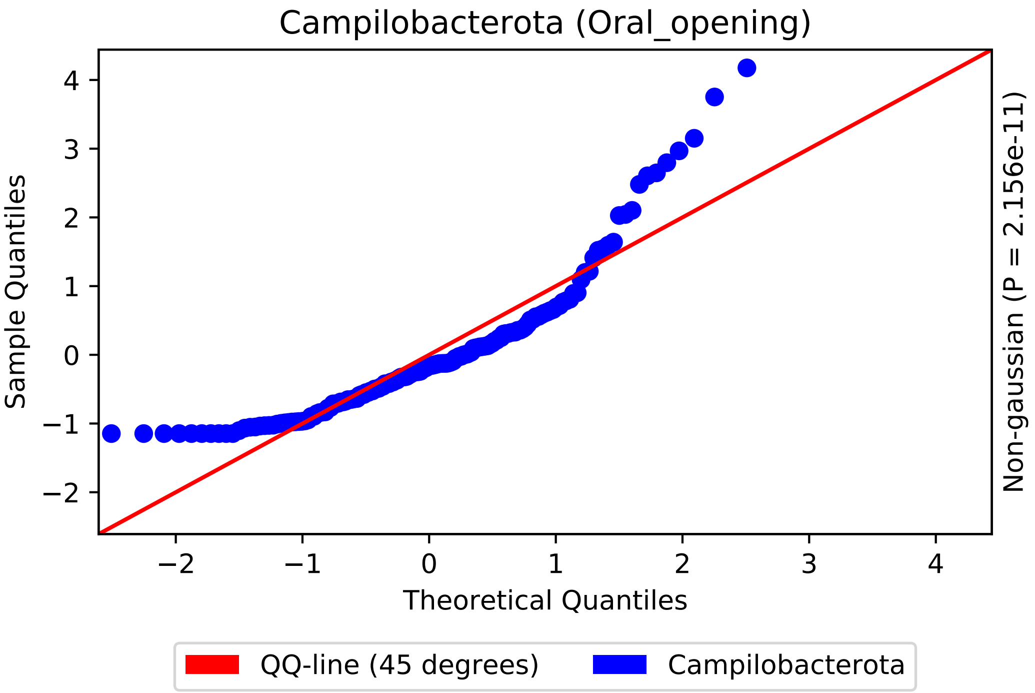

Campilobacterota (Oral_opening)
Sample Quantiles
Theoretical Quantiles
Non-gaussian (P = 2.156e-11)
4
3
2
1
0
−1
−2
−2
−1
0
1
2
3
4
QQ-line (45 degrees)
Campilobacterota

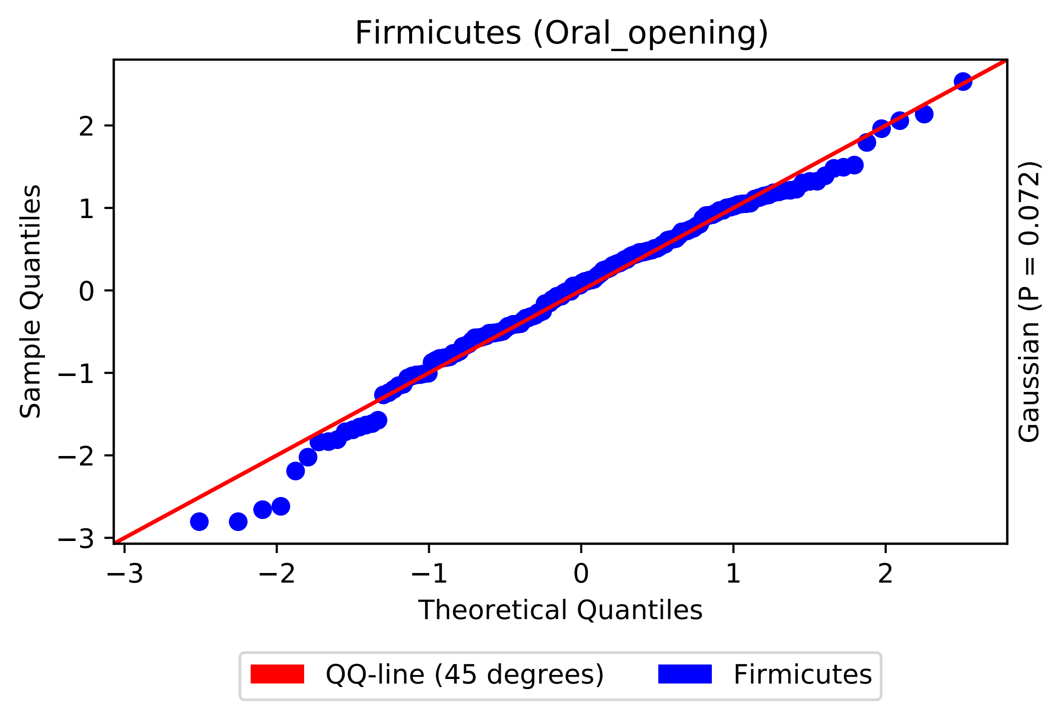
Firmicutes (Oral_opening)
Sample Quantiles
Theoretical Quantiles
Gaussian (P = 0.072)
−3
−2
−1
0
1
2
QQ-line (45 degrees)
Firmicutes

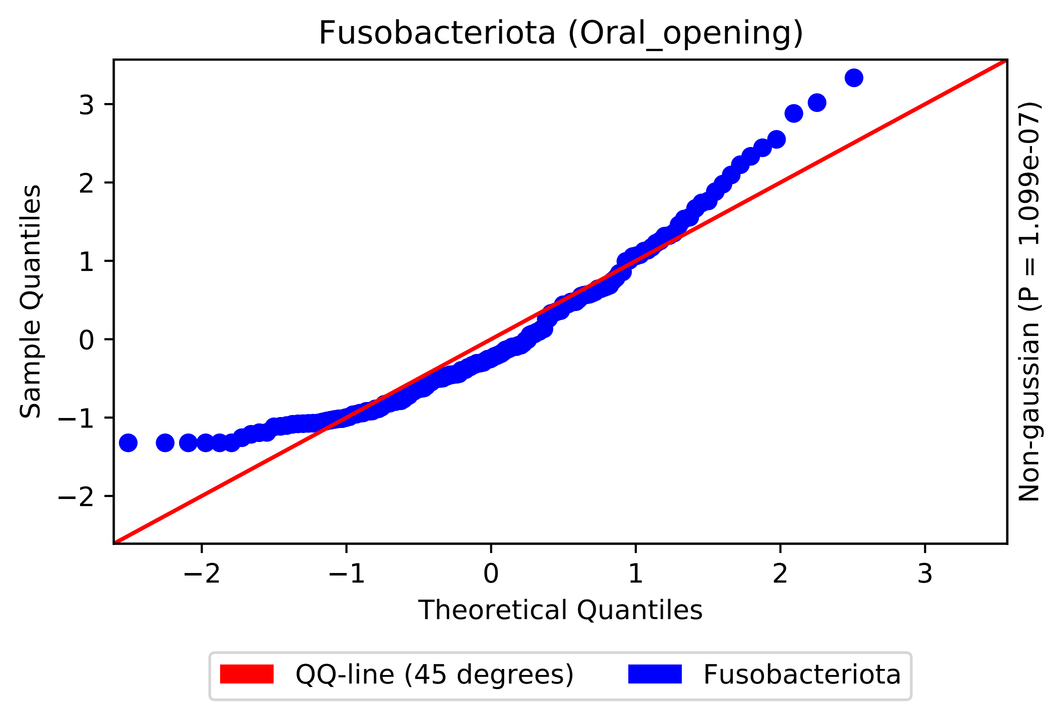

Fusobacteriota (Oral_opening)
Sample Quantiles
Theoretical Quantiles
Non-gaussian (P = 1.099e-07)
−2
−1
0
1
2
3
QQ-line (45 degrees)
Fusobacteriota

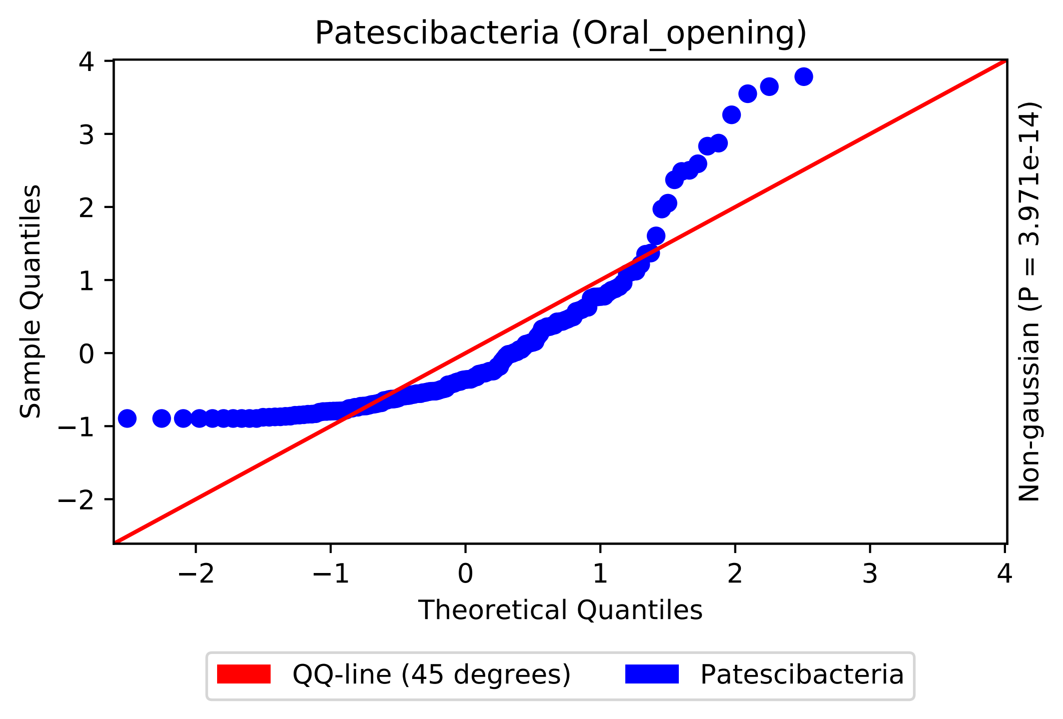
Patescibacteria (Oral_opening)
Sample Quantiles
Theoretical Quantiles
Non-gaussian (P = 3.971e-14)
4
3
2
1
0
−1
−2
−2
−1
0
1
2
3
4
QQ-line (45 degrees)
Patescibacteria

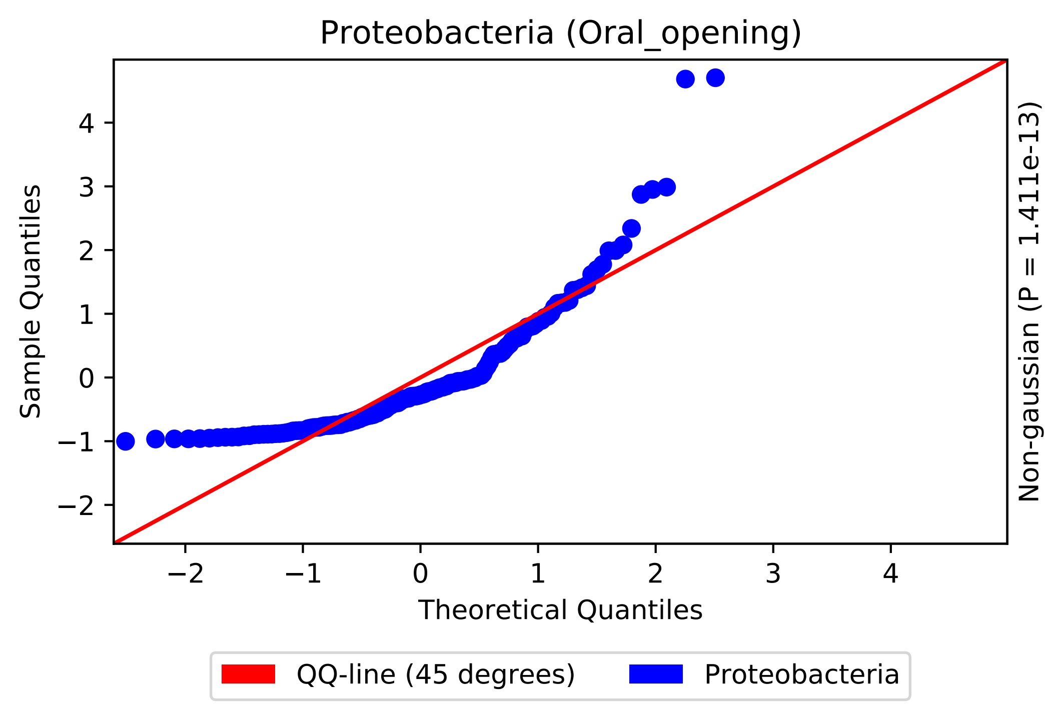

Proteobacteria (Oral_opening)
Sample Quantiles
Theoretical Quantiles
Non-gaussian (P = 1.411e-13)
−2
−1
0
1
2
3
4
QQ-line (45 degrees)
Proteobacteria

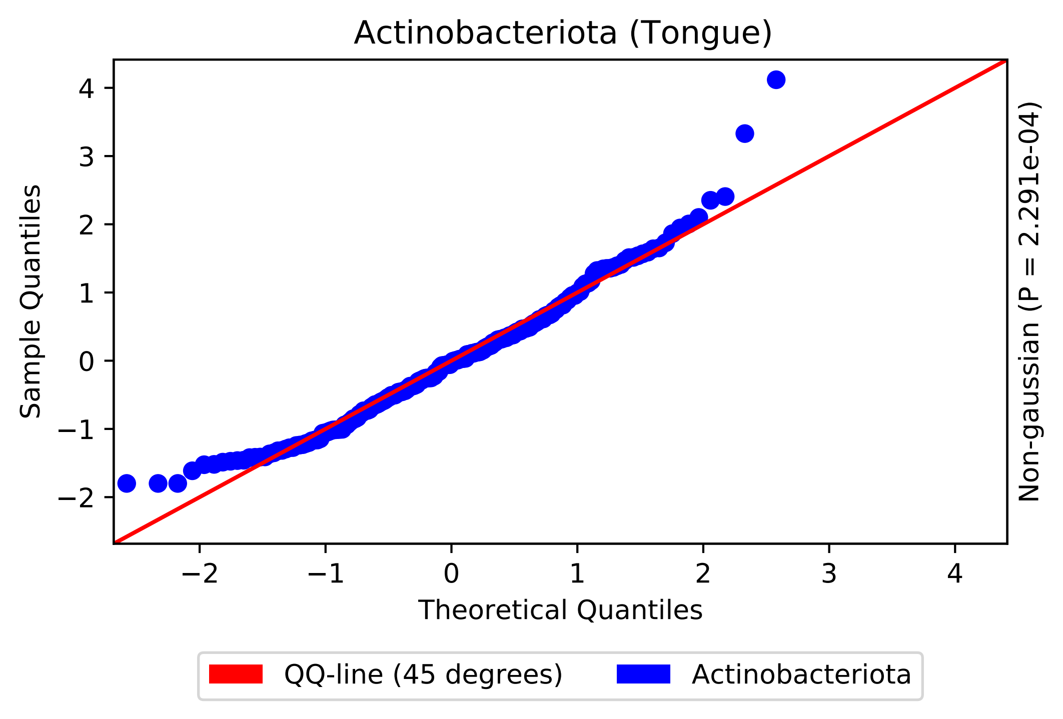

Actinobacteriota (Tongue)
Sample Quantiles
Theoretical Quantiles
Non-gaussian (P = 2.291e-04)
4
3
2
1
0
−1
−2
−2
−1
0
1
2
3
4
QQ-line (45 degrees)
Actinobacteriota

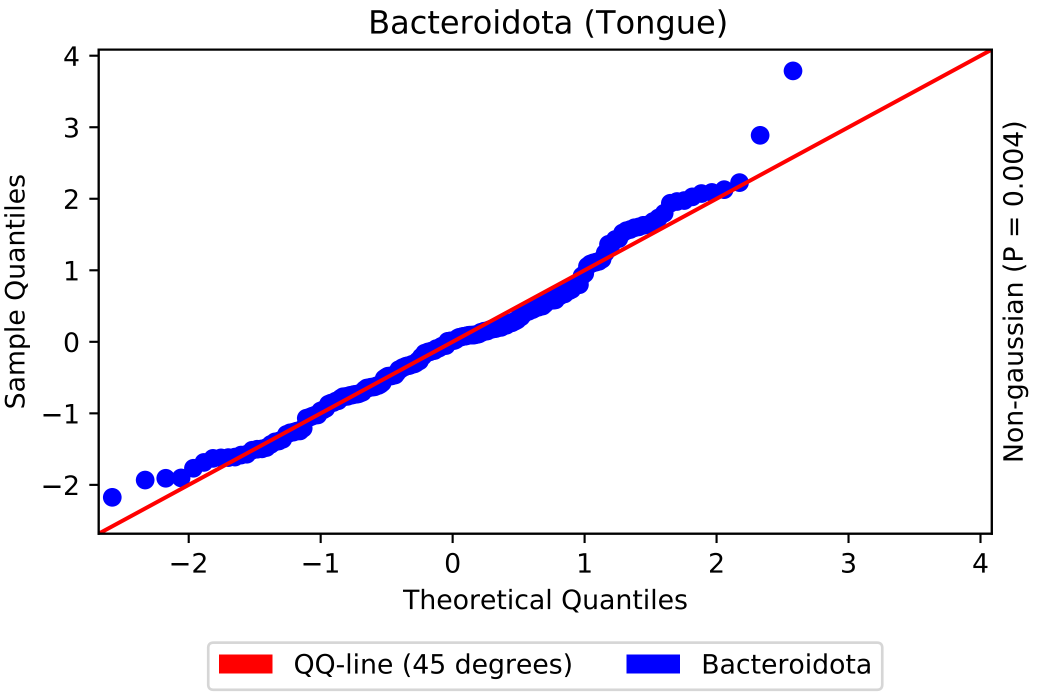
Bacteroidota (Tongue)
Sample Quantiles
Theoretical Quantiles
Non-gaussian (P = 0.004)
QQ-line (45 degrees)
Bacteroidota
4
3
2
1
0
−1
−2
3

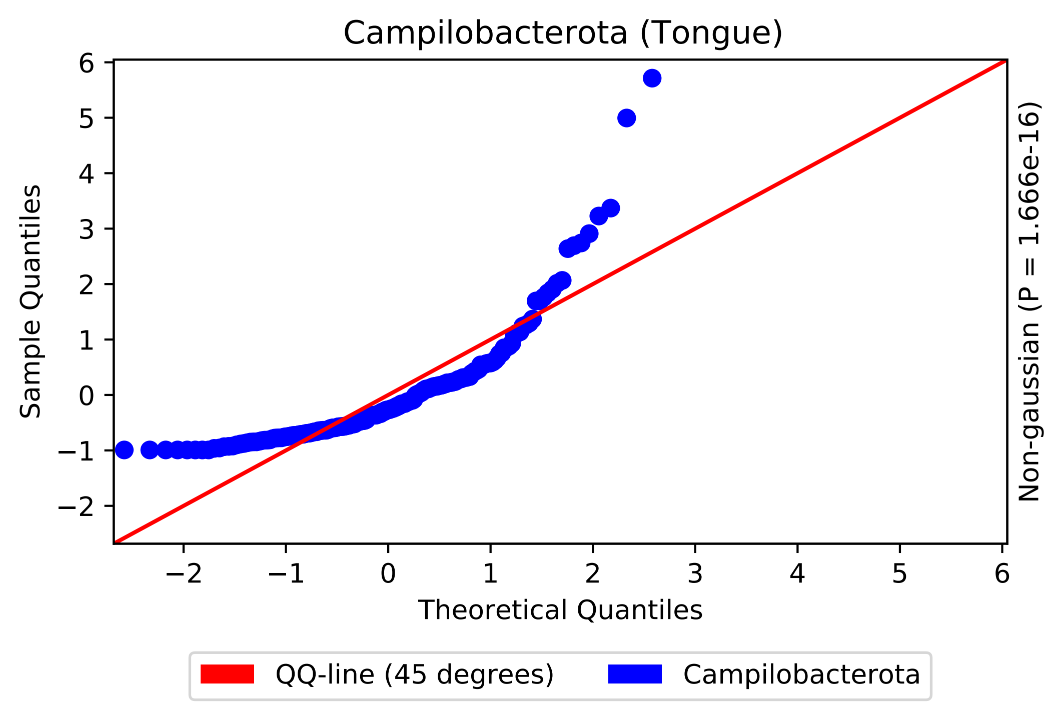
Campilobacterota (Tongue)
Sample Quantiles
Theoretical Quantiles
Non-gaussian (P = 1.666e-16)
QQ-line (45 degrees)
Campilobacterota
6
5
4
3
2
1
0
−1
−2
−2
−1
0
1
2
3
4
5
6

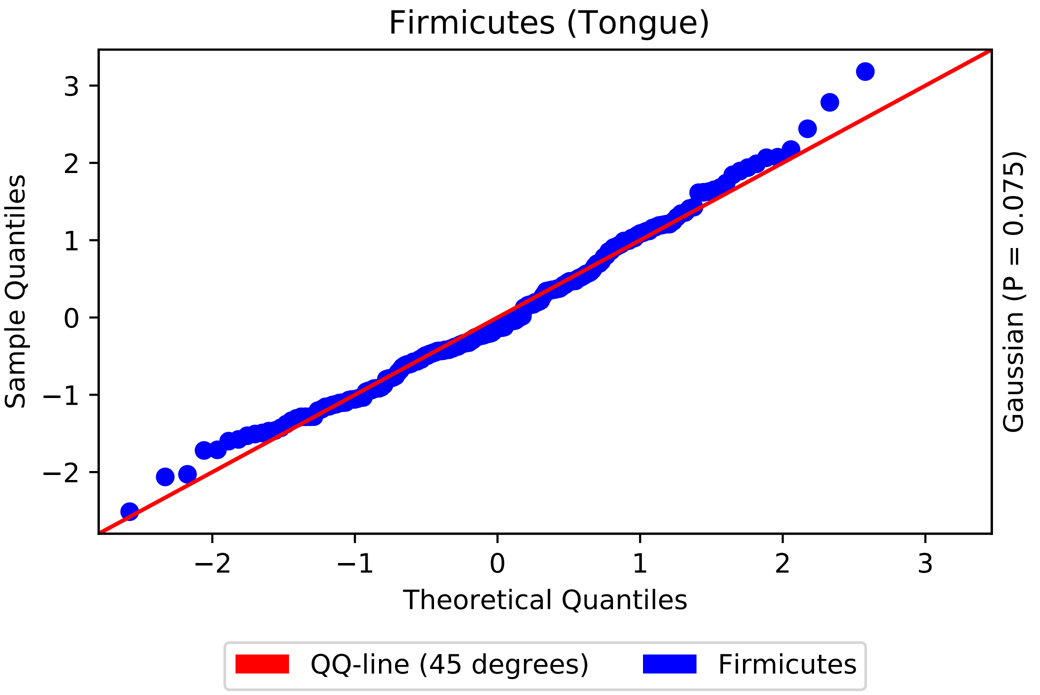

Firmicutes (Tongue)
Sample Quantiles
Theoretical Quantiles
Gaussian (P = 0.075)
QQ-line (45 degrees)
Firmicutes
3
2
1
0
−1
−2
−2
−1
0
1
2
3

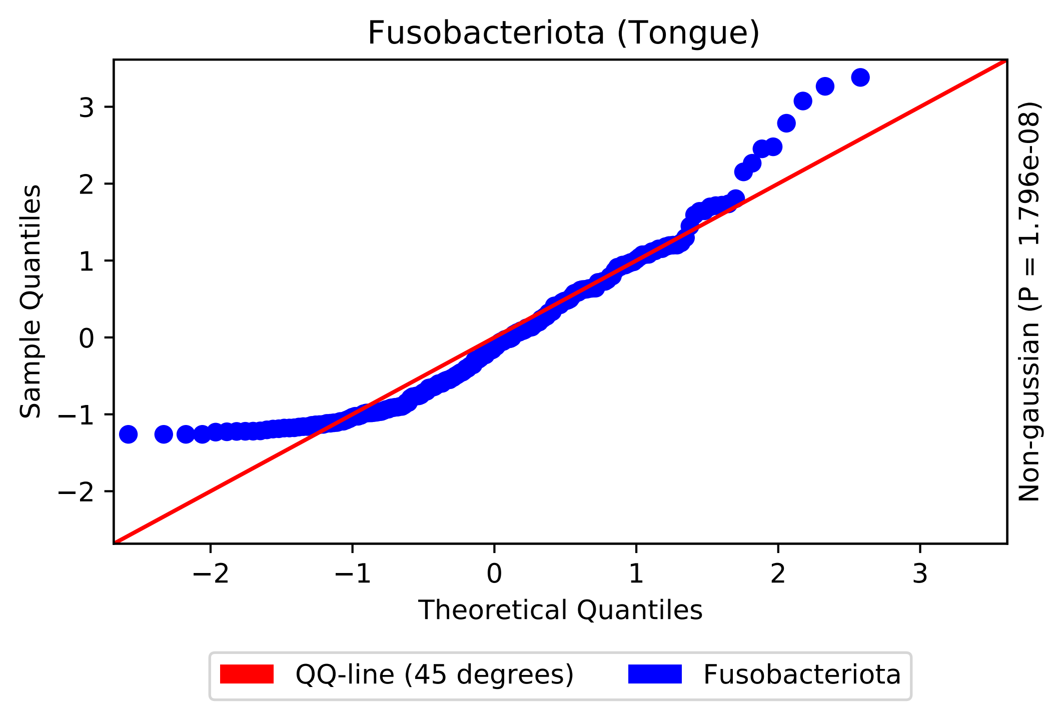

Fusobacteriota (Tongue)
Sample Quantiles
Theoretical Quantiles
Non-gaussian (P = 1.796e-08)
−2
−1
0
1
2
3
QQ-line (45 degrees)
Fusobacteriota

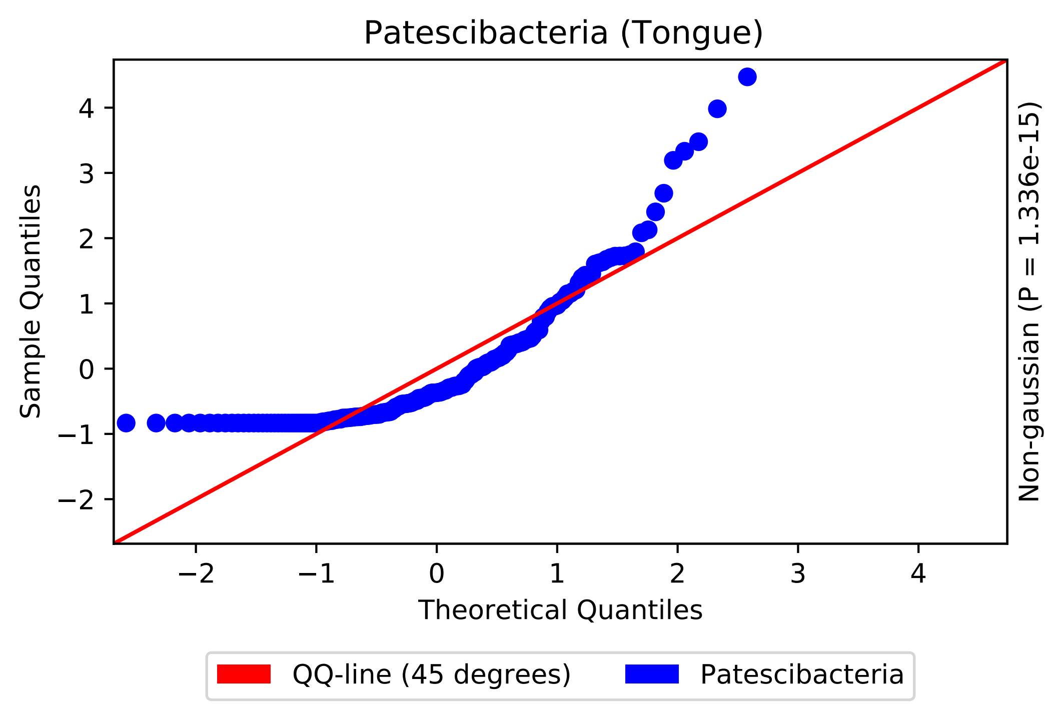

Patescibacteria (Tongue)
Sample Quantiles
Theoretical Quantiles
−2
−1
0
1
2
3
4
Non-gaussian (P = 1.336e-15)
QQ-line (45 degrees)
Patescibacteria

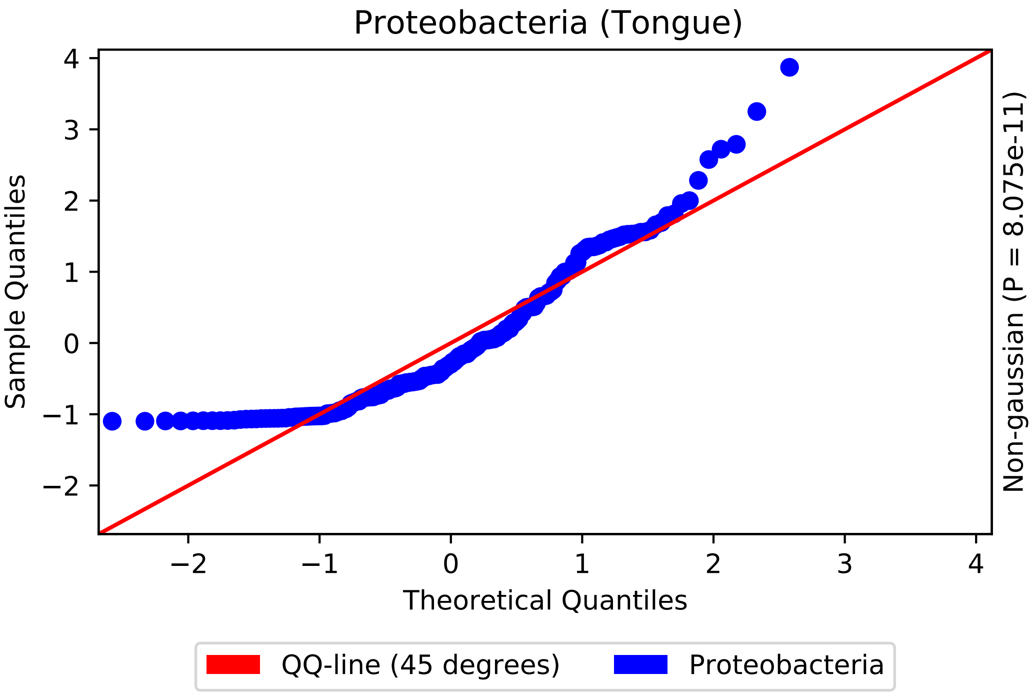

Proteobacteria (Tongue)
Sample Quantiles
Theoretical Quantiles
Non-gaussian (P = 8.075e-11)
QQ-line (45 degrees)
Proteobacteria
4
3
2
1
0
−1
−2
−2
−1
0
1
2
3
4

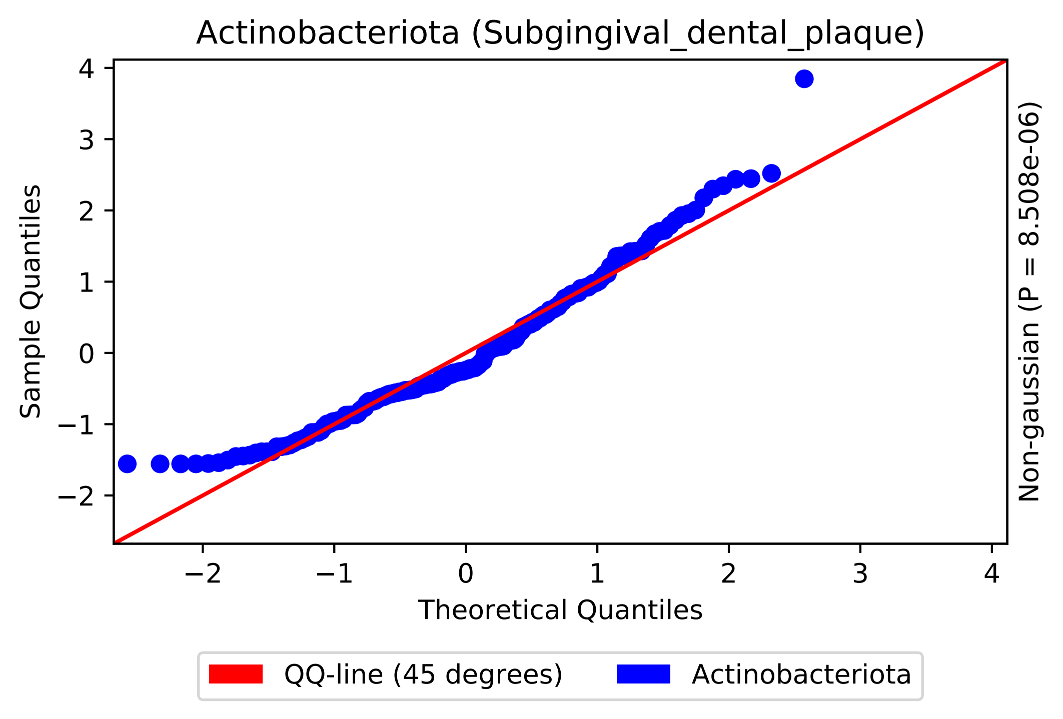

Actinobacteriota (Subgingival_dental_plaque)
Sample Quantiles
Theoretical Quantiles
Non-gaussian (P = 8.508e-06)
4
3
2
1
0
−1
−2
−2
−1
0
1
2
3
4
QQ-line (45 degrees)
Actinobacteriota

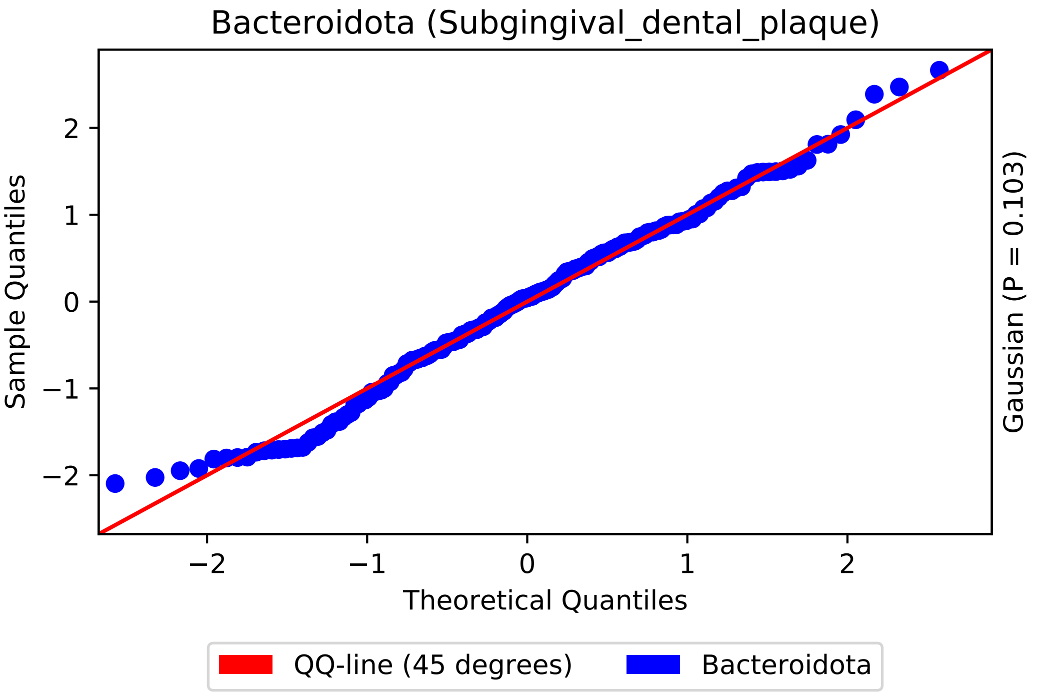
Bacteroidota (Subgingival_dental_plaque)
Sample Quantiles
Theoretical Quantiles
Gaussian (P = 0.103)
−2
−1
0
1
2
QQ-line (45 degrees)
Bacteroidota

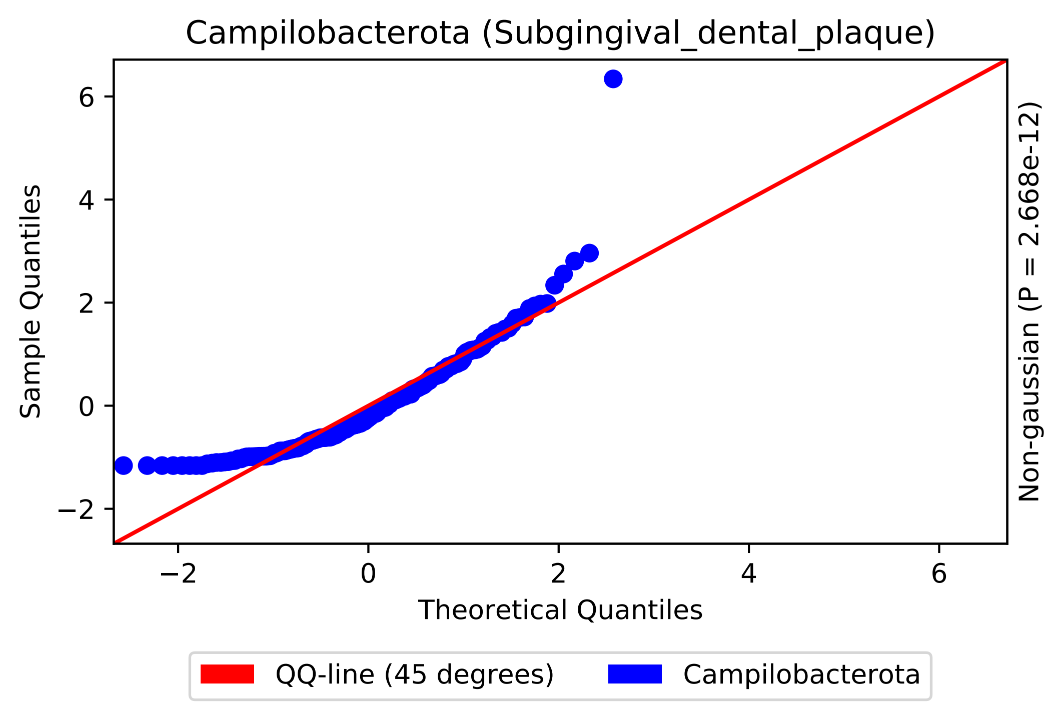
Campilobacterota (Subgingival_dental_plaque)
Sample Quantiles
Theoretical Quantiles
Non-gaussian (P = 2.668e-12)
QQ-line (45 degrees)
Campilobacterota
6
4
2
0
−2
−2
0
2
4
6

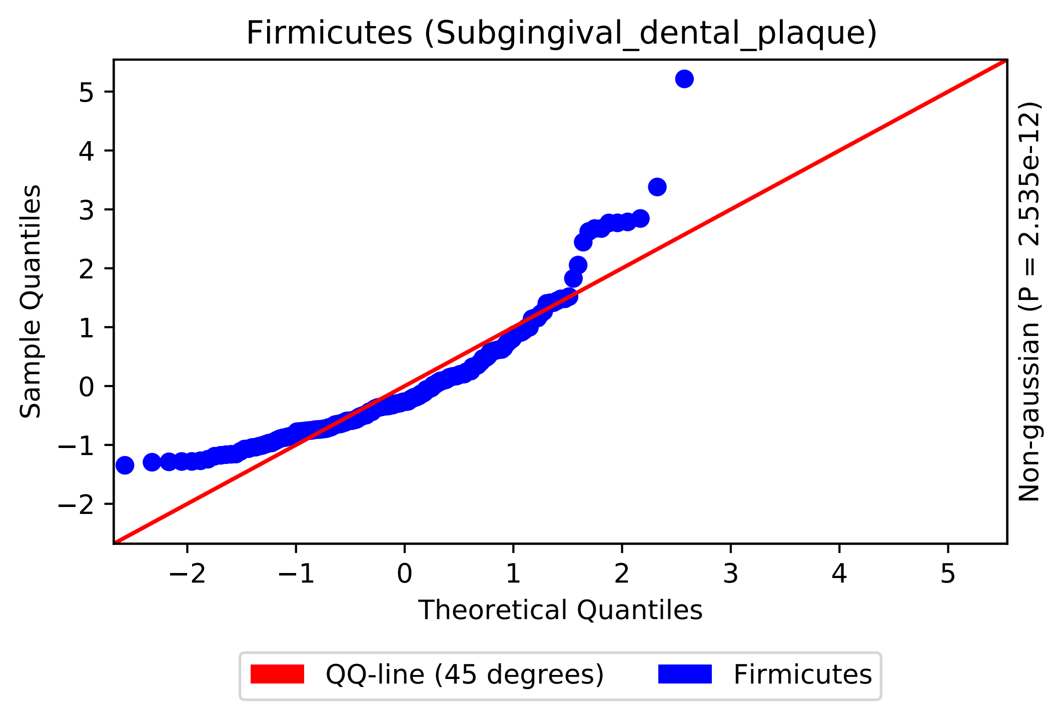
Firmicutes (Subgingival_dental_plaque)
Sample Quantiles
Theoretical Quantiles
Non-gaussian (P = 2.535e-12)
−2
−1
0
1
2
3
4
5
QQ-line (45 degrees)
Firmicutes

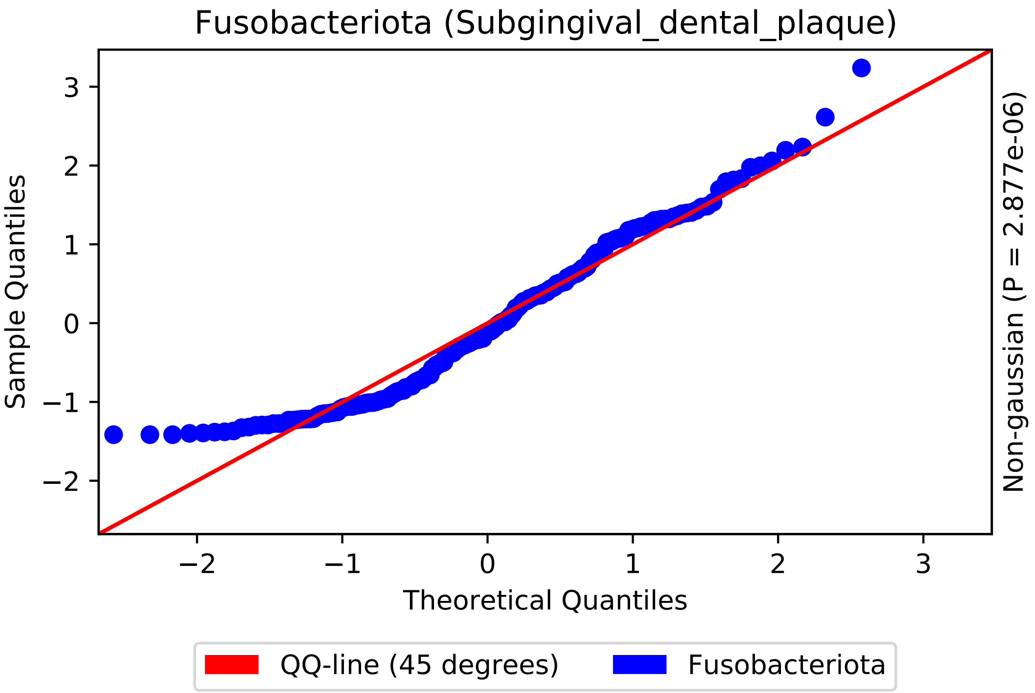
Fusobacteriota (Subgingival_dental_plaque)
Sample Quantiles
Theoretical Quantiles
Non-gaussian (P = 2.877e-06)
−2
−1
0
1
2
3
QQ-line (45 degrees)
Fusobacteriota

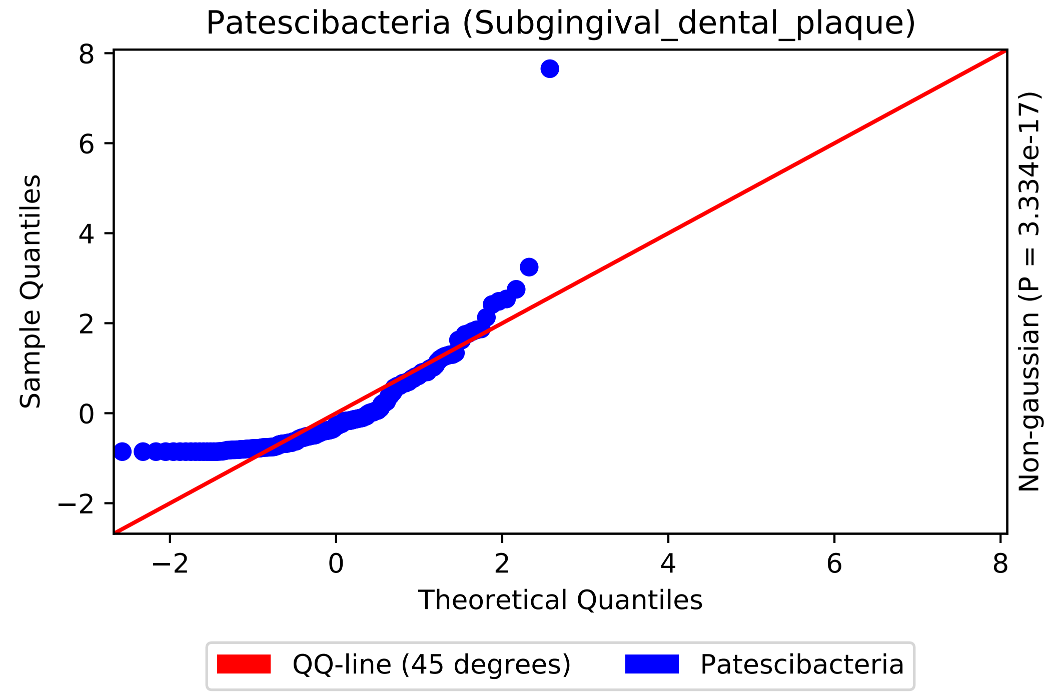
Patescibacteria (Subgingival_dental_plaque)
Sample Quantiles
Theoretical Quantiles
Non-gaussian (P = 3.334e-17)
QQ-line (45 degrees)
Patescibacteria

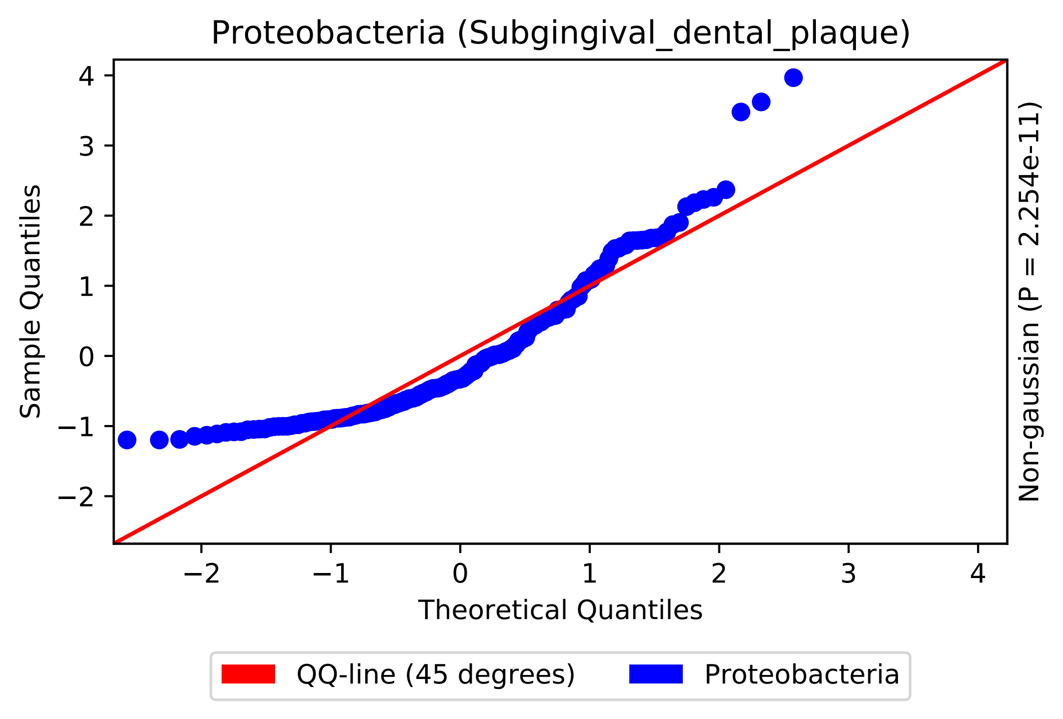
Proteobacteria (Subgingival_dental_plaque)
Sample Quantiles
Theoretical Quantiles
Non-gaussian (P = 2.254e-11)
4
3
2
1
0
−1
−2
−2
−1
0
1
2
3
4
QQ-line (45 degrees)
Proteobacteria

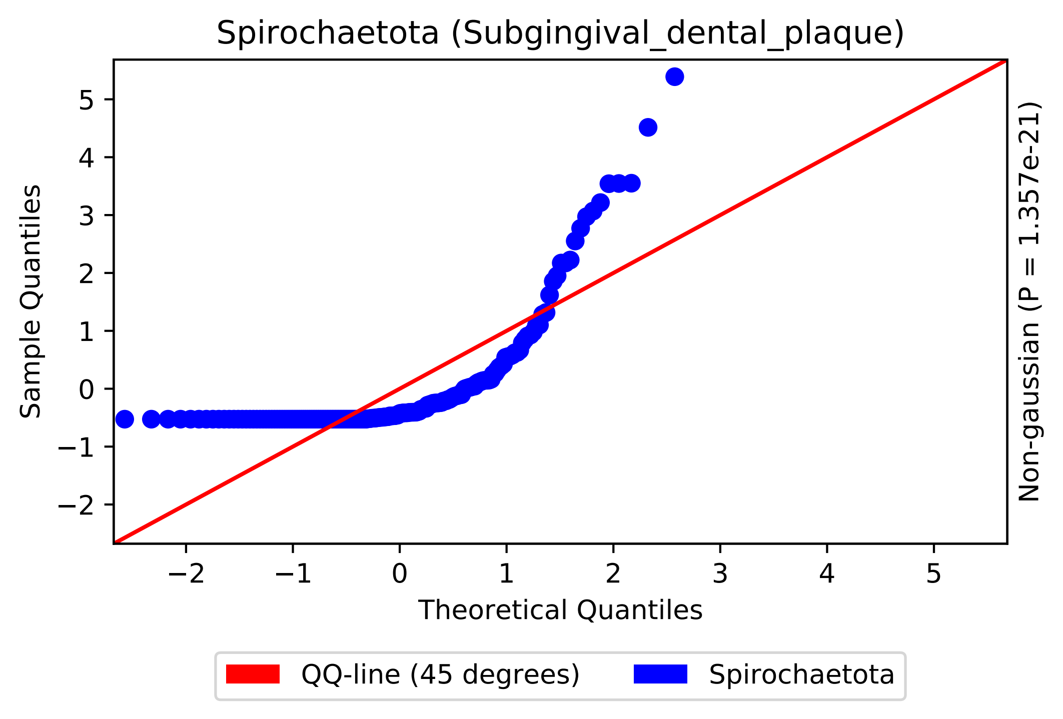
Spirochaetota (Subgingival_dental_plaque)
Sample Quantiles
Theoretical Quantiles
Non-gaussian (P = 1.357e-21)
−2
−1
0
1
2
3
4
5
QQ-line (45 degrees)
Spirochaetota

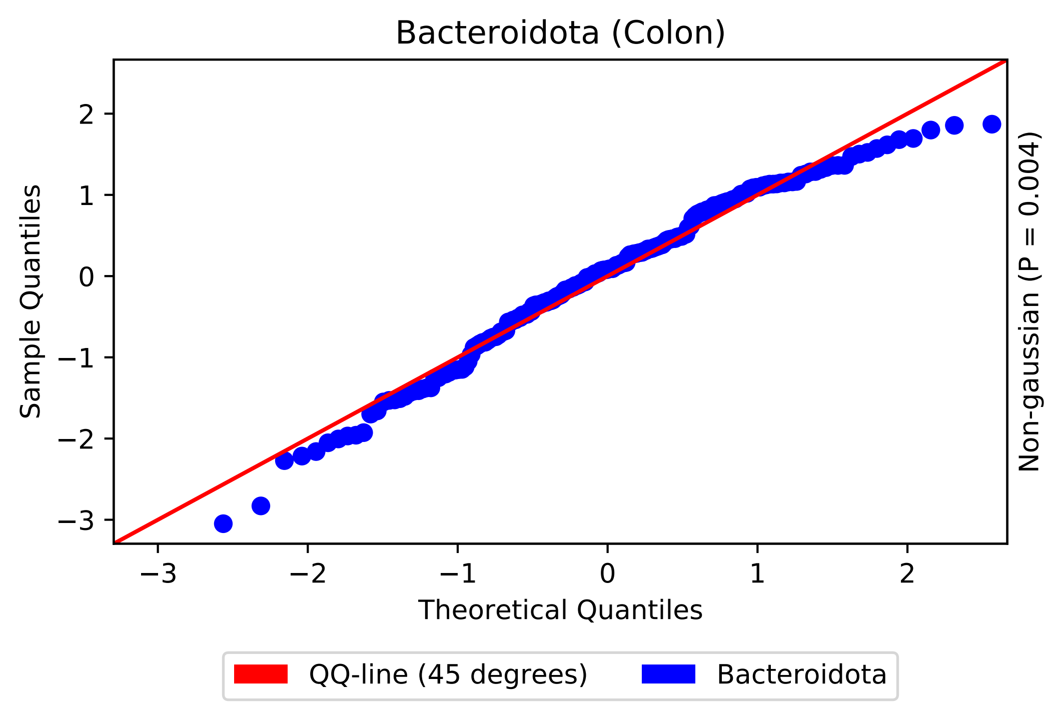
Bacteroidota (Colon)
Sample Quantiles
Theoretical Quantiles
Non-gaussian (P = 0.004)
−3
−2
−1
0
1
2
QQ-line (45 degrees)
Bacteroidota

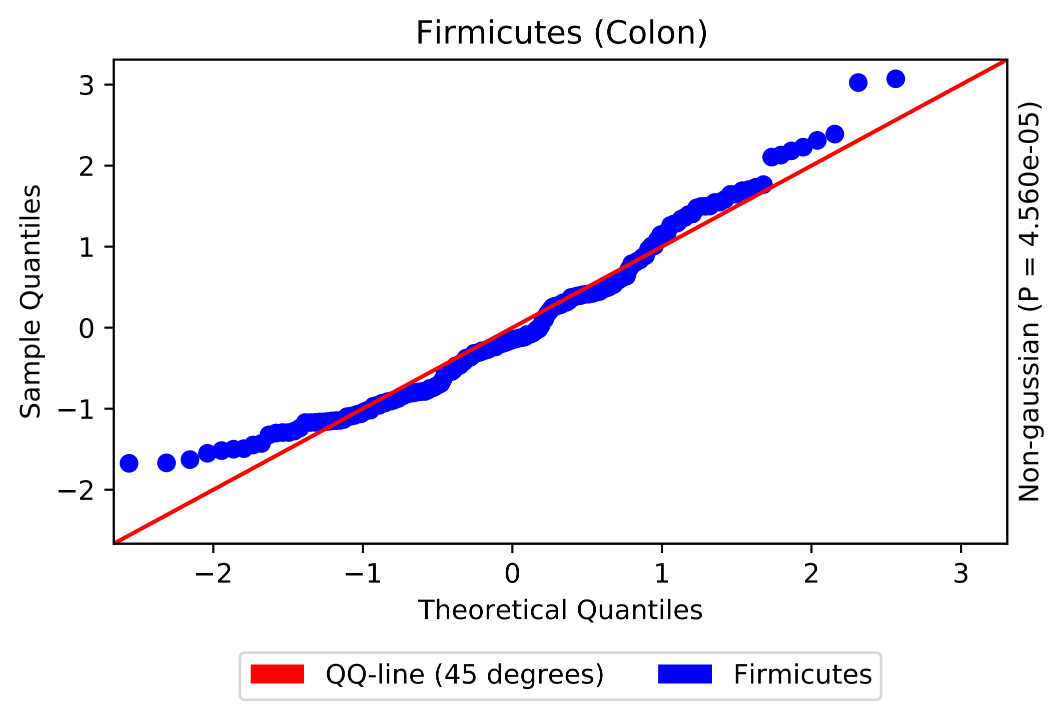

Firmicutes (Colon)
Sample Quantiles
Theoretical Quantiles
Non-gaussian (P = 4.560e-05)
3
2
1
0
−1
−2
−2
−1
0
1
2
3
QQ-line (45 degrees)
Firmicutes

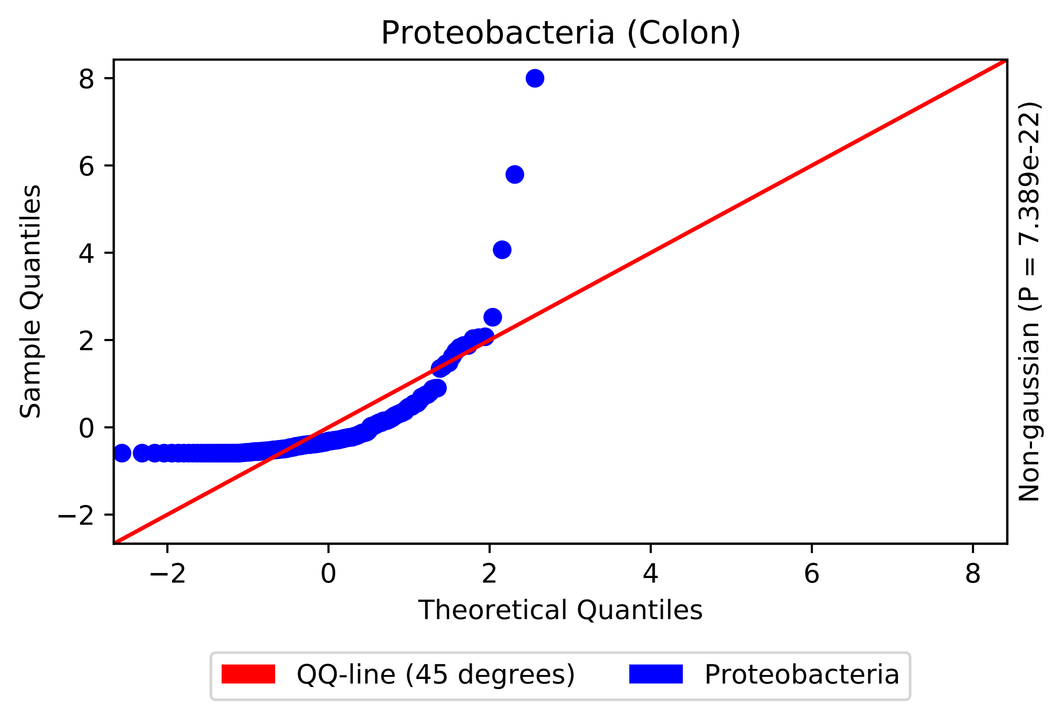
Proteobacteria (Colon)
Sample Quantiles
Theoretical Quantiles
Non-gaussian (P = 7.389e-22)
8
6
4
2
0
−2
−2
0
2
4
6
8
QQ-line (45 degrees)
Proteobacteria

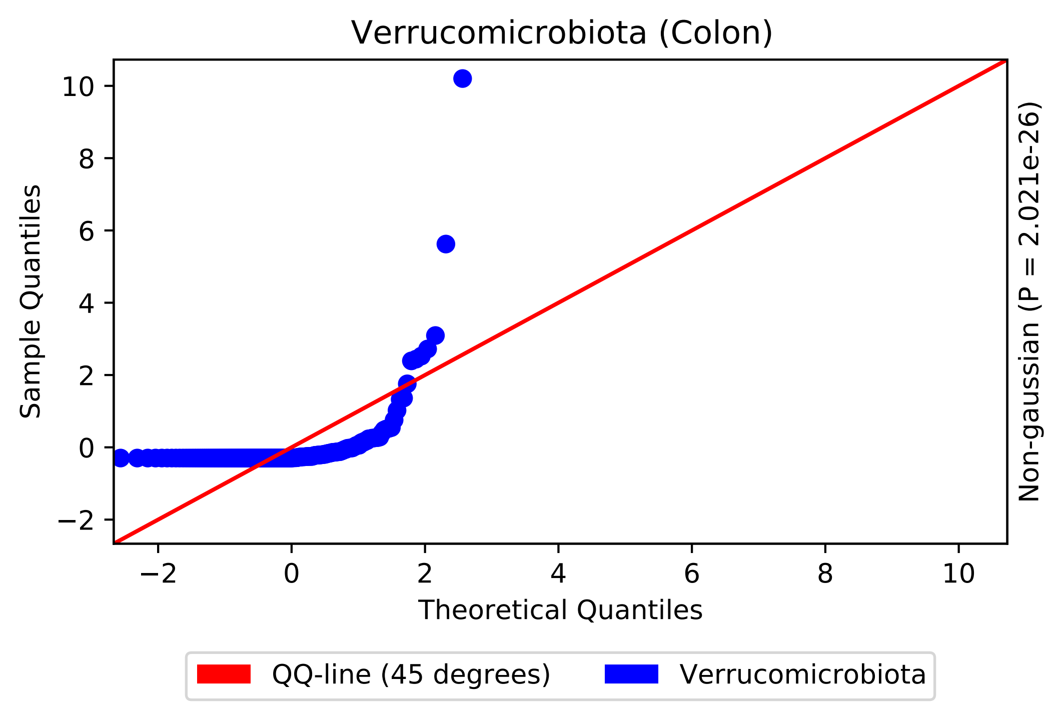
Verrucomicrobiota (Colon)
Sample Quantiles
Theoretical Quantiles
Non-gaussian (P = 2.021e-26)
10
8
6
4
2
0
−2
−2
0
2
4
6
8
10
QQ-line (45 degrees)
Verrucomicrobiota
